# Reanalysis And Integration Of Public Microarray Datasets Reveals Novel Host Genes Modulated In Leprosy

**DOI:** 10.1101/824805

**Authors:** Thyago Leal-Calvo, Milton Ozório Moraes

## Abstract

**Background:** Leprosy is an insidious disease caused primarily by mycobacteria. The difficulties in culturing this slow-growing bacteria together with the chronic progression of the disease have hampered the development of accurate methods for diagnosis. Host gene expression profiling is an important tool to assess overall tissue activity, whether in health or disease conditions. High-throughput gene expression experiments have become popular over the last decade or so, and public databases have been created to easily store and retrieve these data. This has enabled researchers to reuse and reanalyze existing datasets with the aim of generating novel and or more robust information. In this work, after a systematic search, nine microarray datasets evaluating host gene expression in leprosy were reanalyzed and the information was integrated to strengthen evidence of differential expression for several genes.

**Results:** Reanalysis of individual datasets revealed several differentially expressed genes (DEGs). Then, five integration methods were tested, both at the P-value and effect size level. In the end, random effects model (*REM*) and ratio association (*sdef*) were selected as the main methods to pinpoint DEGs. Overall, some classic gene/pathways were found corroborating previous findings and validating this approach for analysis. Also, various original DEGs related to poorly understood processes in leprosy were described. Nevertheless, some of the novel genes have already been associated with leprosy pathogenesis by genetic or functional studies, whilst others are, as yet, unrelated or poorly studied in these contexts.

**Conclusions:** This study reinforces evidences of differential expression of several genes and presents novel genes and pathways associated with leprosy pathogenesis. Altogether, these data are useful in better understanding host responses to the disease and, at the same time, provide a list of potential host biomarkers that could be useful in complementing leprosy diagnosis based on transcriptional levels.

## Background

Leprosy is a chronic infectious disease caused by *Mycobacterium leprae*. This pathogen preferably resides in skin macrophages and Schwann cells in peripheral nerves, which often results in strong inflammatory responses and nerve dysfunction [1–3]. Although curable, the World Health Organization (WHO) reported in 2017 a stagnant number of new leprosy cases globally, of which India and Brazil were the leaders [4]. Despite the effectiveness of treatment, transmission likely occurs prior to diagnosis since it is primarily detected based solely on clinical findings. As yet there is no gold standard method to differentiate asymptomatic infection from disease, and early detection and unambiguous diagnosis is incredibly difficult [5–8]. As exposure to the etiological agent is not sufficient to develop the disease [9], it is assumed that early diagnosis is key in controlling the disease transmission, along with chemo- and immunoprophylactic strategies for high risk individuals, such as household contacts [10]. Leprosy is considered a complex disease, as environmental and host genetic factors can affect the outcome during the different steps of the natural disease course. The stages that can be influenced can include mycobacterial clearance, progression to localized (tuberculoid, TT) or disseminated (lepromatous, LL) forms, and occurrence of reactional episodes (reversal reaction, RR, or erythema nodosum leprosum, ENL) [5,11,12].

To date, large-scale approaches, such genotyping through genomic scans, genome-wide association studies or whole exome sequencing, or expression analysis through microarrays and RNA sequencing, have been connecting several important pathways and genes to the pathogenesis of leprosy [13–16]. High-throughput gene expression analysis can provide valuable information in the identification of genes involved in host responses to infection, to assess disease severity, and to discover biomarkers for both diagnosis and prognosis. These associations between either single nucleotide polymorphisms (SNP) depicted in genetic studies or genes/transcripts found in expression analysis, must then be independently confirmed using mechanistic biological studies [7,16,17]. In this regard, gene functional analysis is also an important aspect in defining host-*M. leprae* interaction [18–20]. The popularity of large-scale studies coupled with the search for scientific reproducibility and open science, has led to the creation of databases for the easy storage and retrieval or public data. NCBI’s Gene Expression Omnibus (GEO), one of the most important databases for gene expression data, has to this date 119,721 entries [21,22]. Despite their comprehensiveness, microarray-based studies may report findings that are not reproducible, or that fail to detect minor differences among samples [23]. The combination of multiple independent studies can therefore increase reliability and generalizability of results [24], especially because manually comparing studies with distinct designs is not a trivial task. Therefore, reanalysis of existing data can yield relevant information that were not of immediate interest to the researchers who initially conducted the experiment. Reanalysis can also be used to independently validate related experiments, resulting in more robust evidences of differential expression [25–27].

In this study, nine host gene expression studies linked to leprosy that had been deposited in GEO were comprehensively reanalyzed and integrated. Overall, the datasets of these experiments included human samples from all clinical classifications of the leprosy spectrum and experimental samples of murine cells. Genes that are consistently modulated across different independent studies were identified, and consequently yielded a significantly reduced number of false-positives and spurious signals than what would be expected by considering only one experiment. Some of these identified genes have been independently reported before, thus acting as analytical positive controls. Pathways that are clearly involved in leprosy pathogenesis, such as type 1 interferon and neutrophil mediated immunity, were enriched with these novel genes and therefore may be useful for constructing molecular signatures to improve diagnosis or be targets for prophylaxis and therapeutics.

## Results

### Search results and individual reanalysis

A systematic search was conducted using specific keywords in GEO to identify leprosy-related datasets (See methods). Up until October 2017, 18 datasets were found, of which 9 (8 human, 1 murine) were selected for reanalysis (Figure 1). Excluding criteria of datasets comprised those measuring non-coding RNAs or pathogen gene expression, studies involving compound stimulus, and/or those with only one sample biological group. Individual analysis of these public datasets revealed many differentially expressed genes (DEGs) across several comparisons within each study, encompassing various leprosy clinical forms and some *in vitro* experimental conditions, such as stimulus with live or sonicated *M. leprae*. As expected, in studies with few biological replicates no DEGs were detected after adjusting P-values for multiple testing. For this reason, the Guerreiro *et al*. (2013) [28] dataset was excluded from subsequent integration analyses. Table 1 summarizes some information from the reanalyzed datasets and their public accession IDs.

**Table 1.** Description of the nine GEO datasets used in this study.

| GEO Accession | Organism | Sample type | Sample groups (n) | Microarray Platform | Comparison | Ref. |
| --- | --- | --- | --- | --- | --- | --- |
| **GSE40950<sup>†</sup>** | *H. sapiens* | Cell culture | Infected (6), Control (6) | Stanford Custom | NA | 28 |
| **GSE35423** | *H. sapiens* | Cell culture | Infected 24h (2), Control 24h (2), Infected 48h (2), Control 48h (2) | Applied Biosystems | In vitro | 14 |
| **GSE24280** | *H. sapiens* | Skin biopsy, blood* | BT (6), MB (6), BT blood (6), MB blood (6), healthy skin (1) | NimbleGen | MB vs. PB<br>LL vs. HS | NA |
| **GSE443** | *H. sapiens* | Skin biopsy | LL (6), BT (5) | HG_U95Av2<br>Affymetrix | MB vs. PB | 80 |
| **GSE17763** | *H. sapiens* | Skin biopsy | LL (7), BT (10), RR (7) | HG-U133 Affymetrix | MB vs. PB | 81 |
| **GSE16844** | *H. sapiens* | Skin biopsy | ENL (6), LL (7) | HG-U133 Affymetrix | LL vs. ENL | 47 |
| **GSE74481** | *H. sapiens* | Skin biopsy | TT (10), BT (10), BB (10), BL (10), LL (4), RR (14), ENL (10), healthy (9) | Agilent SurePrint | MB vs. PB<br>LL vs. HS<br>LL vs. ENL | 82 |
| **GSE100853** | *H. sapiens* | Whole blood culture | Stimulated (51), Mock cultures (51) | Illumina HumanHT | In vitro | 83 |
| **GSE95748** | *M. musculus* | Cell culture | Control (2), Infected 14 days (2), Infected 28 days (2), pSLCs (2) | Affymetrix Mouse | In vitro | 52 |

<sup>†</sup>Dataset excluded from integration/meta-analysis due to no differentially expressed genes during individual analysis. \*Blood samples were not considered in this work, because it was only represented in one dataset at the time. MB: multibacillary leprosy; TT: tuberculoid leprosy; BT: borderline tuberculoid; BB: borderline-borderline; BL: borderline-lepromatous; LL: lepromatous lepromatous; RR: reversal reaction or type 1 reaction (T1R); ENL: erythema nodosum leprosum or type 2 reaction (T2R).

**Fig 1.**
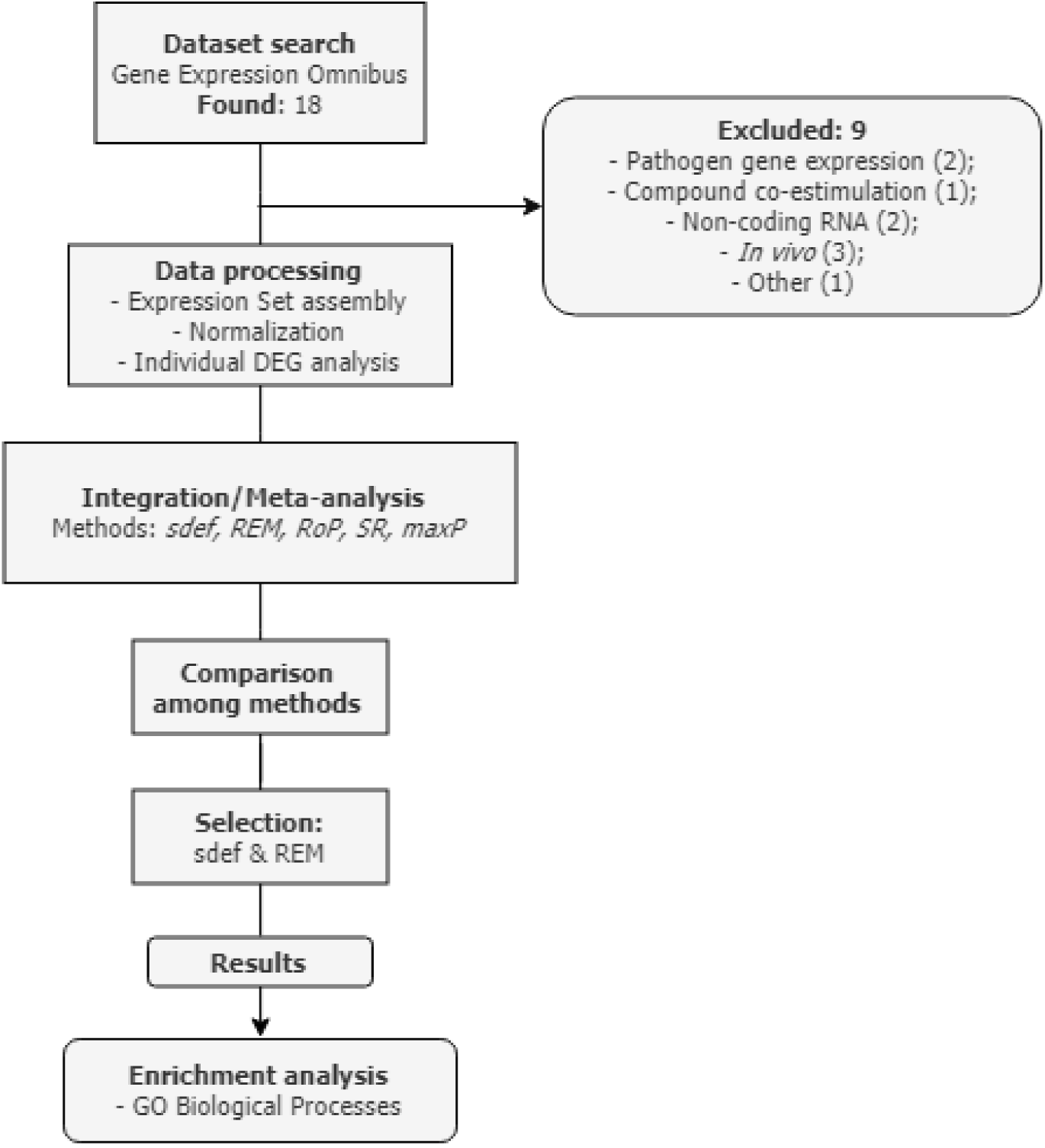
Simplified flowchart of the reanalysis process. Initially, GEO was queried with specific keywords aiming at finding leprosy-related datasets. Further, studies were excluded according to study parameters, such as pathogen transcriptome profiling only, datasets measuring non-coding RNAs, compound stimulus and with insufficient groups or biological replicates. Then, the nine datasets were preprocessed individually using common statistical procedures and thresholds whenever possible, also including available covariates that could confound group-specific differential gene expression. Next, both the lists of DEG with their P-values normalized gene expression matrices were used with integration/meta-analytical tools to summarize evidence of differential expression across studies for related comparisons. Finally, the consolidated DEG lists were used as input in over-represented analysis in order to translate genes to biological processes according to Gene Ontology annotation.

### Common differentially expressed genes by the ratio association (*sdef*) method and meta-analysis tools

After individual differential expression analysis for each of the nine included studies, samples were grouped from the different studies into four comparison categories based on clinical features to find common DEGs, i.e. LL vs. BT (4 independent datasets); LL vs. ENL (2 ind. datasets); LL vs. Control (2 ind. datasets); *in vitro* experiments of Stimulated vs. Control (3 ind. datasets, 6 comparisons) (Table 1 and Figure 2). Hence, only genes common to most or all studies within each category, mapped by Entrezid, were used in the ratio association test, hereafter referred as the *sdef* method, and the meta-analysis procedures (see Methods). According to the publication that defined *sdef*, the method is considered conservative with less type 1 errors in expense of more type 2 errors [29]. Additional gene expression meta-analysis tools from *MetaDE* R package were also used, such as the P-value-based methods, r-th ordered P-value (*rOP*), maximum P-value (*maxP*), and sum of ranks (*SR*), and the random effects model (*REM*), which is based on the effect size.

**Fig 2.**
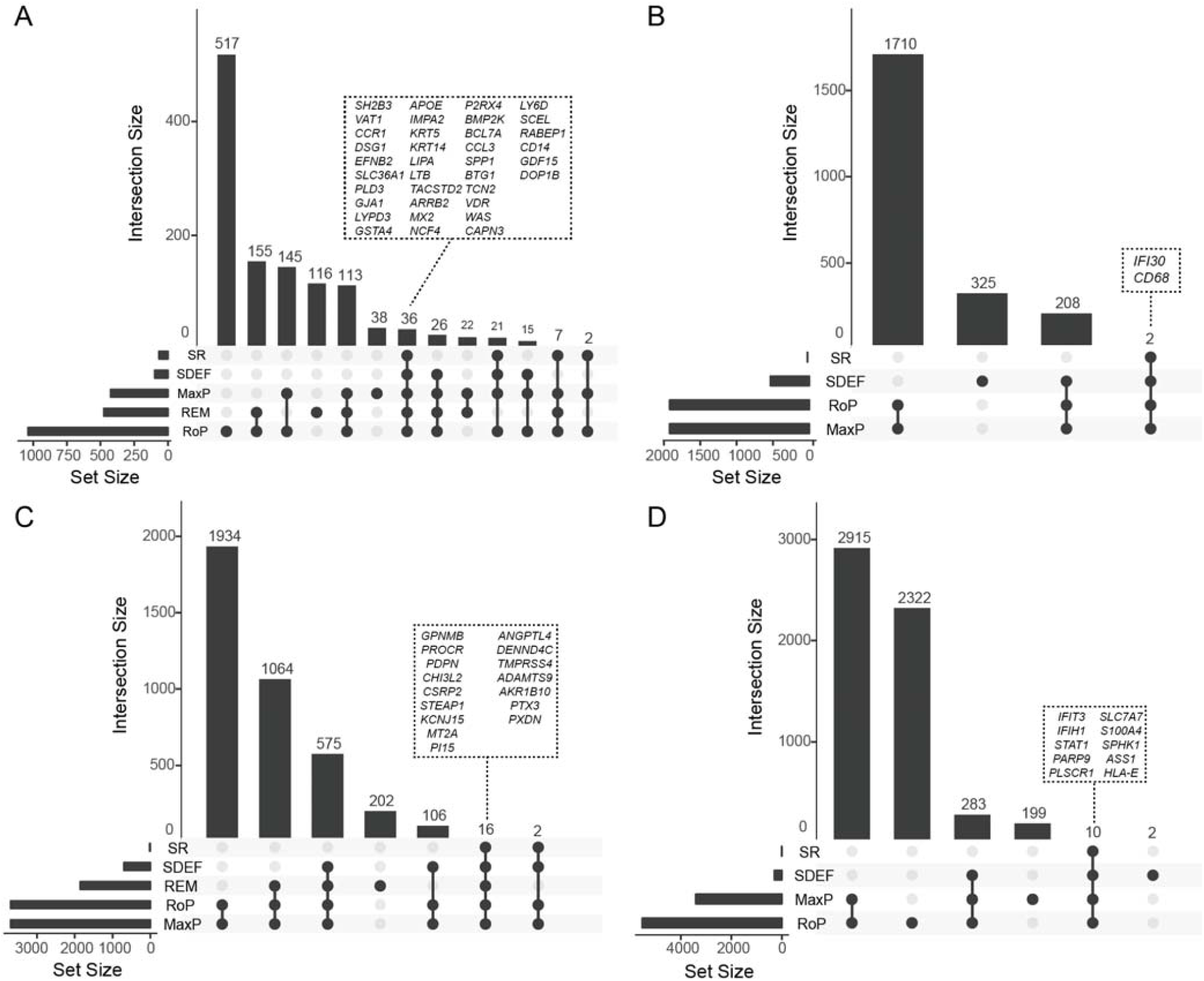
Upset plots showing the number of identified genes from each method and their intersection. Vertical bars show the number common genes (intersection size) from a given set of methods (bottom filled connected circles). Horizontal bars illustrate the total number of genes (set size) output from each integration or meta-analysis method. Dashed boxes contain genes selected in common from all methods. **a** Genes selected for LL vs. BT comparisons (4 independent datasets). **b** LL vs. Control (2 ind. datasets); **c** LL vs. ENL (2 ind. datasets); **d** *in vitro* experiments stimulated vs. Control (3 ind. datasets, 6 comparisons).

Figure 2 shows the number of genes selected, with a false discovery rate (FDR) less than or equal to 0.1, from each integration approach and the intersection between the different methods. The *rOP* and *maxP* methods were more liberal and selected more DEGs for all comparison categories. In the two-study cases, such as LL vs. ENL, the *maxP* and *rOP* are equivalent and selected the same genes (Figure 1bc). As expected, *sdef* selected few genes (98), but the most conservative procedure was the *SR* tool, which resulted in 66 genes in LL vs. BT (Figure 1a), and even fewer for the other comparisons (Figure 1bcd). Overall, the ratio association method (*sdef*) gave intermediate results, selecting more genes than *SR*, but fewer than *REM*, *rOP*, or *maxP*. Unlike all the other approaches, the *REM* method considers the effect size and its sign/direction, where larger effects with the same sign across studies are more likely to be truly differentially expressed. For this reason, the *REM* was used to select genes from LL vs. BT and LL vs. ENL comparisons. However, due to some dataset-specific limitations (See methods section), the *sdef* method was used for LL vs. Control and Stimulated vs. Control (*in vitro*) comparisons (Additional File 1 – Tables S3 and S4; Additional File 2 – Figures S1 and S2).

Table 2 shows the top 20 genes selected from the LL vs. BT comparison using the *REM* approach. With a FDR at 10%, 475 genes were considered statistically significant by this method and 36 were identified by all five tested methods (Figure 1a). Excluding the *SR*, 26 other genes were also considered differentially expressed, totaling 501 genes. Among the top selected DEGs with higher expression in LL than BT skin biopsies are: *CCR1*, *CD14*, *GDF15*, *APOE*, *APOC1*, *P2RX4*, *LIPA*, and *LGALS9*. Whilst for the converse, some of the more expressed in BT than LL are: *GATA3*, *KRT5*, *LTB*, *VDR*, *KRT14*, *DSG*, *SCEL*, *GBP2*, *S100B*, and *NEBL* (Table 2; Additional File 1 – Table S1).

**Table 2.** Top 20 genes selected by the REM from LL vs. BT comparison across four datasets.

| Entrezid | Symbol | std. log<sub>2</sub>FC | 95% CI | FDR | Tau<sup>2</sup> |
| --- | --- | --- | --- | --- | --- |
| 1230 | *CCR1* | 1.872 | [1.217, 2.527] | 0.0001 | 0.000 |
| 3852 | *KRT5* | -2.040 | [-2.782, -1.298] | 0.0001 | 0.087 |
| 7421 | *VDR* | -1.785 | [-2.433, -1.136] | 0.0001 | 0.000 |
| 929 | *CD14* | 1.800 | [1.148, 2.452] | 0.0001 | 0.000 |
| 2941 | *GSTA4* | -1.759 | [-2.406, -1.113] | 0.0001 | 0.000 |
| 2697 | *GJA1* | -1.695 | [-2.332, -1.058] | 0.0002 | 0.000 |
| 605 | *BCL7A* | -1.857 | [-2.557, -1.157] | 0.0002 | 0.058 |
| 1948 | *EFNB2* | -1.653 | [-2.292, -1.013] | 0.0002 | 0.000 |
| 9518 | *GDF15* | 2.077 | [1.275, 2.879] | 0.0002 | 0.174 |
| 5783 | *PTPN13* | -1.702 | [-2.367, -1.037] | 0.0003 | 0.029 |
| 6351 | *CCL4* | 1.616 | [0.984, 2.247] | 0.0003 | 0.000 |
| 4689 | *NCF4* | 1.566 | [0.938, 2.195] | 0.0005 | 0.000 |
| 3613 | *IMPA2* | -1.776 | [-2.491, -1.061] | 0.0005 | 0.090 |
| 1212 | *CLTB* | -1.551 | [-2.177, -0.924] | 0.0005 | 0.000 |
| 3615 | *IMPDH2* | -1.546 | [-2.175, -0.916] | 0.0005 | 0.000 |
| 348 | *APOE* | 1.522 | [0.899, 2.144] | 0.0006 | 0.000 |
| 55556 | *ENOSF1* | 1.528 | [0.897, 2.158] | 0.0007 | 0.005 |
| 2212 | *FCGR2A* | 1.716 | [0.996, 2.436] | 0.0008 | 0.103 |
| 27076 | *LYPD3* | -1.475 | [-2.095, -0.855] | 0.0008 | 0.000 |
| 7291 | *TWIST1* | -1.479 | [-2.101, -0.857] | 0.0008 | 0.000 |

Genes shown are the top 20 with smallest adjusted P-value (FDR). FDR, false discovery ratio. Tau<sup>2</sup> is a represents the between-study variance.

Table 3 shows the top 20 genes selected by the *REM* method for LL vs. ENL comparison from two studies. With a 10% FDR, 1857 genes were selected (Figure 2b, Additional File 1 – Table S2). Of these, 16 were commonly discovered by all five methods, and excluding the *SR*, another 575 genes were common to *REM*, *sdef*, *rOP*, and *maxP* (Figure 2b). Among the top genes found by *REM* that are more expressed in ENL than LL are: *PROCR*, *AKR1B10*, *S100A12*, *PTX3*, *PI15*, *CYP7B1*, *STEAP1*, *LTF*, *ANGPTL4*, and *RNASE2.* Conversely, genes such as *PER3*, *ANKMY2H*, *MOAP1*, *GPNMB*, *LIPA*, *P2RX7*, and *SEPTIN8* have higher expression in LL than ENL skin lesions (Table 3; Additional File 1 – Table S2).

**Table 3.** Top 20 genes selected by the REM from LL vs. ENL comparison across two datasets.

| Entrezid | Symbol | std. log2FC | 95% CI | FDR | Tau2 |
| --- | --- | --- | --- | --- | --- |
| 10544 | *PROCR* | -3.705 | [-4.992, -2.417] | 0.0001 | 0.00 |
| 57016 | *AKR1B10* | -3.940 | [-5.294, -2.586] | 0.0001 | 0.00 |
| 6283 | *S100A12* | -3.853 | [-5.177, -2.529] | 0.0001 | 0.00 |
| 8863 | *PER3* | 3.713 | [2.424, 5.002] | 0.0001 | 0.00 |
| 9420 | *CYP7B1* | -3.571 | [-4.831, -2.311] | 0.0001 | 0.00 |
| 1117 | *CHI3L2* | -3.321 | [-4.527, -2.114] | 0.0001 | 0.00 |
| 57037 | *ANKMY2* | 3.334 | [2.125, 4.544] | 0.0001 | 0.00 |
| 64112 | *MOAP1* | 3.344 | [2.132, 4.556] | 0.0001 | 0.00 |
| 7837 | *PXDN* | -3.446 | [-4.69, -2.202] | 0.0001 | 0.00 |
| 1466 | *CSRP2* | -3.285 | [-4.485, -2.085] | 0.0002 | 0.00 |
| 4502 | *MT2A* | -3.238 | [-4.428, -2.048] | 0.0002 | 0.00 |
| 57178 | *ZMIZ1* | 3.384 | [2.138, 4.629] | 0.0002 | 0.02 |
| 154141 | *MBOAT1* | 3.165 | [1.985, 4.344] | 0.0002 | 0.00 |
| 4495 | *MT1G* | -3.075 | [-4.233, -1.917] | 0.0003 | 0.00 |
| 23034 | *SAMD4A* | 3.061 | [1.905, 4.218] | 0.0003 | 0.00 |
| 56649 | *TMPRSS4* | -3.610 | [-4.98, -2.241] | 0.0003 | 0.13 |
| 10457 | *GPNMB* | 3.003 | [1.852, 4.154] | 0.0003 | 0.00 |
| 51050 | *PI15* | -2.995 | [-4.144, -1.846] | 0.0003 | 0.00 |
| 5806 | *PTX3* | -2.983 | [-4.122, -1.843] | 0.0003 | 0.00 |
| 5879 | *RAC1* | 2.996 | [1.852, 4.14] | 0.0003 | 0.00 |

Genes shown are the top 20 with smallest adjusted P-value (FDR). FDR, false discovery ratio. Tau<sup>2</sup> represents the between-study variance.

### Gene ontology over-representation analysis

After obtaining the lists of DEGs associated in different studies by the *sdef* or *REM* methods, the gene ontology over-representation analysis (ORA) was used to understand the possible biological role of these genes. In addition, to show the most common direction of regulation for genes in a given ontology a score was calculated, where positive scores signify that most of the genes are up-regulated and vice-versa [30].

For the LL vs. BT category, 313 genes with an FDR less than or equal to 0.1 and standardized |log_2_FC| greater than or equal to 1 were used in the analysis. In total, 138 gene ontology (GO) biological processes were enriched with an FDR < 0.1 for this category. The top 20 GO biological processes are shown in Figure 3a (Additional File 3 – Table S5). Several biological processes had genes more expressed in LL than BT, such as ‘innate immune response’, ‘regulation of response to external stimulus’, ‘inflammatory response’, ‘leukocyte migration’, ‘response to lipoprotein particle’ and ‘type 1 interferon signaling pathway’. Whilst, several epithelial ontologies had genes more expressed in BT than LL, like ‘epithelial cell differentiation’, ‘epidermis development’, ‘keratinocyte differentiation’ and ‘cornification’ (Figure 3a; Additional File 3 – Table S5). Figure 3b shows some enriched processes together with the standardized log_2_FC for the annotated genes (Figure 3b; Additional File 3 – Table S5).

**Fig 3.**
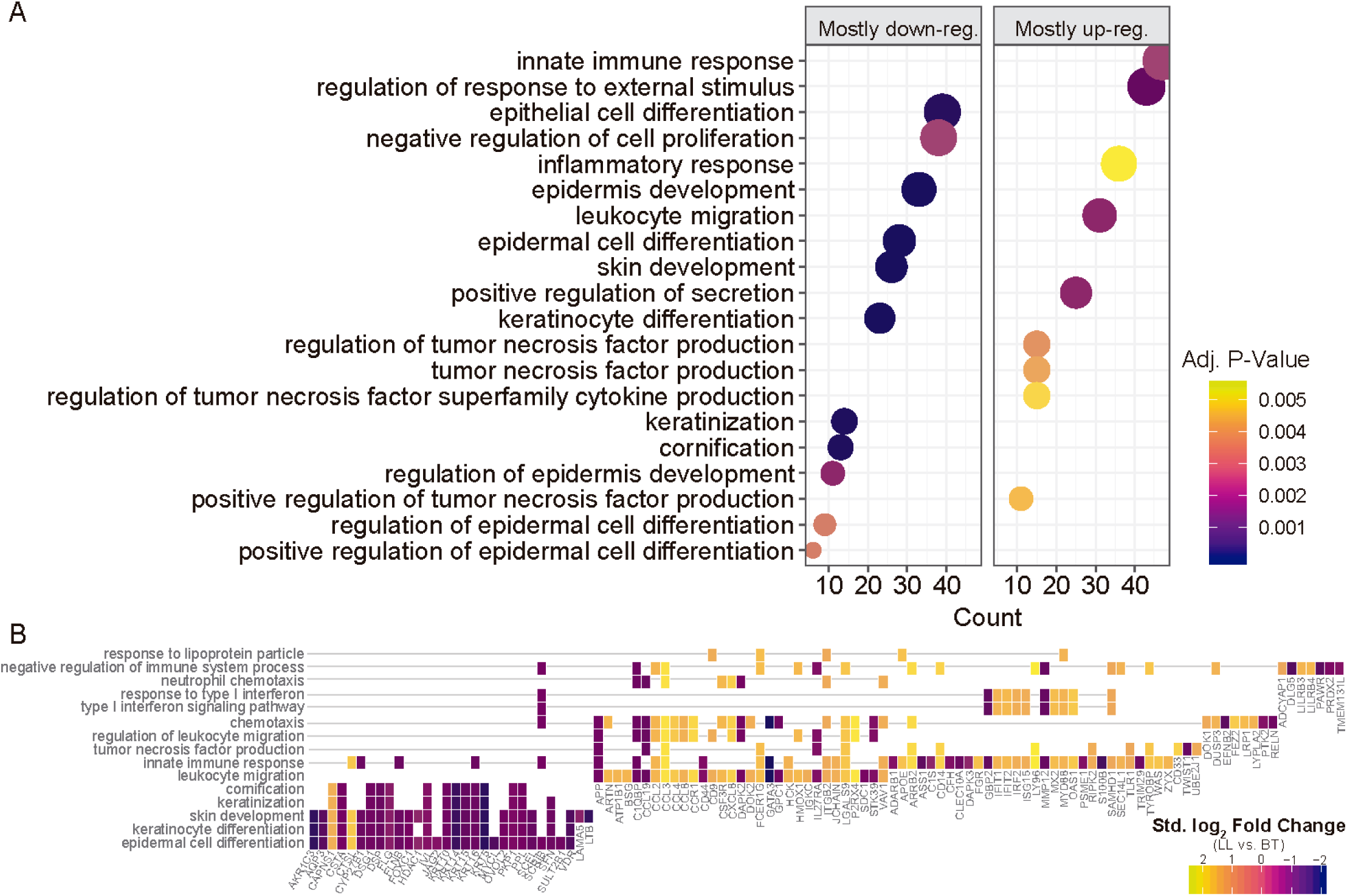
Enriched Gene Ontology biological processes from the LL vs. BT genes selected from random effects model (REM). **a** Top 20 ontologies with adj. P-value < 0.1 enriched from DEG selected by the REM method. X-axis shows the number of genes contained in the ontology and the dot size is proportional to this number. A score was calculated to show if genes within a given ontology were mostly up- or downregulated and are shown separately. **b** Heat plot showing some of the enriched biological processes with their gene members along with the effect size (standardized log_2_ fold change from REM according to Hedges & Olkin estimator).

For the LL vs. ENL comparison, the genes selected by the *REM* method with adjusted P-value (FDR) ≤ 0.01 and standardized |log_2_FC| ≥ 1 were used, which resulted in 1857 genes for enrichment analysis. Nine biological processes were enriched with an FDR < 0.1 and are shown in Figure z4a. Several ontologies were composed of up- and down-regulated (scores around 0.0) genes in LL vs. ENL, especially neutrophil-related processes and ‘iron ion homeostasis’ (Figure 4a; Additional File 3 – Table S6). Interestingly, the ontology ‘phagosome maturation’ contained more genes with higher expression in LL when compared to ENL skin lesions (Figure 4a; Additional File 3 – Table S6).

**Fig 4.**
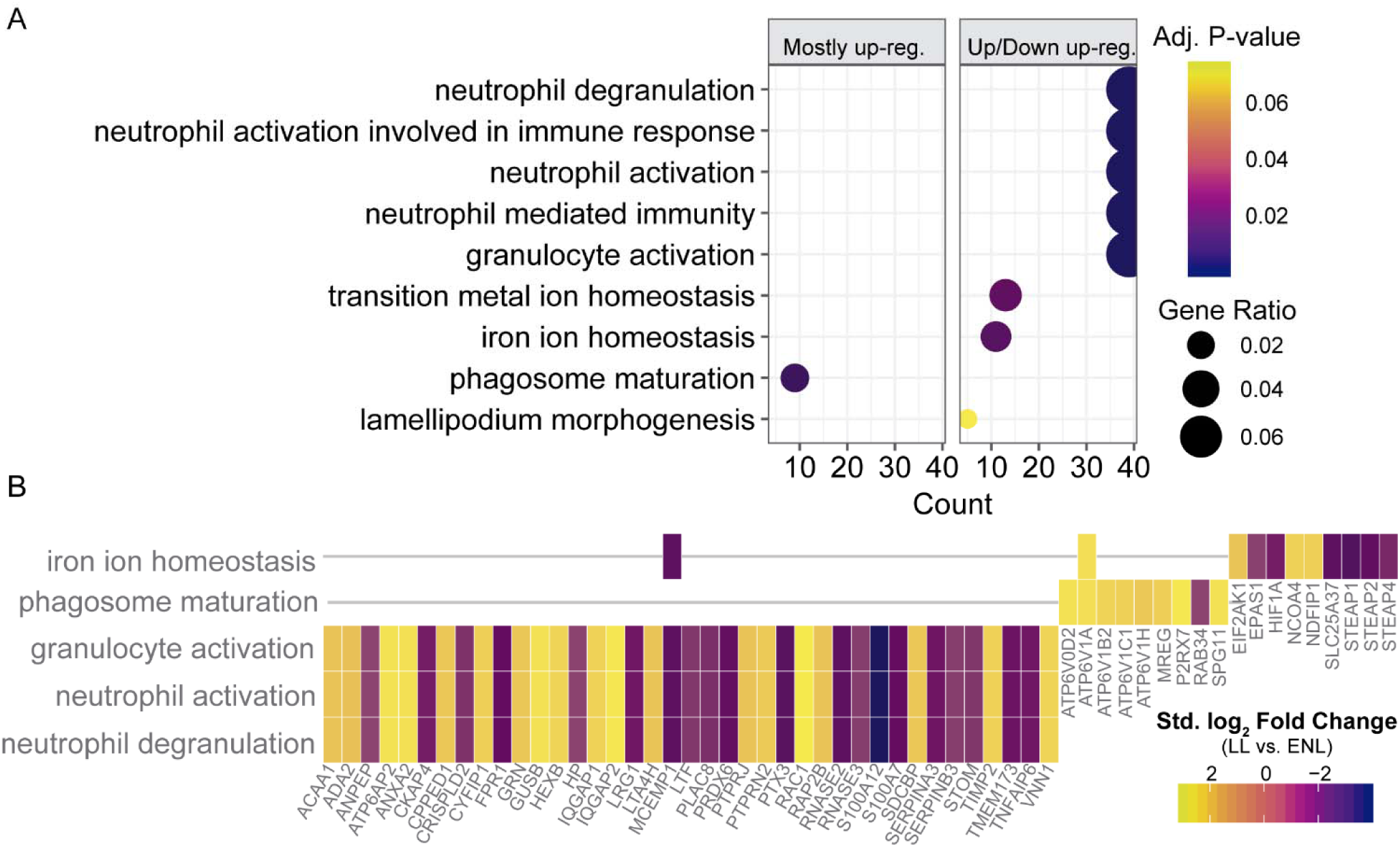
Enriched Gene Ontology biological processes from the LL vs. ENL genes selected from random effects model (REM). **a** Top 9 ontologies with adj. P-value < 0.1 enriched from DEG selected by the REM method. X-axis shows the number of genes contained in the ontology and the dot size is proportional to this number. A score was calculated to show if genes within a given ontology were mostly up- or downregulated and are shown separately. **b** Heat plot showing some of the enriched biological processes with their gene members along with the effect size (standardized log_2_ fold change from REM according to Hedges & Olkin unbiased estimator).

## DISCUSSION

Using publicly available data deposited in GEO, nine microarray independent datasets linked to leprosy were selected and reanalyzed. Individual reanalysis revealed several statistically significant DEGs. In general, the reanalyzed results agreed with those from the original authors, although it was not our aim to individually compare them. Individual results were then integrated using aggregation and meta-analysis statistical tools to find common patterns of expression across different studies on similar biological groups. The integration provided a comprehensive view of the differential expression profile of several genes involved in processes ranging from the innate and adaptive immune system to structural components and metabolic pathways. In addition, future datasets, whether from microarrays or RNA sequencing can also add information regarding the revealed genes and pathways and could even be aggregated to create gene expression signatures representative of leprosy [31].

Defining if a gene is differentially expressed in microarray or RNA-seq studies usually revolves around the adjusted P-value and an cutoff for the fold-change (effect size) [32,33]. Still, null hypothesis testing strongly depends on sample size, and underpowered experiments are expected to have few or no DEGs, depending on the investigated phenomenon [34,35]. For this reason, using multiple independent datasets confers an advantage in classifying genes as differentially expressed.

Nevertheless, only assessing the final published list of DEGs from an individual study further limits common discoveries, as several factors must be accounted for before comparing the final lists. This practice is considered biased because: 1. not all genes screened in one study are also assayed in another one, especially if they are from different microarray platforms and manufacturers; 2. the statistical framework used, as well as the P-value correction method and thresholds, dictates the size of the final list of genes; 3. preprocessing and normalization methods have substantial impact on the resulting list of genes; 4. some annotation identifiers are more biased than others and usually are redundant, especially gene symbols; 5. some datasets do not have their findings reported in peer-reviewed journals [36–39]. For this reason, reanalyzing each study from the raw data using consistent procedures and common statistical settings, accounting for the caveats described, is a good workaround to produces more comparable lists.

In this work, five aggregation/meta-analysis tools were used to screen and select common DEGs, one based on P-values and the other on standardized effect sizes [40]. Through this aggregating approach, several DEGs that were poorly or not previously described in leprosy were discovered. Although the precise role of these genes is still unknown, their importance can be inferred based on gene ontology analysis and previously elucidated pathways deemed relevant in the host response to leprosy. GO enrichment of the lists of consistent (same effect size sign) DEGs from the LL vs. BT context revealed several genes involved with epithelial cell differentiation, skin development, keratinocyte differentiation, and cornification that were down-regulated in multibacillary (LL) compared to paucibacillary (BT) leprosy. Moreover, a similar profile is seen when comparing LL lesions with healthy skin (Additional Files 1 and 2, TableS3, Figure S1), with several genes involved in epidermis development and keratinocyte differentiation being less expressed in lepromatous lesions. Whilst the genes primarily up-regulated in LL vs. BT or LL vs. Control are involved with endocytosis [41], phagocytosis, lysosomes, lipid transport and metabolism [42–44], secretory granules, immune response, iron transport and ferric iron binding [45], neutrophil activation and degranulation [46,47], for which some work has been conducted by other leprosy researchers. Together, these data suggest that in the skin, multiple cell types are involved with the disease, especially neutrophils and perhaps the, as yet, poorly studied keratinocytes [48,49].

Several cornification and skin development genes were extensively down-regulated in multibacillary leprosy (LL). Given that this effect is repeatedly observed in independent samples analyzed by distinct microarray technologies, it is unlikely that this phenomenon was not accurate. Two hypotheses can be formulated to explain this substantial down-regulation of keratinocyte-associated genes. First, this may be the result of inflammatory processes in the skin leading to epidermis thinning [50,51], resulting in lower numbers of keratinocytes being sampled. Second, *M. leprae* may directly or indirectly slow or arrest keratinocyte terminal differentiation during cornification. This could be as either a consequence from immunological activity surrounding keratinocytes or these cells being directly infected and reprogrammed [48,49]. Masaki et al. have already shown a similar phenomenon in which *M. leprae* is capable of reprogramming Schwann cells to de-differentiate into stem-like states in order to survive and disseminate [52–55]. In any case, further experiments are needed to understand this process more precisely.

Interestingly, reanalysis of three studies based on *in vitro* experiments demonstrated common activated genes, although they were generated using different stimuli on distinct cells. Furthermore, enrichment analysis of these genes resulted in several converging ontologies when comparing the results obtained from datasets on human skin biopsies. This approach has strengthened the importance of some previous known pathways, such as: type 1 and 2 interferon pathways [15,56], tumor necrosis factor [57], and NF-kappa B signaling [58,59].

Although gene expression can be validated by other techniques, such as qPCR, the analysis of multiple microarrays from independent samples, performed by different researchers in different laboratories, producing analogous results are a good indicator of true differential gene expression, especially in cases where more than three datasets were examined. However, proper replication of our findings in independent samples and with a different method may still be important to truly estimate gene expression differences [34,37,60].

## Conclusions

In this study nine public microarray datasets regarding leprosy were reanalyzed using a standardized approach, integrating their individual results to uncover differential gene expression signals. Aggregation of the results revealed several genes involved in leprosy pathogenesis that are already being investigated, as well as many novel candidates that may be pivotal in a comprehensive understanding of the host response to the disease. Genes that were more likely to be differentially expressed based on direction of the modulation and statistical significance were selected and combined from multiple independent datasets. Categories were formed by grouping samples from different studies to enable comparisons of differentially expressed genes across the disease spectrum as well as with healthy controls and for *in vitro* studies. Although some comparisons are underrepresented by the number of studies, the reanalyzed results pinpoint some pathways and genes that could be further characterized, for potential diagnostic purposes or for the general understanding of disease immunopathogenesis. In the future, when more datasets become available, we can expand this work and also create context-specific gene signatures for leprosy.

## Methods

### Search, selection and data retrieval

Initially, a manual search of NCBI’s Gene Expression Omnibus (GEO) was conducted with the following string: *“Mycobacterium leprae”[All Fields] OR “M. leprae”[All Fields] OR “leprae” [All Fields] OR (“leprosy”[MeSH Terms] OR leprosy[All Fields]) AND “gse”[Filter].* Search results were examined, and datasets were excluded if they consisted of: a) gene expression only from the pathogen; b) compound co-stimulation; c) only one kind of sample which did not allow any comparison; and, d) probes targeting only non-coding RNAs. No studies were discarded due to publication or experiment date, microarray platform or unavailability of raw data since this would decrease even further the number of datasets available. Whenever possible, raw data were preferentially downloaded; otherwise, the GEO gene expression matrix was used. A check was performed to determine whether samples were duplicated between studies by accessing contents of the available papers and individual/sample descriptions from GEO [23,61]. For this reason, datasets regarding the *in vivo* mouse footpad model were excluded because of possible biological sample duplication (Additional File 1 – GEO Search Results).

### Microarray data analysis

Raw data or expression matrices were imported to R environment software (version 3.4.1 running on Rstudio IDE v. 1.1.383) powered by Bioconductor (v. 1.26.1) libraries, *GEOquery* (v. 2.42.0), *Biobase* (v. 2.36.2) and *limma* (v. 3.32.6) [62–66]. If raw data was available, an *ExpressionSet* object was built with raw files plus phenotype and feature data available from GEO or the original publication. Datasets were preprocessed according to chip technology, where Affymetrix and other single-channel technologies were normalized with a Robust Multichip Average (RMA) method using the *affy* R package (v. 1.54.0) [67,68]. Two-channel data were background-corrected using the *normexp* method with a custom offset value, followed by within array normalization (Shrunk Robust Splines), and a final quantile normalization between arrays (*limma*) [69,70]. Multidimensional scaling (MDS) plots and Principal Component Analysis (PCA) were performed to assess the structure of data. Duplicated genes (mapped by Entrezid) were filtered out according to the smallest mean expression across all samples. Differential gene expression analysis was done by fitting gene-wise linear models and empirical Bayes moderated t-statistics, both implemented in *limma* [63,71]. Whenever available, covariates and batch information were included in the linear model. Resulting P-values were adjusted for multiple testing as proposed by Benjamini and Hochberg’s (BH) method [72] to control the FDR. The P-value distribution was visually inspected with histograms. Furthermore, the *biomaRt* (v. 2.32.1) R package was used to convert mouse genes to human orthologues Entrezid identifiers [73]. Finally, all ribosomal protein-coding genes were excluded from the processed expression matrices.

### Common differentially expressed genes from independent studies by the association ratio method (*sdef*)

To find an intersection of genes differentially expressed across studies, but for related comparisons, a frequentist and Bayesian association ratio analysis was employed [29]. Briefly, the chance association of genes from similar experiments is tested by a ratio measuring the relative increase of genes in common from different experiments with respect to the number expected by chance, that is, under the hypothesis of independence. Then, the statistical significance of this ratio is assessed by Monte Carlo permutation; also, a joint model of the experiments is formulated in a Bayesian framework. Four categories were created including different studies comparing similar groups; then, for each category, the ratio of observed to expected probability of genes to be in common was calculated by the frequentist and Bayesian methods, as proposed by the *sdef* (v. 1.6) R package [74]. For this analysis, all unique genes (Entrezid) with their corresponding nominal P-values were used. After obtaining the common DEGs list with the Bayesian approach, which considers an alpha threshold that maximizes the number of DEGs without the credibility interval including 1 (not rejecting the null), the nominal P-values were replaced by their FDR-adjusted counterparts calculated in *limma*. Thus, the association ratio method was used as an aggregation list tool.

### Meta-analysis for gene expression

The developers of *sdef* proposed that this method is rather conservative, aiming for fewer false positives when compared to other methods [29,74]. For this reason, in this study the *sdef* approach was compared with more standard gene expression meta-analysis tools. For this, the R *MetaDE* library was used [75], which implements 12 major meta-analysis methods for differential gene expression. Besides the maximum P-value (*maxP*), r-th ordered p-value (*rOP*), and sum of ranks (*SR*) P-value-based methods, the random effects model (*REM*) which is based on effect sizes was also tested [40,76,77]. The *REM* method requires at least two samples per group to compute the effect size, due to this, the comparison LL vs. Control was not used, as one of the datasets only contained one healthy sample (GSE24280). Furthermore, given that the *in vitro* datasets were generated with different stimuli (live or sonicated *M. leprae*) and on different cells (Schwann cells, whole-blood), it was expected that there would be different modulation direction and, therefore, the *REM* method would be too conservative by penalizing discordant genes, even though they may have been strongly expressed irrespective of their direction.

### Gene ontology functional enrichment analysis

Gene ontology over-representation analysis (ORA) was used to better understand the biological role of the resulting gene lists. For each comparison category, the DEG lists were tested for enriched functional categories according to the GO Biological Process (BP), and P-values were calculated based on the hypergeometric distribution [78]. P-values were also adjusted for multiple testing using the BH method [72] (*clusterProfiler* v. 3.4.4, [79]). For LL vs. BT and LL vs. ENL comparisons, the selected genes used were from the *REM* method with a standardized |log_2_FC| ≥ 1 and an FDR < 0.1 or 0.01. For the LL vs. Control, the genes from the *sdef* analysis with same gene expression direction in both studies were used. As for the *in vitro* comparison, the genes that resulted from the *sdef* approach with a median |log_2_FC| ≥ 0.5 (41% relative increased expression) were used. Since studies for the *in vitro* comparison were heterogeneous in experimental design, a filter for concordance modulation direction was not applied. Significant ontologies were visualized with custom functions from *clusterProfiler* R package [79]. A score was also calculated [30] to depict the proportion of up- or down-regulated genes that compose a given ontology; where positive scores mean most of the genes are up-regulated in a given ontology and vice-versa (Eq. 1).

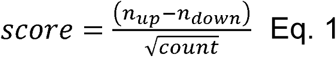

### Inspection of gene lists

The effect size (log_2_FC or standardized log_2_FC), direction of gene regulation (up- or down-regulation), and adjusted P-values are collectively important metrics to rank critical genes for further investigation. Therefore, all genes that were found to be associated among independent studies were tabulated with their FDR-adjusted P-values, standardized log_2_FC (from *REM*) or median log_2_FC (from *sdef*), and number of related comparisons in which the gene was significantly modulated (FDR ≤ 0.1). These results are shown partially in the main text and fully within the supplemental files.

## List of abbreviations

*sdef*: ratio association method
*REM*: random effects model
*rOP*: r-th ordered p-value
*maxP*: maximum P-value
*SR*: sum of ranks
MB: multibacillary leprosy
PB: paucibacillary leprosy
DEG: differentially expressed gene
FDR: false discovery rate
BH: Benjamini-Hochberg
LL: lepromatous lepromatous leprosy
ENL: erythema nodosum leprosum
BT: borderline-tuberculoid
FC: fold change
GEO: Gene Expression Omnibus
NCBI: National Center for Biotechnology Information

## Additional Files

**Additional File 1 – Supplementary tables S1-S4 and GEO search results.** Full table of results for the integration/meta-analysis with FDR ≤ 0.1. This file contains four .XLS spreadsheets with results for the LL vs. Control and Stimulated vs. Control (*in vitro*) categories not presented within main text. One .PDF containing the 18 results from the *GEOquery*. (.ZIP 2.14 MB)

**Additional File 2 – Supplementary figures S1-S2 + REM forest plots.** Enrichment analysis for the LL vs. Control and Stimulated vs. Control (*in vitro*) categories and forest plots for DEGs from LL vs. BT and LL vs. ENL random effects model (*REM*) estimates. (four .PDF files, .ZIP 2.99 MB)

**Additional File 3 – Supplementary tables S5-S8.** Four .XLS spreadsheets containing full enrichment results for all categories analyzed including the numeric score and its categorical label. (.ZIP 661 KB)

## Declarations

### Ethics approval and consent to participate

Please refer to original reports for more information on ethical protocols adopted.

### Consent for publication

Not applicable

### Availability of data and material

All datasets analyzed during the current study are available under NCBI’s Gene Expression Omnibus repository accessions: GSE40950 [28], GSE35423 [14], GSE24280 [unknown], GSE443 [80], GSE17763 [81], GSE16844 [47], GSE74481 [82], GSE95748[52], and GSE100853 [83].

All R source code used in the analyzes are readily available at GitHub (https://github.com/thyagoleal/leprosy_reanalysis_paper).

### Authors’ contributions

TLC analyzed the study, interpreted data and drafted the manuscript.

MOM conceptualized the study, interpreted results and reviewed the manuscript.

