## Supplementary tables S1-S4 and GEO search results for "Reanalysis And Integration Of Public Microarray Datasets Reveals Novel Host Genes Modulated In Leprosy": geo_search_results.pdf (from another version of this preprint: version 2, 2019-12-07)

1. Differential Expression of MicroRNAs in Leprosy Skin Lesions
(Submitter supplied) In this study, a comprehensive assessment of human miRNA was performed on leprosy skin lesions using DNA chip microarrays, which included the entire spectrum of the disease along with its reactional states. Sixty-eight samples from leprotic lesions (10TT, 10BT, 10BB, 10BL, 4LL, 14R1, and 10R2) and nine skin biopsies from healthy individuals were used as controls (CC) (ages ranged from 06 to 83 years, 48 were male and 29 female). more...
Organism: Homo sapiens
Type: Expression profiling by array
Platform: GPL15018 77 Samples
FTP download: GEO (TXT) ftp://ftp.ncbi.nlm.nih.gov/geo/series/GSE102nnn/GSE102314/
Series Accession: GSE102314 ID: 200102314

2. Deciphering the genetic control of gene expression following Mycobacterium leprae antigens stimulation
(Submitter supplied) Leprosy is a human infectious disease caused by Mycobacterium leprae. A strong host genetic contribution to leprosy susceptibility is well established. However, the modulation of the transcriptional response to infection and the mechanism(s) of disease control are poorly understood. To address this gap in knowledge of leprosy pathogenicity, we conducted a genome-wide search for expression quantitative trait loci (eQTL) that are associated with transcript variation â€“â€“ before and after stimulation with M. more...
Organism: Homo sapiens
Type: Expression profiling by array
Platform: GPL15988 102 Samples
FTP download: GEO (TXT) ftp://ftp.ncbi.nlm.nih.gov/geo/series/GSE100nnn/GSE100853/
Series Accession: GSE100853 ID: 200100853

3. Expression data from control and Mycobacterium leprae-infected mouse Schwann cells
(Submitter supplied) Mycobacterium leprae infect mouse Schwann cells in vitro, reprogram the Schwann cells, and induce mesenchymal stem cell-like cells. In this reprogramming, a vast number of stem cell-related genes, especially those with transcription factor activities are differentially regulated We used microarrays to detail the global program of gene expression underlying the Mycobacterial infection and identified distinct classes of differentially-regulated genes during this process.
Organism: Mus musculus
Type: Expression profiling by array
Platform: GPL1261 8 Samples
FTP download: GEO (CEL) ftp://ftp.ncbi.nlm.nih.gov/geo/series/GSE95nnn/GSE95748/
Series Accession: GSE95748 ID: 200095748

4. Integration of cell type deconvolution with immune pathways identifies gene networks of host defense and immunopathology in leprosy
(Submitter supplied) Transcriptome profiles derived from the site of human disease has led to the identification of genes that contribute to pathogenesis, yet the complex mixture of cell types in these lesions has been an obstacle for defining specific mechanisms. Leprosy provides an outstanding model to study host defense and pathogenesis in a human infectious disease, given its clinical spectrum which interrelates with the host immunologic and pathologic responses. more...
Organism: Homo sapiens
Type: Expression profiling by array
Platform: GPL570 6 Samples
FTP download: GEO (CEL) ftp://ftp.ncbi.nlm.nih.gov/geo/series/GSE84nnn/GSE84893/
Series Accession: GSE84893 ID: 200084893

5. Analysis of RNA Based Transcriptome of Mycobacterium leprae genome from Environmental and Clinical Samples
(Submitter supplied) BACKGROUND: This study was conducted to analyse M. leprae gene expression profiles in biopsy of leprosy patient and in environmental soil samples. Investigation was carried out to find out the expression profile for the minimal gene set necessary for growth of environmental M. leprae and its genes potentially require for infection and pathogenesis of leprosy. METHOD: RNA was extracted from human biopsy and environmental soil samples and total RNA obtained by subtractive hybridization of rRNA, was reverse

transcribed and amplified using random primers. more...
Organism: Mycobacterium leprae
Type: Expression profiling by genome tiling array
Platform: GPL20100 4 Samples
FTP download: GEO (TXT) ftp://ftp.ncbi.nlm.nih.gov/geo/series/GSE68nnn/GSE68192/
Series Accession: GSE68192 ID: 200068192

6. Genome-wide screening of mRNA expression in leprosy patients
(Submitter supplied) In this study, a comprehensive assessment of human mRNA was performed on leprosy skin lesions using DNA chip microarrays, which included the entire spectrum of the disease along with its reactional states. Sixty-six samples from leprotic lesions (10TT, 10BT, 10BB, 10BL, 4LL, 14R1, and 10R2) and nine skin biopsies from healthy individuals were used as controls (CC) (ages ranged from 06 to 83 years, 48 were male and 29 female). more...
Organism: Homo sapiens
Type: Expression profiling by array
Platform: GPL14550 76 Samples
FTP download: GEO (TXT) ftp://ftp.ncbi.nlm.nih.gov/geo/series/GSE74nnn/GSE74481/
Series Accession: GSE74481 ID: 200074481

7. Microarray analysis of IL-10 stimulated adherent peripheral blood mononuclear cells
(Submitter supplied) The immune mechanisms that control resistance vs. susceptibility to mycobacterial infection in humans were investigated by studying leprosy skin lesions, the site where the battle between the host and the pathogen is joined. Using an integrative genomics approach, we found an inverse correlation between of IFN-beta and IFN-gamma gene expression programs at the site of disease. The Type II IFN, IFN-gamma and its downstream vitamin D-dependent antimicrobial genes were preferentially expressed in the lesions from patients with the self-healing tuberculoid form of the disease and mediated antimicrobial activity against the pathogen, Mycobacterium leprae in vitro. more...
Organism: Homo sapiens
Type: Expression profiling by array
Dataset: GDS4551 Platform: GPL570 8 Samples
FTP download: GEO (CEL) ftp://ftp.ncbi.nlm.nih.gov/geo/series/GSE43nnn/GSE43700/
Series Accession: GSE43700 ID: 200043700

8. Effect of live Mycobacterium leprae infection on global gene expression of primary human Schwann cells at 24 hours
(Submitter supplied) In an attempt to understand M. leprae interaction with the human host, Stanford Genomics HEEBO Arrays were use to identify modulated genes in primary human Schwann cells infected with live M. leprae at a MOI of 100:1 for 24 hours.
Organism: Homo sapiens
Type: Expression profiling by array
Platform: GPL16068 6 Samples
FTP download: GEO (GPR) ftp://ftp.ncbi.nlm.nih.gov/geo/series/GSE40nnn/GSE40950/
Series Accession: GSE40950 ID: 200040950

9. Effect of live Mycobacterium leprae on primary human Schwann cell gene expression
(Submitter supplied) In attempt to understand M.leprae interaction with the human host, Applied Biosystems microarrays containing 30,865 probles were used to identify modulated genes in primary human Schwann Cells (SC) infected with live M. leprae at two early time points, 24 and 48 hours.
Organism: Homo sapiens
Type: Expression profiling by array
Platform: GPL2986 8 Samples
FTP download: GEO (TXT) ftp://ftp.ncbi.nlm.nih.gov/geo/series/GSE35nnn/GSE35423/
Series Accession: GSE35423 ID: 200035423

10. miRNA expression profile in leprosy skin biopsy specimens
(Submitter supplied) Skin biopsy specimens of skin lesions were profiled for miRNA expression. In this study, we indentified miRNA species that were differentially expressed in the skin lesions of either the lepromatous or

tuberculoid forms of leprosy. One miRNA species, hsa-mir-21, found in the lepromatous lesions was capable of downregulating the vitamin D-dependent antimicrobial pathway.
Organism: Homo sapiens
Type: Non-coding RNA profiling by array
Platform: GPL14777 11 Samples
FTP download: GEO (CEL, TXT) ftp://ftp.ncbi.nlm.nih.gov/geo/series/GSE33nnn/GSE33192/
Series Accession: GSE33192 ID: 200033192

11. Transcriptome analysis in Human leprosy patients
(Submitter supplied) Leprosy is classified from paucibacillary leprosy (TT and BT) to multibacillary leprosy (BB, BL, and LL). In our study, we focus on selecting or classfying the significant genes to discriminate between paucibacillary and multibacillary group and the important genes to distinguish between blood and the corresponding tissue in patients.
Organism: Homo sapiens
Type: Expression profiling by array
Platform: GPL10191 25 Samples
FTP download: GEO (CALLS, PAIR) ftp://ftp.ncbi.nlm.nih.gov/geo/series/GSE24nnn/GSE24280/
Series Accession: GSE24280 ID: 200024280

12. Comparison of Gene expression pattern between heat-killed M.leprae and live M. leprae-infected peritoneal macrophage in vivo model
(Submitter supplied) Our goal is to discriminate specific genes in live M.leprae-infected peritoneal macrophages in comparison to heat-killed M.leprae infected peritoneal macrophages using microarray.
Organism: Mus musculus
Type: Expression profiling by array
Platform: GPL4134 16 Samples
FTP download: GEO (TXT) ftp://ftp.ncbi.nlm.nih.gov/geo/series/GSE23nnn/GSE23749/
Series Accession: GSE23749 ID: 200023749

13. Transcriptome analysis in Mycobacterium leprae (M. leprae)-infected granuloma of BALB/c nu/nu mice
(Submitter supplied) Our object is to characterize the distinguish gene enrichment group in skin of Mycobacterium leprae (M. leprae)-infected footpads compared to that of Mycobacterium leprae (M. leprae) non-infected footpads.
Organism: Mus musculus
Type: Expression profiling by array
Platform: GPL6105 6 Samples
FTP download: GEO (TXT) ftp://ftp.ncbi.nlm.nih.gov/geo/series/GSE24nnn/GSE24120/
Series Accession: GSE24120 ID: 200024120

14. Profiling of Gene expression in Mycobacterium leprae (M. leprae)-infected footpad of BALB/c nu/nu mice
(Submitter supplied) Our goal is to understand the mechanism of granuloma formation in molecular level using Mycobacterium leprae (M. leprae)-infected footpads.
Organism: Mus musculus
Type: Expression profiling by array
Platform: GPL10650 6 Samples
FTP download: GEO (TXT) ftp://ftp.ncbi.nlm.nih.gov/geo/series/GSE22nnn/GSE22784/
Series Accession: GSE22784 ID: 200022784

15. Divergence of macrophage phagocytic and antimicrobial programs in leprosy
(Submitter supplied) Effective innate immunity against many microbial pathogens requires macrophage programs that upregulate phagocytosis and direct antimicrobial pathways, two functions generally assumed to be coordinately regulated. Here the regulation of these key functions was investigated in human blood-derived macrophages. IL-10 induced the phagocytic pathway, including CD209 and scavenger receptors, resulting in phagocytosis of mycobacteria and oxLDL. more...
Organism: Homo sapiens
Type: Expression profiling by array
Platform: GPL570 24 Samples
FTP download: GEO (CEL, CHP) ftp://ftp.ncbi.nlm.nih.gov/geo/series/GSE17nnn/GSE17763/

Series Accession: GSE17763 ID: 200017763

16. Implications of high level pseudogene transcription in Mycobacterium leprae
(Submitter supplied) Approximately half of M. lepraeâ€™s transcriptome consists of inactive gene products. This has an impact on overall energy and resource consumption without potential benefit to this organism. However, multiple translational â€˜silencingâ€™ mechanisms are present, reducing additional energy and resource expenditure required for protein production from these transcripts. The Mycobacterium leprae genome has less than 50% coding capacity and 1,133 pseudogenes. more...
Organism: Mycobacterium leprae
Type: Expression profiling by array
Platform: GPL8883 4 Samples
FTP download: GEO ftp://ftp.ncbi.nlm.nih.gov/geo/series/GSE17nnn/GSE17191/
Series Accession: GSE17191 ID: 200017191

17. Integrated pathways for neutrophil recruitment and inflammation in leprosy
(Submitter supplied) Neutrophil recruitment is pivotal to host defense against microbial infection, but also contributes to the immunopathology of disease. We investigated the mechanism of neutrophil recruitment in human infectious disease by bioinformatic pathways analysis of the gene expression profiles in the skin lesions of leprosy. In erythema nodosum leprosum (ENL), which occurs in patients with lepromatous leprosy (L-lep), and is characterized by neutrophil infiltration in lesions, the most overrepresented biologic functional group was 'cell movement' including E-selectin, which was coordinately regulated with IL-1beta. more...
Organism: Homo sapiens
Type: Expression profiling by array
Platform: GPL570 13 Samples
FTP download: GEO (CEL, CHP) ftp://ftp.ncbi.nlm.nih.gov/geo/series/GSE16nnn/GSE16844/
Series Accession: GSE16844 ID: 200016844

18. Leprosy lesion gene expression
(Submitter supplied) Leprosy presents as a clinical and immunological spectrum of disease. With the use of gene expression profiling, we observed that a distinction in gene expression correlates with and accurately classifies the clinical form of the disease. Genes belonging to the leukocyte immunoglobulin-like receptor (LIR) family were significantly up-regulated in lesions of lepromatous patients suffering from the disseminated form of the infection. more...
Organism: Homo sapiens
Type: Expression profiling by array
Dataset: GDS358 Platform: GPL8300 11 Samples
FTP download: GEO ftp://ftp.ncbi.nlm.nih.gov/geo/series/GSEnnn/GSE443/
Series Accession: GSE443 ID: 200000443
