## Supplementary figures S1-S2 + REM forest plots for "Reanalysis And Integration Of Public Microarray Datasets Reveals Novel Host Genes Modulated In Leprosy": FigS2.pdf (from another version of this preprint: version 2, 2019-12-07)

Figure S2 - Enriched biological processes for the Stimulated vs. Mock *in vitro* datasets (selected by sdef)

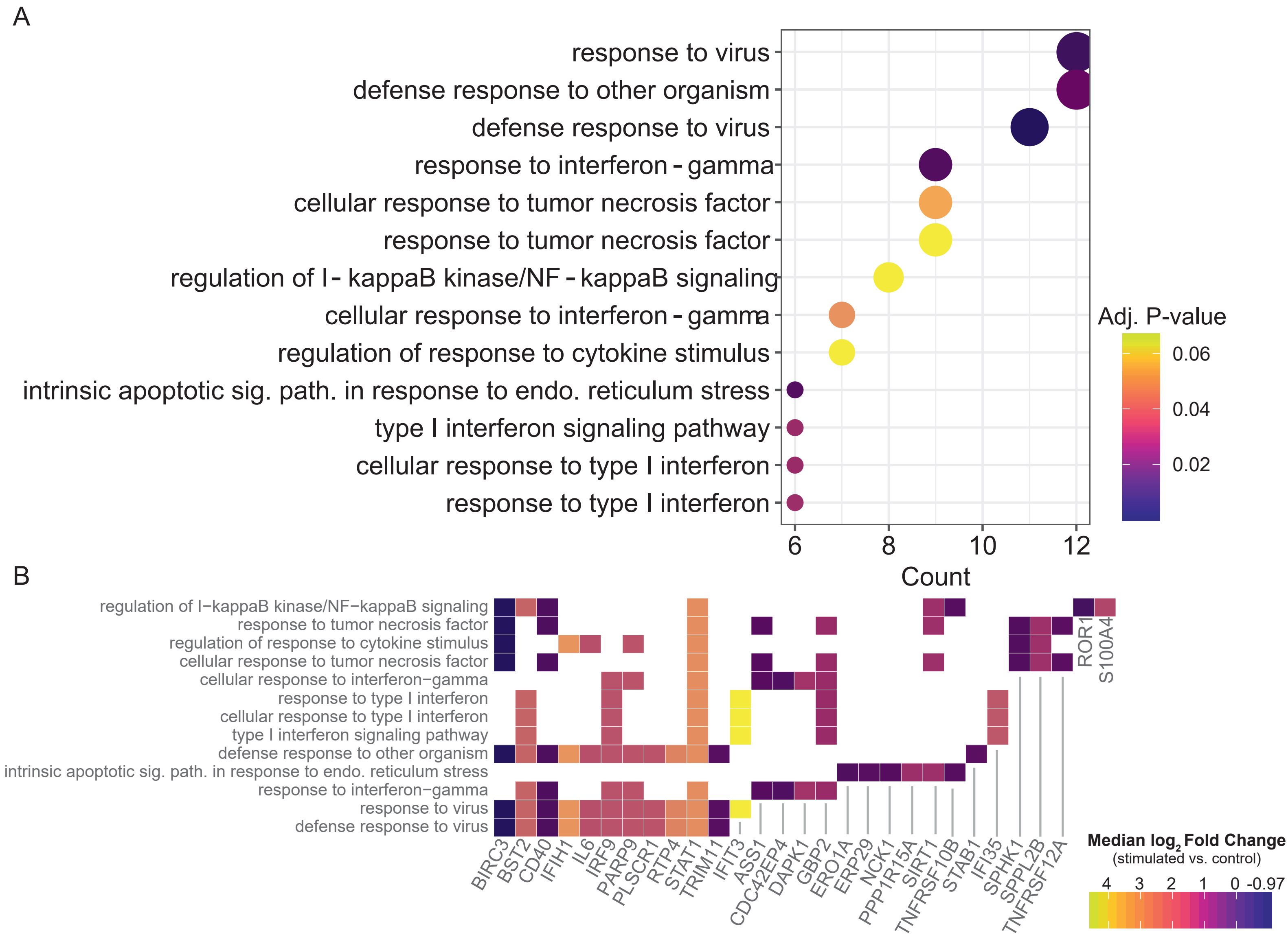
