## Supplementary figures S1-S2 + REM forest plots for "Reanalysis And Integration Of Public Microarray Datasets Reveals Novel Host Genes Modulated In Leprosy": LLvsBT_REM_forestplot.pdf (from another version of this preprint: version 2, 2019-12-07)

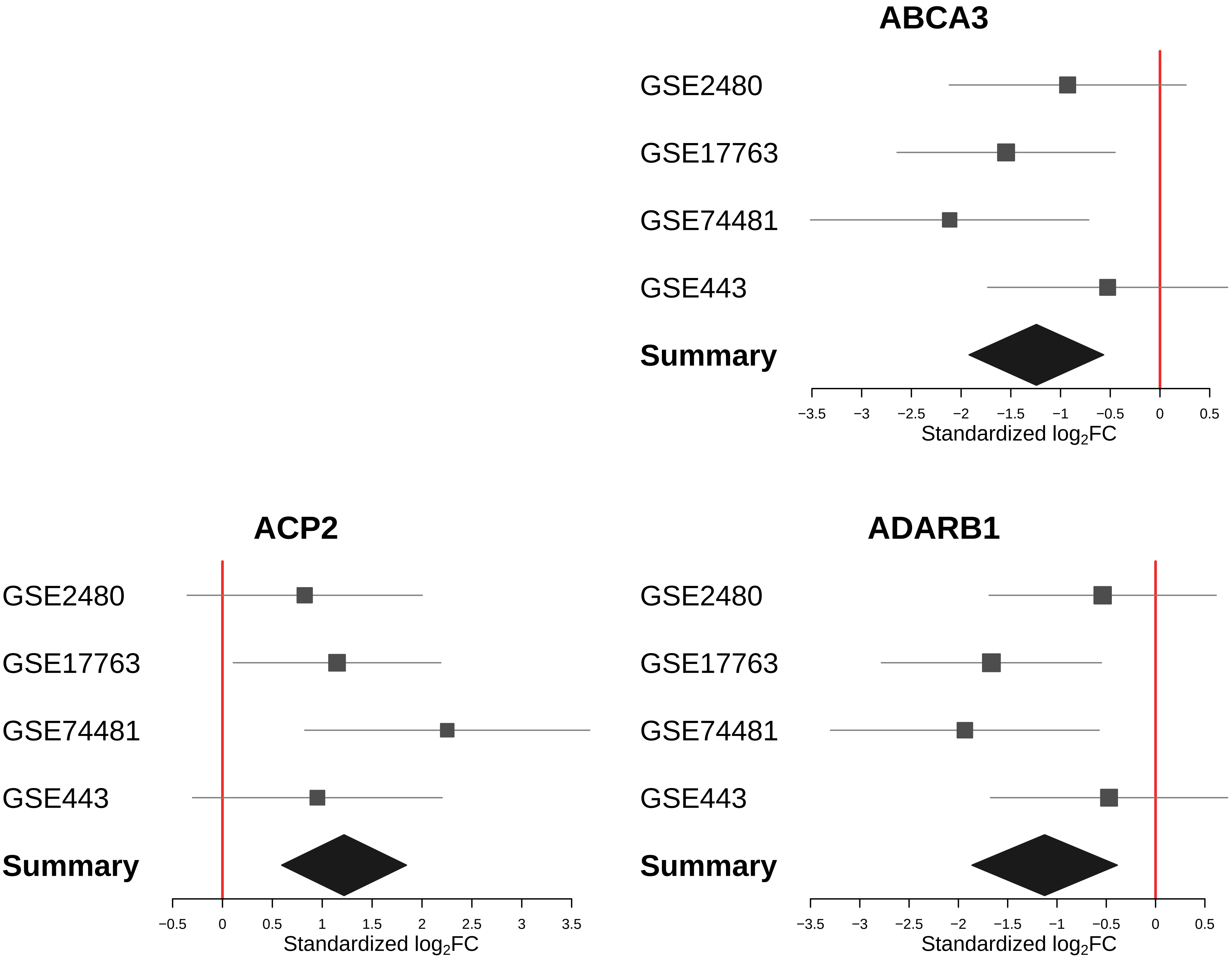

ABCA3
GSE2480
GSE17763
GSE74481
GSE443
Summary
−3.5
−3
−2.5
−2
−1.5
−1
−0.5
0
0.5
Standardized log2FC
ACP2
GSE2480
GSE17763
GSE74481
GSE443
Summary
−0.5
0
0.5
1
1.5
2
2.5
3
3.5
Standardized log2FC
ADARB1
GSE2480
GSE17763
GSE74481
GSE443
Summary
−3.5
−3
−2.5
−2
−1.5
−1
−0.5
0
0.5
Standardized log2FC

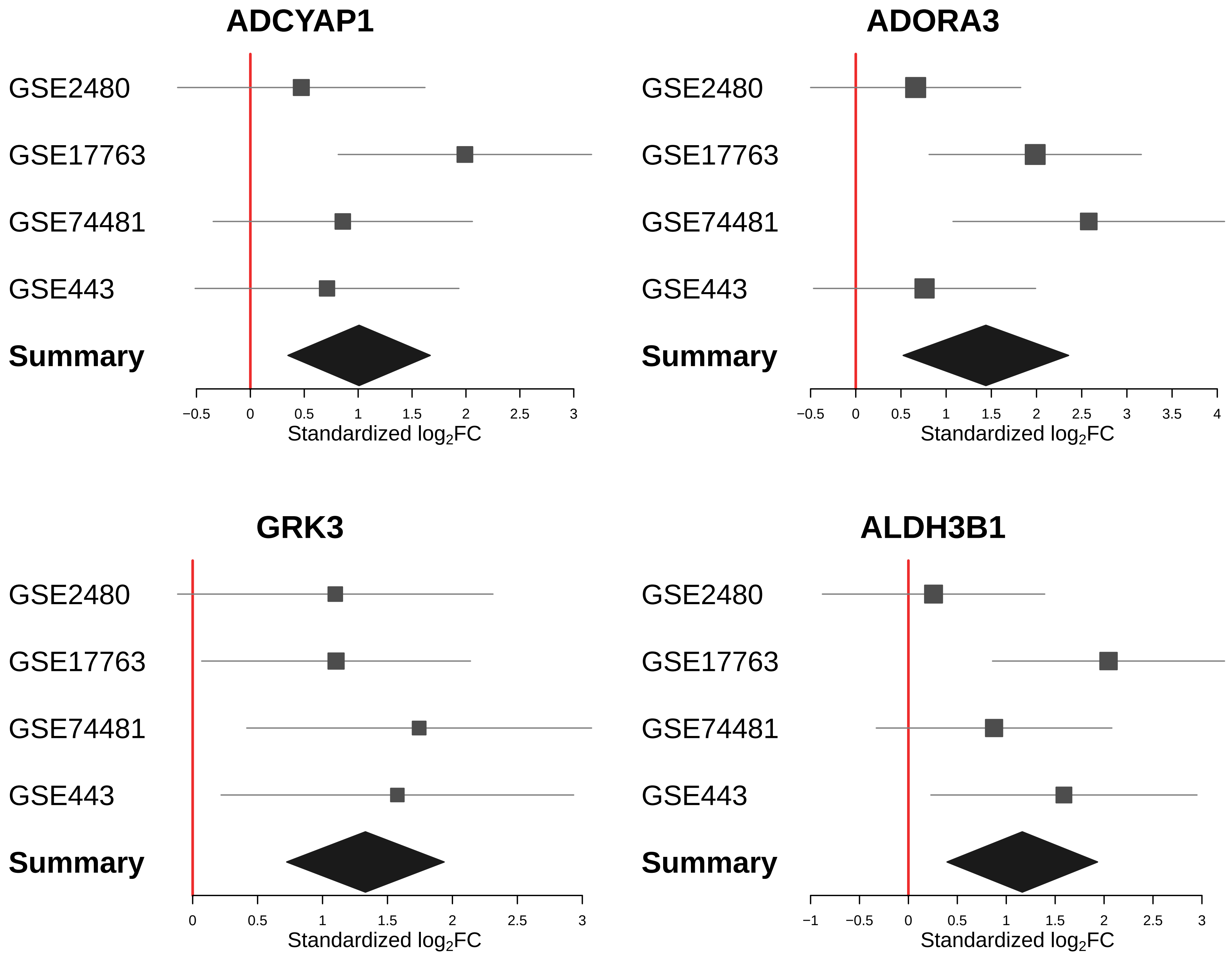
ADCYAP1
GSE2480
GSE17763
GSE74481
GSE443
Summary
−0.5 0 0.5 1 1.5 2 2.5 3
Standardized log2FC
ADORA3
GSE2480
GSE17763
GSE74481
GSE443
Summary
−0.5 0 0.5 1 1.5 2 2.5 3 3.5 4
Standardized log2FC
GRK3
GSE2480
GSE17763
GSE74481
GSE443
Summary
0 0.5 1 1.5 2 2.5 3
Standardized log2FC
ALDH3B1
GSE2480
GSE17763
GSE74481
GSE443
Summary
−1 −0.5 0 0.5 1 1.5 2 2.5 3
Standardized log2FC

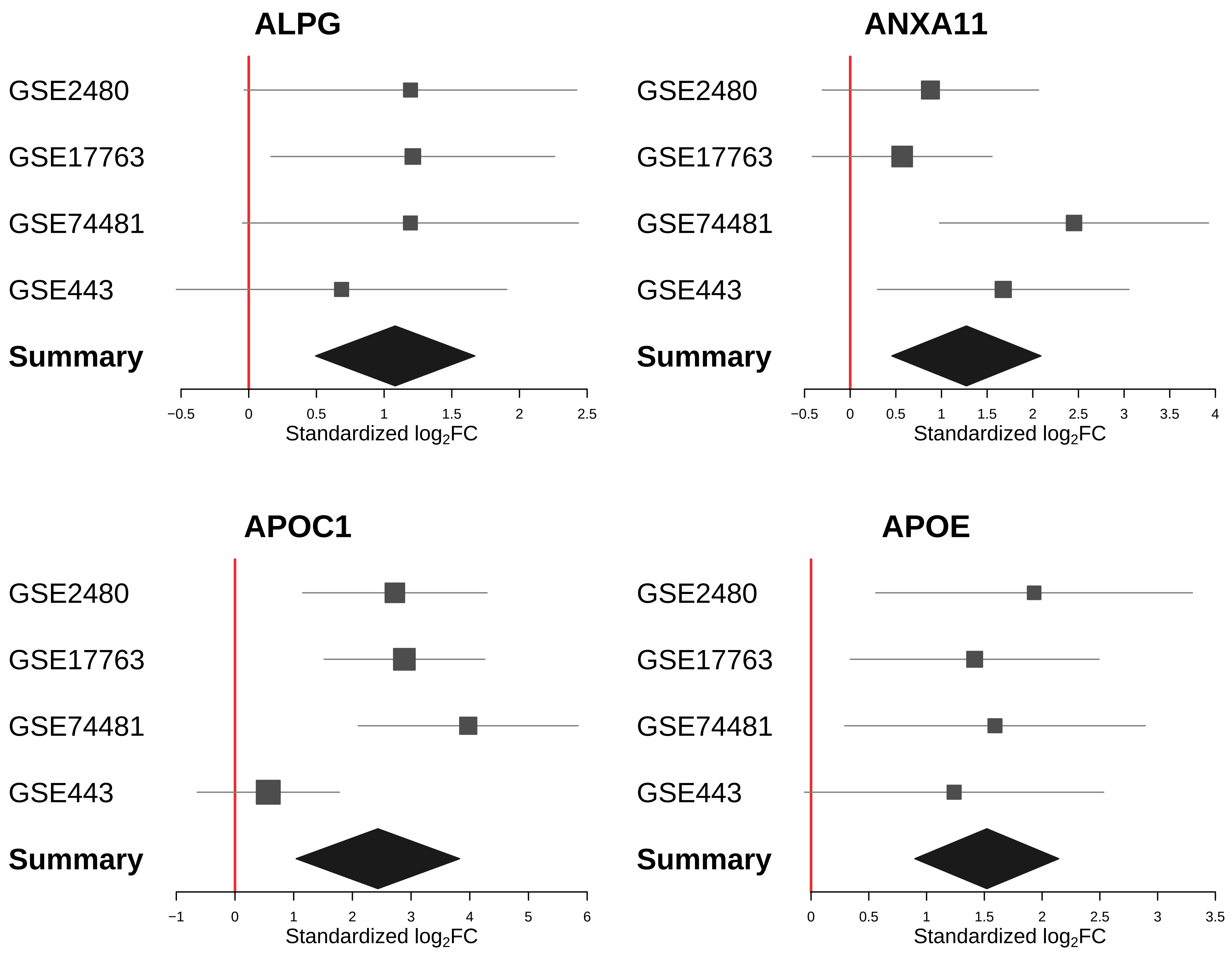

ALPG
ANXA11
APOC1
APOE
GSE2480
GSE17763
GSE74481
GSE443
Summary
Standardized log2FC
−0.5 0 0.5 1 1.5 2 2.5
−0.5 0 0.5 1 1.5 2 2.5 3 3.5 4
−1 0 1 2 3 4 5 6
0 0.5 1 1.5 2 2.5 3 3.5

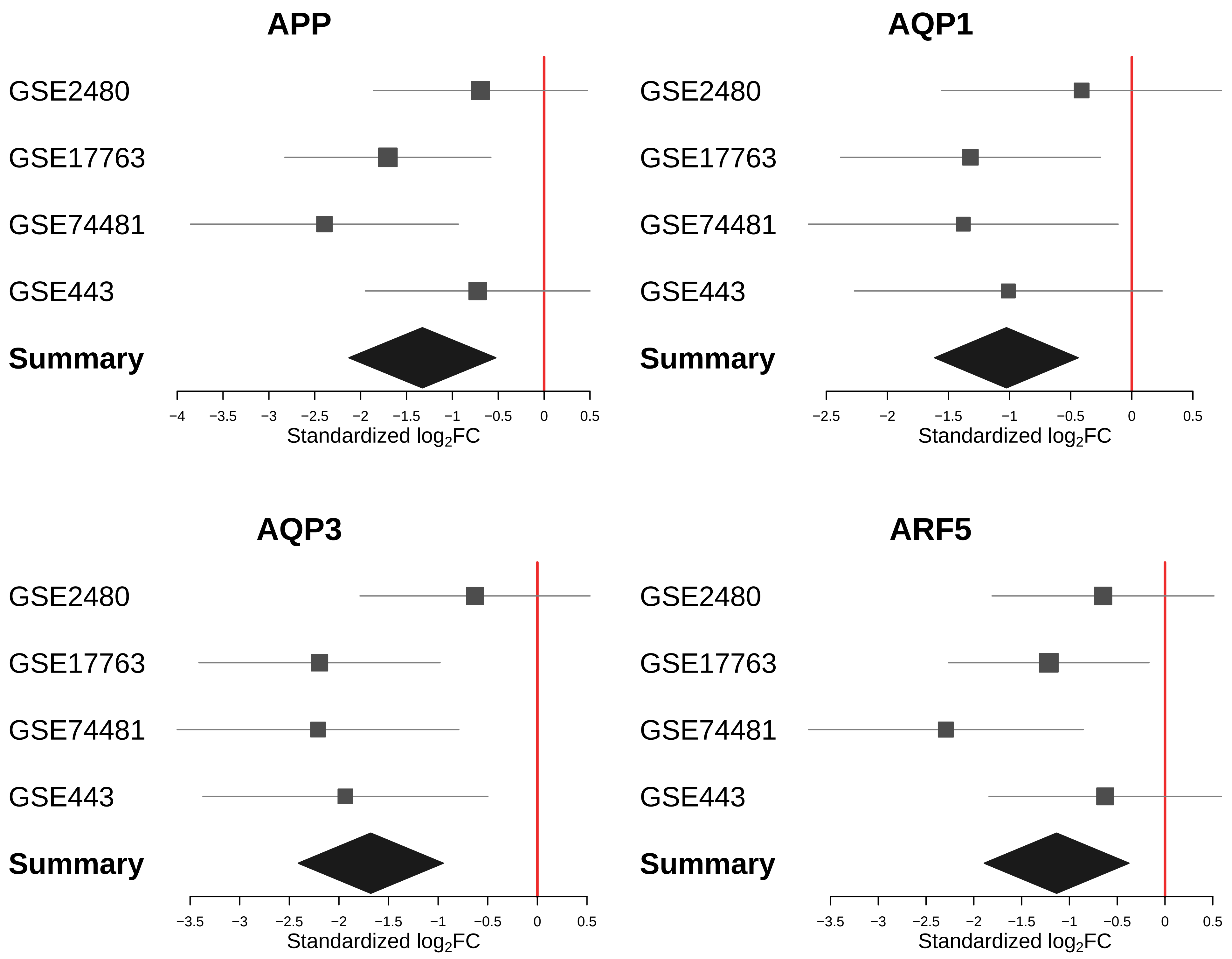

APP
AQP1
AQP3
ARF5
GSE2480
GSE17763
GSE74481
GSE443
Summary
Standardized $log_2FC$
−4
−3.5
−3
−2.5
−2
−1.5
−1
−0.5
0
0.5

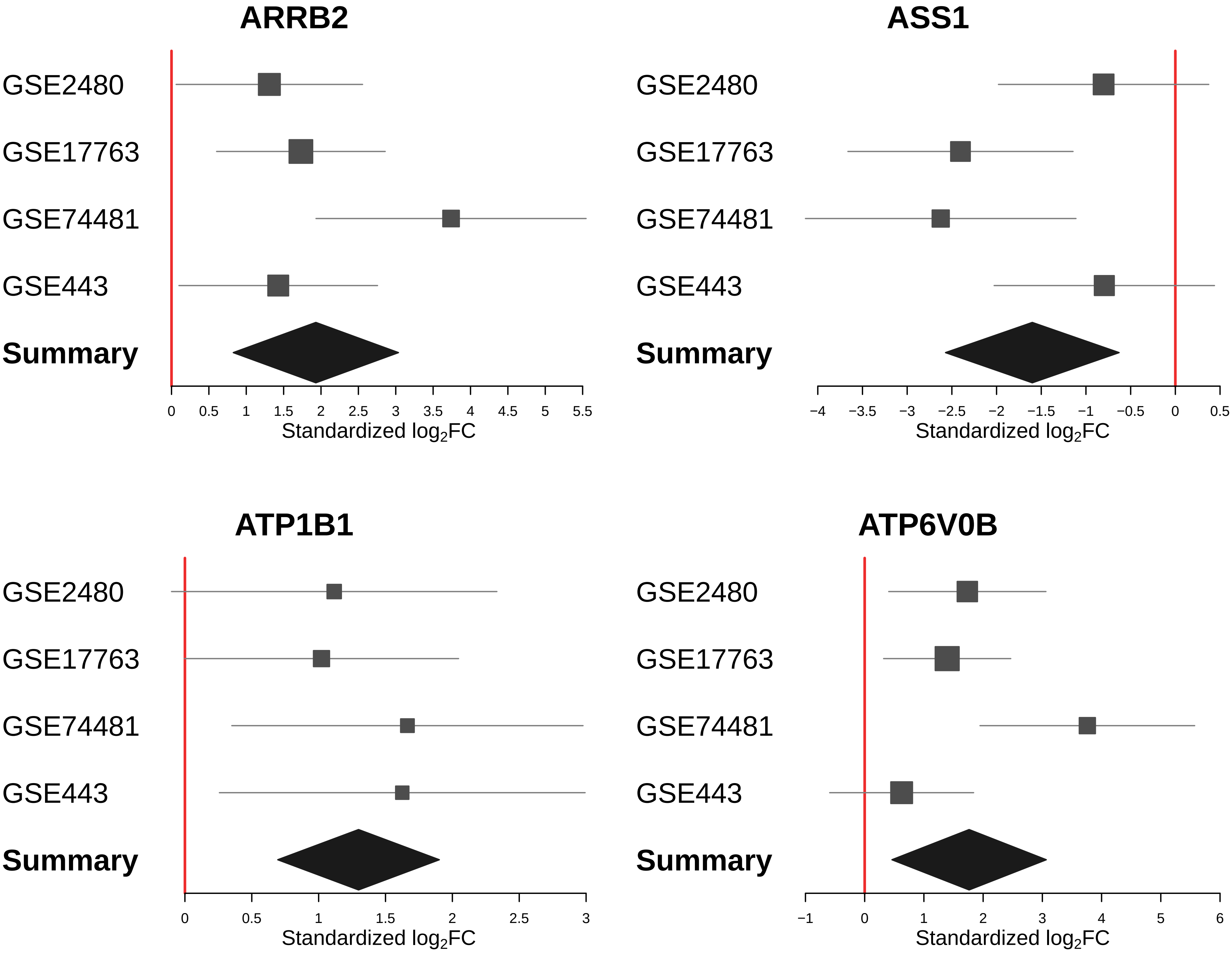
ARRB2
GSE2480
GSE17763
GSE74481
GSE443
Summary
0 0.5 1 1.5 2 2.5 3 3.5 4 4.5 5 5.5
Standardized log2FC
ASS1
GSE2480
GSE17763
GSE74481
GSE443
Summary
−4 −3.5 −3 −2.5 −2 −1.5 −1 −0.5 0 0.5
Standardized log2FC
ATP1B1
GSE2480
GSE17763
GSE74481
GSE443
Summary
0 0.5 1 1.5 2 2.5 3
Standardized log2FC
ATP6V0B
GSE2480
GSE17763
GSE74481
GSE443
Summary
−1 0 1 2 3 4 5 6
Standardized log2FC

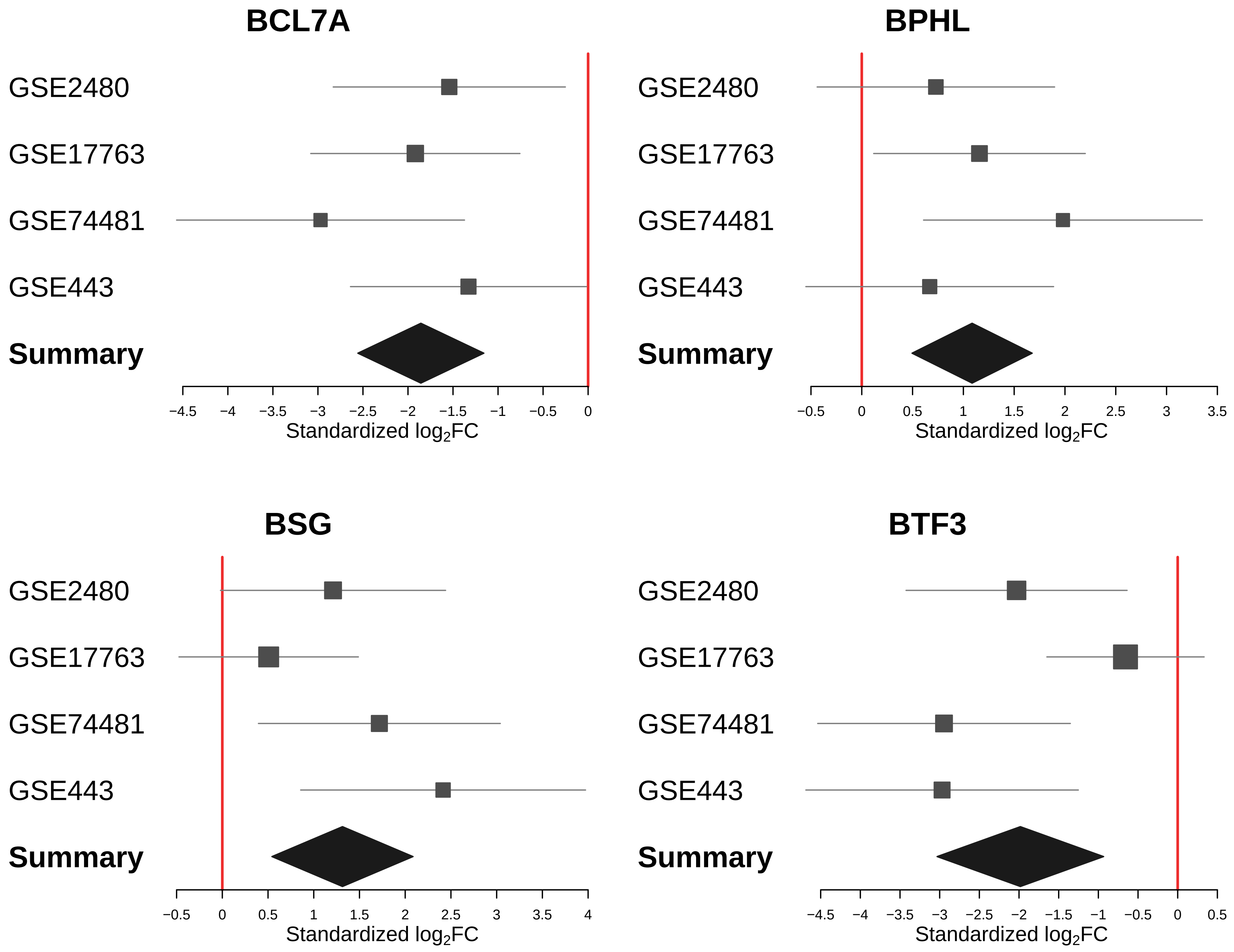
BCL7A
GSE2480
GSE17763
GSE74481
GSE443
Summary
−4.5 −4 −3.5 −3 −2.5 −2 −1.5 −1 −0.5 0
Standardized $log_2FC$
BPHL
GSE2480
GSE17763
GSE74481
GSE443
Summary
−0.5 0 0.5 1 1.5 2 2.5 3 3.5
Standardized $log_2FC$
BSG
GSE2480
GSE17763
GSE74481
GSE443
Summary
−0.5 0 0.5 1 1.5 2 2.5 3 3.5 4
Standardized $log_2FC$
BTF3
GSE2480
GSE17763
GSE74481
GSE443
Summary
−4.5 −4 −3.5 −3 −2.5 −2 −1.5 −1 −0.5 0 0.5
Standardized $log_2FC$

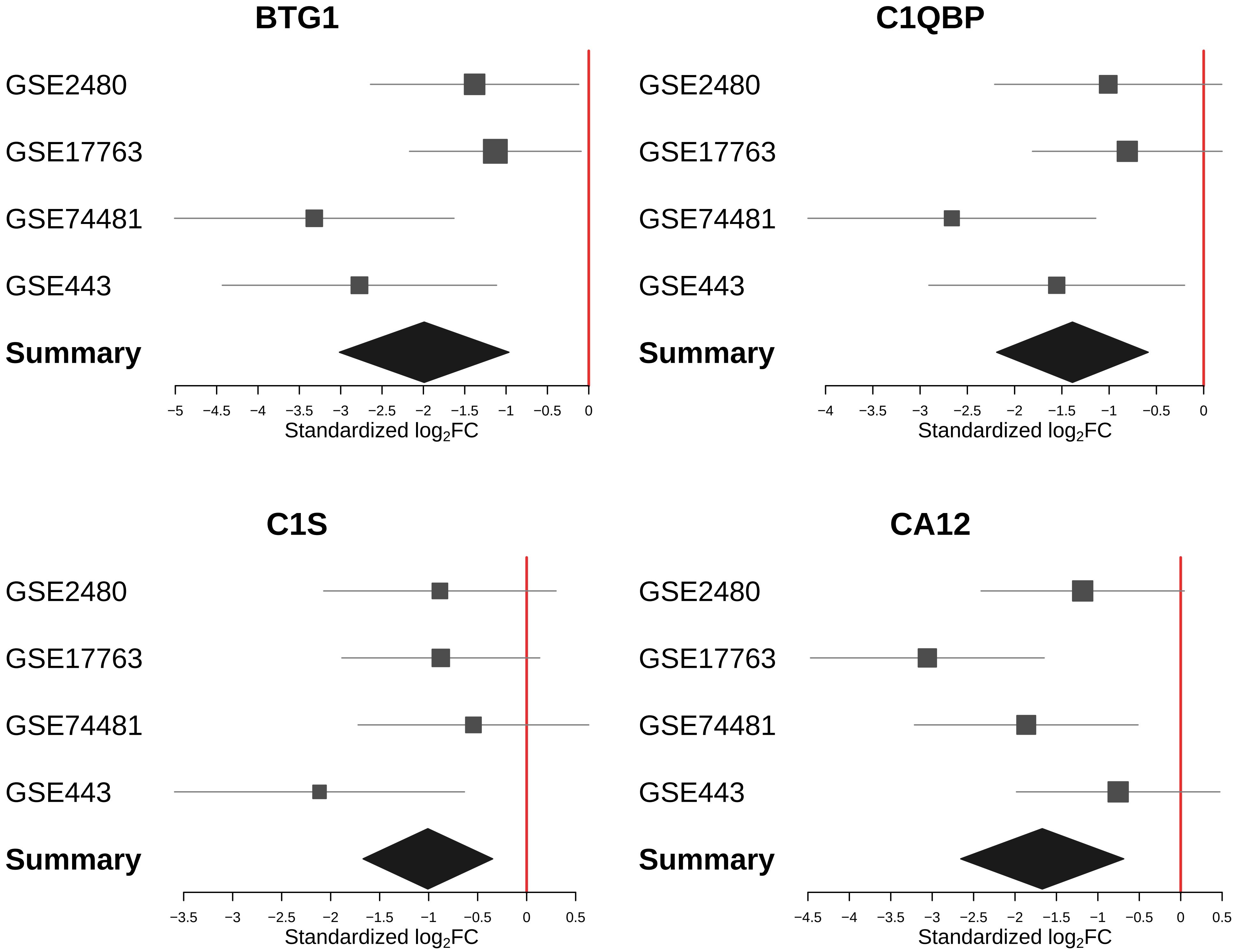
BTG1
GSE2480
GSE17763
GSE74481
GSE443
Summary
−5 −4.5 −4 −3.5 −3 −2.5 −2 −1.5 −1 −0.5 0
Standardized $log_2FC$
C1QBP
GSE2480
GSE17763
GSE74481
GSE443
Summary
−4 −3.5 −3 −2.5 −2 −1.5 −1 −0.5 0
Standardized $log_2FC$
C1S
GSE2480
GSE17763
GSE74481
GSE443
Summary
−3.5 −3 −2.5 −2 −1.5 −1 −0.5 0 0.5
Standardized $log_2FC$
CA12
GSE2480
GSE17763
GSE74481
GSE443
Summary
−4.5 −4 −3.5 −3 −2.5 −2 −1.5 −1 −0.5 0 0.5
Standardized $log_2FC$

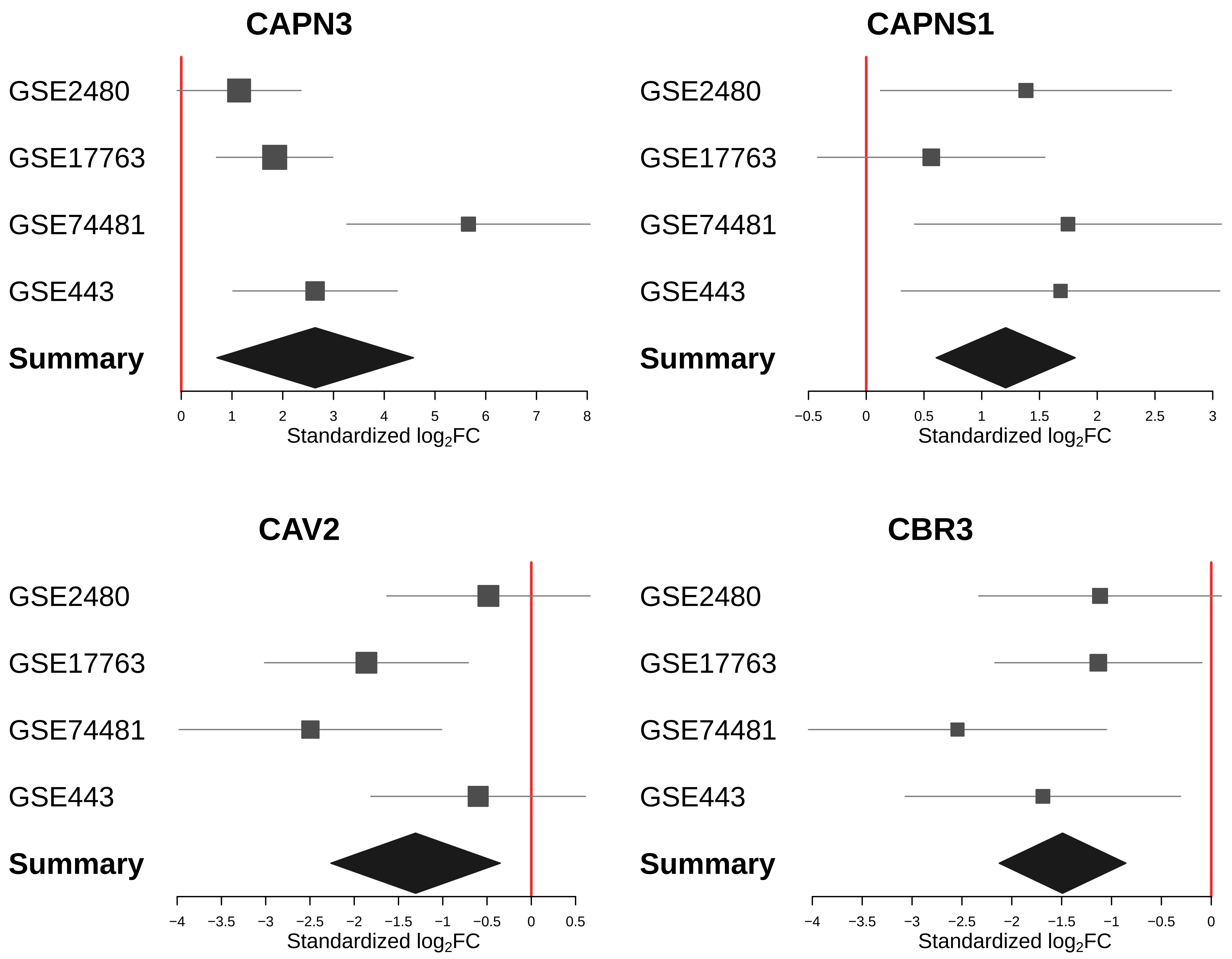

CAPN3
CAPNS1
CAV2
CBR3
GSE2480
GSE17763
GSE74481
GSE443
Summary
Standardized log2FC

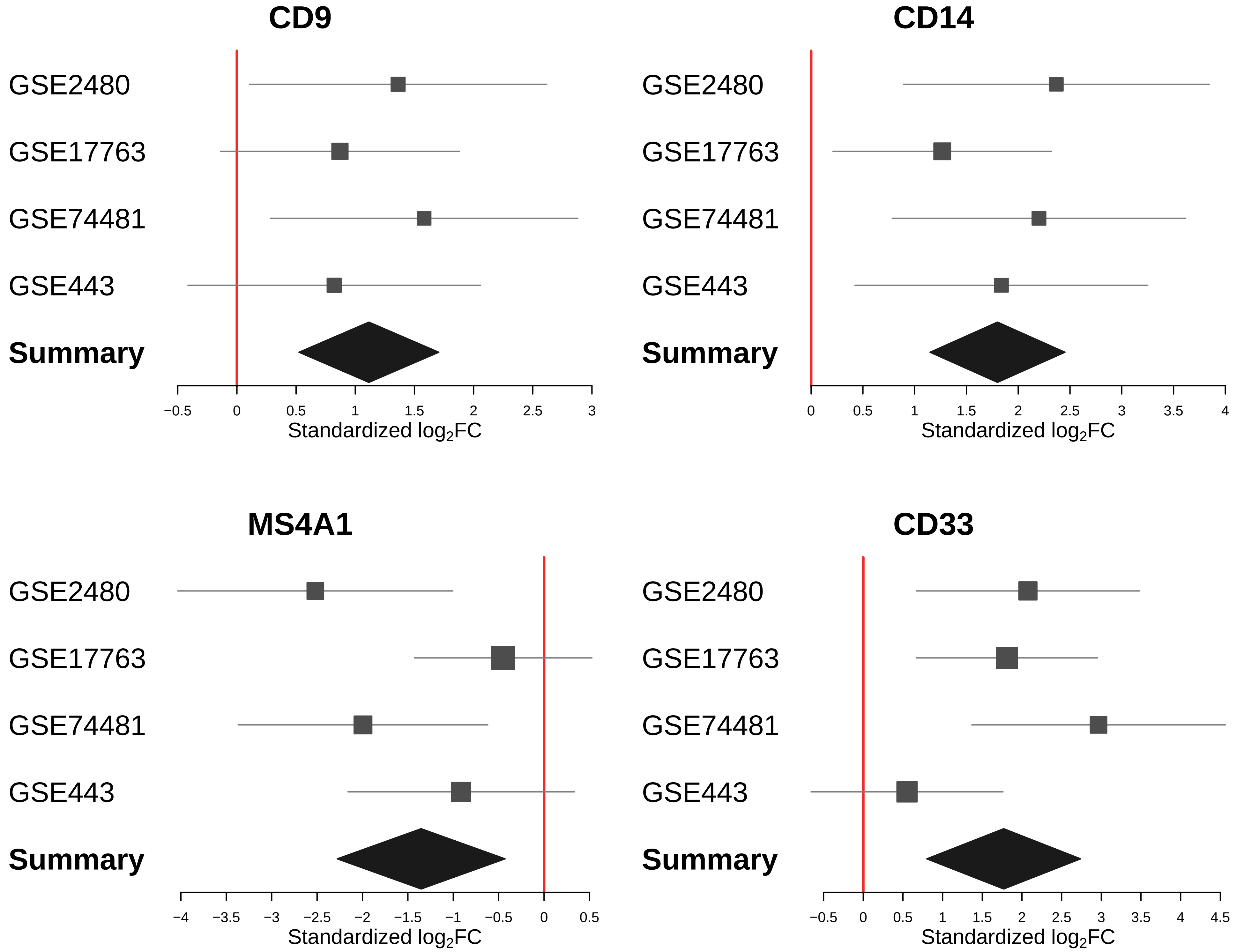

CD9
CD14
MS4A1
CD33
GSE2480
GSE17763
GSE74481
GSE443
Summary
Standardized log2FC

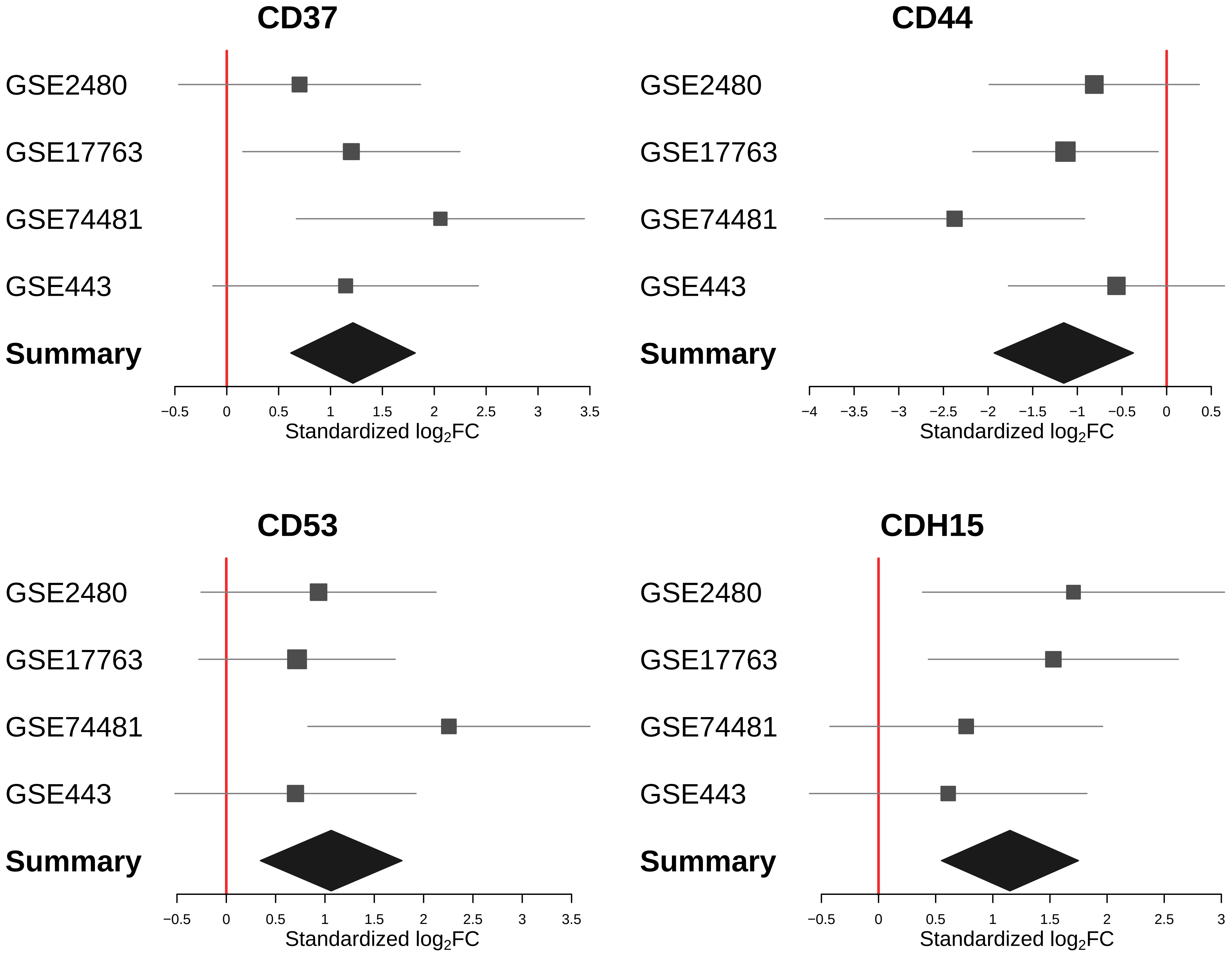

CD37
GSE2480
GSE17763
GSE74481
GSE443
Summary
−0.5 0 0.5 1 1.5 2 2.5 3 3.5
Standardized log2FC
CD44
GSE2480
GSE17763
GSE74481
GSE443
Summary
−4 −3.5 −3 −2.5 −2 −1.5 −1 −0.5 0 0.5
Standardized log2FC
CD53
GSE2480
GSE17763
GSE74481
GSE443
Summary
−0.5 0 0.5 1 1.5 2 2.5 3 3.5
Standardized log2FC
CDH15
GSE2480
GSE17763
GSE74481
GSE443
Summary
−0.5 0 0.5 1 1.5 2 2.5 3
Standardized log2FC

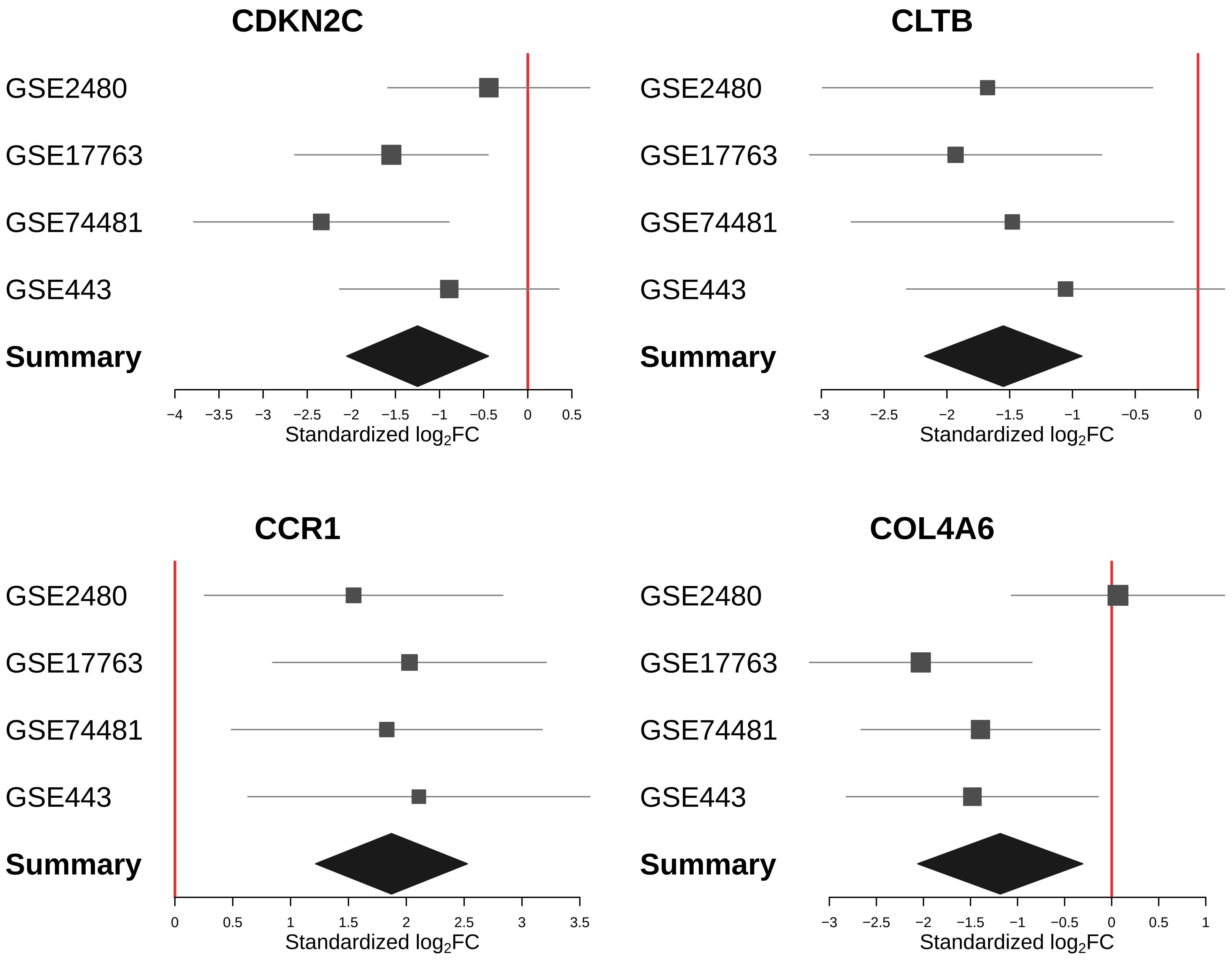
CDKN2C
GSE2480
GSE17763
GSE74481
GSE443
Summary
−4 −3.5 −3 −2.5 −2 −1.5 −1 −0.5 0 0.5
Standardized $\log_2$FC
CLTB
GSE2480
GSE17763
GSE74481
GSE443
Summary
−3 −2.5 −2 −1.5 −1 −0.5 0
Standardized $\log_2$FC
CCR1
GSE2480
GSE17763
GSE74481
GSE443
Summary
0 0.5 1 1.5 2 2.5 3 3.5
Standardized $\log_2$FC
COL4A6
GSE2480
GSE17763
GSE74481
GSE443
Summary
−3 −2.5 −2 −1.5 −1 −0.5 0 0.5 1
Standardized $\log_2$FC

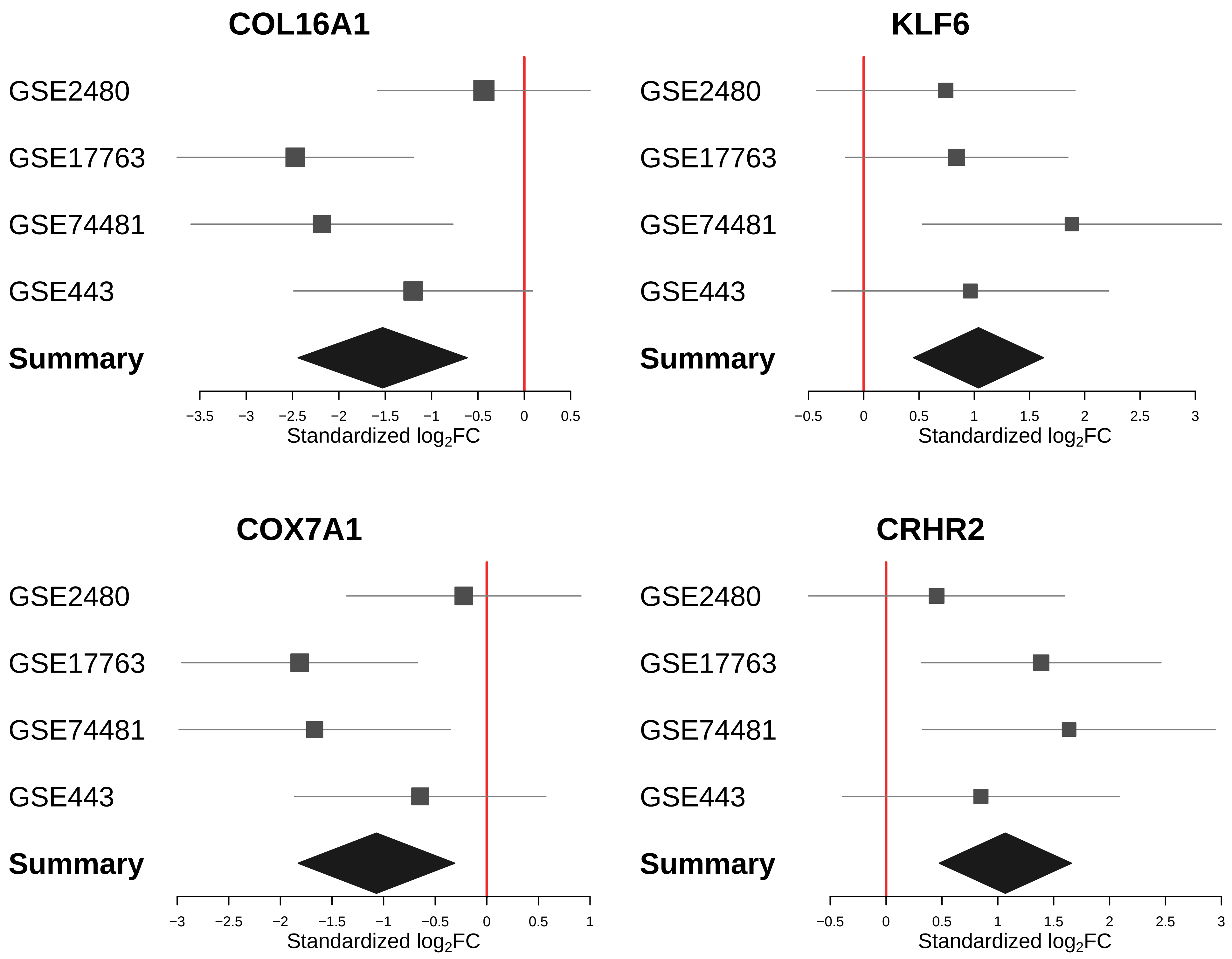

COL16A1
KLF6
COX7A1
CRHR2
GSE2480
GSE17763
GSE74481
GSE443
Summary
Standardized $\log_2$FC
−3.5 −3 −2.5 −2 −1.5 −1 −0.5 0 0.5
−0.5 0 0.5 1 1.5 2 2.5 3
−3 −2.5 −2 −1.5 −1 −0.5 0 0.5 1

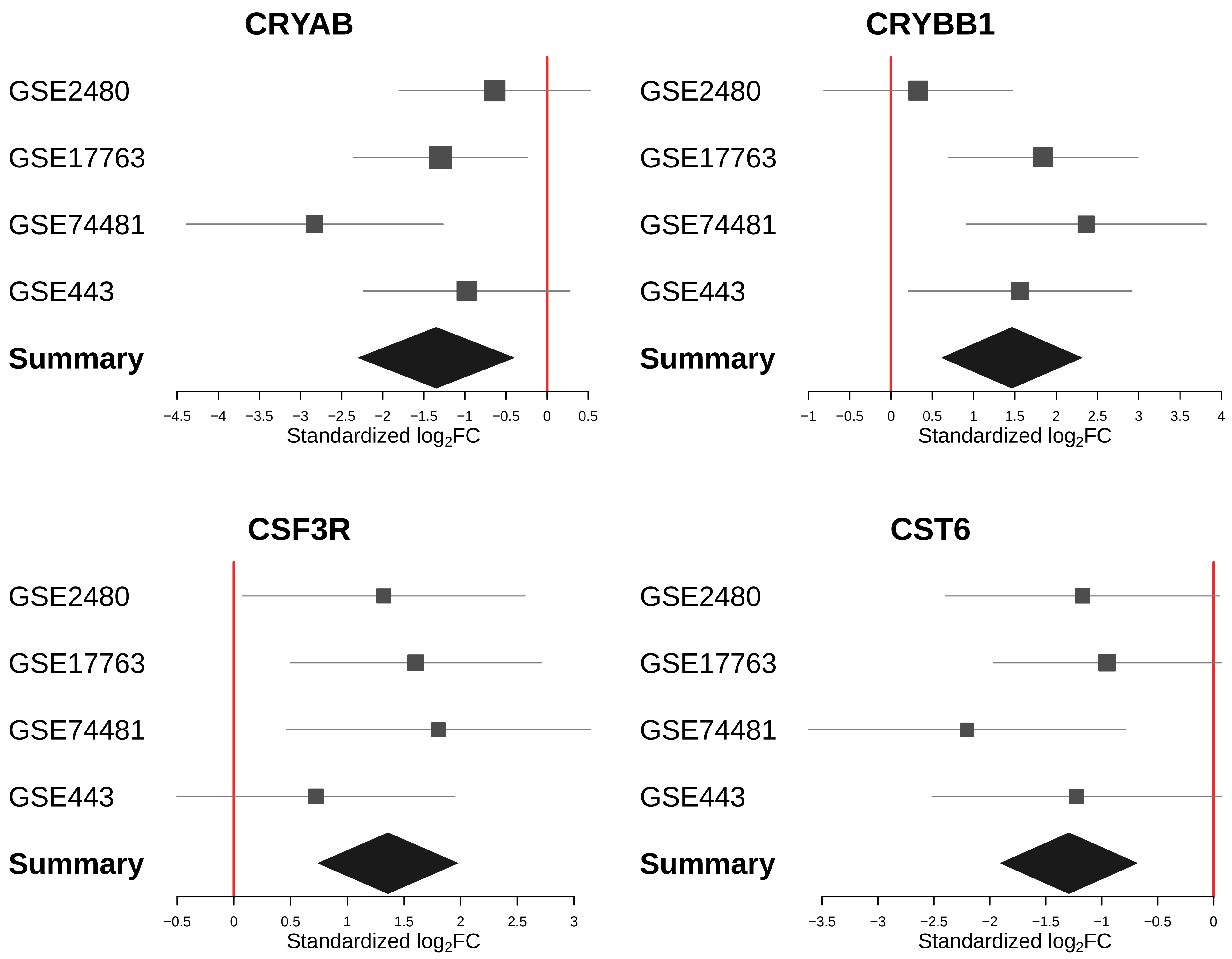

CRYAB
GSE2480
GSE17763
GSE74481
GSE443
Summary
−4.5 −4 −3.5 −3 −2.5 −2 −1.5 −1 −0.5 0 0.5
Standardized $log_2FC$
CRYBB1
GSE2480
GSE17763
GSE74481
GSE443
Summary
−1 −0.5 0 0.5 1 1.5 2 2.5 3 3.5 4
Standardized $log_2FC$
CSF3R
GSE2480
GSE17763
GSE74481
GSE443
Summary
−0.5 0 0.5 1 1.5 2 2.5 3
Standardized $log_2FC$
CST6
GSE2480
GSE17763
GSE74481
GSE443
Summary
−3.5 −3 −2.5 −2 −1.5 −1 −0.5 0
Standardized $log_2FC$

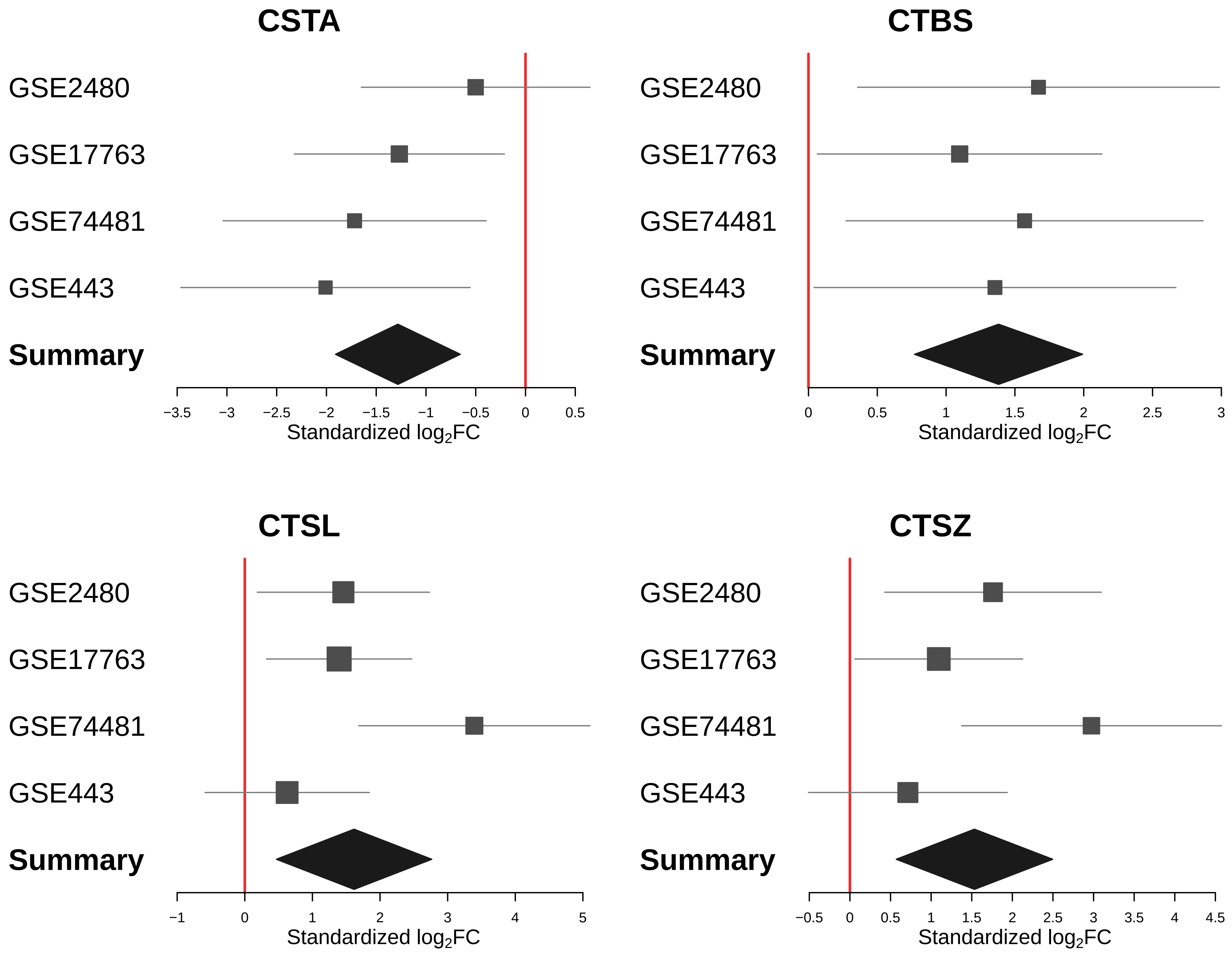
CSTA
CTBS
CTSL
CTSZ
GSE2480
GSE17763
GSE74481
GSE443
Summary
Standardized $log_2FC$
−3.5 −3 −2.5 −2 −1.5 −1 −0.5 0 0.5
0 0.5 1 1.5 2 2.5 3
−1 0 1 2 3 4 5
−0.5 0 0.5 1 1.5 2 2.5 3 3.5 4 4.5

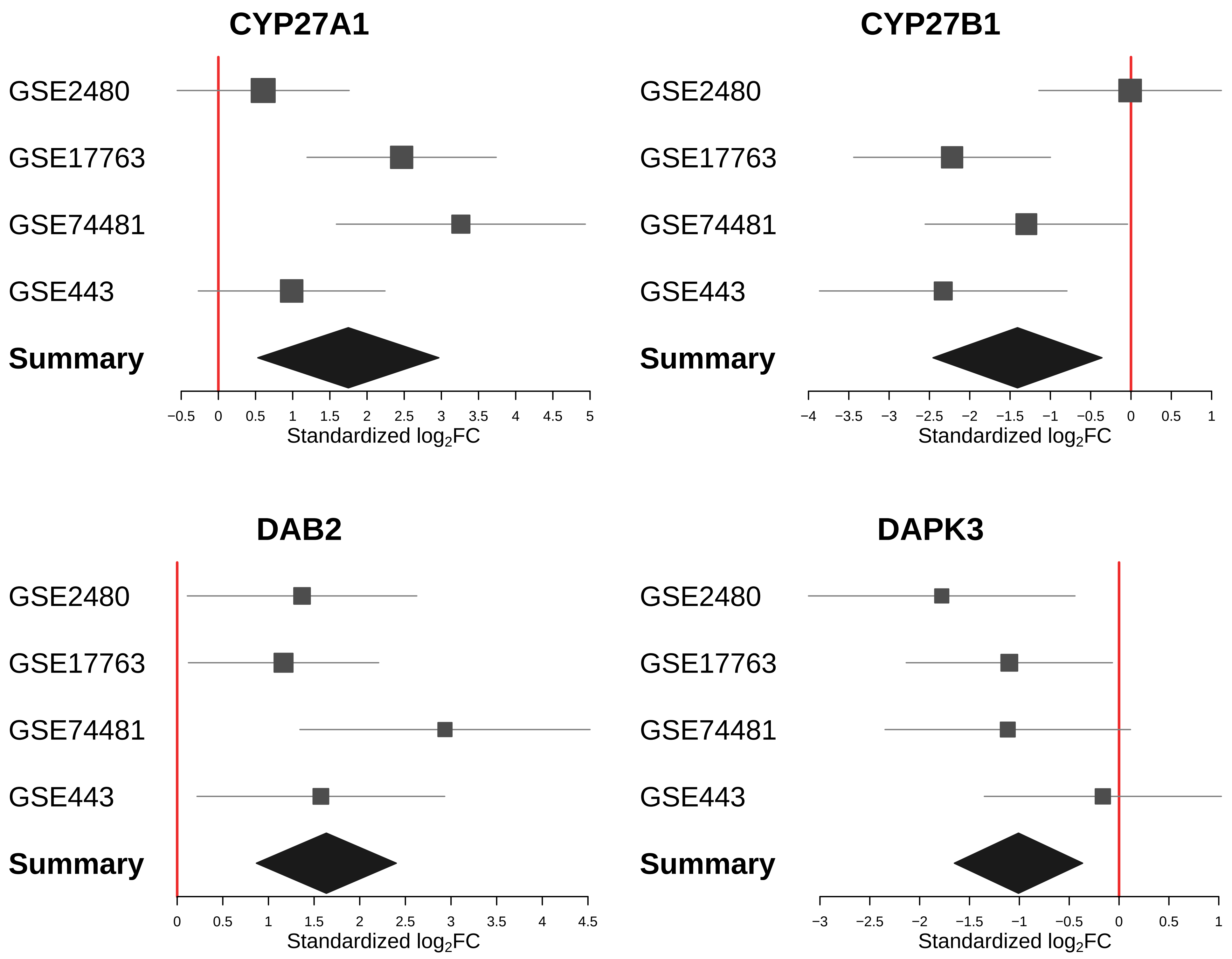
CYP27A1
CYP27B1
DAB2
DAPK3
GSE2480
GSE17763
GSE74481
GSE443
Summary
Standardized log2FC
−0.5 0 0.5 1 1.5 2 2.5 3 3.5 4 4.5 5
−4 −3.5 −3 −2.5 −2 −1.5 −1 −0.5 0 0.5 1
0 0.5 1 1.5 2 2.5 3 3.5 4 4.5
−3 −2.5 −2 −1.5 −1 −0.5 0 0.5 1

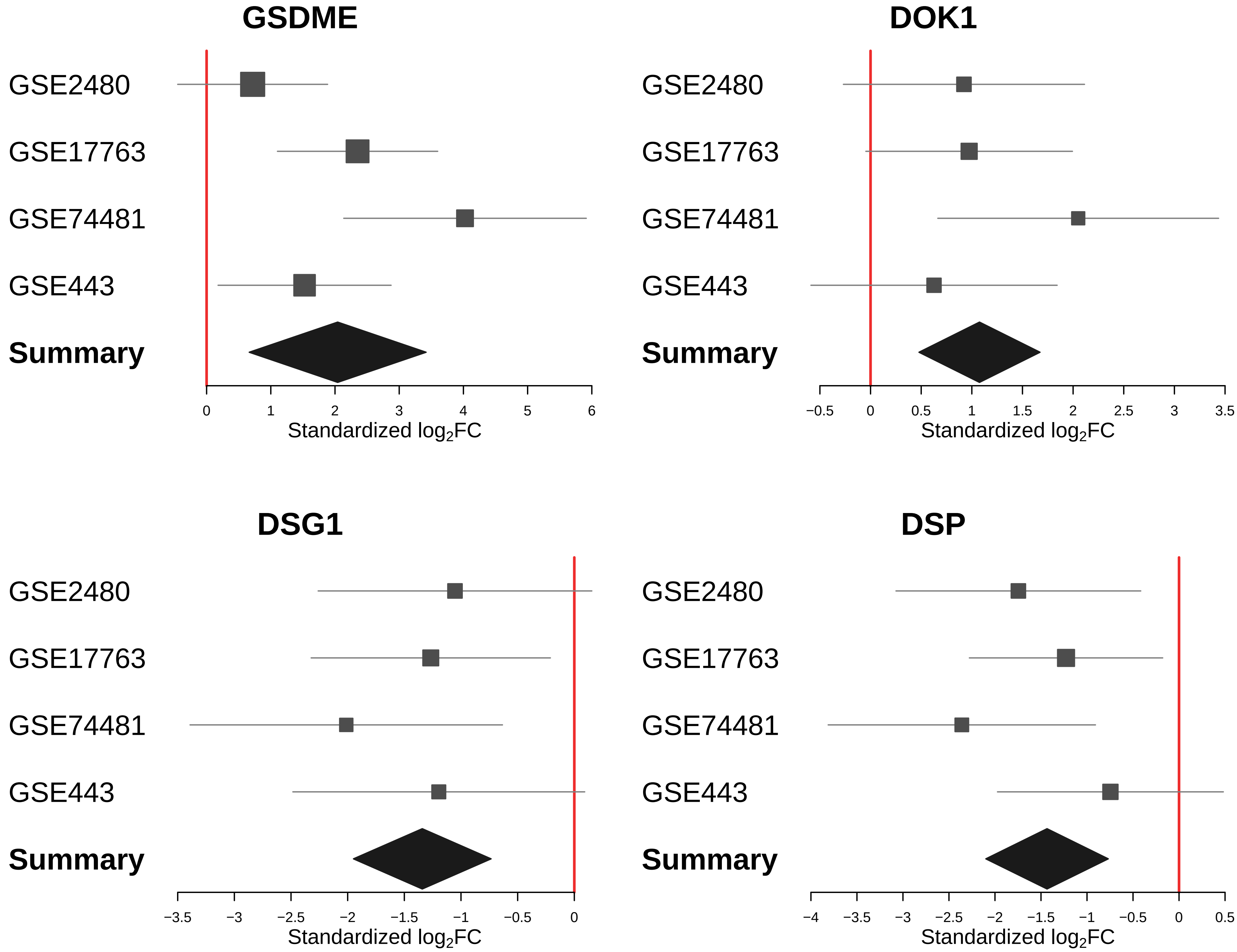
GSDME
DOK1
DSG1
DSP
GSE2480
GSE17763
GSE74481
GSE443
Summary
0 1 2 3 4 5 6
−0.5 0 0.5 1 1.5 2 2.5 3 3.5
−3.5 −3 −2.5 −2 −1.5 −1 −0.5 0
−4 −3.5 −3 −2.5 −2 −1.5 −1 −0.5 0 0.5
Standardized log2FC

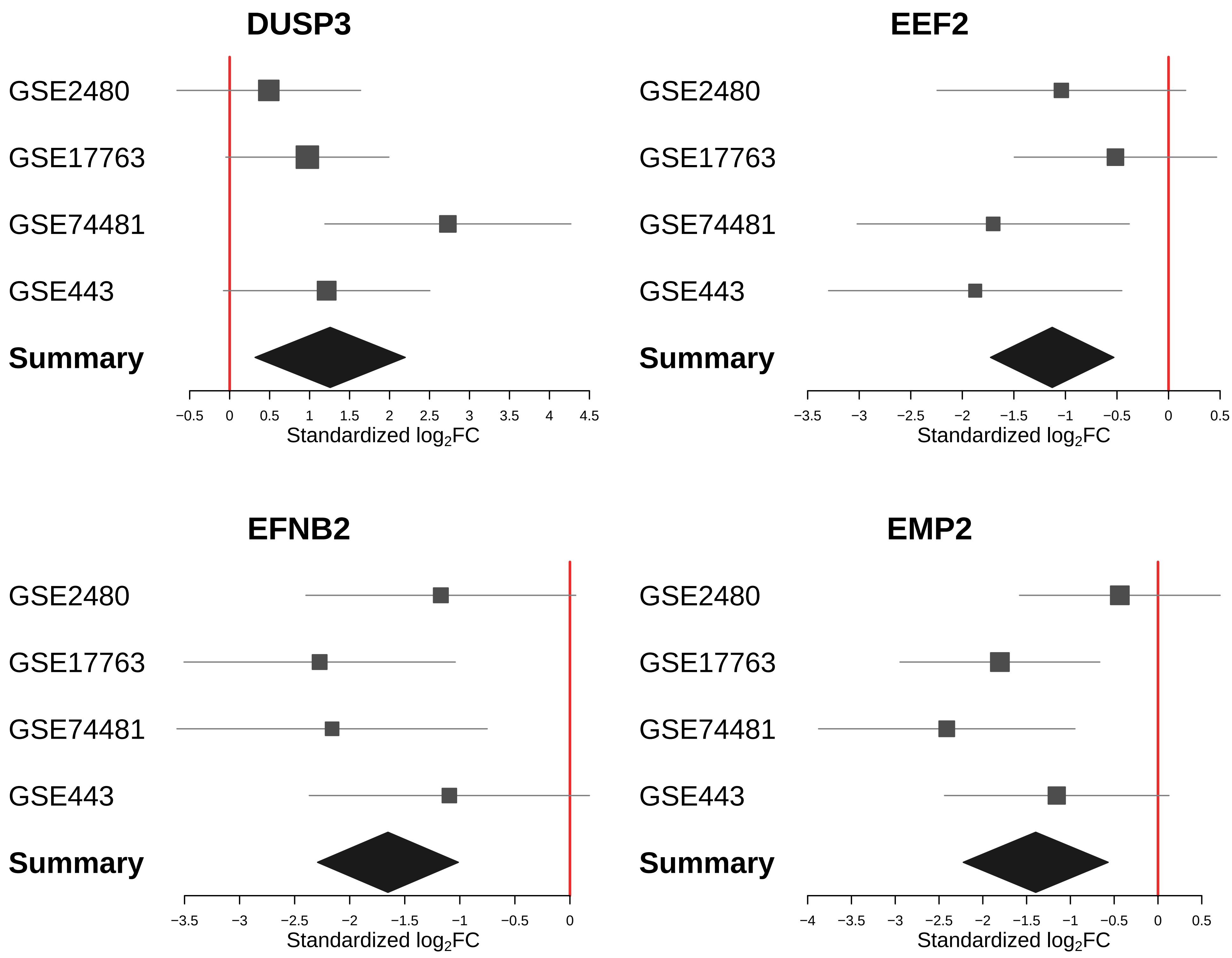
DUSP3
GSE2480
GSE17763
GSE74481
GSE443
Summary
−0.5 0 0.5 1 1.5 2 2.5 3 3.5 4 4.5
Standardized log2FC
EEF2
GSE2480
GSE17763
GSE74481
GSE443
Summary
−3.5 −3 −2.5 −2 −1.5 −1 −0.5 0 0.5
Standardized log2FC
EFNB2
GSE2480
GSE17763
GSE74481
GSE443
Summary
−3.5 −3 −2.5 −2 −1.5 −1 −0.5 0
Standardized log2FC
EMP2
GSE2480
GSE17763
GSE74481
GSE443
Summary
−4 −3.5 −3 −2.5 −2 −1.5 −1 −0.5 0 0.5
Standardized log2FC

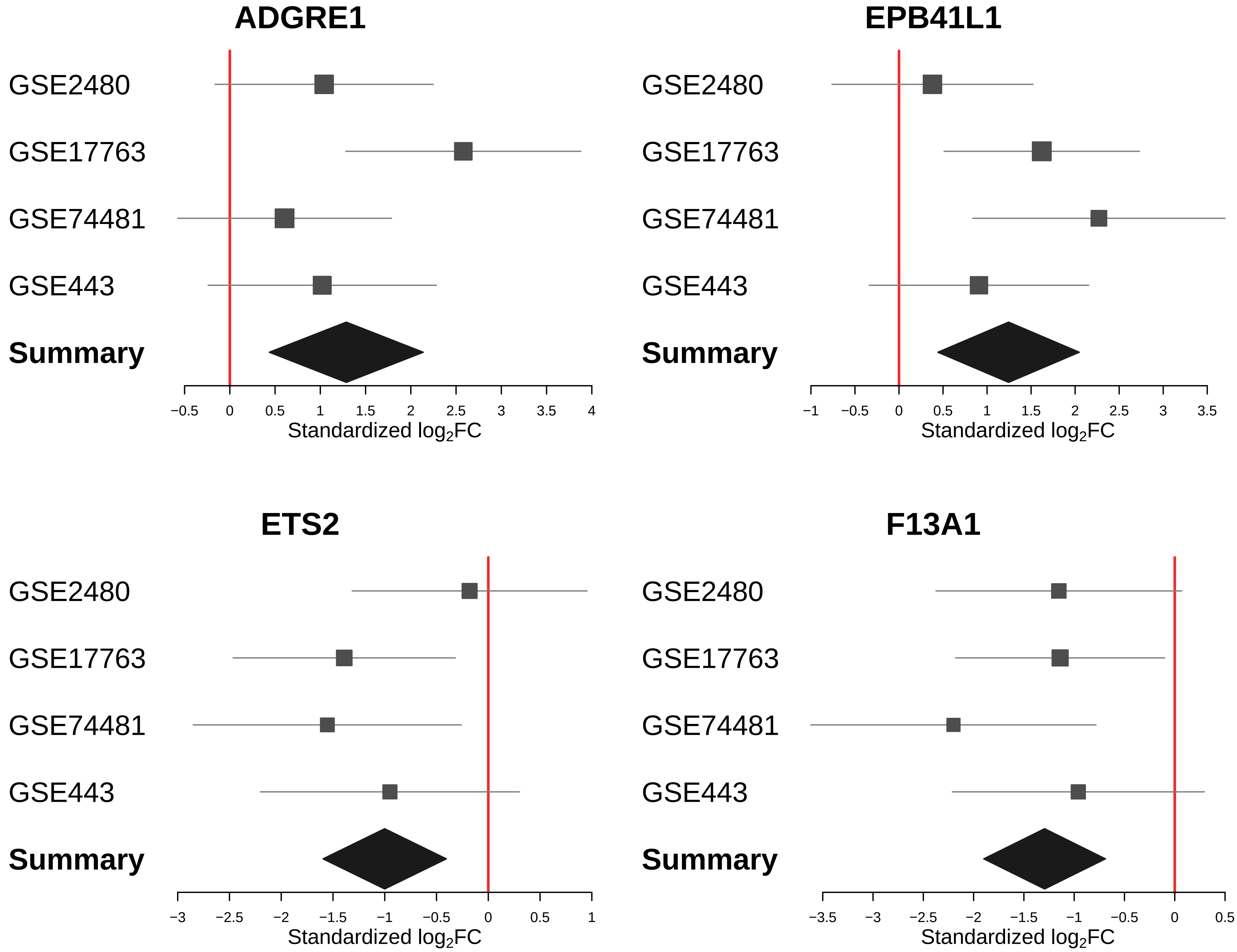
ADGRE1
EPB41L1
ETS2
F13A1
GSE2480
GSE17763
GSE74481
GSE443
Summary
Standardized $log_2FC$
−0.5 0 0.5 1 1.5 2 2.5 3 3.5 4
−1 −0.5 0 0.5 1 1.5 2 2.5 3 3.5
−3 −2.5 −2 −1.5 −1 −0.5 0 0.5 1
−3.5 −3 −2.5 −2 −1.5 −1 −0.5 0 0.5

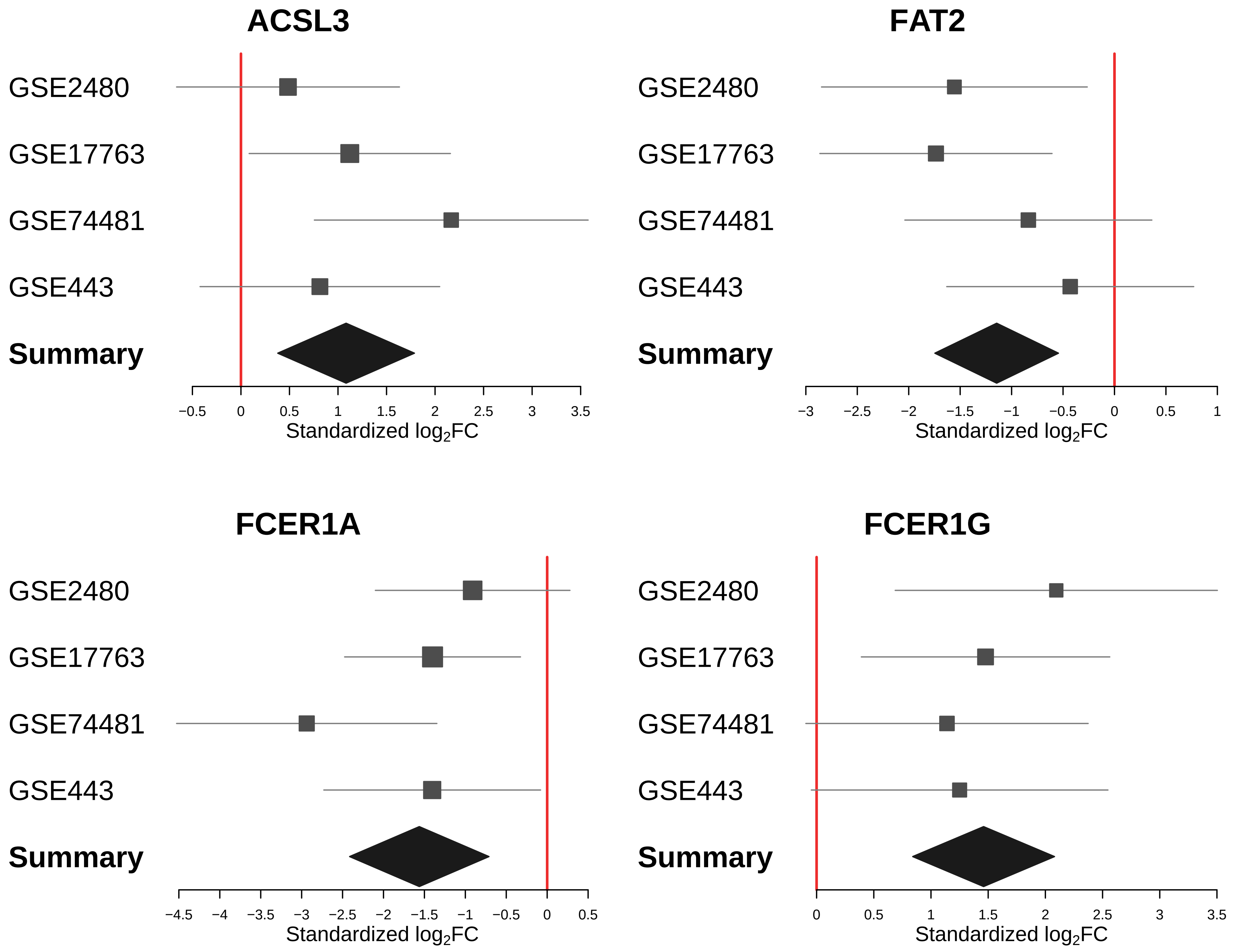

ACSL3
GSE2480
GSE17763
GSE74481
GSE443
Summary
−0.5 0 0.5 1 1.5 2 2.5 3 3.5
Standardized $log_2FC$
FAT2
GSE2480
GSE17763
GSE74481
GSE443
Summary
−3 −2.5 −2 −1.5 −1 −0.5 0 0.5 1
Standardized $log_2FC$
FCER1A
GSE2480
GSE17763
GSE74481
GSE443
Summary
−4.5 −4 −3.5 −3 −2.5 −2 −1.5 −1 −0.5 0 0.5
Standardized $log_2FC$
FCER1G
GSE2480
GSE17763
GSE74481
GSE443
Summary
0 0.5 1 1.5 2 2.5 3 3.5
Standardized $log_2FC$

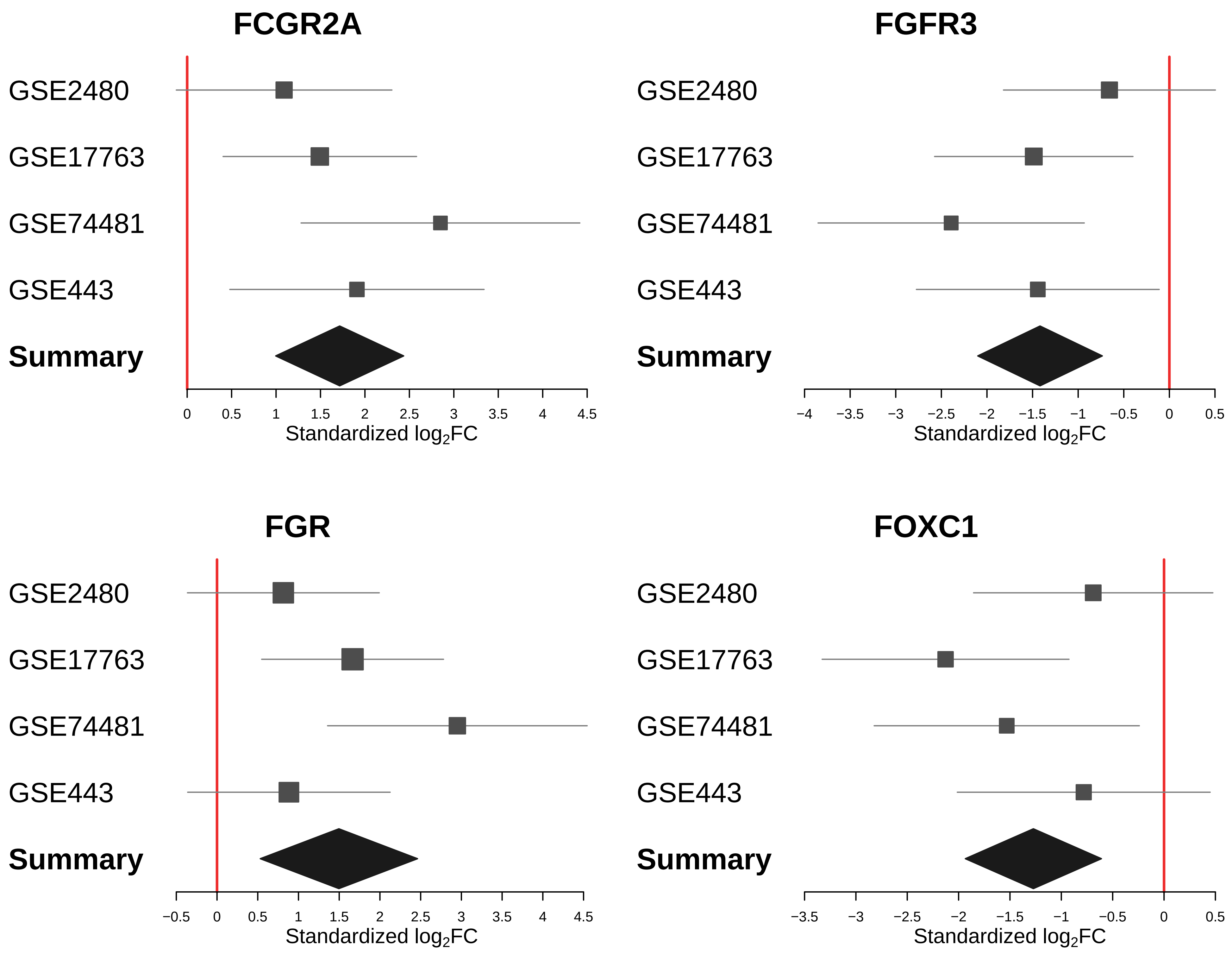
FCGR2A
FGFR3
FGR
FOXC1
GSE2480
GSE17763
GSE74481
GSE443
Summary
Standardized log2FC
0 0.5 1 1.5 2 2.5 3 3.5 4 4.5
−4 −3.5 −3 −2.5 −2 −1.5 −1 −0.5 0 0.5
−0.5 0 0.5 1 1.5 2 2.5 3 3.5 4 4.5
−3.5 −3 −2.5 −2 −1.5 −1 −0.5 0 0.5

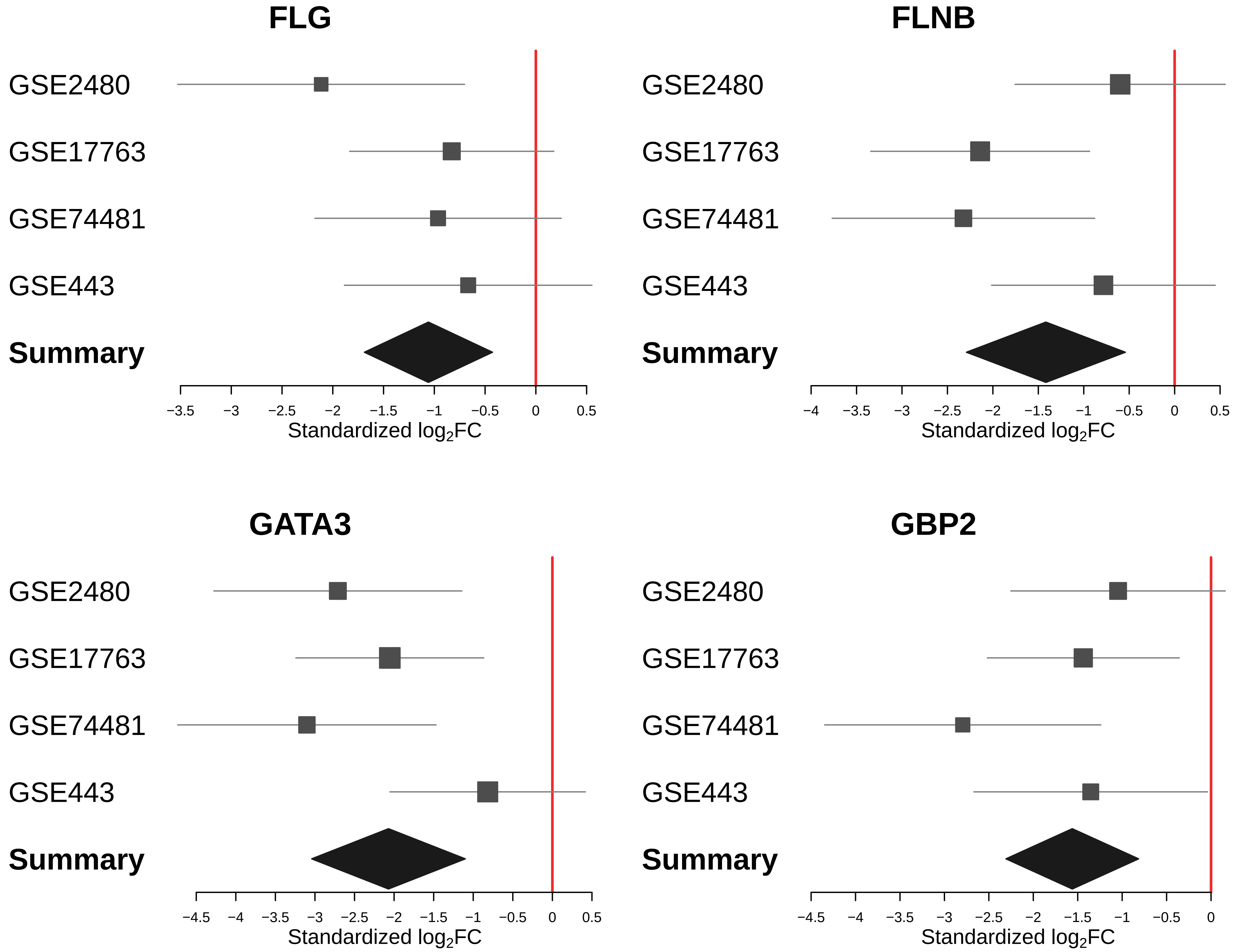
FLG
GSE2480
GSE17763
GSE74481
GSE443
Summary
−3.5 −3 −2.5 −2 −1.5 −1 −0.5 0 0.5
Standardized log2FC
FLNB
GSE2480
GSE17763
GSE74481
GSE443
Summary
−4 −3.5 −3 −2.5 −2 −1.5 −1 −0.5 0 0.5
Standardized log2FC
GATA3
GSE2480
GSE17763
GSE74481
GSE443
Summary
−4.5 −4 −3.5 −3 −2.5 −2 −1.5 −1 −0.5 0 0.5
Standardized log2FC
GBP2
GSE2480
GSE17763
GSE74481
GSE443
Summary
−4.5 −4 −3.5 −3 −2.5 −2 −1.5 −1 −0.5 0
Standardized log2FC

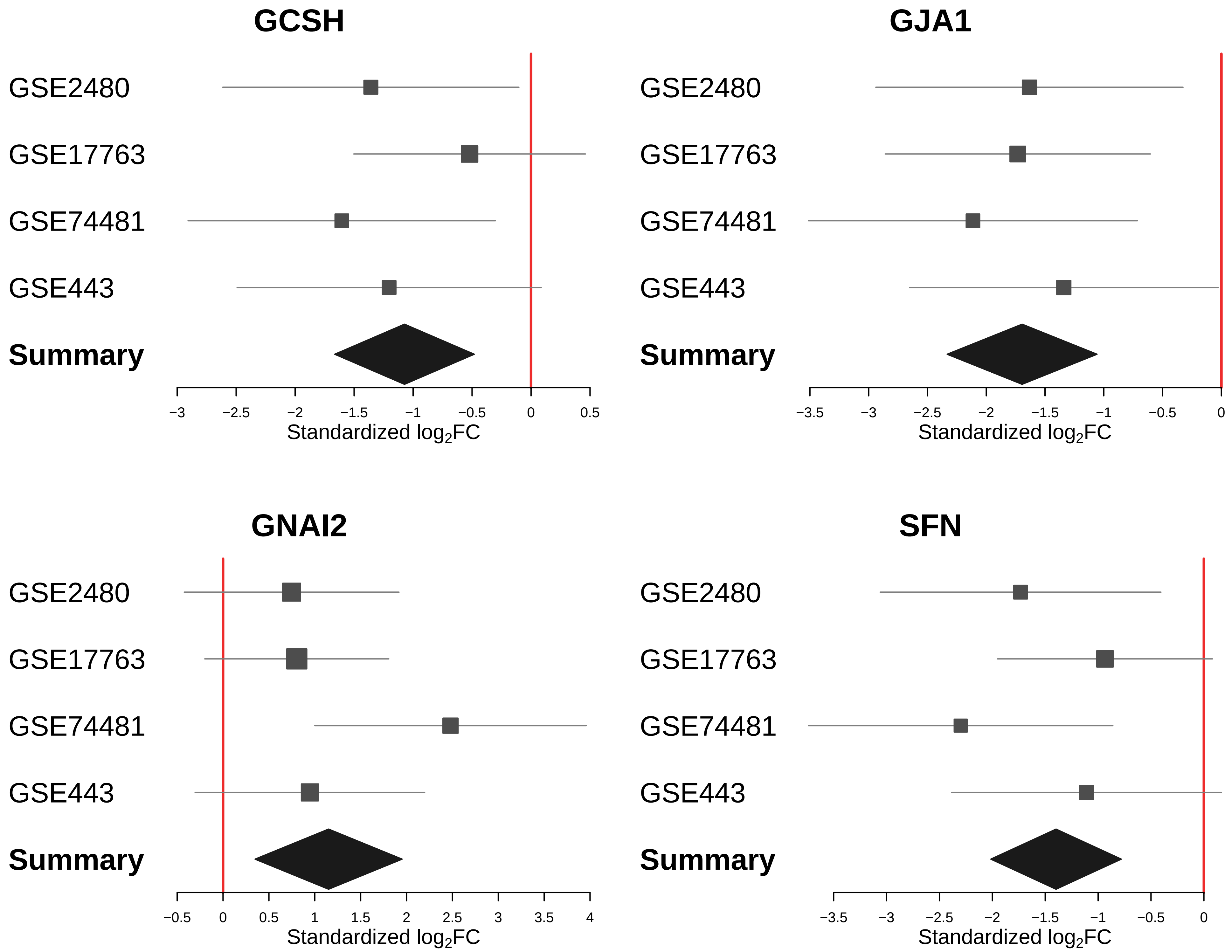

GCSH
GSE2480
GSE17763
GSE74481
GSE443
Summary
−3 −2.5 −2 −1.5 −1 −0.5 0 0.5
Standardized $log_2FC$
GJA1
GSE2480
GSE17763
GSE74481
GSE443
Summary
−3.5 −3 −2.5 −2 −1.5 −1 −0.5 0
Standardized $log_2FC$
GNAI2
GSE2480
GSE17763
GSE74481
GSE443
Summary
−0.5 0 0.5 1 1.5 2 2.5 3 3.5 4
Standardized $log_2FC$
SFN
GSE2480
GSE17763
GSE74481
GSE443
Summary
−3.5 −3 −2.5 −2 −1.5 −1 −0.5 0
Standardized $log_2FC$

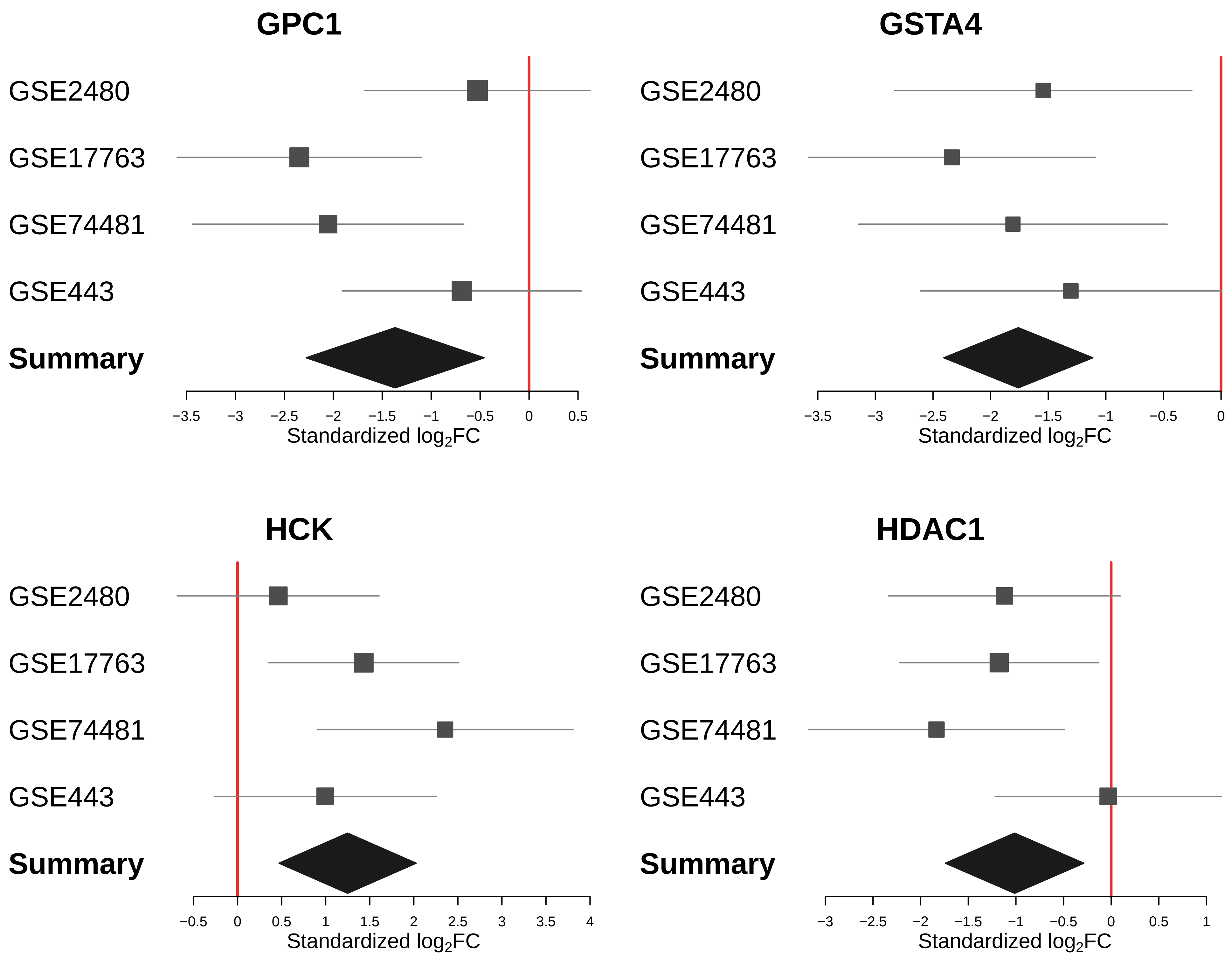
GPC1
GSE2480
GSE17763
GSE74481
GSE443
Summary
−3.5 −3 −2.5 −2 −1.5 −1 −0.5 0 0.5
Standardized log2FC
GSTA4
GSE2480
GSE17763
GSE74481
GSE443
Summary
−3.5 −3 −2.5 −2 −1.5 −1 −0.5 0
Standardized log2FC
HCK
GSE2480
GSE17763
GSE74481
GSE443
Summary
−0.5 0 0.5 1 1.5 2 2.5 3 3.5 4
Standardized log2FC
HDAC1
GSE2480
GSE17763
GSE74481
GSE443
Summary
−3 −2.5 −2 −1.5 −1 −0.5 0 0.5 1
Standardized log2FC

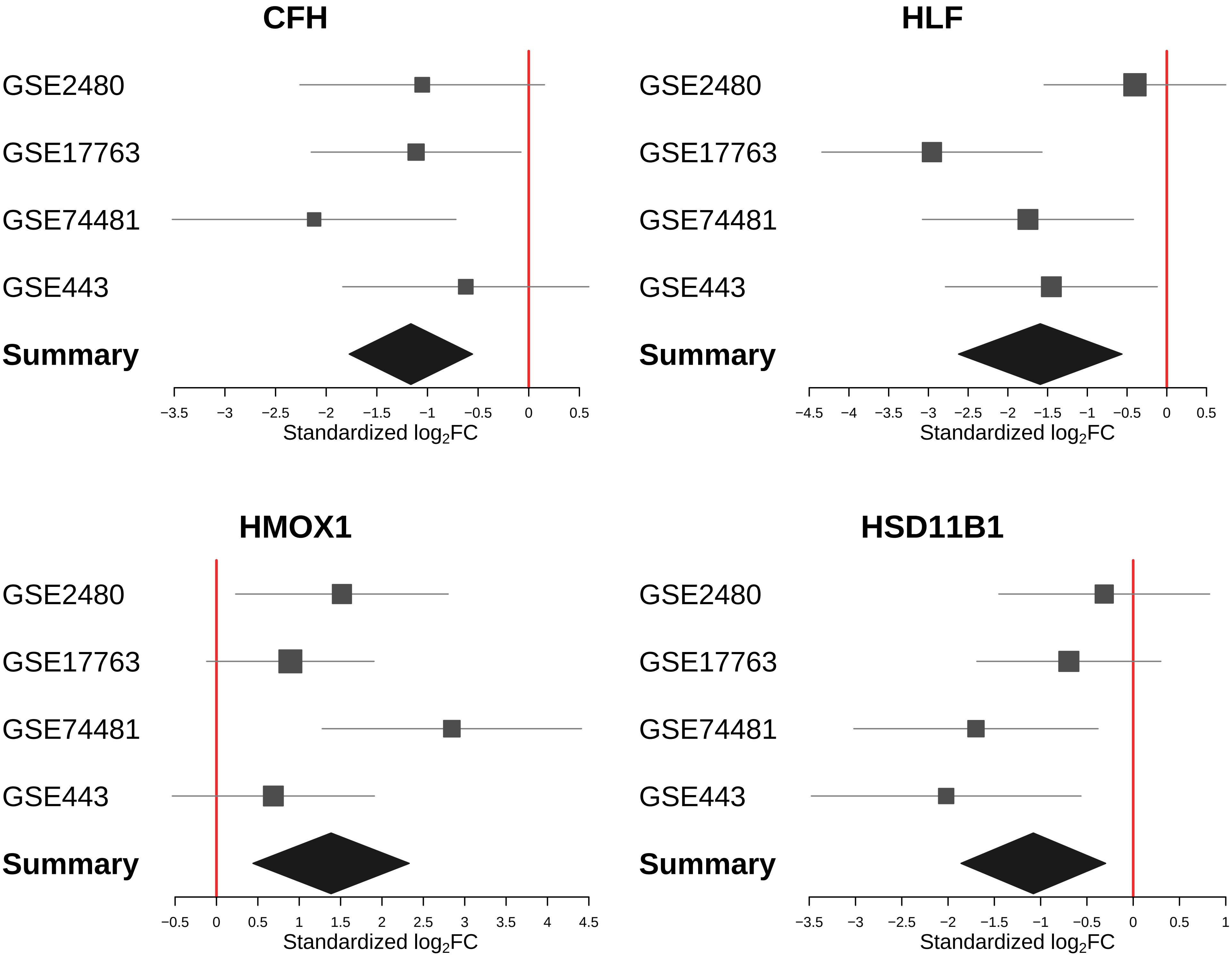

CFH
GSE2480
GSE17763
GSE74481
GSE443
Summary
−3.5 −3 −2.5 −2 −1.5 −1 −0.5 0 0.5
Standardized log2FC
HLF
GSE2480
GSE17763
GSE74481
GSE443
Summary
−4.5 −4 −3.5 −3 −2.5 −2 −1.5 −1 −0.5 0 0.5
Standardized log2FC
HMOX1
GSE2480
GSE17763
GSE74481
GSE443
Summary
−0.5 0 0.5 1 1.5 2 2.5 3 3.5 4 4.5
Standardized log2FC
HSD11B1
GSE2480
GSE17763
GSE74481
GSE443
Summary
−3.5 −3 −2.5 −2 −1.5 −1 −0.5 0 0.5 1
Standardized log2FC

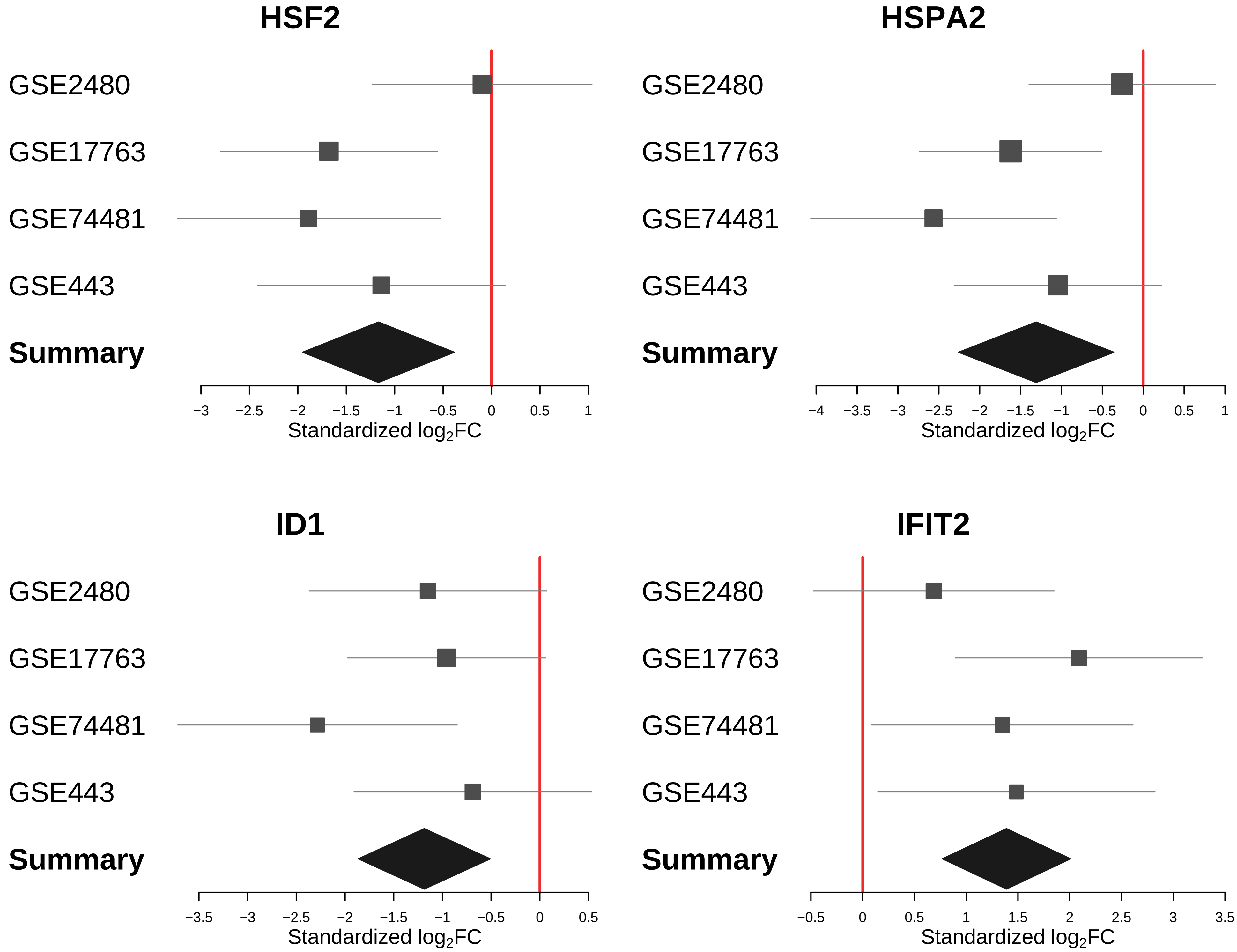
HSF2
HSPA2
ID1
IFIT2
GSE2480
GSE17763
GSE74481
GSE443
Summary
Standardized log2FC
−3 −2.5 −2 −1.5 −1 −0.5 0 0.5 1
−4 −3.5 −3 −2.5 −2 −1.5 −1 −0.5 0 0.5 1
−3.5 −3 −2.5 −2 −1.5 −1 −0.5 0 0.5
−0.5 0 0.5 1 1.5 2 2.5 3 3.5

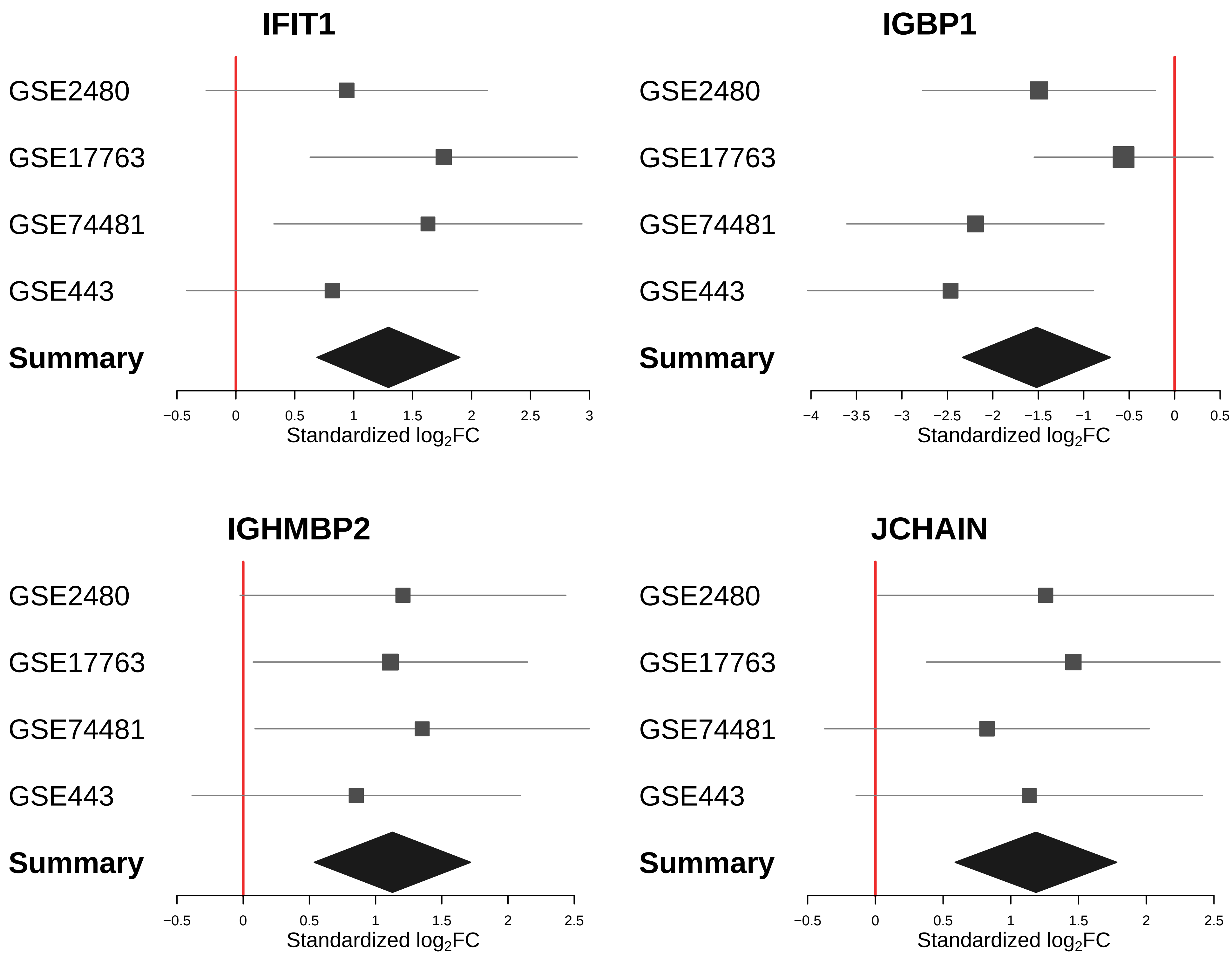
IFIT1
GSE2480
GSE17763
GSE74481
GSE443
Summary
−0.5 0 0.5 1 1.5 2 2.5 3
Standardized log2FC
IGBP1
GSE2480
GSE17763
GSE74481
GSE443
Summary
−4 −3.5 −3 −2.5 −2 −1.5 −1 −0.5 0 0.5
Standardized log2FC
IGHMBP2
GSE2480
GSE17763
GSE74481
GSE443
Summary
−0.5 0 0.5 1 1.5 2 2.5
Standardized log2FC
JCHAIN
GSE2480
GSE17763
GSE74481
GSE443
Summary
−0.5 0 0.5 1 1.5 2 2.5
Standardized log2FC

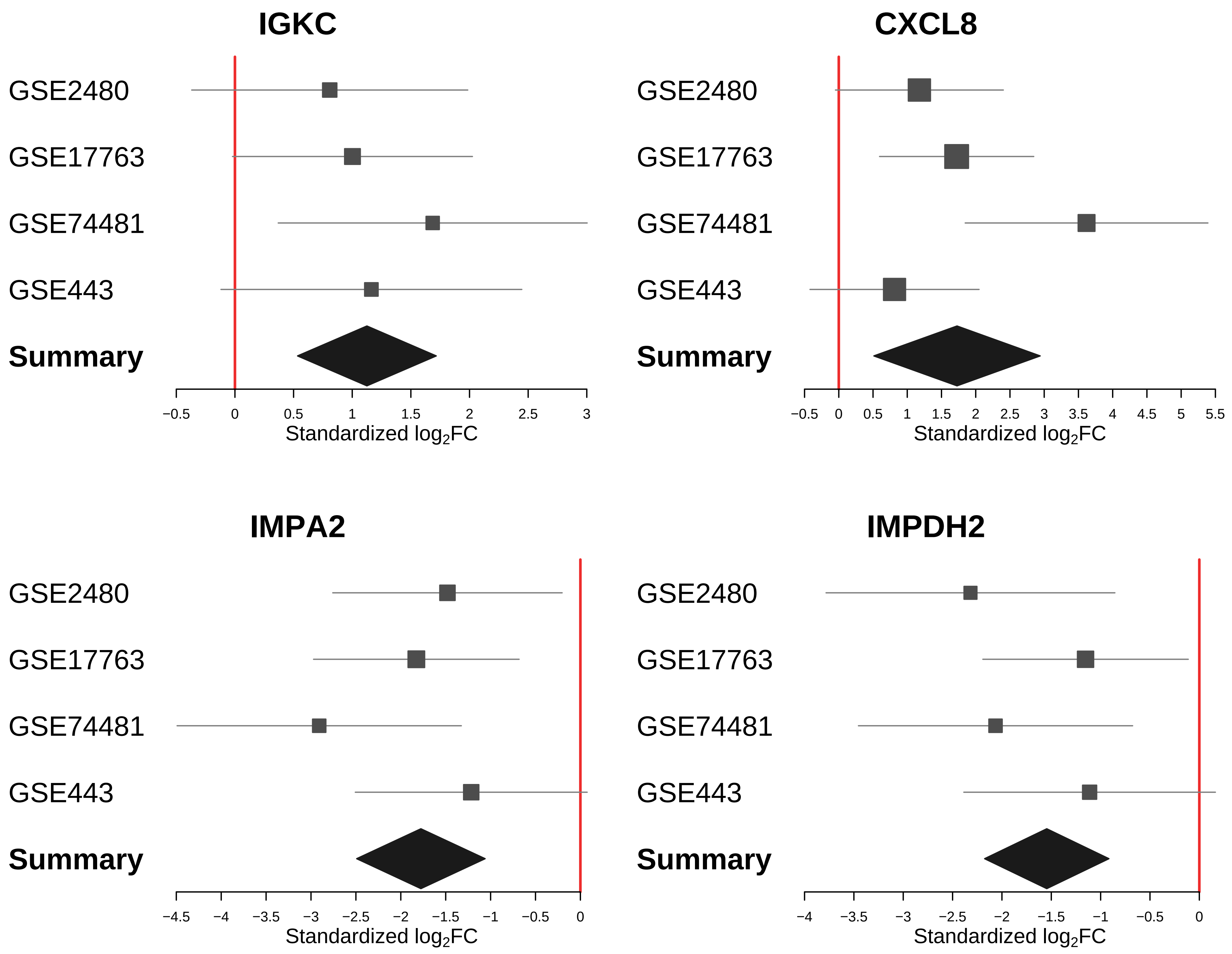

IGKC
CXCL8
IMPA2
IMPDH2
GSE2480
GSE17763
GSE74481
GSE443
Summary
Standardized $log_2FC$
−0.5 0 0.5 1 1.5 2 2.5 3
−0.5 0 0.5 1 1.5 2 2.5 3 3.5 4 4.5 5 5.5
−4.5 −4 −3.5 −3 −2.5 −2 −1.5 −1 −0.5 0
−4 −3.5 −3 −2.5 −2 −1.5 −1 −0.5 0

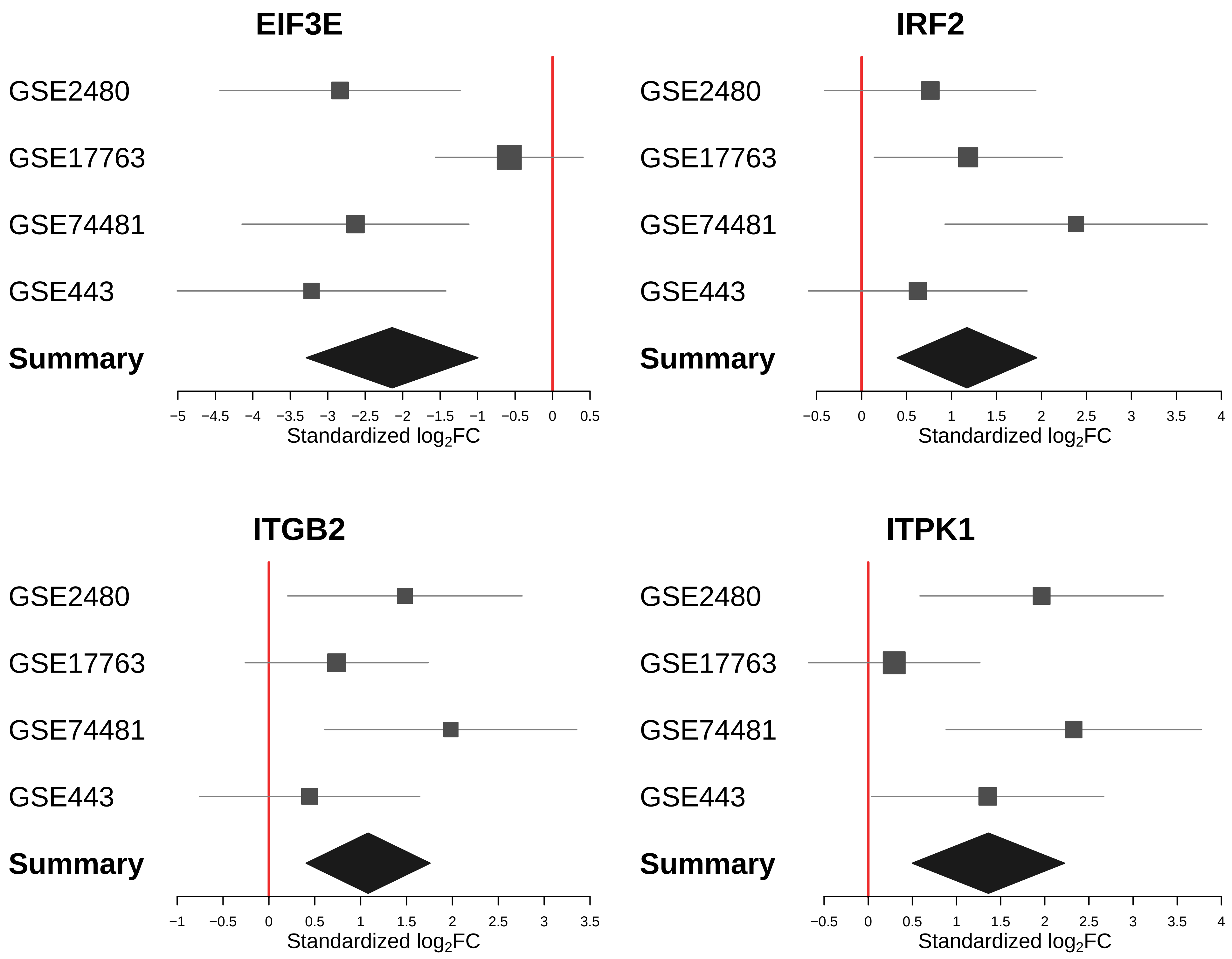
EIF3E
IRF2
ITGB2
ITPK1
GSE2480
GSE17763
GSE74481
GSE443
Summary
−5 −4.5 −4 −3.5 −3 −2.5 −2 −1.5 −1 −0.5 0 0.5
−0.5 0 0.5 1 1.5 2 2.5 3 3.5 4
−1 −0.5 0 0.5 1 1.5 2 2.5 3 3.5
Standardized log2FC

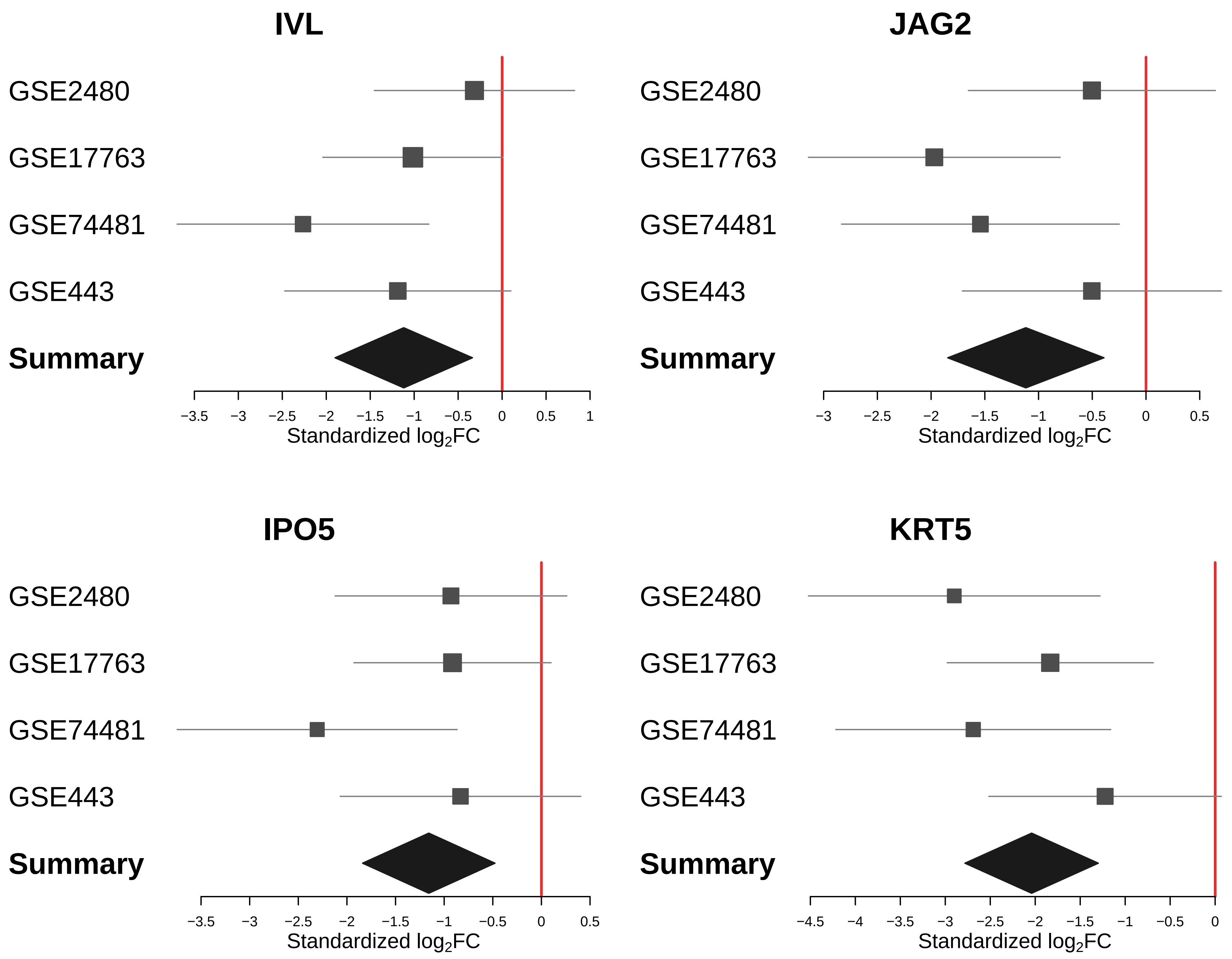

IVL
GSE2480
GSE17763
GSE74481
GSE443
Summary
−3.5 −3 −2.5 −2 −1.5 −1 −0.5 0 0.5 1
Standardized $log_2FC$
JAG2
GSE2480
GSE17763
GSE74481
GSE443
Summary
−3 −2.5 −2 −1.5 −1 −0.5 0 0.5
Standardized $log_2FC$
IPO5
GSE2480
GSE17763
GSE74481
GSE443
Summary
−3.5 −3 −2.5 −2 −1.5 −1 −0.5 0 0.5
Standardized $log_2FC$
KRT5
GSE2480
GSE17763
GSE74481
GSE443
Summary
−4.5 −4 −3.5 −3 −2.5 −2 −1.5 −1 −0.5 0
Standardized $log_2FC$

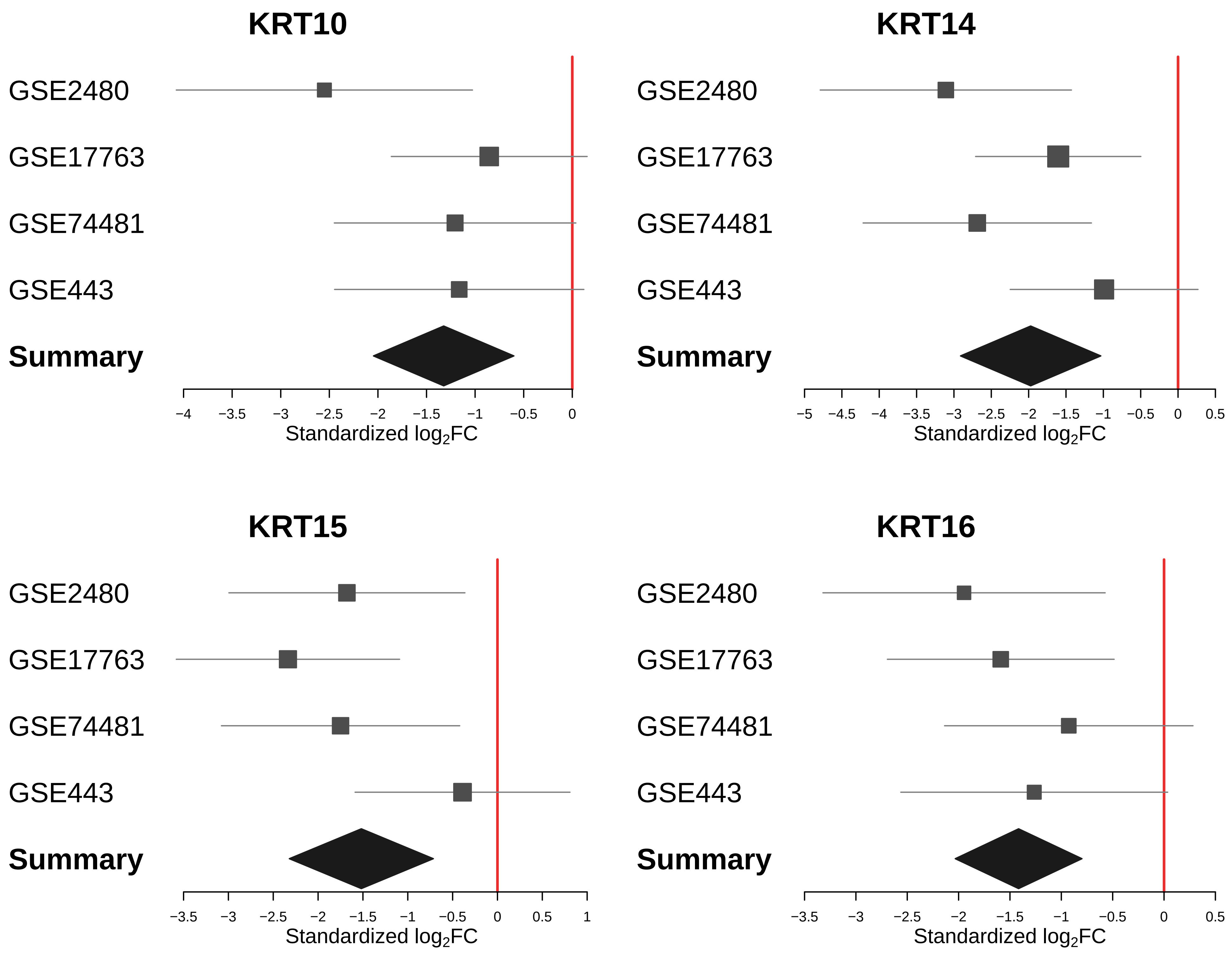
KRT10
GSE2480
GSE17763
GSE74481
GSE443
Summary
−4 −3.5 −3 −2.5 −2 −1.5 −1 −0.5 0
Standardized log2FC
KRT14
GSE2480
GSE17763
GSE74481
GSE443
Summary
−5 −4.5 −4 −3.5 −3 −2.5 −2 −1.5 −1 −0.5 0 0.5
Standardized log2FC
KRT15
GSE2480
GSE17763
GSE74481
GSE443
Summary
−3.5 −3 −2.5 −2 −1.5 −1 −0.5 0 0.5 1
Standardized log2FC
KRT16
GSE2480
GSE17763
GSE74481
GSE443
Summary
−3.5 −3 −2.5 −2 −1.5 −1 −0.5 0 0.5
Standardized log2FC

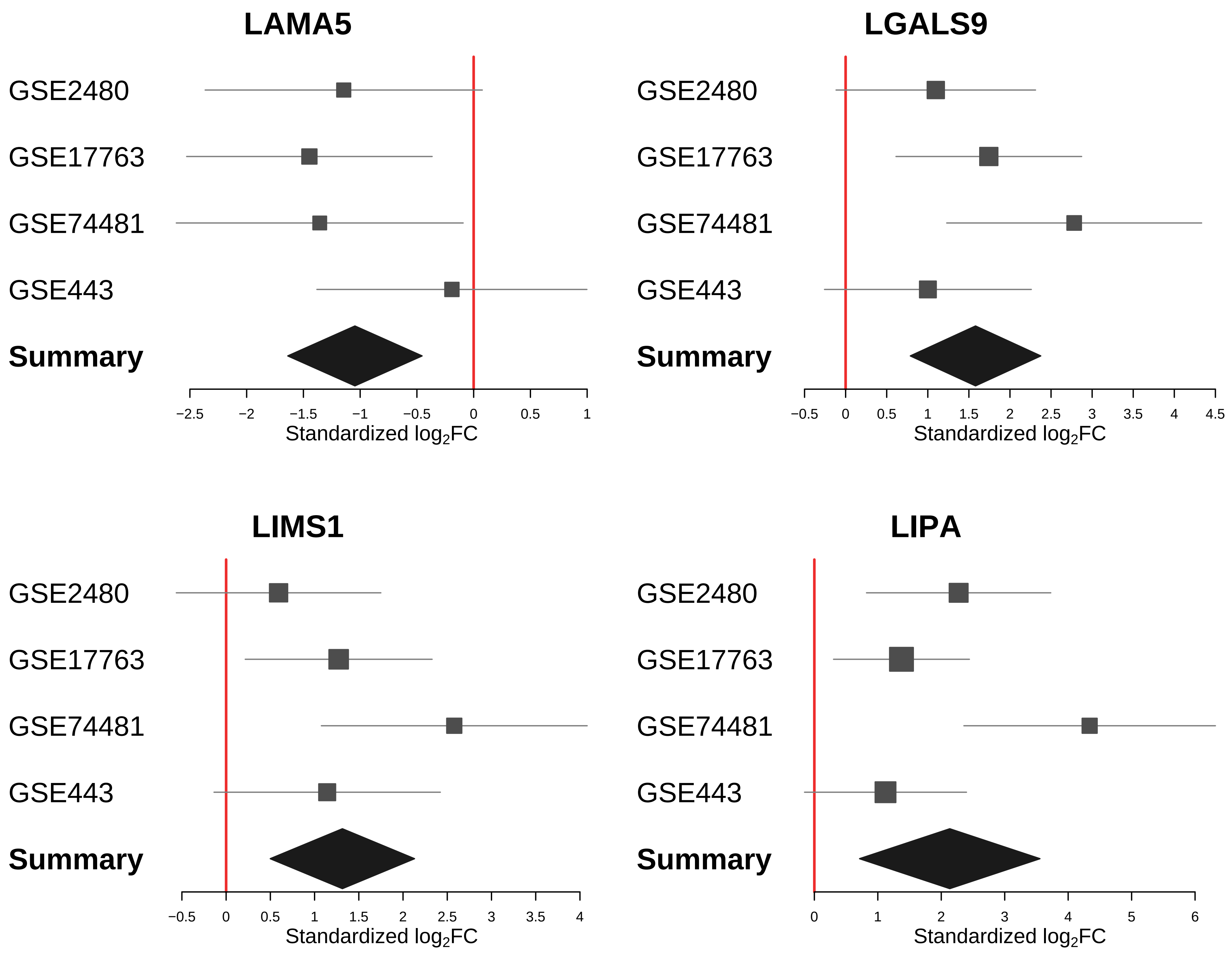

LAMA5
GSE2480
GSE17763
GSE74481
GSE443
Summary
−2.5 −2 −1.5 −1 −0.5 0 0.5 1
Standardized log2FC
LGALS9
GSE2480
GSE17763
GSE74481
GSE443
Summary
−0.5 0 0.5 1 1.5 2 2.5 3 3.5 4 4.5
Standardized log2FC
LIMS1
GSE2480
GSE17763
GSE74481
GSE443
Summary
−0.5 0 0.5 1 1.5 2 2.5 3 3.5 4
Standardized log2FC
LIPA
GSE2480
GSE17763
GSE74481
GSE443
Summary
0 1 2 3 4 5 6
Standardized log2FC

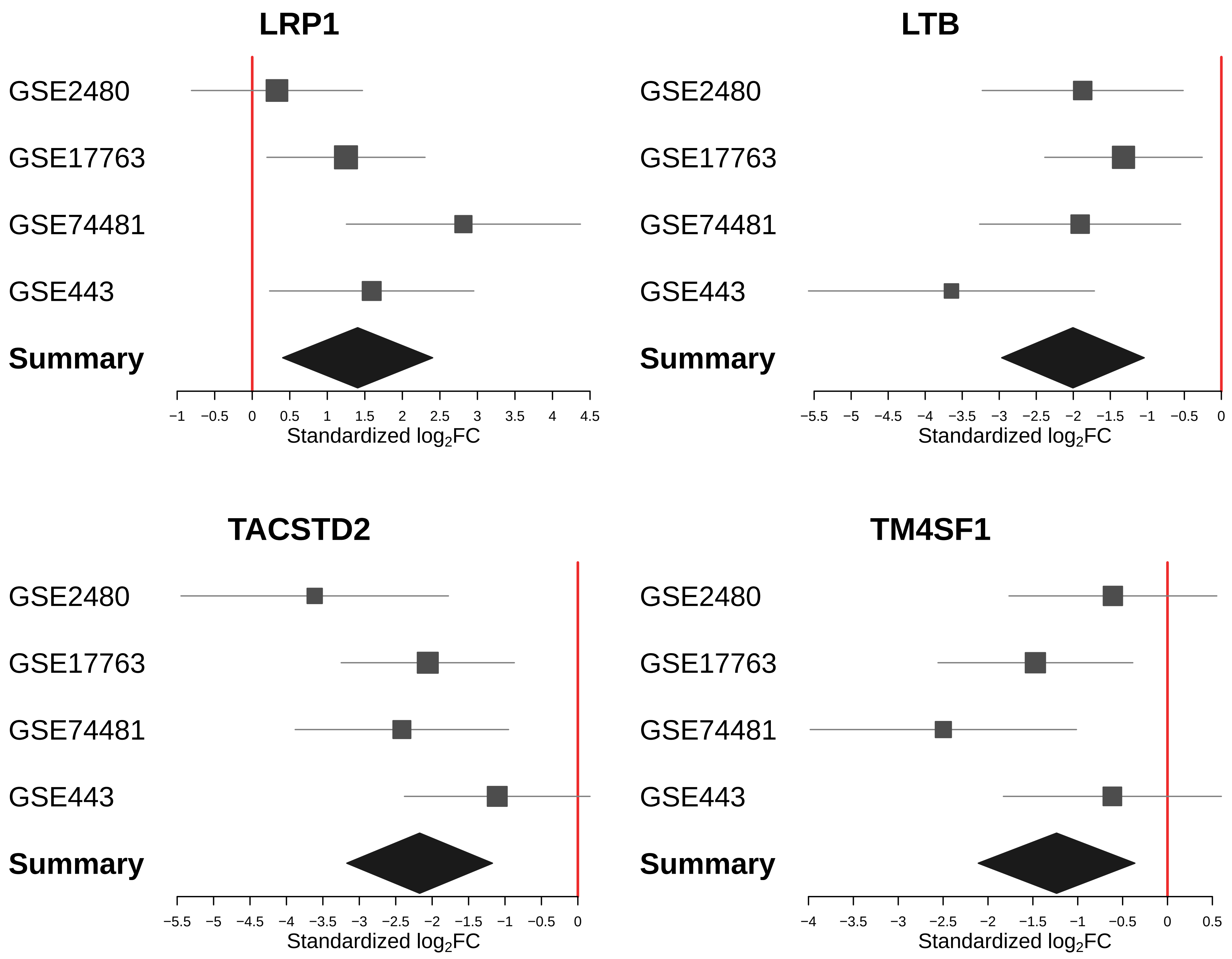

LRP1
LTB
TACSTD2
TM4SF1
GSE2480
GSE17763
GSE74481
GSE443
Summary
Standardized log2FC

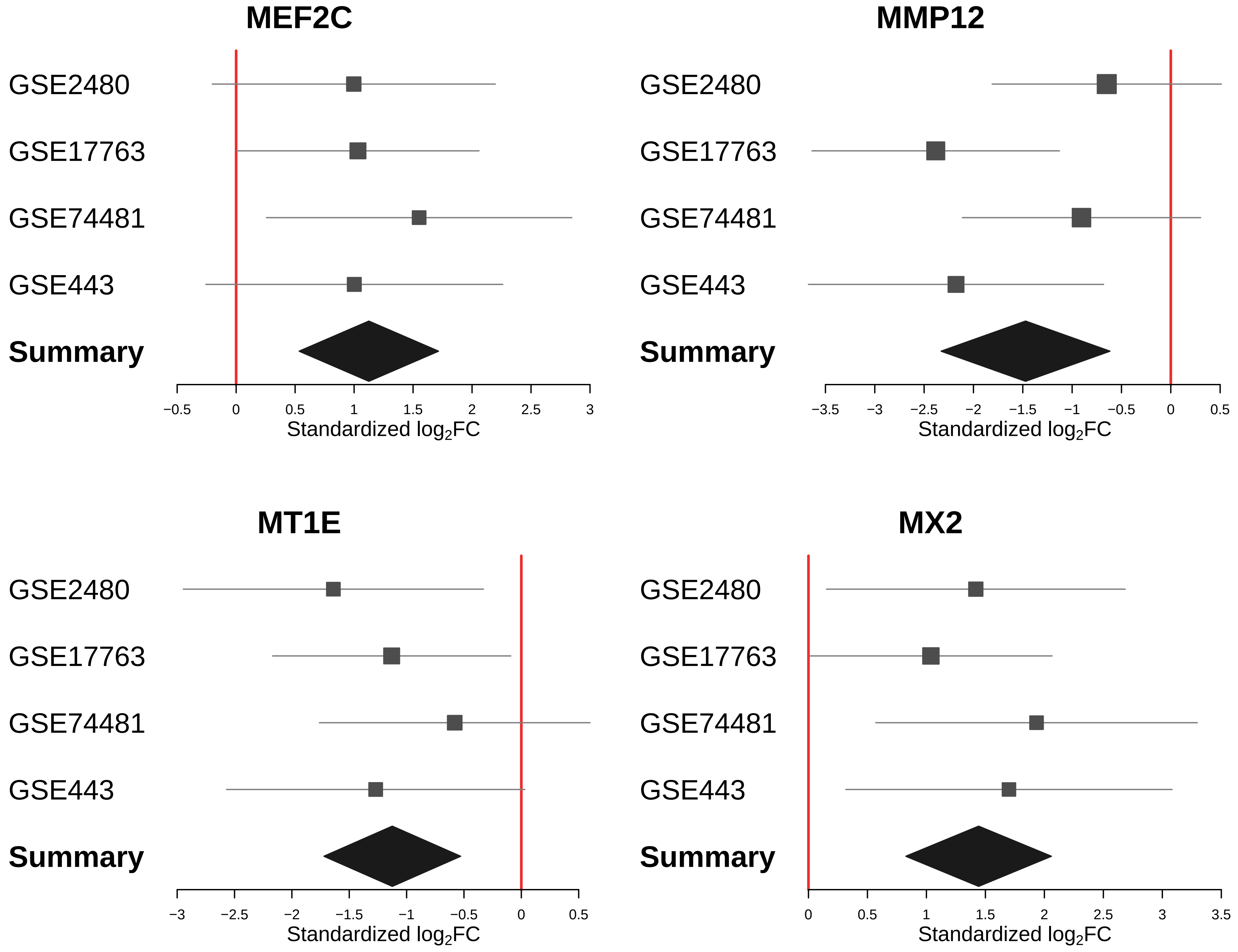
MEF2C
GSE2480
GSE17763
GSE74481
GSE443
Summary
−0.5 0 0.5 1 1.5 2 2.5 3
Standardized $log_2FC$
MMP12
GSE2480
GSE17763
GSE74481
GSE443
Summary
−3.5 −3 −2.5 −2 −1.5 −1 −0.5 0 0.5
Standardized $log_2FC$
MT1E
GSE2480
GSE17763
GSE74481
GSE443
Summary
−3 −2.5 −2 −1.5 −1 −0.5 0 0.5
Standardized $log_2FC$
MX2
GSE2480
GSE17763
GSE74481
GSE443
Summary
0 0.5 1 1.5 2 2.5 3 3.5
Standardized $log_2FC$

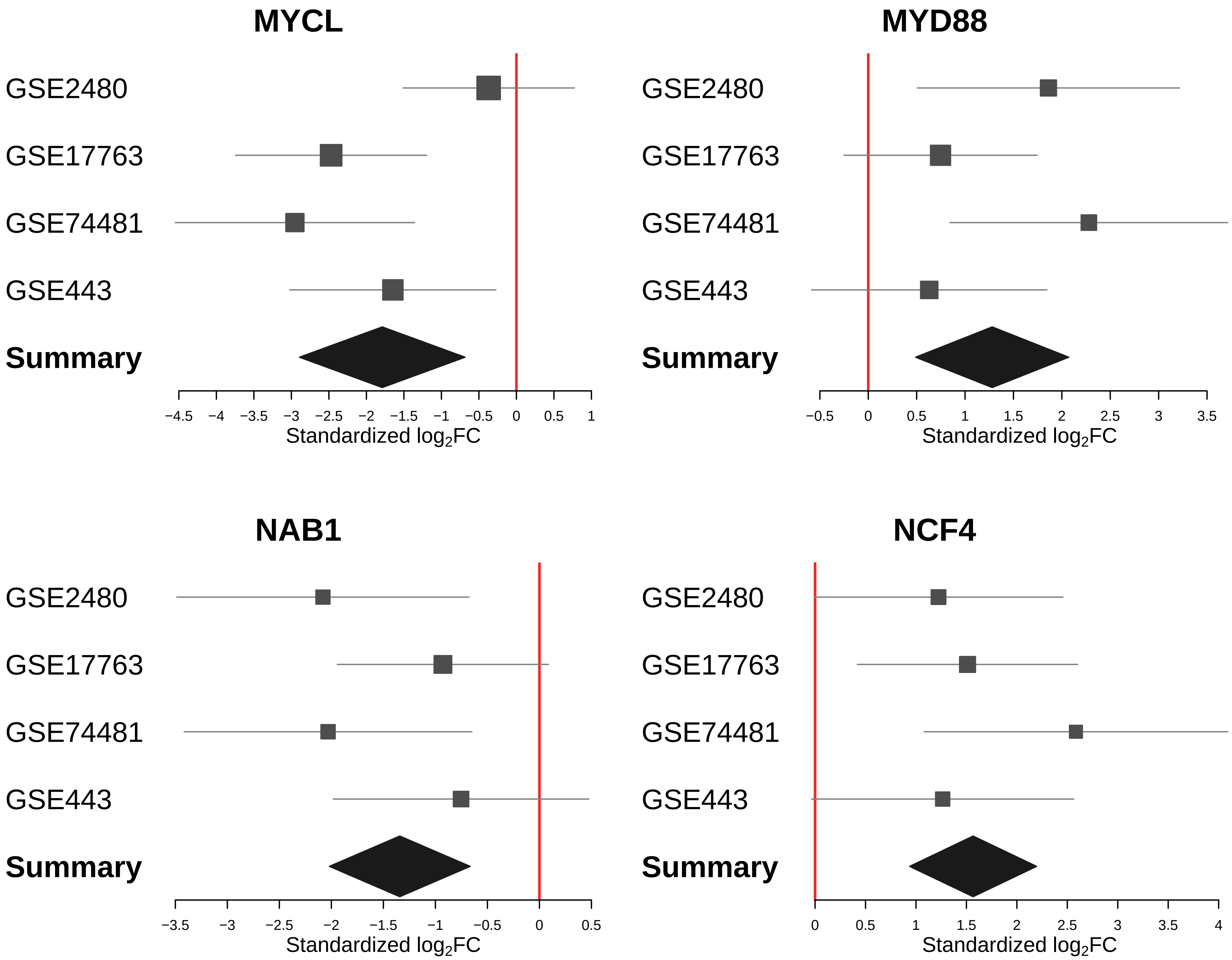
MYCL
GSE2480
GSE17763
GSE74481
GSE443
Summary
−4.5 −4 −3.5 −3 −2.5 −2 −1.5 −1 −0.5 0 0.5 1
Standardized $log_2FC$
MYD88
GSE2480
GSE17763
GSE74481
GSE443
Summary
−0.5 0 0.5 1 1.5 2 2.5 3 3.5
Standardized $log_2FC$
NAB1
GSE2480
GSE17763
GSE74481
GSE443
Summary
−3.5 −3 −2.5 −2 −1.5 −1 −0.5 0 0.5
Standardized $log_2FC$
NCF4
GSE2480
GSE17763
GSE74481
GSE443
Summary
0 0.5 1 1.5 2 2.5 3 3.5 4
Standardized $log_2FC$

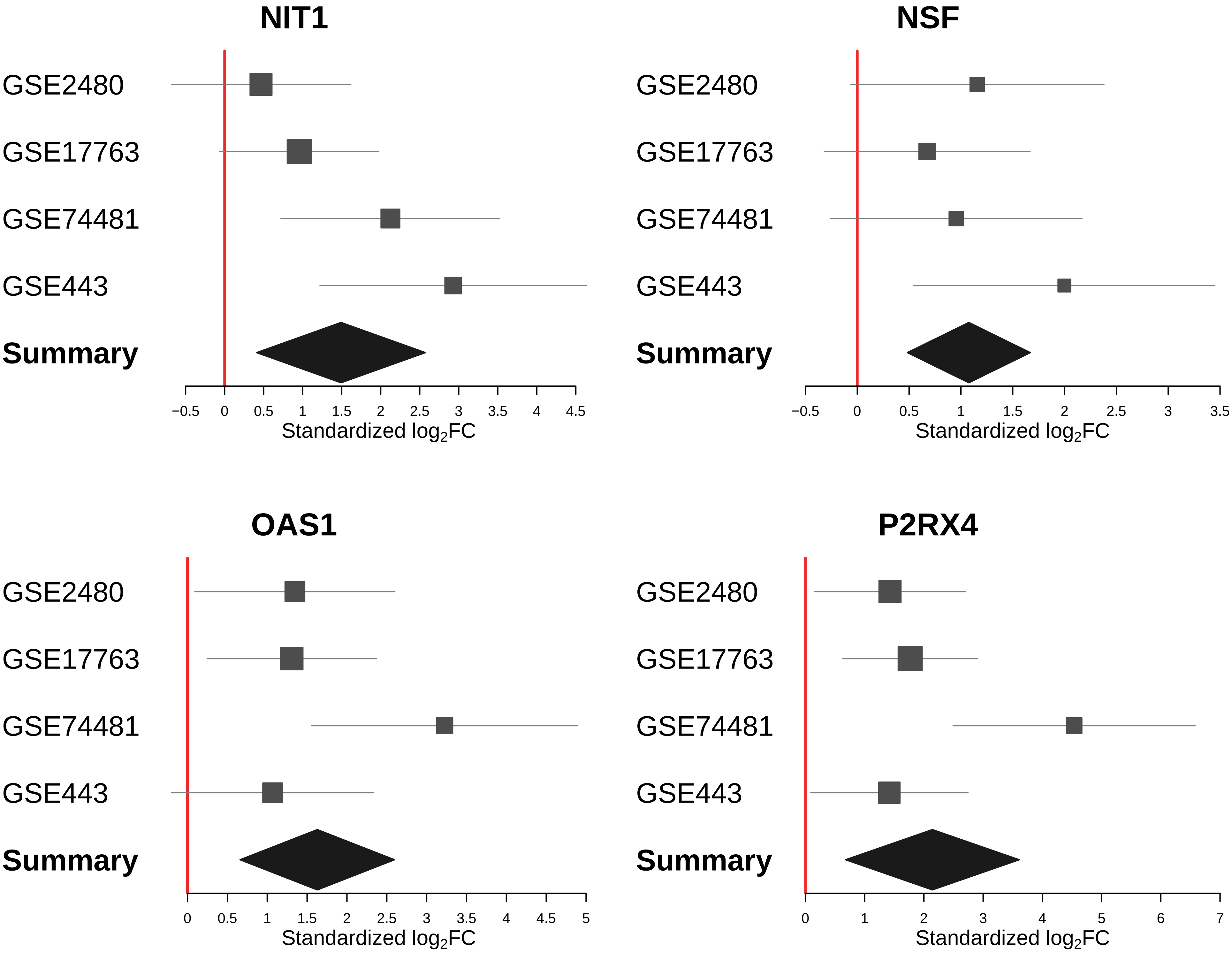
NIT1
GSE2480
GSE17763
GSE74481
GSE443
Summary
−0.5 0 0.5 1 1.5 2 2.5 3 3.5 4 4.5
Standardized log2FC
NSF
GSE2480
GSE17763
GSE74481
GSE443
Summary
−0.5 0 0.5 1 1.5 2 2.5 3 3.5
Standardized log2FC
OAS1
GSE2480
GSE17763
GSE74481
GSE443
Summary
0 0.5 1 1.5 2 2.5 3 3.5 4 4.5 5
Standardized log2FC
P2RX4
GSE2480
GSE17763
GSE74481
GSE443
Summary
0 1 2 3 4 5 6 7
Standardized log2FC

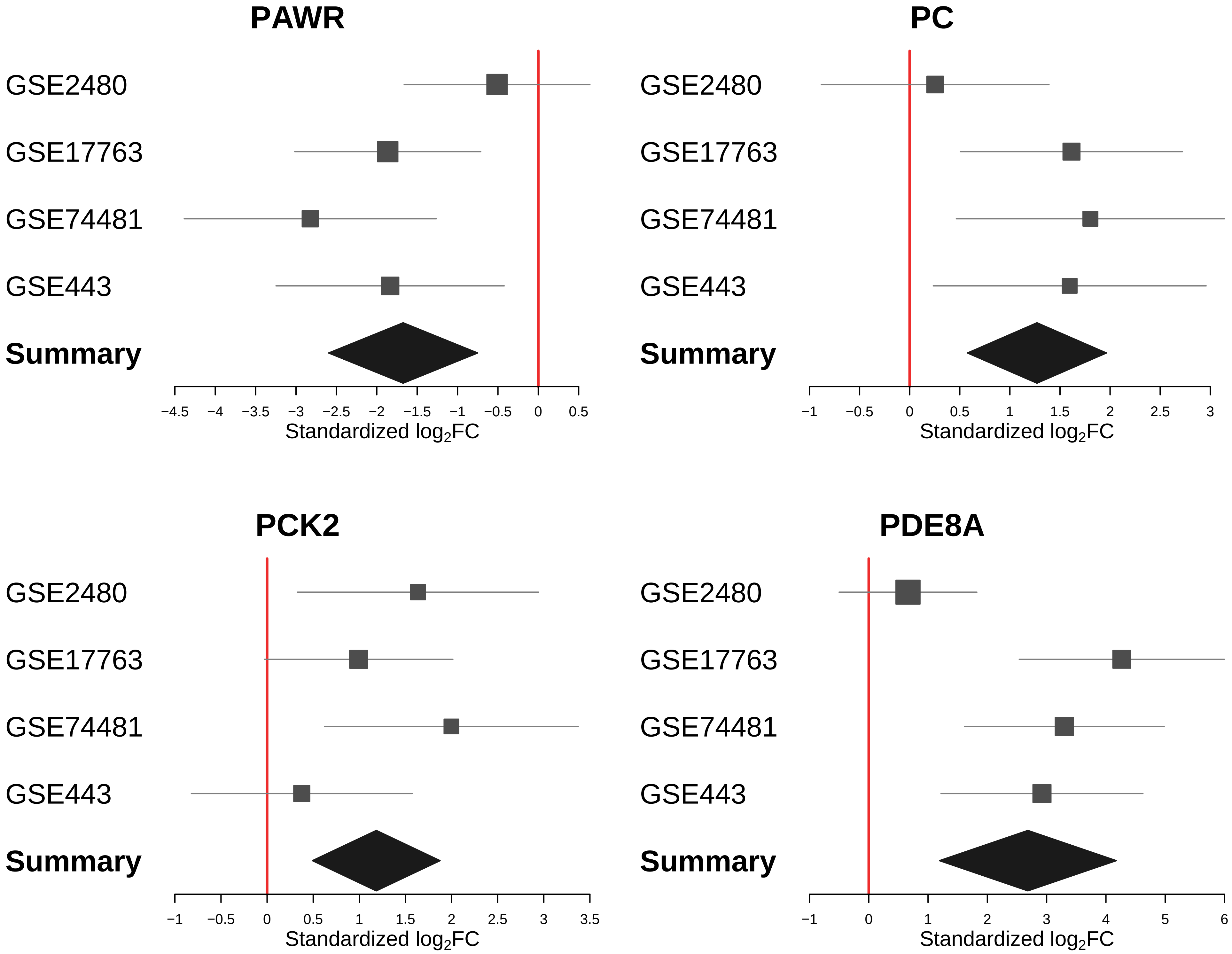
PAWR
GSE2480
GSE17763
GSE74481
GSE443
Summary
−4.5 −4 −3.5 −3 −2.5 −2 −1.5 −1 −0.5 0 0.5
Standardized log2FC
PC
GSE2480
GSE17763
GSE74481
GSE443
Summary
−1 −0.5 0 0.5 1 1.5 2 2.5 3
Standardized log2FC
PCK2
GSE2480
GSE17763
GSE74481
GSE443
Summary
−1 −0.5 0 0.5 1 1.5 2 2.5 3 3.5
Standardized log2FC
PDE8A
GSE2480
GSE17763
GSE74481
GSE443
Summary
−1 0 1 2 3 4 5 6
Standardized log2FC

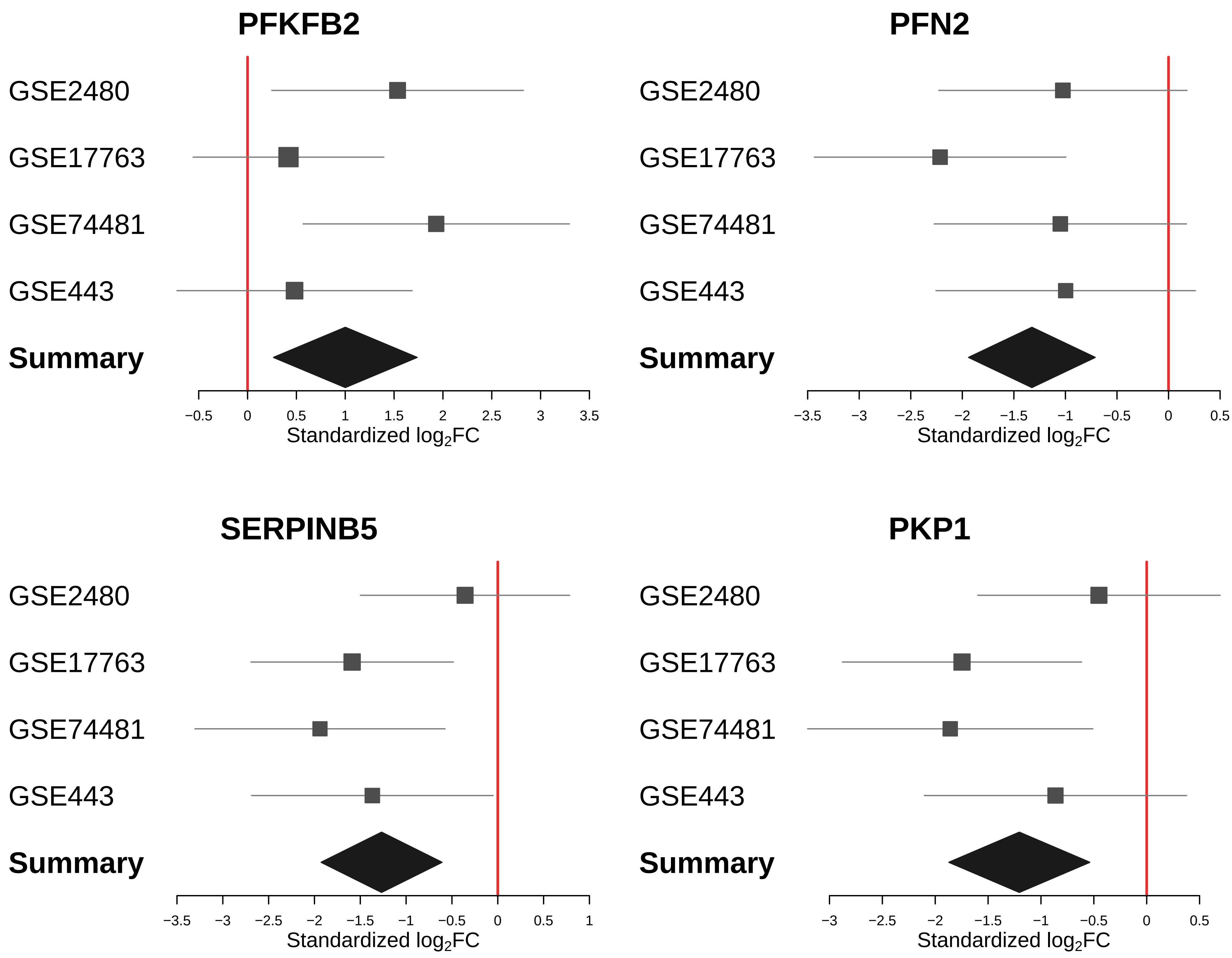
PFKFB2
GSE2480
GSE17763
GSE74481
GSE443
Summary
−0.5 0 0.5 1 1.5 2 2.5 3 3.5
Standardized $log_2FC$
PFN2
GSE2480
GSE17763
GSE74481
GSE443
Summary
−3.5 −3 −2.5 −2 −1.5 −1 −0.5 0 0.5
Standardized $log_2FC$
SERPINB5
GSE2480
GSE17763
GSE74481
GSE443
Summary
−3.5 −3 −2.5 −2 −1.5 −1 −0.5 0 0.5 1
Standardized $log_2FC$
PKP1
GSE2480
GSE17763
GSE74481
GSE443
Summary
−3 −2.5 −2 −1.5 −1 −0.5 0 0.5
Standardized $log_2FC$

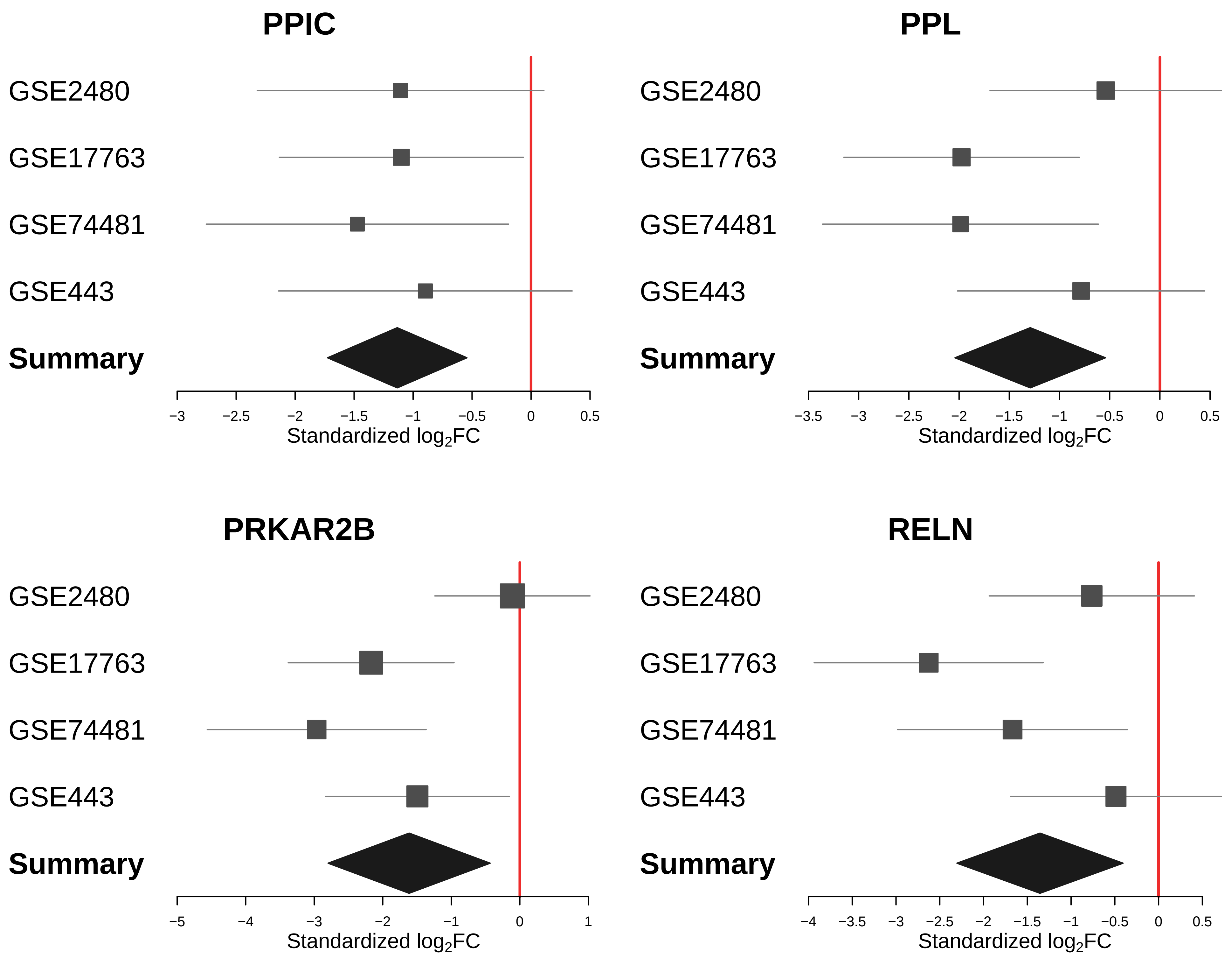
PPIC
GSE2480
GSE17763
GSE74481
GSE443
Summary
−3 −2.5 −2 −1.5 −1 −0.5 0 0.5
Standardized log2FC
PPL
GSE2480
GSE17763
GSE74481
GSE443
Summary
−3.5 −3 −2.5 −2 −1.5 −1 −0.5 0 0.5
Standardized log2FC
PRKAR2B
GSE2480
GSE17763
GSE74481
GSE443
Summary
−5 −4 −3 −2 −1 0 1
Standardized log2FC
RELN
GSE2480
GSE17763
GSE74481
GSE443
Summary
−4 −3.5 −3 −2.5 −2 −1.5 −1 −0.5 0 0.5
Standardized log2FC

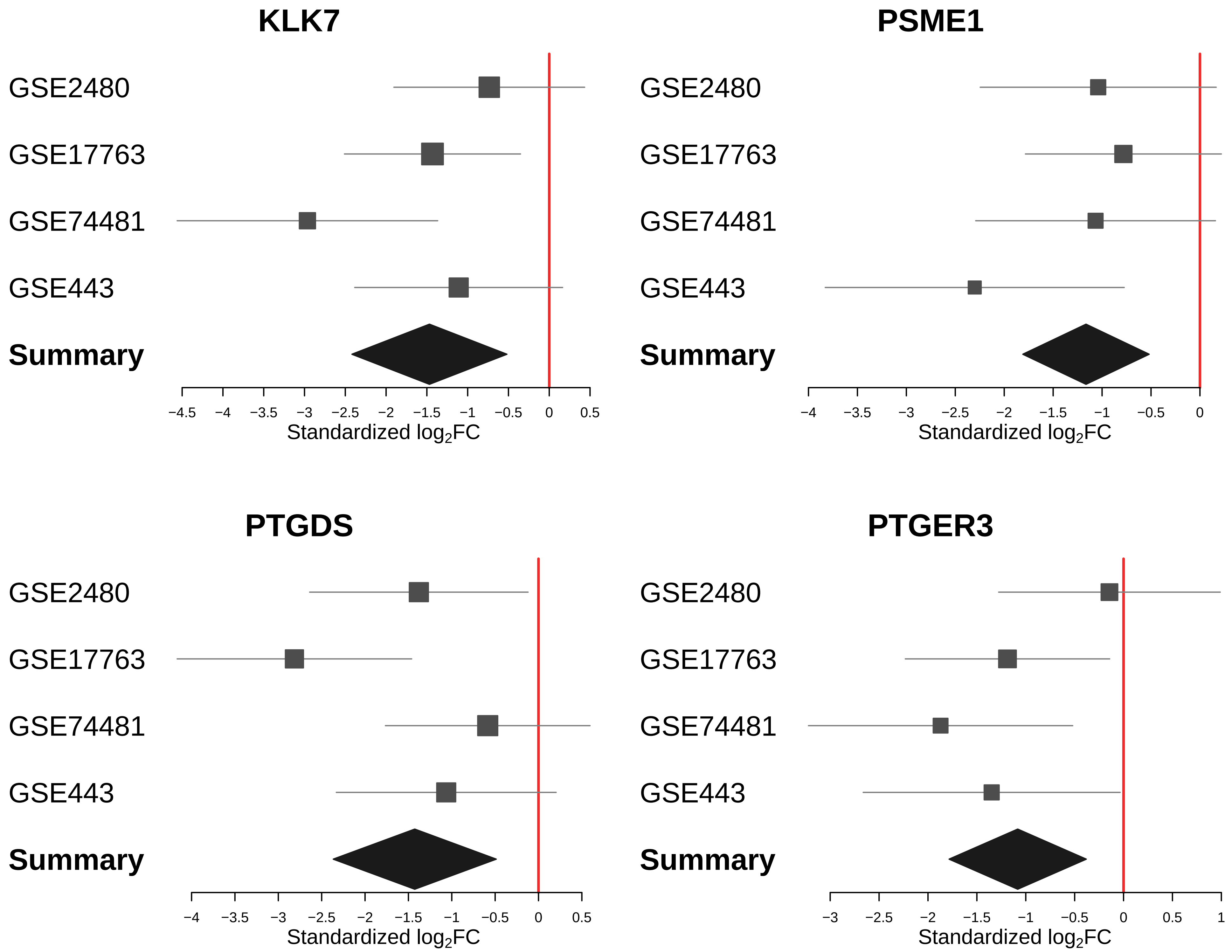

KLK7
GSE2480
GSE17763
GSE74481
GSE443
Summary
−4.5 −4 −3.5 −3 −2.5 −2 −1.5 −1 −0.5 0 0.5
Standardized $log_2FC$
PSME1
GSE2480
GSE17763
GSE74481
GSE443
Summary
−4 −3.5 −3 −2.5 −2 −1.5 −1 −0.5 0
Standardized $log_2FC$
PTGDS
GSE2480
GSE17763
GSE74481
GSE443
Summary
−4 −3.5 −3 −2.5 −2 −1.5 −1 −0.5 0 0.5
Standardized $log_2FC$
PTGER3
GSE2480
GSE17763
GSE74481
GSE443
Summary
−3 −2.5 −2 −1.5 −1 −0.5 0 0.5 1
Standardized $log_2FC$

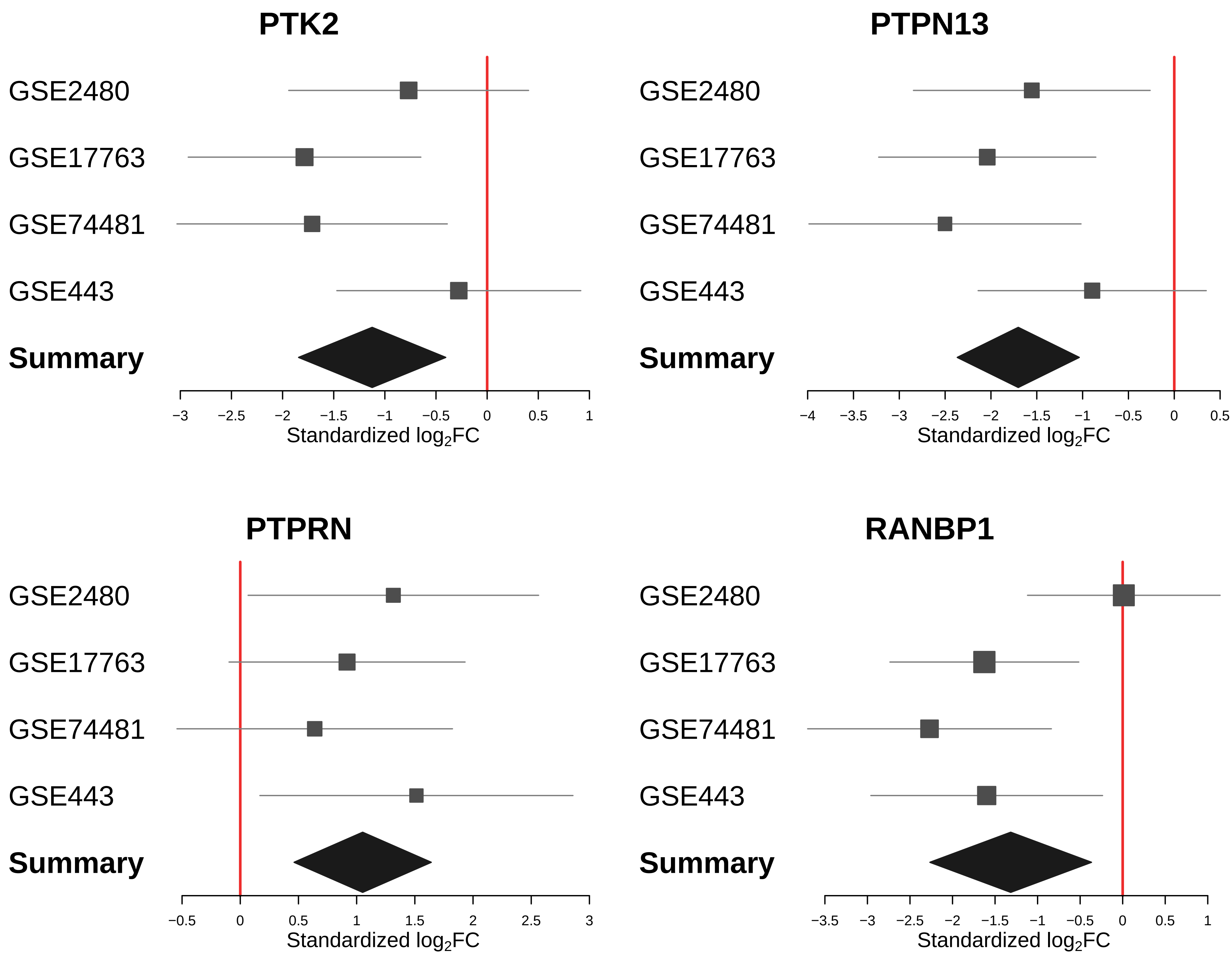

PTK2
PTPN13
PTPRN
RANBP1
GSE2480
GSE17763
GSE74481
GSE443
Summary
Standardized log2FC

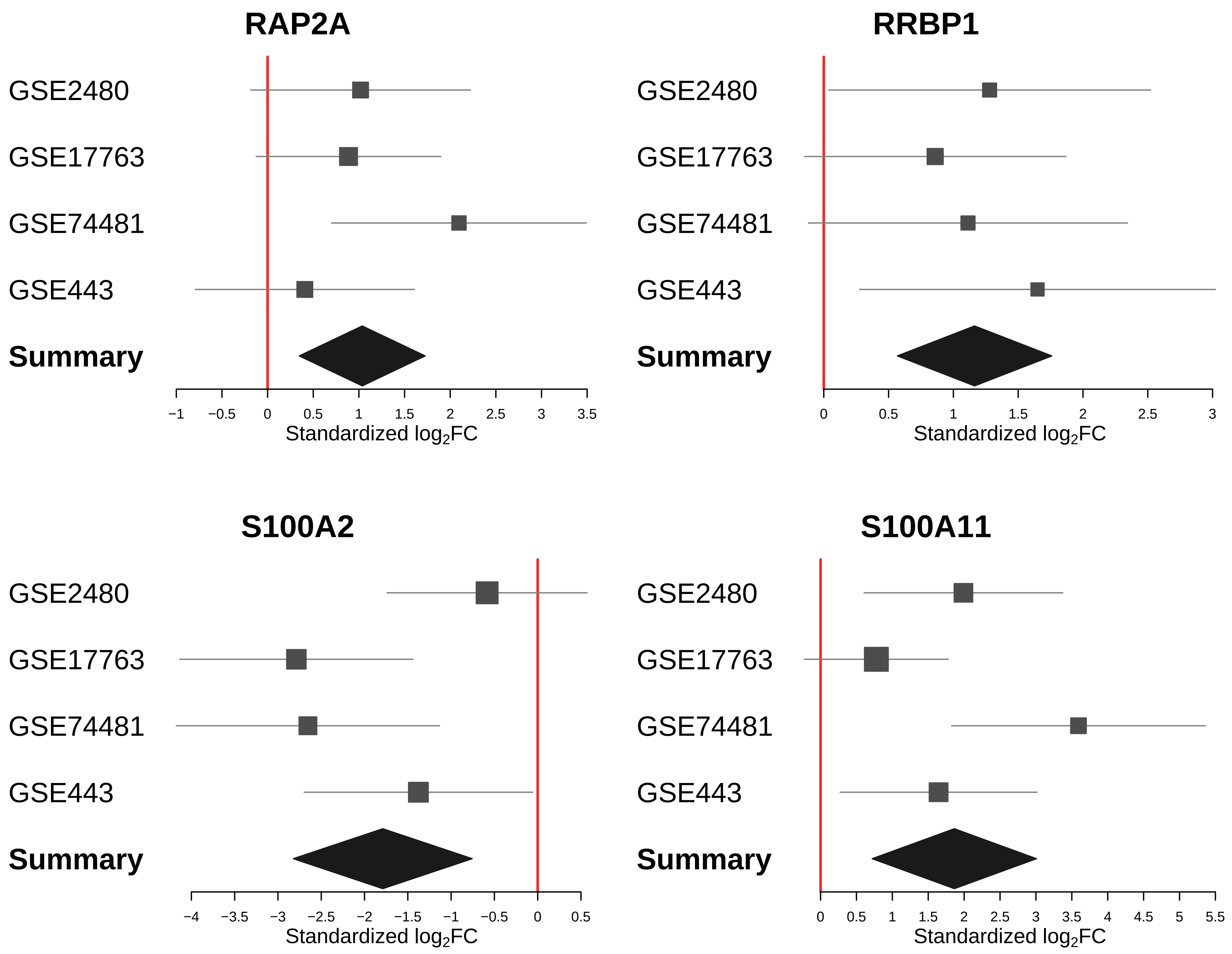

RAP2A
RRBP1
S100A2
S100A11
GSE2480
GSE17763
GSE74481
GSE443
Summary
Standardized $log_2FC$

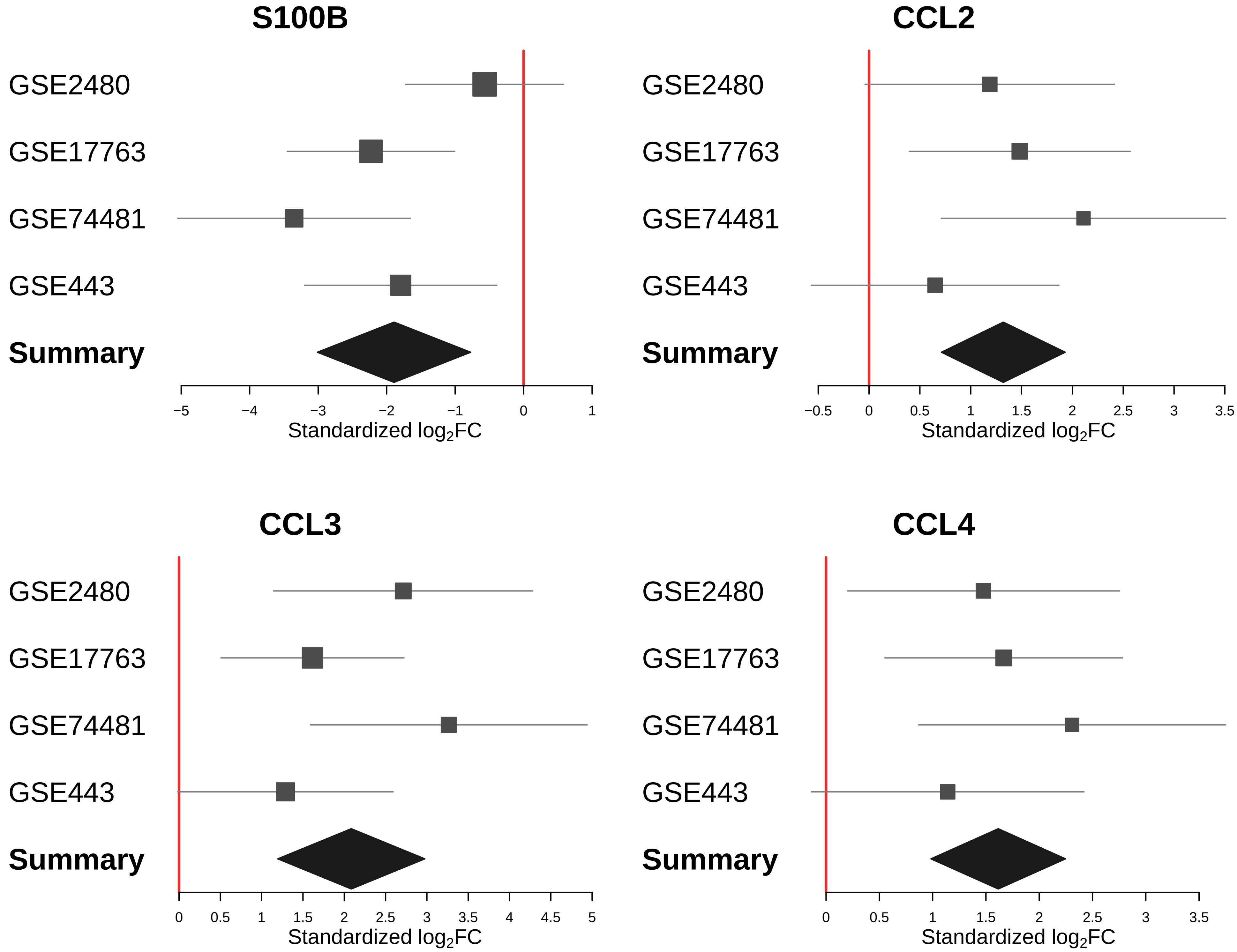

S100B
CCL2
CCL3
CCL4
GSE2480
GSE17763
GSE74481
GSE443
Summary
Standardized log2FC

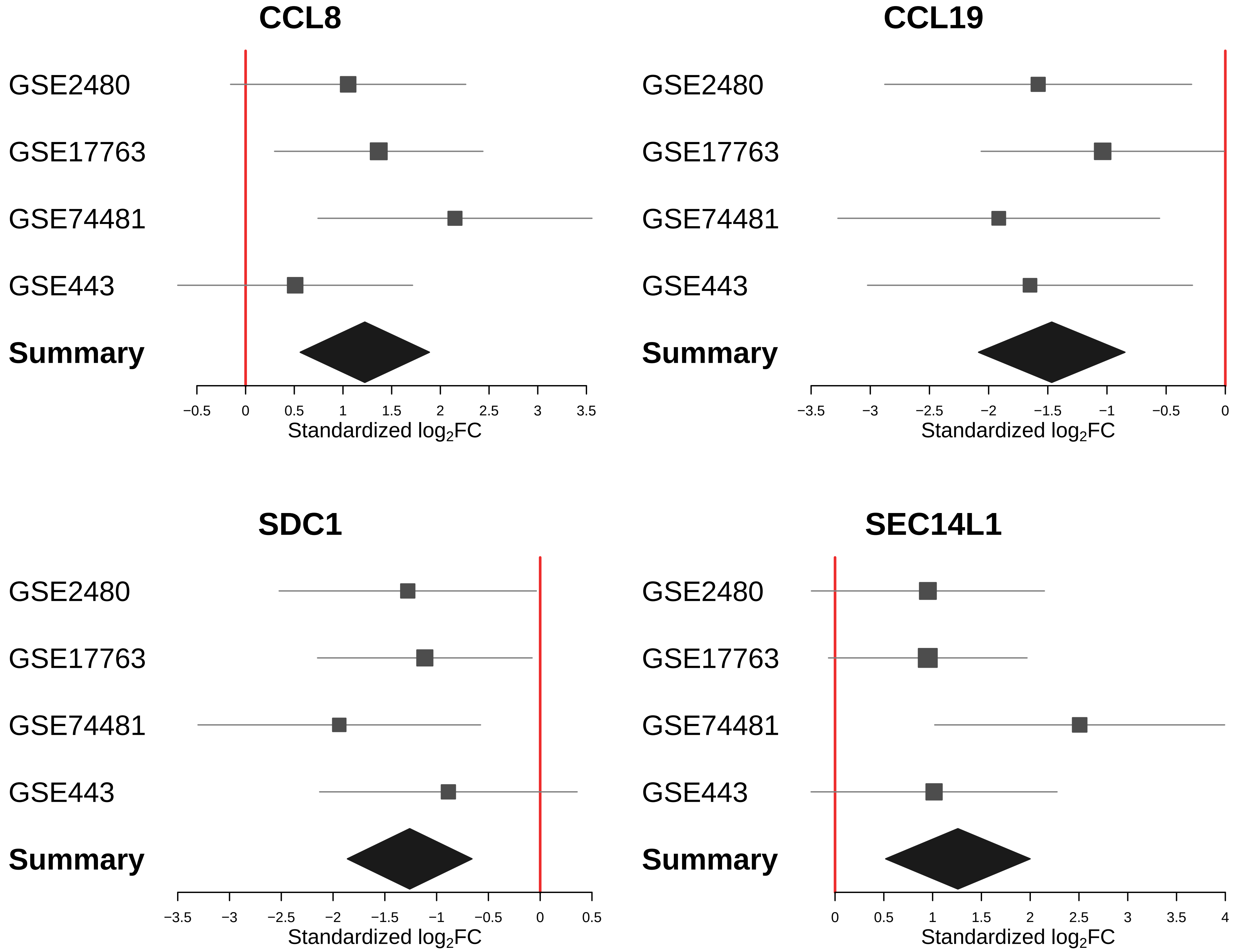

CCL8
CCL19
SDC1
SEC14L1
GSE2480
GSE17763
GSE74481
GSE443
Summary
Standardized log2FC

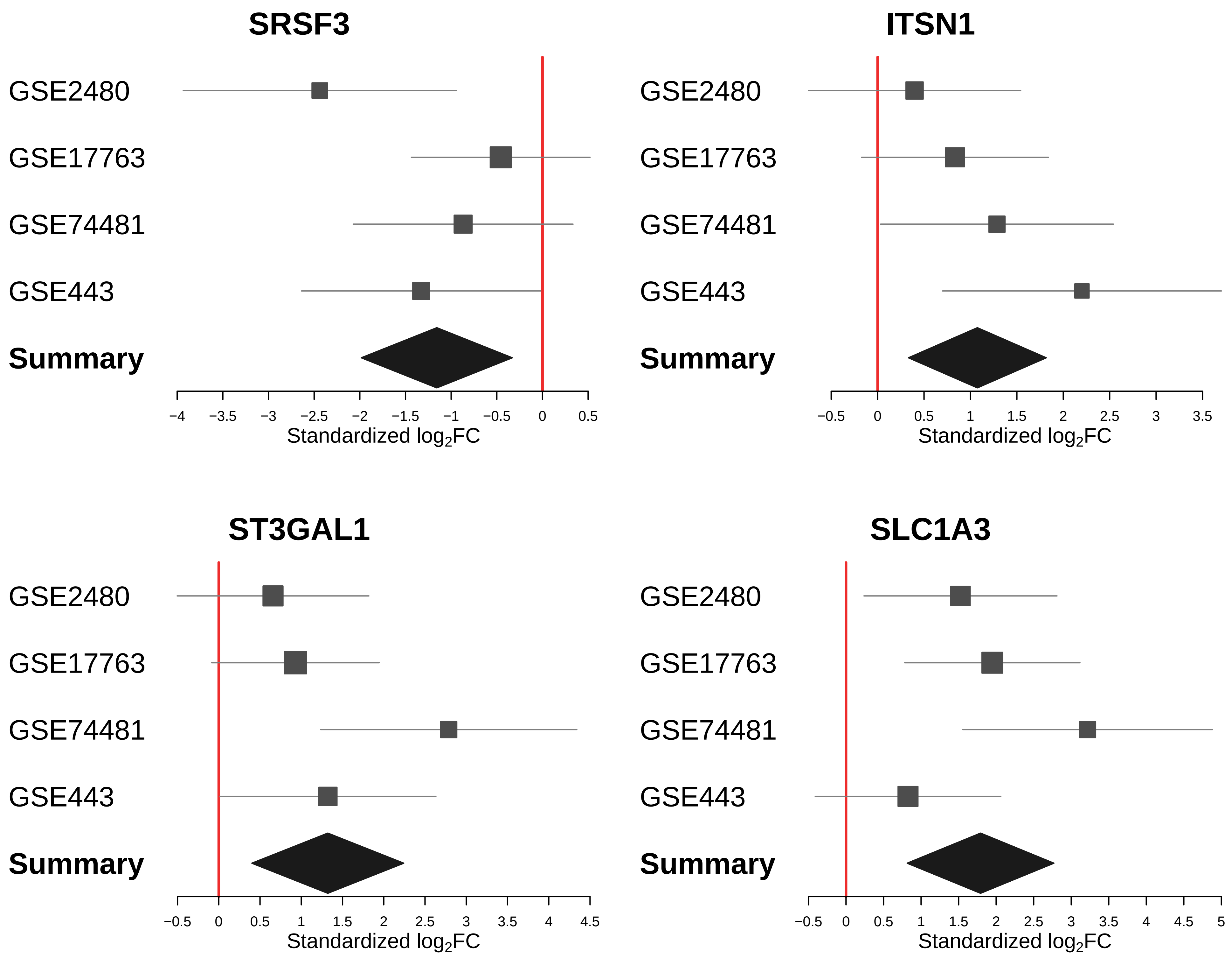
SRSF3
GSE2480
GSE17763
GSE74481
GSE443
Summary
−4 −3.5 −3 −2.5 −2 −1.5 −1 −0.5 0 0.5
Standardized log2FC
ITSN1
GSE2480
GSE17763
GSE74481
GSE443
Summary
−0.5 0 0.5 1 1.5 2 2.5 3 3.5
Standardized log2FC
ST3GAL1
GSE2480
GSE17763
GSE74481
GSE443
Summary
−0.5 0 0.5 1 1.5 2 2.5 3 3.5 4 4.5
Standardized log2FC
SLC1A3
GSE2480
GSE17763
GSE74481
GSE443
Summary
−0.5 0 0.5 1 1.5 2 2.5 3 3.5 4 4.5 5
Standardized log2FC

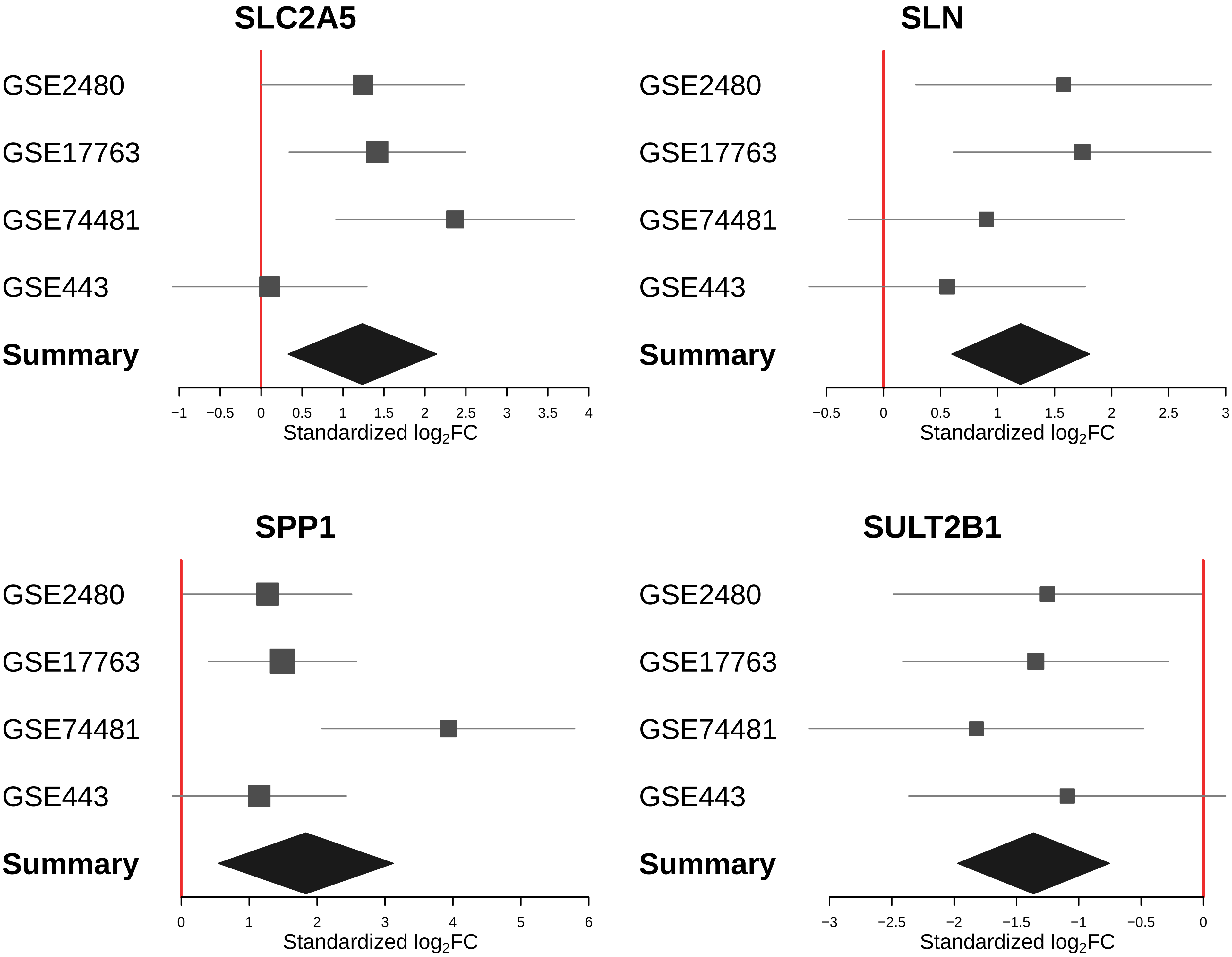
SLC2A5
GSE2480
GSE17763
GSE74481
GSE443
Summary
−1 −0.5 0 0.5 1 1.5 2 2.5 3 3.5 4
Standardized log2FC
SLN
GSE2480
GSE17763
GSE74481
GSE443
Summary
−0.5 0 0.5 1 1.5 2 2.5 3
Standardized log2FC
SPP1
GSE2480
GSE17763
GSE74481
GSE443
Summary
0 1 2 3 4 5 6
Standardized log2FC
SULT2B1
GSE2480
GSE17763
GSE74481
GSE443
Summary
−3 −2.5 −2 −1.5 −1 −0.5 0
Standardized log2FC

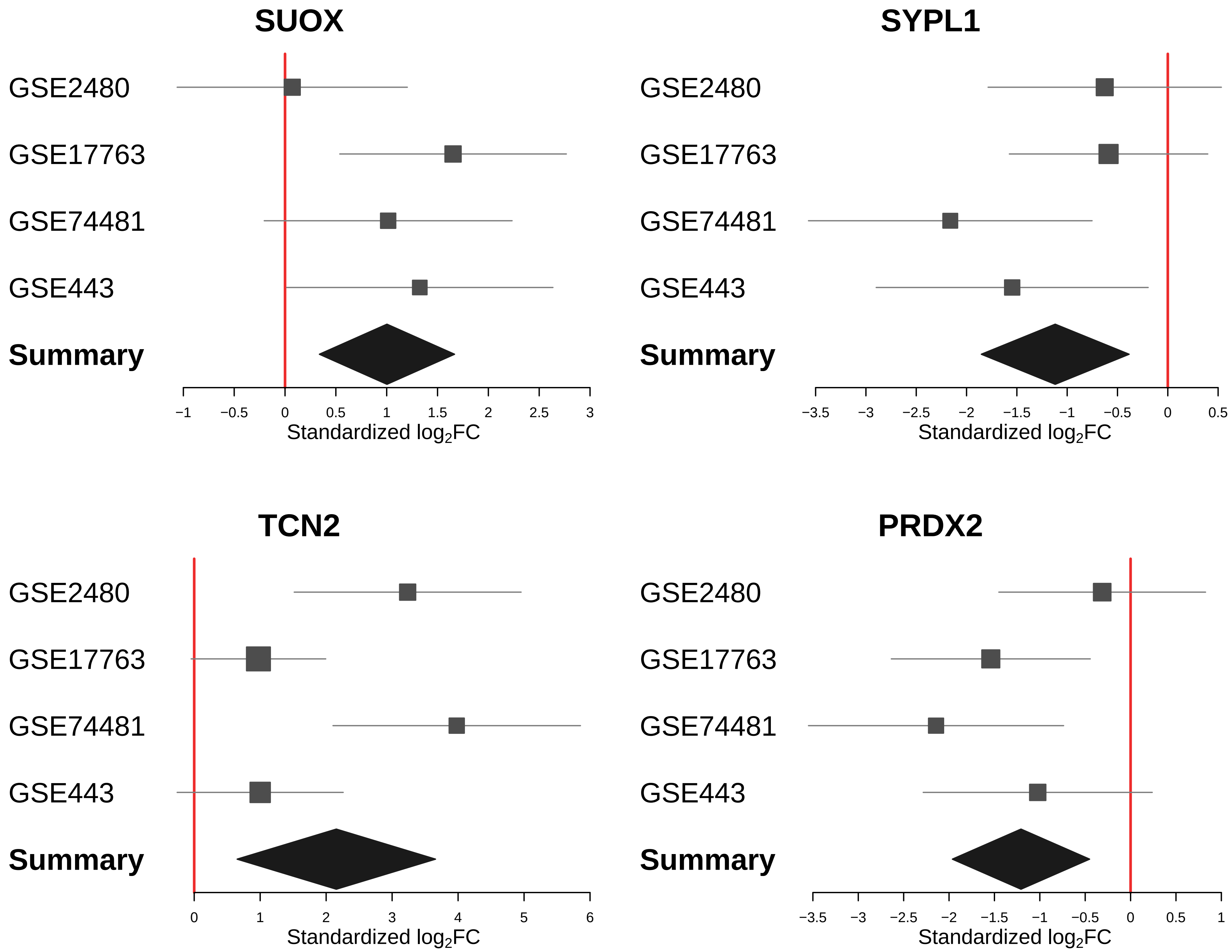
SUOX
GSE2480
GSE17763
GSE74481
GSE443
Summary
−1 −0.5 0 0.5 1 1.5 2 2.5 3
Standardized $log_2FC$
SYPL1
GSE2480
GSE17763
GSE74481
GSE443
Summary
−3.5 −3 −2.5 −2 −1.5 −1 −0.5 0 0.5
Standardized $log_2FC$
TCN2
GSE2480
GSE17763
GSE74481
GSE443
Summary
0 1 2 3 4 5 6
Standardized $log_2FC$
PRDX2
GSE2480
GSE17763
GSE74481
GSE443
Summary
−3.5 −3 −2.5 −2 −1.5 −1 −0.5 0 0.5 1
Standardized $log_2FC$

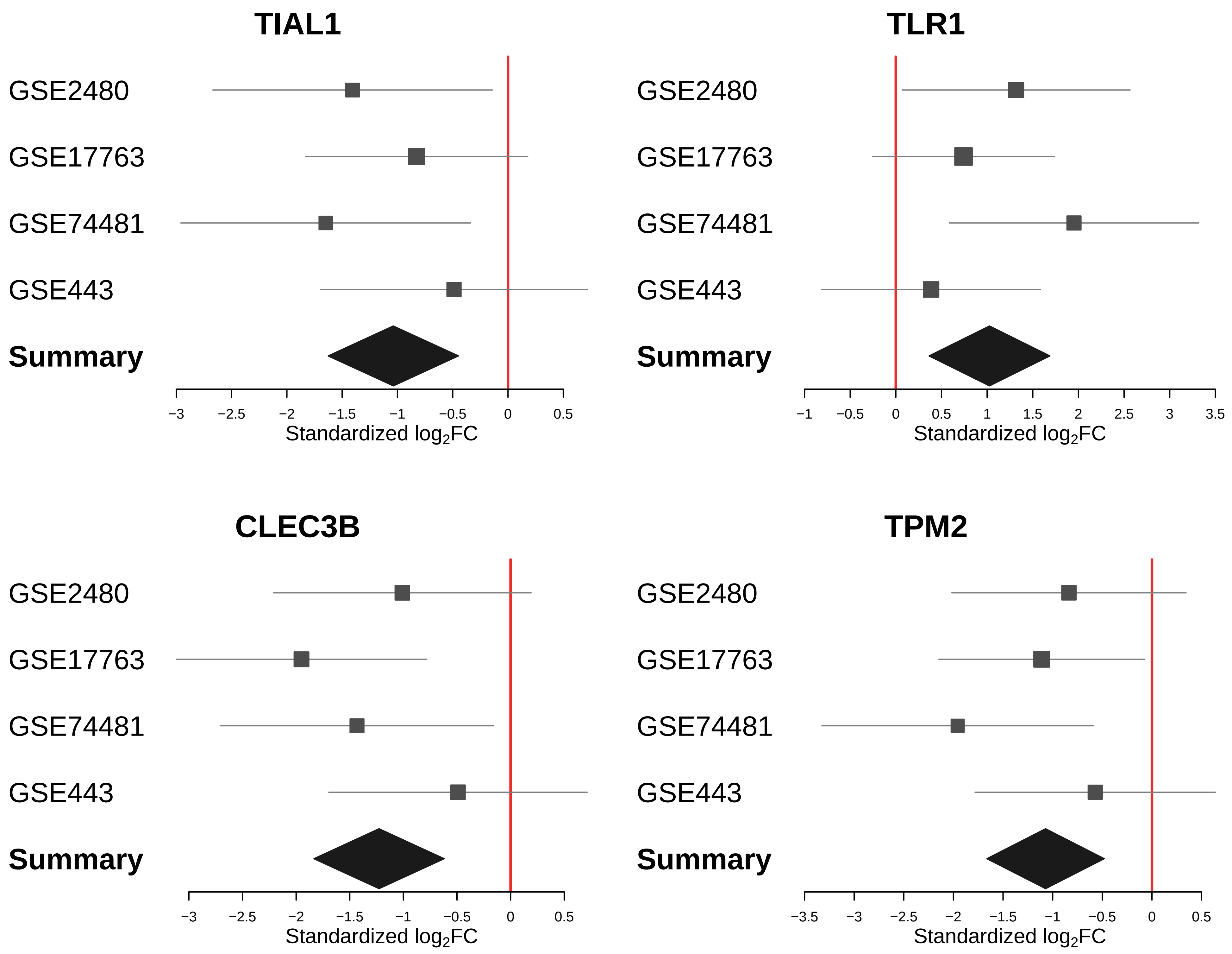
TIAL1
GSE2480
GSE17763
GSE74481
GSE443
Summary
−3
−2.5
−2
−1.5
−1
−0.5
0
0.5
Standardized log2FC
TLR1
GSE2480
GSE17763
GSE74481
GSE443
Summary
−1
−0.5
0
0.5
1
1.5
2
2.5
3
3.5
Standardized log2FC
CLEC3B
GSE2480
GSE17763
GSE74481
GSE443
Summary
−3
−2.5
−2
−1.5
−1
−0.5
0
0.5
Standardized log2FC
TPM2
GSE2480
GSE17763
GSE74481
GSE443
Summary
−3.5
−3
−2.5
−2
−1.5
−1
−0.5
0
0.5
Standardized log2FC

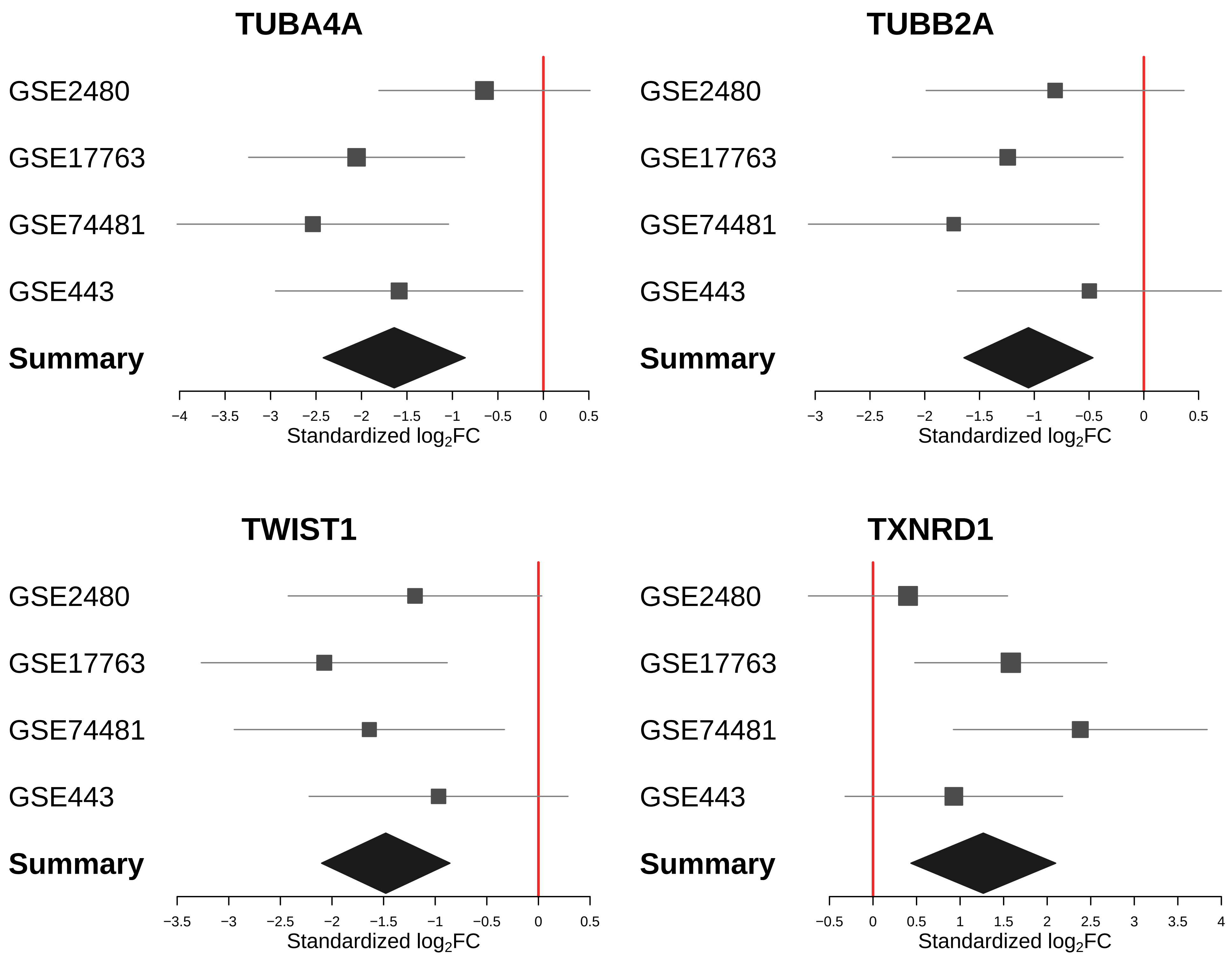

TUBA4A
GSE2480
GSE17763
GSE74481
GSE443
Summary
−4 −3.5 −3 −2.5 −2 −1.5 −1 −0.5 0 0.5
Standardized $log_2FC$
TUBB2A
GSE2480
GSE17763
GSE74481
GSE443
Summary
−3 −2.5 −2 −1.5 −1 −0.5 0 0.5
Standardized $log_2FC$
TWIST1
GSE2480
GSE17763
GSE74481
GSE443
Summary
−3.5 −3 −2.5 −2 −1.5 −1 −0.5 0 0.5
Standardized $log_2FC$
TXNRD1
GSE2480
GSE17763
GSE74481
GSE443
Summary
−0.5 0 0.5 1 1.5 2 2.5 3 3.5 4
Standardized $log_2FC$

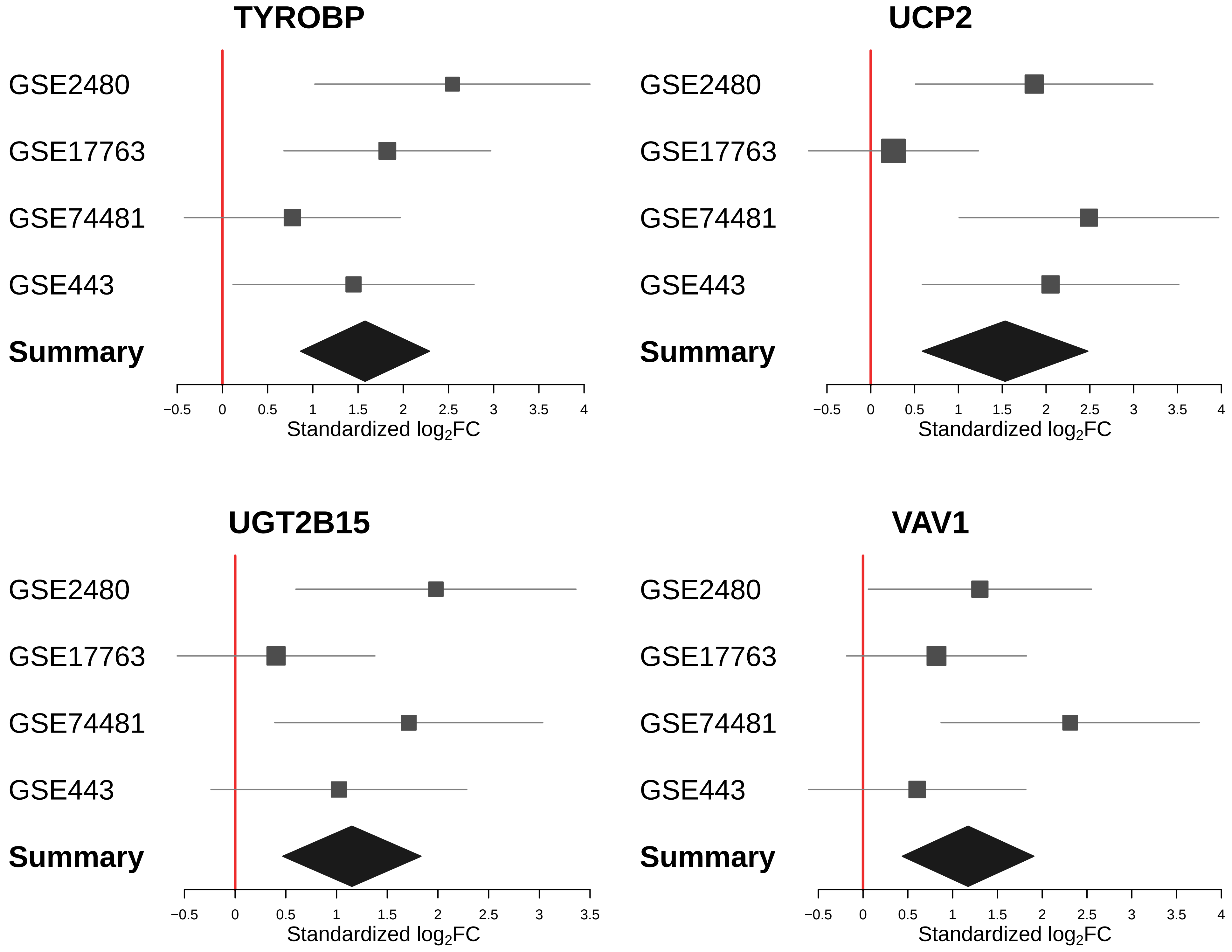
TYROBP
UCP2
UGT2B15
VAV1
GSE2480
GSE17763
GSE74481
GSE443
Summary
−0.5 0 0.5 1 1.5 2 2.5 3 3.5 4
−0.5 0 0.5 1 1.5 2 2.5 3 3.5
Standardized log2FC

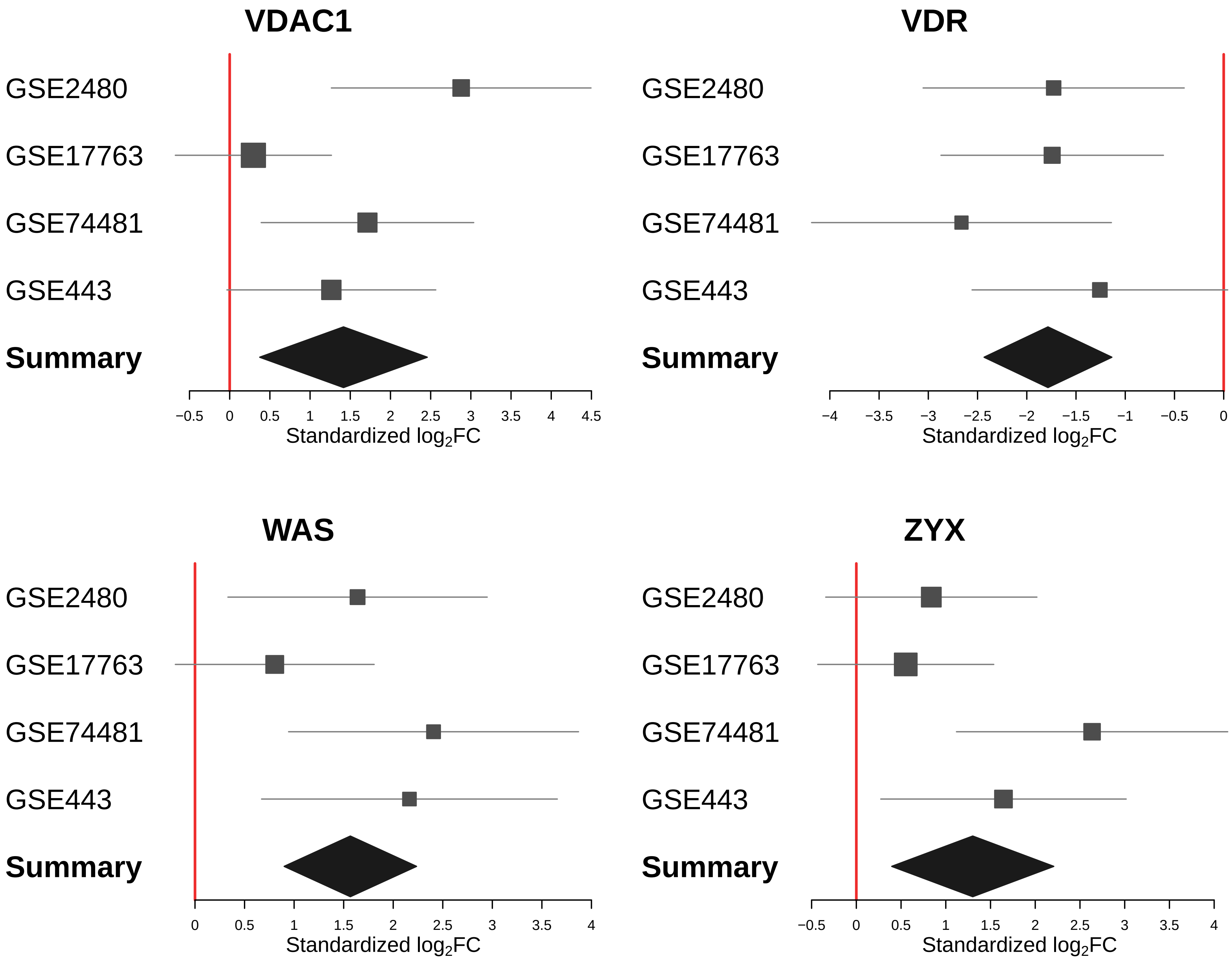
VDAC1
GSE2480
GSE17763
GSE74481
GSE443
Summary
−0.5 0 0.5 1 1.5 2 2.5 3 3.5 4 4.5
Standardized log2FC
VDR
GSE2480
GSE17763
GSE74481
GSE443
Summary
−4 −3.5 −3 −2.5 −2 −1.5 −1 −0.5 0
Standardized log2FC
WAS
GSE2480
GSE17763
GSE74481
GSE443
Summary
0 0.5 1 1.5 2 2.5 3 3.5 4
Standardized log2FC
ZYX
GSE2480
GSE17763
GSE74481
GSE443
Summary
−0.5 0 0.5 1 1.5 2 2.5 3 3.5 4
Standardized log2FC

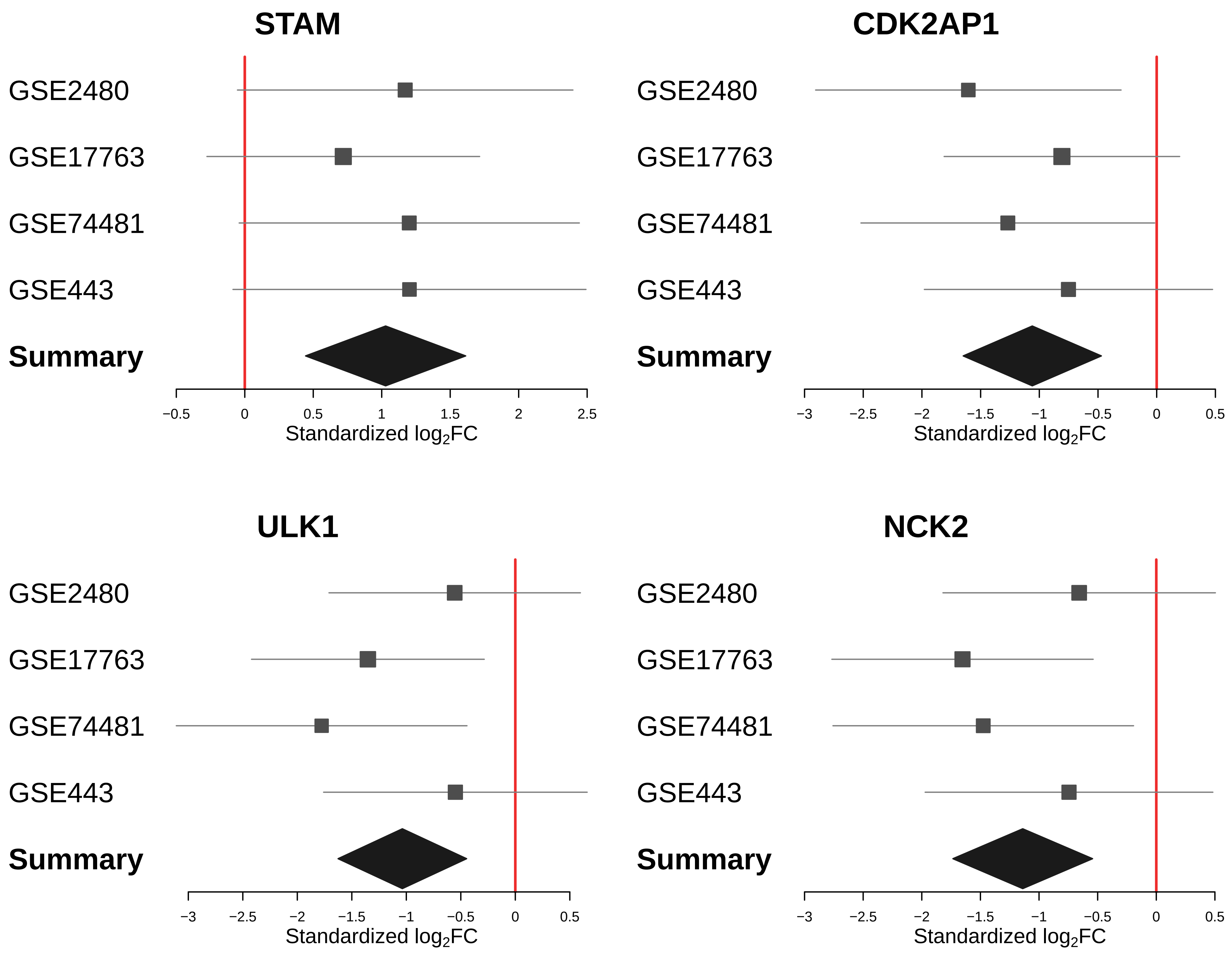

STAM
CDK2AP1
ULK1
NCK2
GSE2480
GSE17763
GSE74481
GSE443
Summary
Standardized log2FC
−0.5
0
0.5
1
1.5
2
2.5
−3
−2.5
−2
−1.5
−1
−0.5
0
0.5

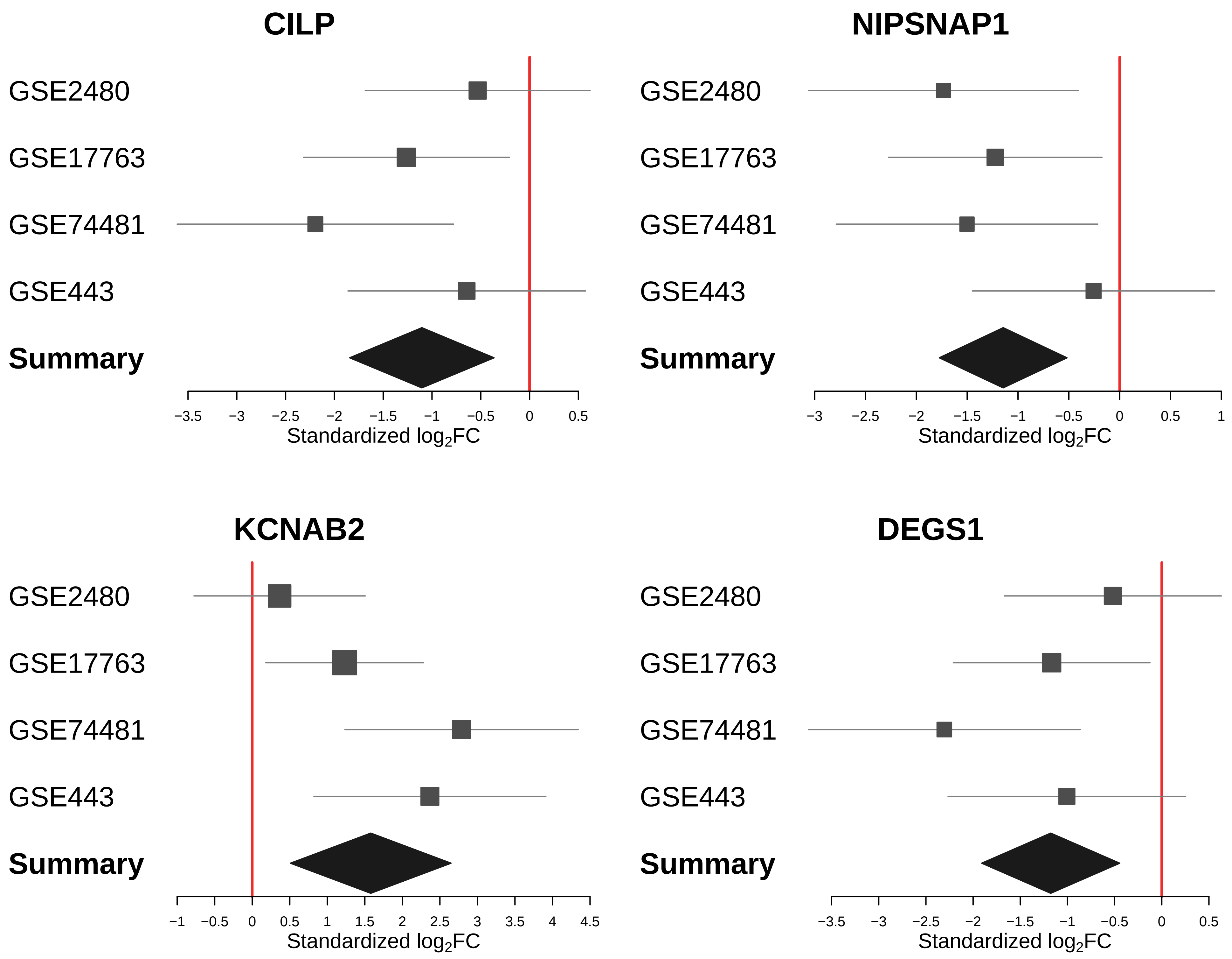
CILP
GSE2480
GSE17763
GSE74481
GSE443
Summary
−3.5 −3 −2.5 −2 −1.5 −1 −0.5 0 0.5
Standardized log2FC
NIPSNAP1
GSE2480
GSE17763
GSE74481
GSE443
Summary
−3 −2.5 −2 −1.5 −1 −0.5 0 0.5 1
Standardized log2FC
KCNAB2
GSE2480
GSE17763
GSE74481
GSE443
Summary
−1 −0.5 0 0.5 1 1.5 2 2.5 3 3.5 4 4.5
Standardized log2FC
DEGS1
GSE2480
GSE17763
GSE74481
GSE443
Summary
−3.5 −3 −2.5 −2 −1.5 −1 −0.5 0 0.5
Standardized log2FC

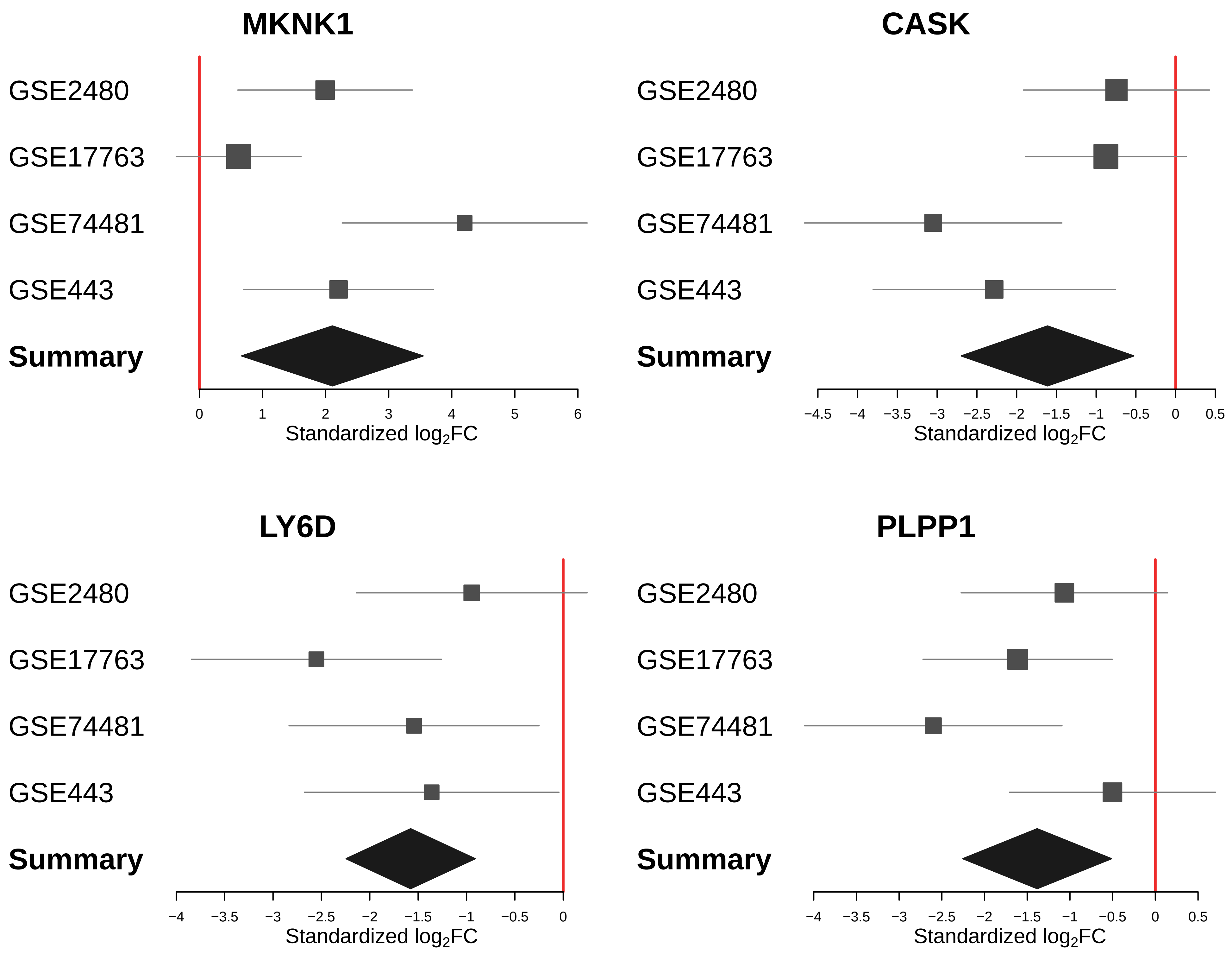
MKNK1
GSE2480
GSE17763
GSE74481
GSE443
Summary
0 1 2 3 4 5 6
Standardized log2FC
CASK
GSE2480
GSE17763
GSE74481
GSE443
Summary
−4.5 −4 −3.5 −3 −2.5 −2 −1.5 −1 −0.5 0 0.5
Standardized log2FC
LY6D
GSE2480
GSE17763
GSE74481
GSE443
Summary
−4 −3.5 −3 −2.5 −2 −1.5 −1 −0.5 0
Standardized log2FC
PLPP1
GSE2480
GSE17763
GSE74481
GSE443
Summary
−4 −3.5 −3 −2.5 −2 −1.5 −1 −0.5 0 0.5
Standardized log2FC

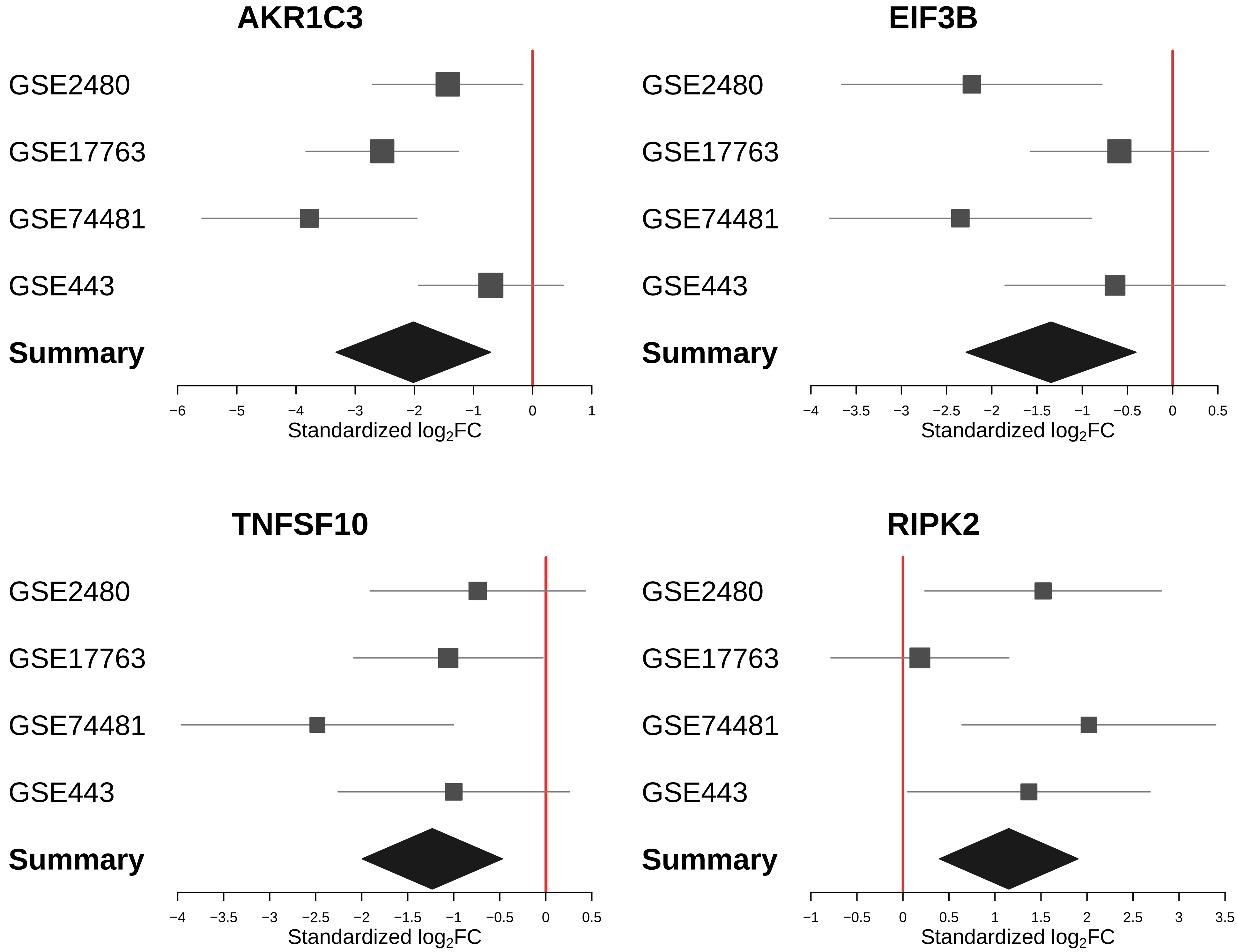
AKR1C3
EIF3B
TNFSF10
RIPK2
GSE2480
GSE17763
GSE74481
GSE443
Summary
Standardized log2FC
−6 −5 −4 −3 −2 −1 0 1
−4 −3.5 −3 −2.5 −2 −1.5 −1 −0.5 0 0.5
−1 −0.5 0 0.5 1 1.5 2 2.5 3 3.5

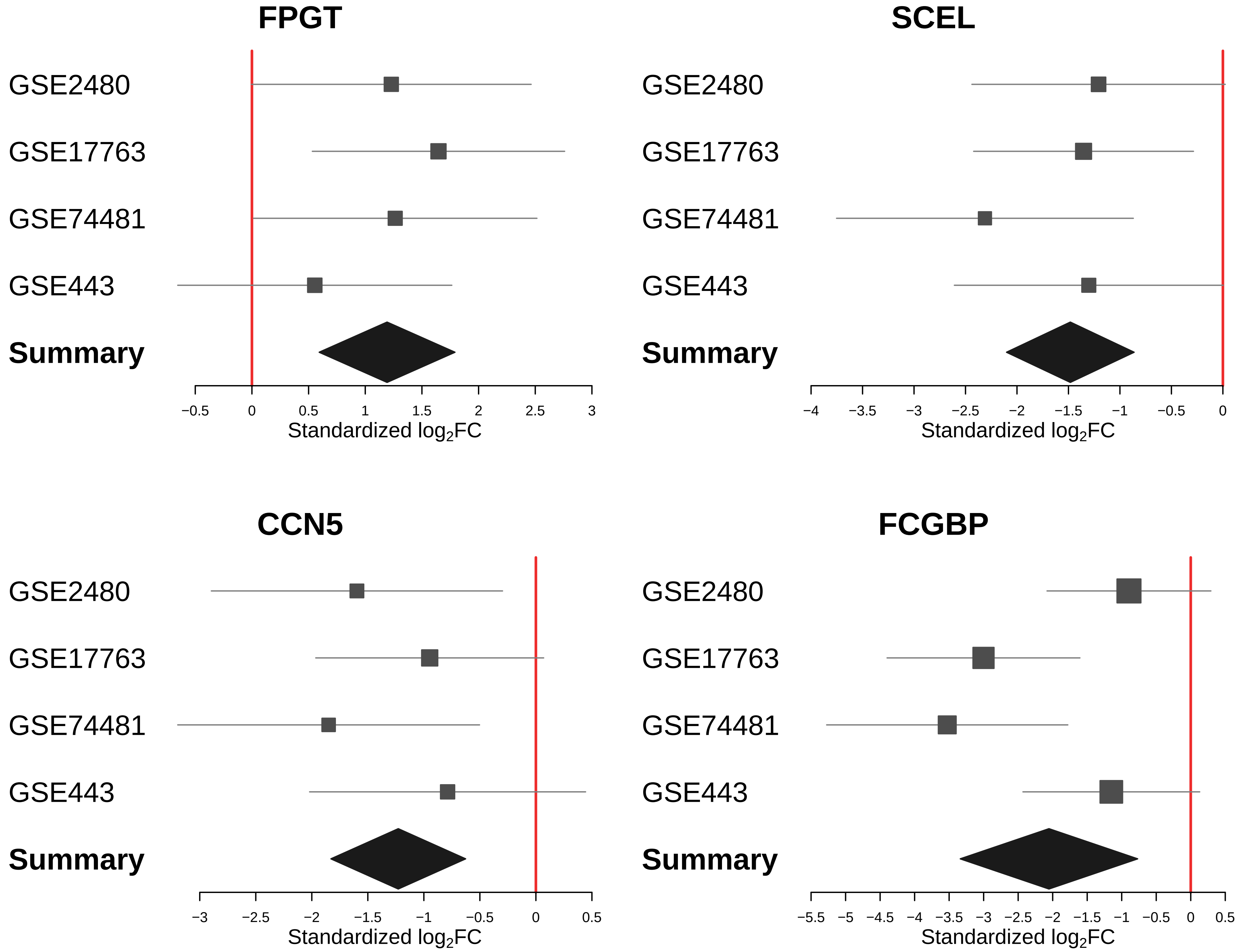
FPGT
SCEL
CCN5
FCGBP
GSE2480
GSE17763
GSE74481
GSE443
Summary
Standardized log2FC

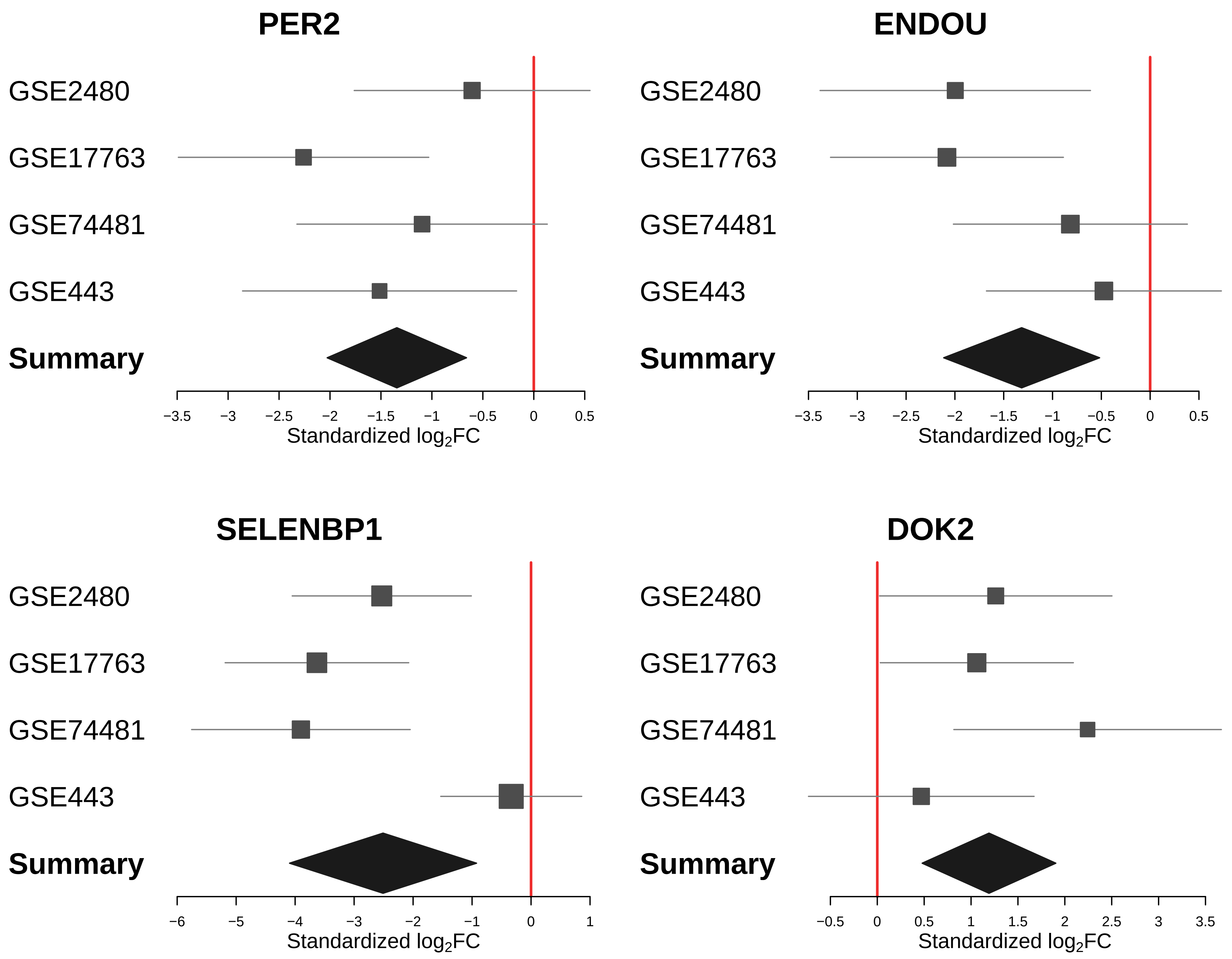

PER2
GSE2480
GSE17763
GSE74481
GSE443
Summary
−3.5 −3 −2.5 −2 −1.5 −1 −0.5 0 0.5
Standardized log2FC
ENDOU
GSE2480
GSE17763
GSE74481
GSE443
Summary
−3.5 −3 −2.5 −2 −1.5 −1 −0.5 0 0.5
Standardized log2FC
SELENBP1
GSE2480
GSE17763
GSE74481
GSE443
Summary
−6 −5 −4 −3 −2 −1 0 1
Standardized log2FC
DOK2
GSE2480
GSE17763
GSE74481
GSE443
Summary
−0.5 0 0.5 1 1.5 2 2.5 3 3.5
Standardized log2FC

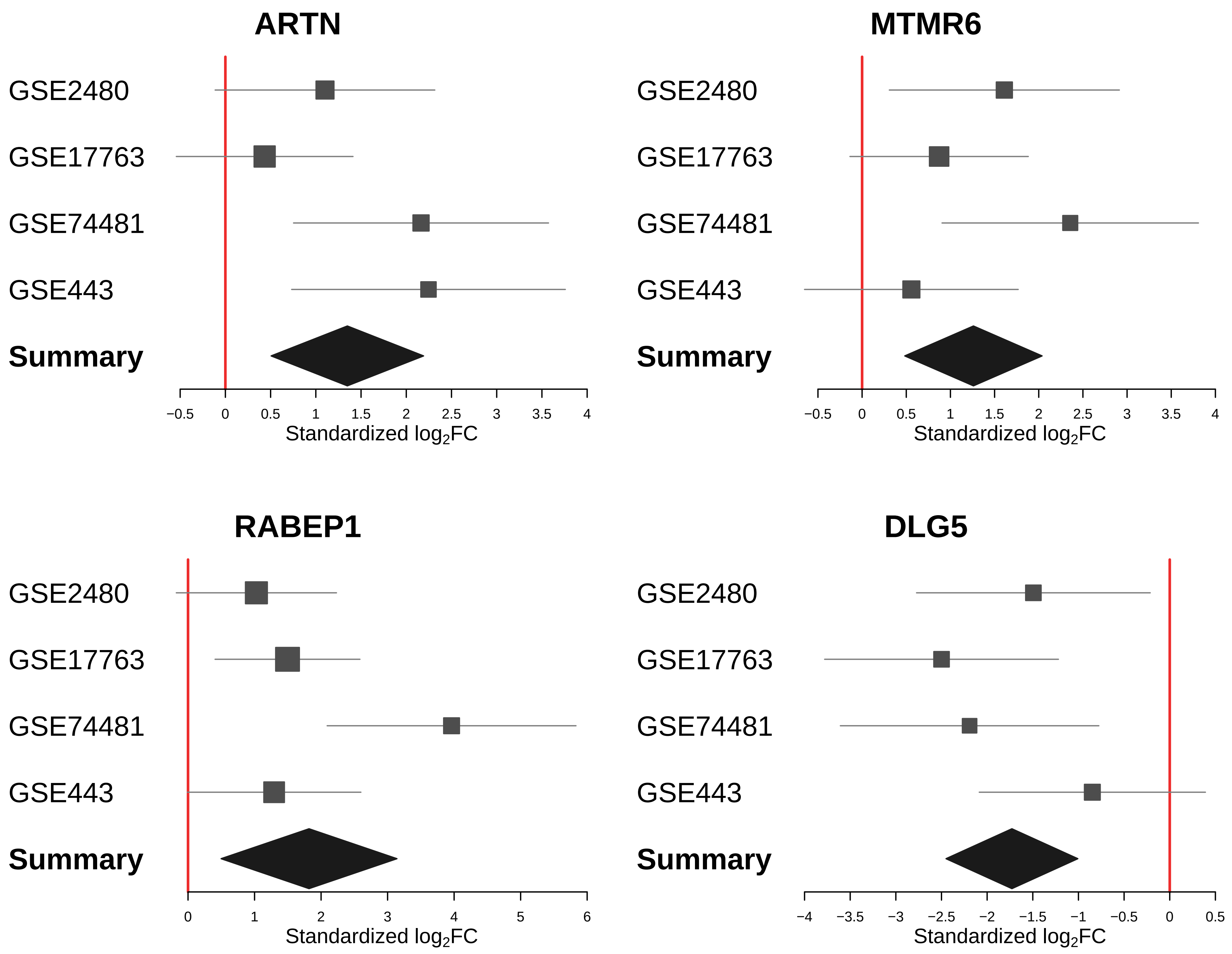
ARTN
MTMR6
RABEP1
DLG5
GSE2480
GSE17763
GSE74481
GSE443
Summary
Standardized log2FC
−0.5 0 0.5 1 1.5 2 2.5 3 3.5 4
0 1 2 3 4 5 6
−4 −3.5 −3 −2.5 −2 −1.5 −1 −0.5 0 0.5

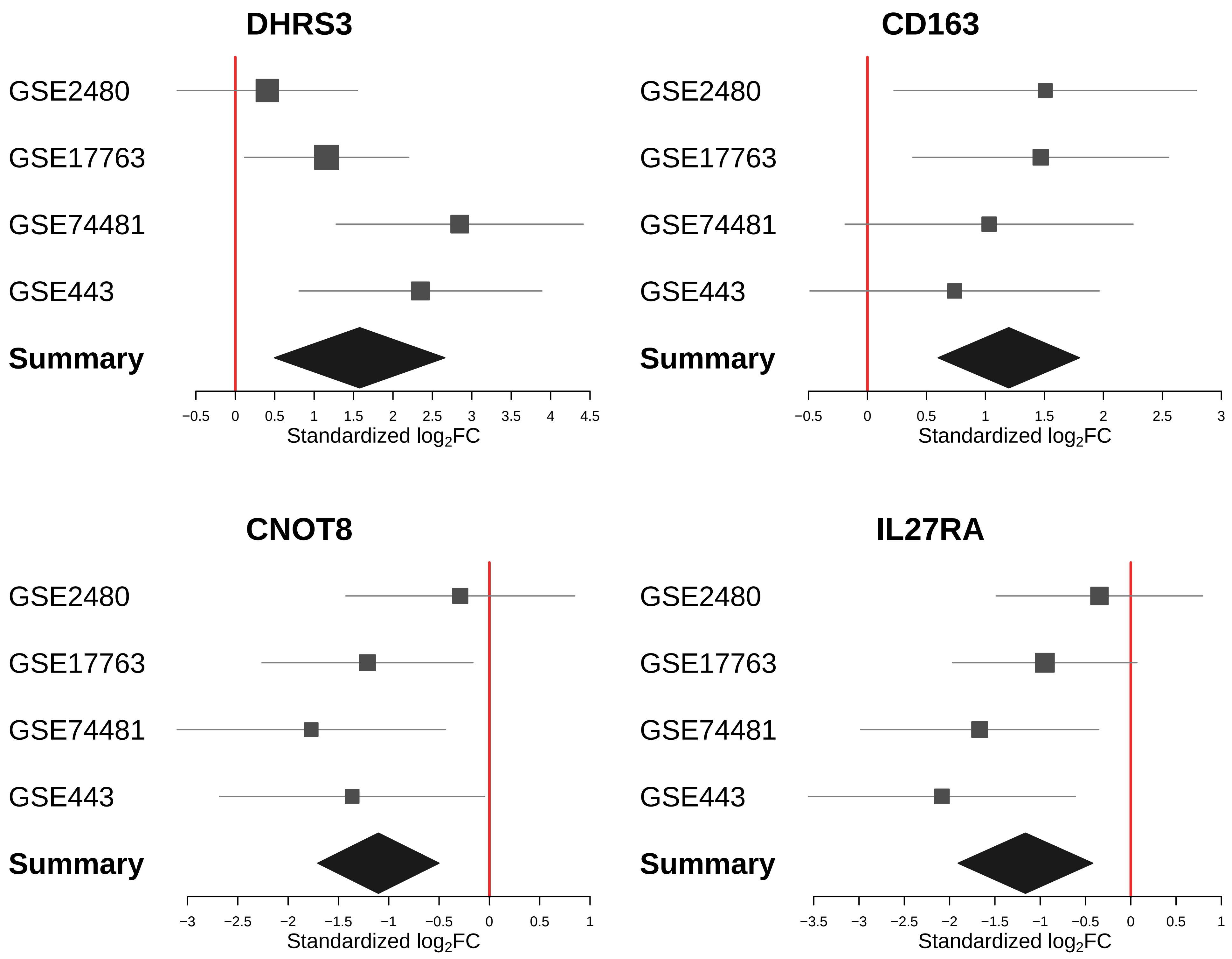
DHRS3
CD163
CNOT8
IL27RA
GSE2480
GSE17763
GSE74481
GSE443
Summary
Standardized log2FC
−0.5
0
0.5
1
1.5
2
2.5
3
3.5
4
4.5
−3.5
−3
−2.5
−2
−1.5
−1
−0.5

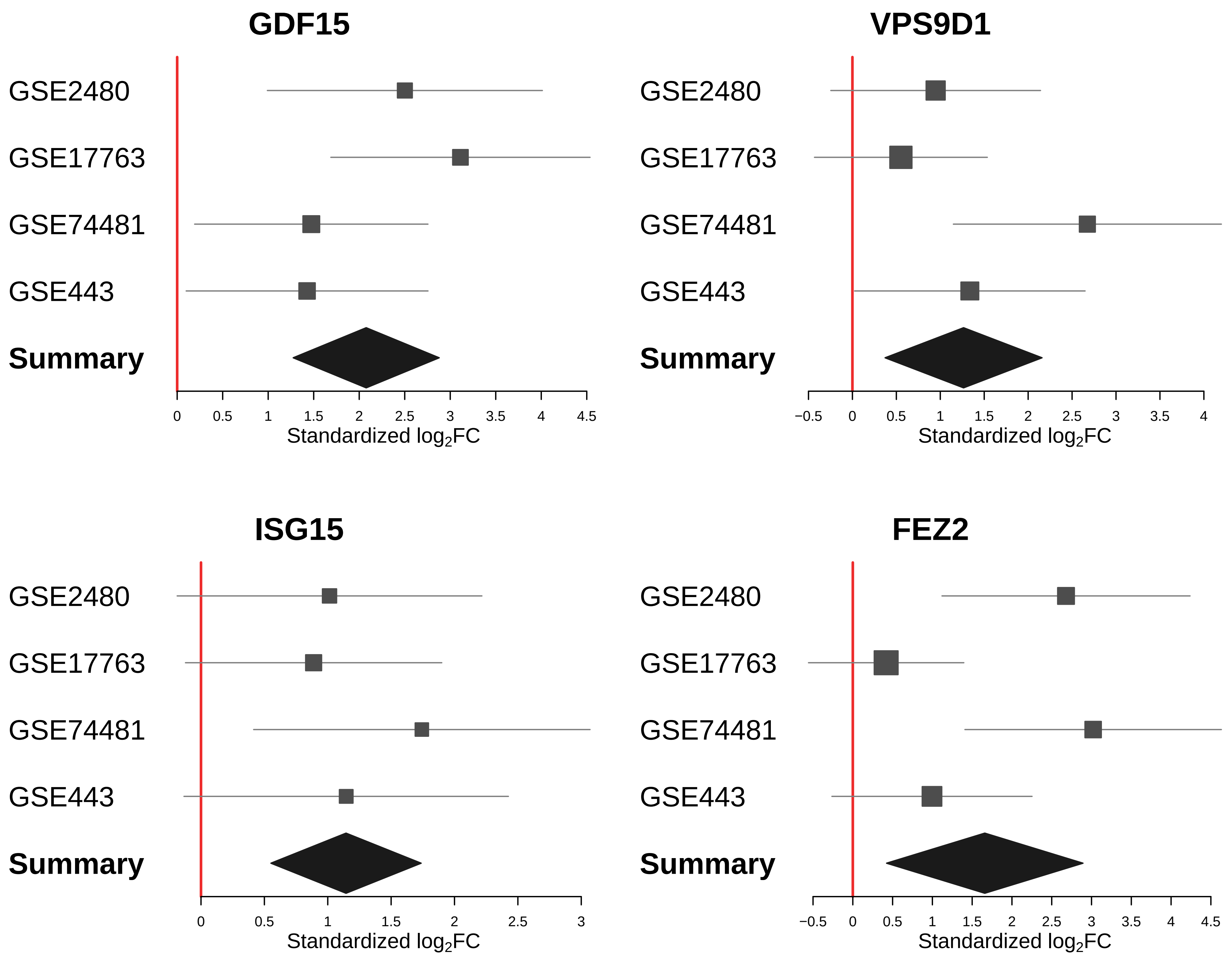
GDF15
GSE2480
GSE17763
GSE74481
GSE443
Summary
0 0.5 1 1.5 2 2.5 3 3.5 4 4.5
Standardized log2FC
VPS9D1
GSE2480
GSE17763
GSE74481
GSE443
Summary
−0.5 0 0.5 1 1.5 2 2.5 3 3.5 4
Standardized log2FC
ISG15
GSE2480
GSE17763
GSE74481
GSE443
Summary
0 0.5 1 1.5 2 2.5 3
Standardized log2FC
FEZ2
GSE2480
GSE17763
GSE74481
GSE443
Summary
−0.5 0 0.5 1 1.5 2 2.5 3 3.5 4 4.5
Standardized log2FC

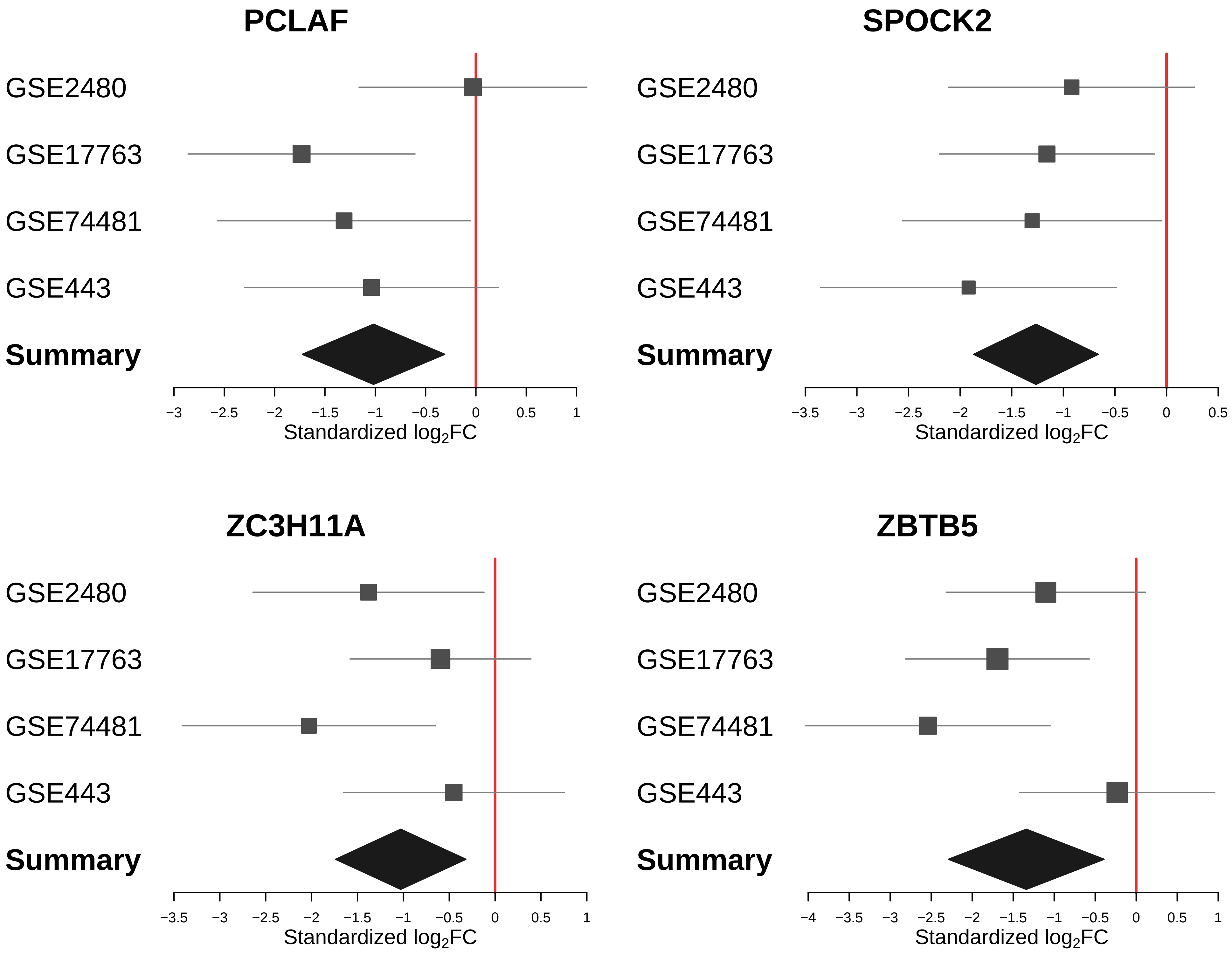
PCLAF
GSE2480
GSE17763
GSE74481
GSE443
Summary
−3 −2.5 −2 −1.5 −1 −0.5 0 0.5 1
Standardized log2FC
SPOCK2
GSE2480
GSE17763
GSE74481
GSE443
Summary
−3.5 −3 −2.5 −2 −1.5 −1 −0.5 0 0.5
Standardized log2FC
ZC3H11A
GSE2480
GSE17763
GSE74481
GSE443
Summary
−3.5 −3 −2.5 −2 −1.5 −1 −0.5 0 0.5 1
Standardized log2FC
ZBTB5
GSE2480
GSE17763
GSE74481
GSE443
Summary
−4 −3.5 −3 −2.5 −2 −1.5 −1 −0.5 0 0.5 1
Standardized log2FC

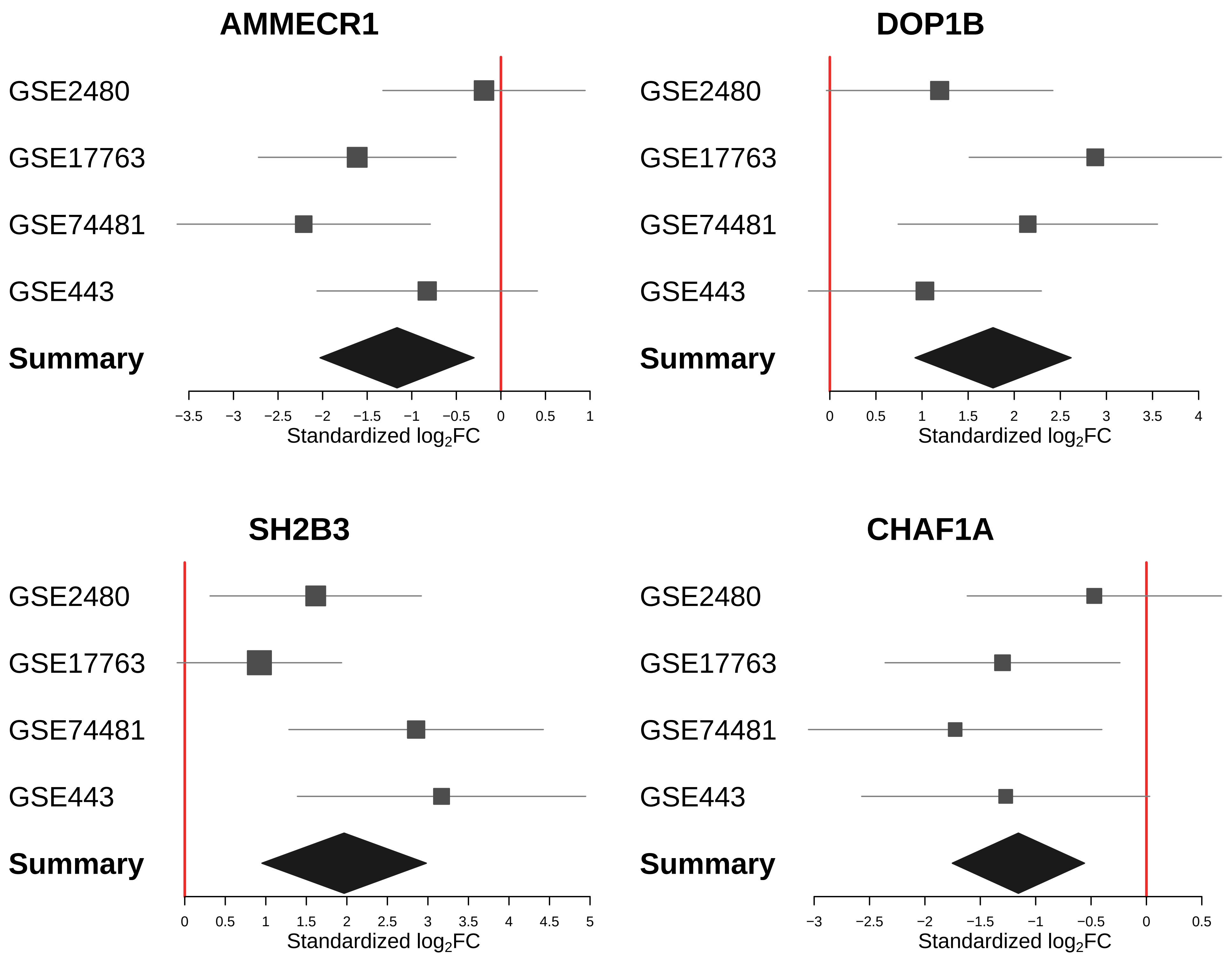
AMMECR1
GSE2480
GSE17763
GSE74481
GSE443
Summary
−3.5 −3 −2.5 −2 −1.5 −1 −0.5 0 0.5 1
Standardized log2FC
DOP1B
GSE2480
GSE17763
GSE74481
GSE443
Summary
0 0.5 1 1.5 2 2.5 3 3.5 4
Standardized log2FC
SH2B3
GSE2480
GSE17763
GSE74481
GSE443
Summary
0 0.5 1 1.5 2 2.5 3 3.5 4 4.5 5
Standardized log2FC
CHAF1A
GSE2480
GSE17763
GSE74481
GSE443
Summary
−3 −2.5 −2 −1.5 −1 −0.5 0 0.5
Standardized log2FC

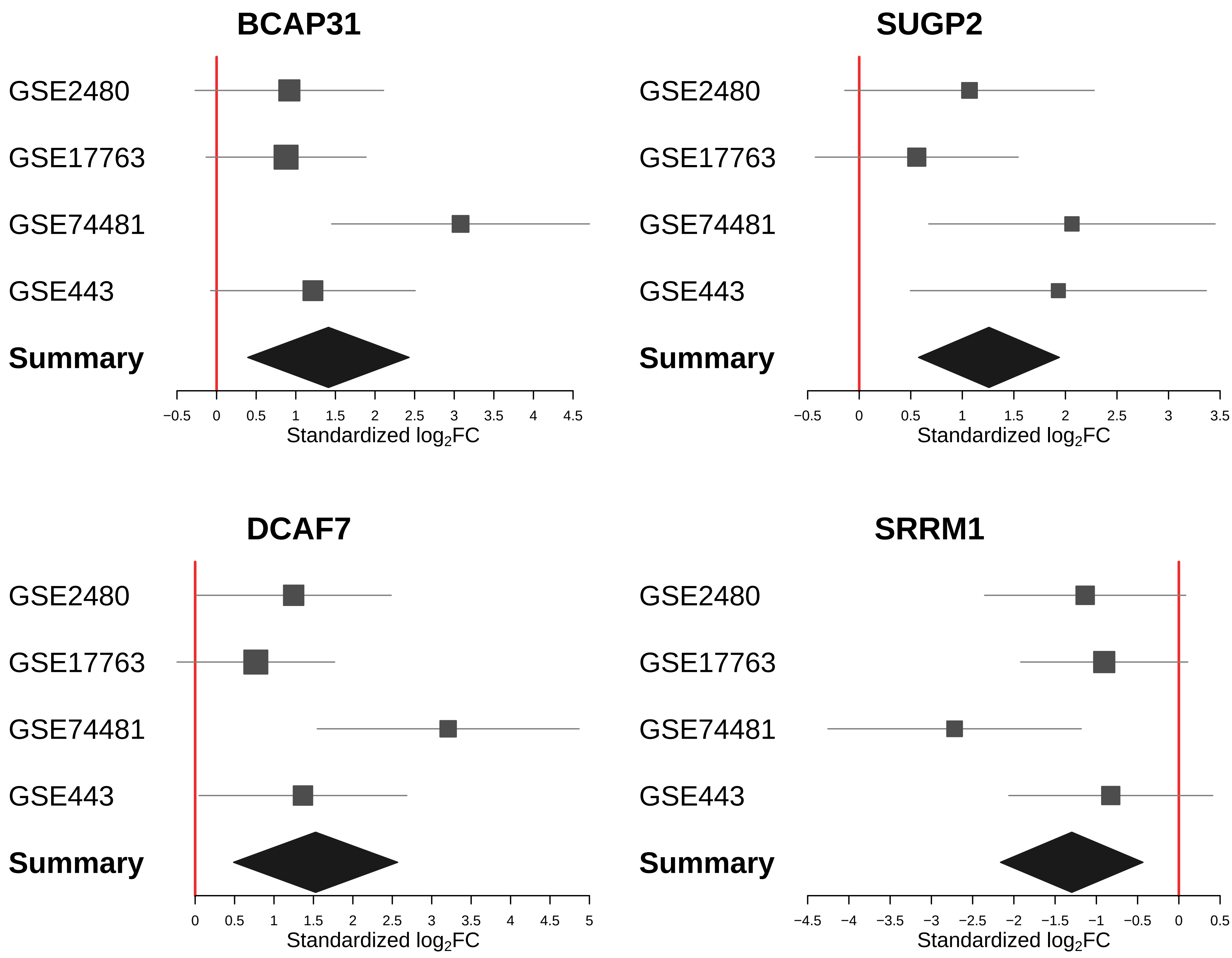

BCAP31
GSE2480
GSE17763
GSE74481
GSE443
Summary
−0.5 0 0.5 1 1.5 2 2.5 3 3.5 4 4.5
Standardized log2FC
SUGP2
GSE2480
GSE17763
GSE74481
GSE443
Summary
−0.5 0 0.5 1 1.5 2 2.5 3 3.5
Standardized log2FC
DCAF7
GSE2480
GSE17763
GSE74481
GSE443
Summary
0 0.5 1 1.5 2 2.5 3 3.5 4 4.5 5
Standardized log2FC
SRRM1
GSE2480
GSE17763
GSE74481
GSE443
Summary
−4.5 −4 −3.5 −3 −2.5 −2 −1.5 −1 −0.5 0 0.5
Standardized log2FC

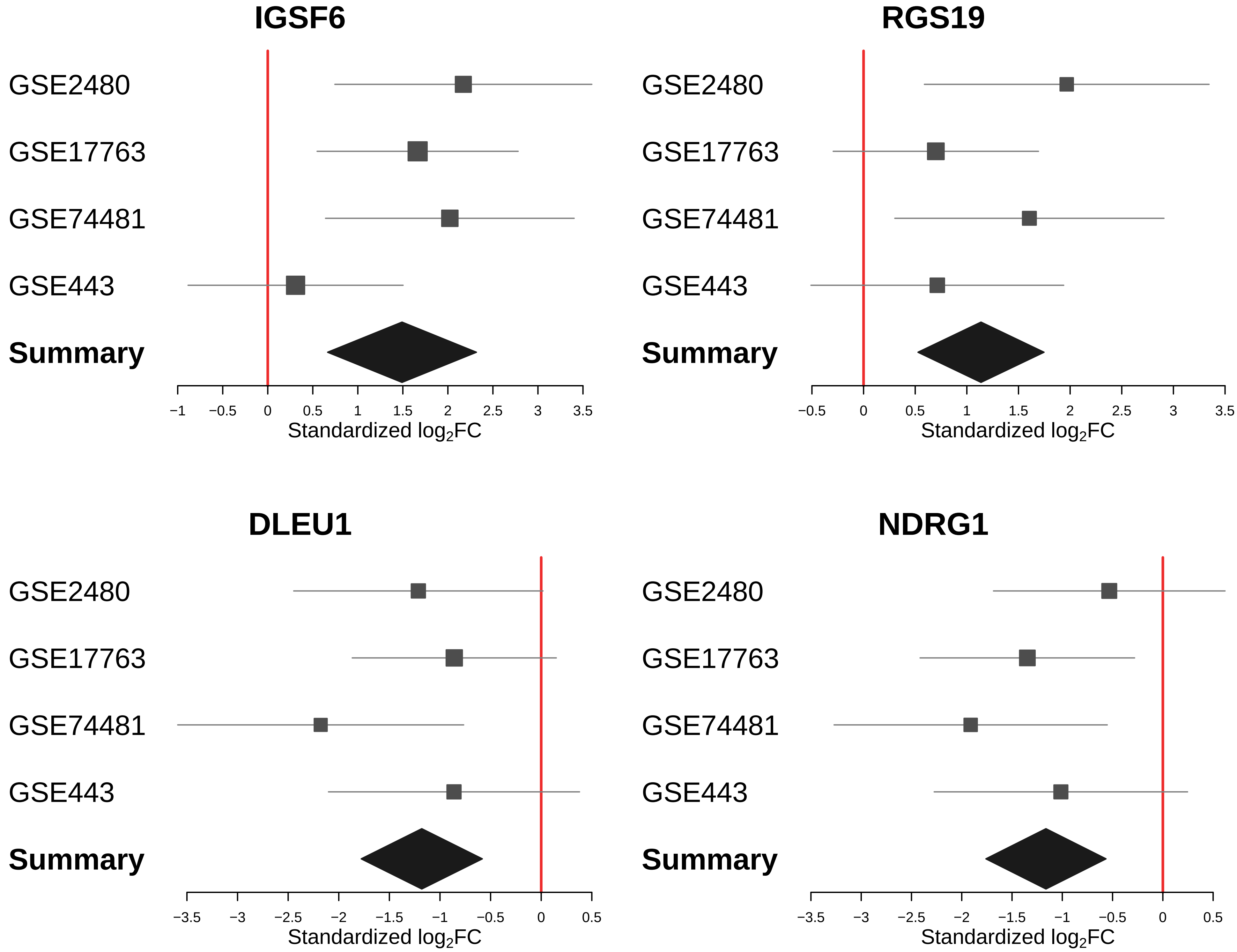

IGSF6
GSE2480
GSE17763
GSE74481
GSE443
Summary
−1 −0.5 0 0.5 1 1.5 2 2.5 3 3.5
Standardized log2FC
RGS19
GSE2480
GSE17763
GSE74481
GSE443
Summary
−0.5 0 0.5 1 1.5 2 2.5 3 3.5
Standardized log2FC
DLEU1
GSE2480
GSE17763
GSE74481
GSE443
Summary
−3.5 −3 −2.5 −2 −1.5 −1 −0.5 0 0.5
Standardized log2FC
NDRG1
GSE2480
GSE17763
GSE74481
GSE443
Summary
−3.5 −3 −2.5 −2 −1.5 −1 −0.5 0 0.5
Standardized log2FC

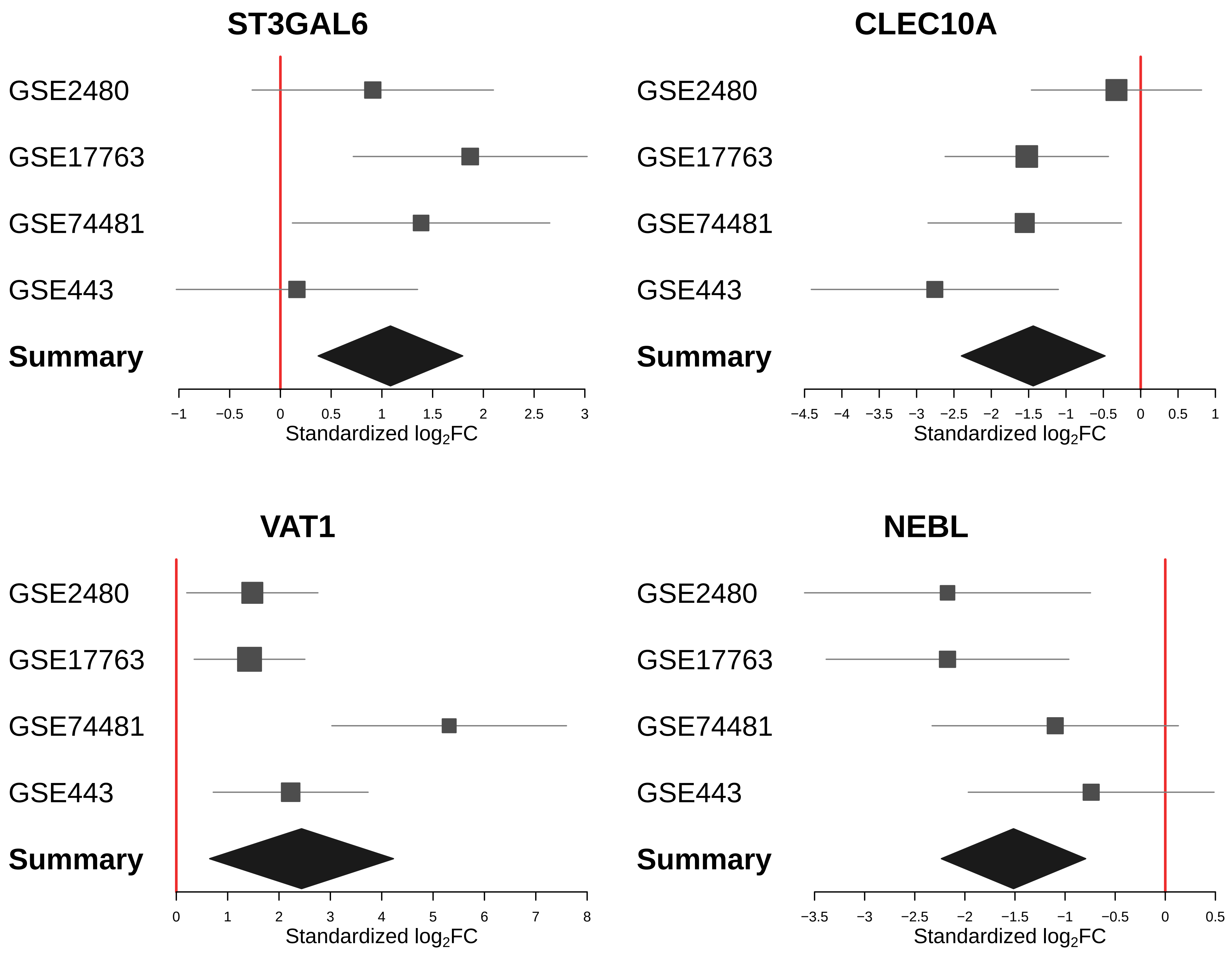

ST3GAL6
GSE2480
GSE17763
GSE74481
GSE443
Summary
−1 −0.5 0 0.5 1 1.5 2 2.5 3
Standardized log2FC
CLEC10A
GSE2480
GSE17763
GSE74481
GSE443
Summary
−4.5 −4 −3.5 −3 −2.5 −2 −1.5 −1 −0.5 0 0.5 1
Standardized log2FC
VAT1
GSE2480
GSE17763
GSE74481
GSE443
Summary
0 1 2 3 4 5 6 7 8
Standardized log2FC
NEBL
GSE2480
GSE17763
GSE74481
GSE443
Summary
−3.5 −3 −2.5 −2 −1.5 −1 −0.5 0 0.5
Standardized log2FC

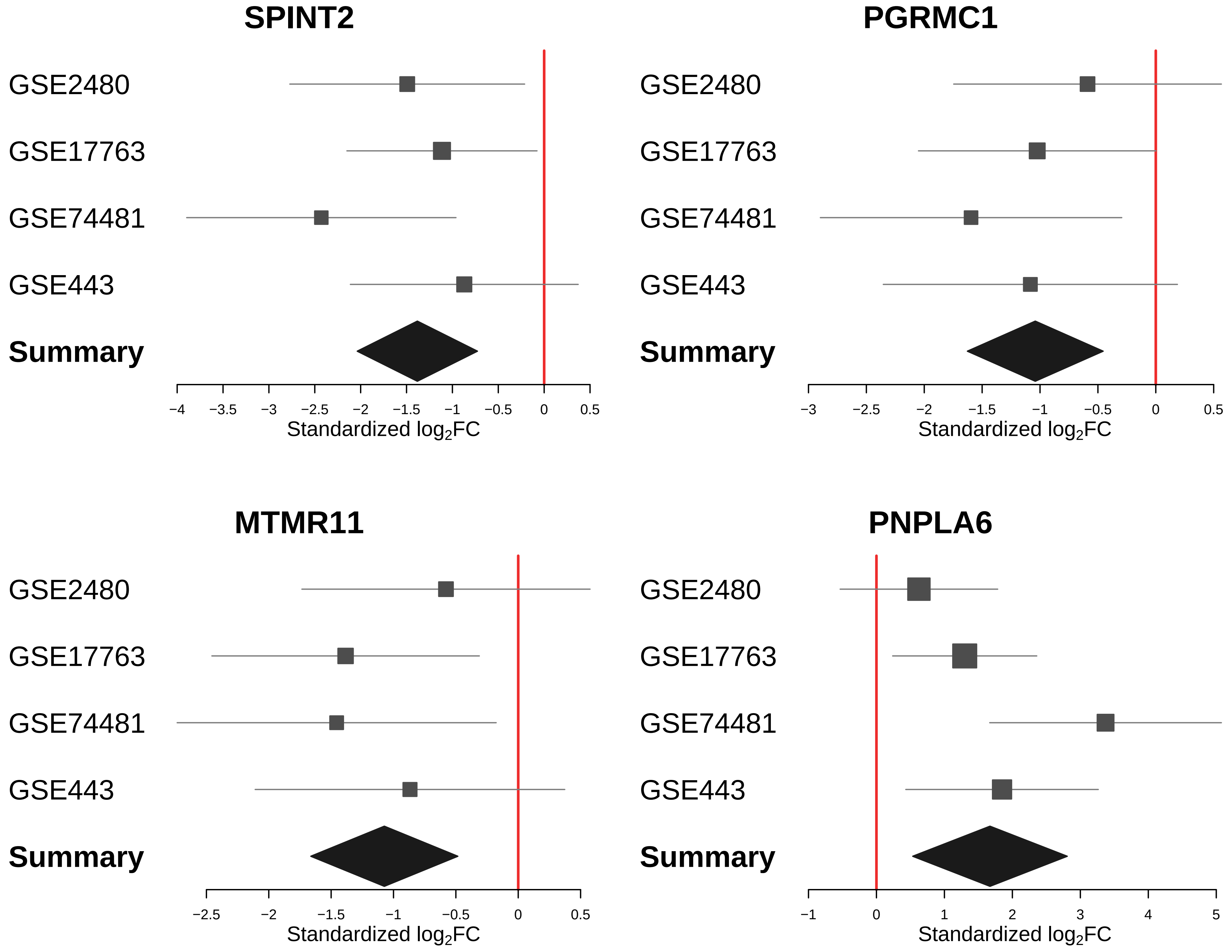

SPINT2
GSE2480
GSE17763
GSE74481
GSE443
Summary
−4 −3.5 −3 −2.5 −2 −1.5 −1 −0.5 0 0.5
Standardized log2FC
PGRMC1
GSE2480
GSE17763
GSE74481
GSE443
Summary
−3 −2.5 −2 −1.5 −1 −0.5 0 0.5
Standardized log2FC
MTMR11
GSE2480
GSE17763
GSE74481
GSE443
Summary
−2.5 −2 −1.5 −1 −0.5 0 0.5
Standardized log2FC
PNPLA6
GSE2480
GSE17763
GSE74481
GSE443
Summary
−1 0 1 2 3 4 5
Standardized log2FC

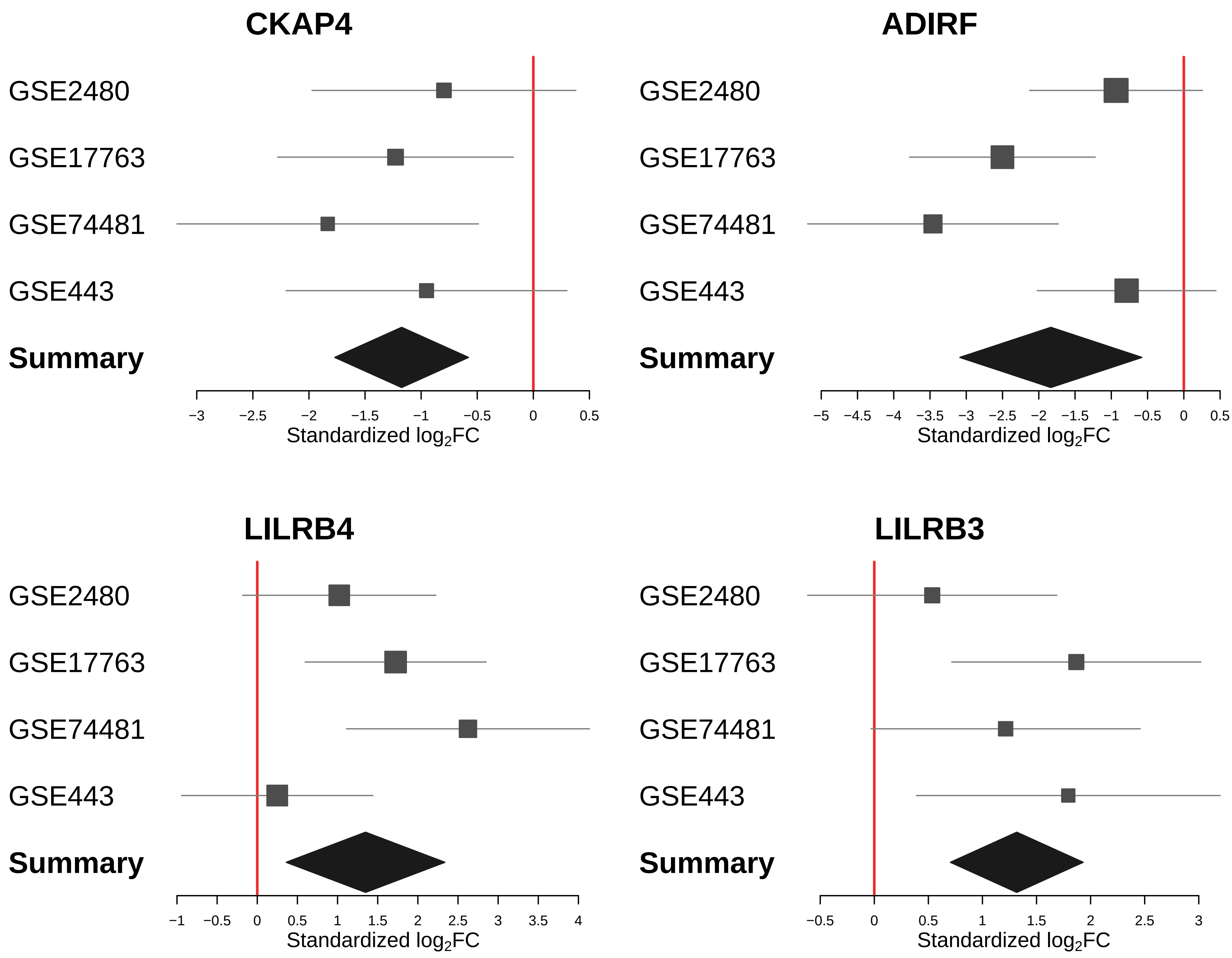
CKAP4
GSE2480
GSE17763
GSE74481
GSE443
Summary
−3 −2.5 −2 −1.5 −1 −0.5 0 0.5
Standardized log2FC
ADIRF
GSE2480
GSE17763
GSE74481
GSE443
Summary
−5 −4.5 −4 −3.5 −3 −2.5 −2 −1.5 −1 −0.5 0 0.5
Standardized log2FC
LILRB4
GSE2480
GSE17763
GSE74481
GSE443
Summary
−1 −0.5 0 0.5 1 1.5 2 2.5 3 3.5 4
Standardized log2FC
LILRB3
GSE2480
GSE17763
GSE74481
GSE443
Summary
−0.5 0 0.5 1 1.5 2 2.5 3
Standardized log2FC

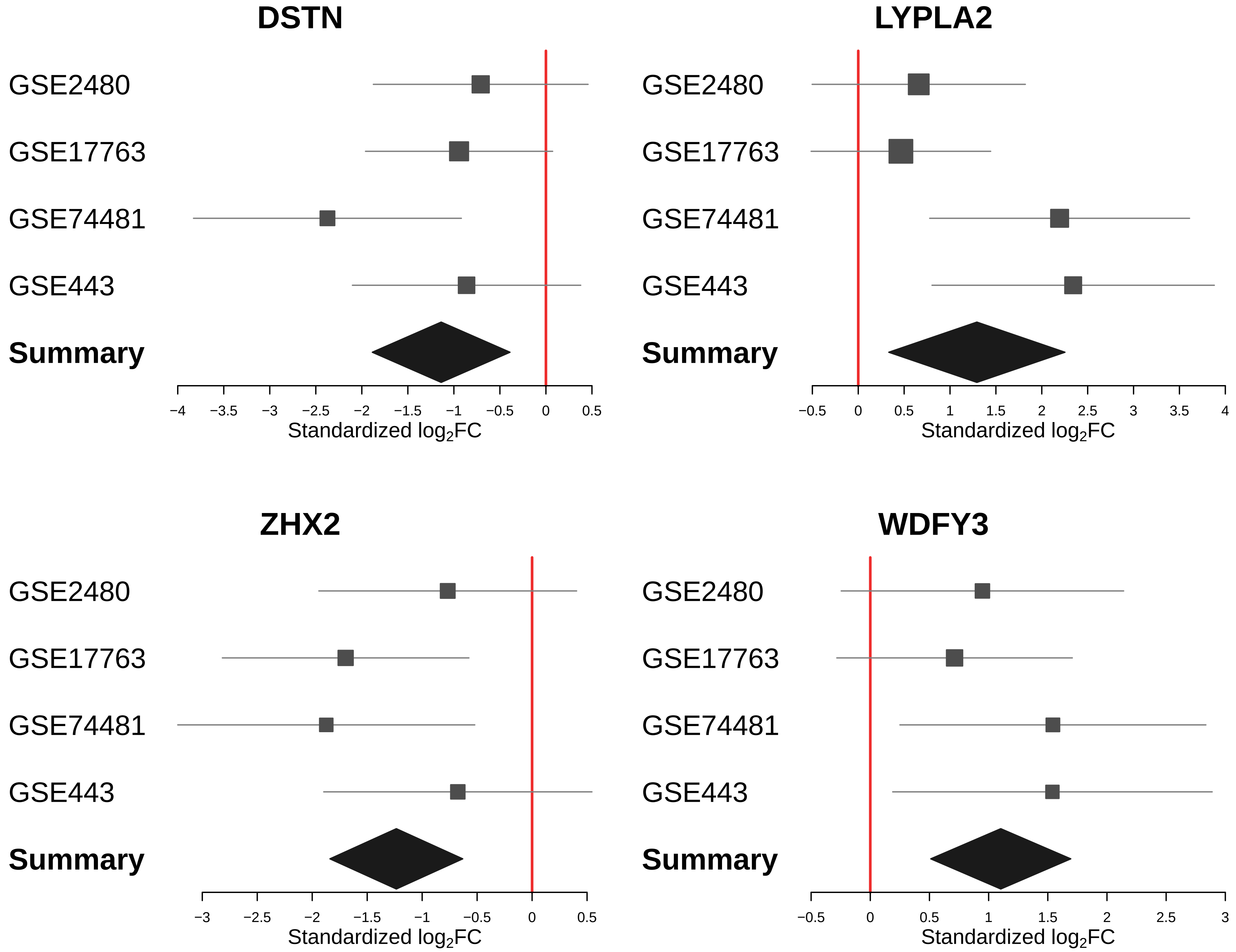

DSTN
GSE2480
GSE17763
GSE74481
GSE443
Summary
−4 −3.5 −3 −2.5 −2 −1.5 −1 −0.5 0 0.5
Standardized log2FC
LYPLA2
GSE2480
GSE17763
GSE74481
GSE443
Summary
−0.5 0 0.5 1 1.5 2 2.5 3 3.5 4
Standardized log2FC
ZHX2
GSE2480
GSE17763
GSE74481
GSE443
Summary
−3 −2.5 −2 −1.5 −1 −0.5 0 0.5
Standardized log2FC
WDFY3
GSE2480
GSE17763
GSE74481
GSE443
Summary
−0.5 0 0.5 1 1.5 2 2.5 3
Standardized log2FC

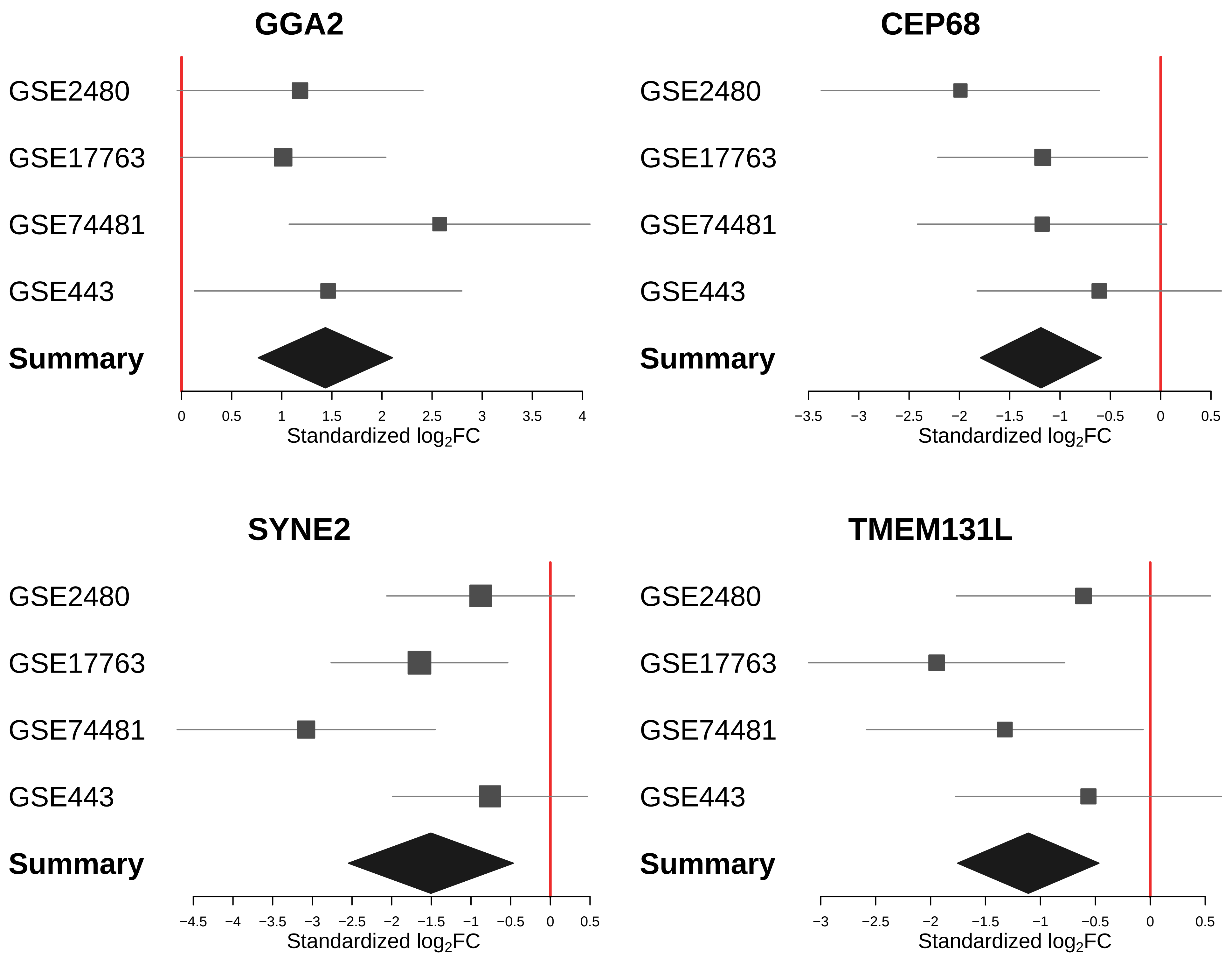
GGA2
GSE2480
GSE17763
GSE74481
GSE443
Summary
0 0.5 1 1.5 2 2.5 3 3.5 4
Standardized log2FC
CEP68
GSE2480
GSE17763
GSE74481
GSE443
Summary
−3.5 −3 −2.5 −2 −1.5 −1 −0.5 0 0.5
Standardized log2FC
SYNE2
GSE2480
GSE17763
GSE74481
GSE443
Summary
−4.5 −4 −3.5 −3 −2.5 −2 −1.5 −1 −0.5 0 0.5
Standardized log2FC
TMEM131L
GSE2480
GSE17763
GSE74481
GSE443
Summary
−3 −2.5 −2 −1.5 −1 −0.5 0 0.5
Standardized log2FC

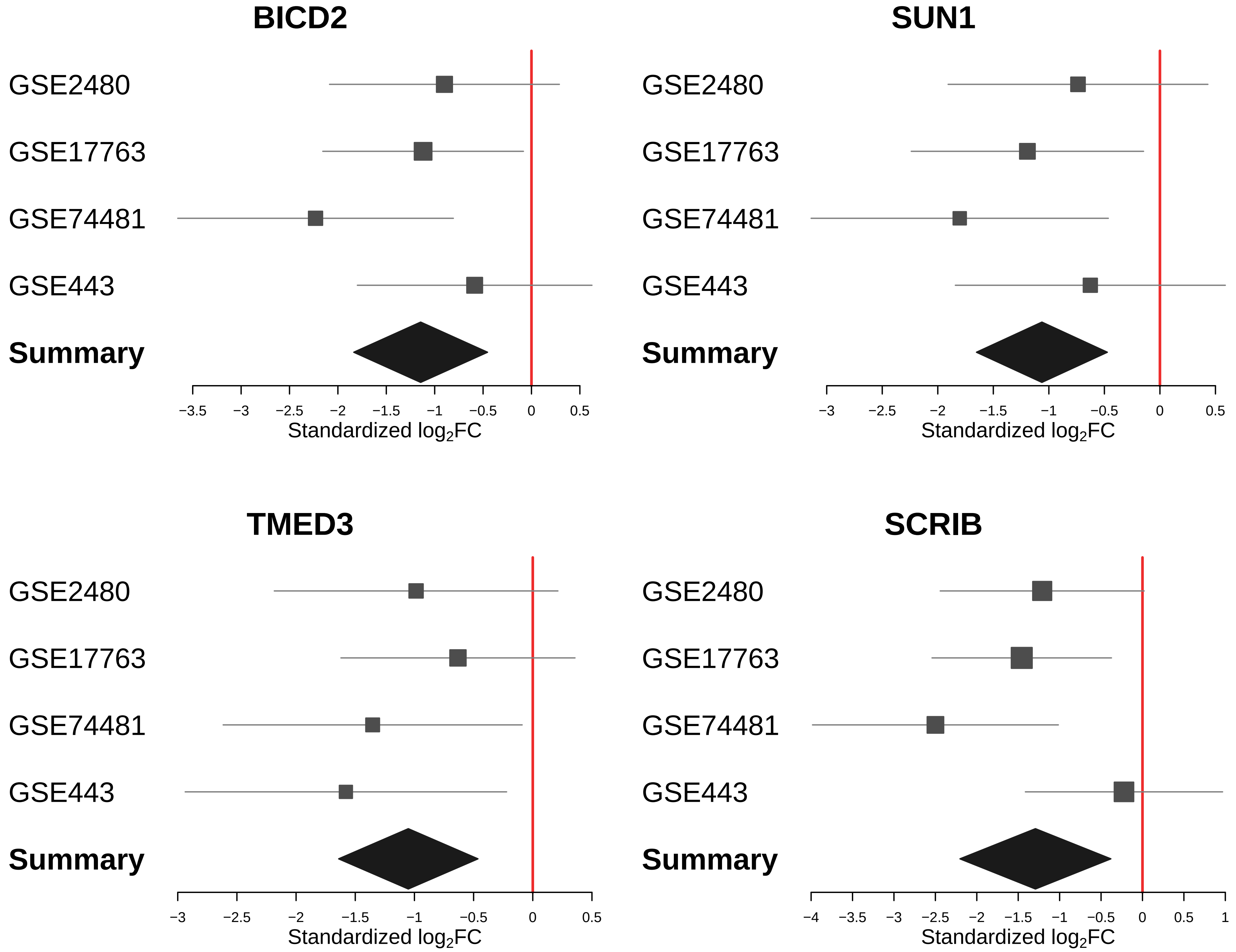
BICD2
GSE2480
GSE17763
GSE74481
GSE443
Summary
−3.5 −3 −2.5 −2 −1.5 −1 −0.5 0 0.5
Standardized $log_2FC$
SUN1
GSE2480
GSE17763
GSE74481
GSE443
Summary
−3 −2.5 −2 −1.5 −1 −0.5 0 0.5
Standardized $log_2FC$
TMED3
GSE2480
GSE17763
GSE74481
GSE443
Summary
−3 −2.5 −2 −1.5 −1 −0.5 0 0.5
Standardized $log_2FC$
SCRIB
GSE2480
GSE17763
GSE74481
GSE443
Summary
−4 −3.5 −3 −2.5 −2 −1.5 −1 −0.5 0 0.5 1
Standardized $log_2FC$

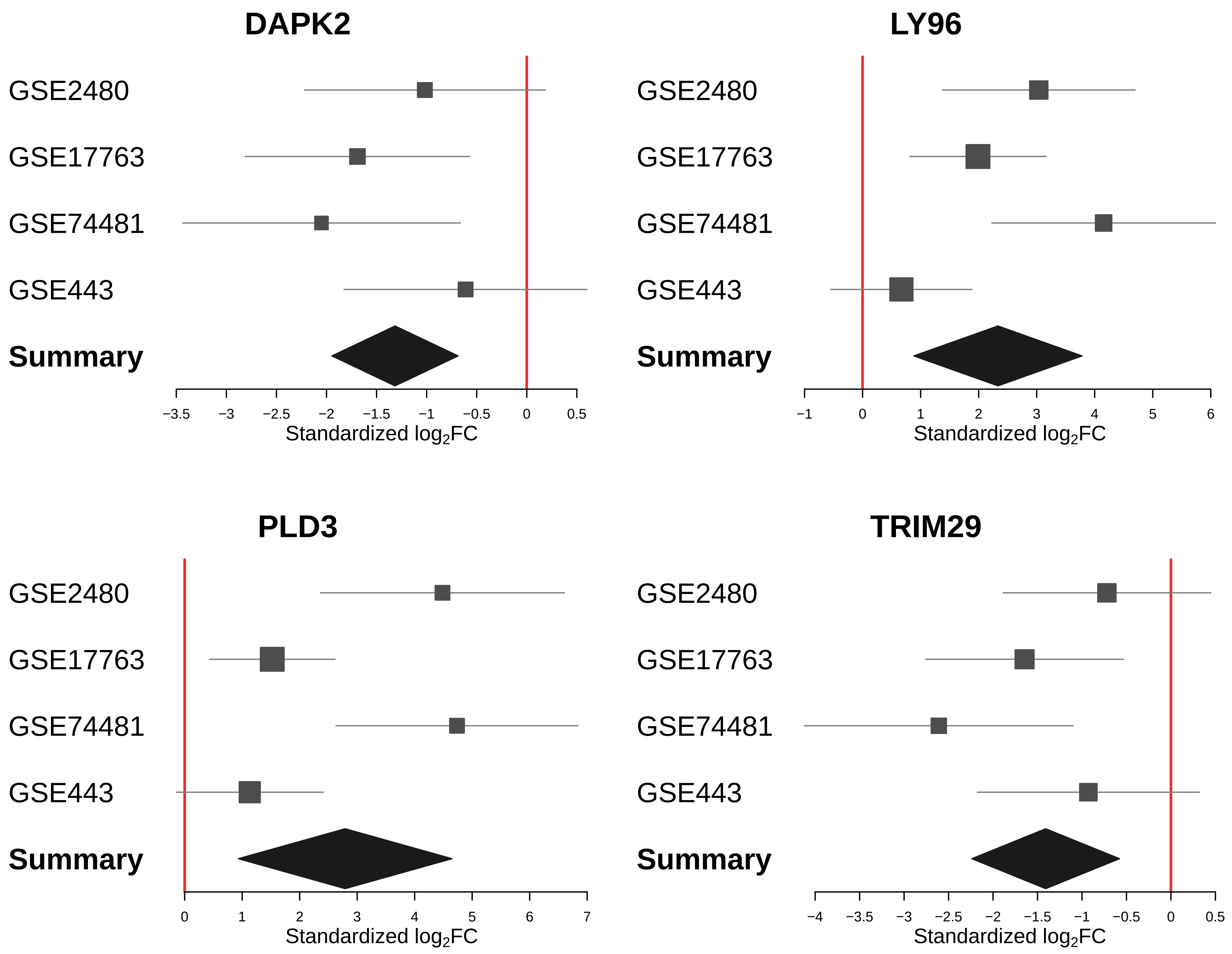
DAPK2
LY96
PLD3
TRIM29
GSE2480
GSE17763
GSE74481
GSE443
Summary
−3.5 −3 −2.5 −2 −1.5 −1 −0.5 0 0.5
−1 0 1 2 3 4 5 6
0 1 2 3 4 5 6 7
−4 −3.5 −3 −2.5 −2 −1.5 −1 −0.5 0 0.5
Standardized $log_2FC$

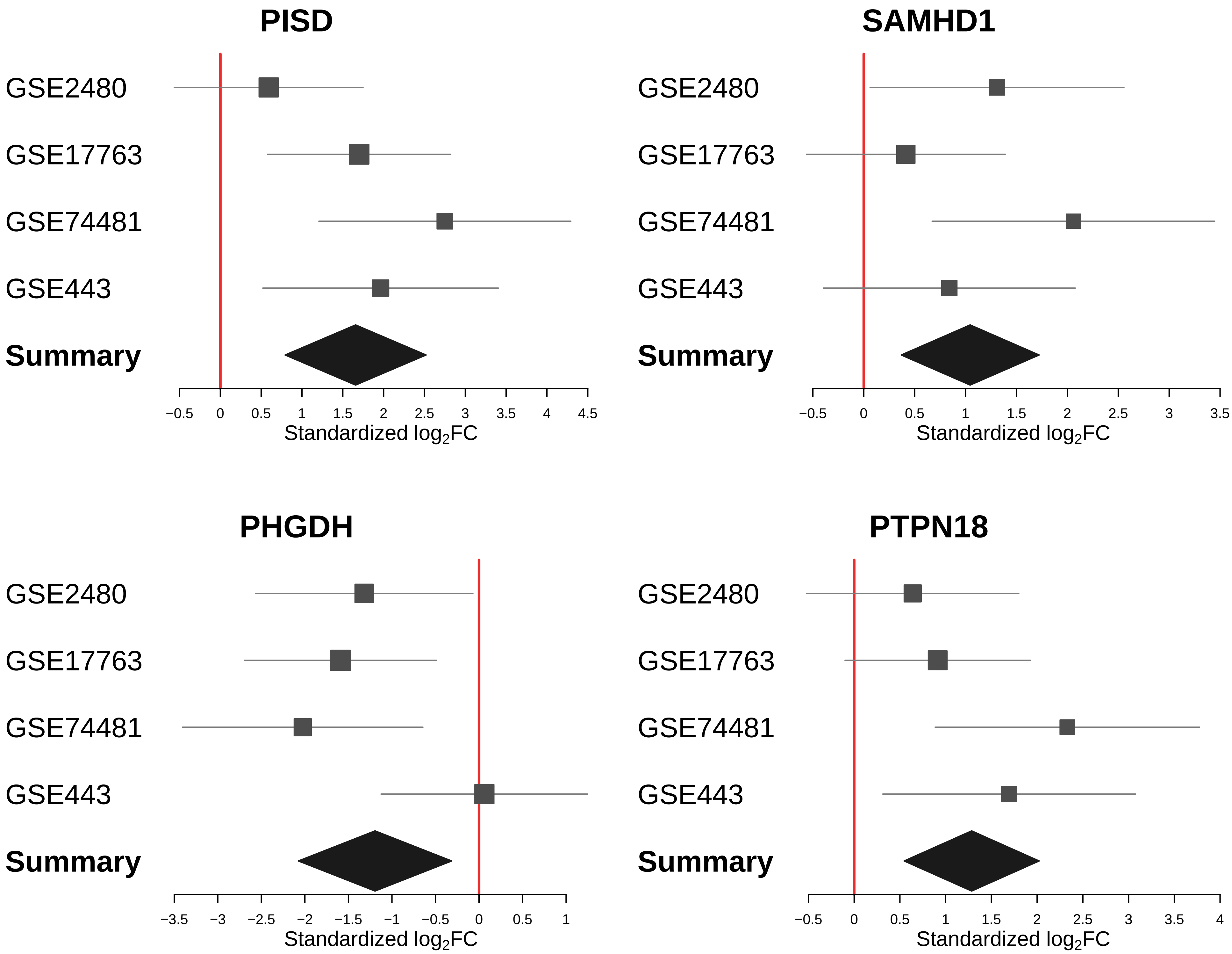
PISD
GSE2480
GSE17763
GSE74481
GSE443
Summary
−0.5 0 0.5 1 1.5 2 2.5 3 3.5 4 4.5
Standardized log2FC
SAMHD1
GSE2480
GSE17763
GSE74481
GSE443
Summary
−0.5 0 0.5 1 1.5 2 2.5 3 3.5
Standardized log2FC
PHGDH
GSE2480
GSE17763
GSE74481
GSE443
Summary
−3.5 −3 −2.5 −2 −1.5 −1 −0.5 0 0.5 1
Standardized log2FC
PTPN18
GSE2480
GSE17763
GSE74481
GSE443
Summary
−0.5 0 0.5 1 1.5 2 2.5 3 3.5 4
Standardized log2FC

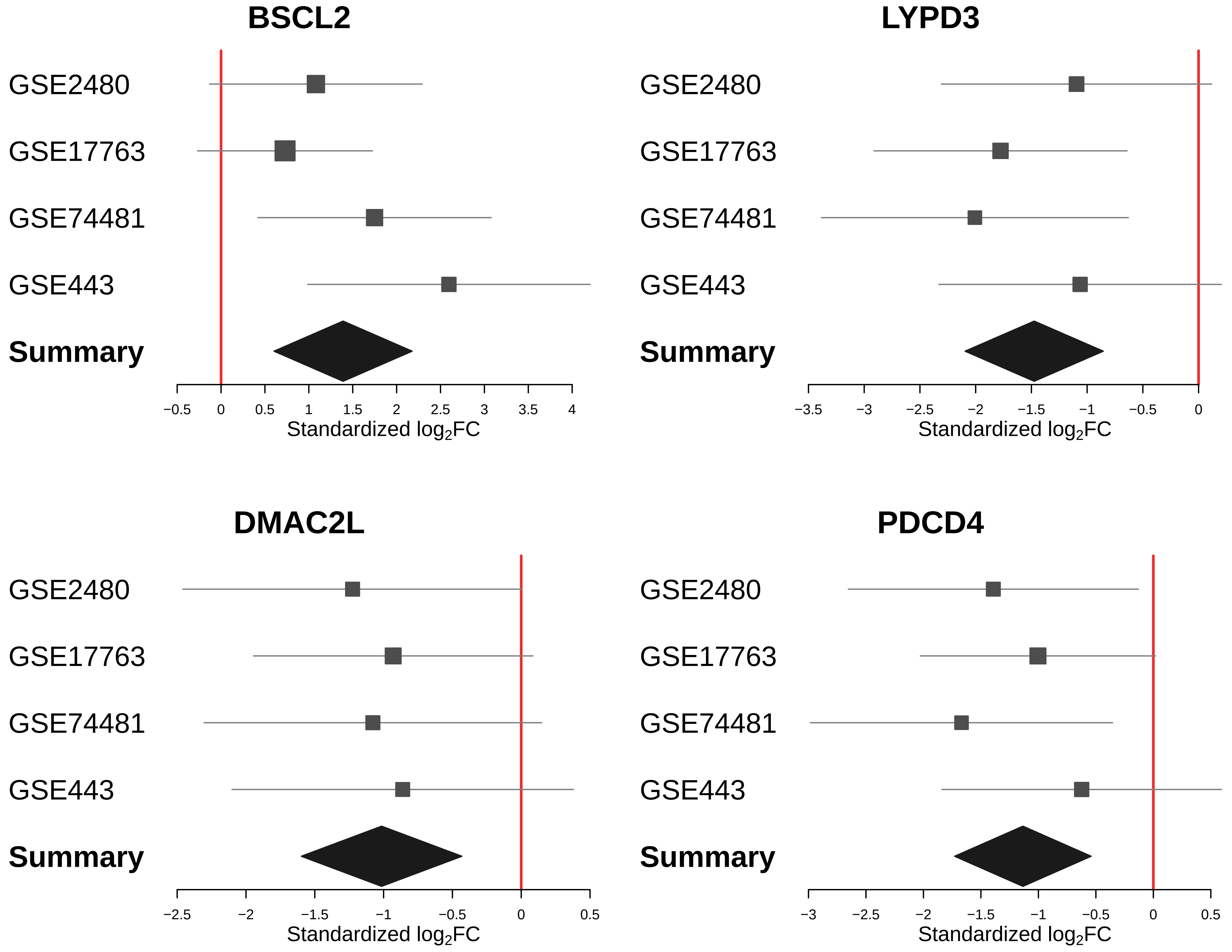

BSCL2
GSE2480
GSE17763
GSE74481
GSE443
Summary
−0.5 0 0.5 1 1.5 2 2.5 3 3.5 4
Standardized log2FC
LYPD3
GSE2480
GSE17763
GSE74481
GSE443
Summary
−3.5 −3 −2.5 −2 −1.5 −1 −0.5 0
Standardized log2FC
DMAC2L
GSE2480
GSE17763
GSE74481
GSE443
Summary
−2.5 −2 −1.5 −1 −0.5 0 0.5
Standardized log2FC
PDCD4
GSE2480
GSE17763
GSE74481
GSE443
Summary
−3 −2.5 −2 −1.5 −1 −0.5 0 0.5
Standardized log2FC

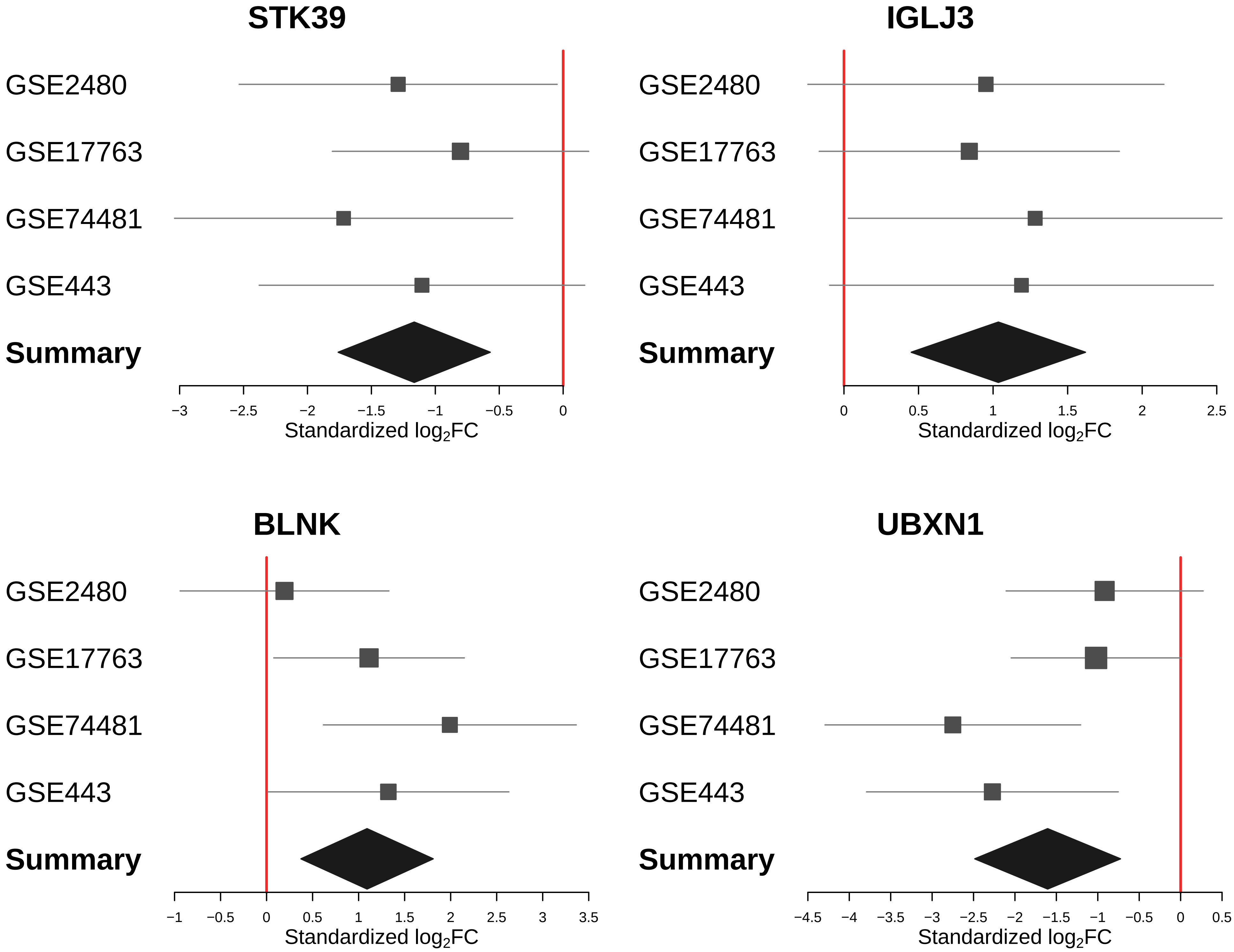
STK39
GSE2480
GSE17763
GSE74481
GSE443
Summary
−3 −2.5 −2 −1.5 −1 −0.5 0
Standardized log2FC
IGLJ3
GSE2480
GSE17763
GSE74481
GSE443
Summary
0 0.5 1 1.5 2 2.5
Standardized log2FC
BLNK
GSE2480
GSE17763
GSE74481
GSE443
Summary
−1 −0.5 0 0.5 1 1.5 2 2.5 3 3.5
Standardized log2FC
UBXN1
GSE2480
GSE17763
GSE74481
GSE443
Summary
−4.5 −4 −3.5 −3 −2.5 −2 −1.5 −1 −0.5 0 0.5
Standardized log2FC

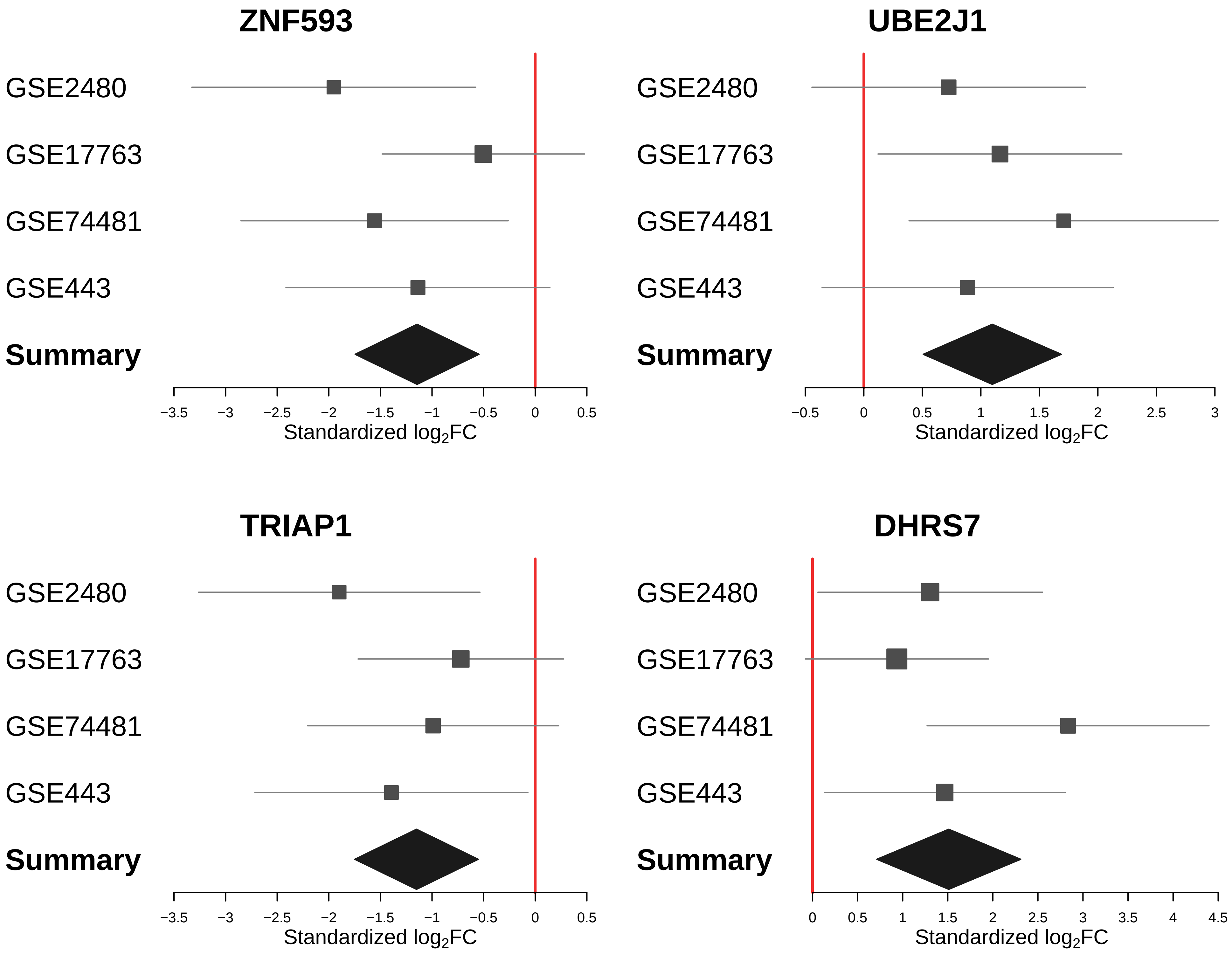
ZNF593
GSE2480
GSE17763
GSE74481
GSE443
Summary
−3.5 −3 −2.5 −2 −1.5 −1 −0.5 0 0.5
Standardized log2FC
UBE2J1
GSE2480
GSE17763
GSE74481
GSE443
Summary
−0.5 0 0.5 1 1.5 2 2.5 3
Standardized log2FC
TRIAP1
GSE2480
GSE17763
GSE74481
GSE443
Summary
−3.5 −3 −2.5 −2 −1.5 −1 −0.5 0 0.5
Standardized log2FC
DHRS7
GSE2480
GSE17763
GSE74481
GSE443
Summary
0 0.5 1 1.5 2 2.5 3 3.5 4 4.5
Standardized log2FC

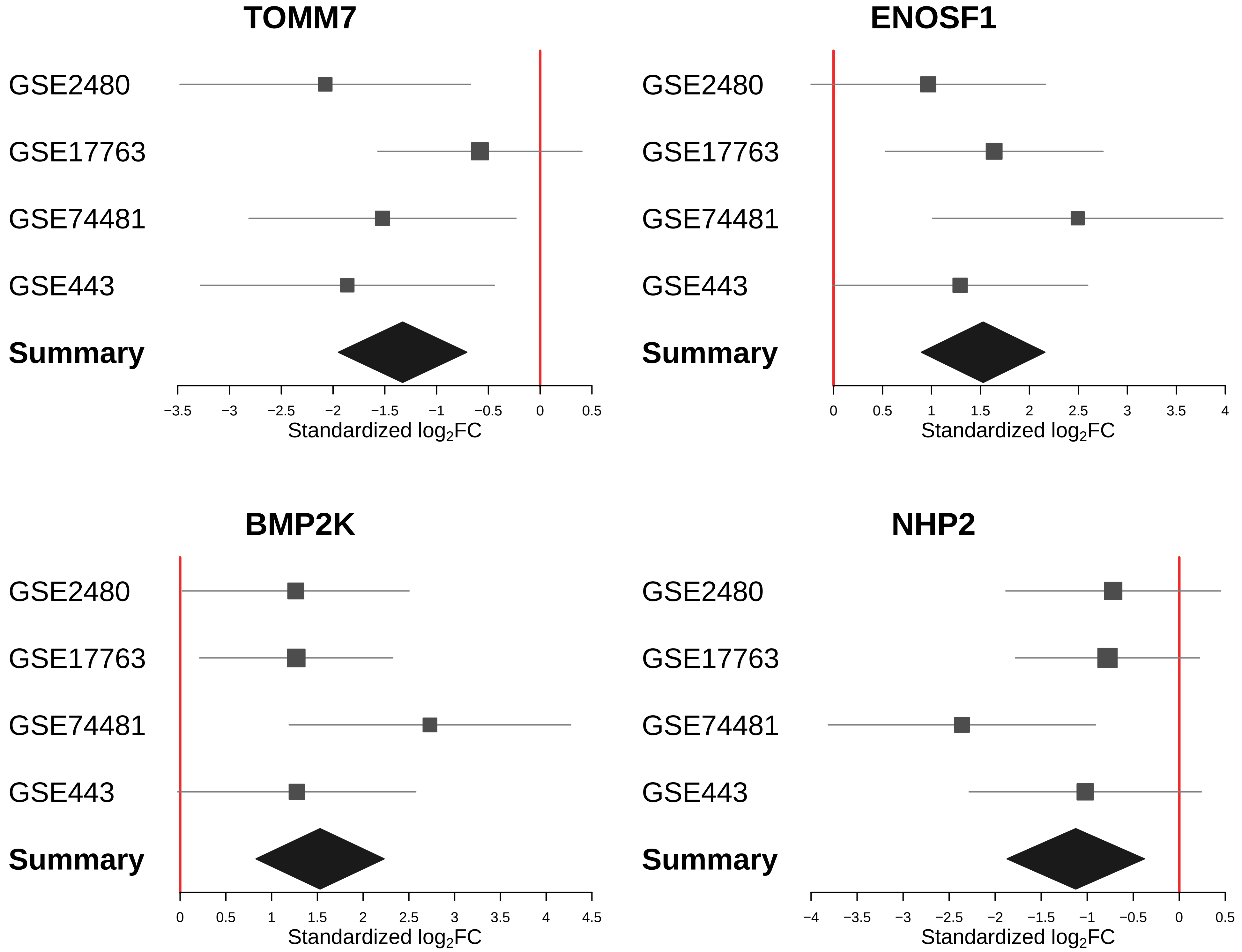
TOMM7
GSE2480
GSE17763
GSE74481
GSE443
Summary
−3.5 −3 −2.5 −2 −1.5 −1 −0.5 0 0.5
Standardized $log_2FC$
ENOSF1
GSE2480
GSE17763
GSE74481
GSE443
Summary
0 0.5 1 1.5 2 2.5 3 3.5 4
Standardized $log_2FC$
BMP2K
GSE2480
GSE17763
GSE74481
GSE443
Summary
0 0.5 1 1.5 2 2.5 3 3.5 4 4.5
Standardized $log_2FC$
NHP2
GSE2480
GSE17763
GSE74481
GSE443
Summary
−4 −3.5 −3 −2.5 −2 −1.5 −1 −0.5 0 0.5
Standardized $log_2FC$

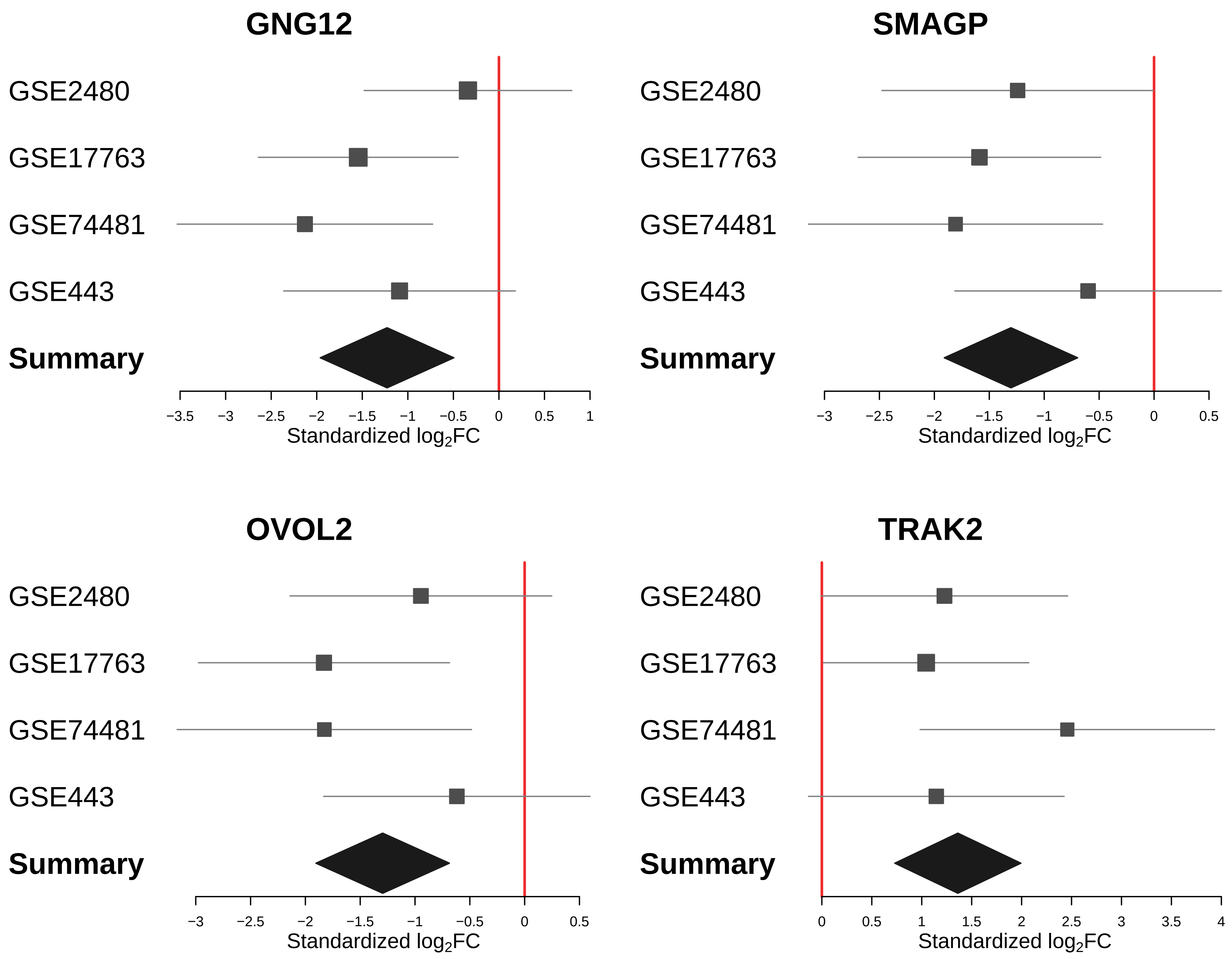
GNG12
GSE2480
GSE17763
GSE74481
GSE443
Summary
−3.5 −3 −2.5 −2 −1.5 −1 −0.5 0 0.5 1
Standardized log2FC
SMAGP
GSE2480
GSE17763
GSE74481
GSE443
Summary
−3 −2.5 −2 −1.5 −1 −0.5 0 0.5
Standardized log2FC
OVOL2
GSE2480
GSE17763
GSE74481
GSE443
Summary
−3 −2.5 −2 −1.5 −1 −0.5 0 0.5
Standardized log2FC
TRAK2
GSE2480
GSE17763
GSE74481
GSE443
Summary
0 0.5 1 1.5 2 2.5 3 3.5 4
Standardized log2FC

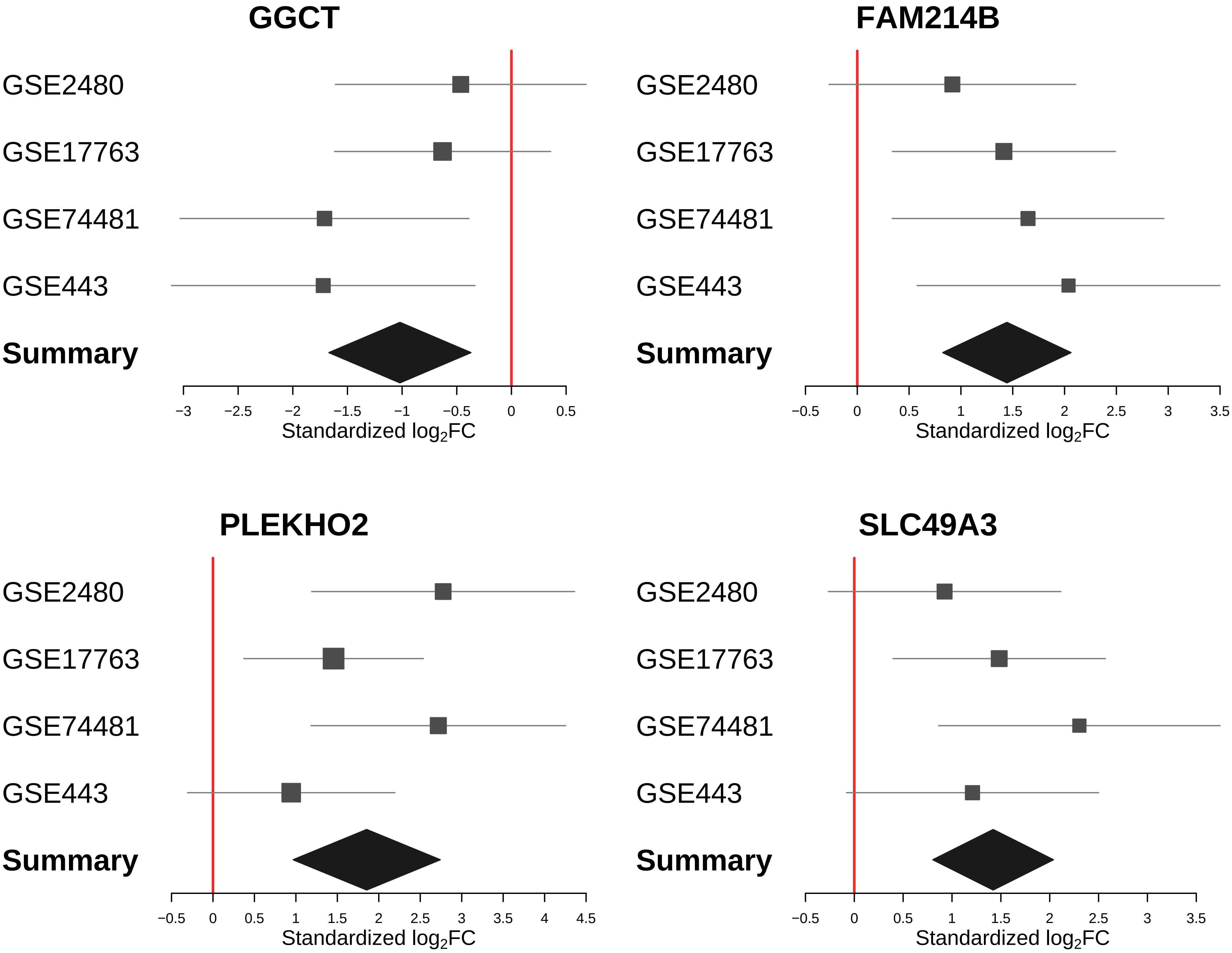
GGCT
GSE2480
GSE17763
GSE74481
GSE443
Summary
−3
−2.5
−2
−1.5
−1
−0.5
0
0.5
Standardized log2FC
FAM214B
GSE2480
GSE17763
GSE74481
GSE443
Summary
−0.5
0
0.5
1
1.5
2
2.5
3
3.5
Standardized log2FC
PLEKHO2
GSE2480
GSE17763
GSE74481
GSE443
Summary
−0.5
0
0.5
1
1.5
2
2.5
3
3.5
4
4.5
Standardized log2FC
SLC49A3
GSE2480
GSE17763
GSE74481
GSE443
Summary
−0.5
0
0.5
1
1.5
2
2.5
3
3.5
Standardized log2FC

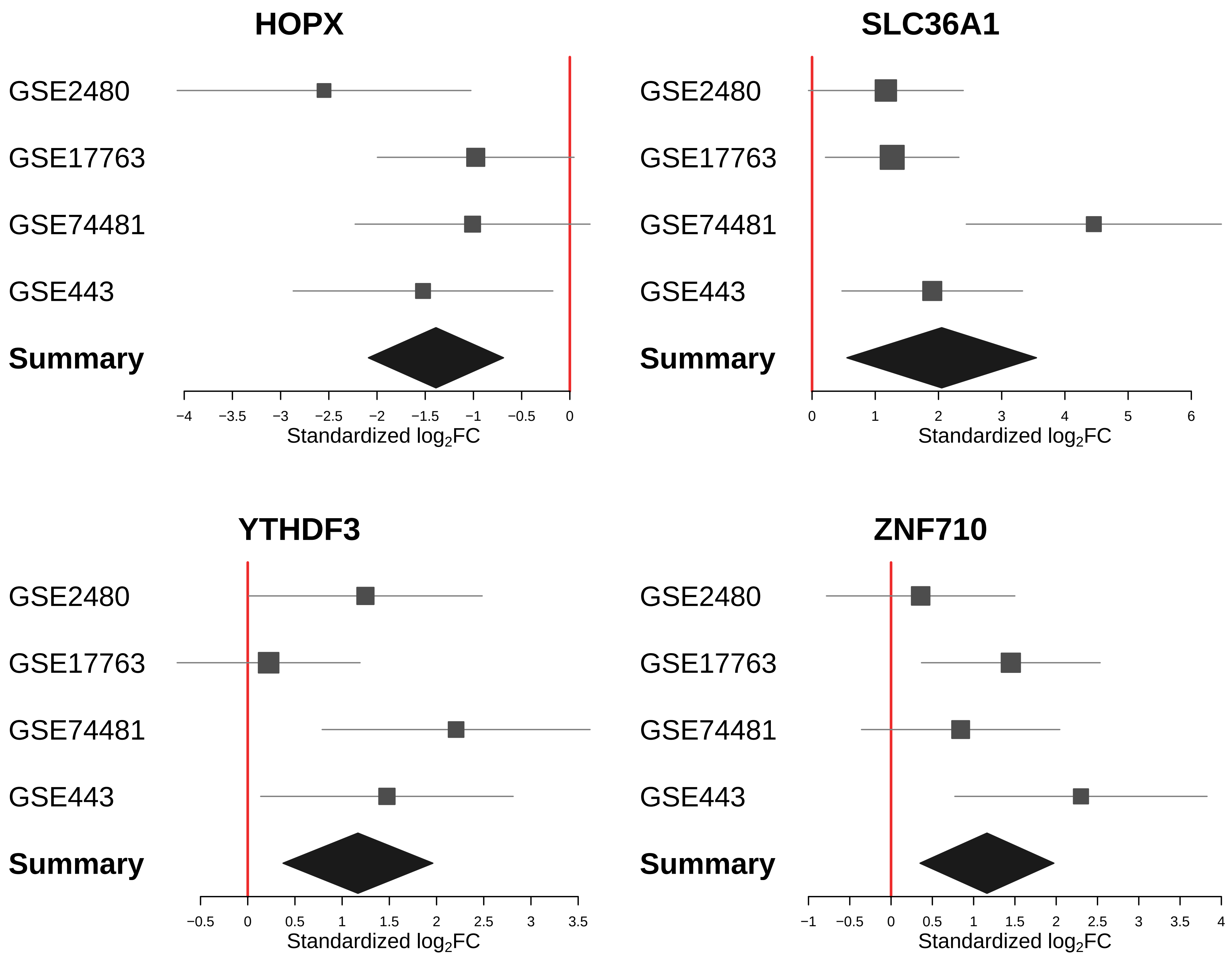

HOPX
SLC36A1
YTHDF3
ZNF710
GSE2480
GSE17763
GSE74481
GSE443
Summary
Standardized log2FC

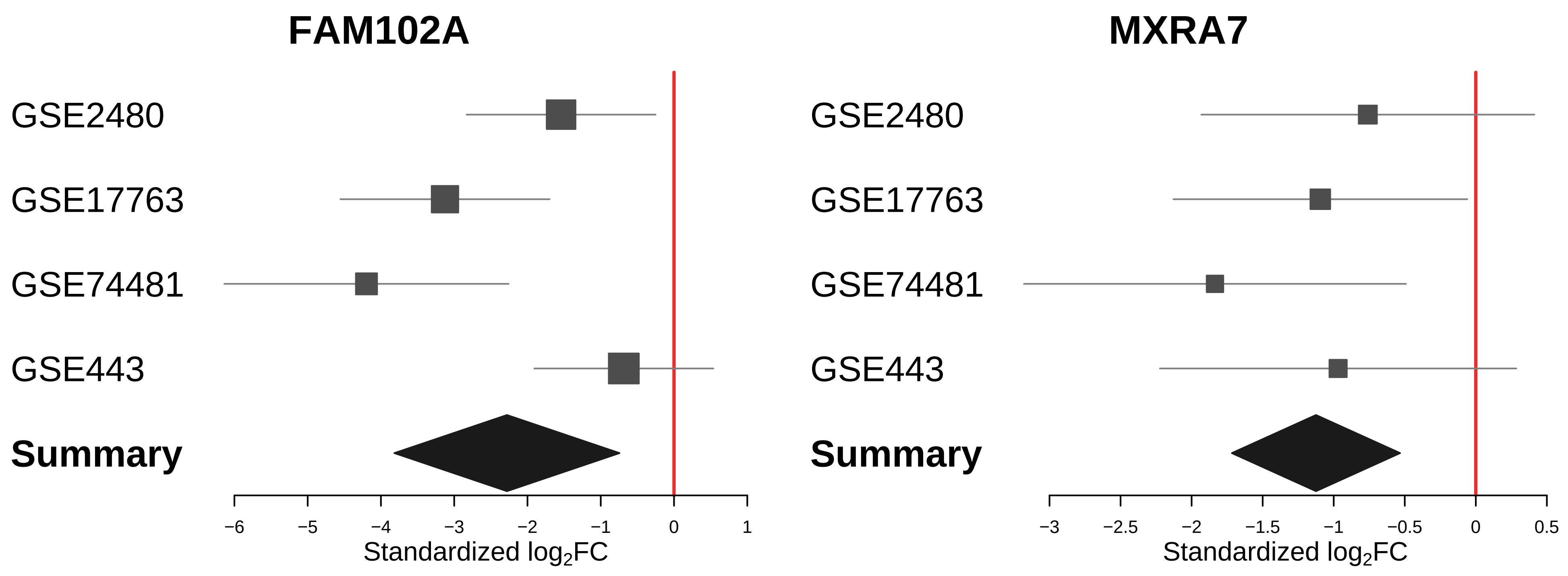
FAM102A
GSE2480
GSE17763
GSE74481
GSE443
Summary
−6 −5 −4 −3 −2 −1 0 1
Standardized log2FC
MXRA7
GSE2480
GSE17763
GSE74481
GSE443
Summary
−3 −2.5 −2 −1.5 −1 −0.5 0 0.5
Standardized log2FC
