## Supplementary figures S1-S2 + REM forest plots for "Reanalysis And Integration Of Public Microarray Datasets Reveals Novel Host Genes Modulated In Leprosy": LLvsENL_REM_forestplot.pdf (from another version of this preprint: version 2, 2019-12-07)

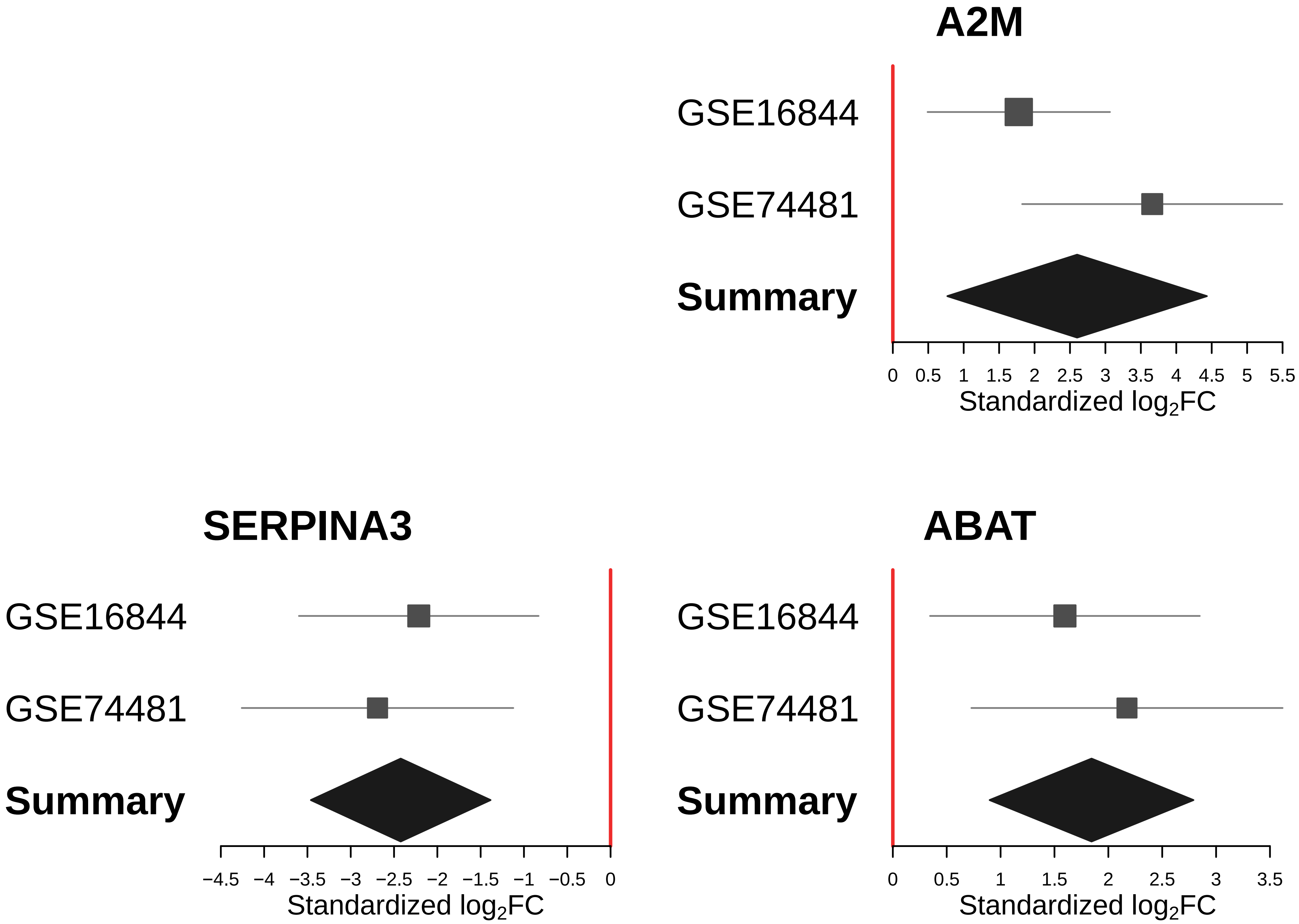
A2M
GSE16844
GSE74481
Summary
0 0.5 1 1.5 2 2.5 3 3.5 4 4.5 5 5.5
Standardized log2FC
SERPINA3
GSE16844
GSE74481
Summary
−4.5 −4 −3.5 −3 −2.5 −2 −1.5 −1 −0.5 0
Standardized log2FC
ABAT
GSE16844
GSE74481
Summary
0 0.5 1 1.5 2 2.5 3 3.5
Standardized log2FC

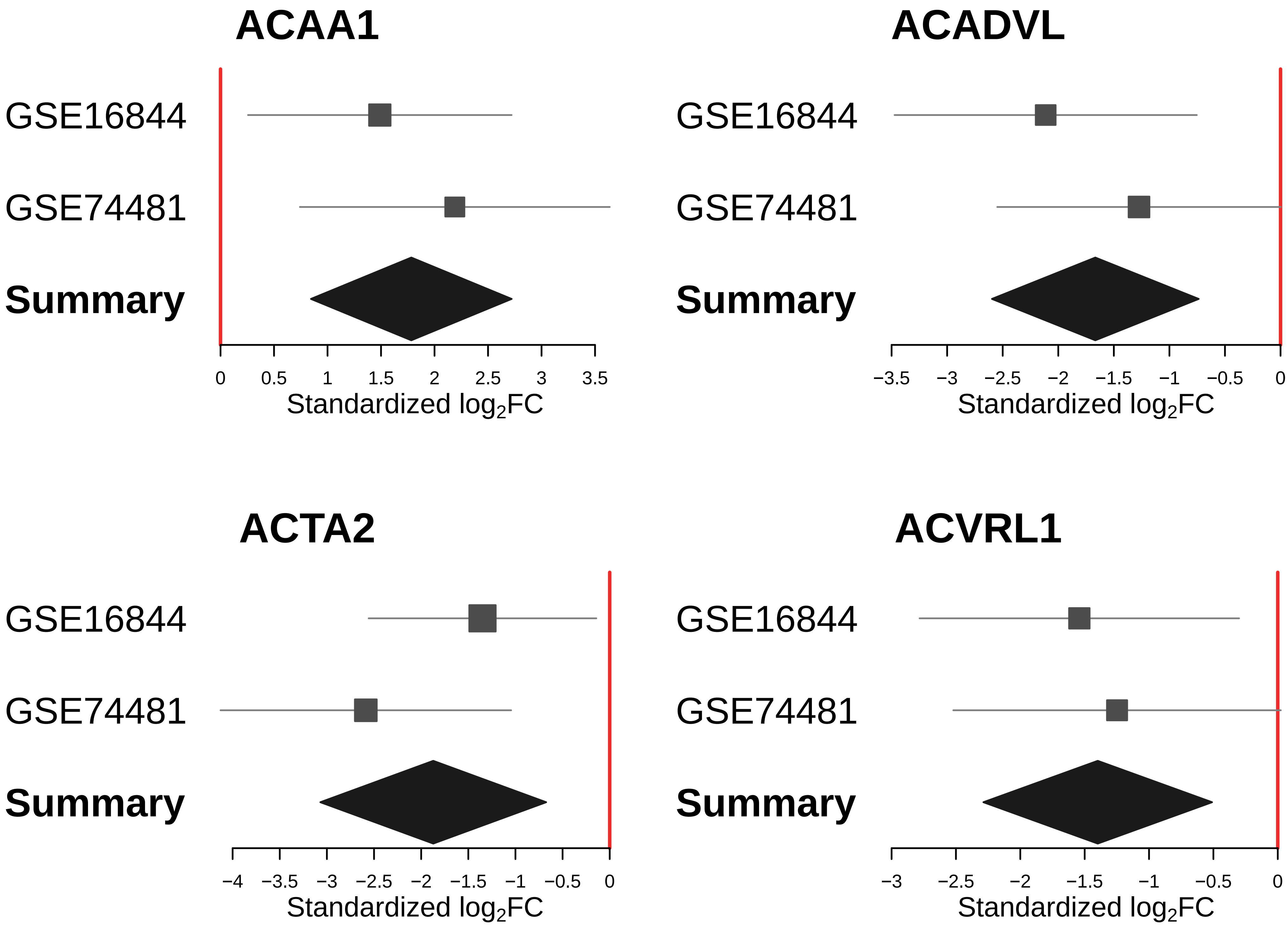

ACAA1
GSE16844
GSE74481
Summary
0 0.5 1 1.5 2 2.5 3 3.5
Standardized log2FC
ACADVL
GSE16844
GSE74481
Summary
−3.5 −3 −2.5 −2 −1.5 −1 −0.5 0
Standardized log2FC
ACTA2
GSE16844
GSE74481
Summary
−4 −3.5 −3 −2.5 −2 −1.5 −1 −0.5 0
Standardized log2FC
ACVRL1
GSE16844
GSE74481
Summary
−3 −2.5 −2 −1.5 −1 −0.5 0
Standardized log2FC

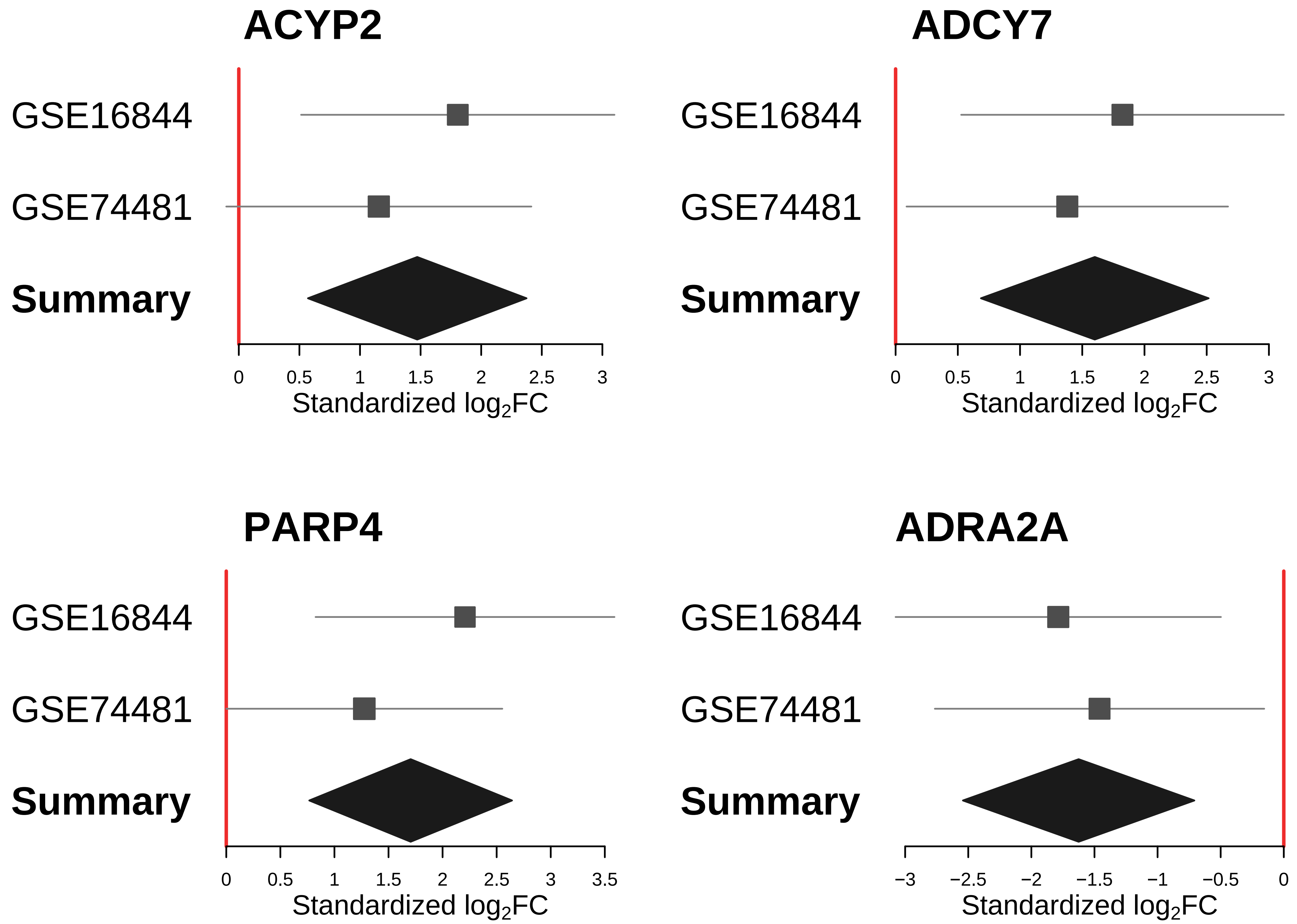

ACYP2
GSE16844
GSE74481
Summary
0 0.5 1 1.5 2 2.5 3
Standardized log2FC
ADCY7
GSE16844
GSE74481
Summary
0 0.5 1 1.5 2 2.5 3
Standardized log2FC
PARP4
GSE16844
GSE74481
Summary
0 0.5 1 1.5 2 2.5 3 3.5
Standardized log2FC
ADRA2A
GSE16844
GSE74481
Summary
−3 −2.5 −2 −1.5 −1 −0.5 0
Standardized log2FC

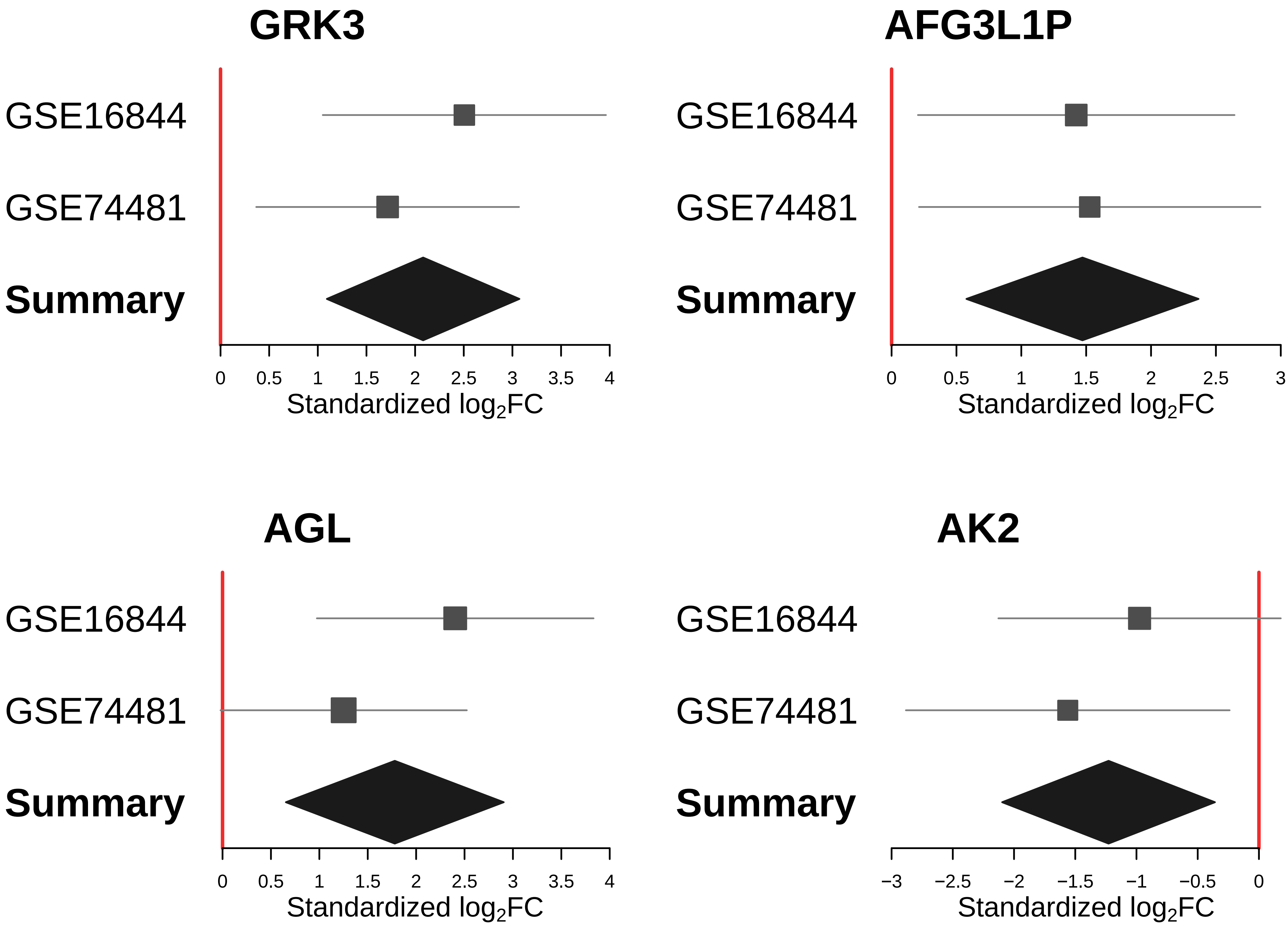
GRK3
GSE16844
GSE74481
Summary
0 0.5 1 1.5 2 2.5 3 3.5 4
Standardized log2FC
AFG3L1P
GSE16844
GSE74481
Summary
0 0.5 1 1.5 2 2.5 3
Standardized log2FC
AGL
GSE16844
GSE74481
Summary
0 0.5 1 1.5 2 2.5 3 3.5 4
Standardized log2FC
AK2
GSE16844
GSE74481
Summary
−3 −2.5 −2 −1.5 −1 −0.5 0
Standardized log2FC

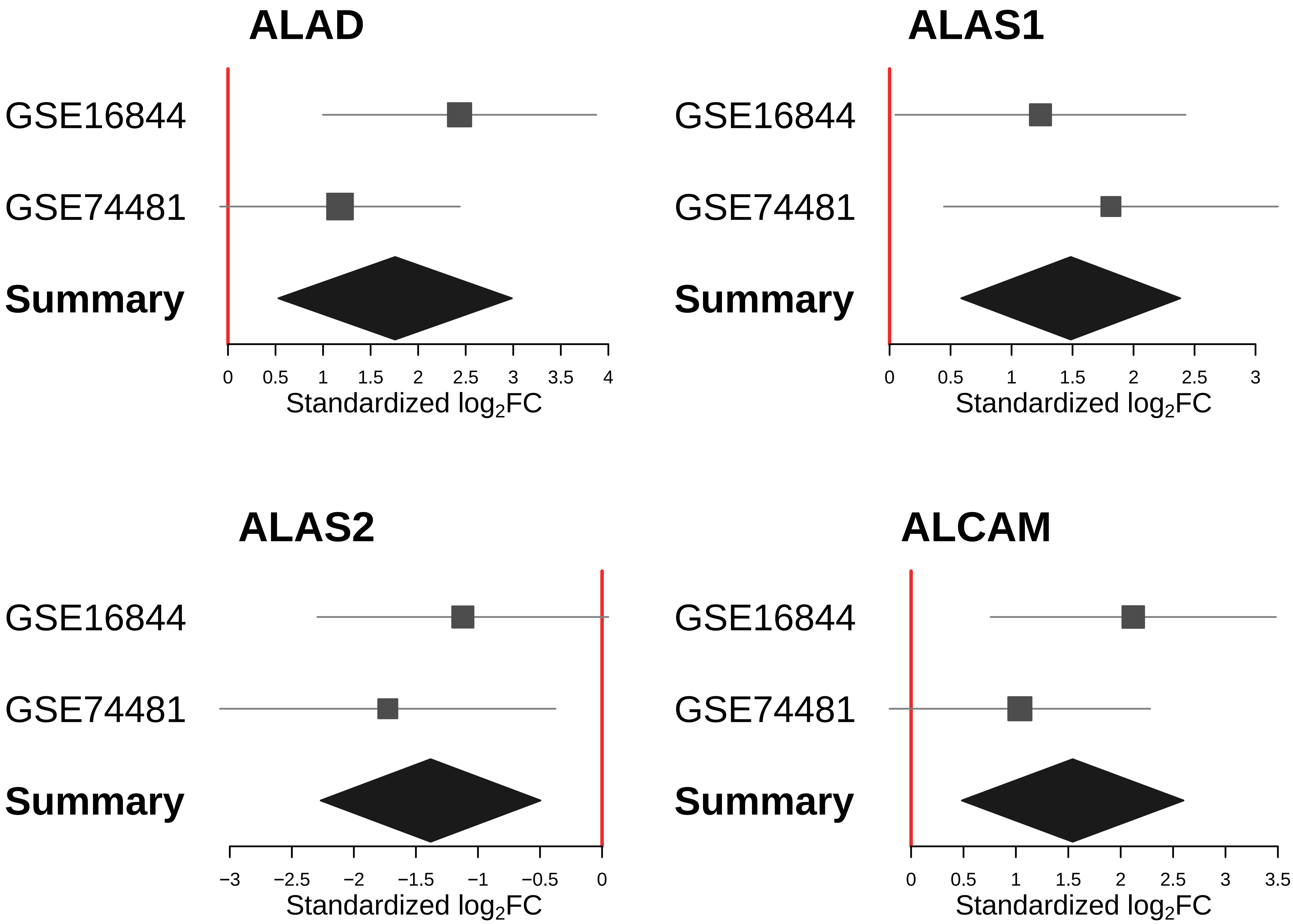
ALAD
GSE16844
GSE74481
Summary
0 0.5 1 1.5 2 2.5 3 3.5 4
Standardized log2FC
ALAS1
GSE16844
GSE74481
Summary
0 0.5 1 1.5 2 2.5 3
Standardized log2FC
ALAS2
GSE16844
GSE74481
Summary
−3 −2.5 −2 −1.5 −1 −0.5 0
Standardized log2FC
ALCAM
GSE16844
GSE74481
Summary
0 0.5 1 1.5 2 2.5 3 3.5
Standardized log2FC

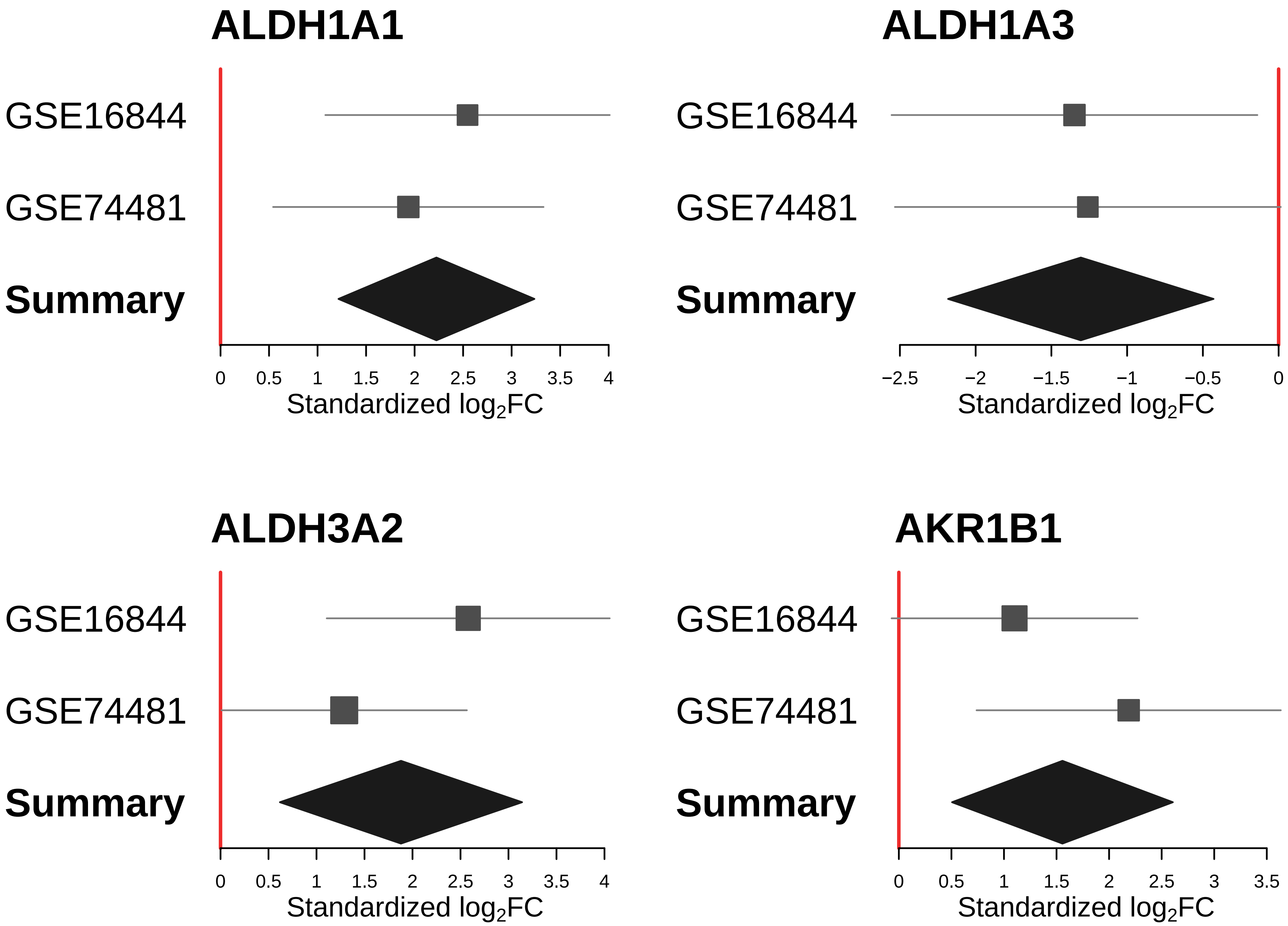
ALDH1A1
GSE16844
GSE74481
Summary
0
0.5
1
1.5
2
2.5
3
3.5
4
Standardized log2FC
ALDH1A3
GSE16844
GSE74481
Summary
−2.5
−2
−1.5
−1
−0.5
0
Standardized log2FC
ALDH3A2
GSE16844
GSE74481
Summary
0
0.5
1
1.5
2
2.5
3
3.5
4
Standardized log2FC
AKR1B1
GSE16844
GSE74481
Summary
0
0.5
1
1.5
2
2.5
3
3.5
Standardized log2FC

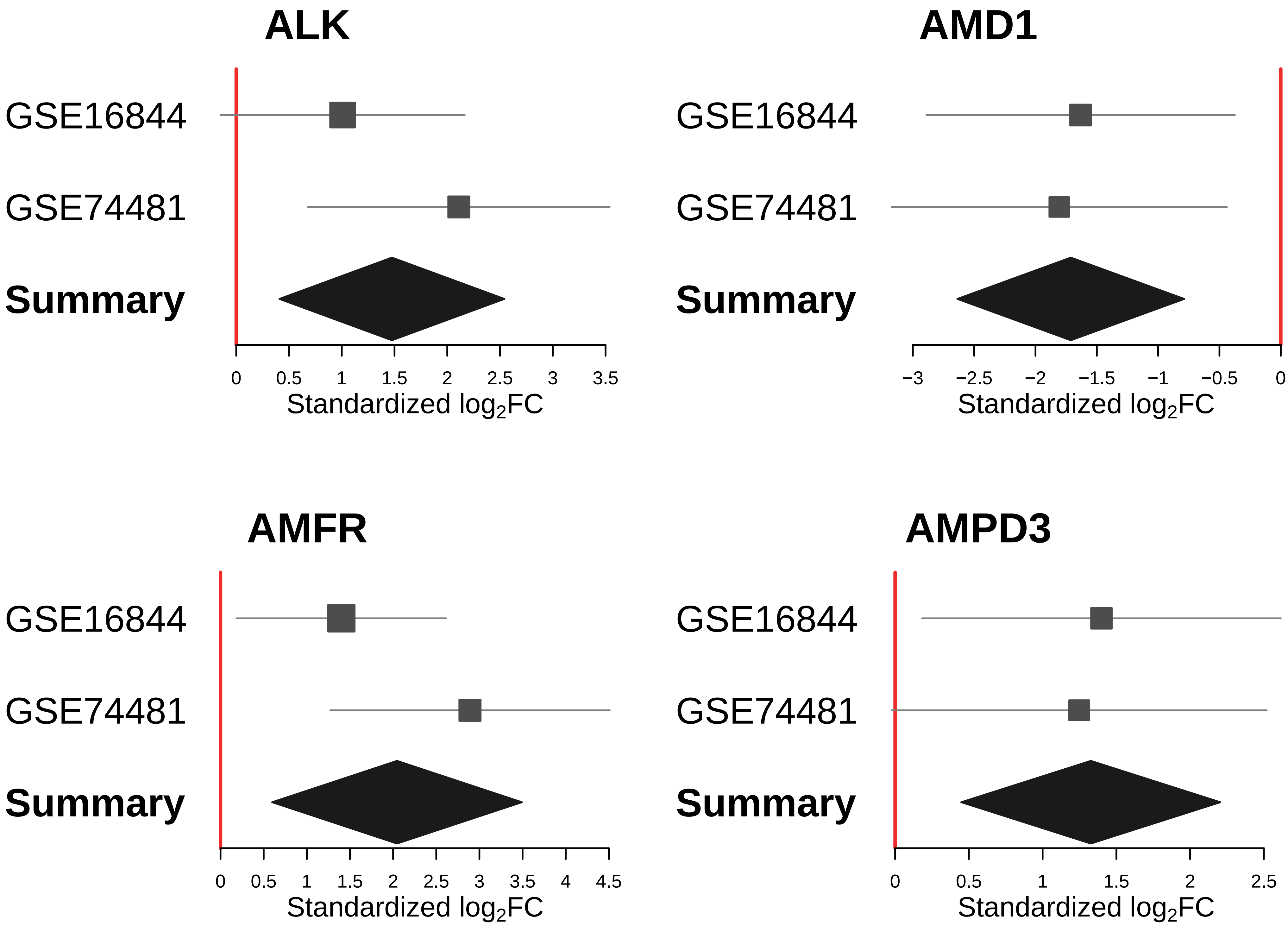
ALK
GSE16844
GSE74481
Summary
0 0.5 1 1.5 2 2.5 3 3.5
Standardized log2FC
AMD1
GSE16844
GSE74481
Summary
−3 −2.5 −2 −1.5 −1 −0.5 0
Standardized log2FC
AMFR
GSE16844
GSE74481
Summary
0 0.5 1 1.5 2 2.5 3 3.5 4 4.5
Standardized log2FC
AMPD3
GSE16844
GSE74481
Summary
0 0.5 1 1.5 2 2.5
Standardized log2FC

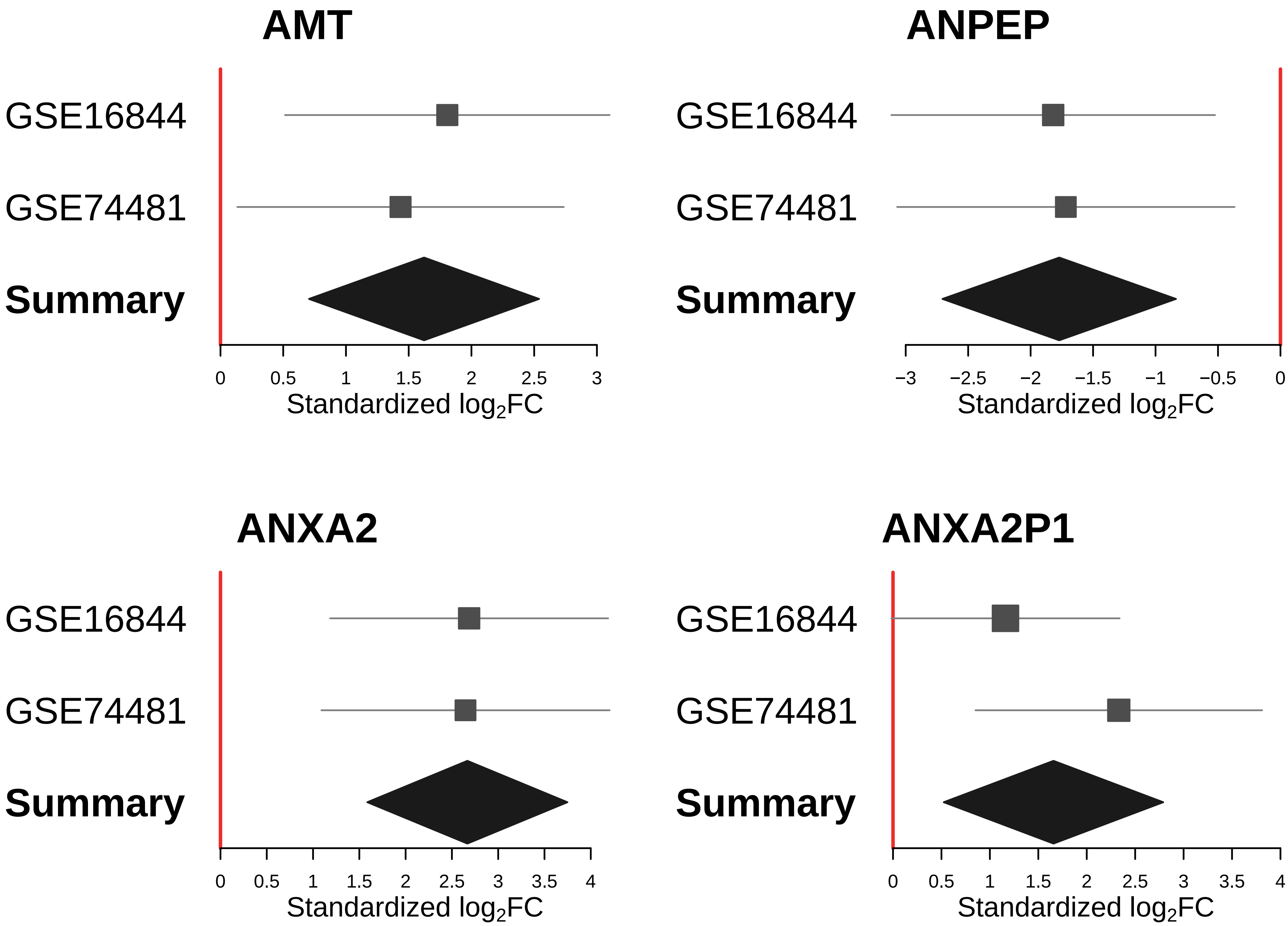
AMT
GSE16844
GSE74481
Summary
0 0.5 1 1.5 2 2.5 3
Standardized log2FC
ANPEP
GSE16844
GSE74481
Summary
−3 −2.5 −2 −1.5 −1 −0.5 0
Standardized log2FC
ANXA2
GSE16844
GSE74481
Summary
0 0.5 1 1.5 2 2.5 3 3.5 4
Standardized log2FC
ANXA2P1
GSE16844
GSE74481
Summary
0 0.5 1 1.5 2 2.5 3 3.5 4
Standardized log2FC

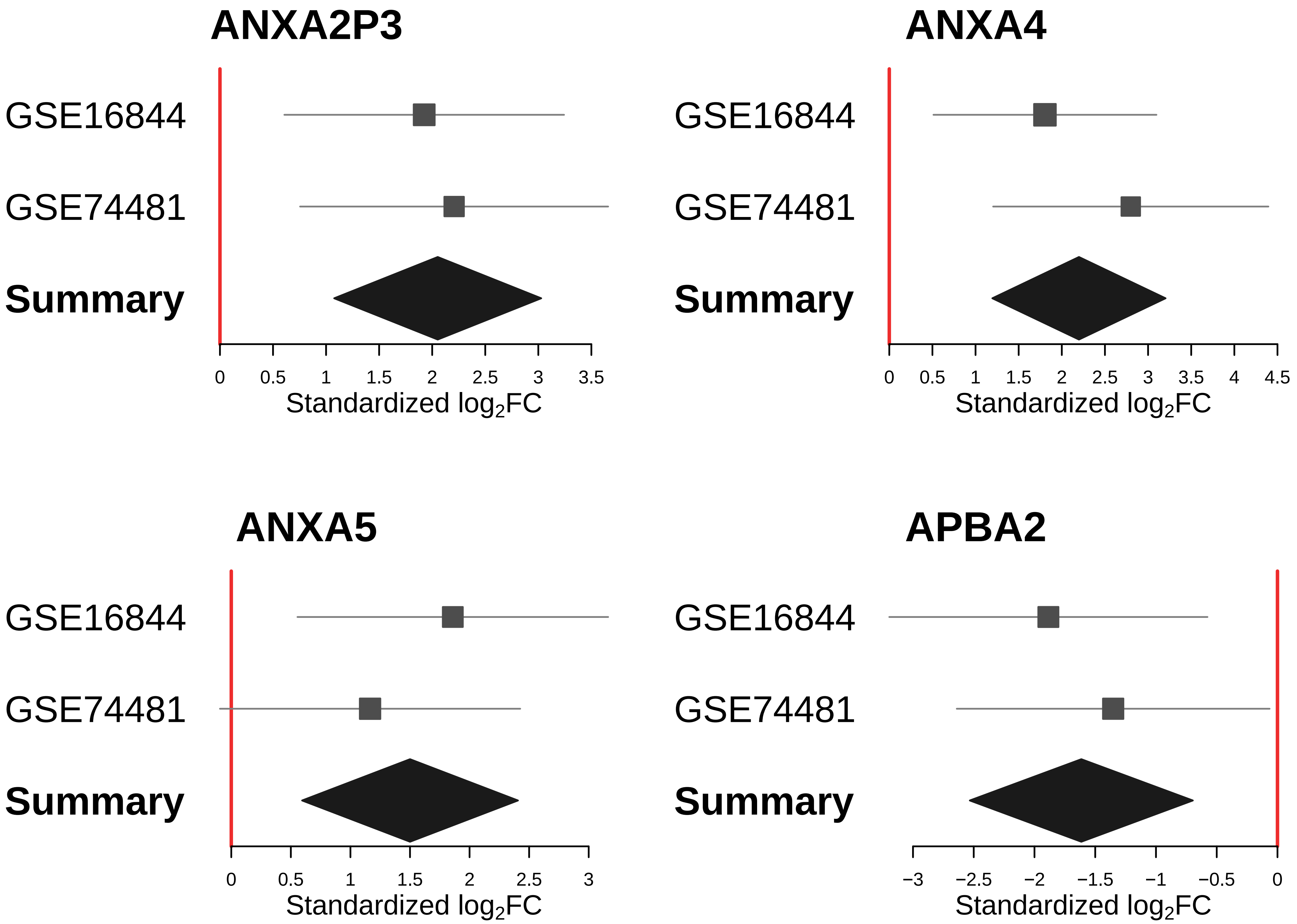
ANXA2P3
GSE16844
GSE74481
Summary
0
0.5
1
1.5
2
2.5
3
3.5
Standardized log2FC
ANXA4
GSE16844
GSE74481
Summary
0
0.5
1
1.5
2
2.5
3
3.5
4
4.5
Standardized log2FC
ANXA5
GSE16844
GSE74481
Summary
0
0.5
1
1.5
2
2.5
3
Standardized log2FC
APBA2
GSE16844
GSE74481
Summary
−3
−2.5
−2
−1.5
−1
−0.5
0
Standardized log2FC

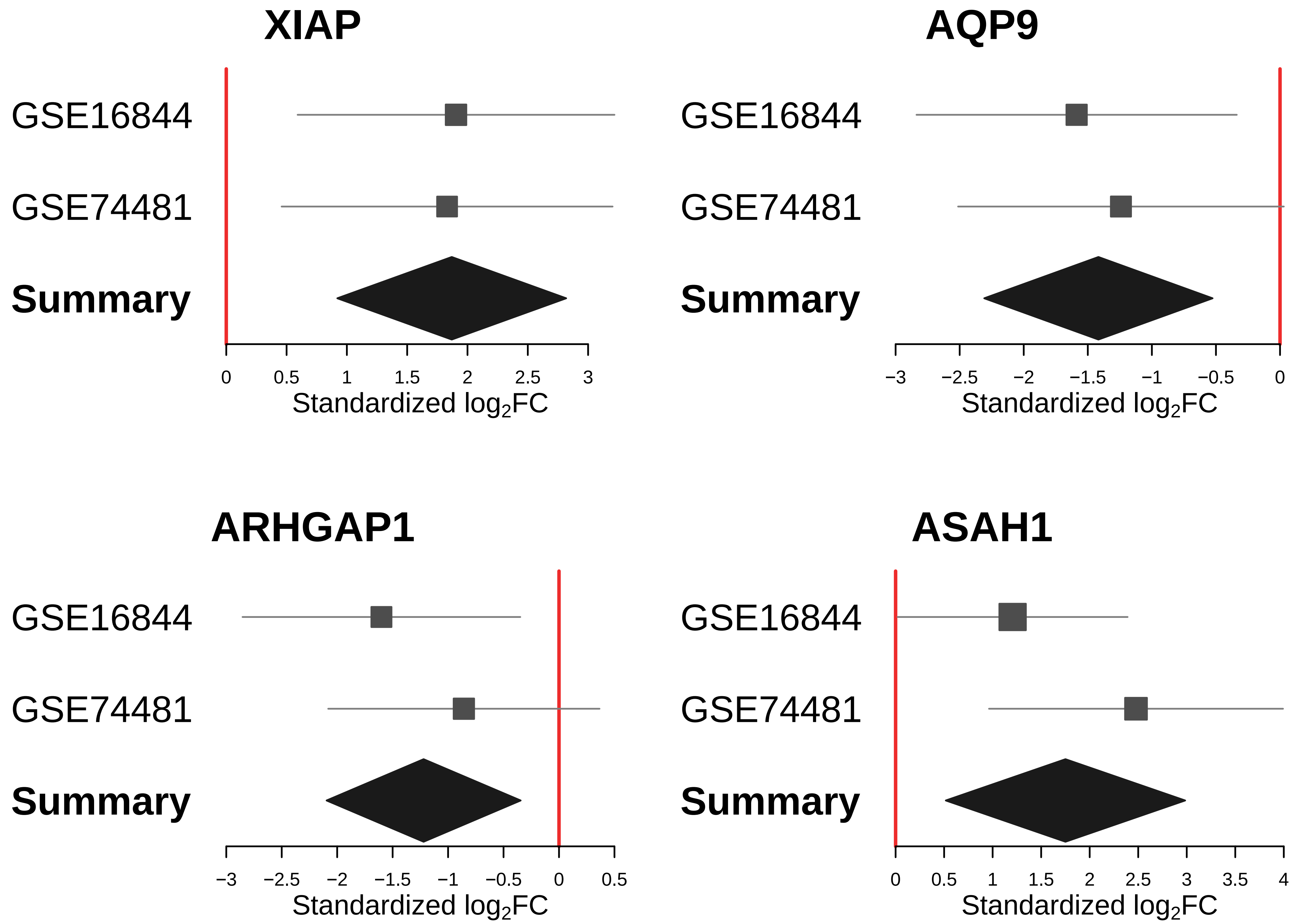

XIAP
GSE16844
GSE74481
Summary
0
0.5
1
1.5
2
2.5
3
Standardized log2FC
AQP9
GSE16844
GSE74481
Summary
−3
−2.5
−2
−1.5
−1
−0.5
0
Standardized log2FC
ARHGAP1
GSE16844
GSE74481
Summary
−3
−2.5
−2
−1.5
−1
−0.5
0
0.5
Standardized log2FC
ASAH1
GSE16844
GSE74481
Summary
0
0.5
1
1.5
2
2.5
3
3.5
4
Standardized log2FC

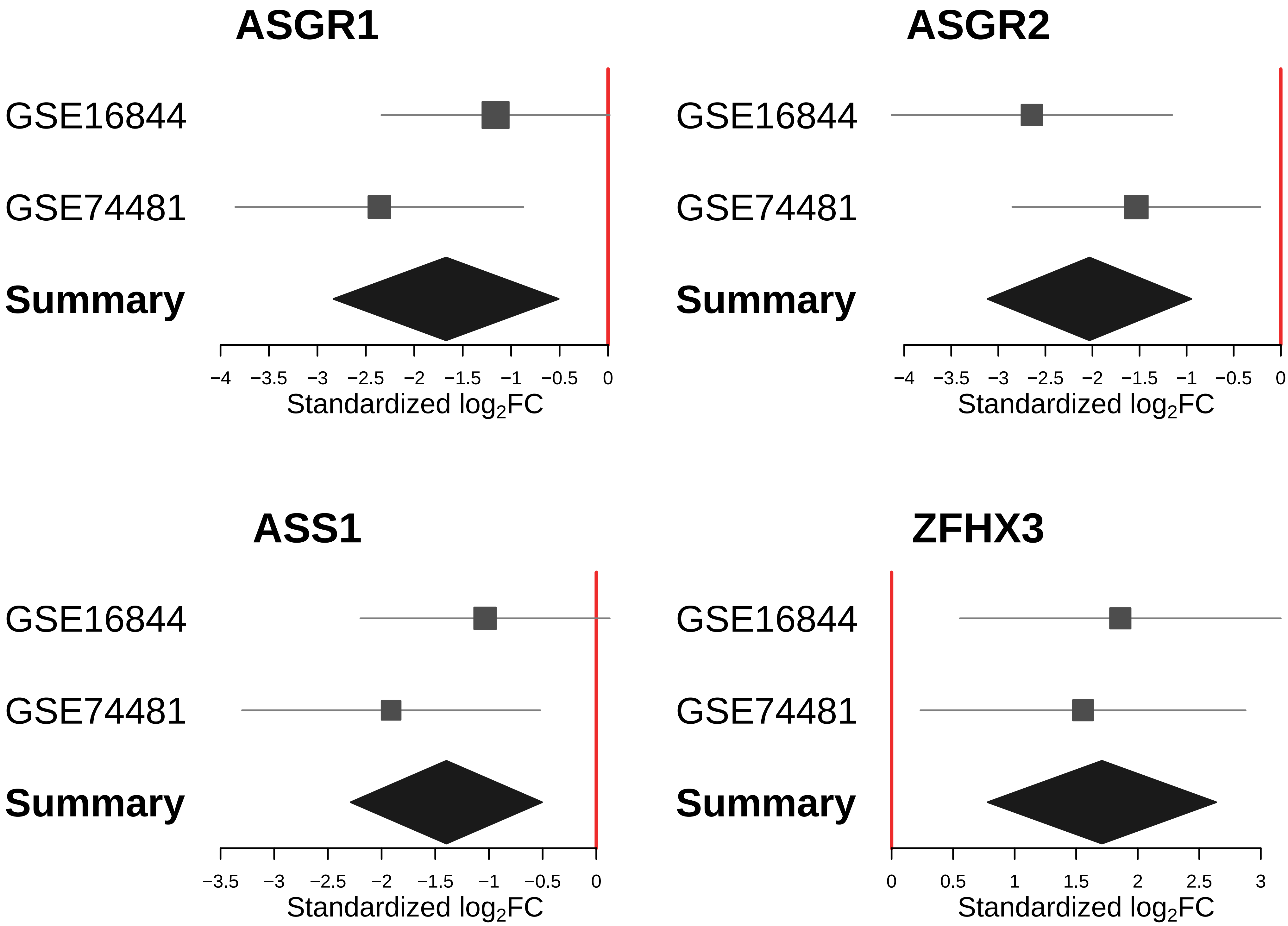
ASGR1
GSE16844
GSE74481
Summary
−4 −3.5 −3 −2.5 −2 −1.5 −1 −0.5 0
Standardized $log_2FC$
ASGR2
GSE16844
GSE74481
Summary
−4 −3.5 −3 −2.5 −2 −1.5 −1 −0.5 0
Standardized $log_2FC$
ASS1
GSE16844
GSE74481
Summary
−3.5 −3 −2.5 −2 −1.5 −1 −0.5 0
Standardized $log_2FC$
ZFHX3
GSE16844
GSE74481
Summary
0 0.5 1 1.5 2 2.5 3
Standardized $log_2FC$

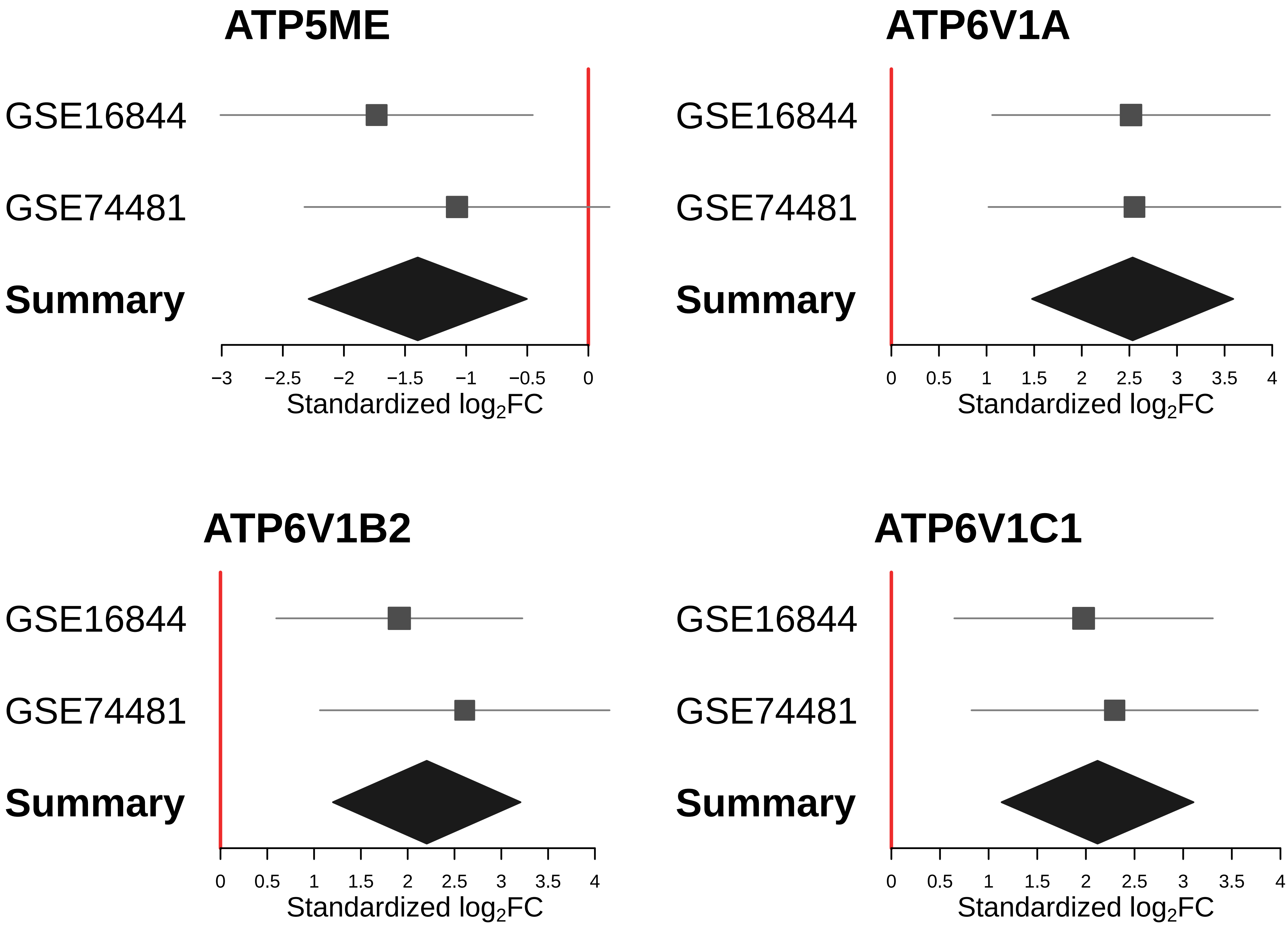
ATP5ME
GSE16844
GSE74481
Summary
−3
−2.5
−2
−1.5
−1
−0.5
0
Standardized log2FC
ATP6V1A
GSE16844
GSE74481
Summary
0
0.5
1
1.5
2
2.5
3
3.5
4
Standardized log2FC
ATP6V1B2
GSE16844
GSE74481
Summary
0
0.5
1
1.5
2
2.5
3
3.5
4
Standardized log2FC
ATP6V1C1
GSE16844
GSE74481
Summary
0
0.5
1
1.5
2
2.5
3
3.5
4
Standardized log2FC

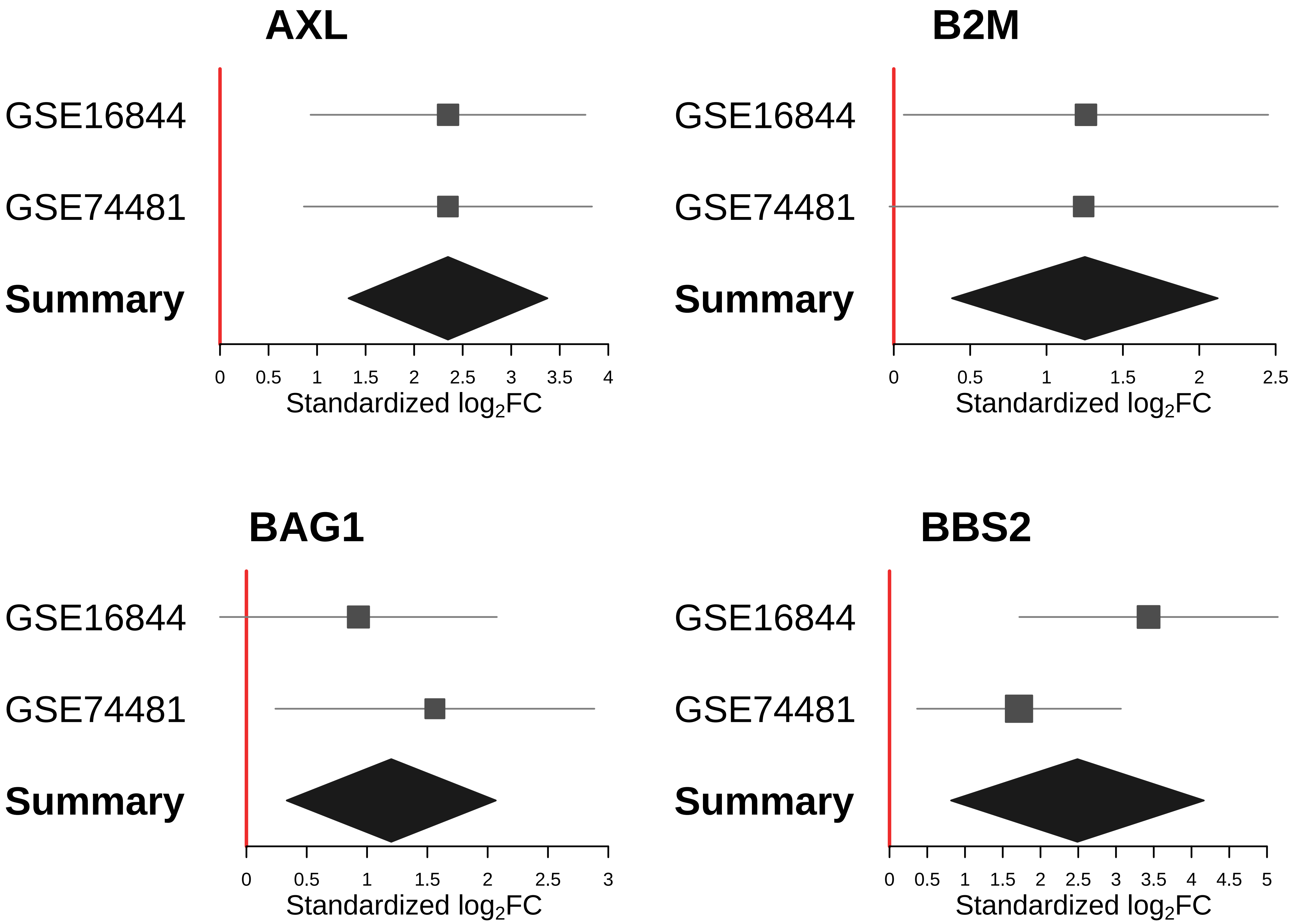
AXL
GSE16844
GSE74481
Summary
0 0.5 1 1.5 2 2.5 3 3.5 4
Standardized log2FC
B2M
GSE16844
GSE74481
Summary
0 0.5 1 1.5 2 2.5
Standardized log2FC
BAG1
GSE16844
GSE74481
Summary
0 0.5 1 1.5 2 2.5 3
Standardized log2FC
BBS2
GSE16844
GSE74481
Summary
0 0.5 1 1.5 2 2.5 3 3.5 4 4.5 5
Standardized log2FC

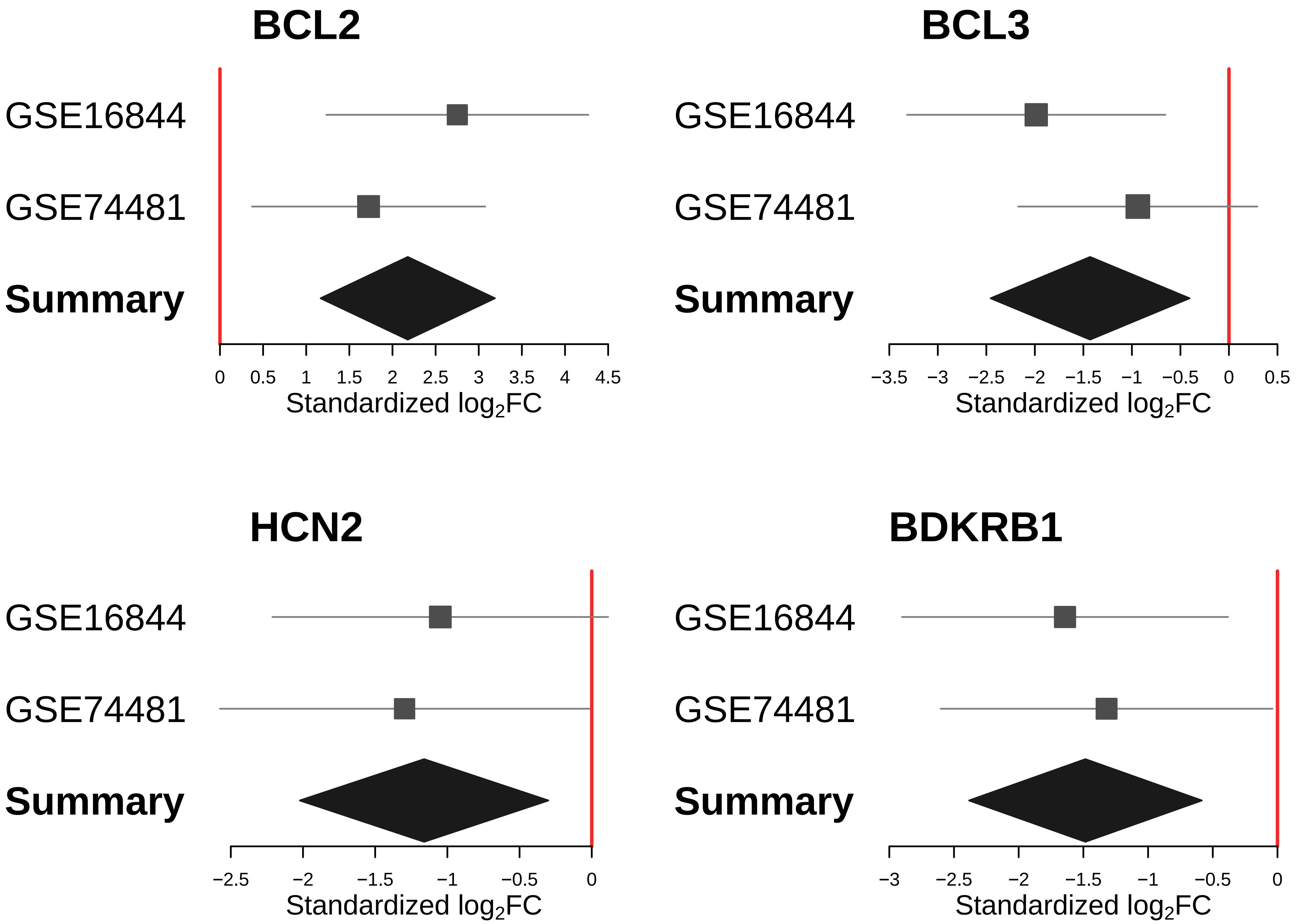
BCL2
GSE16844
GSE74481
Summary
0 0.5 1 1.5 2 2.5 3 3.5 4 4.5
Standardized log2FC
BCL3
GSE16844
GSE74481
Summary
−3.5 −3 −2.5 −2 −1.5 −1 −0.5 0 0.5
Standardized log2FC
HCN2
GSE16844
GSE74481
Summary
−2.5 −2 −1.5 −1 −0.5 0
Standardized log2FC
BDKRB1
GSE16844
GSE74481
Summary
−3 −2.5 −2 −1.5 −1 −0.5 0
Standardized log2FC

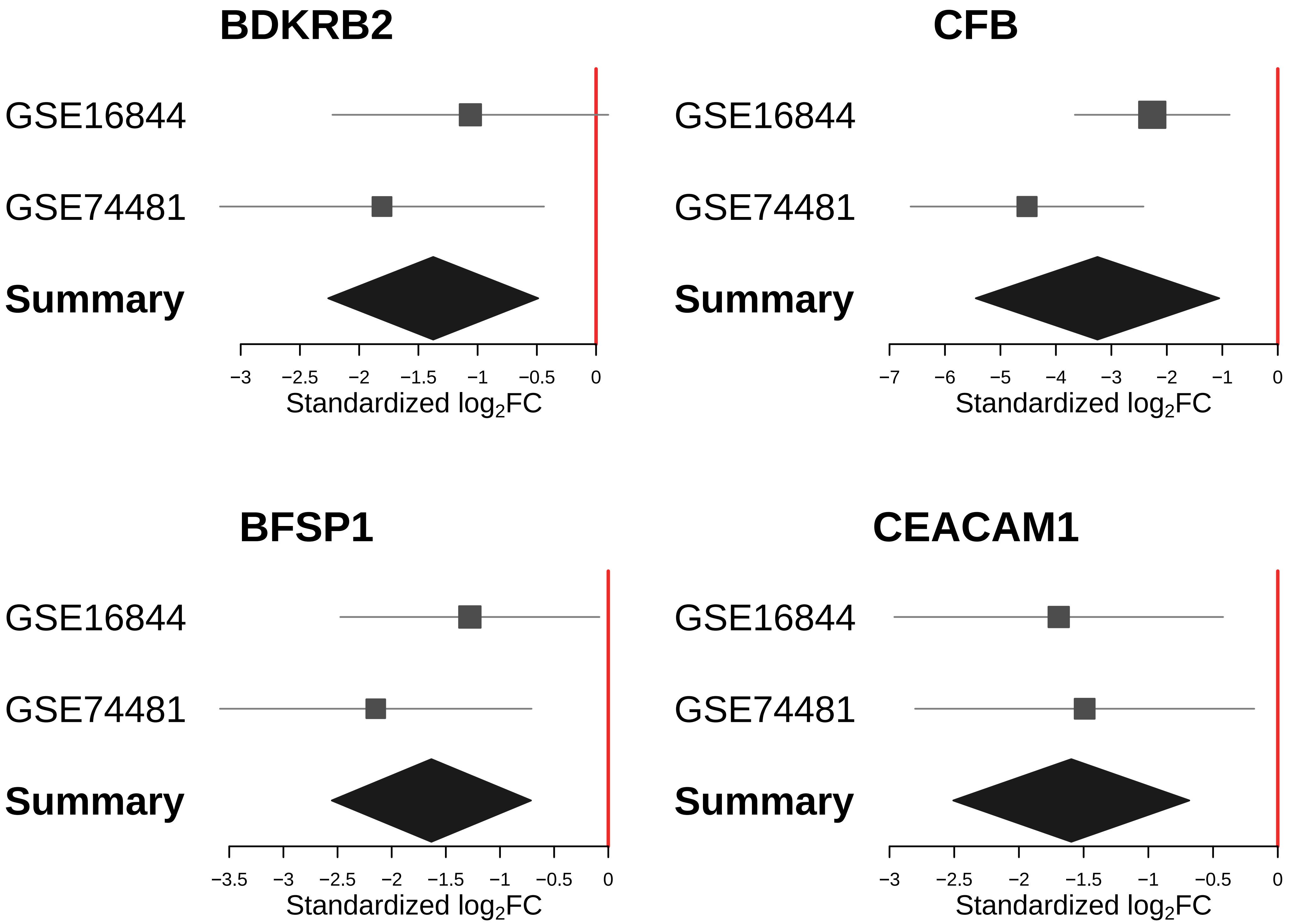
BDKRB2
GSE16844
GSE74481
Summary
−3 −2.5 −2 −1.5 −1 −0.5 0
Standardized log2FC
CFB
GSE16844
GSE74481
Summary
−7 −6 −5 −4 −3 −2 −1 0
Standardized log2FC
BFSP1
GSE16844
GSE74481
Summary
−3.5 −3 −2.5 −2 −1.5 −1 −0.5 0
Standardized log2FC
CEACAM1
GSE16844
GSE74481
Summary
−3 −2.5 −2 −1.5 −1 −0.5 0
Standardized log2FC

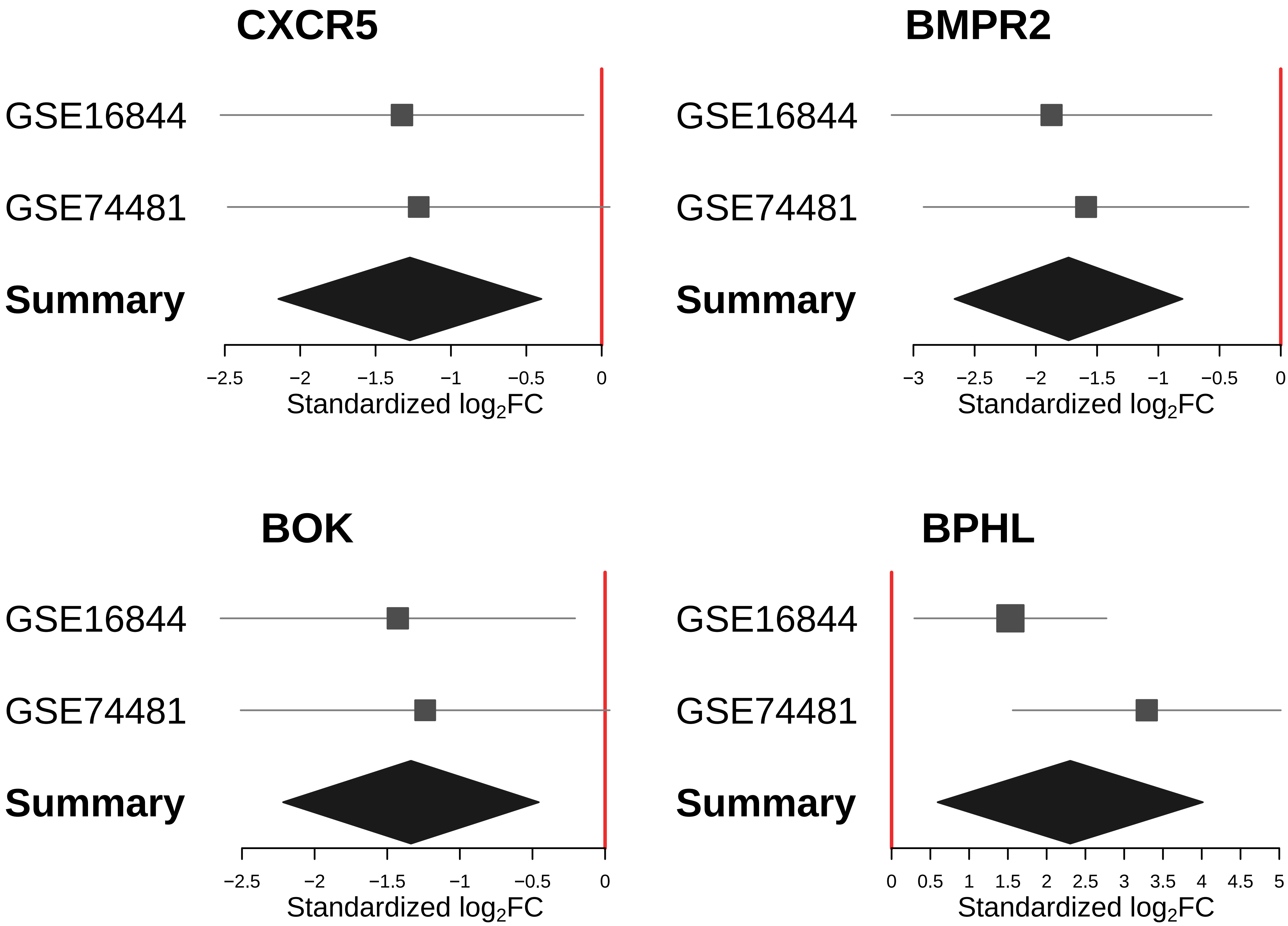

CXCR5
GSE16844
GSE74481
Summary
−2.5
−2
−1.5
−1
−0.5
0
Standardized log2FC
BMPR2
GSE16844
GSE74481
Summary
−3
−2.5
−2
−1.5
−1
−0.5
0
Standardized log2FC
BOK
GSE16844
GSE74481
Summary
−2.5
−2
−1.5
−1
−0.5
0
Standardized log2FC
BPHL
GSE16844
GSE74481
Summary
0
0.5
1
1.5
2
2.5
3
3.5
4
4.5
5
Standardized log2FC

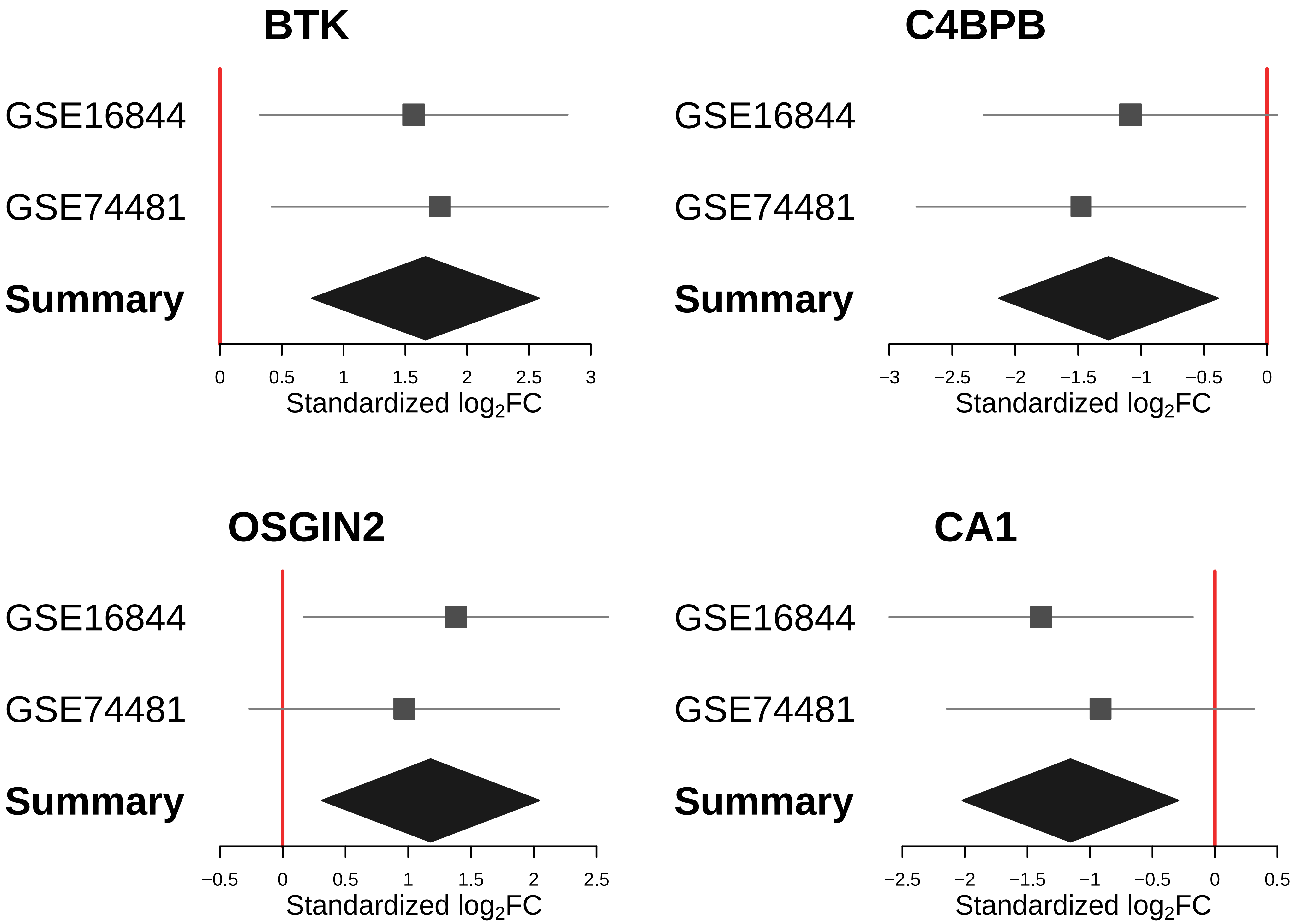

BTK
GSE16844
GSE74481
Summary
0 0.5 1 1.5 2 2.5 3
Standardized log2FC
C4BPB
GSE16844
GSE74481
Summary
−3 −2.5 −2 −1.5 −1 −0.5 0
Standardized log2FC
OSGIN2
GSE16844
GSE74481
Summary
−0.5 0 0.5 1 1.5 2 2.5
Standardized log2FC
CA1
GSE16844
GSE74481
Summary
−2.5 −2 −1.5 −1 −0.5 0 0.5
Standardized log2FC

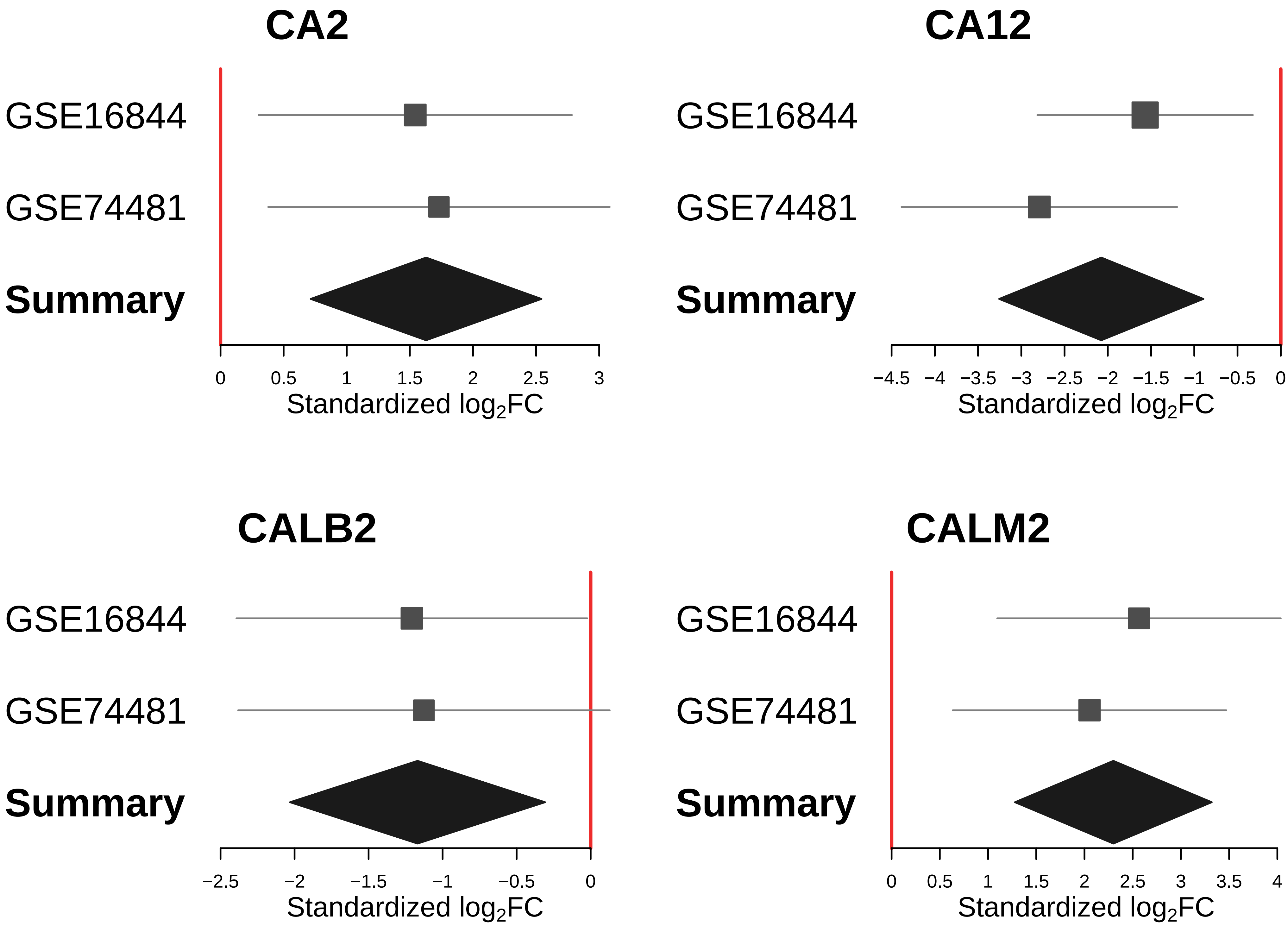

CA2
GSE16844
GSE74481
Summary
0
0.5
1
1.5
2
2.5
3
Standardized log2FC
CA12
GSE16844
GSE74481
Summary
−4.5
−4
−3.5
−3
−2.5
−2
−1.5
−1
−0.5
0
Standardized log2FC
CALB2
GSE16844
GSE74481
Summary
−2.5
−2
−1.5
−1
−0.5
0
Standardized log2FC
CALM2
GSE16844
GSE74481
Summary
0
0.5
1
1.5
2
2.5
3
3.5
4
Standardized log2FC

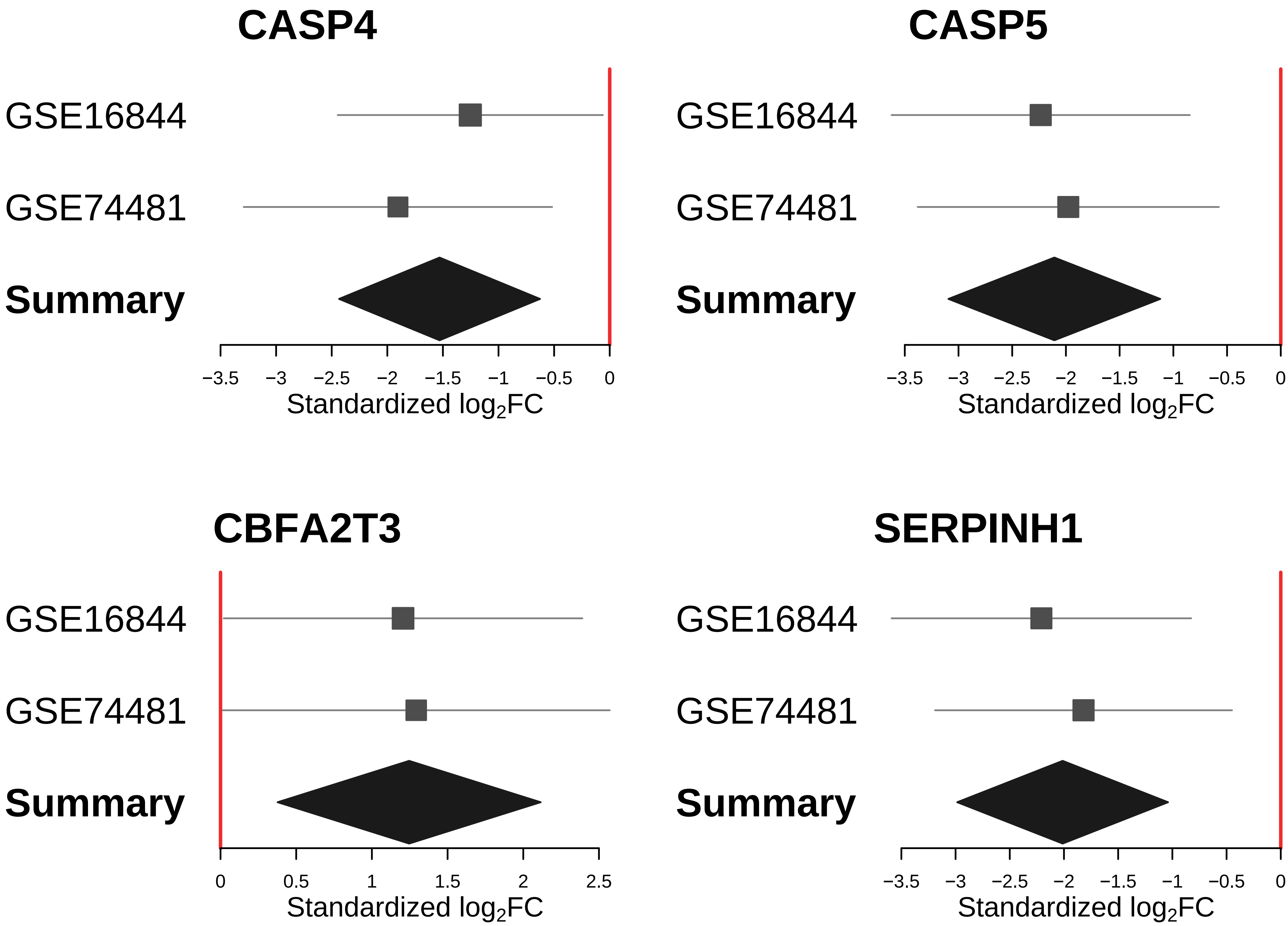

CASP4
GSE16844
GSE74481
Summary
−3.5
−3
−2.5
−2
−1.5
−1
−0.5
0
Standardized log2FC
CASP5
GSE16844
GSE74481
Summary
−3.5
−3
−2.5
−2
−1.5
−1
−0.5
0
Standardized log2FC
CBFA2T3
GSE16844
GSE74481
Summary
0
0.5
1
1.5
2
2.5
Standardized log2FC
SERPINH1
GSE16844
GSE74481
Summary
−3.5
−3
−2.5
−2
−1.5
−1
−0.5
0
Standardized log2FC

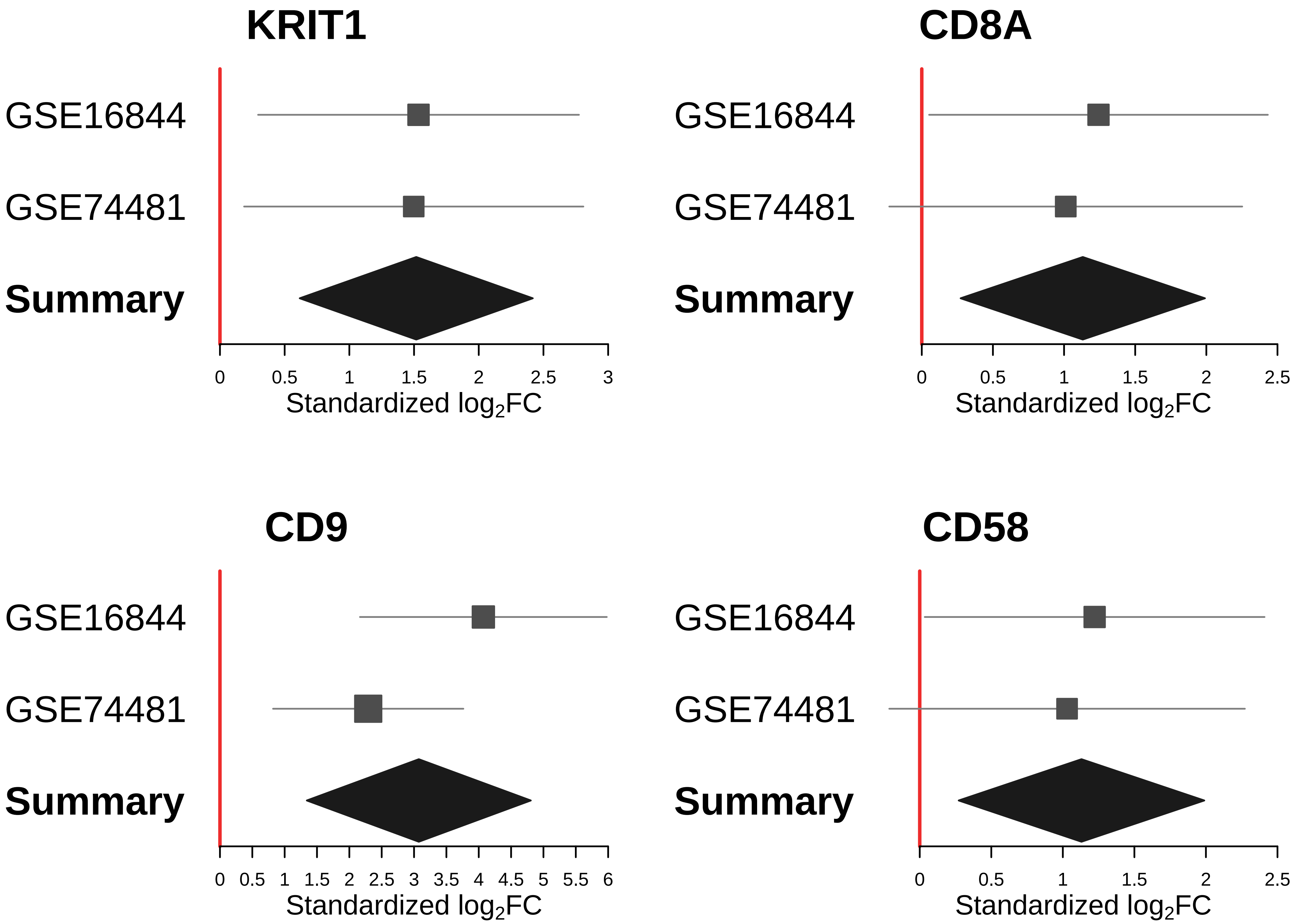
KRIT1
GSE16844
GSE74481
Summary
0 0.5 1 1.5 2 2.5 3
Standardized $log_2FC$
CD8A
GSE16844
GSE74481
Summary
0 0.5 1 1.5 2 2.5
Standardized $log_2FC$
CD9
GSE16844
GSE74481
Summary
0 0.5 1 1.5 2 2.5 3 3.5 4 4.5 5 5.5 6
Standardized $log_2FC$
CD58
GSE16844
GSE74481
Summary
0 0.5 1 1.5 2 2.5
Standardized $log_2FC$

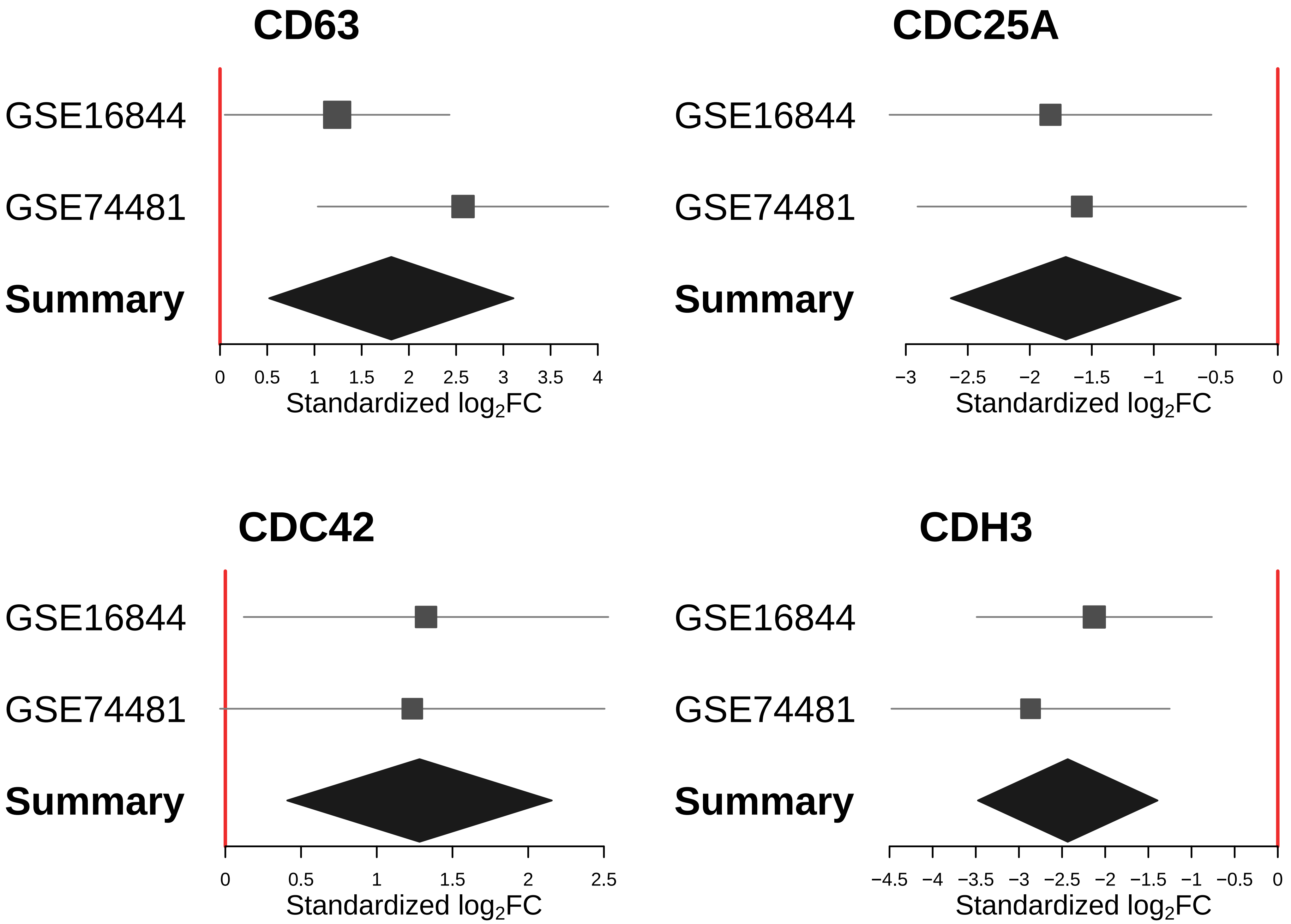
CD63
GSE16844
GSE74481
Summary
0 0.5 1 1.5 2 2.5 3 3.5 4
Standardized log2FC
CDC25A
GSE16844
GSE74481
Summary
−3 −2.5 −2 −1.5 −1 −0.5 0
Standardized log2FC
CDC42
GSE16844
GSE74481
Summary
0 0.5 1 1.5 2 2.5
Standardized log2FC
CDH3
GSE16844
GSE74481
Summary
−4.5 −4 −3.5 −3 −2.5 −2 −1.5 −1 −0.5 0
Standardized log2FC

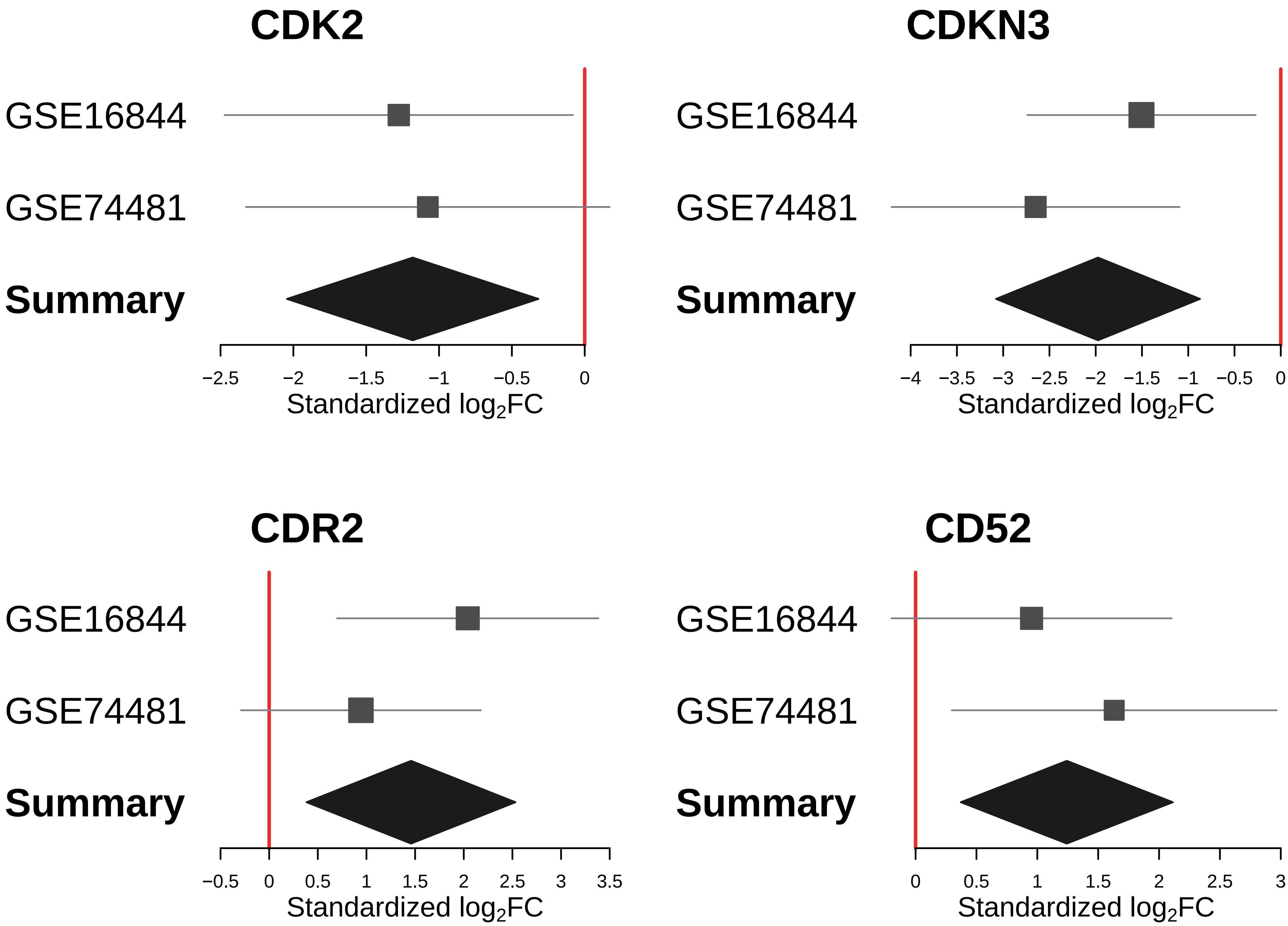
CDK2
GSE16844
GSE74481
Summary
−2.5 −2 −1.5 −1 −0.5 0
Standardized $log_2FC$
CDKN3
GSE16844
GSE74481
Summary
−4 −3.5 −3 −2.5 −2 −1.5 −1 −0.5 0
Standardized $log_2FC$
CDR2
GSE16844
GSE74481
Summary
−0.5 0 0.5 1 1.5 2 2.5 3 3.5
Standardized $log_2FC$
CD52
GSE16844
GSE74481
Summary
0 0.5 1 1.5 2 2.5 3
Standardized $log_2FC$

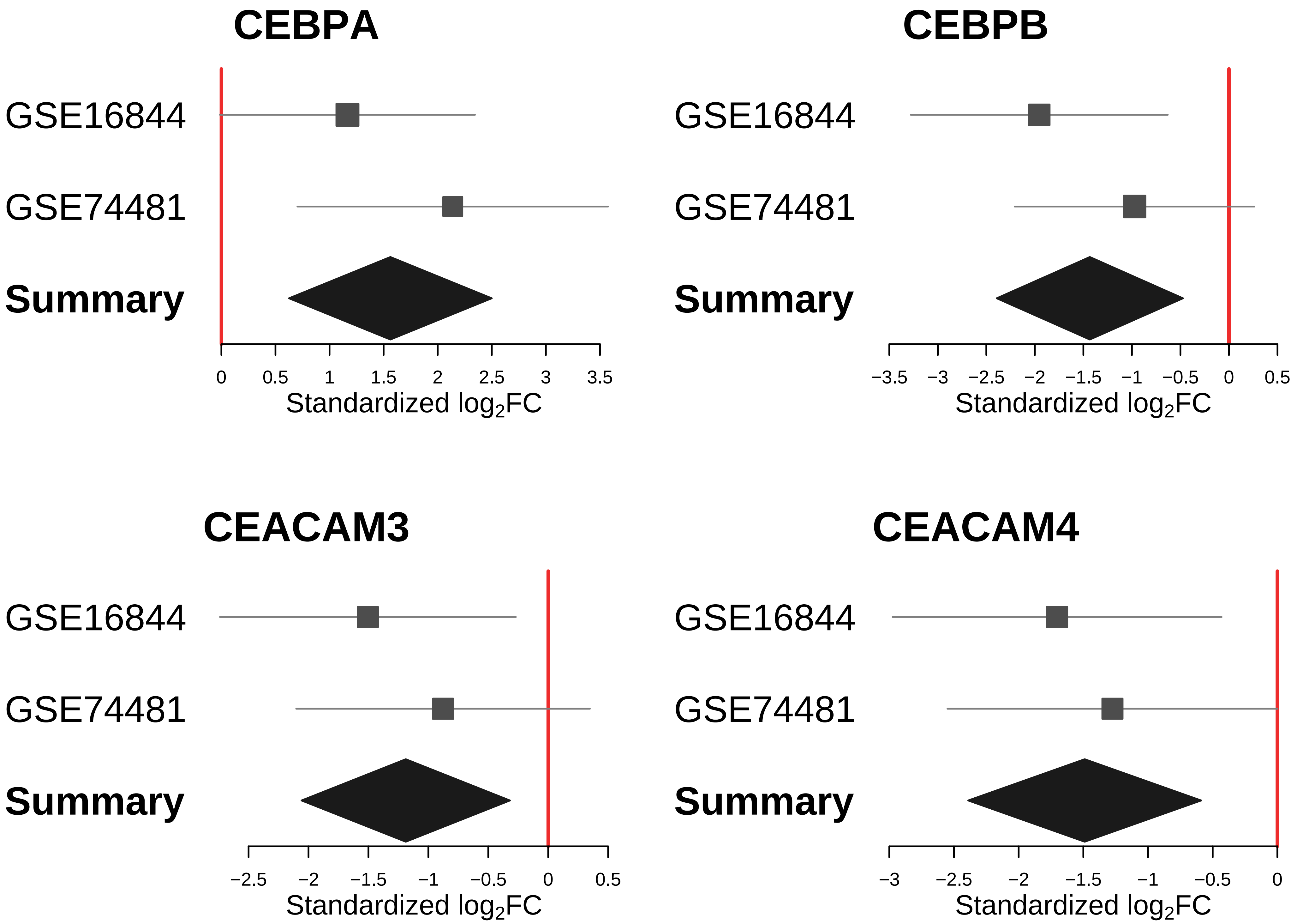
CEBPA
GSE16844
GSE74481
Summary
0 0.5 1 1.5 2 2.5 3 3.5
Standardized log2FC
CEBPB
GSE16844
GSE74481
Summary
−3.5 −3 −2.5 −2 −1.5 −1 −0.5 0 0.5
Standardized log2FC
CEACAM3
GSE16844
GSE74481
Summary
−2.5 −2 −1.5 −1 −0.5 0 0.5
Standardized log2FC
CEACAM4
GSE16844
GSE74481
Summary
−3 −2.5 −2 −1.5 −1 −0.5 0
Standardized log2FC

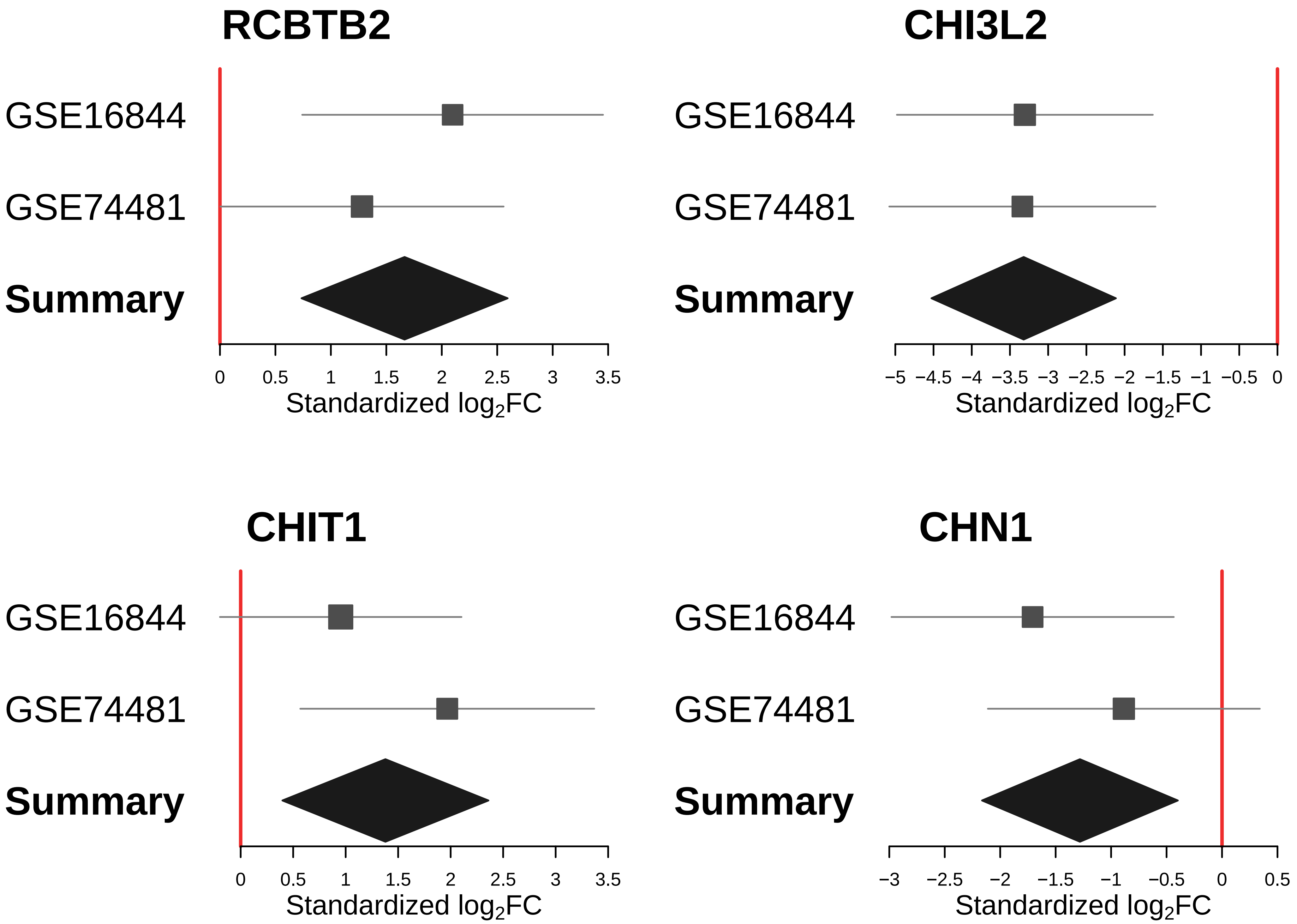

RCBTB2
GSE16844
GSE74481
Summary
0
0.5
1
1.5
2
2.5
3
3.5
Standardized log2FC
CHI3L2
GSE16844
GSE74481
Summary
−5
−4.5
−4
−3.5
−3
−2.5
−2
−1.5
−1
−0.5
0
Standardized log2FC
CHIT1
GSE16844
GSE74481
Summary
0
0.5
1
1.5
2
2.5
3
3.5
Standardized log2FC
CHN1
GSE16844
GSE74481
Summary
−3
−2.5
−2
−1.5
−1
−0.5
0
0.5
Standardized log2FC

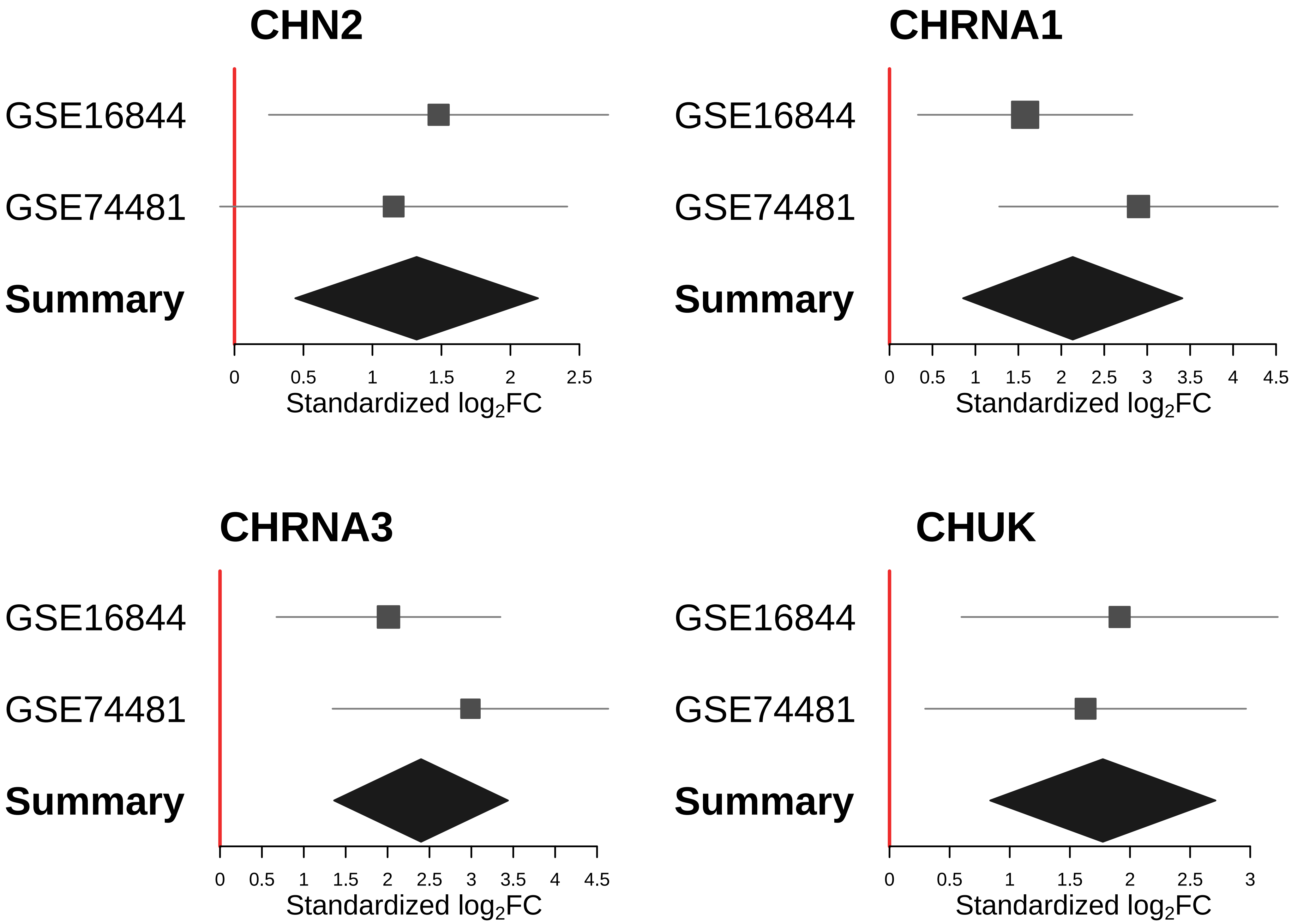

CHN2
GSE16844
GSE74481
Summary
0
0.5
1
1.5
2
2.5
Standardized log2FC
CHRNA1
GSE16844
GSE74481
Summary
0
0.5
1
1.5
2
2.5
3
3.5
4
4.5
Standardized log2FC
CHRNA3
GSE16844
GSE74481
Summary
0
0.5
1
1.5
2
2.5
3
3.5
4
4.5
Standardized log2FC
CHUK
GSE16844
GSE74481
Summary
0
0.5
1
1.5
2
2.5
3
Standardized log2FC

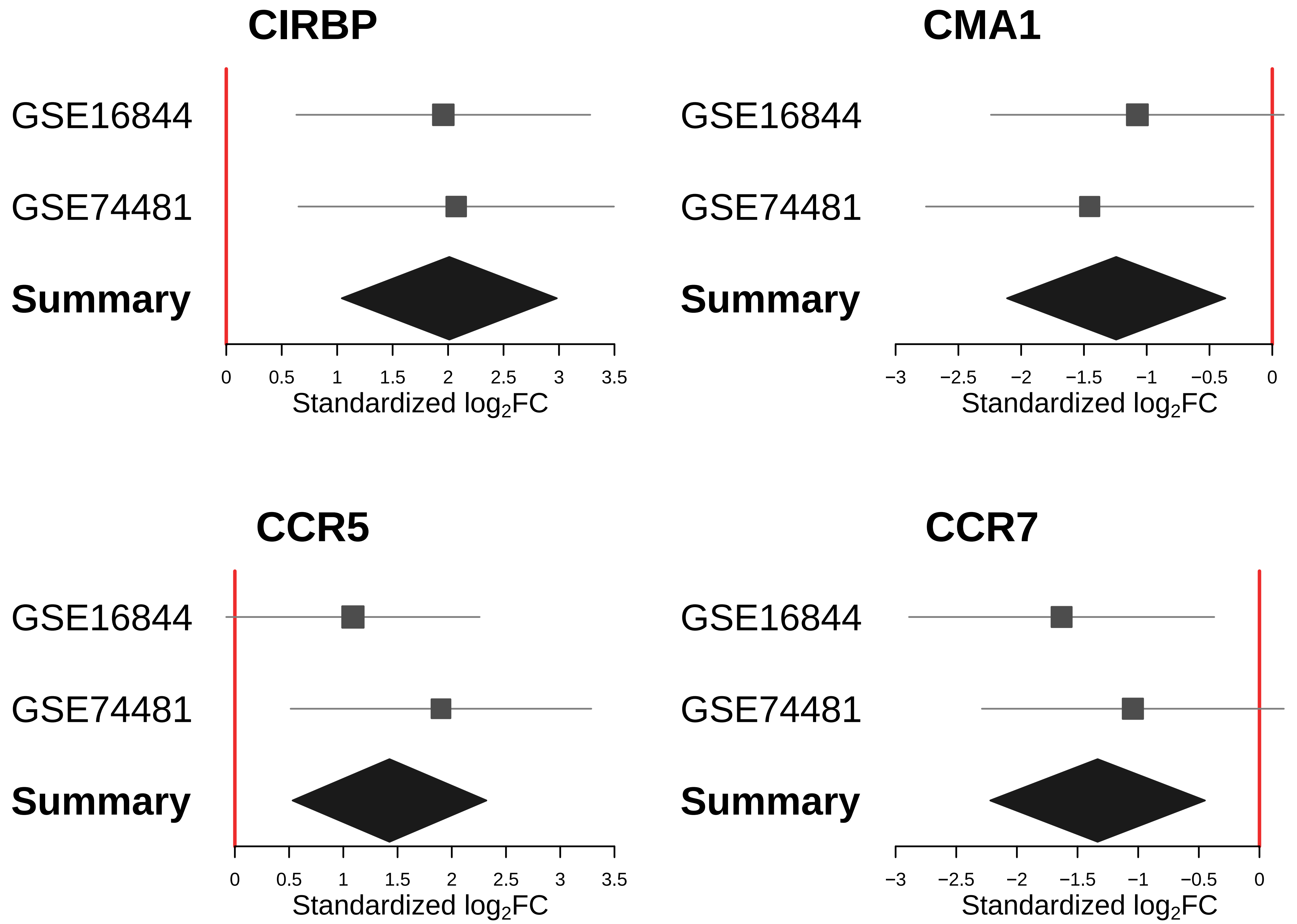
CIRBP
GSE16844
GSE74481
Summary
0 0.5 1 1.5 2 2.5 3 3.5
Standardized log2FC
CMA1
GSE16844
GSE74481
Summary
−3 −2.5 −2 −1.5 −1 −0.5 0
Standardized log2FC
CCR5
GSE16844
GSE74481
Summary
0 0.5 1 1.5 2 2.5 3 3.5
Standardized log2FC
CCR7
GSE16844
GSE74481
Summary
−3 −2.5 −2 −1.5 −1 −0.5 0
Standardized log2FC

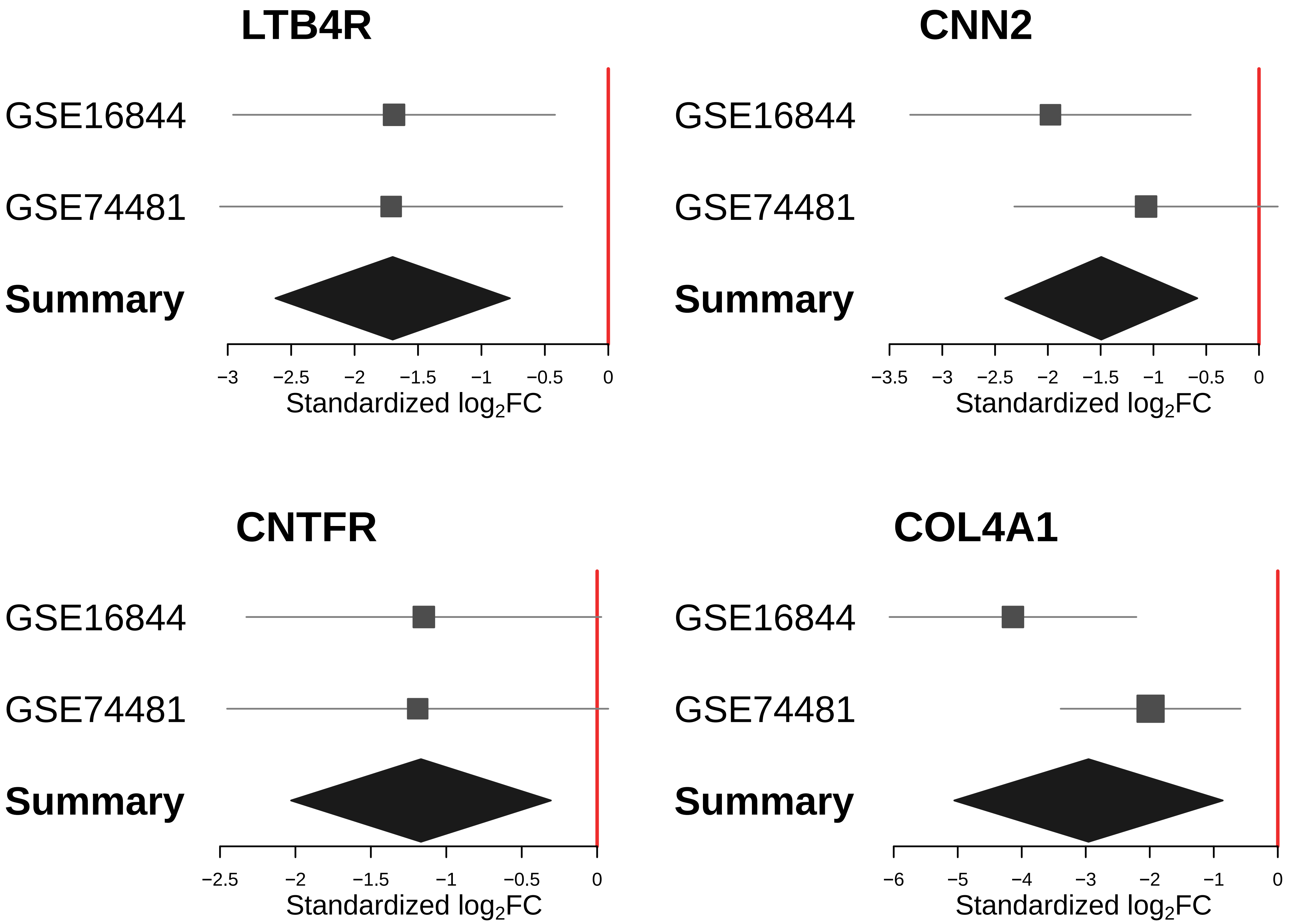

LTB4R
GSE16844
GSE74481
Summary
−3
−2.5
−2
−1.5
−1
−0.5
0
Standardized $log_2FC$
CNN2
GSE16844
GSE74481
Summary
−3.5
−3
−2.5
−2
−1.5
−1
−0.5
0
Standardized $log_2FC$
CNTFR
GSE16844
GSE74481
Summary
−2.5
−2
−1.5
−1
−0.5
0
Standardized $log_2FC$
COL4A1
GSE16844
GSE74481
Summary
−6
−5
−4
−3
−2
−1
0
Standardized $log_2FC$

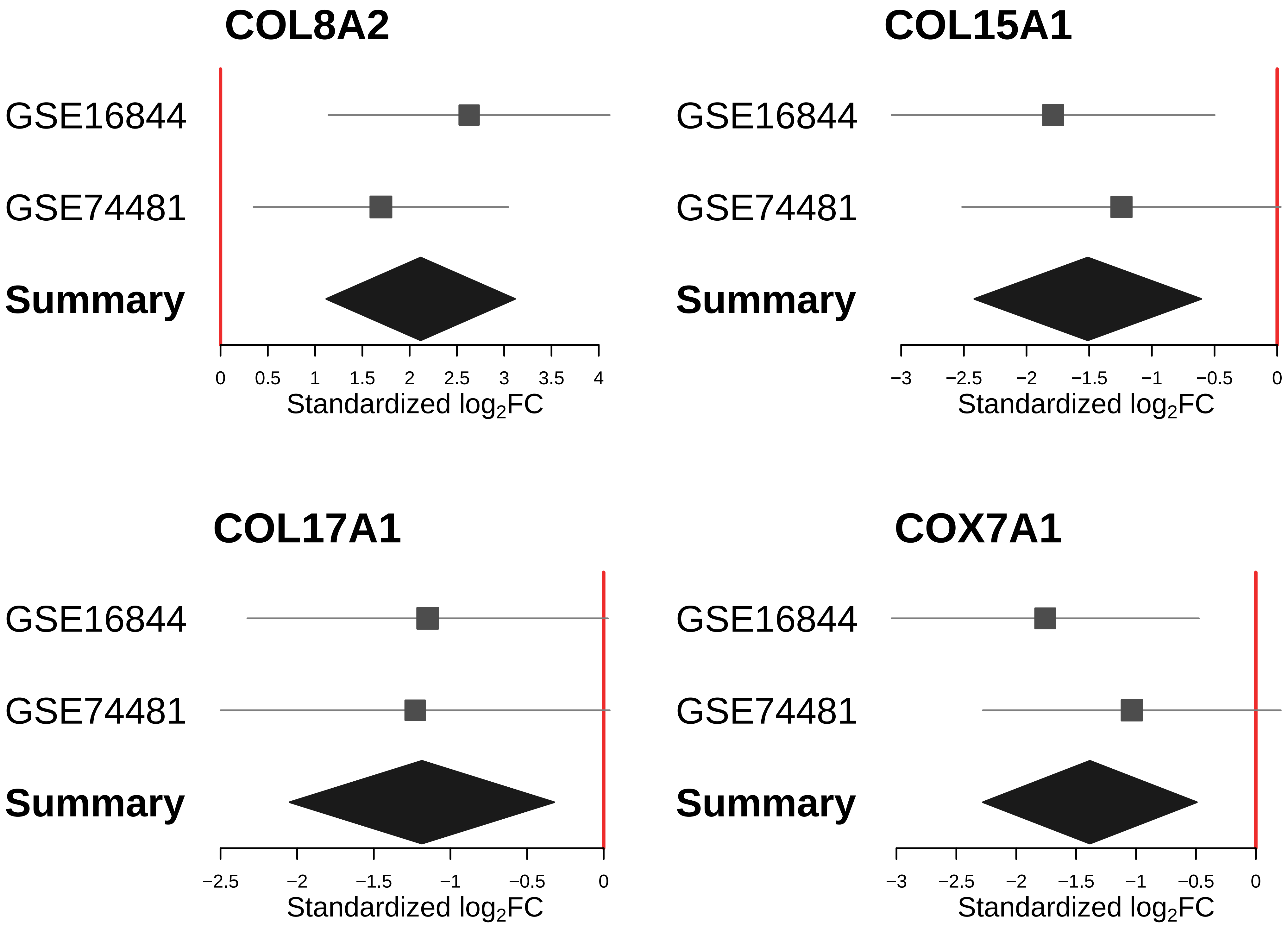
COL8A2
GSE16844
GSE74481
Summary
0 0.5 1 1.5 2 2.5 3 3.5 4
Standardized log2FC
COL15A1
GSE16844
GSE74481
Summary
−3 −2.5 −2 −1.5 −1 −0.5 0
Standardized log2FC
COL17A1
GSE16844
GSE74481
Summary
−2.5 −2 −1.5 −1 −0.5 0
Standardized log2FC
COX7A1
GSE16844
GSE74481
Summary
−3 −2.5 −2 −1.5 −1 −0.5 0
Standardized log2FC

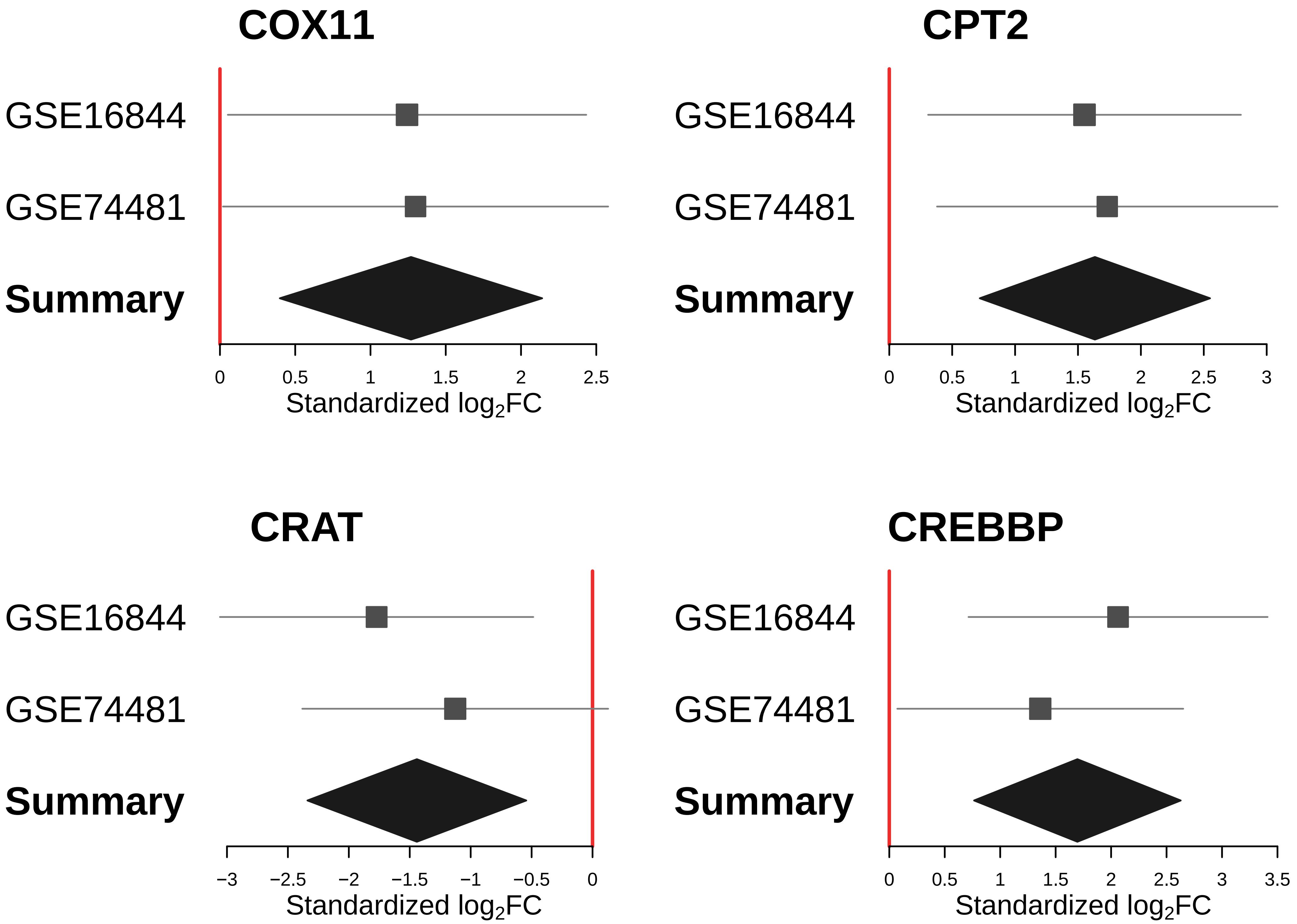

COX11
GSE16844
GSE74481
Summary
0
0.5
1
1.5
2
2.5
Standardized log2FC
CPT2
GSE16844
GSE74481
Summary
0
0.5
1
1.5
2
2.5
3
Standardized log2FC
CRAT
GSE16844
GSE74481
Summary
−3
−2.5
−2
−1.5
−1
−0.5
0
Standardized log2FC
CREBBP
GSE16844
GSE74481
Summary
0
0.5
1
1.5
2
2.5
3
3.5
Standardized log2FC

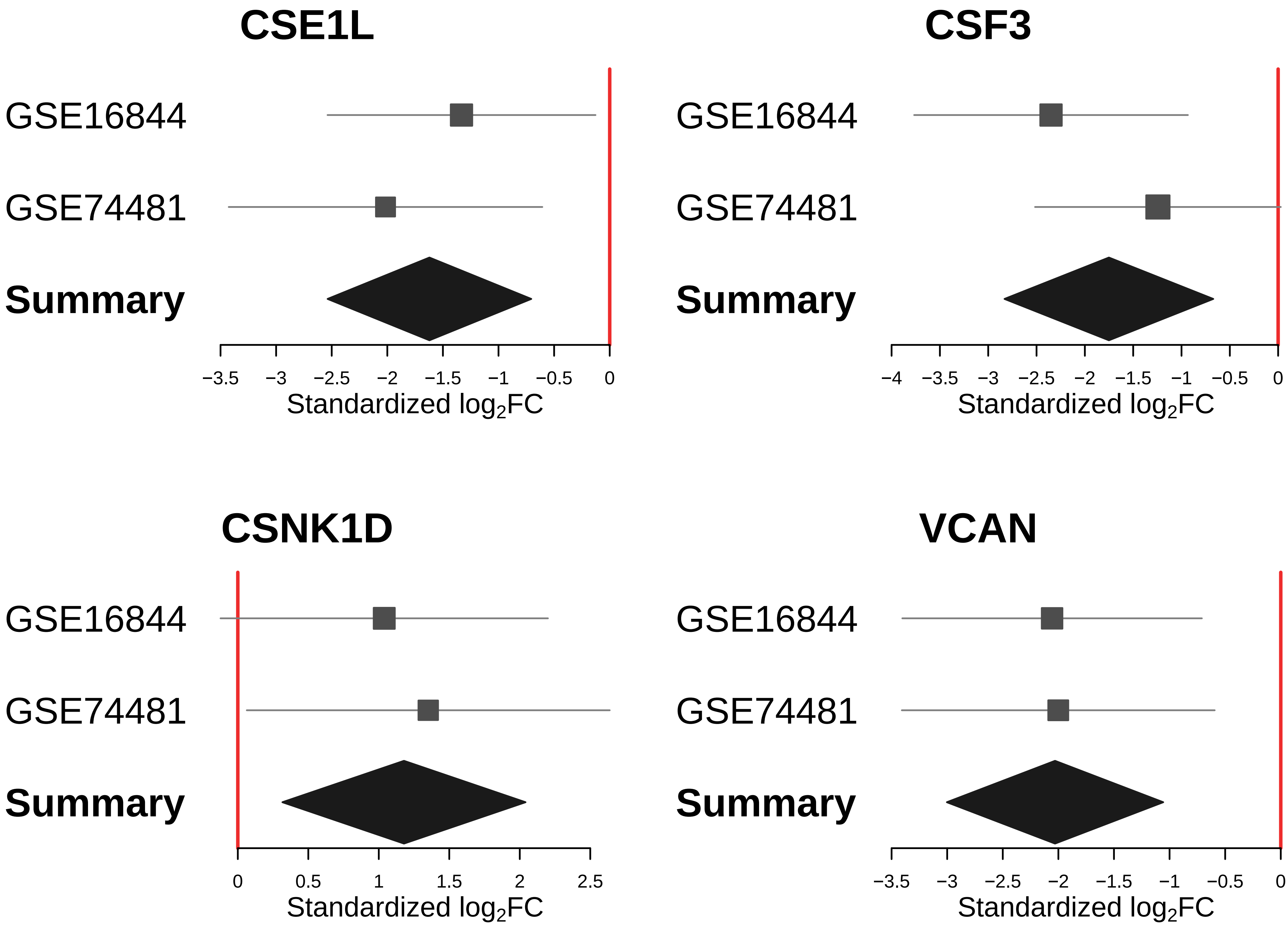
CSE1L
GSE16844
GSE74481
Summary
−3.5
−3
−2.5
−2
−1.5
−1
−0.5
0
Standardized log2FC
CSF3
GSE16844
GSE74481
Summary
−4
−3.5
−3
−2.5
−2
−1.5
−1
−0.5
0
Standardized log2FC
CSNK1D
GSE16844
GSE74481
Summary
0
0.5
1
1.5
2
2.5
Standardized log2FC
VCAN
GSE16844
GSE74481
Summary
−3.5
−3
−2.5
−2
−1.5
−1
−0.5
0
Standardized log2FC

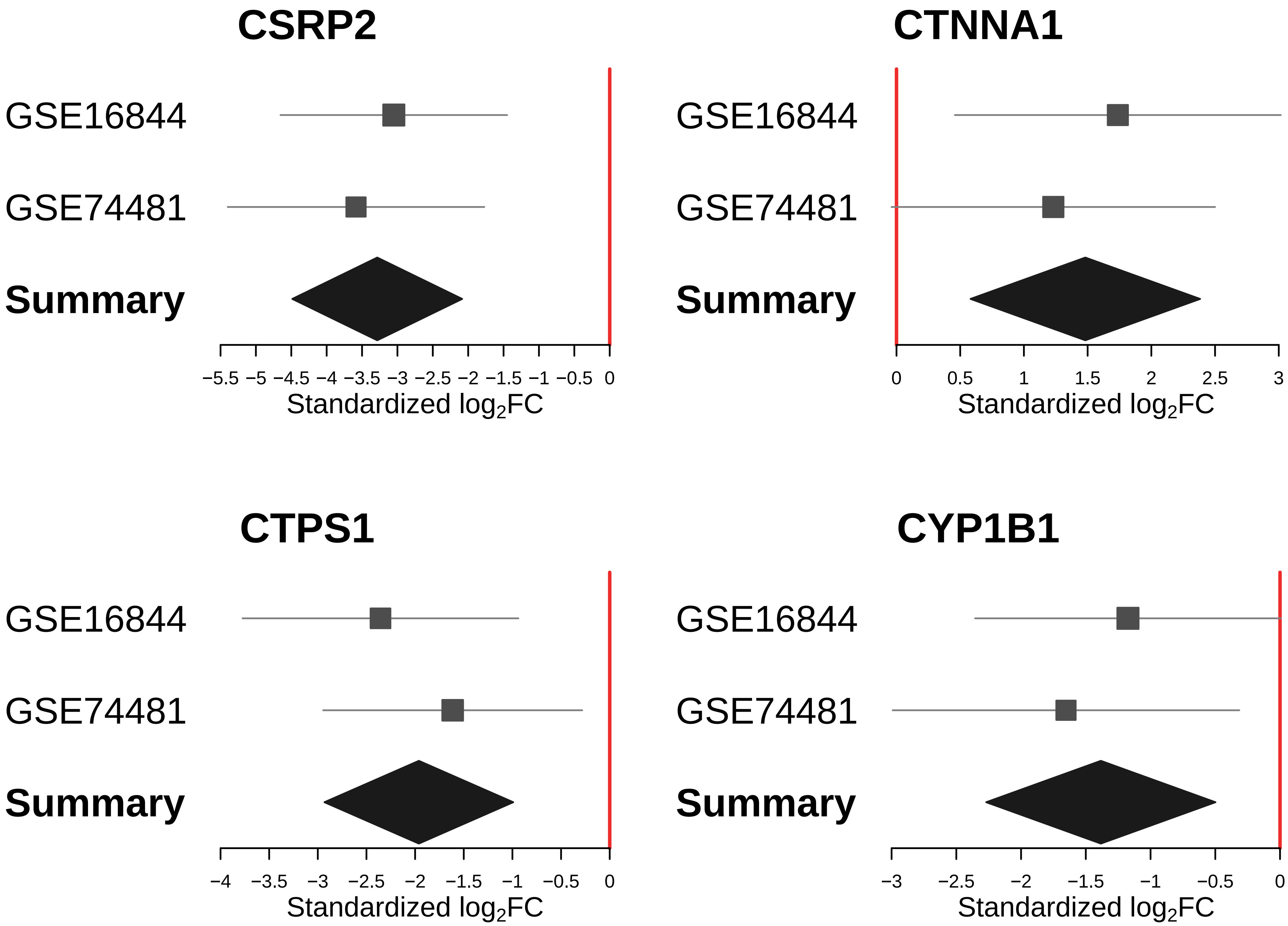

CSRP2
GSE16844
GSE74481
Summary
−5.5 −5 −4.5 −4 −3.5 −3 −2.5 −2 −1.5 −1 −0.5 0
Standardized log2FC
CTNNA1
GSE16844
GSE74481
Summary
0 0.5 1 1.5 2 2.5 3
Standardized log2FC
CTPS1
GSE16844
GSE74481
Summary
−4 −3.5 −3 −2.5 −2 −1.5 −1 −0.5 0
Standardized log2FC
CYP1B1
GSE16844
GSE74481
Summary
−3 −2.5 −2 −1.5 −1 −0.5 0
Standardized log2FC

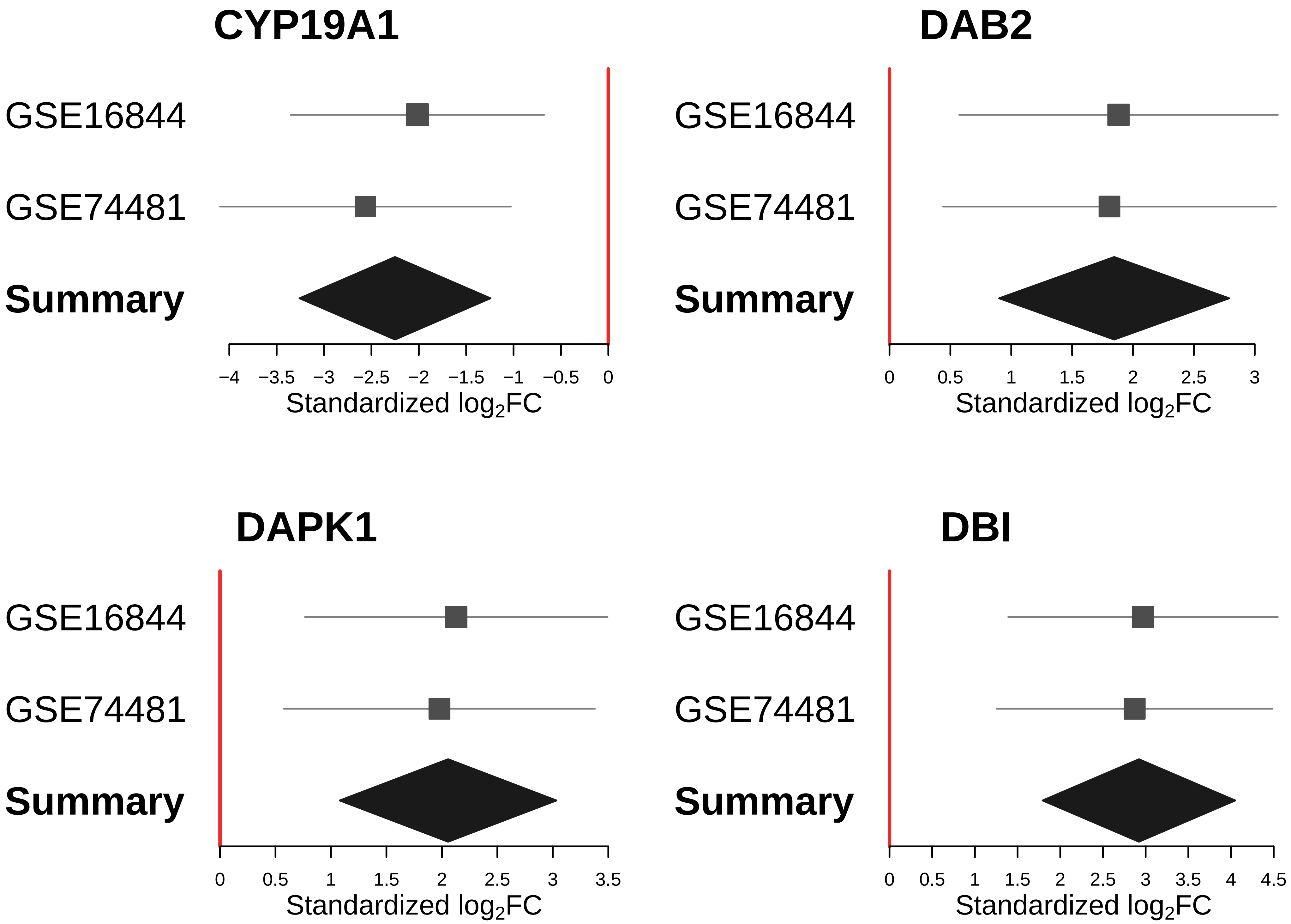
CYP19A1
DAB2
DAPK1
DBI
GSE16844
GSE74481
Summary
−4
−3.5
−3
−2.5
−2
−1.5
−1
−0.5
0
0.5
1
1.5
2
2.5
3
3.5
4
4.5
Standardized log2FC

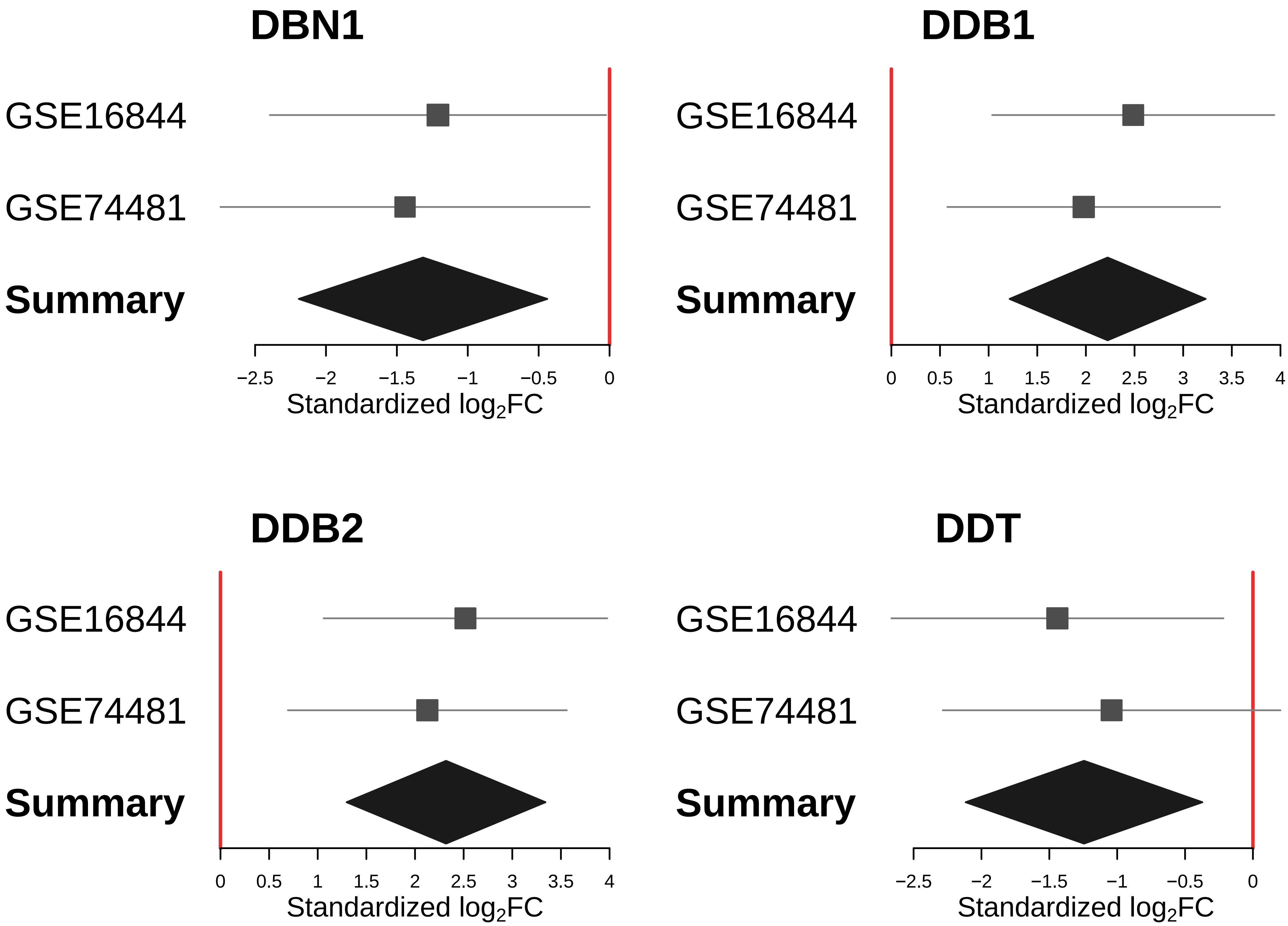
DBN1
GSE16844
GSE74481
Summary
−2.5 −2 −1.5 −1 −0.5 0
Standardized log2FC
DDB1
GSE16844
GSE74481
Summary
0 0.5 1 1.5 2 2.5 3 3.5 4
Standardized log2FC
DDB2
GSE16844
GSE74481
Summary
0 0.5 1 1.5 2 2.5 3 3.5 4
Standardized log2FC
DDT
GSE16844
GSE74481
Summary
−2.5 −2 −1.5 −1 −0.5 0
Standardized log2FC

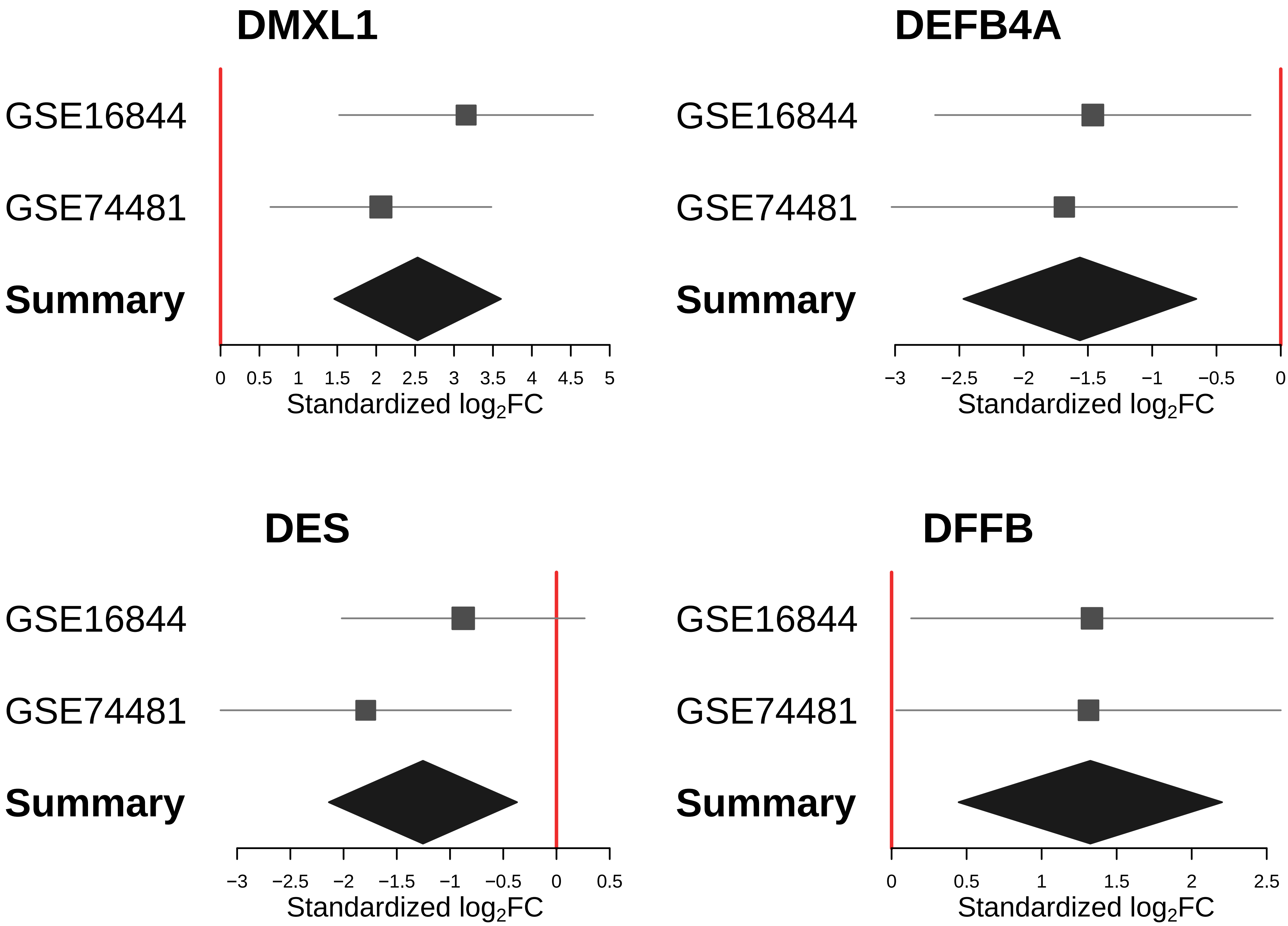
DMXL1
GSE16844
GSE74481
Summary
0 0.5 1 1.5 2 2.5 3 3.5 4 4.5 5
Standardized log2FC
DEFB4A
GSE16844
GSE74481
Summary
−3 −2.5 −2 −1.5 −1 −0.5 0
Standardized log2FC
DES
GSE16844
GSE74481
Summary
−3 −2.5 −2 −1.5 −1 −0.5 0 0.5
Standardized log2FC
DFFB
GSE16844
GSE74481
Summary
0 0.5 1 1.5 2 2.5
Standardized log2FC

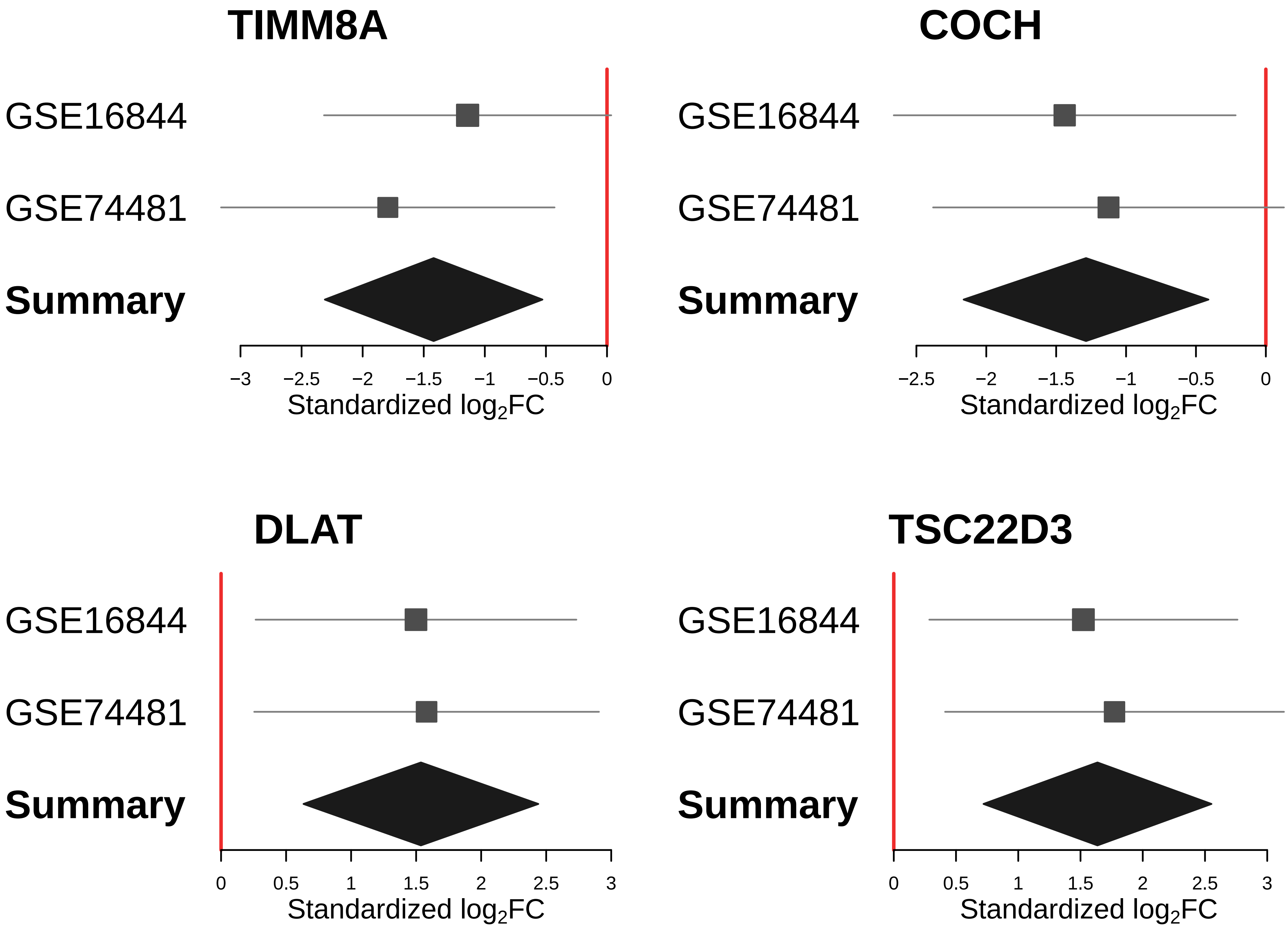

TIMM8A
COCH
DLAT
TSC22D3
GSE16844
GSE74481
Summary
−3
−2.5
−2
−1.5
−1
−0.5
0
0.5
1
1.5
2
2.5
3
Standardized log2FC

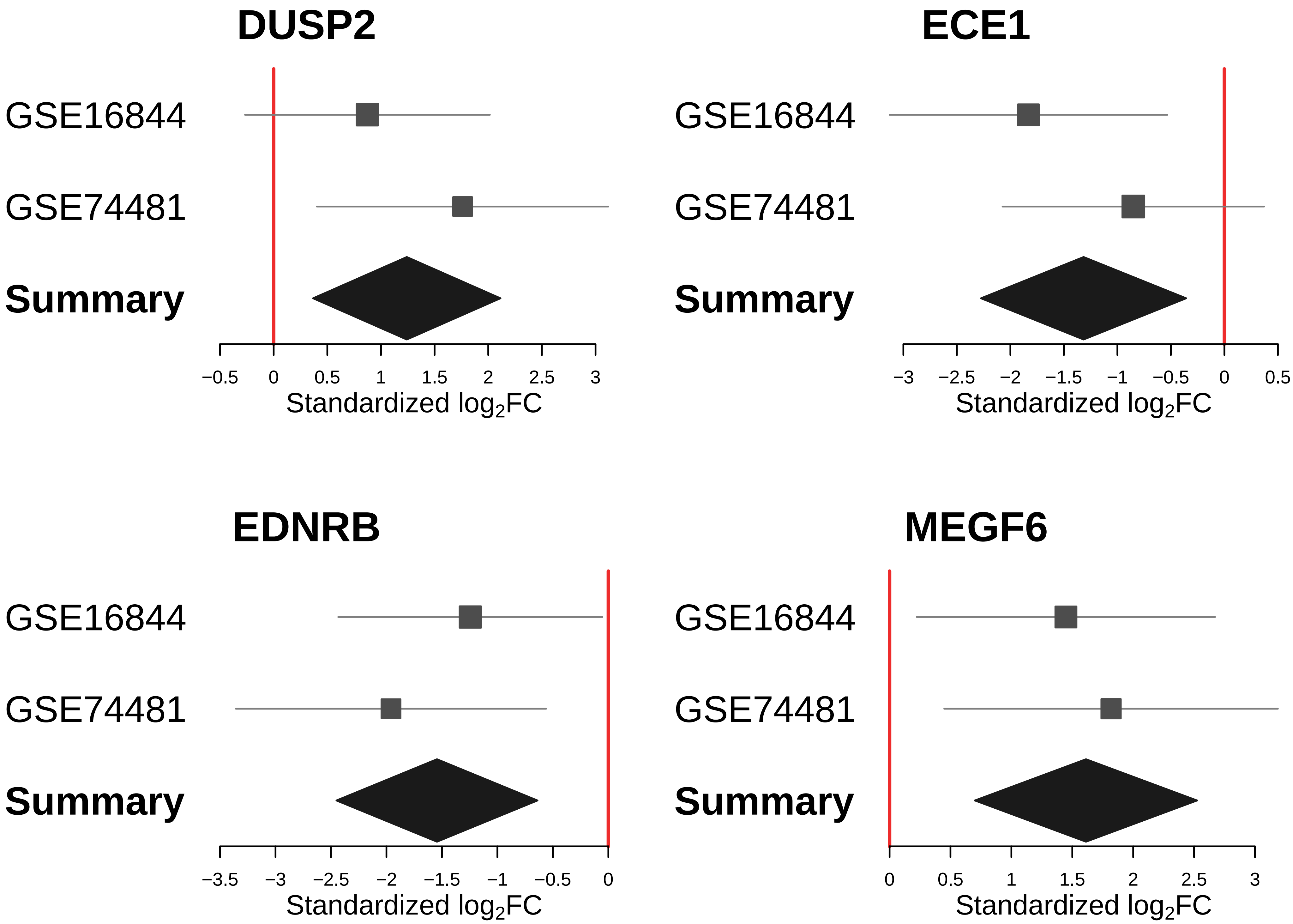
DUSP2
GSE16844
GSE74481
Summary
−0.5 0 0.5 1 1.5 2 2.5 3
Standardized log2FC
ECE1
GSE16844
GSE74481
Summary
−3 −2.5 −2 −1.5 −1 −0.5 0 0.5
Standardized log2FC
EDNRB
GSE16844
GSE74481
Summary
−3.5 −3 −2.5 −2 −1.5 −1 −0.5 0
Standardized log2FC
MEGF6
GSE16844
GSE74481
Summary
0 0.5 1 1.5 2 2.5 3
Standardized log2FC

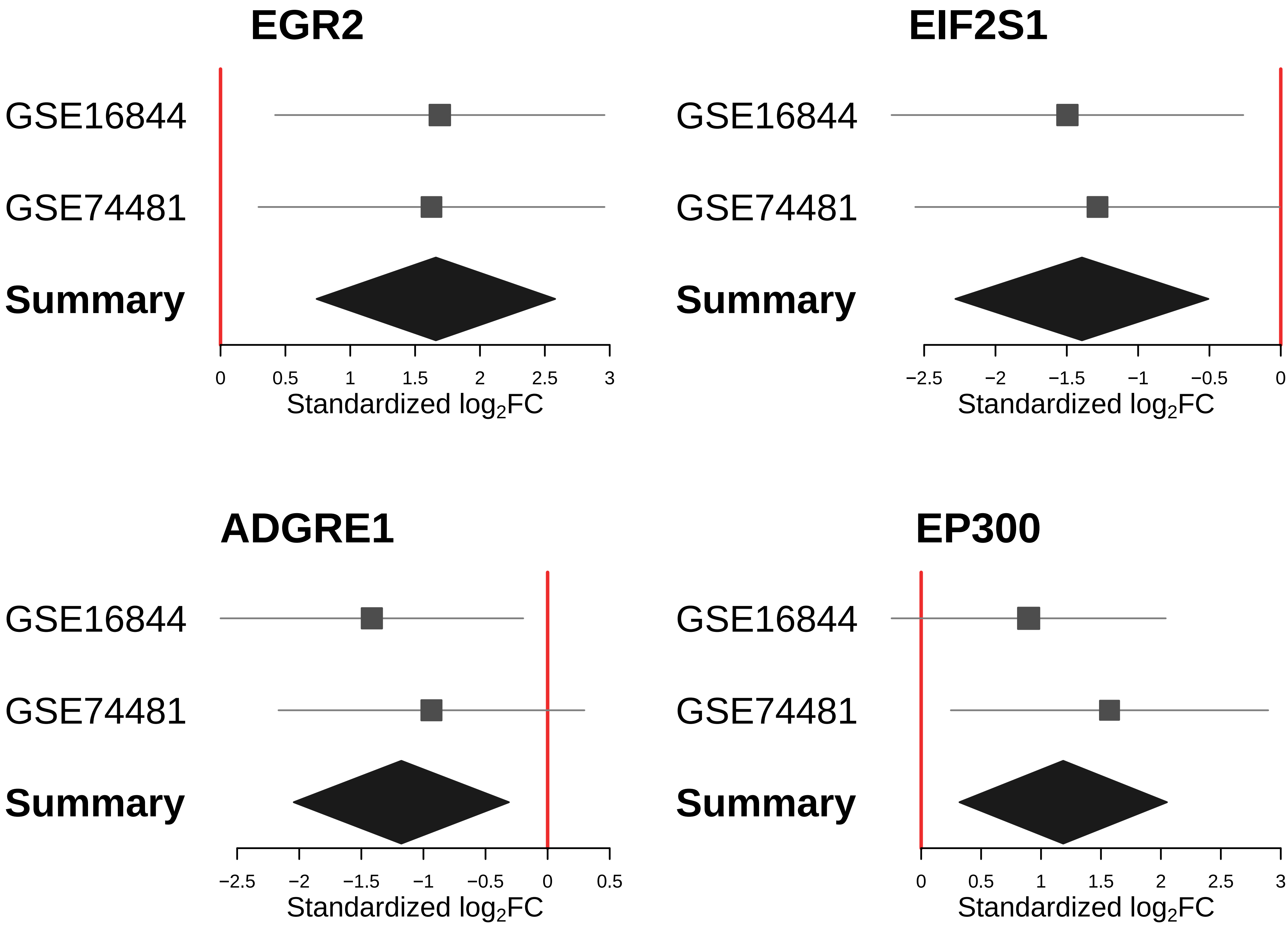

EGR2
GSE16844
GSE74481
Summary
0 0.5 1 1.5 2 2.5 3
Standardized log2FC
EIF2S1
GSE16844
GSE74481
Summary
−2.5 −2 −1.5 −1 −0.5 0
Standardized log2FC
ADGRE1
GSE16844
GSE74481
Summary
−2.5 −2 −1.5 −1 −0.5 0 0.5
Standardized log2FC
EP300
GSE16844
GSE74481
Summary
0 0.5 1 1.5 2 2.5 3
Standardized log2FC

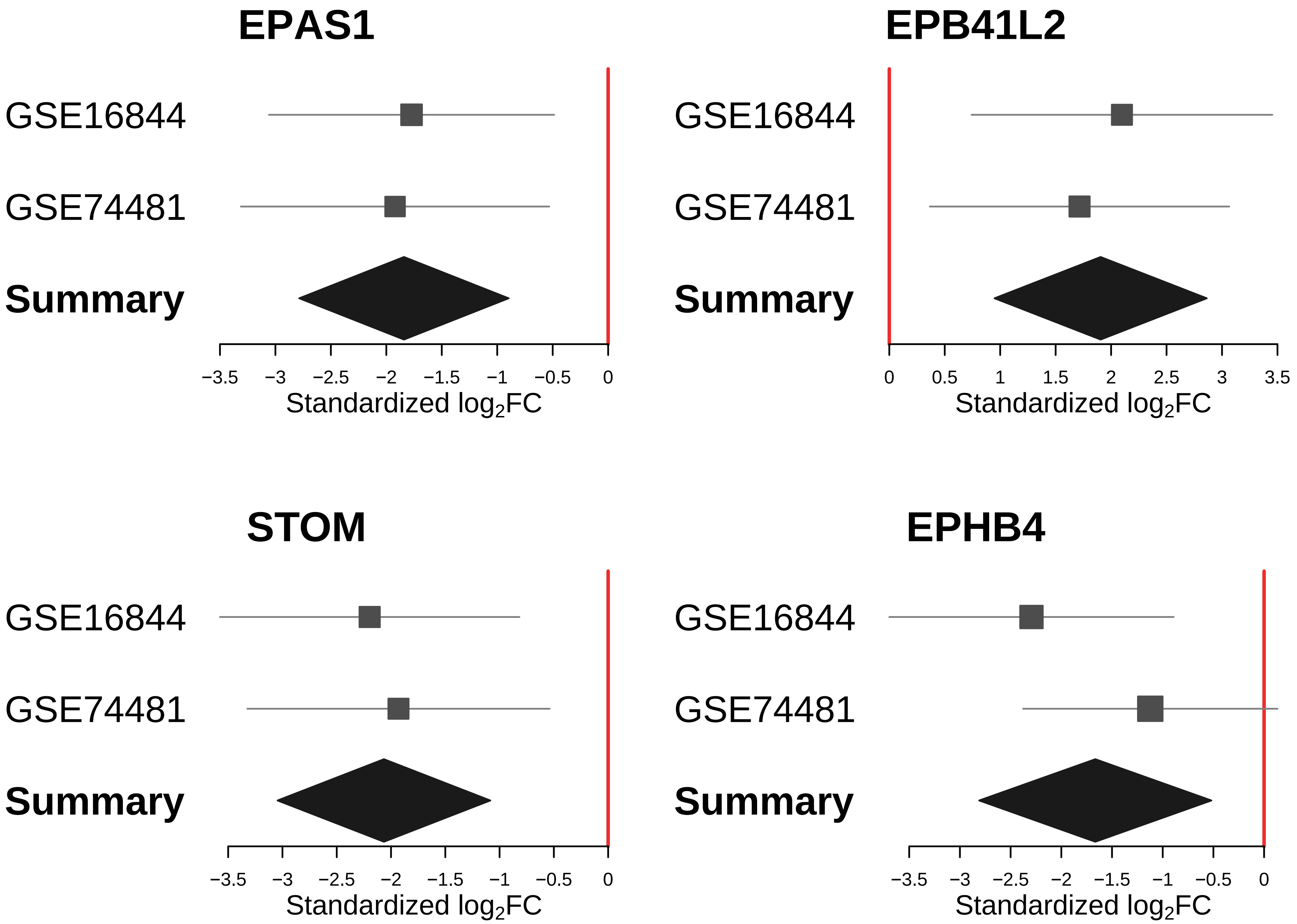
EPAS1
GSE16844
GSE74481
Summary
−3.5 −3 −2.5 −2 −1.5 −1 −0.5 0
Standardized log2FC
EPB41L2
GSE16844
GSE74481
Summary
0 0.5 1 1.5 2 2.5 3 3.5
Standardized log2FC
STOM
GSE16844
GSE74481
Summary
−3.5 −3 −2.5 −2 −1.5 −1 −0.5 0
Standardized log2FC
EPHB4
GSE16844
GSE74481
Summary
−3.5 −3 −2.5 −2 −1.5 −1 −0.5 0
Standardized log2FC

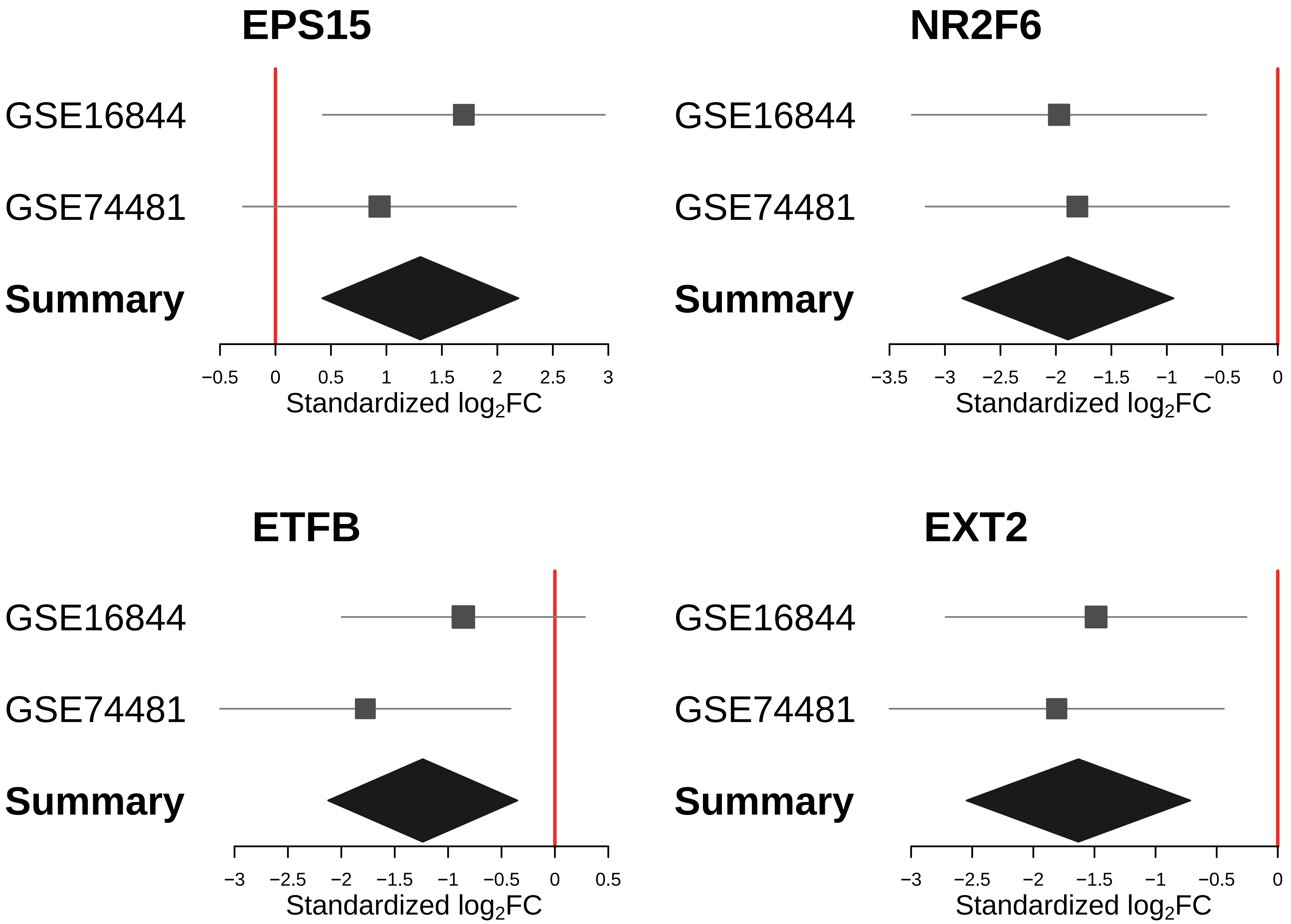
EPS15
GSE16844
GSE74481
Summary
−0.5 0 0.5 1 1.5 2 2.5 3
Standardized log2FC
NR2F6
GSE16844
GSE74481
Summary
−3.5 −3 −2.5 −2 −1.5 −1 −0.5 0
Standardized log2FC
ETFB
GSE16844
GSE74481
Summary
−3 −2.5 −2 −1.5 −1 −0.5 0 0.5
Standardized log2FC
EXT2
GSE16844
GSE74481
Summary
−3 −2.5 −2 −1.5 −1 −0.5 0
Standardized log2FC

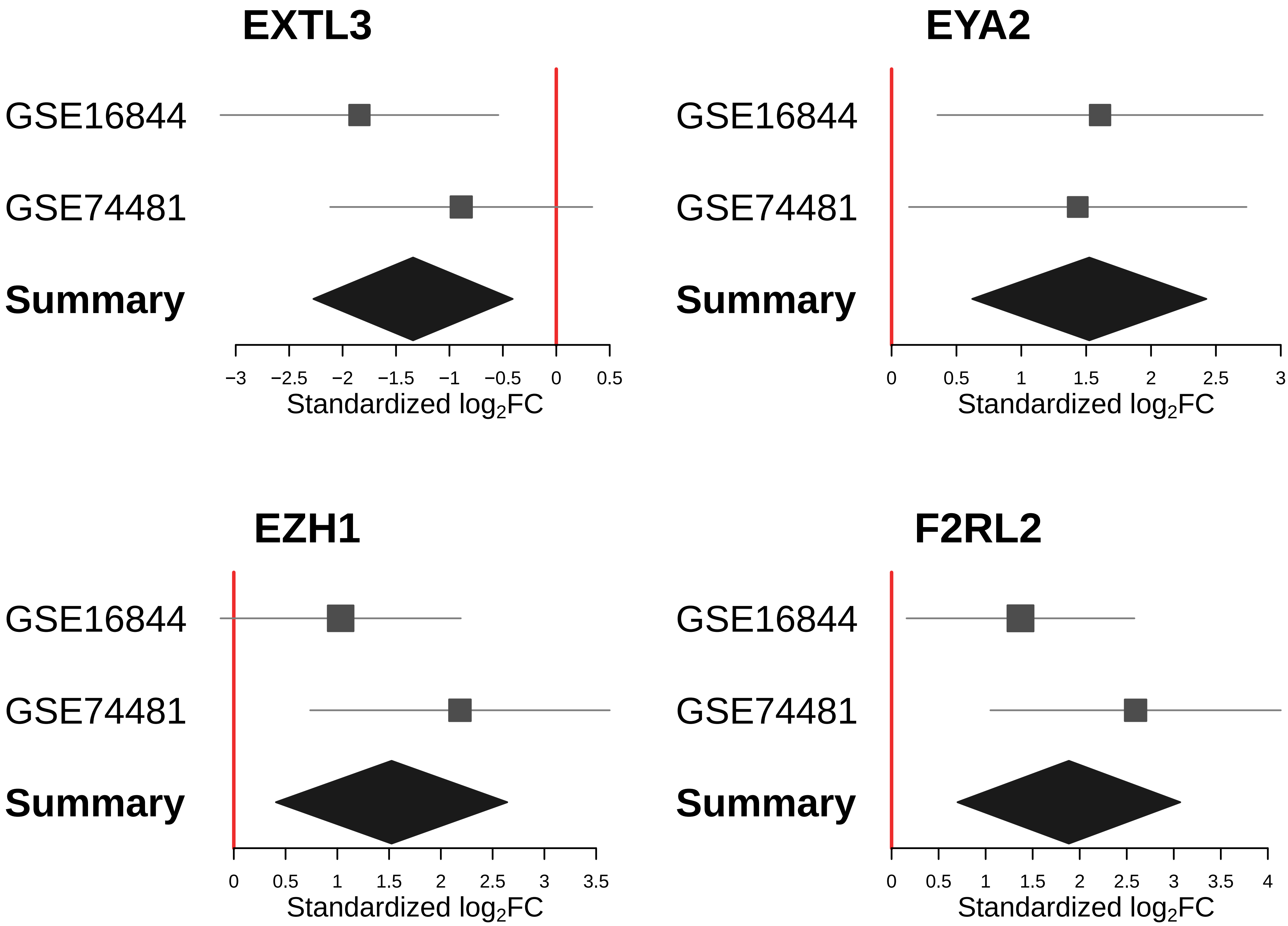

EXTL3
GSE16844
GSE74481
Summary
−3 −2.5 −2 −1.5 −1 −0.5 0 0.5
Standardized log2FC
EYA2
GSE16844
GSE74481
Summary
0 0.5 1 1.5 2 2.5 3
Standardized log2FC
EZH1
GSE16844
GSE74481
Summary
0 0.5 1 1.5 2 2.5 3 3.5
Standardized log2FC
F2RL2
GSE16844
GSE74481
Summary
0 0.5 1 1.5 2 2.5 3 3.5 4
Standardized log2FC

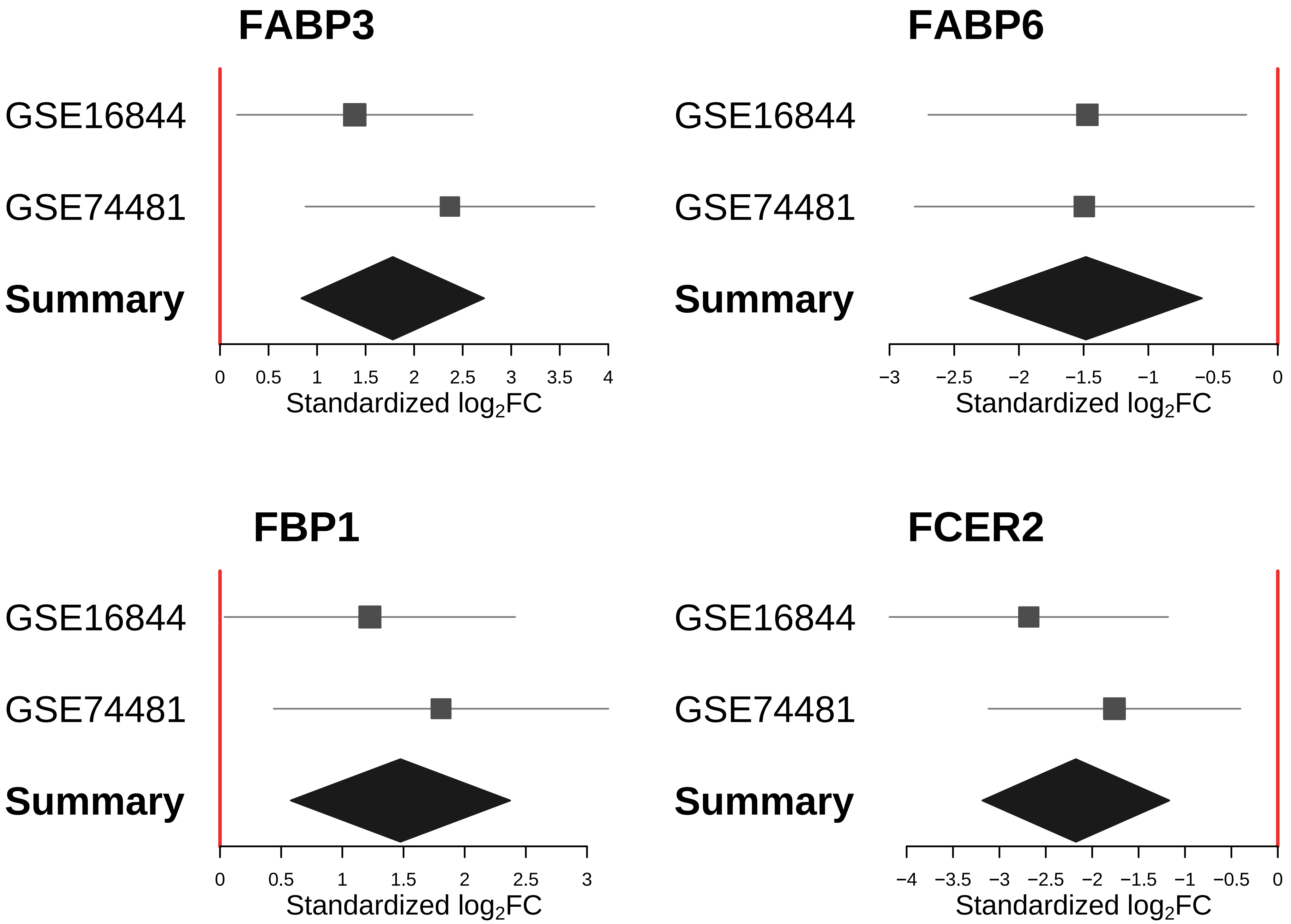
FABP3
GSE16844
GSE74481
Summary
0
0.5
1
1.5
2
2.5
3
3.5
4
Standardized log2FC
FABP6
GSE16844
GSE74481
Summary
−3
−2.5
−2
−1.5
−1
−0.5
0
Standardized log2FC
FBP1
GSE16844
GSE74481
Summary
0
0.5
1
1.5
2
2.5
3
Standardized log2FC
FCER2
GSE16844
GSE74481
Summary
−4
−3.5
−3
−2.5
−2
−1.5
−1
−0.5
0
Standardized log2FC

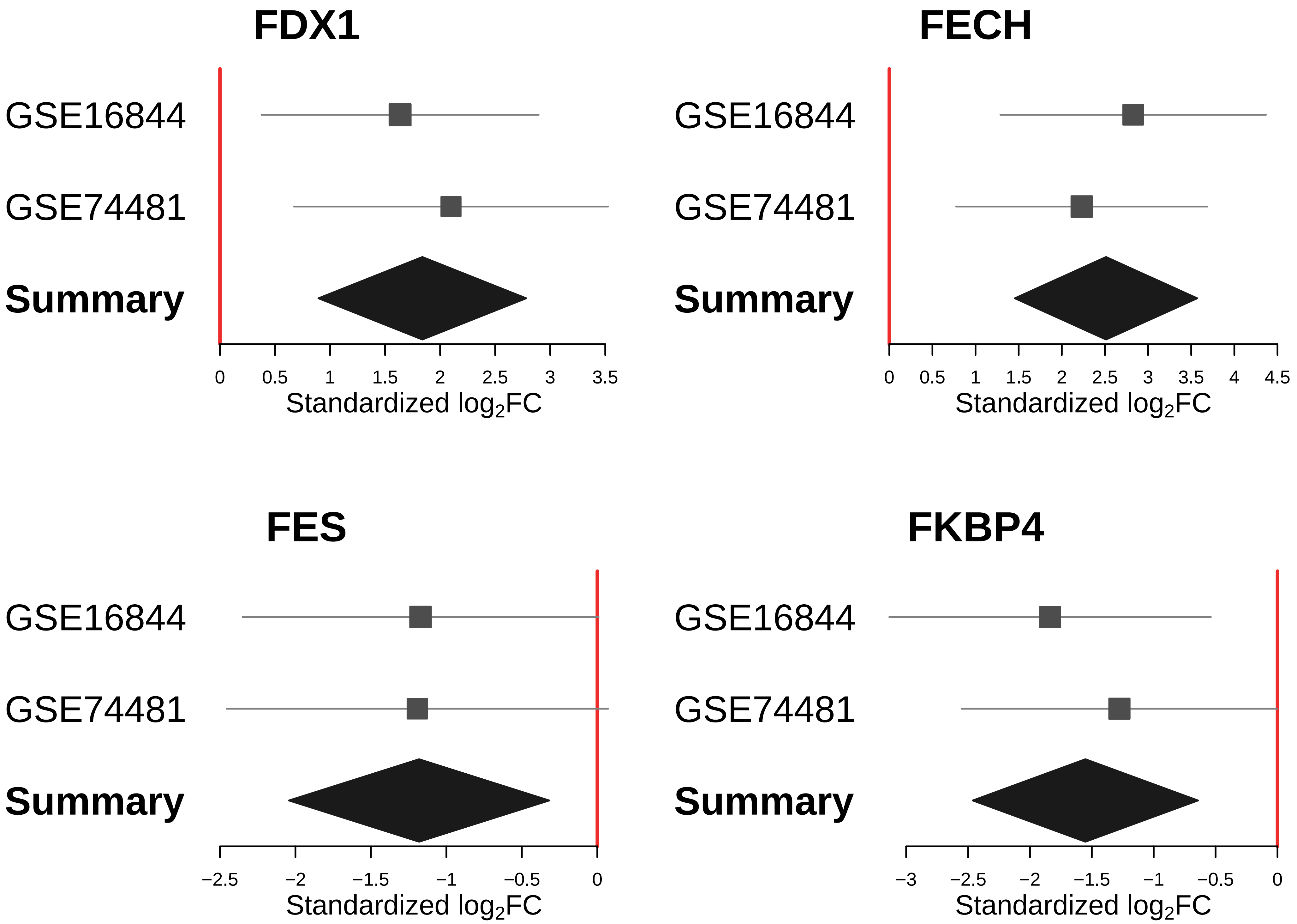
FDX1
GSE16844
GSE74481
Summary
0
0.5
1
1.5
2
2.5
3
3.5
Standardized log2FC
FECH
GSE16844
GSE74481
Summary
0
0.5
1
1.5
2
2.5
3
3.5
4
4.5
Standardized log2FC
FES
GSE16844
GSE74481
Summary
−2.5
−2
−1.5
−1
−0.5
0
Standardized log2FC
FKBP4
GSE16844
GSE74481
Summary
−3
−2.5
−2
−1.5
−1
−0.5
0
Standardized log2FC

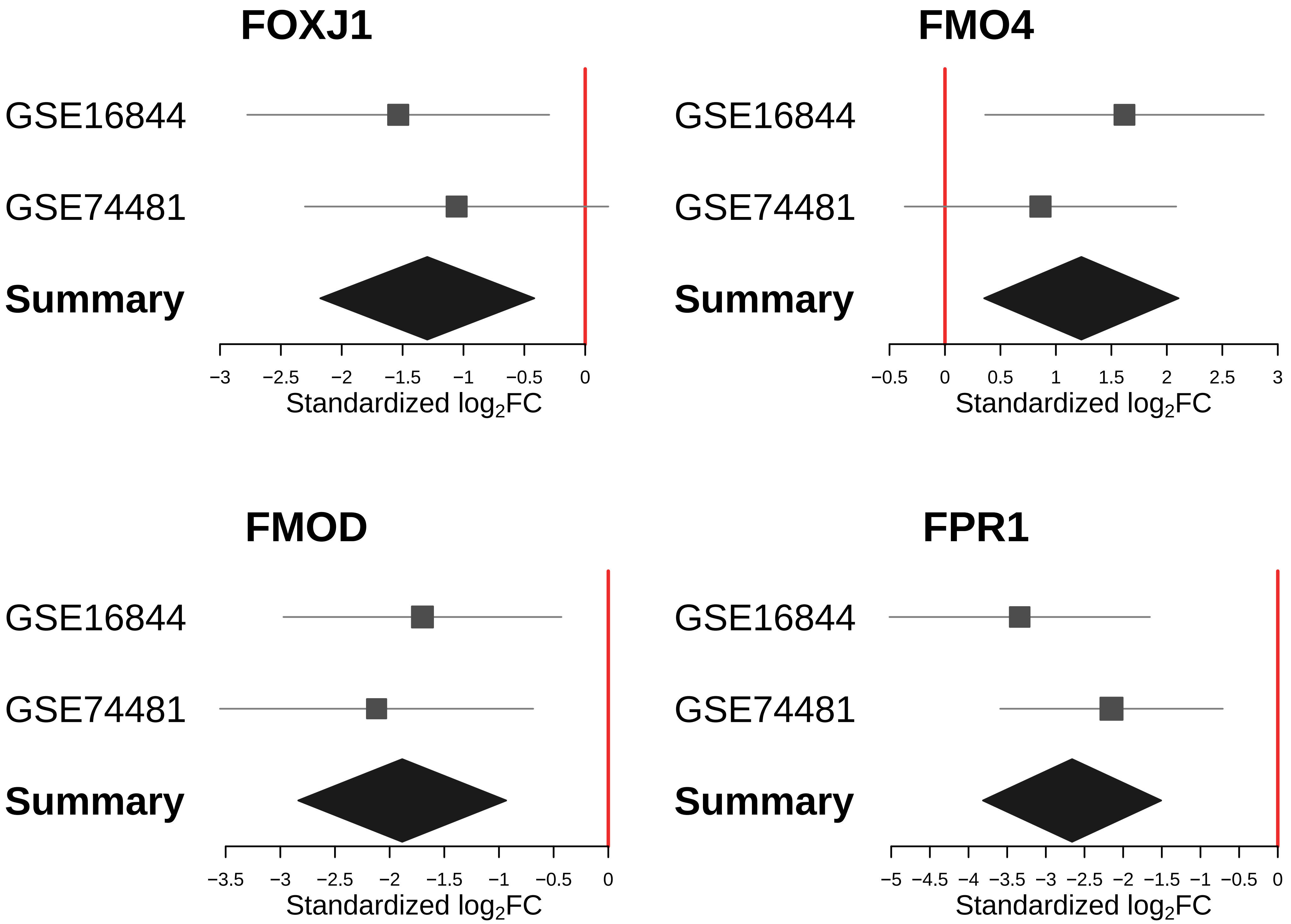
FOXJ1
GSE16844
GSE74481
Summary
−3 −2.5 −2 −1.5 −1 −0.5 0
Standardized log2FC
FMO4
GSE16844
GSE74481
Summary
−0.5 0 0.5 1 1.5 2 2.5 3
Standardized log2FC
FMOD
GSE16844
GSE74481
Summary
−3.5 −3 −2.5 −2 −1.5 −1 −0.5 0
Standardized log2FC
FPR1
GSE16844
GSE74481
Summary
−5 −4.5 −4 −3.5 −3 −2.5 −2 −1.5 −1 −0.5 0
Standardized log2FC

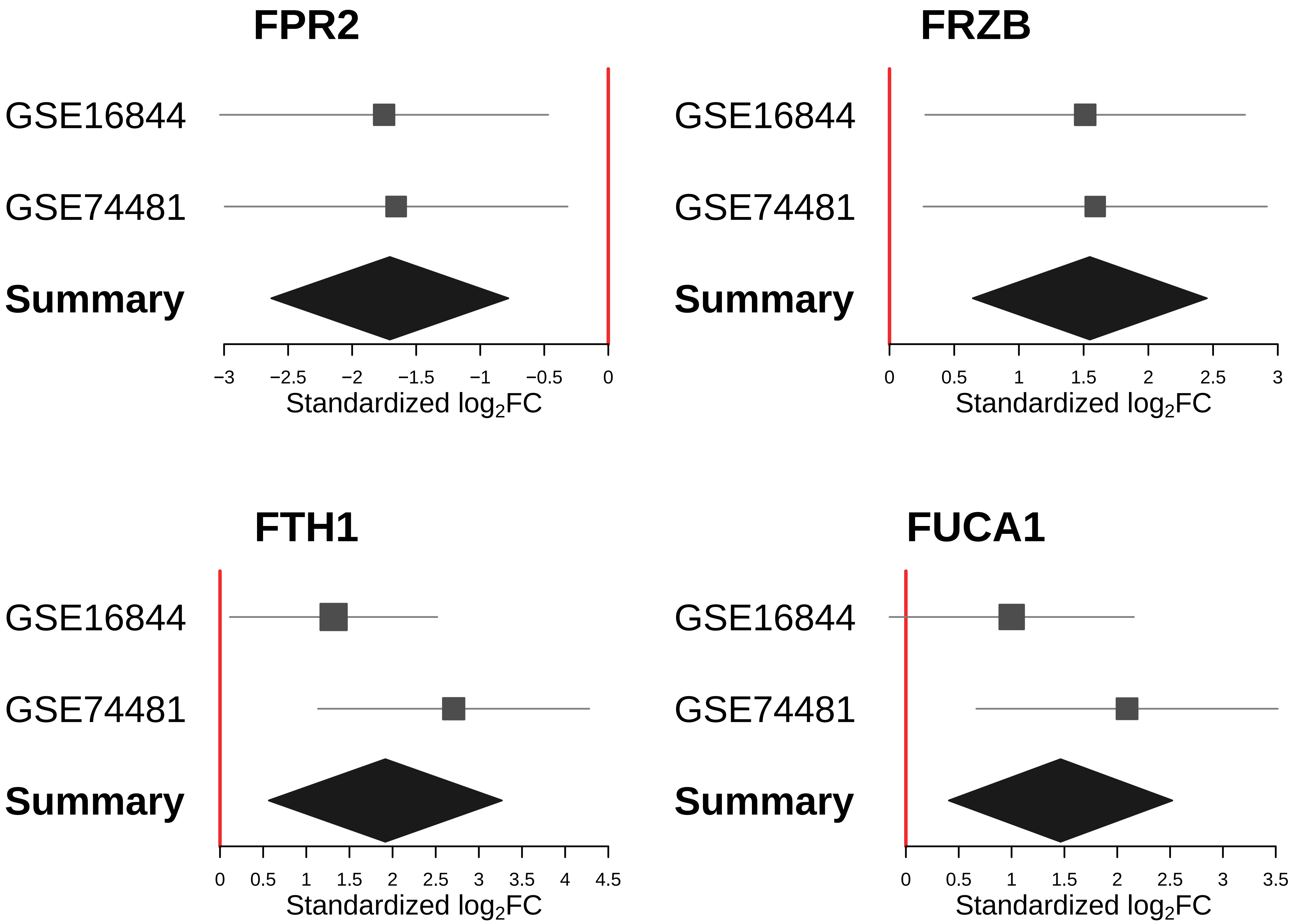
FPR2
GSE16844
GSE74481
Summary
−3 −2.5 −2 −1.5 −1 −0.5 0
Standardized log2FC
FRZB
GSE16844
GSE74481
Summary
0 0.5 1 1.5 2 2.5 3
Standardized log2FC
FTH1
GSE16844
GSE74481
Summary
0 0.5 1 1.5 2 2.5 3 3.5 4 4.5
Standardized log2FC
FUCA1
GSE16844
GSE74481
Summary
0 0.5 1 1.5 2 2.5 3 3.5
Standardized log2FC

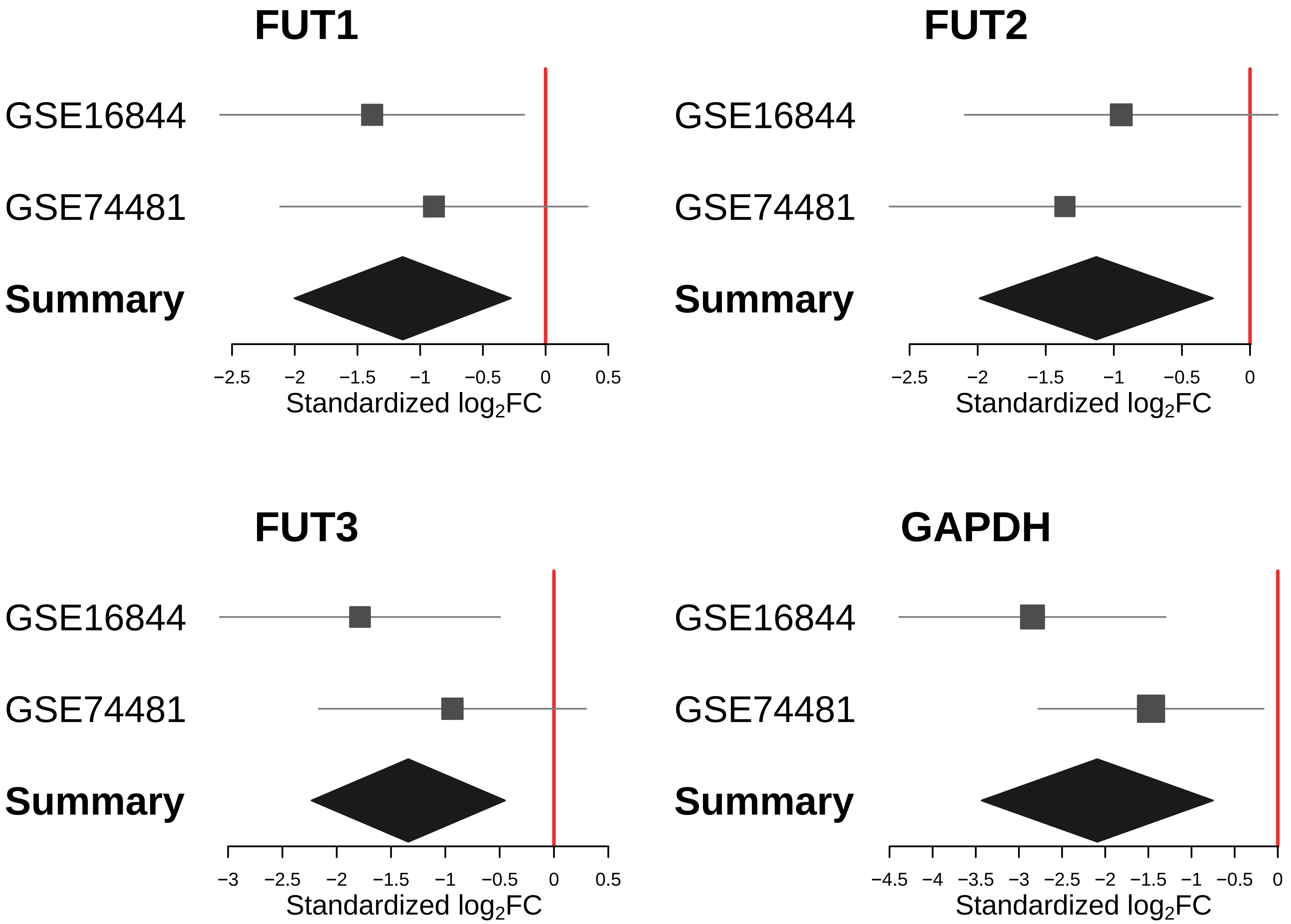
FUT1
GSE16844
GSE74481
Summary
−2.5 −2 −1.5 −1 −0.5 0 0.5
Standardized log2FC
FUT2
GSE16844
GSE74481
Summary
−2.5 −2 −1.5 −1 −0.5 0
Standardized log2FC
FUT3
GSE16844
GSE74481
Summary
−3 −2.5 −2 −1.5 −1 −0.5 0 0.5
Standardized log2FC
GAPDH
GSE16844
GSE74481
Summary
−4.5 −4 −3.5 −3 −2.5 −2 −1.5 −1 −0.5 0
Standardized log2FC

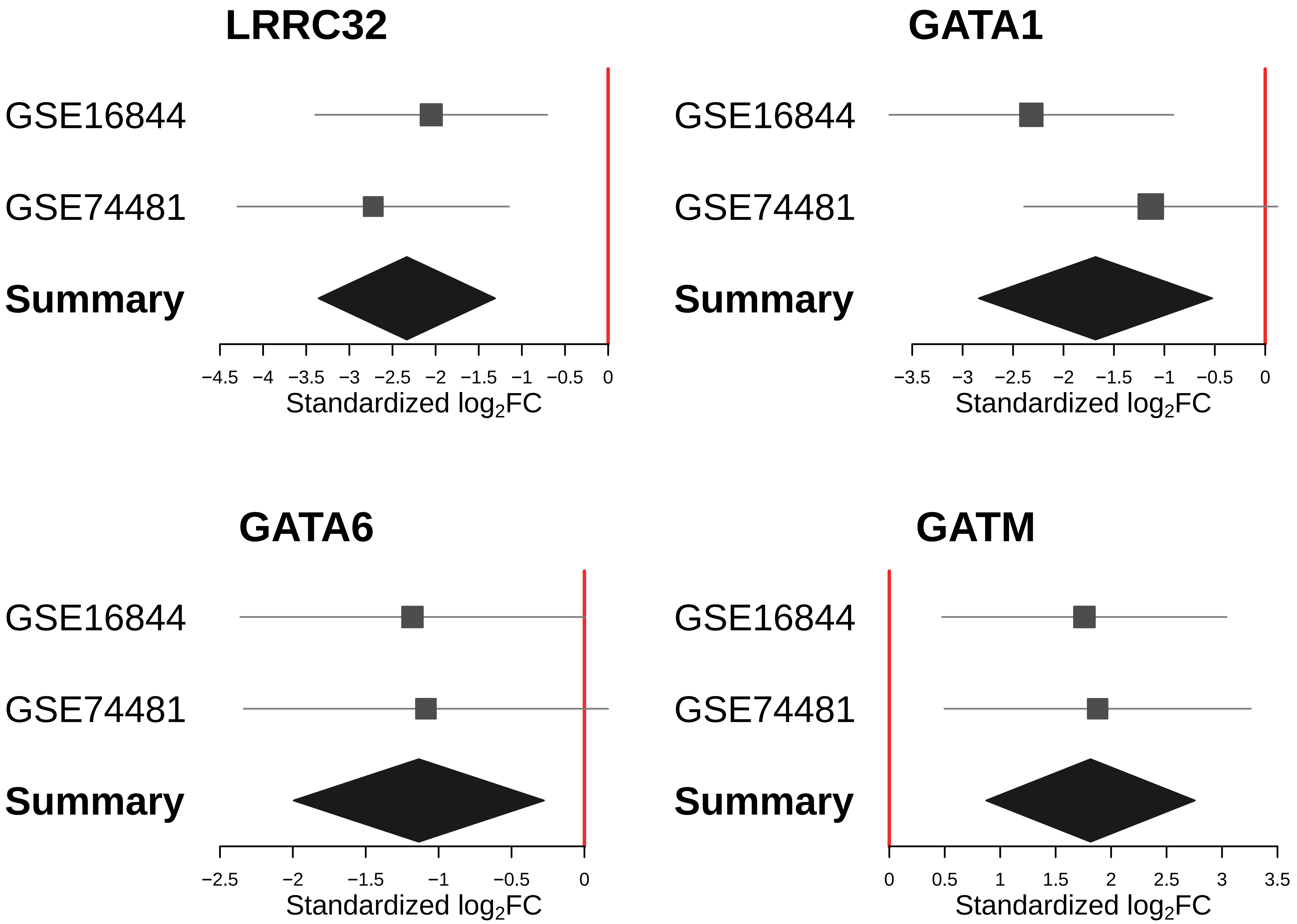
LRRC32
GSE16844
GSE74481
Summary
−4.5 −4 −3.5 −3 −2.5 −2 −1.5 −1 −0.5 0
Standardized log2FC
GATA1
GSE16844
GSE74481
Summary
−3.5 −3 −2.5 −2 −1.5 −1 −0.5 0
Standardized log2FC
GATA6
GSE16844
GSE74481
Summary
−2.5 −2 −1.5 −1 −0.5 0
Standardized log2FC
GATM
GSE16844
GSE74481
Summary
0 0.5 1 1.5 2 2.5 3 3.5
Standardized log2FC

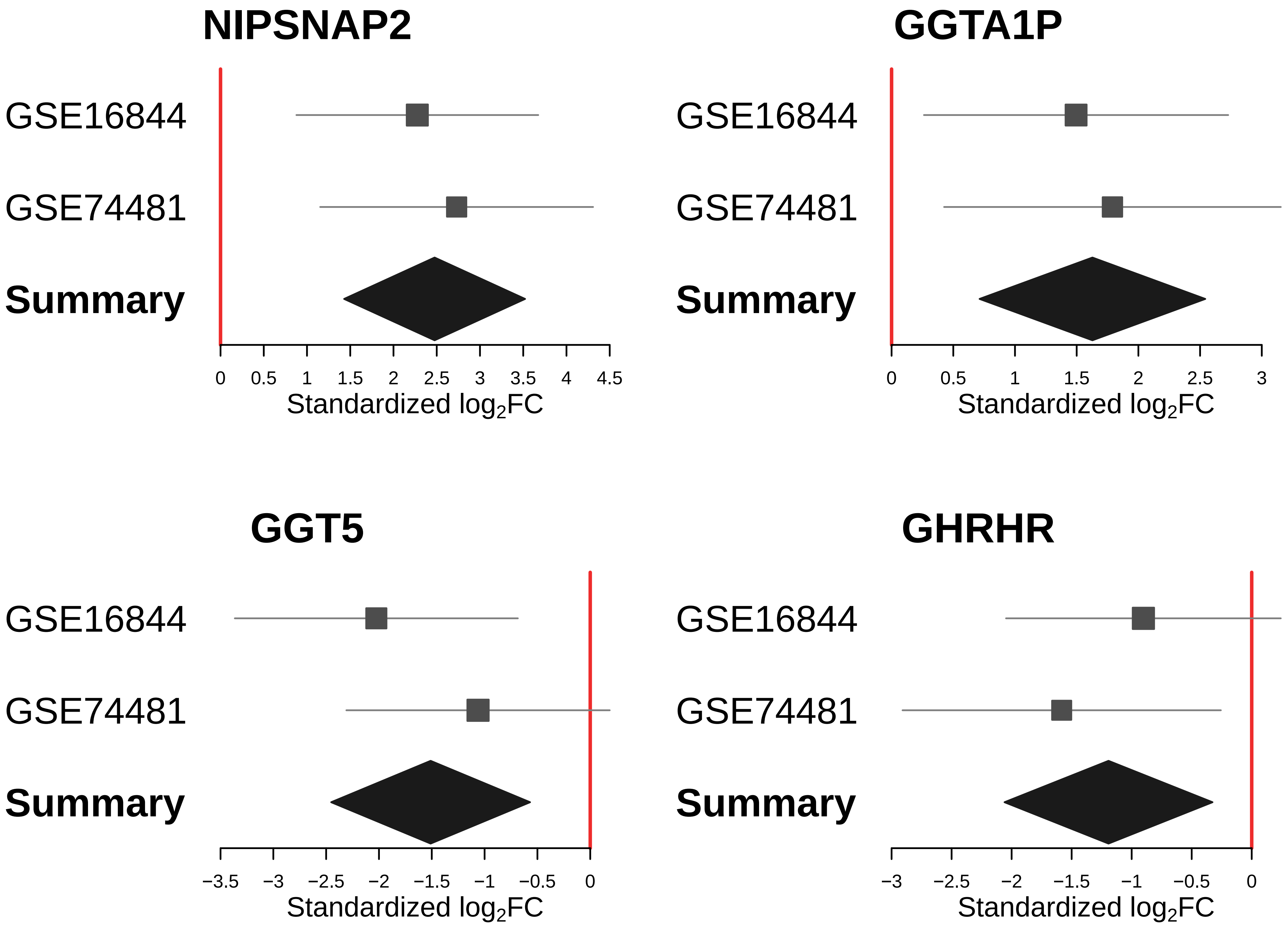
NIPSNAP2
GSE16844
GSE74481
Summary
0 0.5 1 1.5 2 2.5 3 3.5 4 4.5
Standardized log2FC
GGTA1P
GSE16844
GSE74481
Summary
0 0.5 1 1.5 2 2.5 3
Standardized log2FC
GGT5
GSE16844
GSE74481
Summary
−3.5 −3 −2.5 −2 −1.5 −1 −0.5 0
Standardized log2FC
GHRHR
GSE16844
GSE74481
Summary
−3 −2.5 −2 −1.5 −1 −0.5 0
Standardized log2FC

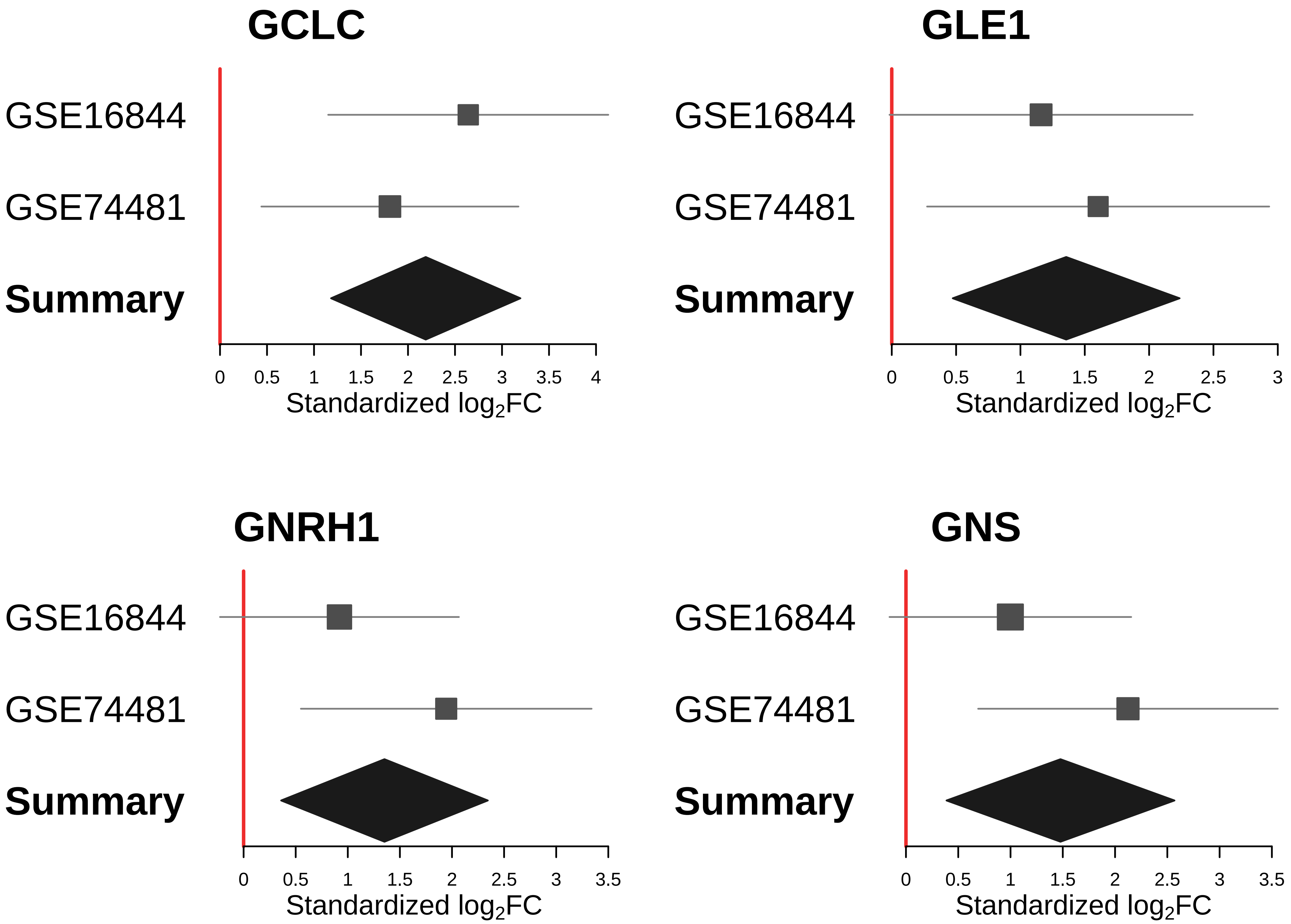

GCLC
GSE16844
GSE74481
Summary
0 0.5 1 1.5 2 2.5 3 3.5 4
Standardized log2FC
GLE1
GSE16844
GSE74481
Summary
0 0.5 1 1.5 2 2.5 3
Standardized log2FC
GNRH1
GSE16844
GSE74481
Summary
0 0.5 1 1.5 2 2.5 3 3.5
Standardized log2FC
GNS
GSE16844
GSE74481
Summary
0 0.5 1 1.5 2 2.5 3 3.5
Standardized log2FC

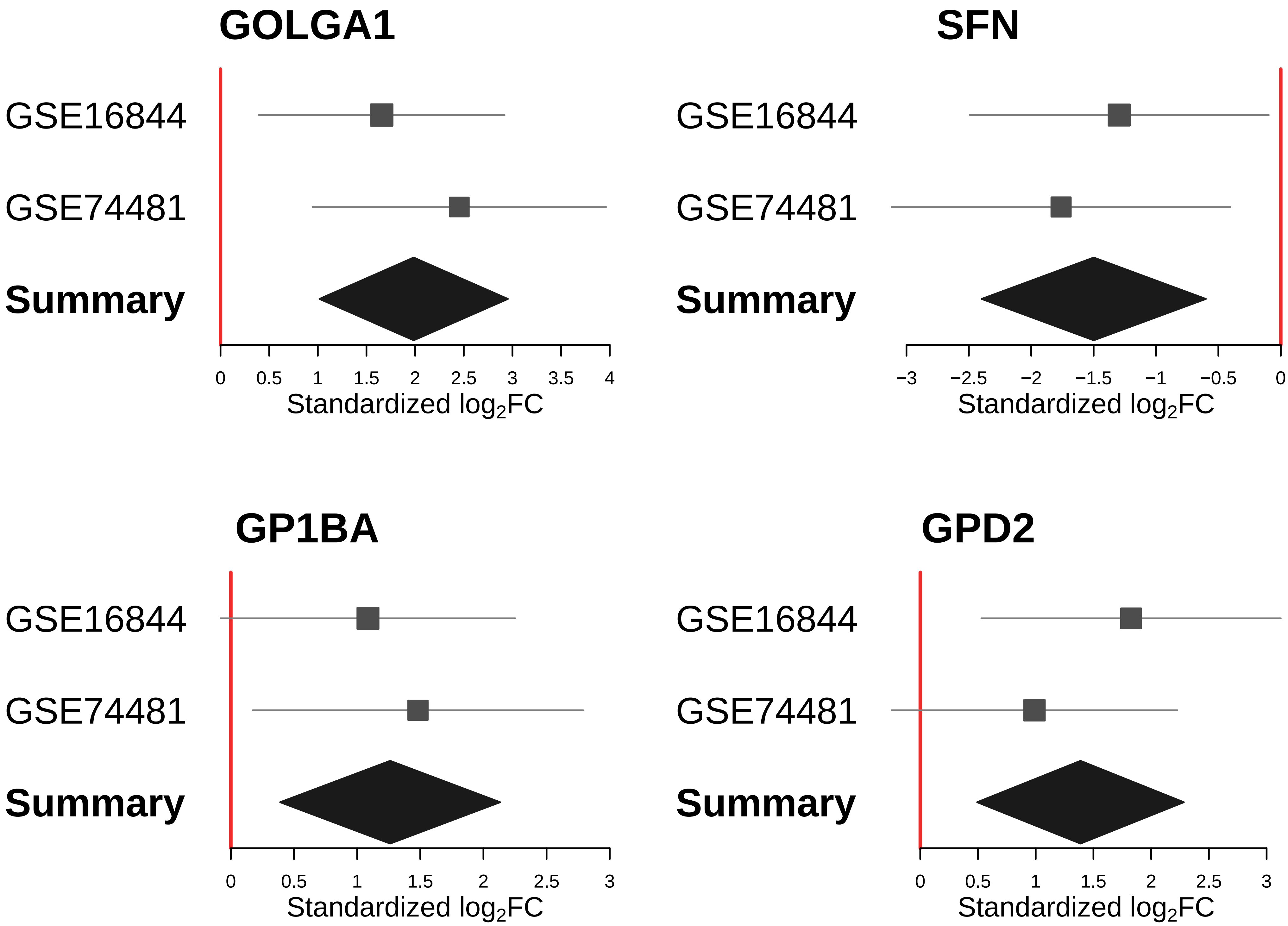
GOLGA1
GSE16844
GSE74481
Summary
0 0.5 1 1.5 2 2.5 3 3.5 4
Standardized log2FC
SFN
GSE16844
GSE74481
Summary
−3 −2.5 −2 −1.5 −1 −0.5 0
Standardized log2FC
GP1BA
GSE16844
GSE74481
Summary
0 0.5 1 1.5 2 2.5 3
Standardized log2FC
GPD2
GSE16844
GSE74481
Summary
0 0.5 1 1.5 2 2.5 3
Standardized log2FC

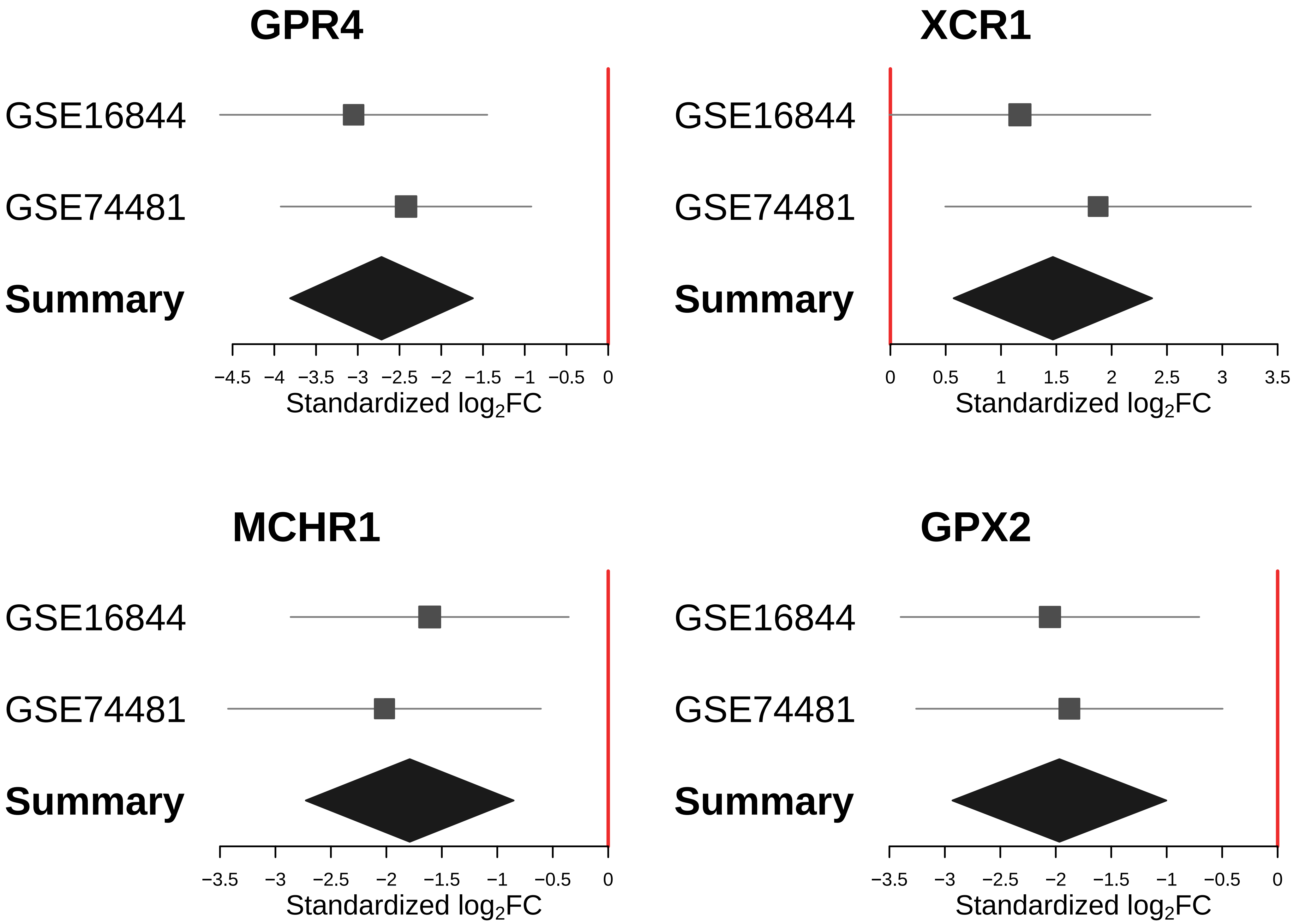
GPR4
GSE16844
GSE74481
Summary
−4.5 −4 −3.5 −3 −2.5 −2 −1.5 −1 −0.5 0
Standardized log2FC
XCR1
GSE16844
GSE74481
Summary
0 0.5 1 1.5 2 2.5 3 3.5
Standardized log2FC
MCHR1
GSE16844
GSE74481
Summary
−3.5 −3 −2.5 −2 −1.5 −1 −0.5 0
Standardized log2FC
GPX2
GSE16844
GSE74481
Summary
−3.5 −3 −2.5 −2 −1.5 −1 −0.5 0
Standardized log2FC

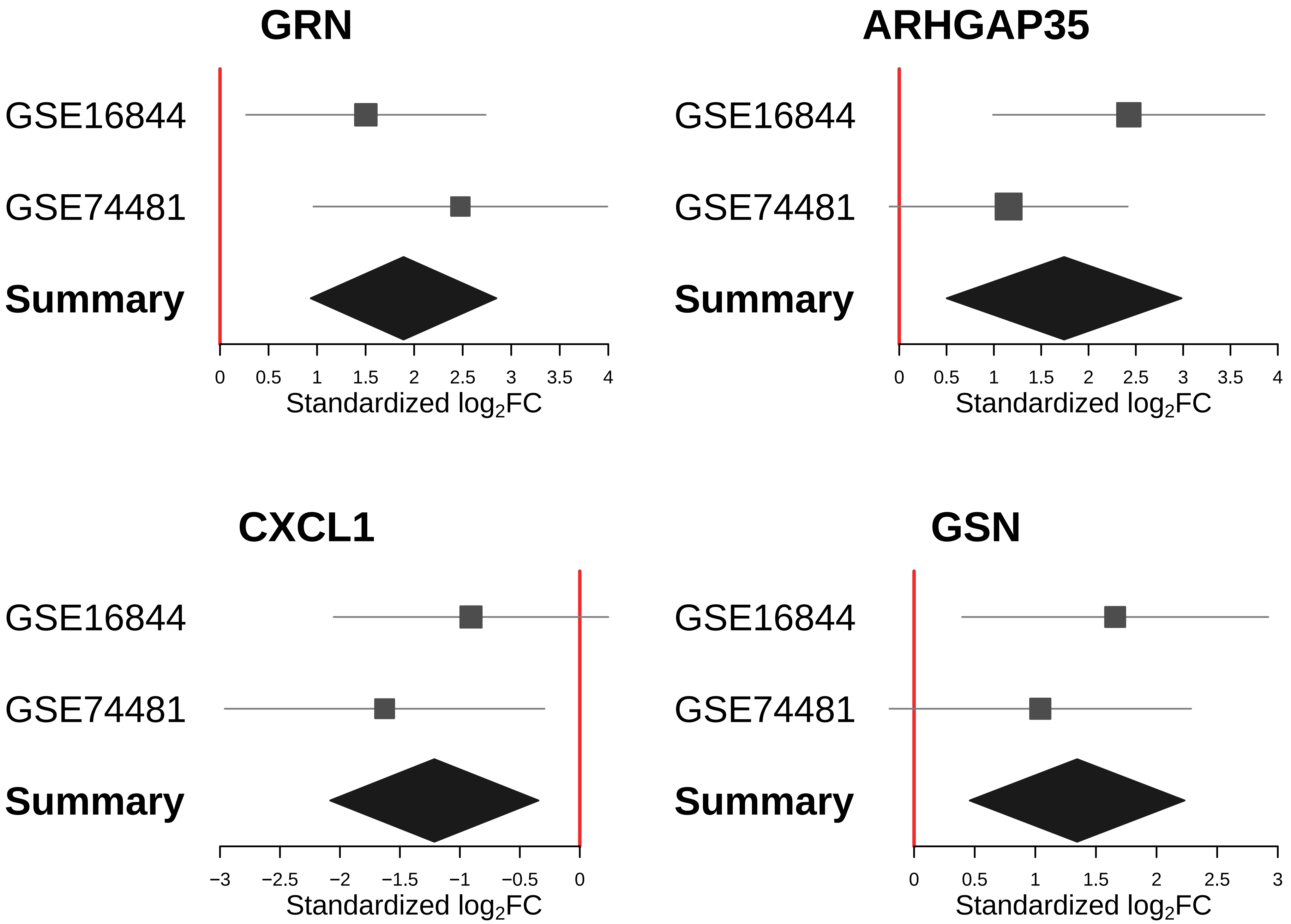
GRN
GSE16844
GSE74481
Summary
0 0.5 1 1.5 2 2.5 3 3.5 4
Standardized log2FC
ARHGAP35
GSE16844
GSE74481
Summary
0 0.5 1 1.5 2 2.5 3 3.5 4
Standardized log2FC
CXCL1
GSE16844
GSE74481
Summary
−3 −2.5 −2 −1.5 −1 −0.5 0
Standardized log2FC
GSN
GSE16844
GSE74481
Summary
0 0.5 1 1.5 2 2.5 3
Standardized log2FC

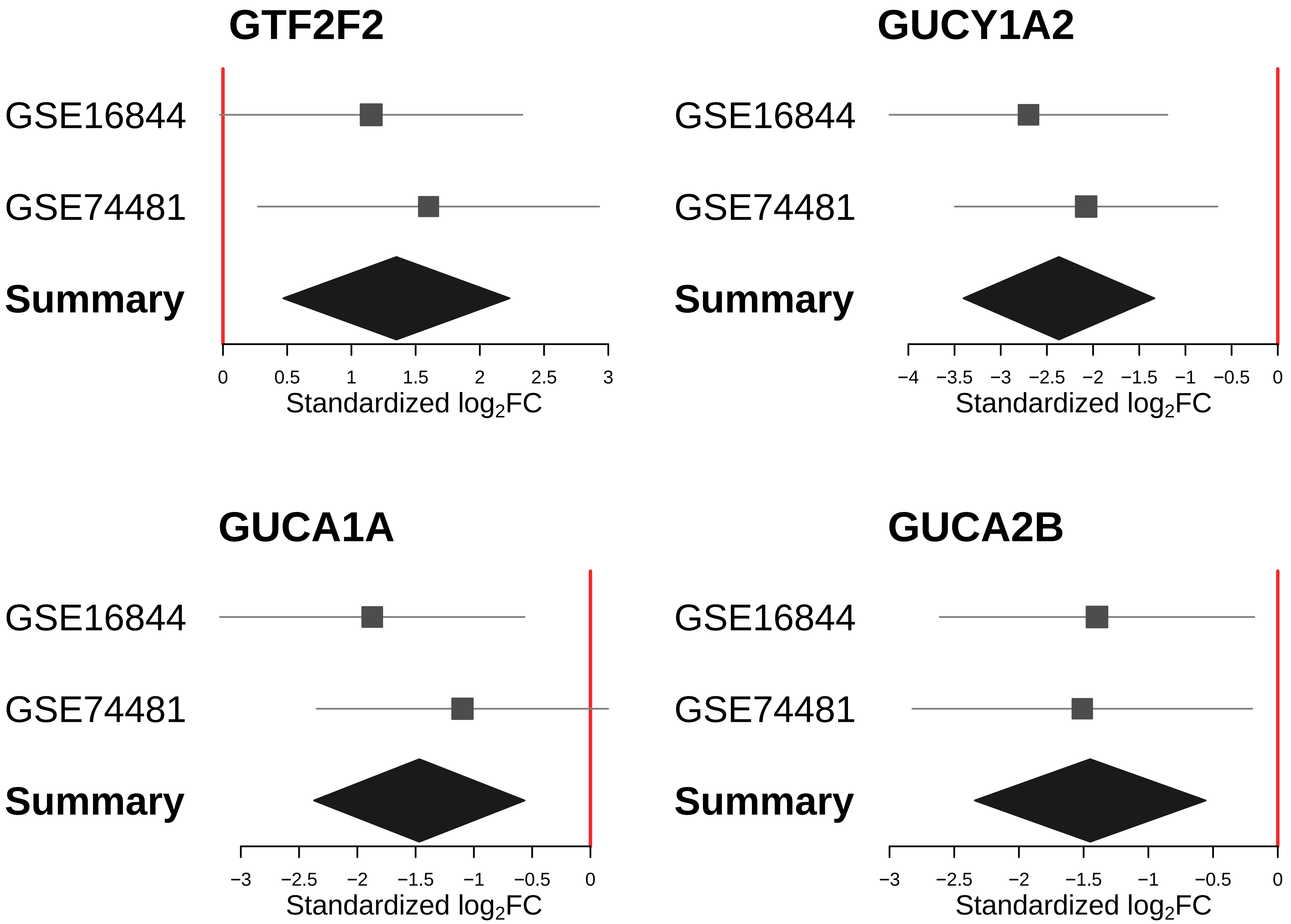
GTF2F2
GSE16844
GSE74481
Summary
0 0.5 1 1.5 2 2.5 3
Standardized log2FC
GUCY1A2
GSE16844
GSE74481
Summary
−4 −3.5 −3 −2.5 −2 −1.5 −1 −0.5 0
Standardized log2FC
GUCA1A
GSE16844
GSE74481
Summary
−3 −2.5 −2 −1.5 −1 −0.5 0
Standardized log2FC
GUCA2B
GSE16844
GSE74481
Summary
−3 −2.5 −2 −1.5 −1 −0.5 0
Standardized log2FC

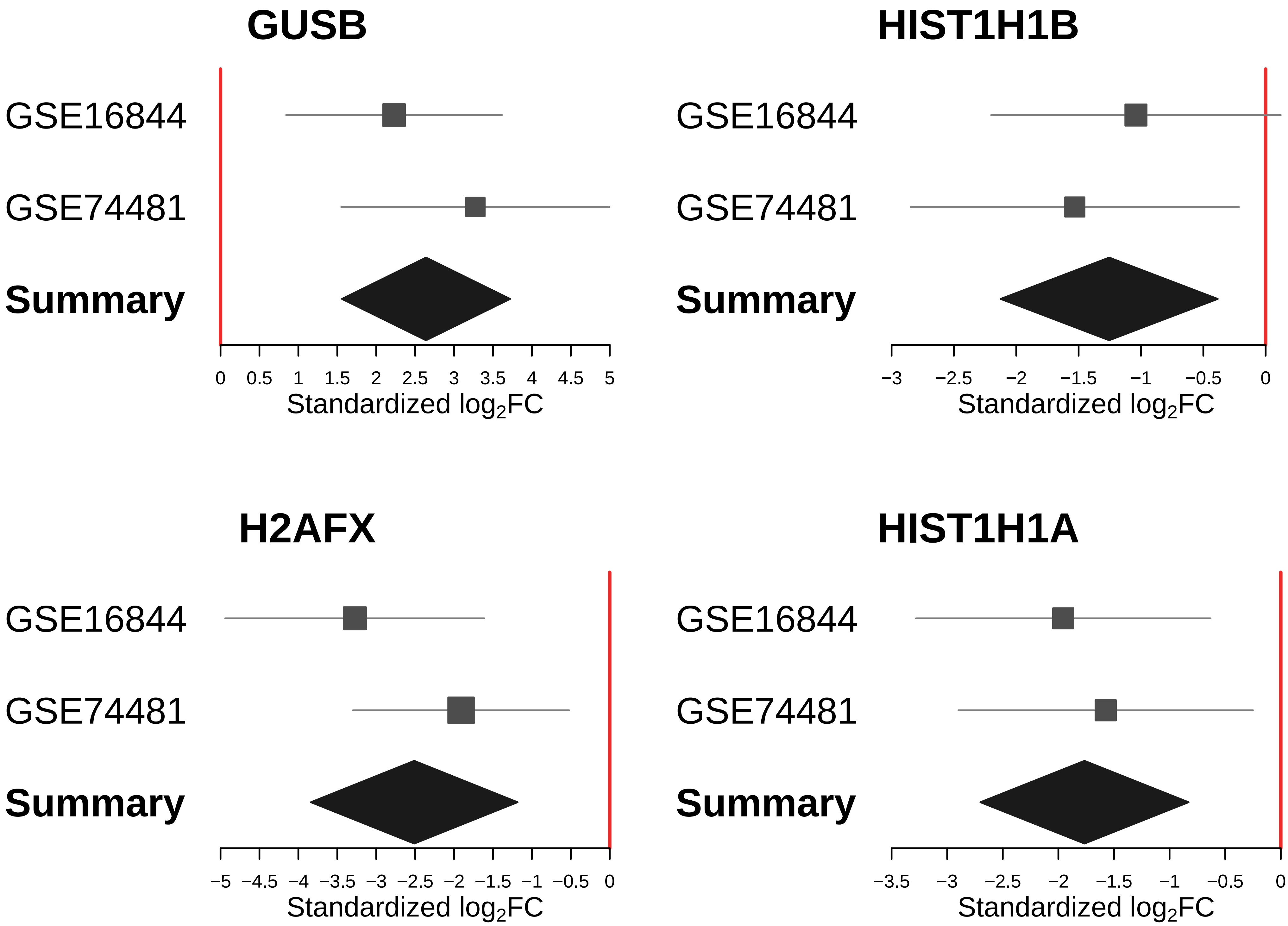
GUSB
GSE16844
GSE74481
Summary
0 0.5 1 1.5 2 2.5 3 3.5 4 4.5 5
Standardized log2FC
HIST1H1B
GSE16844
GSE74481
Summary
−3 −2.5 −2 −1.5 −1 −0.5 0
Standardized log2FC
H2AFX
GSE16844
GSE74481
Summary
−5 −4.5 −4 −3.5 −3 −2.5 −2 −1.5 −1 −0.5 0
Standardized log2FC
HIST1H1A
GSE16844
GSE74481
Summary
−3.5 −3 −2.5 −2 −1.5 −1 −0.5 0
Standardized log2FC

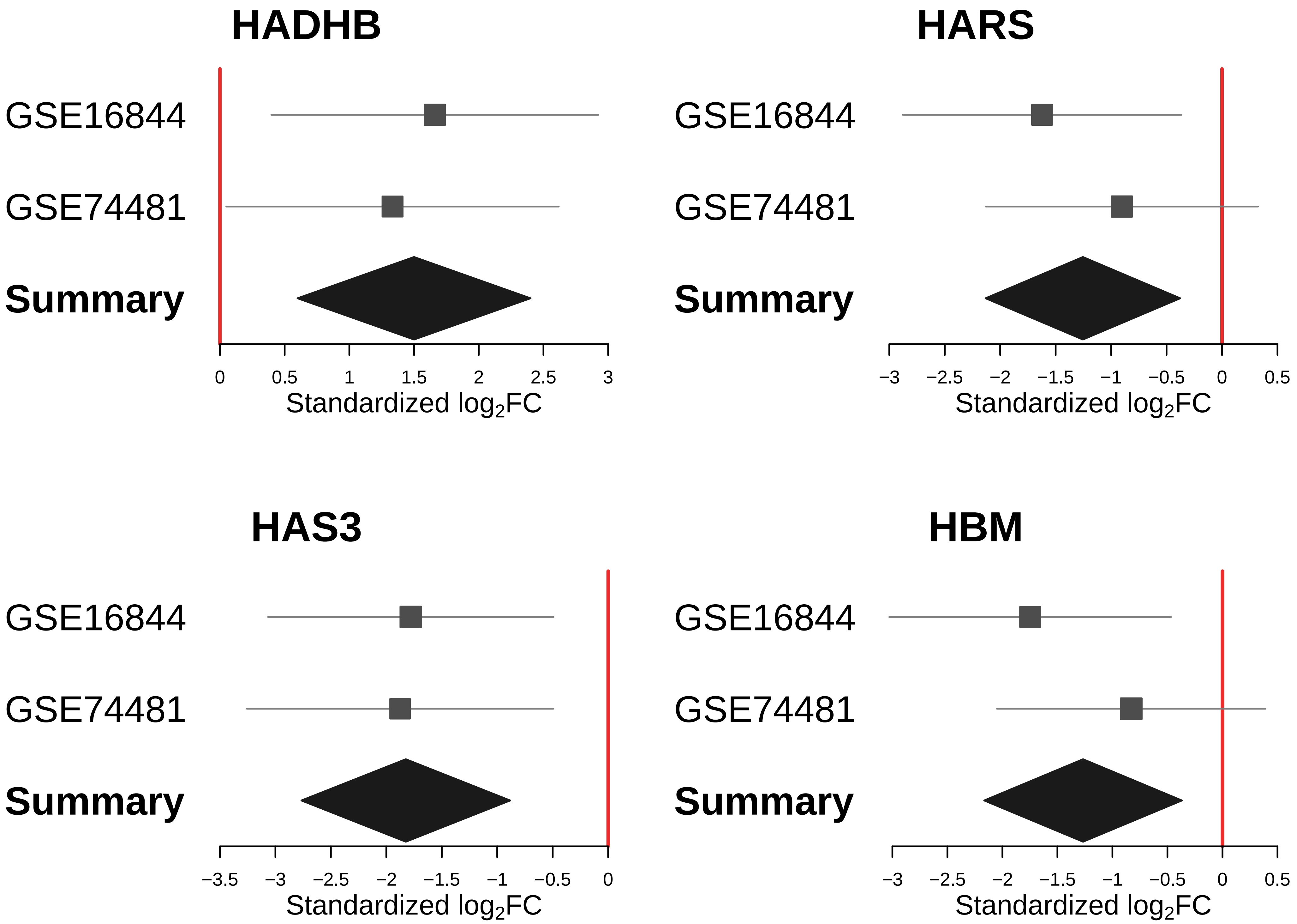
HADHB
GSE16844
GSE74481
Summary
0 0.5 1 1.5 2 2.5 3
Standardized log2FC
HARS
GSE16844
GSE74481
Summary
−3 −2.5 −2 −1.5 −1 −0.5 0 0.5
Standardized log2FC
HAS3
GSE16844
GSE74481
Summary
−3.5 −3 −2.5 −2 −1.5 −1 −0.5 0
Standardized log2FC
HBM
GSE16844
GSE74481
Summary
−3 −2.5 −2 −1.5 −1 −0.5 0 0.5
Standardized log2FC

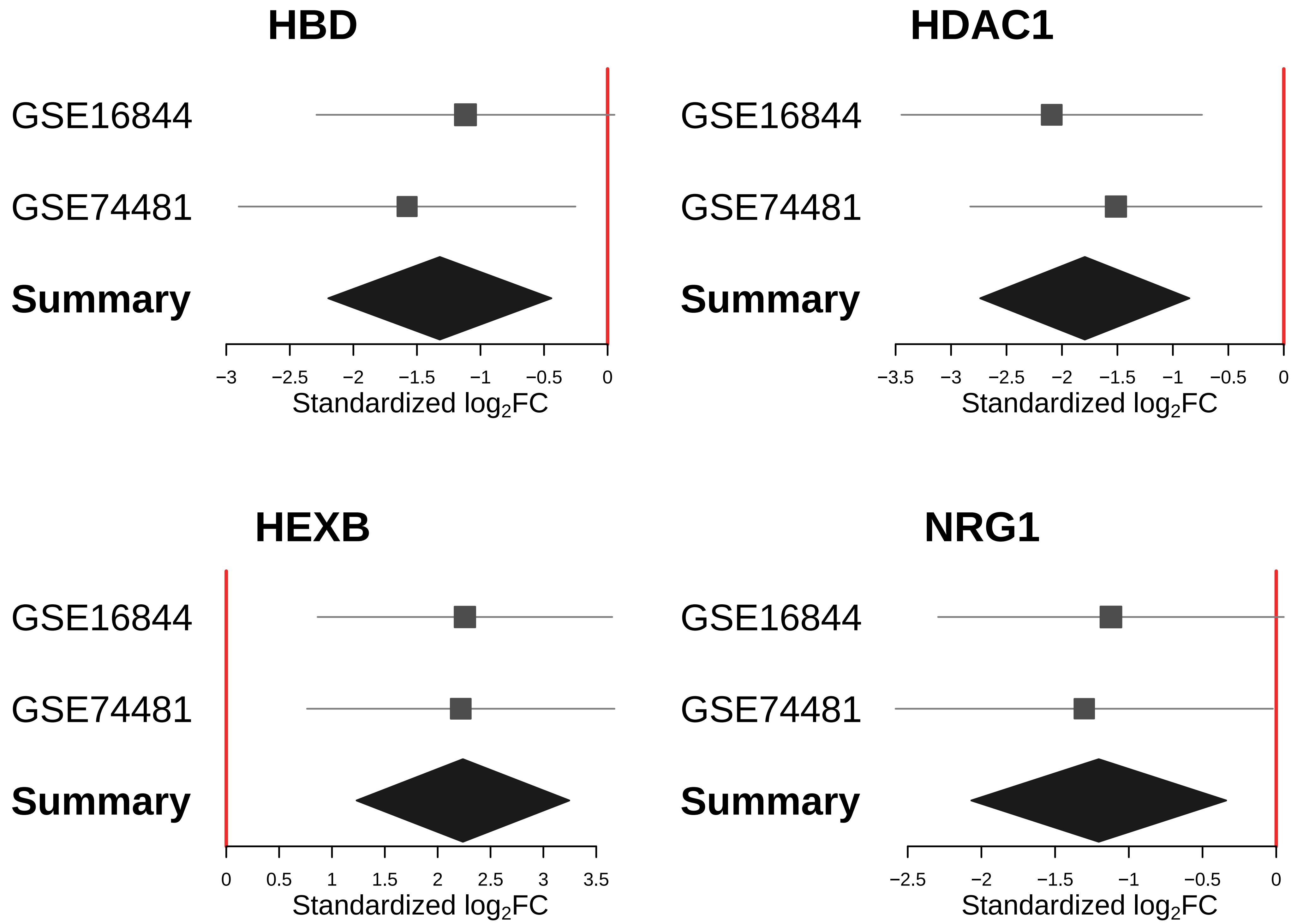
HBD
GSE16844
GSE74481
Summary
−3 −2.5 −2 −1.5 −1 −0.5 0
Standardized log2FC
HDAC1
GSE16844
GSE74481
Summary
−3.5 −3 −2.5 −2 −1.5 −1 −0.5 0
Standardized log2FC
HEXB
GSE16844
GSE74481
Summary
0 0.5 1 1.5 2 2.5 3 3.5
Standardized log2FC
NRG1
GSE16844
GSE74481
Summary
−2.5 −2 −1.5 −1 −0.5 0
Standardized log2FC

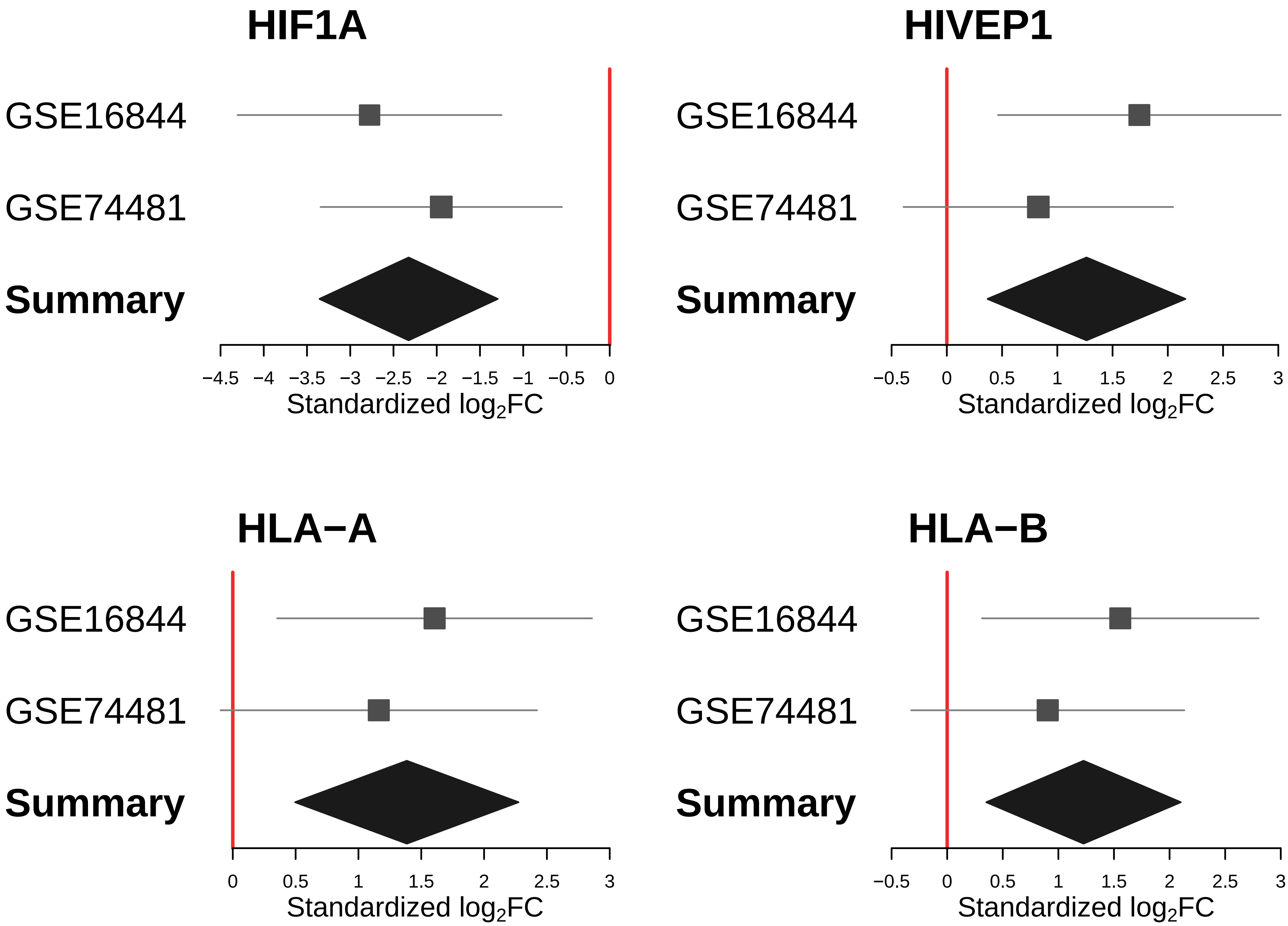
HIF1A
GSE16844
GSE74481
Summary
−4.5 −4 −3.5 −3 −2.5 −2 −1.5 −1 −0.5 0
Standardized log2FC
HIVEP1
GSE16844
GSE74481
Summary
−0.5 0 0.5 1 1.5 2 2.5 3
Standardized log2FC
HLA−A
GSE16844
GSE74481
Summary
0 0.5 1 1.5 2 2.5 3
Standardized log2FC
HLA−B
GSE16844
GSE74481
Summary
−0.5 0 0.5 1 1.5 2 2.5 3
Standardized log2FC

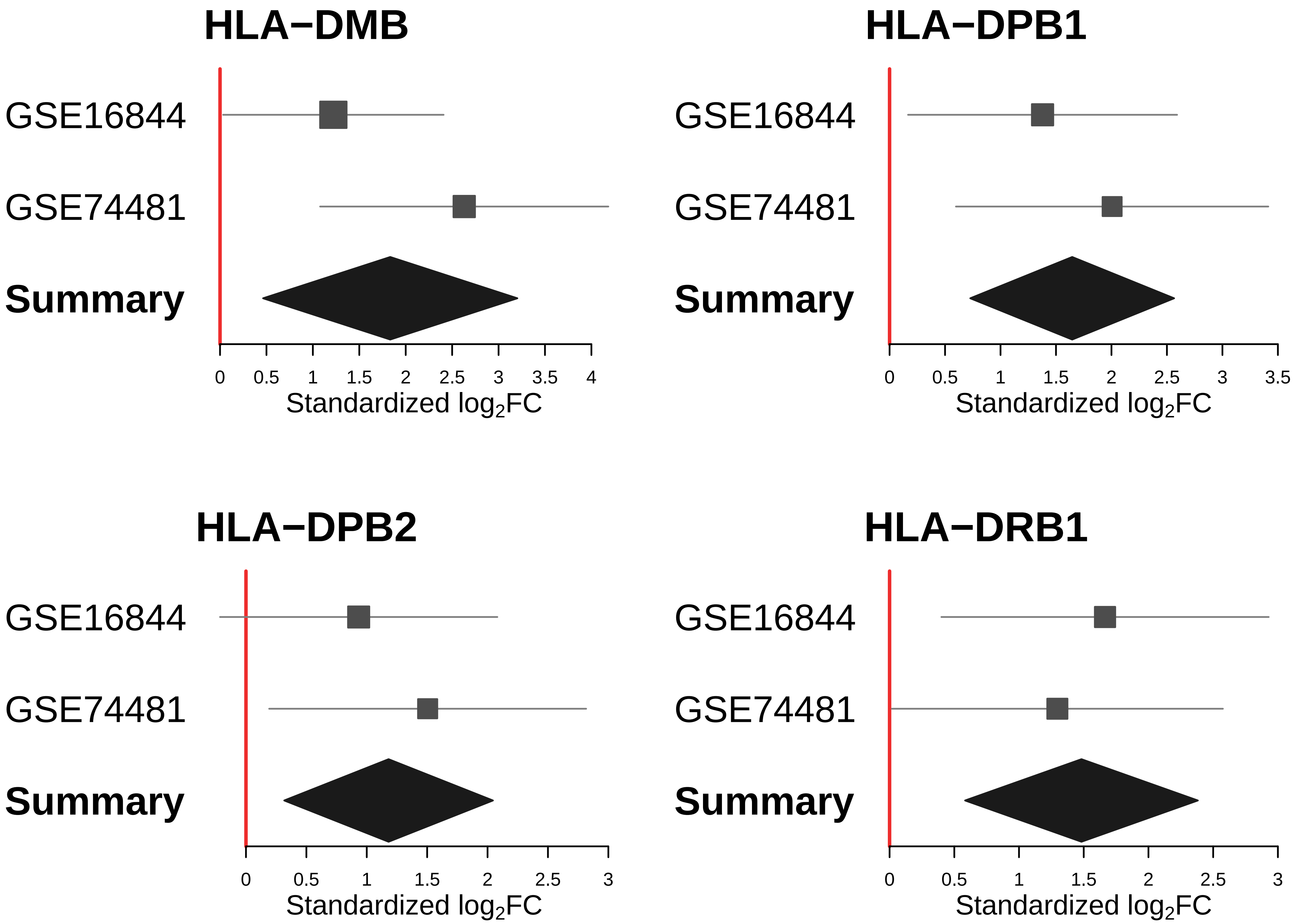
HLA−DMB
HLA−DPB1
HLA−DPB2
HLA−DRB1
GSE16844
GSE74481
Summary
0 0.5 1 1.5 2 2.5 3 3.5 4
0 0.5 1 1.5 2 2.5 3 3.5
0 0.5 1 1.5 2 2.5 3
Standardized log2FC

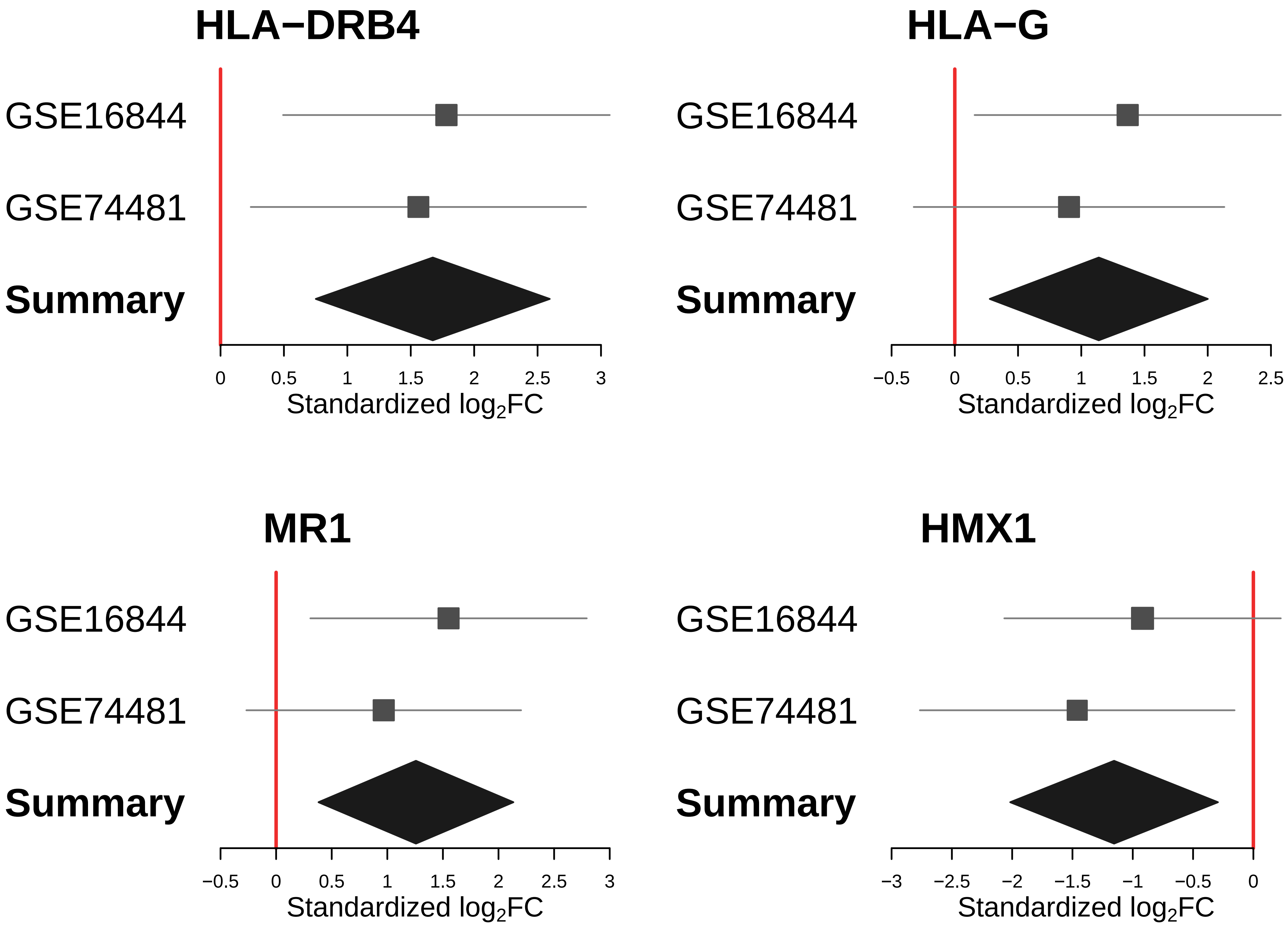

HLA−DRB4
GSE16844
GSE74481
Summary
0 0.5 1 1.5 2 2.5 3
Standardized log2FC
HLA−G
GSE16844
GSE74481
Summary
−0.5 0 0.5 1 1.5 2 2.5
Standardized log2FC
MR1
GSE16844
GSE74481
Summary
−0.5 0 0.5 1 1.5 2 2.5 3
Standardized log2FC
HMX1
GSE16844
GSE74481
Summary
−3 −2.5 −2 −1.5 −1 −0.5 0
Standardized log2FC

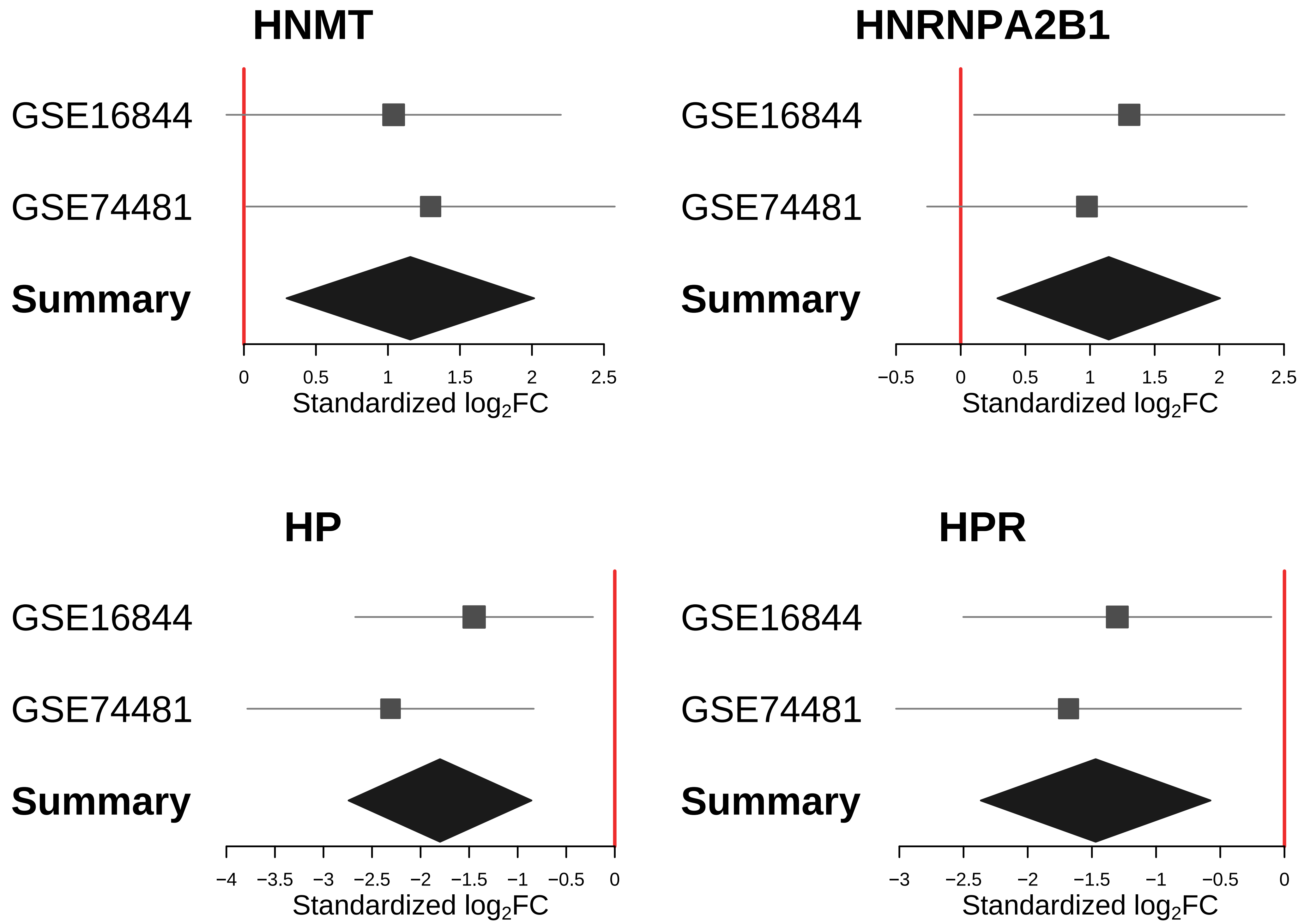
HNMT
GSE16844
GSE74481
Summary
0
0.5
1
1.5
2
2.5
Standardized log2FC
HNRNPA2B1
GSE16844
GSE74481
Summary
−0.5
0
0.5
1
1.5
2
2.5
Standardized log2FC
HP
GSE16844
GSE74481
Summary
−4
−3.5
−3
−2.5
−2
−1.5
−1
−0.5
0
Standardized log2FC
HPR
GSE16844
GSE74481
Summary
−3
−2.5
−2
−1.5
−1
−0.5
0
Standardized log2FC

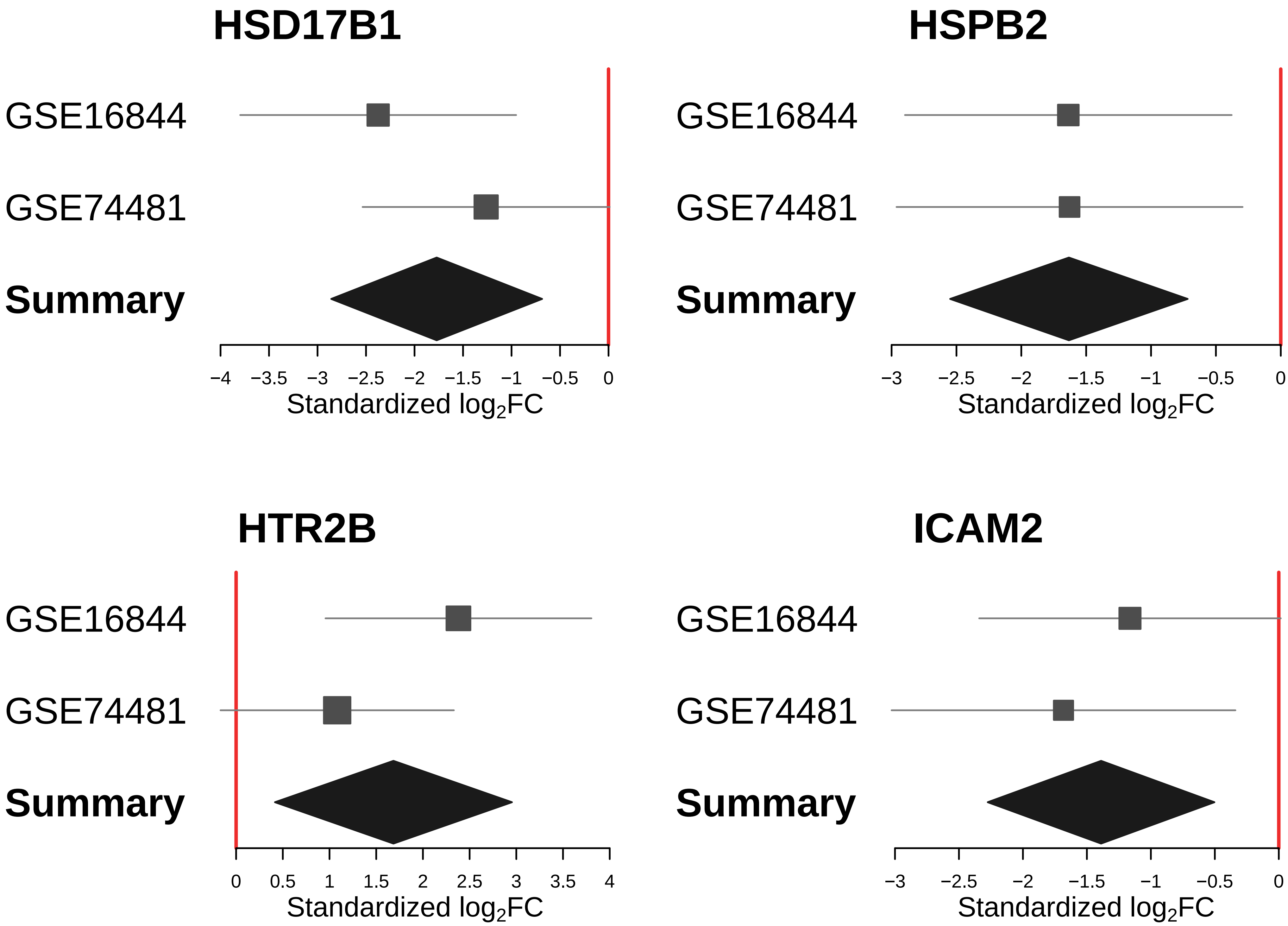
HSD17B1
GSE16844
GSE74481
Summary
−4 −3.5 −3 −2.5 −2 −1.5 −1 −0.5 0
Standardized log2FC
HSPB2
GSE16844
GSE74481
Summary
−3 −2.5 −2 −1.5 −1 −0.5 0
Standardized log2FC
HTR2B
GSE16844
GSE74481
Summary
0 0.5 1 1.5 2 2.5 3 3.5 4
Standardized log2FC
ICAM2
GSE16844
GSE74481
Summary
−3 −2.5 −2 −1.5 −1 −0.5 0
Standardized log2FC

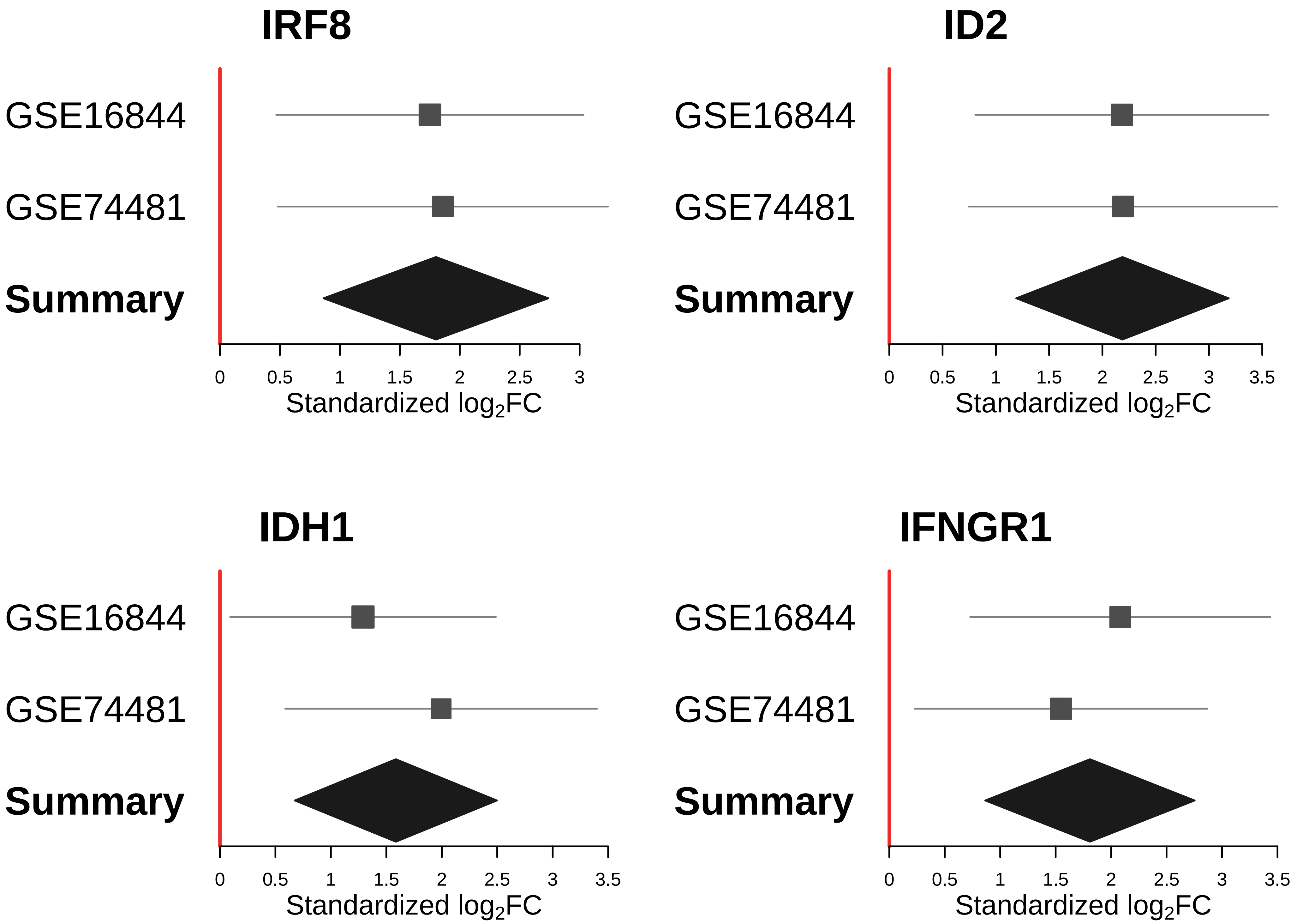

IRF8
GSE16844
GSE74481
Summary
0 0.5 1 1.5 2 2.5 3
Standardized log2FC
ID2
GSE16844
GSE74481
Summary
0 0.5 1 1.5 2 2.5 3 3.5
Standardized log2FC
IDH1
GSE16844
GSE74481
Summary
0 0.5 1 1.5 2 2.5 3 3.5
Standardized log2FC
IFNGR1
GSE16844
GSE74481
Summary
0 0.5 1 1.5 2 2.5 3 3.5
Standardized log2FC

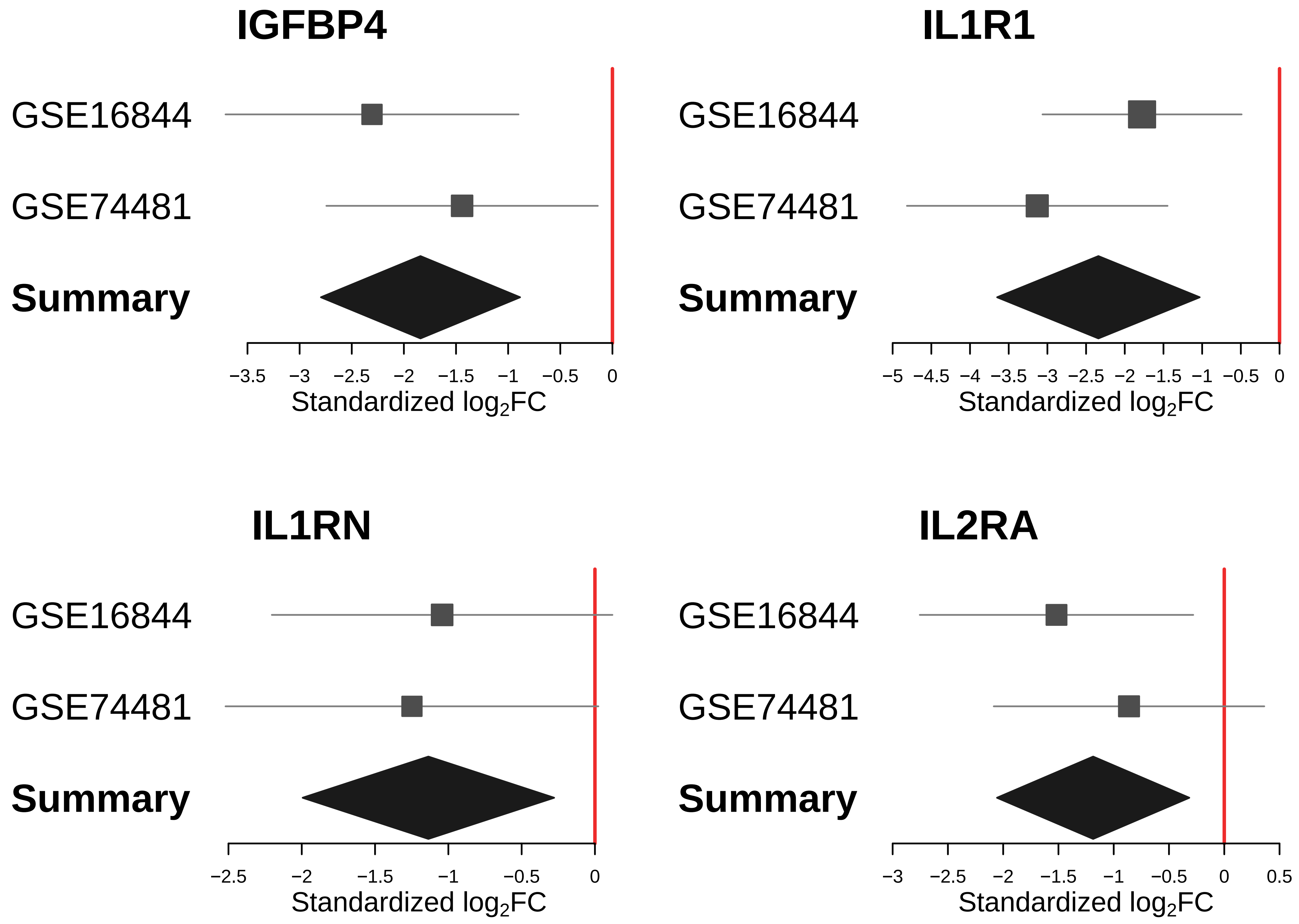
IGFBP4
GSE16844
GSE74481
Summary
−3.5 −3 −2.5 −2 −1.5 −1 −0.5 0
Standardized log2FC
IL1R1
GSE16844
GSE74481
Summary
−5 −4.5 −4 −3.5 −3 −2.5 −2 −1.5 −1 −0.5 0
Standardized log2FC
IL1RN
GSE16844
GSE74481
Summary
−2.5 −2 −1.5 −1 −0.5 0
Standardized log2FC
IL2RA
GSE16844
GSE74481
Summary
−3 −2.5 −2 −1.5 −1 −0.5 0 0.5
Standardized log2FC

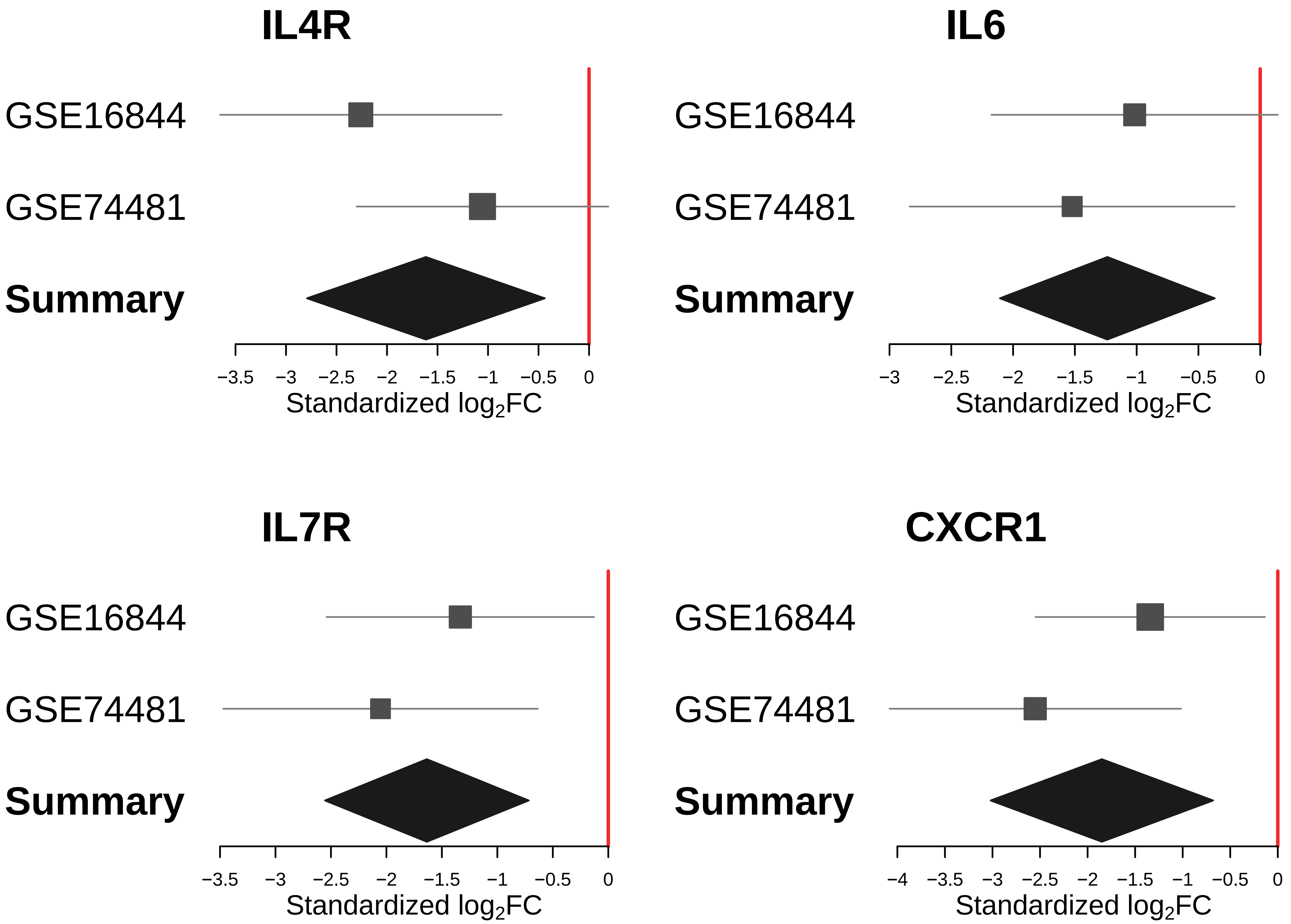
IL4R
GSE16844
GSE74481
Summary
−3.5 −3 −2.5 −2 −1.5 −1 −0.5 0
Standardized log2FC
IL6
GSE16844
GSE74481
Summary
−3 −2.5 −2 −1.5 −1 −0.5 0
Standardized log2FC
IL7R
GSE16844
GSE74481
Summary
−3.5 −3 −2.5 −2 −1.5 −1 −0.5 0
Standardized log2FC
CXCR1
GSE16844
GSE74481
Summary
−4 −3.5 −3 −2.5 −2 −1.5 −1 −0.5 0
Standardized log2FC

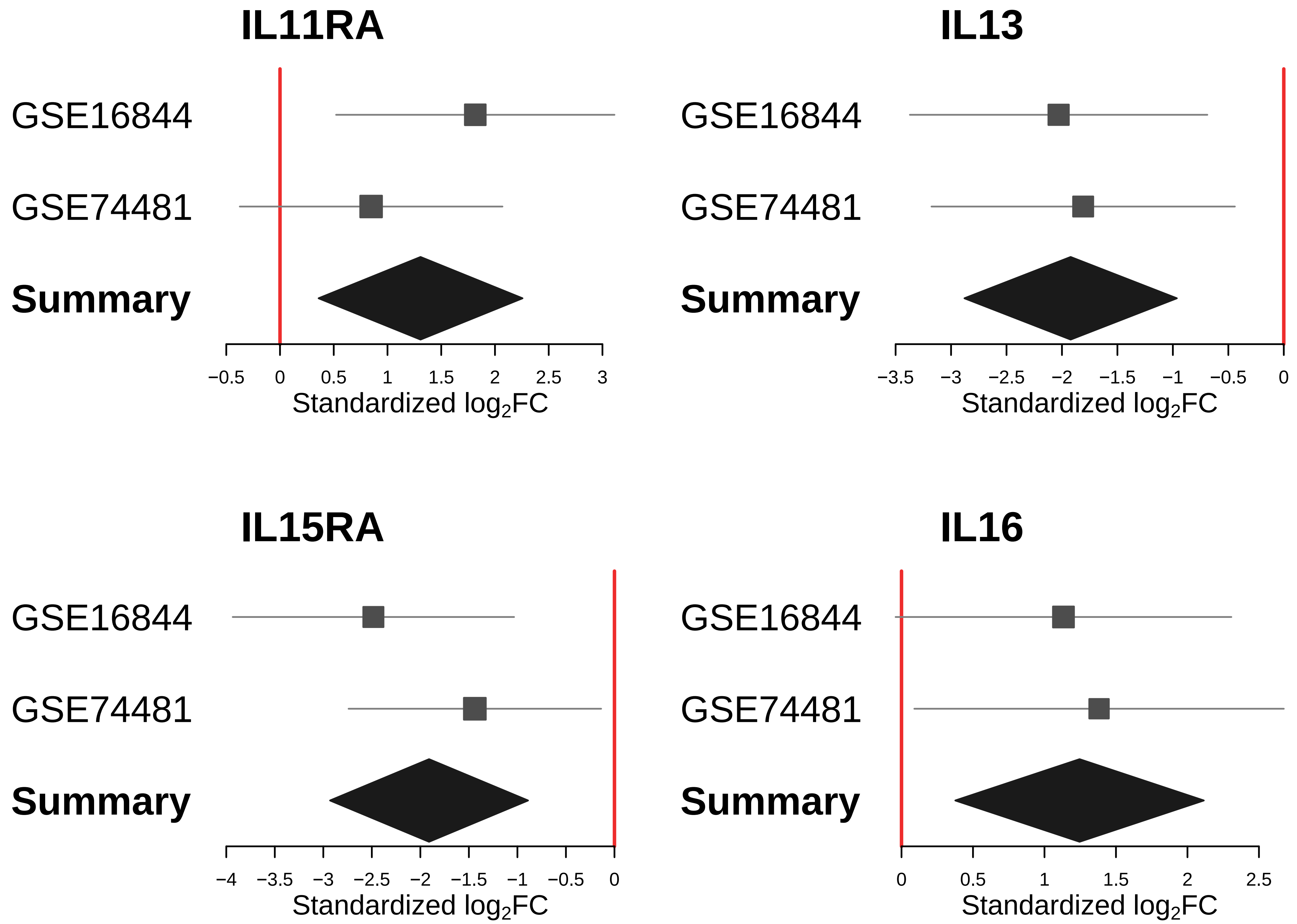
IL11RA
GSE16844
GSE74481
Summary
−0.5 0 0.5 1 1.5 2 2.5 3
Standardized $\log_2$FC
IL13
GSE16844
GSE74481
Summary
−3.5 −3 −2.5 −2 −1.5 −1 −0.5 0
Standardized $\log_2$FC
IL15RA
GSE16844
GSE74481
Summary
−4 −3.5 −3 −2.5 −2 −1.5 −1 −0.5 0
Standardized $\log_2$FC
IL16
GSE16844
GSE74481
Summary
0 0.5 1 1.5 2 2.5
Standardized $\log_2$FC

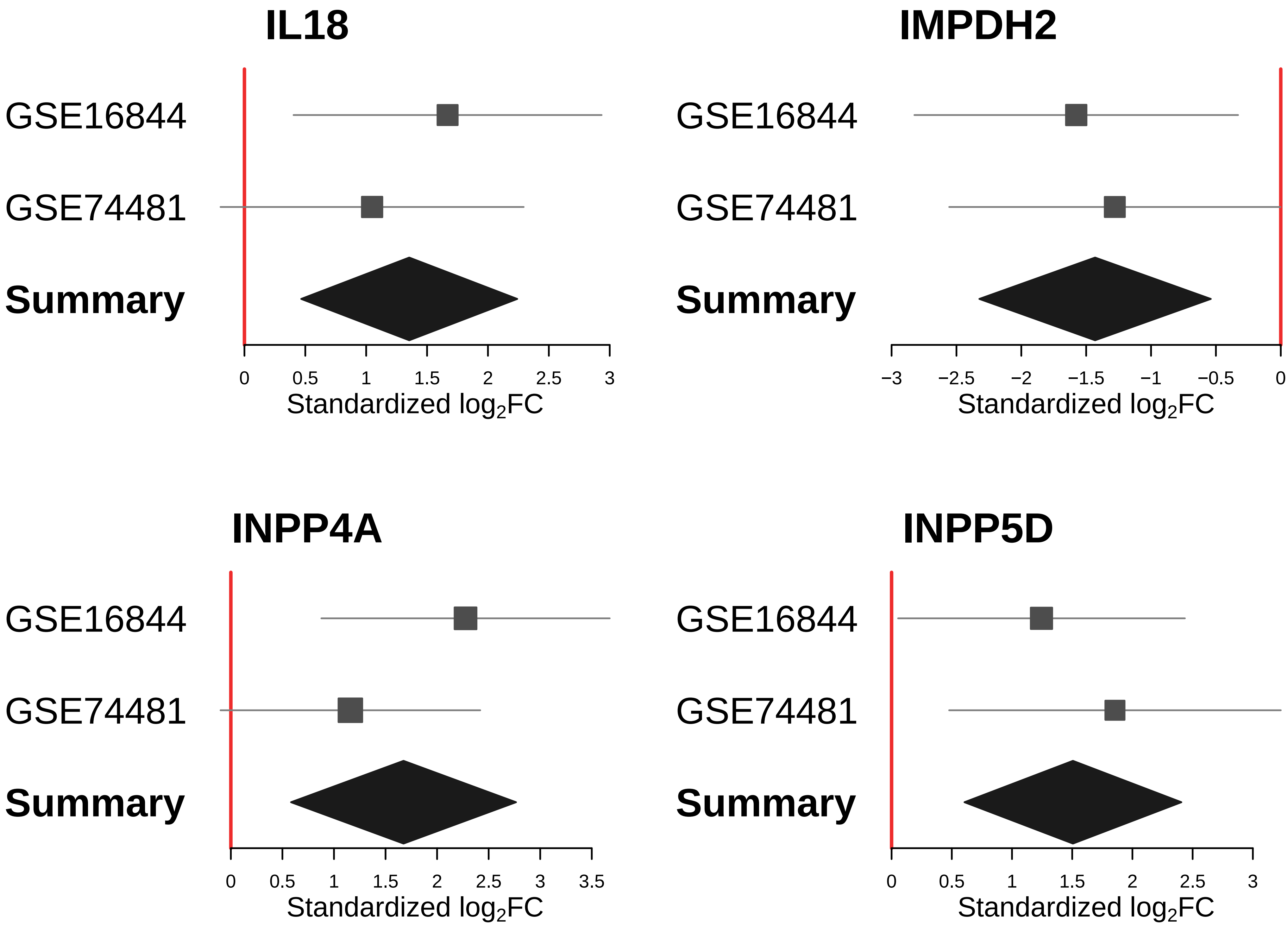

IL18
GSE16844
GSE74481
Summary
0 0.5 1 1.5 2 2.5 3
Standardized log2FC
IMPDH2
GSE16844
GSE74481
Summary
−3 −2.5 −2 −1.5 −1 −0.5 0
Standardized log2FC
INPP4A
GSE16844
GSE74481
Summary
0 0.5 1 1.5 2 2.5 3 3.5
Standardized log2FC
INPP5D
GSE16844
GSE74481
Summary
0 0.5 1 1.5 2 2.5 3
Standardized log2FC

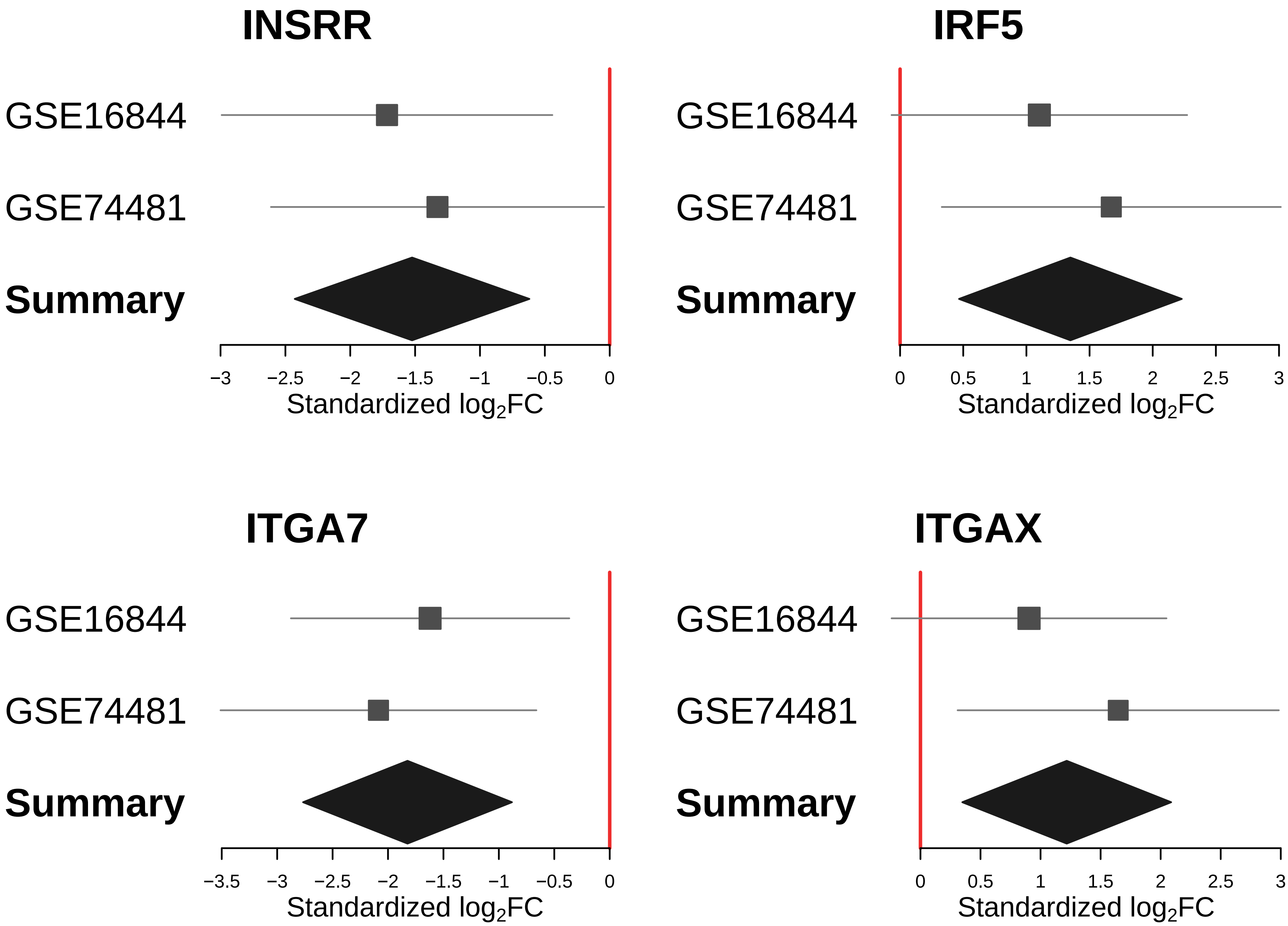

INSRR
GSE16844
GSE74481
Summary
−3
−2.5
−2
−1.5
−1
−0.5
0
Standardized log2FC
IRF5
GSE16844
GSE74481
Summary
0
0.5
1
1.5
2
2.5
3
Standardized log2FC
ITGA7
GSE16844
GSE74481
Summary
−3.5
−3
−2.5
−2
−1.5
−1
−0.5
0
Standardized log2FC
ITGAX
GSE16844
GSE74481
Summary
0
0.5
1
1.5
2
2.5
3
Standardized log2FC

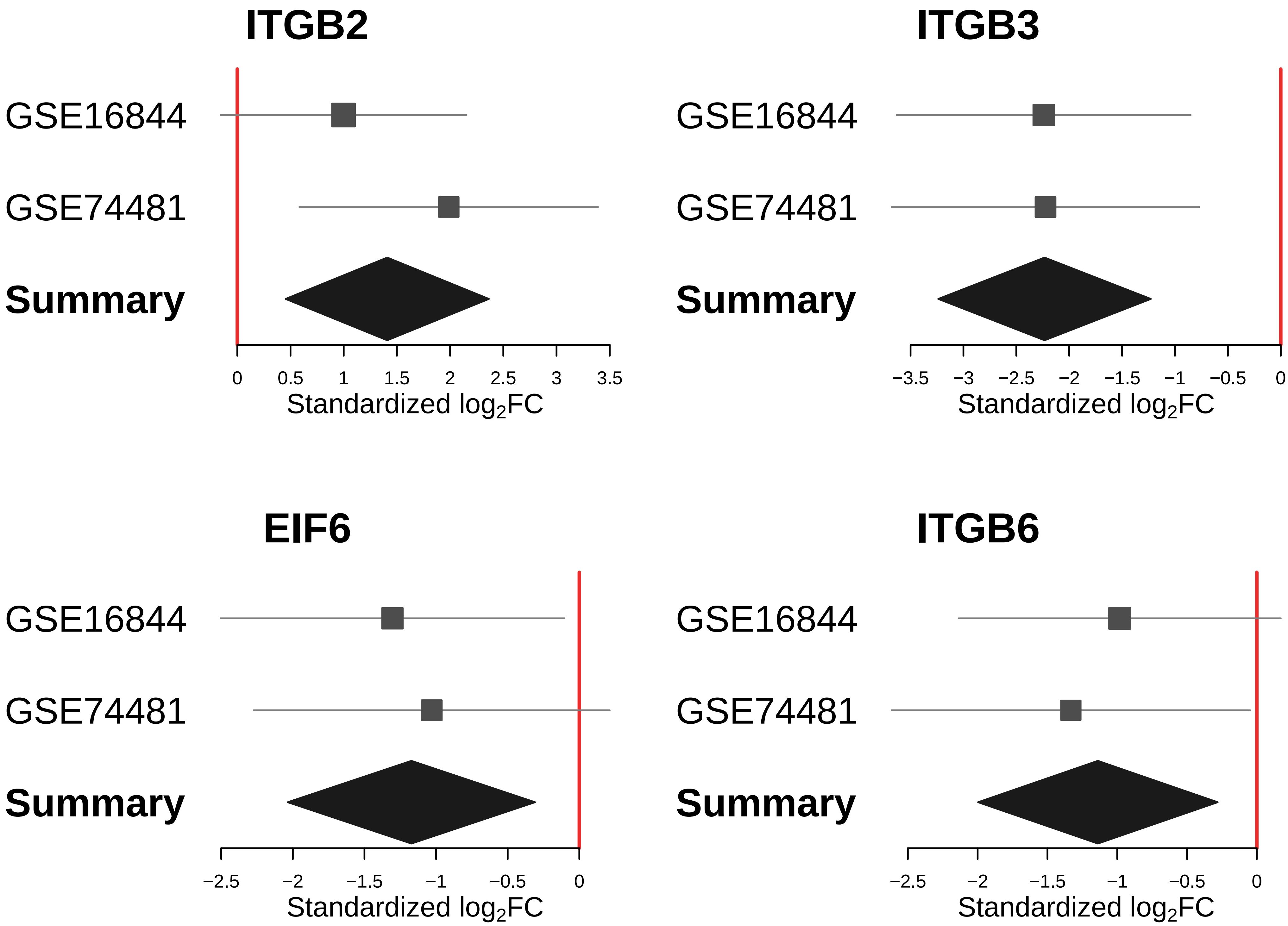

ITGB2
GSE16844
GSE74481
Summary
0
0.5
1
1.5
2
2.5
3
3.5
Standardized log2FC
ITGB3
GSE16844
GSE74481
Summary
−3.5
−3
−2.5
−2
−1.5
−1
−0.5
0
Standardized log2FC
EIF6
GSE16844
GSE74481
Summary
−2.5
−2
−1.5
−1
−0.5
0
Standardized log2FC
ITGB6
GSE16844
GSE74481
Summary
−2.5
−2
−1.5
−1
−0.5
0
Standardized log2FC

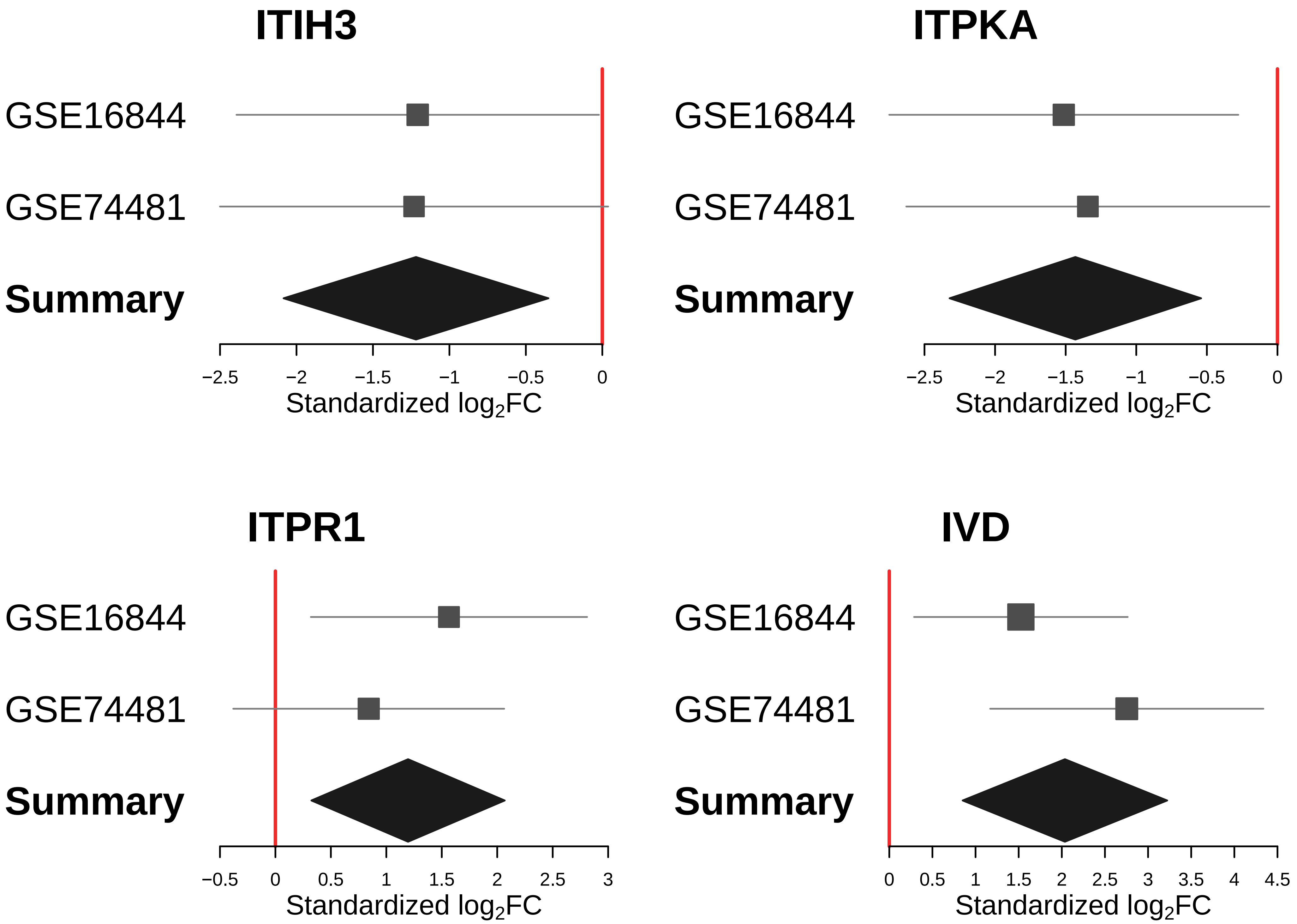
ITIH3
GSE16844
GSE74481
Summary
−2.5 −2 −1.5 −1 −0.5 0
Standardized log2FC
ITPKA
GSE16844
GSE74481
Summary
−2.5 −2 −1.5 −1 −0.5 0
Standardized log2FC
ITPR1
GSE16844
GSE74481
Summary
−0.5 0 0.5 1 1.5 2 2.5 3
Standardized log2FC
IVD
GSE16844
GSE74481
Summary
0 0.5 1 1.5 2 2.5 3 3.5 4 4.5
Standardized log2FC

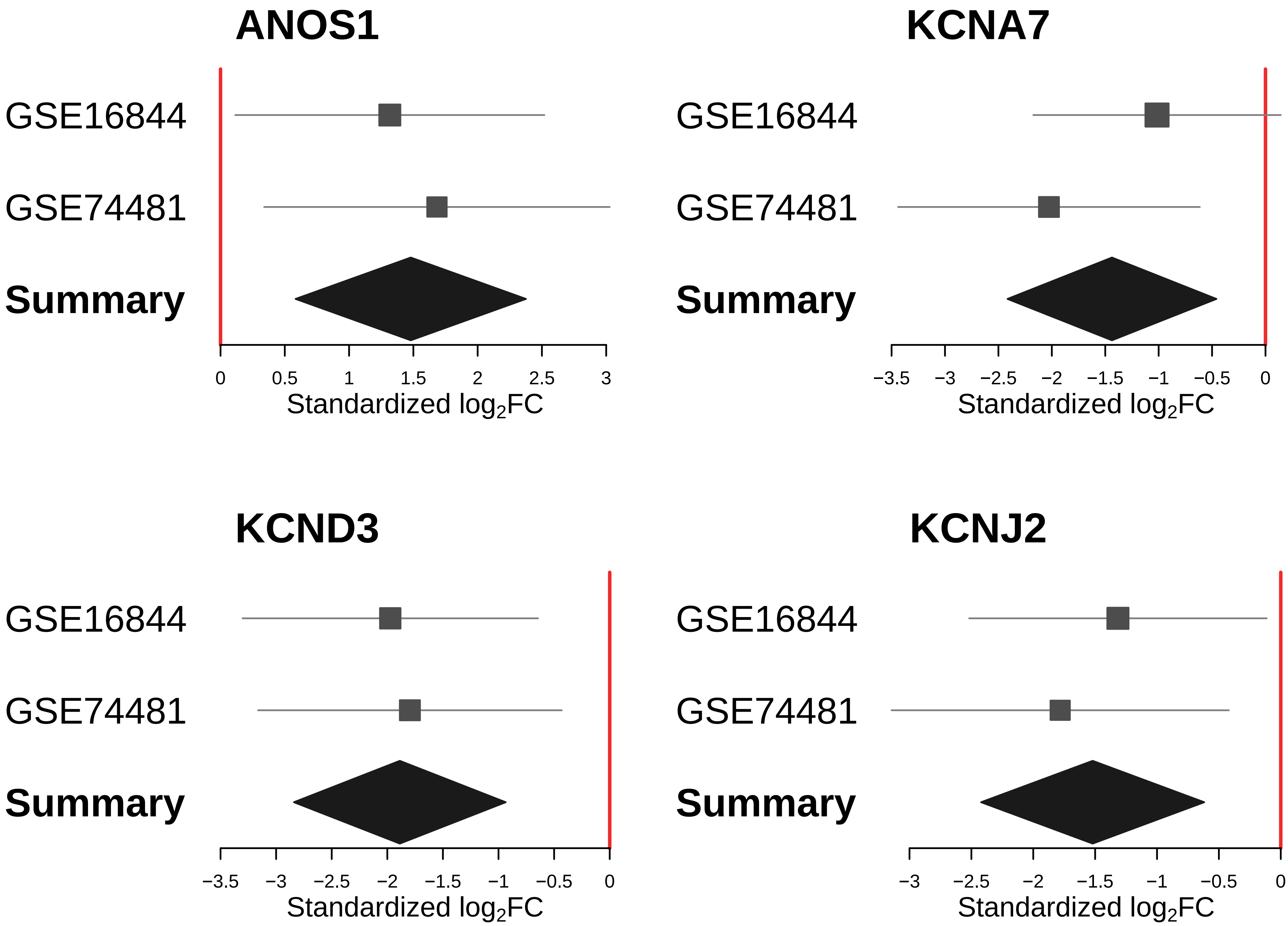
ANOS1
GSE16844
GSE74481
Summary
0
0.5
1
1.5
2
2.5
3
Standardized log2FC
KCNA7
GSE16844
GSE74481
Summary
−3.5
−3
−2.5
−2
−1.5
−1
−0.5
0
Standardized log2FC
KCND3
GSE16844
GSE74481
Summary
−3.5
−3
−2.5
−2
−1.5
−1
−0.5
0
Standardized log2FC
KCNJ2
GSE16844
GSE74481
Summary
−3
−2.5
−2
−1.5
−1
−0.5
0
Standardized log2FC

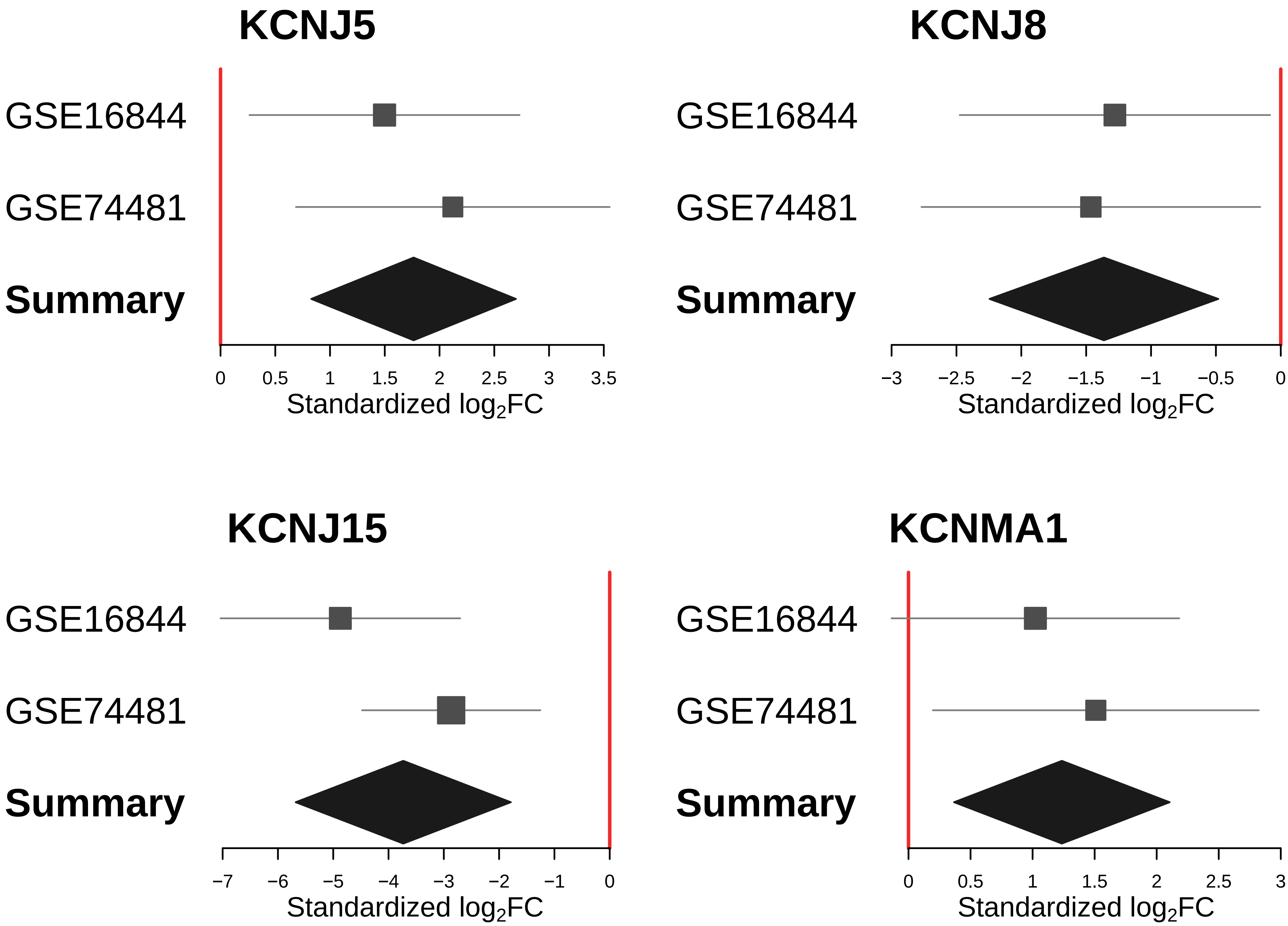
KCNJ5
GSE16844
GSE74481
Summary
0 0.5 1 1.5 2 2.5 3 3.5
Standardized log2FC
KCNJ8
GSE16844
GSE74481
Summary
−3 −2.5 −2 −1.5 −1 −0.5 0
Standardized log2FC
KCNJ15
GSE16844
GSE74481
Summary
−7 −6 −5 −4 −3 −2 −1 0
Standardized log2FC
KCNMA1
GSE16844
GSE74481
Summary
0 0.5 1 1.5 2 2.5 3
Standardized log2FC

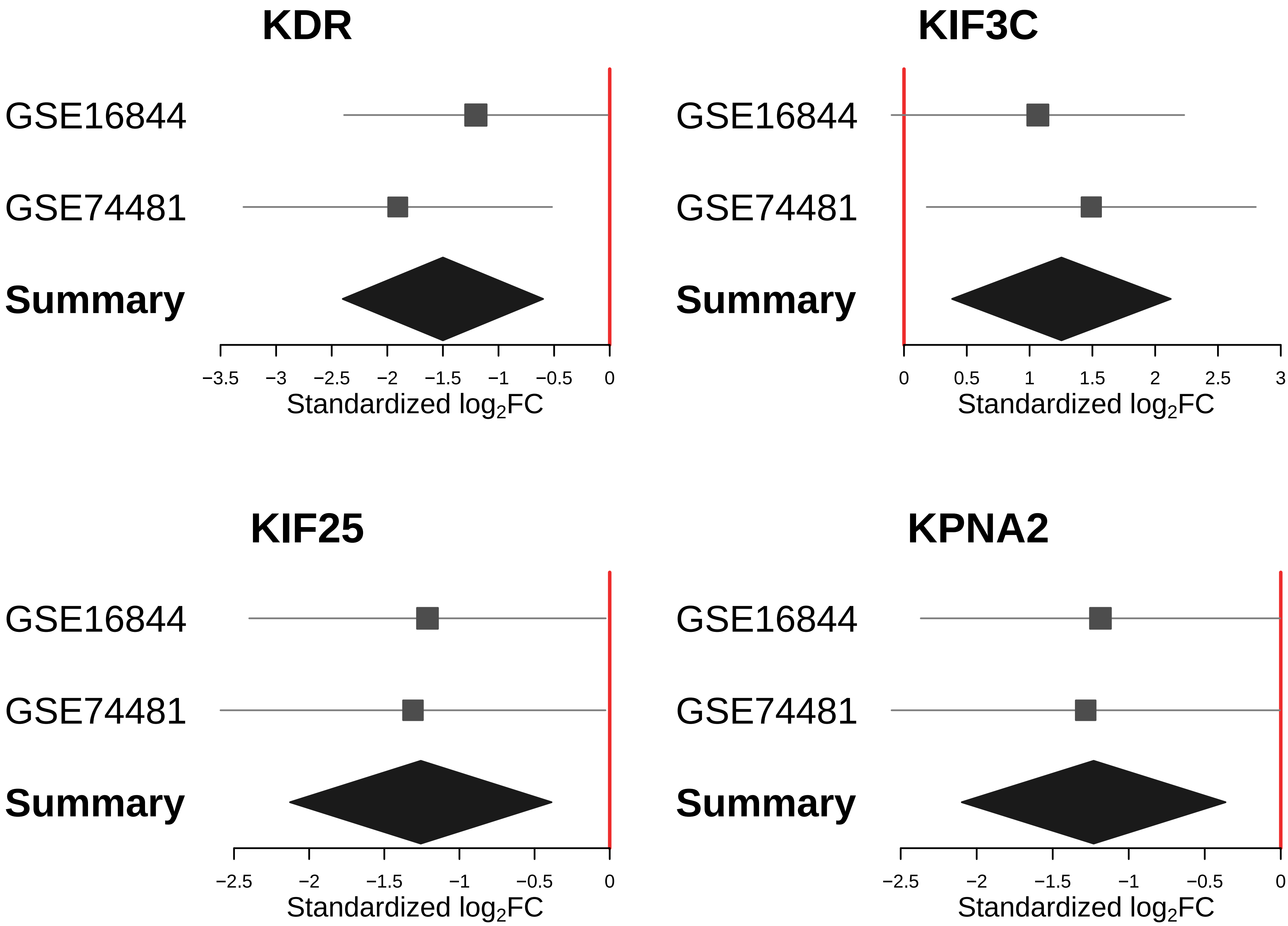

KDR
GSE16844
GSE74481
Summary
−3.5 −3 −2.5 −2 −1.5 −1 −0.5 0
Standardized log2FC
KIF3C
GSE16844
GSE74481
Summary
0 0.5 1 1.5 2 2.5 3
Standardized log2FC
KIF25
GSE16844
GSE74481
Summary
−2.5 −2 −1.5 −1 −0.5 0
Standardized log2FC
KPNA2
GSE16844
GSE74481
Summary
−2.5 −2 −1.5 −1 −0.5 0
Standardized log2FC

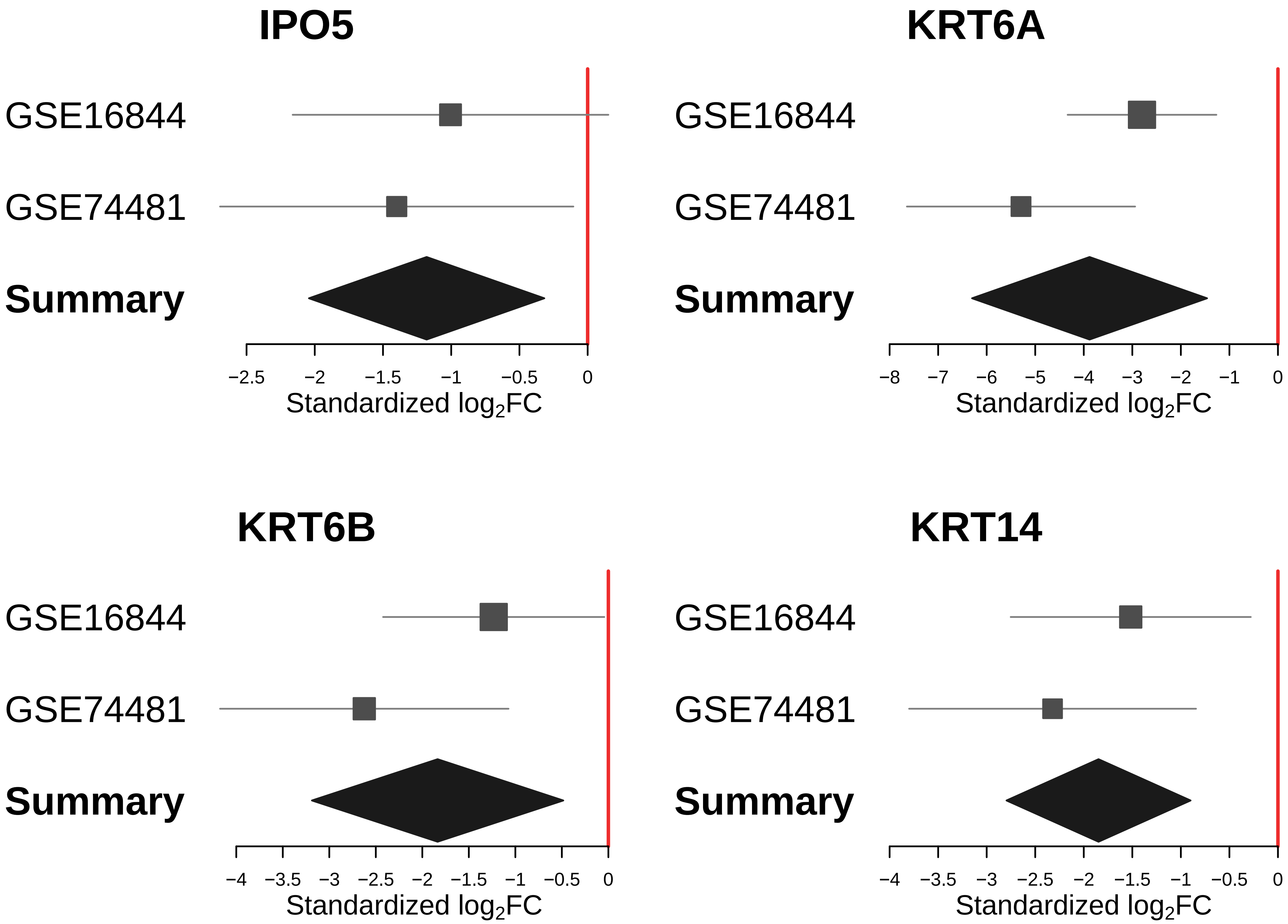
IPO5
GSE16844
GSE74481
Summary
−2.5 −2 −1.5 −1 −0.5 0
Standardized log2FC
KRT6A
GSE16844
GSE74481
Summary
−8 −7 −6 −5 −4 −3 −2 −1 0
Standardized log2FC
KRT6B
GSE16844
GSE74481
Summary
−4 −3.5 −3 −2.5 −2 −1.5 −1 −0.5 0
Standardized log2FC
KRT14
GSE16844
GSE74481
Summary
−4 −3.5 −3 −2.5 −2 −1.5 −1 −0.5 0
Standardized log2FC

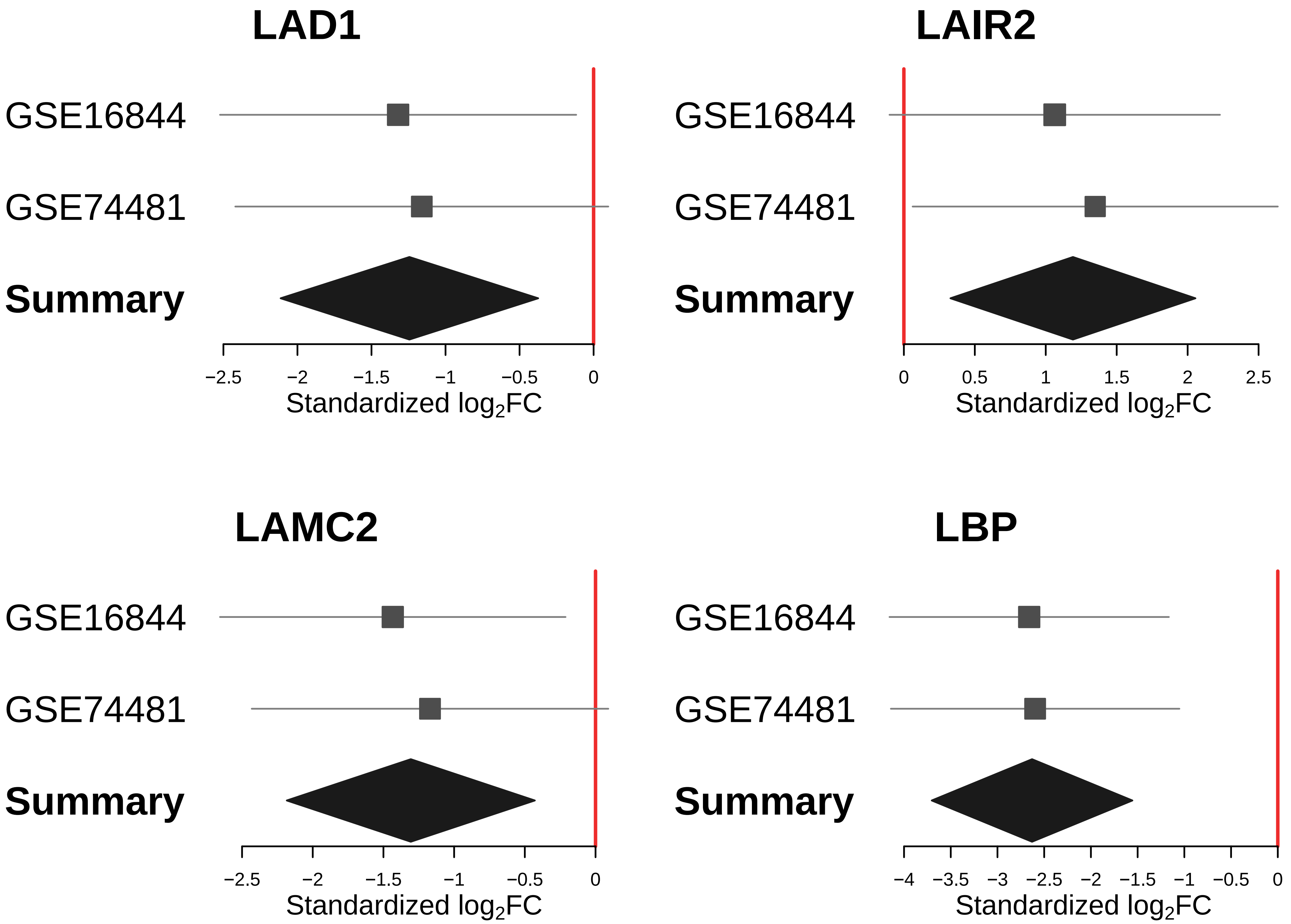
LAD1
GSE16844
GSE74481
Summary
−2.5 −2 −1.5 −1 −0.5 0
Standardized log2FC
LAIR2
GSE16844
GSE74481
Summary
0 0.5 1 1.5 2 2.5
Standardized log2FC
LAMC2
GSE16844
GSE74481
Summary
−2.5 −2 −1.5 −1 −0.5 0
Standardized log2FC
LBP
GSE16844
GSE74481
Summary
−4 −3.5 −3 −2.5 −2 −1.5 −1 −0.5 0
Standardized log2FC

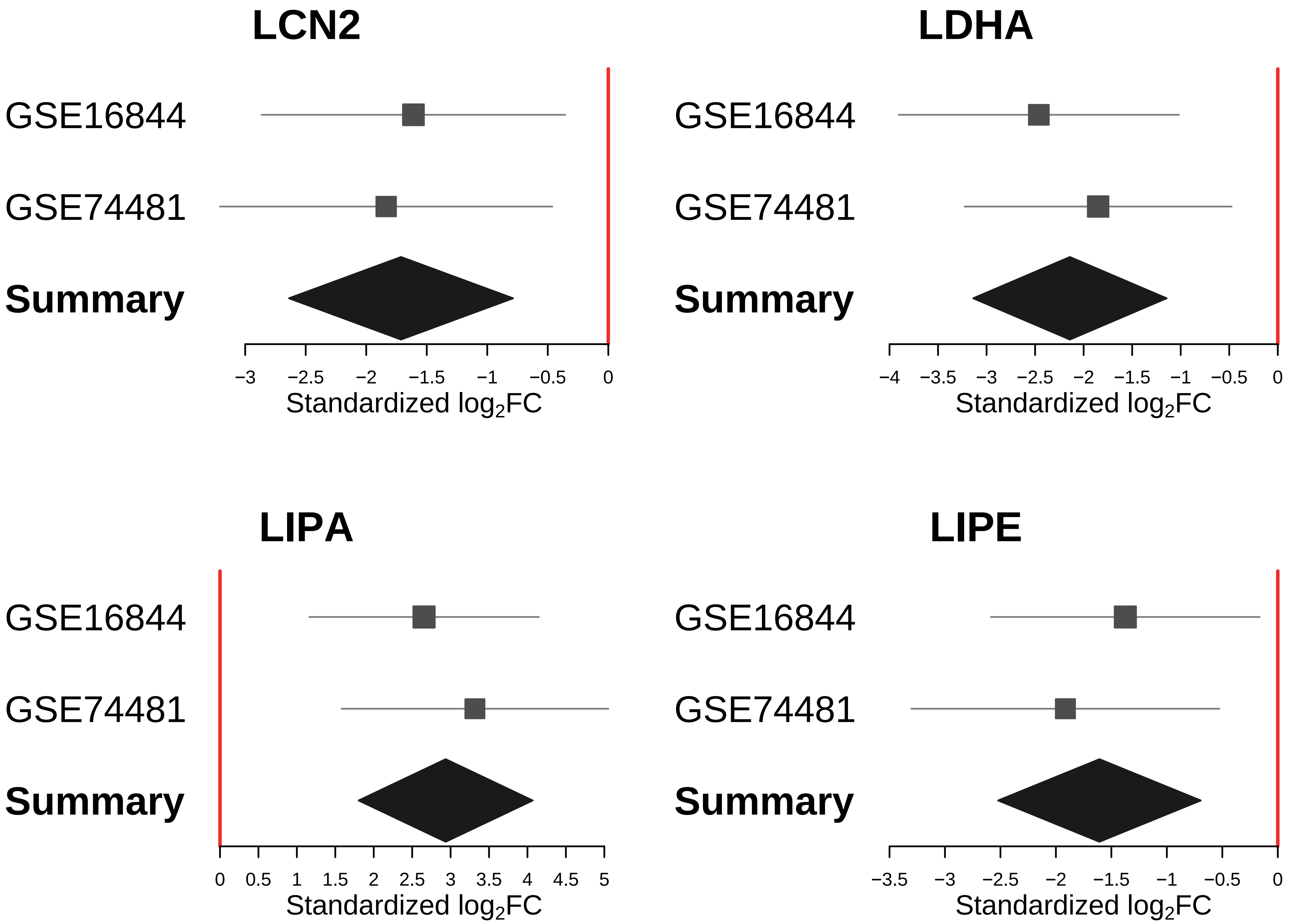
LCN2
GSE16844
GSE74481
Summary
−3 −2.5 −2 −1.5 −1 −0.5 0
Standardized log2FC
LDHA
GSE16844
GSE74481
Summary
−4 −3.5 −3 −2.5 −2 −1.5 −1 −0.5 0
Standardized log2FC
LIPA
GSE16844
GSE74481
Summary
0 0.5 1 1.5 2 2.5 3 3.5 4 4.5 5
Standardized log2FC
LIPE
GSE16844
GSE74481
Summary
−3.5 −3 −2.5 −2 −1.5 −1 −0.5 0
Standardized log2FC

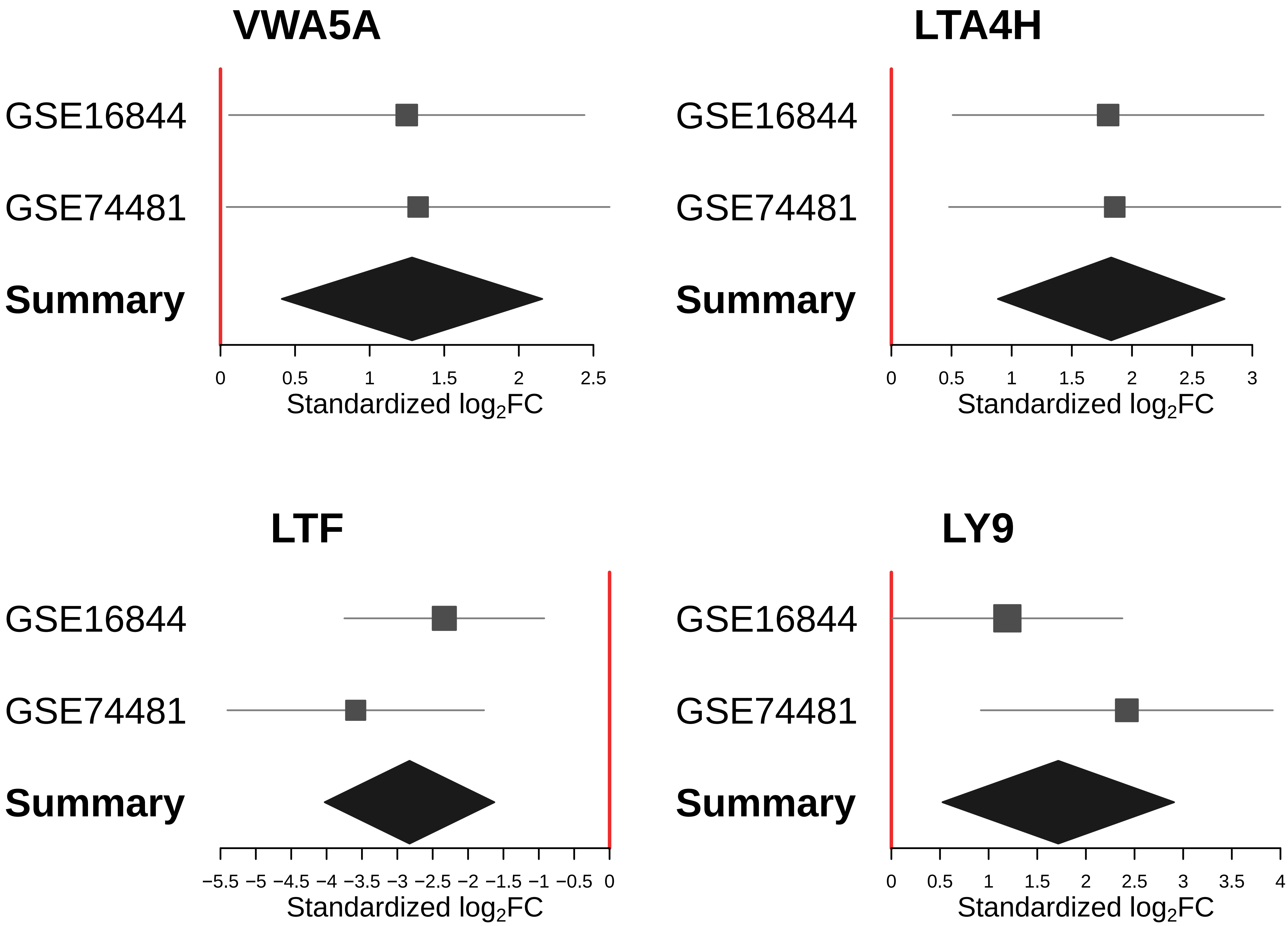
VWA5A
GSE16844
GSE74481
Summary
0
0.5
1
1.5
2
2.5
Standardized log2FC
LTA4H
GSE16844
GSE74481
Summary
0
0.5
1
1.5
2
2.5
3
Standardized log2FC
LTF
GSE16844
GSE74481
Summary
−5.5
−5
−4.5
−4
−3.5
−3
−2.5
−2
−1.5
−1
−0.5
0
Standardized log2FC
LY9
GSE16844
GSE74481
Summary
0
0.5
1
1.5
2
2.5
3
3.5
4
Standardized log2FC

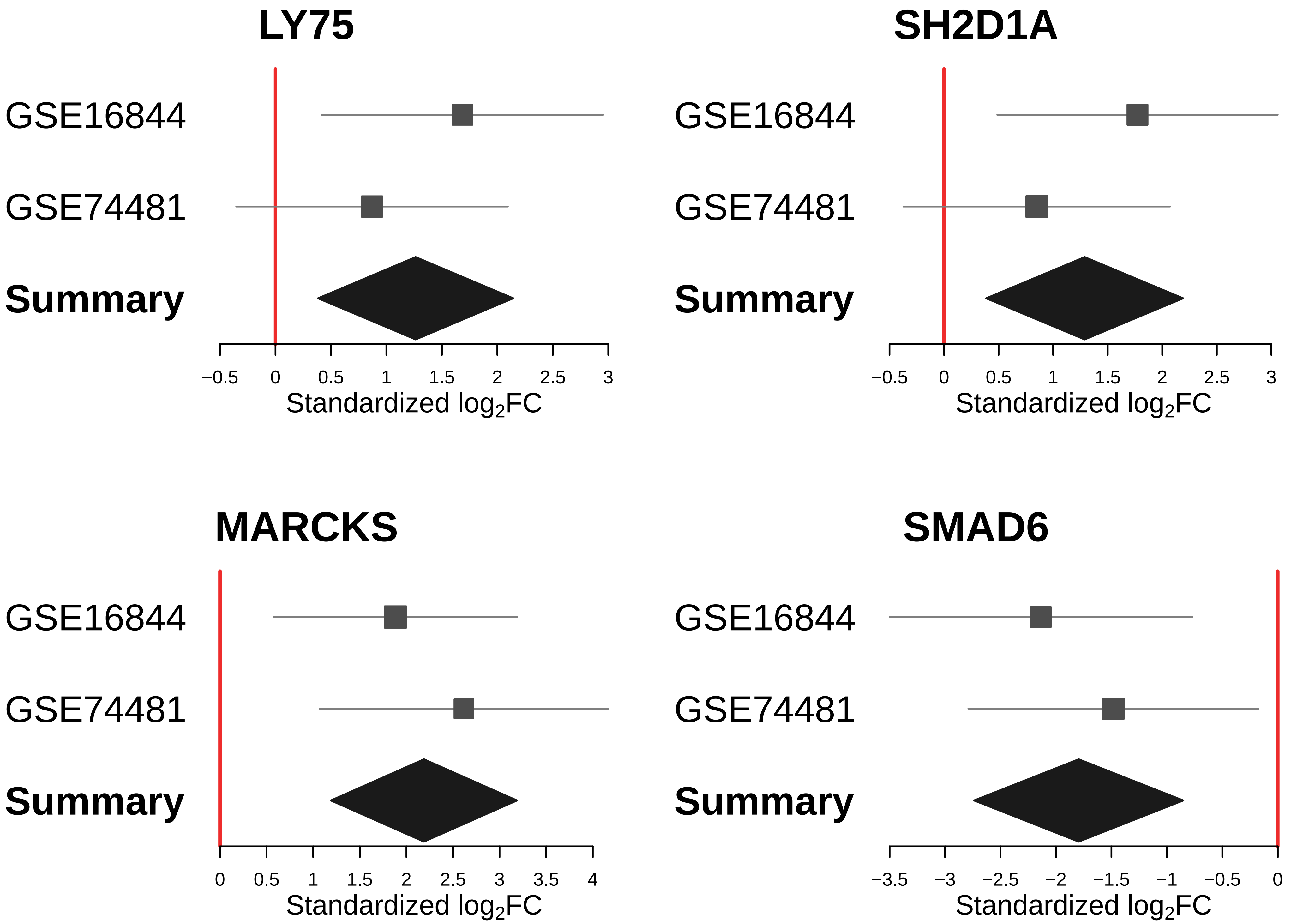
LY75
GSE16844
GSE74481
Summary
−0.5 0 0.5 1 1.5 2 2.5 3
Standardized log2FC
SH2D1A
GSE16844
GSE74481
Summary
−0.5 0 0.5 1 1.5 2 2.5 3
Standardized log2FC
MARCKS
GSE16844
GSE74481
Summary
0 0.5 1 1.5 2 2.5 3 3.5 4
Standardized log2FC
SMAD6
GSE16844
GSE74481
Summary
−3.5 −3 −2.5 −2 −1.5 −1 −0.5 0
Standardized log2FC

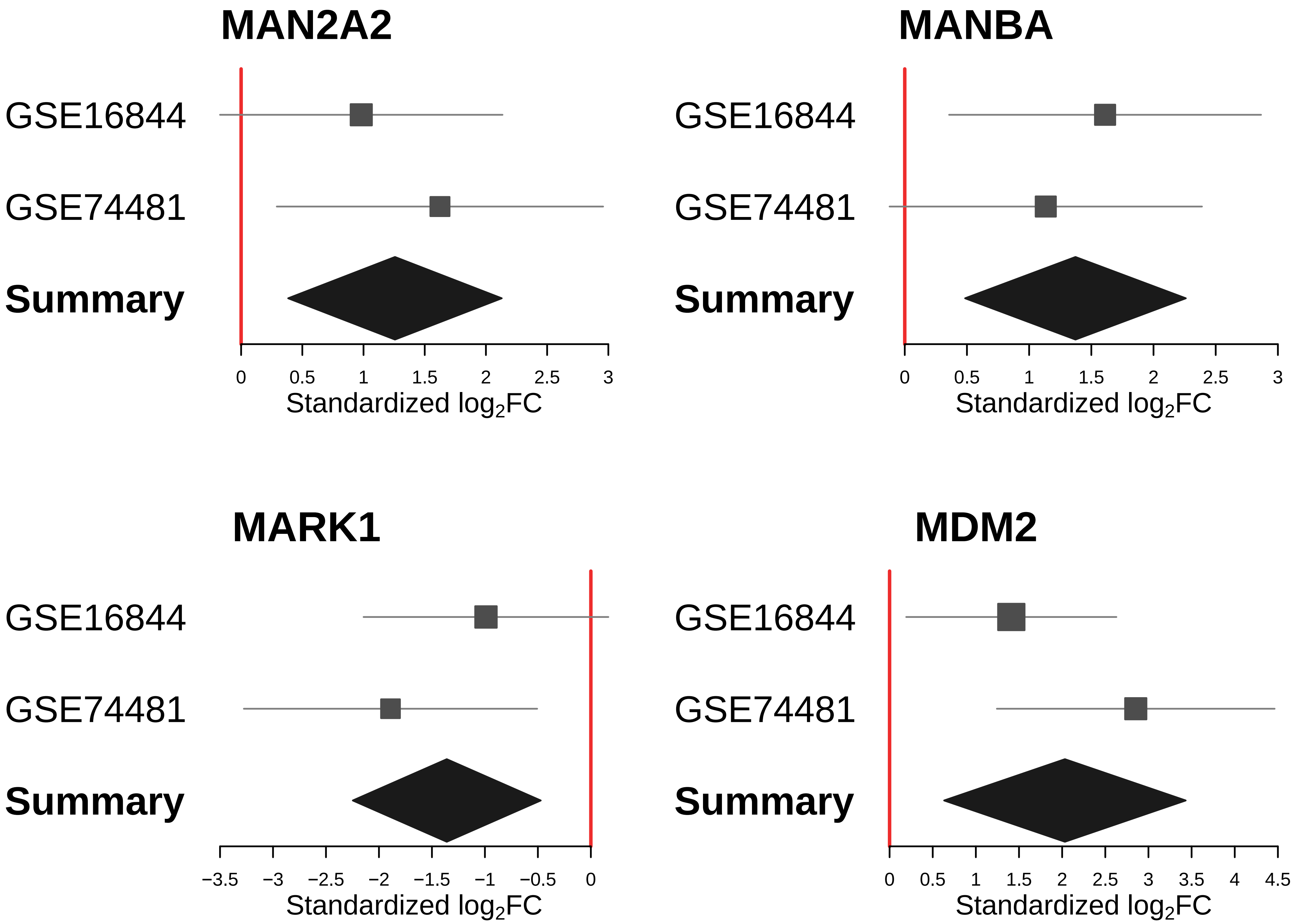
MAN2A2
GSE16844
GSE74481
Summary
0
0.5
1
1.5
2
2.5
3
Standardized log2FC
MANBA
GSE16844
GSE74481
Summary
0
0.5
1
1.5
2
2.5
3
Standardized log2FC
MARK1
GSE16844
GSE74481
Summary
−3.5
−3
−2.5
−2
−1.5
−1
−0.5
0
Standardized log2FC
MDM2
GSE16844
GSE74481
Summary
0
0.5
1
1.5
2
2.5
3
3.5
4
4.5
Standardized log2FC

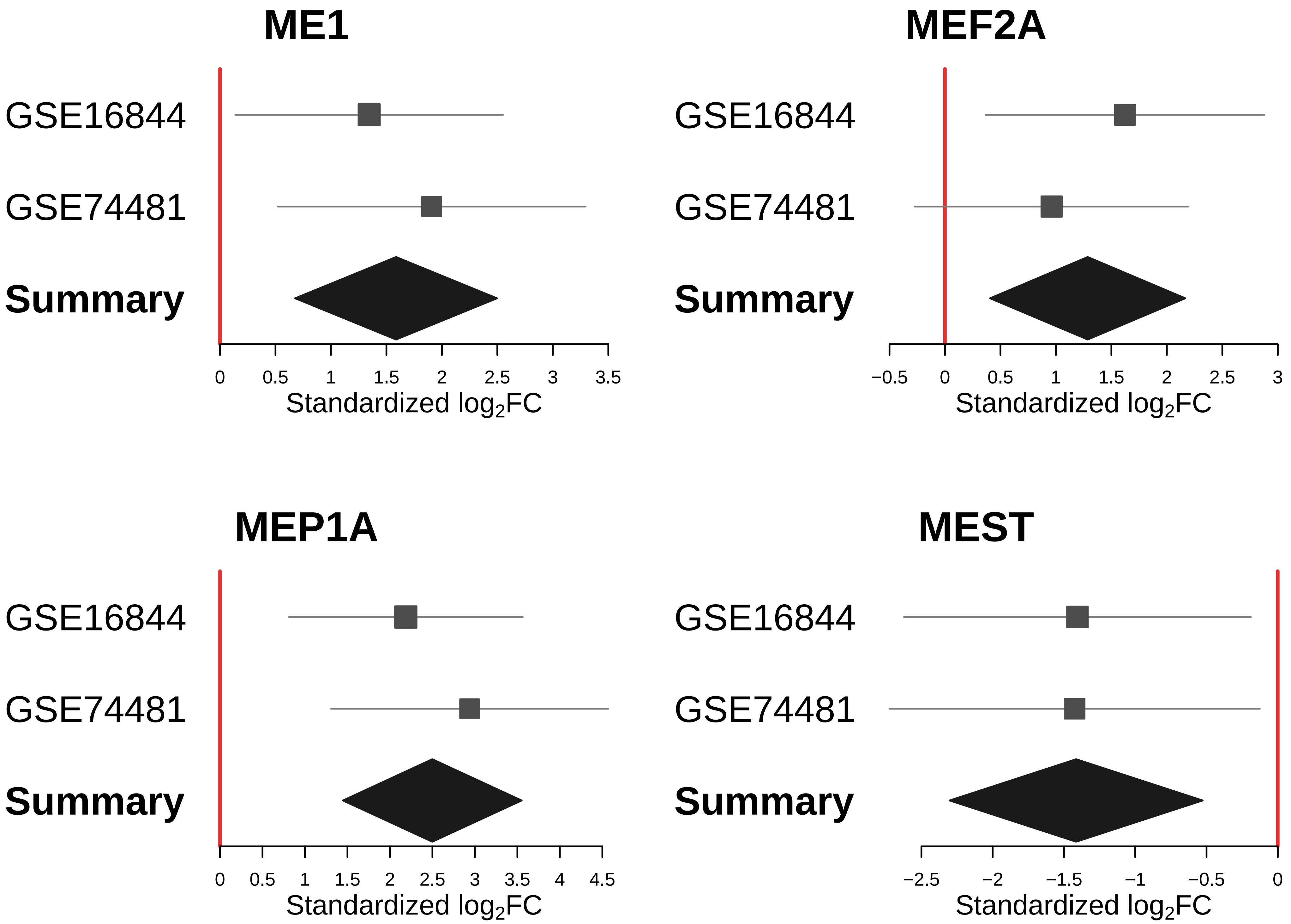
ME1
GSE16844
GSE74481
Summary
0
0.5
1
1.5
2
2.5
3
3.5
Standardized log2FC
MEF2A
GSE16844
GSE74481
Summary
−0.5
0
0.5
1
1.5
2
2.5
3
Standardized log2FC
MEP1A
GSE16844
GSE74481
Summary
0
0.5
1
1.5
2
2.5
3
3.5
4
4.5
Standardized log2FC
MEST
GSE16844
GSE74481
Summary
−2.5
−2
−1.5
−1
−0.5
0
Standardized log2FC

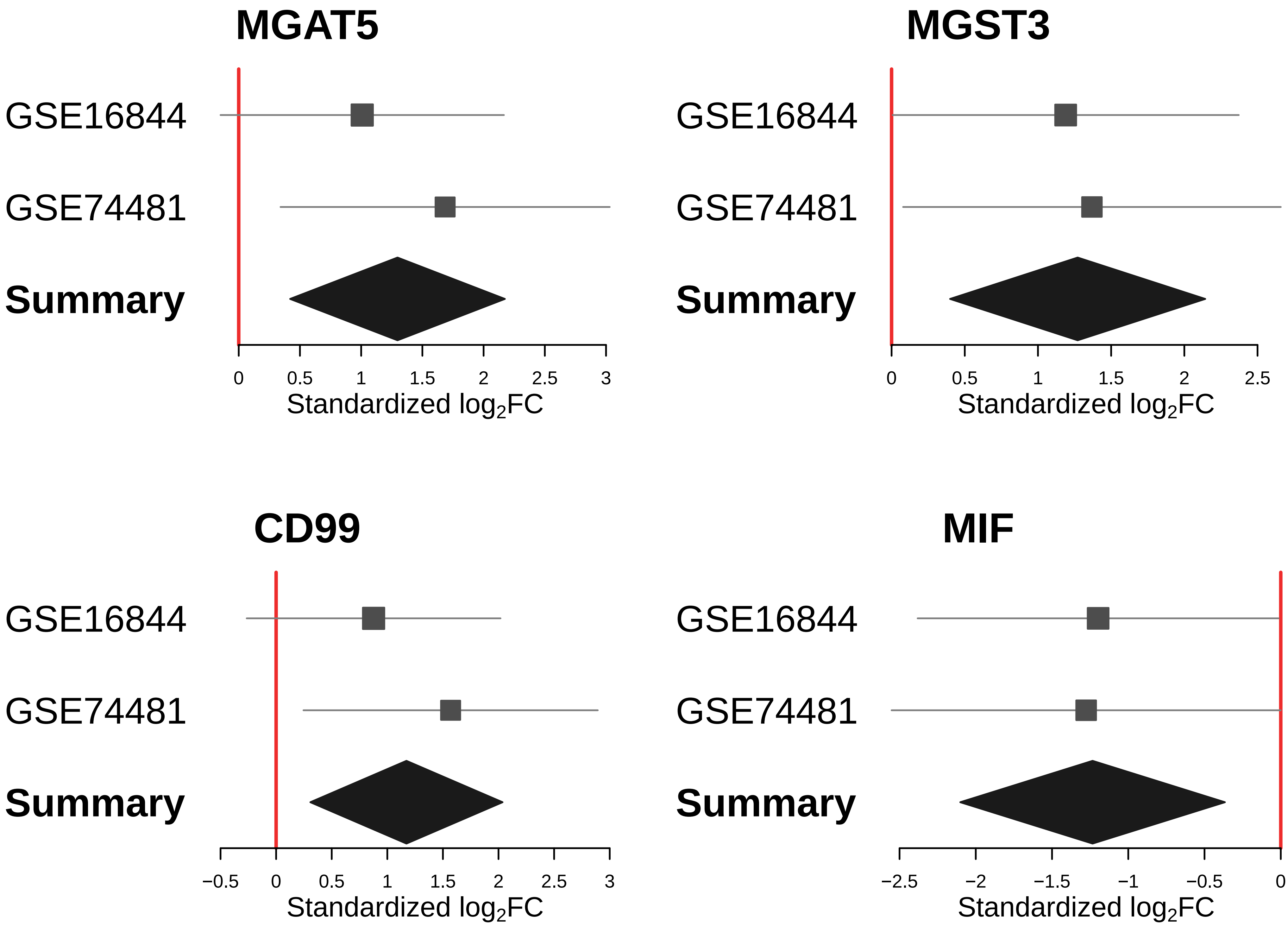
MGAT5
GSE16844
GSE74481
Summary
0
0.5
1
1.5
2
2.5
3
Standardized log2FC
MGST3
GSE16844
GSE74481
Summary
0
0.5
1
1.5
2
2.5
Standardized log2FC
CD99
GSE16844
GSE74481
Summary
−0.5
0
0.5
1
1.5
2
2.5
3
Standardized log2FC
MIF
GSE16844
GSE74481
Summary
−2.5
−2
−1.5
−1
−0.5
0
Standardized log2FC

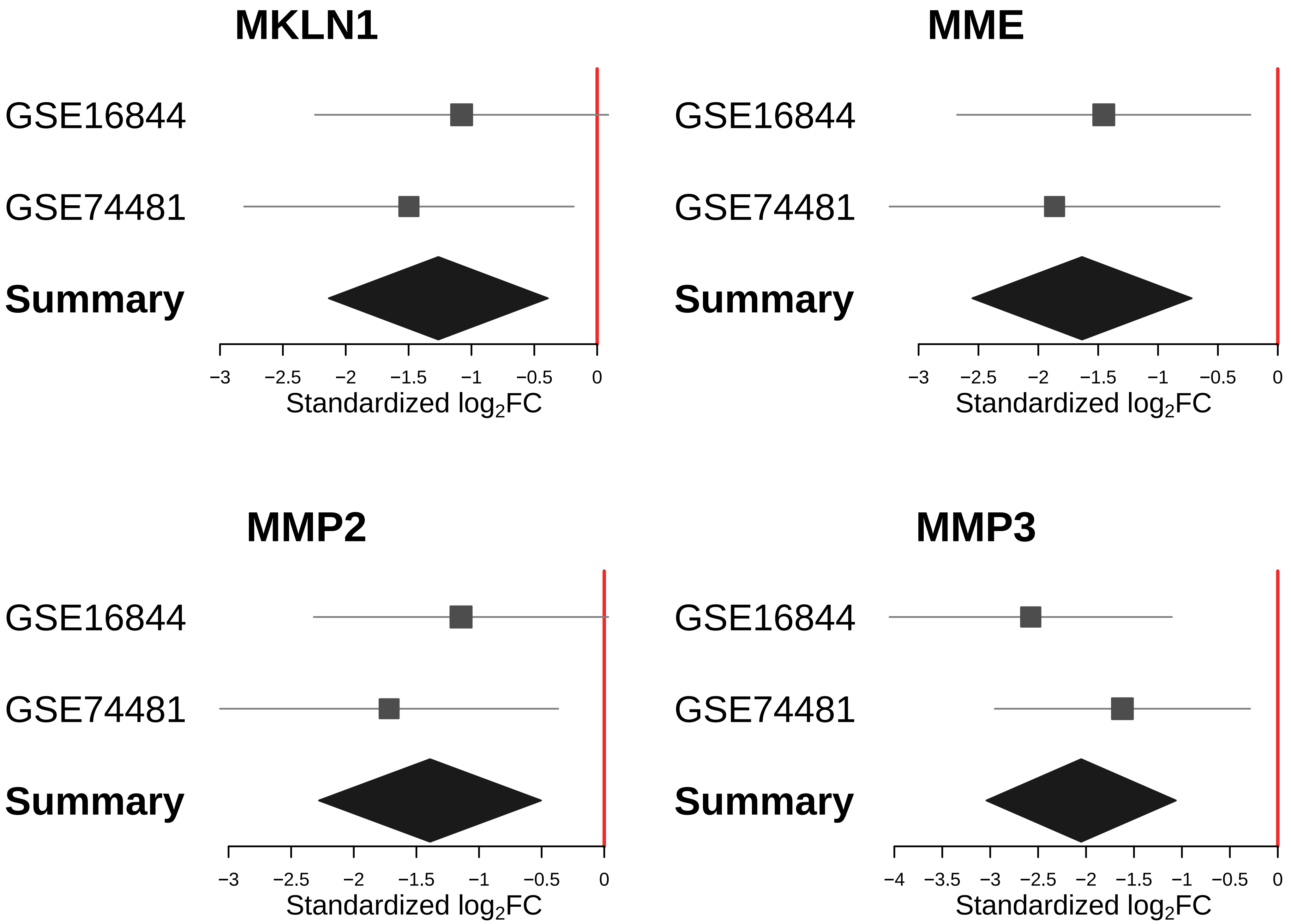

MKLN1
GSE16844
GSE74481
Summary
−3
−2.5
−2
−1.5
−1
−0.5
0
Standardized log2FC
MME
GSE16844
GSE74481
Summary
−3
−2.5
−2
−1.5
−1
−0.5
0
Standardized log2FC
MMP2
GSE16844
GSE74481
Summary
−3
−2.5
−2
−1.5
−1
−0.5
0
Standardized log2FC
MMP3
GSE16844
GSE74481
Summary
−4
−3.5
−3
−2.5
−2
−1.5
−1
−0.5
0
Standardized log2FC

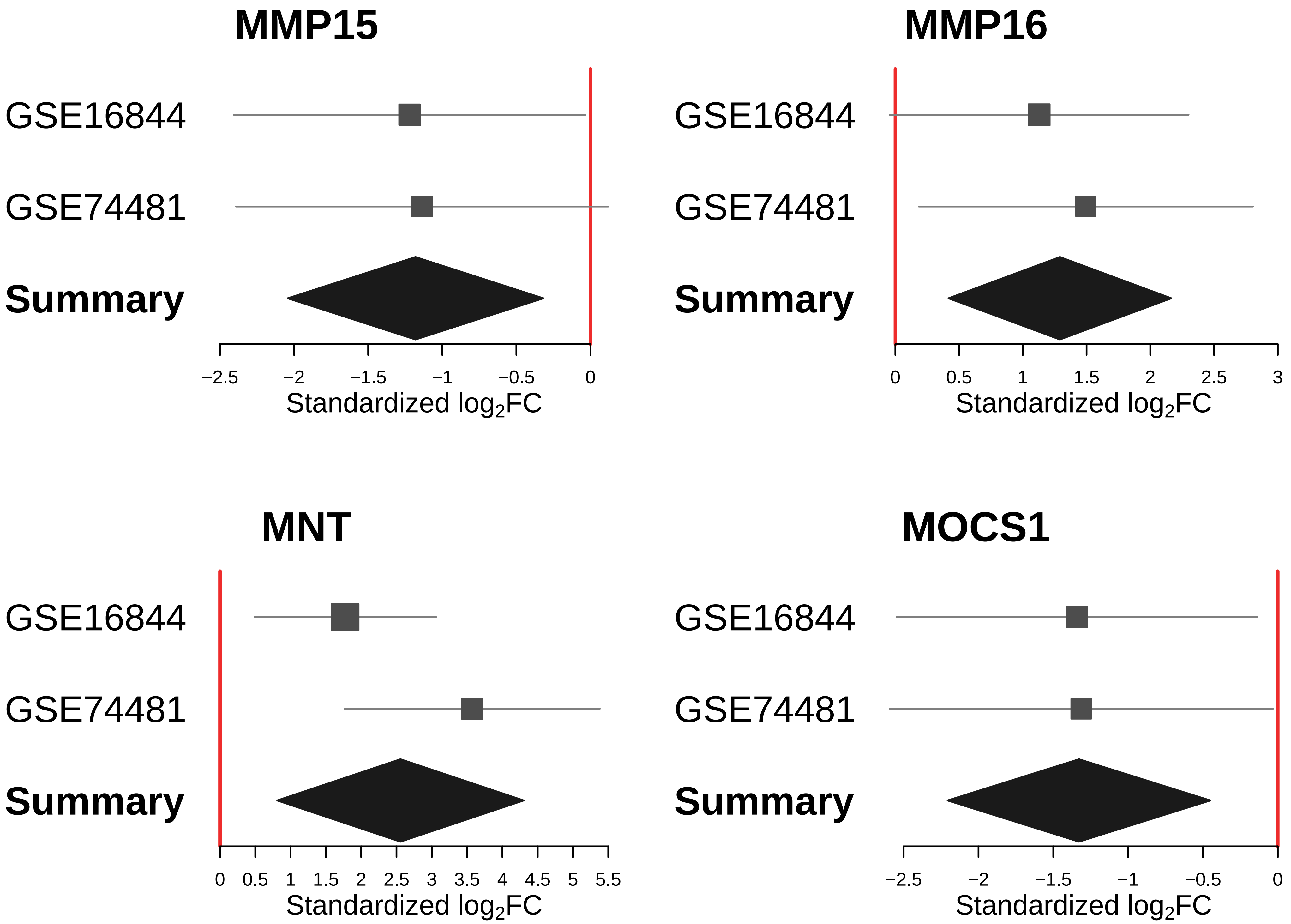
MMP15
GSE16844
GSE74481
Summary
−2.5 −2 −1.5 −1 −0.5 0
Standardized log2FC
MMP16
GSE16844
GSE74481
Summary
0 0.5 1 1.5 2 2.5 3
Standardized log2FC
MNT
GSE16844
GSE74481
Summary
0 0.5 1 1.5 2 2.5 3 3.5 4 4.5 5 5.5
Standardized log2FC
MOCS1
GSE16844
GSE74481
Summary
−2.5 −2 −1.5 −1 −0.5 0
Standardized log2FC

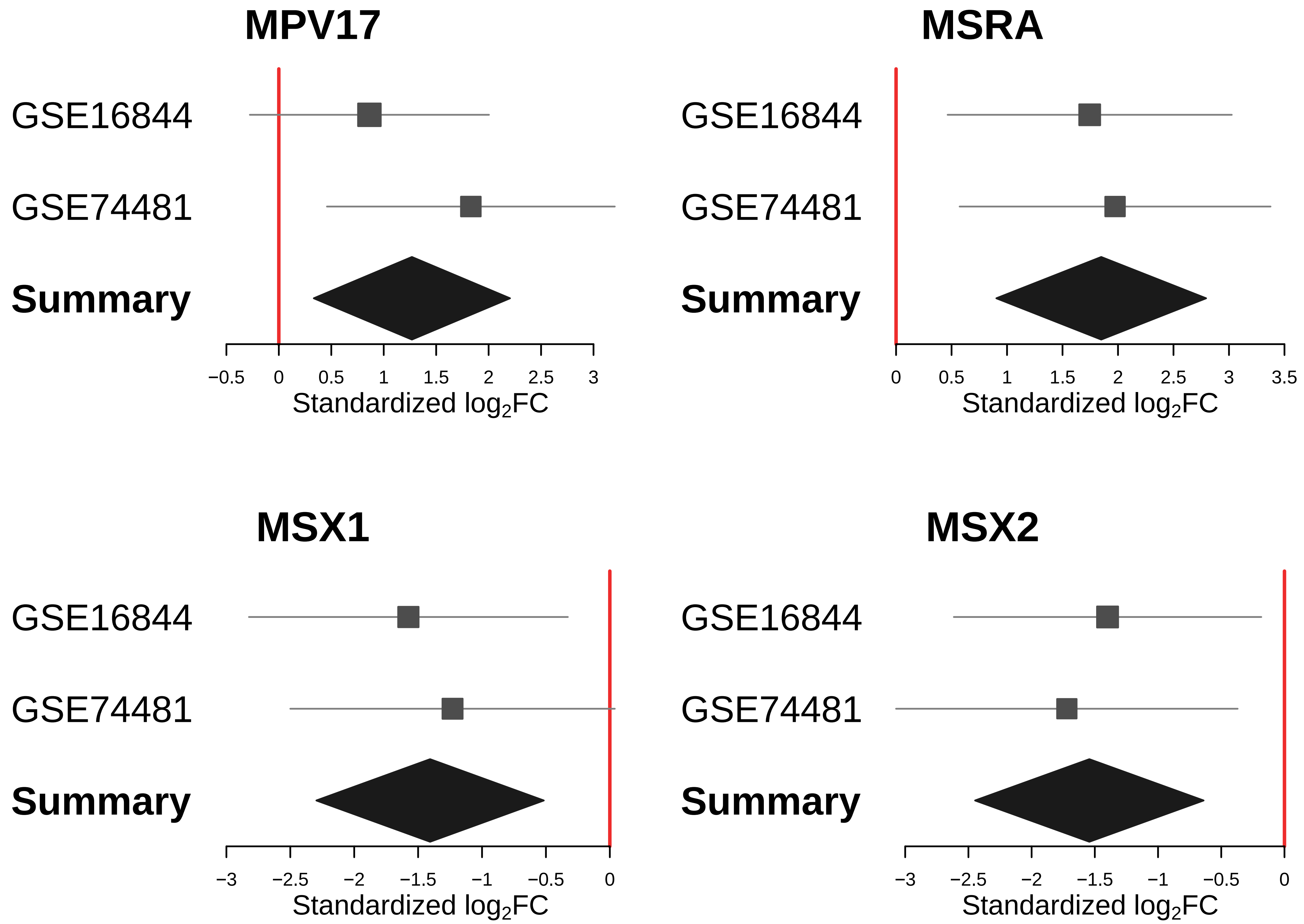
MPV17
GSE16844
GSE74481
Summary
−0.5 0 0.5 1 1.5 2 2.5 3
Standardized log2FC
MSRA
GSE16844
GSE74481
Summary
0 0.5 1 1.5 2 2.5 3 3.5
Standardized log2FC
MSX1
GSE16844
GSE74481
Summary
−3 −2.5 −2 −1.5 −1 −0.5 0
Standardized log2FC
MSX2
GSE16844
GSE74481
Summary
−3 −2.5 −2 −1.5 −1 −0.5 0
Standardized log2FC

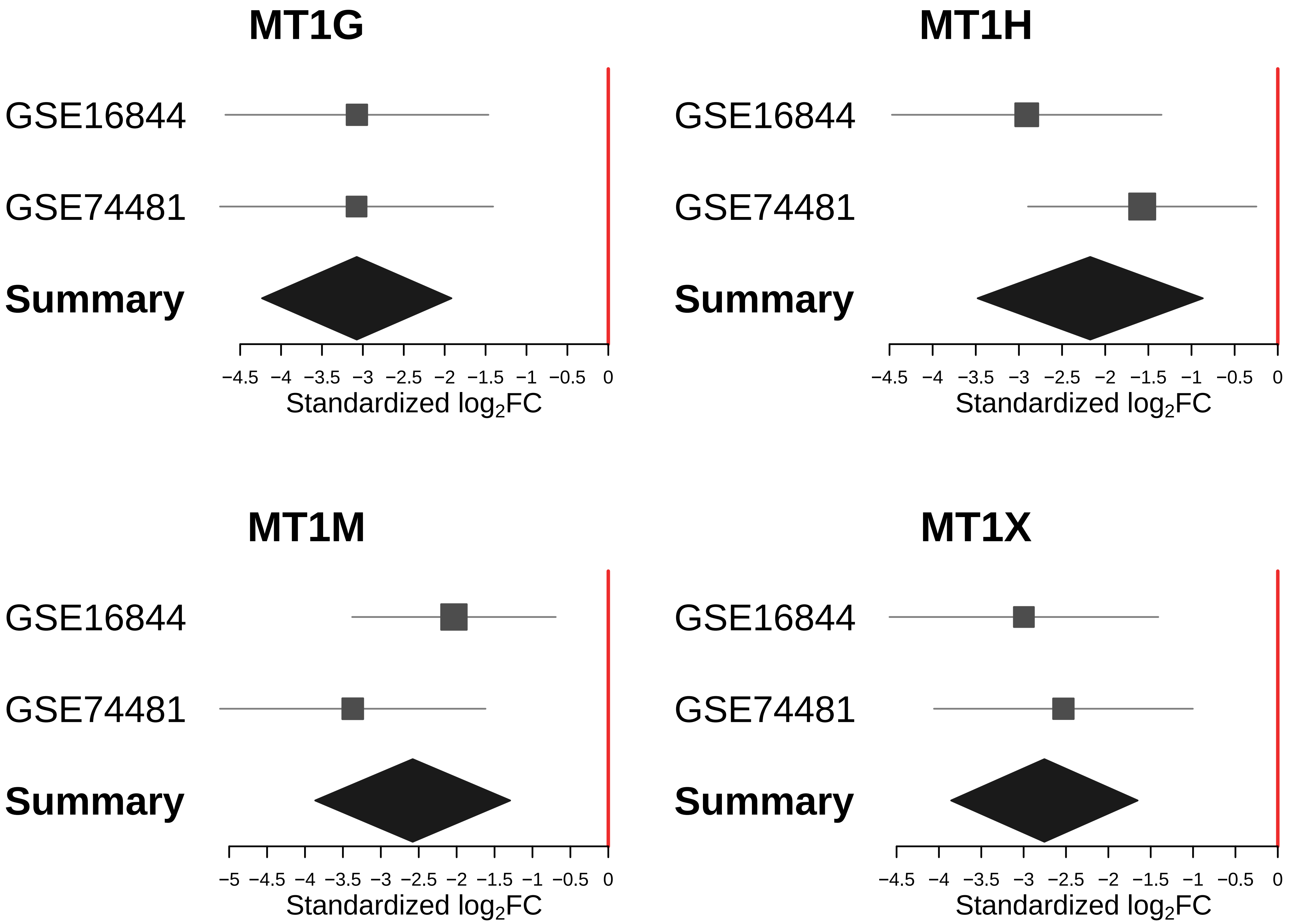
MT1G
GSE16844
GSE74481
Summary
−4.5 −4 −3.5 −3 −2.5 −2 −1.5 −1 −0.5 0
Standardized log2FC
MT1H
GSE16844
GSE74481
Summary
−4.5 −4 −3.5 −3 −2.5 −2 −1.5 −1 −0.5 0
Standardized log2FC
MT1M
GSE16844
GSE74481
Summary
−5 −4.5 −4 −3.5 −3 −2.5 −2 −1.5 −1 −0.5 0
Standardized log2FC
MT1X
GSE16844
GSE74481
Summary
−4.5 −4 −3.5 −3 −2.5 −2 −1.5 −1 −0.5 0
Standardized log2FC

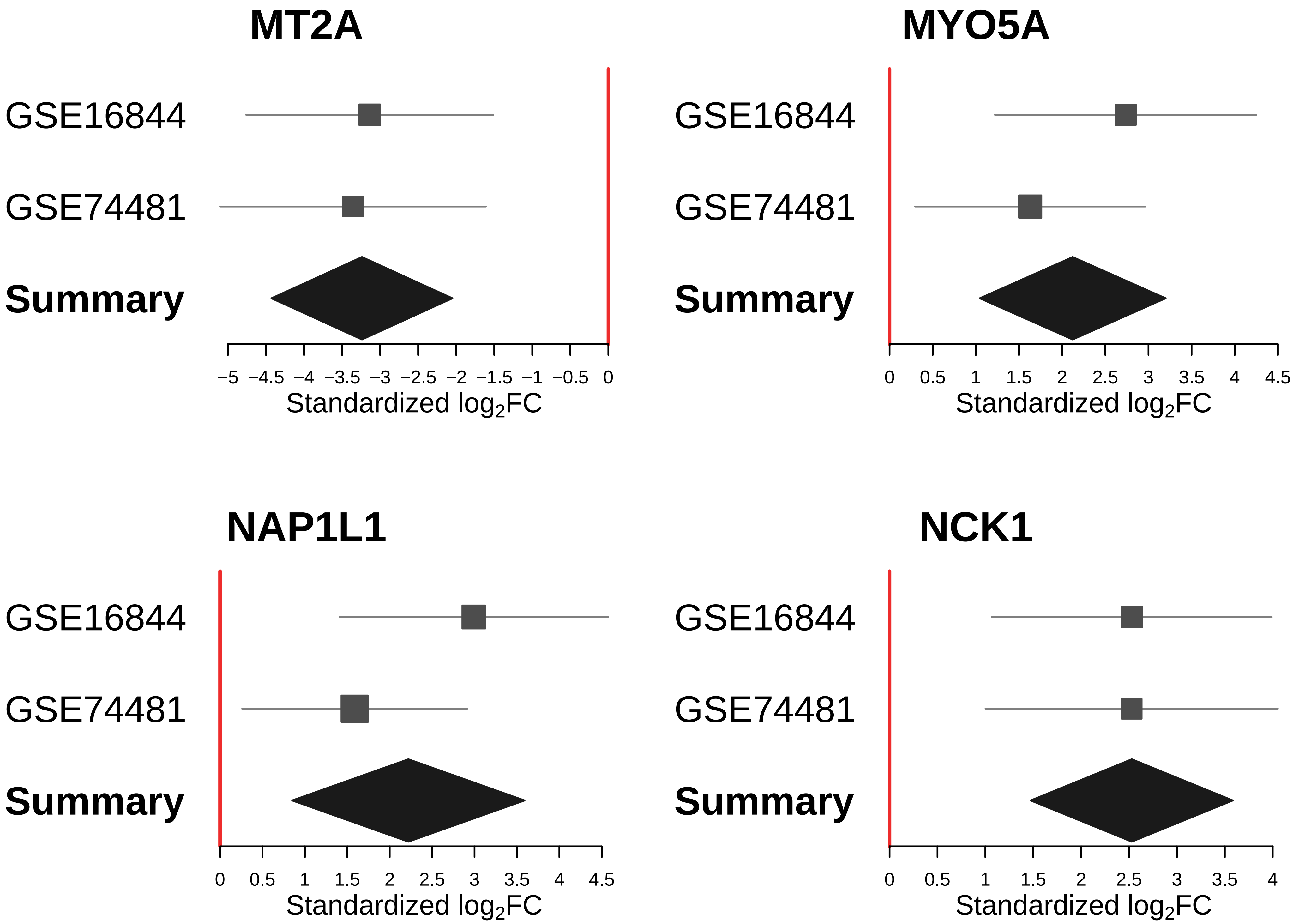
MT2A
GSE16844
GSE74481
Summary
−5 −4.5 −4 −3.5 −3 −2.5 −2 −1.5 −1 −0.5 0
Standardized $log_2FC$
MYO5A
GSE16844
GSE74481
Summary
0 0.5 1 1.5 2 2.5 3 3.5 4 4.5
Standardized $log_2FC$
NAP1L1
GSE16844
GSE74481
Summary
0 0.5 1 1.5 2 2.5 3 3.5 4 4.5
Standardized $log_2FC$
NCK1
GSE16844
GSE74481
Summary
0 0.5 1 1.5 2 2.5 3 3.5 4
Standardized $log_2FC$

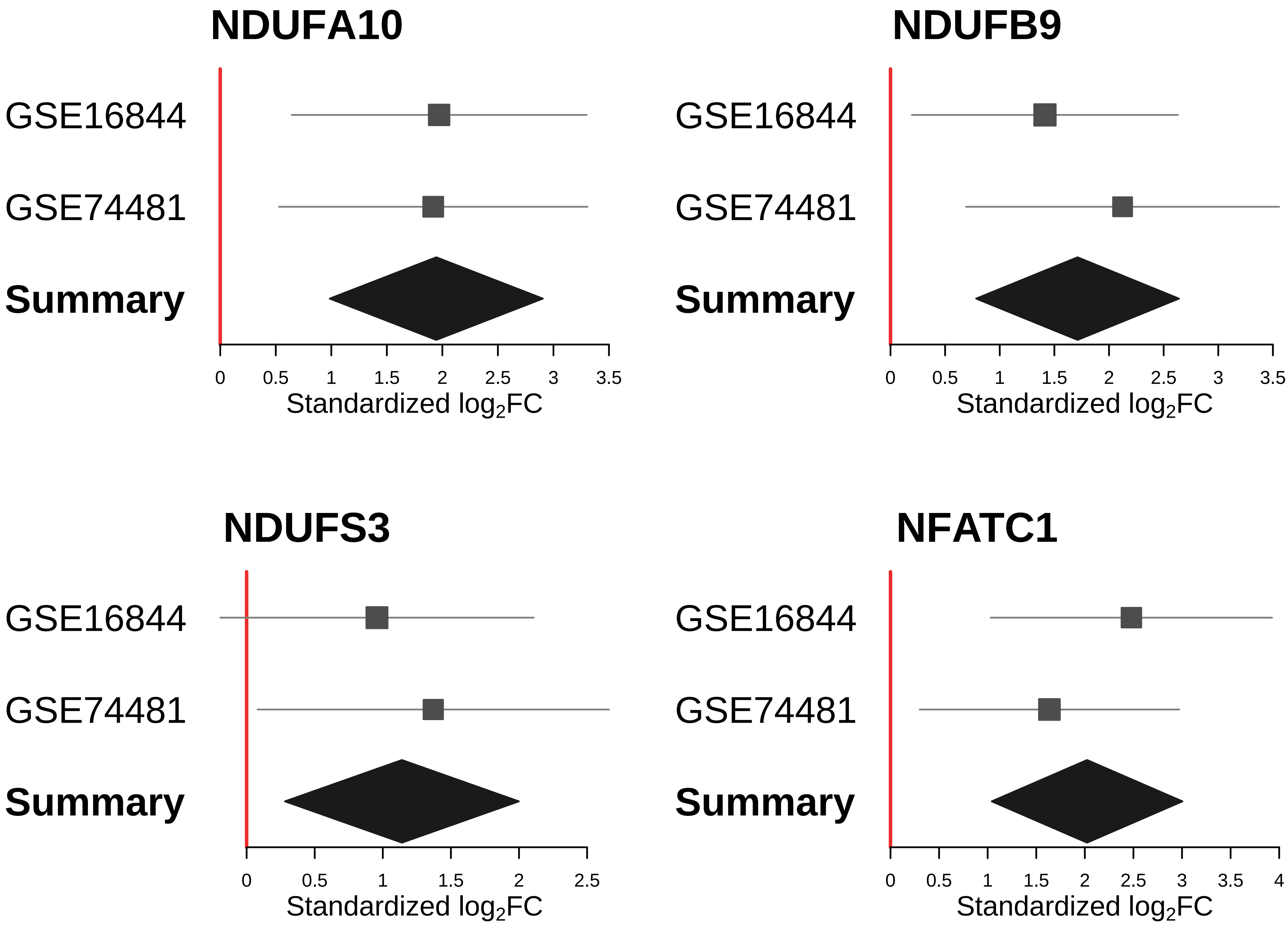

NDUFA10
GSE16844
GSE74481
Summary
0
0.5
1
1.5
2
2.5
3
3.5
Standardized log2FC
NDUFB9
GSE16844
GSE74481
Summary
0
0.5
1
1.5
2
2.5
3
3.5
Standardized log2FC
NDUFS3
GSE16844
GSE74481
Summary
0
0.5
1
1.5
2
2.5
Standardized log2FC
NFATC1
GSE16844
GSE74481
Summary
0
0.5
1
1.5
2
2.5
3
3.5
4
Standardized log2FC

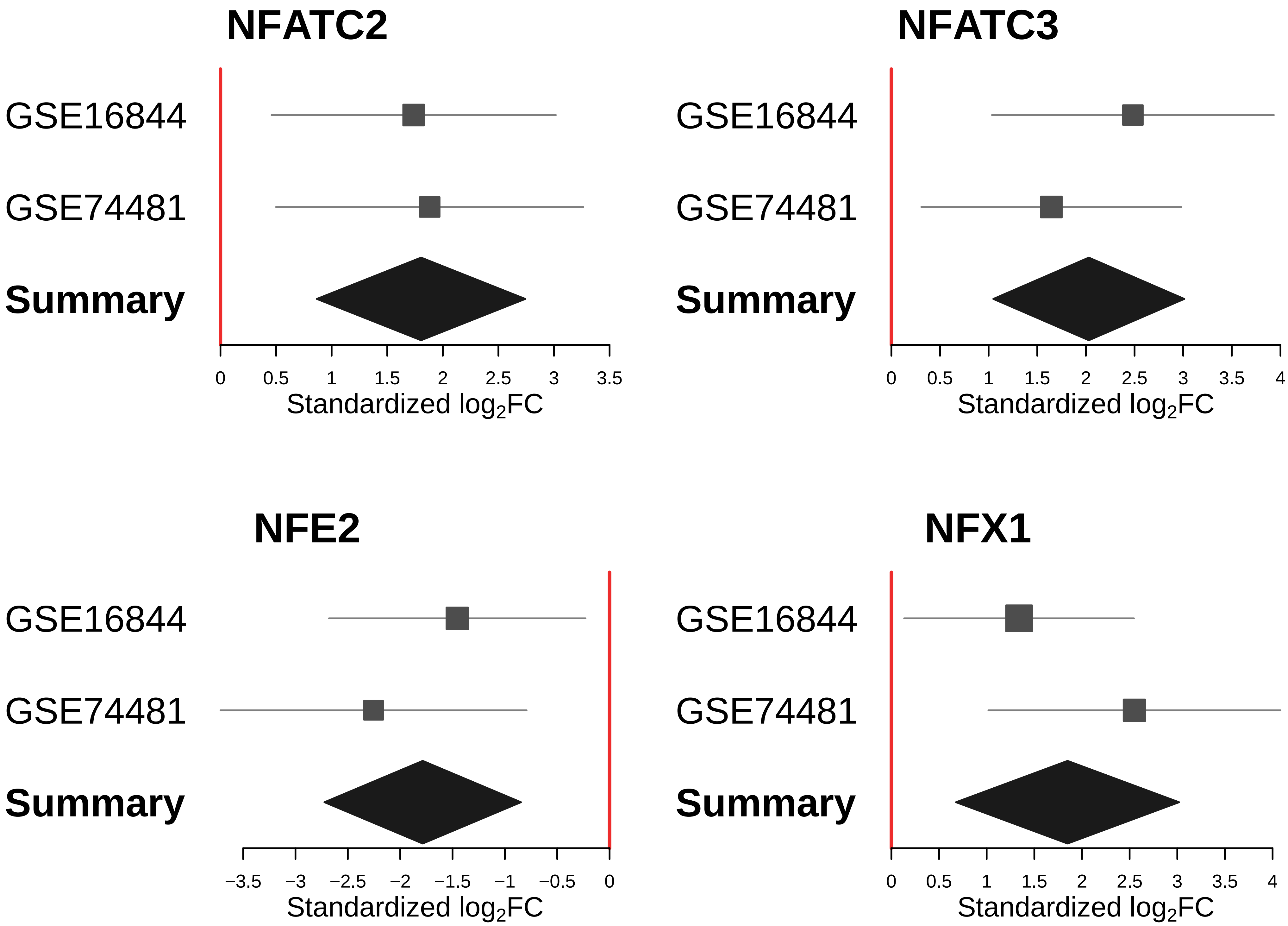
NFATC2
GSE16844
GSE74481
Summary
0 0.5 1 1.5 2 2.5 3 3.5
Standardized log2FC
NFATC3
GSE16844
GSE74481
Summary
0 0.5 1 1.5 2 2.5 3 3.5 4
Standardized log2FC
NFE2
GSE16844
GSE74481
Summary
−3.5 −3 −2.5 −2 −1.5 −1 −0.5 0
Standardized log2FC
NFX1
GSE16844
GSE74481
Summary
0 0.5 1 1.5 2 2.5 3 3.5 4
Standardized log2FC

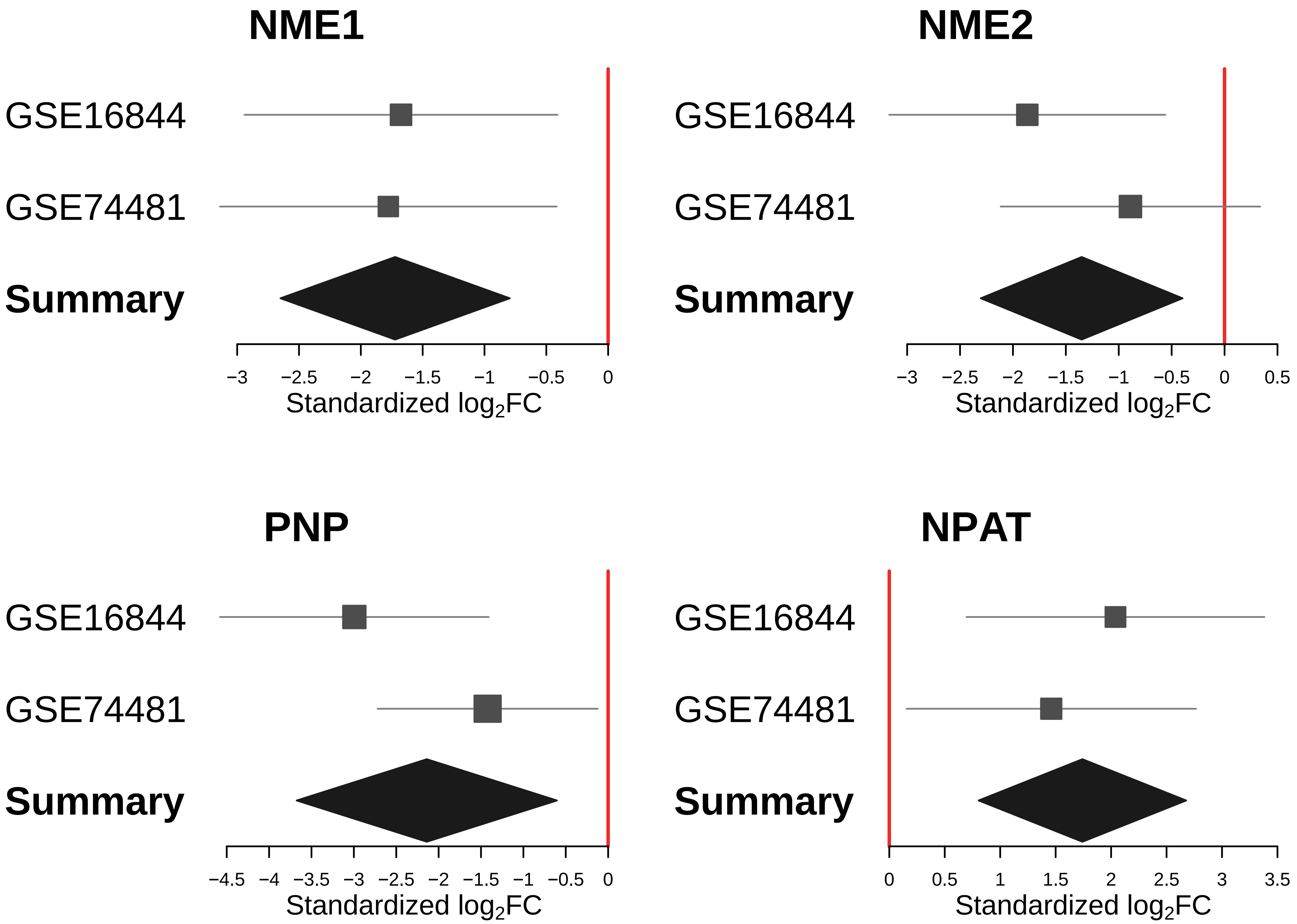
NME1
GSE16844
GSE74481
Summary
−3 −2.5 −2 −1.5 −1 −0.5 0
Standardized log2FC
NME2
GSE16844
GSE74481
Summary
−3 −2.5 −2 −1.5 −1 −0.5 0 0.5
Standardized log2FC
PNP
GSE16844
GSE74481
Summary
−4.5 −4 −3.5 −3 −2.5 −2 −1.5 −1 −0.5 0
Standardized log2FC
NPAT
GSE16844
GSE74481
Summary
0 0.5 1 1.5 2 2.5 3 3.5
Standardized log2FC

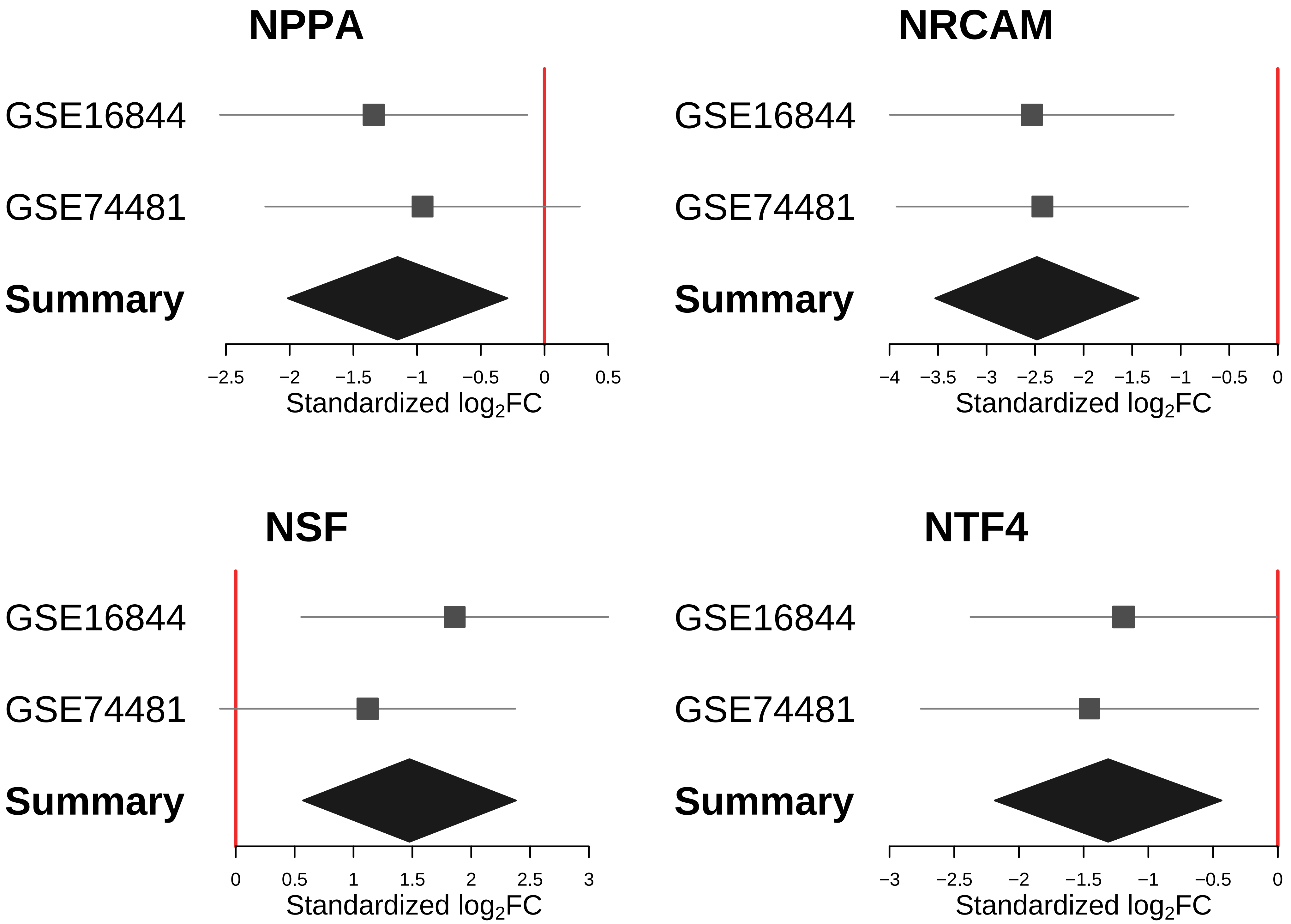
NPPA
GSE16844
GSE74481
Summary
−2.5 −2 −1.5 −1 −0.5 0 0.5
Standardized log2FC
NRCAM
GSE16844
GSE74481
Summary
−4 −3.5 −3 −2.5 −2 −1.5 −1 −0.5 0
Standardized log2FC
NSF
GSE16844
GSE74481
Summary
0 0.5 1 1.5 2 2.5 3
Standardized log2FC
NTF4
GSE16844
GSE74481
Summary
−3 −2.5 −2 −1.5 −1 −0.5 0
Standardized log2FC

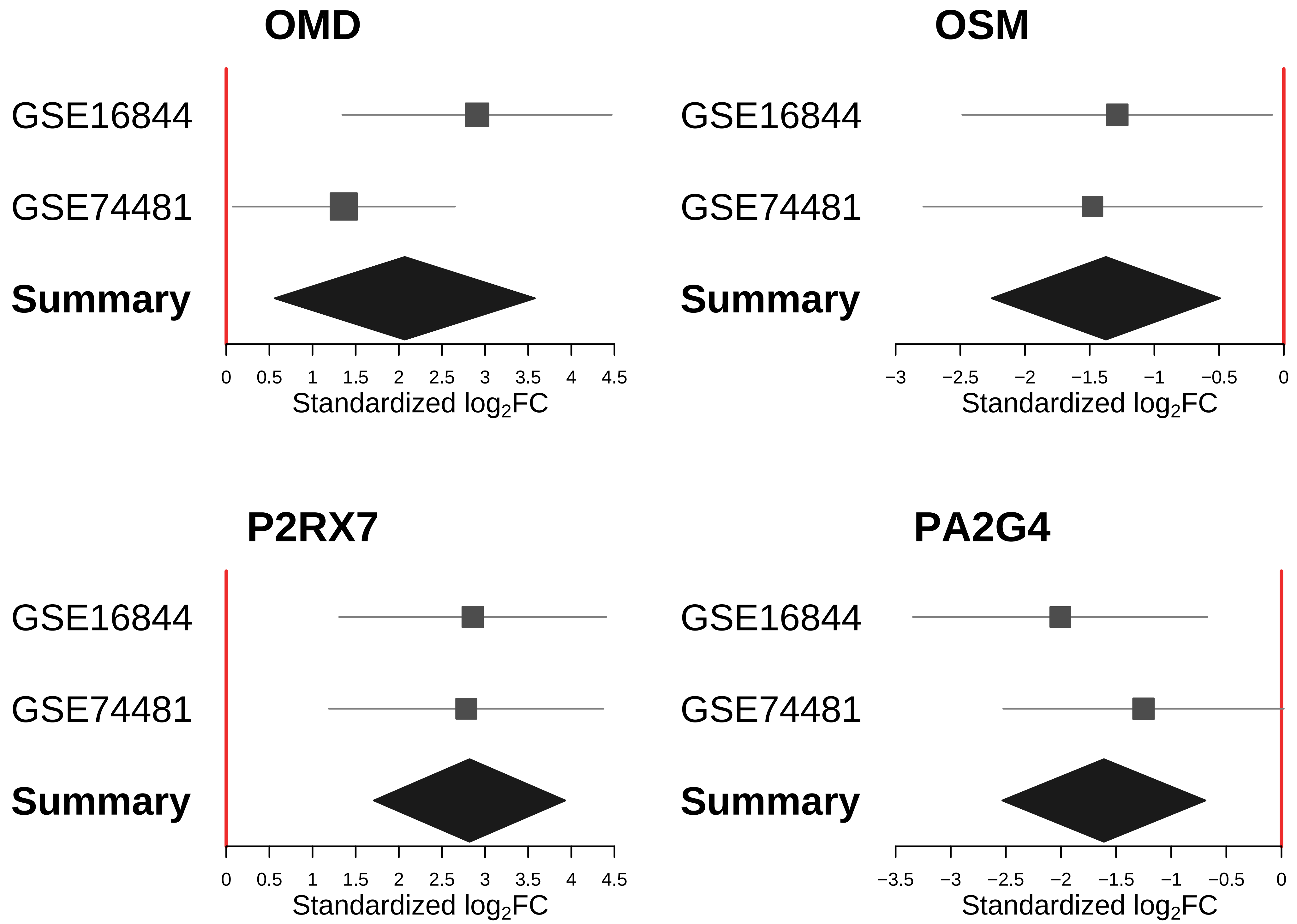
OMD
GSE16844
GSE74481
Summary
0 0.5 1 1.5 2 2.5 3 3.5 4 4.5
Standardized log2FC
OSM
GSE16844
GSE74481
Summary
−3 −2.5 −2 −1.5 −1 −0.5 0
Standardized log2FC
P2RX7
GSE16844
GSE74481
Summary
0 0.5 1 1.5 2 2.5 3 3.5 4 4.5
Standardized log2FC
PA2G4
GSE16844
GSE74481
Summary
−3.5 −3 −2.5 −2 −1.5 −1 −0.5 0
Standardized log2FC

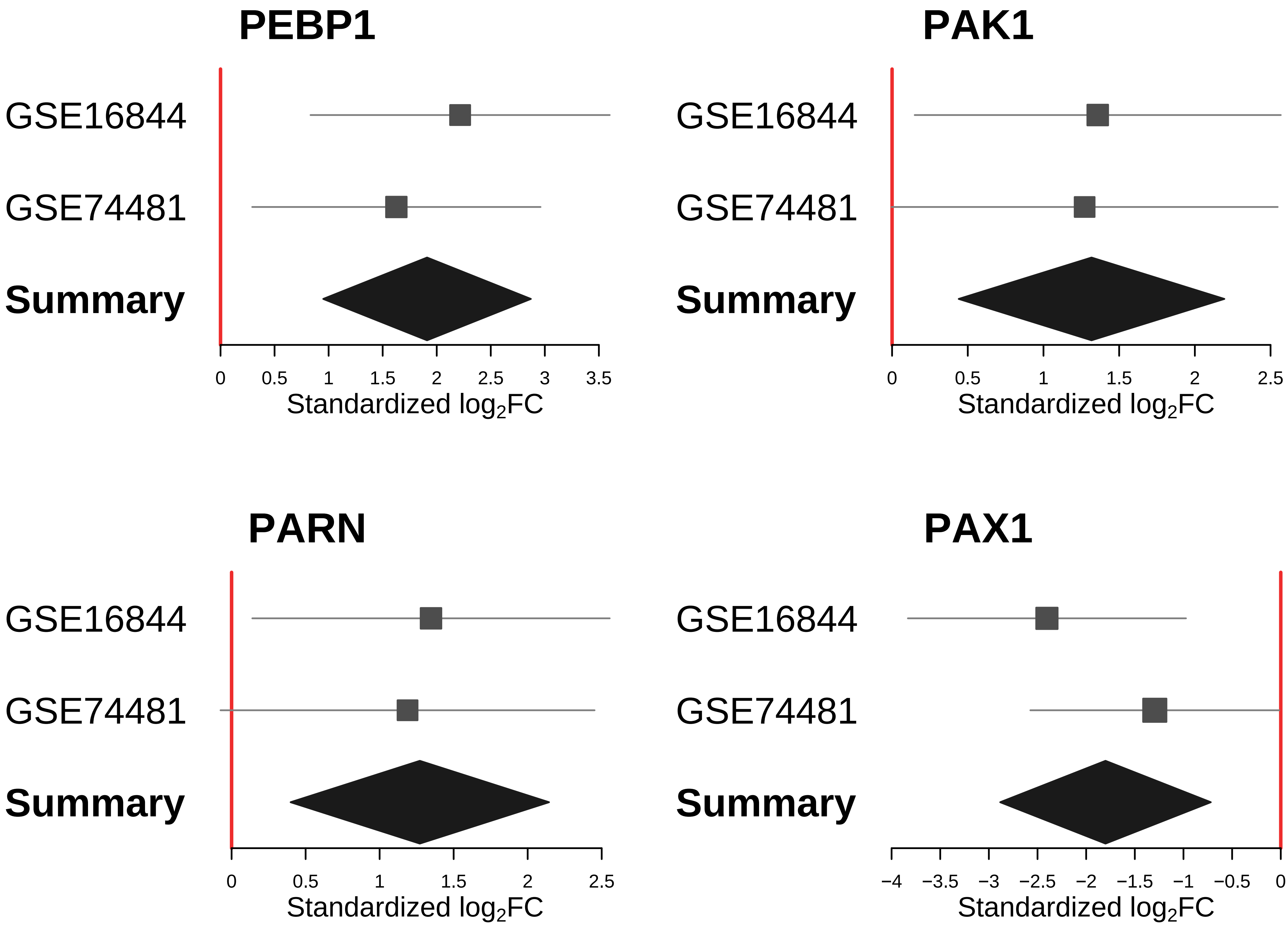
PEBP1
GSE16844
GSE74481
Summary
0 0.5 1 1.5 2 2.5 3 3.5
Standardized log2FC
PAK1
GSE16844
GSE74481
Summary
0 0.5 1 1.5 2 2.5
Standardized log2FC
PARN
GSE16844
GSE74481
Summary
0 0.5 1 1.5 2 2.5
Standardized log2FC
PAX1
GSE16844
GSE74481
Summary
−4 −3.5 −3 −2.5 −2 −1.5 −1 −0.5 0
Standardized log2FC

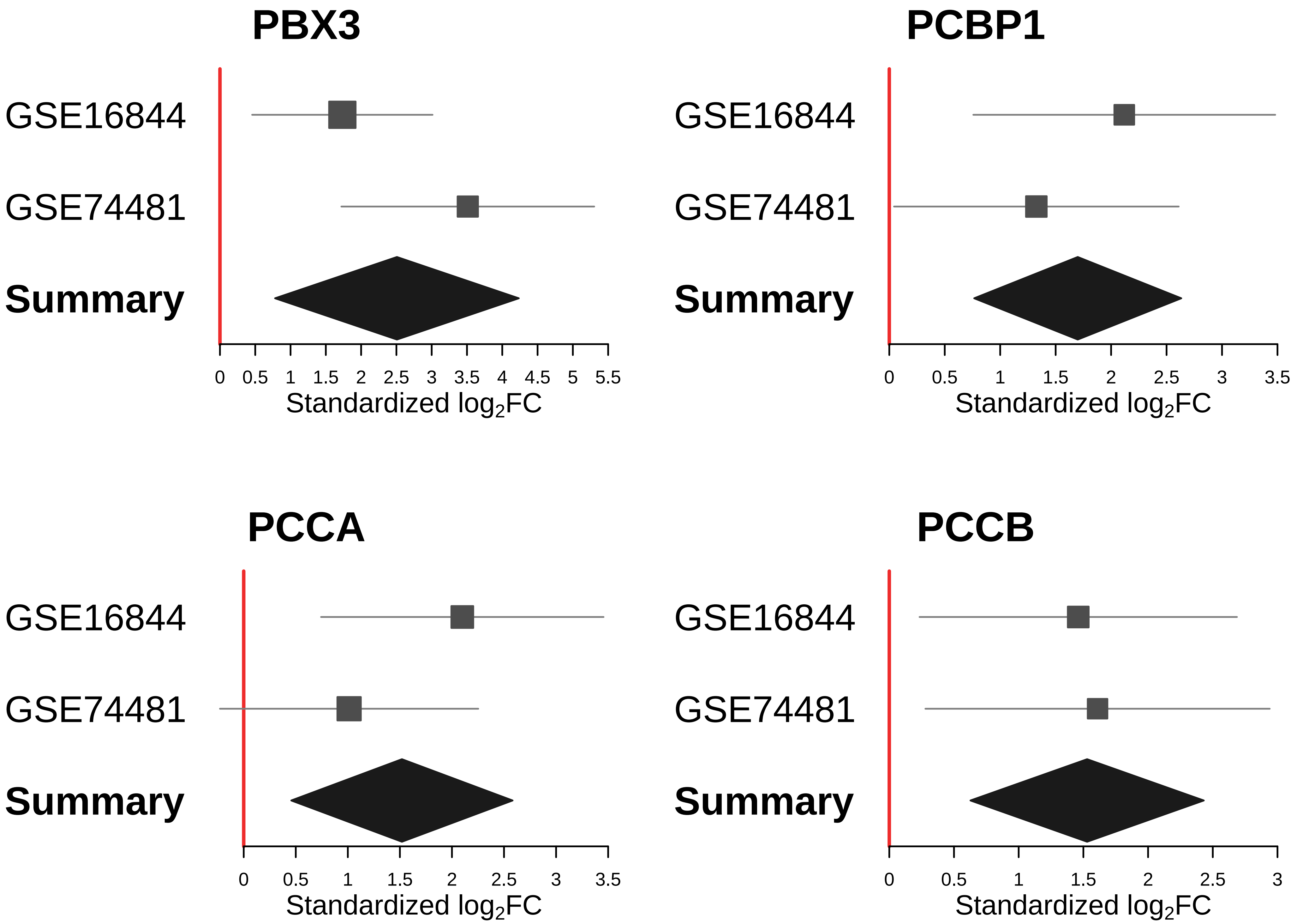
PBX3
GSE16844
GSE74481
Summary
0 0.5 1 1.5 2 2.5 3 3.5 4 4.5 5 5.5
Standardized $log_2FC$
PCBP1
GSE16844
GSE74481
Summary
0 0.5 1 1.5 2 2.5 3 3.5
Standardized $log_2FC$
PCCA
GSE16844
GSE74481
Summary
0 0.5 1 1.5 2 2.5 3 3.5
Standardized $log_2FC$
PCCB
GSE16844
GSE74481
Summary
0 0.5 1 1.5 2 2.5 3
Standardized $log_2FC$

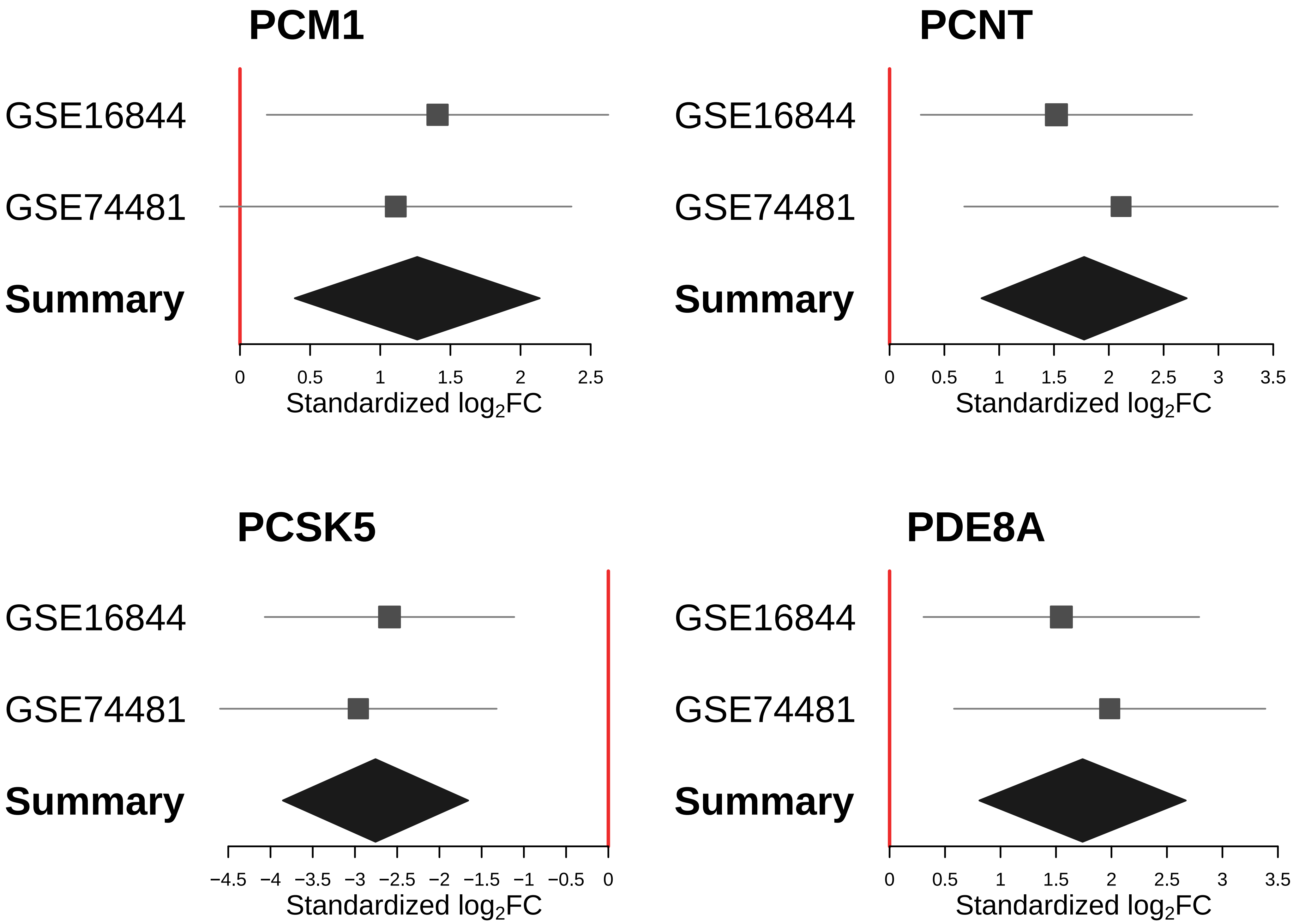
PCM1
GSE16844
GSE74481
Summary
0
0.5
1
1.5
2
2.5
Standardized log2FC
PCNT
GSE16844
GSE74481
Summary
0
0.5
1
1.5
2
2.5
3
3.5
Standardized log2FC
PCSK5
GSE16844
GSE74481
Summary
−4.5
−4
−3.5
−3
−2.5
−2
−1.5
−1
−0.5
0
Standardized log2FC
PDE8A
GSE16844
GSE74481
Summary
0
0.5
1
1.5
2
2.5
3
3.5
Standardized log2FC

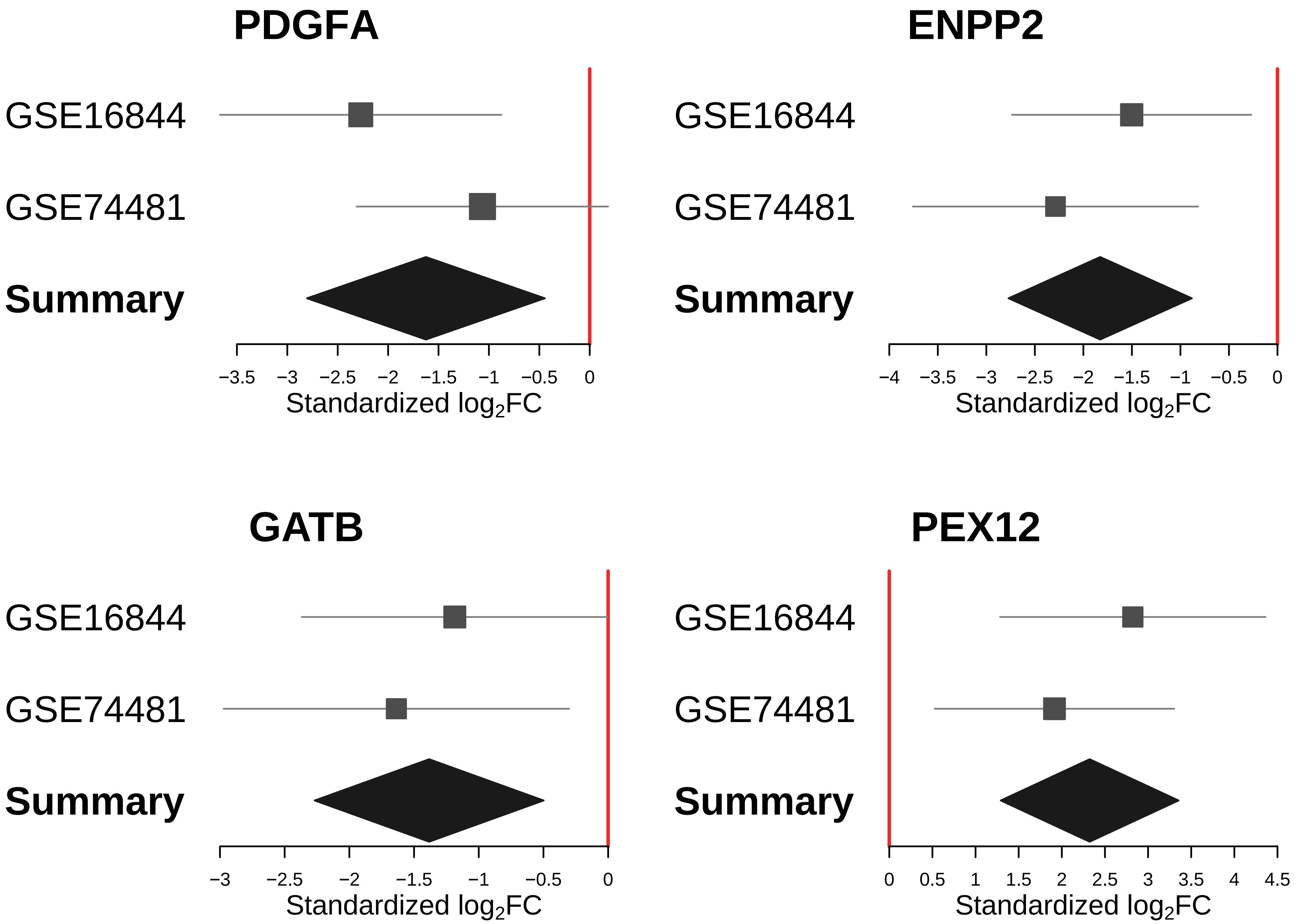
PDGFA
GSE16844
GSE74481
Summary
−3.5 −3 −2.5 −2 −1.5 −1 −0.5 0
Standardized log2FC
ENPP2
GSE16844
GSE74481
Summary
−4 −3.5 −3 −2.5 −2 −1.5 −1 −0.5 0
Standardized log2FC
GATB
GSE16844
GSE74481
Summary
−3 −2.5 −2 −1.5 −1 −0.5 0
Standardized log2FC
PEX12
GSE16844
GSE74481
Summary
0 0.5 1 1.5 2 2.5 3 3.5 4 4.5
Standardized log2FC

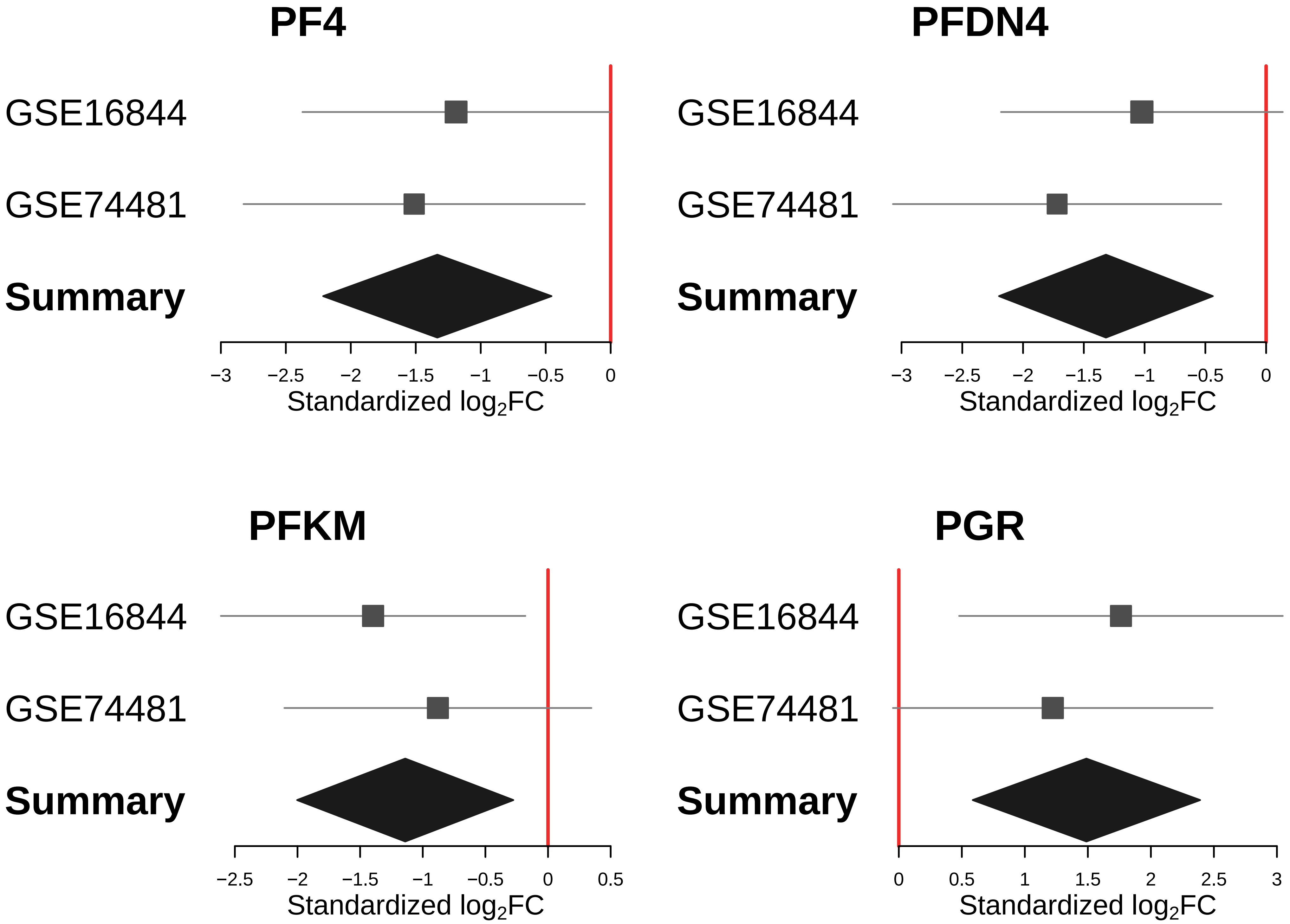
PF4
GSE16844
GSE74481
Summary
−3
−2.5
−2
−1.5
−1
−0.5
0
Standardized $log_2FC$
PFDN4
GSE16844
GSE74481
Summary
−3
−2.5
−2
−1.5
−1
−0.5
0
Standardized $log_2FC$
PFKM
GSE16844
GSE74481
Summary
−2.5
−2
−1.5
−1
−0.5
0
0.5
Standardized $log_2FC$
PGR
GSE16844
GSE74481
Summary
0
0.5
1
1.5
2
2.5
3
Standardized $log_2FC$

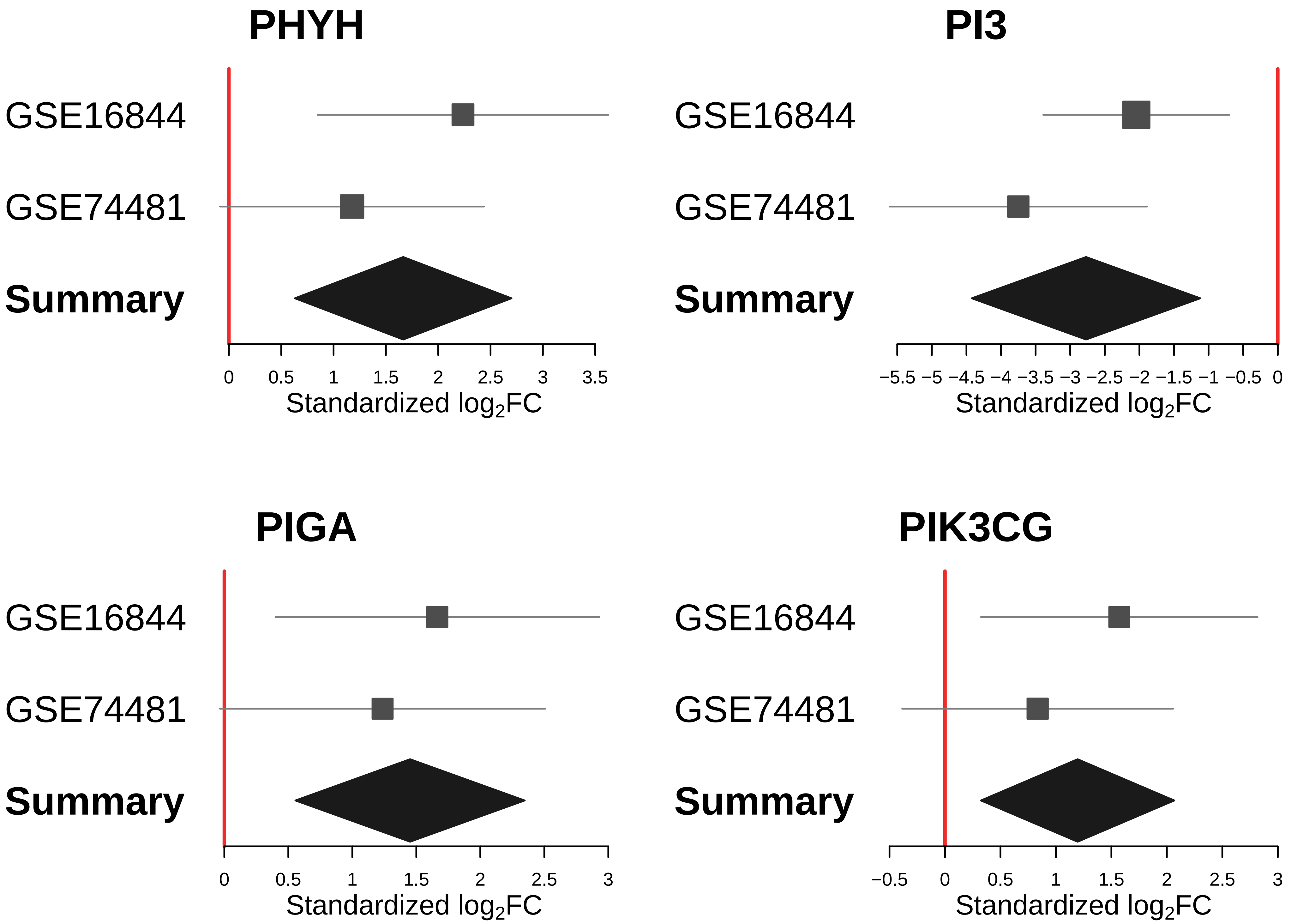
PHYH
GSE16844
GSE74481
Summary
0 0.5 1 1.5 2 2.5 3 3.5
Standardized log2FC
PI3
GSE16844
GSE74481
Summary
−5.5 −5 −4.5 −4 −3.5 −3 −2.5 −2 −1.5 −1 −0.5 0
Standardized log2FC
PIGA
GSE16844
GSE74481
Summary
0 0.5 1 1.5 2 2.5 3
Standardized log2FC
PIK3CG
GSE16844
GSE74481
Summary
−0.5 0 0.5 1 1.5 2 2.5 3
Standardized log2FC

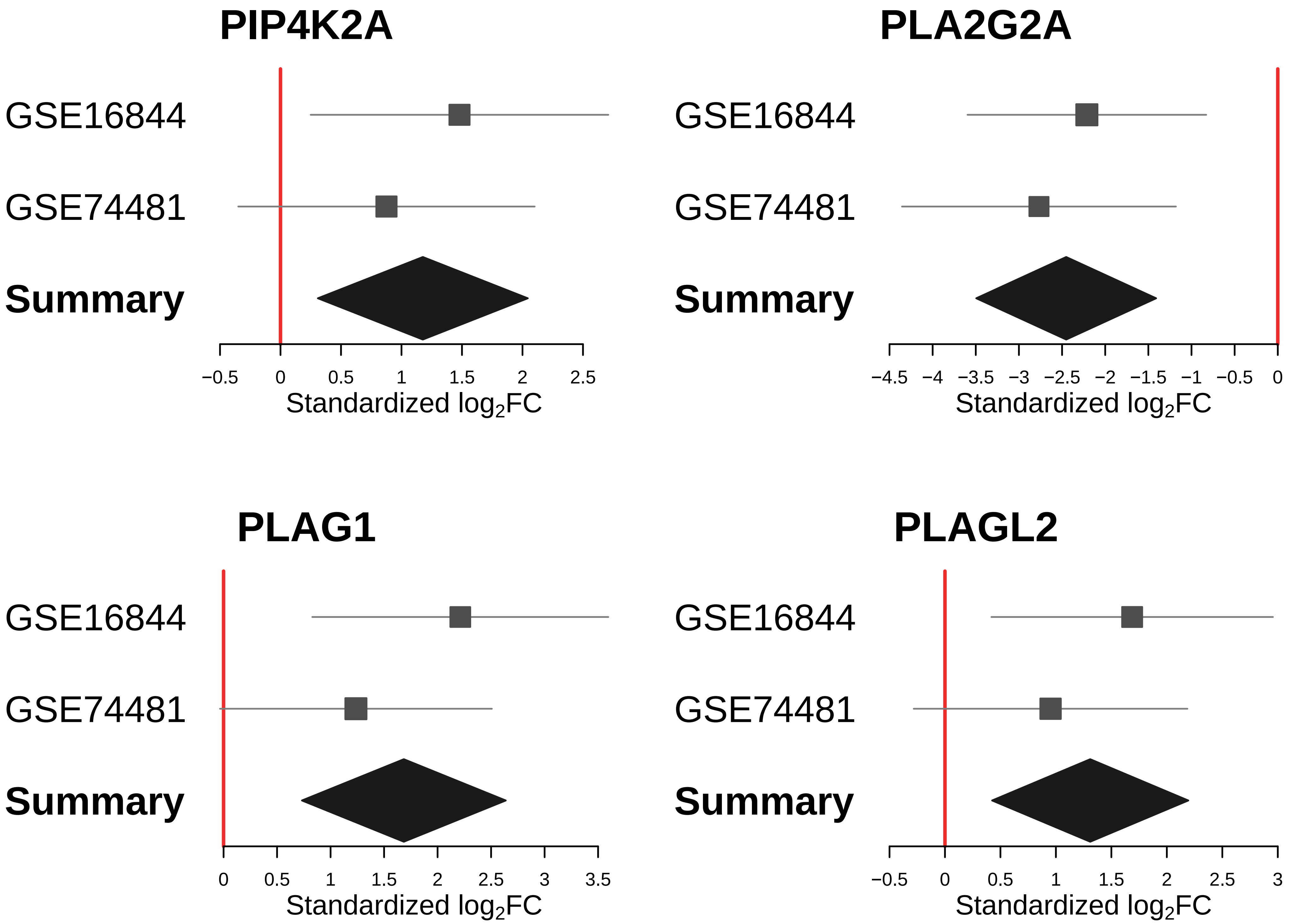
PIP4K2A
GSE16844
GSE74481
Summary
−0.5 0 0.5 1 1.5 2 2.5
Standardized log2FC
PLA2G2A
GSE16844
GSE74481
Summary
−4.5 −4 −3.5 −3 −2.5 −2 −1.5 −1 −0.5 0
Standardized log2FC
PLAG1
GSE16844
GSE74481
Summary
0 0.5 1 1.5 2 2.5 3 3.5
Standardized log2FC
PLAGL2
GSE16844
GSE74481
Summary
−0.5 0 0.5 1 1.5 2 2.5 3
Standardized log2FC

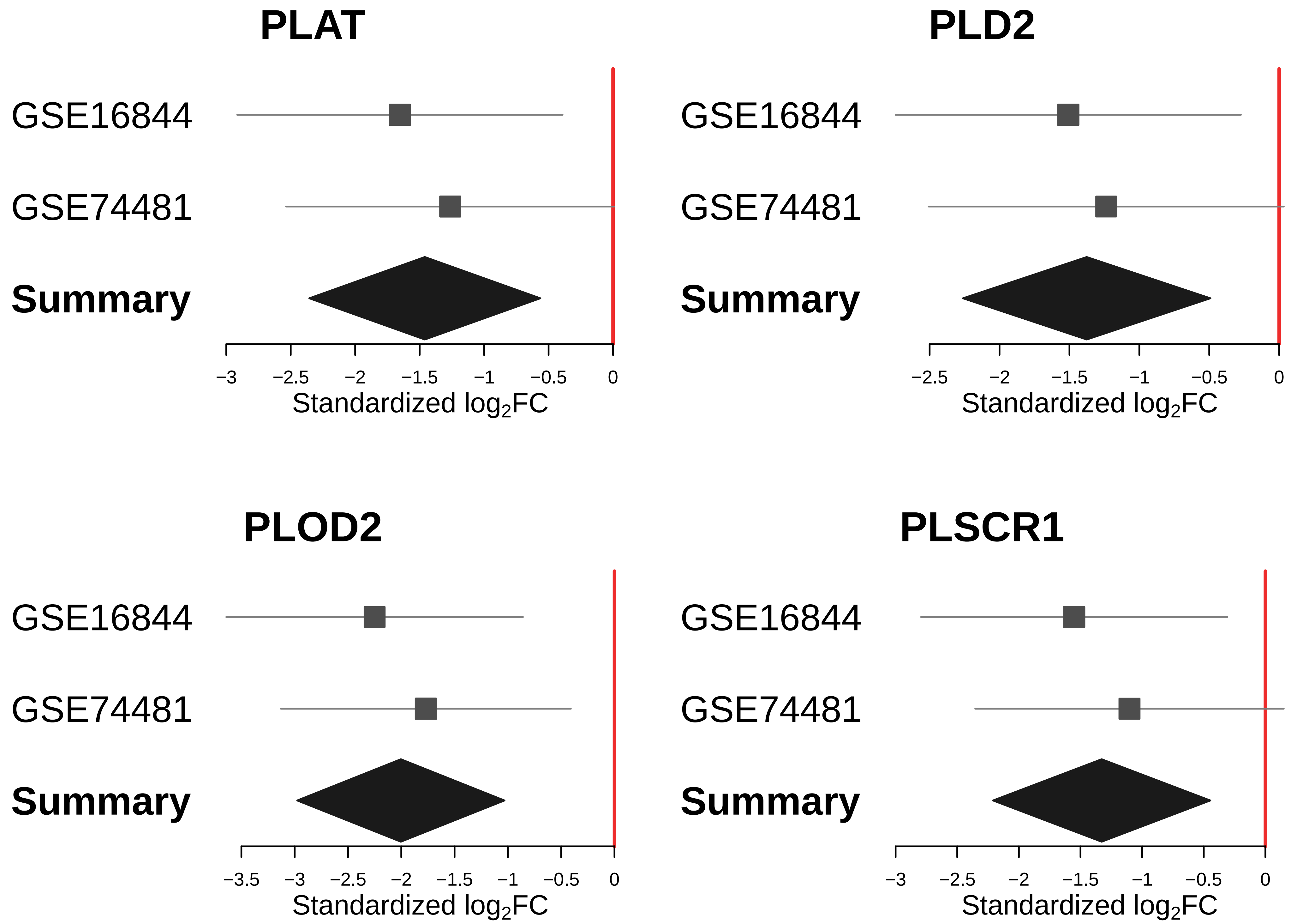

PLAT
GSE16844
GSE74481
Summary
−3 −2.5 −2 −1.5 −1 −0.5 0
Standardized log2FC
PLD2
GSE16844
GSE74481
Summary
−2.5 −2 −1.5 −1 −0.5 0
Standardized log2FC
PLOD2
GSE16844
GSE74481
Summary
−3.5 −3 −2.5 −2 −1.5 −1 −0.5 0
Standardized log2FC
PLSCR1
GSE16844
GSE74481
Summary
−3 −2.5 −2 −1.5 −1 −0.5 0
Standardized log2FC

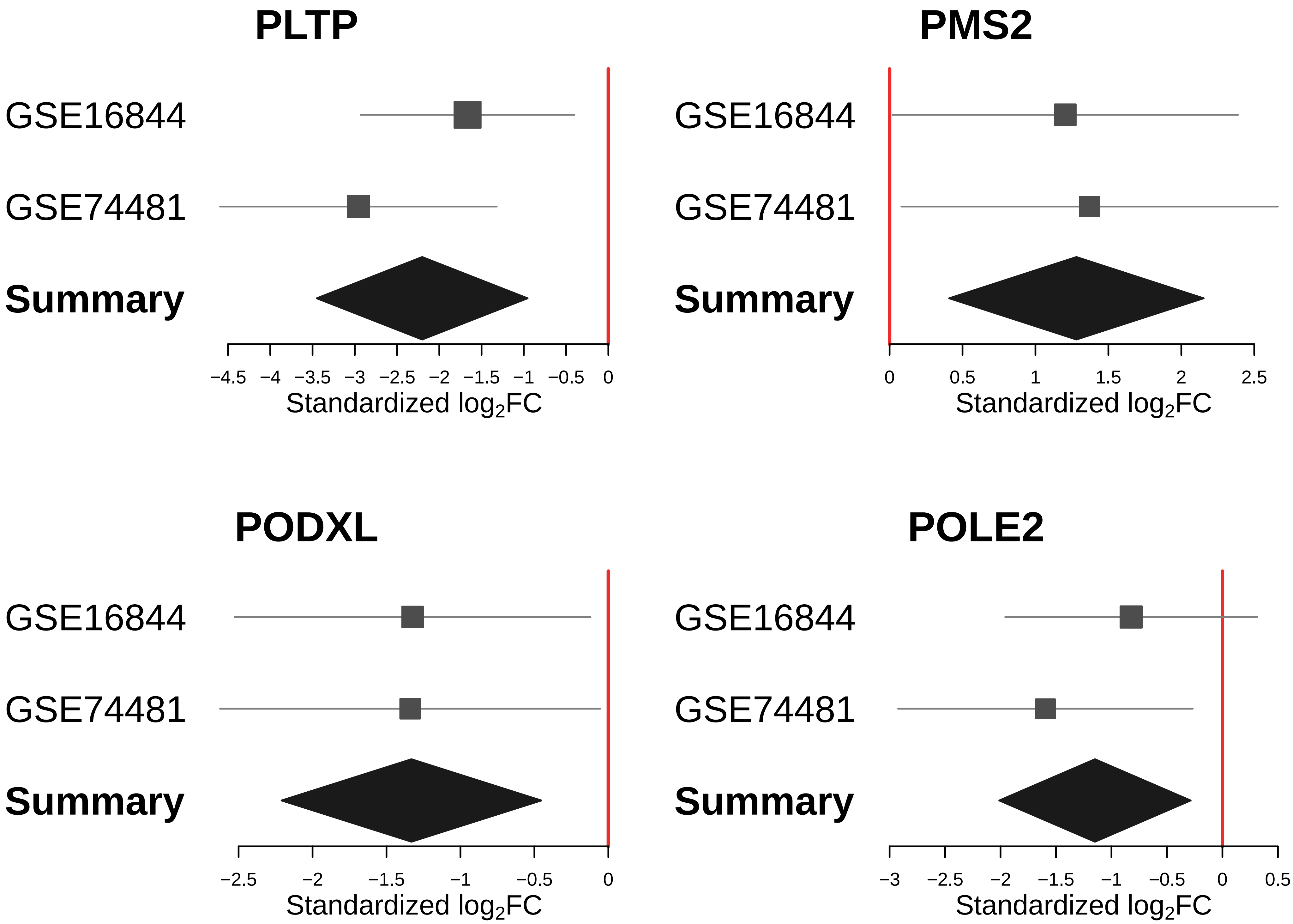
PLTP
GSE16844
GSE74481
Summary
−4.5 −4 −3.5 −3 −2.5 −2 −1.5 −1 −0.5 0
Standardized log2FC
PMS2
GSE16844
GSE74481
Summary
0 0.5 1 1.5 2 2.5
Standardized log2FC
PODXL
GSE16844
GSE74481
Summary
−2.5 −2 −1.5 −1 −0.5 0
Standardized log2FC
POLE2
GSE16844
GSE74481
Summary
−3 −2.5 −2 −1.5 −1 −0.5 0 0.5
Standardized log2FC

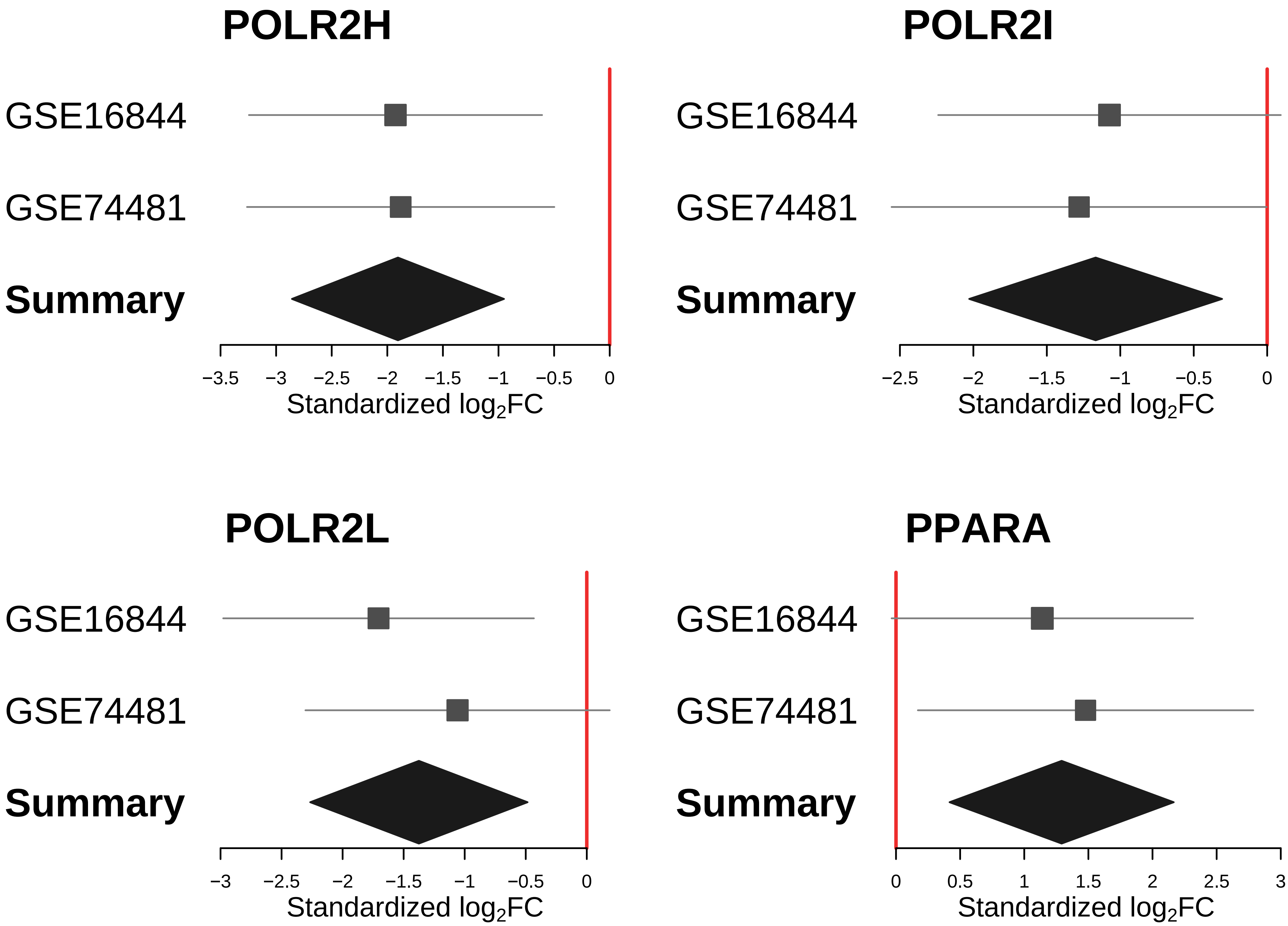
POLR2H
GSE16844
GSE74481
Summary
−3.5 −3 −2.5 −2 −1.5 −1 −0.5 0
Standardized log2FC
POLR2I
GSE16844
GSE74481
Summary
−2.5 −2 −1.5 −1 −0.5 0
Standardized log2FC
POLR2L
GSE16844
GSE74481
Summary
−3 −2.5 −2 −1.5 −1 −0.5 0
Standardized log2FC
PPARA
GSE16844
GSE74481
Summary
0 0.5 1 1.5 2 2.5 3
Standardized log2FC

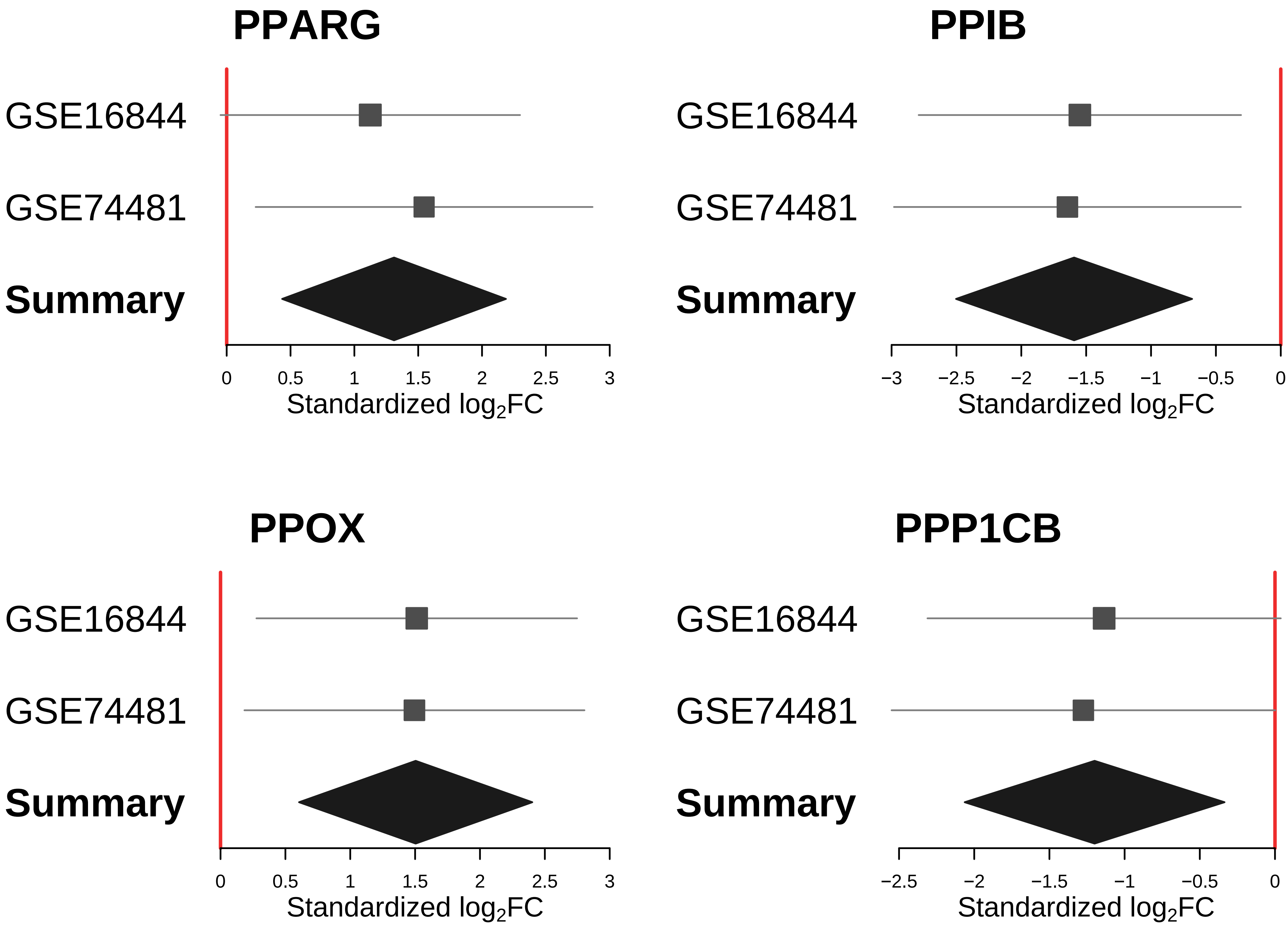
PPARG
GSE16844
GSE74481
Summary
0
0.5
1
1.5
2
2.5
3
Standardized log2FC
PPIB
GSE16844
GSE74481
Summary
−3
−2.5
−2
−1.5
−1
−0.5
0
Standardized log2FC
PPOX
GSE16844
GSE74481
Summary
0
0.5
1
1.5
2
2.5
3
Standardized log2FC
PPP1CB
GSE16844
GSE74481
Summary
−2.5
−2
−1.5
−1
−0.5
0
Standardized log2FC

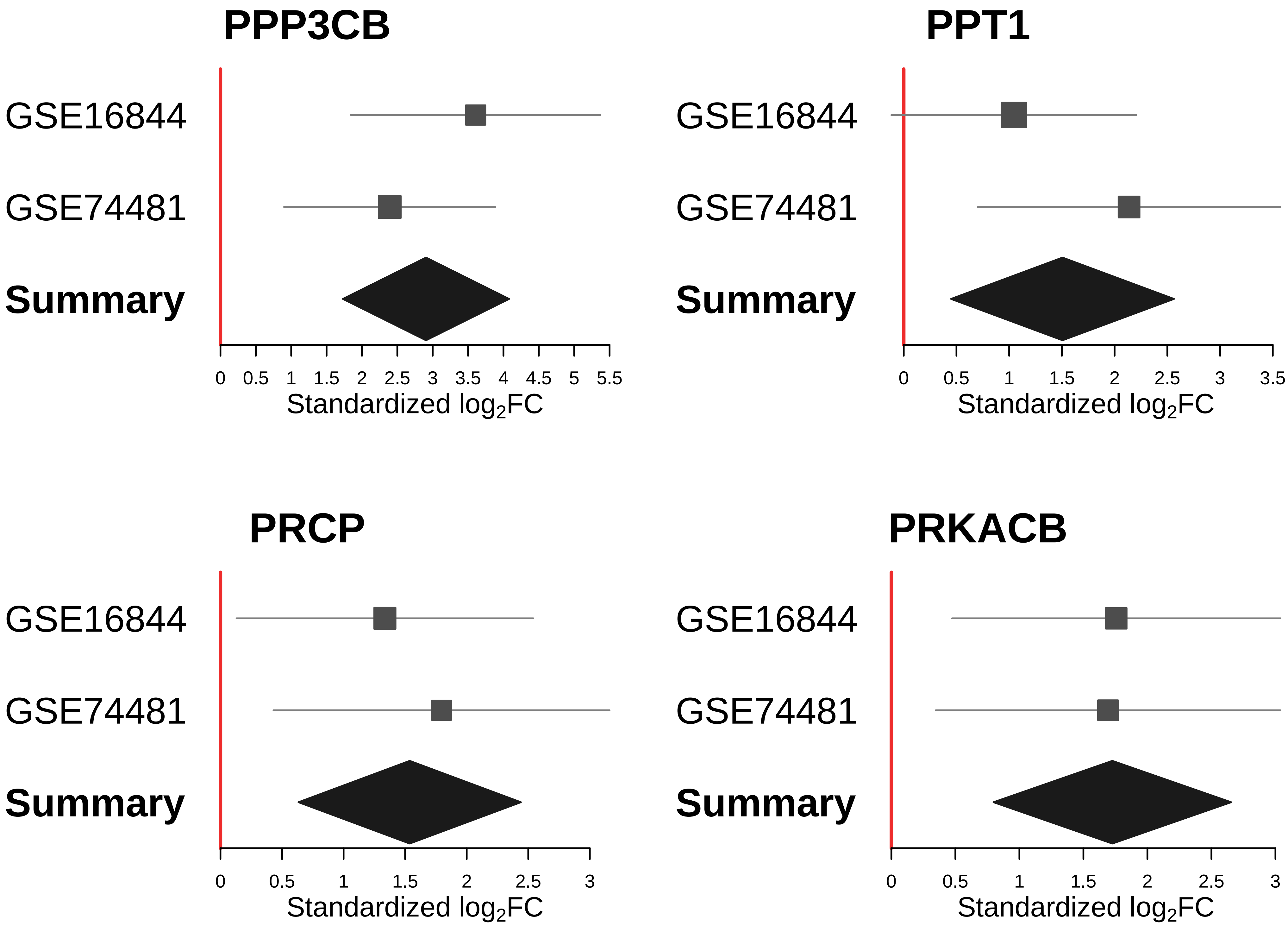
PPP3CB
GSE16844
GSE74481
Summary
0 0.5 1 1.5 2 2.5 3 3.5 4 4.5 5 5.5
Standardized log2FC
PPT1
GSE16844
GSE74481
Summary
0 0.5 1 1.5 2 2.5 3 3.5
Standardized log2FC
PRCP
GSE16844
GSE74481
Summary
0 0.5 1 1.5 2 2.5 3
Standardized log2FC
PRKACB
GSE16844
GSE74481
Summary
0 0.5 1 1.5 2 2.5 3
Standardized log2FC

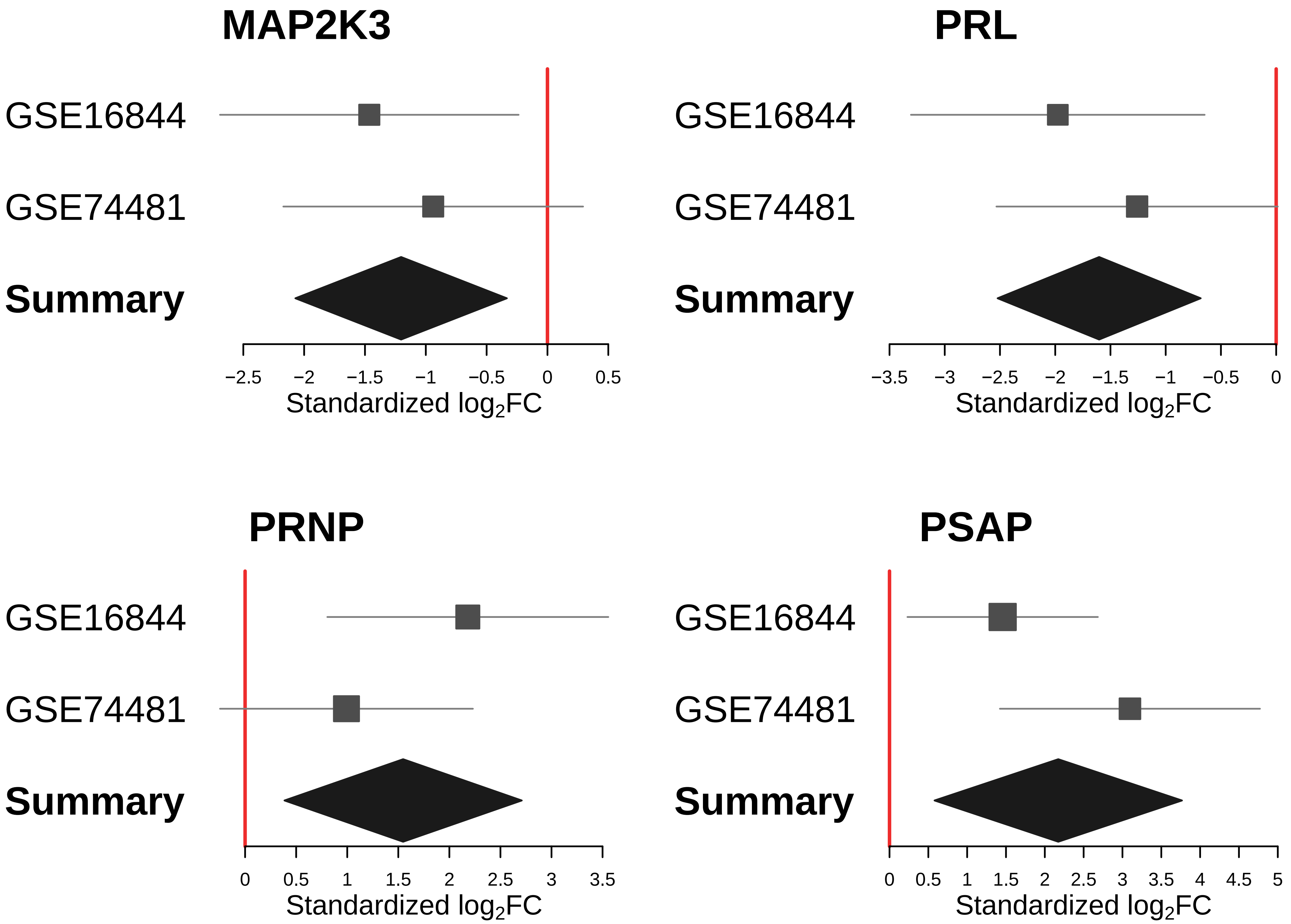
MAP2K3
GSE16844
GSE74481
Summary
−2.5 −2 −1.5 −1 −0.5 0 0.5
Standardized log2FC
PRL
GSE16844
GSE74481
Summary
−3.5 −3 −2.5 −2 −1.5 −1 −0.5 0
Standardized log2FC
PRNP
GSE16844
GSE74481
Summary
0 0.5 1 1.5 2 2.5 3 3.5
Standardized log2FC
PSAP
GSE16844
GSE74481
Summary
0 0.5 1 1.5 2 2.5 3 3.5 4 4.5 5
Standardized log2FC

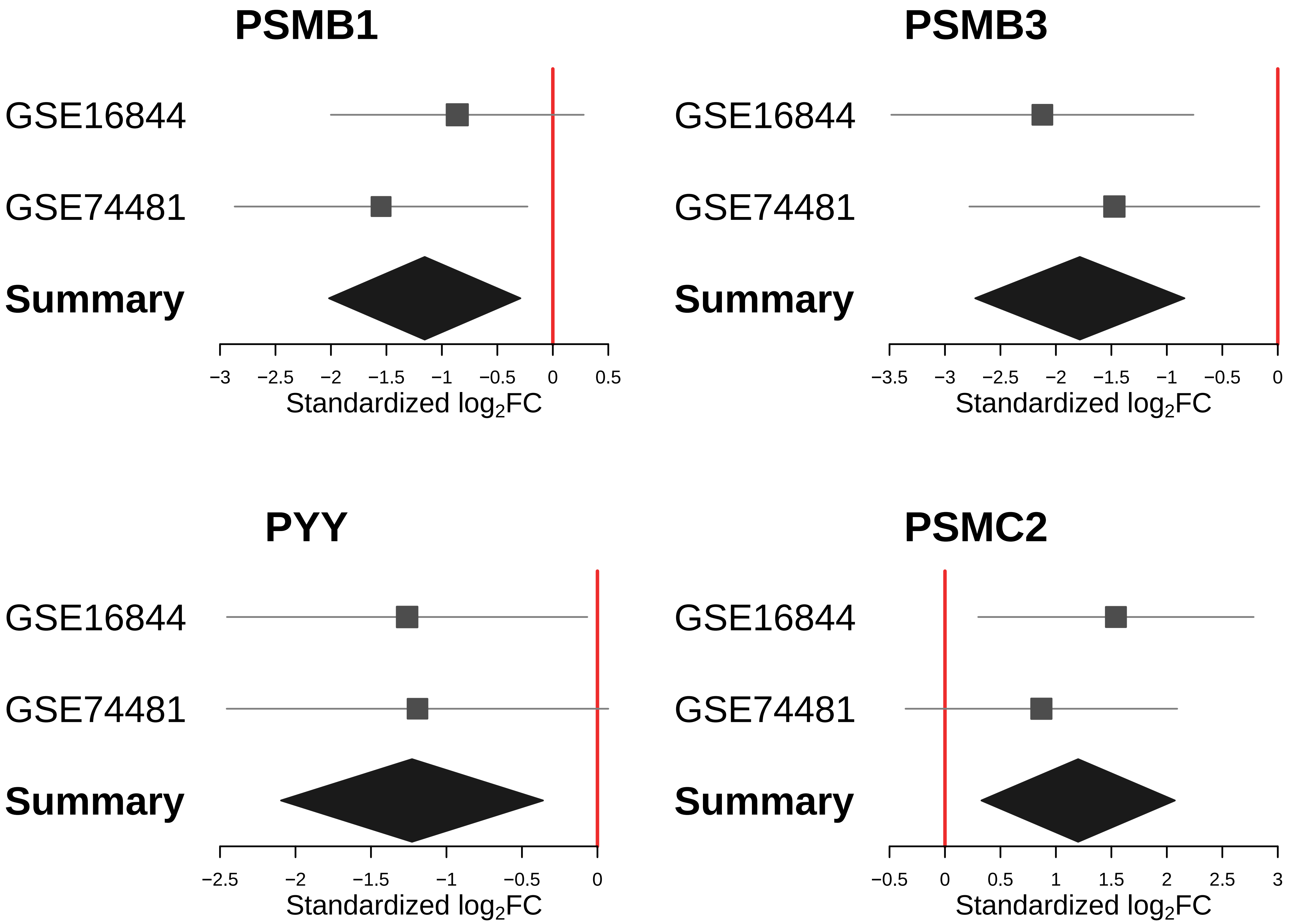
PSMB1
GSE16844
GSE74481
Summary
−3
−2.5
−2
−1.5
−1
−0.5
0
0.5
Standardized log2FC
PSMB3
GSE16844
GSE74481
Summary
−3.5
−3
−2.5
−2
−1.5
−1
−0.5
0
Standardized log2FC
PYY
GSE16844
GSE74481
Summary
−2.5
−2
−1.5
−1
−0.5
0
Standardized log2FC
PSMC2
GSE16844
GSE74481
Summary
−0.5
0
0.5
1
1.5
2
2.5
3
Standardized log2FC

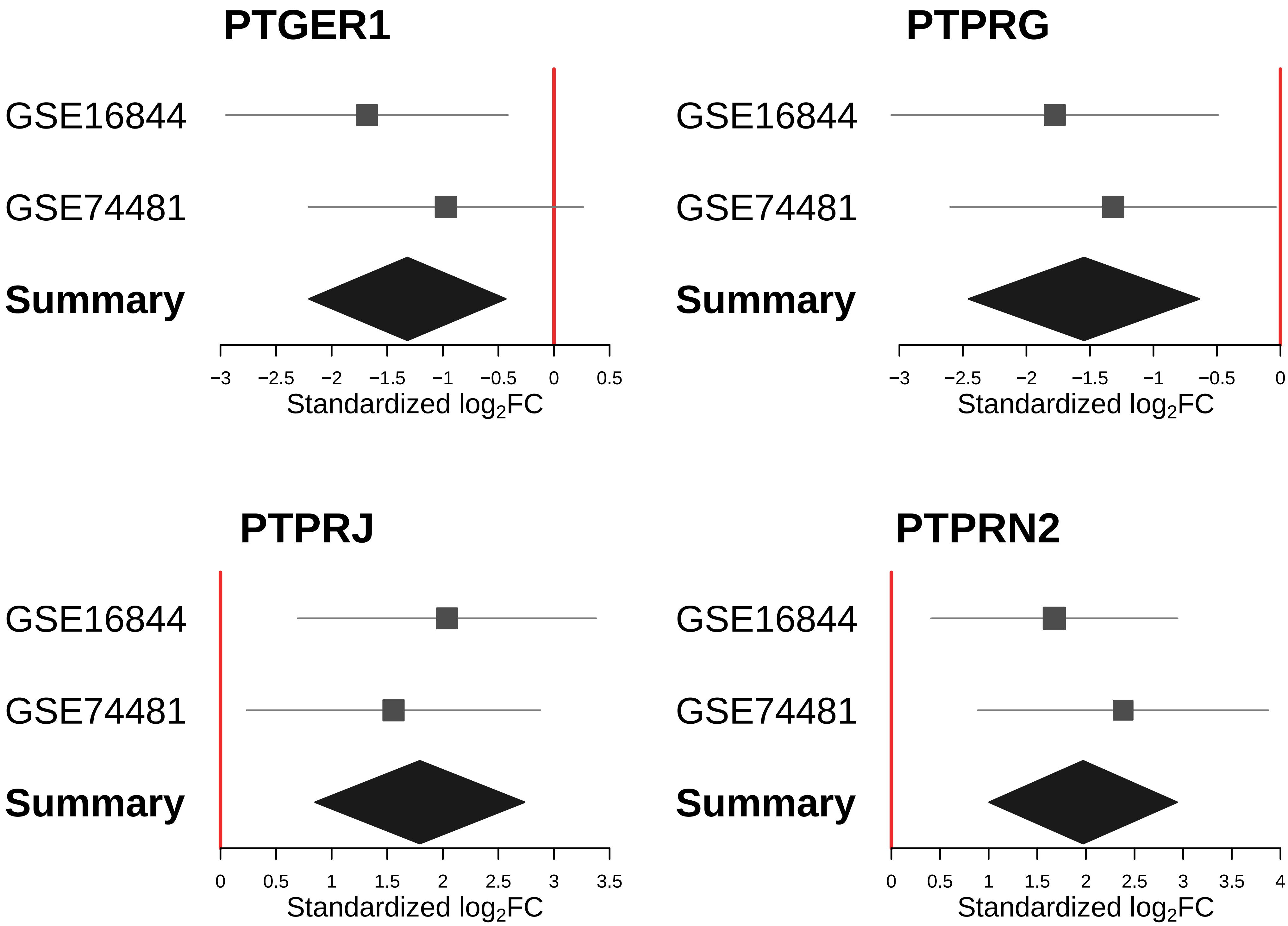

PTGER1
GSE16844
GSE74481
Summary
−3 −2.5 −2 −1.5 −1 −0.5 0 0.5
Standardized log2FC
PTPRG
GSE16844
GSE74481
Summary
−3 −2.5 −2 −1.5 −1 −0.5 0
Standardized log2FC
PTPRJ
GSE16844
GSE74481
Summary
0 0.5 1 1.5 2 2.5 3 3.5
Standardized log2FC
PTPRN2
GSE16844
GSE74481
Summary
0 0.5 1 1.5 2 2.5 3 3.5 4
Standardized log2FC

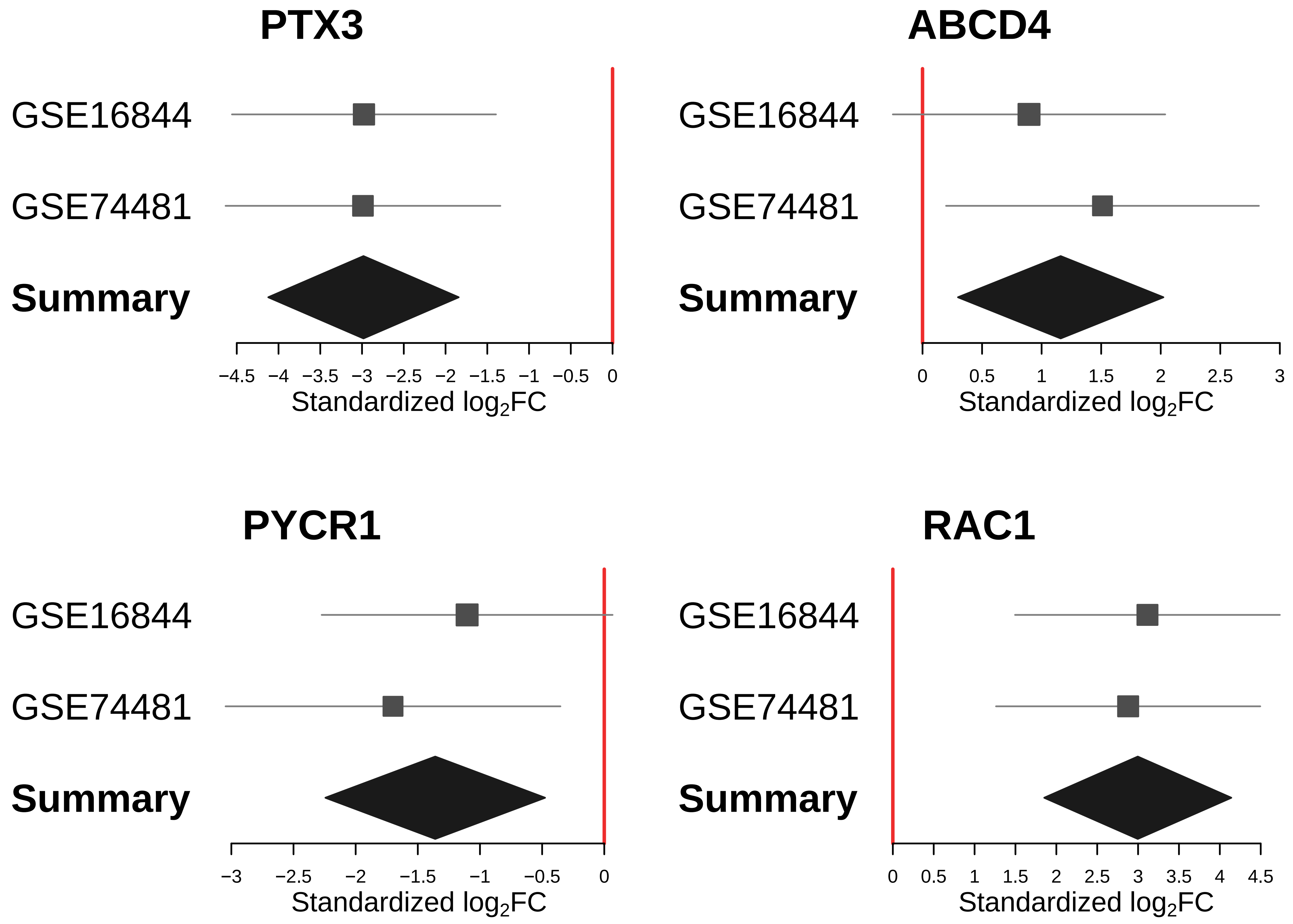
PTX3
GSE16844
GSE74481
Summary
−4.5 −4 −3.5 −3 −2.5 −2 −1.5 −1 −0.5 0
Standardized log2FC
ABCD4
GSE16844
GSE74481
Summary
0 0.5 1 1.5 2 2.5 3
Standardized log2FC
PYCR1
GSE16844
GSE74481
Summary
−3 −2.5 −2 −1.5 −1 −0.5 0
Standardized log2FC
RAC1
GSE16844
GSE74481
Summary
0 0.5 1 1.5 2 2.5 3 3.5 4 4.5
Standardized log2FC

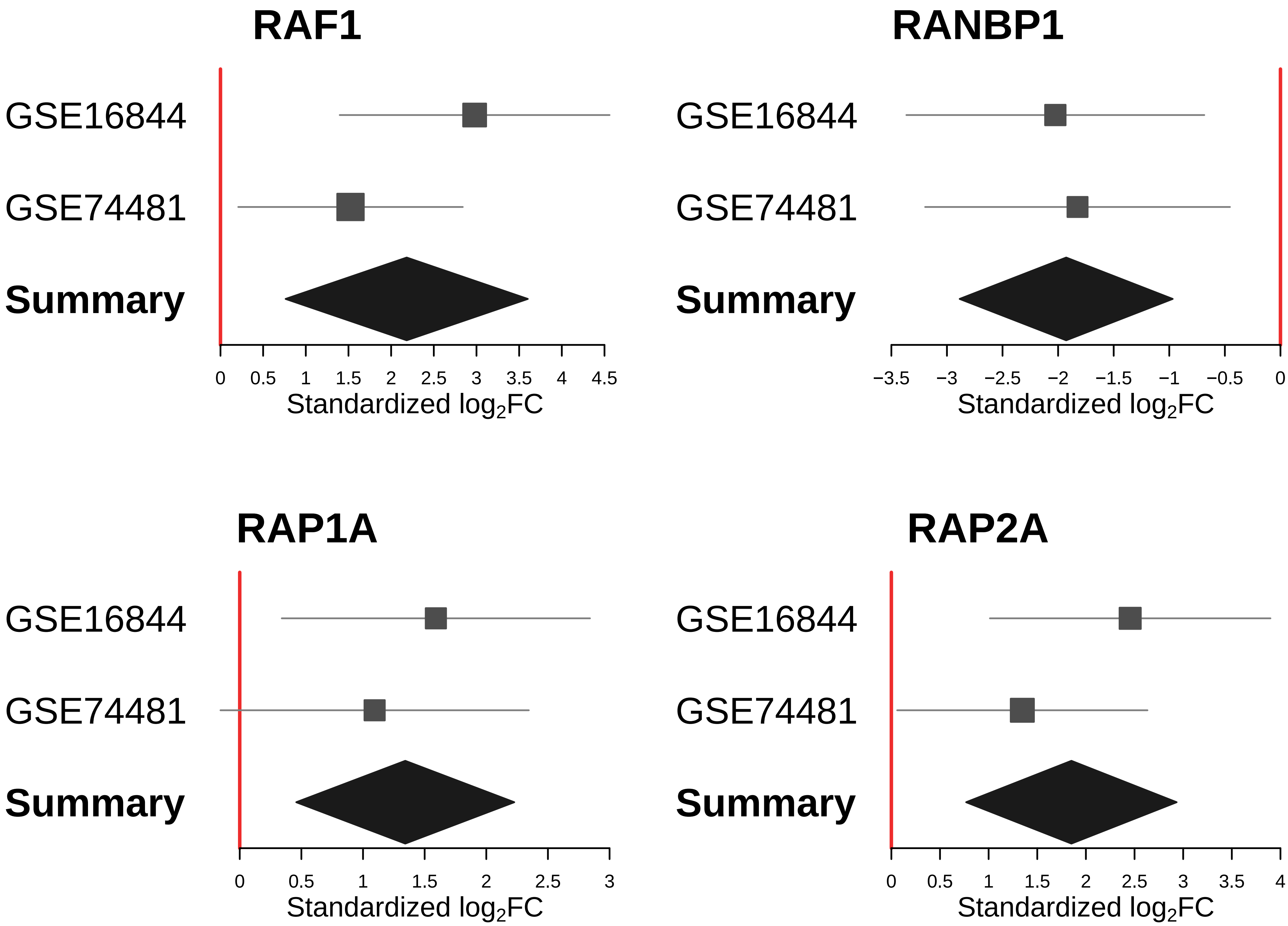
RAF1
GSE16844
GSE74481
Summary
0
0.5
1
1.5
2
2.5
3
3.5
4
4.5
Standardized log2FC
RANBP1
GSE16844
GSE74481
Summary
−3.5
−3
−2.5
−2
−1.5
−1
−0.5
0
Standardized log2FC
RAP1A
GSE16844
GSE74481
Summary
0
0.5
1
1.5
2
2.5
3
Standardized log2FC
RAP2A
GSE16844
GSE74481
Summary
0
0.5
1
1.5
2
2.5
3
3.5
4
Standardized log2FC

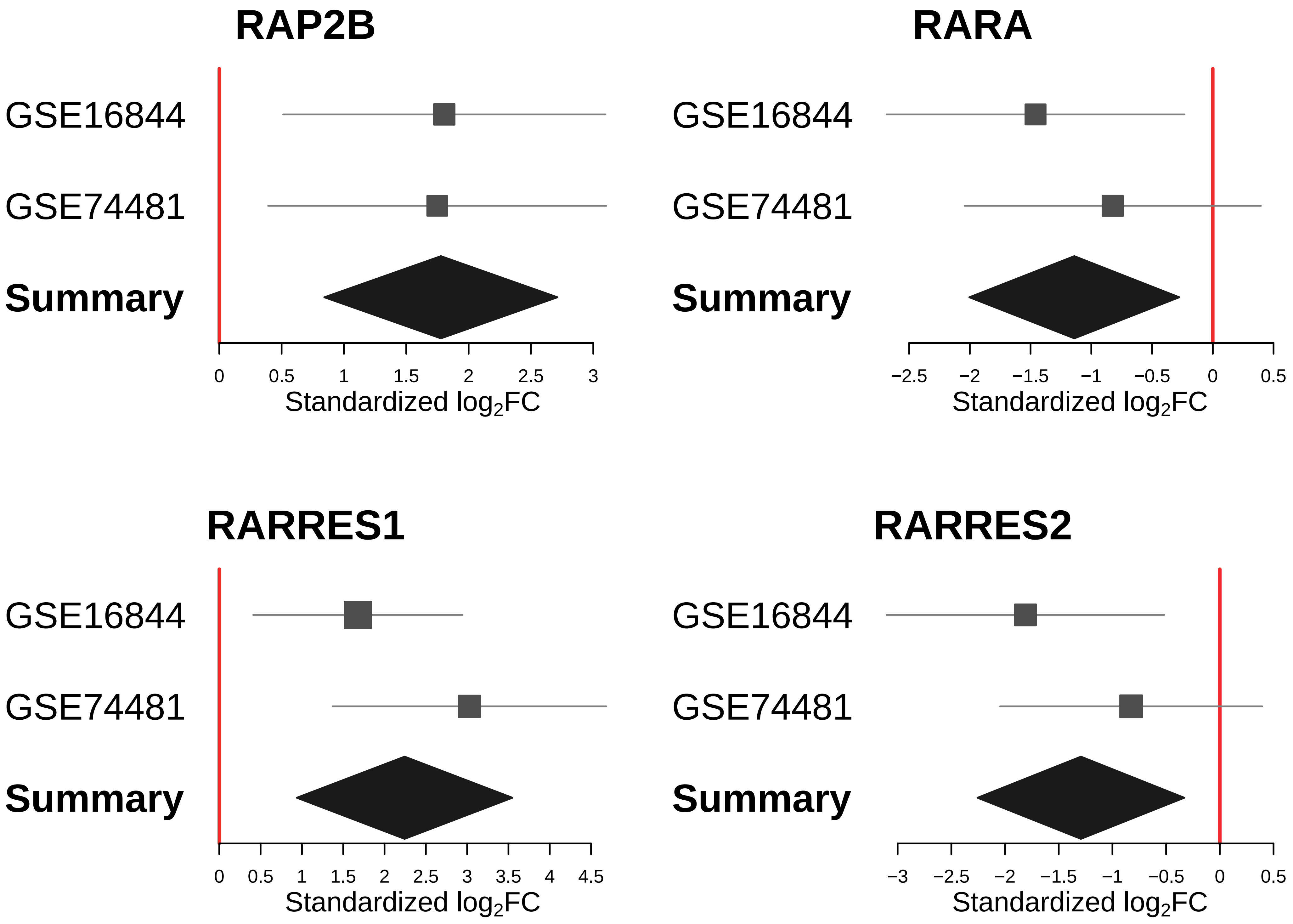
RAP2B
GSE16844
GSE74481
Summary
0
0.5
1
1.5
2
2.5
3
Standardized log2FC
RARA
GSE16844
GSE74481
Summary
−2.5
−2
−1.5
−1
−0.5
0
0.5
Standardized log2FC
RARRES1
GSE16844
GSE74481
Summary
0
0.5
1
1.5
2
2.5
3
3.5
4
4.5
Standardized log2FC
RARRES2
GSE16844
GSE74481
Summary
−3
−2.5
−2
−1.5
−1
−0.5
0
0.5
Standardized log2FC

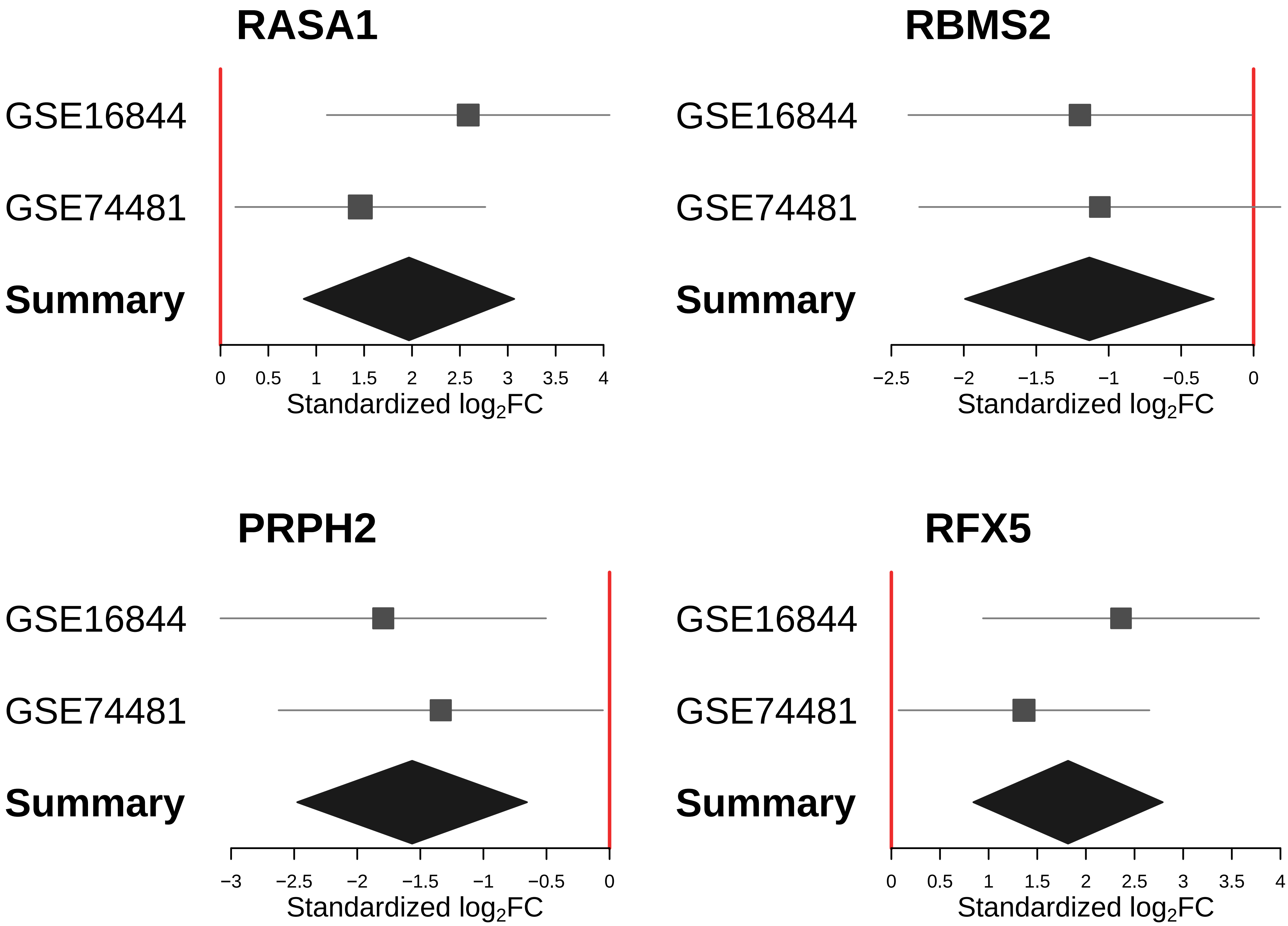
RASA1
GSE16844
GSE74481
Summary
0 0.5 1 1.5 2 2.5 3 3.5 4
Standardized log2FC
RBMS2
GSE16844
GSE74481
Summary
−2.5 −2 −1.5 −1 −0.5 0
Standardized log2FC
PRPH2
GSE16844
GSE74481
Summary
−3 −2.5 −2 −1.5 −1 −0.5 0
Standardized log2FC
RFX5
GSE16844
GSE74481
Summary
0 0.5 1 1.5 2 2.5 3 3.5 4
Standardized log2FC

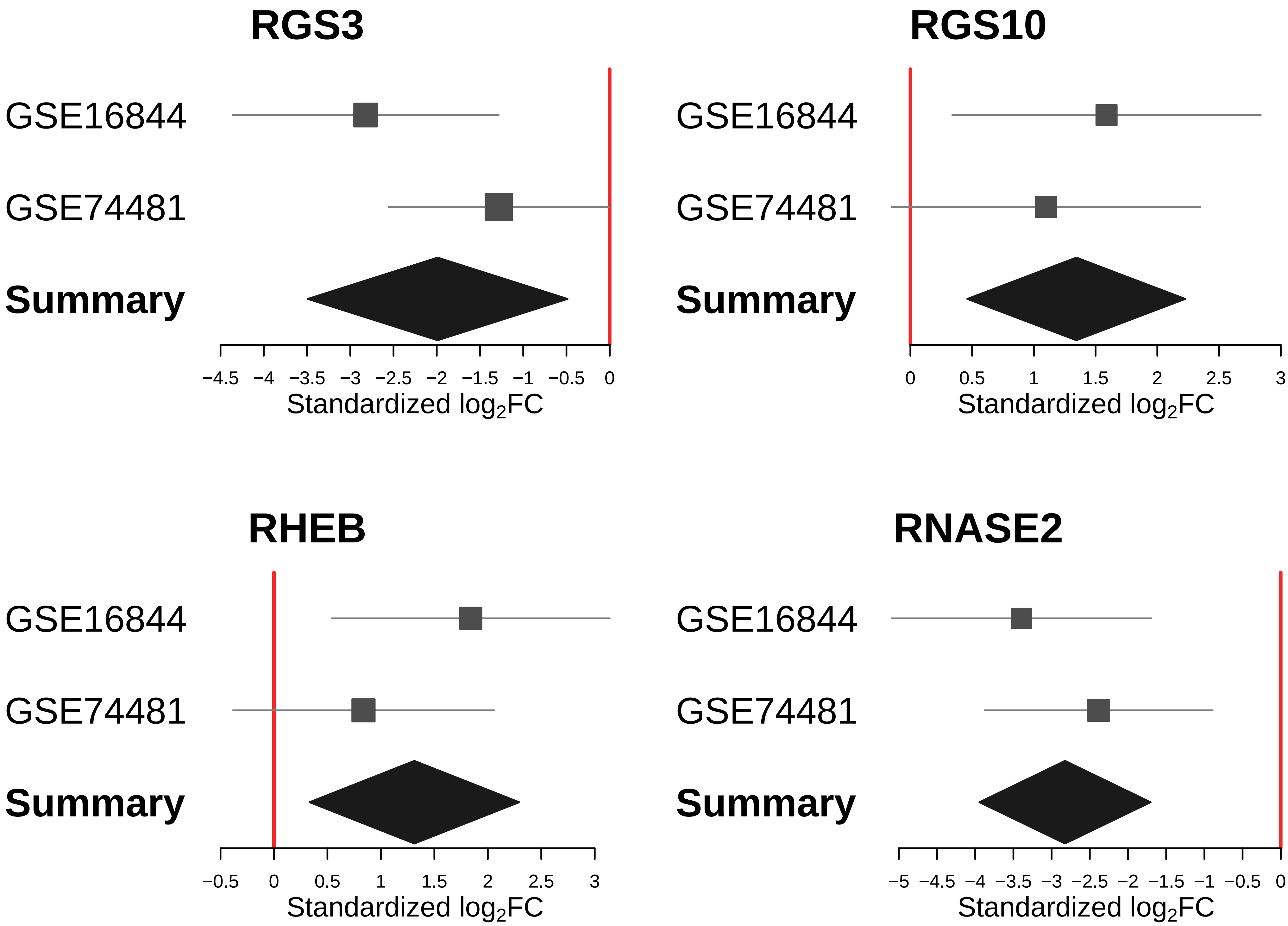
RGS3
GSE16844
GSE74481
Summary
−4.5 −4 −3.5 −3 −2.5 −2 −1.5 −1 −0.5 0
Standardized log2FC
RGS10
GSE16844
GSE74481
Summary
0 0.5 1 1.5 2 2.5 3
Standardized log2FC
RHEB
GSE16844
GSE74481
Summary
−0.5 0 0.5 1 1.5 2 2.5 3
Standardized log2FC
RNASE2
GSE16844
GSE74481
Summary
−5 −4.5 −4 −3.5 −3 −2.5 −2 −1.5 −1 −0.5 0
Standardized log2FC

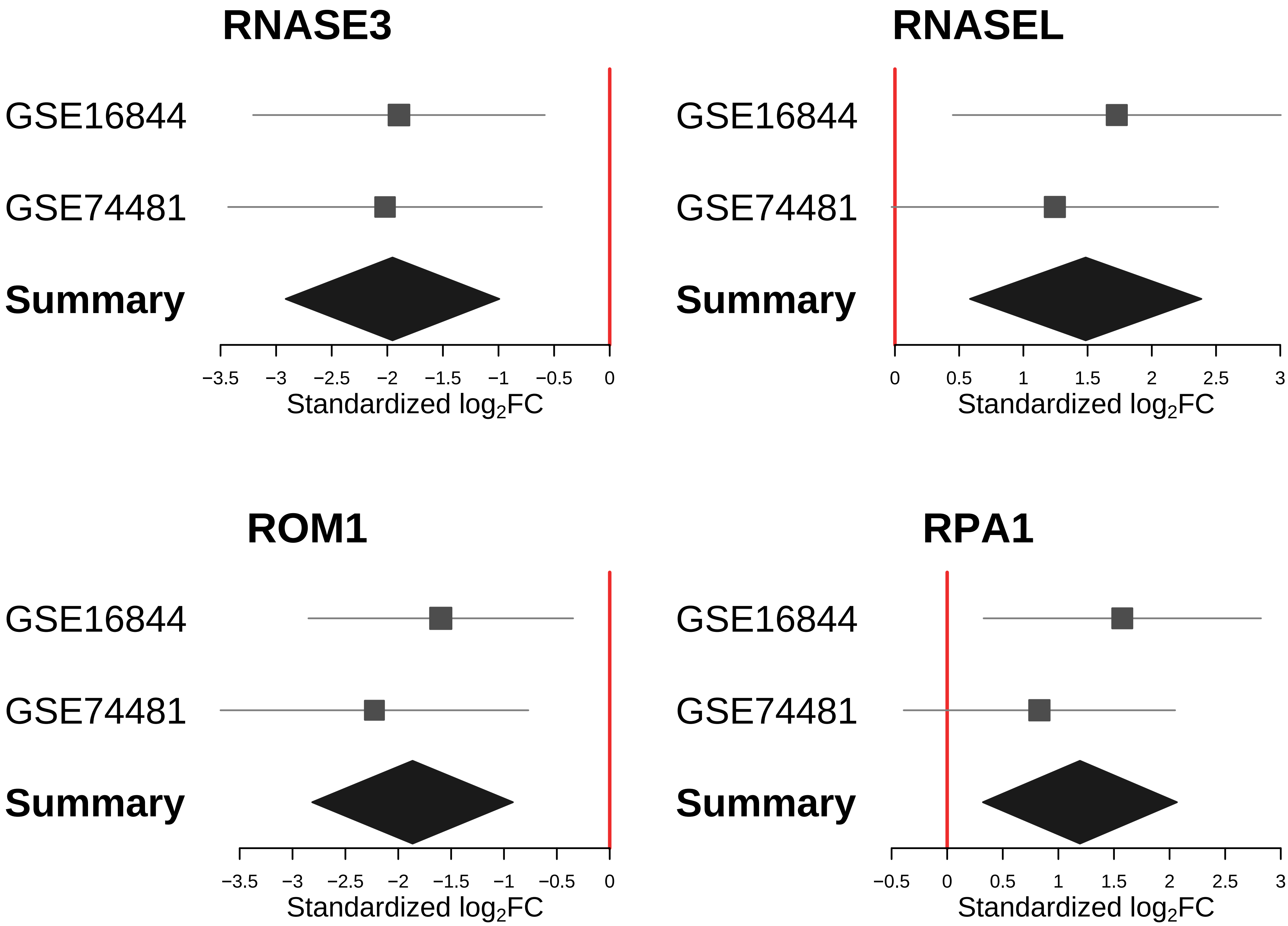
RNASE3
GSE16844
GSE74481
Summary
−3.5 −3 −2.5 −2 −1.5 −1 −0.5 0
Standardized log2FC
RNASEL
GSE16844
GSE74481
Summary
0 0.5 1 1.5 2 2.5 3
Standardized log2FC
ROM1
GSE16844
GSE74481
Summary
−3.5 −3 −2.5 −2 −1.5 −1 −0.5 0
Standardized log2FC
RPA1
GSE16844
GSE74481
Summary
−0.5 0 0.5 1 1.5 2 2.5 3
Standardized log2FC

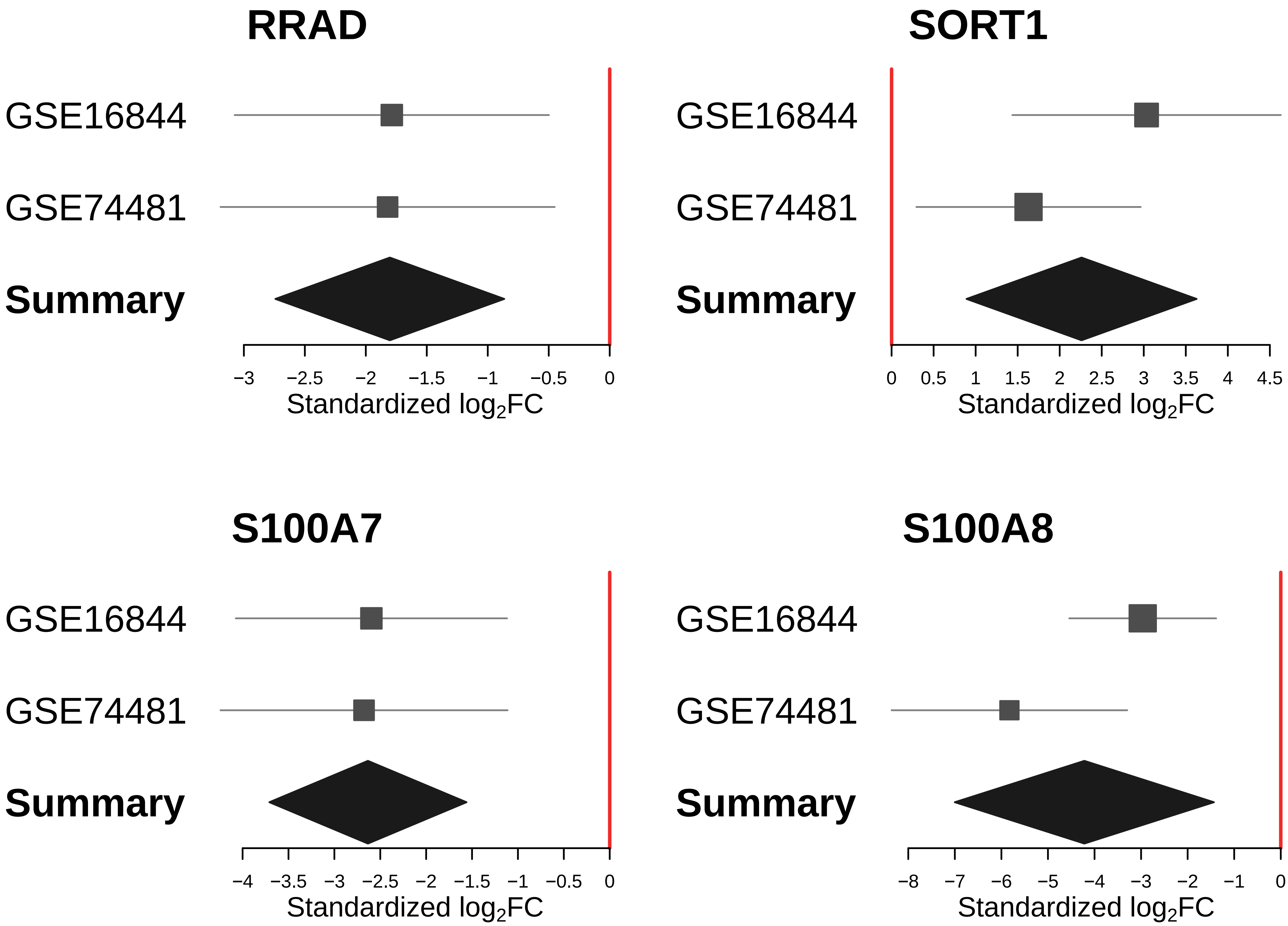

RRAD
GSE16844
GSE74481
Summary
−3 −2.5 −2 −1.5 −1 −0.5 0
Standardized $log_2FC$
SORT1
GSE16844
GSE74481
Summary
0 0.5 1 1.5 2 2.5 3 3.5 4 4.5
Standardized $log_2FC$
S100A7
GSE16844
GSE74481
Summary
−4 −3.5 −3 −2.5 −2 −1.5 −1 −0.5 0
Standardized $log_2FC$
S100A8
GSE16844
GSE74481
Summary
−8 −7 −6 −5 −4 −3 −2 −1 0
Standardized $log_2FC$

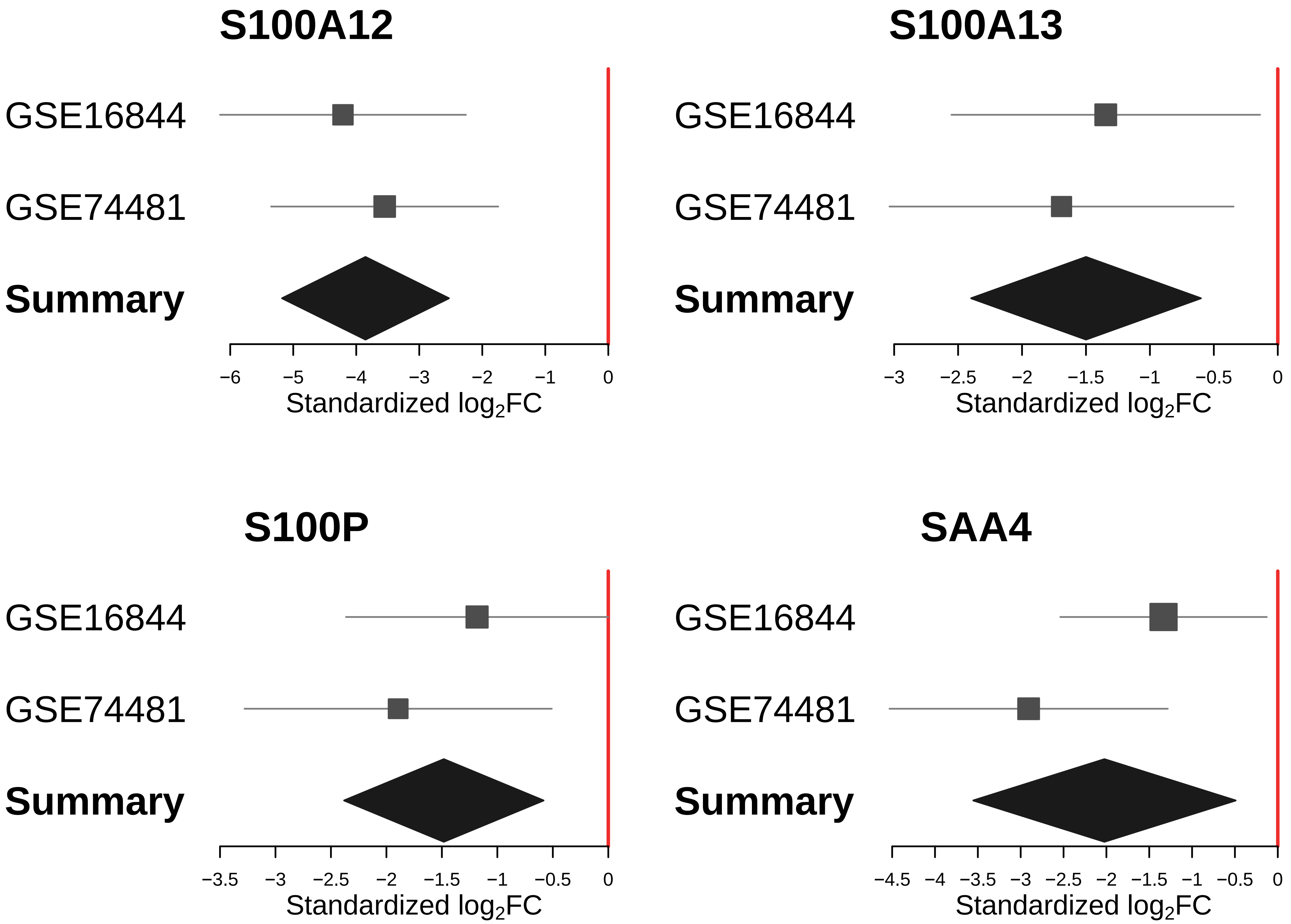
S100A12
GSE16844
GSE74481
Summary
−6 −5 −4 −3 −2 −1 0
Standardized log2FC
S100A13
GSE16844
GSE74481
Summary
−3 −2.5 −2 −1.5 −1 −0.5 0
Standardized log2FC
S100P
GSE16844
GSE74481
Summary
−3.5 −3 −2.5 −2 −1.5 −1 −0.5 0
Standardized log2FC
SAA4
GSE16844
GSE74481
Summary
−4.5 −4 −3.5 −3 −2.5 −2 −1.5 −1 −0.5 0
Standardized log2FC

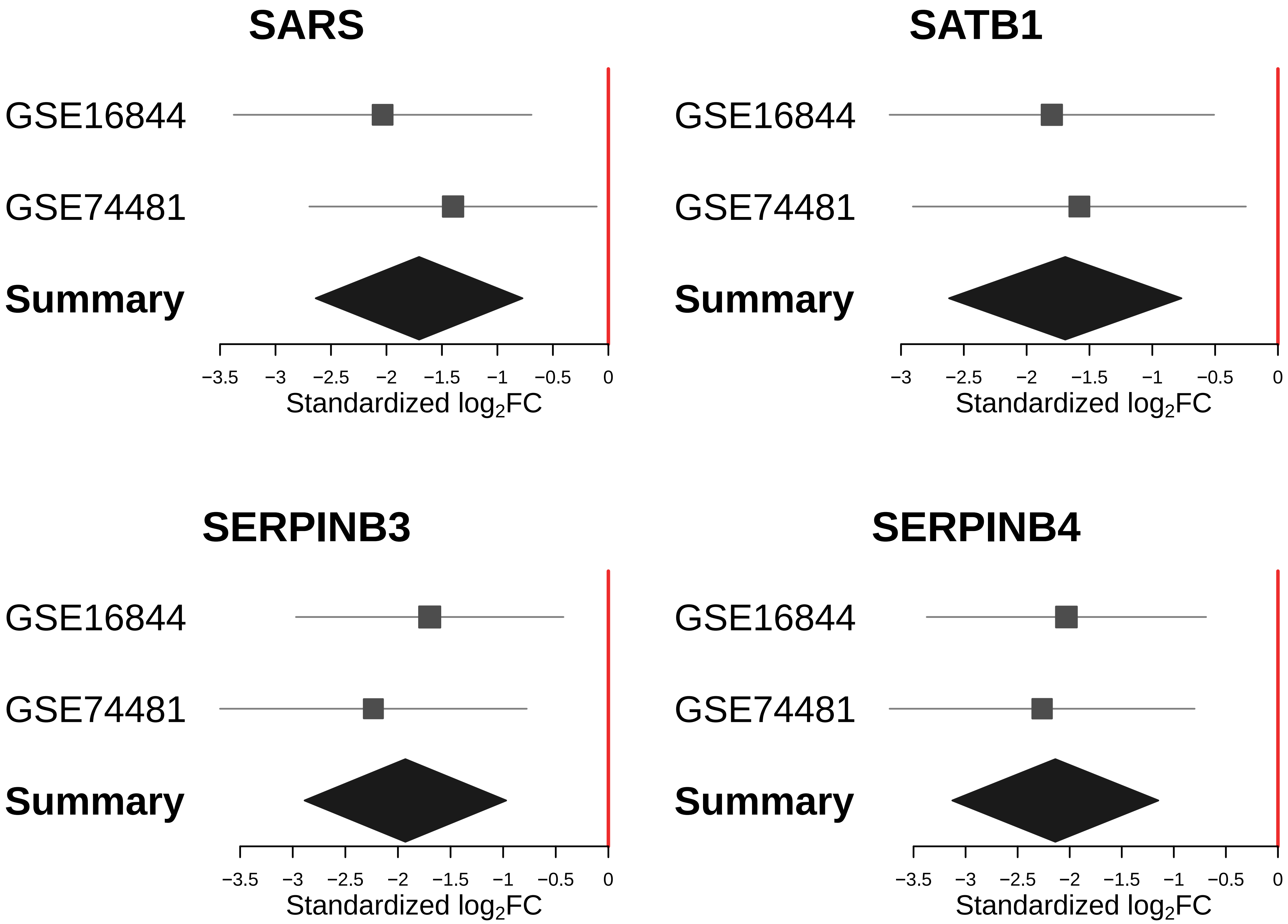
SARS
GSE16844
GSE74481
Summary
−3.5 −3 −2.5 −2 −1.5 −1 −0.5 0
Standardized log2FC
SATB1
GSE16844
GSE74481
Summary
−3 −2.5 −2 −1.5 −1 −0.5 0
Standardized log2FC
SERPINB3
GSE16844
GSE74481
Summary
−3.5 −3 −2.5 −2 −1.5 −1 −0.5 0
Standardized log2FC
SERPINB4
GSE16844
GSE74481
Summary
−3.5 −3 −2.5 −2 −1.5 −1 −0.5 0
Standardized log2FC

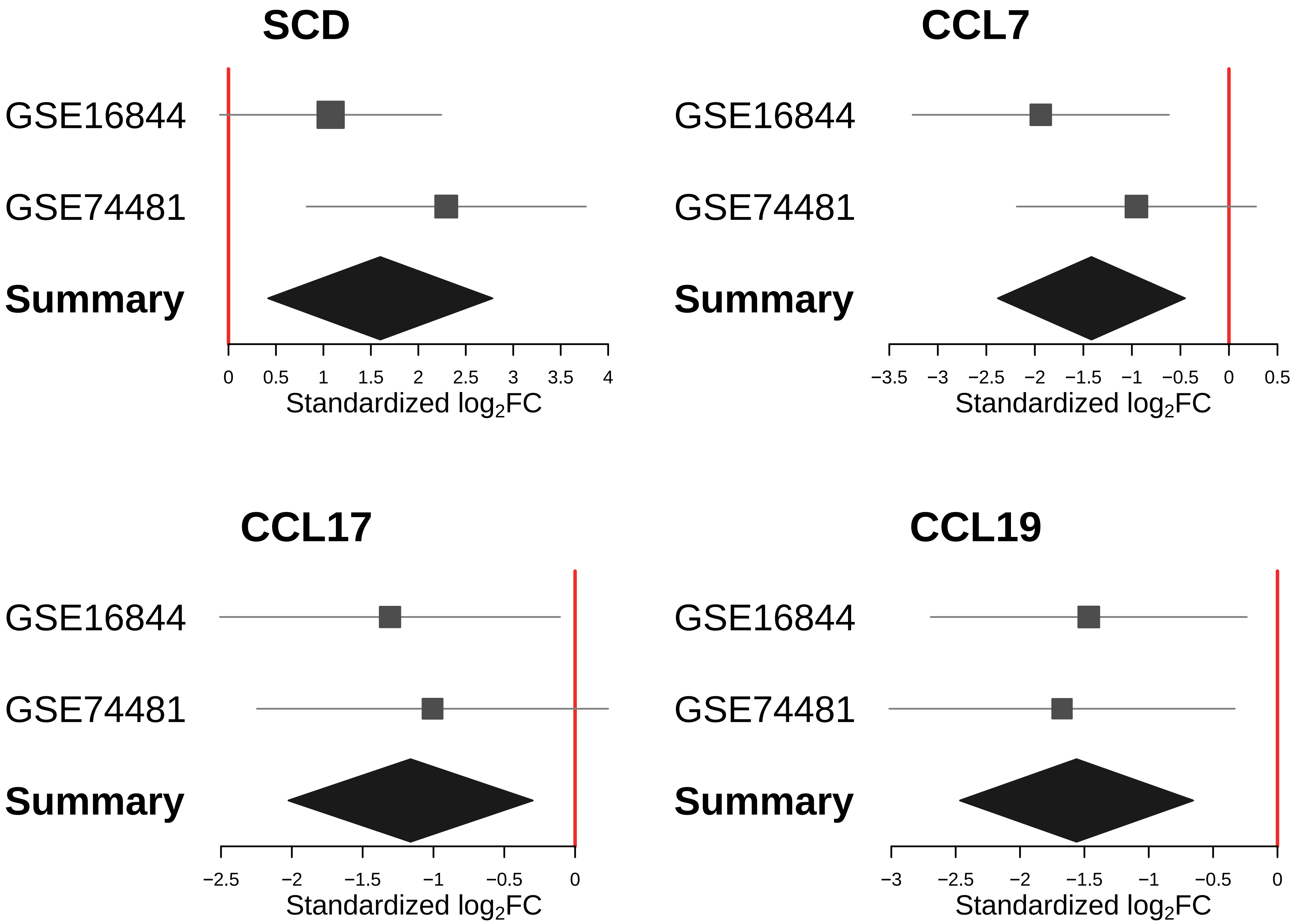

SCD
GSE16844
GSE74481
Summary
0
0.5
1
1.5
2
2.5
3
3.5
4
Standardized log2FC
CCL7
GSE16844
GSE74481
Summary
−3.5
−3
−2.5
−2
−1.5
−1
−0.5
0
0.5
Standardized log2FC
CCL17
GSE16844
GSE74481
Summary
−2.5
−2
−1.5
−1
−0.5
0
Standardized log2FC
CCL19
GSE16844
GSE74481
Summary
−3
−2.5
−2
−1.5
−1
−0.5
0
Standardized log2FC

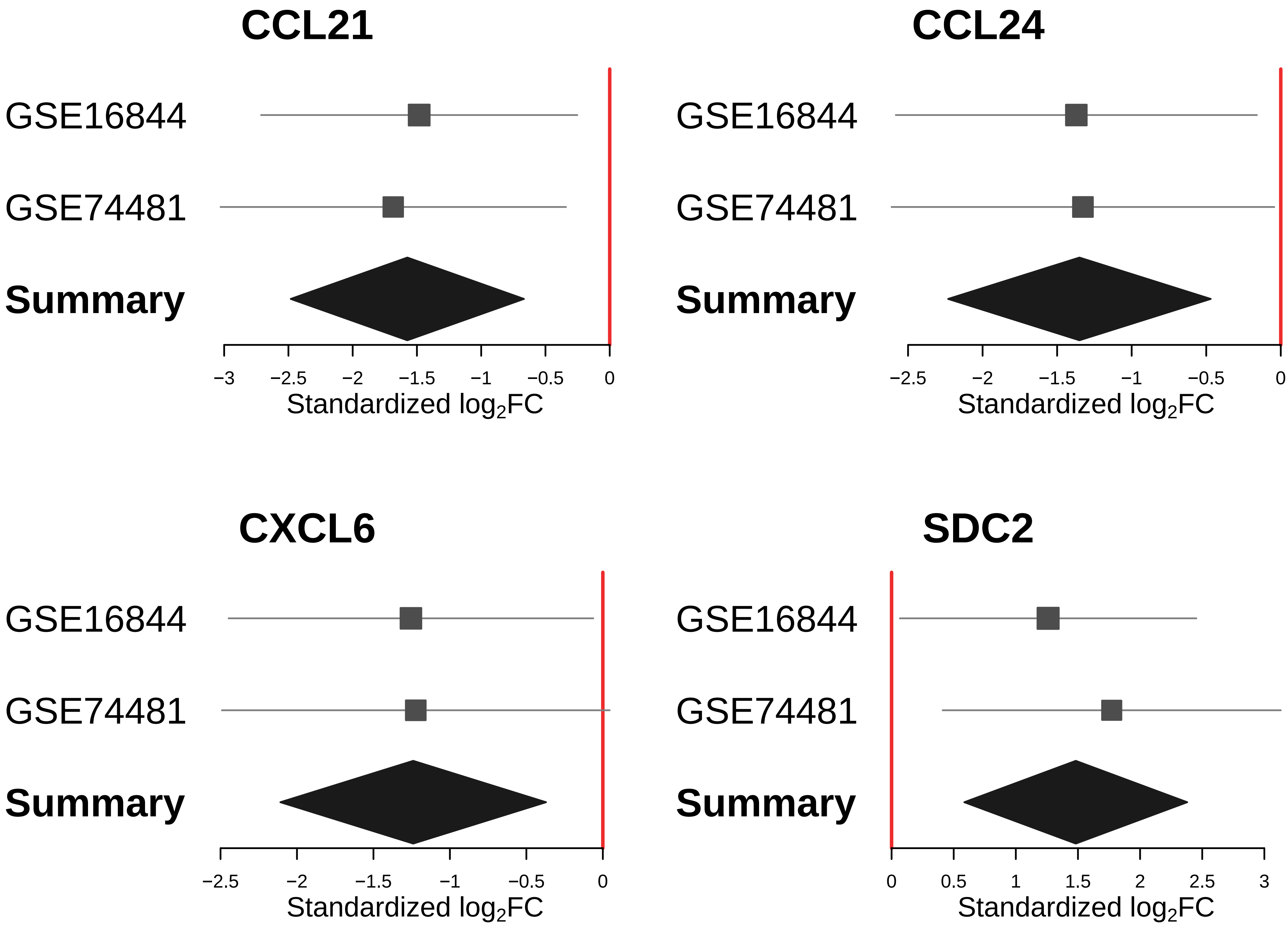

CCL21
GSE16844
GSE74481
Summary
−3 −2.5 −2 −1.5 −1 −0.5 0
Standardized log2FC
CCL24
GSE16844
GSE74481
Summary
−2.5 −2 −1.5 −1 −0.5 0
Standardized log2FC
CXCL6
GSE16844
GSE74481
Summary
−2.5 −2 −1.5 −1 −0.5 0
Standardized log2FC
SDC2
GSE16844
GSE74481
Summary
0 0.5 1 1.5 2 2.5 3
Standardized log2FC

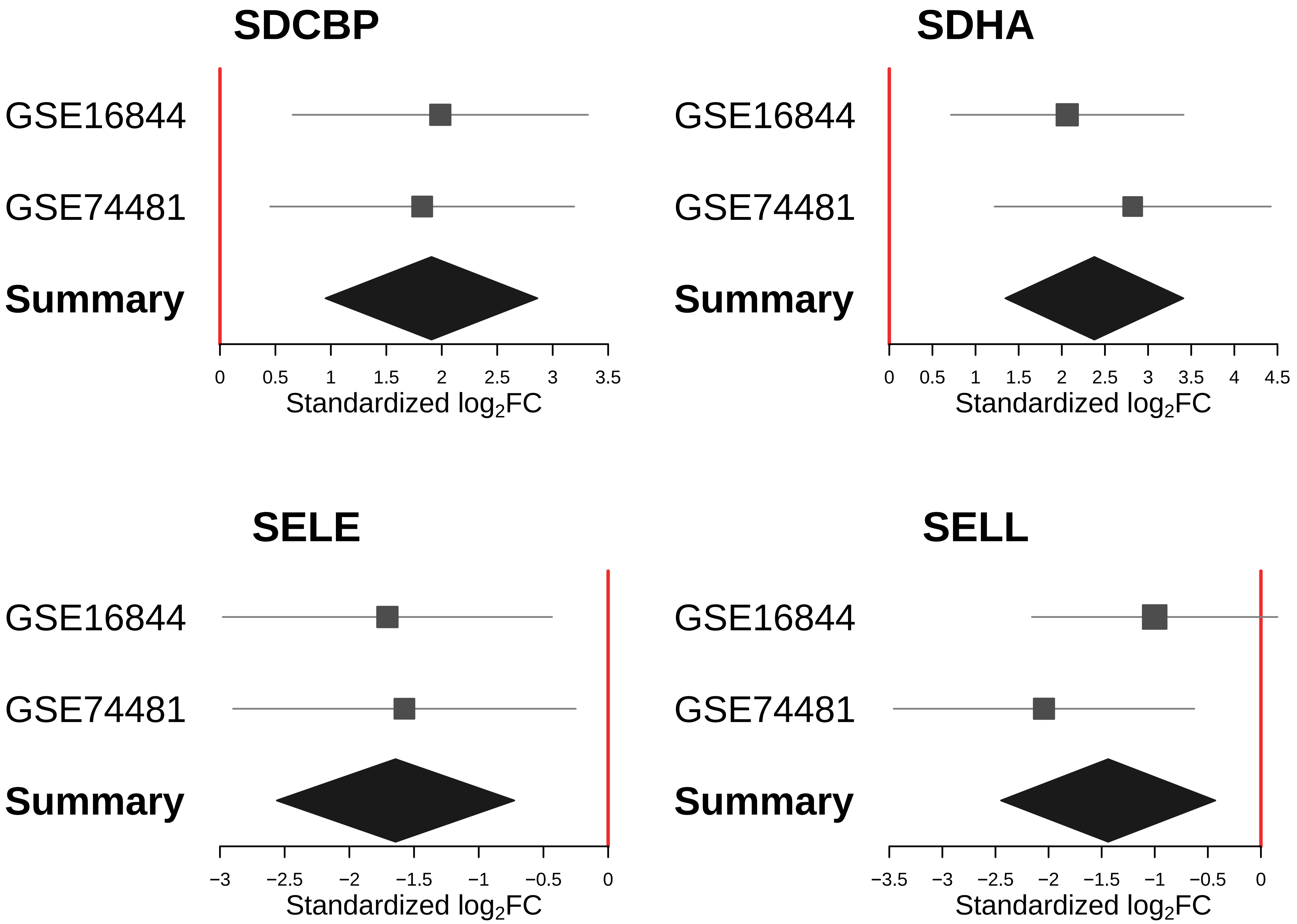
SDCBP
GSE16844
GSE74481
Summary
0
0.5
1
1.5
2
2.5
3
3.5
Standardized log2FC
SDHA
GSE16844
GSE74481
Summary
0
0.5
1
1.5
2
2.5
3
3.5
4
4.5
Standardized log2FC
SELE
GSE16844
GSE74481
Summary
−3
−2.5
−2
−1.5
−1
−0.5
0
Standardized log2FC
SELL
GSE16844
GSE74481
Summary
−3.5
−3
−2.5
−2
−1.5
−1
−0.5
0
Standardized log2FC

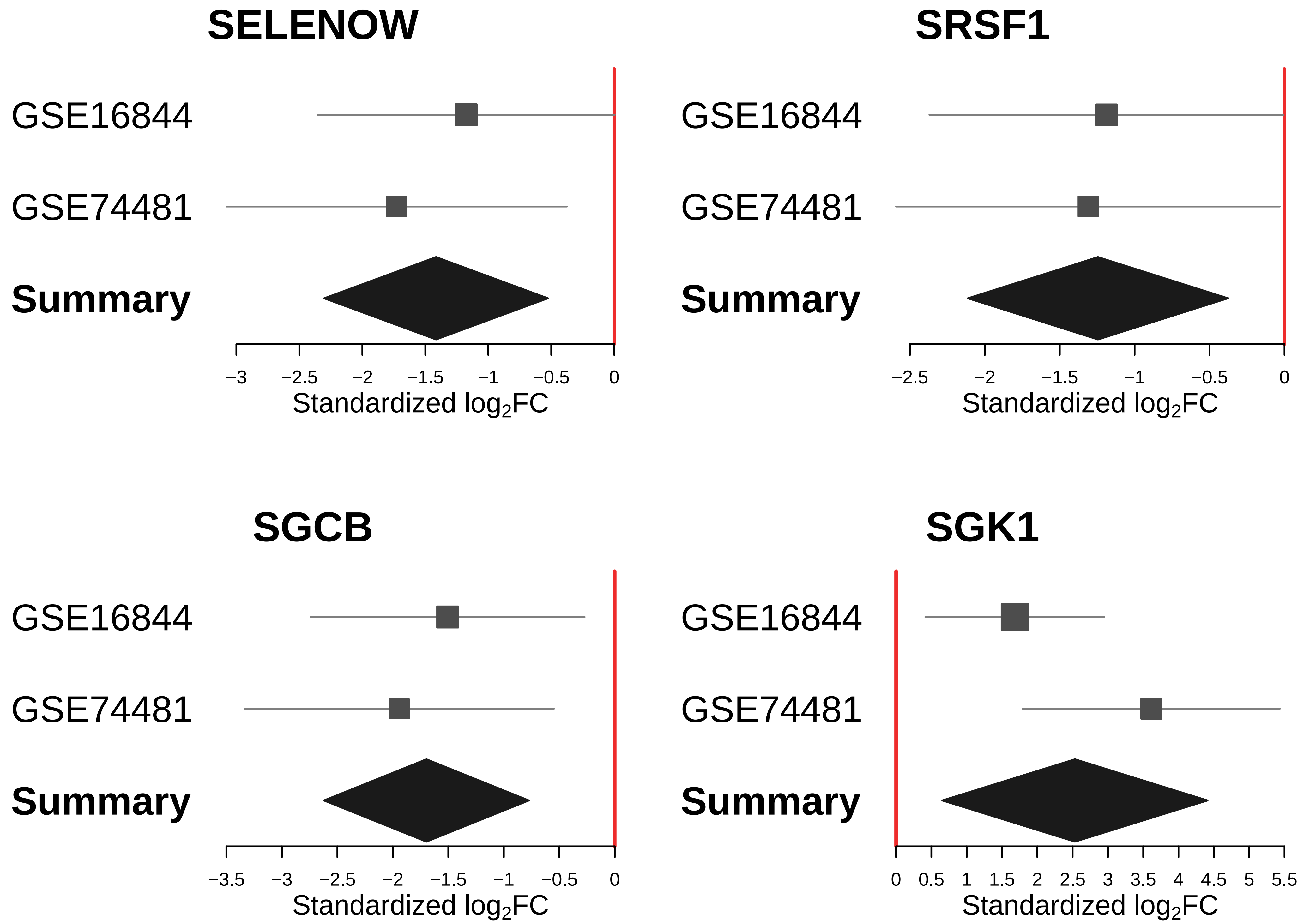
SELENOW
GSE16844
GSE74481
Summary
−3
−2.5
−2
−1.5
−1
−0.5
0
Standardized log2FC
SRSF1
GSE16844
GSE74481
Summary
−2.5
−2
−1.5
−1
−0.5
0
Standardized log2FC
SGCB
GSE16844
GSE74481
Summary
−3.5
−3
−2.5
−2
−1.5
−1
−0.5
0
Standardized log2FC
SGK1
GSE16844
GSE74481
Summary
0
0.5
1
1.5
2
2.5
3
3.5
4
4.5
5
5.5
Standardized log2FC

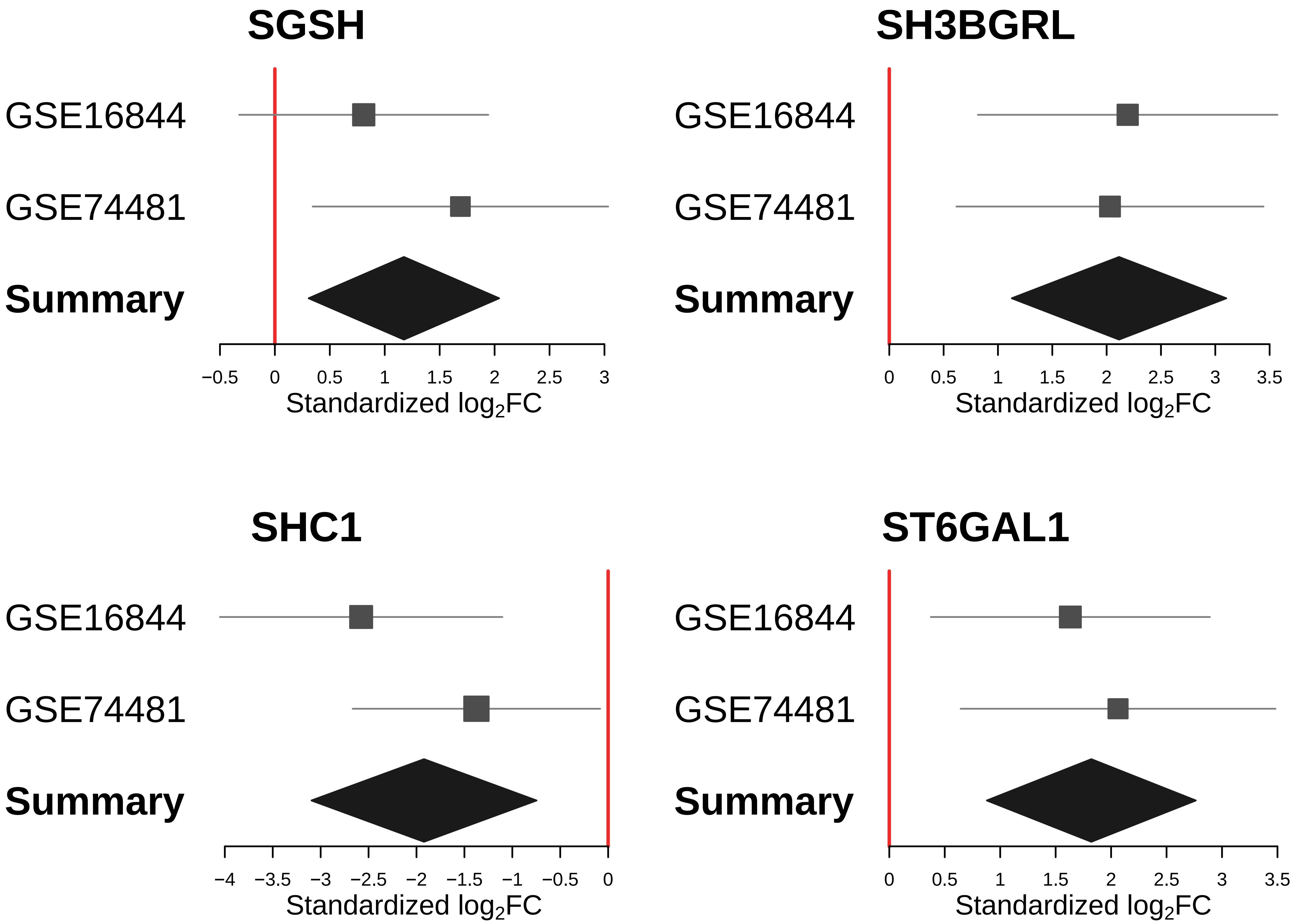

SGSH
GSE16844
GSE74481
Summary
−0.5 0 0.5 1 1.5 2 2.5 3
Standardized log2FC
SH3BGRL
GSE16844
GSE74481
Summary
0 0.5 1 1.5 2 2.5 3 3.5
Standardized log2FC
SHC1
GSE16844
GSE74481
Summary
−4 −3.5 −3 −2.5 −2 −1.5 −1 −0.5 0
Standardized log2FC
ST6GAL1
GSE16844
GSE74481
Summary
0 0.5 1 1.5 2 2.5 3 3.5
Standardized log2FC

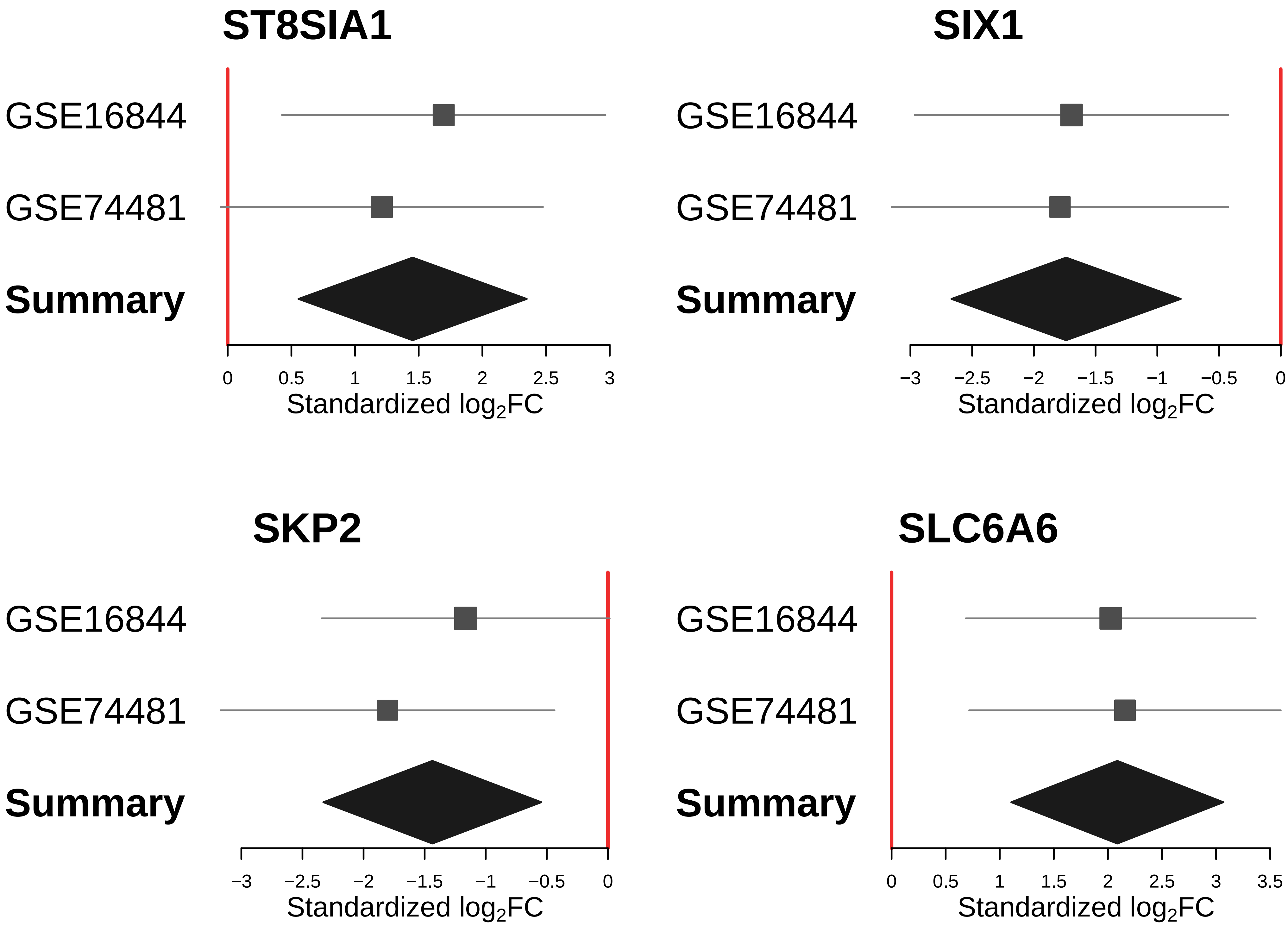
ST8SIA1
GSE16844
GSE74481
Summary
0
0.5
1
1.5
2
2.5
3
Standardized $log_2FC$
SIX1
GSE16844
GSE74481
Summary
−3
−2.5
−2
−1.5
−1
−0.5
0
Standardized $log_2FC$
SKP2
GSE16844
GSE74481
Summary
−3
−2.5
−2
−1.5
−1
−0.5
0
Standardized $log_2FC$
SLC6A6
GSE16844
GSE74481
Summary
0
0.5
1
1.5
2
2.5
3
3.5
Standardized $log_2FC$

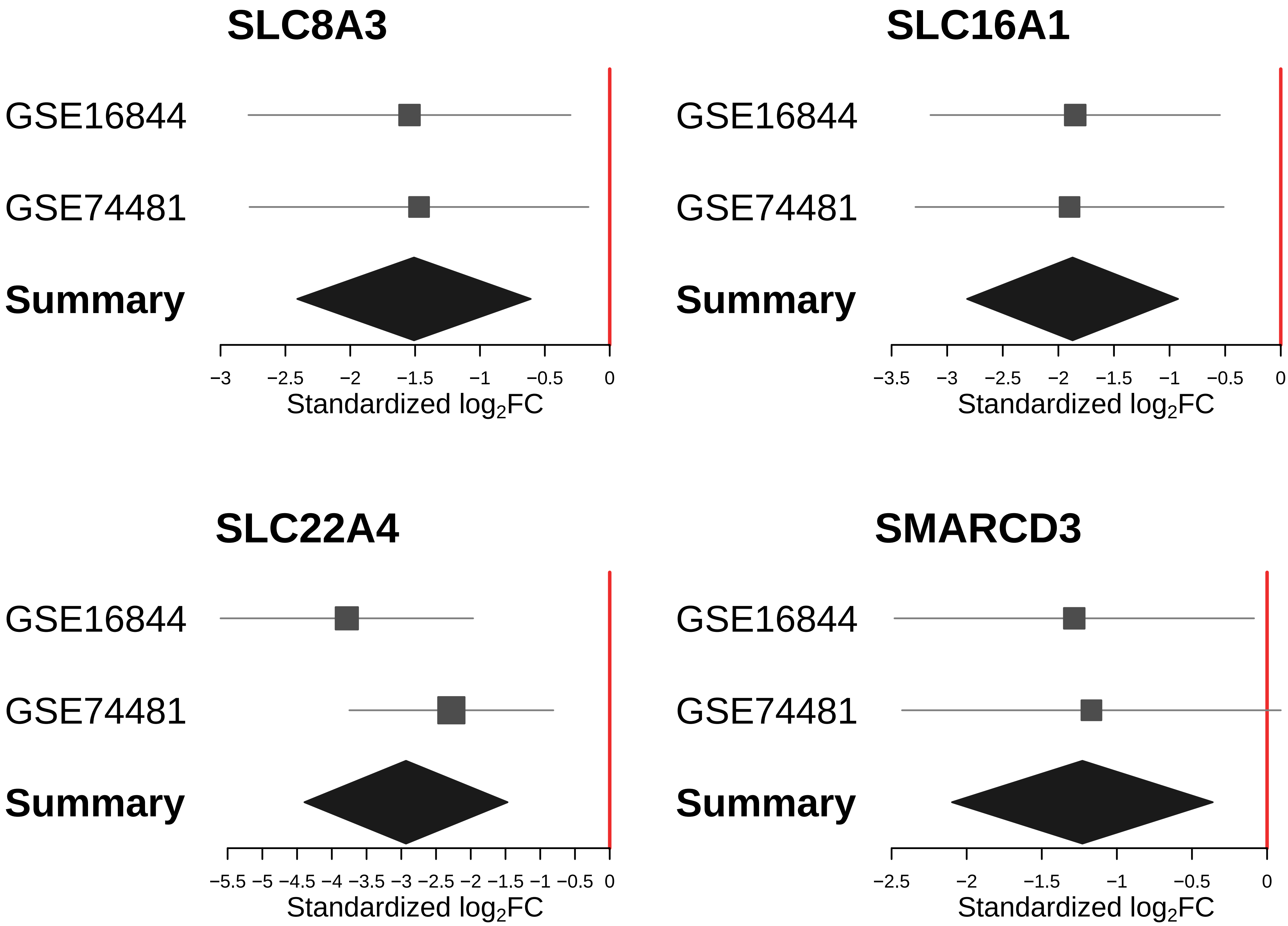
SLC8A3
GSE16844
GSE74481
Summary
−3 −2.5 −2 −1.5 −1 −0.5 0
Standardized $log_2FC$
SLC16A1
GSE16844
GSE74481
Summary
−3.5 −3 −2.5 −2 −1.5 −1 −0.5 0
Standardized $log_2FC$
SLC22A4
GSE16844
GSE74481
Summary
−5.5 −5 −4.5 −4 −3.5 −3 −2.5 −2 −1.5 −1 −0.5 0
Standardized $log_2FC$
SMARCD3
GSE16844
GSE74481
Summary
−2.5 −2 −1.5 −1 −0.5 0
Standardized $log_2FC$

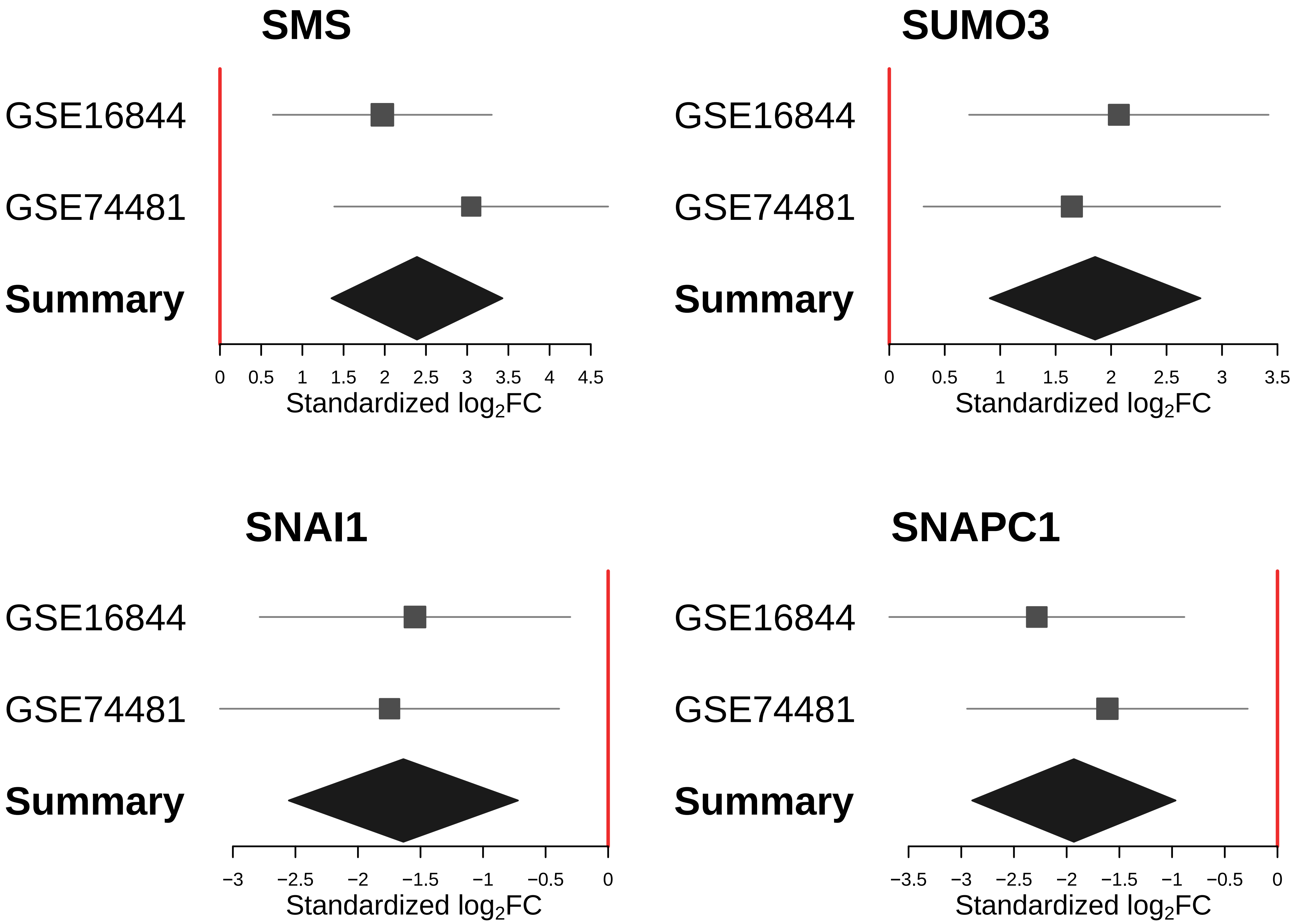
SMS
GSE16844
GSE74481
Summary
0 0.5 1 1.5 2 2.5 3 3.5 4 4.5
Standardized log2FC
SUMO3
GSE16844
GSE74481
Summary
0 0.5 1 1.5 2 2.5 3 3.5
Standardized log2FC
SNAI1
GSE16844
GSE74481
Summary
−3 −2.5 −2 −1.5 −1 −0.5 0
Standardized log2FC
SNAPC1
GSE16844
GSE74481
Summary
−3.5 −3 −2.5 −2 −1.5 −1 −0.5 0
Standardized log2FC

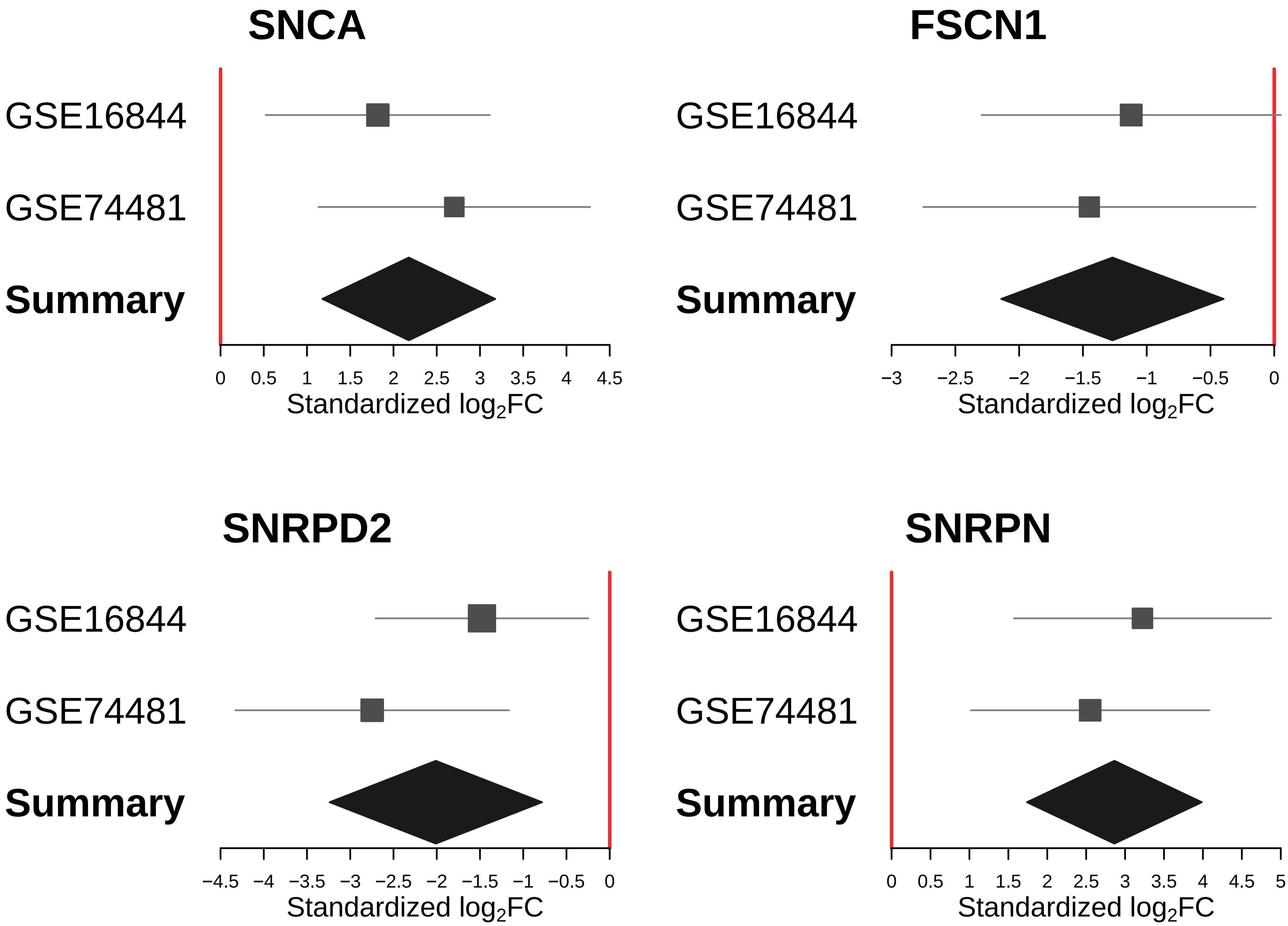
SNCA
GSE16844
GSE74481
Summary
0 0.5 1 1.5 2 2.5 3 3.5 4 4.5
Standardized log2FC
FSCN1
GSE16844
GSE74481
Summary
−3 −2.5 −2 −1.5 −1 −0.5 0
Standardized log2FC
SNRPD2
GSE16844
GSE74481
Summary
−4.5 −4 −3.5 −3 −2.5 −2 −1.5 −1 −0.5 0
Standardized log2FC
SNRPN
GSE16844
GSE74481
Summary
0 0.5 1 1.5 2 2.5 3 3.5 4 4.5 5
Standardized log2FC

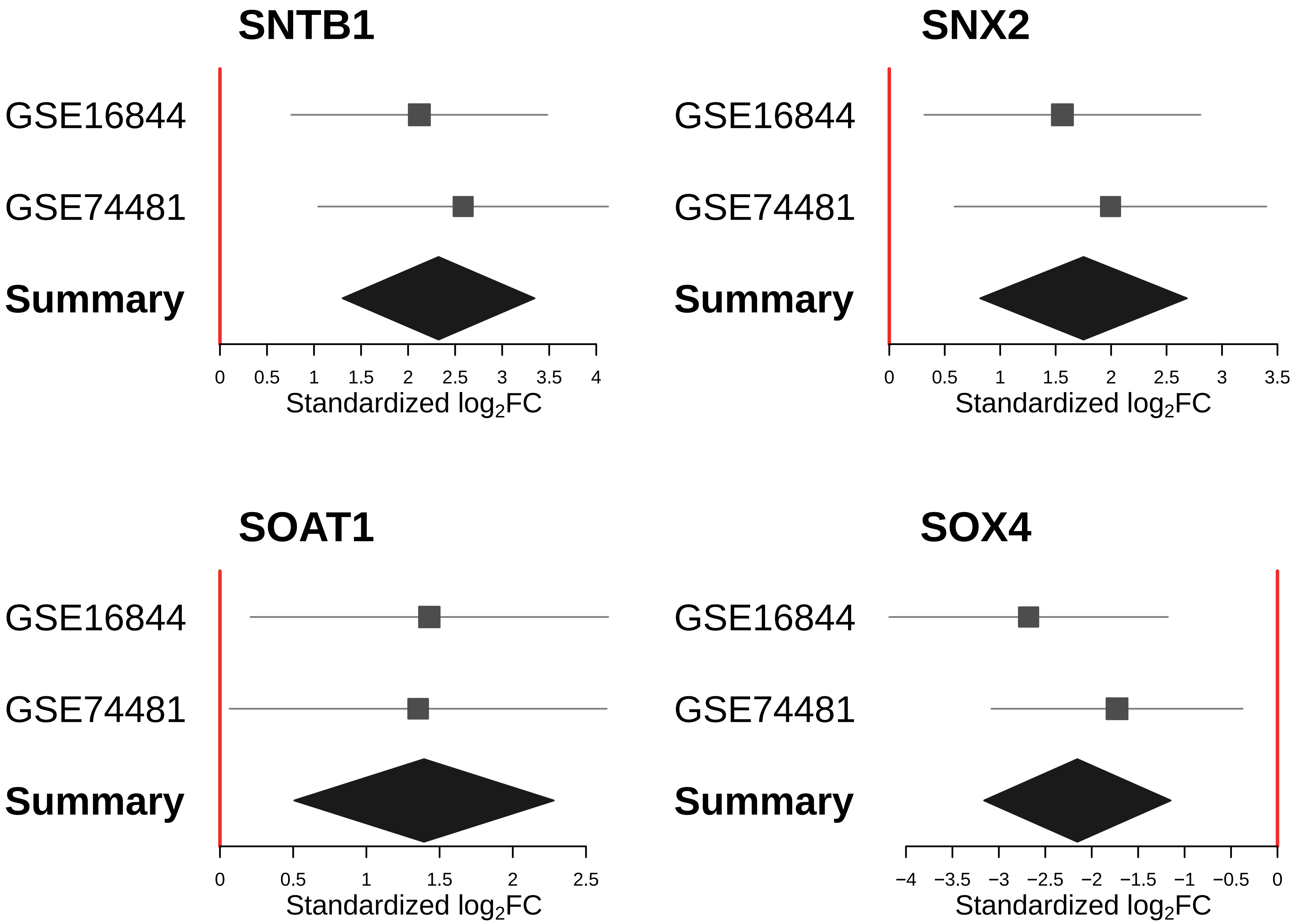
SNTB1
GSE16844
GSE74481
Summary
0 0.5 1 1.5 2 2.5 3 3.5 4
Standardized log2FC
SNX2
GSE16844
GSE74481
Summary
0 0.5 1 1.5 2 2.5 3 3.5
Standardized log2FC
SOAT1
GSE16844
GSE74481
Summary
0 0.5 1 1.5 2 2.5
Standardized log2FC
SOX4
GSE16844
GSE74481
Summary
−4 −3.5 −3 −2.5 −2 −1.5 −1 −0.5 0
Standardized log2FC

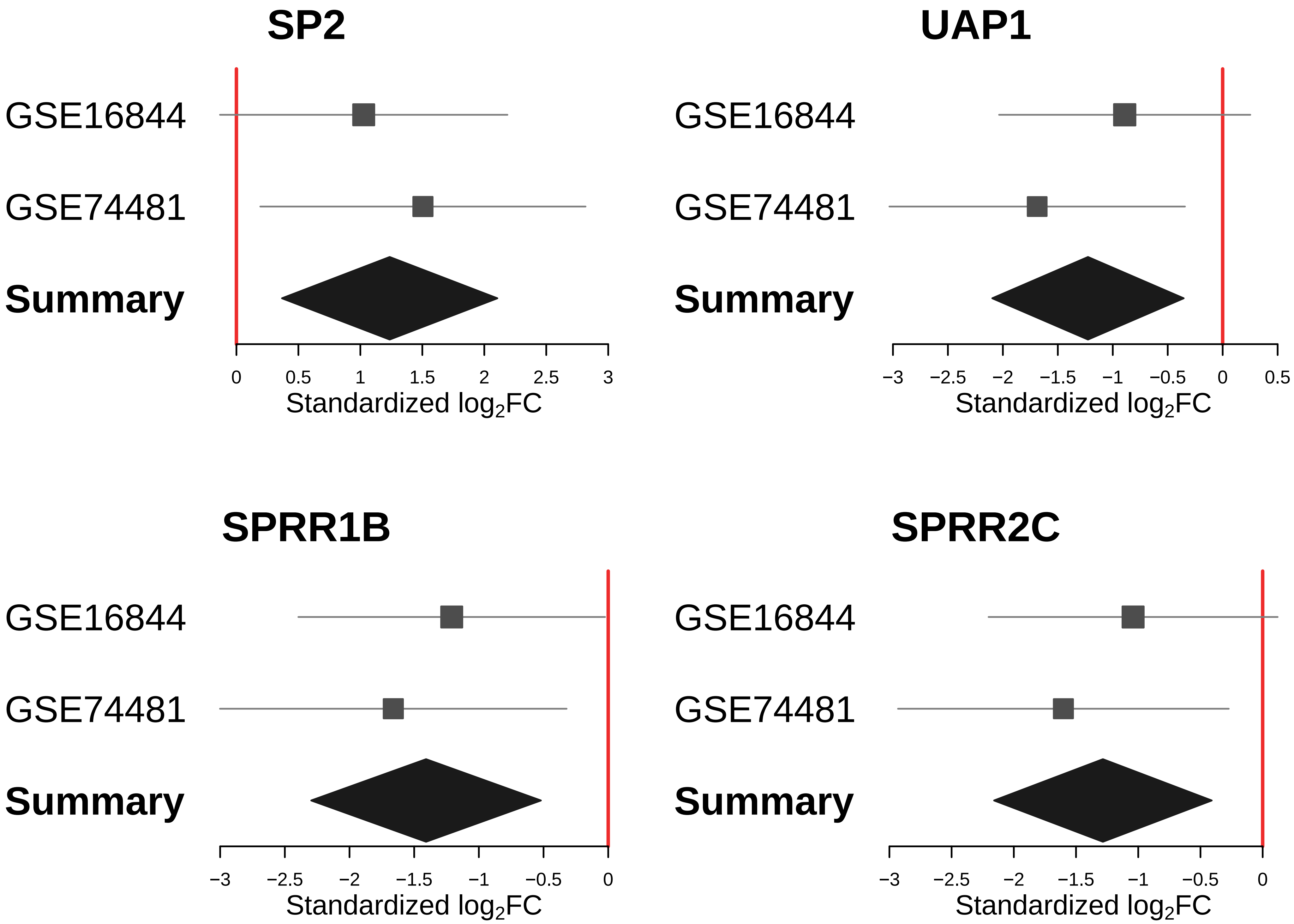
SP2
GSE16844
GSE74481
Summary
0
0.5
1
1.5
2
2.5
3
Standardized log2FC
UAP1
GSE16844
GSE74481
Summary
−3
−2.5
−2
−1.5
−1
−0.5
0
0.5
Standardized log2FC
SPRR1B
GSE16844
GSE74481
Summary
−3
−2.5
−2
−1.5
−1
−0.5
0
Standardized log2FC
SPRR2C
GSE16844
GSE74481
Summary
−3
−2.5
−2
−1.5
−1
−0.5
0
Standardized log2FC

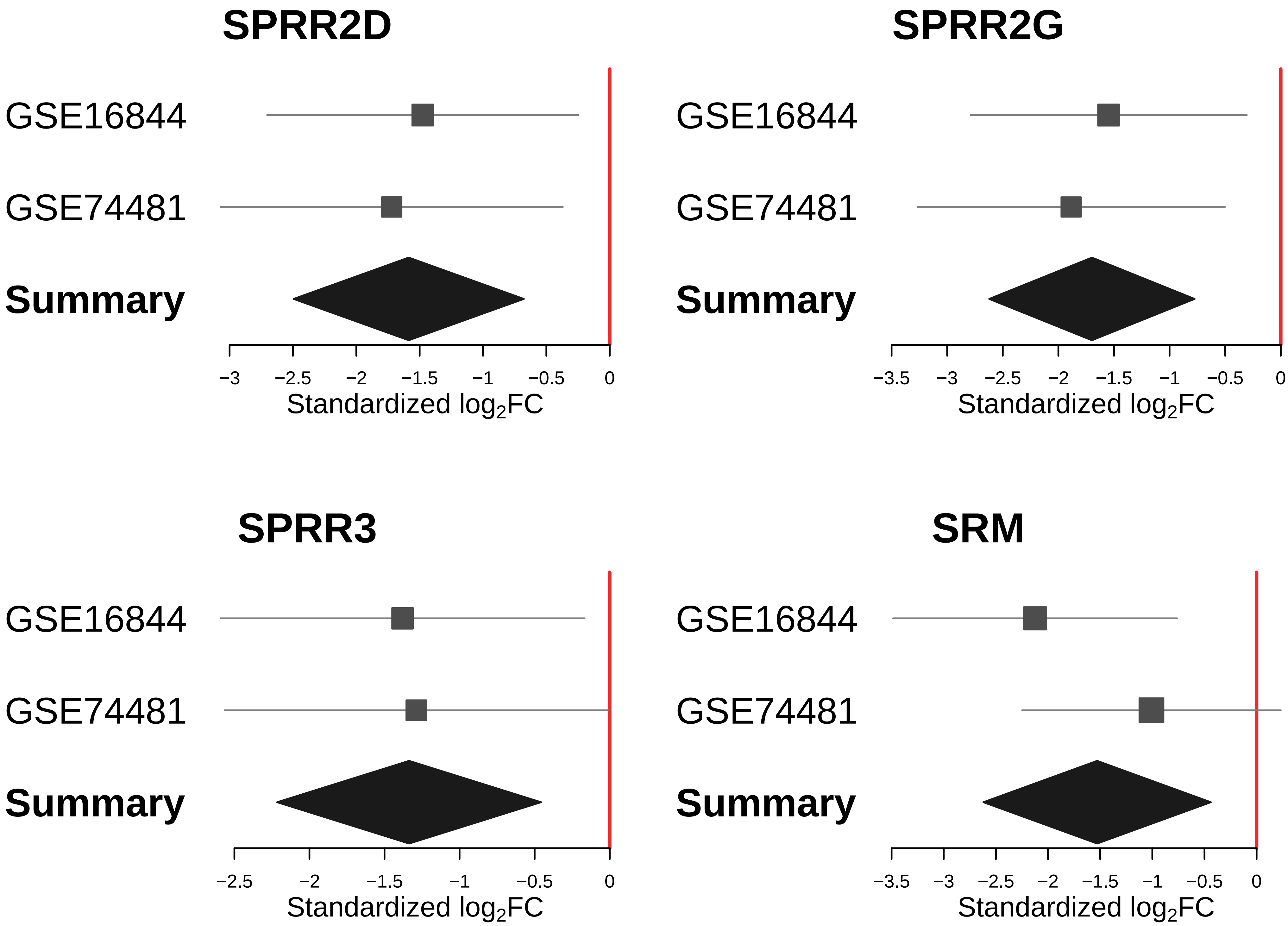
SPRR2D
GSE16844
GSE74481
Summary
−3 −2.5 −2 −1.5 −1 −0.5 0
Standardized log2FC
SPRR2G
GSE16844
GSE74481
Summary
−3.5 −3 −2.5 −2 −1.5 −1 −0.5 0
Standardized log2FC
SPRR3
GSE16844
GSE74481
Summary
−2.5 −2 −1.5 −1 −0.5 0
Standardized log2FC
SRM
GSE16844
GSE74481
Summary
−3.5 −3 −2.5 −2 −1.5 −1 −0.5 0
Standardized log2FC

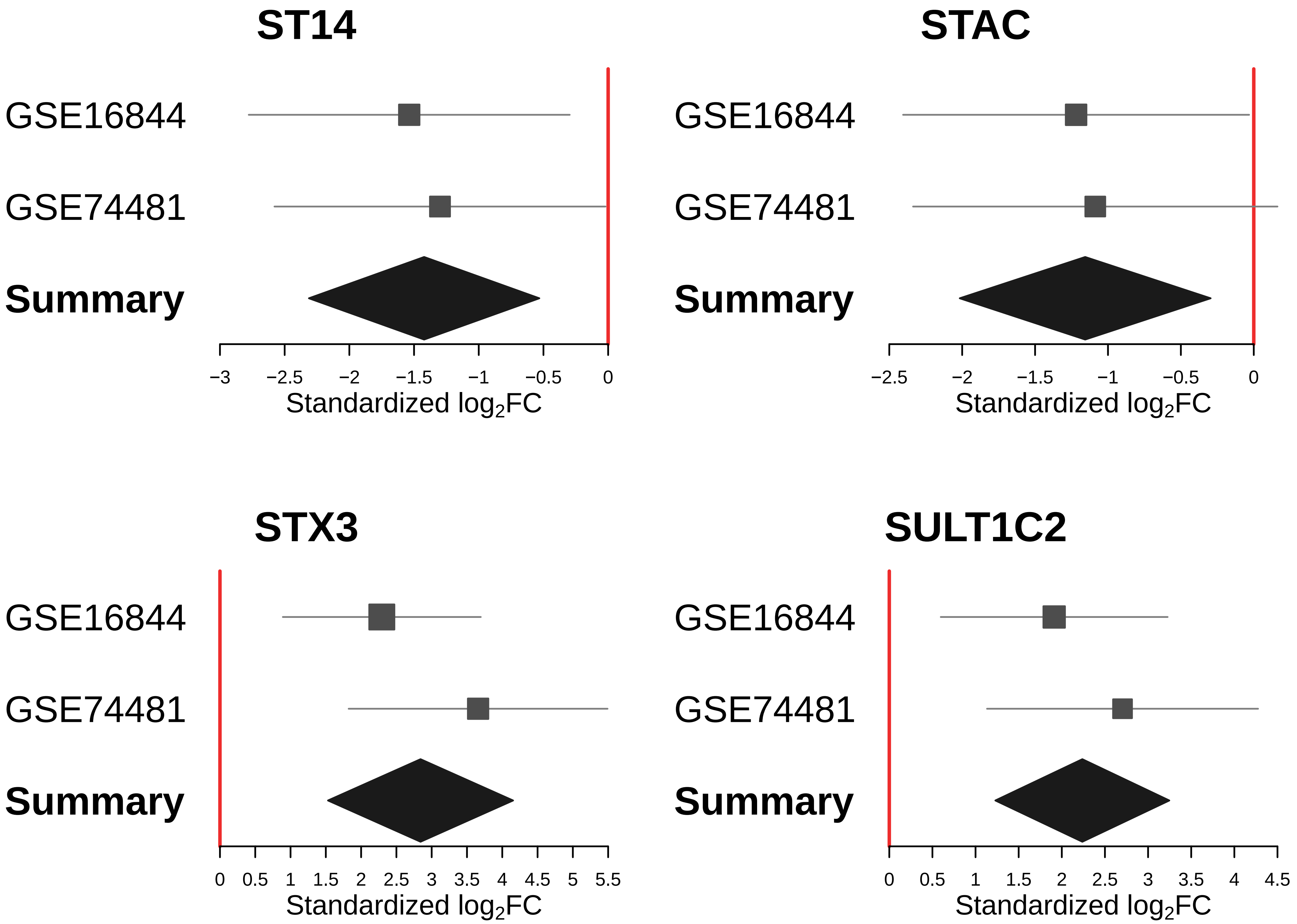

ST14
GSE16844
GSE74481
Summary
−3
−2.5
−2
−1.5
−1
−0.5
0
Standardized $\log_2FC$
STAC
GSE16844
GSE74481
Summary
−2.5
−2
−1.5
−1
−0.5
0
Standardized $\log_2FC$
STX3
GSE16844
GSE74481
Summary
0
0.5
1
1.5
2
2.5
3
3.5
4
4.5
5
5.5
Standardized $\log_2FC$
SULT1C2
GSE16844
GSE74481
Summary
0
0.5
1
1.5
2
2.5
3
3.5
4
4.5
Standardized $\log_2FC$

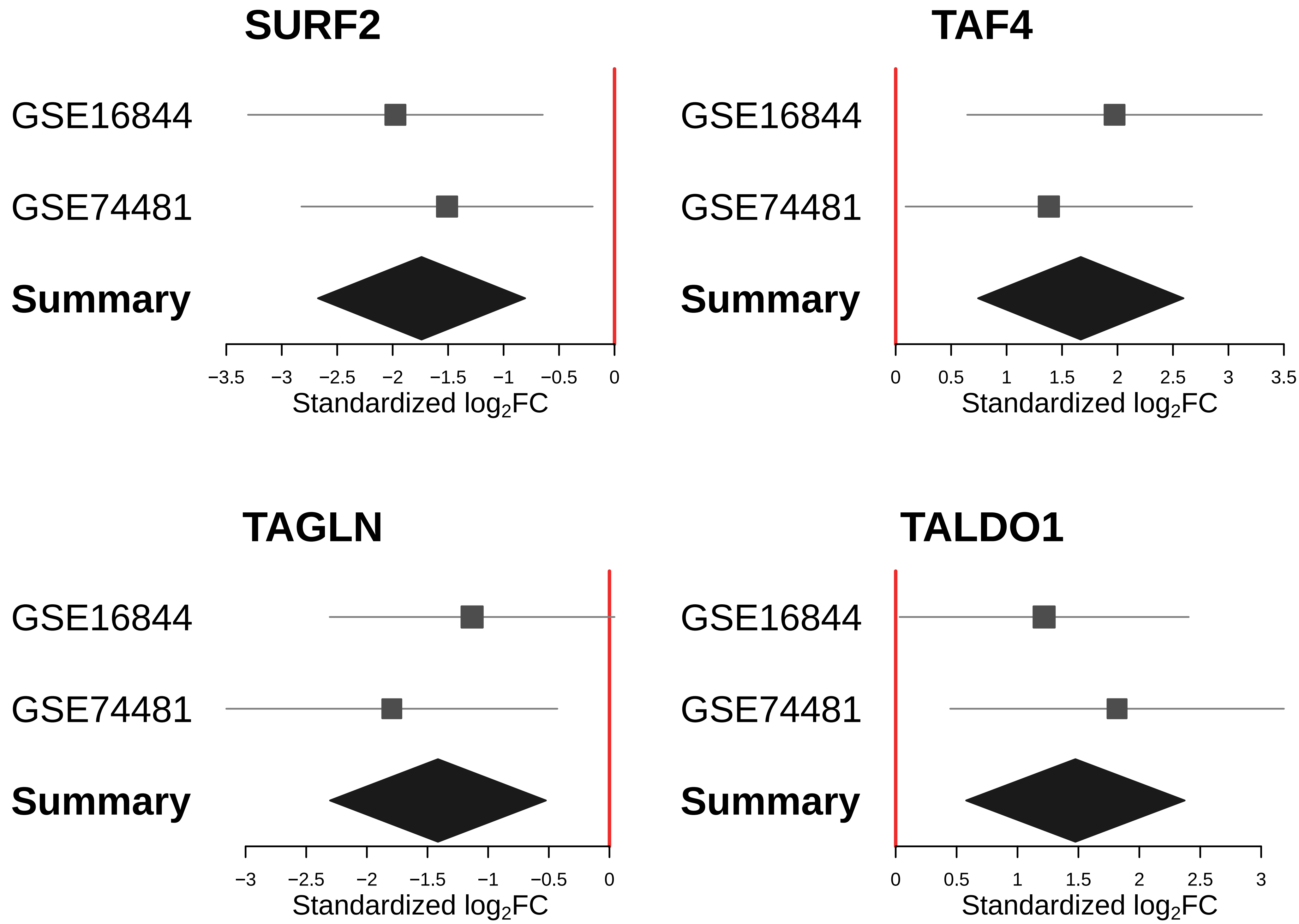
SURF2
GSE16844
GSE74481
Summary
−3.5 −3 −2.5 −2 −1.5 −1 −0.5 0
Standardized log2FC
TAF4
GSE16844
GSE74481
Summary
0 0.5 1 1.5 2 2.5 3 3.5
Standardized log2FC
TAGLN
GSE16844
GSE74481
Summary
−3 −2.5 −2 −1.5 −1 −0.5 0
Standardized log2FC
TALDO1
GSE16844
GSE74481
Summary
0 0.5 1 1.5 2 2.5 3
Standardized log2FC

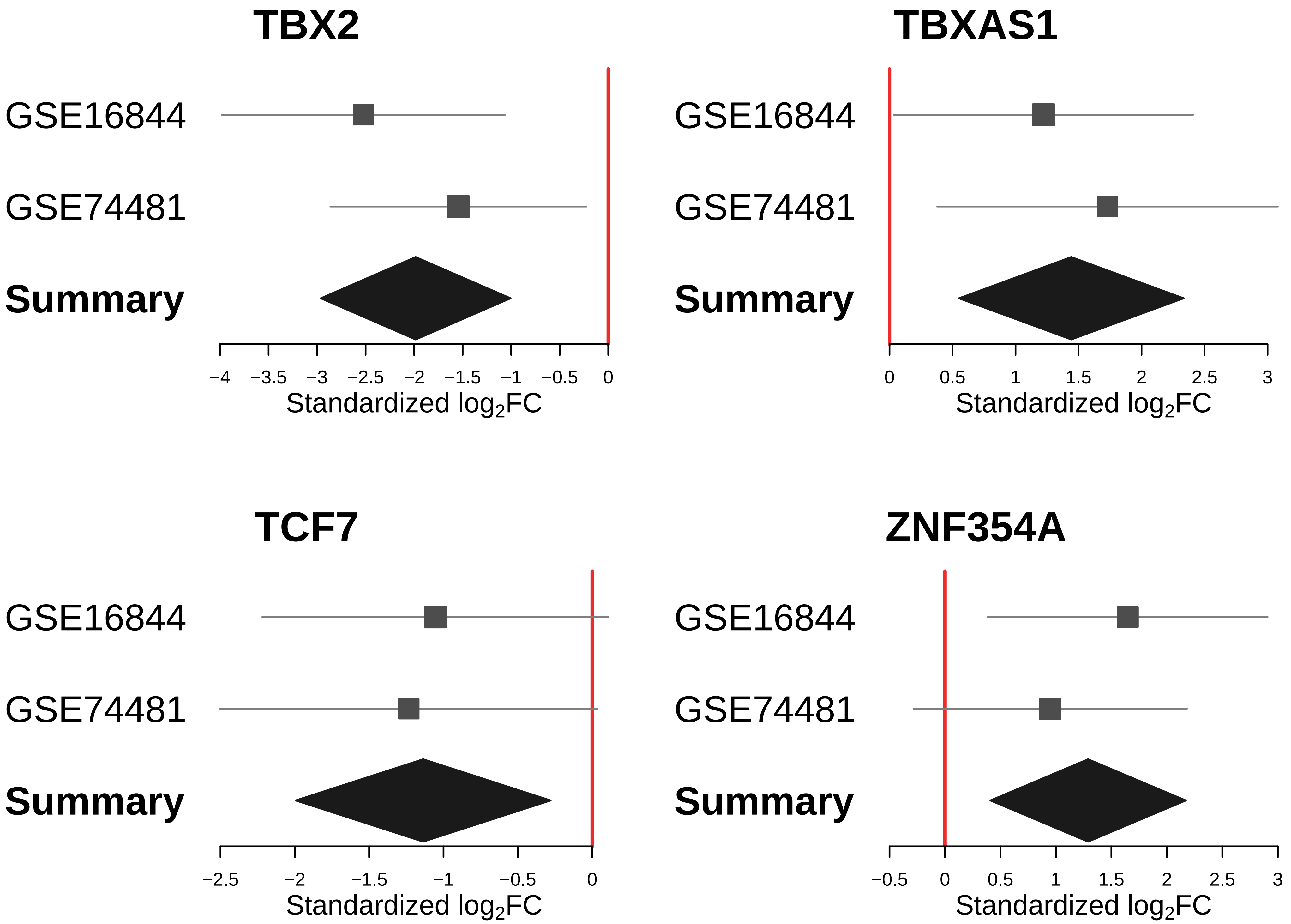
TBX2
GSE16844
GSE74481
Summary
−4 −3.5 −3 −2.5 −2 −1.5 −1 −0.5 0
Standardized log2FC
TBXAS1
GSE16844
GSE74481
Summary
0 0.5 1 1.5 2 2.5 3
Standardized log2FC
TCF7
GSE16844
GSE74481
Summary
−2.5 −2 −1.5 −1 −0.5 0
Standardized log2FC
ZNF354A
GSE16844
GSE74481
Summary
−0.5 0 0.5 1 1.5 2 2.5 3
Standardized log2FC

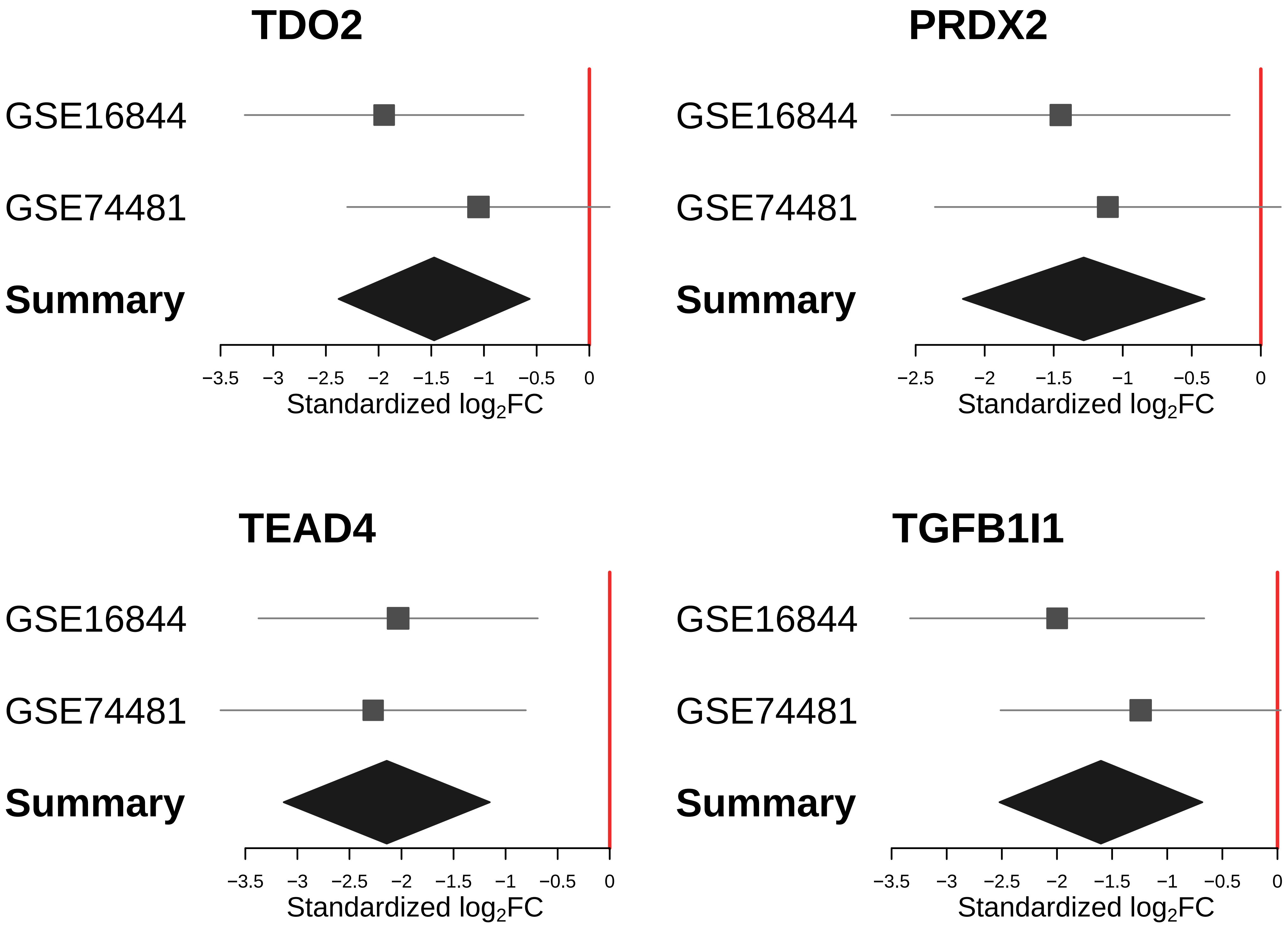
TDO2
PRDX2
TEAD4
TGFB1I1
GSE16844
GSE74481
Summary
−3.5 −3 −2.5 −2 −1.5 −1 −0.5 0
−2.5 −2 −1.5 −1 −0.5 0
Standardized $log_2FC$

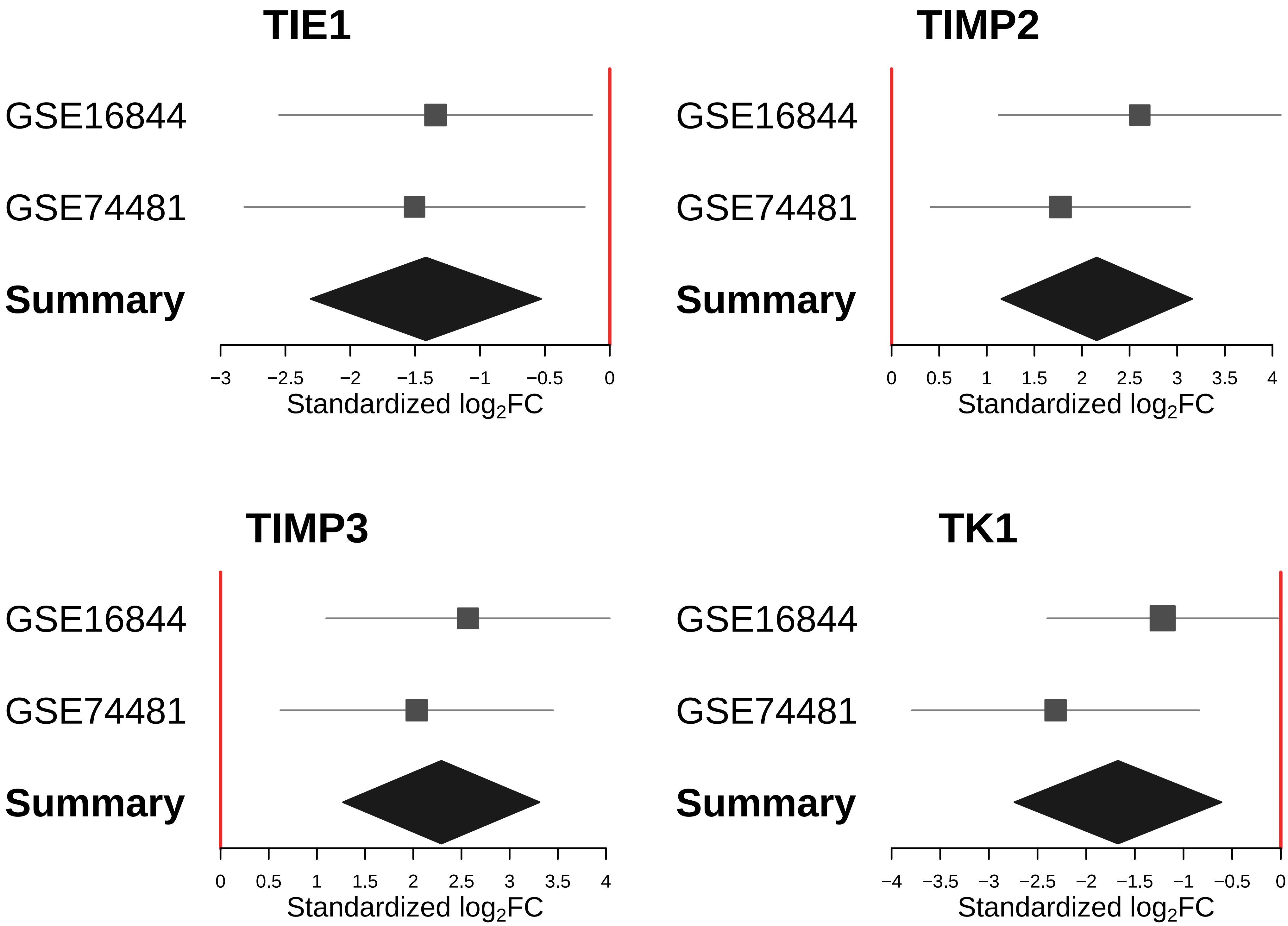
TIE1
GSE16844
GSE74481
Summary
−3 −2.5 −2 −1.5 −1 −0.5 0
Standardized log2FC
TIMP2
GSE16844
GSE74481
Summary
0 0.5 1 1.5 2 2.5 3 3.5 4
Standardized log2FC
TIMP3
GSE16844
GSE74481
Summary
0 0.5 1 1.5 2 2.5 3 3.5 4
Standardized log2FC
TK1
GSE16844
GSE74481
Summary
−4 −3.5 −3 −2.5 −2 −1.5 −1 −0.5 0
Standardized log2FC

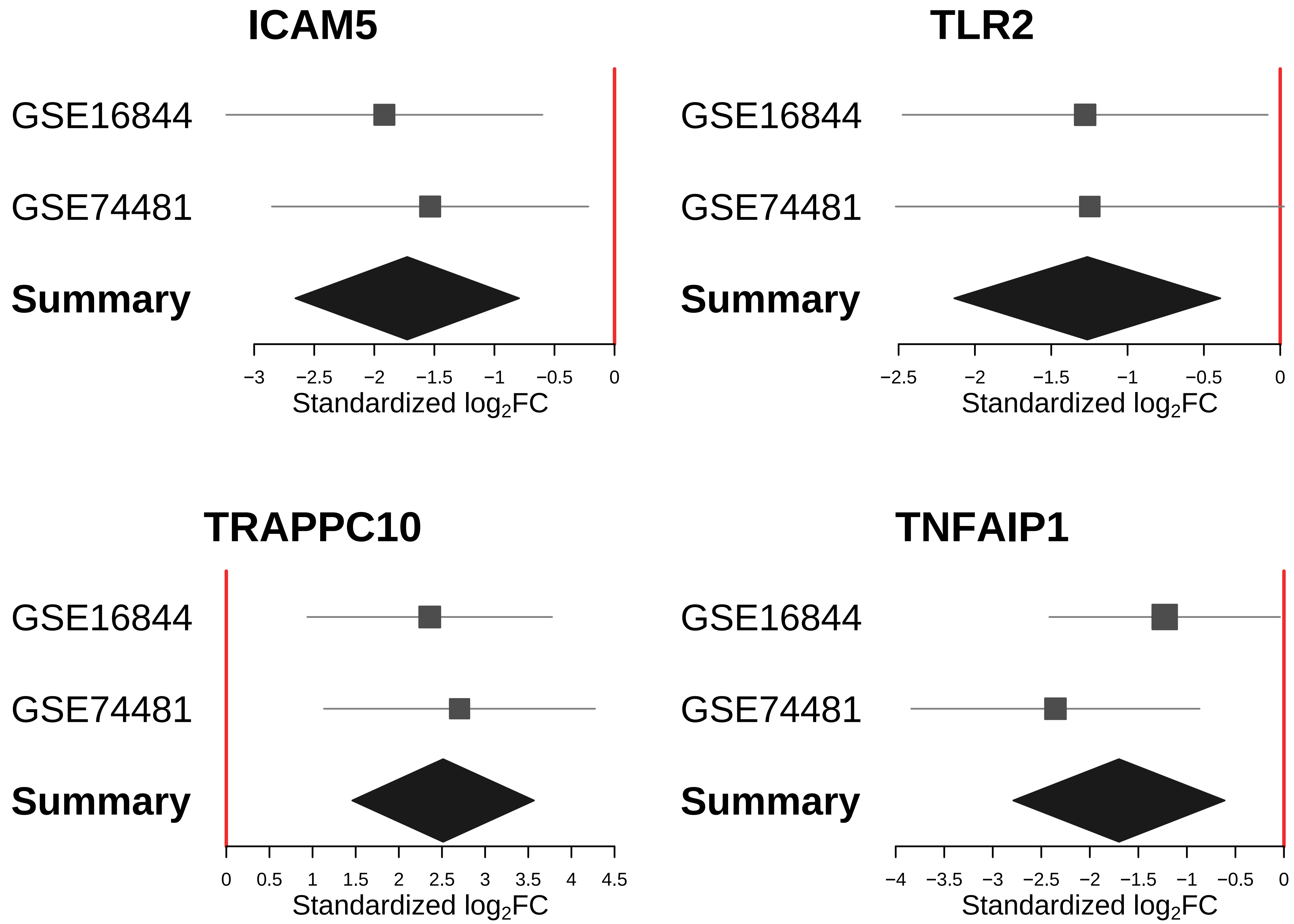

ICAM5
GSE16844
GSE74481
Summary
−3 −2.5 −2 −1.5 −1 −0.5 0
Standardized log2FC
TLR2
GSE16844
GSE74481
Summary
−2.5 −2 −1.5 −1 −0.5 0
Standardized log2FC
TRAPPC10
GSE16844
GSE74481
Summary
0 0.5 1 1.5 2 2.5 3 3.5 4 4.5
Standardized log2FC
TNFAIP1
GSE16844
GSE74481
Summary
−4 −3.5 −3 −2.5 −2 −1.5 −1 −0.5 0
Standardized log2FC

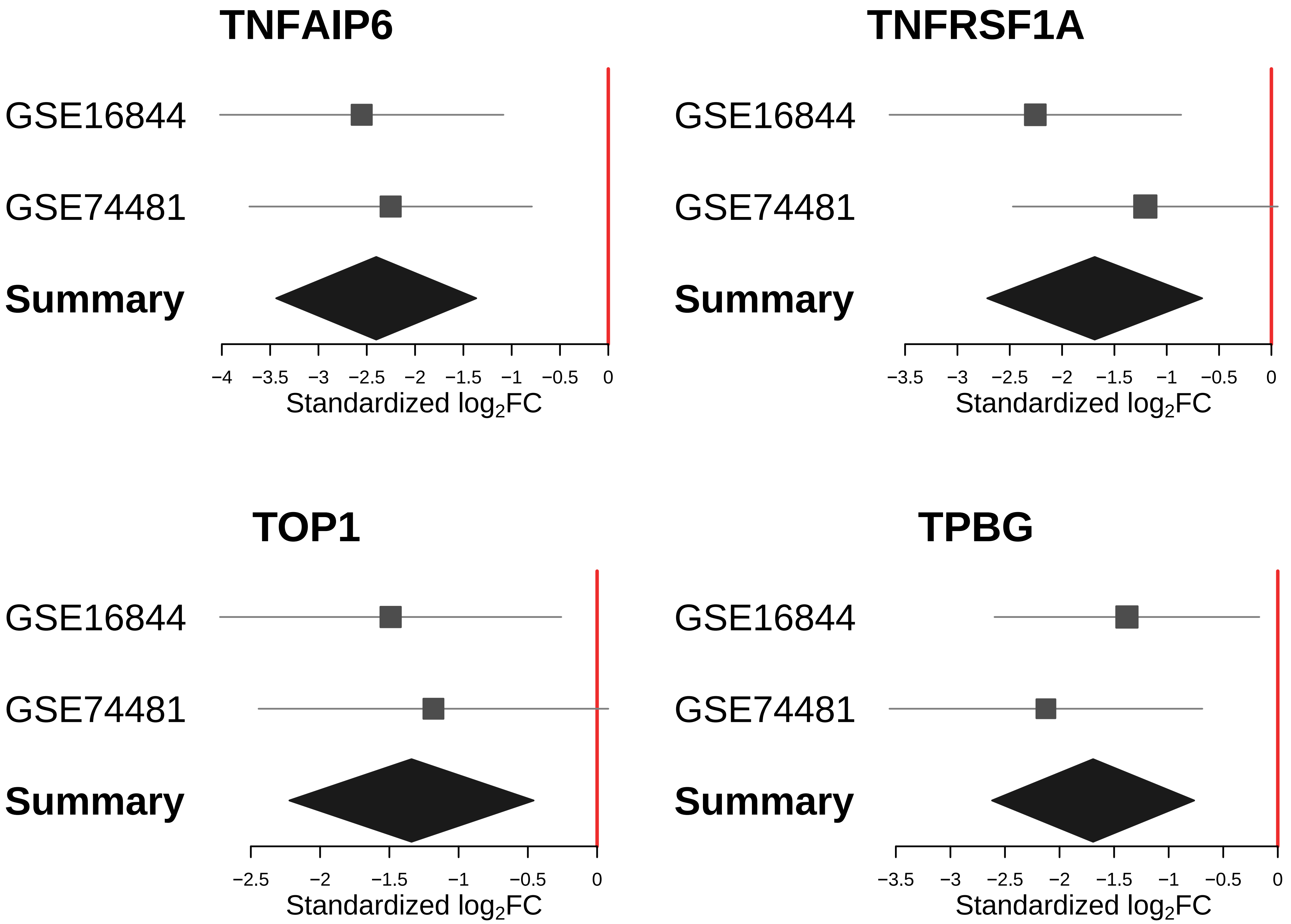
TNFAIP6
GSE16844
GSE74481
Summary
−4 −3.5 −3 −2.5 −2 −1.5 −1 −0.5 0
Standardized log2FC
TNFRSF1A
GSE16844
GSE74481
Summary
−3.5 −3 −2.5 −2 −1.5 −1 −0.5 0
Standardized log2FC
TOP1
GSE16844
GSE74481
Summary
−2.5 −2 −1.5 −1 −0.5 0
Standardized log2FC
TPBG
GSE16844
GSE74481
Summary
−3.5 −3 −2.5 −2 −1.5 −1 −0.5 0
Standardized log2FC

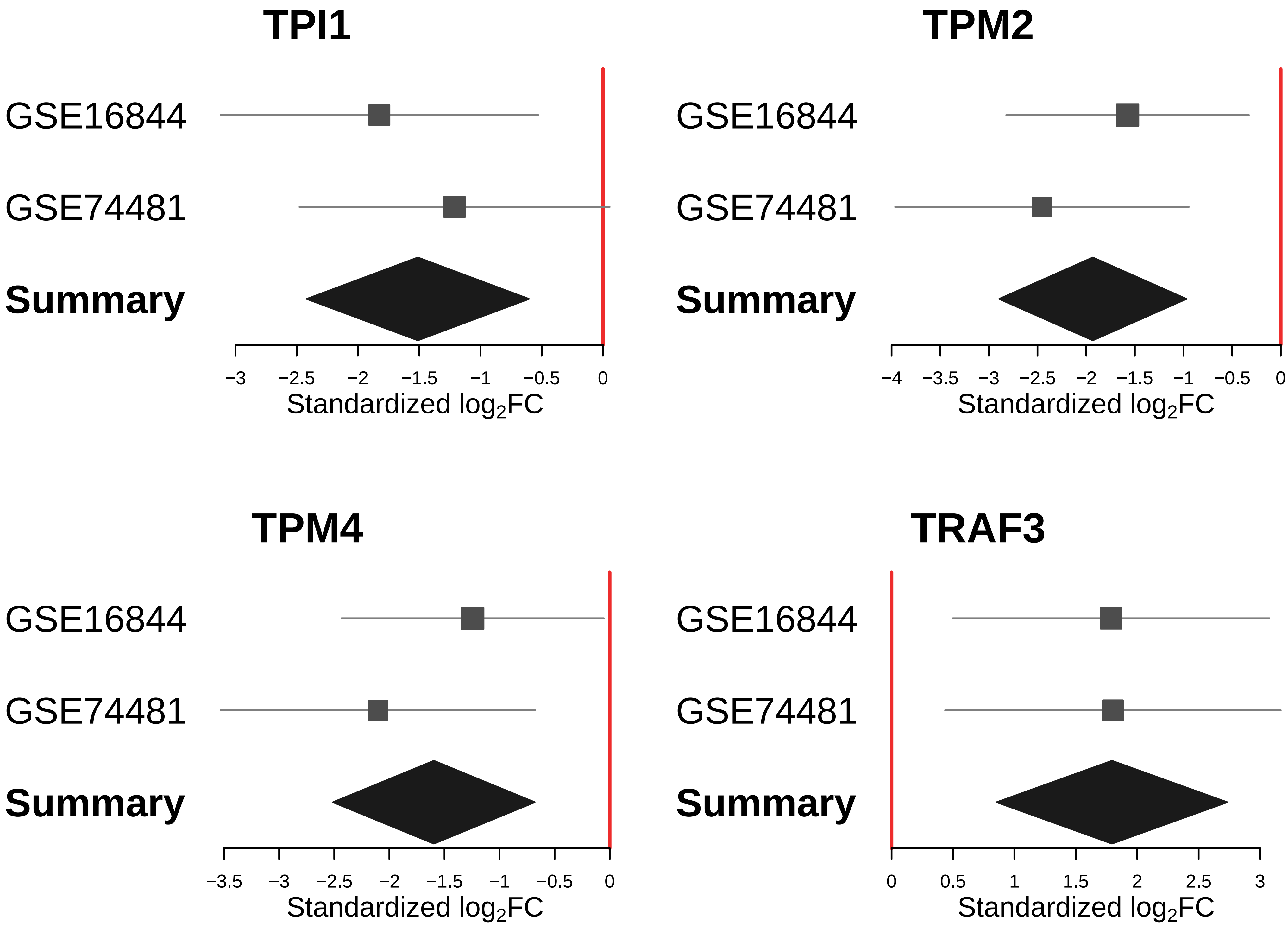
TPI1
GSE16844
GSE74481
Summary
−3
−2.5
−2
−1.5
−1
−0.5
0
Standardized log2FC
TPM2
GSE16844
GSE74481
Summary
−4
−3.5
−3
−2.5
−2
−1.5
−1
−0.5
0
Standardized log2FC
TPM4
GSE16844
GSE74481
Summary
−3.5
−3
−2.5
−2
−1.5
−1
−0.5
0
Standardized log2FC
TRAF3
GSE16844
GSE74481
Summary
0
0.5
1
1.5
2
2.5
3
Standardized log2FC

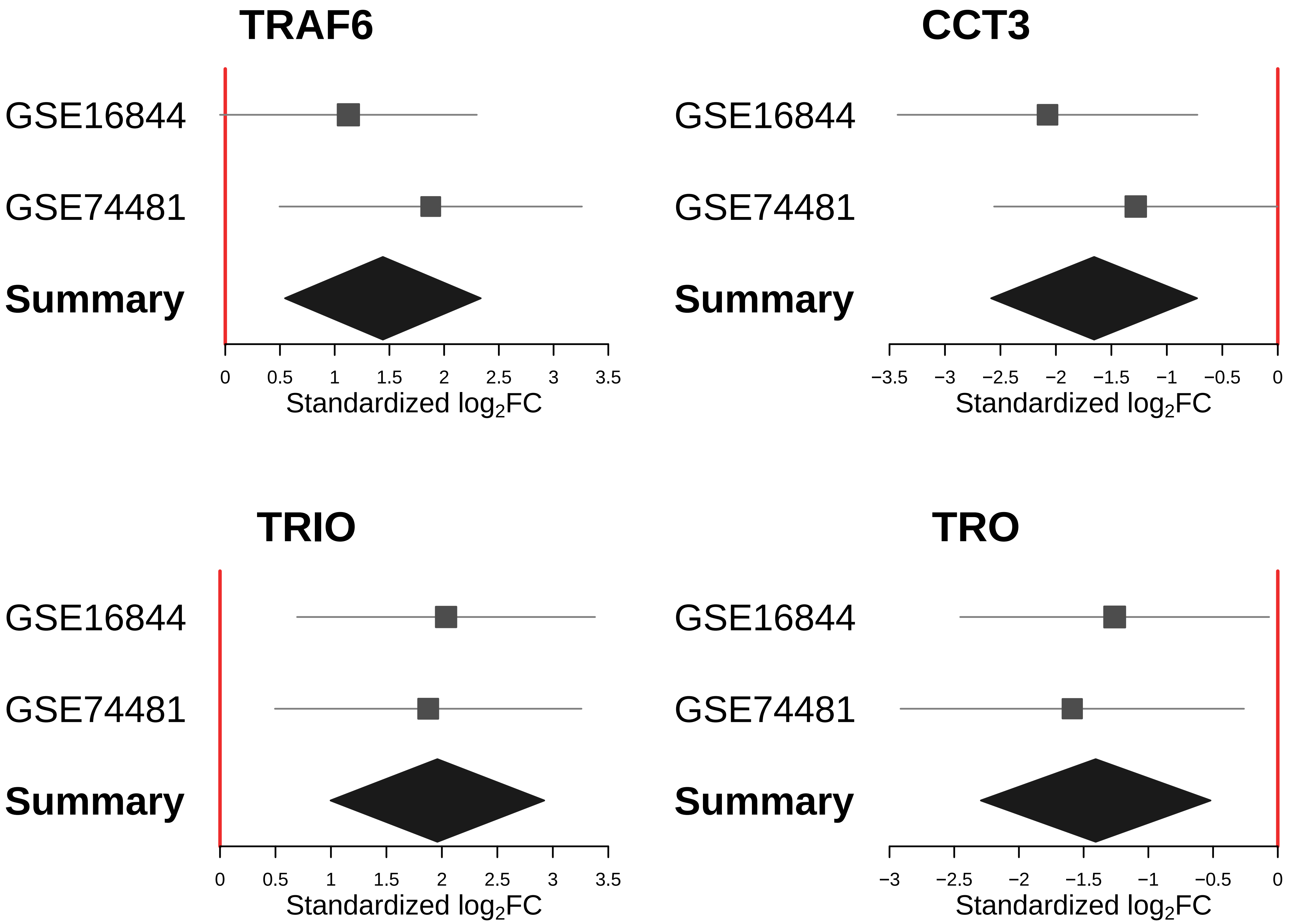
TRAF6
GSE16844
GSE74481
Summary
0
0.5
1
1.5
2
2.5
3
3.5
Standardized log2FC
CCT3
GSE16844
GSE74481
Summary
−3.5
−3
−2.5
−2
−1.5
−1
−0.5
0
Standardized log2FC
TRIO
GSE16844
GSE74481
Summary
0
0.5
1
1.5
2
2.5
3
3.5
Standardized log2FC
TRO
GSE16844
GSE74481
Summary
−3
−2.5
−2
−1.5
−1
−0.5
0
Standardized log2FC

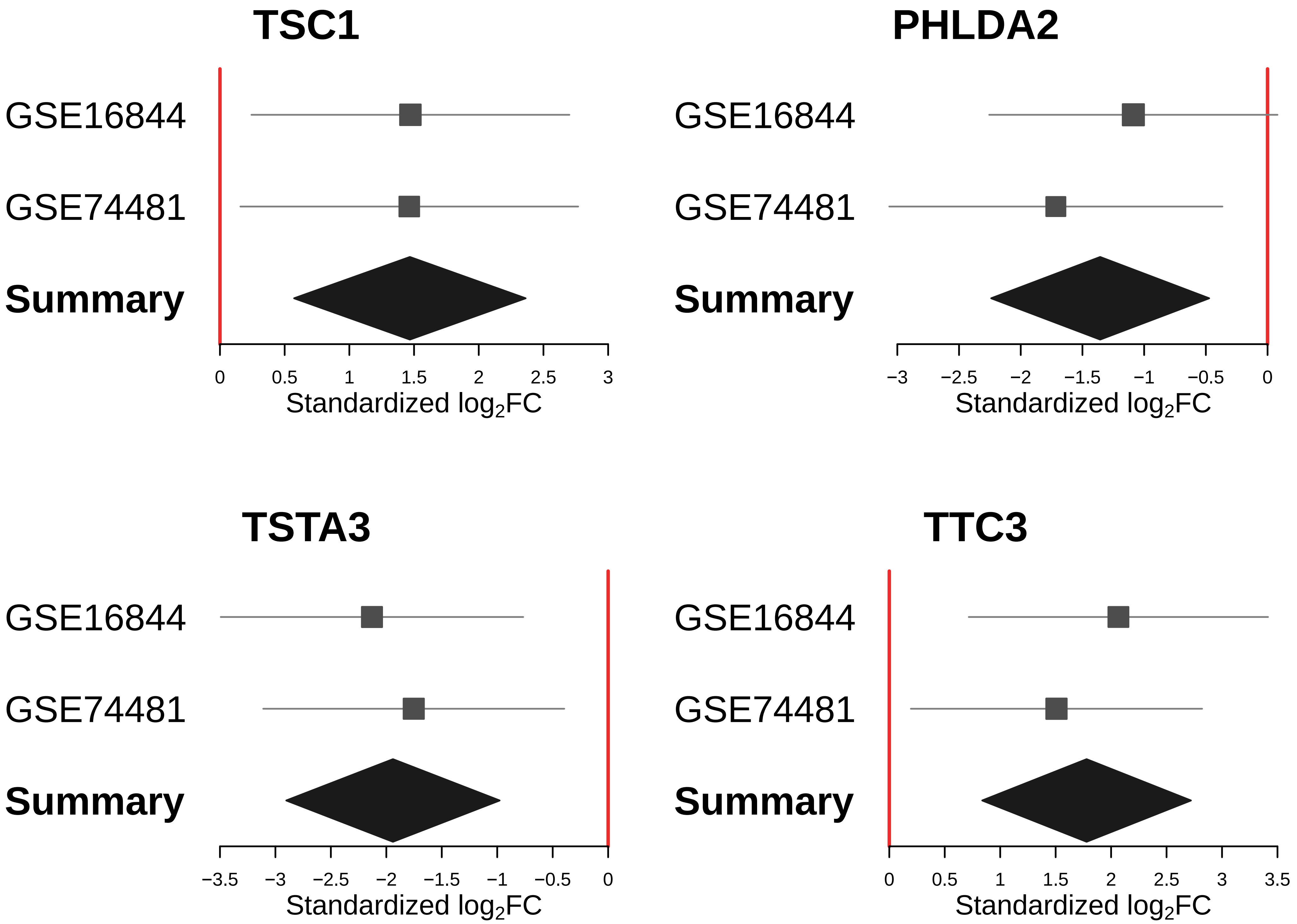
TSC1
GSE16844
GSE74481
Summary
0 0.5 1 1.5 2 2.5 3
Standardized log2FC
PHLDA2
GSE16844
GSE74481
Summary
−3 −2.5 −2 −1.5 −1 −0.5 0
Standardized log2FC
TSTA3
GSE16844
GSE74481
Summary
−3.5 −3 −2.5 −2 −1.5 −1 −0.5 0
Standardized log2FC
TTC3
GSE16844
GSE74481
Summary
0 0.5 1 1.5 2 2.5 3 3.5
Standardized log2FC

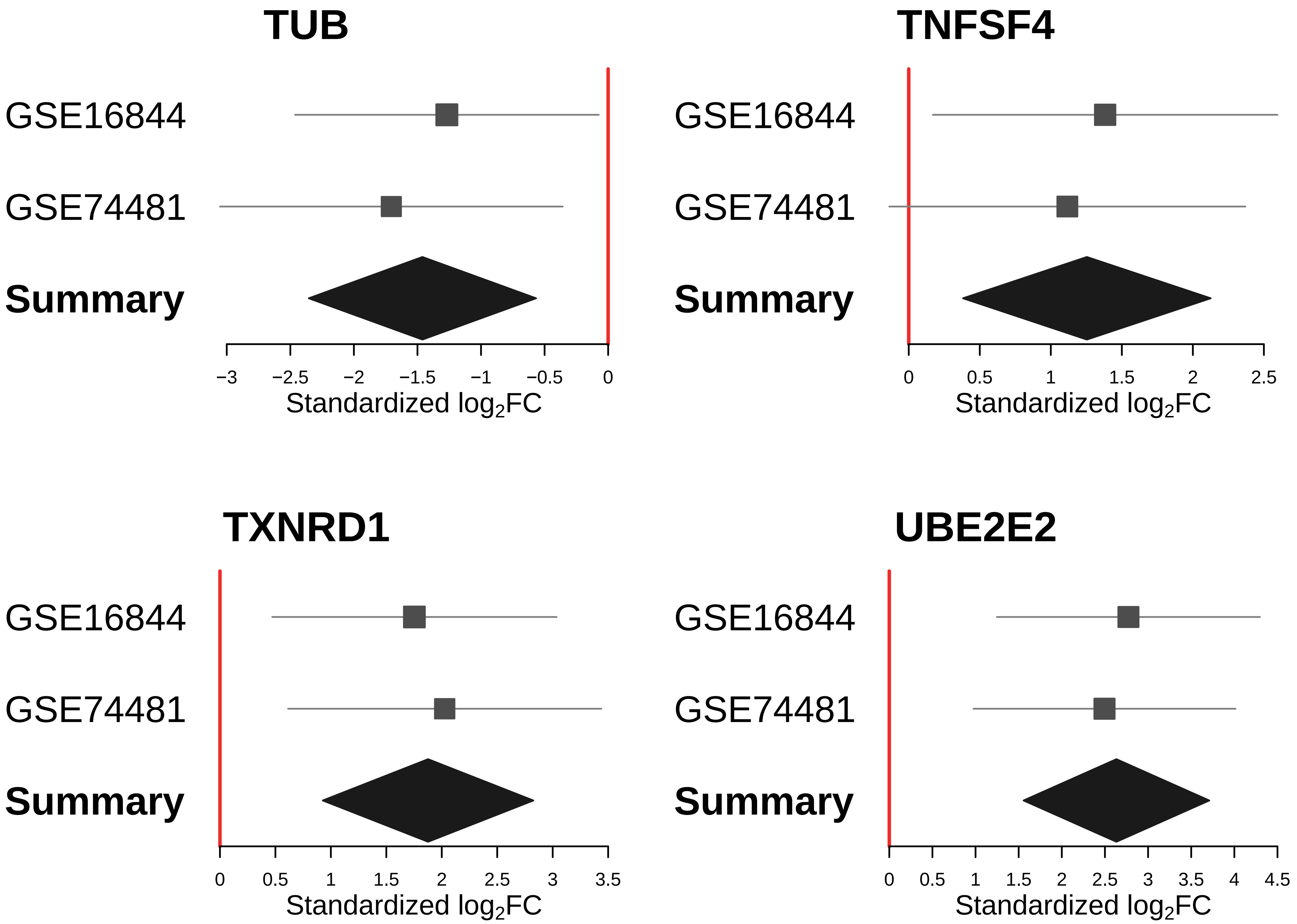

TUB
GSE16844
GSE74481
Summary
−3 −2.5 −2 −1.5 −1 −0.5 0
Standardized log2FC
TNFSF4
GSE16844
GSE74481
Summary
0 0.5 1 1.5 2 2.5
Standardized log2FC
TXNRD1
GSE16844
GSE74481
Summary
0 0.5 1 1.5 2 2.5 3 3.5
Standardized log2FC
UBE2E2
GSE16844
GSE74481
Summary
0 0.5 1 1.5 2 2.5 3 3.5 4 4.5
Standardized log2FC

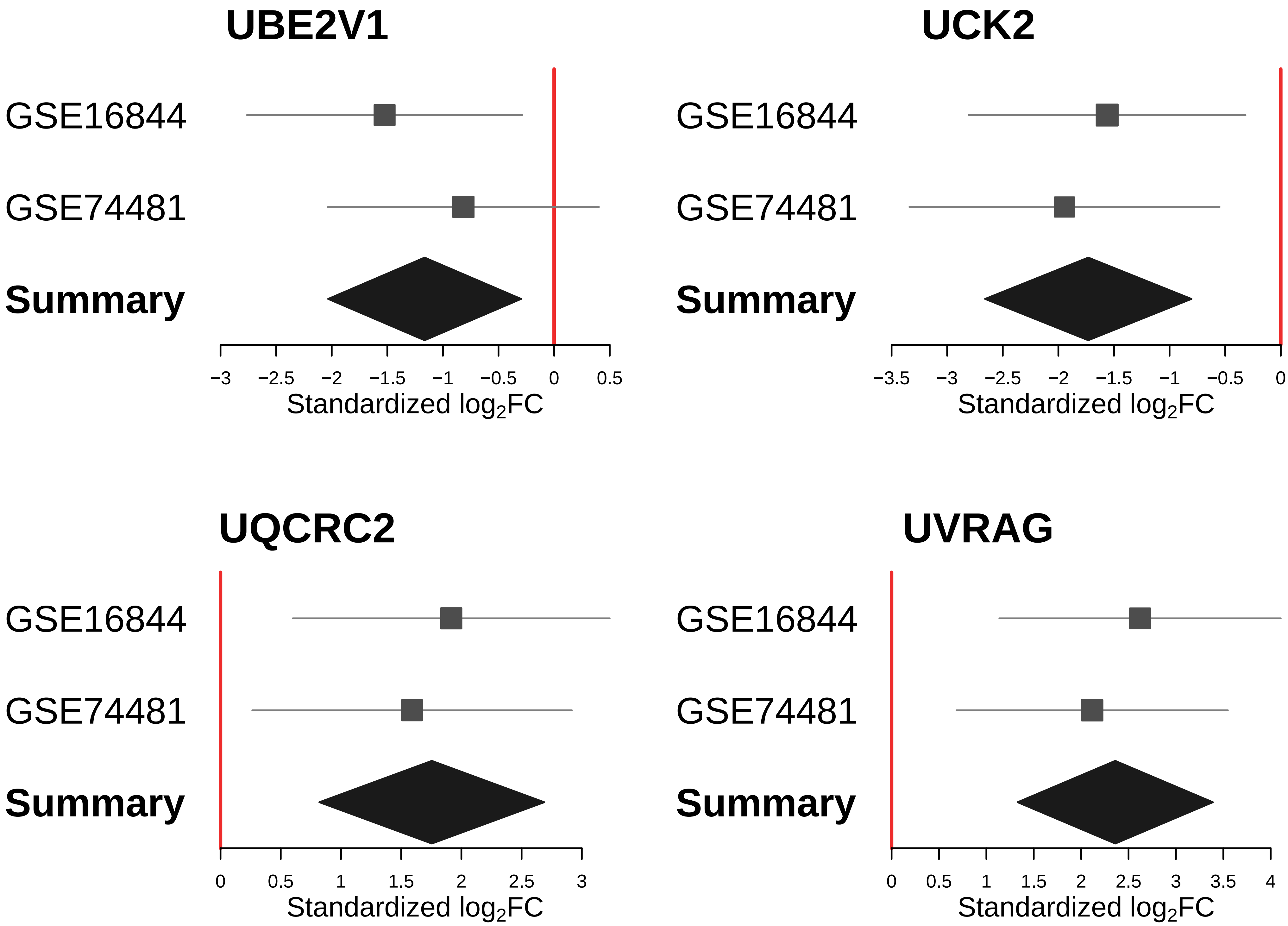
UBE2V1
GSE16844
GSE74481
Summary
−3 −2.5 −2 −1.5 −1 −0.5 0 0.5
Standardized log2FC
UCK2
GSE16844
GSE74481
Summary
−3.5 −3 −2.5 −2 −1.5 −1 −0.5 0
Standardized log2FC
UQCRC2
GSE16844
GSE74481
Summary
0 0.5 1 1.5 2 2.5 3
Standardized log2FC
UVRAG
GSE16844
GSE74481
Summary
0 0.5 1 1.5 2 2.5 3 3.5 4
Standardized log2FC

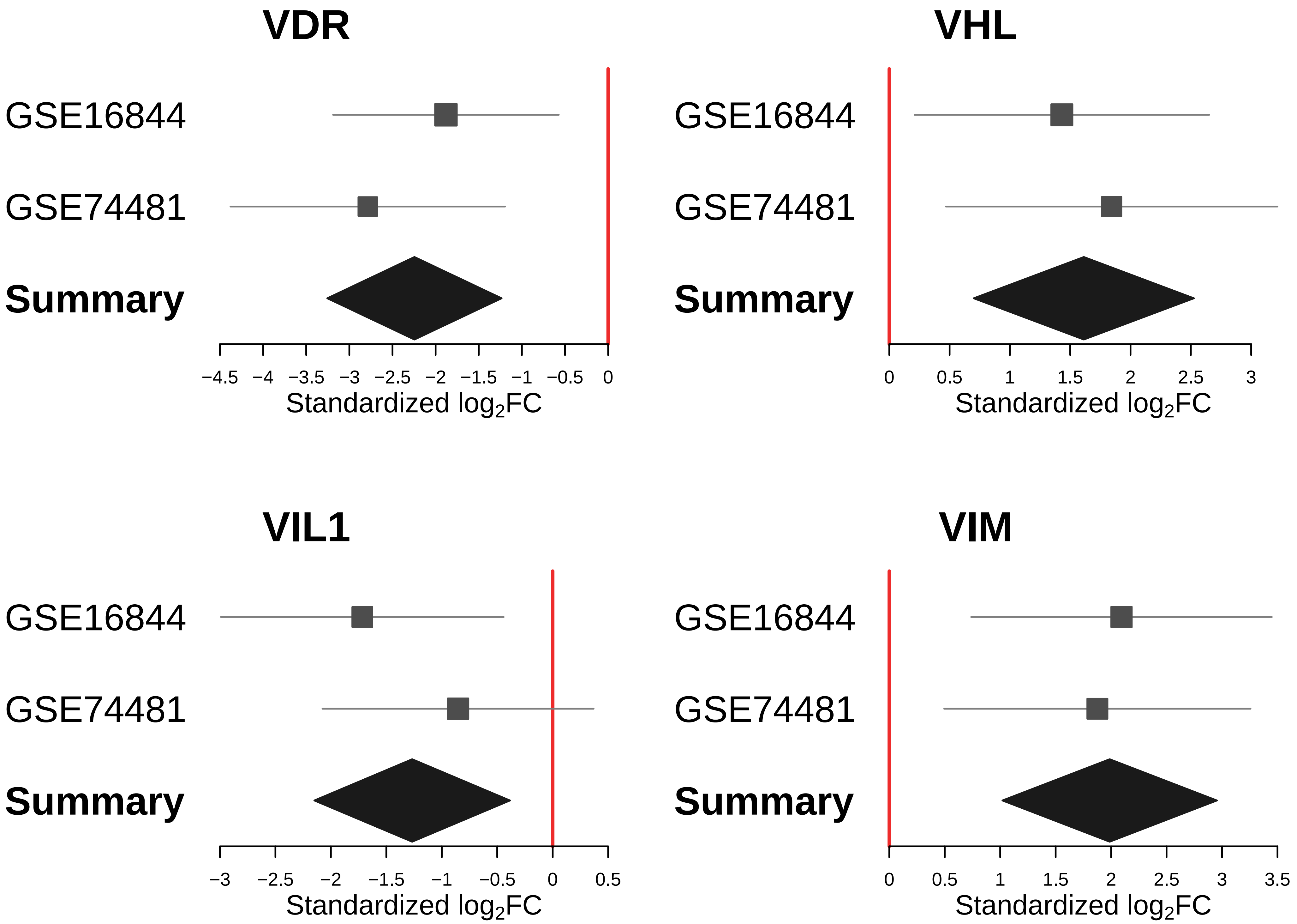
VDR
GSE16844
GSE74481
Summary
−4.5 −4 −3.5 −3 −2.5 −2 −1.5 −1 −0.5 0
Standardized $log_2FC$
VHL
GSE16844
GSE74481
Summary
0 0.5 1 1.5 2 2.5 3
Standardized $log_2FC$
VIL1
GSE16844
GSE74481
Summary
−3 −2.5 −2 −1.5 −1 −0.5 0 0.5
Standardized $log_2FC$
VIM
GSE16844
GSE74481
Summary
0 0.5 1 1.5 2 2.5 3 3.5
Standardized $log_2FC$

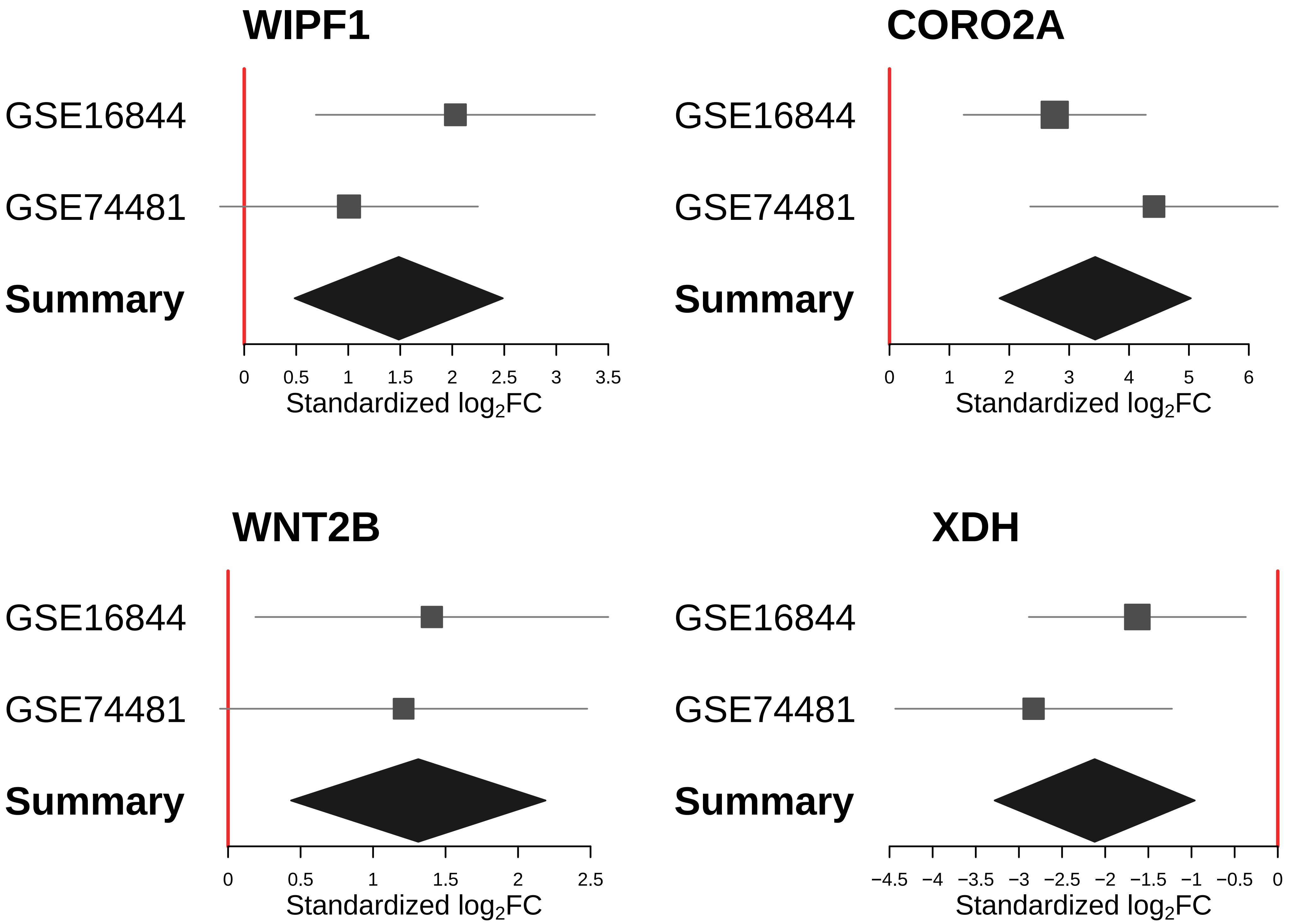
WIPF1
GSE16844
GSE74481
Summary
0 0.5 1 1.5 2 2.5 3 3.5
Standardized log2FC
CORO2A
GSE16844
GSE74481
Summary
0 1 2 3 4 5 6
Standardized log2FC
WNT2B
GSE16844
GSE74481
Summary
0 0.5 1 1.5 2 2.5
Standardized log2FC
XDH
GSE16844
GSE74481
Summary
−4.5 −4 −3.5 −3 −2.5 −2 −1.5 −1 −0.5 0
Standardized log2FC

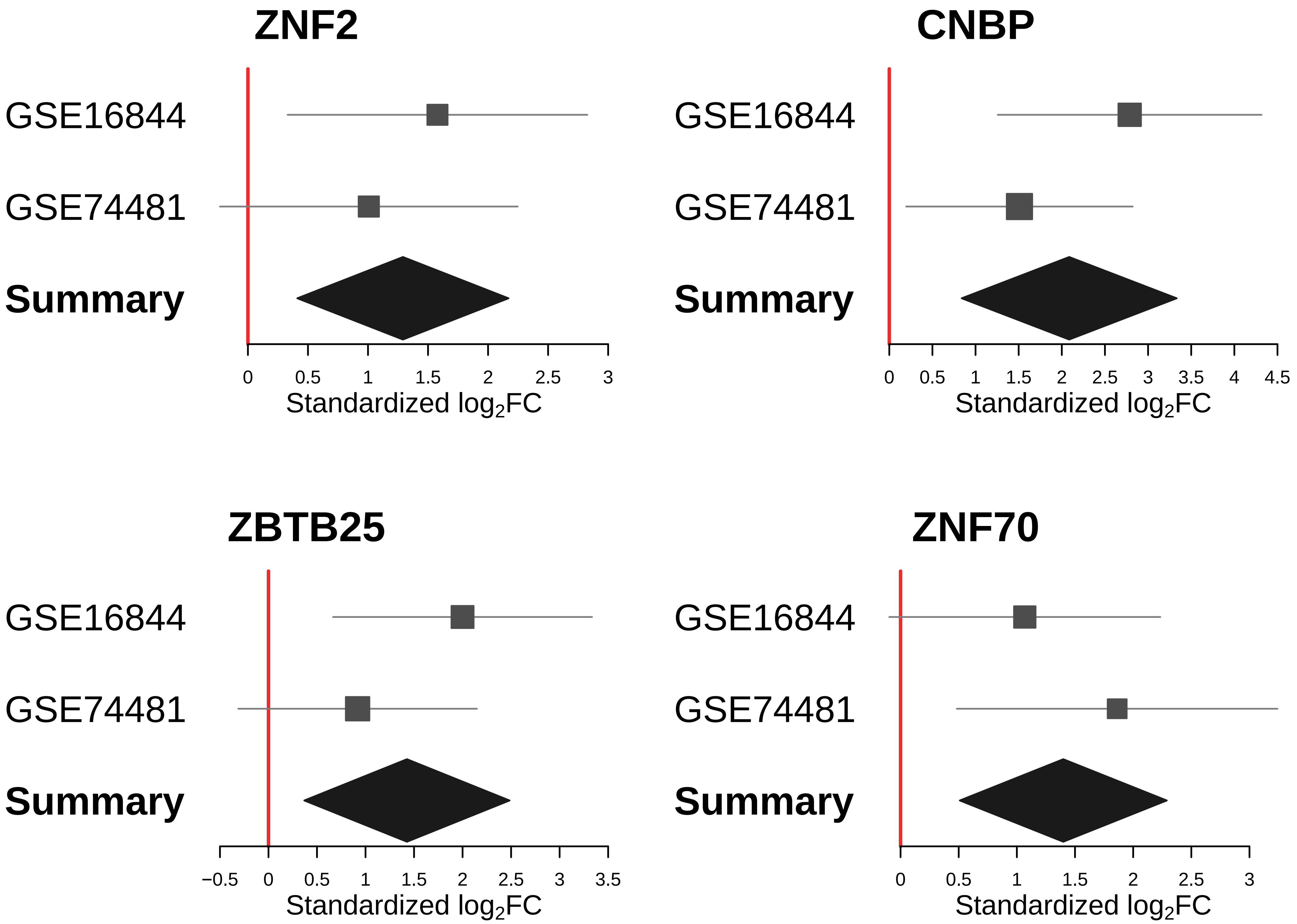

ZNF2
GSE16844
GSE74481
Summary
0
0.5
1
1.5
2
2.5
3
Standardized log2FC
CNBP
GSE16844
GSE74481
Summary
0
0.5
1
1.5
2
2.5
3
3.5
4
4.5
Standardized log2FC
ZBTB25
GSE16844
GSE74481
Summary
−0.5
0
0.5
1
1.5
2
2.5
3
3.5
Standardized log2FC
ZNF70
GSE16844
GSE74481
Summary
0
0.5
1
1.5
2
2.5
3
Standardized log2FC

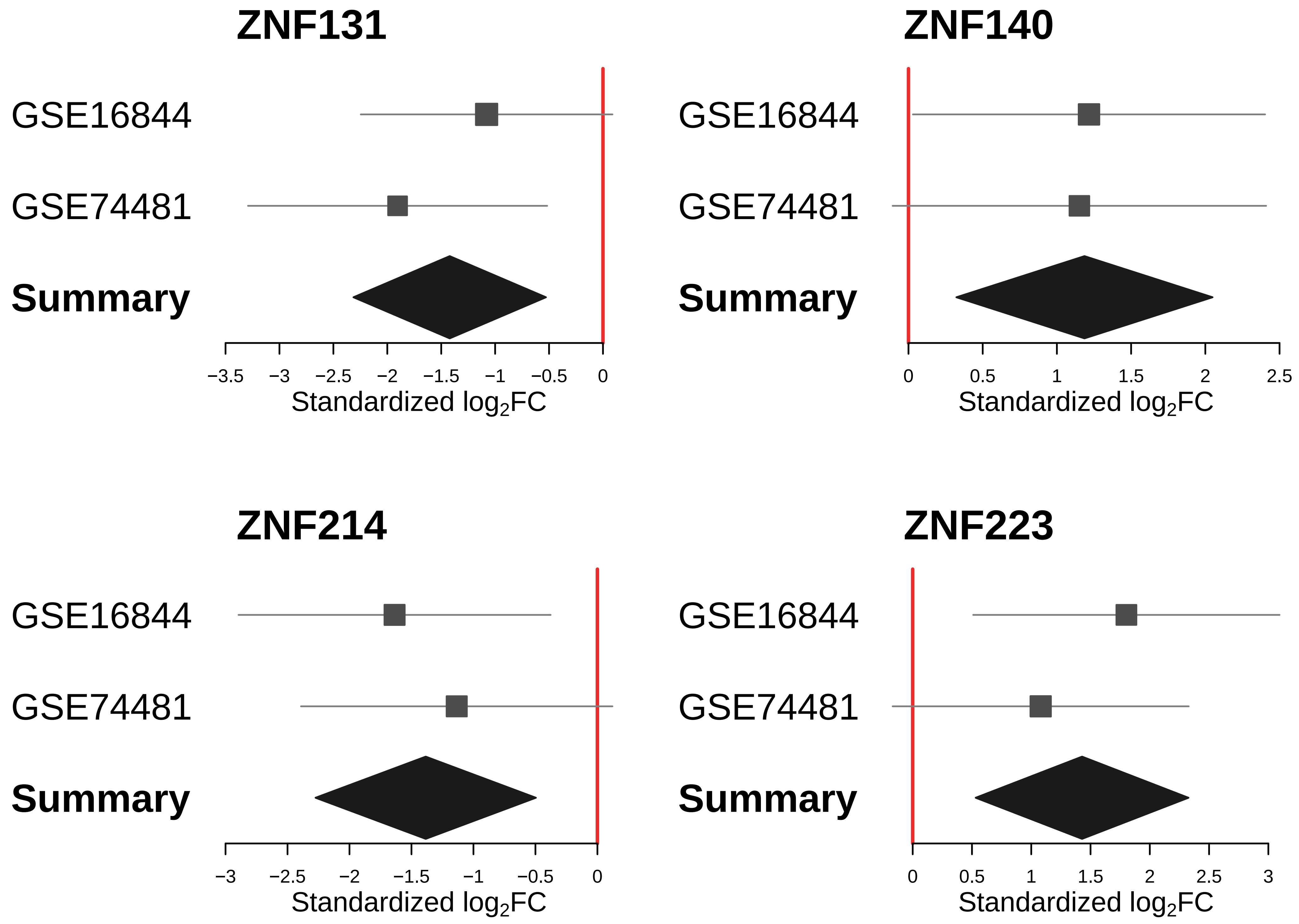

ZNF131
GSE16844
GSE74481
Summary
−3.5 −3 −2.5 −2 −1.5 −1 −0.5 0
Standardized log2FC
ZNF140
GSE16844
GSE74481
Summary
0 0.5 1 1.5 2 2.5
Standardized log2FC
ZNF214
GSE16844
GSE74481
Summary
−3 −2.5 −2 −1.5 −1 −0.5 0
Standardized log2FC
ZNF223
GSE16844
GSE74481
Summary
0 0.5 1 1.5 2 2.5 3
Standardized log2FC

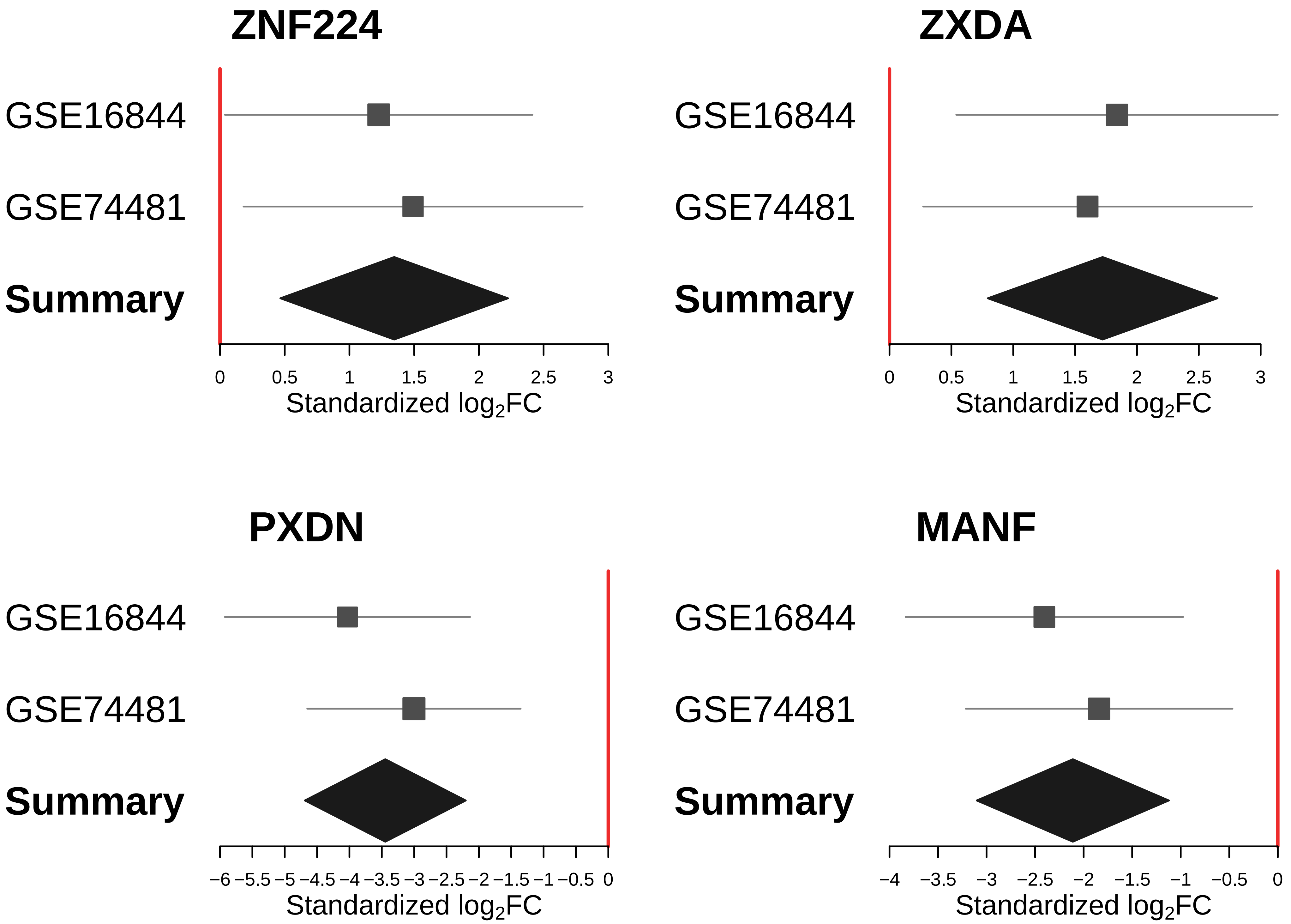
ZNF224
GSE16844
GSE74481
Summary
0 0.5 1 1.5 2 2.5 3
Standardized log2FC
ZXDA
GSE16844
GSE74481
Summary
0 0.5 1 1.5 2 2.5 3
Standardized log2FC
PXDN
GSE16844
GSE74481
Summary
−6 −5.5 −5 −4.5 −4 −3.5 −3 −2.5 −2 −1.5 −1 −0.5 0
Standardized log2FC
MANF
GSE16844
GSE74481
Summary
−4 −3.5 −3 −2.5 −2 −1.5 −1 −0.5 0
Standardized log2FC

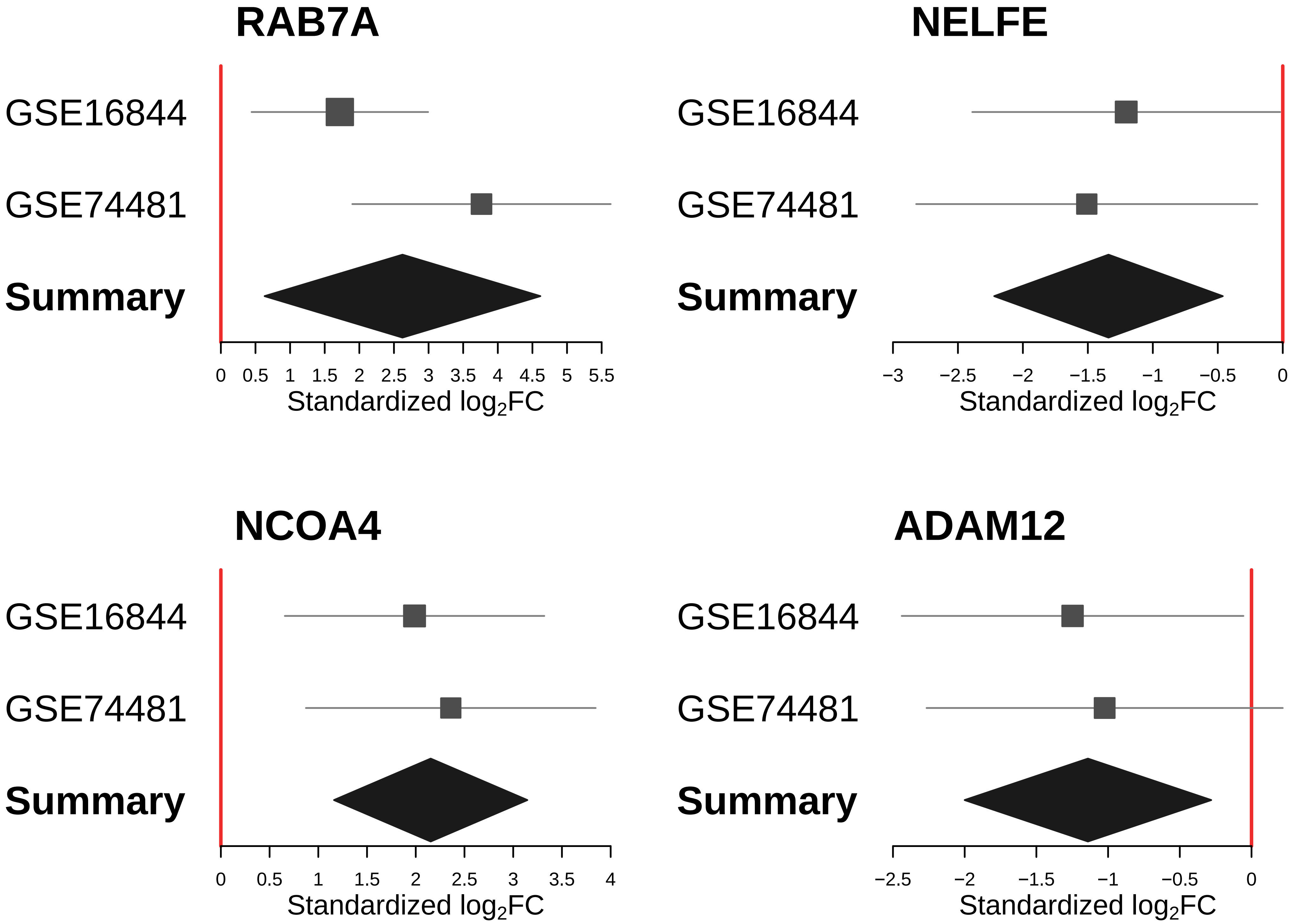
RAB7A
GSE16844
GSE74481
Summary
0 0.5 1 1.5 2 2.5 3 3.5 4 4.5 5 5.5
Standardized log2FC
NELFE
GSE16844
GSE74481
Summary
−3 −2.5 −2 −1.5 −1 −0.5 0
Standardized log2FC
NCOA4
GSE16844
GSE74481
Summary
0 0.5 1 1.5 2 2.5 3 3.5 4
Standardized log2FC
ADAM12
GSE16844
GSE74481
Summary
−2.5 −2 −1.5 −1 −0.5 0
Standardized log2FC

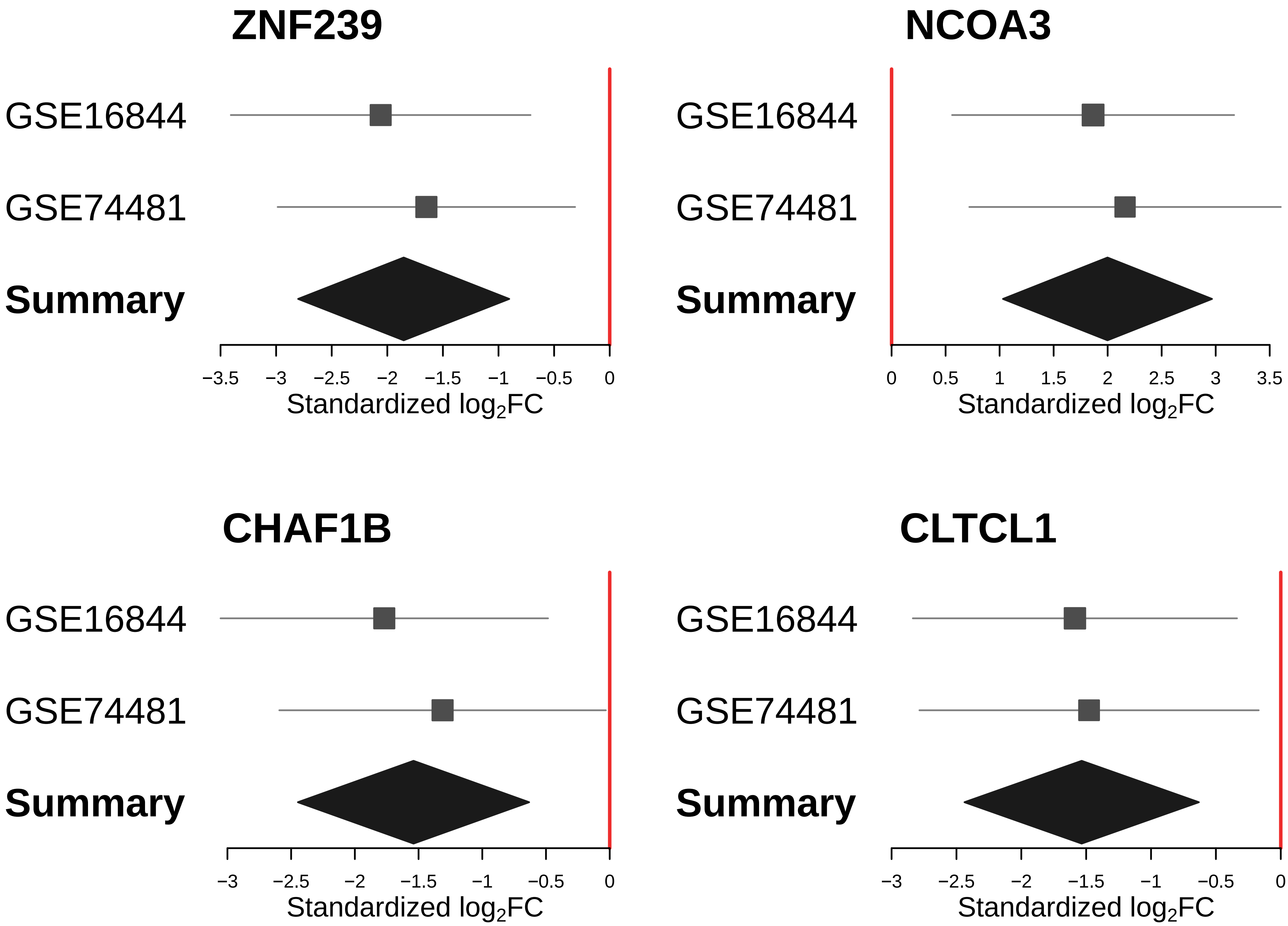
ZNF239
GSE16844
GSE74481
Summary
−3.5 −3 −2.5 −2 −1.5 −1 −0.5 0
Standardized $log_2FC$
NCOA3
GSE16844
GSE74481
Summary
0 0.5 1 1.5 2 2.5 3 3.5
Standardized $log_2FC$
CHAF1B
GSE16844
GSE74481
Summary
−3 −2.5 −2 −1.5 −1 −0.5 0
Standardized $log_2FC$
CLTCL1
GSE16844
GSE74481
Summary
−3 −2.5 −2 −1.5 −1 −0.5 0
Standardized $log_2FC$

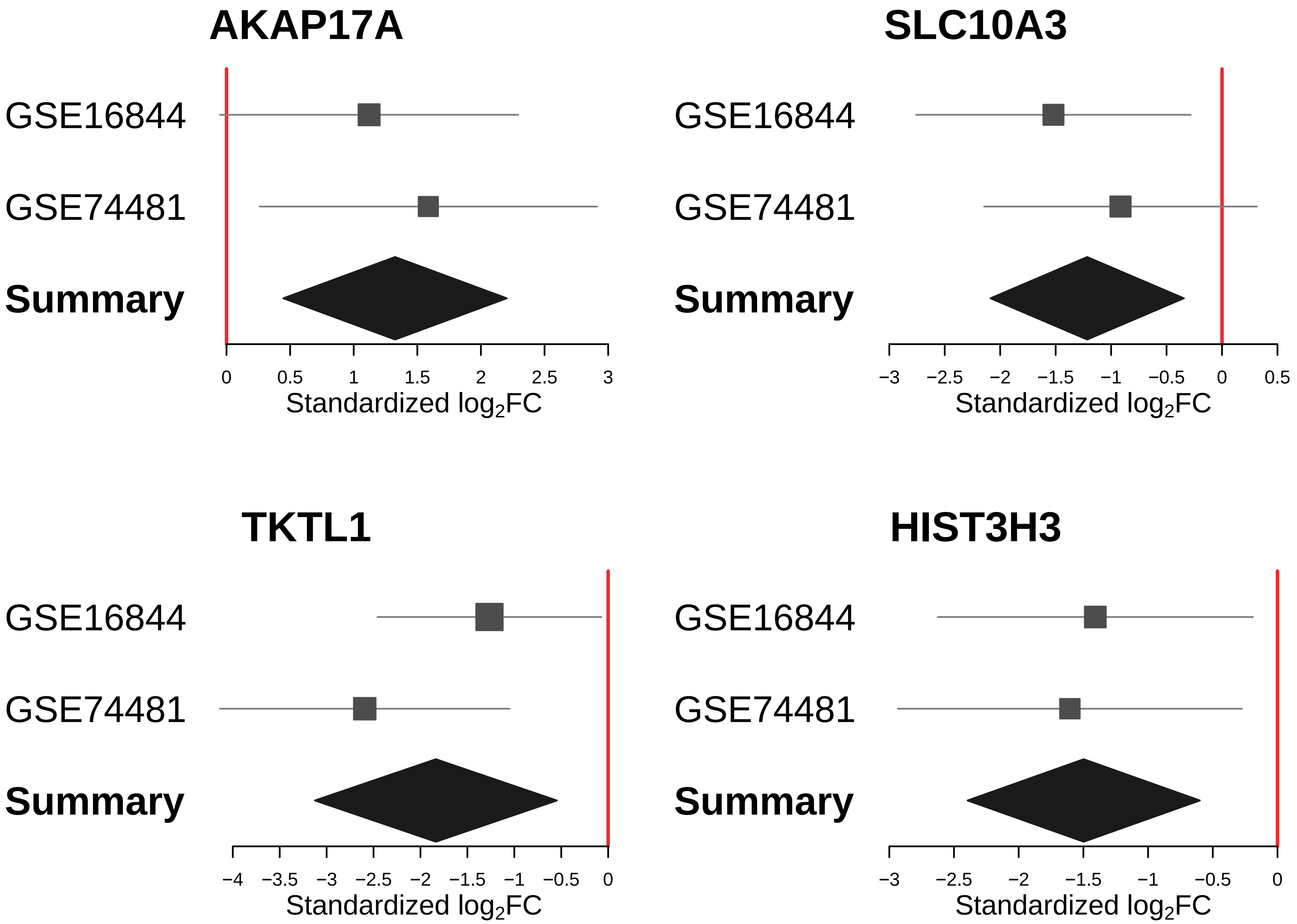
AKAP17A
GSE16844
GSE74481
Summary
0
0.5
1
1.5
2
2.5
3
Standardized log2FC
SLC10A3
GSE16844
GSE74481
Summary
−3
−2.5
−2
−1.5
−1
−0.5
0
0.5
Standardized log2FC
TKTL1
GSE16844
GSE74481
Summary
−4
−3.5
−3
−2.5
−2
−1.5
−1
−0.5
0
Standardized log2FC
HIST3H3
GSE16844
GSE74481
Summary
−3
−2.5
−2
−1.5
−1
−0.5
0
Standardized log2FC

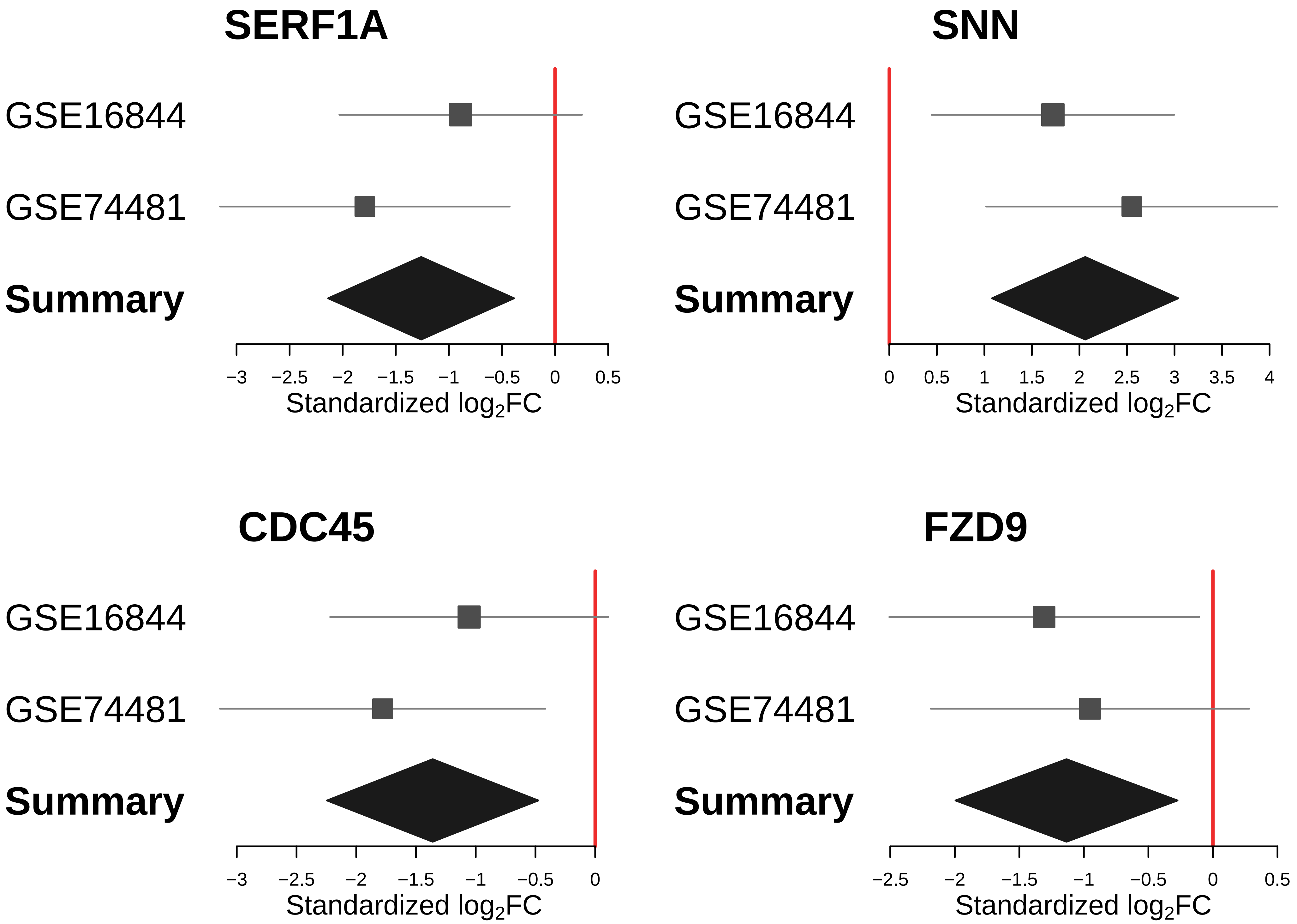

SERF1A
GSE16844
GSE74481
Summary
−3 −2.5 −2 −1.5 −1 −0.5 0 0.5
Standardized log2FC
SNN
GSE16844
GSE74481
Summary
0 0.5 1 1.5 2 2.5 3 3.5 4
Standardized log2FC
CDC45
GSE16844
GSE74481
Summary
−3 −2.5 −2 −1.5 −1 −0.5 0
Standardized log2FC
FZD9
GSE16844
GSE74481
Summary
−2.5 −2 −1.5 −1 −0.5 0 0.5
Standardized log2FC

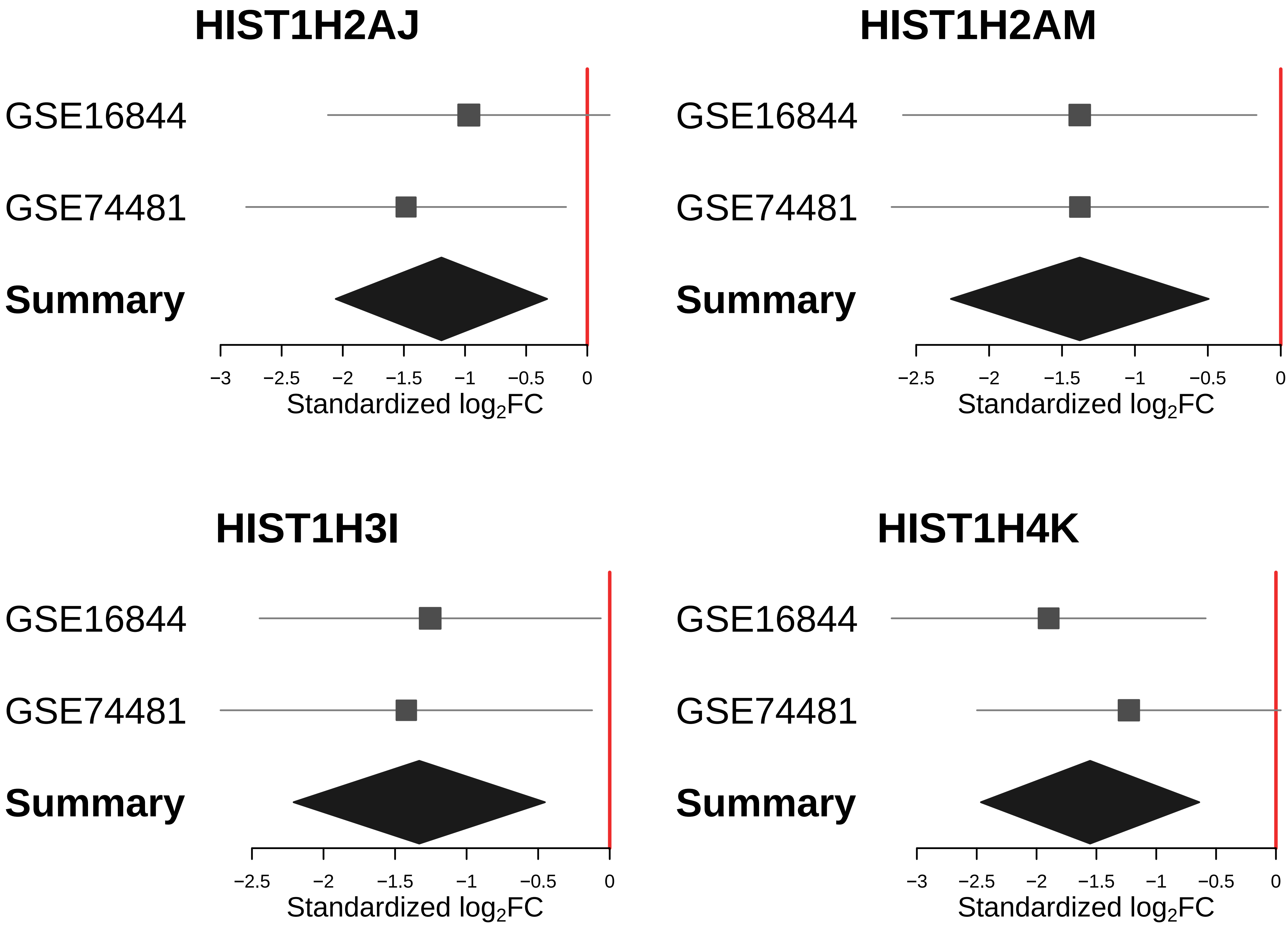
HIST1H2AJ
GSE16844
GSE74481
Summary
−3 −2.5 −2 −1.5 −1 −0.5 0
Standardized log2FC
HIST1H2AM
GSE16844
GSE74481
Summary
−2.5 −2 −1.5 −1 −0.5 0
Standardized log2FC
HIST1H3I
GSE16844
GSE74481
Summary
−2.5 −2 −1.5 −1 −0.5 0
Standardized log2FC
HIST1H4K
GSE16844
GSE74481
Summary
−3 −2.5 −2 −1.5 −1 −0.5 0
Standardized log2FC

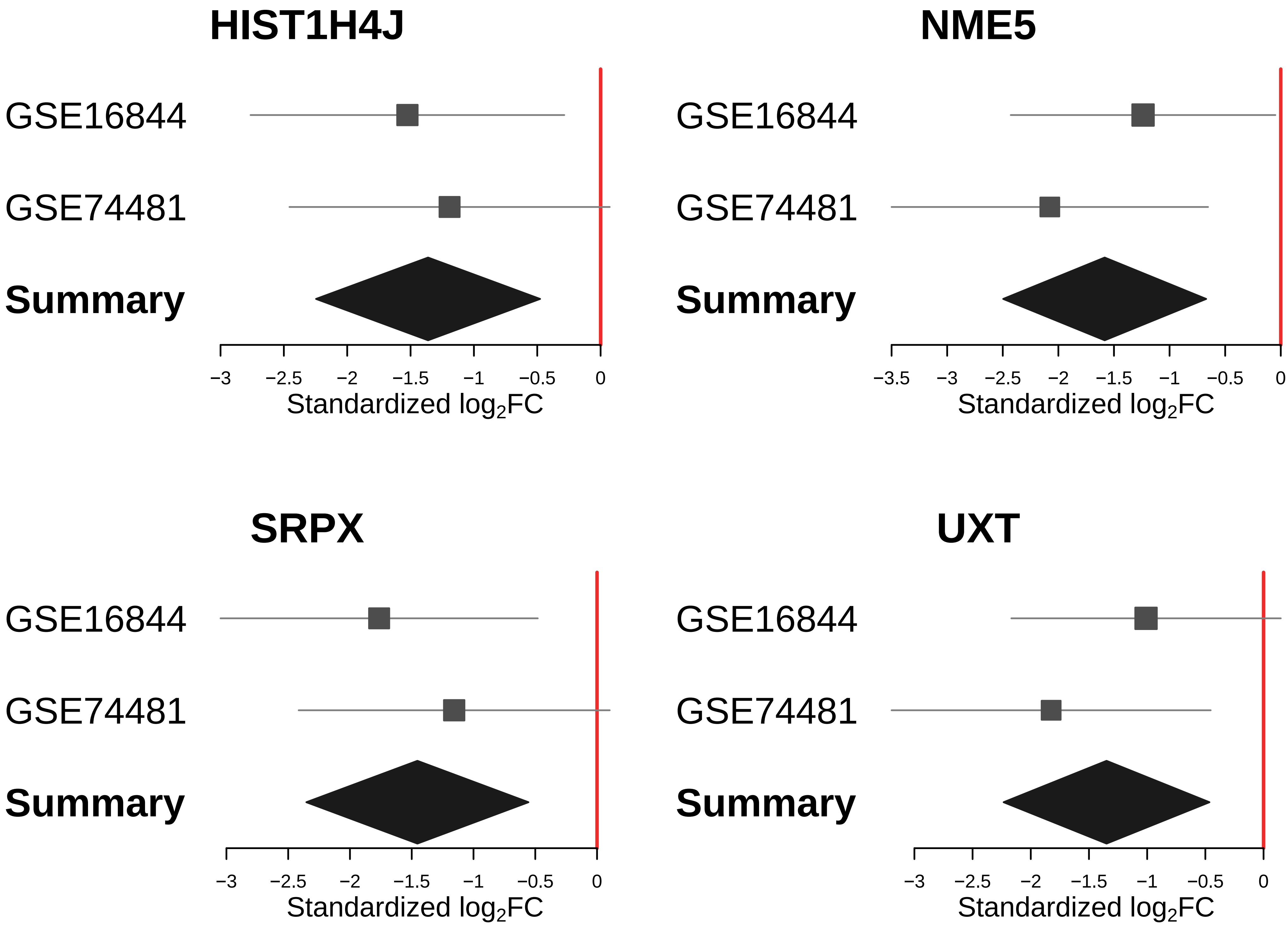
HIST1H4J
GSE16844
GSE74481
Summary
−3 −2.5 −2 −1.5 −1 −0.5 0
Standardized log2FC
NME5
GSE16844
GSE74481
Summary
−3.5 −3 −2.5 −2 −1.5 −1 −0.5 0
Standardized log2FC
SRPX
GSE16844
GSE74481
Summary
−3 −2.5 −2 −1.5 −1 −0.5 0
Standardized log2FC
UXT
GSE16844
GSE74481
Summary
−3 −2.5 −2 −1.5 −1 −0.5 0
Standardized log2FC

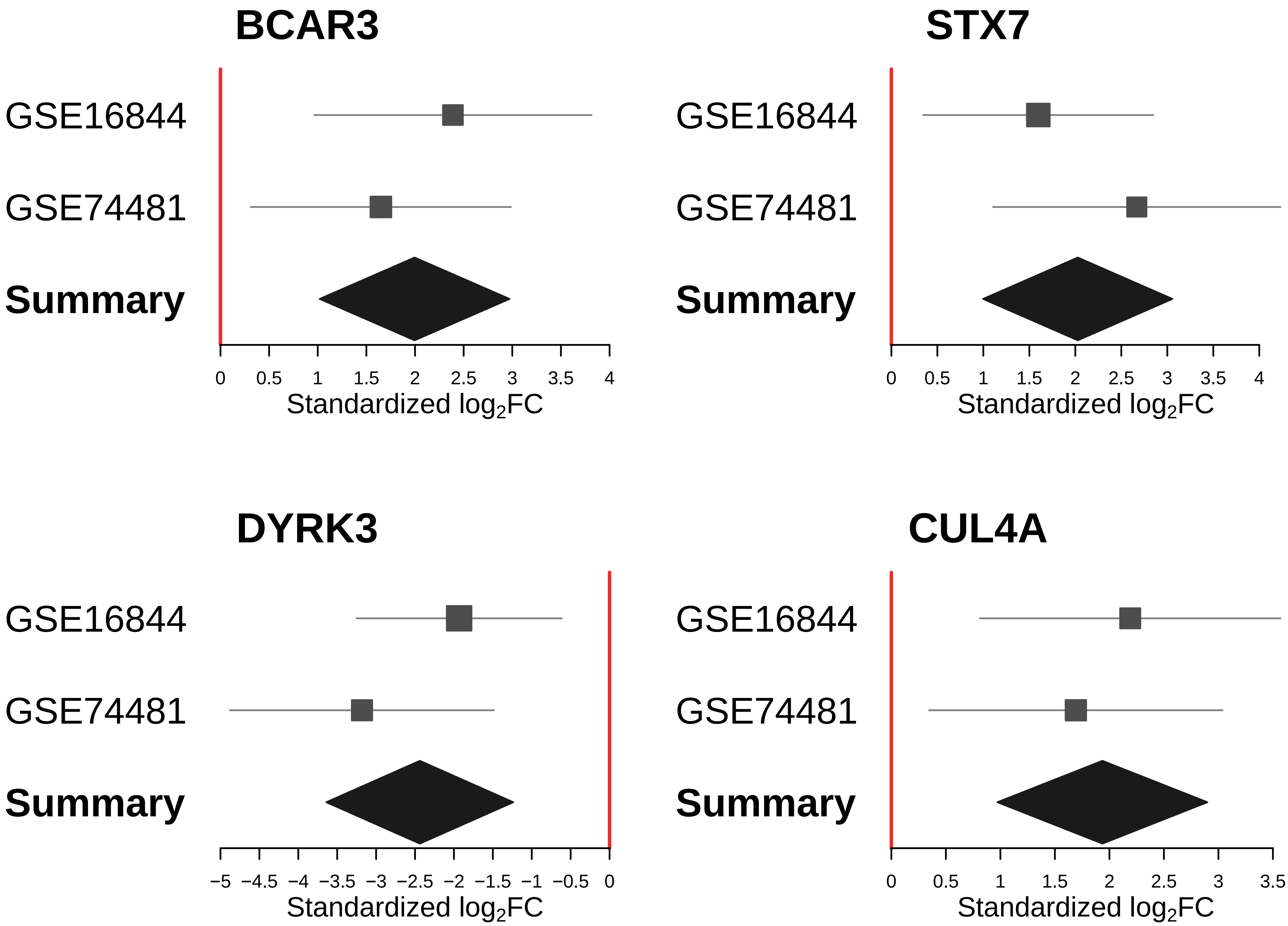

BCAR3
GSE16844
GSE74481
Summary
0 0.5 1 1.5 2 2.5 3 3.5 4
Standardized log2FC
STX7
GSE16844
GSE74481
Summary
0 0.5 1 1.5 2 2.5 3 3.5 4
Standardized log2FC
DYRK3
GSE16844
GSE74481
Summary
−5 −4.5 −4 −3.5 −3 −2.5 −2 −1.5 −1 −0.5 0
Standardized log2FC
CUL4A
GSE16844
GSE74481
Summary
0 0.5 1 1.5 2 2.5 3 3.5
Standardized log2FC

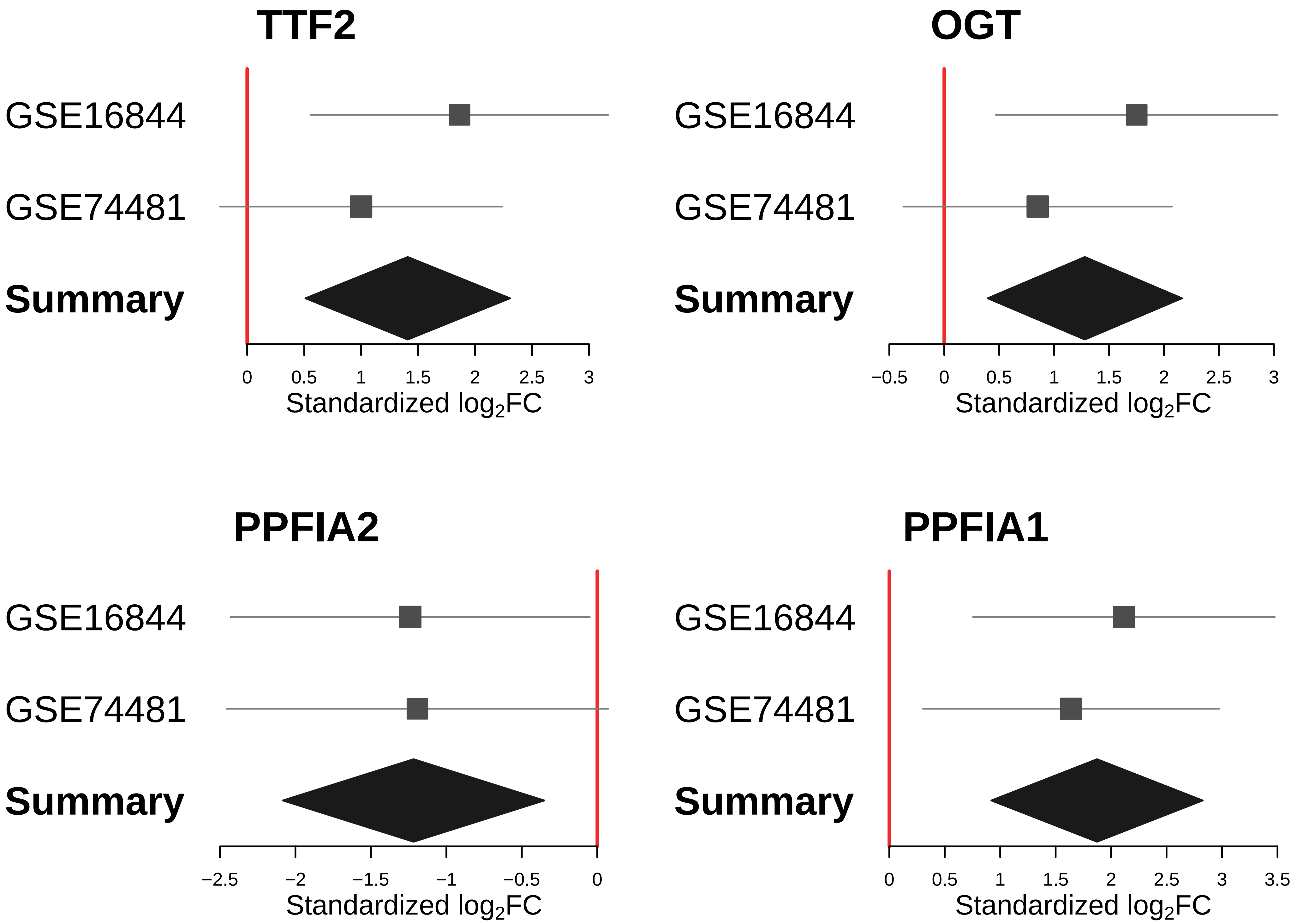
TTF2
GSE16844
GSE74481
Summary
0
0.5
1
1.5
2
2.5
3
Standardized log2FC
OGT
GSE16844
GSE74481
Summary
−0.5
0
0.5
1
1.5
2
2.5
3
Standardized log2FC
PPFIA2
GSE16844
GSE74481
Summary
−2.5
−2
−1.5
−1
−0.5
0
Standardized log2FC
PPFIA1
GSE16844
GSE74481
Summary
0
0.5
1
1.5
2
2.5
3
3.5
Standardized log2FC

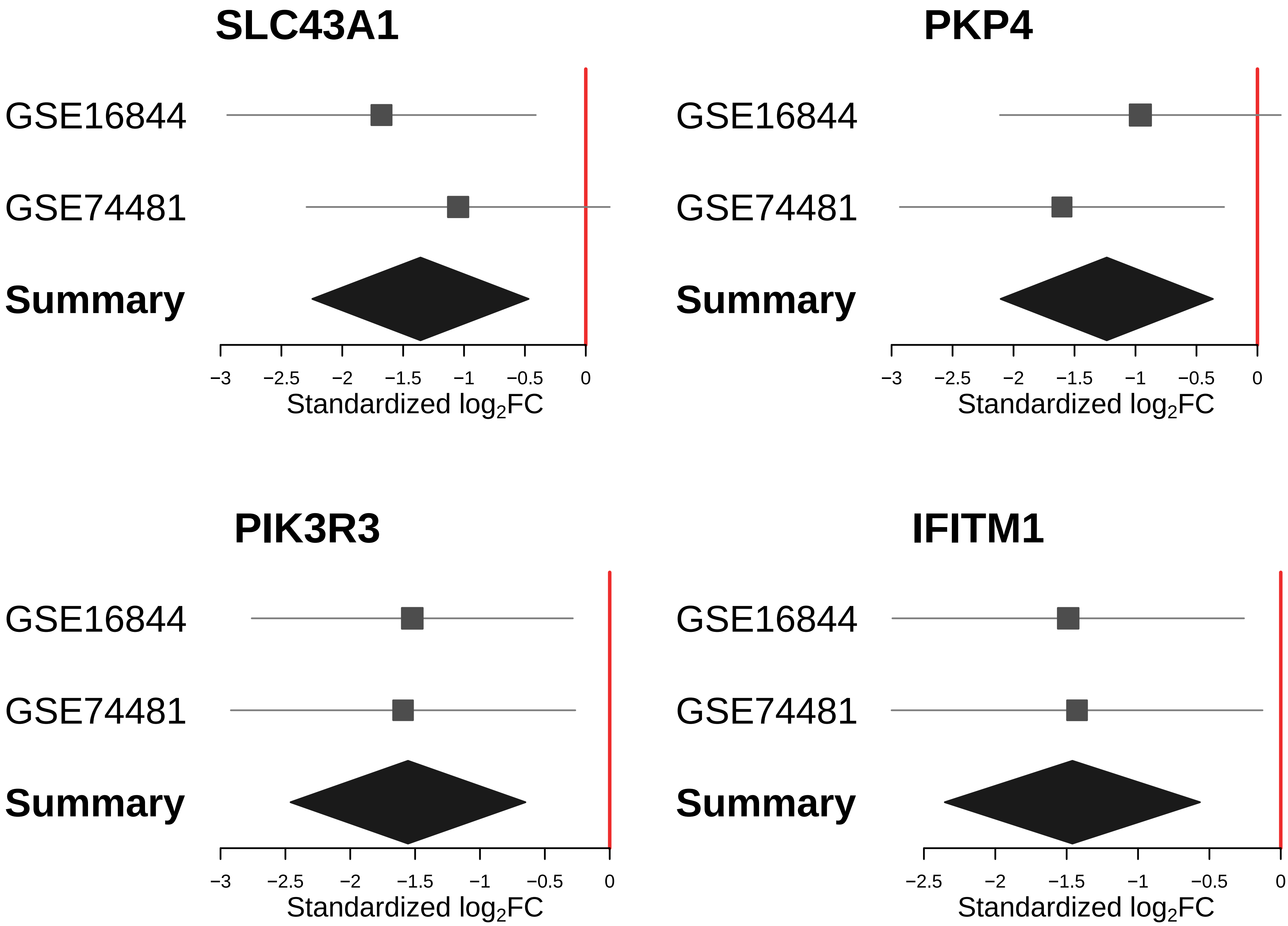

SLC43A1
GSE16844
GSE74481
Summary
−3 −2.5 −2 −1.5 −1 −0.5 0
Standardized $log_2FC$
PKP4
GSE16844
GSE74481
Summary
−3 −2.5 −2 −1.5 −1 −0.5 0
Standardized $log_2FC$
PIK3R3
GSE16844
GSE74481
Summary
−3 −2.5 −2 −1.5 −1 −0.5 0
Standardized $log_2FC$
IFITM1
GSE16844
GSE74481
Summary
−2.5 −2 −1.5 −1 −0.5 0
Standardized $log_2FC$

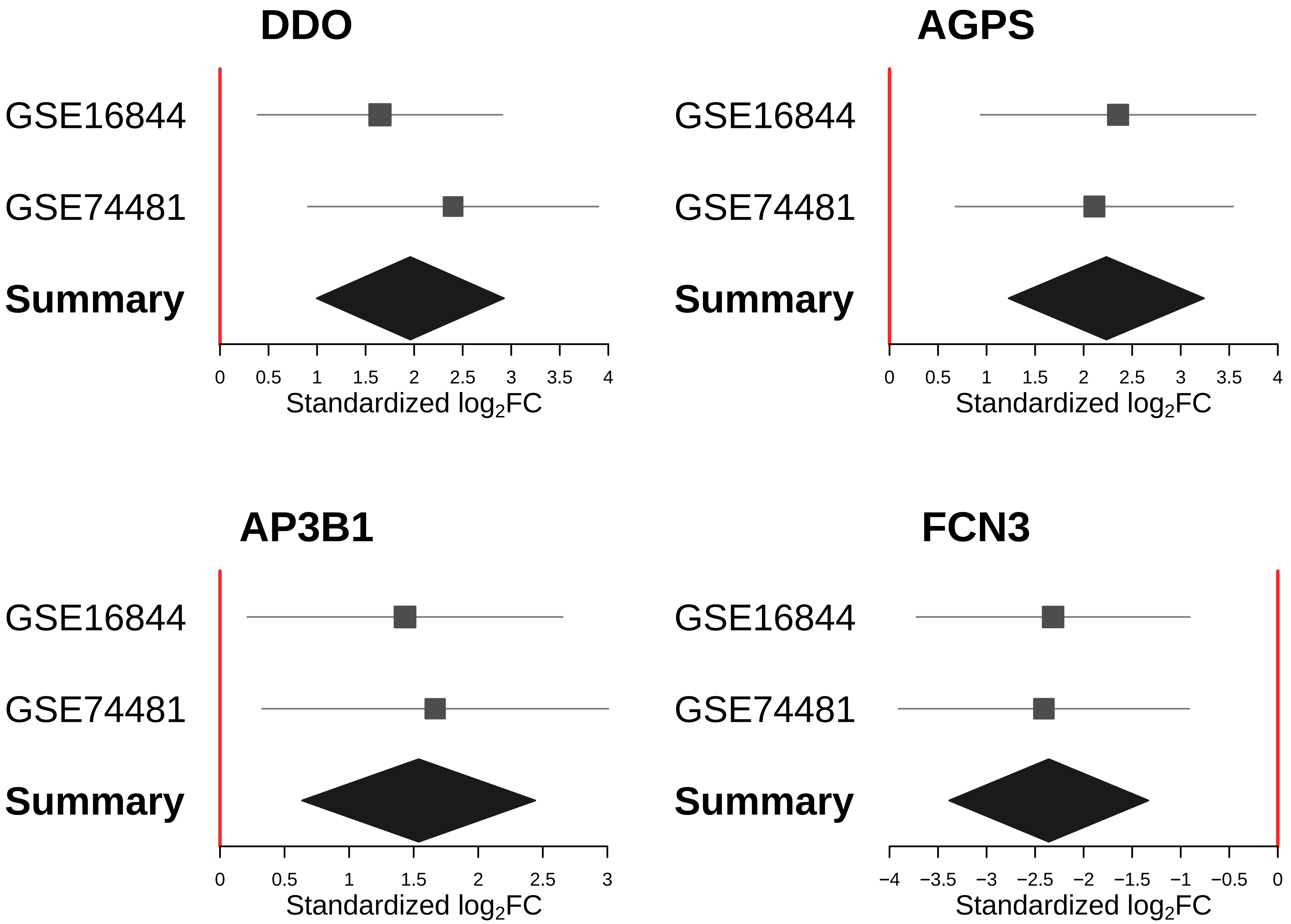
DDO
GSE16844
GSE74481
Summary
0 0.5 1 1.5 2 2.5 3 3.5 4
Standardized log2FC
AGPS
GSE16844
GSE74481
Summary
0 0.5 1 1.5 2 2.5 3 3.5 4
Standardized log2FC
AP3B1
GSE16844
GSE74481
Summary
0 0.5 1 1.5 2 2.5 3
Standardized log2FC
FCN3
GSE16844
GSE74481
Summary
−4 −3.5 −3 −2.5 −2 −1.5 −1 −0.5 0
Standardized log2FC

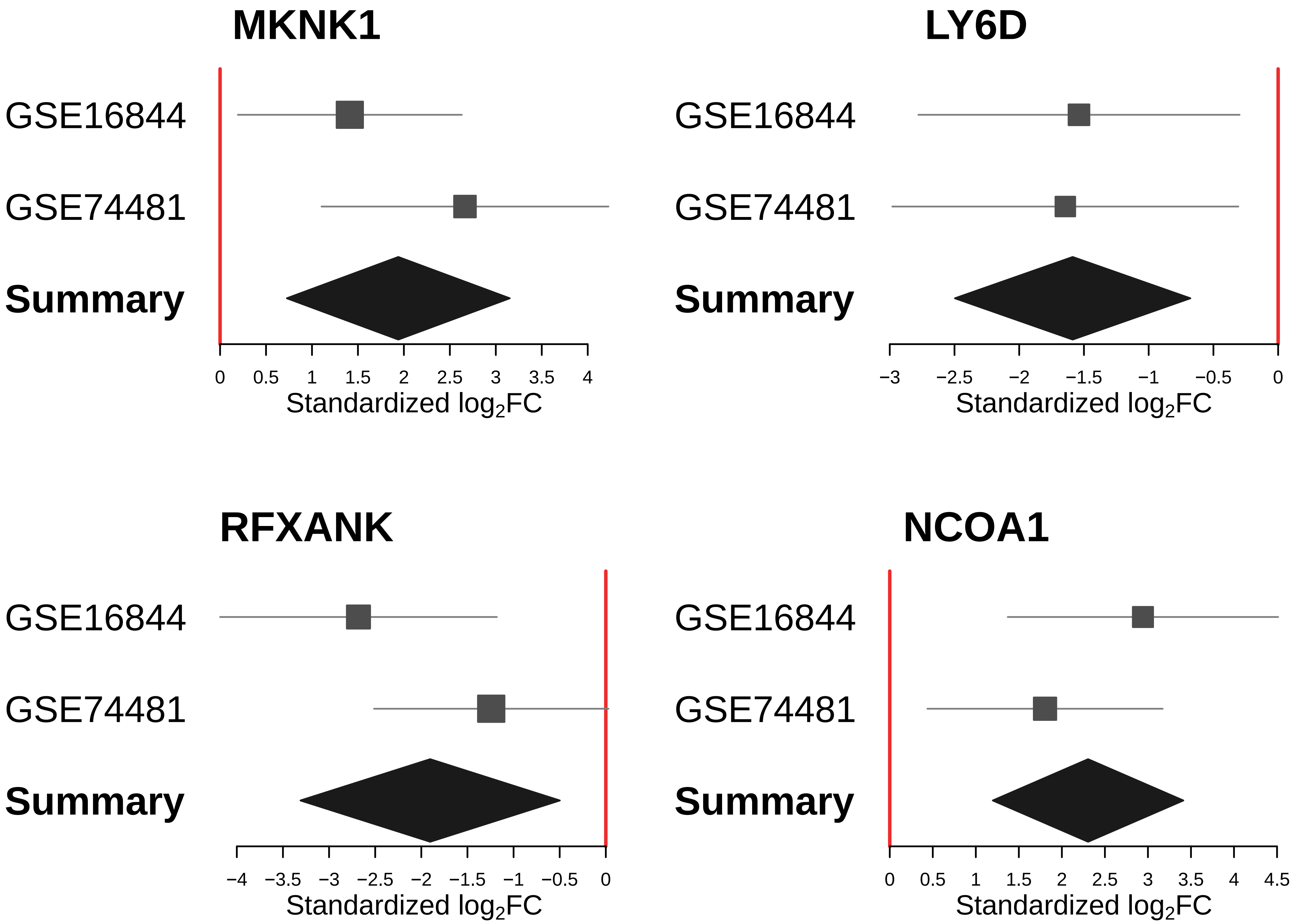

MKNK1
GSE16844
GSE74481
Summary
0 0.5 1 1.5 2 2.5 3 3.5 4
Standardized log2FC
LY6D
GSE16844
GSE74481
Summary
−3 −2.5 −2 −1.5 −1 −0.5 0
Standardized log2FC
RFXANK
GSE16844
GSE74481
Summary
−4 −3.5 −3 −2.5 −2 −1.5 −1 −0.5 0
Standardized log2FC
NCOA1
GSE16844
GSE74481
Summary
0 0.5 1 1.5 2 2.5 3 3.5 4 4.5
Standardized log2FC

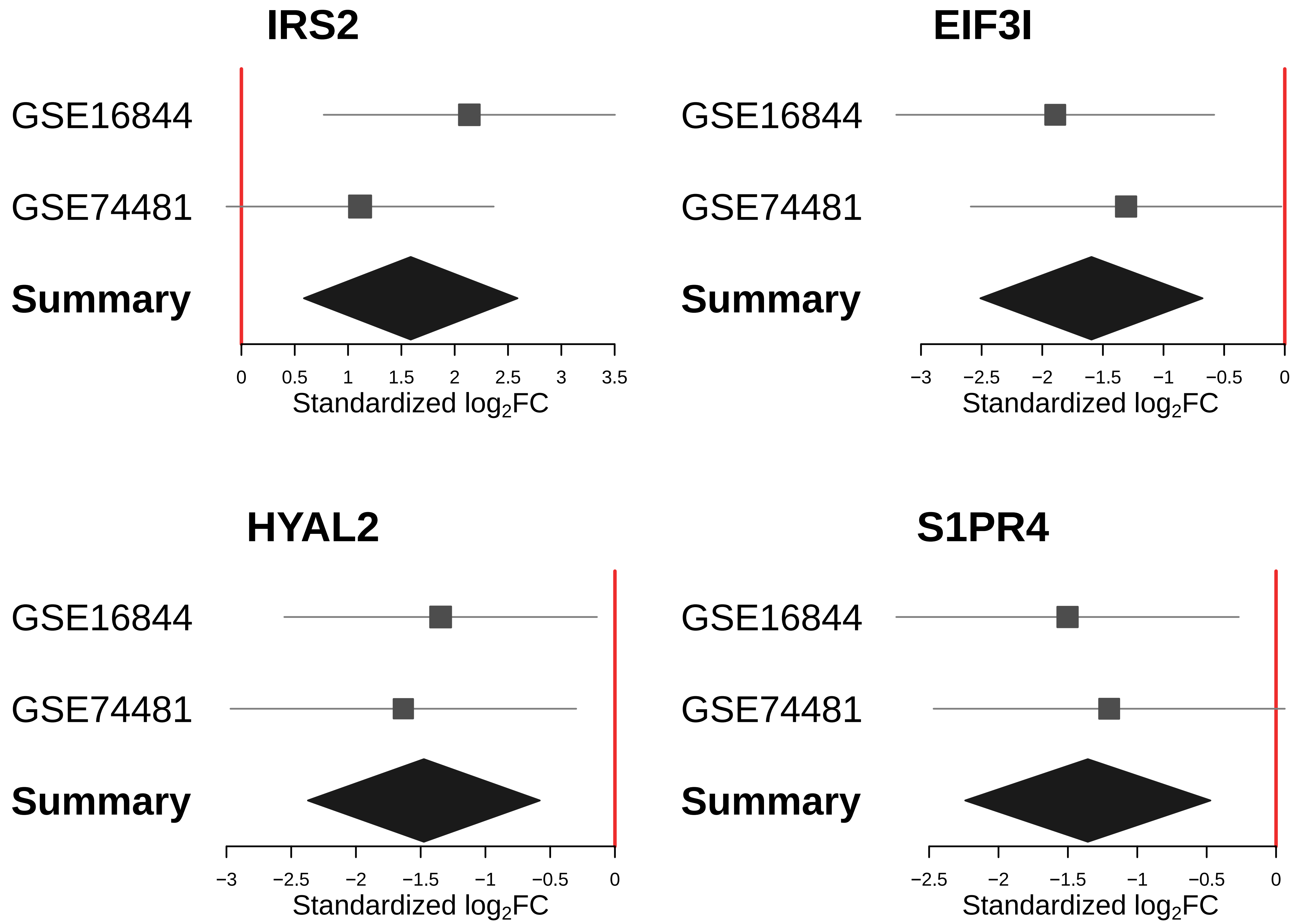
IRS2
GSE16844
GSE74481
Summary
0
0.5
1
1.5
2
2.5
3
3.5
Standardized log2FC
EIF3I
GSE16844
GSE74481
Summary
−3
−2.5
−2
−1.5
−1
−0.5
0
Standardized log2FC
HYAL2
GSE16844
GSE74481
Summary
−3
−2.5
−2
−1.5
−1
−0.5
0
Standardized log2FC
S1PR4
GSE16844
GSE74481
Summary
−2.5
−2
−1.5
−1
−0.5
0
Standardized log2FC

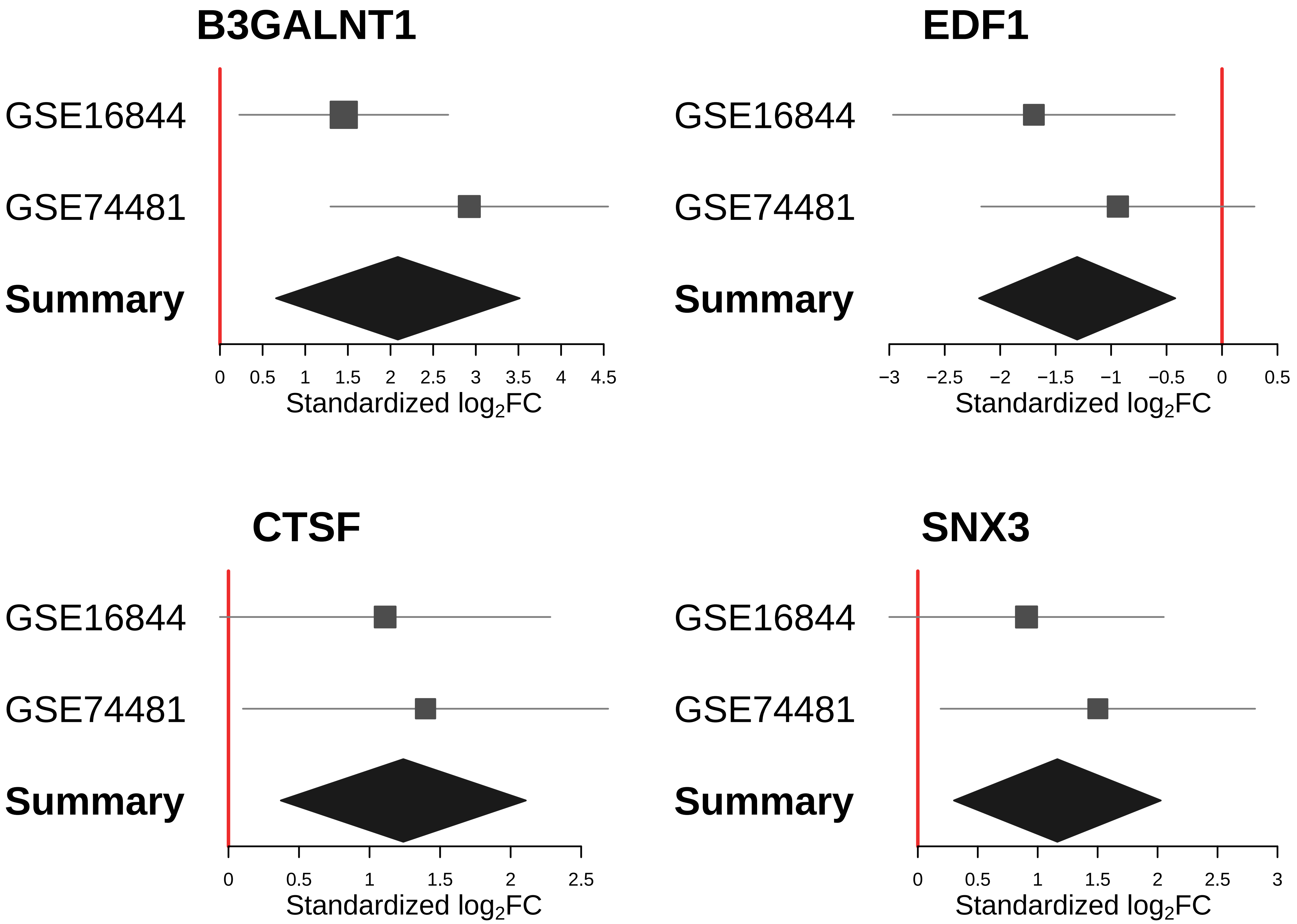
B3GALNT1
GSE16844
GSE74481
Summary
0 0.5 1 1.5 2 2.5 3 3.5 4 4.5
Standardized log2FC
EDF1
GSE16844
GSE74481
Summary
−3 −2.5 −2 −1.5 −1 −0.5 0 0.5
Standardized log2FC
CTSF
GSE16844
GSE74481
Summary
0 0.5 1 1.5 2 2.5
Standardized log2FC
SNX3
GSE16844
GSE74481
Summary
0 0.5 1 1.5 2 2.5 3
Standardized log2FC

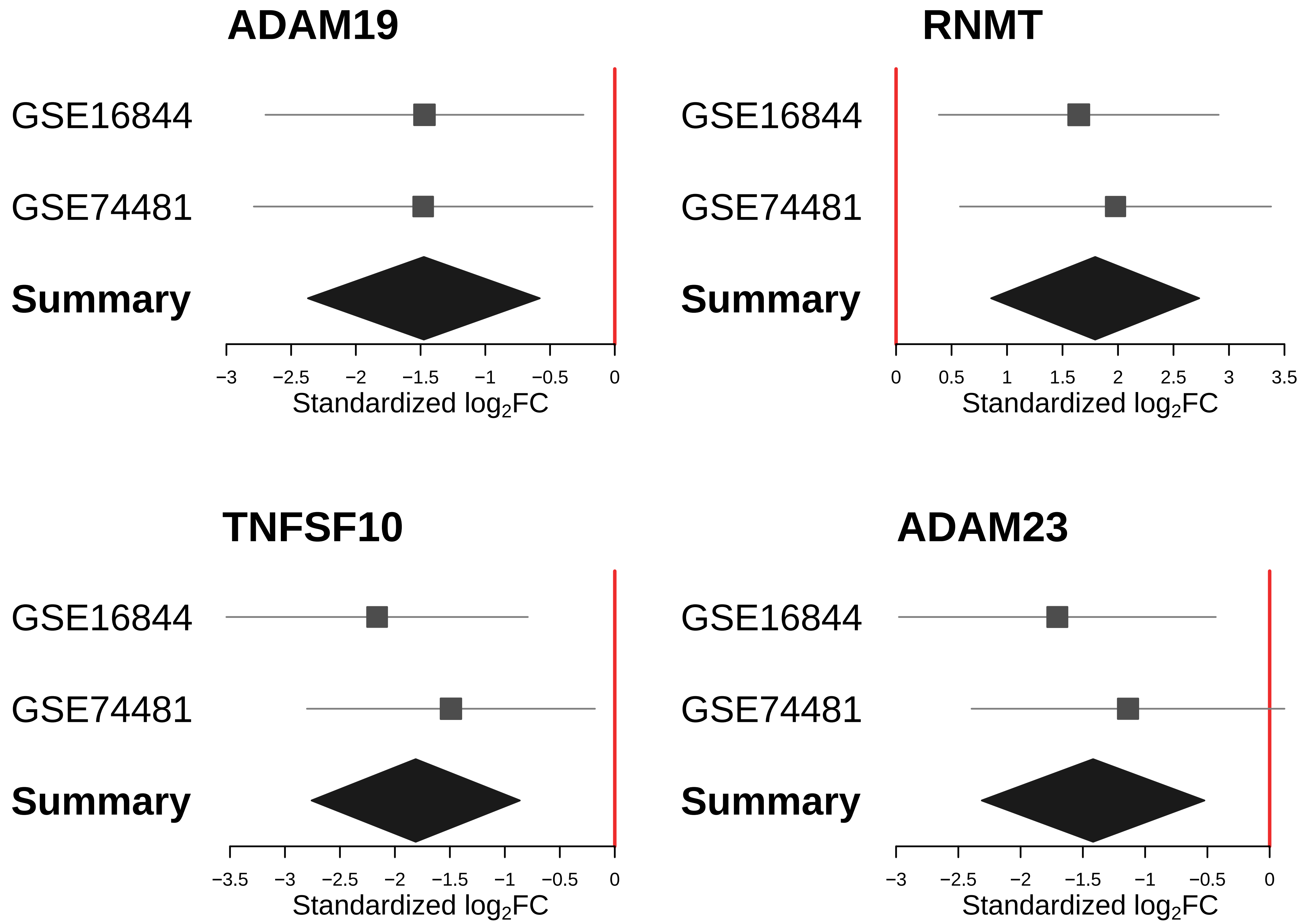
ADAM19
GSE16844
GSE74481
Summary
−3
−2.5
−2
−1.5
−1
−0.5
0
Standardized log2FC
RNMT
GSE16844
GSE74481
Summary
0
0.5
1
1.5
2
2.5
3
3.5
Standardized log2FC
TNFSF10
GSE16844
GSE74481
Summary
−3.5
−3
−2.5
−2
−1.5
−1
−0.5
0
Standardized log2FC
ADAM23
GSE16844
GSE74481
Summary
−3
−2.5
−2
−1.5
−1
−0.5
0
Standardized log2FC

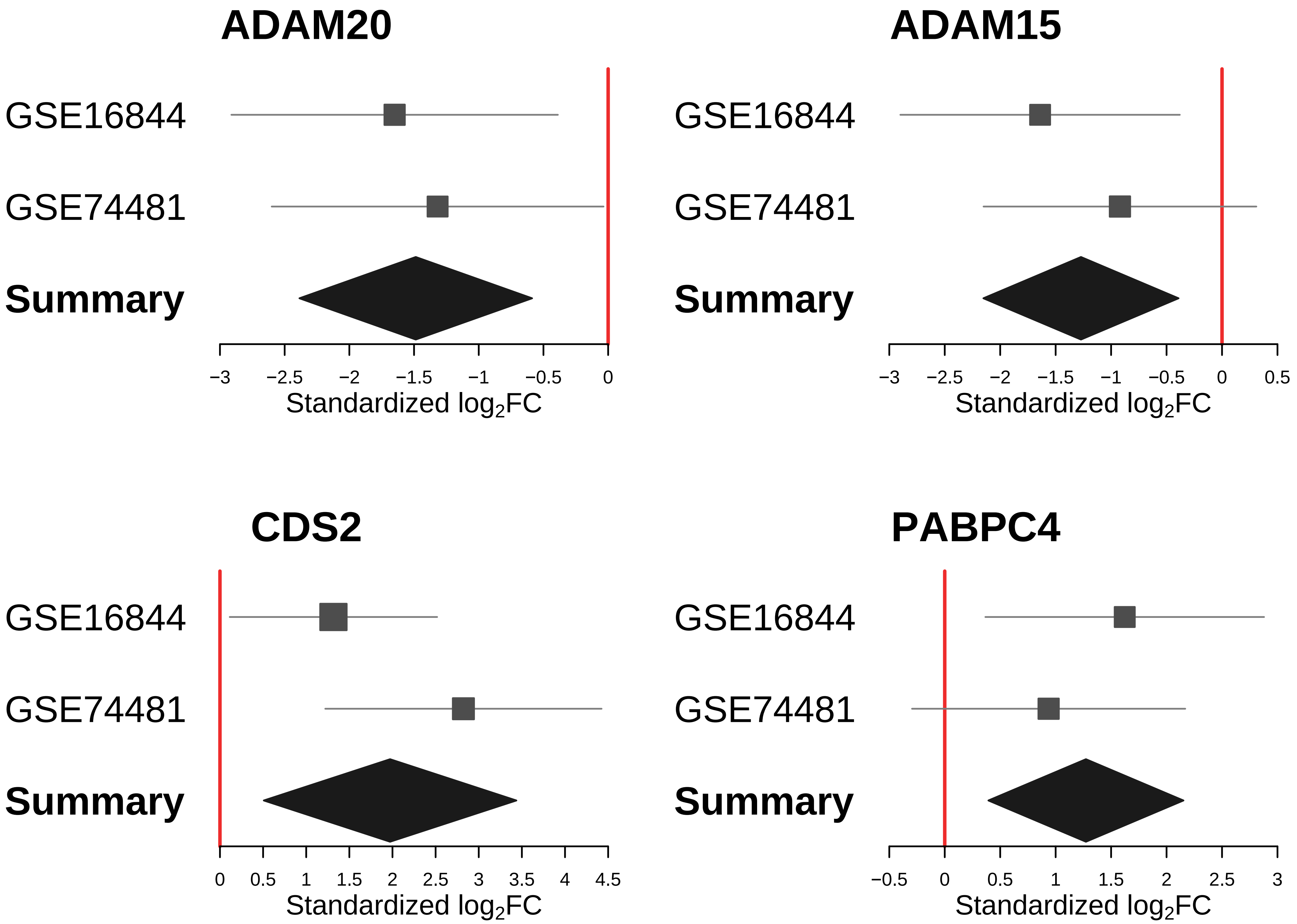

ADAM20
GSE16844
GSE74481
Summary
−3 −2.5 −2 −1.5 −1 −0.5 0
Standardized log2FC
ADAM15
GSE16844
GSE74481
Summary
−3 −2.5 −2 −1.5 −1 −0.5 0 0.5
Standardized log2FC
CDS2
GSE16844
GSE74481
Summary
0 0.5 1 1.5 2 2.5 3 3.5 4 4.5
Standardized log2FC
PABPC4
GSE16844
GSE74481
Summary
−0.5 0 0.5 1 1.5 2 2.5 3
Standardized log2FC

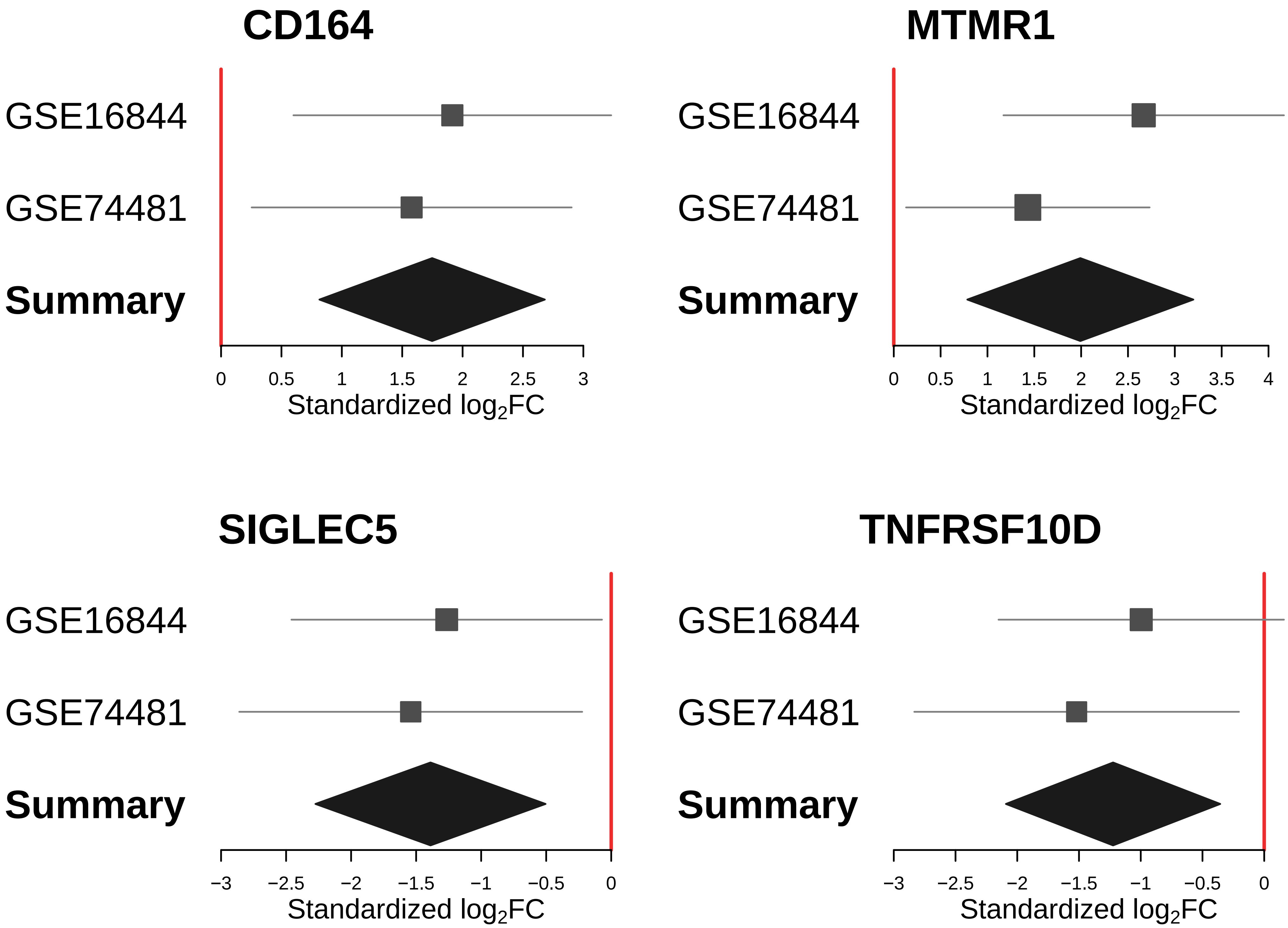
CD164
GSE16844
GSE74481
Summary
0
0.5
1
1.5
2
2.5
3
Standardized $log_2$FC
MTMR1
GSE16844
GSE74481
Summary
0
0.5
1
1.5
2
2.5
3
3.5
4
Standardized $log_2$FC
SIGLEC5
GSE16844
GSE74481
Summary
−3
−2.5
−2
−1.5
−1
−0.5
0
Standardized $log_2$FC
TNFRSF10D
GSE16844
GSE74481
Summary
−3
−2.5
−2
−1.5
−1
−0.5
0
Standardized $log_2$FC

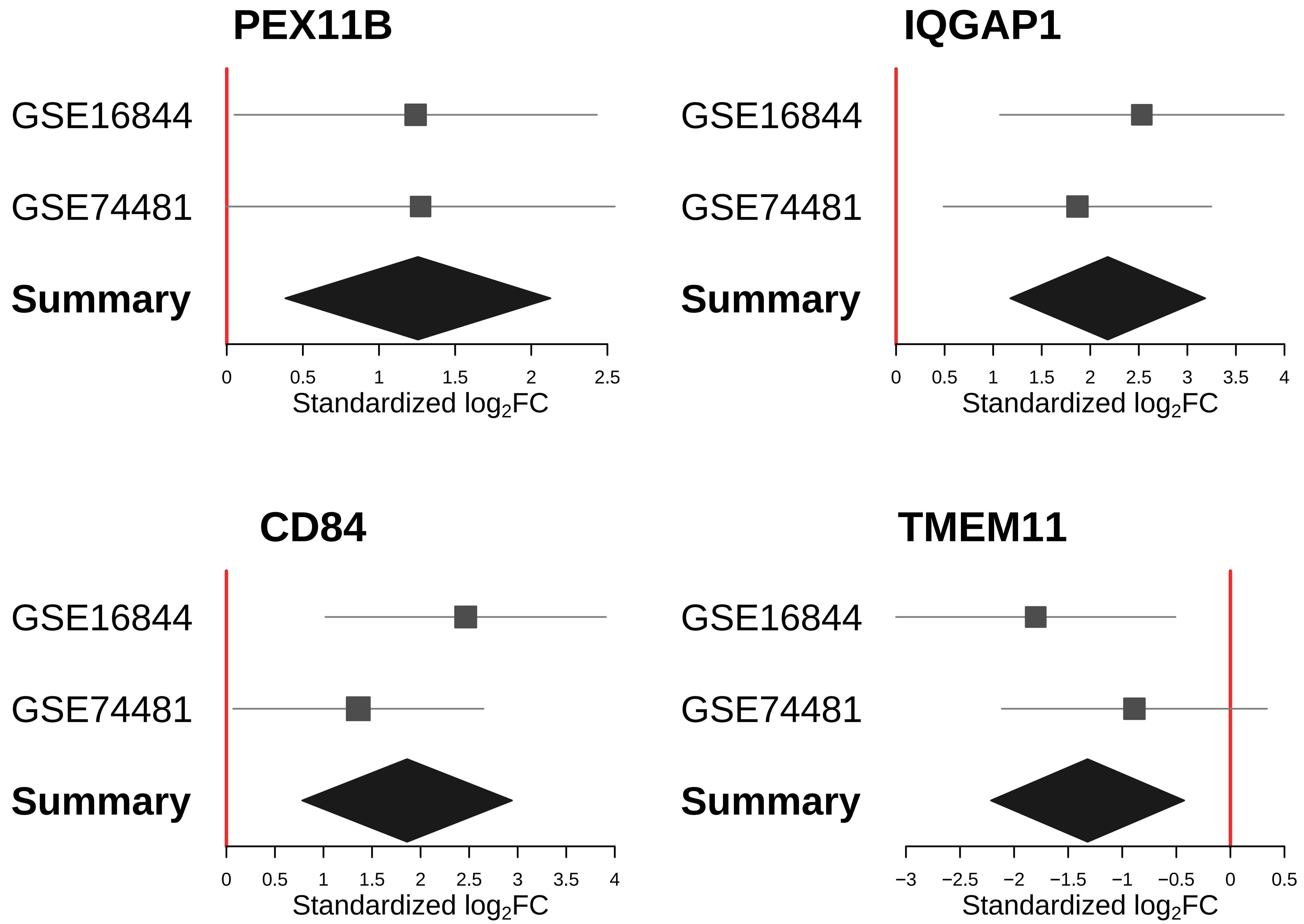
PEX11B
GSE16844
GSE74481
Summary
0
0.5
1
1.5
2
2.5
Standardized log2FC
IQGAP1
GSE16844
GSE74481
Summary
0
0.5
1
1.5
2
2.5
3
3.5
4
Standardized log2FC
CD84
GSE16844
GSE74481
Summary
0
0.5
1
1.5
2
2.5
3
3.5
4
Standardized log2FC
TMEM11
GSE16844
GSE74481
Summary
−3
−2.5
−2
−1.5
−1
−0.5
0
0.5
Standardized log2FC

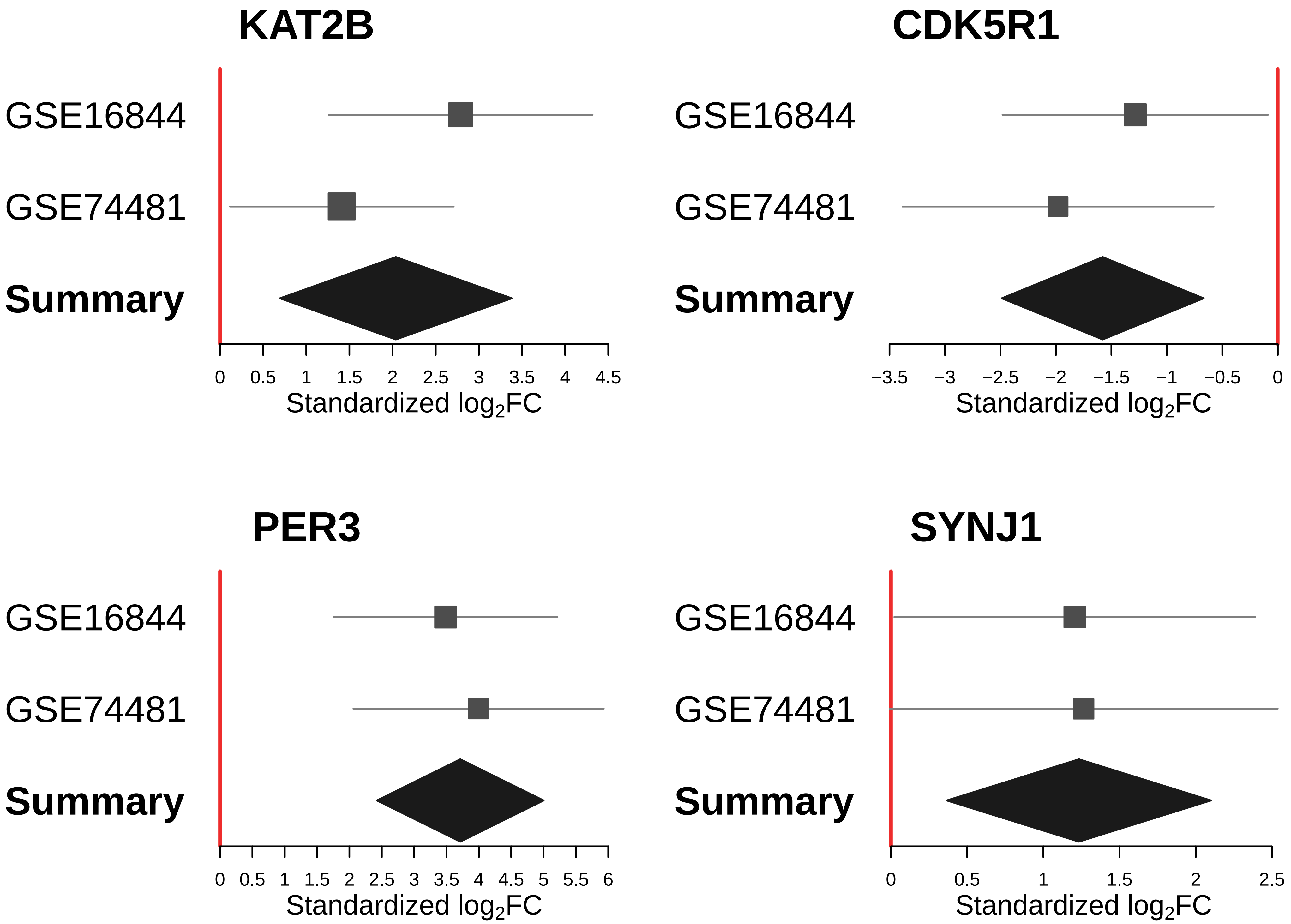
KAT2B
GSE16844
GSE74481
Summary
0 0.5 1 1.5 2 2.5 3 3.5 4 4.5
Standardized log2FC
CDK5R1
GSE16844
GSE74481
Summary
−3.5 −3 −2.5 −2 −1.5 −1 −0.5 0
Standardized log2FC
PER3
GSE16844
GSE74481
Summary
0 0.5 1 1.5 2 2.5 3 3.5 4 4.5 5 5.5 6
Standardized log2FC
SYNJ1
GSE16844
GSE74481
Summary
0 0.5 1 1.5 2 2.5
Standardized log2FC

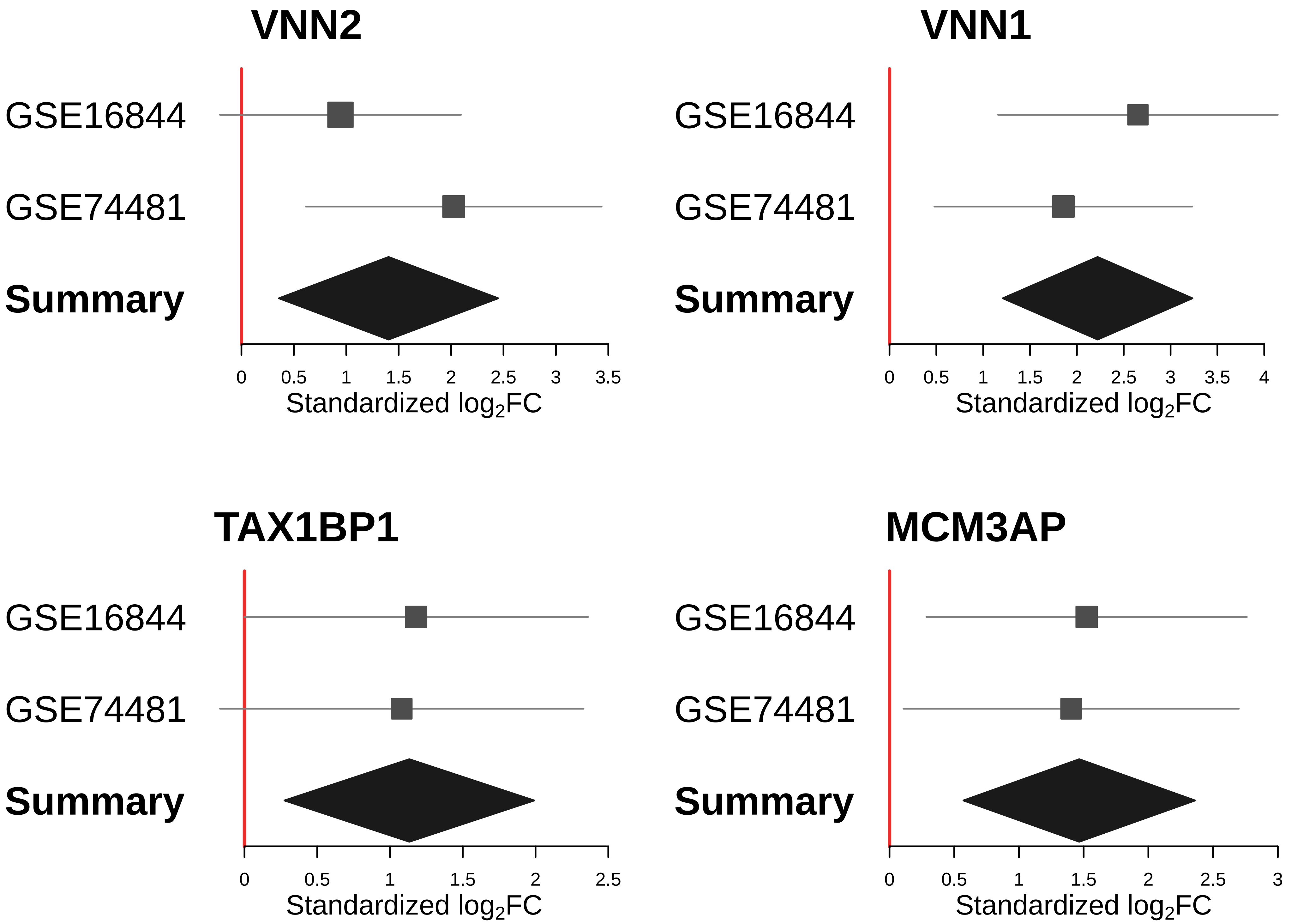

VNN2
GSE16844
GSE74481
Summary
0
0.5
1
1.5
2
2.5
3
3.5
Standardized log2FC
VNN1
GSE16844
GSE74481
Summary
0
0.5
1
1.5
2
2.5
3
3.5
4
Standardized log2FC
TAX1BP1
GSE16844
GSE74481
Summary
0
0.5
1
1.5
2
2.5
Standardized log2FC
MCM3AP
GSE16844
GSE74481
Summary
0
0.5
1
1.5
2
2.5
3
Standardized log2FC

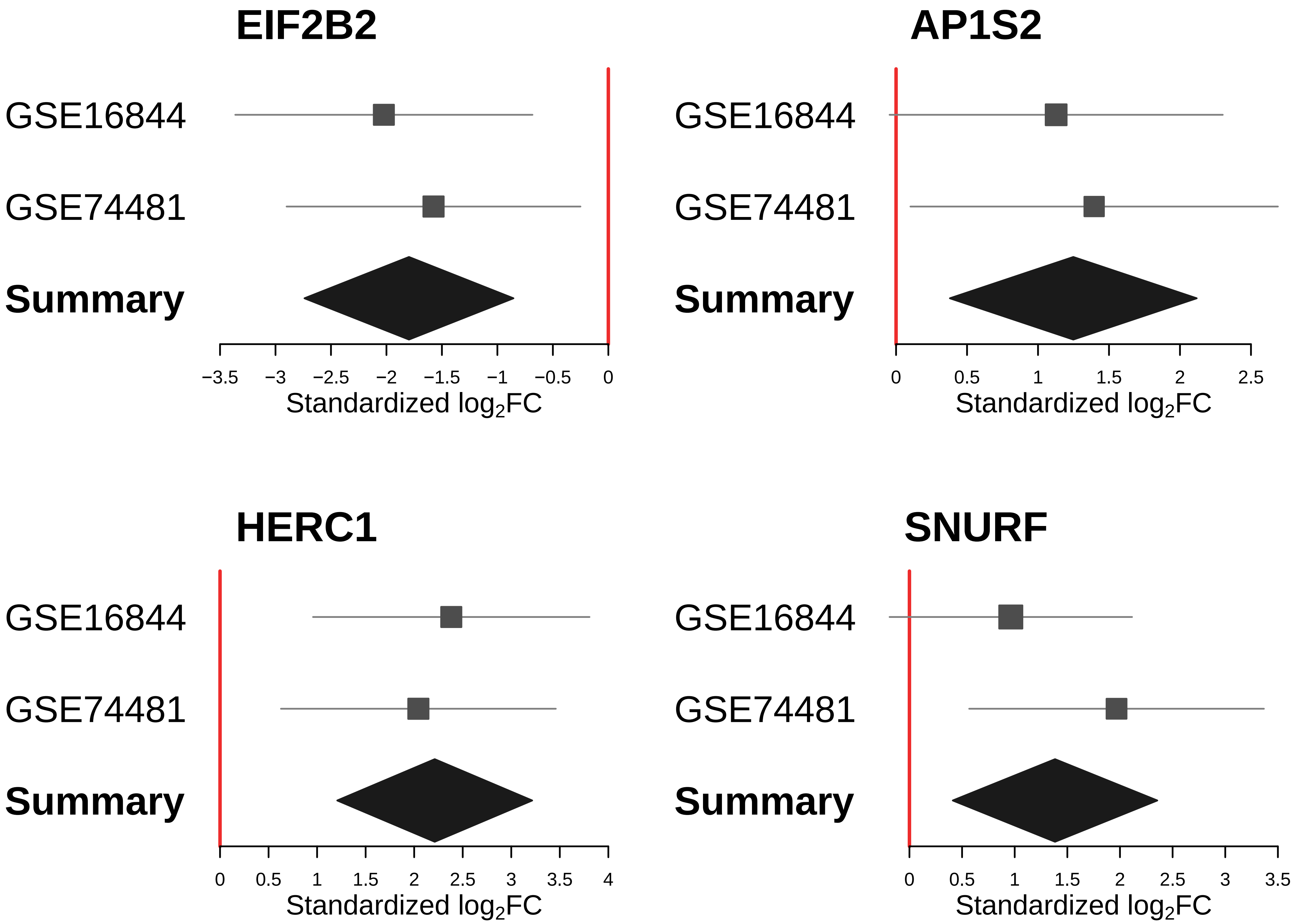
EIF2B2
GSE16844
GSE74481
Summary
−3.5 −3 −2.5 −2 −1.5 −1 −0.5 0
Standardized log2FC
AP1S2
GSE16844
GSE74481
Summary
0 0.5 1 1.5 2 2.5
Standardized log2FC
HERC1
GSE16844
GSE74481
Summary
0 0.5 1 1.5 2 2.5 3 3.5 4
Standardized log2FC
SNURF
GSE16844
GSE74481
Summary
0 0.5 1 1.5 2 2.5 3 3.5
Standardized log2FC

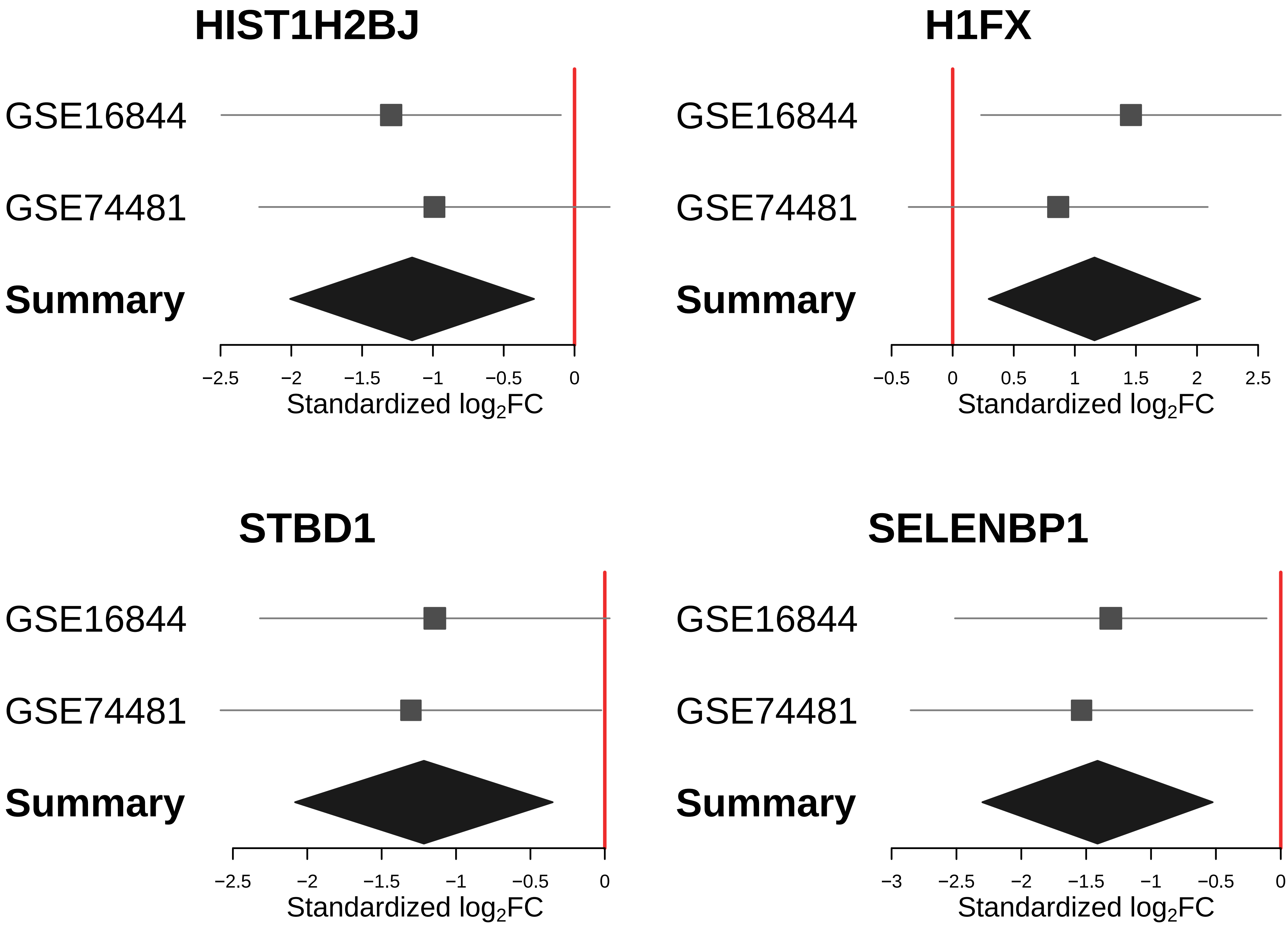
HIST1H2BJ
GSE16844
GSE74481
Summary
−2.5 −2 −1.5 −1 −0.5 0
Standardized log2FC
H1FX
GSE16844
GSE74481
Summary
−0.5 0 0.5 1 1.5 2 2.5
Standardized log2FC
STBD1
GSE16844
GSE74481
Summary
−2.5 −2 −1.5 −1 −0.5 0
Standardized log2FC
SELENBP1
GSE16844
GSE74481
Summary
−3 −2.5 −2 −1.5 −1 −0.5 0
Standardized log2FC

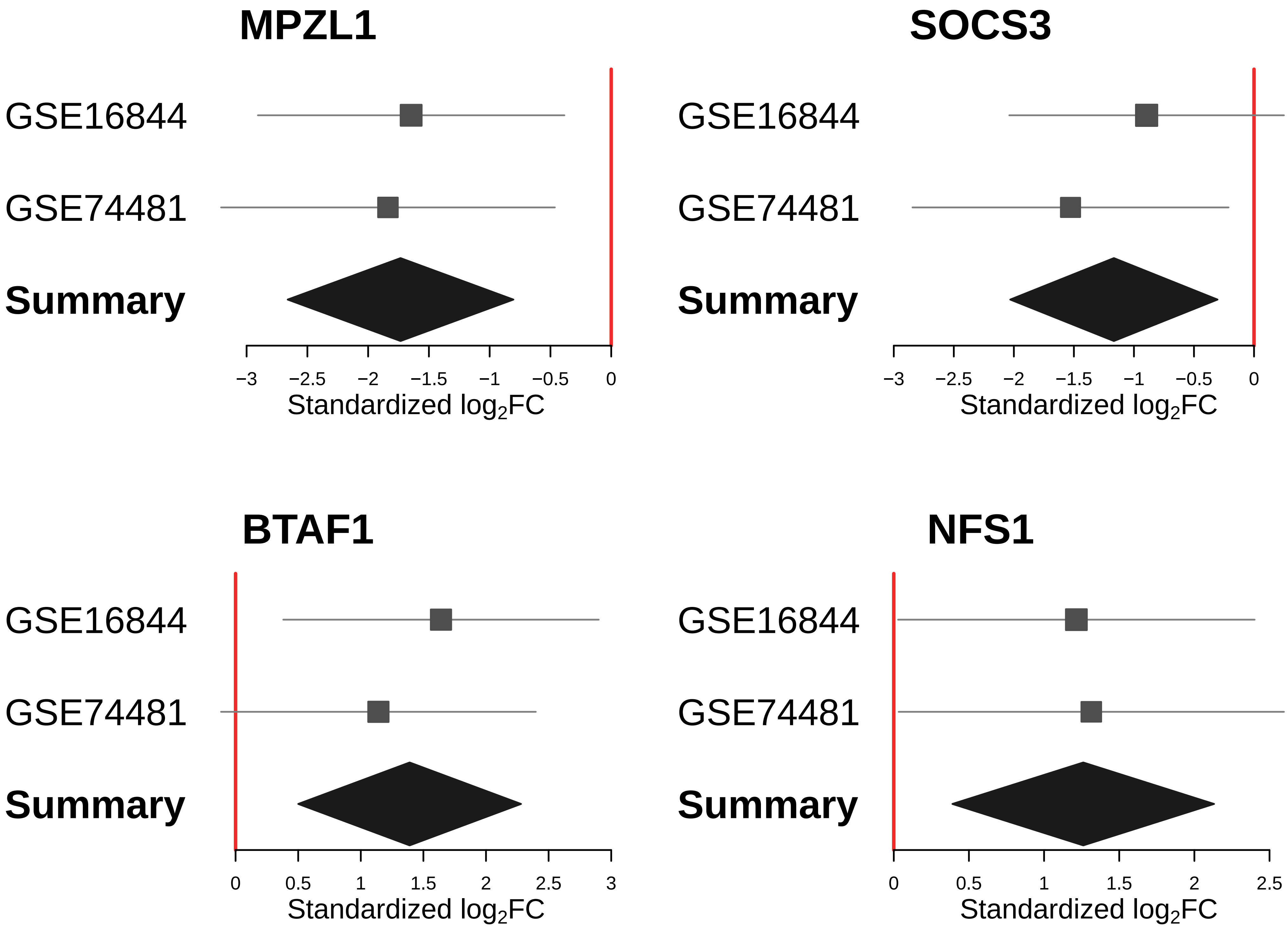

MPZL1
GSE16844
GSE74481
Summary
−3
−2.5
−2
−1.5
−1
−0.5
0
Standardized log2FC
SOCS3
GSE16844
GSE74481
Summary
−3
−2.5
−2
−1.5
−1
−0.5
0
Standardized log2FC
BTAF1
GSE16844
GSE74481
Summary
0
0.5
1
1.5
2
2.5
3
Standardized log2FC
NFS1
GSE16844
GSE74481
Summary
0
0.5
1
1.5
2
2.5
Standardized log2FC

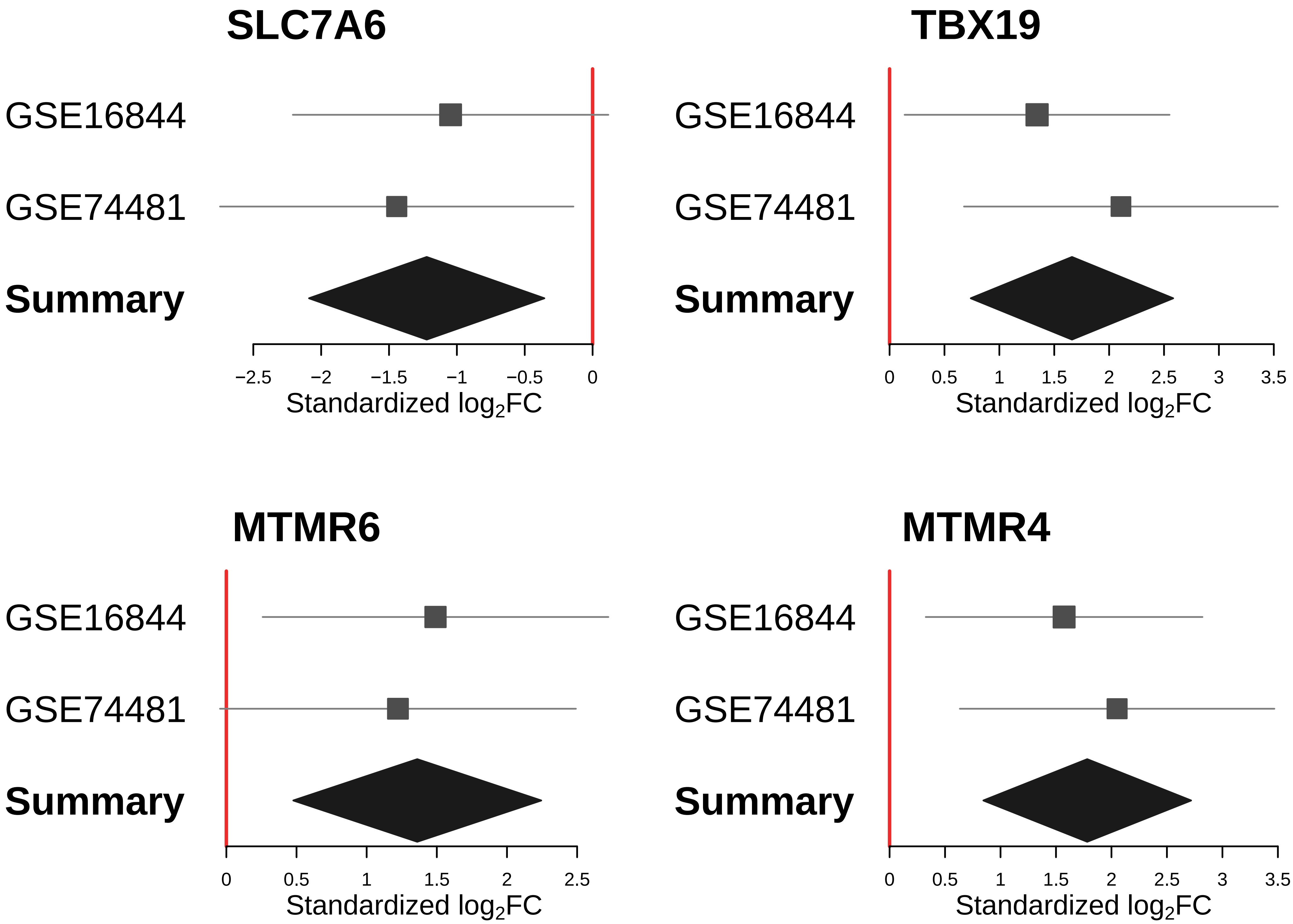
SLC7A6
GSE16844
GSE74481
Summary
−2.5 −2 −1.5 −1 −0.5 0
Standardized log2FC
TBX19
GSE16844
GSE74481
Summary
0 0.5 1 1.5 2 2.5 3 3.5
Standardized log2FC
MTMR6
GSE16844
GSE74481
Summary
0 0.5 1 1.5 2 2.5
Standardized log2FC
MTMR4
GSE16844
GSE74481
Summary
0 0.5 1 1.5 2 2.5 3 3.5
Standardized log2FC

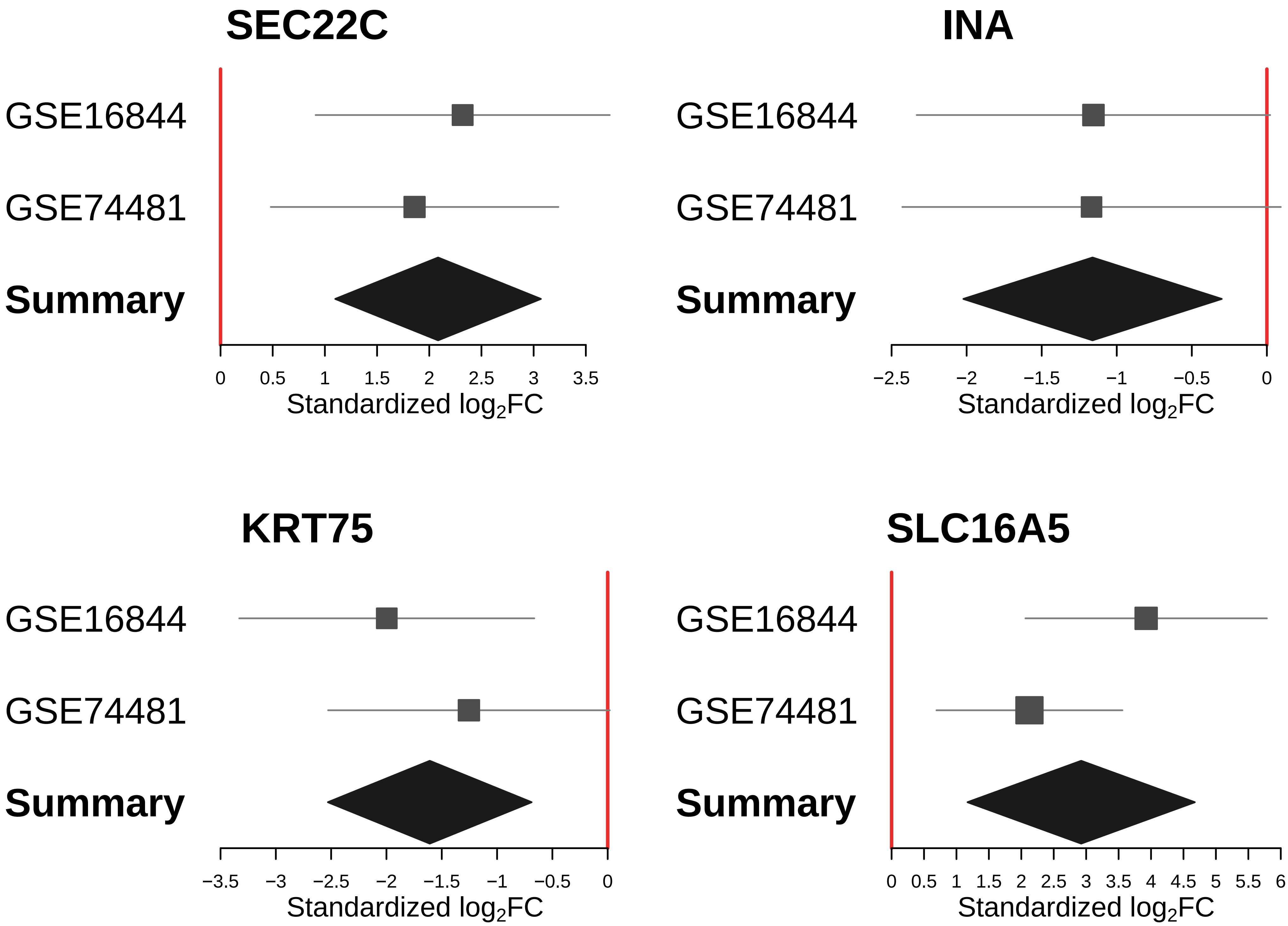

SEC22C
GSE16844
GSE74481
Summary
0
0.5
1
1.5
2
2.5
3
3.5
Standardized log2FC
INA
GSE16844
GSE74481
Summary
−2.5
−2
−1.5
−1
−0.5
0
Standardized log2FC
KRT75
GSE16844
GSE74481
Summary
−3.5
−3
−2.5
−2
−1.5
−1
−0.5
0
Standardized log2FC
SLC16A5
GSE16844
GSE74481
Summary
0
0.5
1
1.5
2
2.5
3
3.5
4
4.5
5
5.5
6
Standardized log2FC

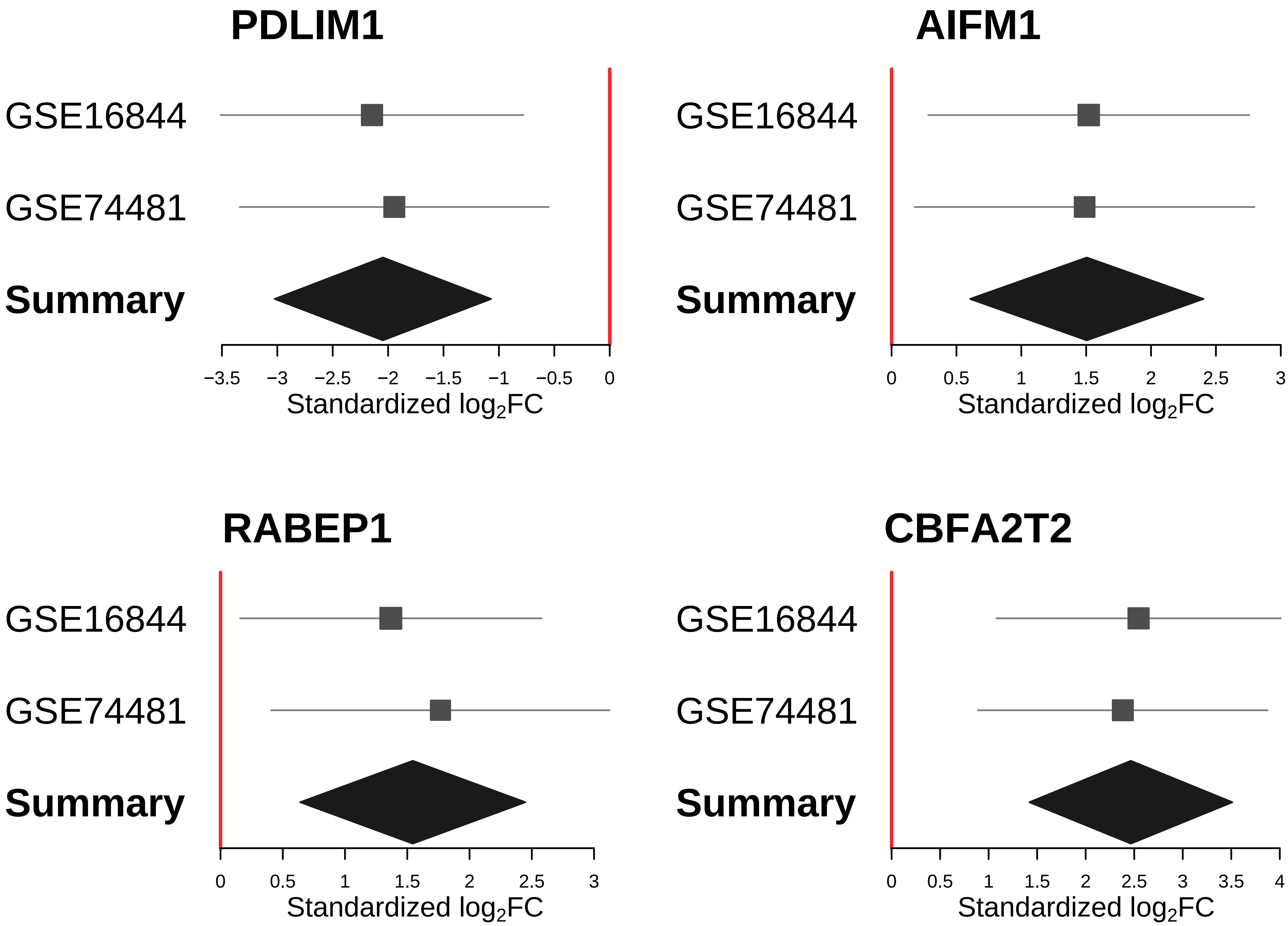
PDLIM1
GSE16844
GSE74481
Summary
−3.5 −3 −2.5 −2 −1.5 −1 −0.5 0
Standardized log2FC
AIFM1
GSE16844
GSE74481
Summary
0 0.5 1 1.5 2 2.5 3
Standardized log2FC
RABEP1
GSE16844
GSE74481
Summary
0 0.5 1 1.5 2 2.5 3
Standardized log2FC
CBFA2T2
GSE16844
GSE74481
Summary
0 0.5 1 1.5 2 2.5 3 3.5 4
Standardized log2FC

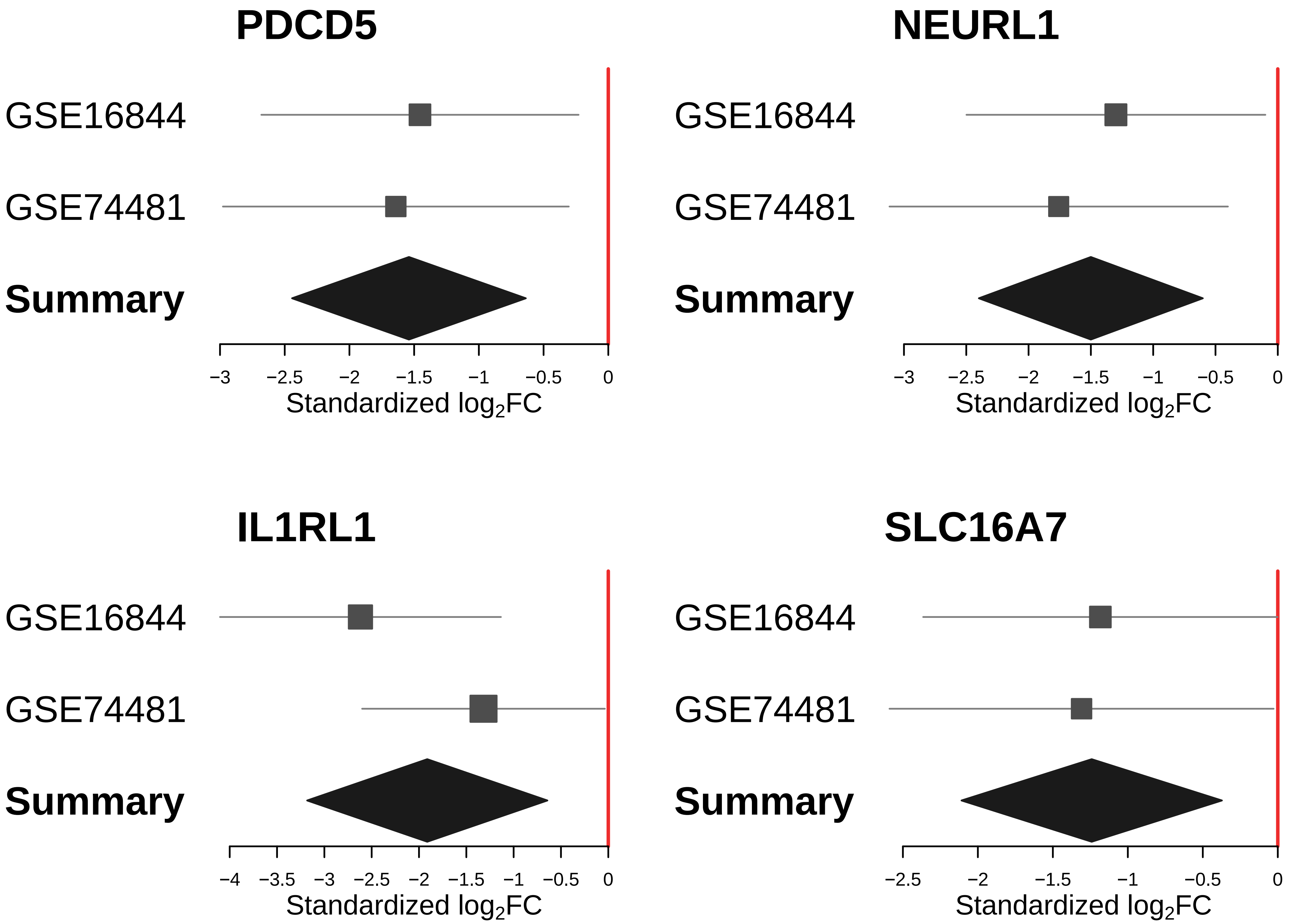
PDCD5
GSE16844
GSE74481
Summary
−3 −2.5 −2 −1.5 −1 −0.5 0
Standardized log2FC
NEURL1
GSE16844
GSE74481
Summary
−3 −2.5 −2 −1.5 −1 −0.5 0
Standardized log2FC
IL1RL1
GSE16844
GSE74481
Summary
−4 −3.5 −3 −2.5 −2 −1.5 −1 −0.5 0
Standardized log2FC
SLC16A7
GSE16844
GSE74481
Summary
−2.5 −2 −1.5 −1 −0.5 0
Standardized log2FC

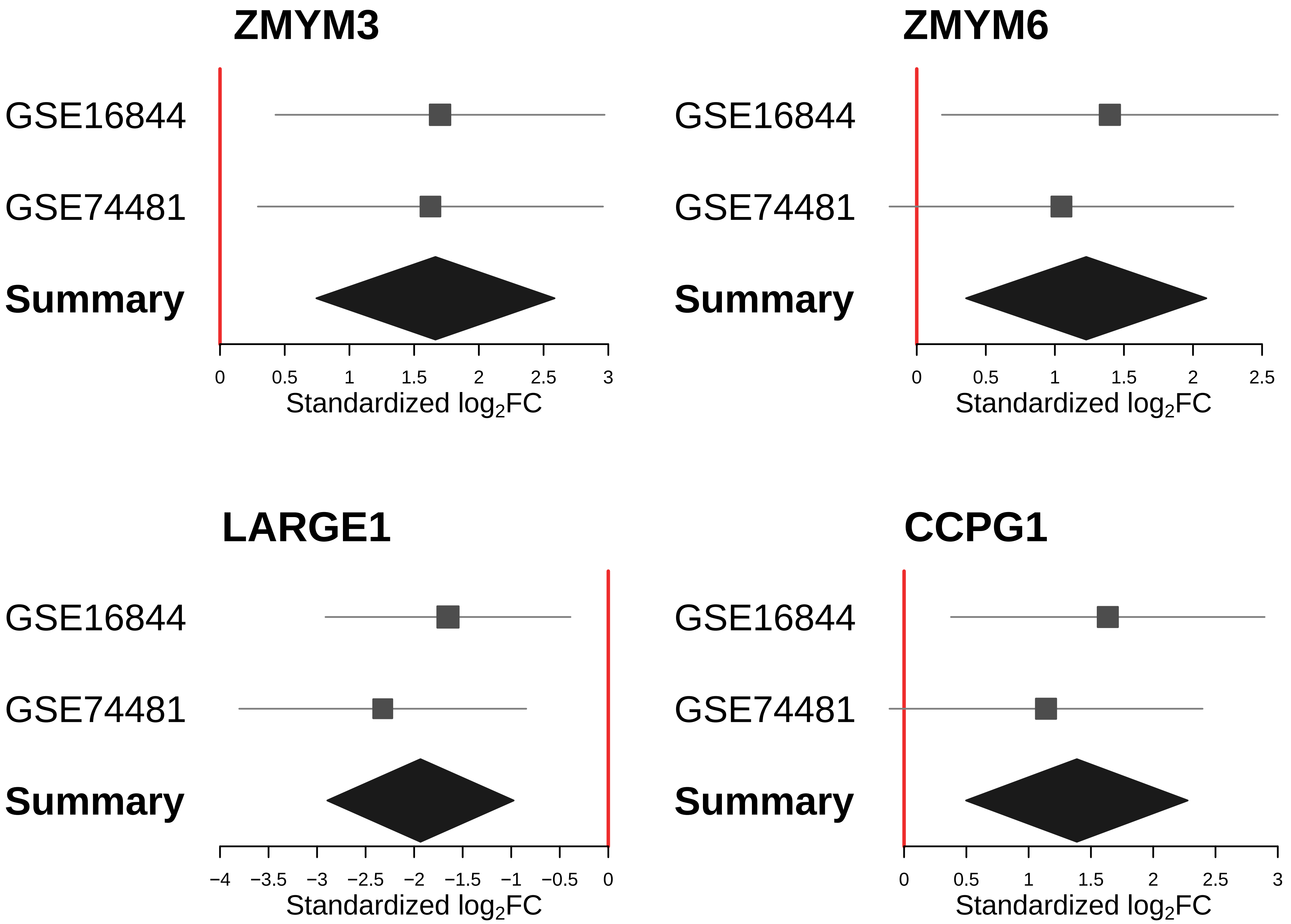
ZMYM3
GSE16844
GSE74481
Summary
0
0.5
1
1.5
2
2.5
3
Standardized log2FC
ZMYM6
GSE16844
GSE74481
Summary
0
0.5
1
1.5
2
2.5
Standardized log2FC
LARGE1
GSE16844
GSE74481
Summary
−4
−3.5
−3
−2.5
−2
−1.5
−1
−0.5
0
Standardized log2FC
CCPG1
GSE16844
GSE74481
Summary
0
0.5
1
1.5
2
2.5
3
Standardized log2FC

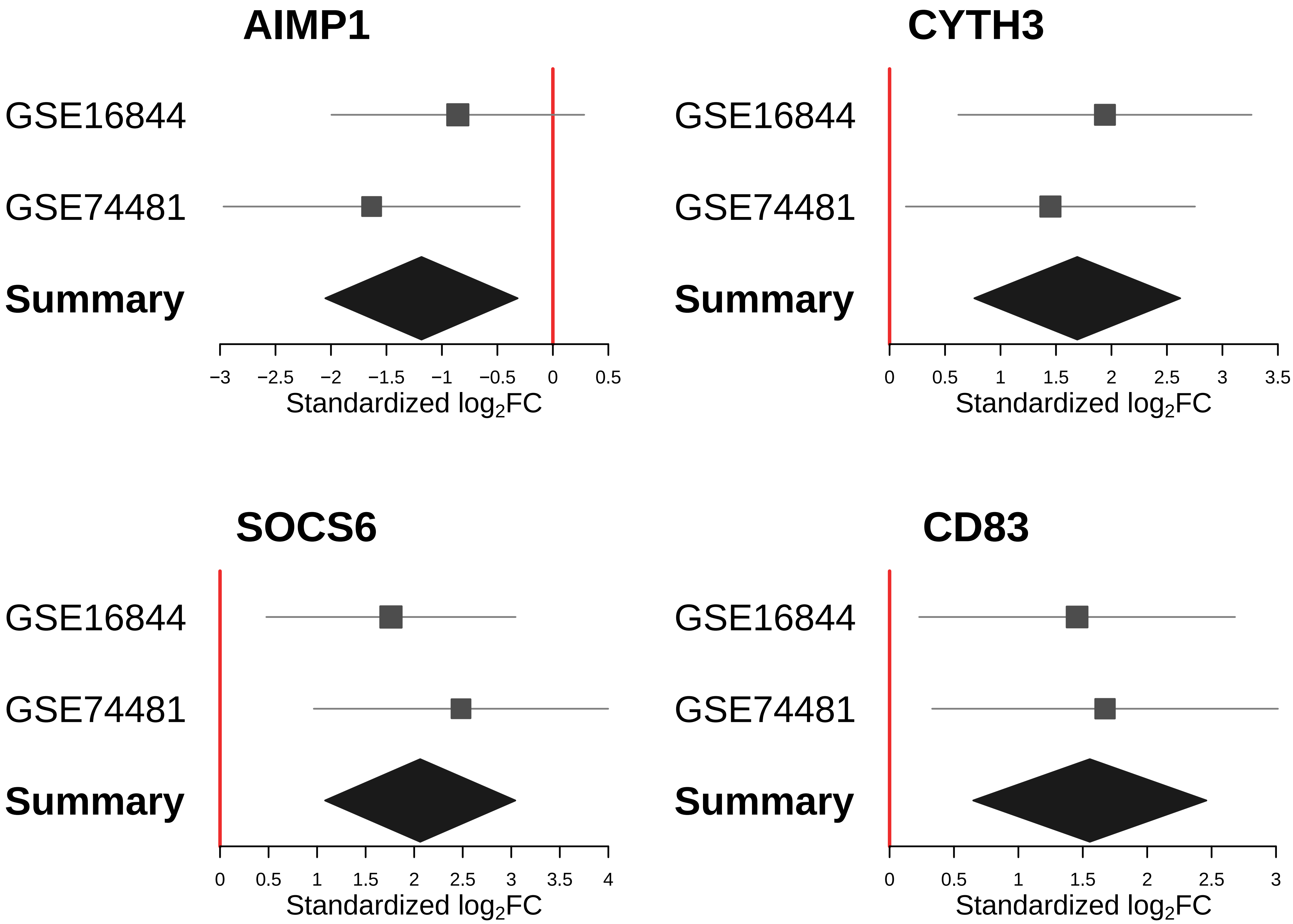
AIMP1
GSE16844
GSE74481
Summary
−3 −2.5 −2 −1.5 −1 −0.5 0 0.5
Standardized log2FC
CYTH3
GSE16844
GSE74481
Summary
0 0.5 1 1.5 2 2.5 3 3.5
Standardized log2FC
SOCS6
GSE16844
GSE74481
Summary
0 0.5 1 1.5 2 2.5 3 3.5 4
Standardized log2FC
CD83
GSE16844
GSE74481
Summary
0 0.5 1 1.5 2 2.5 3
Standardized log2FC

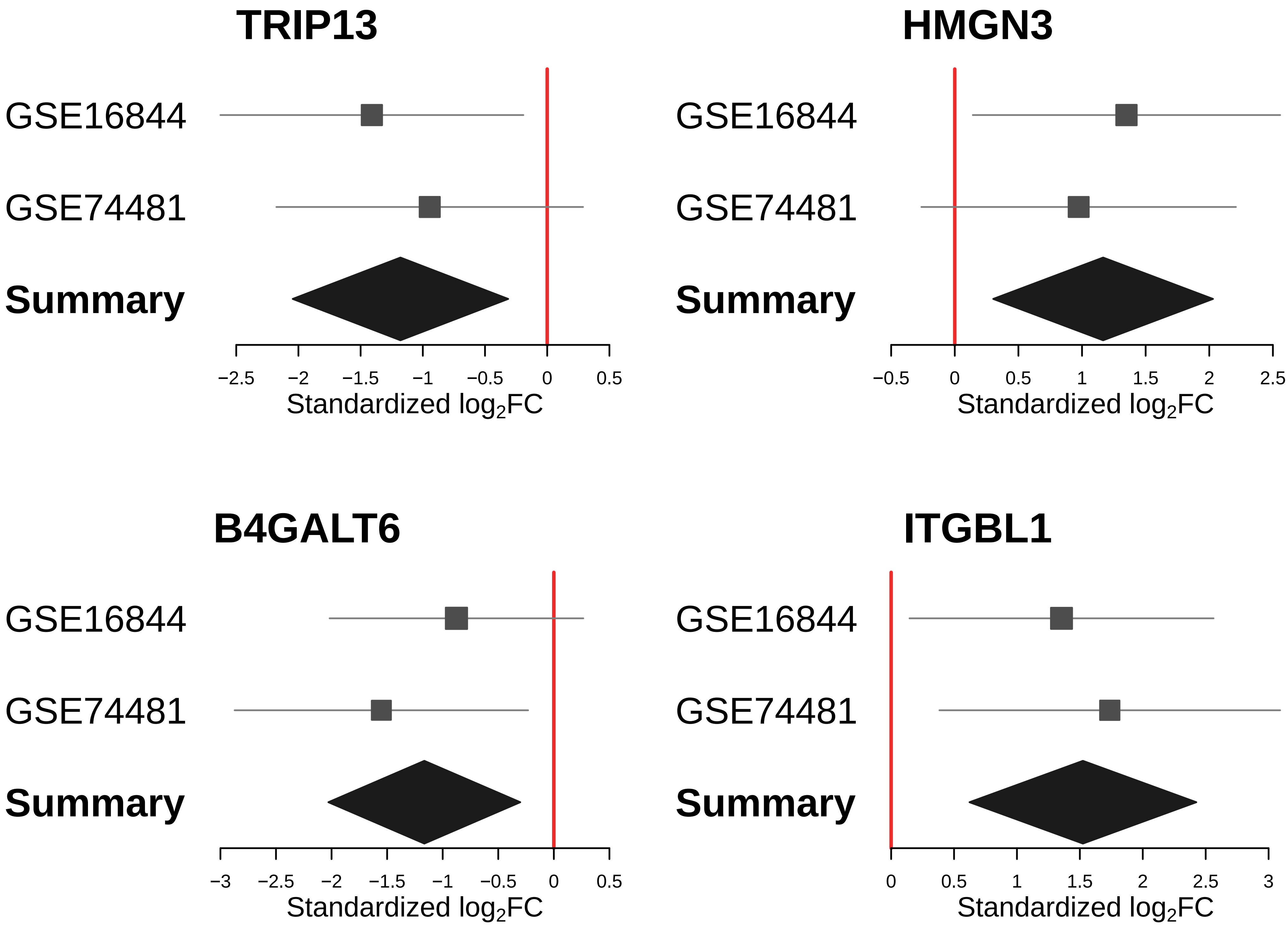
TRIP13
GSE16844
GSE74481
Summary
−2.5 −2 −1.5 −1 −0.5 0 0.5
Standardized log2FC
HMGN3
GSE16844
GSE74481
Summary
−0.5 0 0.5 1 1.5 2 2.5
Standardized log2FC
B4GALT6
GSE16844
GSE74481
Summary
−3 −2.5 −2 −1.5 −1 −0.5 0 0.5
Standardized log2FC
ITGBL1
GSE16844
GSE74481
Summary
0 0.5 1 1.5 2 2.5 3
Standardized log2FC

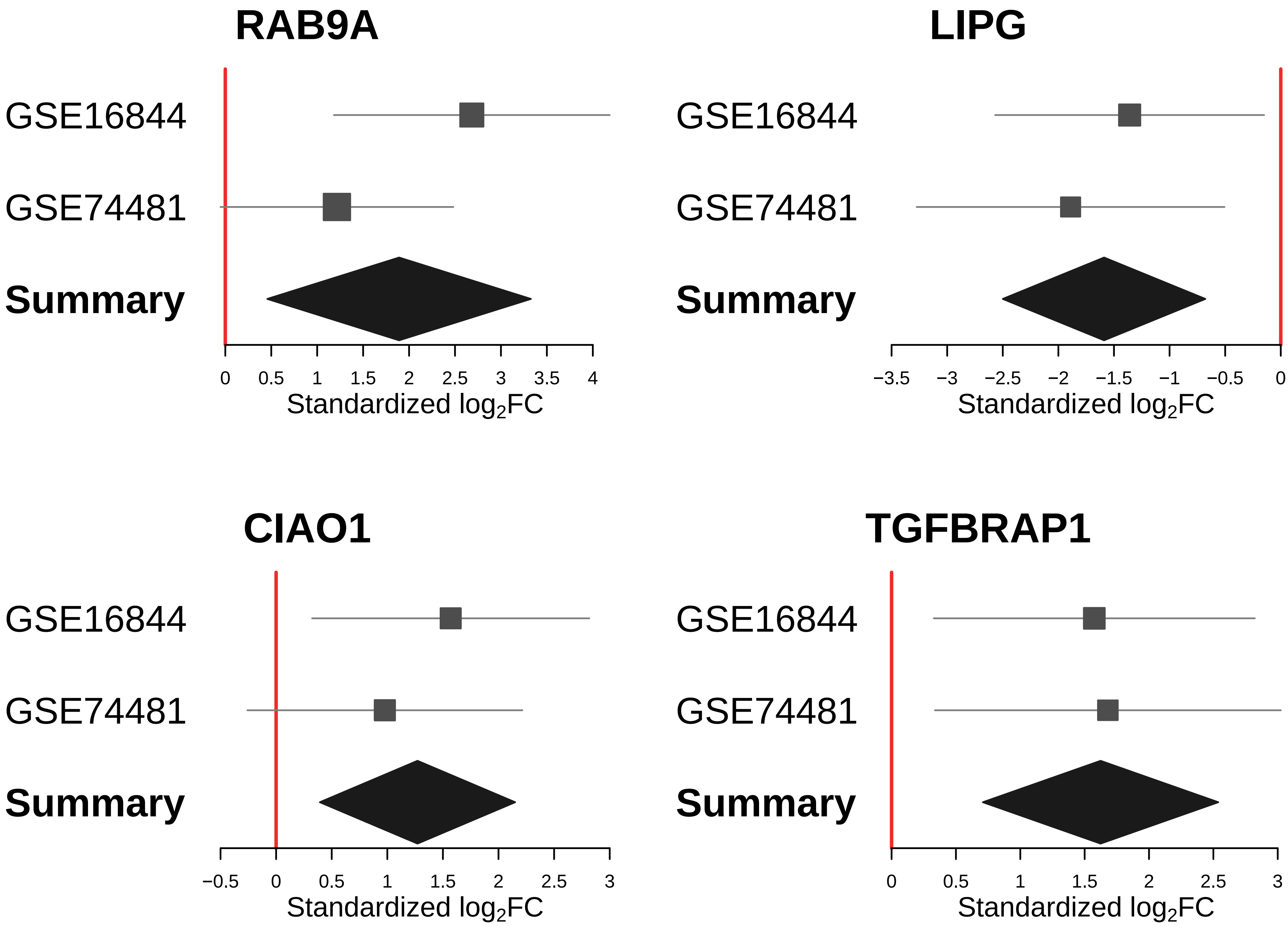
RAB9A
GSE16844
GSE74481
Summary
0
0.5
1
1.5
2
2.5
3
3.5
4
Standardized log2FC
LIPG
GSE16844
GSE74481
Summary
−3.5
−3
−2.5
−2
−1.5
−1
−0.5
0
Standardized log2FC
CIAO1
GSE16844
GSE74481
Summary
−0.5
0
0.5
1
1.5
2
2.5
3
Standardized log2FC
TGFBRAP1
GSE16844
GSE74481
Summary
0
0.5
1
1.5
2
2.5
3
Standardized log2FC

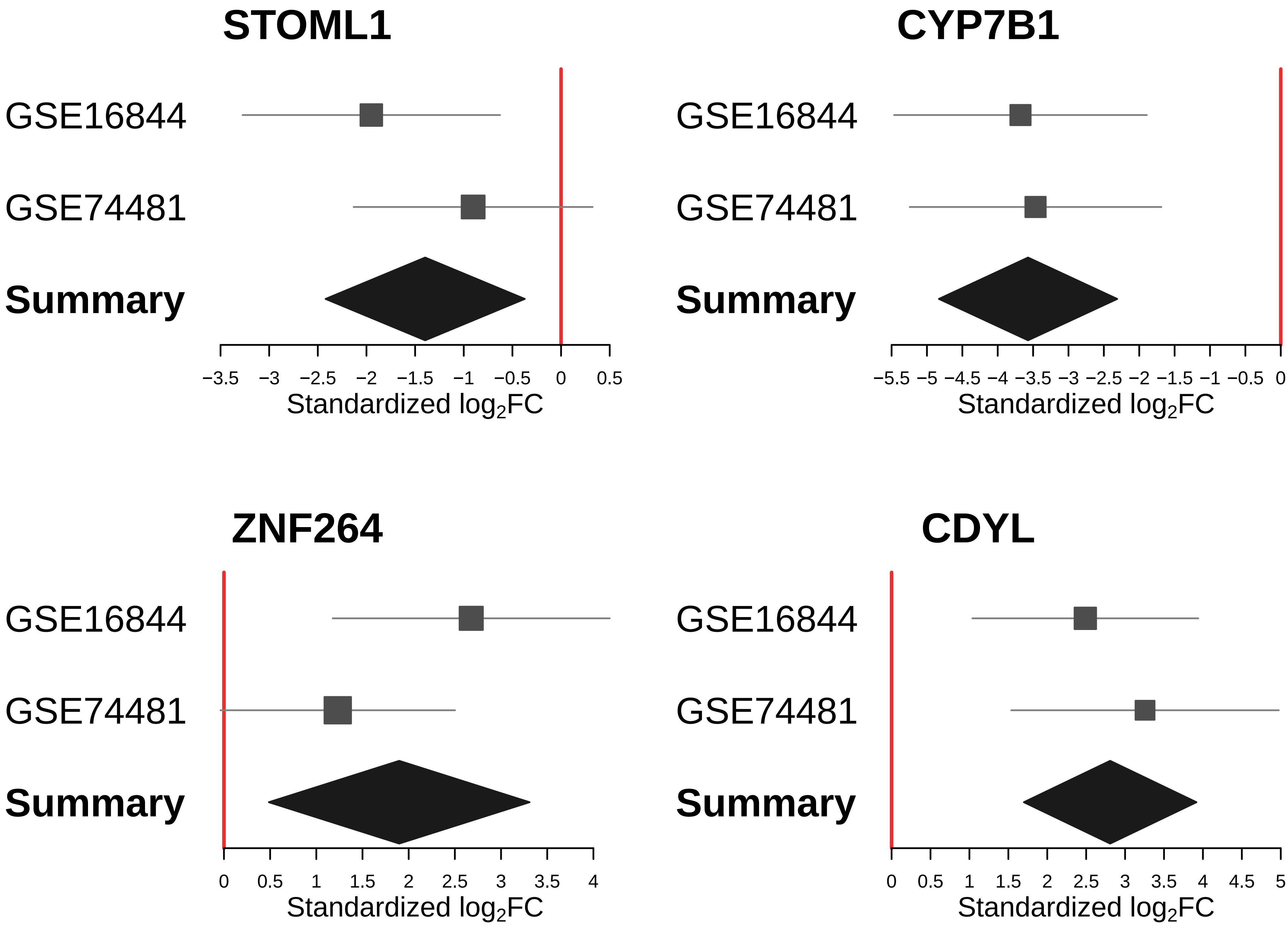
STOML1
GSE16844
GSE74481
Summary
−3.5 −3 −2.5 −2 −1.5 −1 −0.5 0 0.5
Standardized log2FC
CYP7B1
GSE16844
GSE74481
Summary
−5.5 −5 −4.5 −4 −3.5 −3 −2.5 −2 −1.5 −1 −0.5 0
Standardized log2FC
ZNF264
GSE16844
GSE74481
Summary
0 0.5 1 1.5 2 2.5 3 3.5 4
Standardized log2FC
CDYL
GSE16844
GSE74481
Summary
0 0.5 1 1.5 2 2.5 3 3.5 4 4.5 5
Standardized log2FC

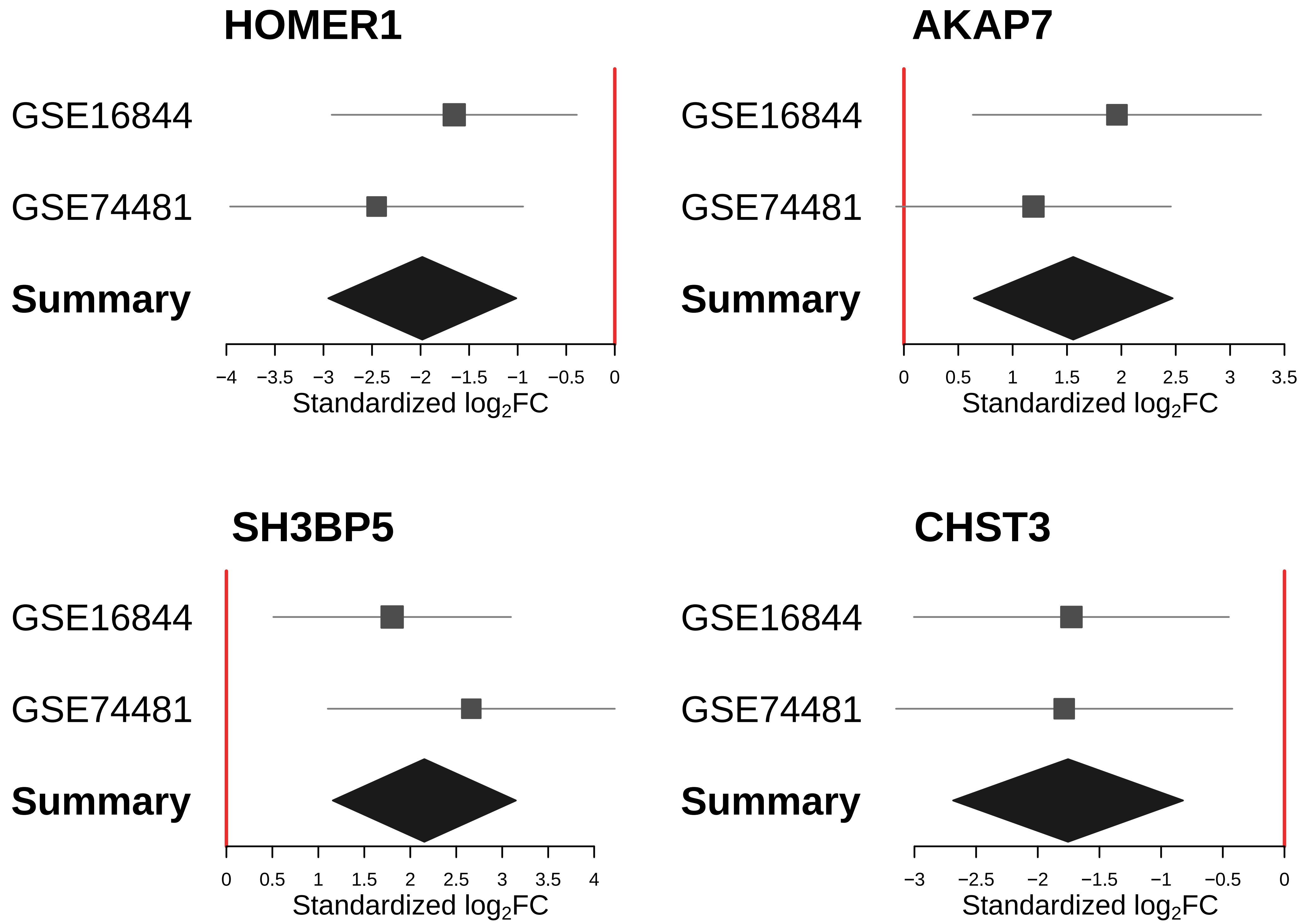
HOMER1
GSE16844
GSE74481
Summary
−4 −3.5 −3 −2.5 −2 −1.5 −1 −0.5 0
Standardized log2FC
AKAP7
GSE16844
GSE74481
Summary
0 0.5 1 1.5 2 2.5 3 3.5
Standardized log2FC
SH3BP5
GSE16844
GSE74481
Summary
0 0.5 1 1.5 2 2.5 3 3.5 4
Standardized log2FC
CHST3
GSE16844
GSE74481
Summary
−3 −2.5 −2 −1.5 −1 −0.5 0
Standardized log2FC

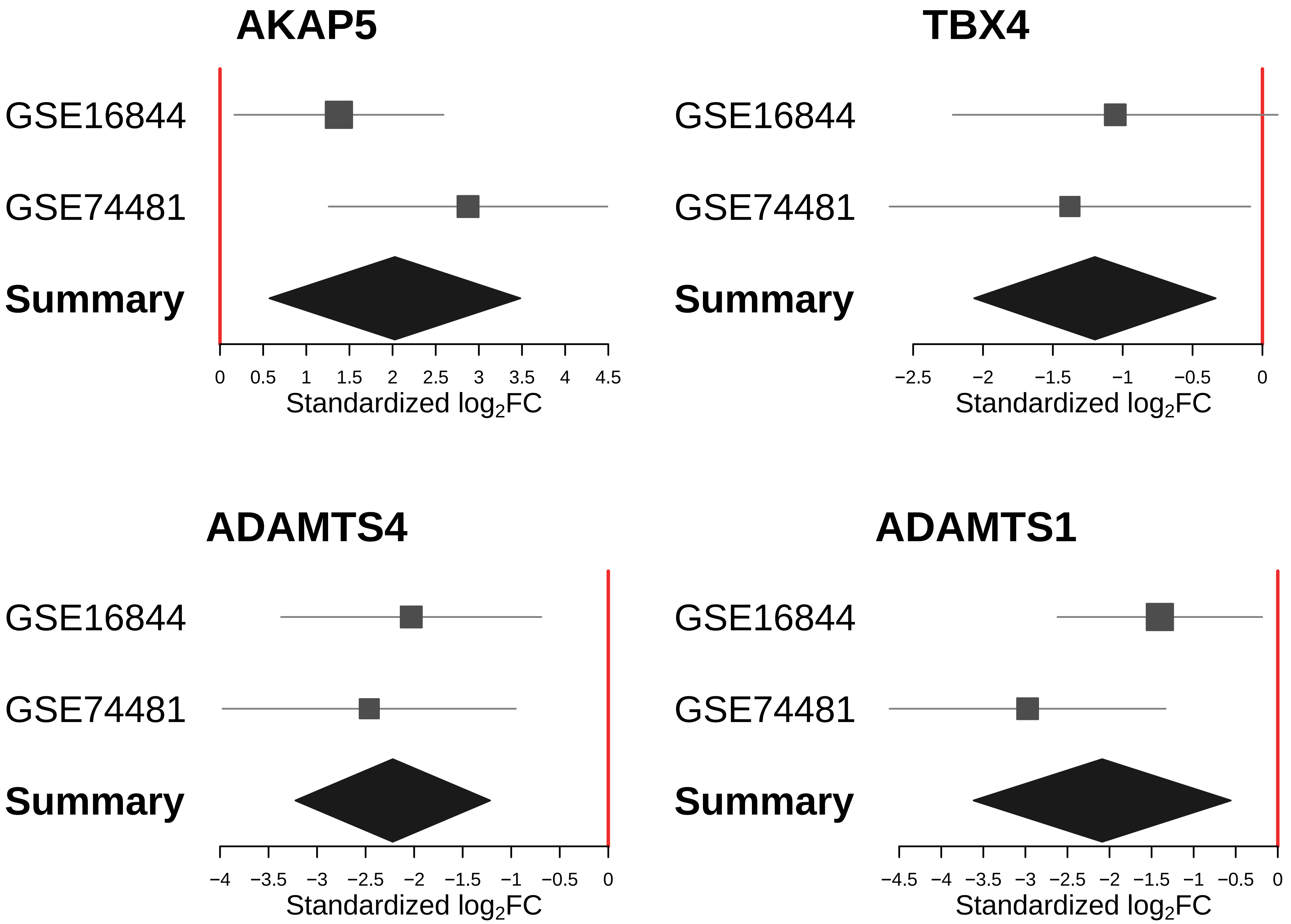
AKAP5
GSE16844
GSE74481
Summary
0 0.5 1 1.5 2 2.5 3 3.5 4 4.5
Standardized log2FC
TBX4
GSE16844
GSE74481
Summary
−2.5 −2 −1.5 −1 −0.5 0
Standardized log2FC
ADAMTS4
GSE16844
GSE74481
Summary
−4 −3.5 −3 −2.5 −2 −1.5 −1 −0.5 0
Standardized log2FC
ADAMTS1
GSE16844
GSE74481
Summary
−4.5 −4 −3.5 −3 −2.5 −2 −1.5 −1 −0.5 0
Standardized log2FC

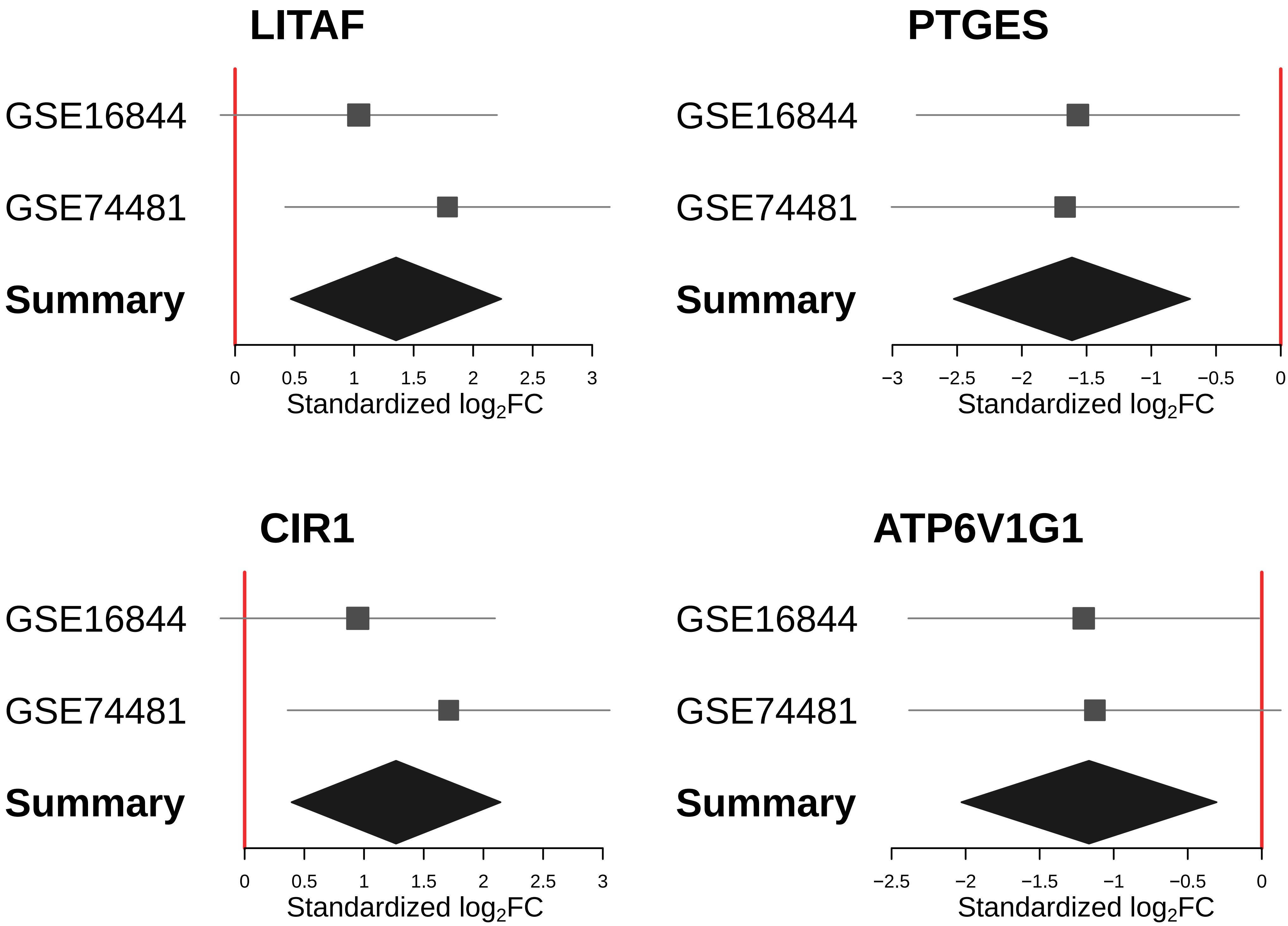
LITAF
GSE16844
GSE74481
Summary
0
0.5
1
1.5
2
2.5
3
Standardized log2FC
PTGES
GSE16844
GSE74481
Summary
−3
−2.5
−2
−1.5
−1
−0.5
0
Standardized log2FC
CIR1
GSE16844
GSE74481
Summary
0
0.5
1
1.5
2
2.5
3
Standardized log2FC
ATP6V1G1
GSE16844
GSE74481
Summary
−2.5
−2
−1.5
−1
−0.5
0
Standardized log2FC

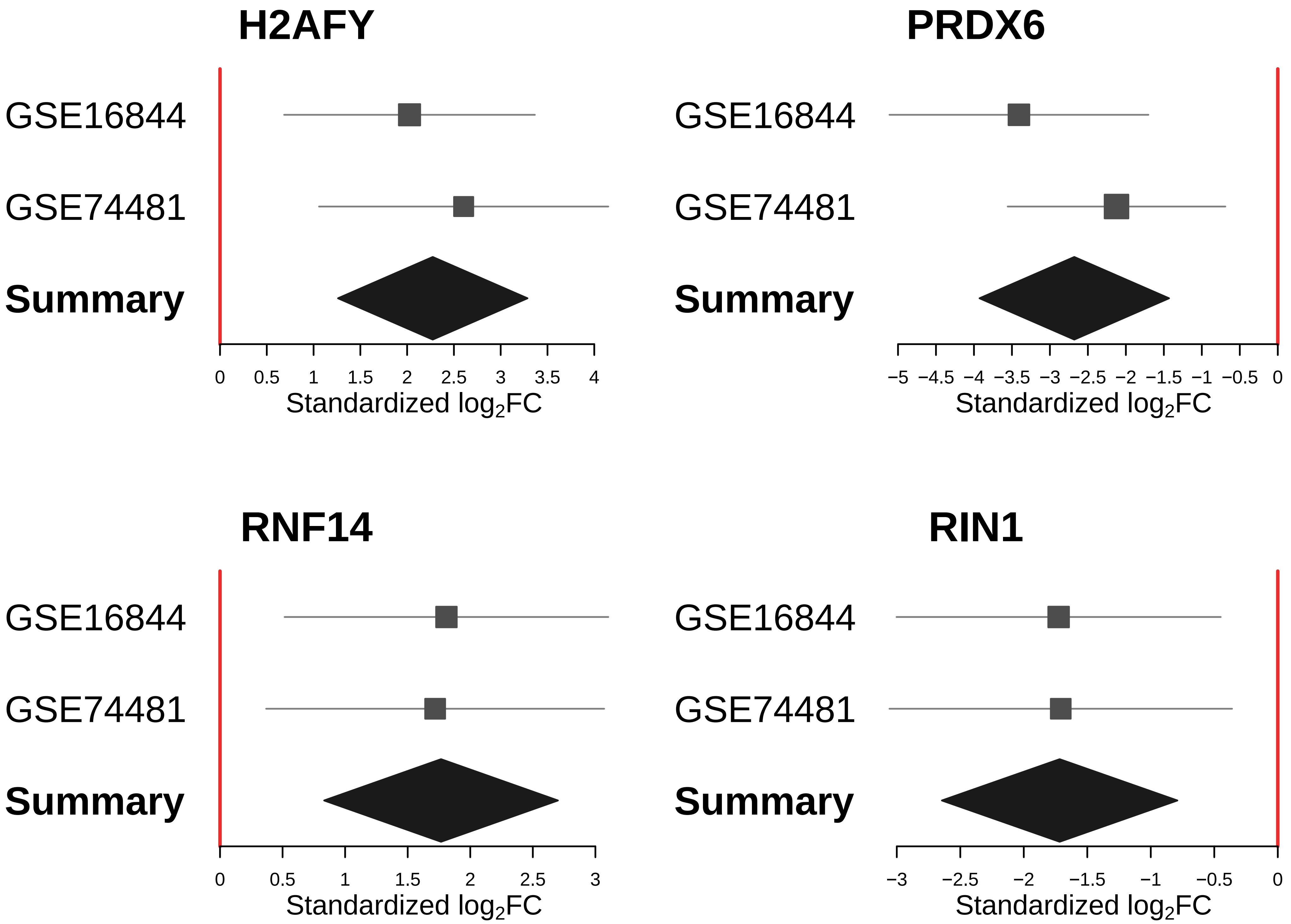
H2AFY
GSE16844
GSE74481
Summary
0 0.5 1 1.5 2 2.5 3 3.5 4
Standardized log2FC
PRDX6
GSE16844
GSE74481
Summary
−5 −4.5 −4 −3.5 −3 −2.5 −2 −1.5 −1 −0.5 0
Standardized log2FC
RNF14
GSE16844
GSE74481
Summary
0 0.5 1 1.5 2 2.5 3
Standardized log2FC
RIN1
GSE16844
GSE74481
Summary
−3 −2.5 −2 −1.5 −1 −0.5 0
Standardized log2FC

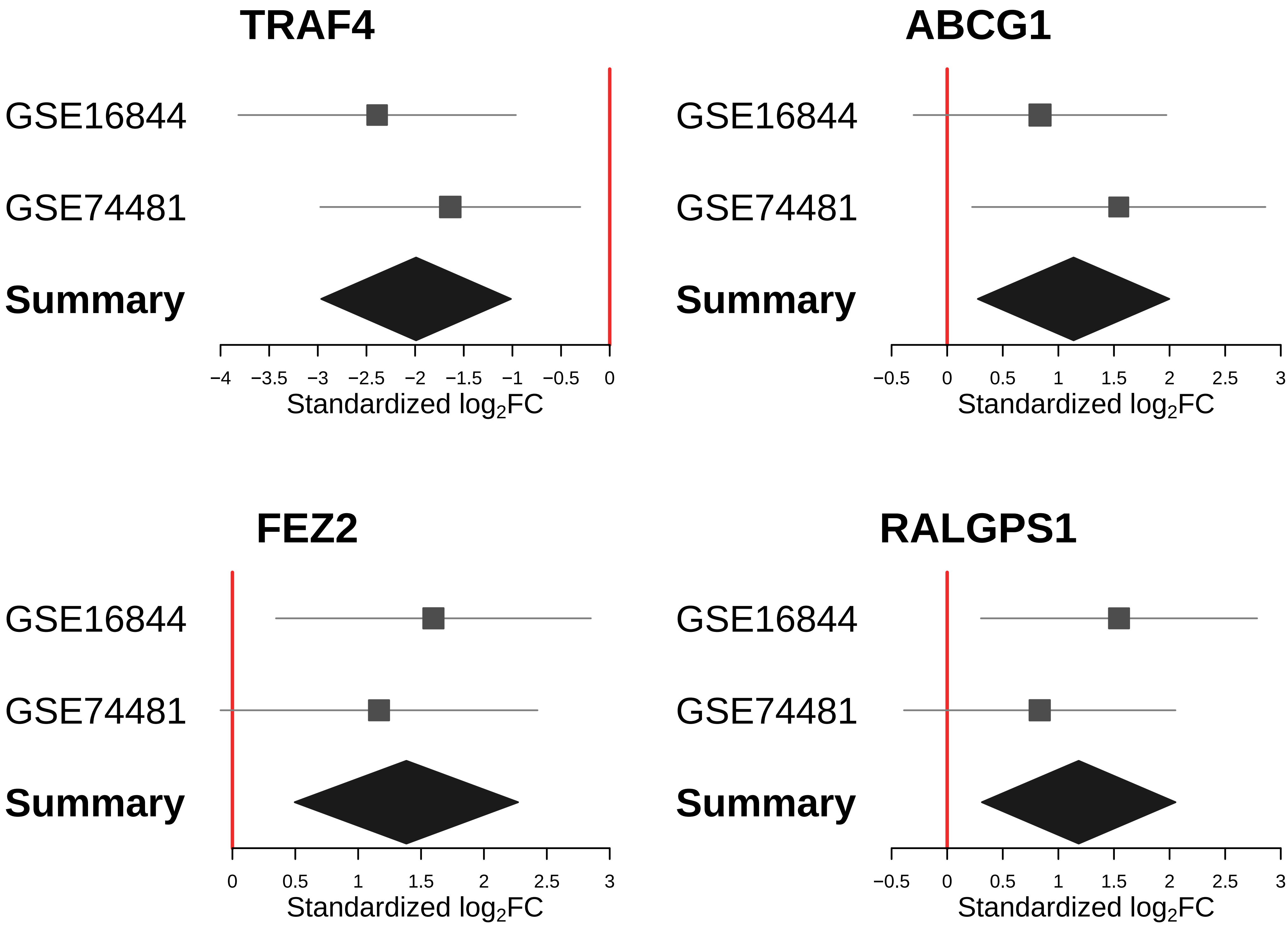
TRAF4
GSE16844
GSE74481
Summary
−4
−3.5
−3
−2.5
−2
−1.5
−1
−0.5
0
Standardized log2FC
ABCG1
GSE16844
GSE74481
Summary
−0.5
0
0.5
1
1.5
2
2.5
3
Standardized log2FC
FEZ2
GSE16844
GSE74481
Summary
0
0.5
1
1.5
2
2.5
3
Standardized log2FC
RALGPS1
GSE16844
GSE74481
Summary
−0.5
0
0.5
1
1.5
2
2.5
3
Standardized log2FC

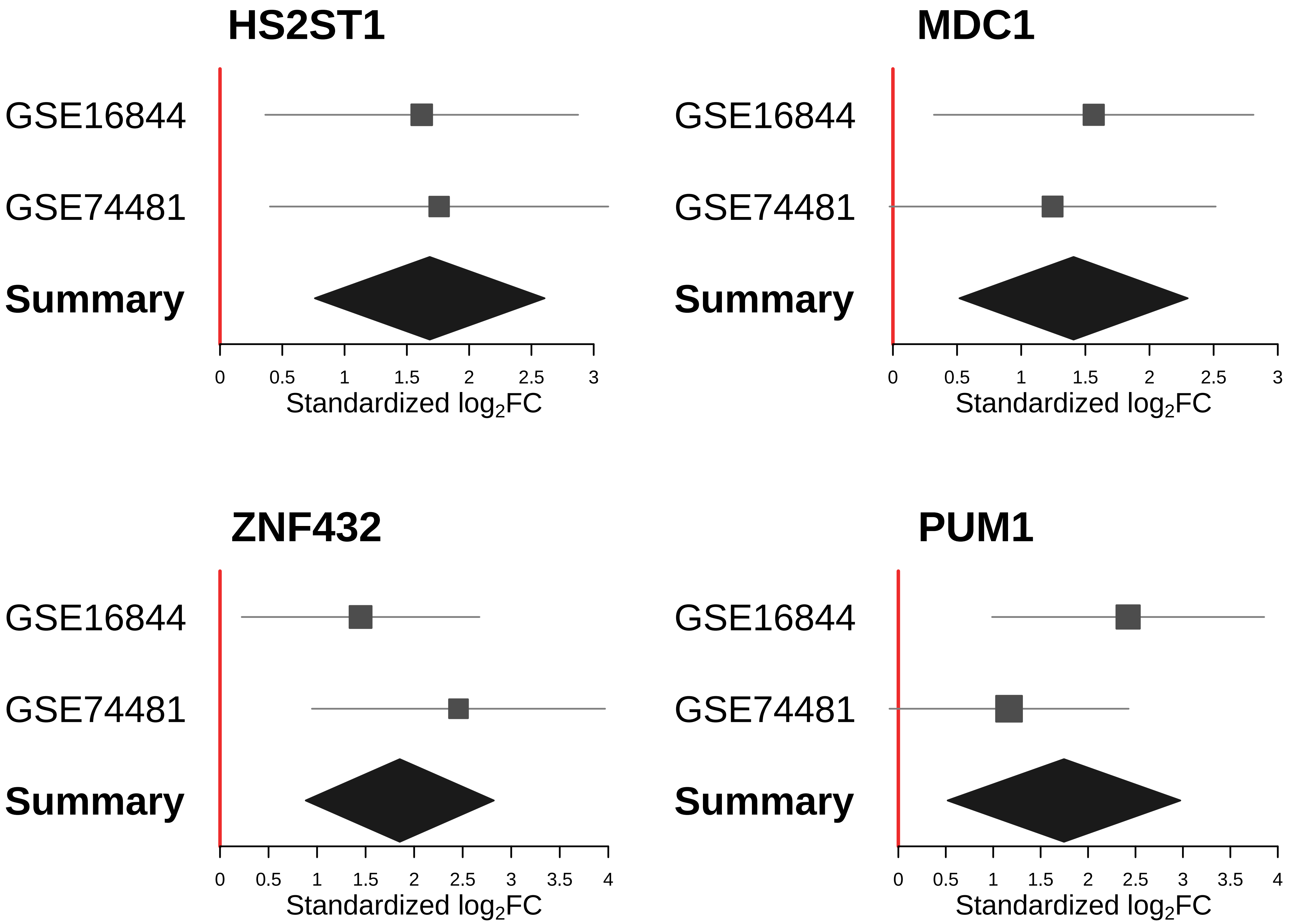
HS2ST1
MDC1
ZNF432
PUM1
GSE16844
GSE74481
Summary
0
0.5
1
1.5
2
2.5
3
3.5
4
Standardized log2FC

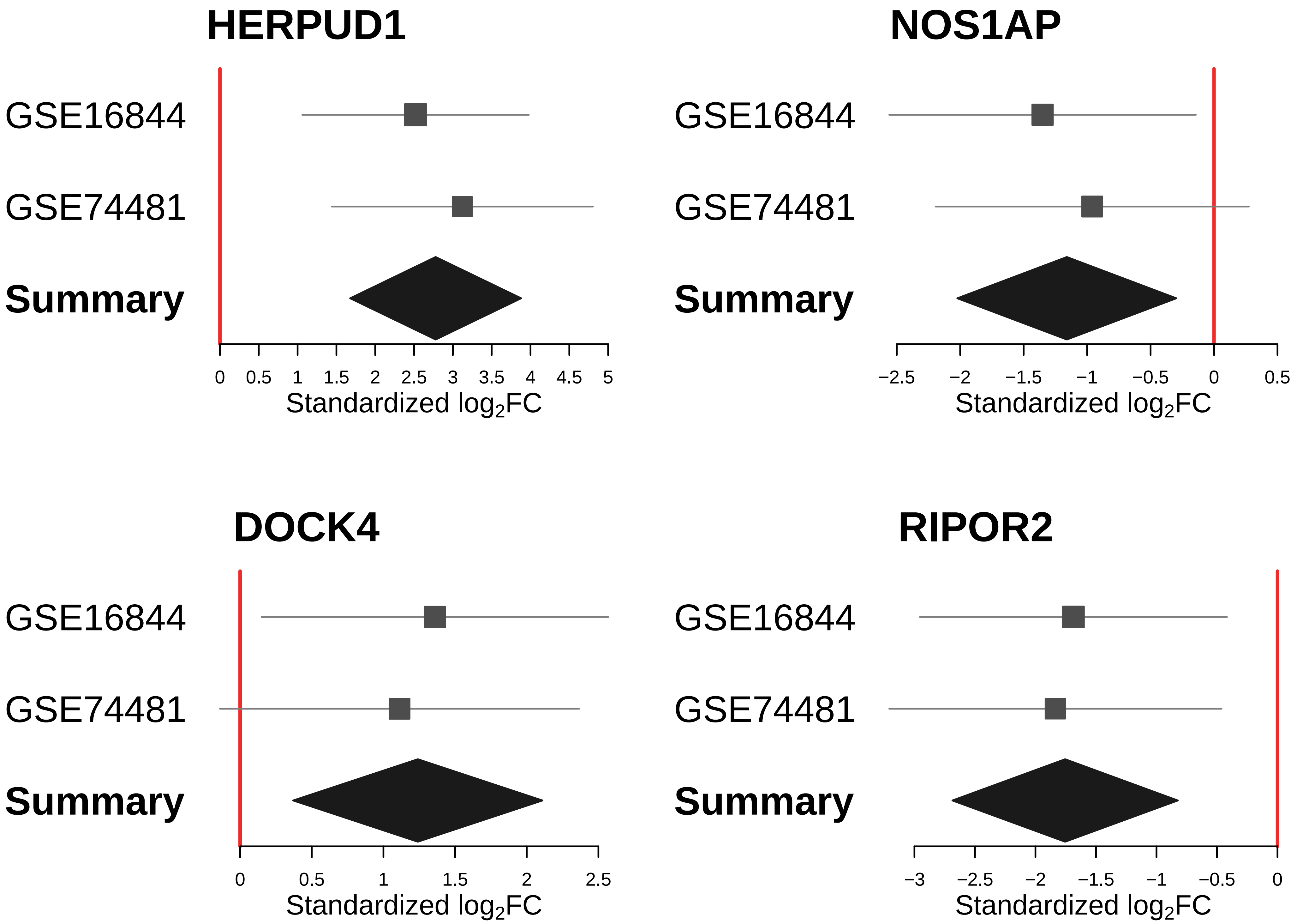
HERPUD1
GSE16844
GSE74481
Summary
0 0.5 1 1.5 2 2.5 3 3.5 4 4.5 5
Standardized log2FC
NOS1AP
GSE16844
GSE74481
Summary
−2.5 −2 −1.5 −1 −0.5 0 0.5
Standardized log2FC
DOCK4
GSE16844
GSE74481
Summary
0 0.5 1 1.5 2 2.5
Standardized log2FC
RIPOR2
GSE16844
GSE74481
Summary
−3 −2.5 −2 −1.5 −1 −0.5 0
Standardized log2FC

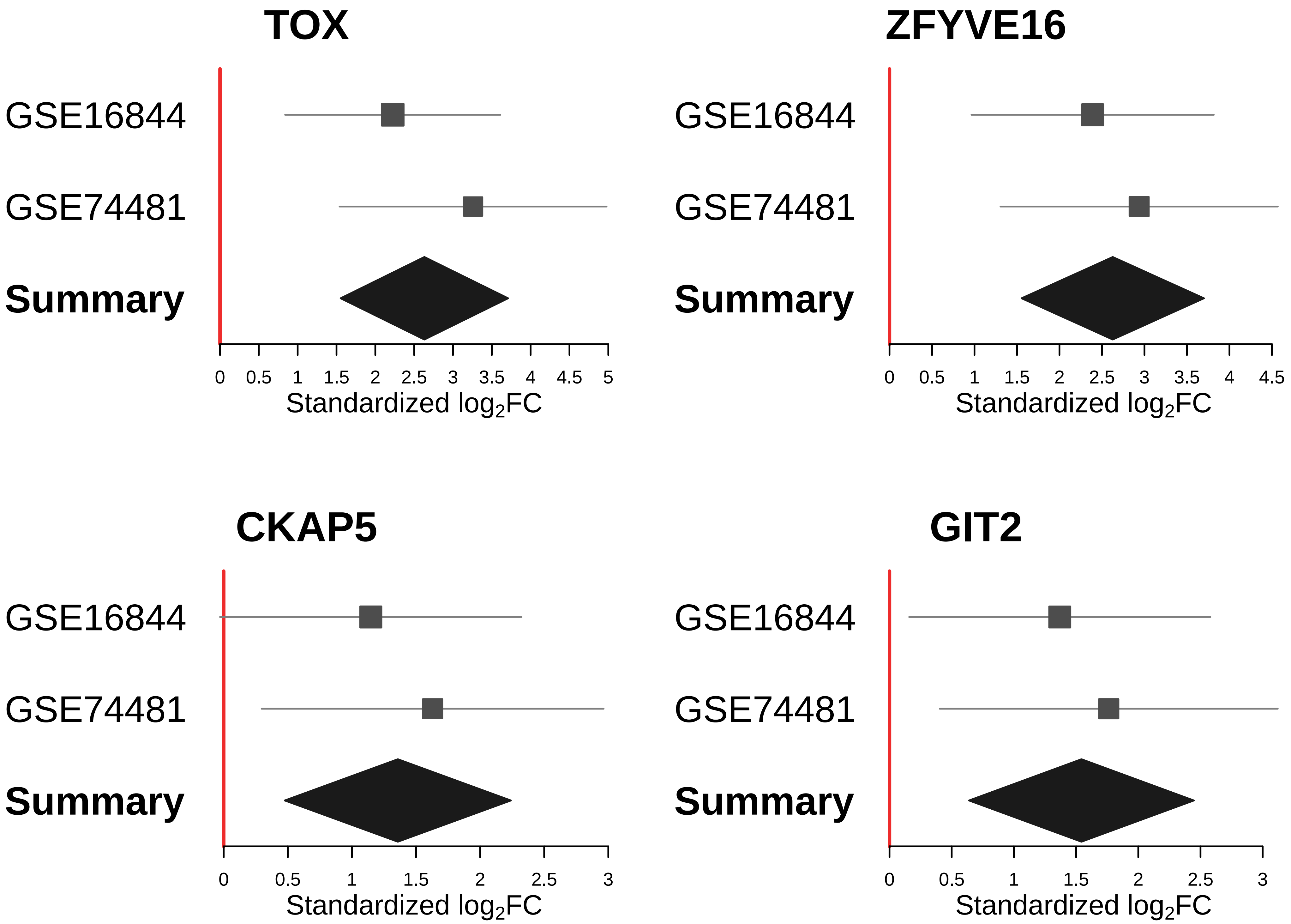
TOX
GSE16844
GSE74481
Summary
0 0.5 1 1.5 2 2.5 3 3.5 4 4.5 5
Standardized $log_2FC$
ZFYVE16
GSE16844
GSE74481
Summary
0 0.5 1 1.5 2 2.5 3 3.5 4 4.5
Standardized $log_2FC$
CKAP5
GSE16844
GSE74481
Summary
0 0.5 1 1.5 2 2.5 3
Standardized $log_2FC$
GIT2
GSE16844
GSE74481
Summary
0 0.5 1 1.5 2 2.5 3
Standardized $log_2FC$

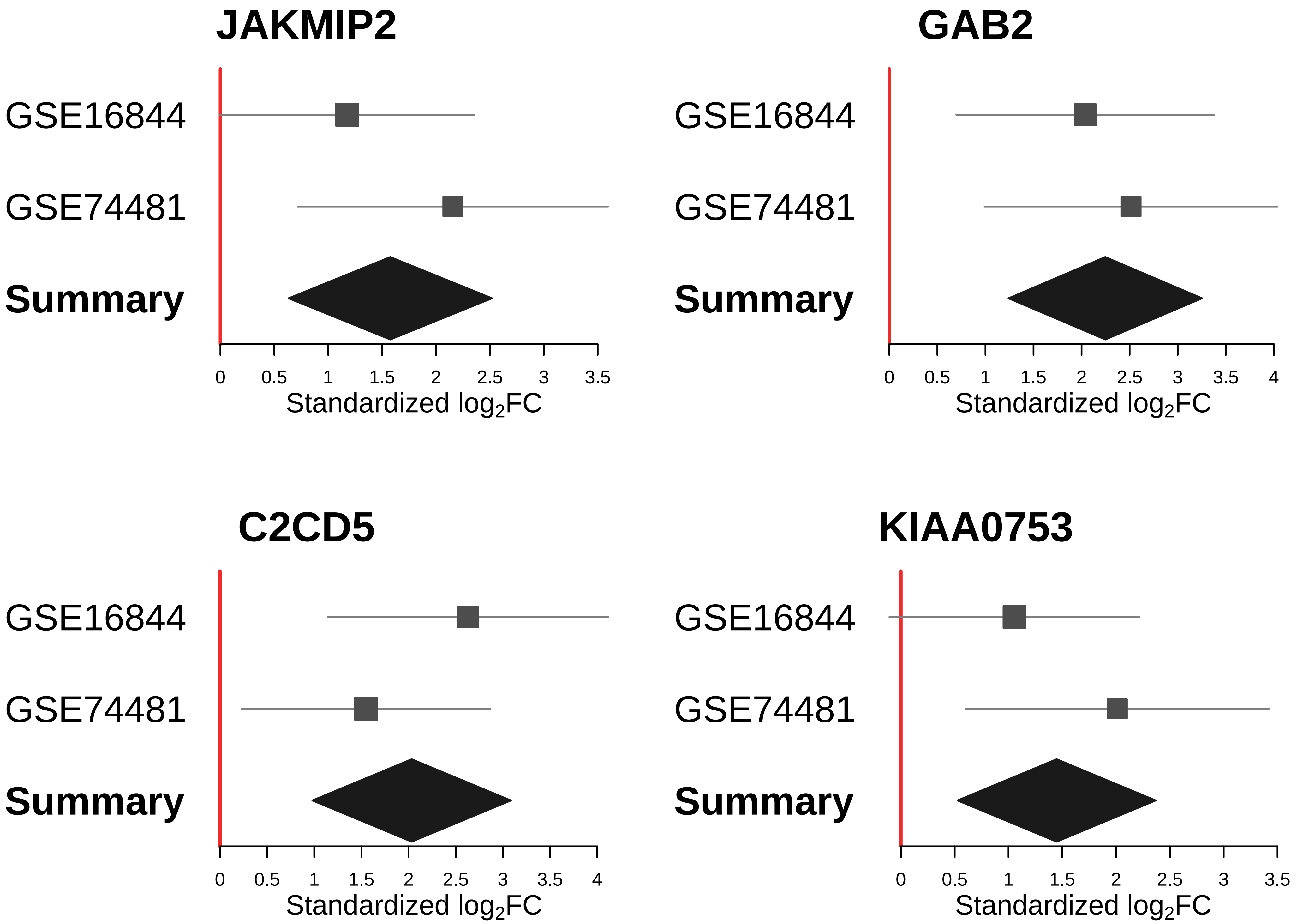
JAKMIP2
GSE16844
GSE74481
Summary
0
0.5
1
1.5
2
2.5
3
3.5
Standardized $log_2FC$
GAB2
GSE16844
GSE74481
Summary
0
0.5
1
1.5
2
2.5
3
3.5
4
Standardized $log_2FC$
C2CD5
GSE16844
GSE74481
Summary
0
0.5
1
1.5
2
2.5
3
3.5
4
Standardized $log_2FC$
KIAA0753
GSE16844
GSE74481
Summary
0
0.5
1
1.5
2
2.5
3
3.5
Standardized $log_2FC$

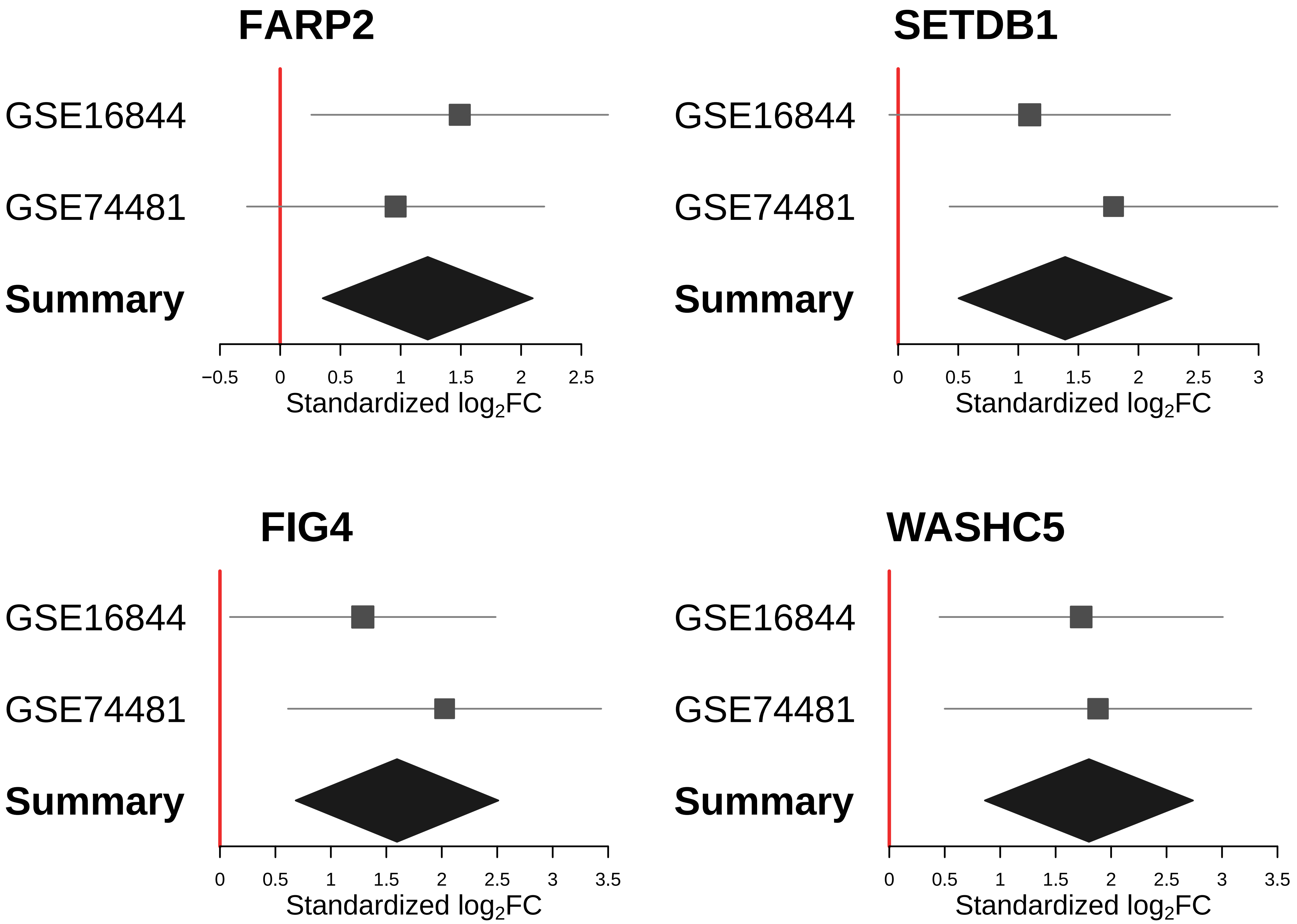
FARP2
GSE16844
GSE74481
Summary
−0.5 0 0.5 1 1.5 2 2.5
Standardized log2FC
SETDB1
GSE16844
GSE74481
Summary
0 0.5 1 1.5 2 2.5 3
Standardized log2FC
FIG4
GSE16844
GSE74481
Summary
0 0.5 1 1.5 2 2.5 3 3.5
Standardized log2FC
WASHC5
GSE16844
GSE74481
Summary
0 0.5 1 1.5 2 2.5 3 3.5
Standardized log2FC

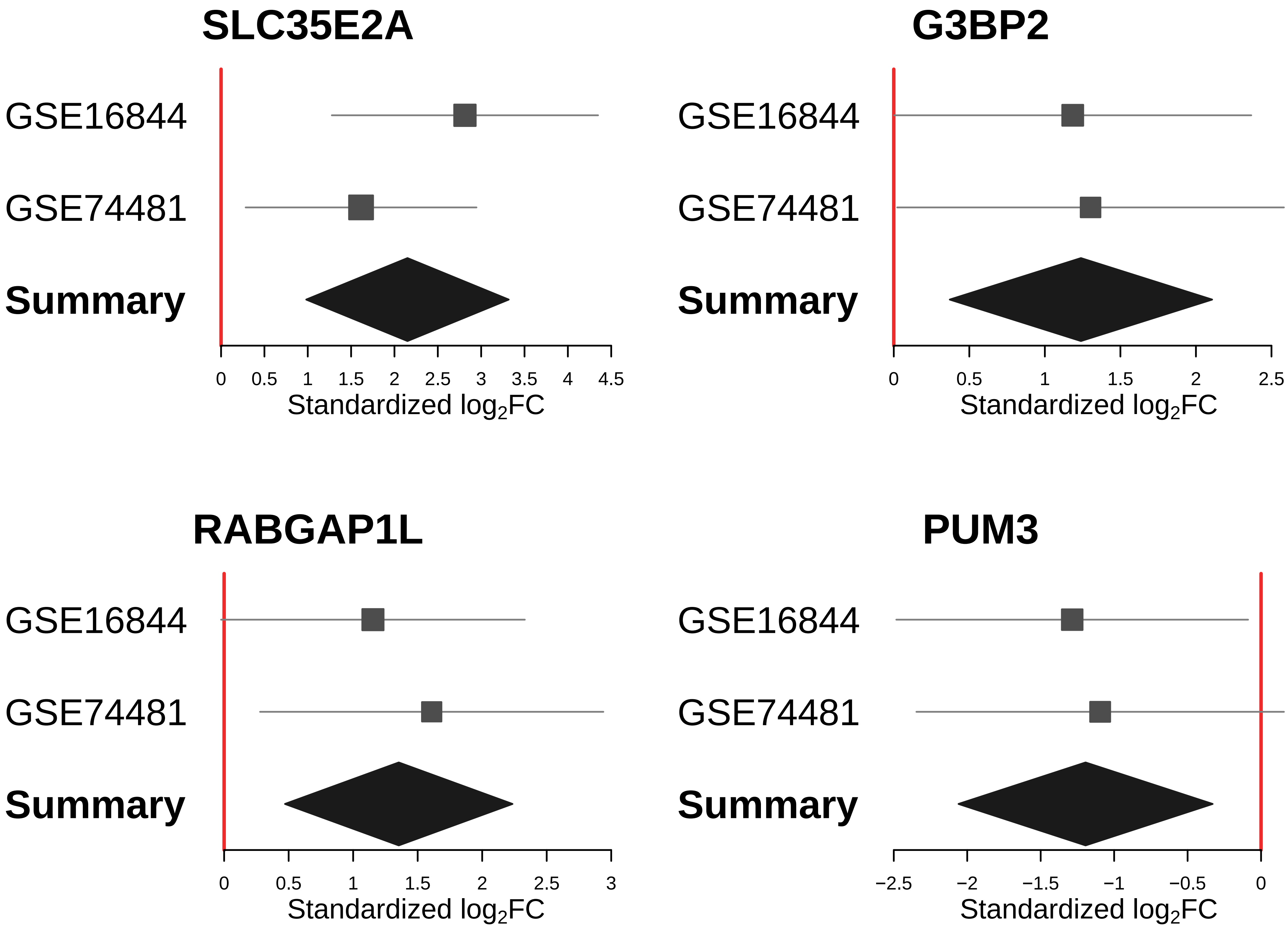

SLC35E2A
GSE16844
GSE74481
Summary
0
0.5
1
1.5
2
2.5
3
3.5
4
4.5
Standardized log2FC
G3BP2
GSE16844
GSE74481
Summary
0
0.5
1
1.5
2
2.5
Standardized log2FC
RABGAP1L
GSE16844
GSE74481
Summary
0
0.5
1
1.5
2
2.5
3
Standardized log2FC
PUM3
GSE16844
GSE74481
Summary
−2.5
−2
−1.5
−1
−0.5
0
Standardized log2FC

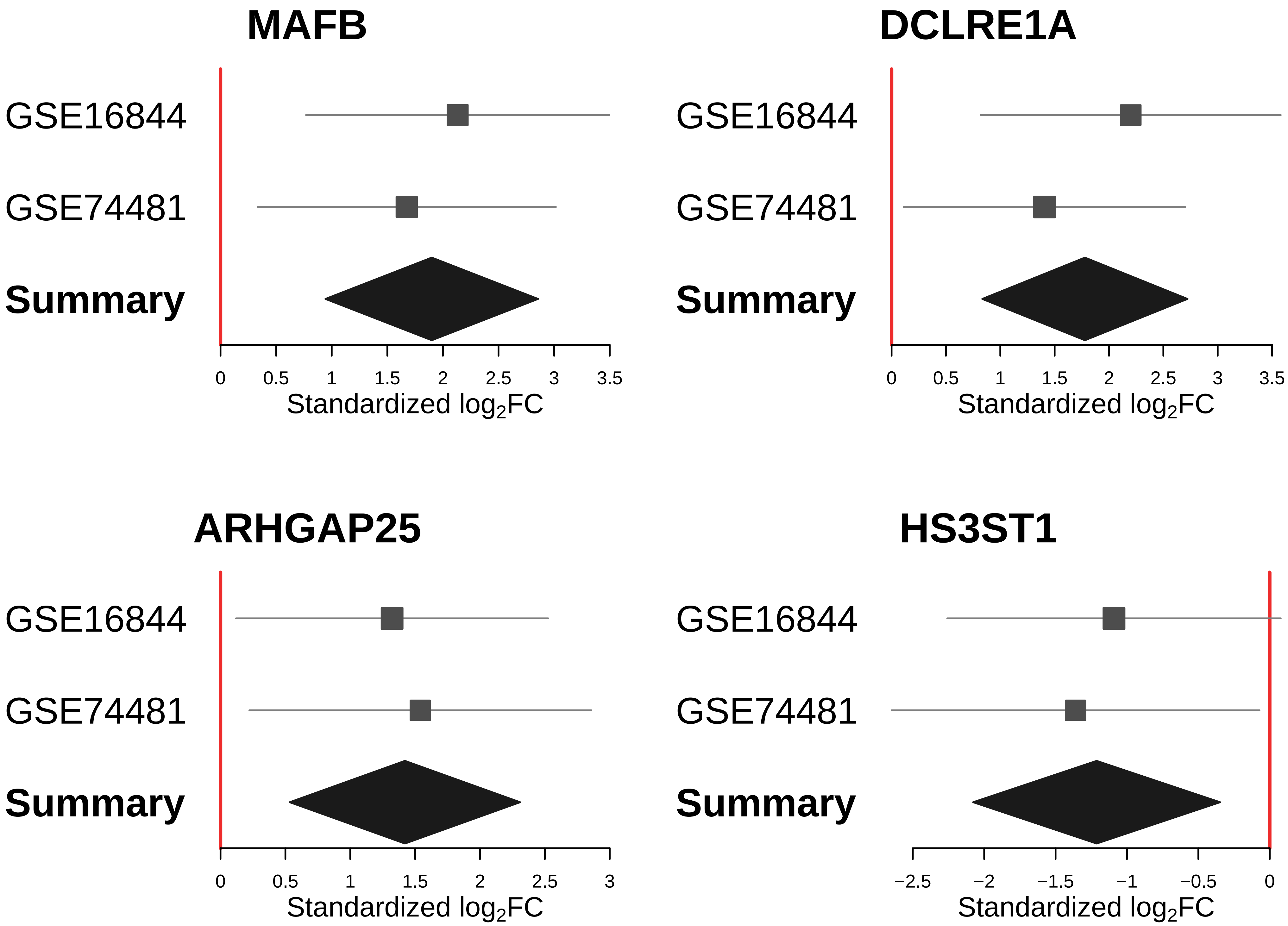
MAFB
GSE16844
GSE74481
Summary
0 0.5 1 1.5 2 2.5 3 3.5
Standardized log2FC
DCLRE1A
GSE16844
GSE74481
Summary
0 0.5 1 1.5 2 2.5 3 3.5
Standardized log2FC
ARHGAP25
GSE16844
GSE74481
Summary
0 0.5 1 1.5 2 2.5 3
Standardized log2FC
HS3ST1
GSE16844
GSE74481
Summary
−2.5 −2 −1.5 −1 −0.5 0
Standardized log2FC

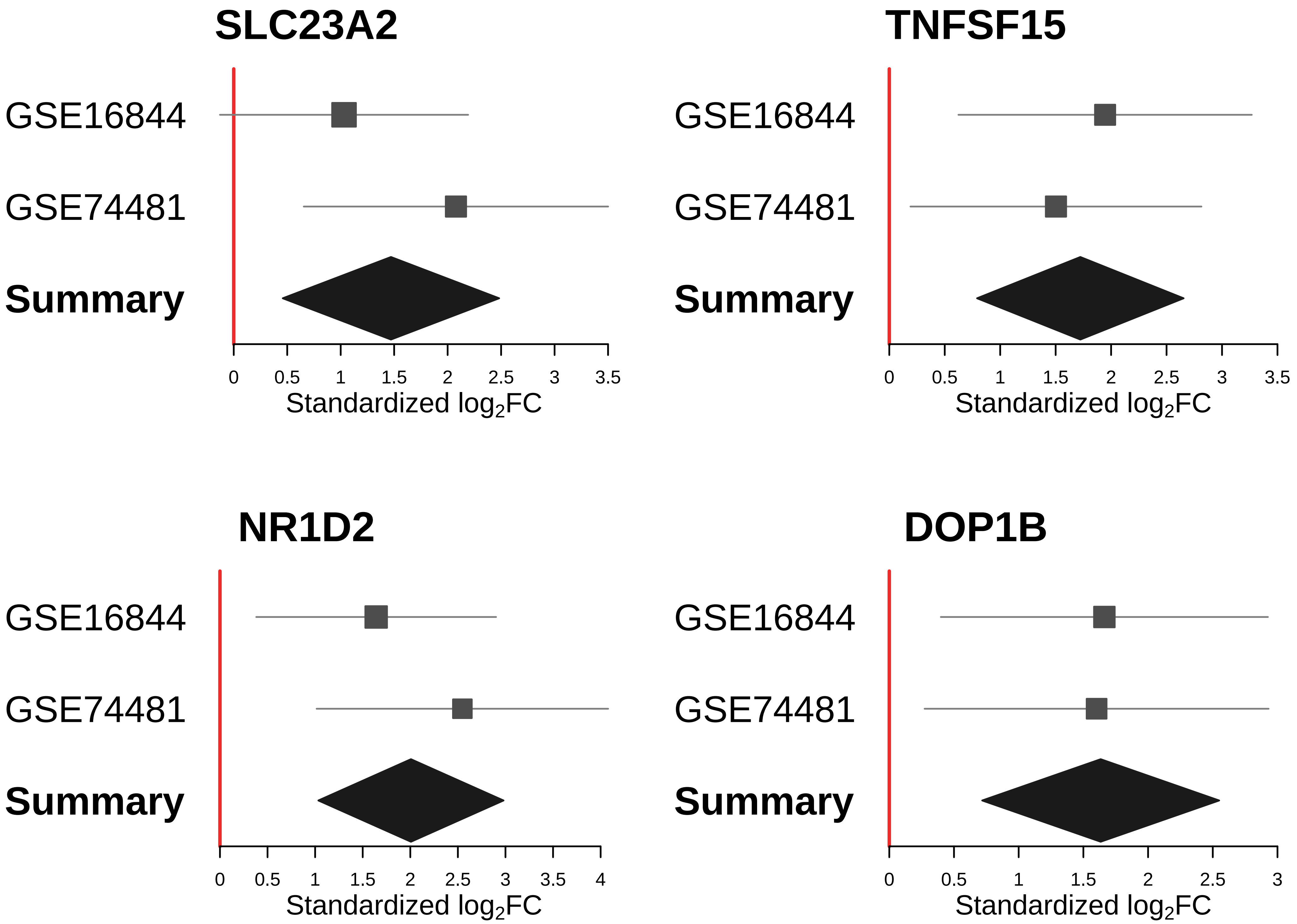
SLC23A2
GSE16844
GSE74481
Summary
0
0.5
1
1.5
2
2.5
3
3.5
Standardized log2FC
TNFSF15
GSE16844
GSE74481
Summary
0
0.5
1
1.5
2
2.5
3
3.5
Standardized log2FC
NR1D2
GSE16844
GSE74481
Summary
0
0.5
1
1.5
2
2.5
3
3.5
4
Standardized log2FC
DOP1B
GSE16844
GSE74481
Summary
0
0.5
1
1.5
2
2.5
3
Standardized log2FC

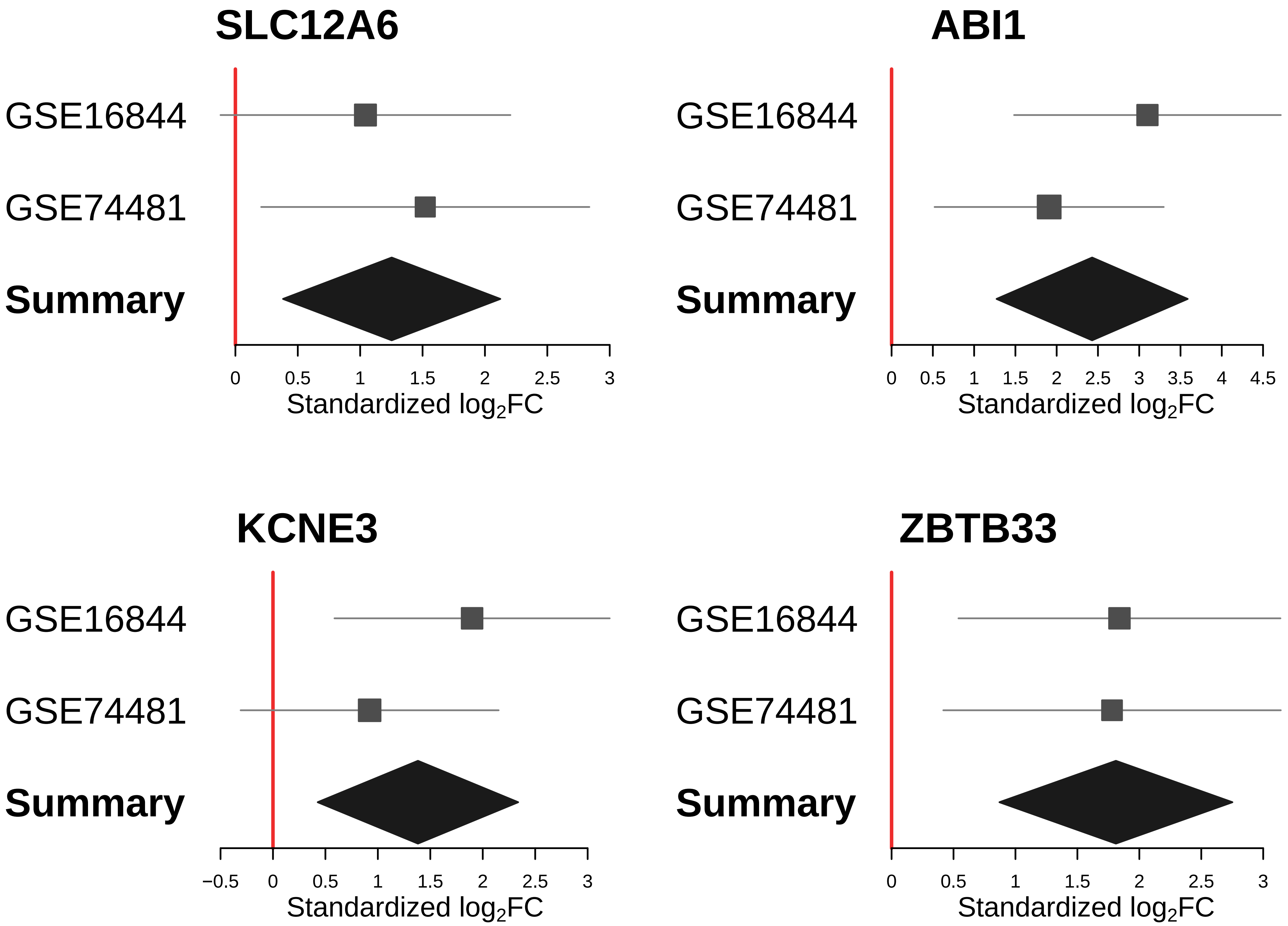
SLC12A6
GSE16844
GSE74481
Summary
0
0.5
1
1.5
2
2.5
3
Standardized $log_2FC$
ABI1
GSE16844
GSE74481
Summary
0
0.5
1
1.5
2
2.5
3
3.5
4
4.5
Standardized $log_2FC$
KCNE3
GSE16844
GSE74481
Summary
−0.5
0
0.5
1
1.5
2
2.5
3
Standardized $log_2FC$
ZBTB33
GSE16844
GSE74481
Summary
0
0.5
1
1.5
2
2.5
3
Standardized $log_2FC$

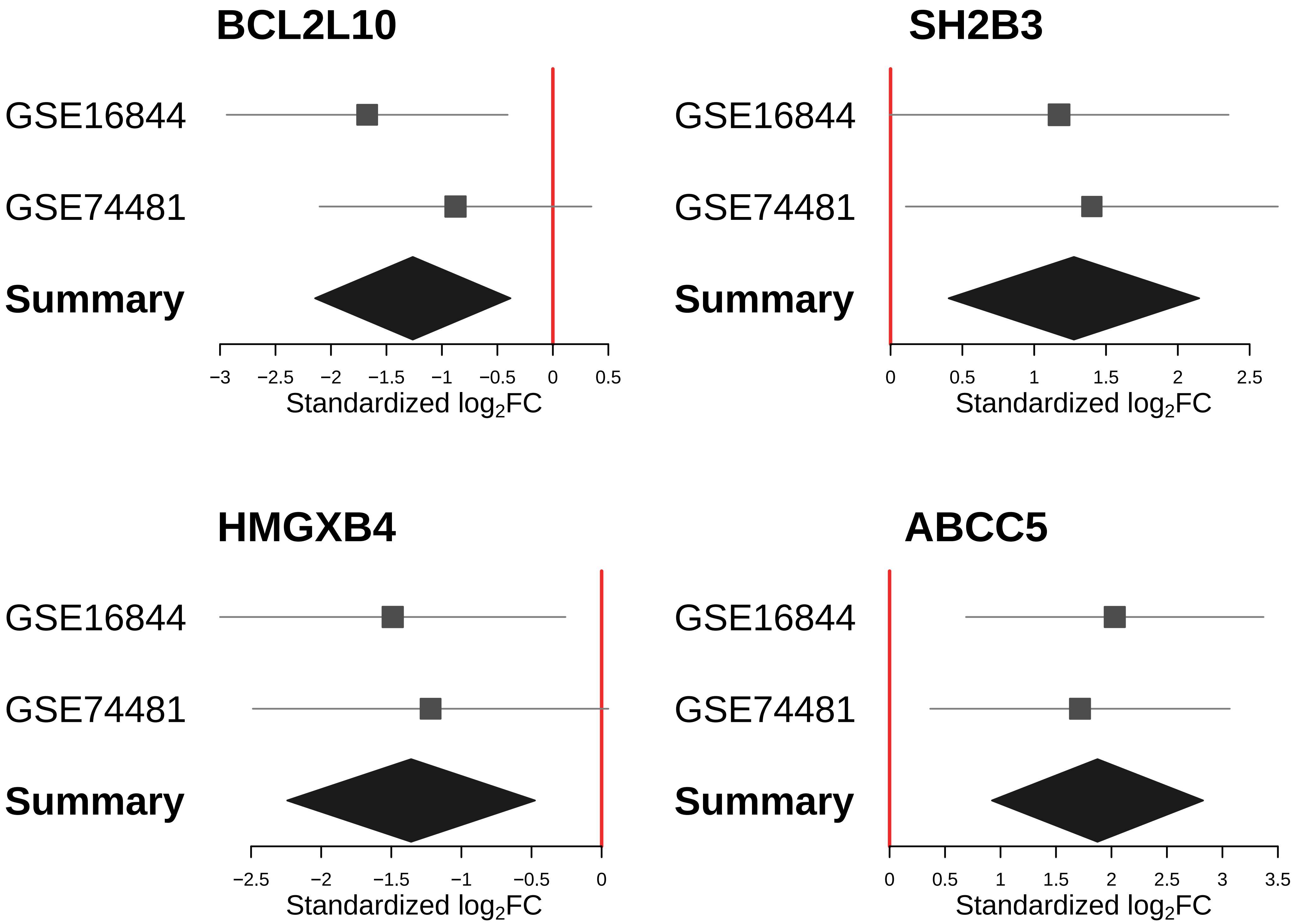
BCL2L10
GSE16844
GSE74481
Summary
−3 −2.5 −2 −1.5 −1 −0.5 0 0.5
Standardized $log_2FC$
SH2B3
GSE16844
GSE74481
Summary
0 0.5 1 1.5 2 2.5
Standardized $log_2FC$
HMGXB4
GSE16844
GSE74481
Summary
−2.5 −2 −1.5 −1 −0.5 0
Standardized $log_2FC$
ABCC5
GSE16844
GSE74481
Summary
0 0.5 1 1.5 2 2.5 3 3.5
Standardized $log_2FC$

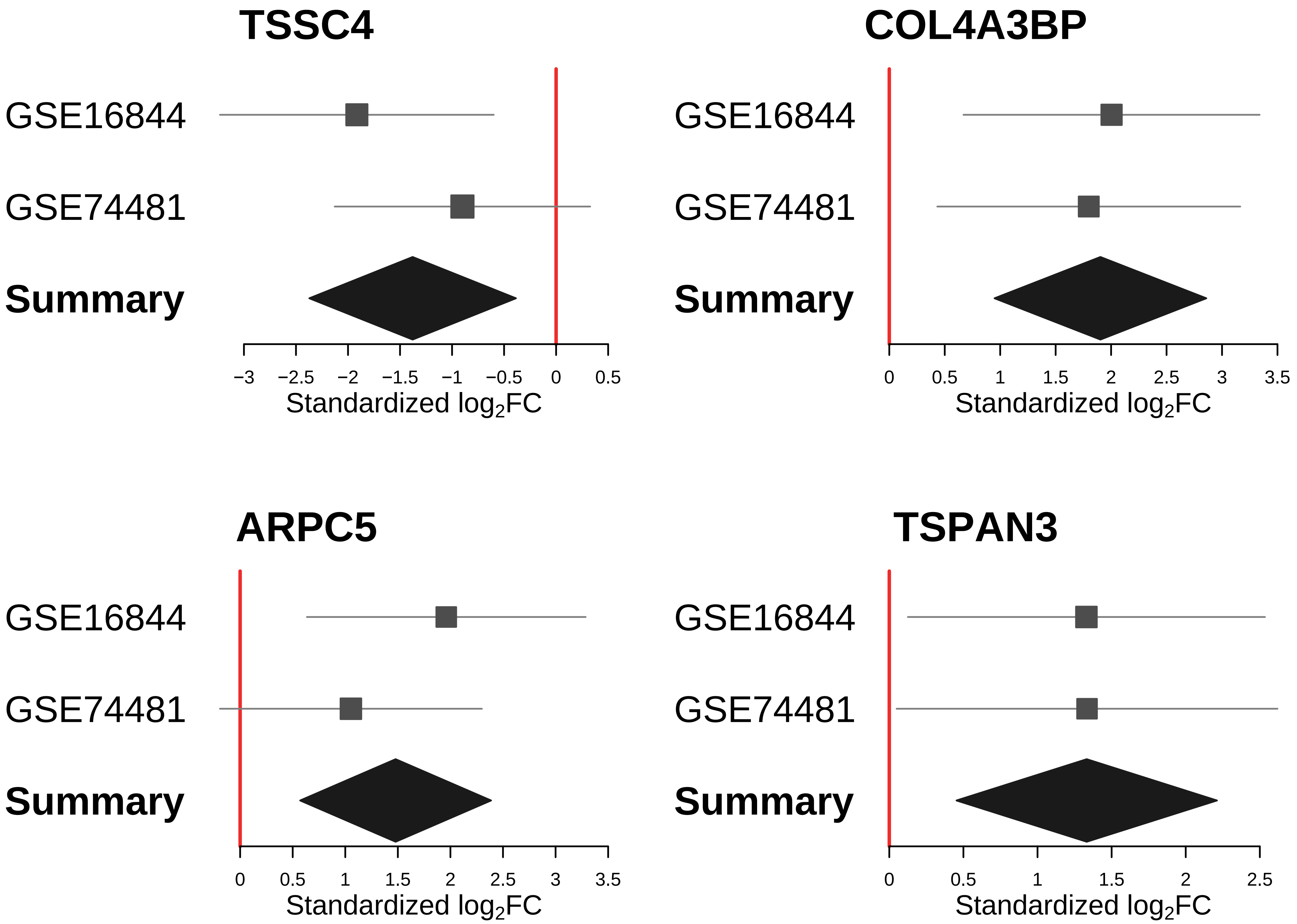
TSSC4
GSE16844
GSE74481
Summary
−3 −2.5 −2 −1.5 −1 −0.5 0 0.5
Standardized $log_2FC$
COL4A3BP
GSE16844
GSE74481
Summary
0 0.5 1 1.5 2 2.5 3 3.5
Standardized $log_2FC$
ARPC5
GSE16844
GSE74481
Summary
0 0.5 1 1.5 2 2.5 3 3.5
Standardized $log_2FC$
TSPAN3
GSE16844
GSE74481
Summary
0 0.5 1 1.5 2 2.5
Standardized $log_2FC$

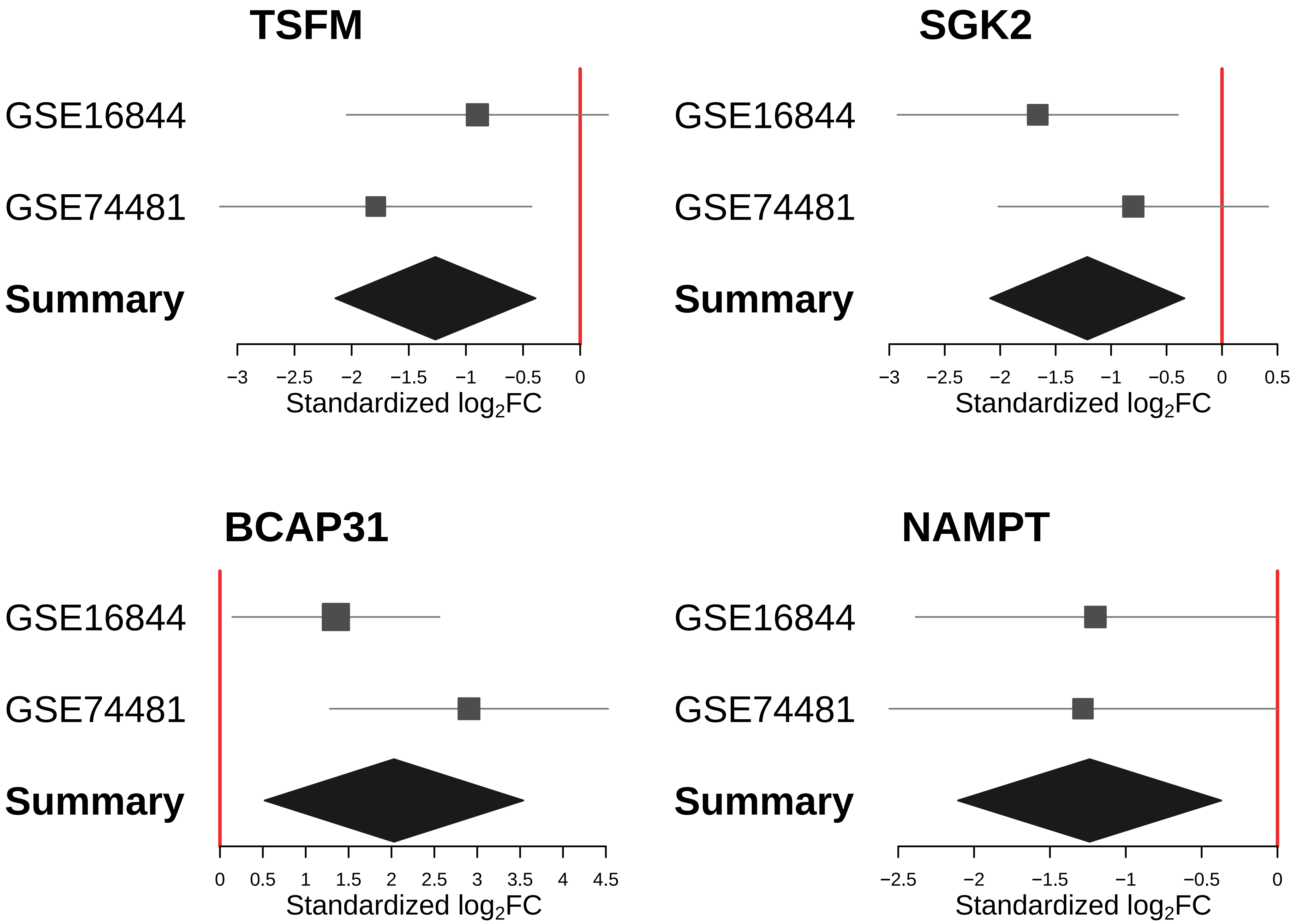
TSFM
GSE16844
GSE74481
Summary
−3 −2.5 −2 −1.5 −1 −0.5 0
Standardized log2FC
SGK2
GSE16844
GSE74481
Summary
−3 −2.5 −2 −1.5 −1 −0.5 0 0.5
Standardized log2FC
BCAP31
GSE16844
GSE74481
Summary
0 0.5 1 1.5 2 2.5 3 3.5 4 4.5
Standardized log2FC
NAMPT
GSE16844
GSE74481
Summary
−2.5 −2 −1.5 −1 −0.5 0
Standardized log2FC

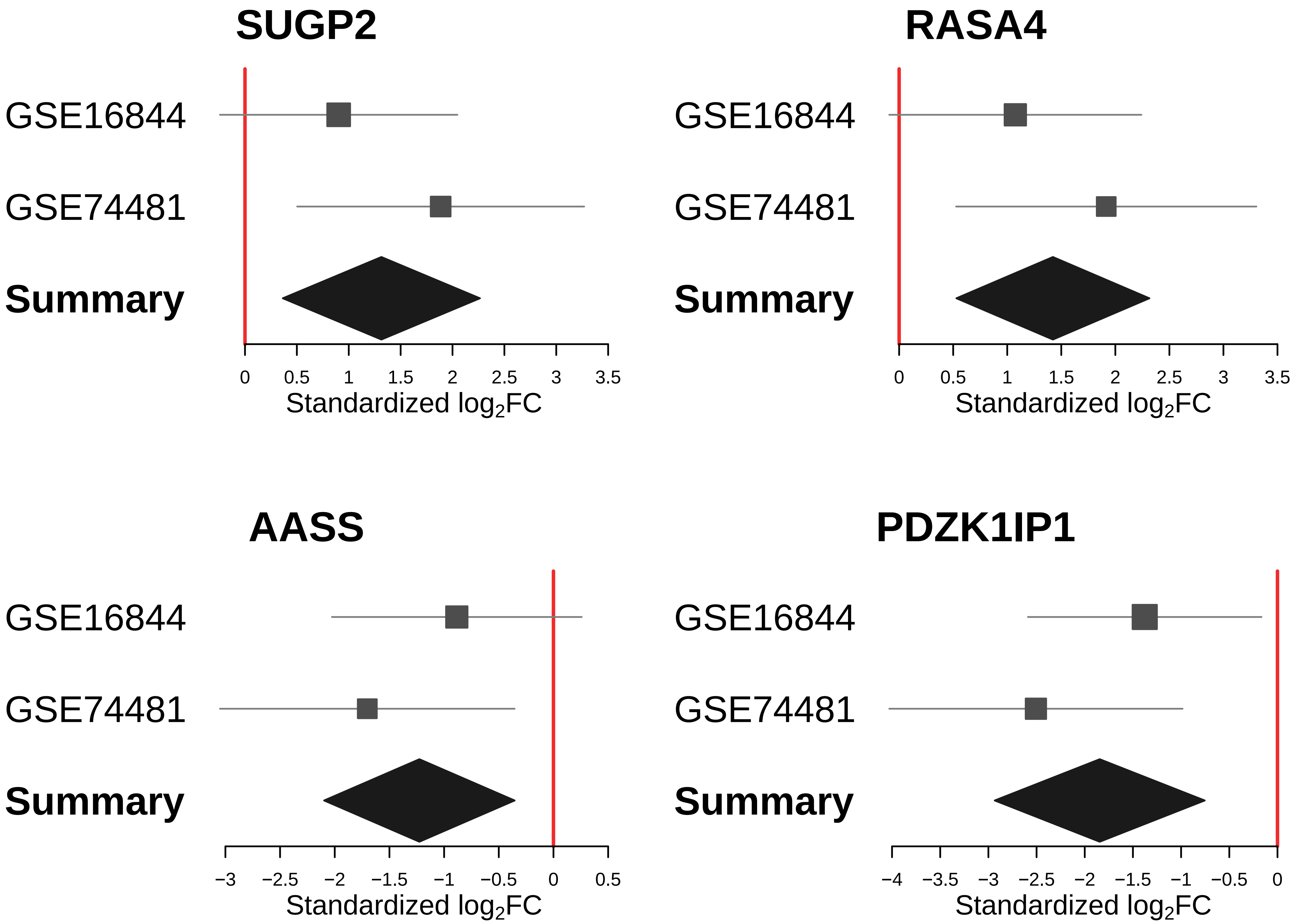

SUGP2
GSE16844
GSE74481
Summary
0 0.5 1 1.5 2 2.5 3 3.5
Standardized log2FC
RASA4
GSE16844
GSE74481
Summary
0 0.5 1 1.5 2 2.5 3 3.5
Standardized log2FC
AASS
GSE16844
GSE74481
Summary
−3 −2.5 −2 −1.5 −1 −0.5 0 0.5
Standardized log2FC
PDZK1IP1
GSE16844
GSE74481
Summary
−4 −3.5 −3 −2.5 −2 −1.5 −1 −0.5 0
Standardized log2FC

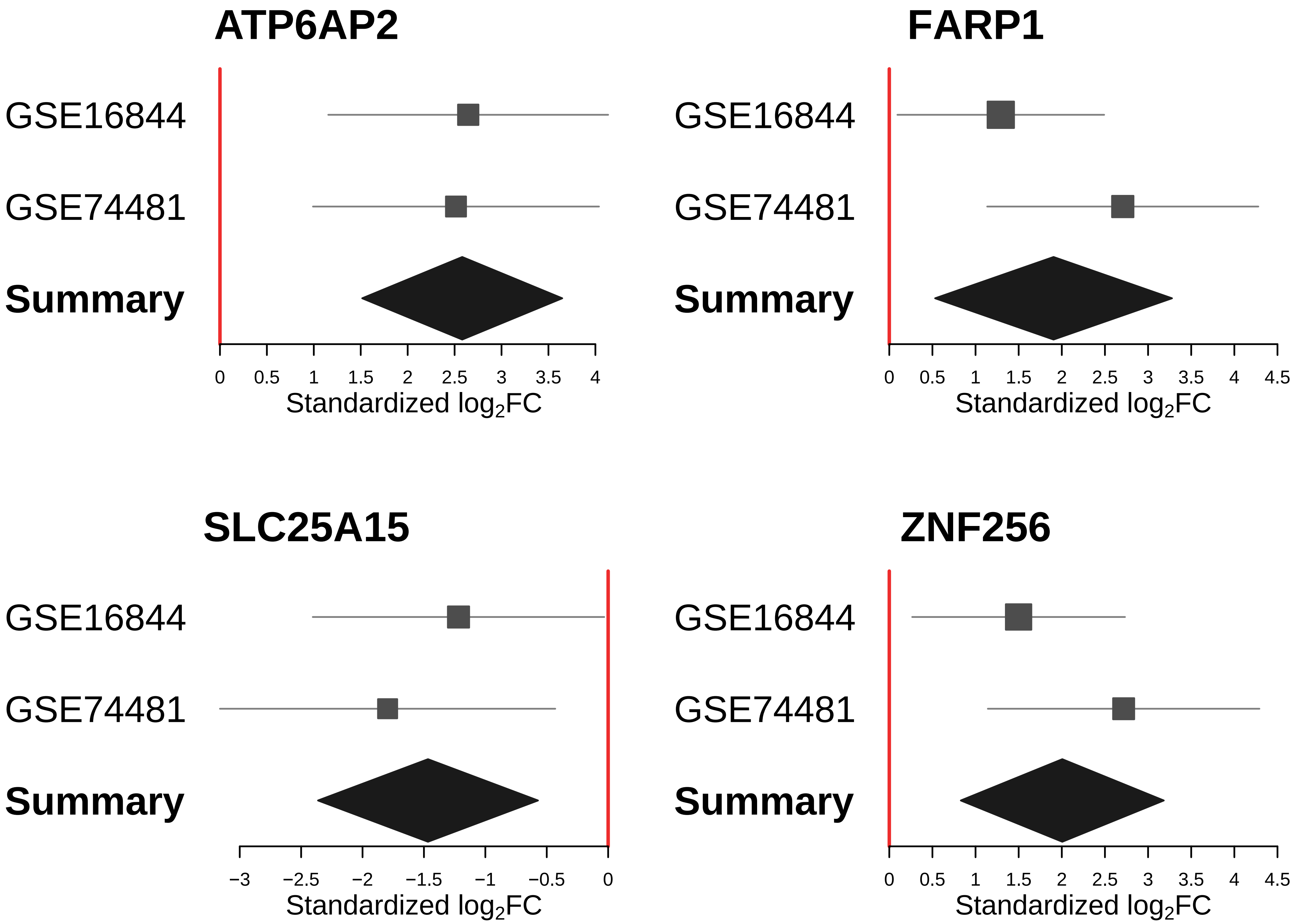

ATP6AP2
GSE16844
GSE74481
Summary
0 0.5 1 1.5 2 2.5 3 3.5 4
Standardized $log_2FC$
FARP1
GSE16844
GSE74481
Summary
0 0.5 1 1.5 2 2.5 3 3.5 4 4.5
Standardized $log_2FC$
SLC25A15
GSE16844
GSE74481
Summary
−3 −2.5 −2 −1.5 −1 −0.5 0
Standardized $log_2FC$
ZNF256
GSE16844
GSE74481
Summary
0 0.5 1 1.5 2 2.5 3 3.5 4 4.5
Standardized $log_2FC$

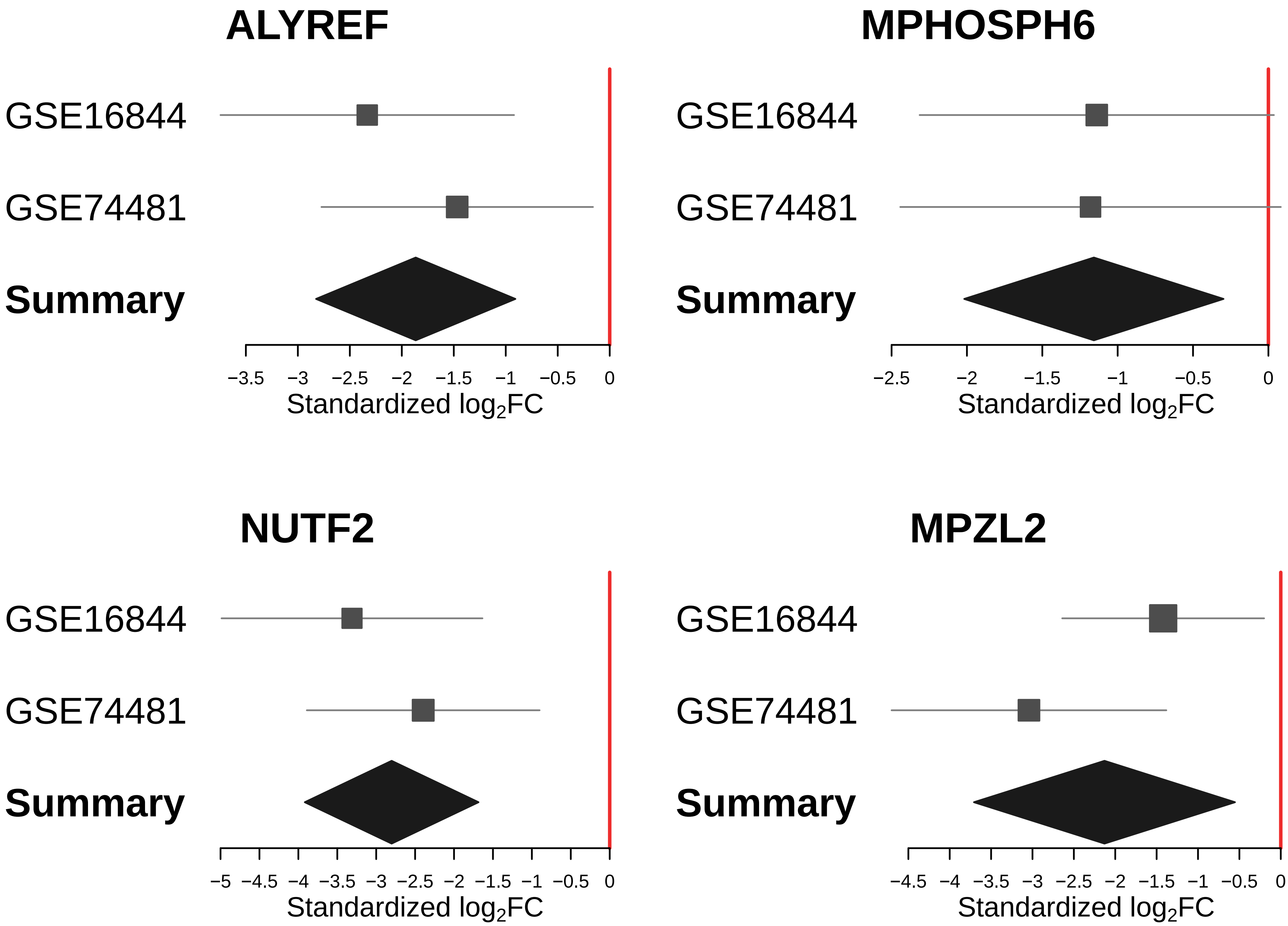
ALYREF
GSE16844
GSE74481
Summary
−3.5 −3 −2.5 −2 −1.5 −1 −0.5 0
Standardized log2FC
MPHOSPH6
GSE16844
GSE74481
Summary
−2.5 −2 −1.5 −1 −0.5 0
Standardized log2FC
NUTF2
GSE16844
GSE74481
Summary
−5 −4.5 −4 −3.5 −3 −2.5 −2 −1.5 −1 −0.5 0
Standardized log2FC
MPZL2
GSE16844
GSE74481
Summary
−4.5 −4 −3.5 −3 −2.5 −2 −1.5 −1 −0.5 0
Standardized log2FC

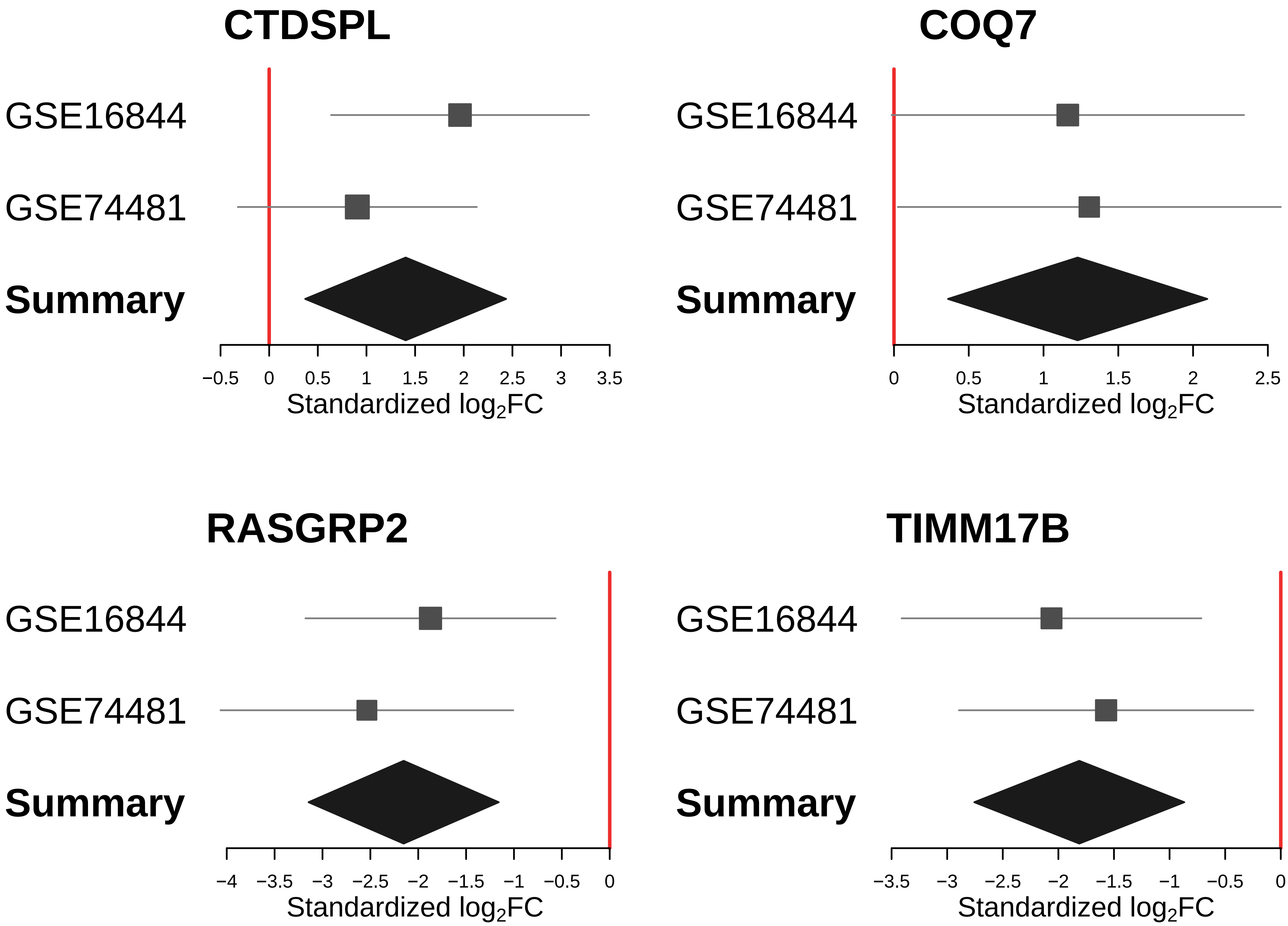
CTDSPL
GSE16844
GSE74481
Summary
−0.5 0 0.5 1 1.5 2 2.5 3 3.5
Standardized log2FC
COQ7
GSE16844
GSE74481
Summary
0 0.5 1 1.5 2 2.5
Standardized log2FC
RASGRP2
GSE16844
GSE74481
Summary
−4 −3.5 −3 −2.5 −2 −1.5 −1 −0.5 0
Standardized log2FC
TIMM17B
GSE16844
GSE74481
Summary
−3.5 −3 −2.5 −2 −1.5 −1 −0.5 0
Standardized log2FC

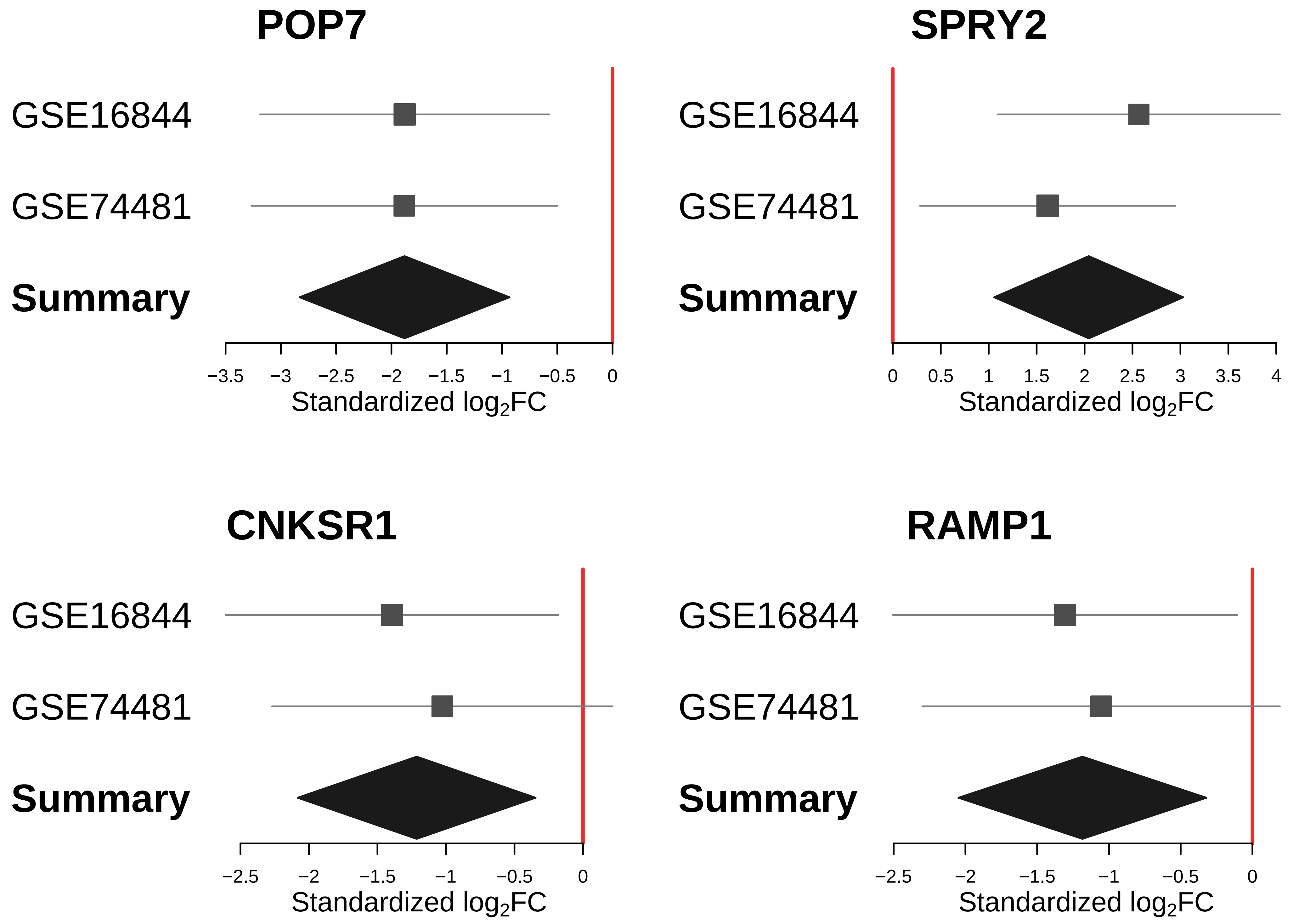
POP7
GSE16844
GSE74481
Summary
−3.5 −3 −2.5 −2 −1.5 −1 −0.5 0
Standardized log2FC
SPRY2
GSE16844
GSE74481
Summary
0 0.5 1 1.5 2 2.5 3 3.5 4
Standardized log2FC
CNKSR1
GSE16844
GSE74481
Summary
−2.5 −2 −1.5 −1 −0.5 0
Standardized log2FC
RAMP1
GSE16844
GSE74481
Summary
−2.5 −2 −1.5 −1 −0.5 0
Standardized log2FC

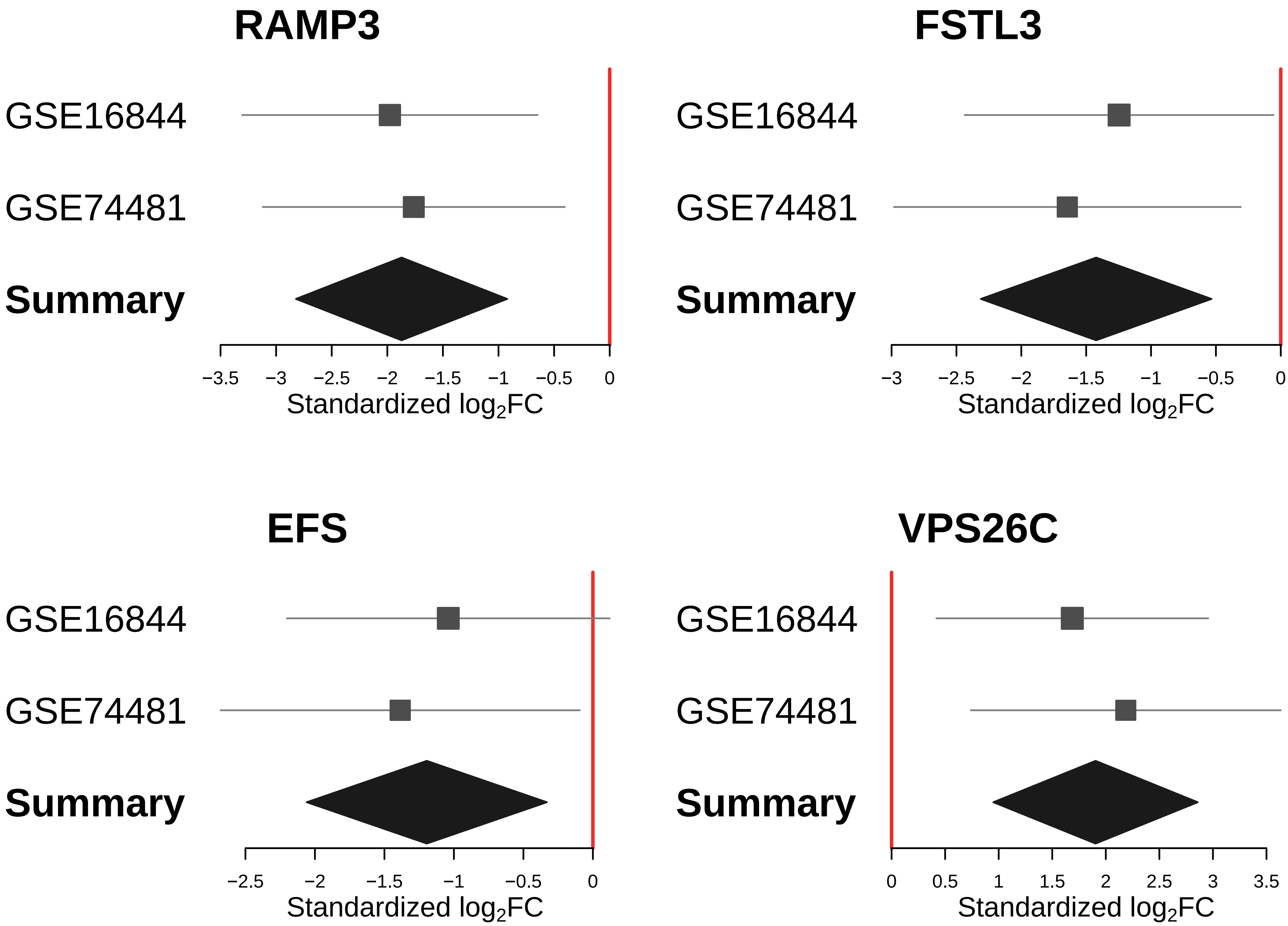
RAMP3
GSE16844
GSE74481
Summary
−3.5 −3 −2.5 −2 −1.5 −1 −0.5 0
Standardized log2FC
FSTL3
GSE16844
GSE74481
Summary
−3 −2.5 −2 −1.5 −1 −0.5 0
Standardized log2FC
EFS
GSE16844
GSE74481
Summary
−2.5 −2 −1.5 −1 −0.5 0
Standardized log2FC
VPS26C
GSE16844
GSE74481
Summary
0 0.5 1 1.5 2 2.5 3 3.5
Standardized log2FC

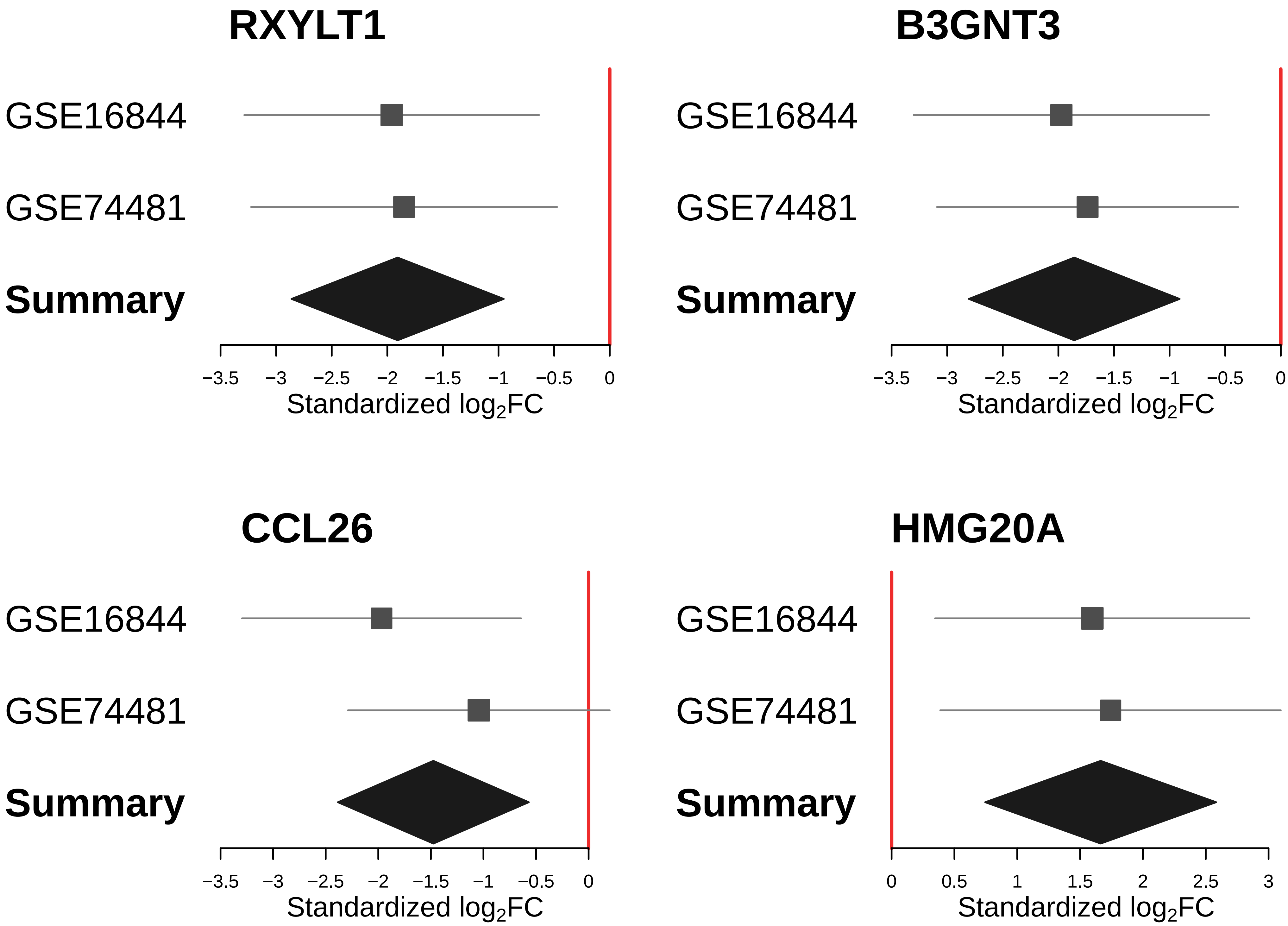

RXYLT1
GSE16844
GSE74481
Summary
−3.5 −3 −2.5 −2 −1.5 −1 −0.5 0
Standardized log2FC
B3GNT3
GSE16844
GSE74481
Summary
−3.5 −3 −2.5 −2 −1.5 −1 −0.5 0
Standardized log2FC
CCL26
GSE16844
GSE74481
Summary
−3.5 −3 −2.5 −2 −1.5 −1 −0.5 0
Standardized log2FC
HMG20A
GSE16844
GSE74481
Summary
0 0.5 1 1.5 2 2.5 3
Standardized log2FC

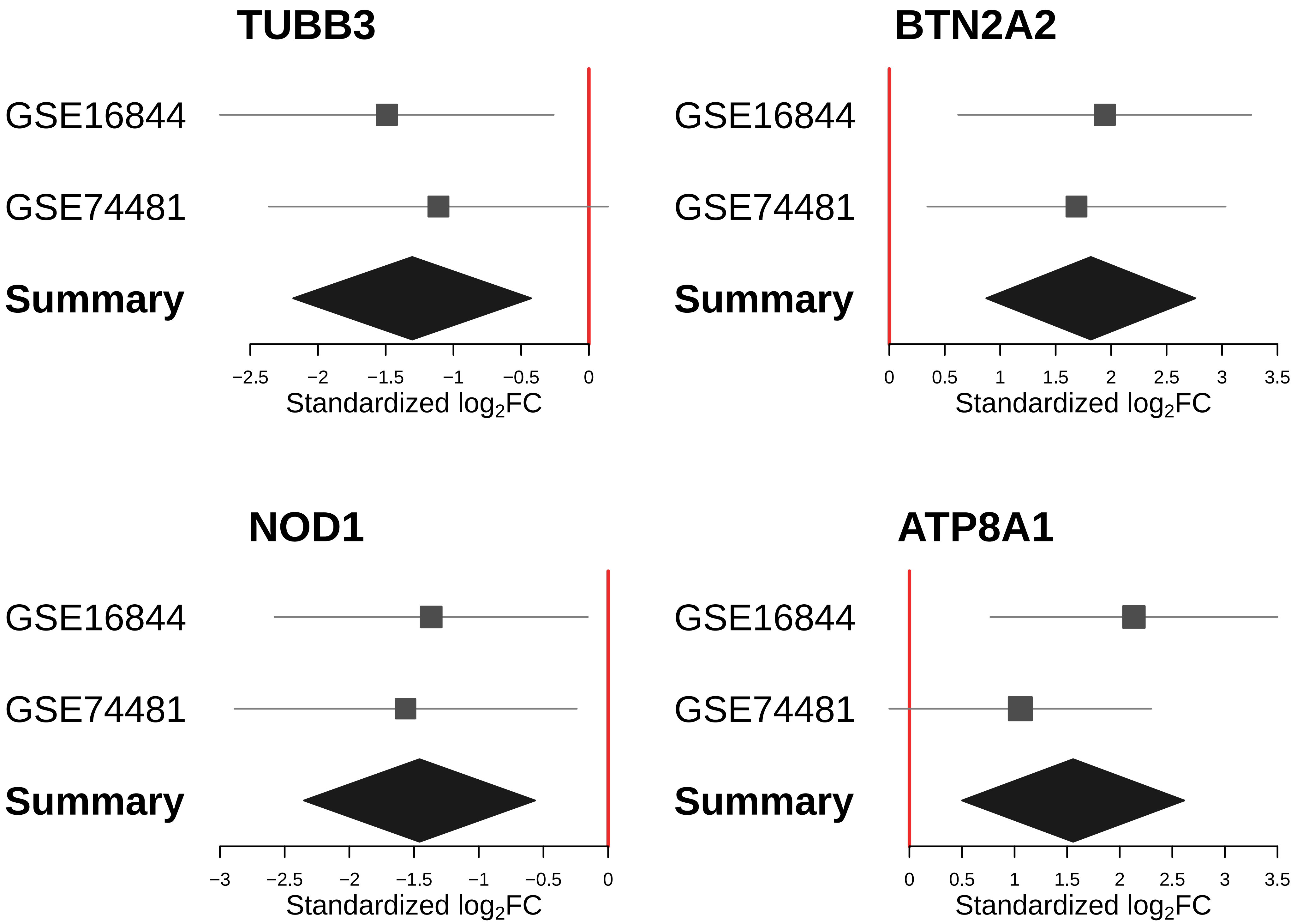
TUBB3
GSE16844
GSE74481
Summary
−2.5
−2
−1.5
−1
−0.5
0
Standardized log2FC
BTN2A2
GSE16844
GSE74481
Summary
0
0.5
1
1.5
2
2.5
3
3.5
Standardized log2FC
NOD1
GSE16844
GSE74481
Summary
−3
−2.5
−2
−1.5
−1
−0.5
0
Standardized log2FC
ATP8A1
GSE16844
GSE74481
Summary
0
0.5
1
1.5
2
2.5
3
3.5
Standardized log2FC

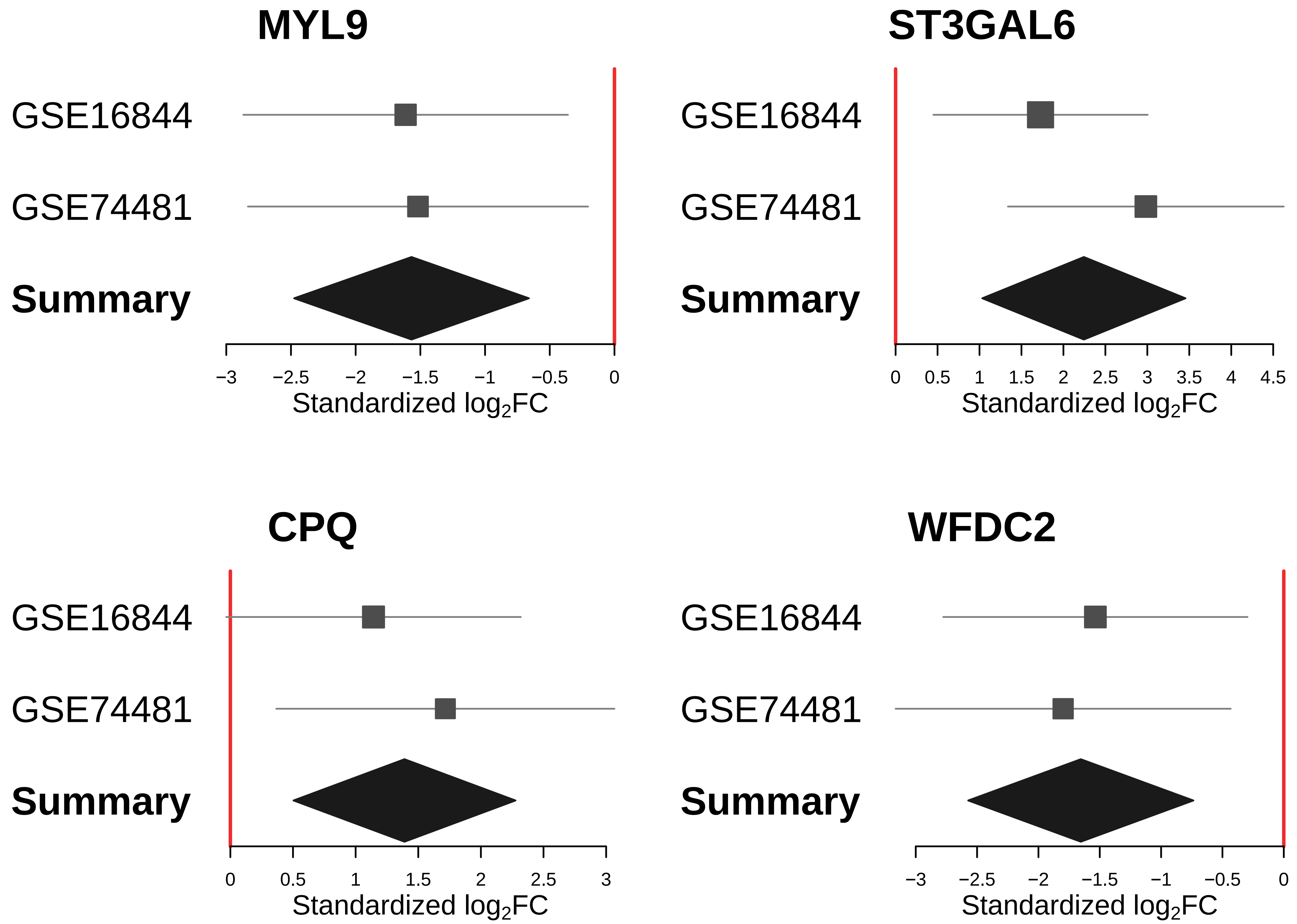
MYL9
ST3GAL6
CPQ
WFDC2
GSE16844
GSE74481
Summary
−3
−2.5
−2
−1.5
−1
−0.5
0
0.5
1
1.5
2
2.5
3
3.5
4
4.5
Standardized log2FC

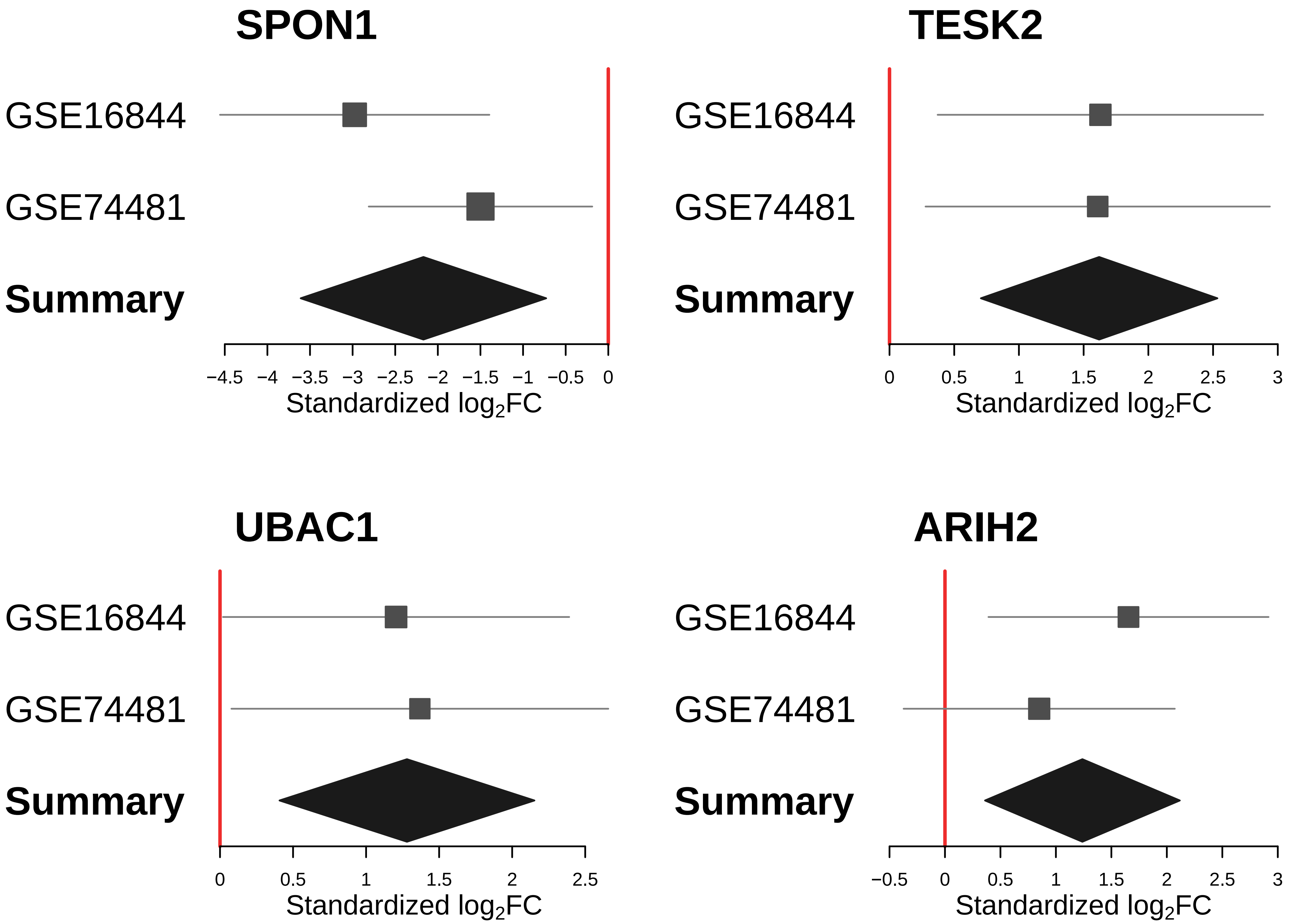
SPON1
GSE16844
GSE74481
Summary
−4.5 −4 −3.5 −3 −2.5 −2 −1.5 −1 −0.5 0
Standardized log2FC
TESK2
GSE16844
GSE74481
Summary
0 0.5 1 1.5 2 2.5 3
Standardized log2FC
UBAC1
GSE16844
GSE74481
Summary
0 0.5 1 1.5 2 2.5
Standardized log2FC
ARIH2
GSE16844
GSE74481
Summary
−0.5 0 0.5 1 1.5 2 2.5 3
Standardized log2FC

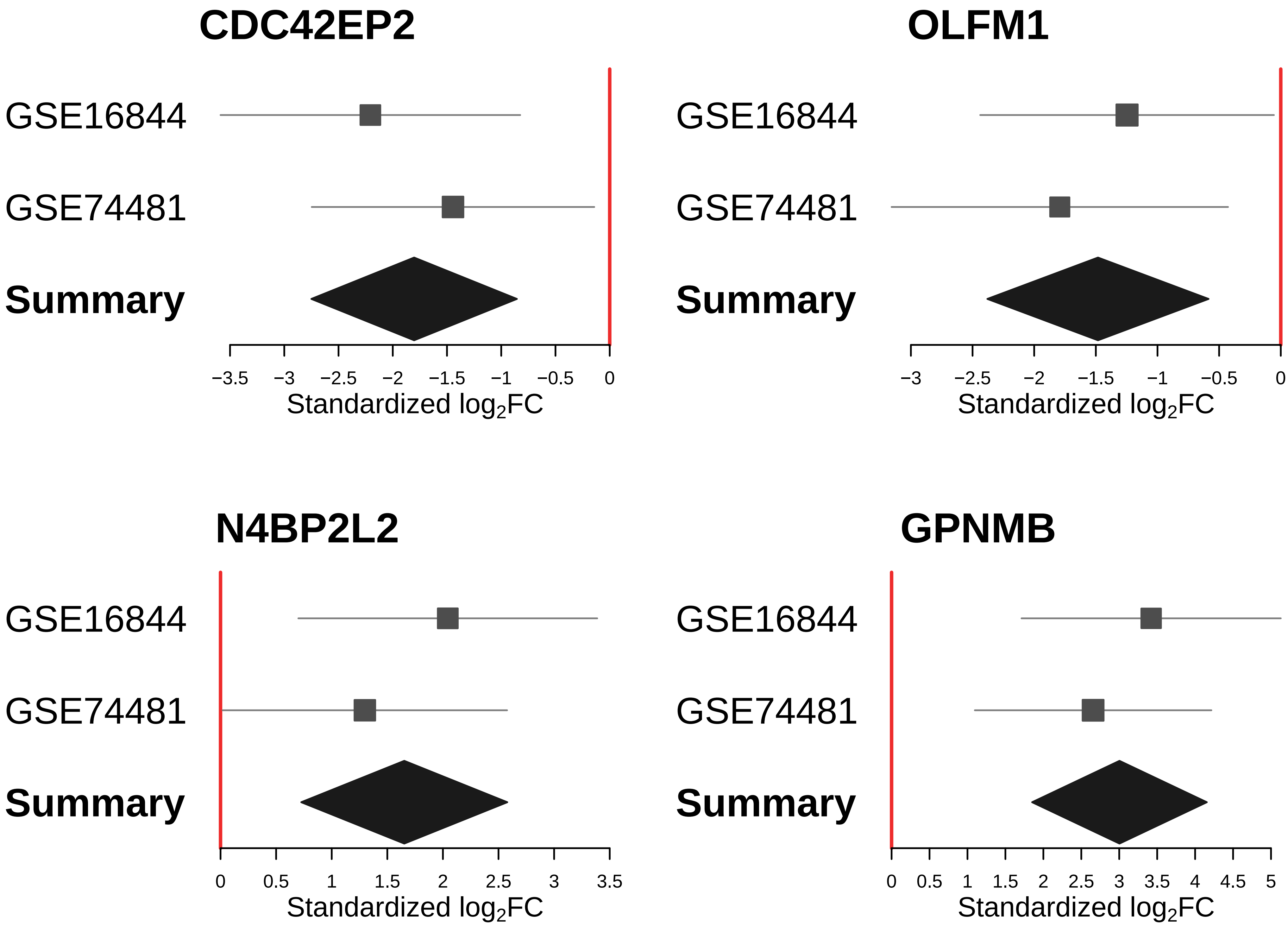
CDC42EP2
GSE16844
GSE74481
Summary
−3.5 −3 −2.5 −2 −1.5 −1 −0.5 0
Standardized log2FC
OLFM1
GSE16844
GSE74481
Summary
−3 −2.5 −2 −1.5 −1 −0.5 0
Standardized log2FC
N4BP2L2
GSE16844
GSE74481
Summary
0 0.5 1 1.5 2 2.5 3 3.5
Standardized log2FC
GPNMB
GSE16844
GSE74481
Summary
0 0.5 1 1.5 2 2.5 3 3.5 4 4.5 5
Standardized log2FC

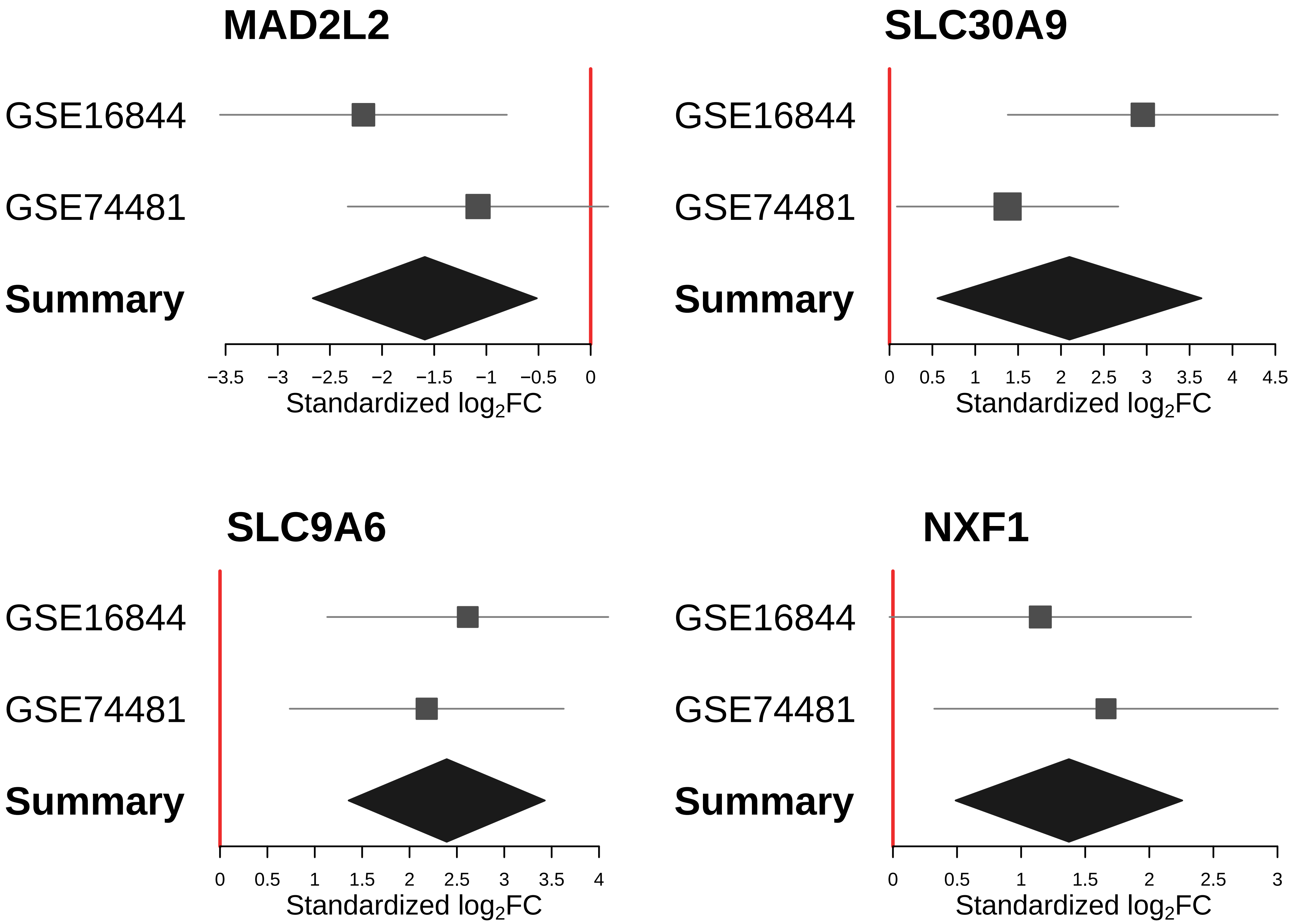
MAD2L2
GSE16844
GSE74481
Summary
−3.5 −3 −2.5 −2 −1.5 −1 −0.5 0
Standardized log2FC
SLC30A9
GSE16844
GSE74481
Summary
0 0.5 1 1.5 2 2.5 3 3.5 4 4.5
Standardized log2FC
SLC9A6
GSE16844
GSE74481
Summary
0 0.5 1 1.5 2 2.5 3 3.5 4
Standardized log2FC
NXF1
GSE16844
GSE74481
Summary
0 0.5 1 1.5 2 2.5 3
Standardized log2FC

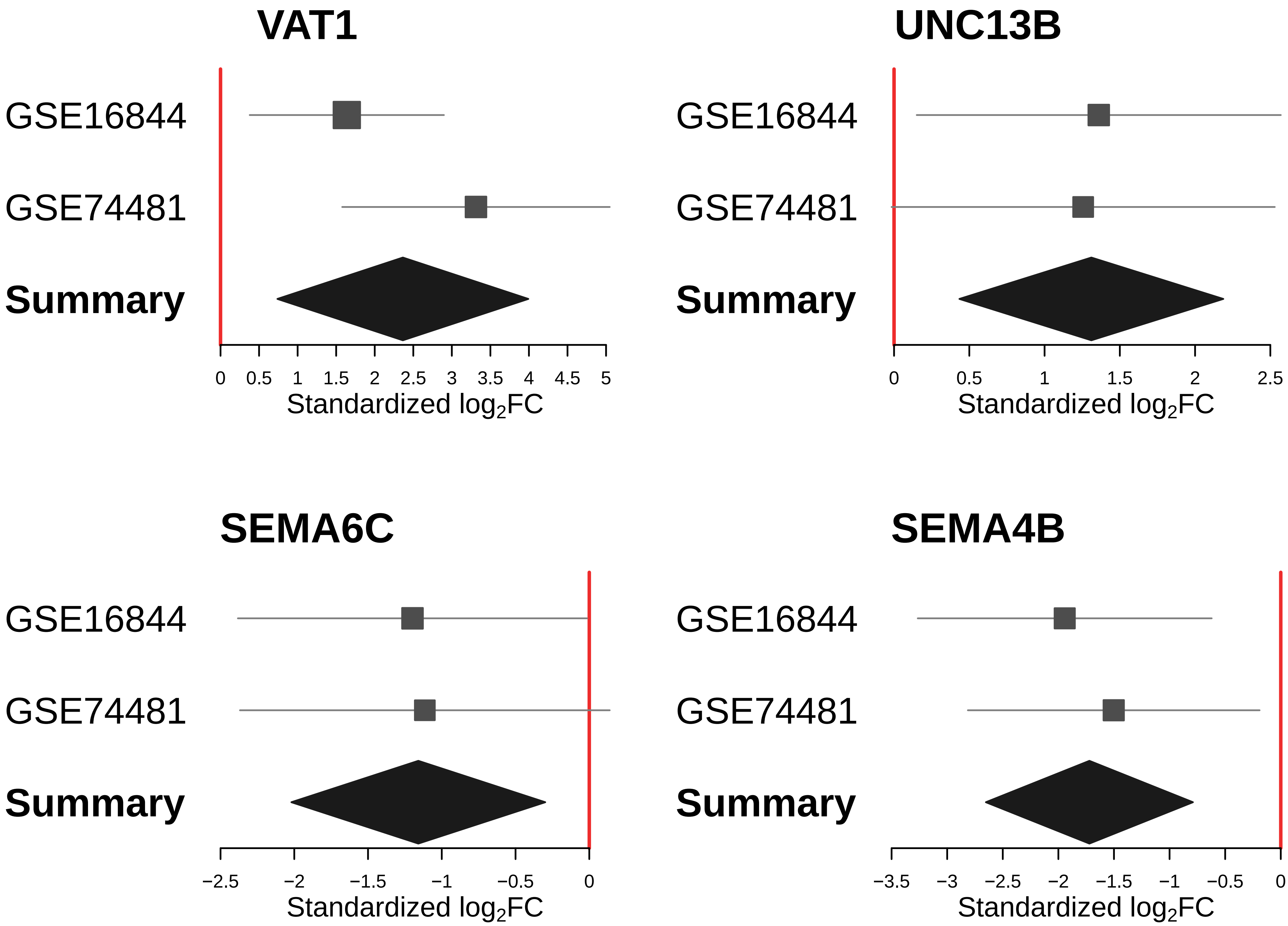
VAT1
GSE16844
GSE74481
Summary
0 0.5 1 1.5 2 2.5 3 3.5 4 4.5 5
Standardized log2FC
UNC13B
GSE16844
GSE74481
Summary
0 0.5 1 1.5 2 2.5
Standardized log2FC
SEMA6C
GSE16844
GSE74481
Summary
−2.5 −2 −1.5 −1 −0.5 0
Standardized log2FC
SEMA4B
GSE16844
GSE74481
Summary
−3.5 −3 −2.5 −2 −1.5 −1 −0.5 0
Standardized log2FC

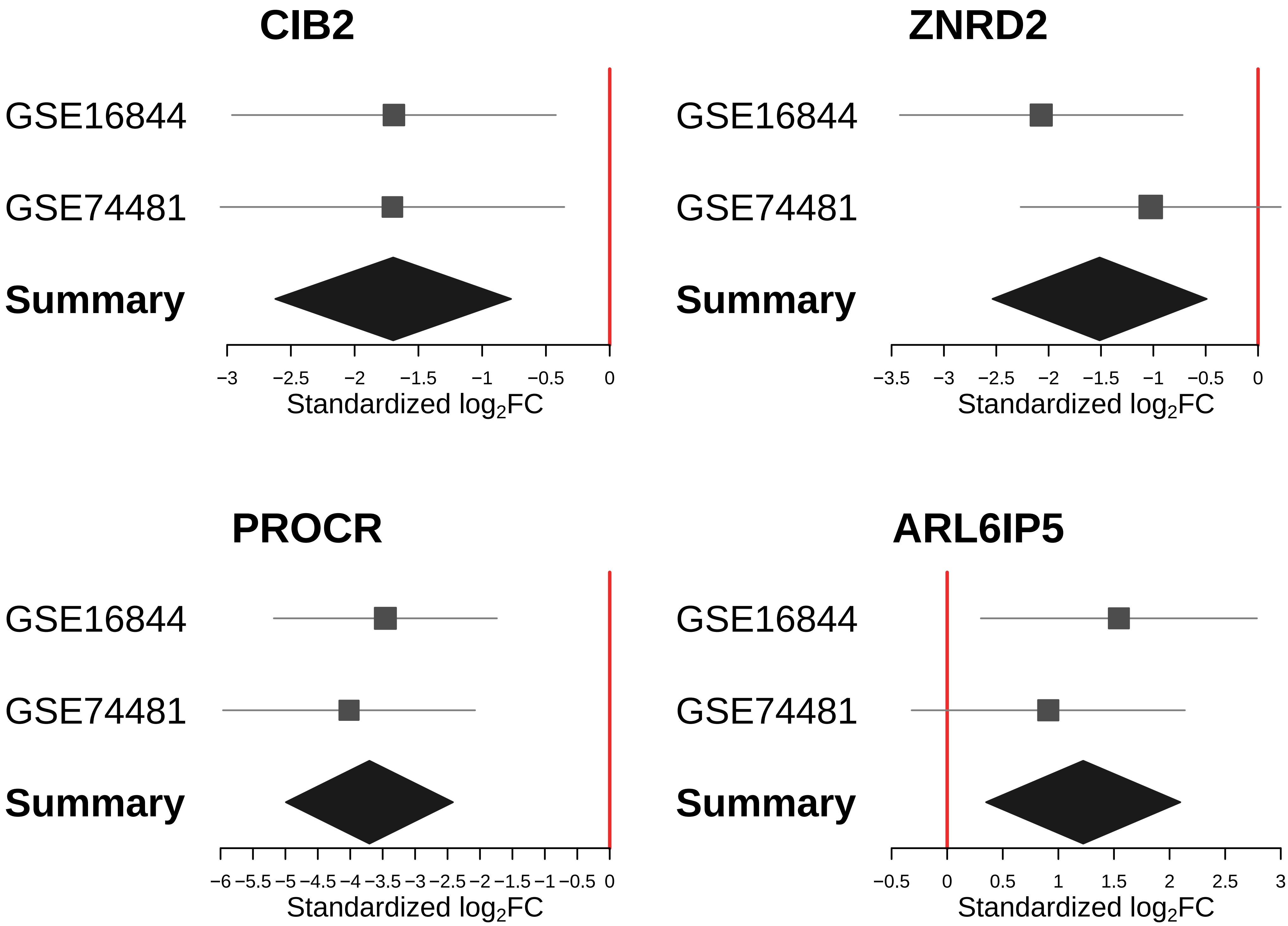
CIB2
GSE16844
GSE74481
Summary
−3 −2.5 −2 −1.5 −1 −0.5 0
Standardized log2FC
ZNRD2
GSE16844
GSE74481
Summary
−3.5 −3 −2.5 −2 −1.5 −1 −0.5 0
Standardized log2FC
PROCR
GSE16844
GSE74481
Summary
−6 −5.5 −5 −4.5 −4 −3.5 −3 −2.5 −2 −1.5 −1 −0.5 0
Standardized log2FC
ARL6IP5
GSE16844
GSE74481
Summary
−0.5 0 0.5 1 1.5 2 2.5 3
Standardized log2FC

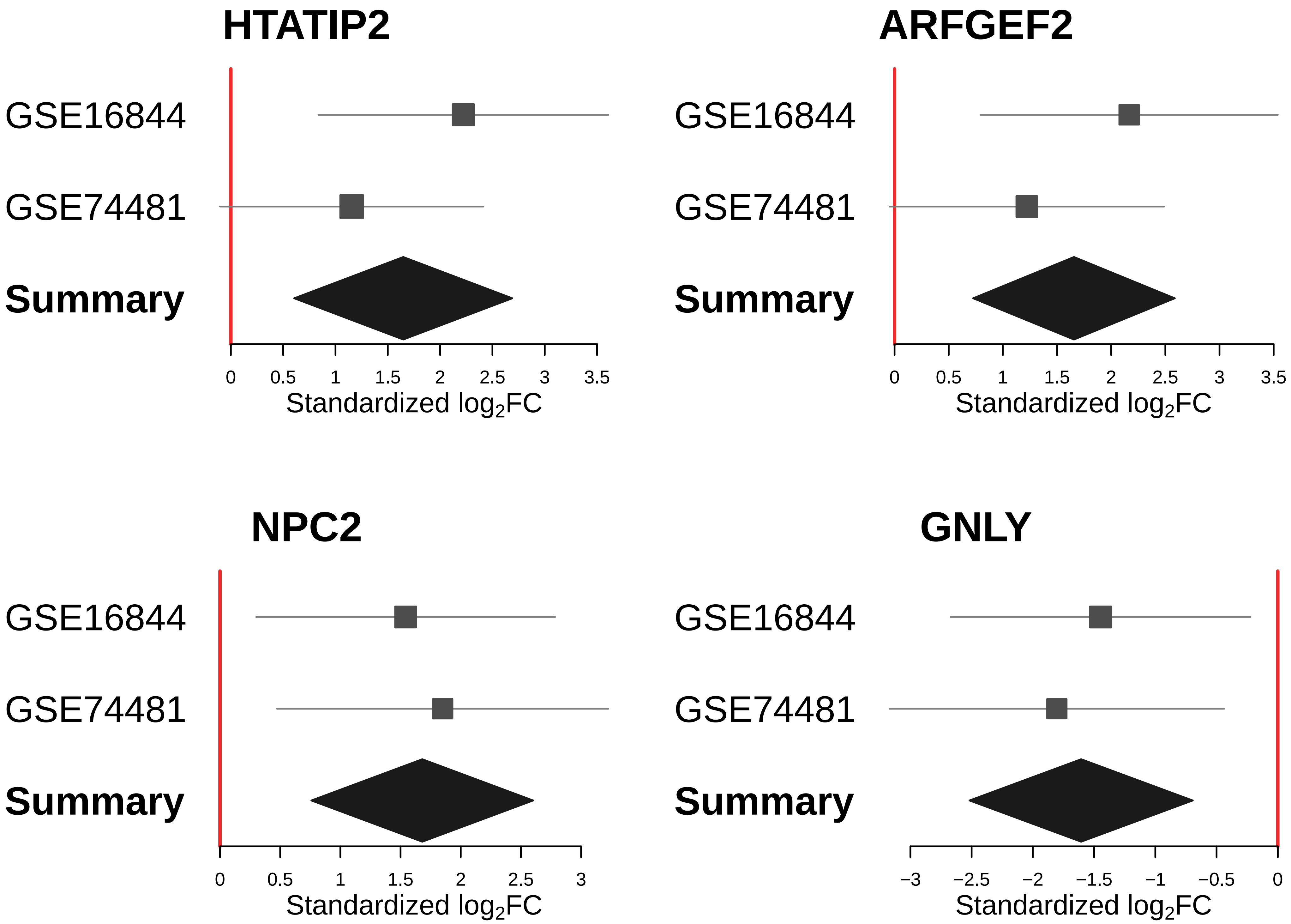

HTATIP2
GSE16844
GSE74481
Summary
0 0.5 1 1.5 2 2.5 3 3.5
Standardized log2FC
ARFGEF2
GSE16844
GSE74481
Summary
0 0.5 1 1.5 2 2.5 3 3.5
Standardized log2FC
NPC2
GSE16844
GSE74481
Summary
0 0.5 1 1.5 2 2.5 3
Standardized log2FC
GNLY
GSE16844
GSE74481
Summary
−3 −2.5 −2 −1.5 −1 −0.5 0
Standardized log2FC

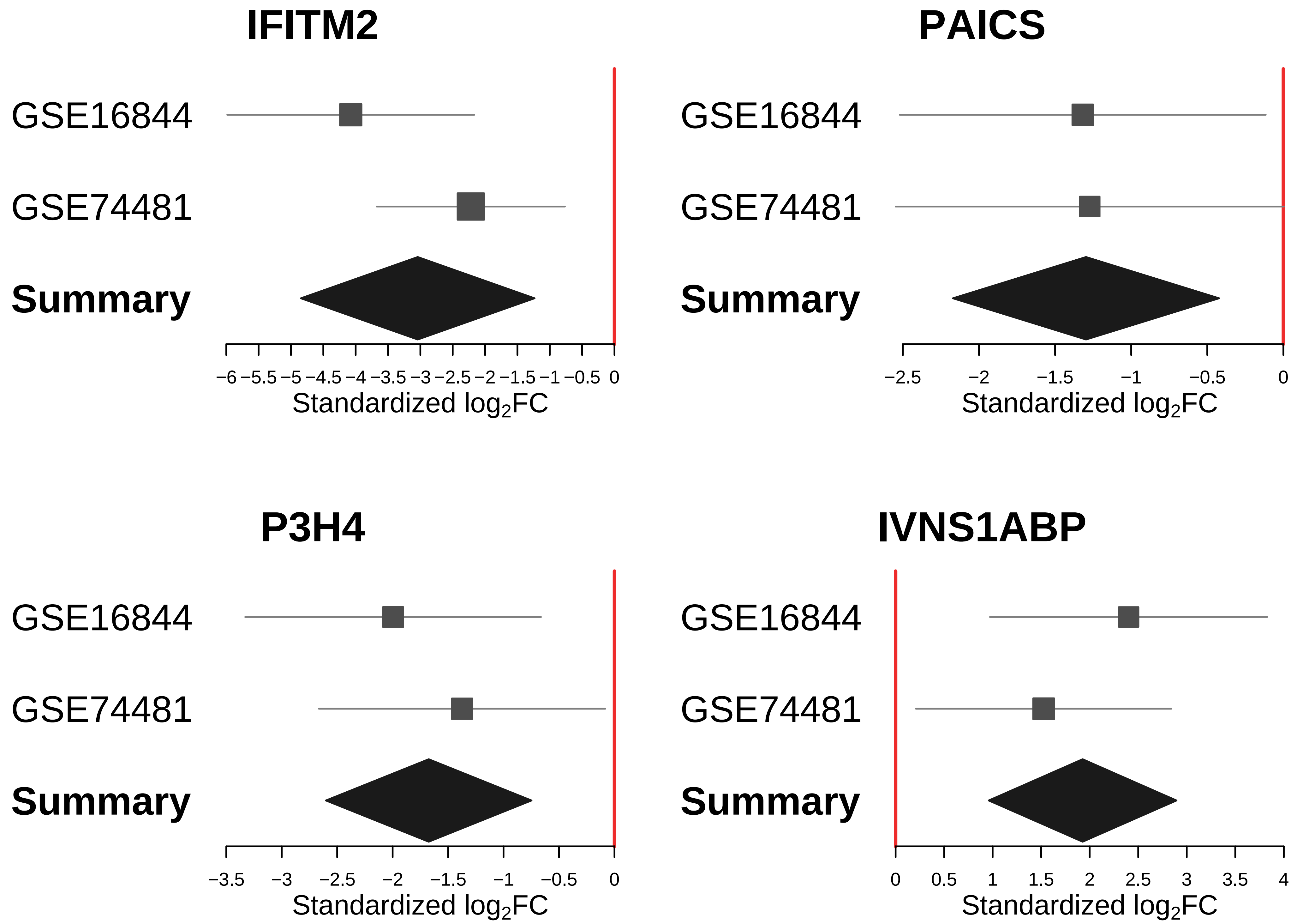
IFITM2
PAICS
P3H4
IVNS1ABP
GSE16844
GSE74481
Summary
Standardized log2FC
−6 −5.5 −5 −4.5 −4 −3.5 −3 −2.5 −2 −1.5 −1 −0.5 0
−2.5 −2 −1.5 −1 −0.5 0
−3.5 −3 −2.5 −2 −1.5 −1 −0.5 0
0 0.5 1 1.5 2 2.5 3 3.5 4

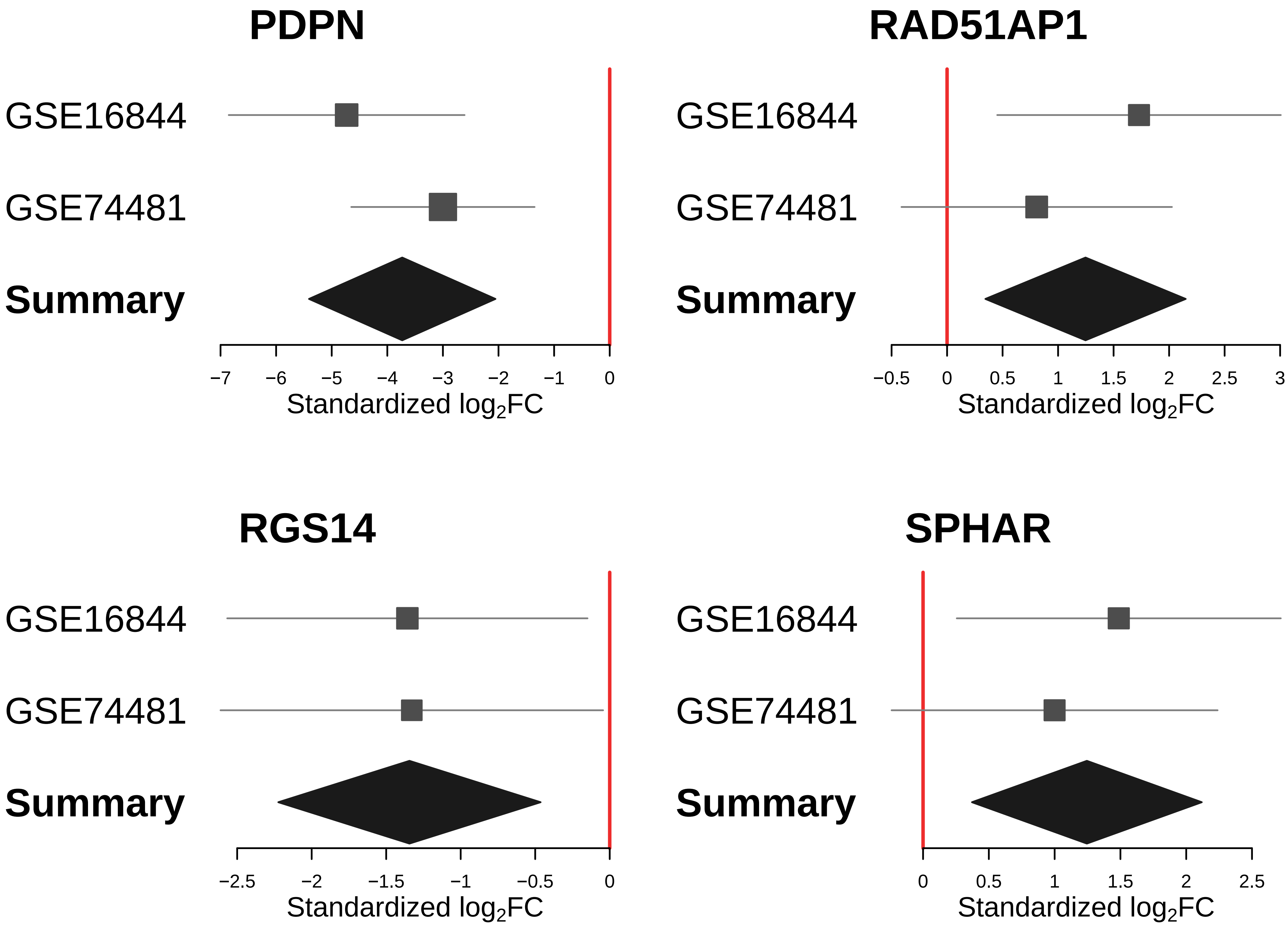
PDPN
GSE16844
GSE74481
Summary
−7 −6 −5 −4 −3 −2 −1 0
Standardized log2FC
RAD51AP1
GSE16844
GSE74481
Summary
−0.5 0 0.5 1 1.5 2 2.5 3
Standardized log2FC
RGS14
GSE16844
GSE74481
Summary
−2.5 −2 −1.5 −1 −0.5 0
Standardized log2FC
SPHAR
GSE16844
GSE74481
Summary
0 0.5 1 1.5 2 2.5
Standardized log2FC

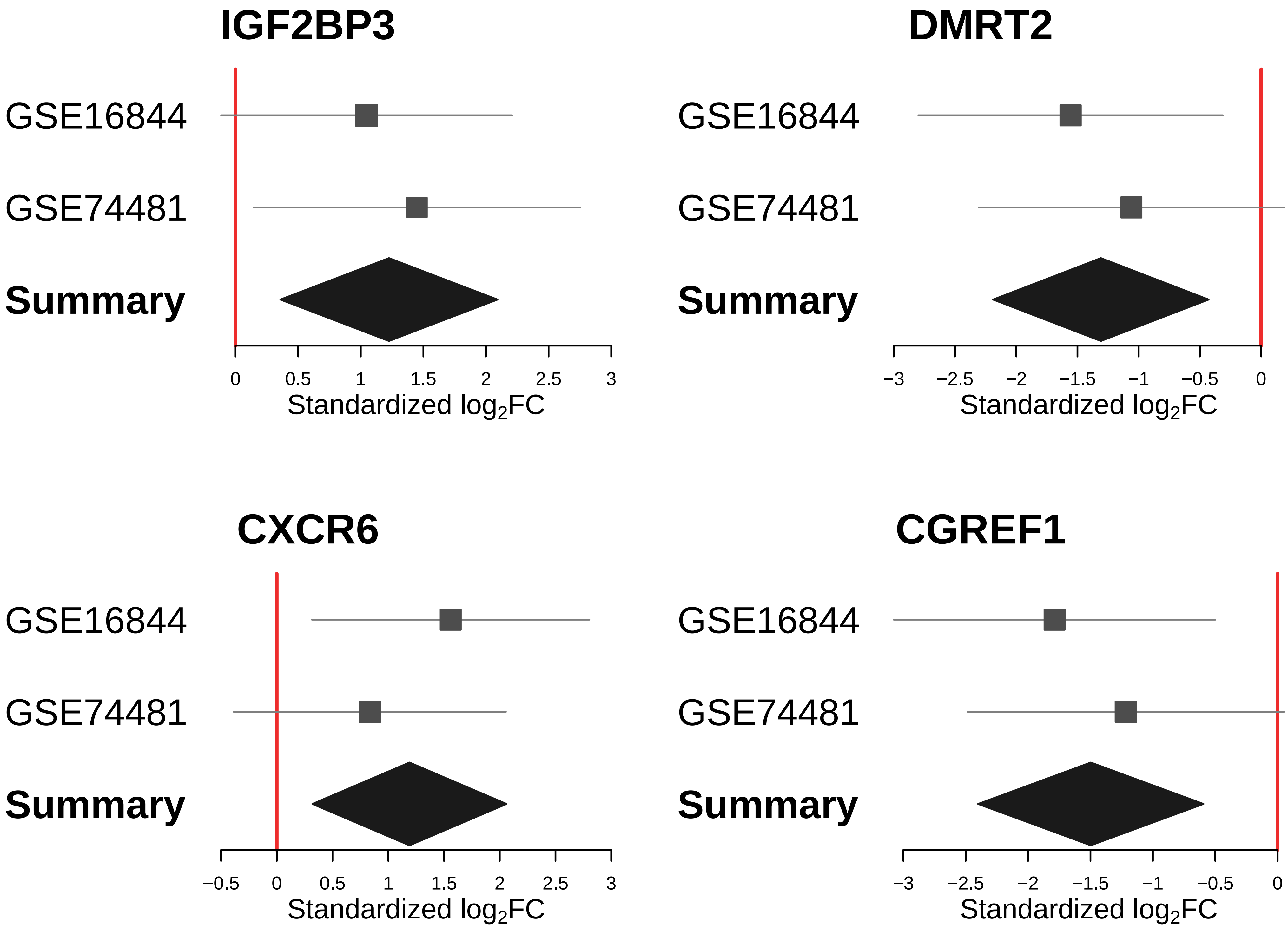
IGF2BP3
GSE16844
GSE74481
Summary
0
0.5
1
1.5
2
2.5
3
Standardized log2FC
DMRT2
GSE16844
GSE74481
Summary
−3
−2.5
−2
−1.5
−1
−0.5
0
Standardized log2FC
CXCR6
GSE16844
GSE74481
Summary
−0.5
0
0.5
1
1.5
2
2.5
3
Standardized log2FC
CGREF1
GSE16844
GSE74481
Summary
−3
−2.5
−2
−1.5
−1
−0.5
0
Standardized log2FC

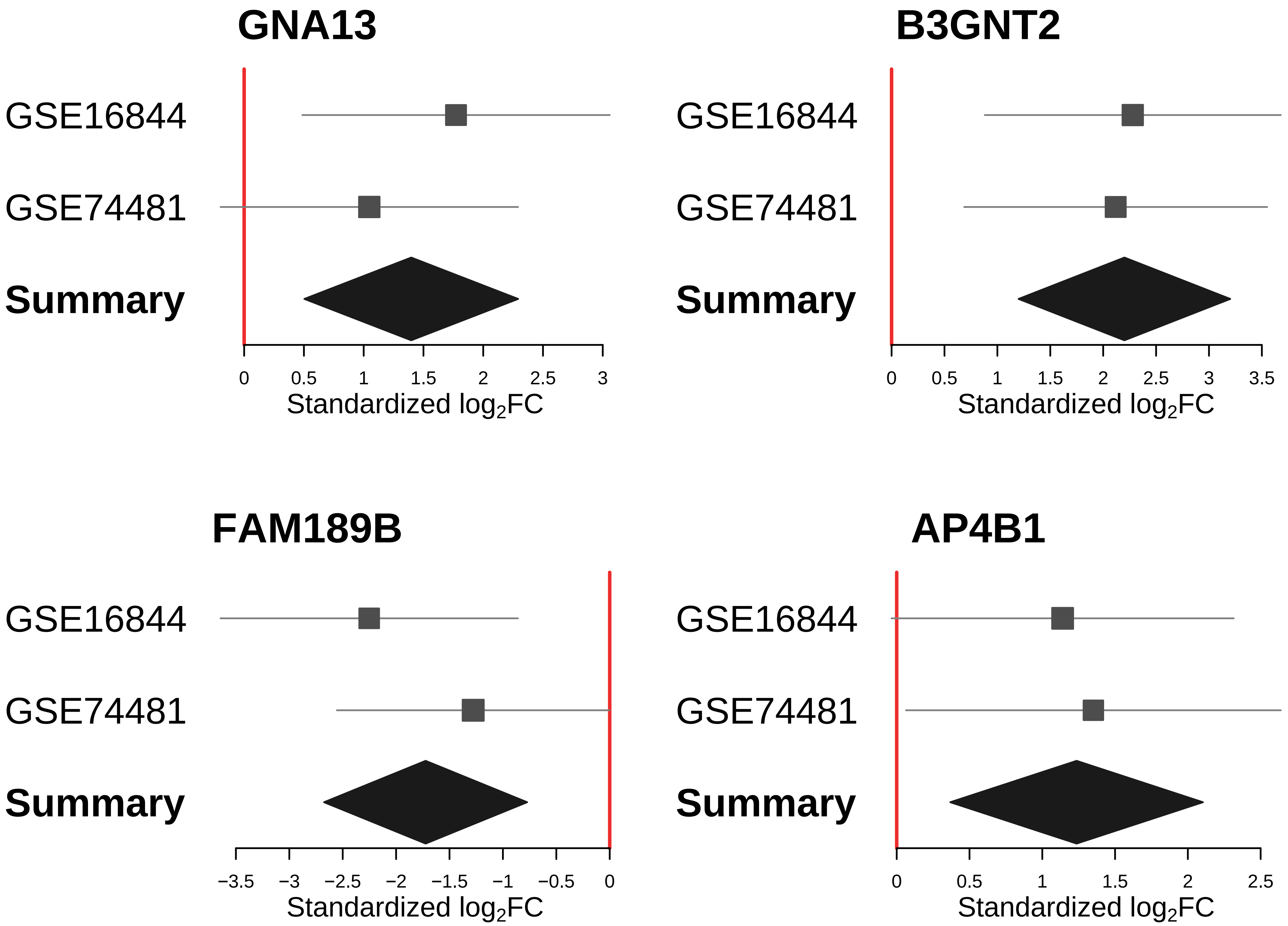

GNA13
GSE16844
GSE74481
Summary
0
0.5
1
1.5
2
2.5
3
Standardized log2FC
B3GNT2
GSE16844
GSE74481
Summary
0
0.5
1
1.5
2
2.5
3
3.5
Standardized log2FC
FAM189B
GSE16844
GSE74481
Summary
−3.5
−3
−2.5
−2
−1.5
−1
−0.5
0
Standardized log2FC
AP4B1
GSE16844
GSE74481
Summary
0
0.5
1
1.5
2
2.5
Standardized log2FC

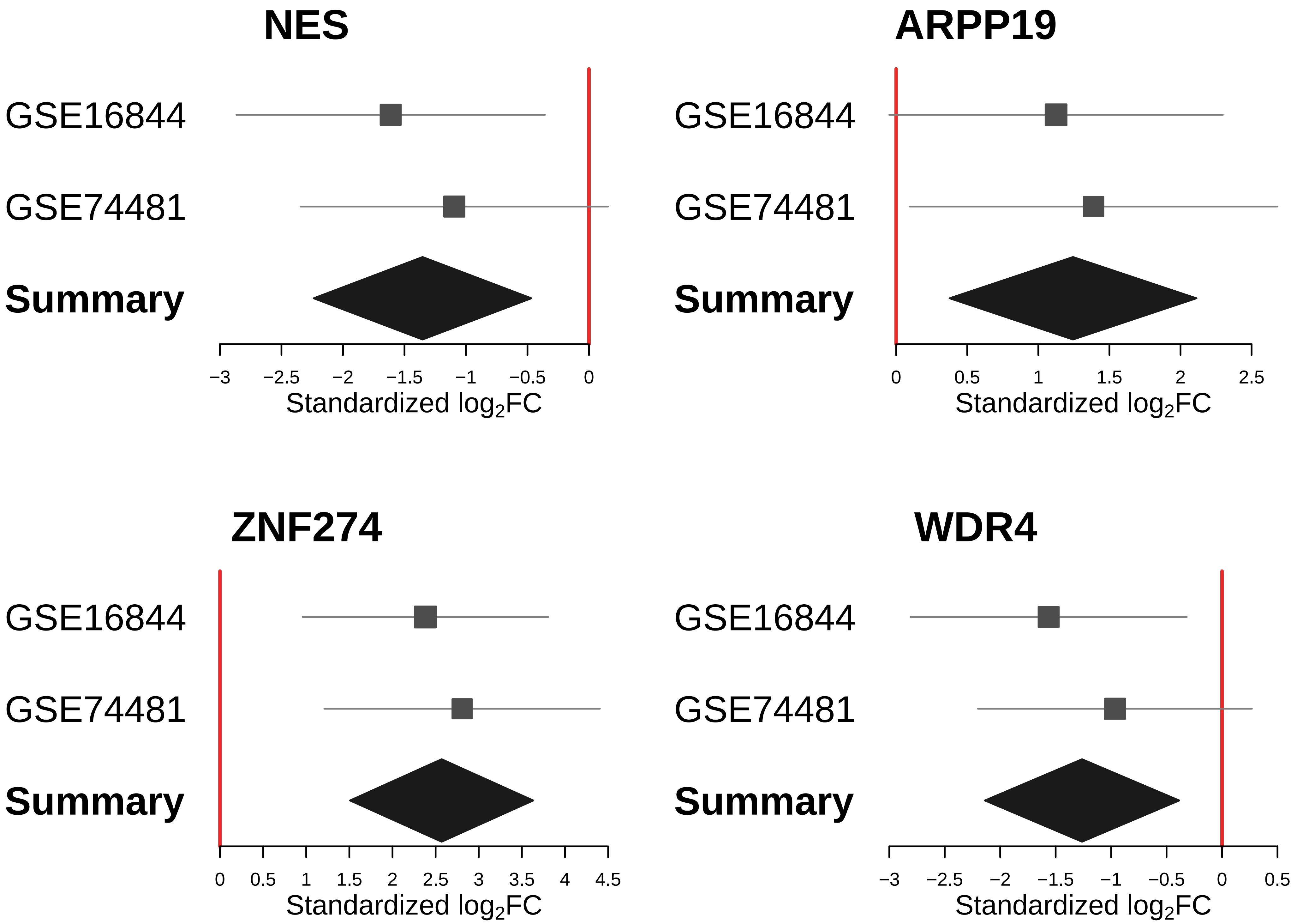
NES
GSE16844
GSE74481
Summary
−3
−2.5
−2
−1.5
−1
−0.5
0
Standardized log2FC
ARPP19
GSE16844
GSE74481
Summary
0
0.5
1
1.5
2
2.5
Standardized log2FC
ZNF274
GSE16844
GSE74481
Summary
0
0.5
1
1.5
2
2.5
3
3.5
4
4.5
Standardized log2FC
WDR4
GSE16844
GSE74481
Summary
−3
−2.5
−2
−1.5
−1
−0.5
0
0.5
Standardized log2FC

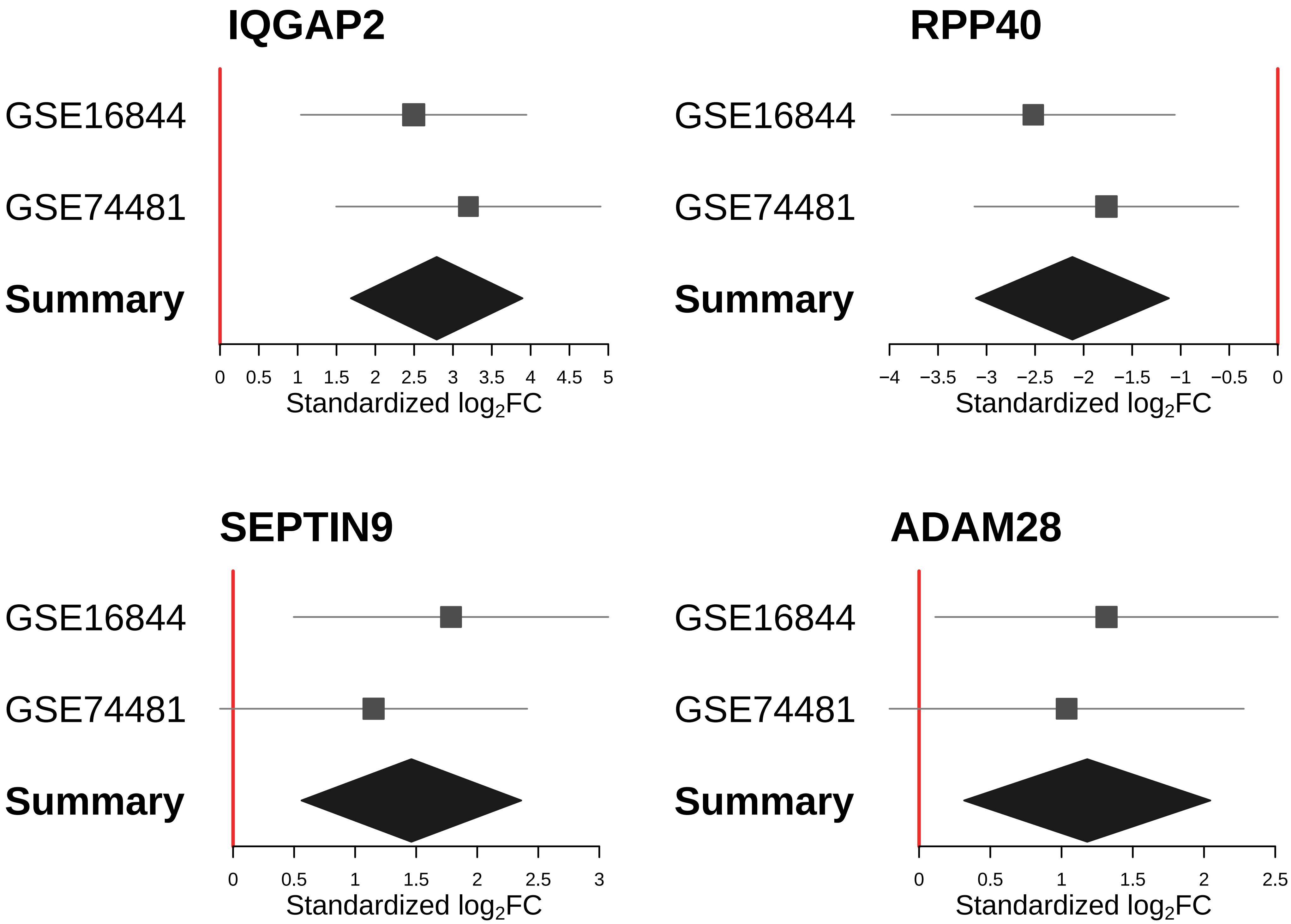

IQGAP2
GSE16844
GSE74481
Summary
0 0.5 1 1.5 2 2.5 3 3.5 4 4.5 5
Standardized log2FC
RPP40
GSE16844
GSE74481
Summary
−4 −3.5 −3 −2.5 −2 −1.5 −1 −0.5 0
Standardized log2FC
SEPTIN9
GSE16844
GSE74481
Summary
0 0.5 1 1.5 2 2.5 3
Standardized log2FC
ADAM28
GSE16844
GSE74481
Summary
0 0.5 1 1.5 2 2.5
Standardized log2FC

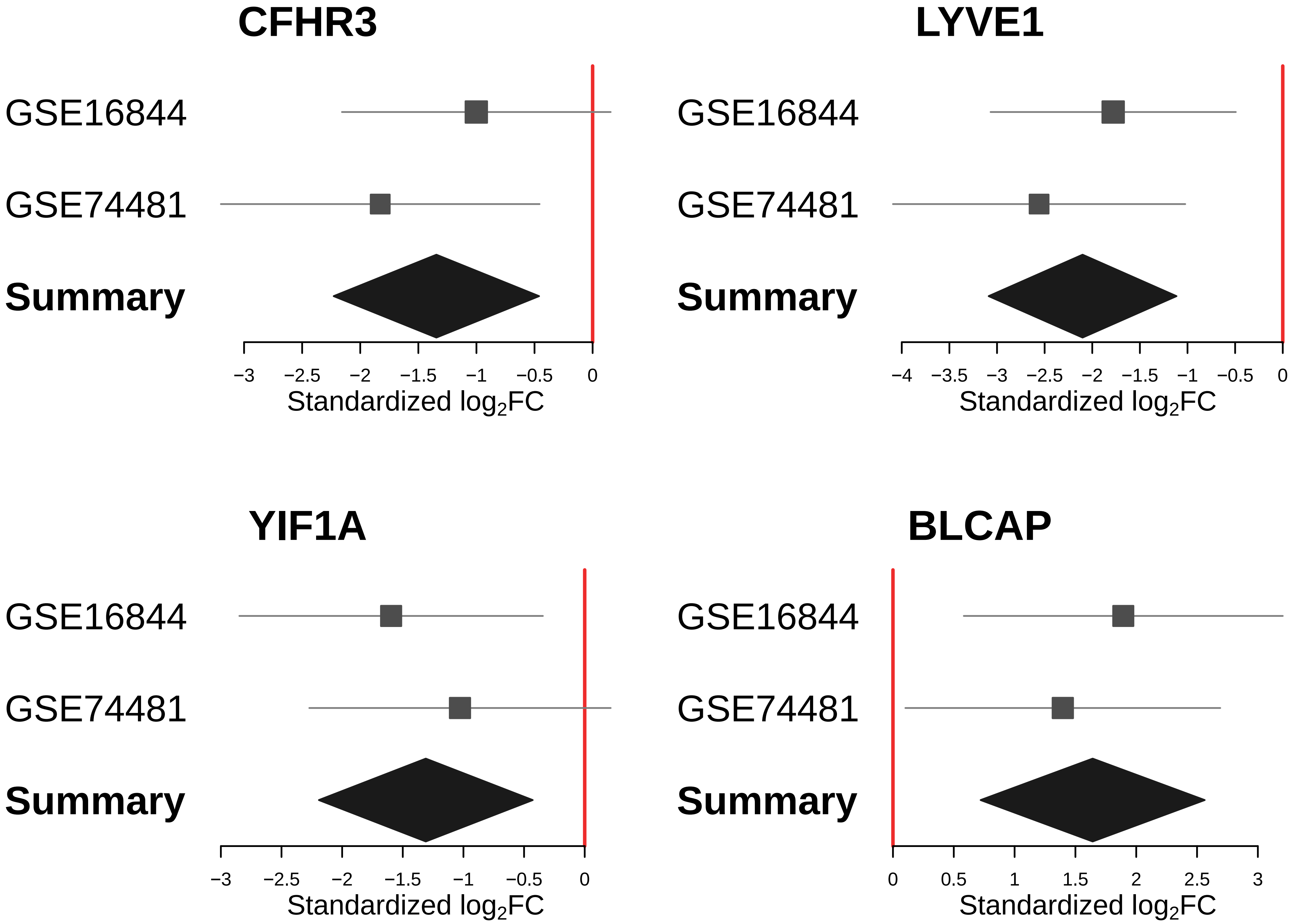

CFHR3
GSE16844
GSE74481
Summary
−3 −2.5 −2 −1.5 −1 −0.5 0
Standardized log2FC
LYVE1
GSE16844
GSE74481
Summary
−4 −3.5 −3 −2.5 −2 −1.5 −1 −0.5 0
Standardized log2FC
YIF1A
GSE16844
GSE74481
Summary
−3 −2.5 −2 −1.5 −1 −0.5 0
Standardized log2FC
BLCAP
GSE16844
GSE74481
Summary
0 0.5 1 1.5 2 2.5 3
Standardized log2FC

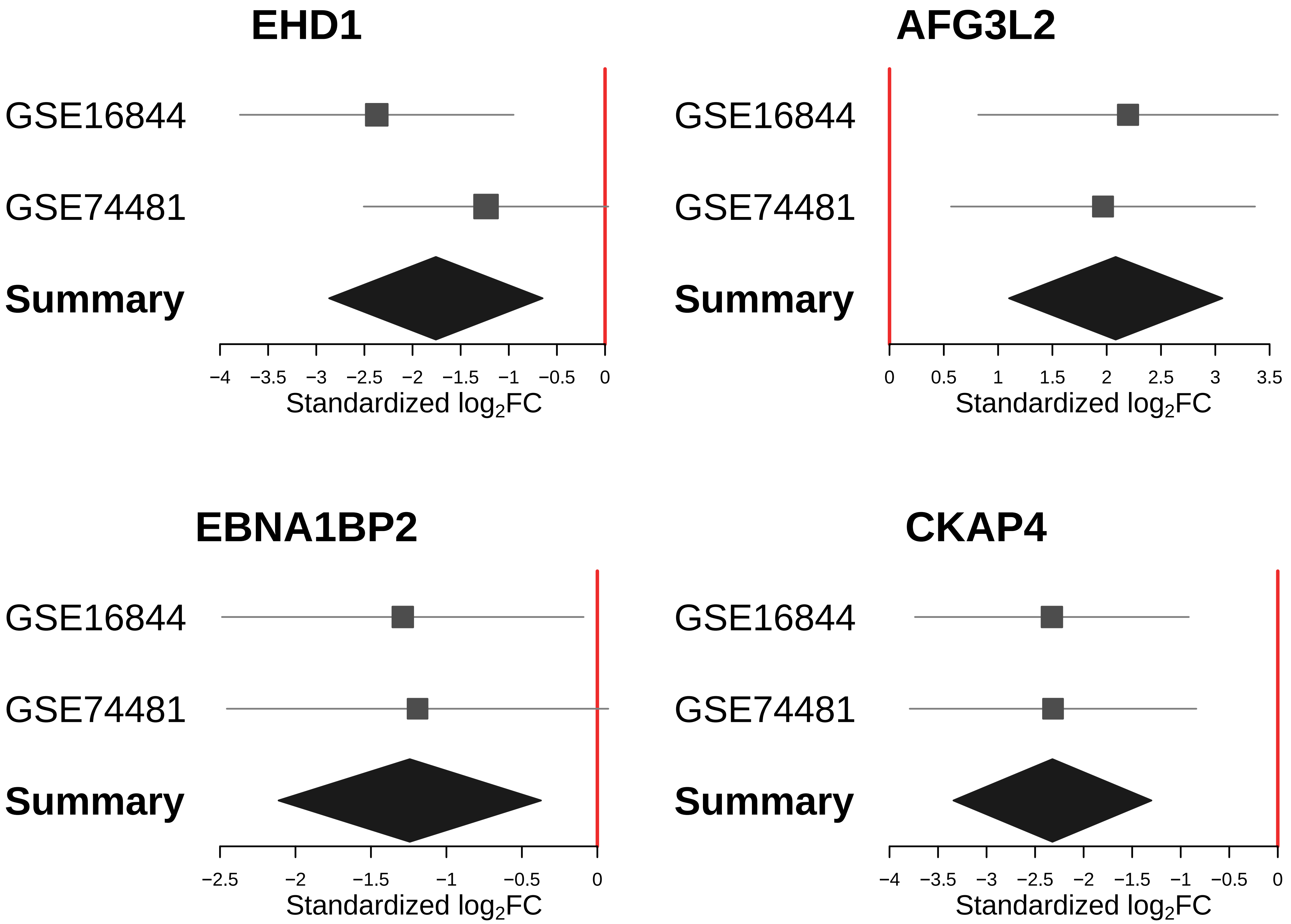
EHD1
GSE16844
GSE74481
Summary
−4 −3.5 −3 −2.5 −2 −1.5 −1 −0.5 0
Standardized log2FC
AFG3L2
GSE16844
GSE74481
Summary
0 0.5 1 1.5 2 2.5 3 3.5
Standardized log2FC
EBNA1BP2
GSE16844
GSE74481
Summary
−2.5 −2 −1.5 −1 −0.5 0
Standardized log2FC
CKAP4
GSE16844
GSE74481
Summary
−4 −3.5 −3 −2.5 −2 −1.5 −1 −0.5 0
Standardized log2FC

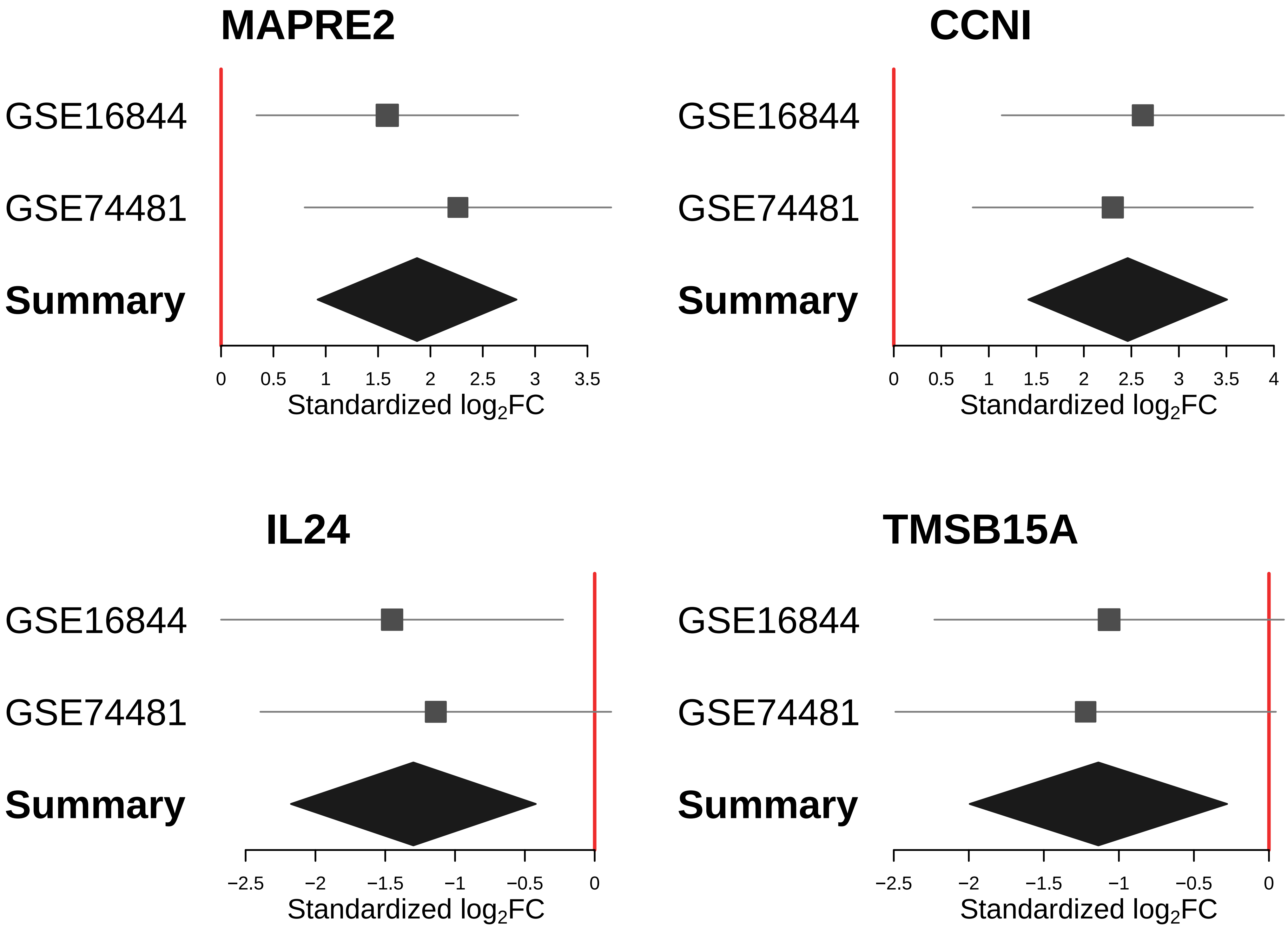
MAPRE2
GSE16844
GSE74481
Summary
0
0.5
1
1.5
2
2.5
3
3.5
Standardized log2FC
CCNI
GSE16844
GSE74481
Summary
0
0.5
1
1.5
2
2.5
3
3.5
4
Standardized log2FC
IL24
GSE16844
GSE74481
Summary
−2.5
−2
−1.5
−1
−0.5
0
Standardized log2FC
TMSB15A
GSE16844
GSE74481
Summary
−2.5
−2
−1.5
−1
−0.5
0
Standardized log2FC

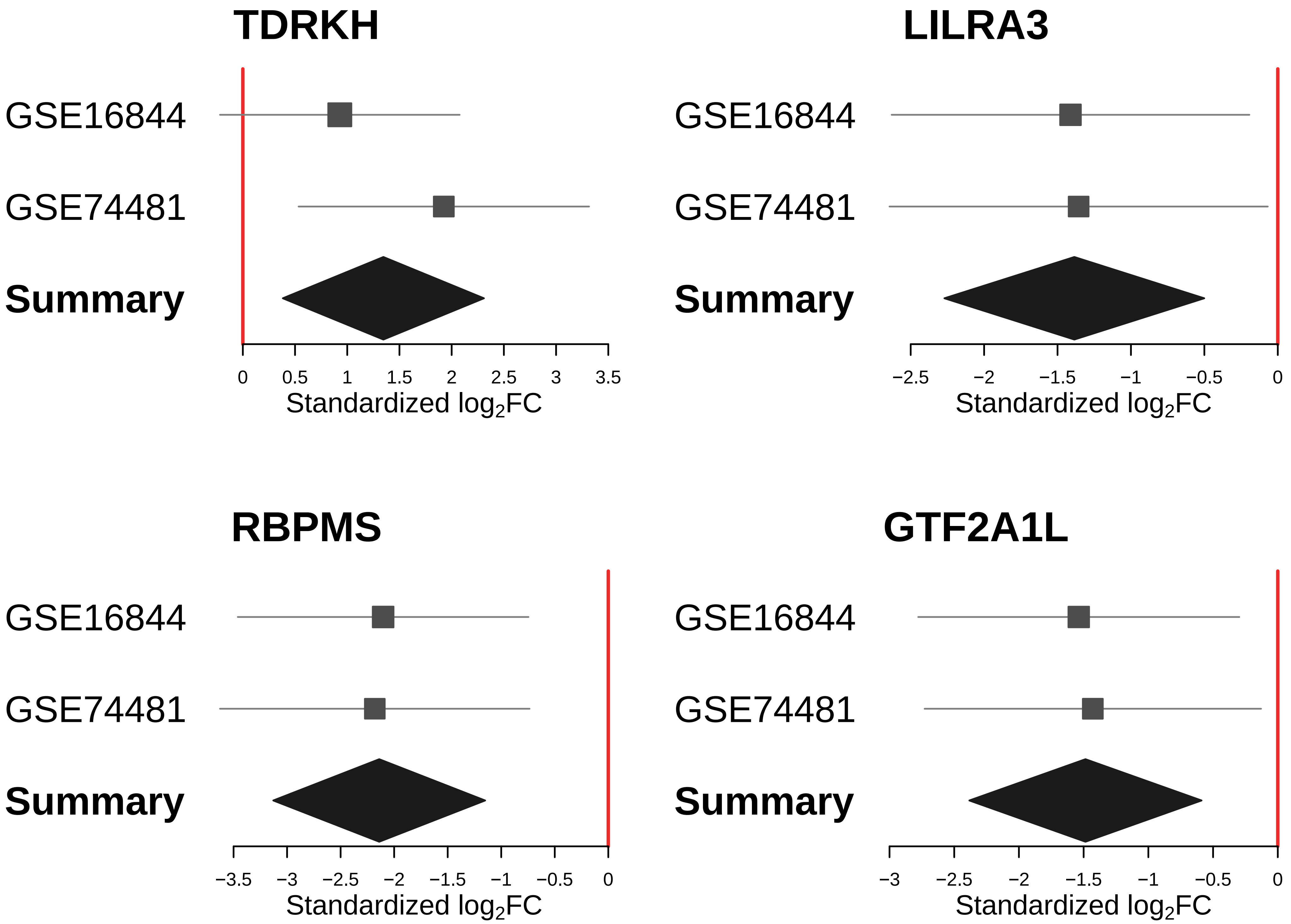

TDRKH
GSE16844
GSE74481
Summary
0 0.5 1 1.5 2 2.5 3 3.5
Standardized log2FC
LILRA3
GSE16844
GSE74481
Summary
−2.5 −2 −1.5 −1 −0.5 0
Standardized log2FC
RBPMS
GSE16844
GSE74481
Summary
−3.5 −3 −2.5 −2 −1.5 −1 −0.5 0
Standardized log2FC
GTF2A1L
GSE16844
GSE74481
Summary
−3 −2.5 −2 −1.5 −1 −0.5 0
Standardized log2FC

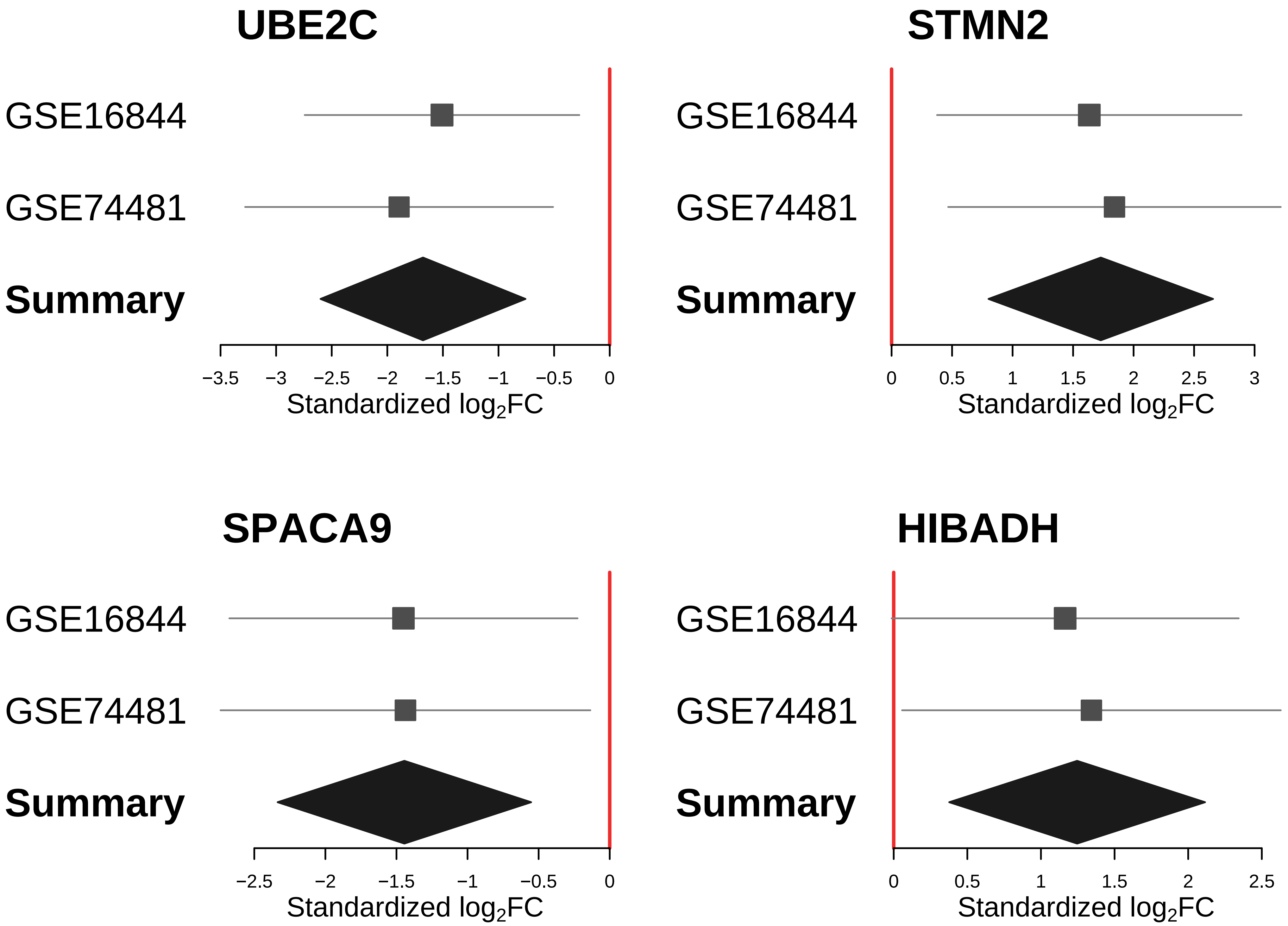

UBE2C
GSE16844
GSE74481
Summary
−3.5 −3 −2.5 −2 −1.5 −1 −0.5 0
Standardized log2FC
STMN2
GSE16844
GSE74481
Summary
0 0.5 1 1.5 2 2.5 3
Standardized log2FC
SPACA9
GSE16844
GSE74481
Summary
−2.5 −2 −1.5 −1 −0.5 0
Standardized log2FC
HIBADH
GSE16844
GSE74481
Summary
0 0.5 1 1.5 2 2.5
Standardized log2FC

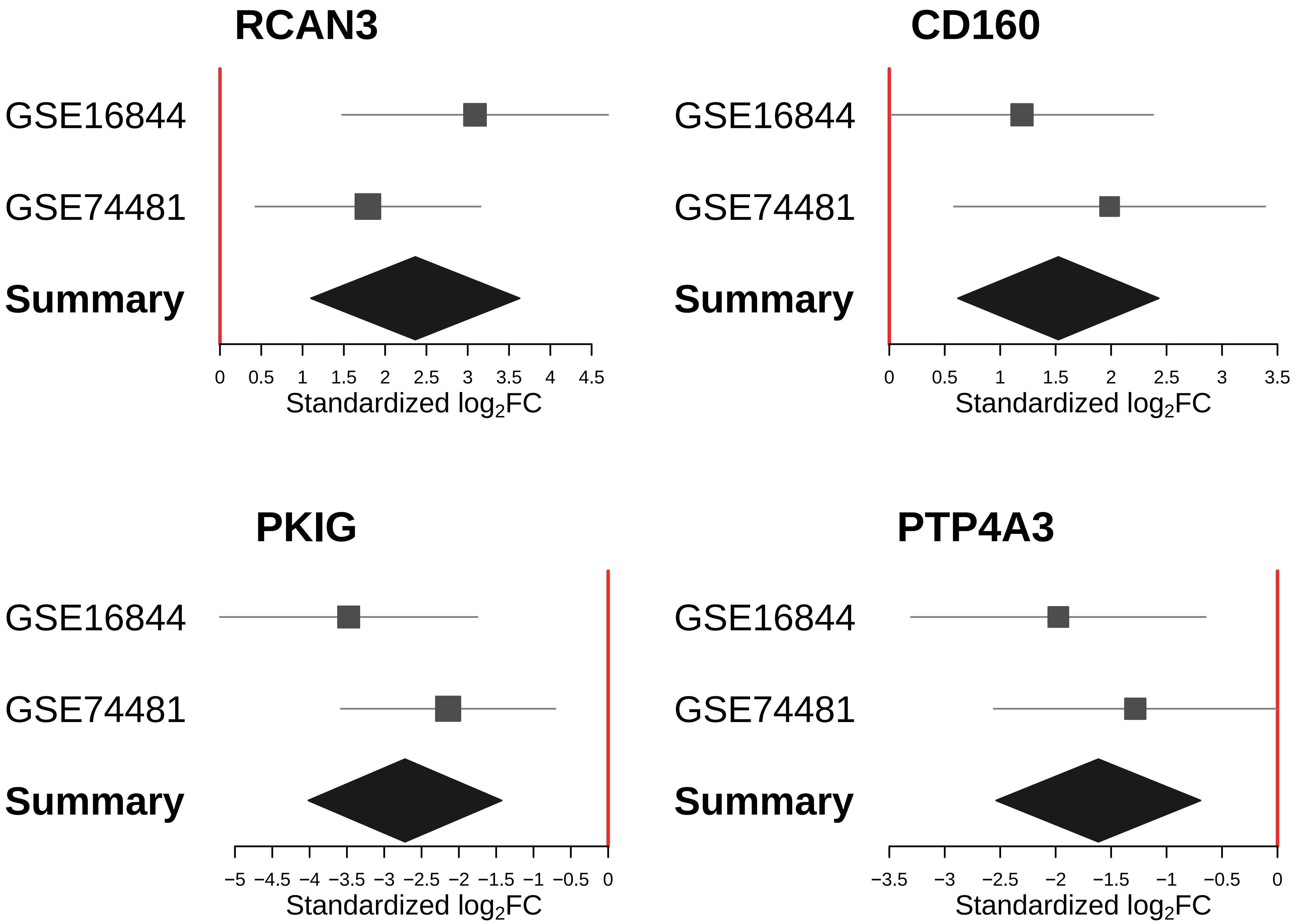
RCAN3
GSE16844
GSE74481
Summary
0 0.5 1 1.5 2 2.5 3 3.5 4 4.5
Standardized log2FC
CD160
GSE16844
GSE74481
Summary
0 0.5 1 1.5 2 2.5 3 3.5
Standardized log2FC
PKIG
GSE16844
GSE74481
Summary
−5 −4.5 −4 −3.5 −3 −2.5 −2 −1.5 −1 −0.5 0
Standardized log2FC
PTP4A3
GSE16844
GSE74481
Summary
−3.5 −3 −2.5 −2 −1.5 −1 −0.5 0
Standardized log2FC

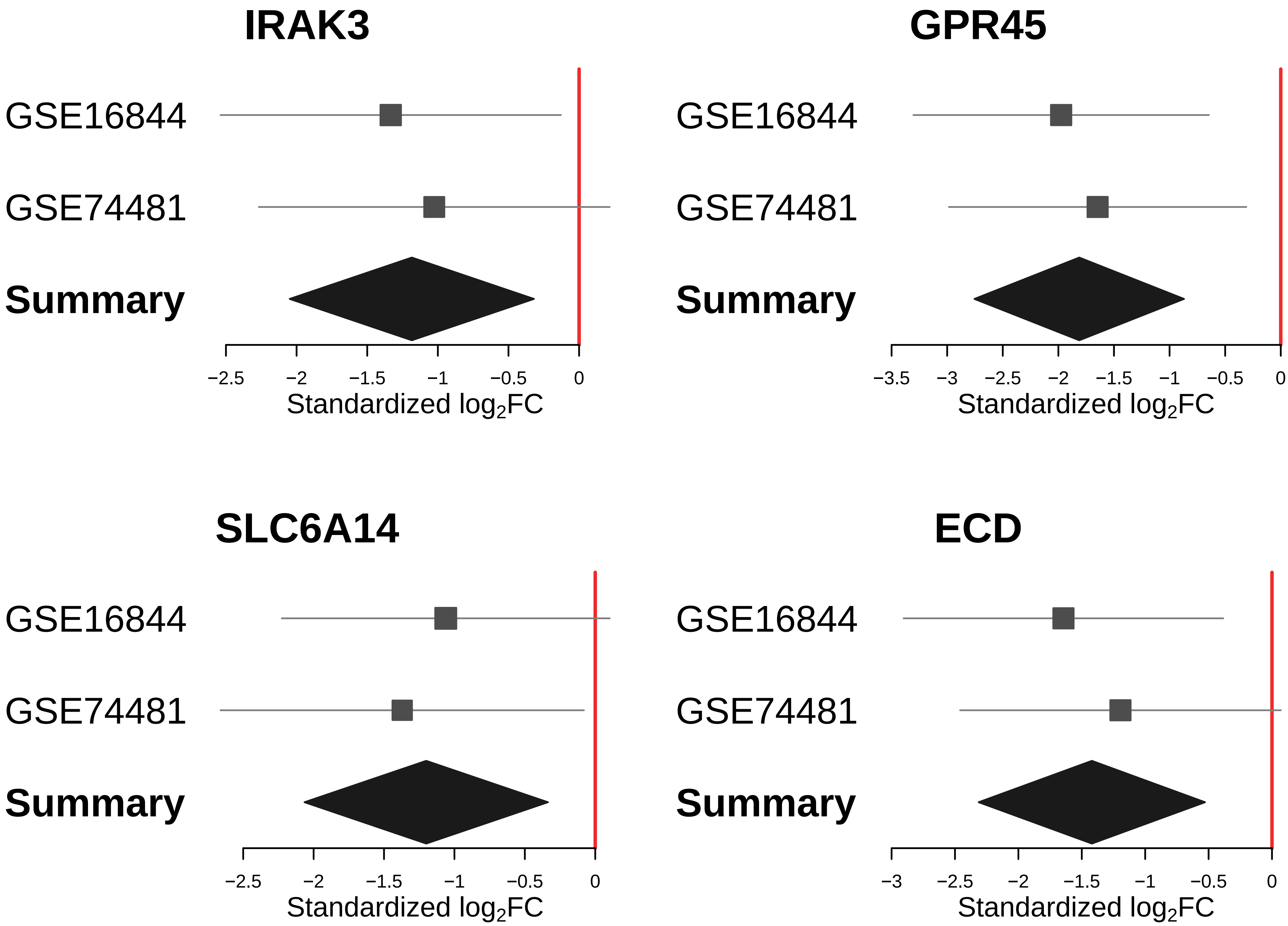

IRAK3
GSE16844
GSE74481
Summary
−2.5
−2
−1.5
−1
−0.5
0
Standardized log2FC
GPR45
GSE16844
GSE74481
Summary
−3.5
−3
−2.5
−2
−1.5
−1
−0.5
0
Standardized log2FC
SLC6A14
GSE16844
GSE74481
Summary
−2.5
−2
−1.5
−1
−0.5
0
Standardized log2FC
ECD
GSE16844
GSE74481
Summary
−3
−2.5
−2
−1.5
−1
−0.5
0
Standardized log2FC

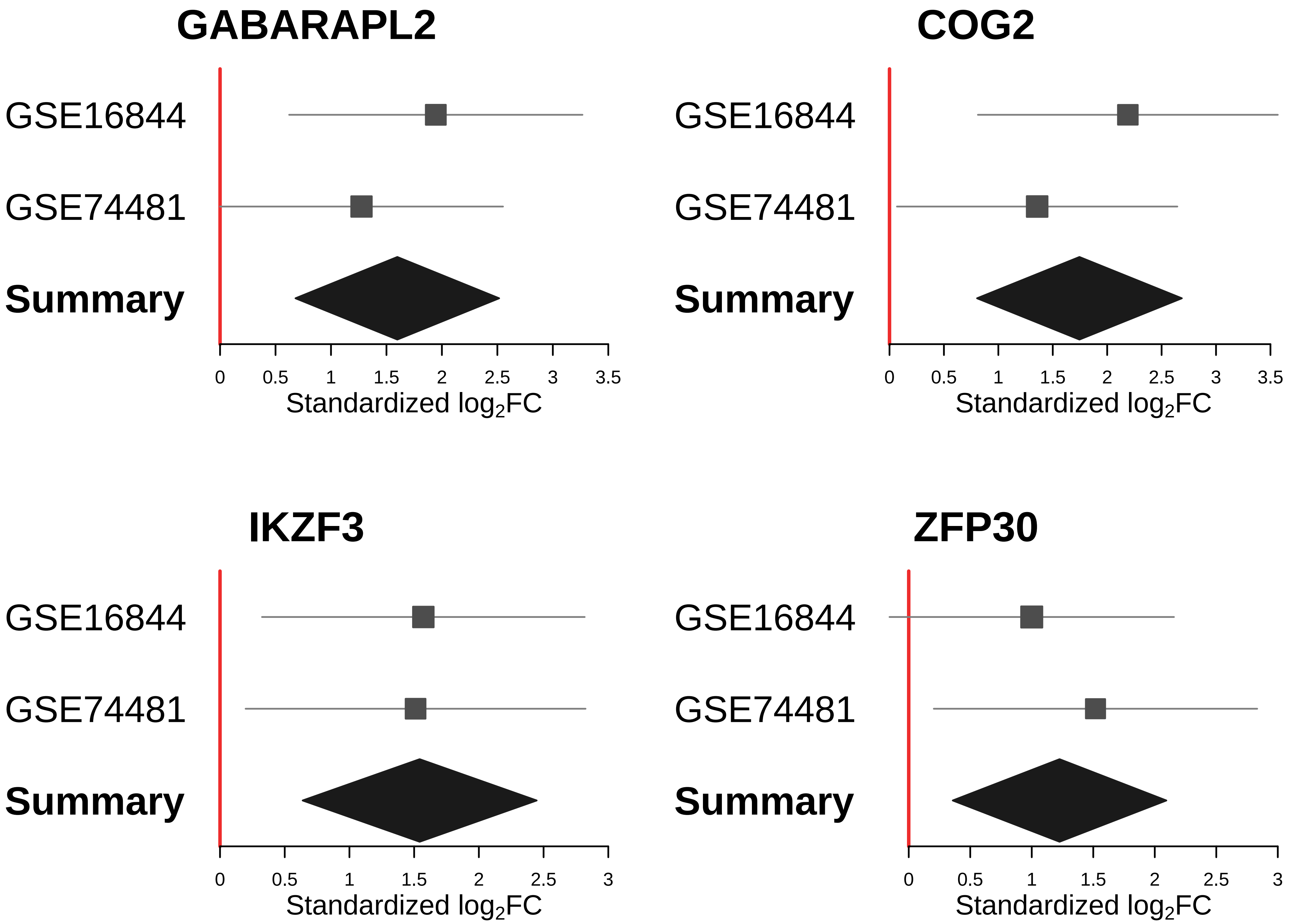
GABARAPL2
COG2
IKZF3
ZFP30
GSE16844
GSE74481
Summary
0
0.5
1
1.5
2
2.5
3
3.5
Standardized log2FC

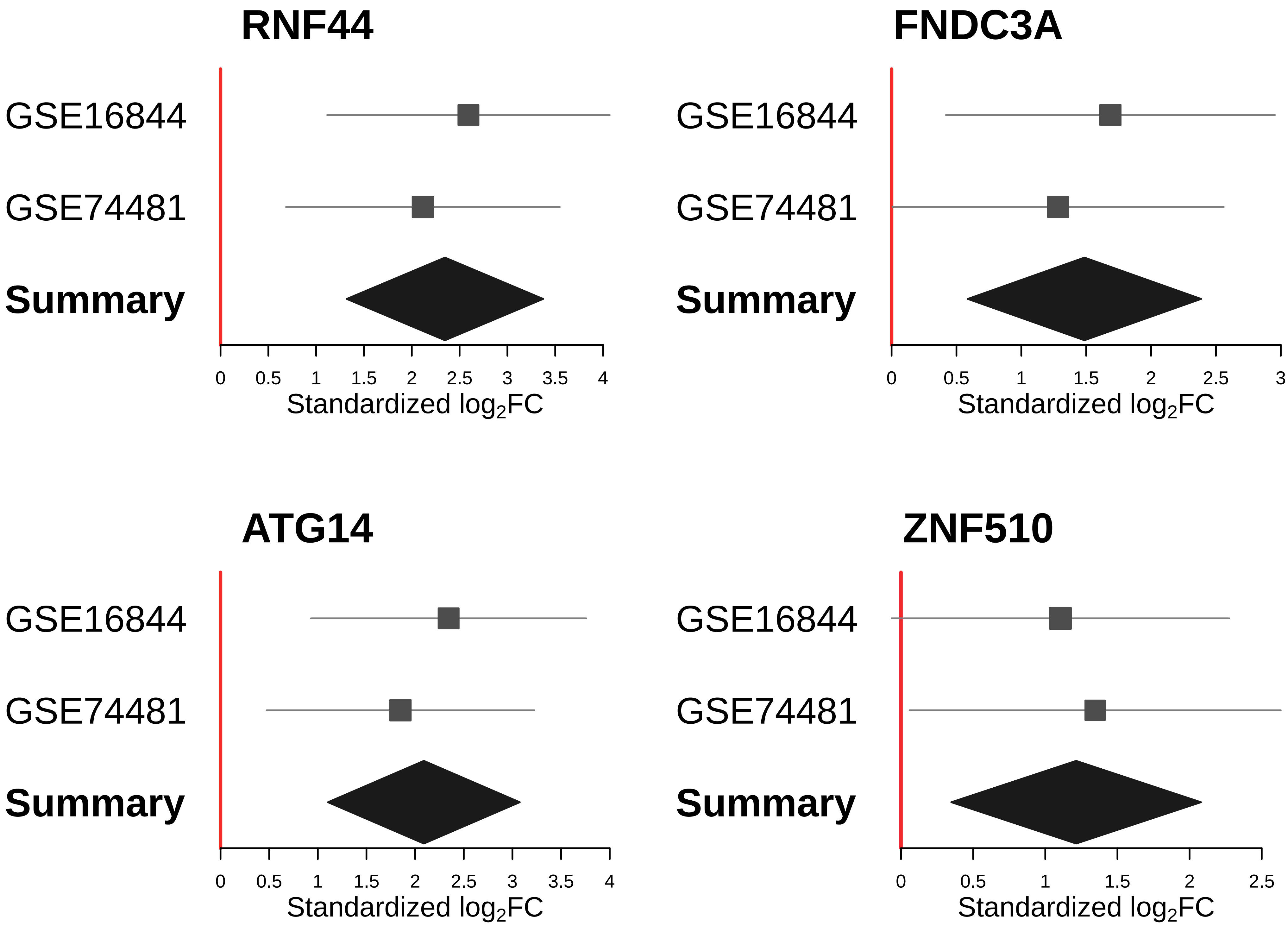
RNF44
FNDC3A
ATG14
ZNF510
GSE16844
GSE74481
Summary
0 0.5 1 1.5 2 2.5 3 3.5 4
0 0.5 1 1.5 2 2.5 3
0 0.5 1 1.5 2 2.5
Standardized $log_2FC$

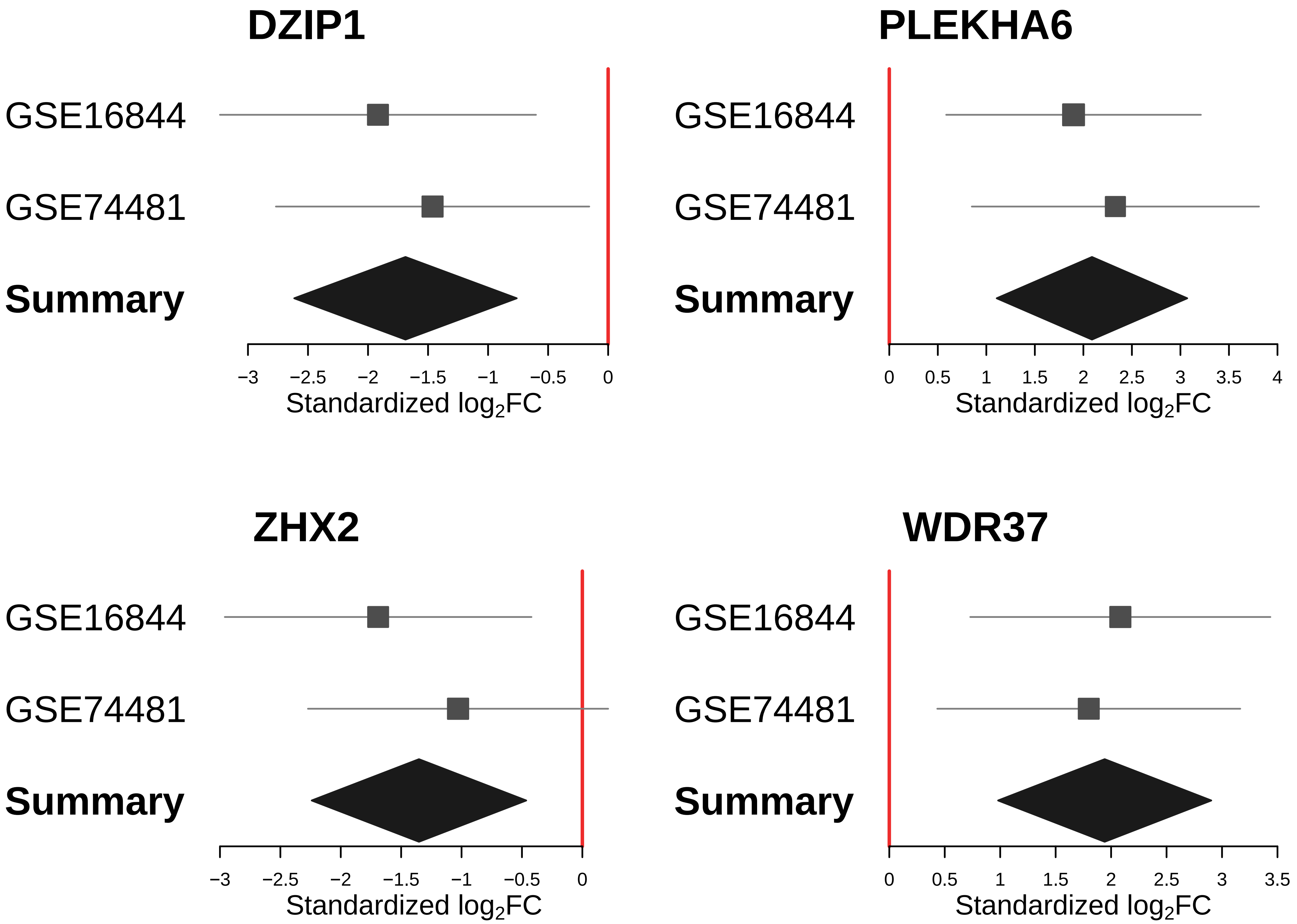
DZIP1
GSE16844
GSE74481
Summary
−3 −2.5 −2 −1.5 −1 −0.5 0
Standardized log2FC
PLEKHA6
GSE16844
GSE74481
Summary
0 0.5 1 1.5 2 2.5 3 3.5 4
Standardized log2FC
ZHX2
GSE16844
GSE74481
Summary
−3 −2.5 −2 −1.5 −1 −0.5 0
Standardized log2FC
WDR37
GSE16844
GSE74481
Summary
0 0.5 1 1.5 2 2.5 3 3.5
Standardized log2FC

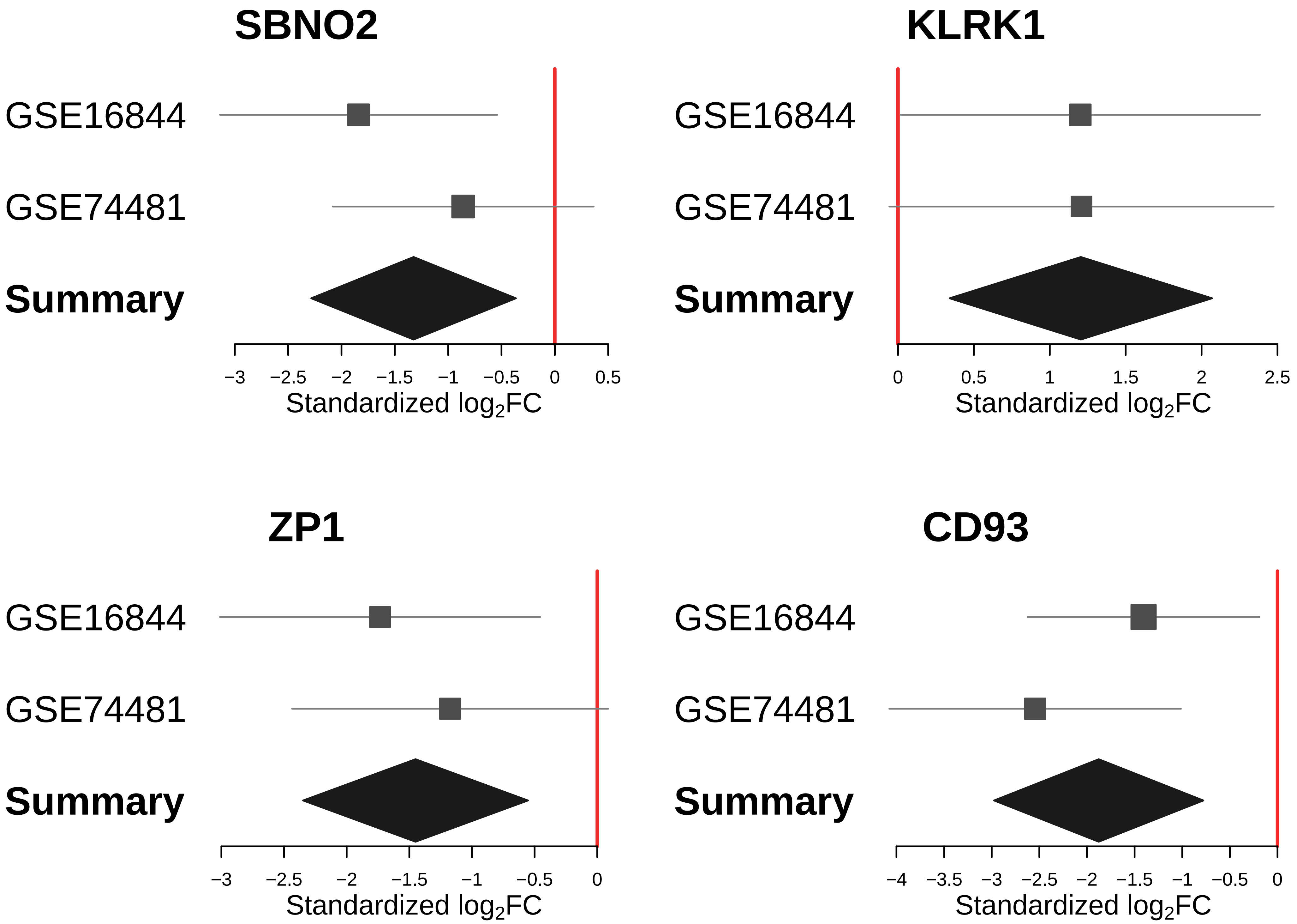
SBNO2
GSE16844
GSE74481
Summary
−3 −2.5 −2 −1.5 −1 −0.5 0 0.5
Standardized log2FC
KLRK1
GSE16844
GSE74481
Summary
0 0.5 1 1.5 2 2.5
Standardized log2FC
ZP1
GSE16844
GSE74481
Summary
−3 −2.5 −2 −1.5 −1 −0.5 0
Standardized log2FC
CD93
GSE16844
GSE74481
Summary
−4 −3.5 −3 −2.5 −2 −1.5 −1 −0.5 0
Standardized log2FC

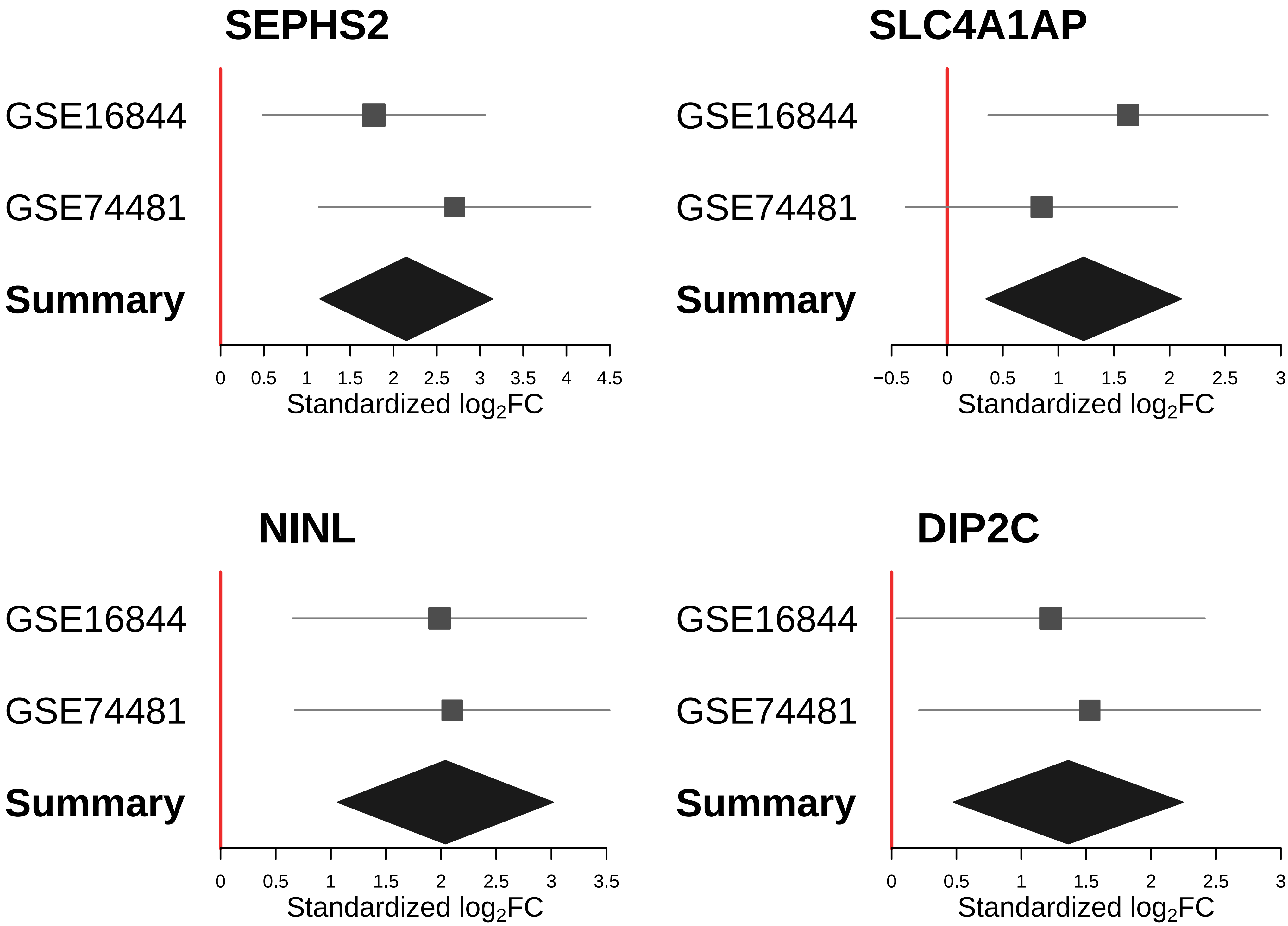
SEPHS2
GSE16844
GSE74481
Summary
0 0.5 1 1.5 2 2.5 3 3.5 4 4.5
Standardized log2FC
SLC4A1AP
GSE16844
GSE74481
Summary
−0.5 0 0.5 1 1.5 2 2.5 3
Standardized log2FC
NINL
GSE16844
GSE74481
Summary
0 0.5 1 1.5 2 2.5 3 3.5
Standardized log2FC
DIP2C
GSE16844
GSE74481
Summary
0 0.5 1 1.5 2 2.5 3
Standardized log2FC

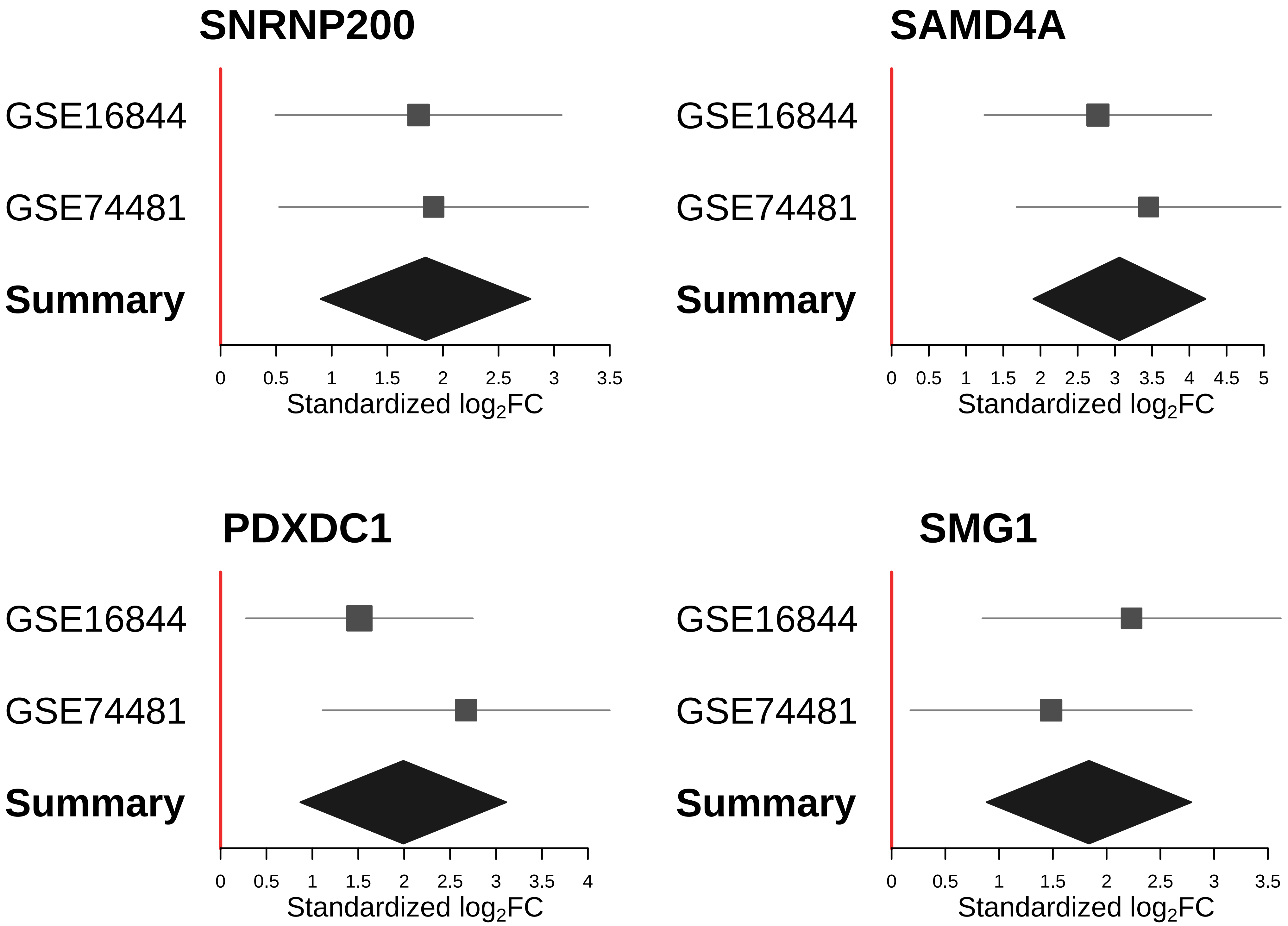
SNRNP200
GSE16844
GSE74481
Summary
0 0.5 1 1.5 2 2.5 3 3.5
Standardized log2FC
SAMD4A
GSE16844
GSE74481
Summary
0 0.5 1 1.5 2 2.5 3 3.5 4 4.5 5
Standardized log2FC
PDXDC1
GSE16844
GSE74481
Summary
0 0.5 1 1.5 2 2.5 3 3.5 4
Standardized log2FC
SMG1
GSE16844
GSE74481
Summary
0 0.5 1 1.5 2 2.5 3 3.5
Standardized log2FC

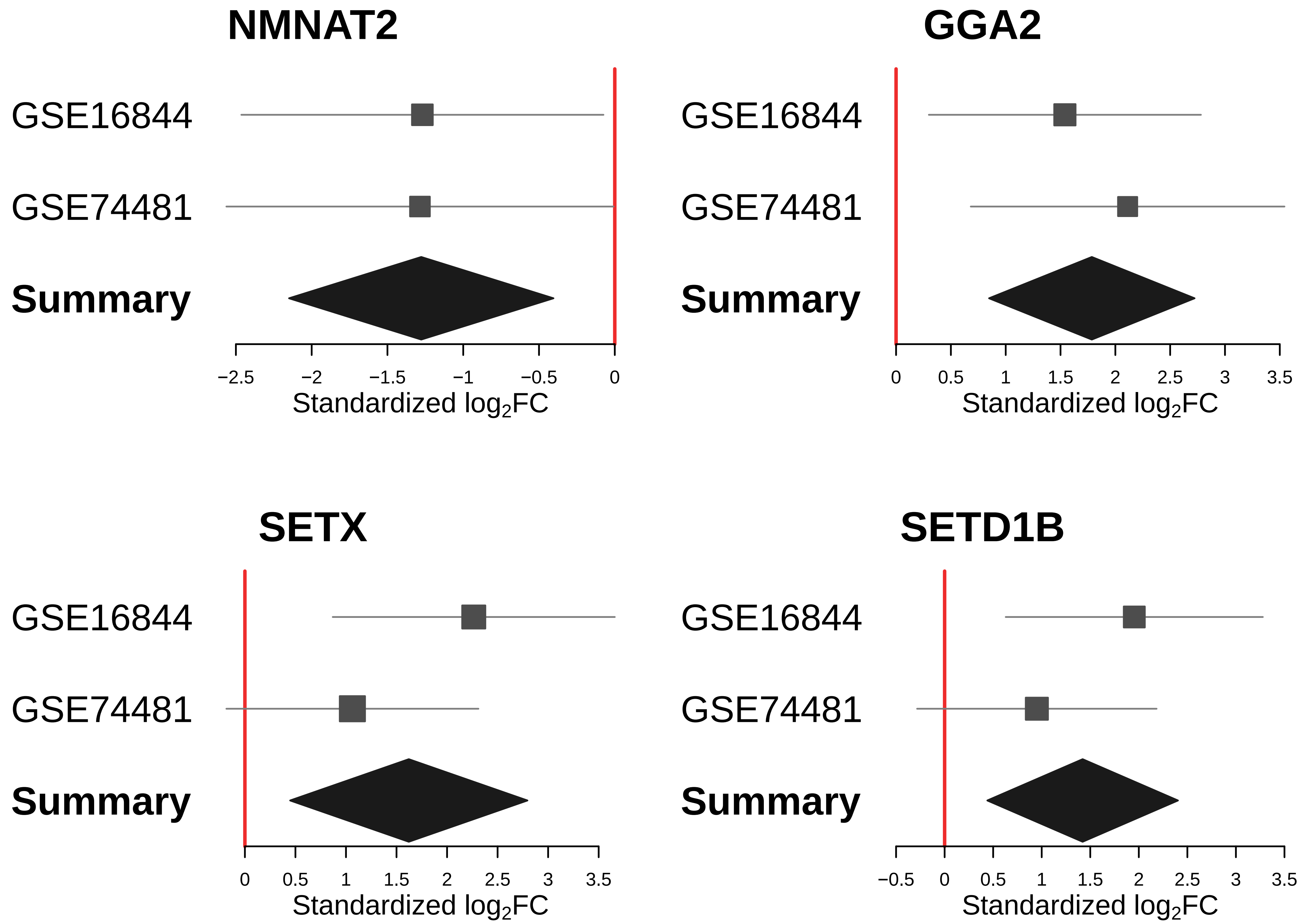
NMNAT2
GSE16844
GSE74481
Summary
−2.5 −2 −1.5 −1 −0.5 0
Standardized log2FC
GGA2
GSE16844
GSE74481
Summary
0 0.5 1 1.5 2 2.5 3 3.5
Standardized log2FC
SETX
GSE16844
GSE74481
Summary
0 0.5 1 1.5 2 2.5 3 3.5
Standardized log2FC
SETD1B
GSE16844
GSE74481
Summary
−0.5 0 0.5 1 1.5 2 2.5 3 3.5
Standardized log2FC

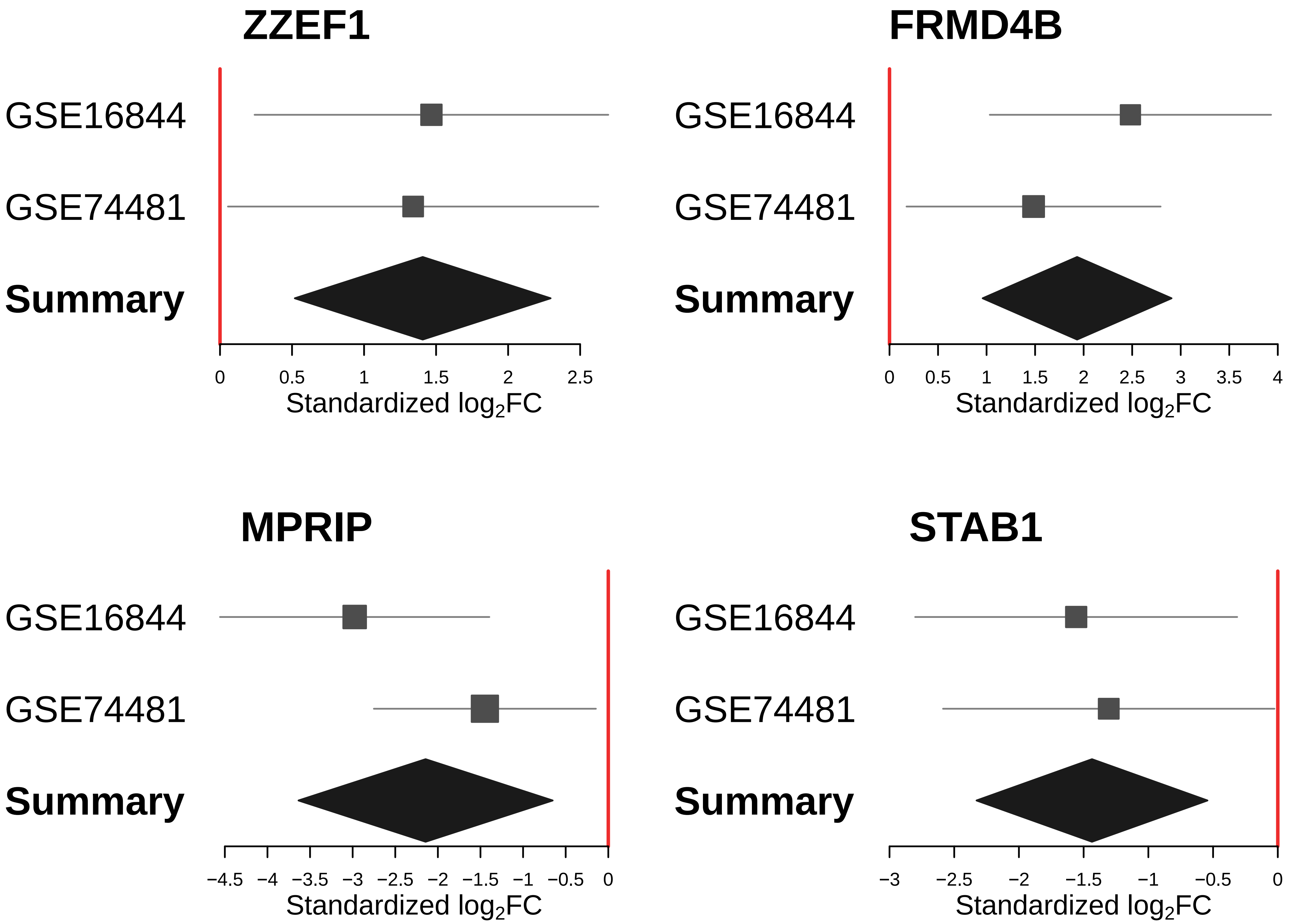

ZZEF1
GSE16844
GSE74481
Summary
0 0.5 1 1.5 2 2.5
Standardized log2FC
FRMD4B
GSE16844
GSE74481
Summary
0 0.5 1 1.5 2 2.5 3 3.5 4
Standardized log2FC
MPRIP
GSE16844
GSE74481
Summary
−4.5 −4 −3.5 −3 −2.5 −2 −1.5 −1 −0.5 0
Standardized log2FC
STAB1
GSE16844
GSE74481
Summary
−3 −2.5 −2 −1.5 −1 −0.5 0
Standardized log2FC

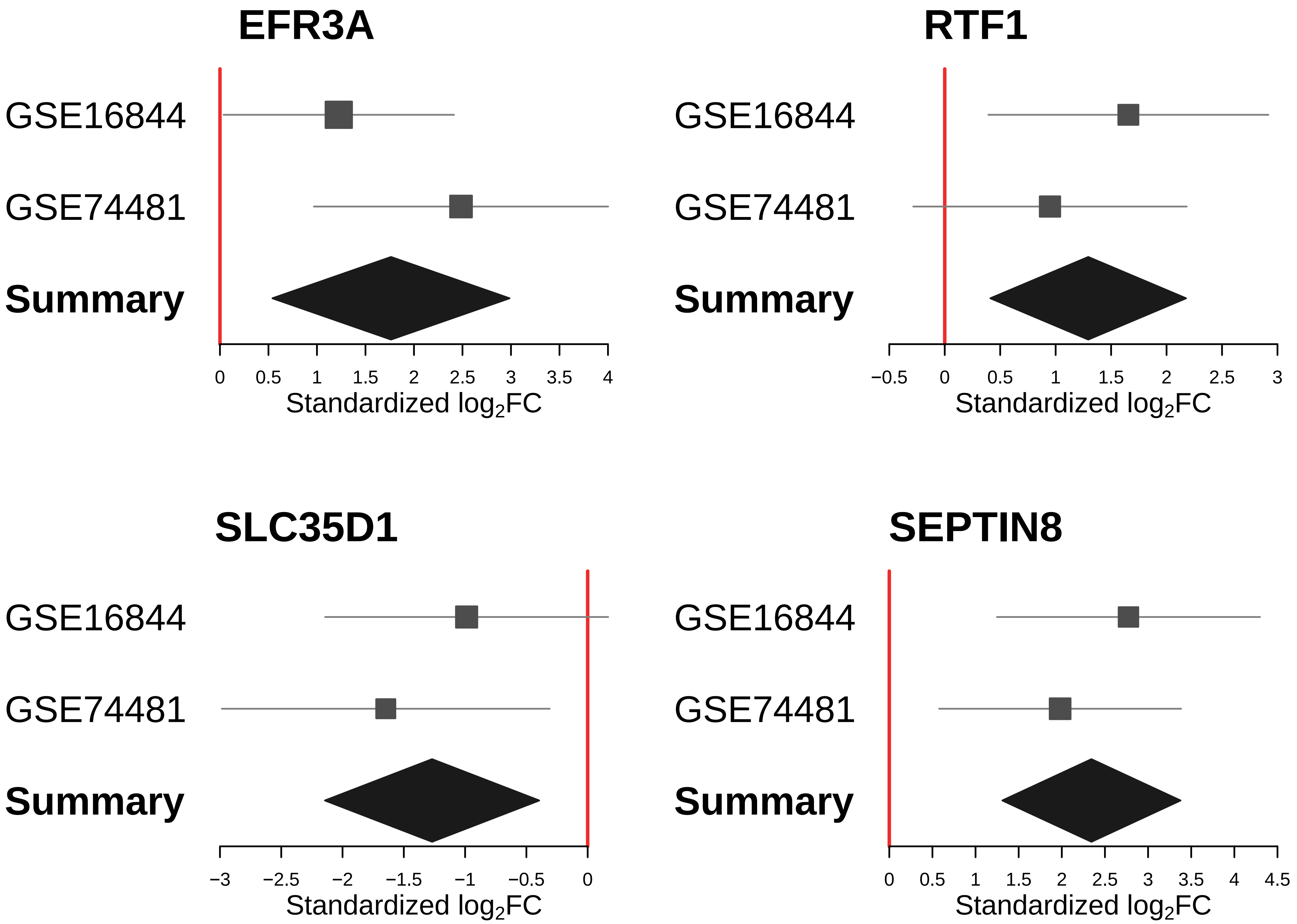

EFR3A
GSE16844
GSE74481
Summary
0 0.5 1 1.5 2 2.5 3 3.5 4
Standardized log2FC
RTF1
GSE16844
GSE74481
Summary
−0.5 0 0.5 1 1.5 2 2.5 3
Standardized log2FC
SLC35D1
GSE16844
GSE74481
Summary
−3 −2.5 −2 −1.5 −1 −0.5 0
Standardized log2FC
SEPTIN8
GSE16844
GSE74481
Summary
0 0.5 1 1.5 2 2.5 3 3.5 4 4.5
Standardized log2FC

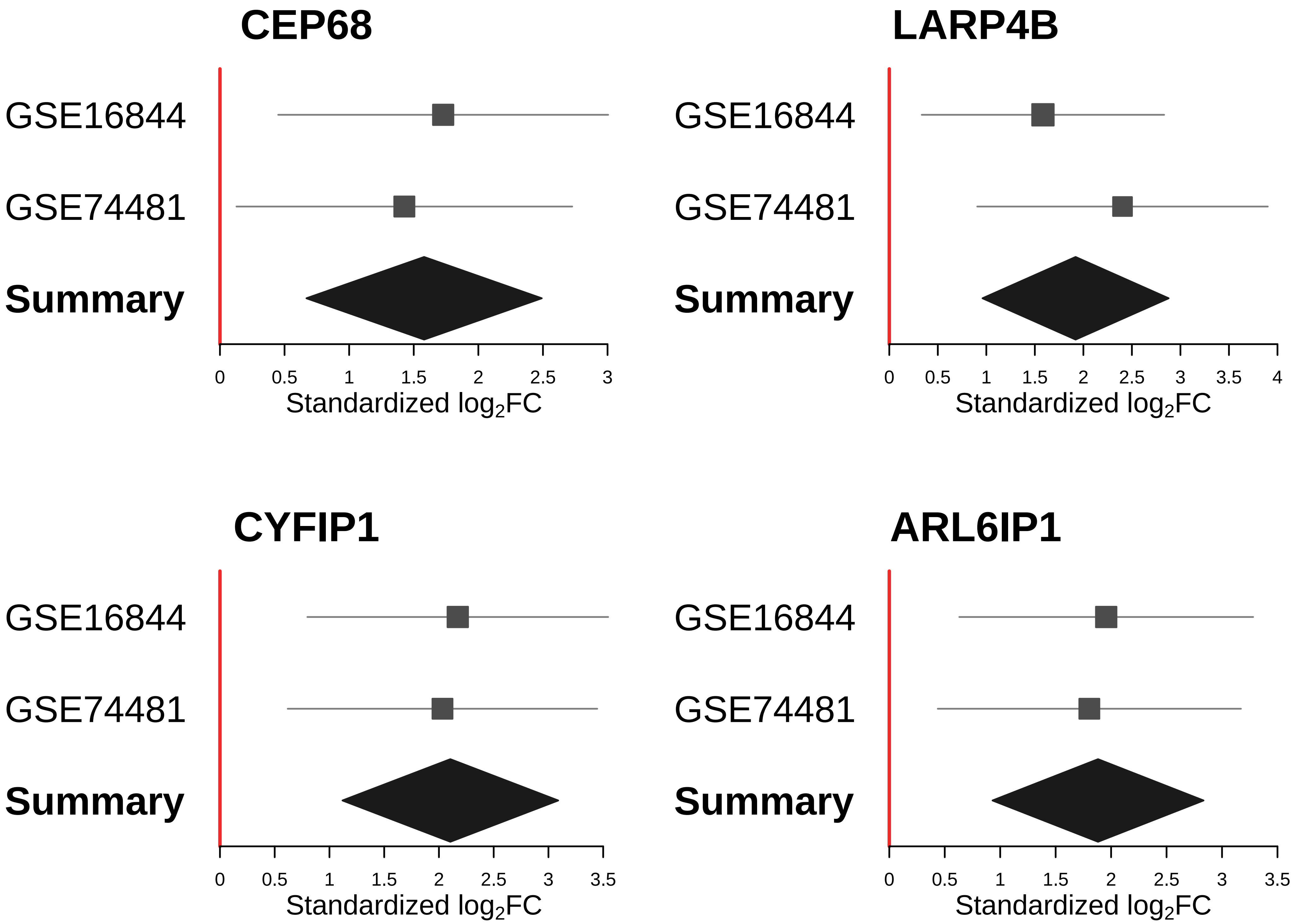
CEP68
GSE16844
GSE74481
Summary
0
0.5
1
1.5
2
2.5
3
Standardized log2FC
LARP4B
GSE16844
GSE74481
Summary
0
0.5
1
1.5
2
2.5
3
3.5
4
Standardized log2FC
CYFIP1
GSE16844
GSE74481
Summary
0
0.5
1
1.5
2
2.5
3
3.5
Standardized log2FC
ARL6IP1
GSE16844
GSE74481
Summary
0
0.5
1
1.5
2
2.5
3
3.5
Standardized log2FC

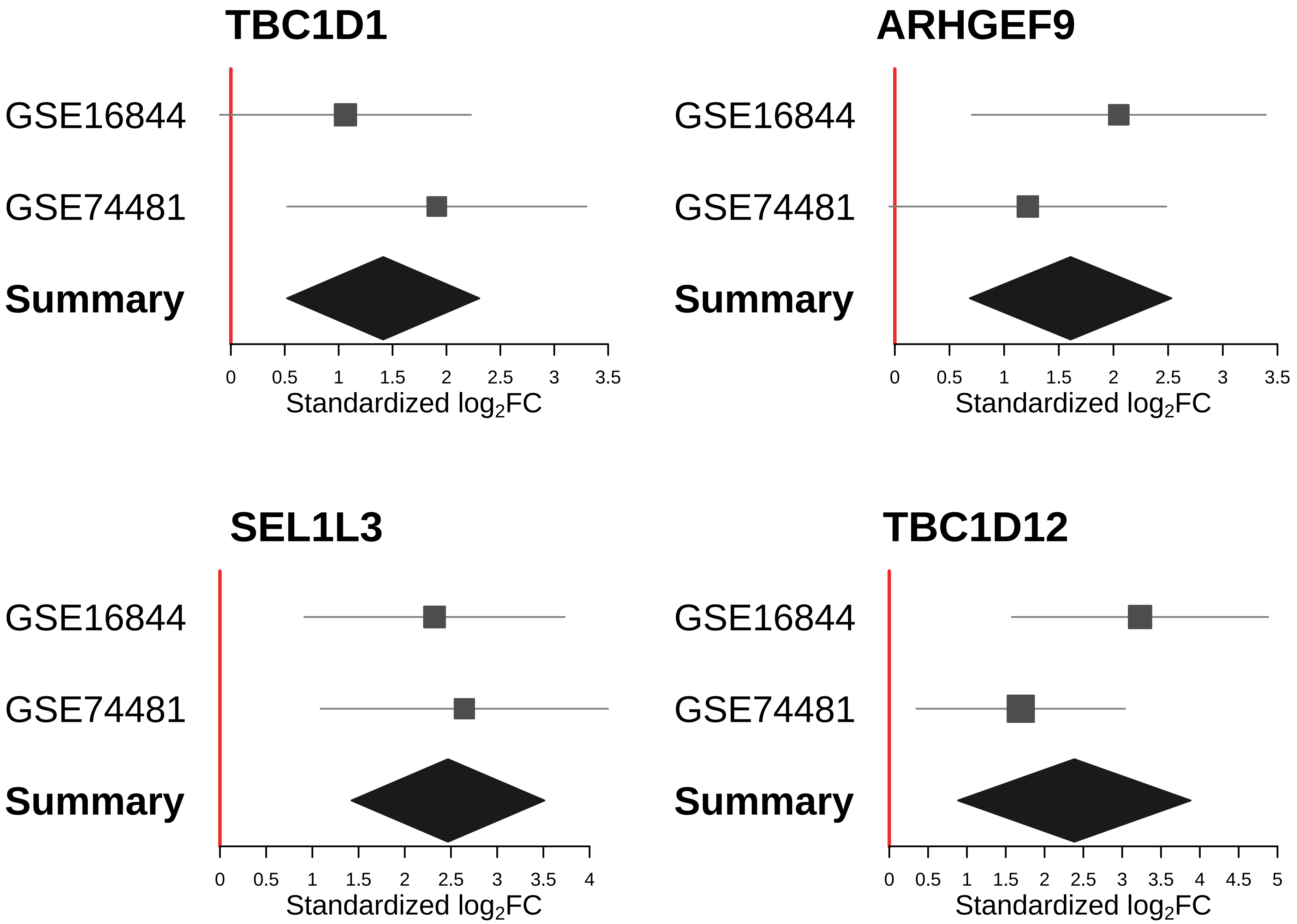
TBC1D1
GSE16844
GSE74481
Summary
0 0.5 1 1.5 2 2.5 3 3.5
Standardized log2FC
ARHGEF9
GSE16844
GSE74481
Summary
0 0.5 1 1.5 2 2.5 3 3.5
Standardized log2FC
SEL1L3
GSE16844
GSE74481
Summary
0 0.5 1 1.5 2 2.5 3 3.5 4
Standardized log2FC
TBC1D12
GSE16844
GSE74481
Summary
0 0.5 1 1.5 2 2.5 3 3.5 4 4.5 5
Standardized log2FC

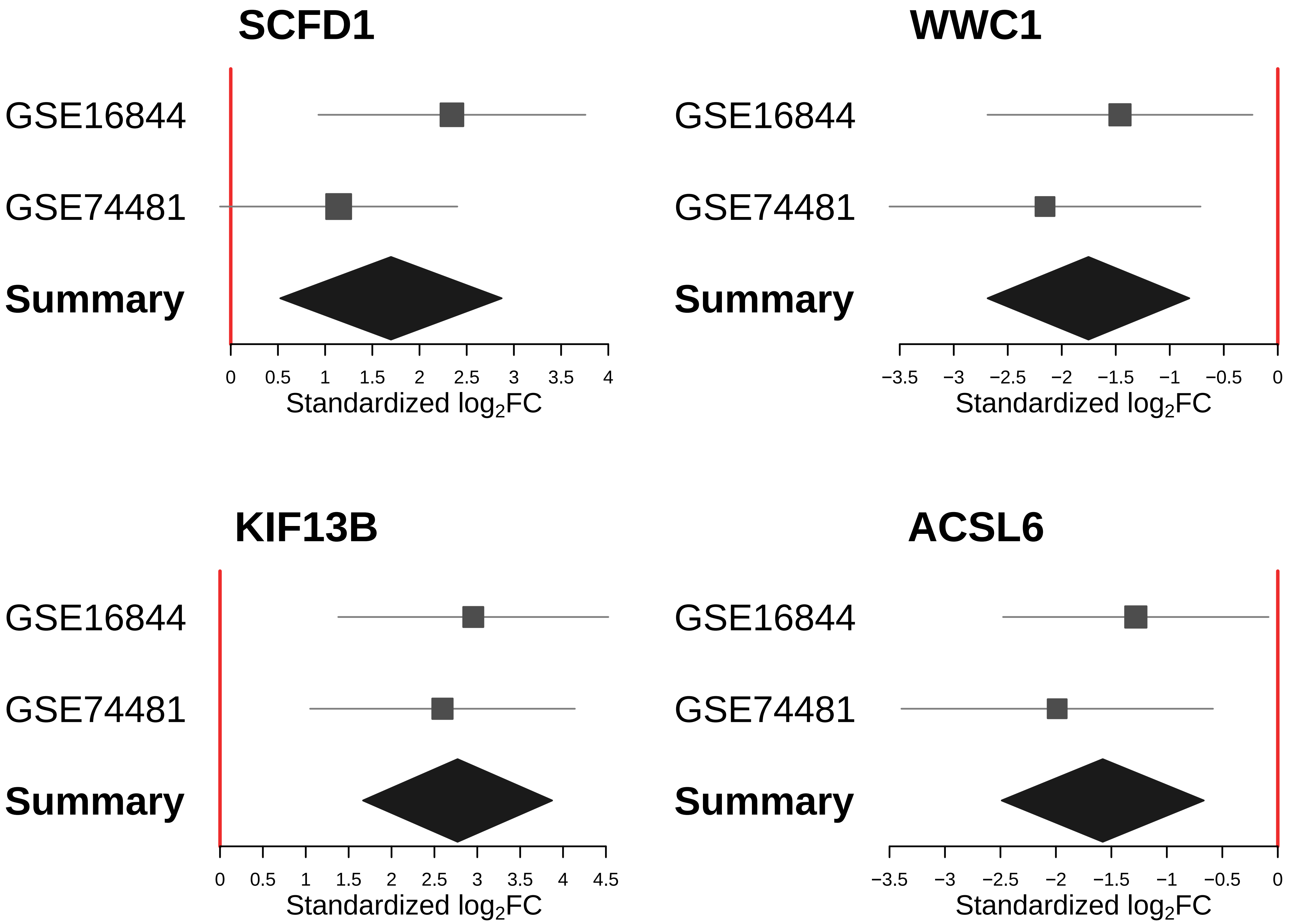

SCFD1
GSE16844
GSE74481
Summary
0 0.5 1 1.5 2 2.5 3 3.5 4
Standardized log2FC
WWC1
GSE16844
GSE74481
Summary
−3.5 −3 −2.5 −2 −1.5 −1 −0.5 0
Standardized log2FC
KIF13B
GSE16844
GSE74481
Summary
0 0.5 1 1.5 2 2.5 3 3.5 4 4.5
Standardized log2FC
ACSL6
GSE16844
GSE74481
Summary
−3.5 −3 −2.5 −2 −1.5 −1 −0.5 0
Standardized log2FC

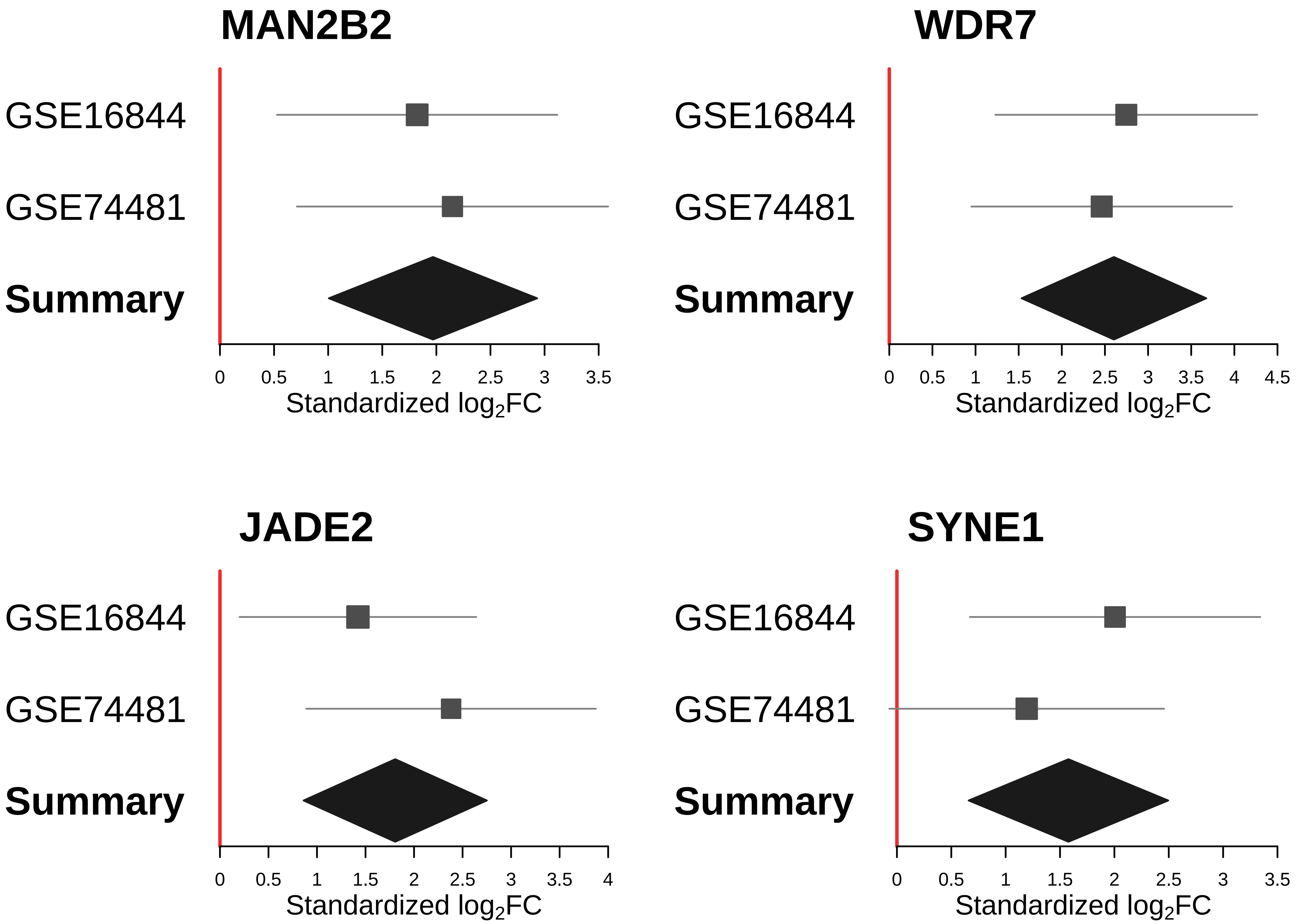
MAN2B2
GSE16844
GSE74481
Summary
0 0.5 1 1.5 2 2.5 3 3.5
Standardized $log_2FC$
WDR7
GSE16844
GSE74481
Summary
0 0.5 1 1.5 2 2.5 3 3.5 4 4.5
Standardized $log_2FC$
JADE2
GSE16844
GSE74481
Summary
0 0.5 1 1.5 2 2.5 3 3.5 4
Standardized $log_2FC$
SYNE1
GSE16844
GSE74481
Summary
0 0.5 1 1.5 2 2.5 3 3.5
Standardized $log_2FC$

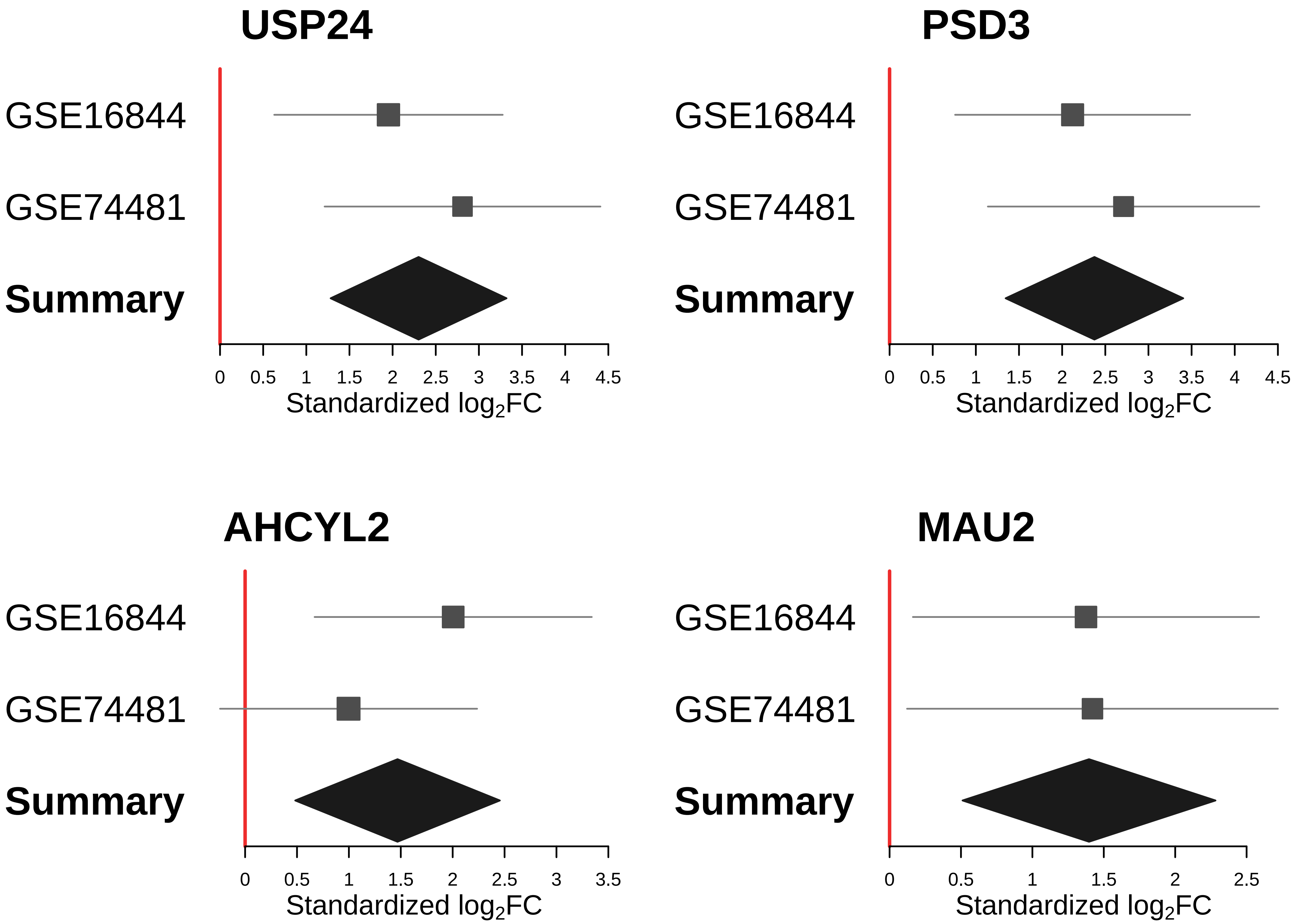

USP24
GSE16844
GSE74481
Summary
0
0.5
1
1.5
2
2.5
3
3.5
4
4.5
Standardized $log_2FC$
PSD3
GSE16844
GSE74481
Summary
0
0.5
1
1.5
2
2.5
3
3.5
4
4.5
Standardized $log_2FC$
AHCYL2
GSE16844
GSE74481
Summary
0
0.5
1
1.5
2
2.5
3
3.5
Standardized $log_2FC$
MAU2
GSE16844
GSE74481
Summary
0
0.5
1
1.5
2
2.5
Standardized $log_2FC$

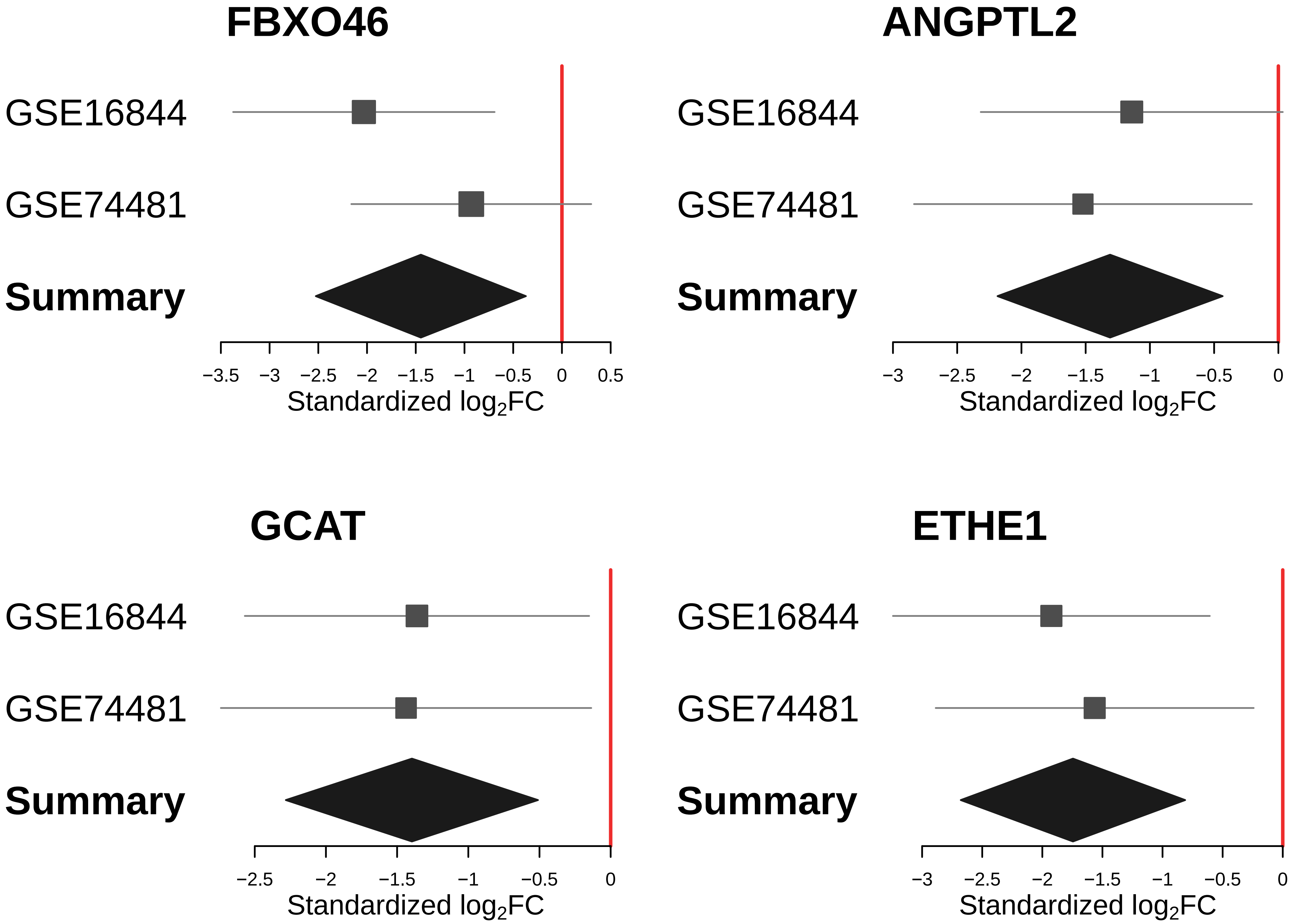
FBXO46
GSE16844
GSE74481
Summary
−3.5 −3 −2.5 −2 −1.5 −1 −0.5 0 0.5
Standardized $log_2FC$
ANGPTL2
GSE16844
GSE74481
Summary
−3 −2.5 −2 −1.5 −1 −0.5 0
Standardized $log_2FC$
GCAT
GSE16844
GSE74481
Summary
−2.5 −2 −1.5 −1 −0.5 0
Standardized $log_2FC$
ETHE1
GSE16844
GSE74481
Summary
−3 −2.5 −2 −1.5 −1 −0.5 0
Standardized $log_2FC$

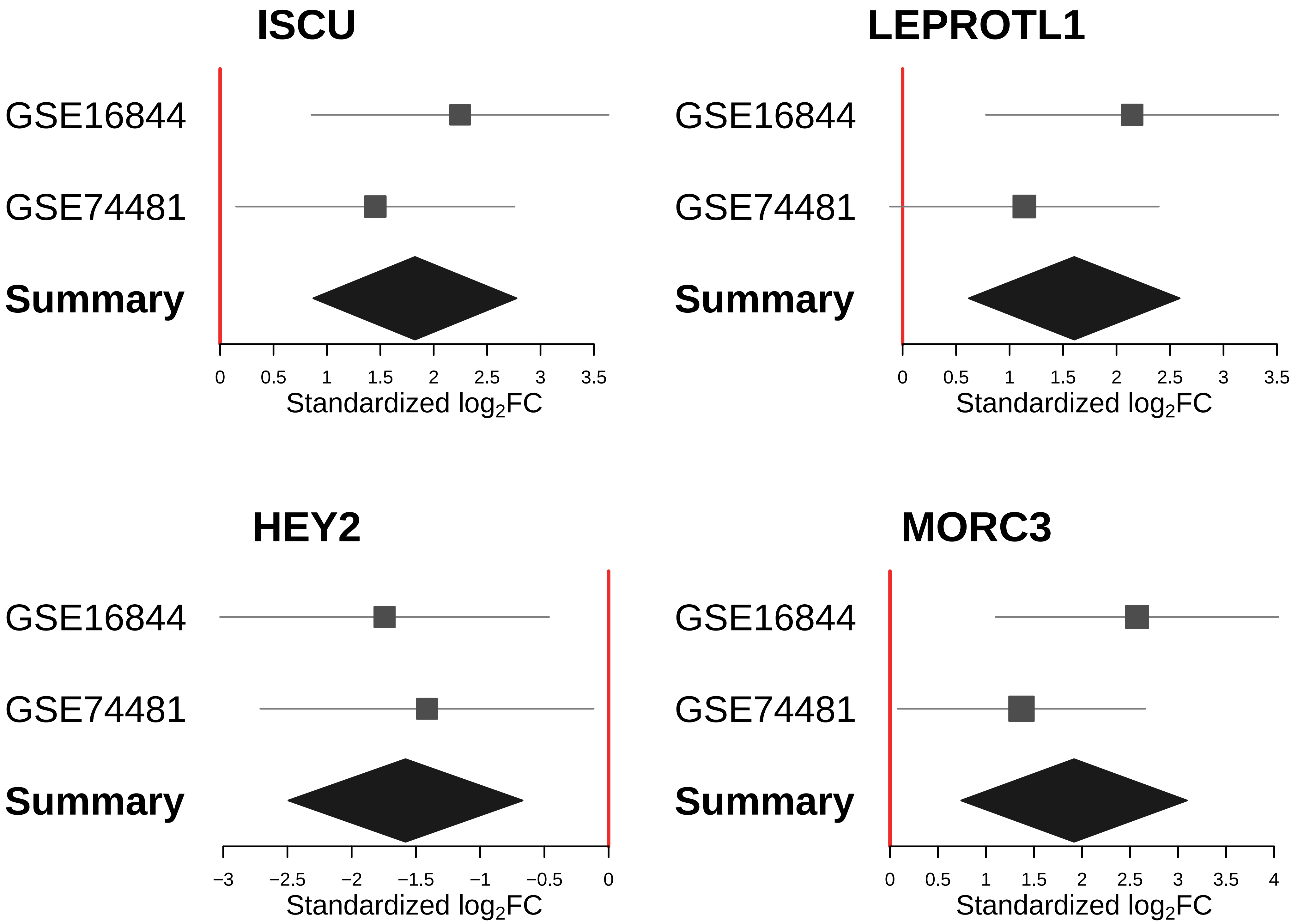
ISCU
GSE16844
GSE74481
Summary
0 0.5 1 1.5 2 2.5 3 3.5
Standardized log2FC
LEPROTL1
GSE16844
GSE74481
Summary
0 0.5 1 1.5 2 2.5 3 3.5
Standardized log2FC
HEY2
GSE16844
GSE74481
Summary
−3 −2.5 −2 −1.5 −1 −0.5 0
Standardized log2FC
MORC3
GSE16844
GSE74481
Summary
0 0.5 1 1.5 2 2.5 3 3.5 4
Standardized log2FC

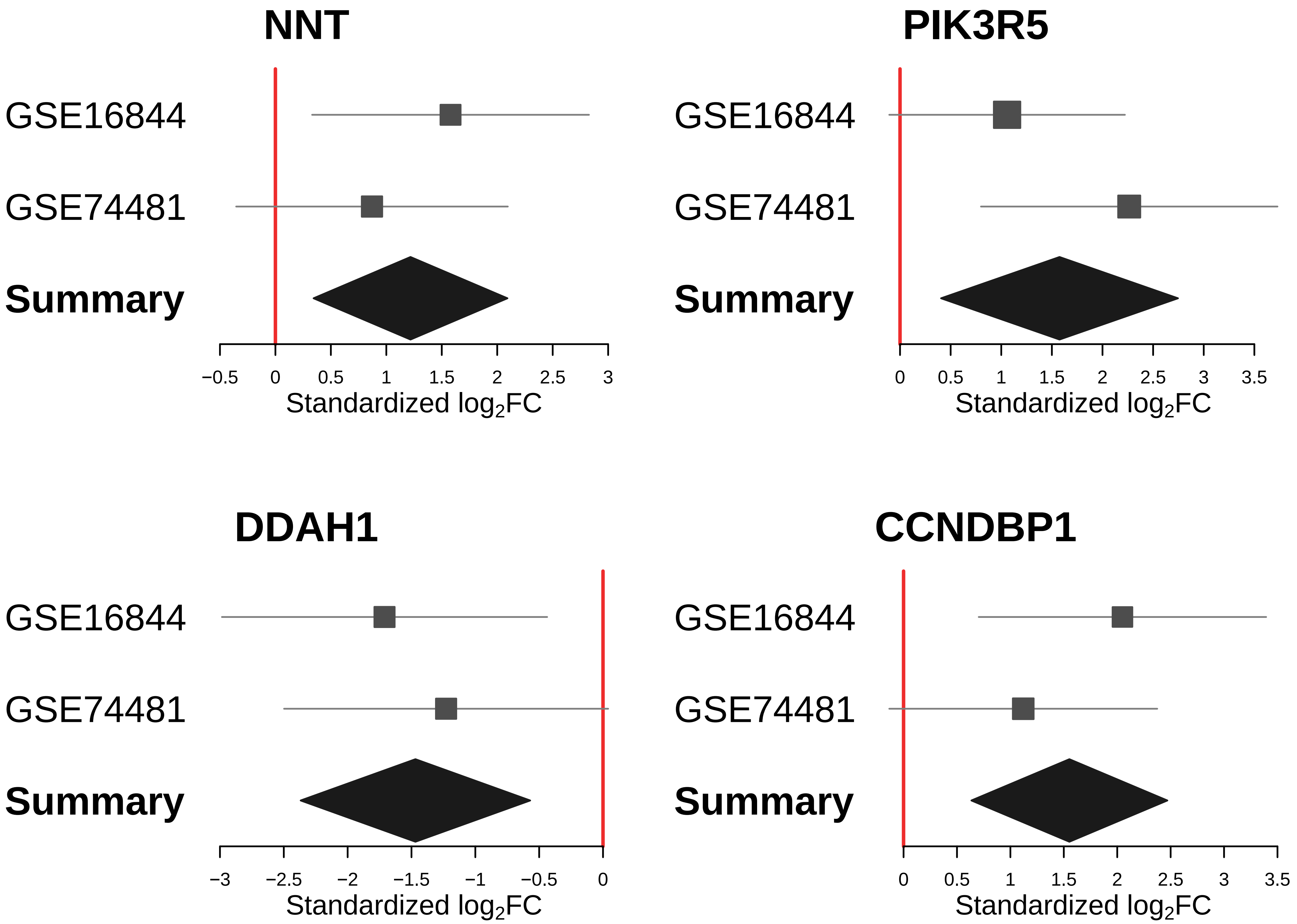
NNT
GSE16844
GSE74481
Summary
−0.5 0 0.5 1 1.5 2 2.5 3
Standardized log2FC
PIK3R5
GSE16844
GSE74481
Summary
0 0.5 1 1.5 2 2.5 3 3.5
Standardized log2FC
DDAH1
GSE16844
GSE74481
Summary
−3 −2.5 −2 −1.5 −1 −0.5 0
Standardized log2FC
CCNDBP1
GSE16844
GSE74481
Summary
0 0.5 1 1.5 2 2.5 3 3.5
Standardized log2FC

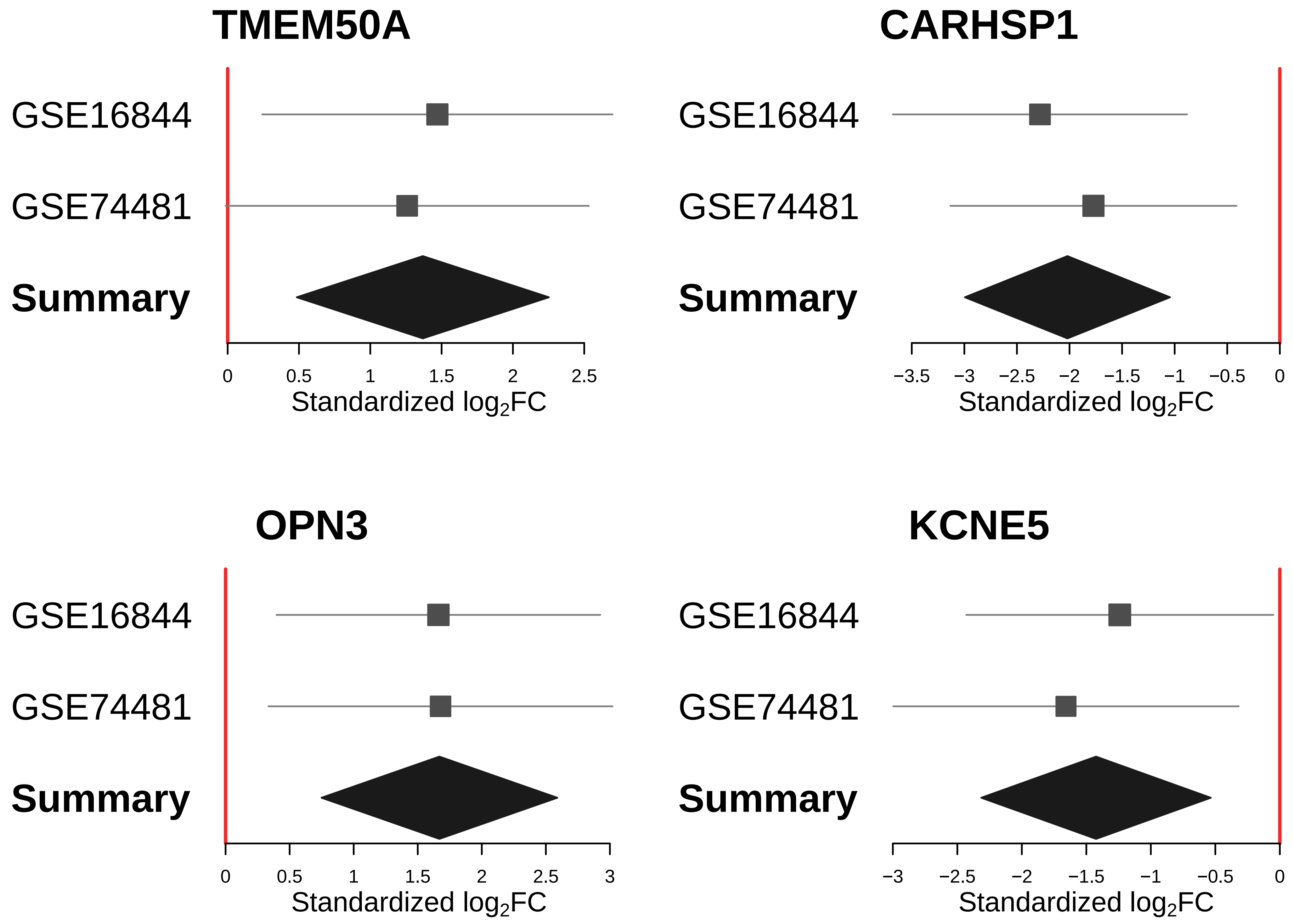

TMEM50A
GSE16844
GSE74481
Summary
0
0.5
1
1.5
2
2.5
Standardized log2FC
CARHSP1
GSE16844
GSE74481
Summary
−3.5
−3
−2.5
−2
−1.5
−1
−0.5
0
Standardized log2FC
OPN3
GSE16844
GSE74481
Summary
0
0.5
1
1.5
2
2.5
3
Standardized log2FC
KCNE5
GSE16844
GSE74481
Summary
−3
−2.5
−2
−1.5
−1
−0.5
0
Standardized log2FC

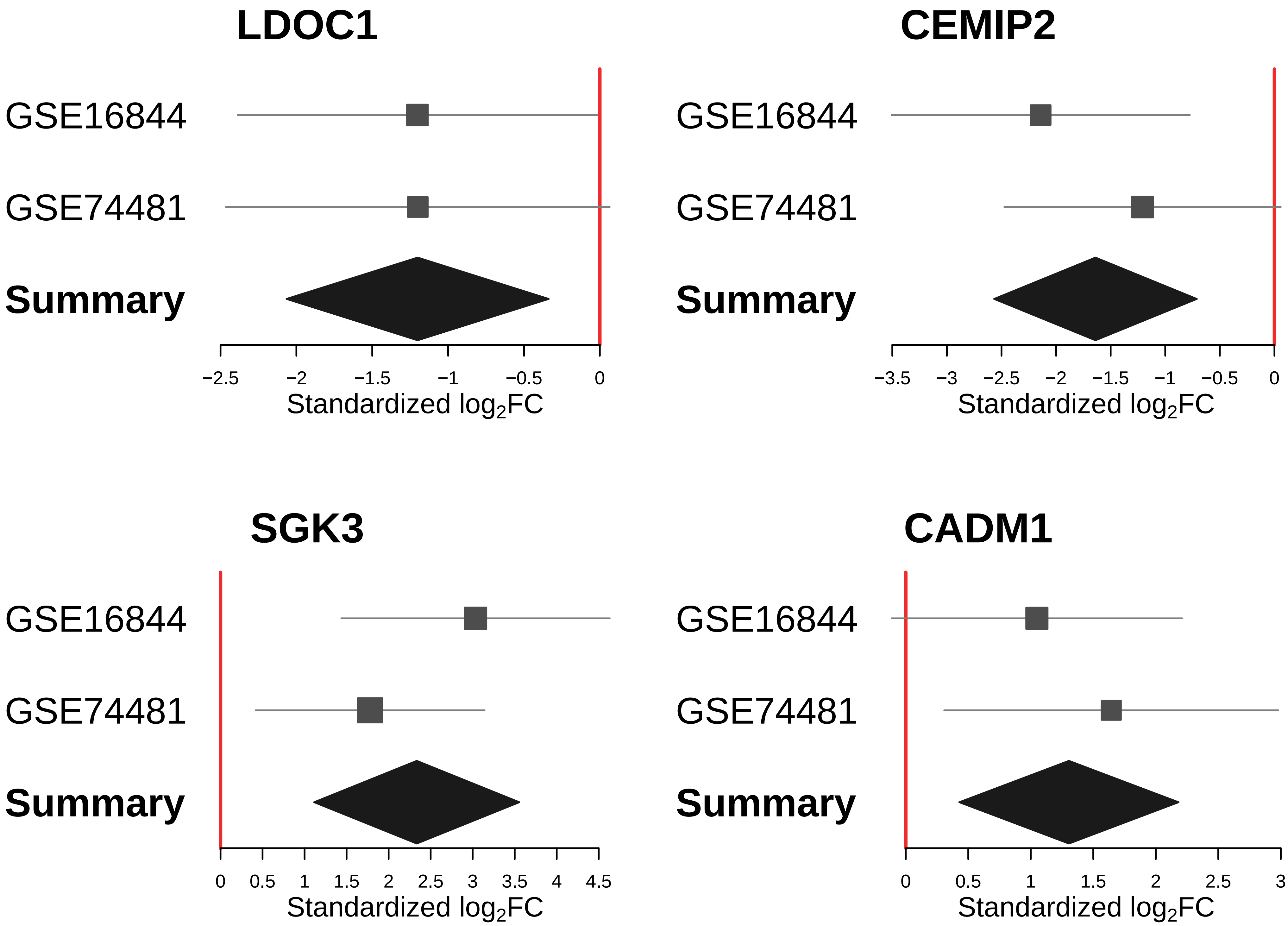
LDOC1
GSE16844
GSE74481
Summary
−2.5 −2 −1.5 −1 −0.5 0
Standardized log2FC
CEMIP2
GSE16844
GSE74481
Summary
−3.5 −3 −2.5 −2 −1.5 −1 −0.5 0
Standardized log2FC
SGK3
GSE16844
GSE74481
Summary
0 0.5 1 1.5 2 2.5 3 3.5 4 4.5
Standardized log2FC
CADM1
GSE16844
GSE74481
Summary
0 0.5 1 1.5 2 2.5 3
Standardized log2FC

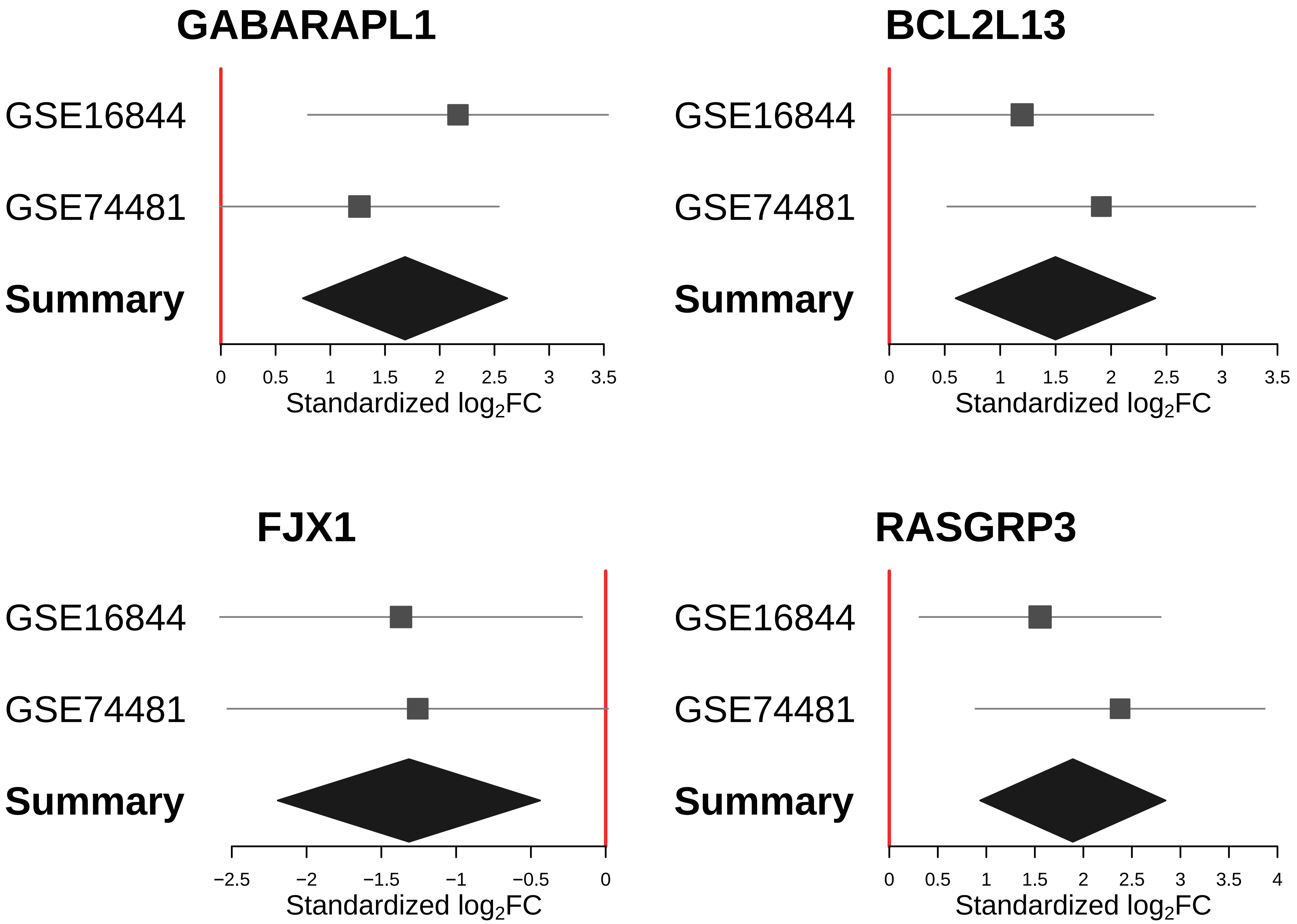
GABARAPL1
GSE16844
GSE74481
Summary
0 0.5 1 1.5 2 2.5 3 3.5
Standardized log2FC
BCL2L13
GSE16844
GSE74481
Summary
0 0.5 1 1.5 2 2.5 3 3.5
Standardized log2FC
FJX1
GSE16844
GSE74481
Summary
−2.5 −2 −1.5 −1 −0.5 0
Standardized log2FC
RASGRP3
GSE16844
GSE74481
Summary
0 0.5 1 1.5 2 2.5 3 3.5 4
Standardized log2FC

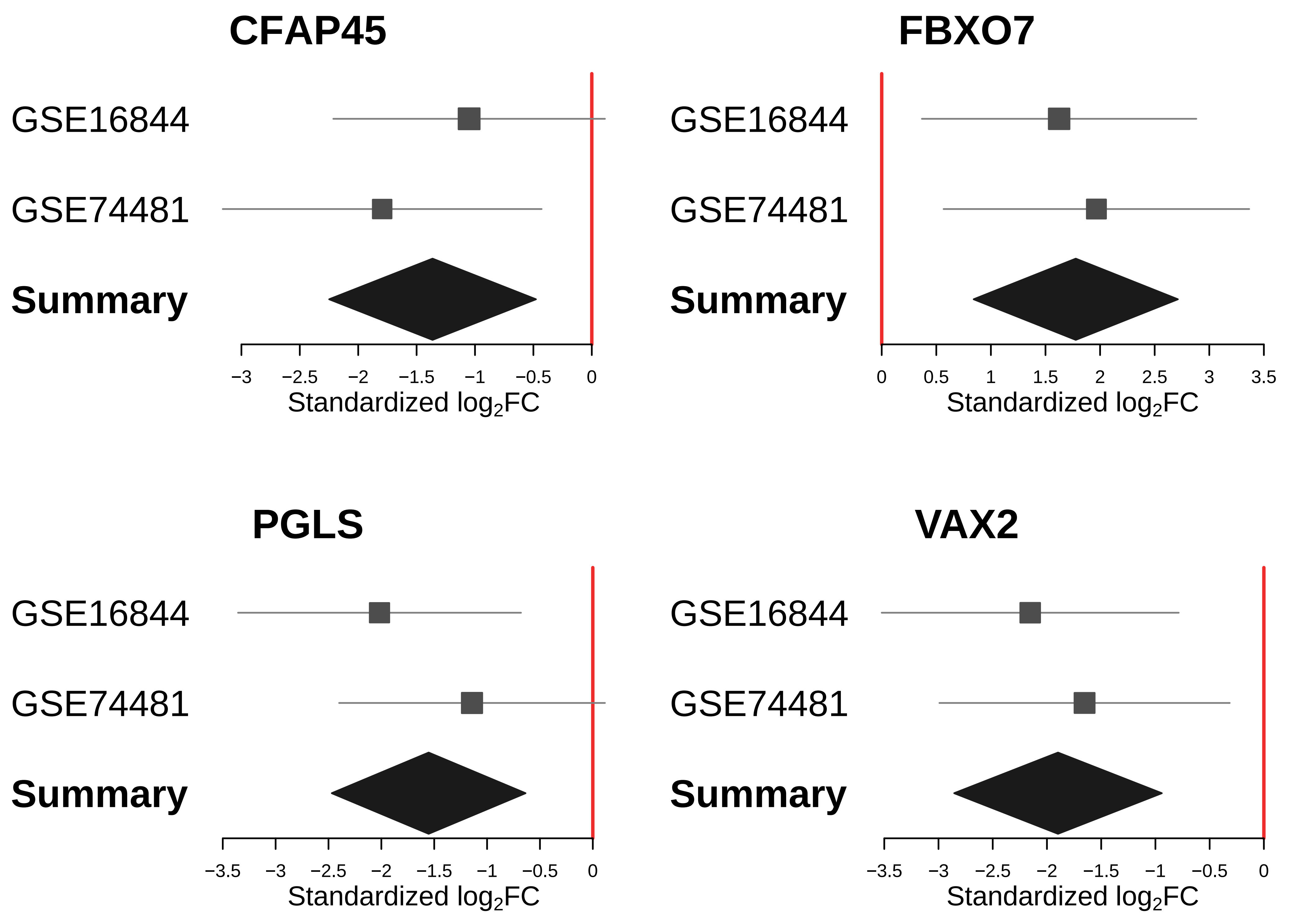
CFAP45
GSE16844
GSE74481
Summary
−3
−2.5
−2
−1.5
−1
−0.5
0
Standardized log2FC
FBXO7
GSE16844
GSE74481
Summary
0
0.5
1
1.5
2
2.5
3
3.5
Standardized log2FC
PGLS
GSE16844
GSE74481
Summary
−3.5
−3
−2.5
−2
−1.5
−1
−0.5
0
Standardized log2FC
VAX2
GSE16844
GSE74481
Summary
−3.5
−3
−2.5
−2
−1.5
−1
−0.5
0
Standardized log2FC

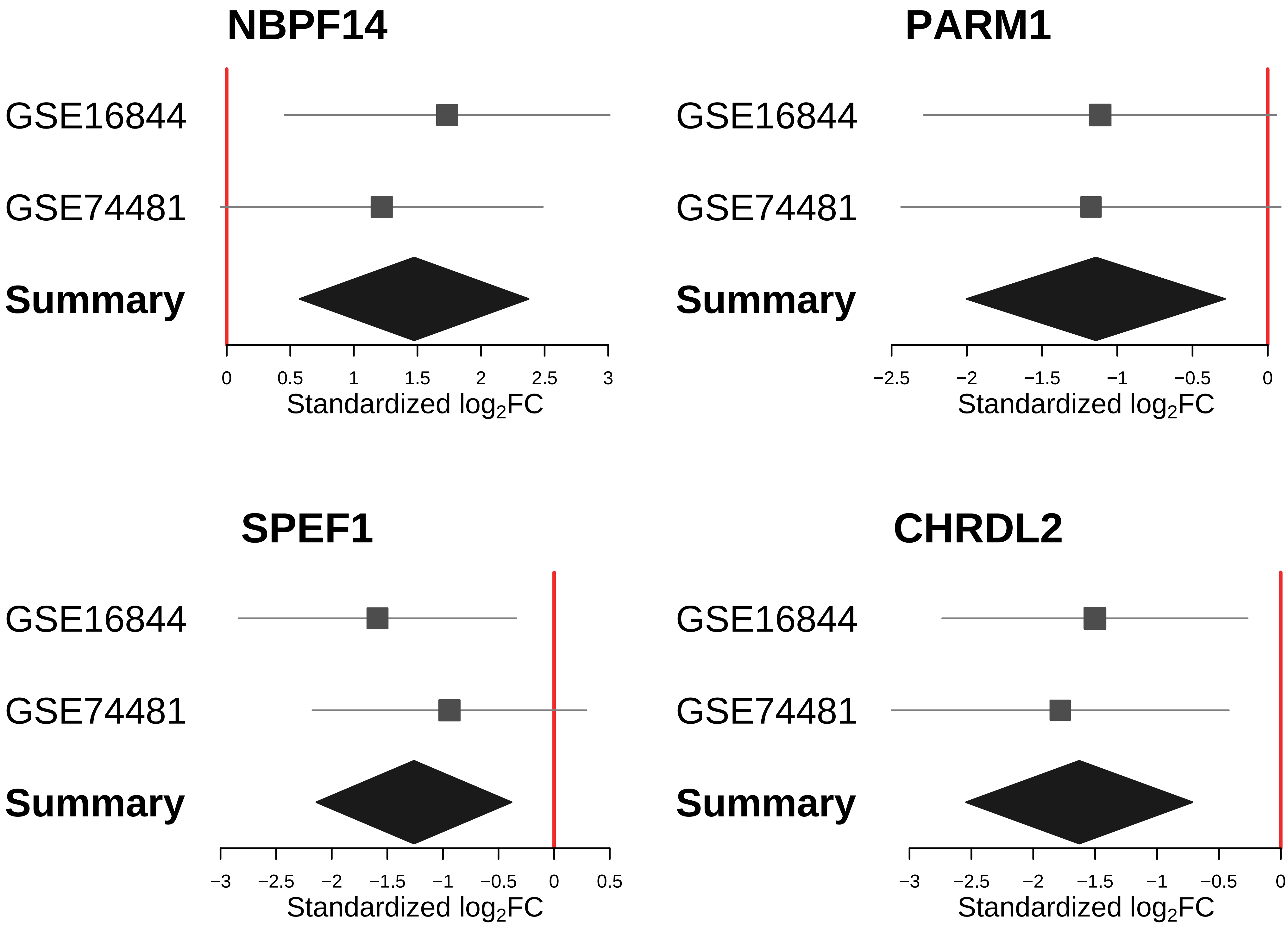
NBPF14
GSE16844
GSE74481
Summary
0
0.5
1
1.5
2
2.5
3
Standardized $log_2FC$
PARM1
GSE16844
GSE74481
Summary
−2.5
−2
−1.5
−1
−0.5
0
Standardized $log_2FC$
SPEF1
GSE16844
GSE74481
Summary
−3
−2.5
−2
−1.5
−1
−0.5
0
0.5
Standardized $log_2FC$
CHRDL2
GSE16844
GSE74481
Summary
−3
−2.5
−2
−1.5
−1
−0.5
0
Standardized $log_2FC$

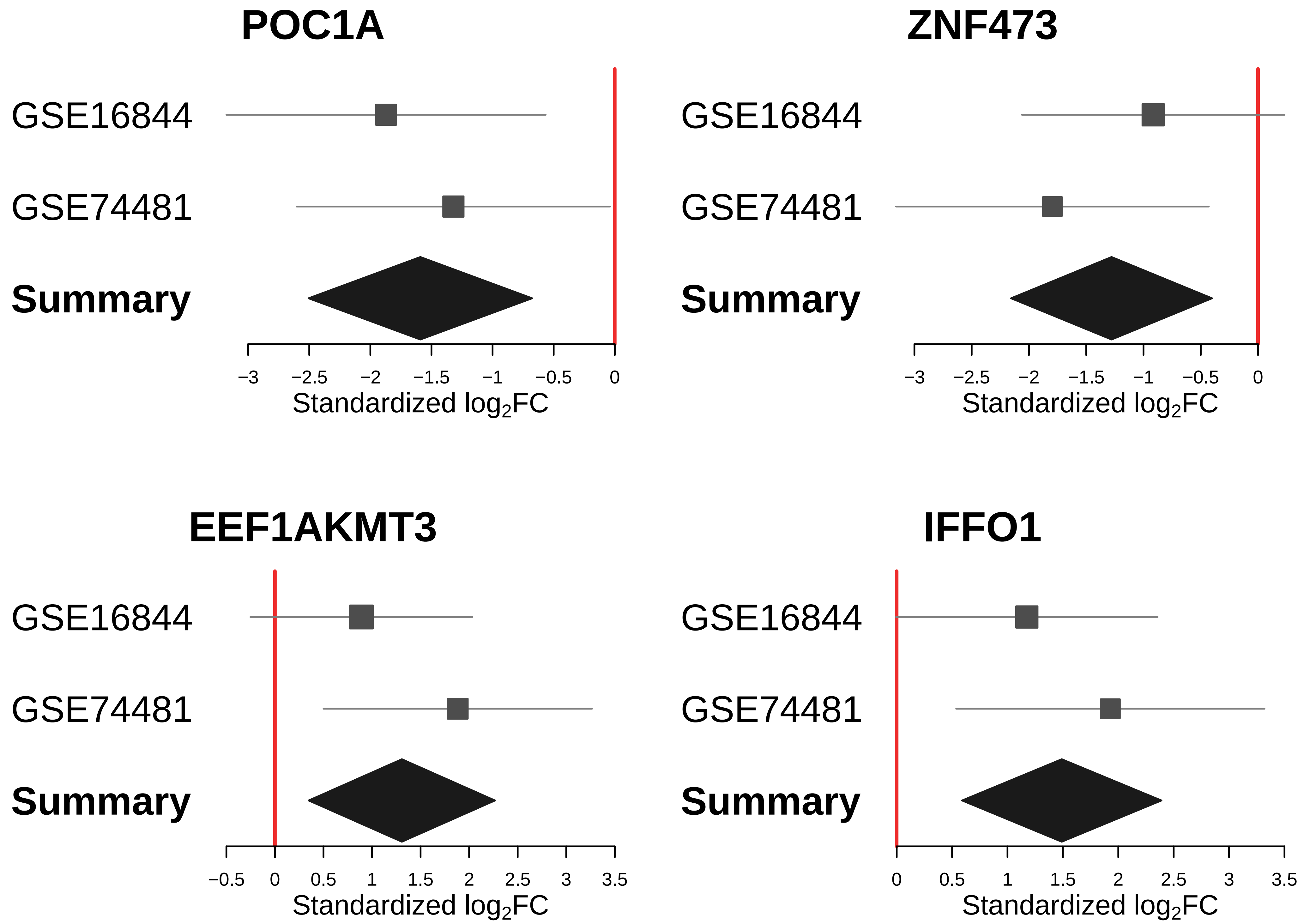
POC1A
GSE16844
GSE74481
Summary
−3 −2.5 −2 −1.5 −1 −0.5 0
Standardized $log_2FC$
ZNF473
GSE16844
GSE74481
Summary
−3 −2.5 −2 −1.5 −1 −0.5 0
Standardized $log_2FC$
EEF1AKMT3
GSE16844
GSE74481
Summary
−0.5 0 0.5 1 1.5 2 2.5 3 3.5
Standardized $log_2FC$
IFFO1
GSE16844
GSE74481
Summary
0 0.5 1 1.5 2 2.5 3 3.5
Standardized $log_2FC$

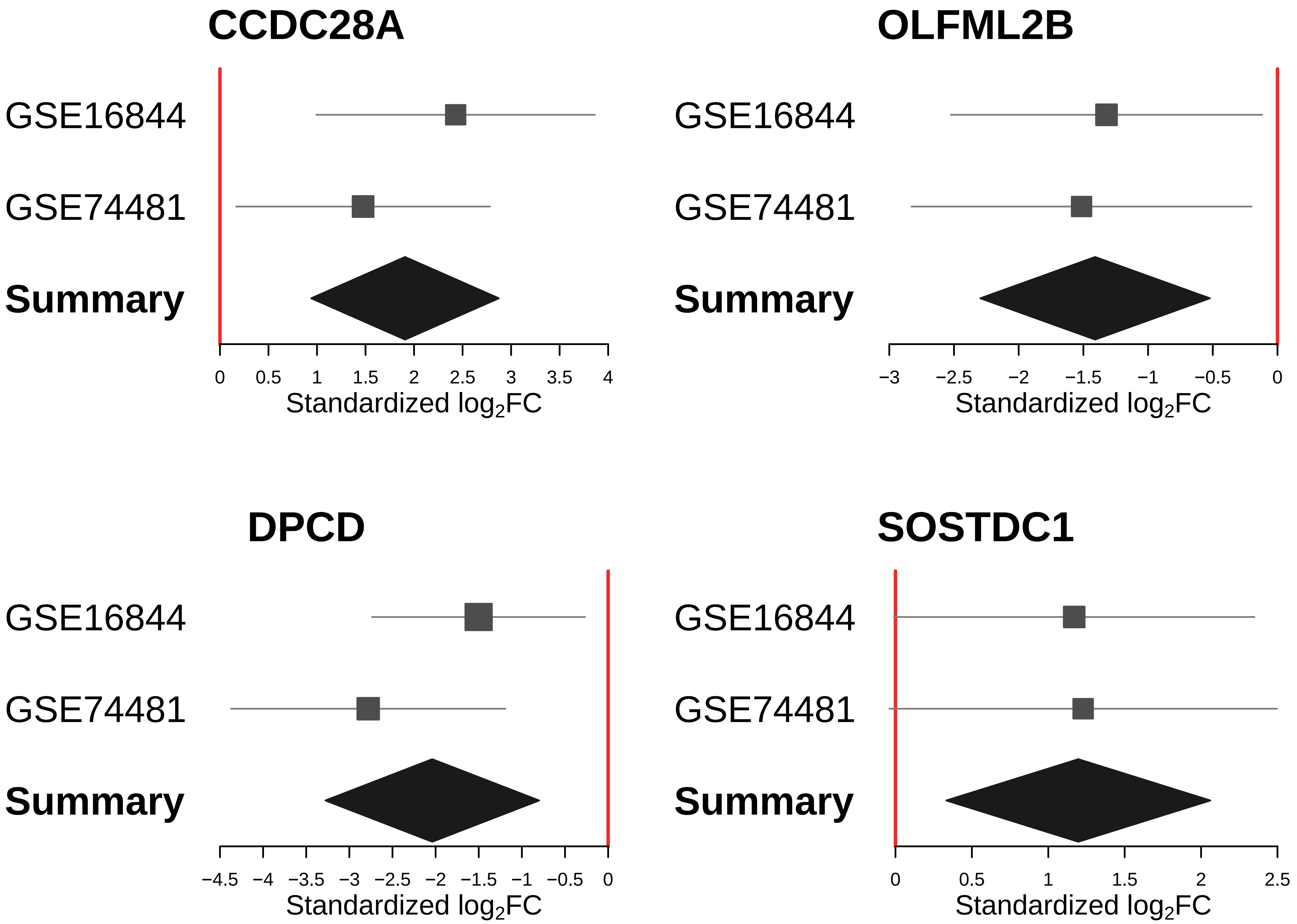
CCDC28A
GSE16844
GSE74481
Summary
0 0.5 1 1.5 2 2.5 3 3.5 4
Standardized $log_2FC$
OLFML2B
GSE16844
GSE74481
Summary
−3 −2.5 −2 −1.5 −1 −0.5 0
Standardized $log_2FC$
DPCD
GSE16844
GSE74481
Summary
−4.5 −4 −3.5 −3 −2.5 −2 −1.5 −1 −0.5 0
Standardized $log_2FC$
SOSTDC1
GSE16844
GSE74481
Summary
0 0.5 1 1.5 2 2.5
Standardized $log_2FC$

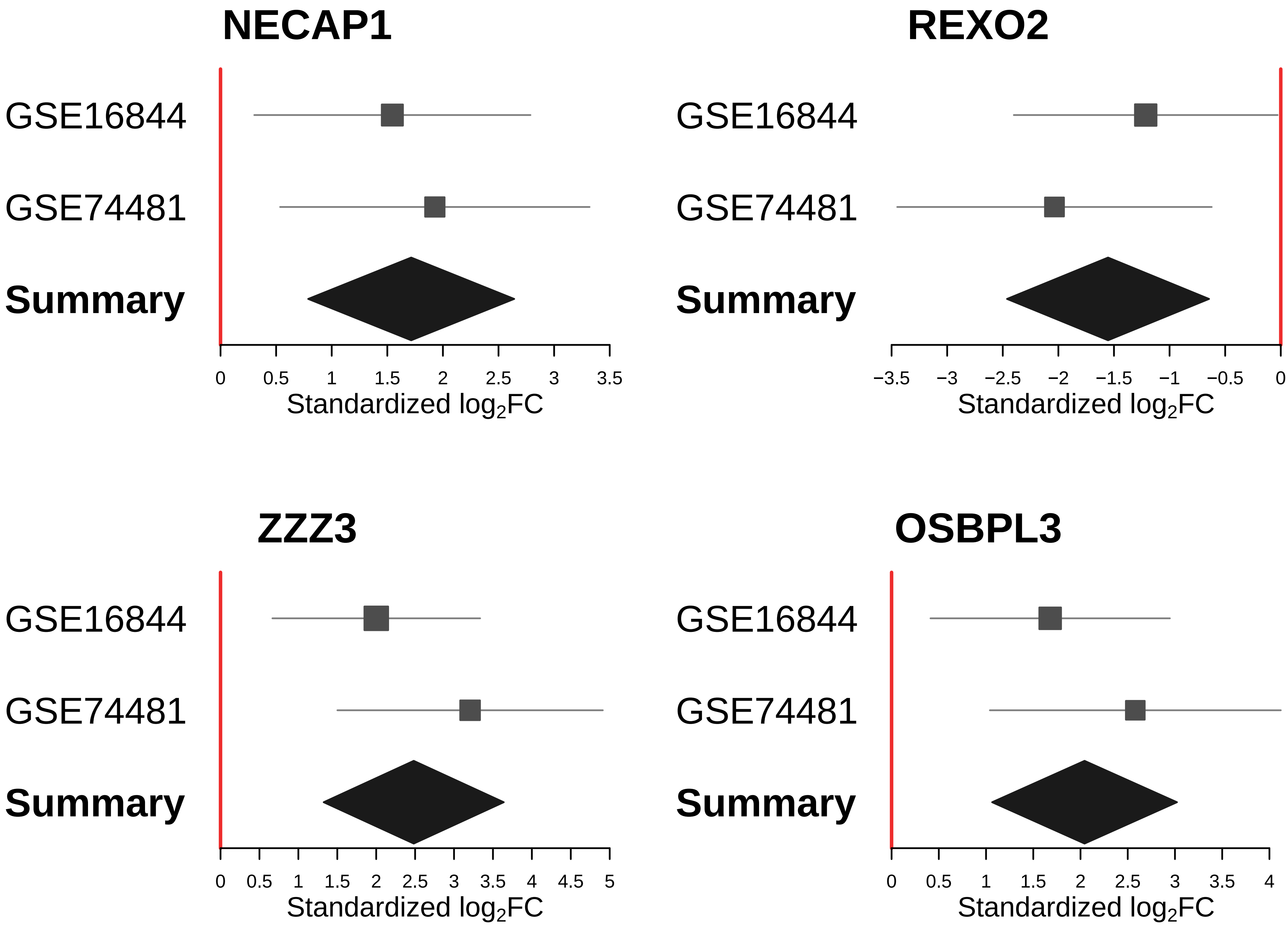
NECAP1
REXO2
ZZZ3
OSBPL3
GSE16844
GSE74481
Summary
Standardized $\log_2$FC
0 0.5 1 1.5 2 2.5 3 3.5
−3.5 −3 −2.5 −2 −1.5 −1 −0.5 0
0 0.5 1 1.5 2 2.5 3 3.5 4 4.5 5
0 0.5 1 1.5 2 2.5 3 3.5 4

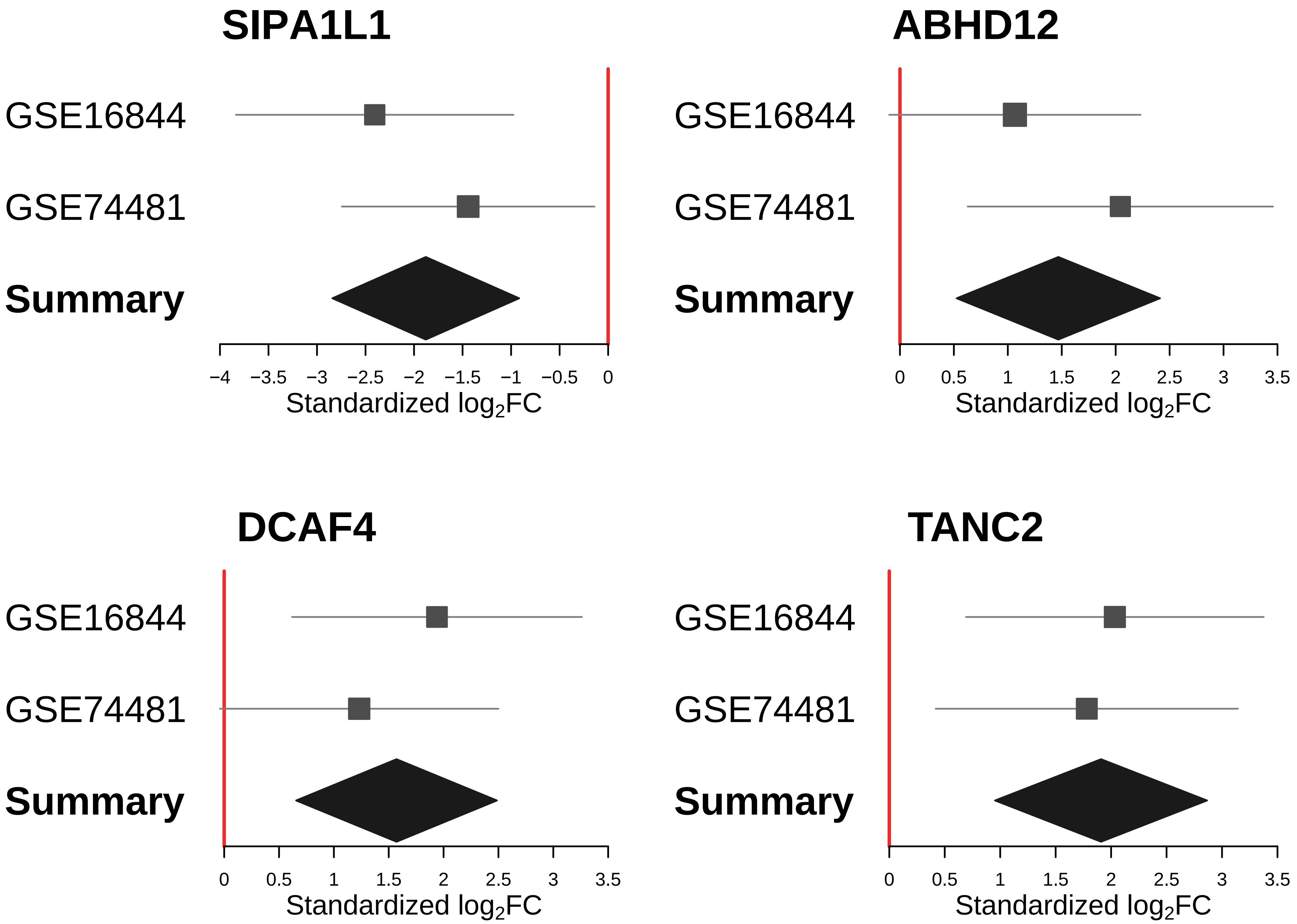

SIPA1L1
GSE16844
GSE74481
Summary
−4 −3.5 −3 −2.5 −2 −1.5 −1 −0.5 0
Standardized $log_2FC$
ABHD12
GSE16844
GSE74481
Summary
0 0.5 1 1.5 2 2.5 3 3.5
Standardized $log_2FC$
DCAF4
GSE16844
GSE74481
Summary
0 0.5 1 1.5 2 2.5 3 3.5
Standardized $log_2FC$
TANC2
GSE16844
GSE74481
Summary
0 0.5 1 1.5 2 2.5 3 3.5
Standardized $log_2FC$

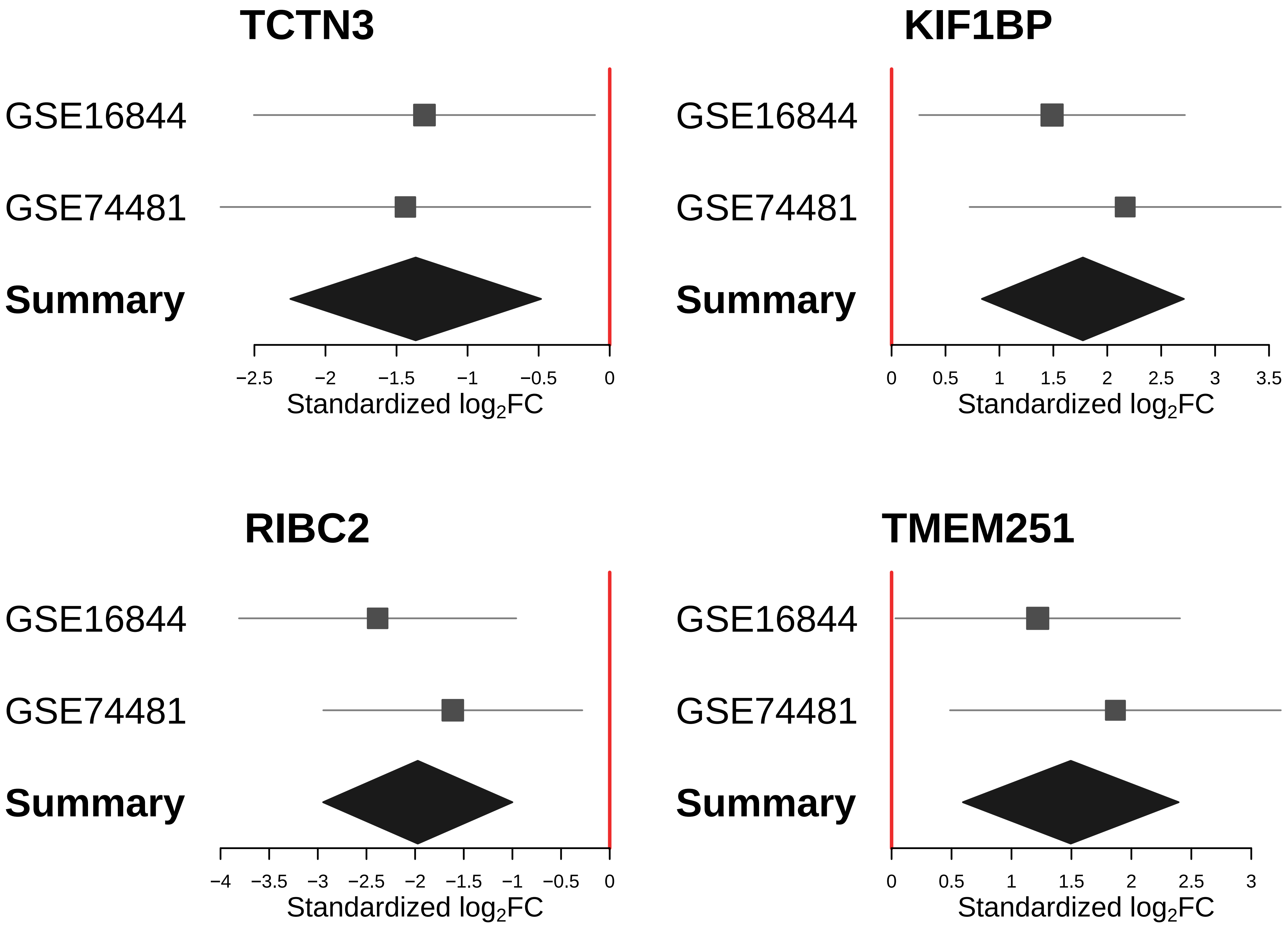
TCTN3
GSE16844
GSE74481
Summary
−2.5 −2 −1.5 −1 −0.5 0
Standardized log2FC
KIF1BP
GSE16844
GSE74481
Summary
0 0.5 1 1.5 2 2.5 3 3.5
Standardized log2FC
RIBC2
GSE16844
GSE74481
Summary
−4 −3.5 −3 −2.5 −2 −1.5 −1 −0.5 0
Standardized log2FC
TMEM251
GSE16844
GSE74481
Summary
0 0.5 1 1.5 2 2.5 3
Standardized log2FC

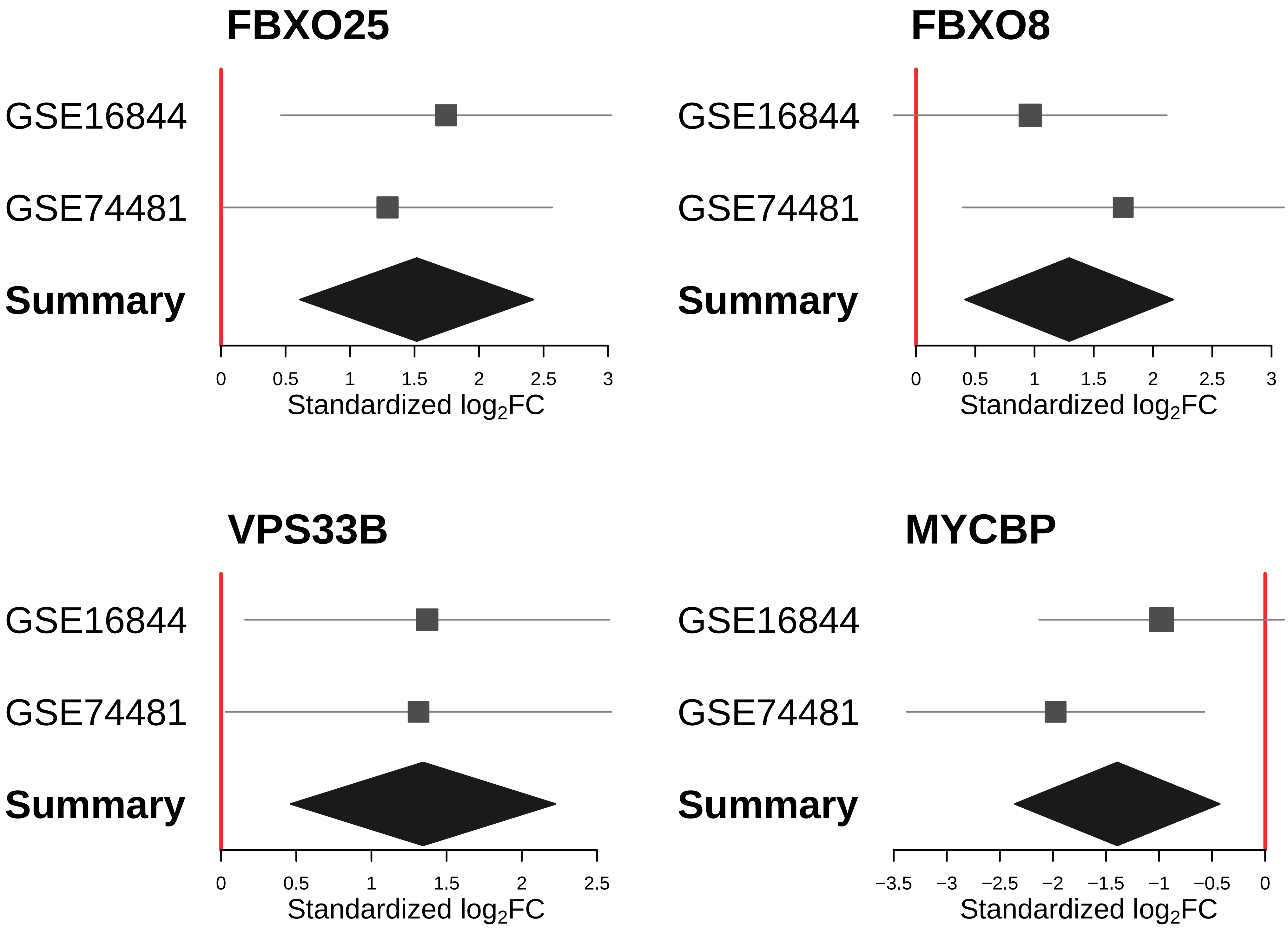
FBXO25
GSE16844
GSE74481
Summary
0
0.5
1
1.5
2
2.5
3
Standardized log2FC
FBXO8
GSE16844
GSE74481
Summary
0
0.5
1
1.5
2
2.5
3
Standardized log2FC
VPS33B
GSE16844
GSE74481
Summary
0
0.5
1
1.5
2
2.5
Standardized log2FC
MYCBP
GSE16844
GSE74481
Summary
−3.5
−3
−2.5
−2
−1.5
−1
−0.5
0
Standardized log2FC

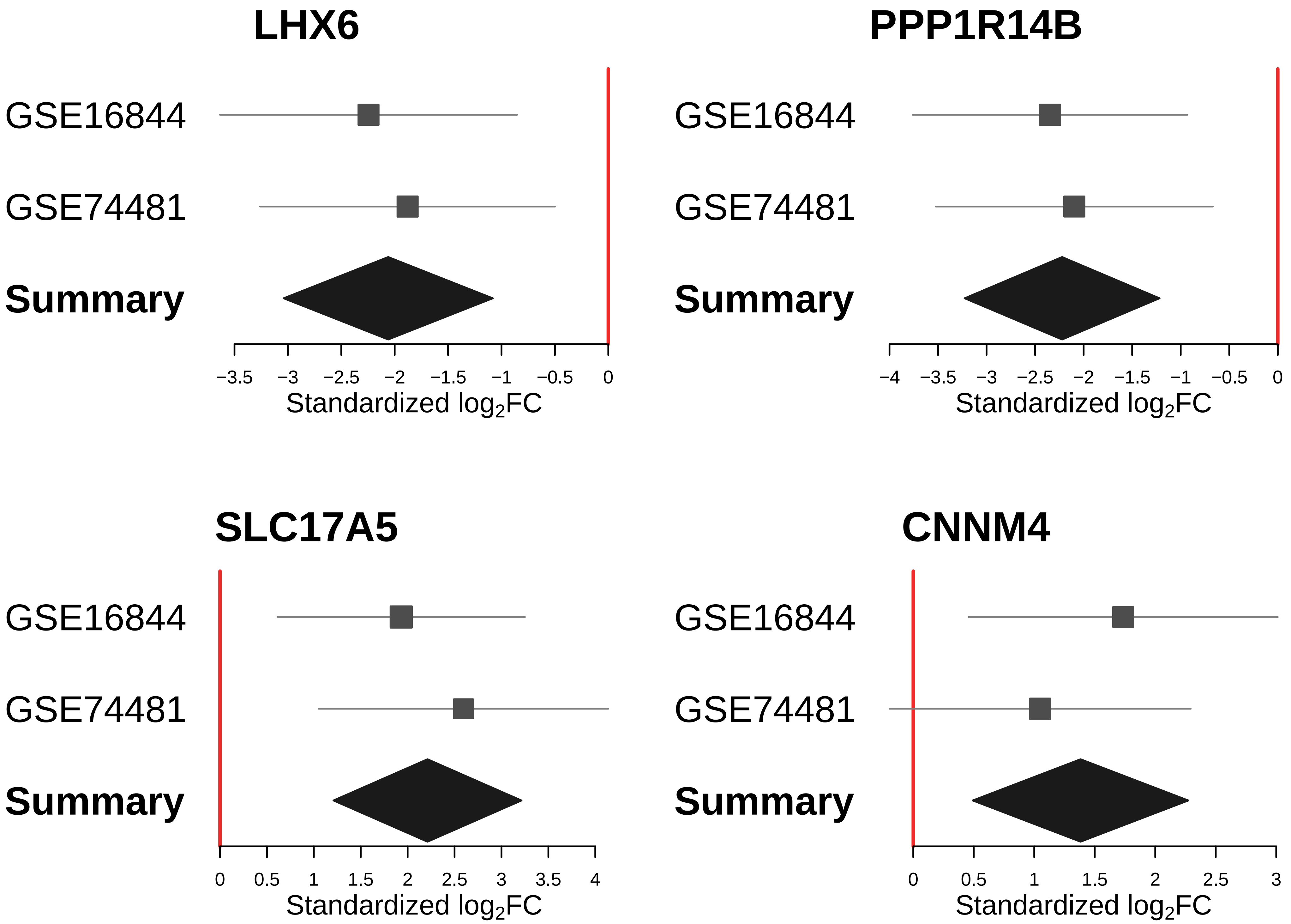
LHX6
GSE16844
GSE74481
Summary
−3.5 −3 −2.5 −2 −1.5 −1 −0.5 0
Standardized log2FC
PPP1R14B
GSE16844
GSE74481
Summary
−4 −3.5 −3 −2.5 −2 −1.5 −1 −0.5 0
Standardized log2FC
SLC17A5
GSE16844
GSE74481
Summary
0 0.5 1 1.5 2 2.5 3 3.5 4
Standardized log2FC
CNNM4
GSE16844
GSE74481
Summary
0 0.5 1 1.5 2 2.5 3
Standardized log2FC

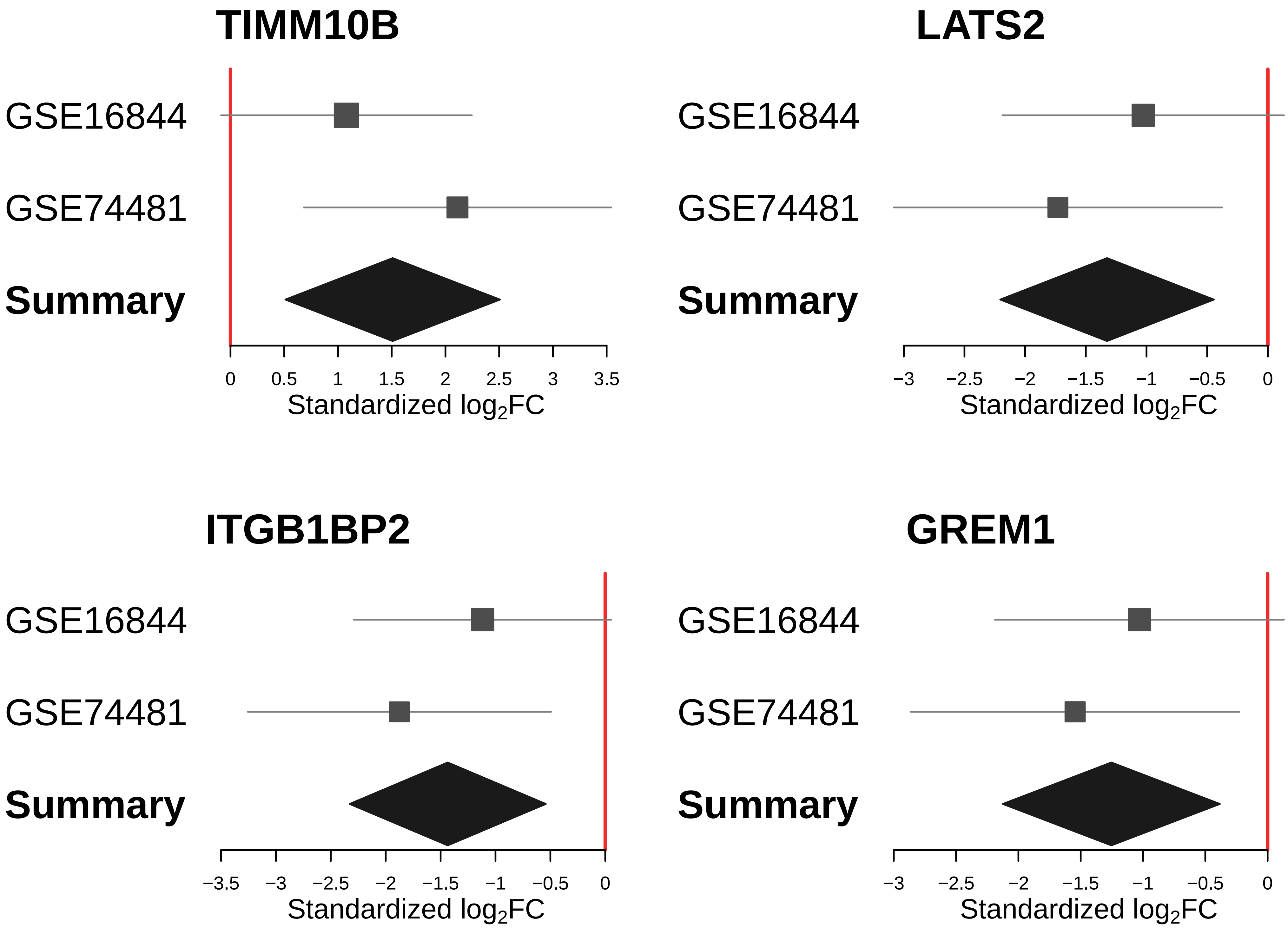
TIMM10B
GSE16844
GSE74481
Summary
0
0.5
1
1.5
2
2.5
3
3.5
Standardized log2FC
LATS2
GSE16844
GSE74481
Summary
−3
−2.5
−2
−1.5
−1
−0.5
0
Standardized log2FC
ITGB1BP2
GSE16844
GSE74481
Summary
−3.5
−3
−2.5
−2
−1.5
−1
−0.5
0
Standardized log2FC
GREM1
GSE16844
GSE74481
Summary
−3
−2.5
−2
−1.5
−1
−0.5
0
Standardized log2FC

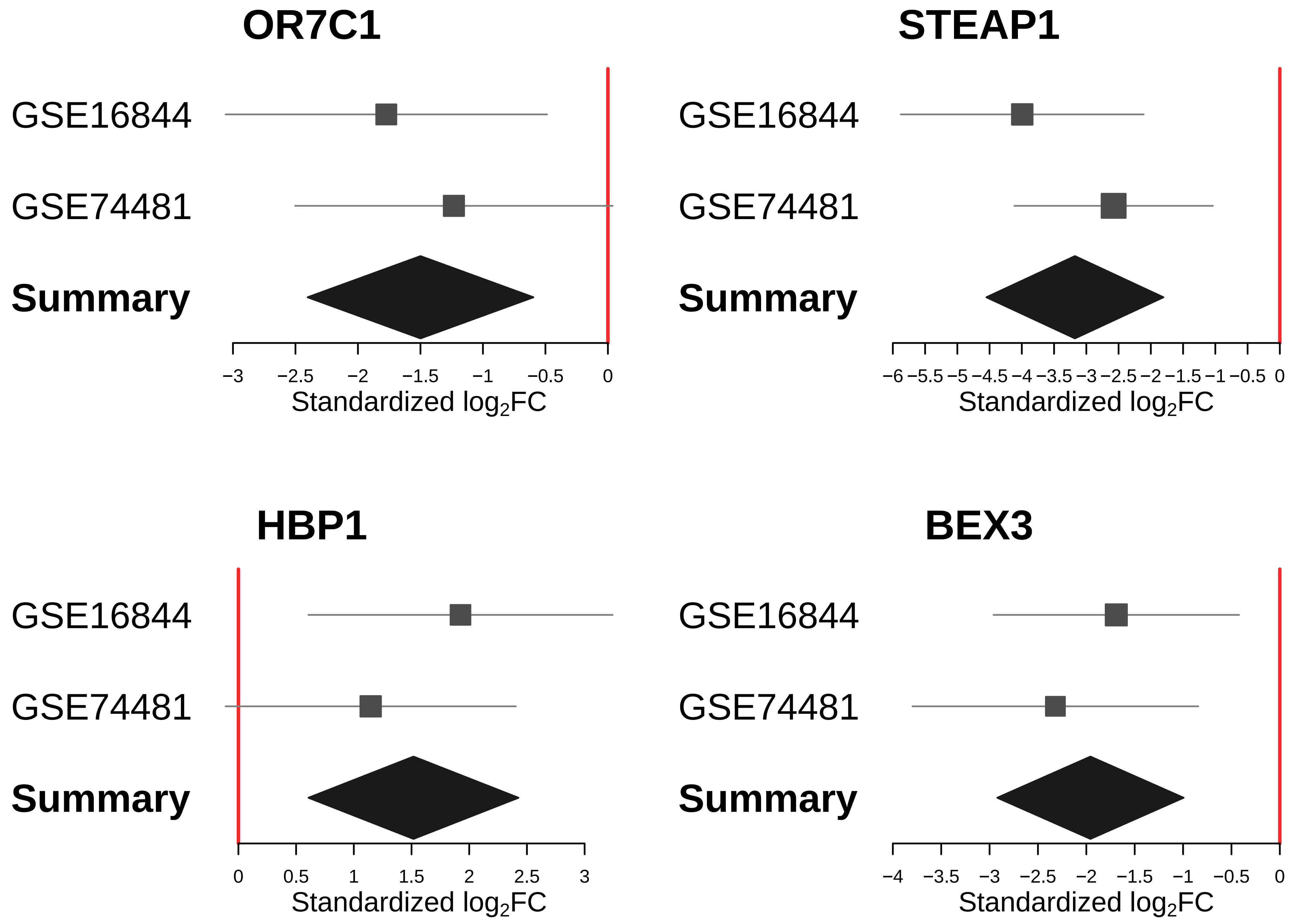

OR7C1
GSE16844
GSE74481
Summary
−3
−2.5
−2
−1.5
−1
−0.5
0
Standardized log2FC
STEAP1
GSE16844
GSE74481
Summary
−6
−5.5
−5
−4.5
−4
−3.5
−3
−2.5
−2
−1.5
−1
−0.5
0
Standardized log2FC
HBP1
GSE16844
GSE74481
Summary
0
0.5
1
1.5
2
2.5
3
Standardized log2FC
BEX3
GSE16844
GSE74481
Summary
−4
−3.5
−3
−2.5
−2
−1.5
−1
−0.5
0
Standardized log2FC

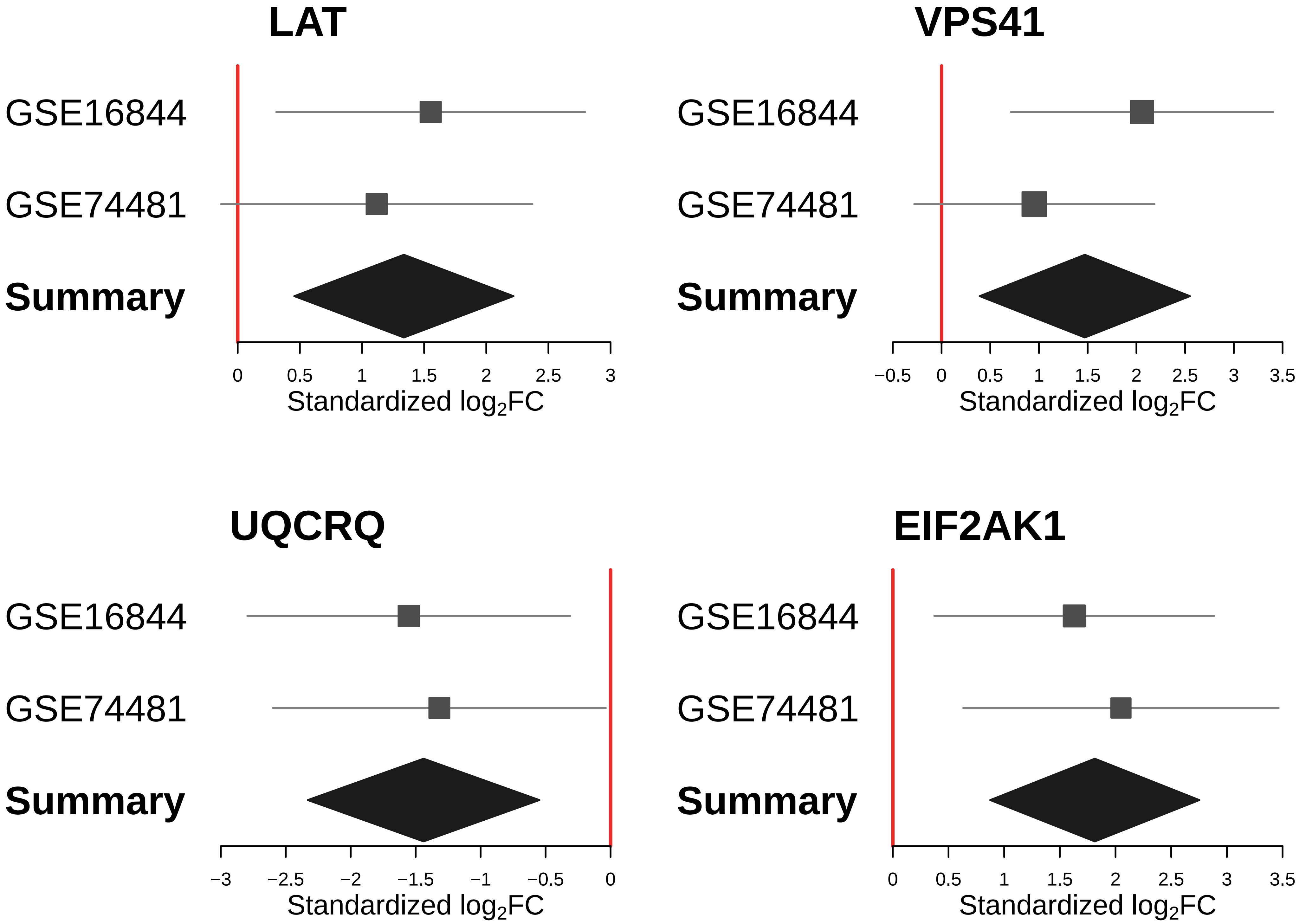

LAT
GSE16844
GSE74481
Summary
0
0.5
1
1.5
2
2.5
3
Standardized log2FC
VPS41
GSE16844
GSE74481
Summary
−0.5
0
0.5
1
1.5
2
2.5
3
3.5
Standardized log2FC
UQCRQ
GSE16844
GSE74481
Summary
−3
−2.5
−2
−1.5
−1
−0.5
0
Standardized log2FC
EIF2AK1
GSE16844
GSE74481
Summary
0
0.5
1
1.5
2
2.5
3
3.5
Standardized log2FC

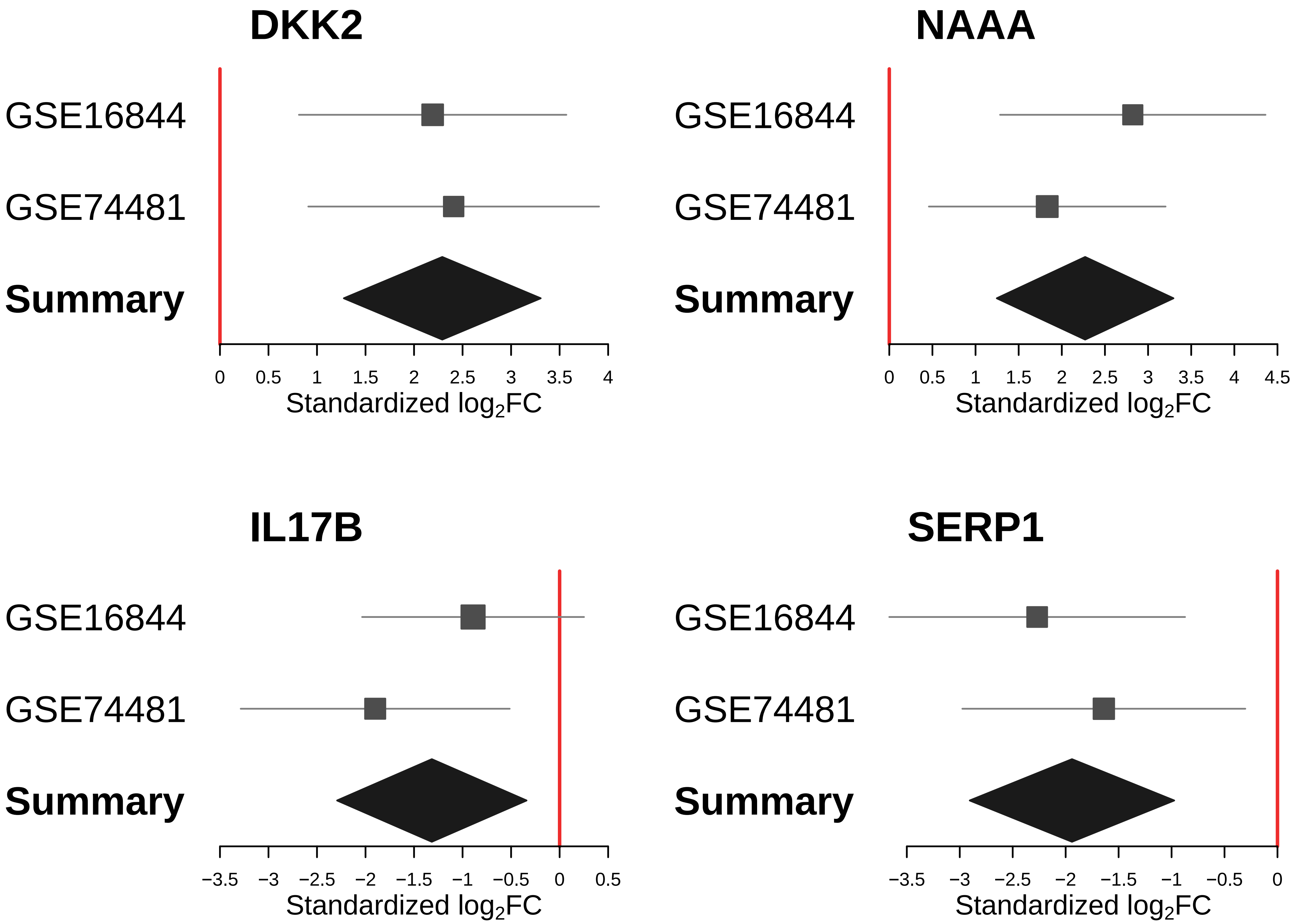
DKK2
GSE16844
GSE74481
Summary
0 0.5 1 1.5 2 2.5 3 3.5 4
Standardized log2FC
NAAA
GSE16844
GSE74481
Summary
0 0.5 1 1.5 2 2.5 3 3.5 4 4.5
Standardized log2FC
IL17B
GSE16844
GSE74481
Summary
−3.5 −3 −2.5 −2 −1.5 −1 −0.5 0 0.5
Standardized log2FC
SERP1
GSE16844
GSE74481
Summary
−3.5 −3 −2.5 −2 −1.5 −1 −0.5 0
Standardized log2FC

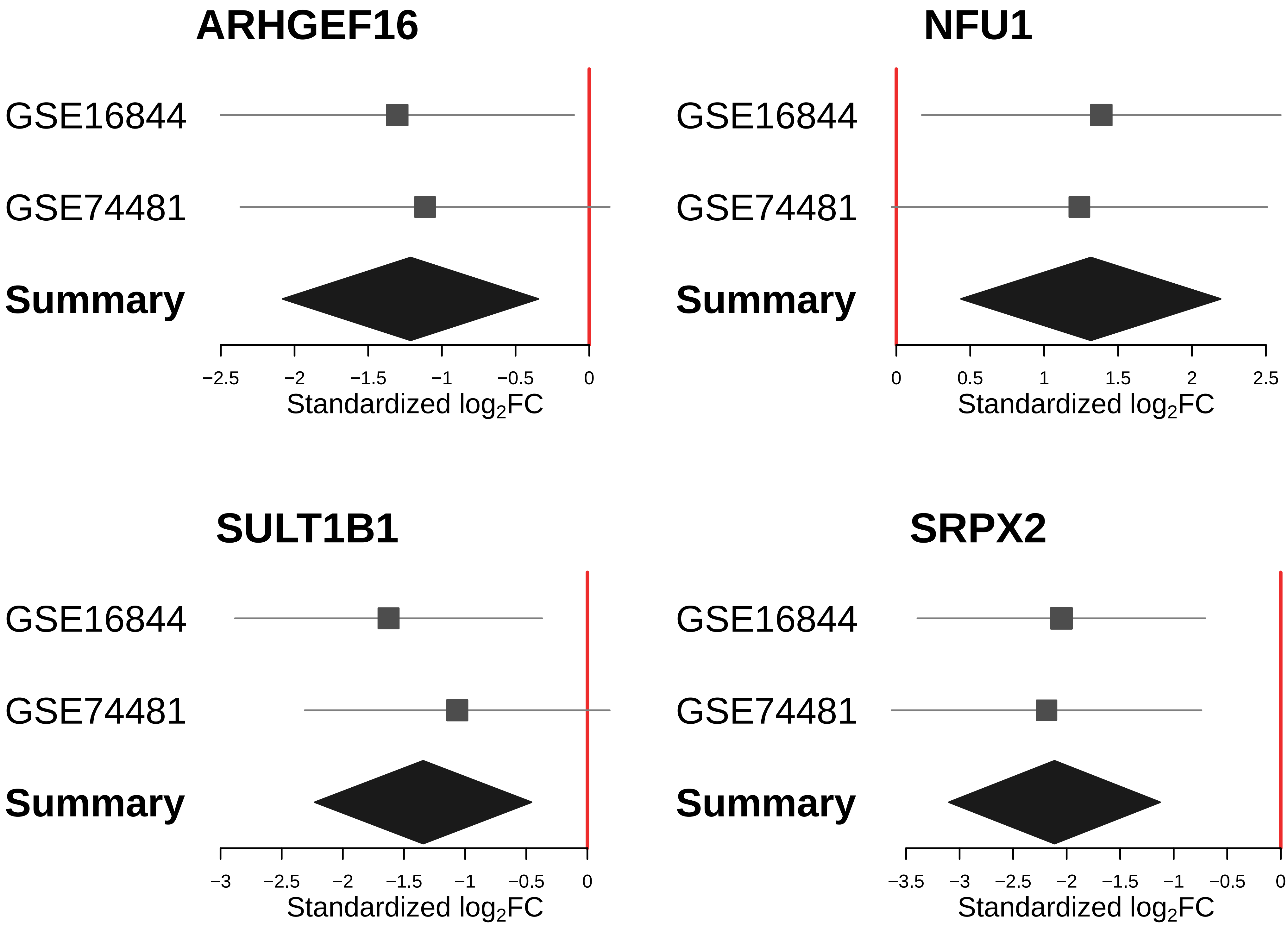
ARHGEF16
GSE16844
GSE74481
Summary
−2.5 −2 −1.5 −1 −0.5 0
Standardized log2FC
NFU1
GSE16844
GSE74481
Summary
0 0.5 1 1.5 2 2.5
Standardized log2FC
SULT1B1
GSE16844
GSE74481
Summary
−3 −2.5 −2 −1.5 −1 −0.5 0
Standardized log2FC
SRPX2
GSE16844
GSE74481
Summary
−3.5 −3 −2.5 −2 −1.5 −1 −0.5 0
Standardized log2FC

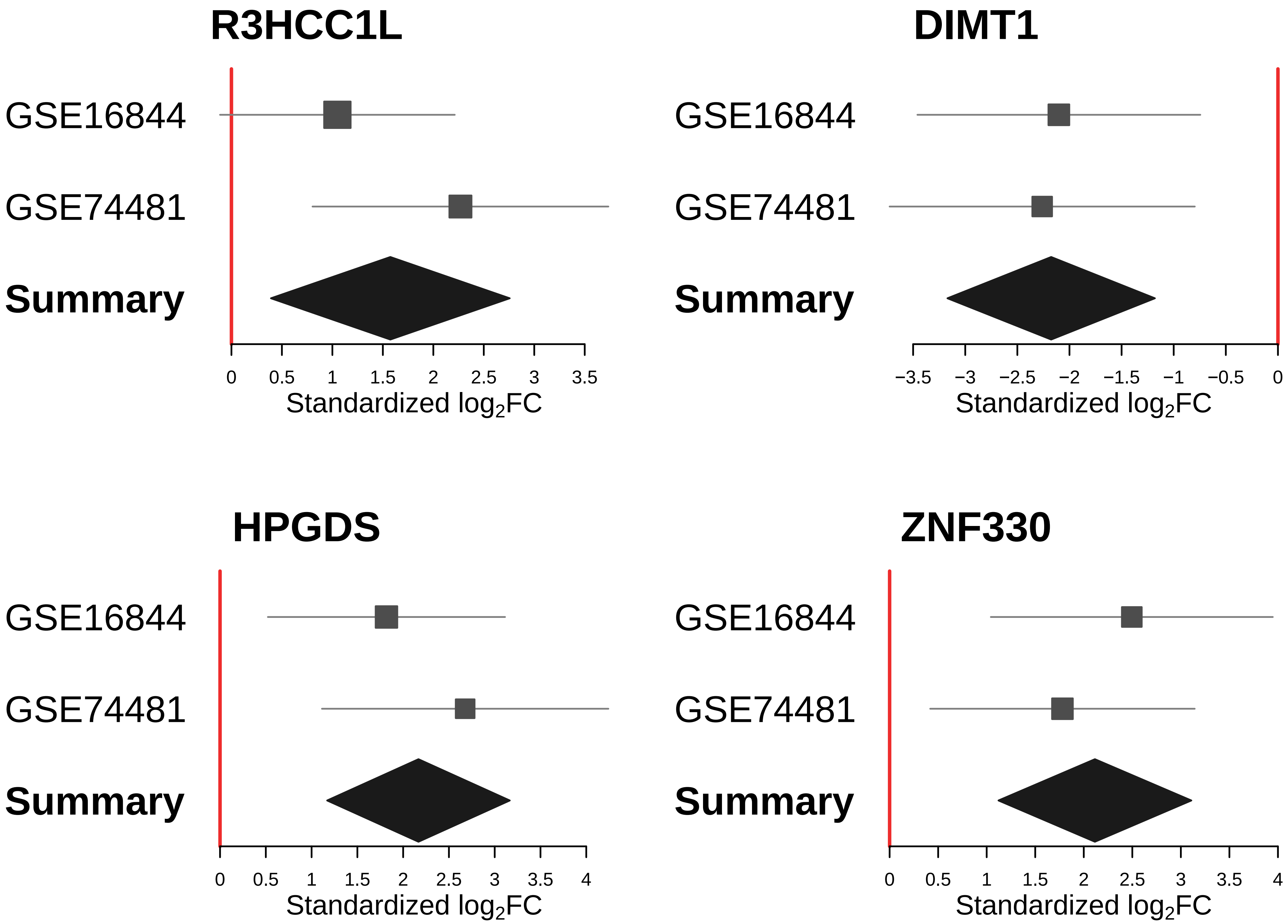
R3HCC1L
GSE16844
GSE74481
Summary
0 0.5 1 1.5 2 2.5 3 3.5
Standardized log2FC
DIMT1
GSE16844
GSE74481
Summary
−3.5 −3 −2.5 −2 −1.5 −1 −0.5 0
Standardized log2FC
HPGDS
GSE16844
GSE74481
Summary
0 0.5 1 1.5 2 2.5 3 3.5 4
Standardized log2FC
ZNF330
GSE16844
GSE74481
Summary
0 0.5 1 1.5 2 2.5 3 3.5 4
Standardized log2FC

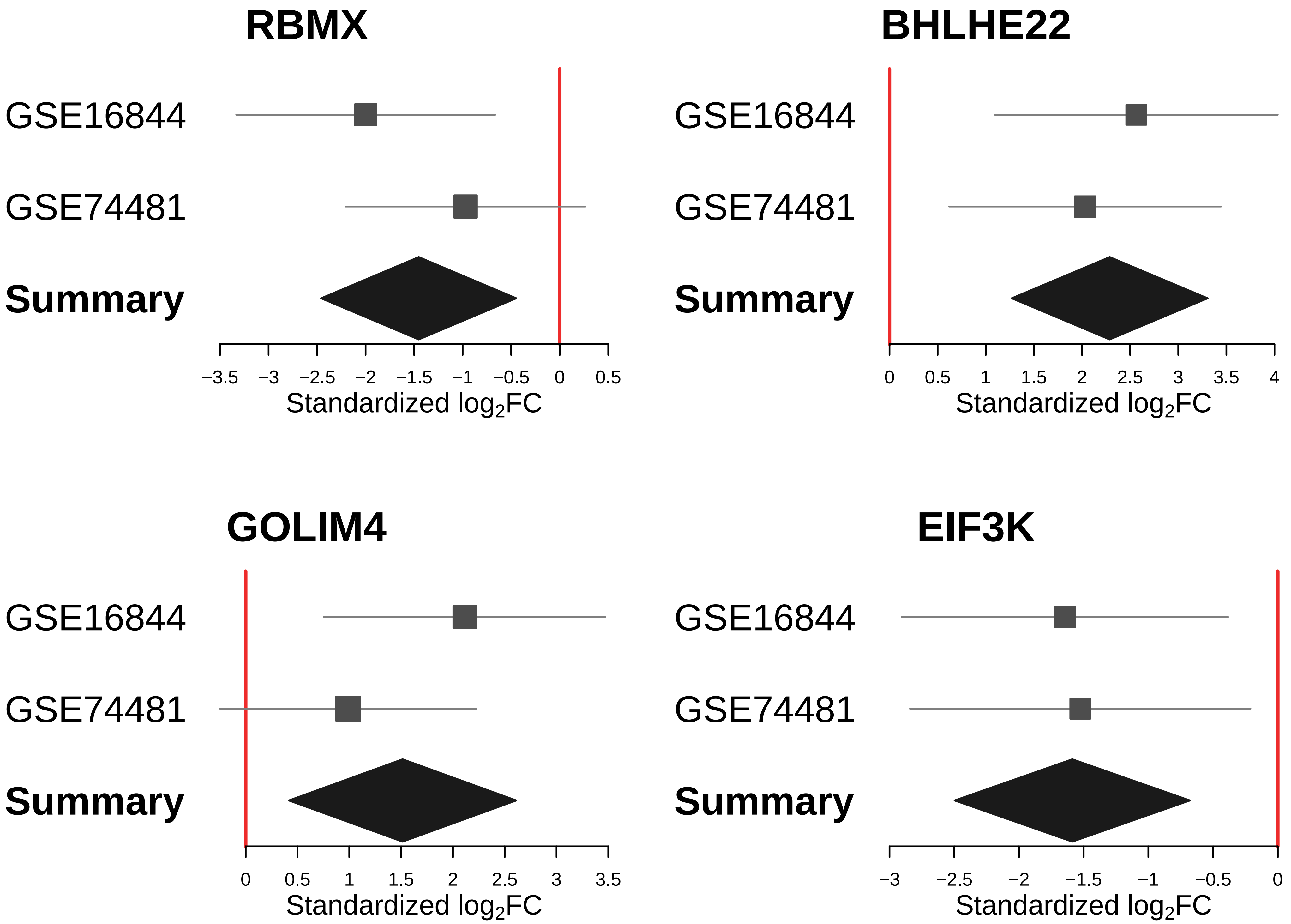
RBMX
GSE16844
GSE74481
Summary
−3.5 −3 −2.5 −2 −1.5 −1 −0.5 0 0.5
Standardized log2FC
BHLHE22
GSE16844
GSE74481
Summary
0 0.5 1 1.5 2 2.5 3 3.5 4
Standardized log2FC
GOLIM4
GSE16844
GSE74481
Summary
0 0.5 1 1.5 2 2.5 3 3.5
Standardized log2FC
EIF3K
GSE16844
GSE74481
Summary
−3 −2.5 −2 −1.5 −1 −0.5 0
Standardized log2FC

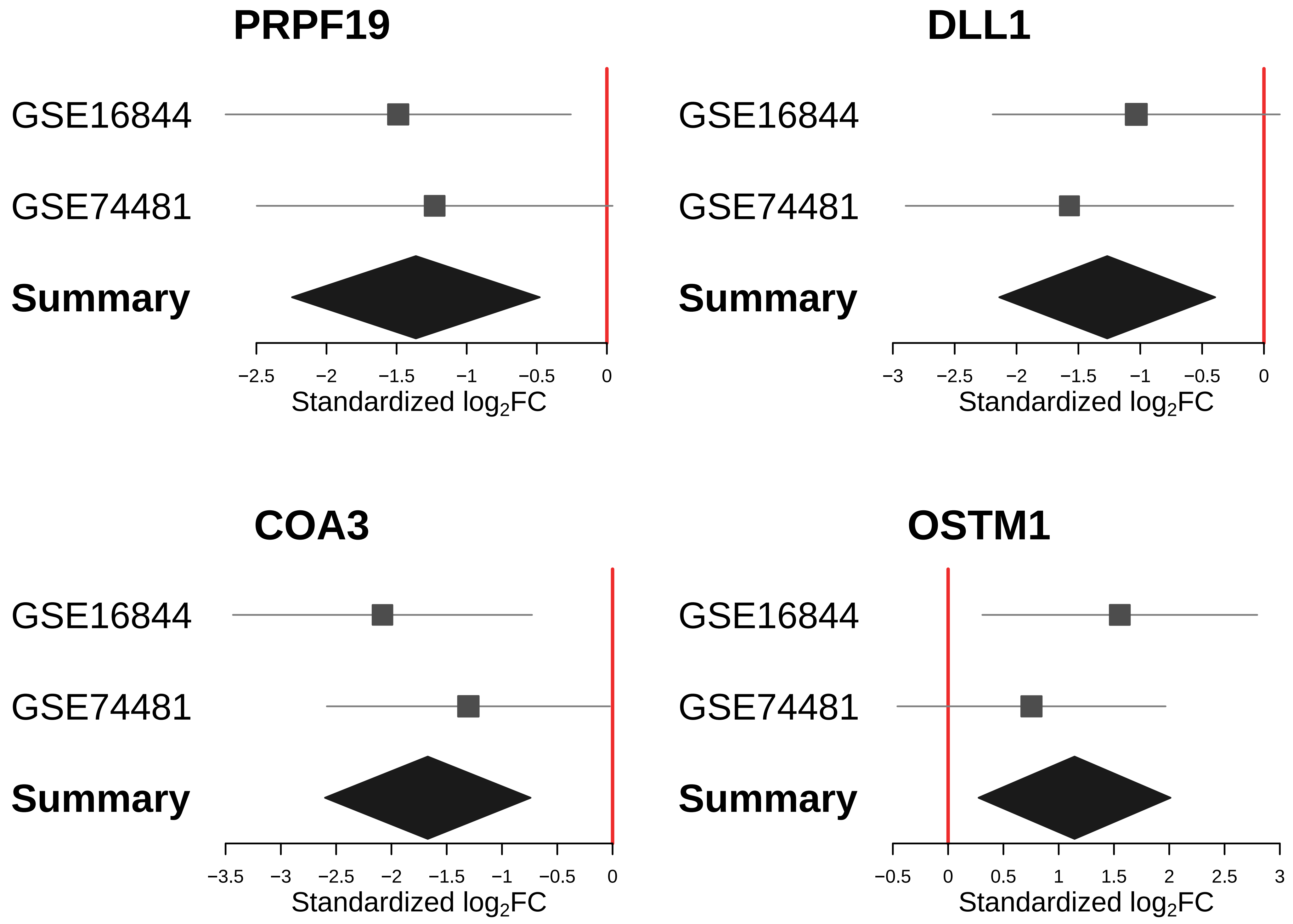
PRPF19
GSE16844
GSE74481
Summary
−2.5 −2 −1.5 −1 −0.5 0
Standardized log2FC
DLL1
GSE16844
GSE74481
Summary
−3 −2.5 −2 −1.5 −1 −0.5 0
Standardized log2FC
COA3
GSE16844
GSE74481
Summary
−3.5 −3 −2.5 −2 −1.5 −1 −0.5 0
Standardized log2FC
OSTM1
GSE16844
GSE74481
Summary
−0.5 0 0.5 1 1.5 2 2.5 3
Standardized log2FC

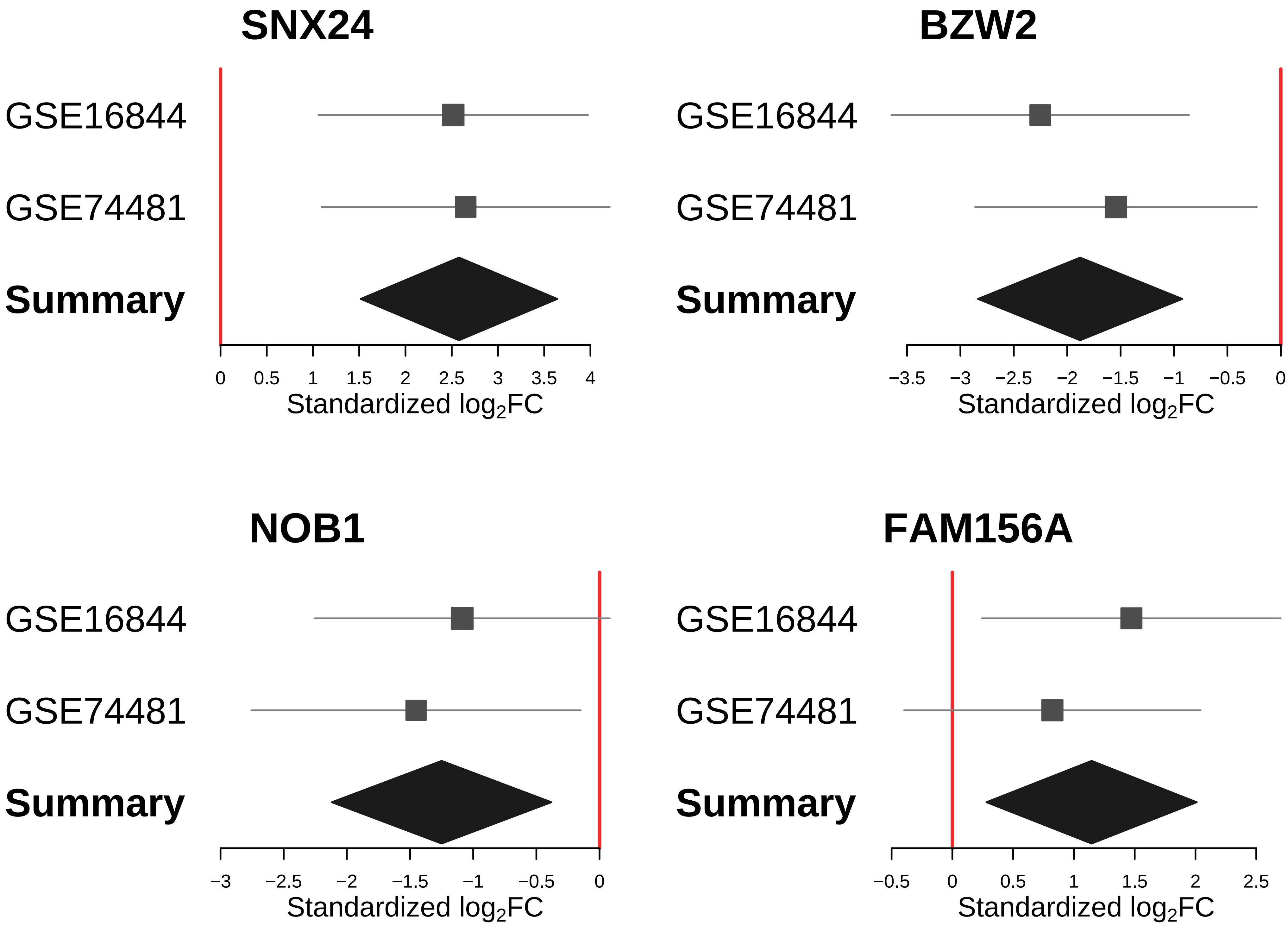

SNX24
GSE16844
GSE74481
Summary
0 0.5 1 1.5 2 2.5 3 3.5 4
Standardized log2FC
BZW2
GSE16844
GSE74481
Summary
−3.5 −3 −2.5 −2 −1.5 −1 −0.5 0
Standardized log2FC
NOB1
GSE16844
GSE74481
Summary
−3 −2.5 −2 −1.5 −1 −0.5 0
Standardized log2FC
FAM156A
GSE16844
GSE74481
Summary
−0.5 0 0.5 1 1.5 2 2.5
Standardized log2FC

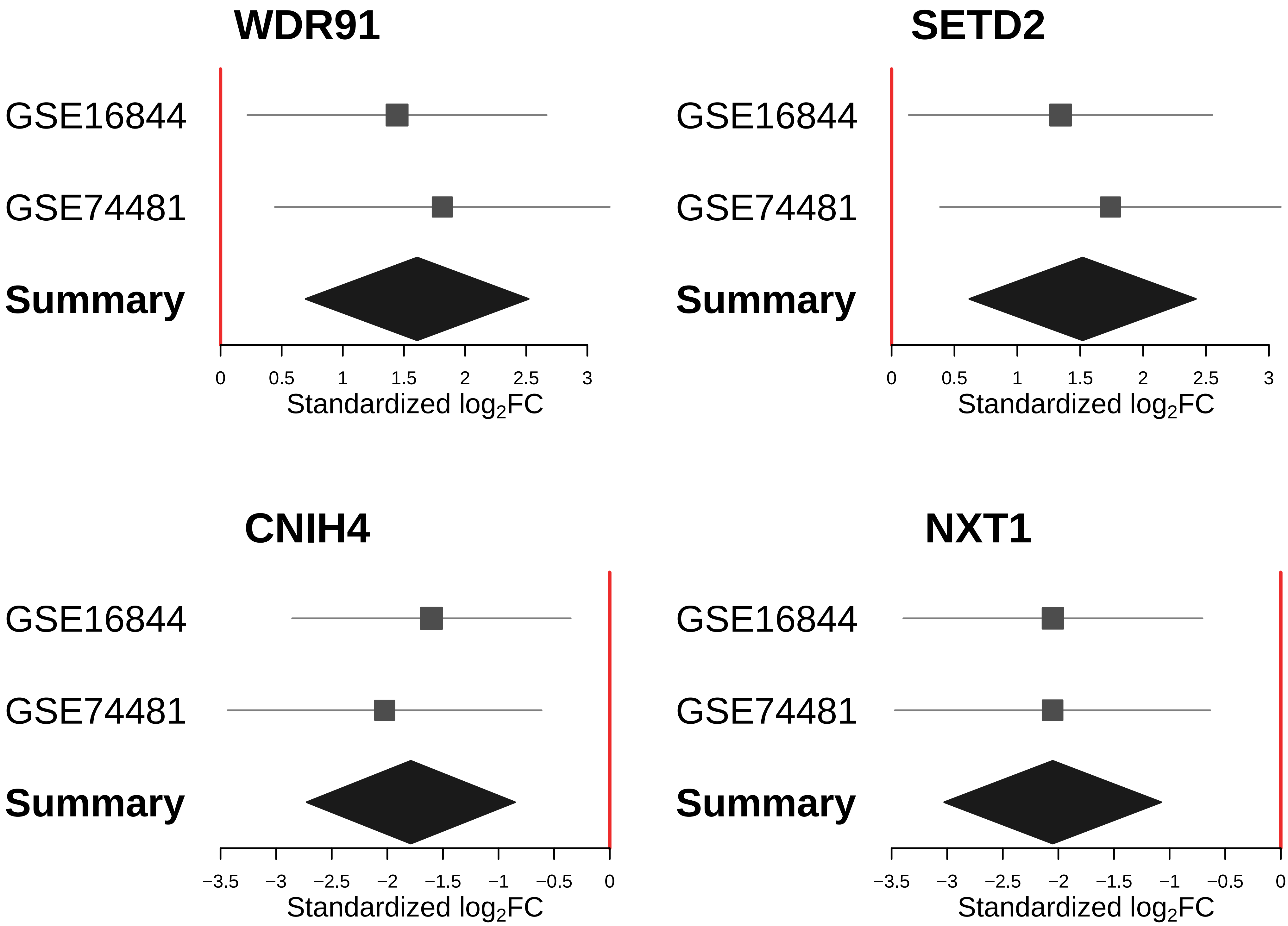
WDR91
GSE16844
GSE74481
Summary
0 0.5 1 1.5 2 2.5 3
Standardized log2FC
SETD2
GSE16844
GSE74481
Summary
0 0.5 1 1.5 2 2.5 3
Standardized log2FC
CNIH4
GSE16844
GSE74481
Summary
−3.5 −3 −2.5 −2 −1.5 −1 −0.5 0
Standardized log2FC
NXT1
GSE16844
GSE74481
Summary
−3.5 −3 −2.5 −2 −1.5 −1 −0.5 0
Standardized log2FC

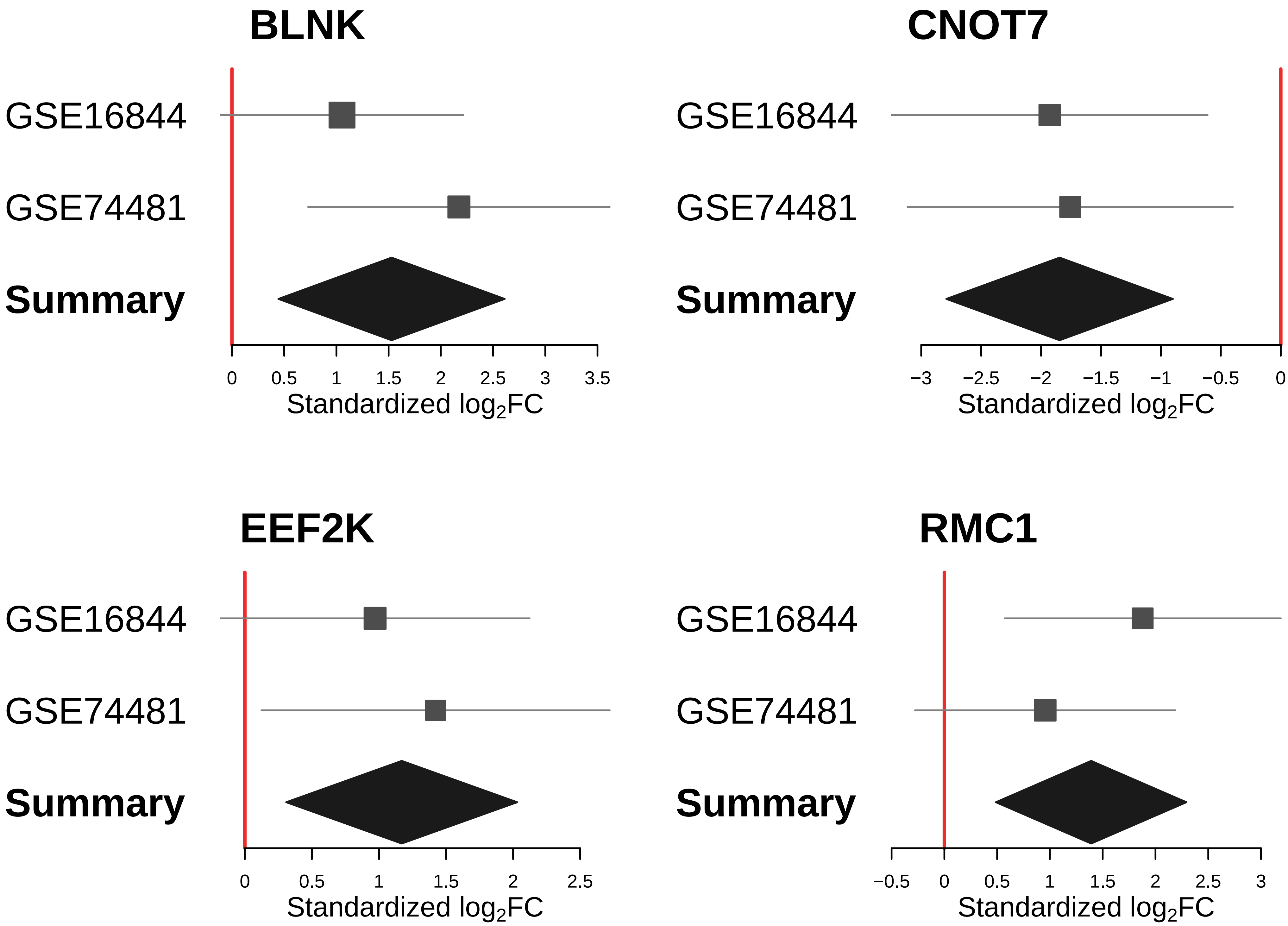

BLNK
GSE16844
GSE74481
Summary
0 0.5 1 1.5 2 2.5 3 3.5
Standardized log2FC
CNOT7
GSE16844
GSE74481
Summary
−3 −2.5 −2 −1.5 −1 −0.5 0
Standardized log2FC
EEF2K
GSE16844
GSE74481
Summary
0 0.5 1 1.5 2 2.5
Standardized log2FC
RMC1
GSE16844
GSE74481
Summary
−0.5 0 0.5 1 1.5 2 2.5 3
Standardized log2FC

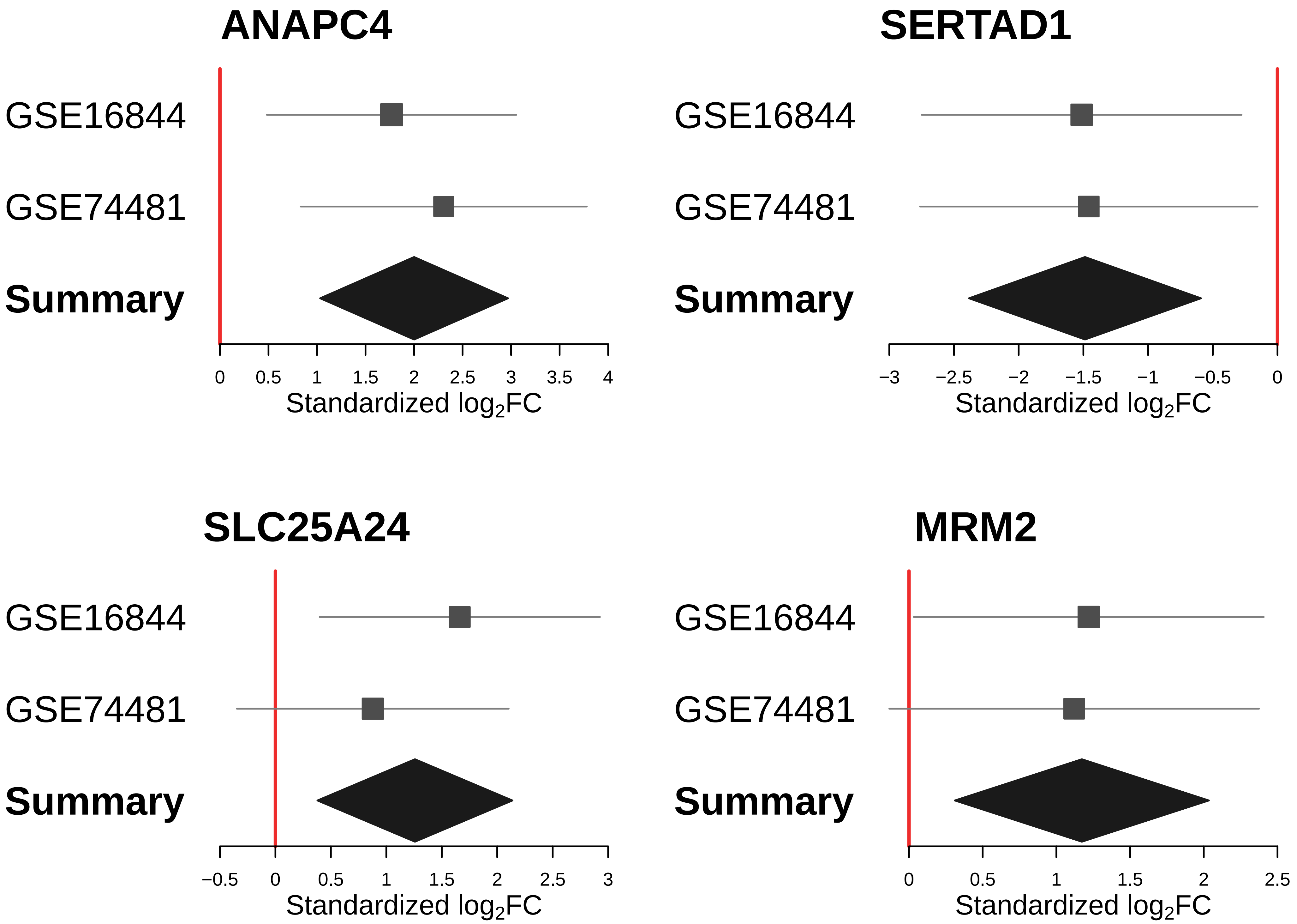
ANAPC4
GSE16844
GSE74481
Summary
0 0.5 1 1.5 2 2.5 3 3.5 4
Standardized log2FC
SERTAD1
GSE16844
GSE74481
Summary
−3 −2.5 −2 −1.5 −1 −0.5 0
Standardized log2FC
SLC25A24
GSE16844
GSE74481
Summary
−0.5 0 0.5 1 1.5 2 2.5 3
Standardized log2FC
MRM2
GSE16844
GSE74481
Summary
0 0.5 1 1.5 2 2.5
Standardized log2FC

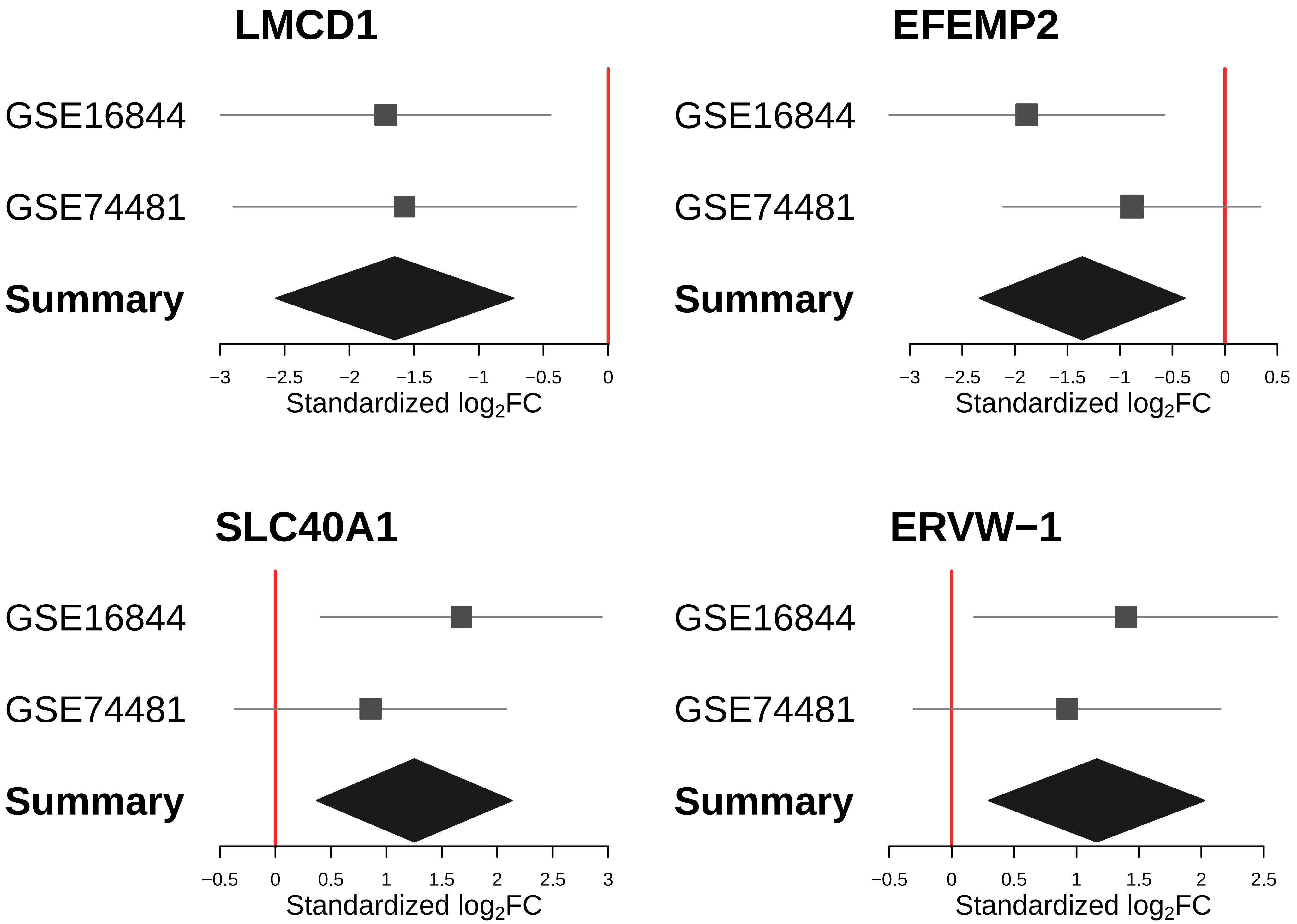
LMCD1
GSE16844
GSE74481
Summary
−3 −2.5 −2 −1.5 −1 −0.5 0
Standardized log2FC
EFEMP2
GSE16844
GSE74481
Summary
−3 −2.5 −2 −1.5 −1 −0.5 0 0.5
Standardized log2FC
SLC40A1
GSE16844
GSE74481
Summary
−0.5 0 0.5 1 1.5 2 2.5 3
Standardized log2FC
ERVW−1
GSE16844
GSE74481
Summary
−0.5 0 0.5 1 1.5 2 2.5
Standardized log2FC

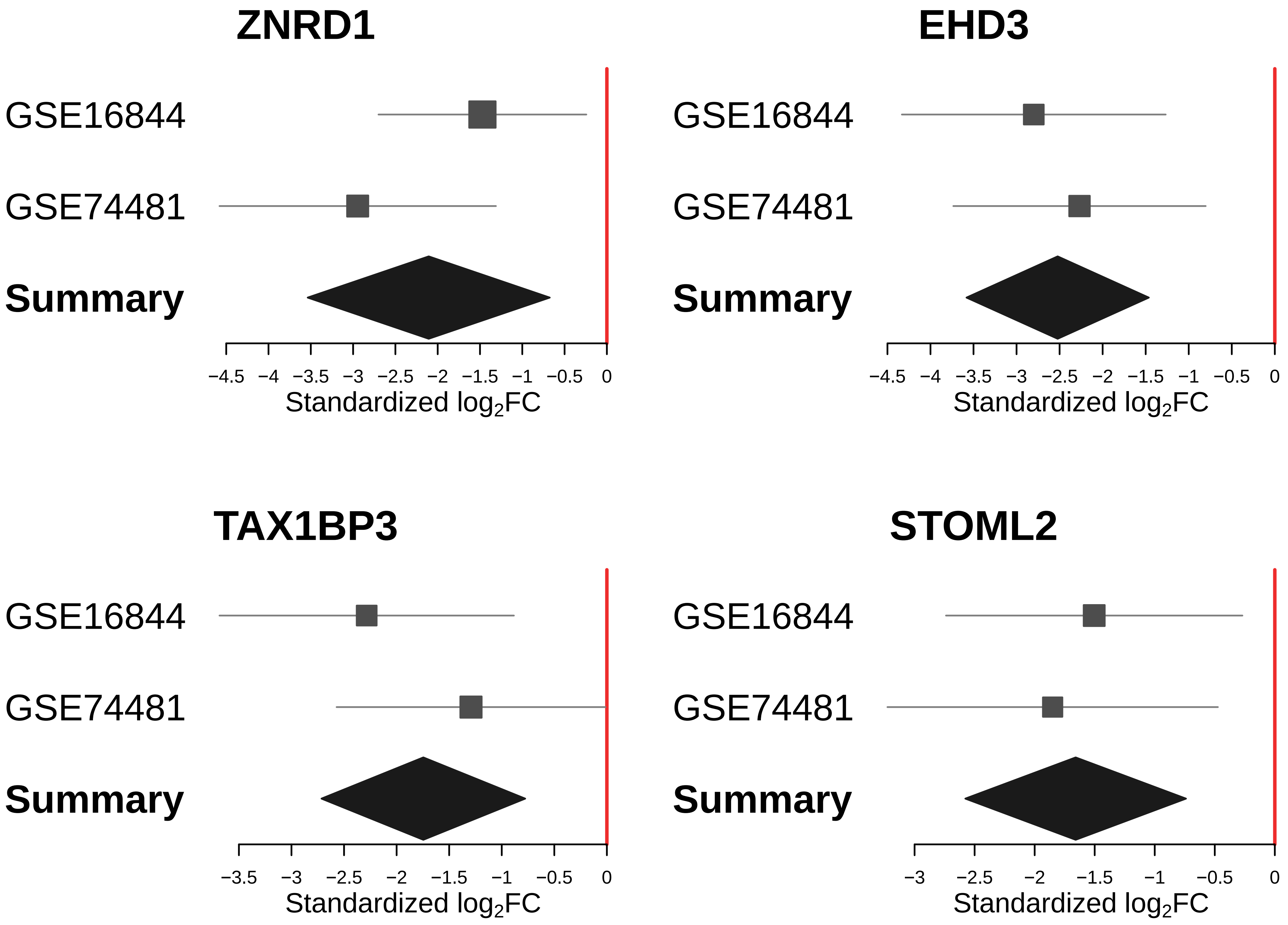
ZNRD1
GSE16844
GSE74481
Summary
−4.5 −4 −3.5 −3 −2.5 −2 −1.5 −1 −0.5 0
Standardized $log_2FC$
EHD3
GSE16844
GSE74481
Summary
−4.5 −4 −3.5 −3 −2.5 −2 −1.5 −1 −0.5 0
Standardized $log_2FC$
TAX1BP3
GSE16844
GSE74481
Summary
−3.5 −3 −2.5 −2 −1.5 −1 −0.5 0
Standardized $log_2FC$
STOML2
GSE16844
GSE74481
Summary
−3 −2.5 −2 −1.5 −1 −0.5 0
Standardized $log_2FC$

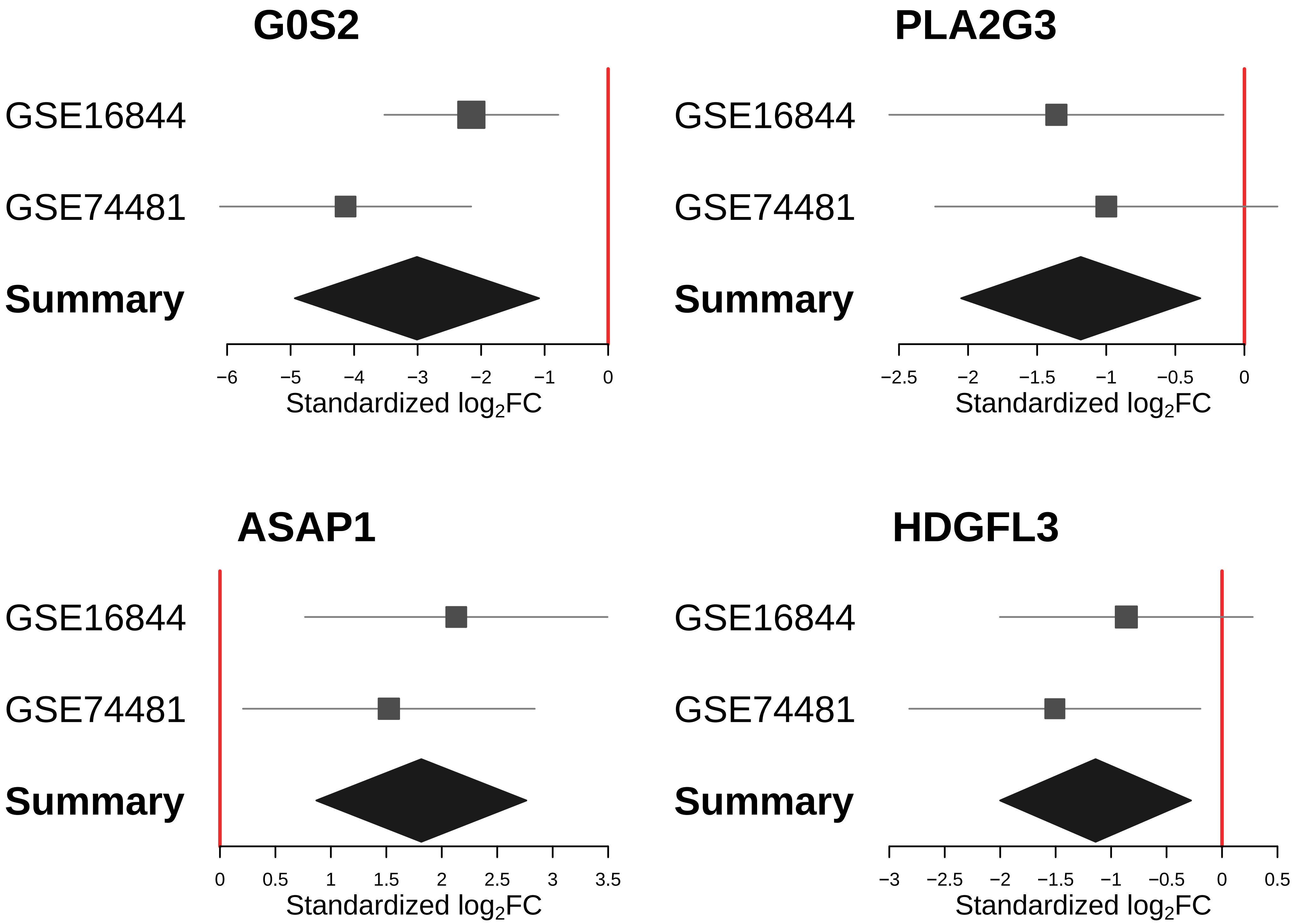

G0S2
GSE16844
GSE74481
Summary
−6
−5
−4
−3
−2
−1
0
Standardized log2FC
PLA2G3
GSE16844
GSE74481
Summary
−2.5
−2
−1.5
−1
−0.5
0
Standardized log2FC
ASAP1
GSE16844
GSE74481
Summary
0
0.5
1
1.5
2
2.5
3
3.5
Standardized log2FC
HDGFL3
GSE16844
GSE74481
Summary
−3
−2.5
−2
−1.5
−1
−0.5
0
0.5
Standardized log2FC

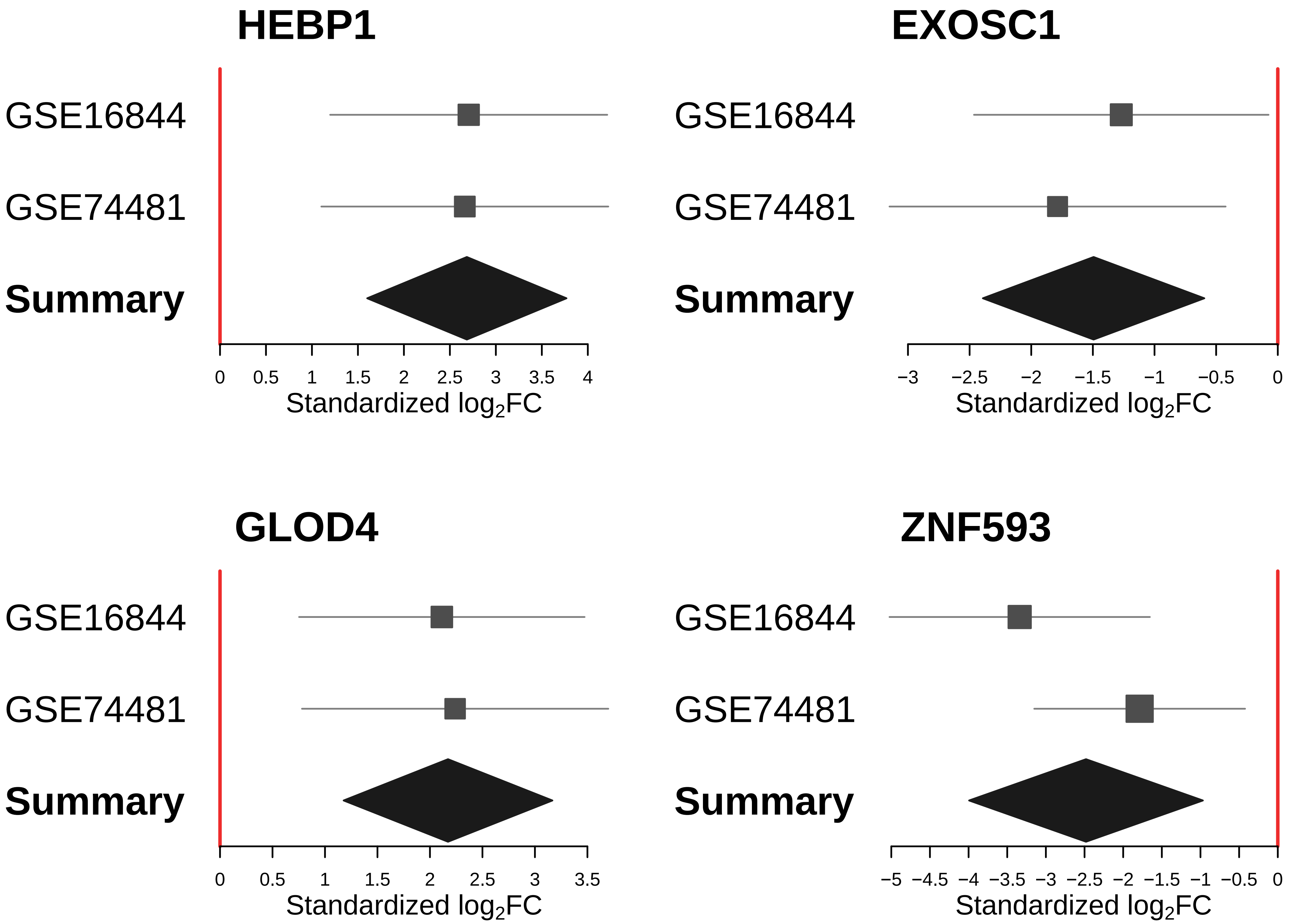
HEBP1
GSE16844
GSE74481
Summary
0
0.5
1
1.5
2
2.5
3
3.5
4
Standardized log2FC
EXOSC1
GSE16844
GSE74481
Summary
−3
−2.5
−2
−1.5
−1
−0.5
0
Standardized log2FC
GLOD4
GSE16844
GSE74481
Summary
0
0.5
1
1.5
2
2.5
3
3.5
Standardized log2FC
ZNF593
GSE16844
GSE74481
Summary
−5
−4.5
−4
−3.5
−3
−2.5
−2
−1.5
−1
−0.5
0
Standardized log2FC

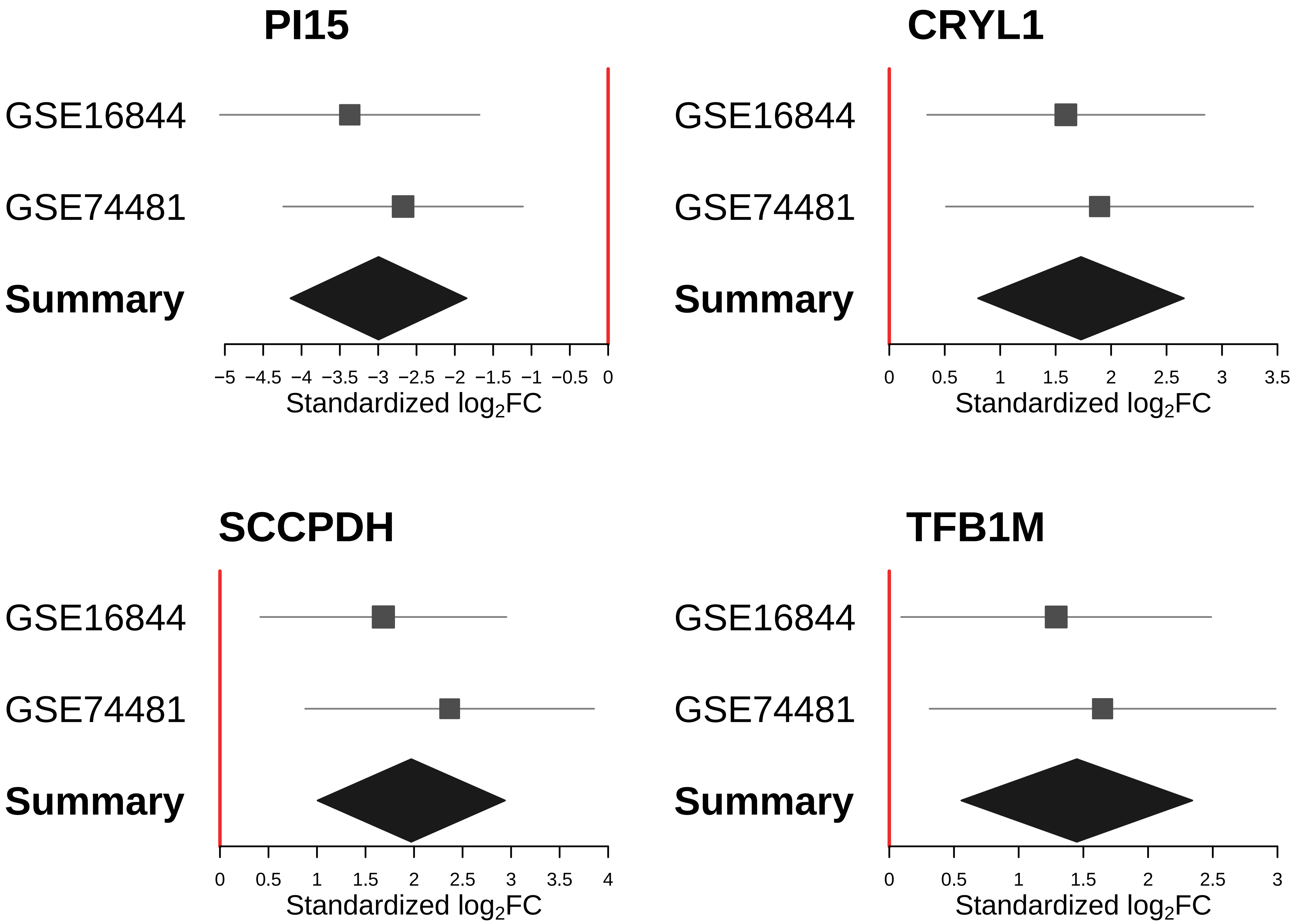
PI15
GSE16844
GSE74481
Summary
−5 −4.5 −4 −3.5 −3 −2.5 −2 −1.5 −1 −0.5 0
Standardized log2FC
CRYL1
GSE16844
GSE74481
Summary
0 0.5 1 1.5 2 2.5 3 3.5
Standardized log2FC
SCCPDH
GSE16844
GSE74481
Summary
0 0.5 1 1.5 2 2.5 3 3.5 4
Standardized log2FC
TFB1M
GSE16844
GSE74481
Summary
0 0.5 1 1.5 2 2.5 3
Standardized log2FC

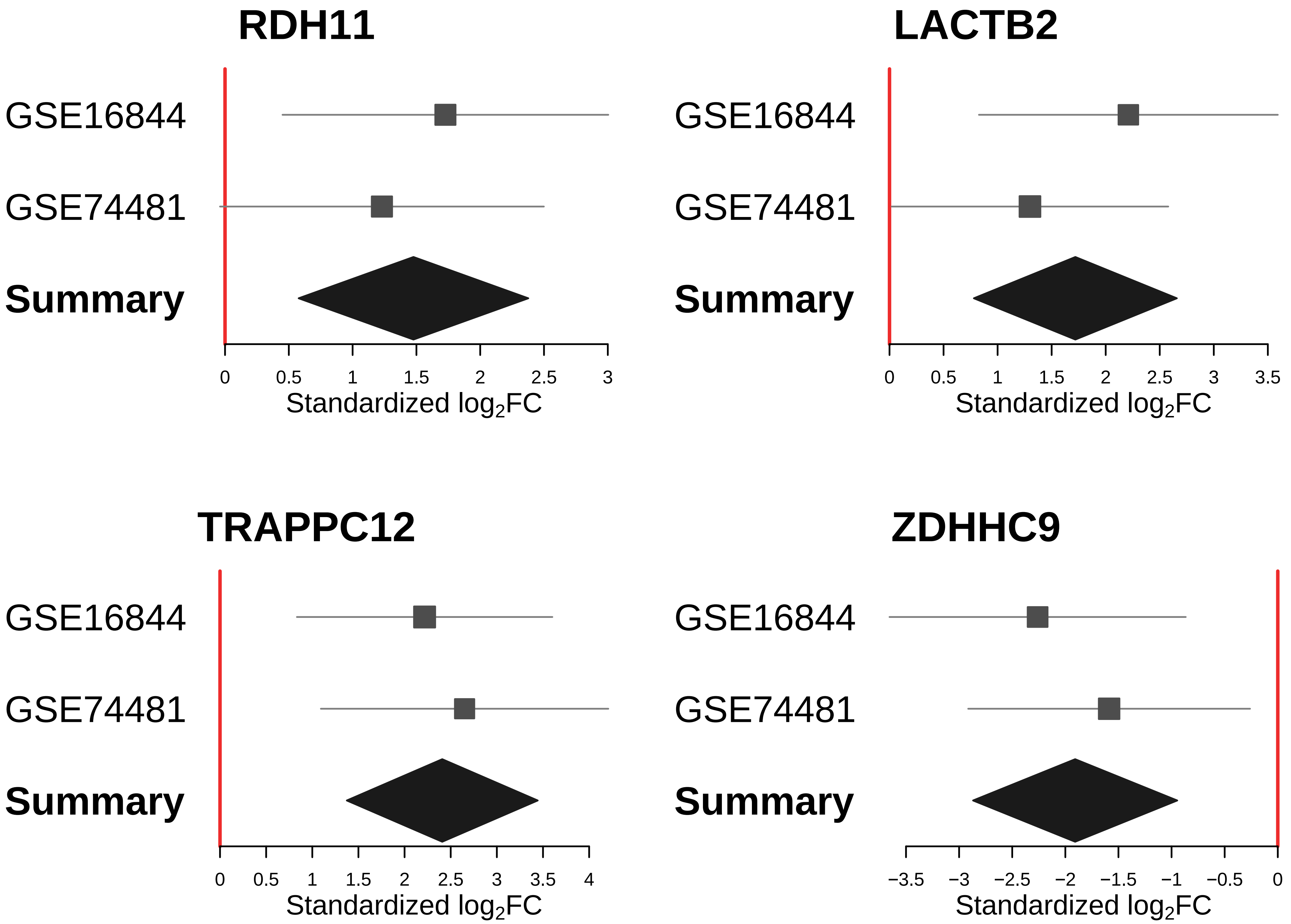
RDH11
GSE16844
GSE74481
Summary
0 0.5 1 1.5 2 2.5 3
Standardized log2FC
LACTB2
GSE16844
GSE74481
Summary
0 0.5 1 1.5 2 2.5 3 3.5
Standardized log2FC
TRAPPC12
GSE16844
GSE74481
Summary
0 0.5 1 1.5 2 2.5 3 3.5 4
Standardized log2FC
ZDHHC9
GSE16844
GSE74481
Summary
−3.5 −3 −2.5 −2 −1.5 −1 −0.5 0
Standardized log2FC

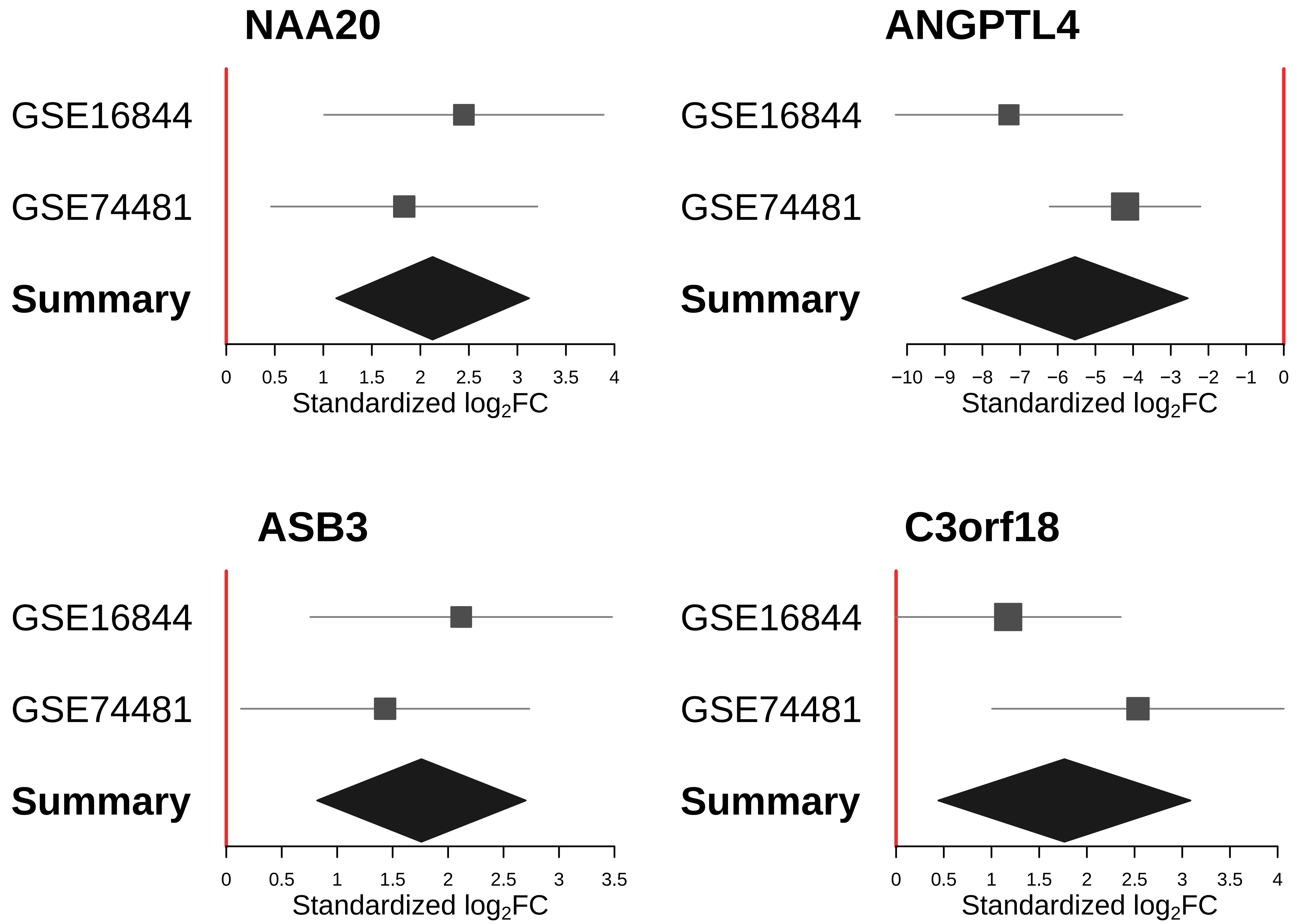
NAA20
GSE16844
GSE74481
Summary
0 0.5 1 1.5 2 2.5 3 3.5 4
Standardized log2FC
ANGPTL4
GSE16844
GSE74481
Summary
−10 −9 −8 −7 −6 −5 −4 −3 −2 −1 0
Standardized log2FC
ASB3
GSE16844
GSE74481
Summary
0 0.5 1 1.5 2 2.5 3 3.5
Standardized log2FC
C3orf18
GSE16844
GSE74481
Summary
0 0.5 1 1.5 2 2.5 3 3.5 4
Standardized log2FC

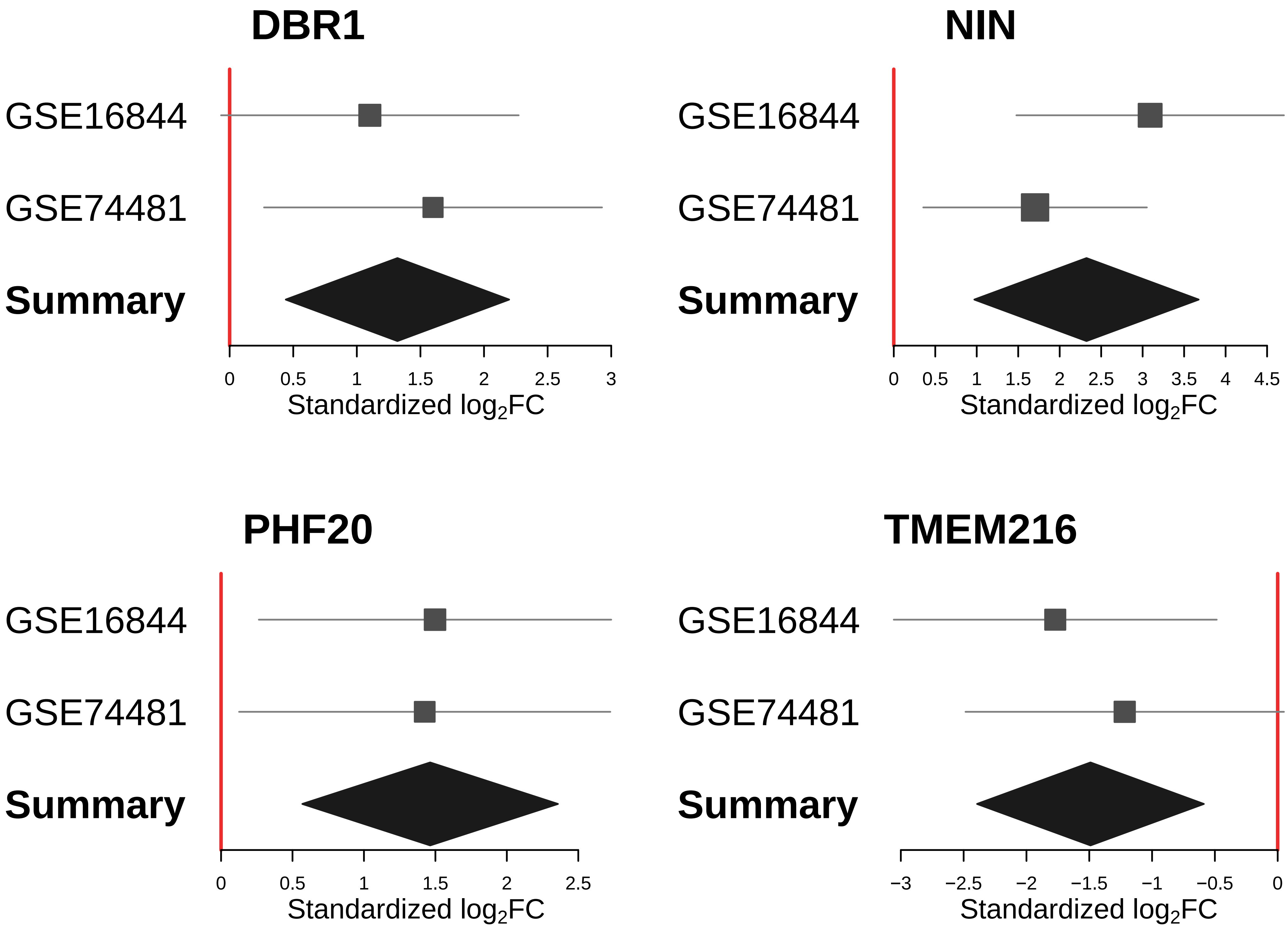
DBR1
GSE16844
GSE74481
Summary
0 0.5 1 1.5 2 2.5 3
Standardized log2FC
NIN
GSE16844
GSE74481
Summary
0 0.5 1 1.5 2 2.5 3 3.5 4 4.5
Standardized log2FC
PHF20
GSE16844
GSE74481
Summary
0 0.5 1 1.5 2 2.5
Standardized log2FC
TMEM216
GSE16844
GSE74481
Summary
−3 −2.5 −2 −1.5 −1 −0.5 0
Standardized log2FC

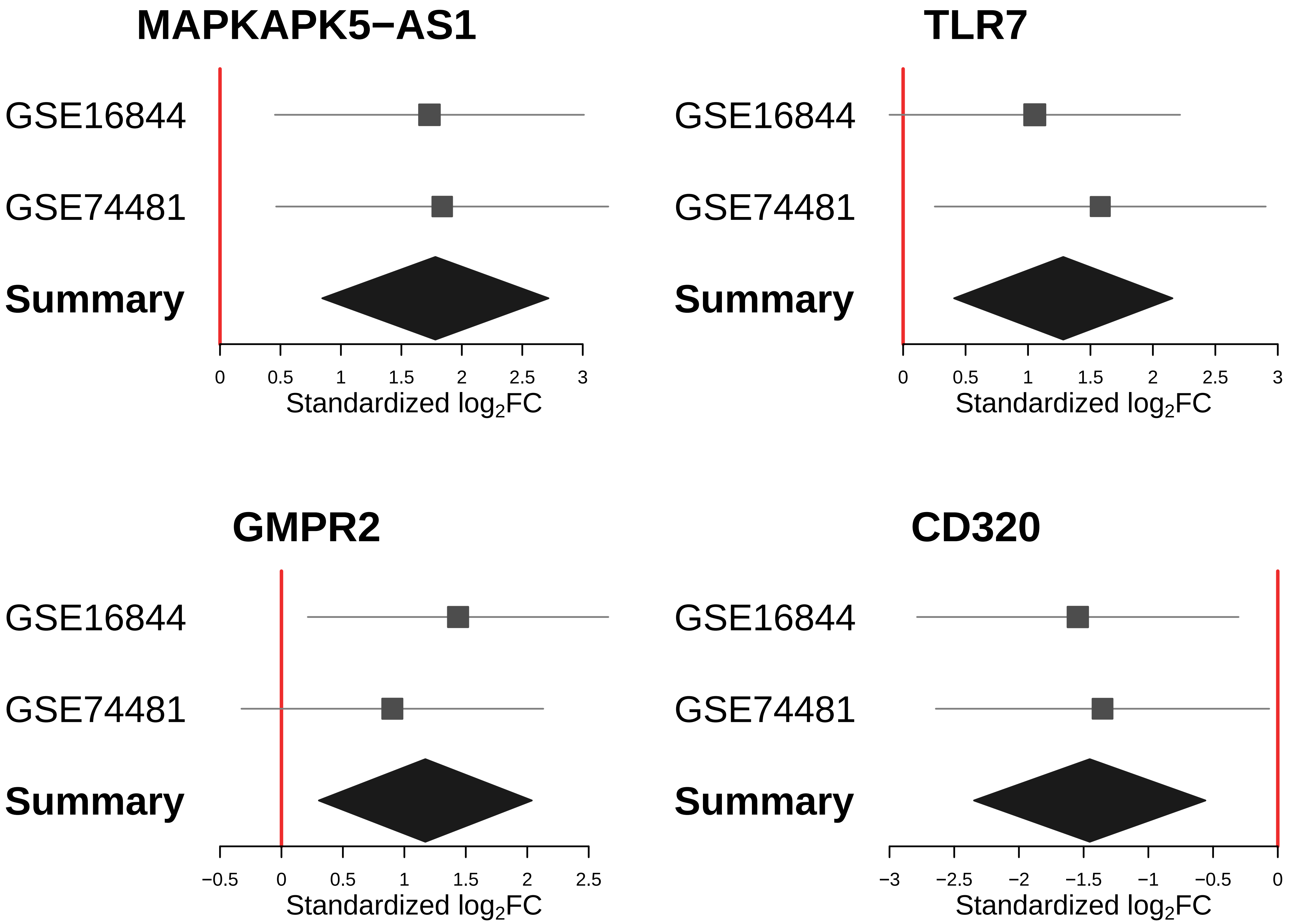
MAPKAPK5−AS1
TLR7
GMPR2
CD320
GSE16844
GSE74481
Summary
0 0.5 1 1.5 2 2.5 3
−0.5 0 0.5 1 1.5 2 2.5
−3 −2.5 −2 −1.5 −1 −0.5 0
Standardized log2FC

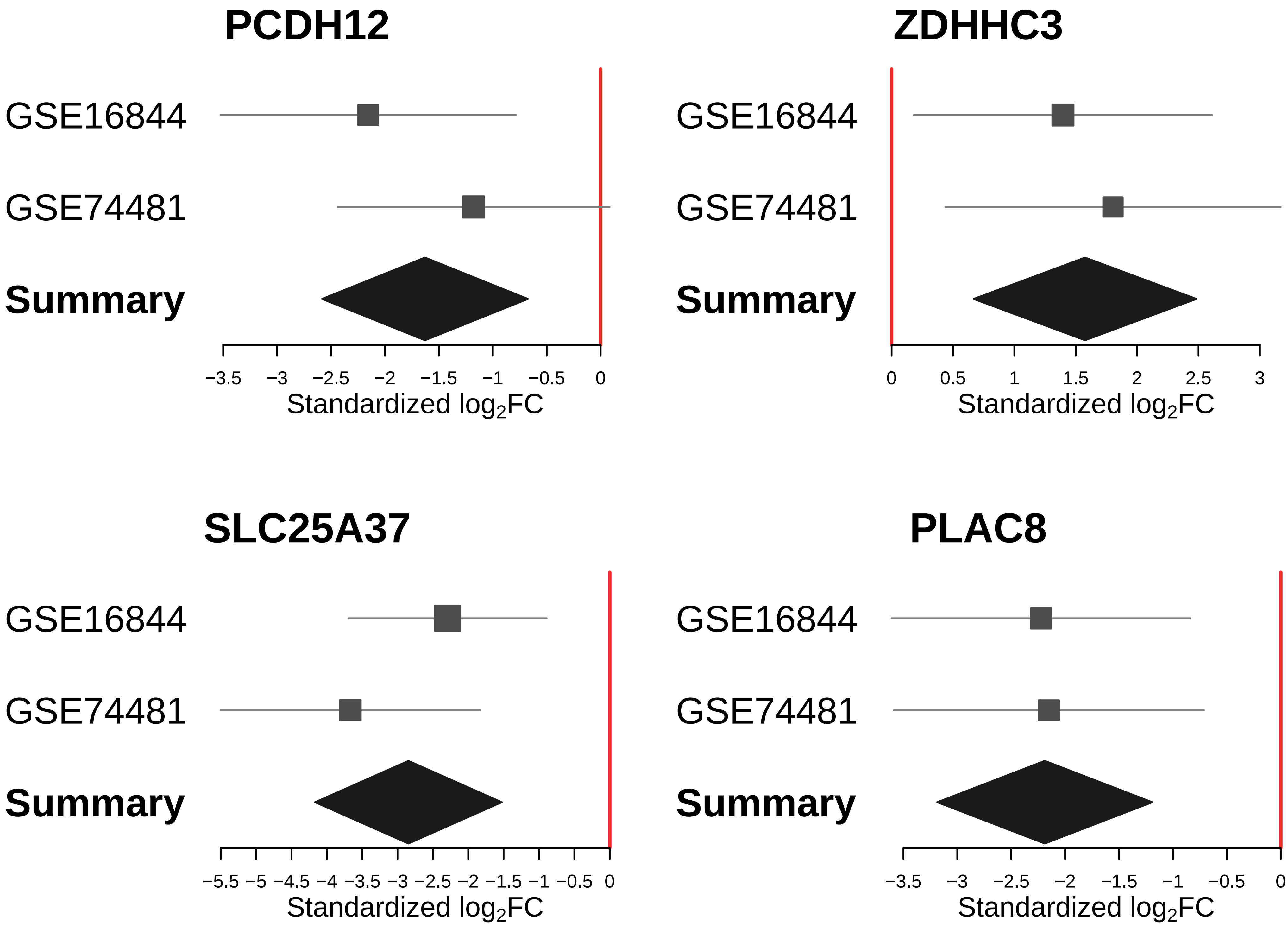
PCDH12
GSE16844
GSE74481
Summary
−3.5 −3 −2.5 −2 −1.5 −1 −0.5 0
Standardized log2FC
ZDHHC3
GSE16844
GSE74481
Summary
0 0.5 1 1.5 2 2.5 3
Standardized log2FC
SLC25A37
GSE16844
GSE74481
Summary
−5.5 −5 −4.5 −4 −3.5 −3 −2.5 −2 −1.5 −1 −0.5 0
Standardized log2FC
PLAC8
GSE16844
GSE74481
Summary
−3.5 −3 −2.5 −2 −1.5 −1 −0.5 0
Standardized log2FC

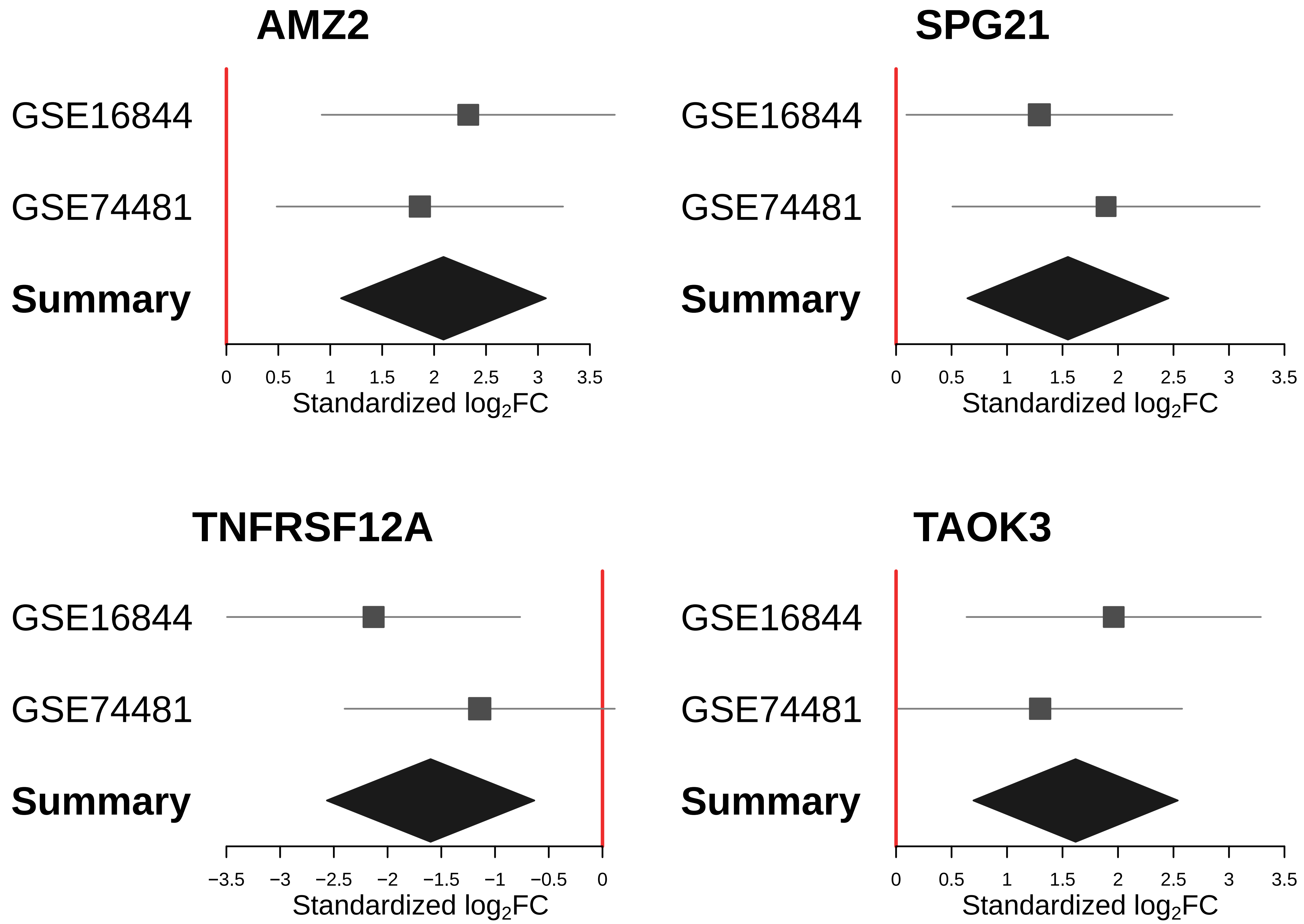
AMZ2
GSE16844
GSE74481
Summary
0
0.5
1
1.5
2
2.5
3
3.5
Standardized log2FC
SPG21
GSE16844
GSE74481
Summary
0
0.5
1
1.5
2
2.5
3
3.5
Standardized log2FC
TNFRSF12A
GSE16844
GSE74481
Summary
−3.5
−3
−2.5
−2
−1.5
−1
−0.5
0
Standardized log2FC
TAOK3
GSE16844
GSE74481
Summary
0
0.5
1
1.5
2
2.5
3
3.5
Standardized log2FC

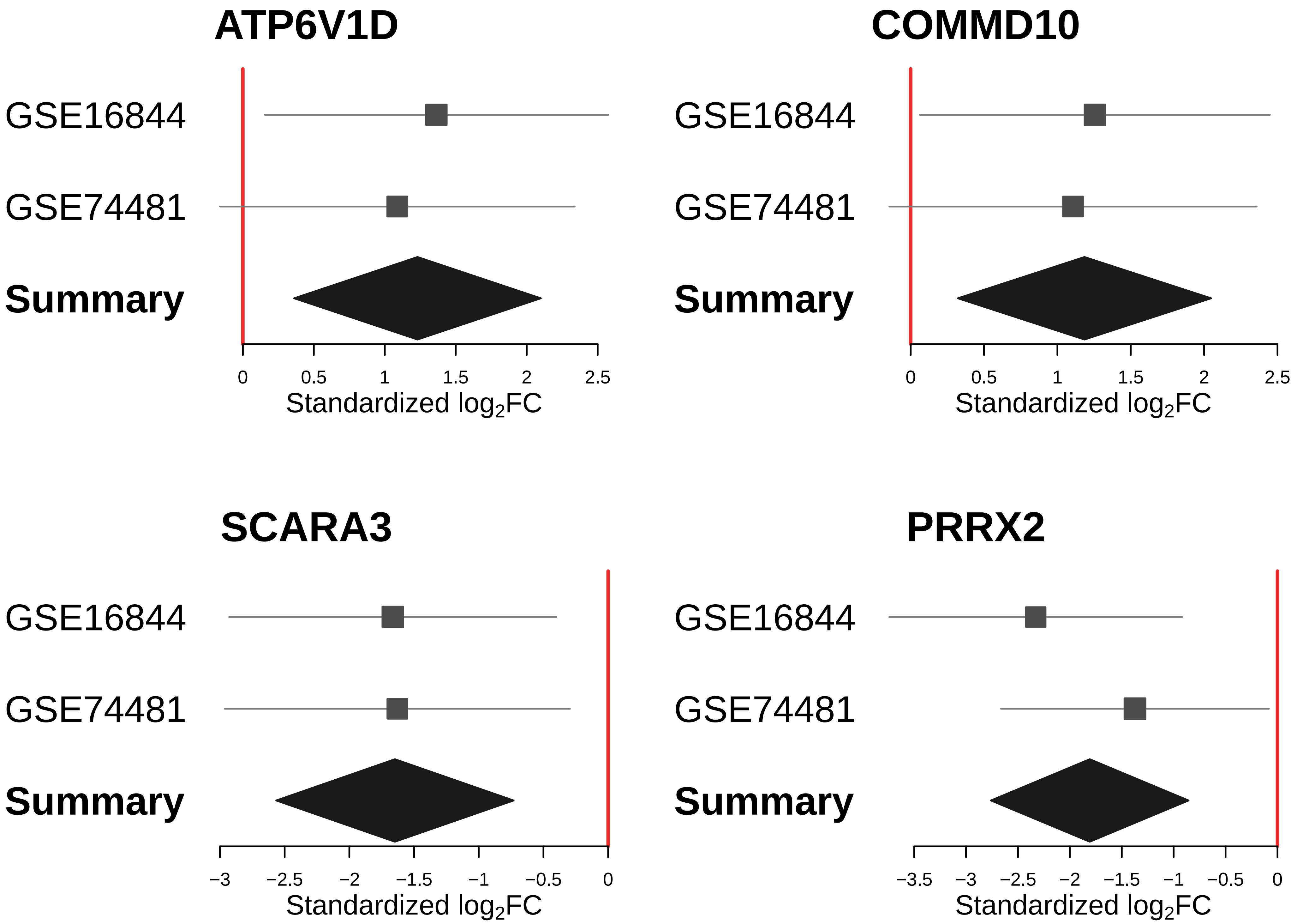

ATP6V1D
GSE16844
GSE74481
Summary
0
0.5
1
1.5
2
2.5
Standardized log2FC
COMMD10
GSE16844
GSE74481
Summary
0
0.5
1
1.5
2
2.5
Standardized log2FC
SCARA3
GSE16844
GSE74481
Summary
−3
−2.5
−2
−1.5
−1
−0.5
0
Standardized log2FC
PRRX2
GSE16844
GSE74481
Summary
−3.5
−3
−2.5
−2
−1.5
−1
−0.5
0
Standardized log2FC

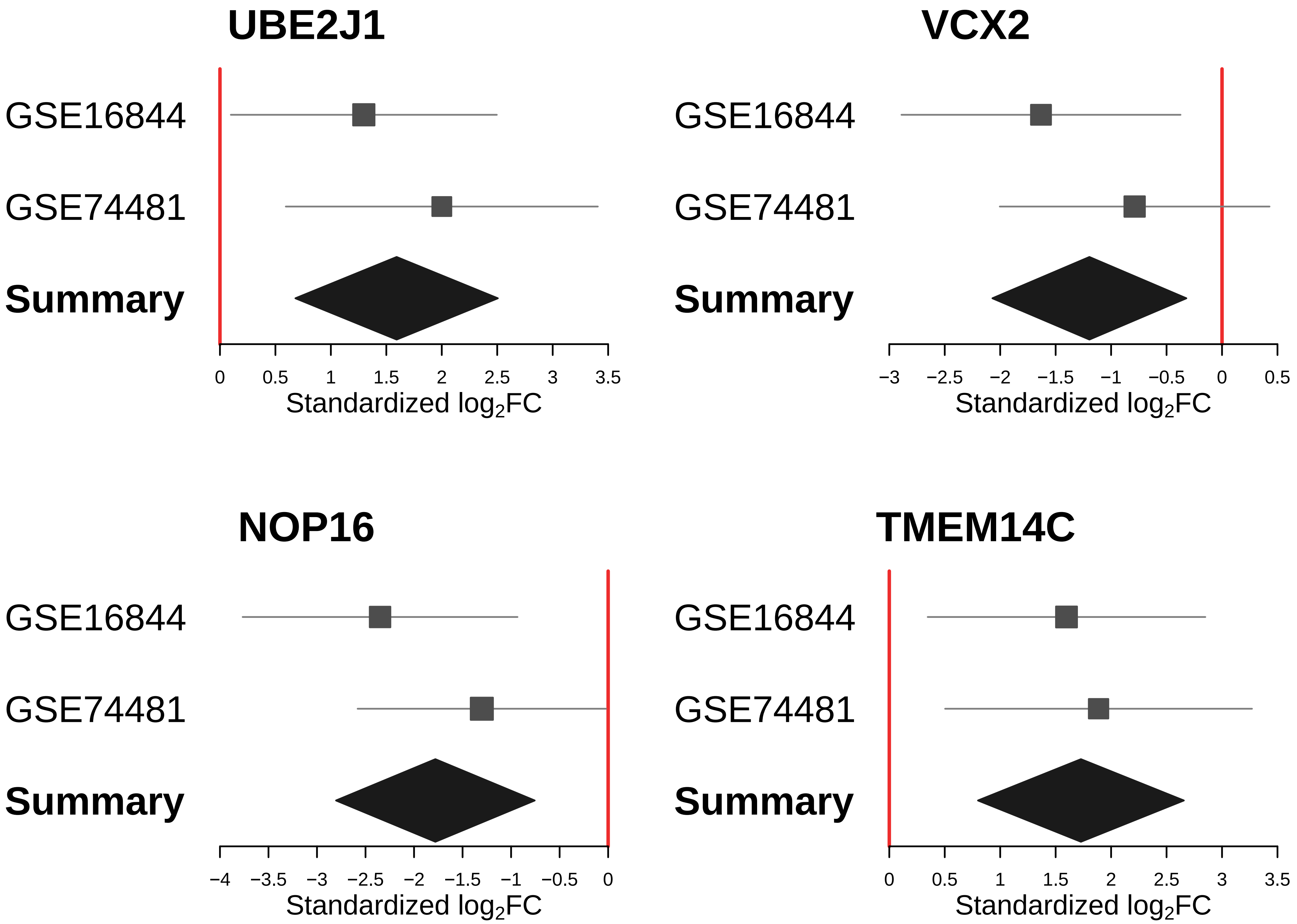
UBE2J1
GSE16844
GSE74481
Summary
0 0.5 1 1.5 2 2.5 3 3.5
Standardized log2FC
VCX2
GSE16844
GSE74481
Summary
−3 −2.5 −2 −1.5 −1 −0.5 0 0.5
Standardized log2FC
NOP16
GSE16844
GSE74481
Summary
−4 −3.5 −3 −2.5 −2 −1.5 −1 −0.5 0
Standardized log2FC
TMEM14C
GSE16844
GSE74481
Summary
0 0.5 1 1.5 2 2.5 3 3.5
Standardized log2FC

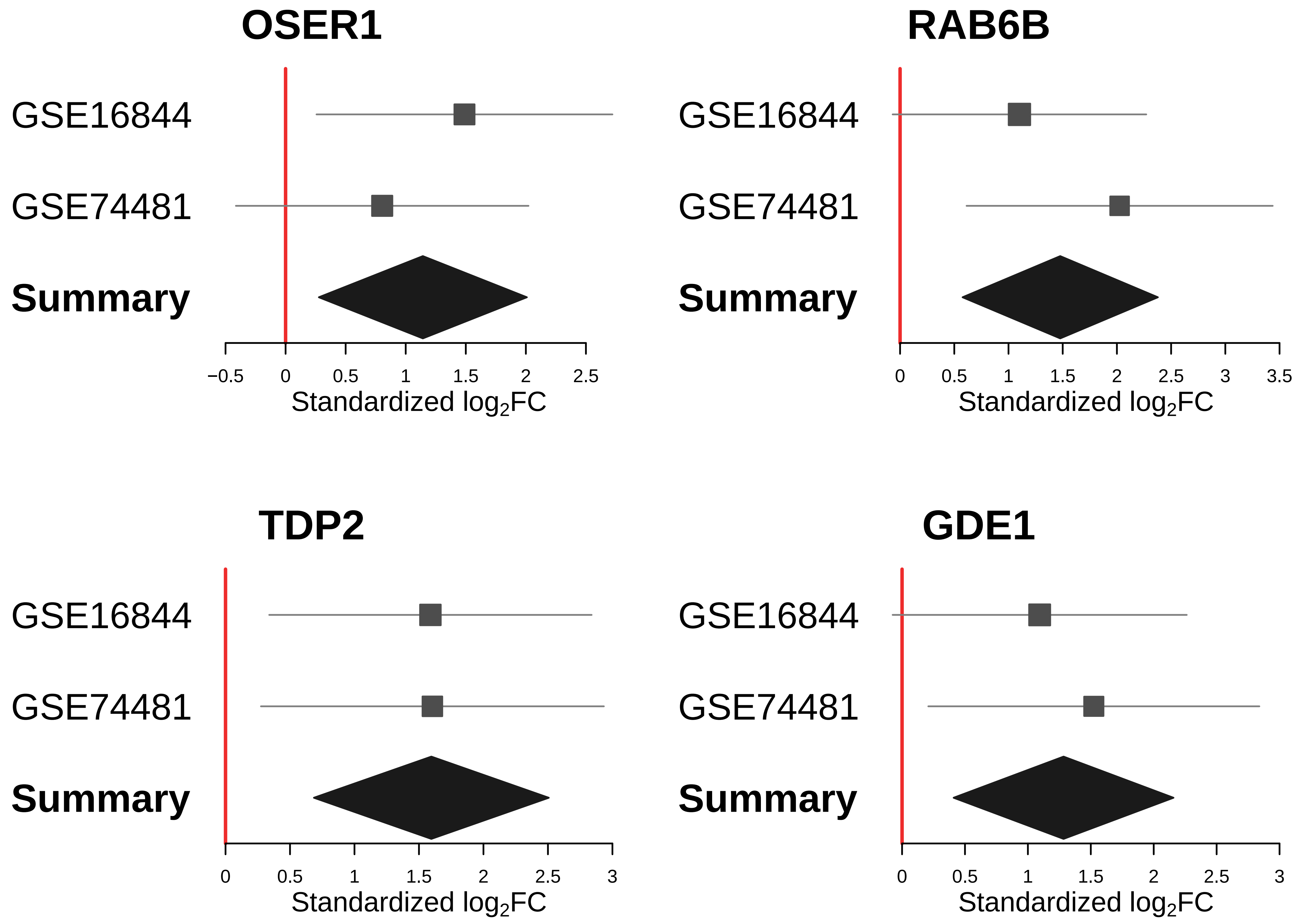

OSER1
GSE16844
GSE74481
Summary
−0.5
0
0.5
1
1.5
2
2.5
Standardized log2FC
RAB6B
GSE16844
GSE74481
Summary
0
0.5
1
1.5
2
2.5
3
3.5
Standardized log2FC
TDP2
GSE16844
GSE74481
Summary
0
0.5
1
1.5
2
2.5
3
Standardized log2FC
GDE1
GSE16844
GSE74481
Summary
0
0.5
1
1.5
2
2.5
3
Standardized log2FC

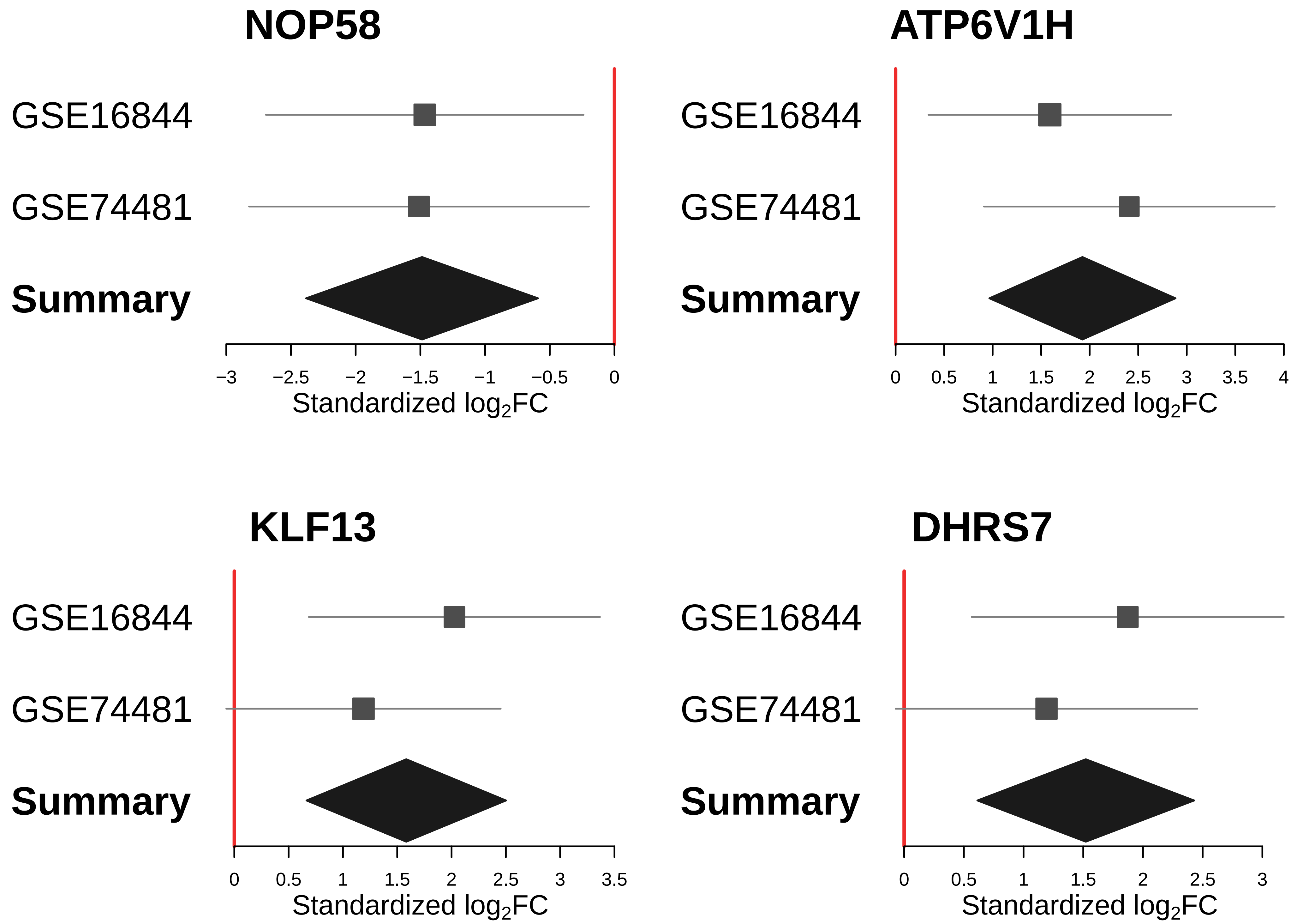

NOP58
GSE16844
GSE74481
Summary
−3
−2.5
−2
−1.5
−1
−0.5
0
Standardized log2FC
ATP6V1H
GSE16844
GSE74481
Summary
0
0.5
1
1.5
2
2.5
3
3.5
4
Standardized log2FC
KLF13
GSE16844
GSE74481
Summary
0
0.5
1
1.5
2
2.5
3
3.5
Standardized log2FC
DHRS7
GSE16844
GSE74481
Summary
0
0.5
1
1.5
2
2.5
3
Standardized log2FC

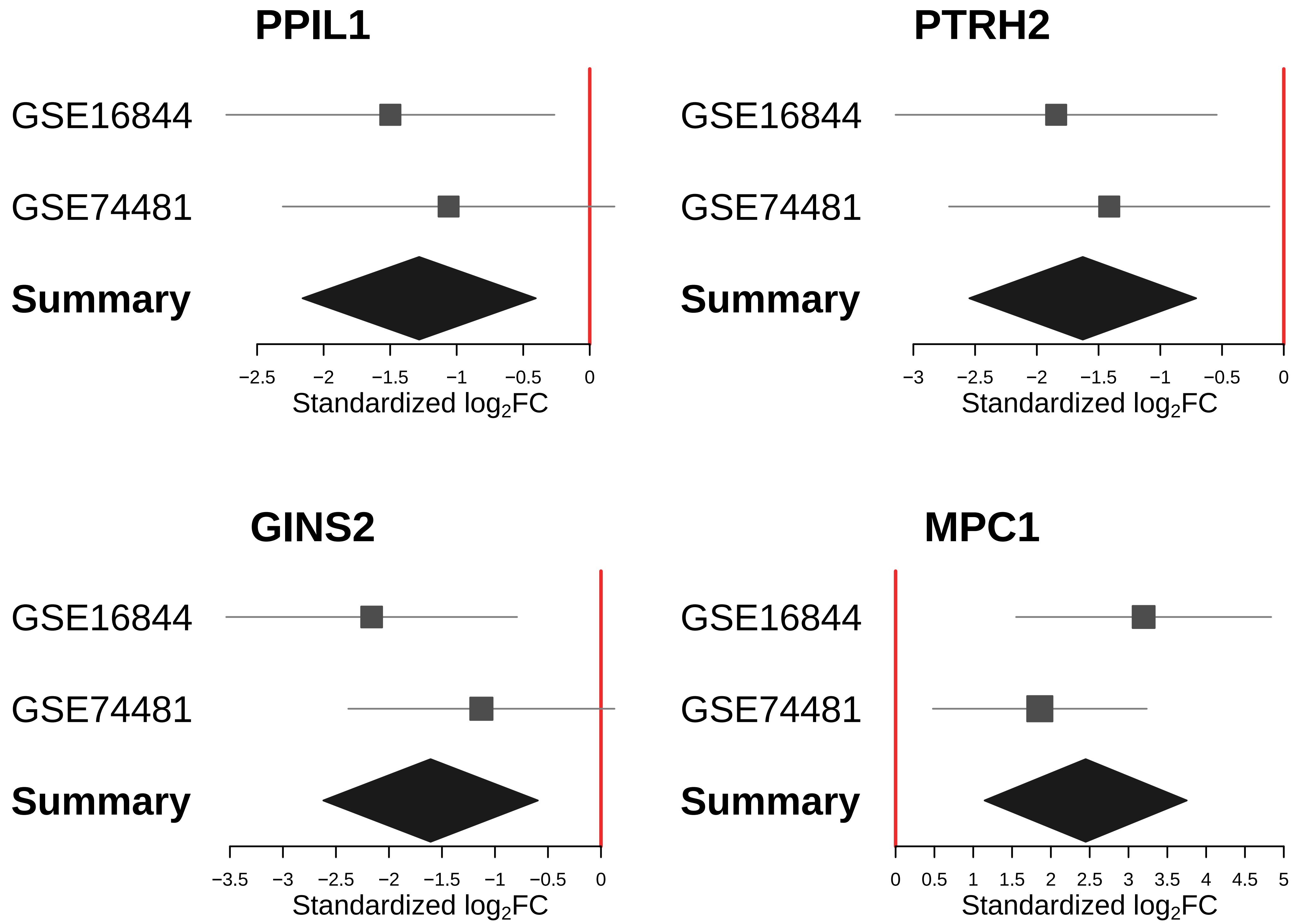
PPIL1
GSE16844
GSE74481
Summary
−2.5 −2 −1.5 −1 −0.5 0
Standardized $log_2FC$
PTRH2
GSE16844
GSE74481
Summary
−3 −2.5 −2 −1.5 −1 −0.5 0
Standardized $log_2FC$
GINS2
GSE16844
GSE74481
Summary
−3.5 −3 −2.5 −2 −1.5 −1 −0.5 0
Standardized $log_2FC$
MPC1
GSE16844
GSE74481
Summary
0 0.5 1 1.5 2 2.5 3 3.5 4 4.5 5
Standardized $log_2FC$

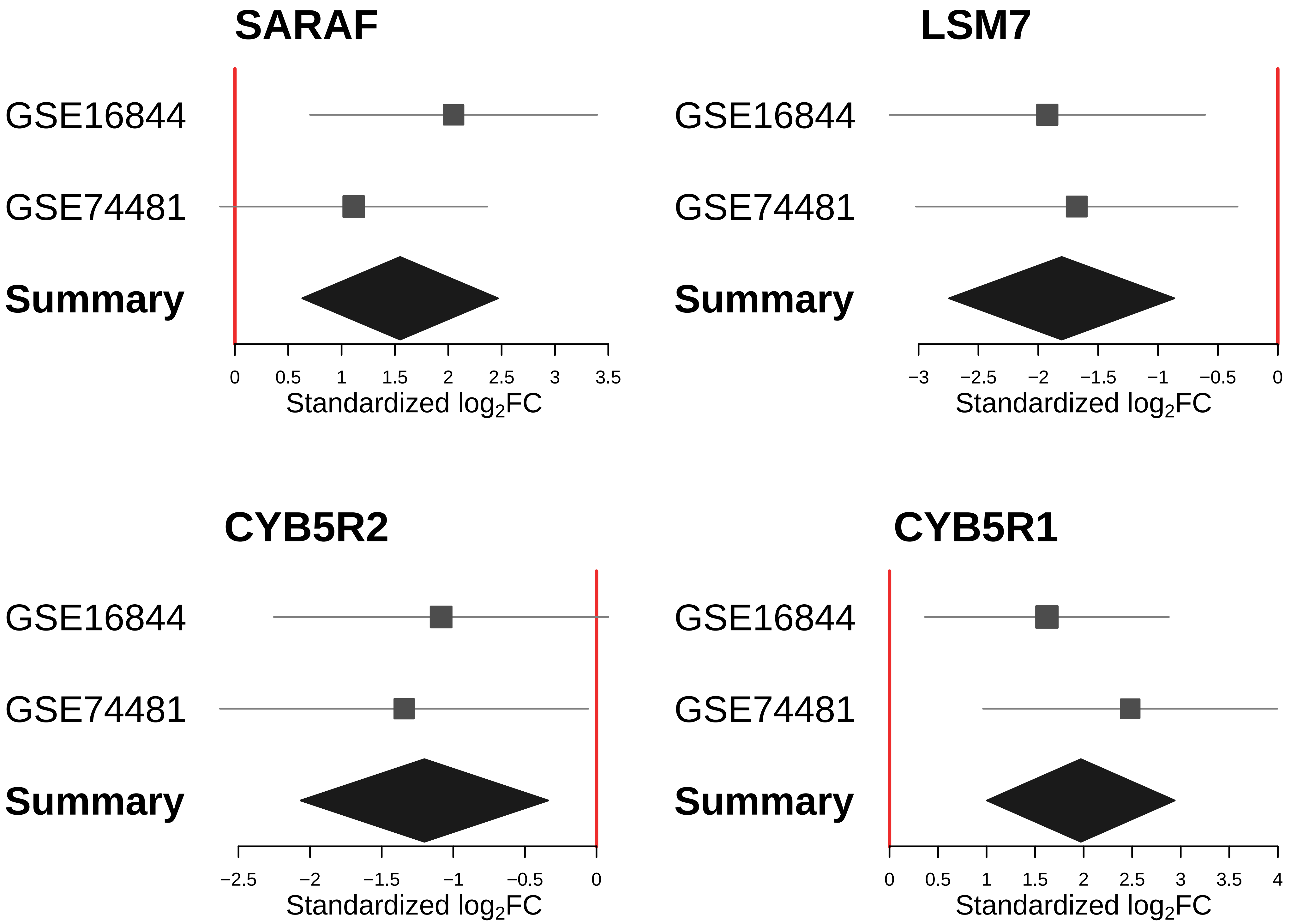
SARAF
GSE16844
GSE74481
Summary
0 0.5 1 1.5 2 2.5 3 3.5
Standardized log2FC
LSM7
GSE16844
GSE74481
Summary
−3 −2.5 −2 −1.5 −1 −0.5 0
Standardized log2FC
CYB5R2
GSE16844
GSE74481
Summary
−2.5 −2 −1.5 −1 −0.5 0
Standardized log2FC
CYB5R1
GSE16844
GSE74481
Summary
0 0.5 1 1.5 2 2.5 3 3.5 4
Standardized log2FC

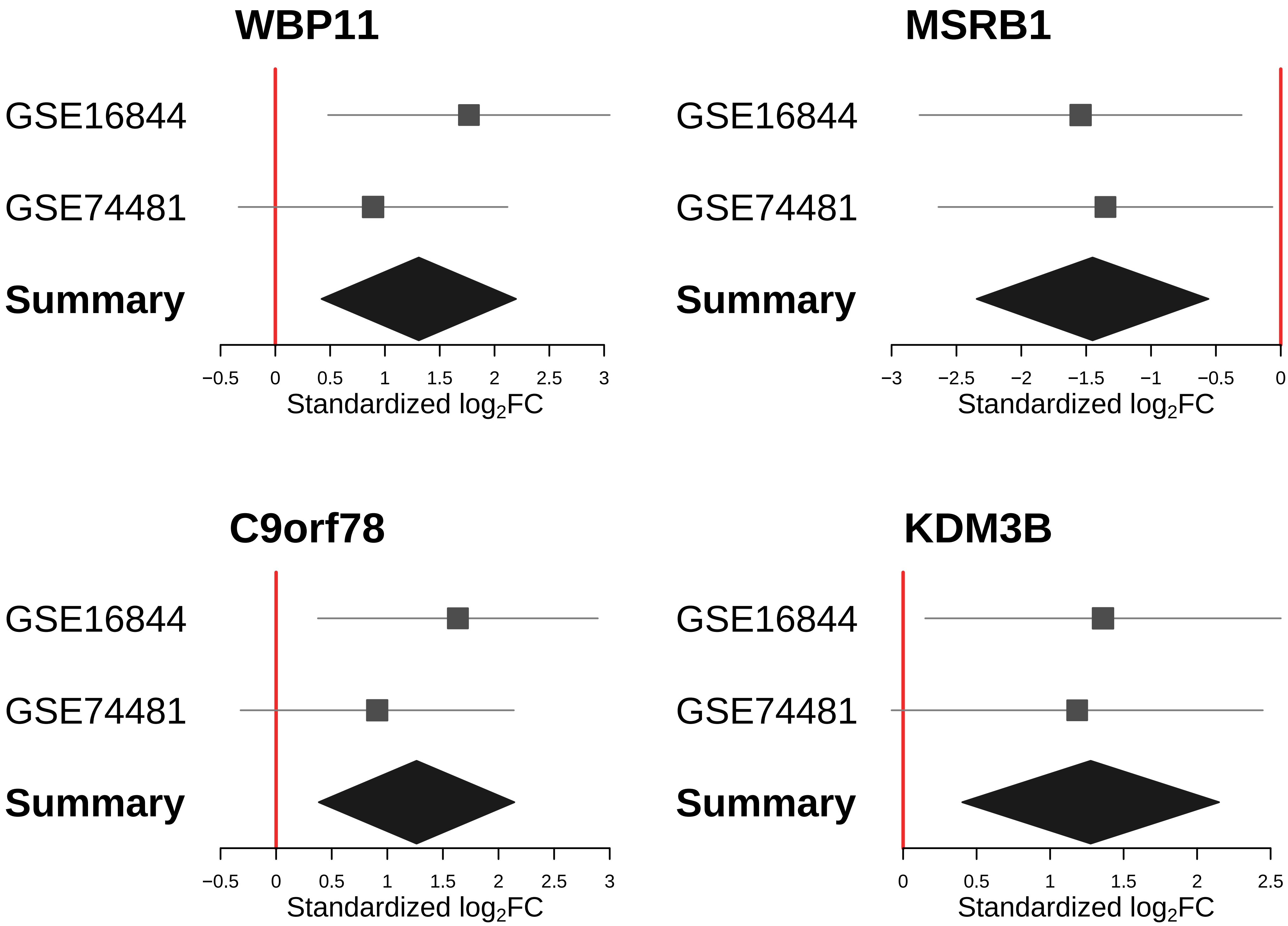
WBP11
GSE16844
GSE74481
Summary
−0.5 0 0.5 1 1.5 2 2.5 3
Standardized log2FC
MSRB1
GSE16844
GSE74481
Summary
−3 −2.5 −2 −1.5 −1 −0.5 0
Standardized log2FC
C9orf78
GSE16844
GSE74481
Summary
−0.5 0 0.5 1 1.5 2 2.5 3
Standardized log2FC
KDM3B
GSE16844
GSE74481
Summary
0 0.5 1 1.5 2 2.5
Standardized log2FC

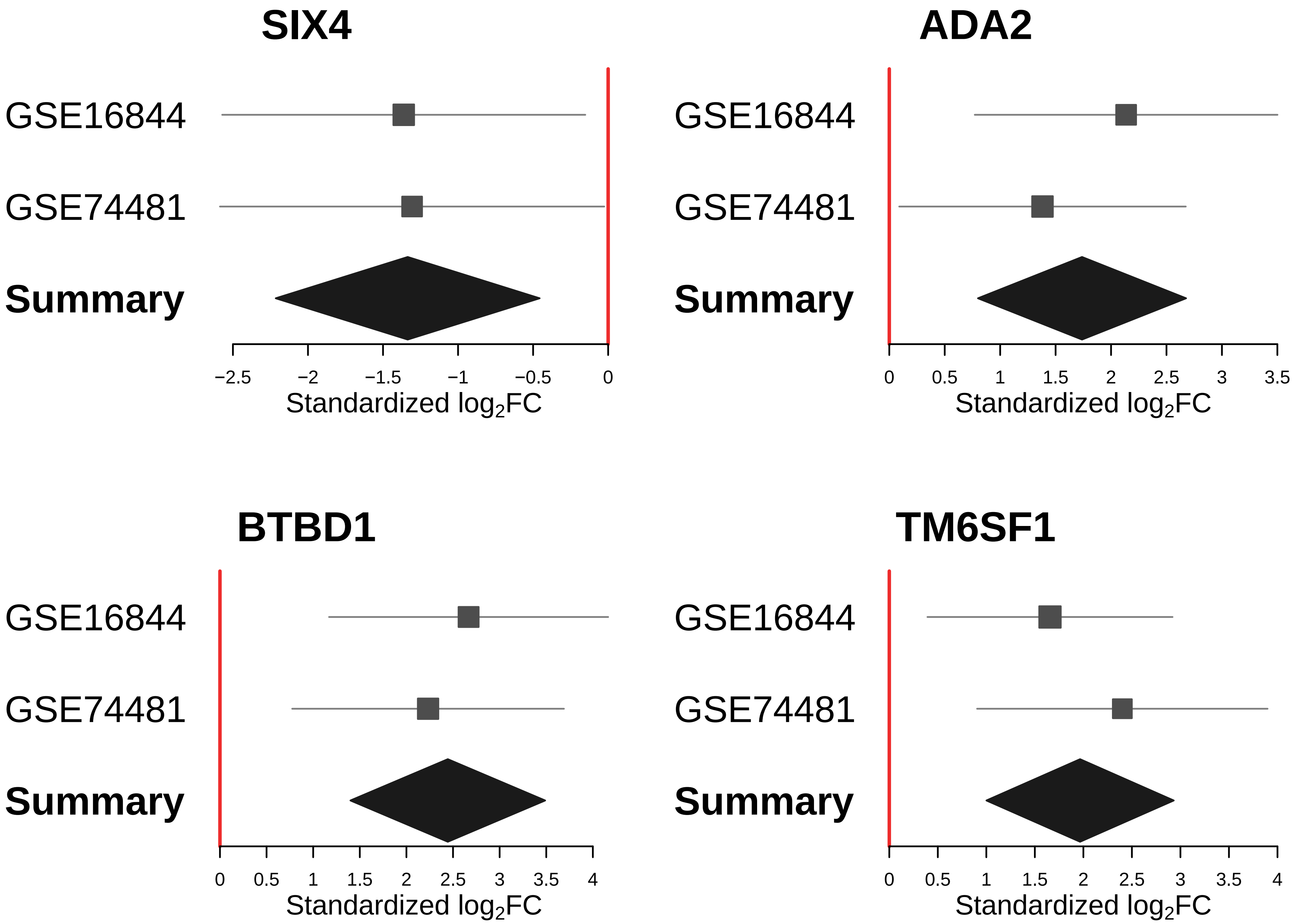

SIX4
GSE16844
GSE74481
Summary
−2.5
−2
−1.5
−1
−0.5
0
Standardized log2FC
ADA2
GSE16844
GSE74481
Summary
0
0.5
1
1.5
2
2.5
3
3.5
Standardized log2FC
BTBD1
GSE16844
GSE74481
Summary
0
0.5
1
1.5
2
2.5
3
3.5
4
Standardized log2FC
TM6SF1
GSE16844
GSE74481
Summary
0
0.5
1
1.5
2
2.5
3
3.5
4
Standardized log2FC

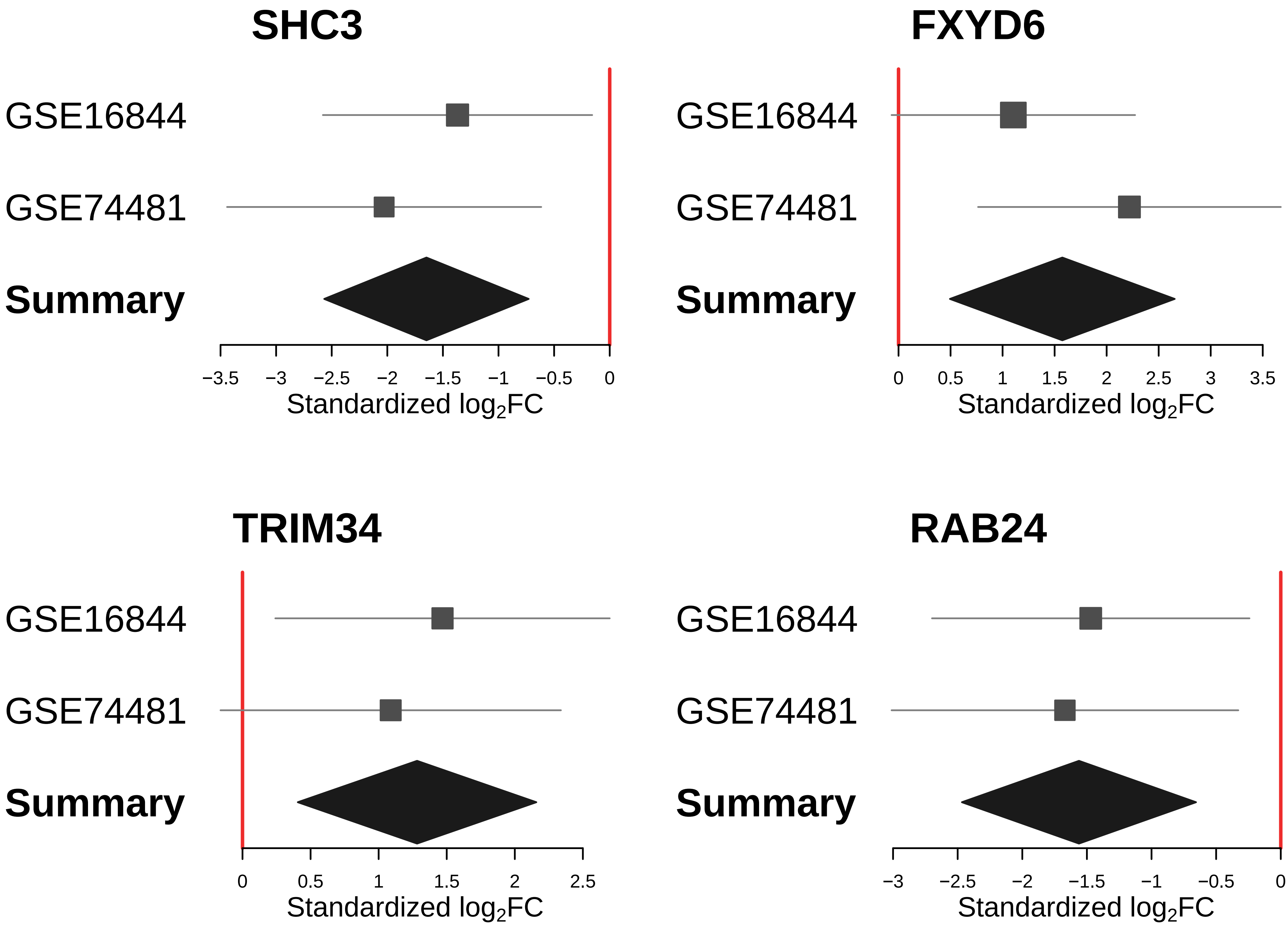
SHC3
GSE16844
GSE74481
Summary
−3.5 −3 −2.5 −2 −1.5 −1 −0.5 0
Standardized log2FC
FXYD6
GSE16844
GSE74481
Summary
0 0.5 1 1.5 2 2.5 3 3.5
Standardized log2FC
TRIM34
GSE16844
GSE74481
Summary
0 0.5 1 1.5 2 2.5
Standardized log2FC
RAB24
GSE16844
GSE74481
Summary
−3 −2.5 −2 −1.5 −1 −0.5 0
Standardized log2FC

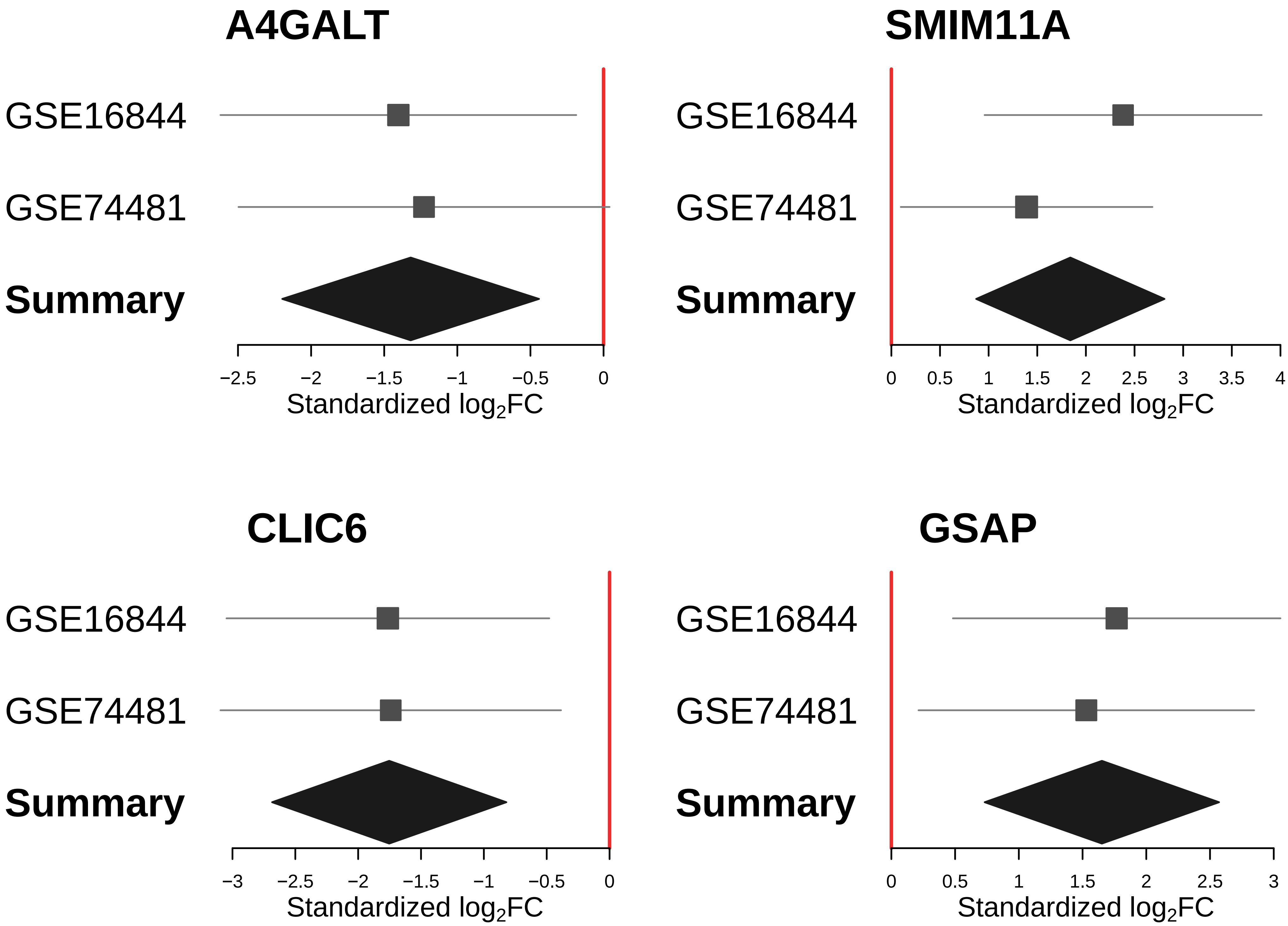

A4GALT
GSE16844
GSE74481
Summary
−2.5 −2 −1.5 −1 −0.5 0
Standardized log2FC
SMIM11A
GSE16844
GSE74481
Summary
0 0.5 1 1.5 2 2.5 3 3.5 4
Standardized log2FC
CLIC6
GSE16844
GSE74481
Summary
−3 −2.5 −2 −1.5 −1 −0.5 0
Standardized log2FC
GSAP
GSE16844
GSE74481
Summary
0 0.5 1 1.5 2 2.5 3
Standardized log2FC

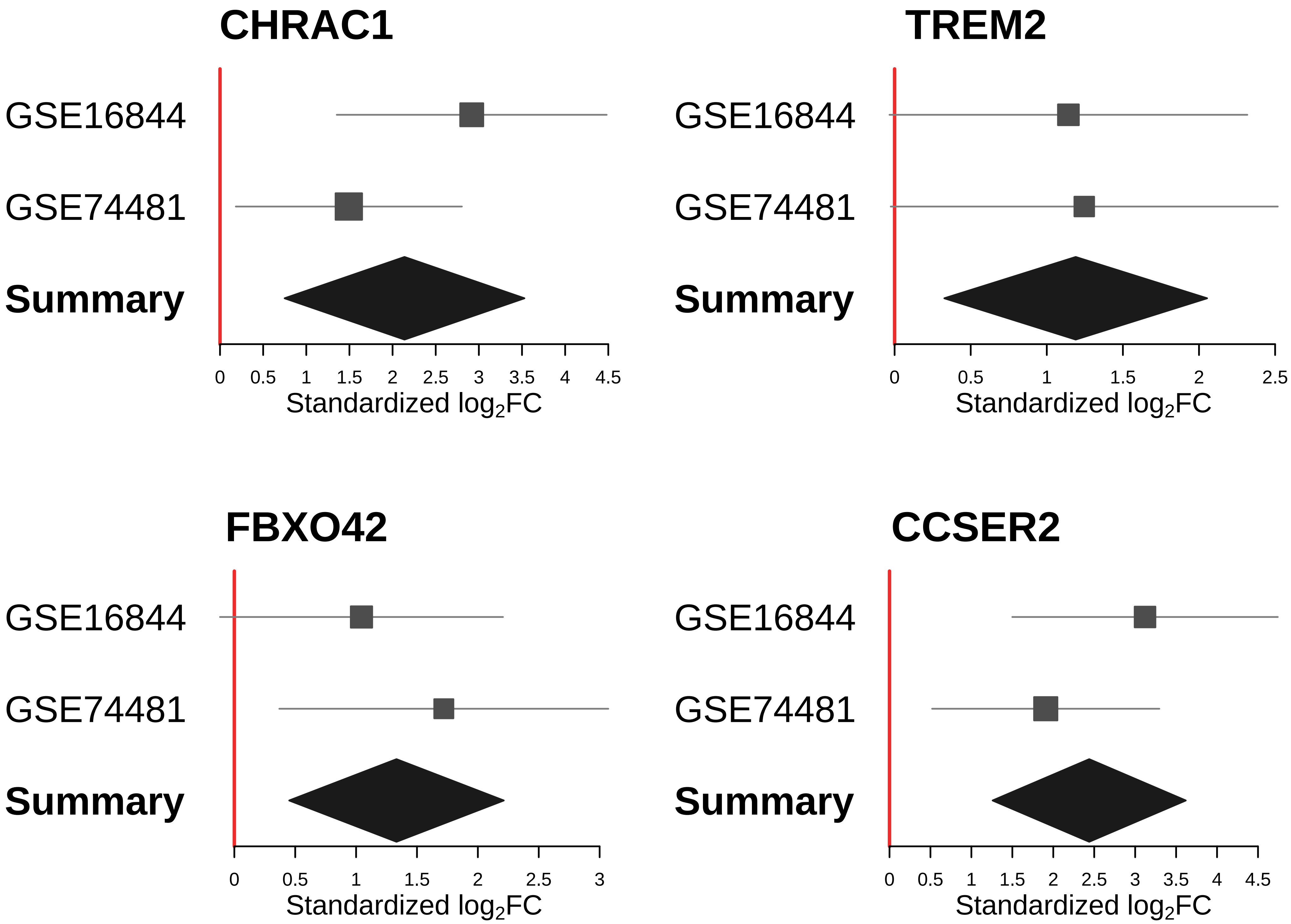

CHRAC1
GSE16844
GSE74481
Summary
0 0.5 1 1.5 2 2.5 3 3.5 4 4.5
Standardized $log_2FC$
TREM2
GSE16844
GSE74481
Summary
0 0.5 1 1.5 2 2.5
Standardized $log_2FC$
FBXO42
GSE16844
GSE74481
Summary
0 0.5 1 1.5 2 2.5 3
Standardized $log_2FC$
CCSER2
GSE16844
GSE74481
Summary
0 0.5 1 1.5 2 2.5 3 3.5 4 4.5
Standardized $log_2FC$

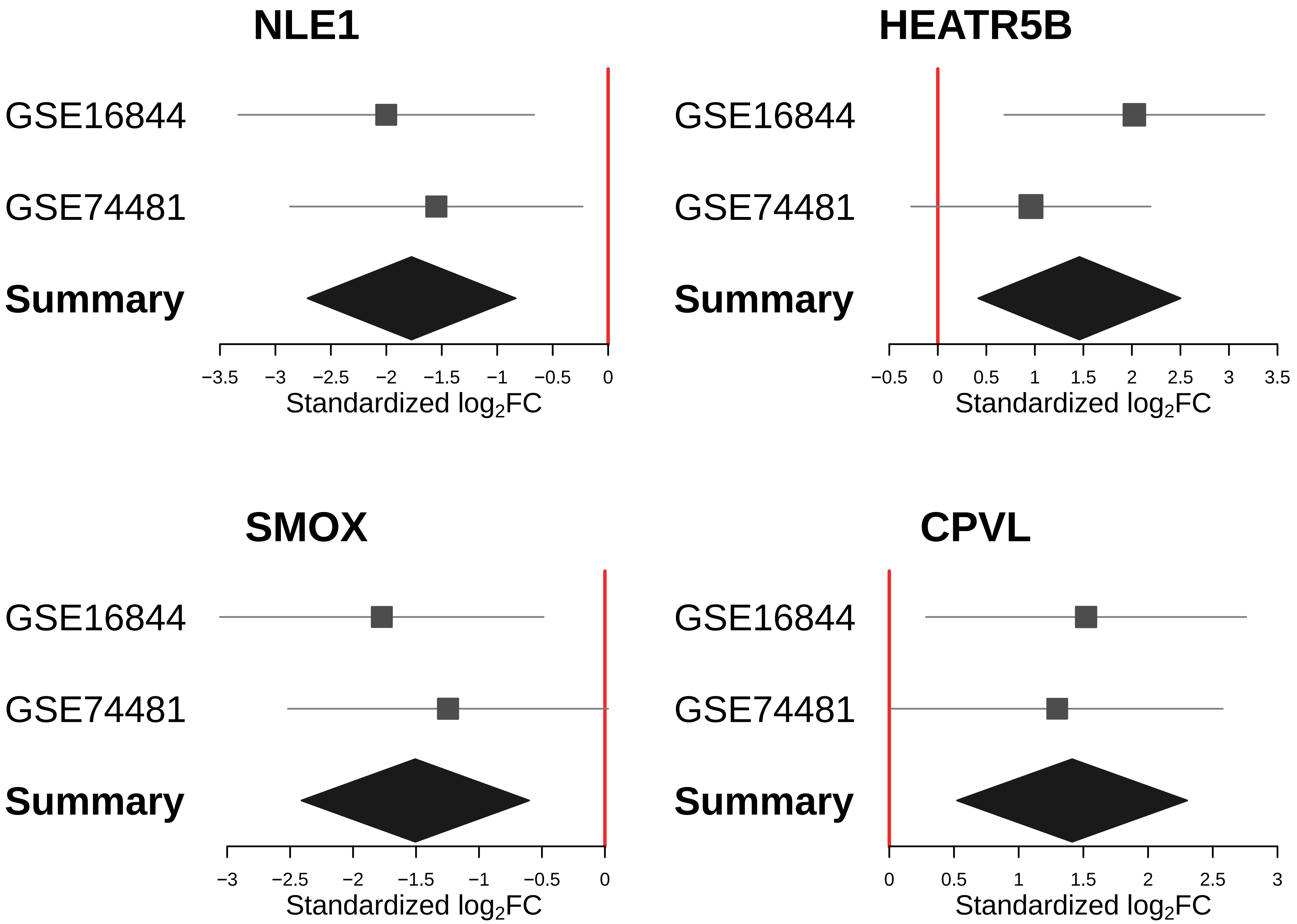
NLE1
GSE16844
GSE74481
Summary
−3.5
−3
−2.5
−2
−1.5
−1
−0.5
0
Standardized log2FC
HEATR5B
GSE16844
GSE74481
Summary
−0.5
0
0.5
1
1.5
2
2.5
3
3.5
Standardized log2FC
SMOX
GSE16844
GSE74481
Summary
−3
−2.5
−2
−1.5
−1
−0.5
0
Standardized log2FC
CPVL
GSE16844
GSE74481
Summary
0
0.5
1
1.5
2
2.5
3
Standardized log2FC

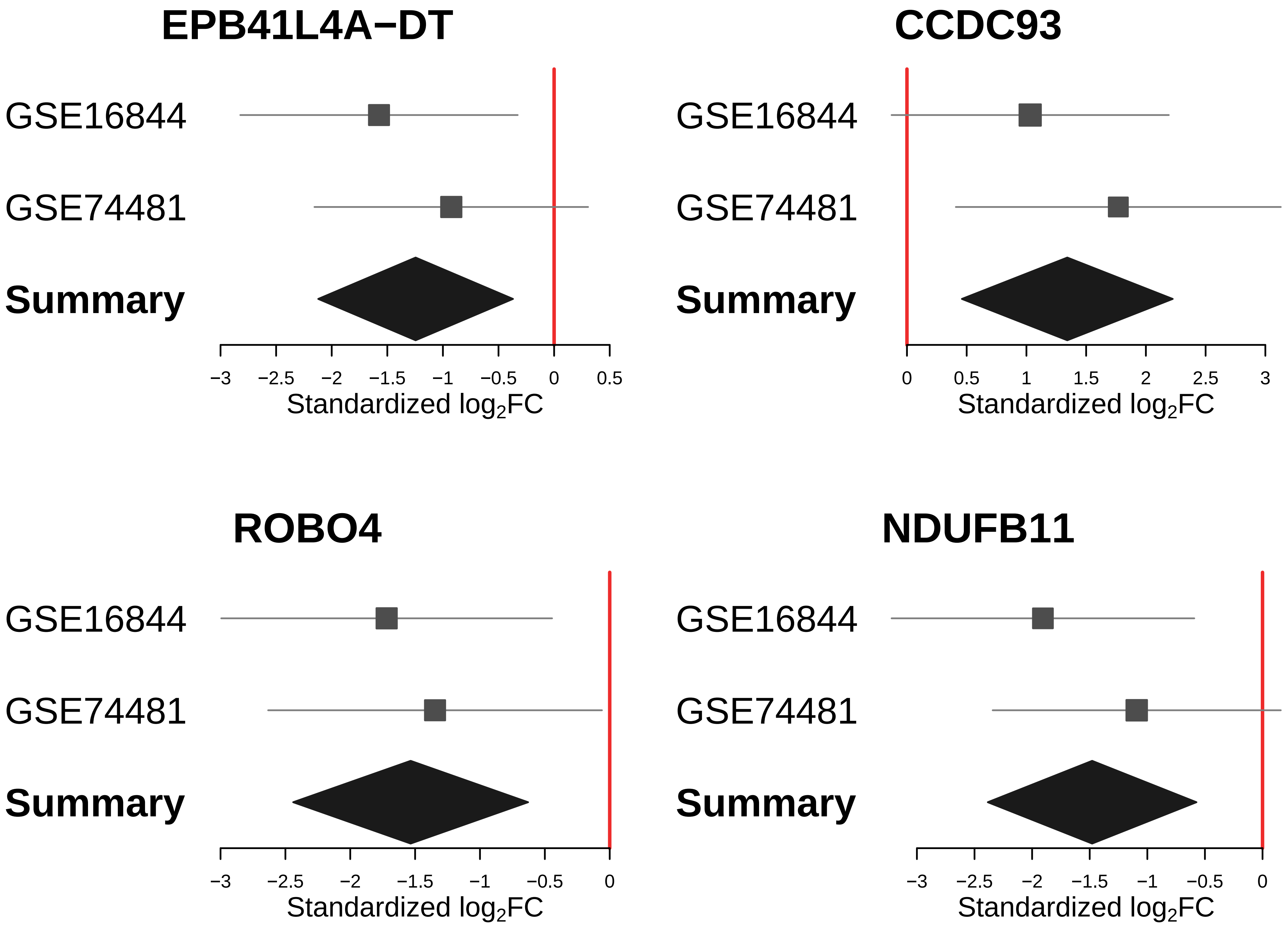
EPB41L4A−DT
GSE16844
GSE74481
Summary
−3
−2.5
−2
−1.5
−1
−0.5
0
0.5
Standardized log2FC
CCDC93
GSE16844
GSE74481
Summary
0
0.5
1
1.5
2
2.5
3
Standardized log2FC
ROBO4
GSE16844
GSE74481
Summary
−3
−2.5
−2
−1.5
−1
−0.5
0
Standardized log2FC
NDUFB11
GSE16844
GSE74481
Summary
−3
−2.5
−2
−1.5
−1
−0.5
0
Standardized log2FC

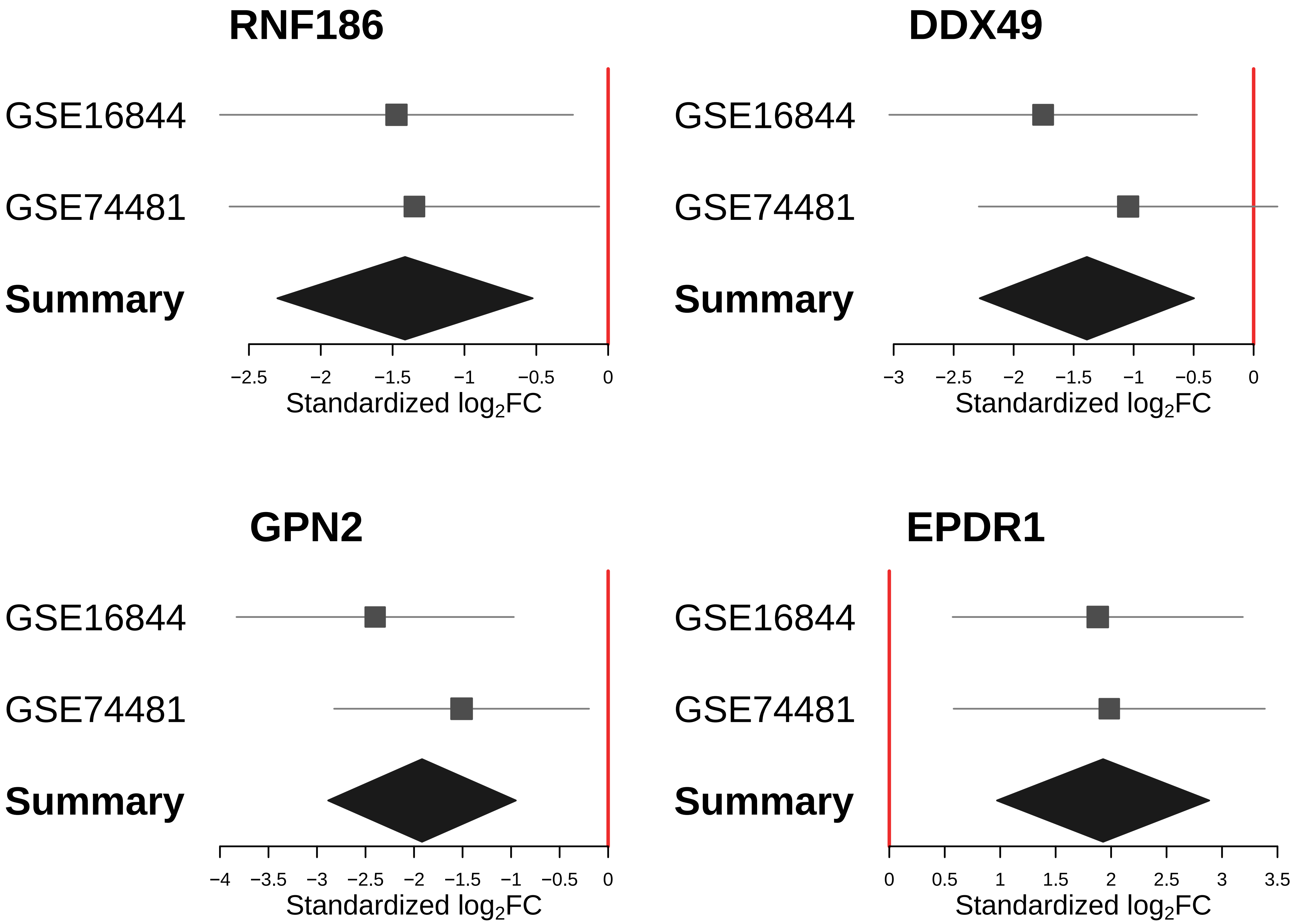
RNF186
GSE16844
GSE74481
Summary
−2.5
−2
−1.5
−1
−0.5
0
Standardized log2FC
DDX49
GSE16844
GSE74481
Summary
−3
−2.5
−2
−1.5
−1
−0.5
0
Standardized log2FC
GPN2
GSE16844
GSE74481
Summary
−4
−3.5
−3
−2.5
−2
−1.5
−1
−0.5
0
Standardized log2FC
EPDR1
GSE16844
GSE74481
Summary
0
0.5
1
1.5
2
2.5
3
3.5
Standardized log2FC

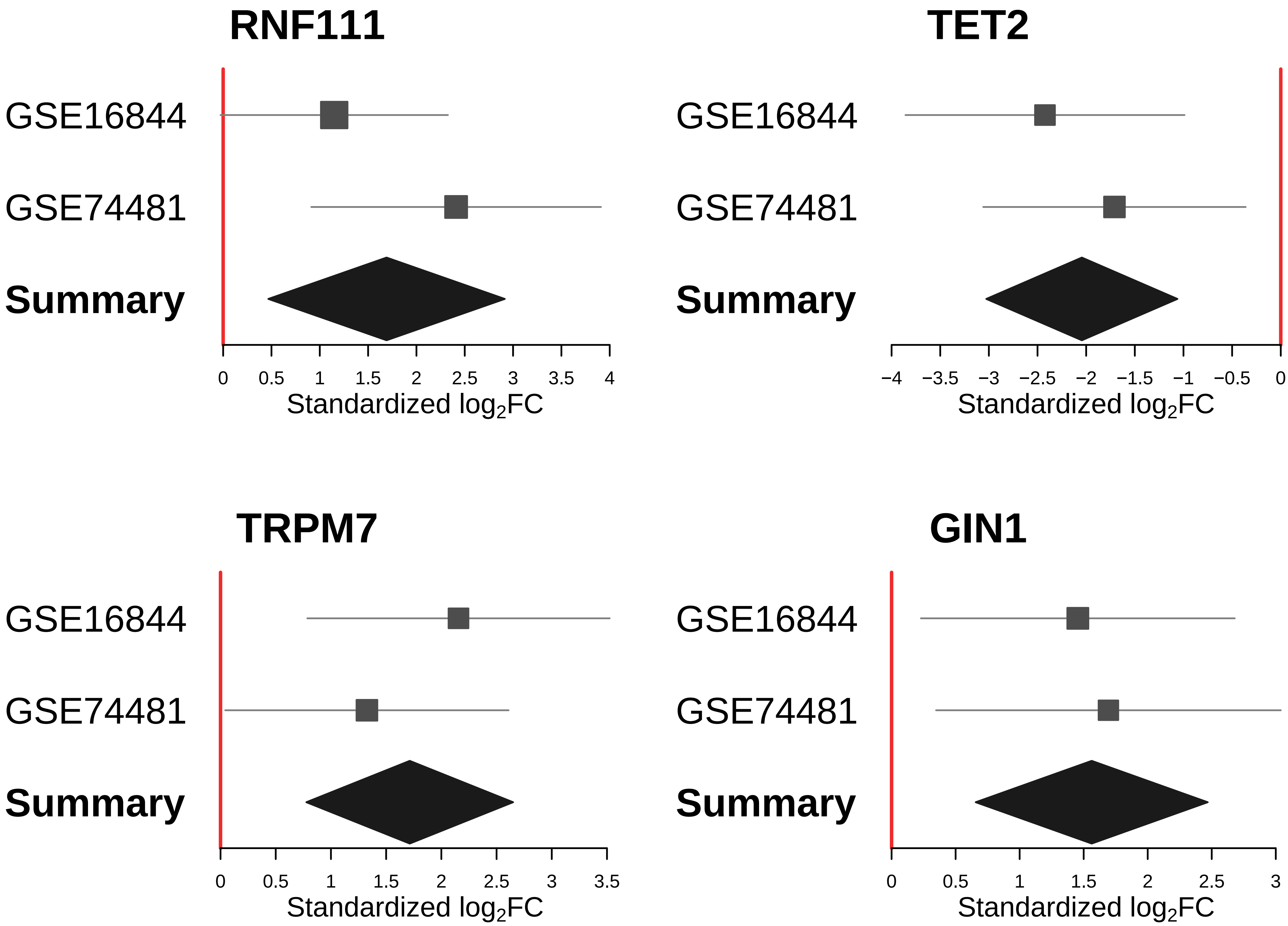
RNF111
GSE16844
GSE74481
Summary
0 0.5 1 1.5 2 2.5 3 3.5 4
Standardized log2FC
TET2
GSE16844
GSE74481
Summary
−4 −3.5 −3 −2.5 −2 −1.5 −1 −0.5 0
Standardized log2FC
TRPM7
GSE16844
GSE74481
Summary
0 0.5 1 1.5 2 2.5 3 3.5
Standardized log2FC
GIN1
GSE16844
GSE74481
Summary
0 0.5 1 1.5 2 2.5 3
Standardized log2FC

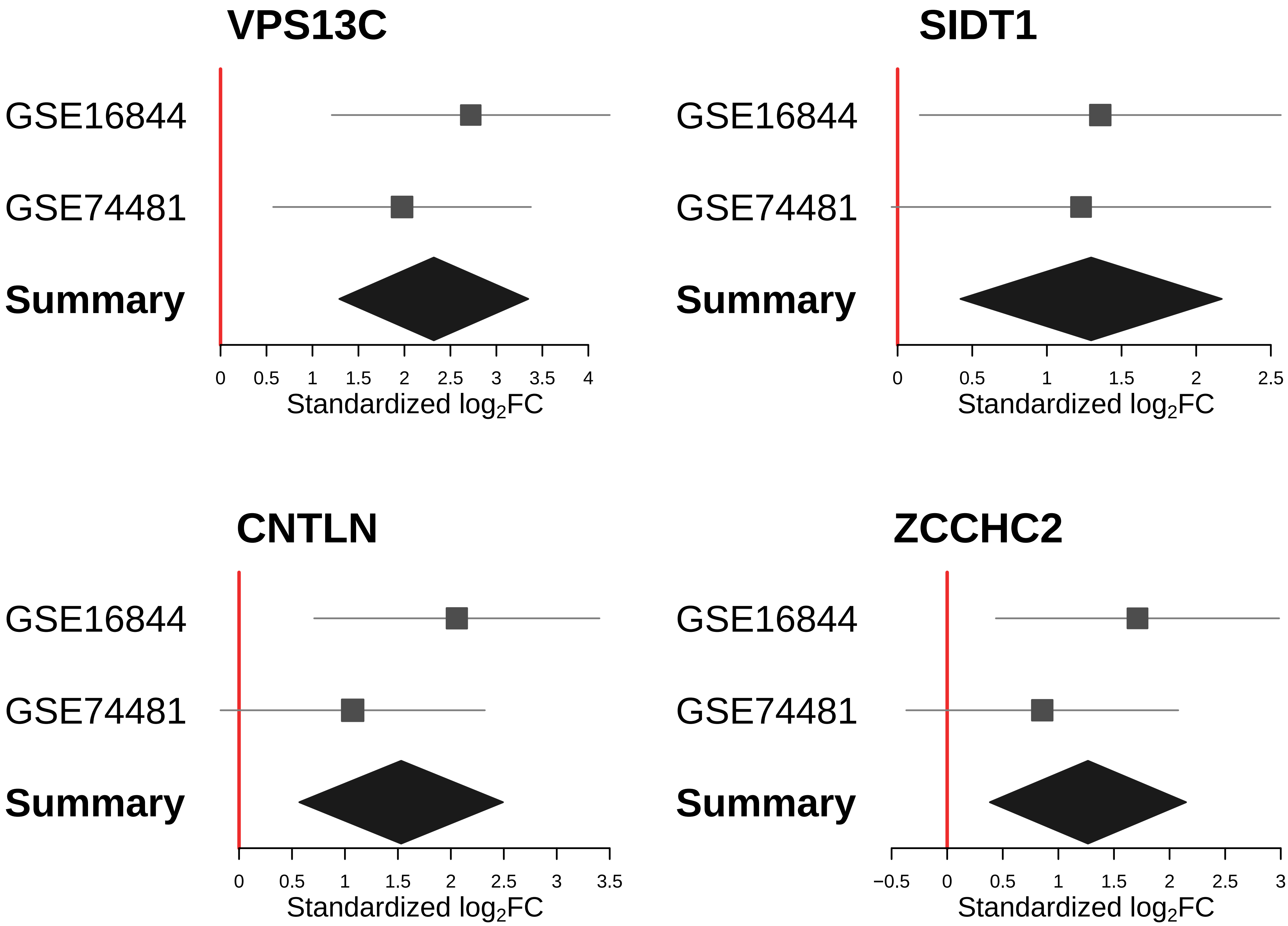
VPS13C
GSE16844
GSE74481
Summary
0 0.5 1 1.5 2 2.5 3 3.5 4
Standardized log2FC
SIDT1
GSE16844
GSE74481
Summary
0 0.5 1 1.5 2 2.5
Standardized log2FC
CNTLN
GSE16844
GSE74481
Summary
0 0.5 1 1.5 2 2.5 3 3.5
Standardized log2FC
ZCCHC2
GSE16844
GSE74481
Summary
−0.5 0 0.5 1 1.5 2 2.5 3
Standardized log2FC

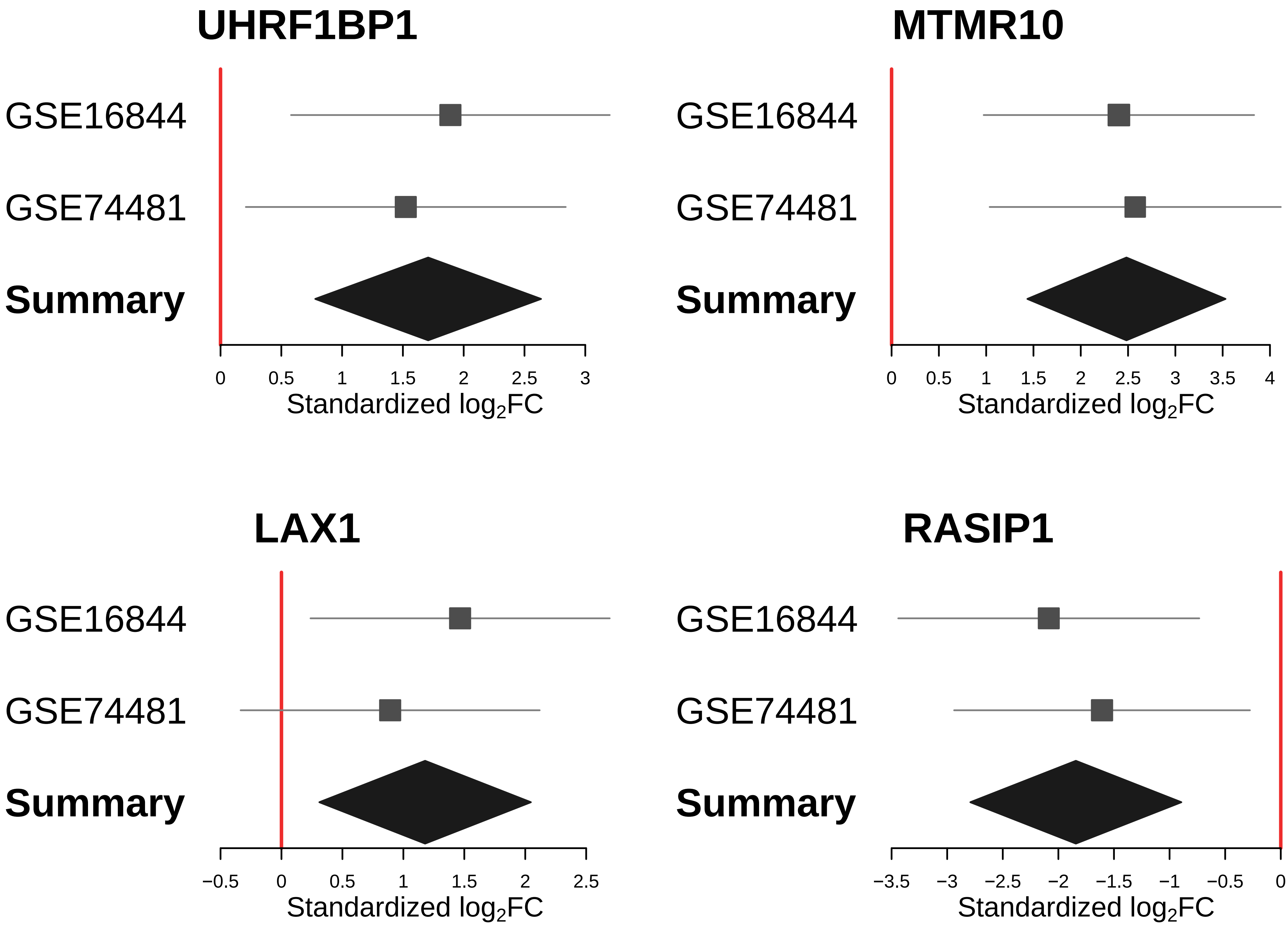
UHRF1BP1
GSE16844
GSE74481
Summary
0
0.5
1
1.5
2
2.5
3
Standardized log2FC
MTMR10
GSE16844
GSE74481
Summary
0
0.5
1
1.5
2
2.5
3
3.5
4
Standardized log2FC
LAX1
GSE16844
GSE74481
Summary
−0.5
0
0.5
1
1.5
2
2.5
Standardized log2FC
RASIP1
GSE16844
GSE74481
Summary
−3.5
−3
−2.5
−2
−1.5
−1
−0.5
0
Standardized log2FC

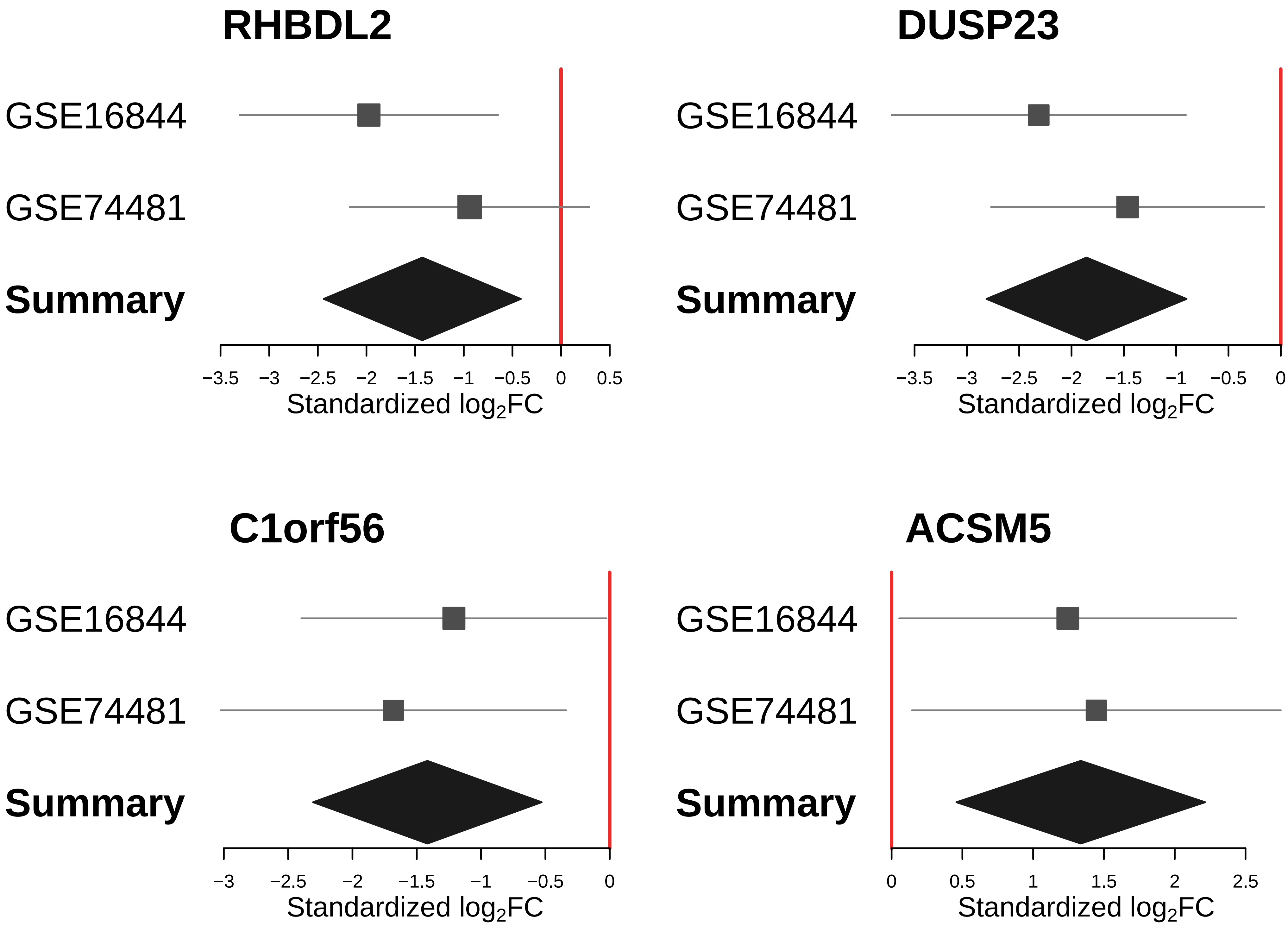
RHBDL2
GSE16844
GSE74481
Summary
−3.5 −3 −2.5 −2 −1.5 −1 −0.5 0 0.5
Standardized log2FC
DUSP23
GSE16844
GSE74481
Summary
−3.5 −3 −2.5 −2 −1.5 −1 −0.5 0
Standardized log2FC
C1orf56
GSE16844
GSE74481
Summary
−3 −2.5 −2 −1.5 −1 −0.5 0
Standardized log2FC
ACSM5
GSE16844
GSE74481
Summary
0 0.5 1 1.5 2 2.5
Standardized log2FC

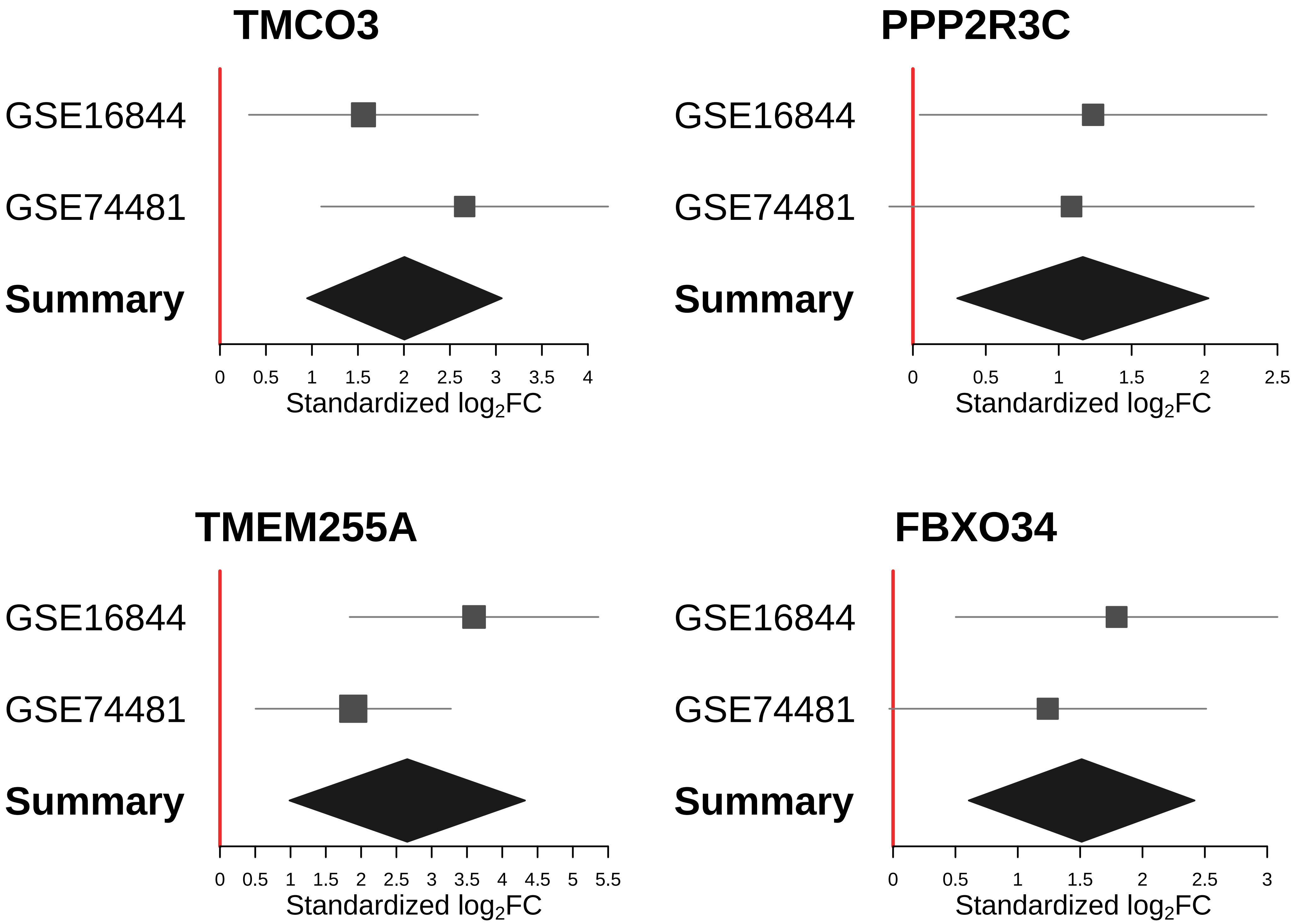
TMCO3
GSE16844
GSE74481
Summary
0 0.5 1 1.5 2 2.5 3 3.5 4
Standardized log2FC
PPP2R3C
GSE16844
GSE74481
Summary
0 0.5 1 1.5 2 2.5
Standardized log2FC
TMEM255A
GSE16844
GSE74481
Summary
0 0.5 1 1.5 2 2.5 3 3.5 4 4.5 5 5.5
Standardized log2FC
FBXO34
GSE16844
GSE74481
Summary
0 0.5 1 1.5 2 2.5 3
Standardized log2FC

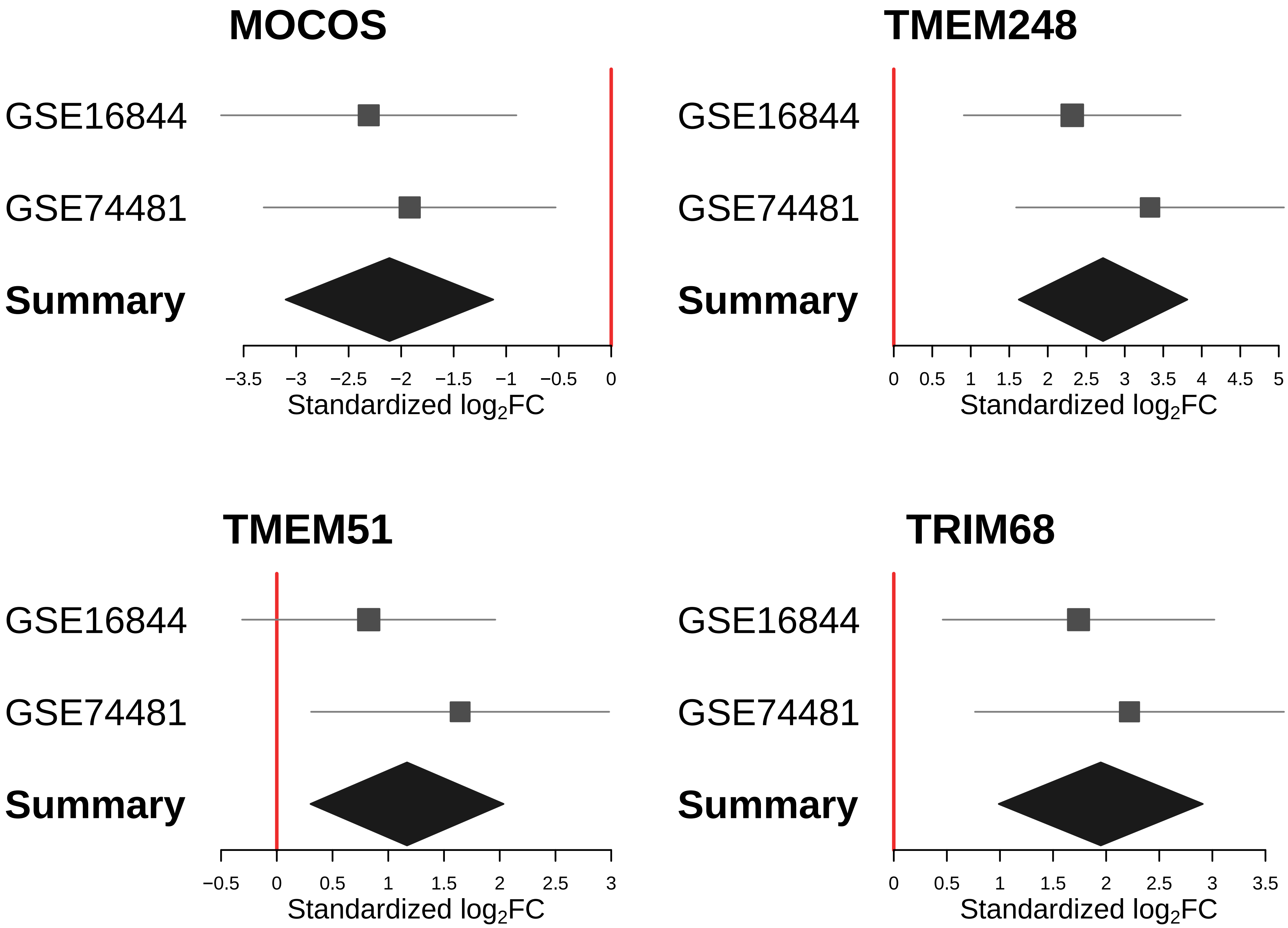

MOCOS
GSE16844
GSE74481
Summary
−3.5 −3 −2.5 −2 −1.5 −1 −0.5 0
Standardized log2FC
TMEM248
GSE16844
GSE74481
Summary
0 0.5 1 1.5 2 2.5 3 3.5 4 4.5 5
Standardized log2FC
TMEM51
GSE16844
GSE74481
Summary
−0.5 0 0.5 1 1.5 2 2.5 3
Standardized log2FC
TRIM68
GSE16844
GSE74481
Summary
0 0.5 1 1.5 2 2.5 3 3.5
Standardized log2FC

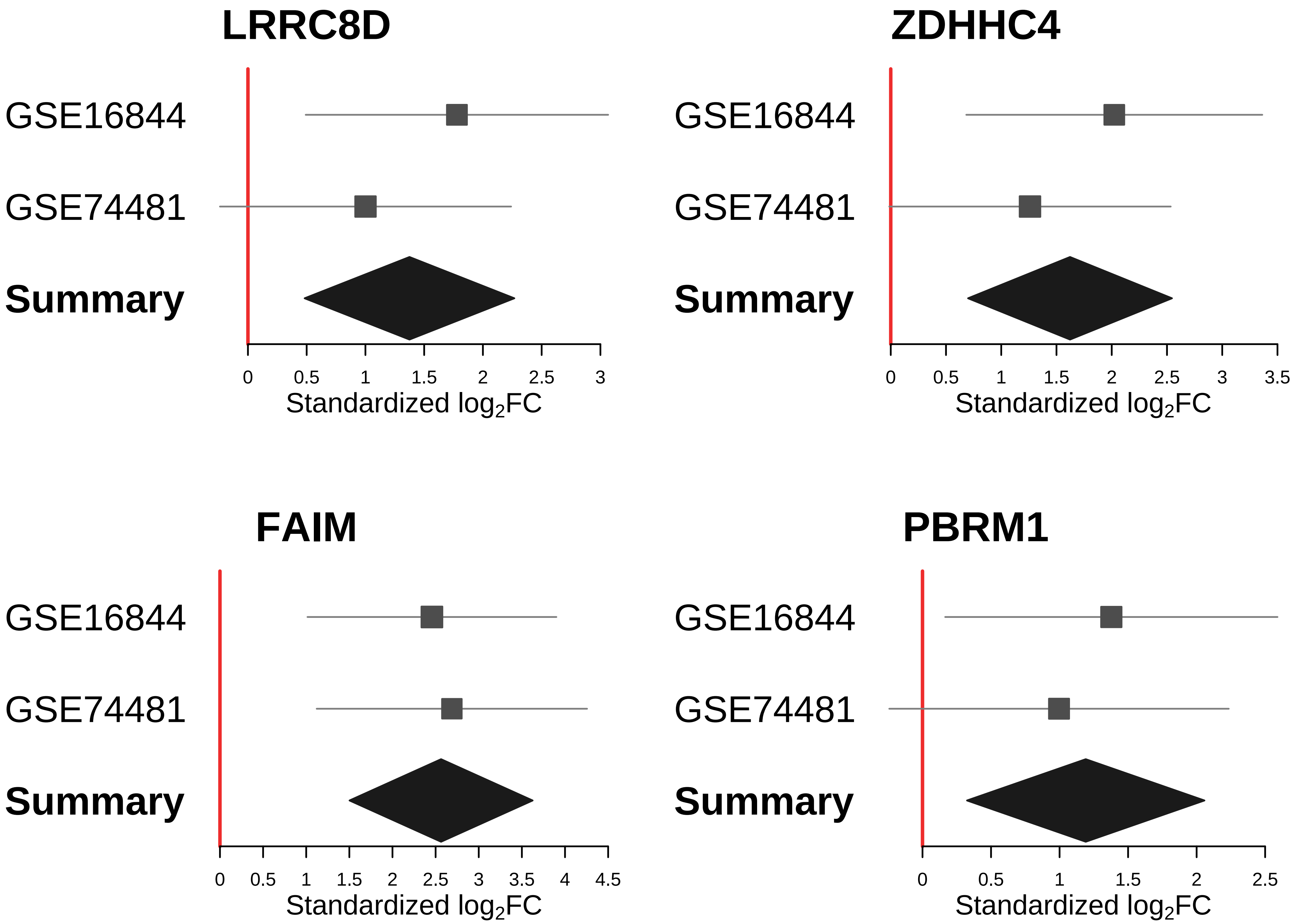
LRRC8D
GSE16844
GSE74481
Summary
0 0.5 1 1.5 2 2.5 3
Standardized log2FC
ZDHHC4
GSE16844
GSE74481
Summary
0 0.5 1 1.5 2 2.5 3 3.5
Standardized log2FC
FAIM
GSE16844
GSE74481
Summary
0 0.5 1 1.5 2 2.5 3 3.5 4 4.5
Standardized log2FC
PBRM1
GSE16844
GSE74481
Summary
0 0.5 1 1.5 2 2.5
Standardized log2FC

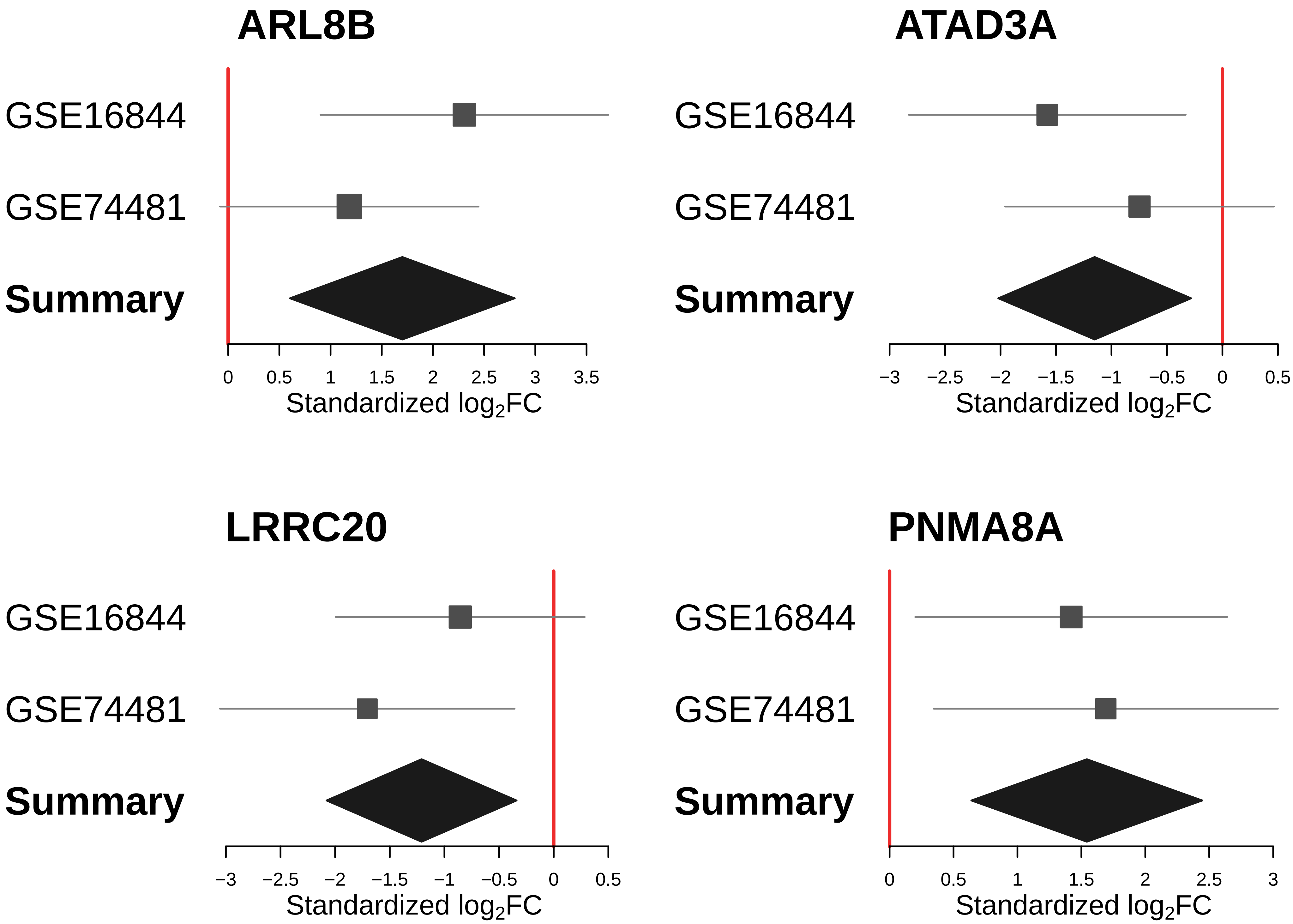
ARL8B
GSE16844
GSE74481
Summary
0
0.5
1
1.5
2
2.5
3
3.5
Standardized log2FC
ATAD3A
GSE16844
GSE74481
Summary
−3
−2.5
−2
−1.5
−1
−0.5
0
0.5
Standardized log2FC
LRRC20
GSE16844
GSE74481
Summary
−3
−2.5
−2
−1.5
−1
−0.5
0
0.5
Standardized log2FC
PNMA8A
GSE16844
GSE74481
Summary
0
0.5
1
1.5
2
2.5
3
Standardized log2FC

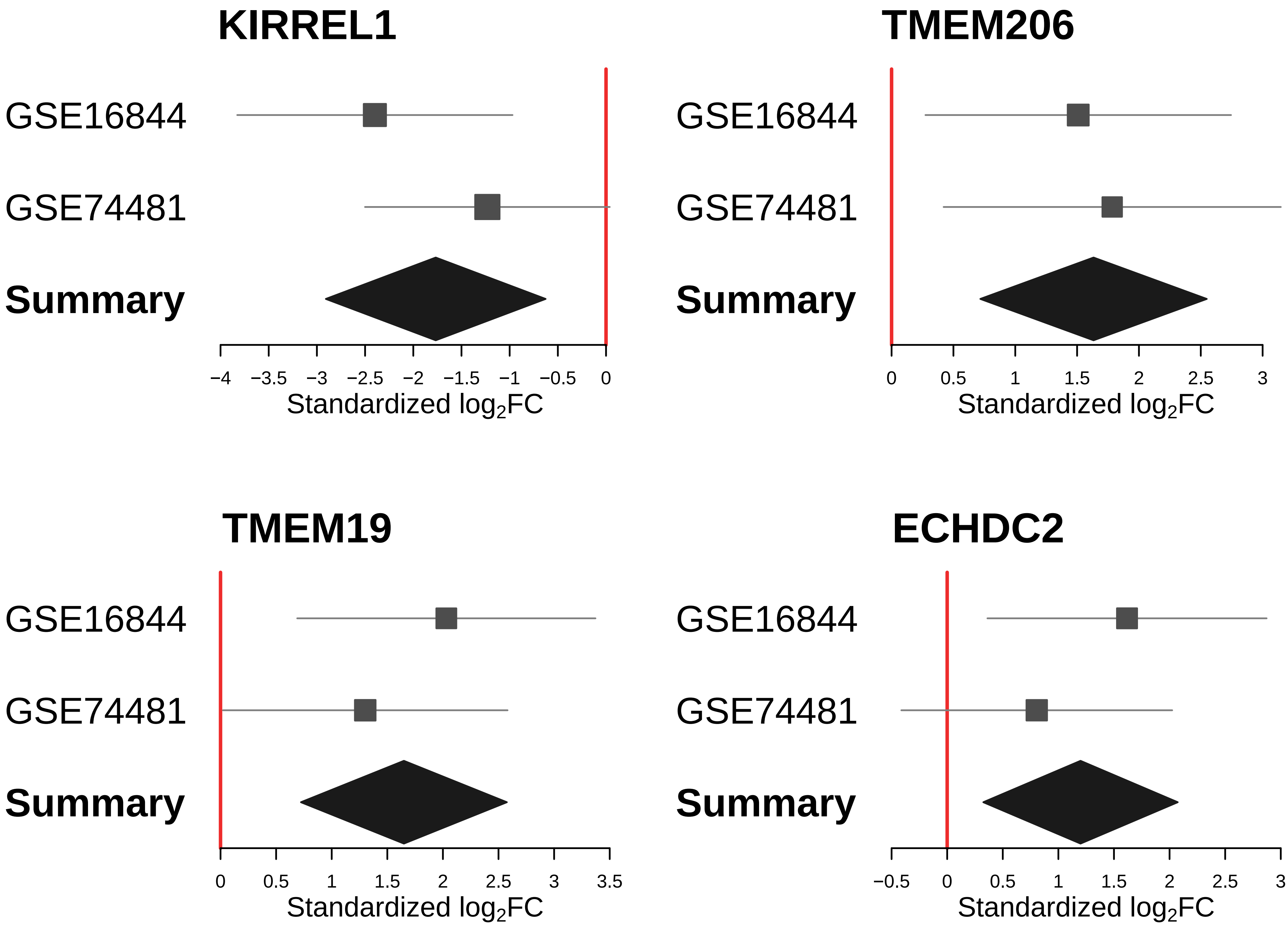
KIRREL1
GSE16844
GSE74481
Summary
−4 −3.5 −3 −2.5 −2 −1.5 −1 −0.5 0
Standardized log2FC
TMEM206
GSE16844
GSE74481
Summary
0 0.5 1 1.5 2 2.5 3
Standardized log2FC
TMEM19
GSE16844
GSE74481
Summary
0 0.5 1 1.5 2 2.5 3 3.5
Standardized log2FC
ECHDC2
GSE16844
GSE74481
Summary
−0.5 0 0.5 1 1.5 2 2.5 3
Standardized log2FC

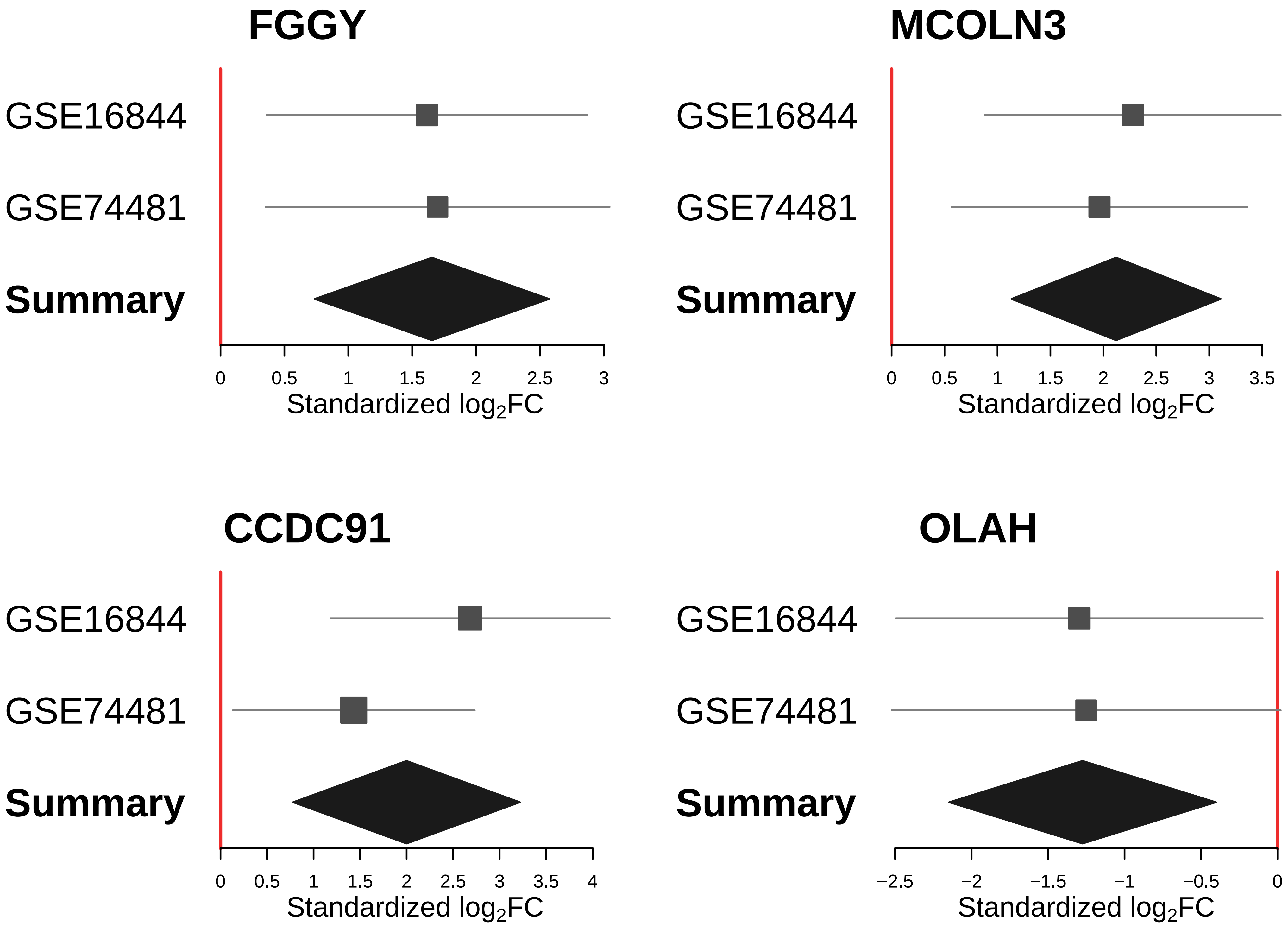
FGGY
GSE16844
GSE74481
Summary
0
0.5
1
1.5
2
2.5
3
Standardized $log_2FC$
MCOLN3
GSE16844
GSE74481
Summary
0
0.5
1
1.5
2
2.5
3
3.5
Standardized $log_2FC$
CCDC91
GSE16844
GSE74481
Summary
0
0.5
1
1.5
2
2.5
3
3.5
4
Standardized $log_2FC$
OLAH
GSE16844
GSE74481
Summary
−2.5
−2
−1.5
−1
−0.5
0
Standardized $log_2FC$

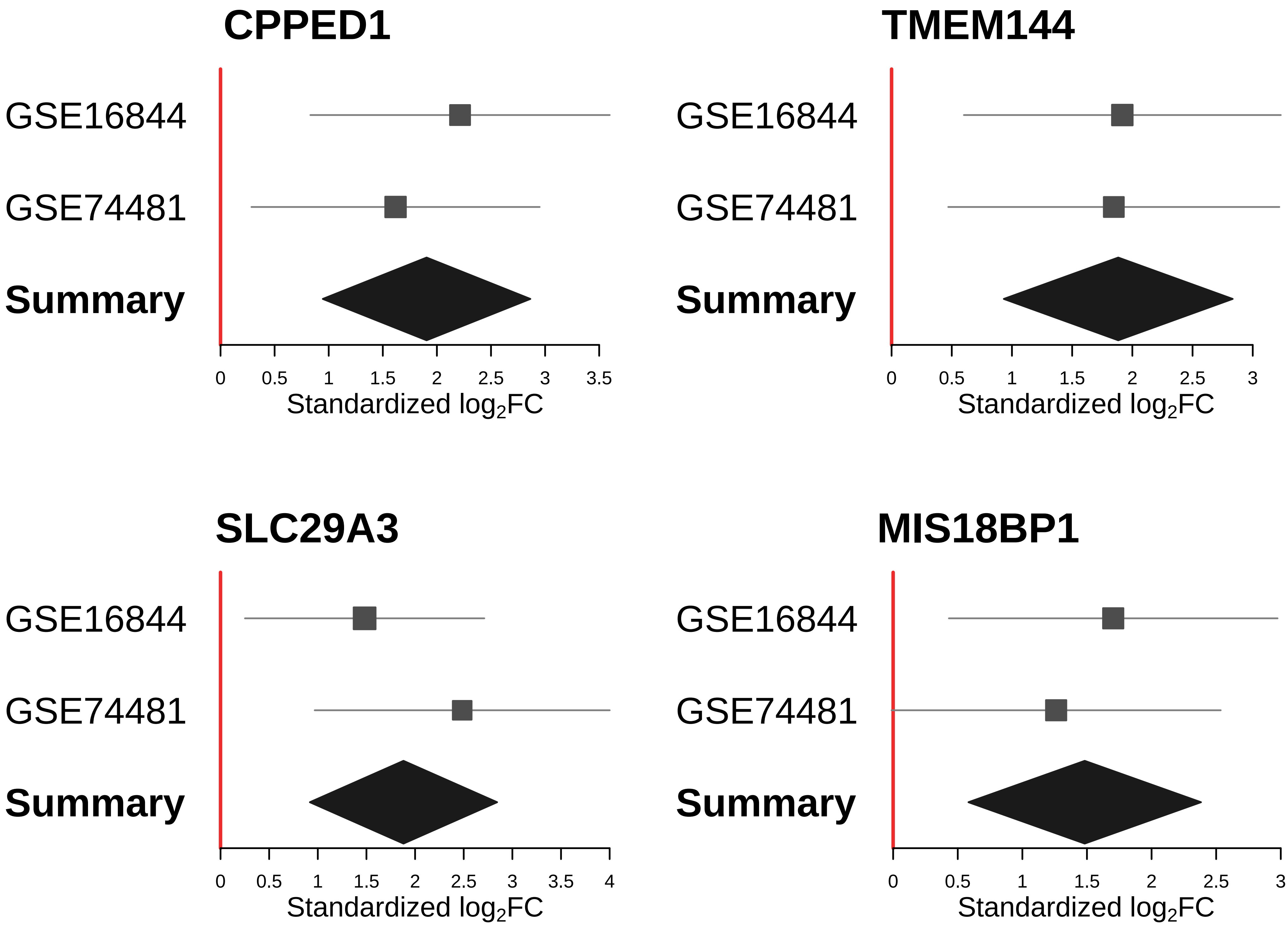
CPPED1
GSE16844
GSE74481
Summary
0
0.5
1
1.5
2
2.5
3
3.5
Standardized $log_2FC$
TMEM144
GSE16844
GSE74481
Summary
0
0.5
1
1.5
2
2.5
3
Standardized $log_2FC$
SLC29A3
GSE16844
GSE74481
Summary
0
0.5
1
1.5
2
2.5
3
3.5
4
Standardized $log_2FC$
MIS18BP1
GSE16844
GSE74481
Summary
0
0.5
1
1.5
2
2.5
3
Standardized $log_2FC$

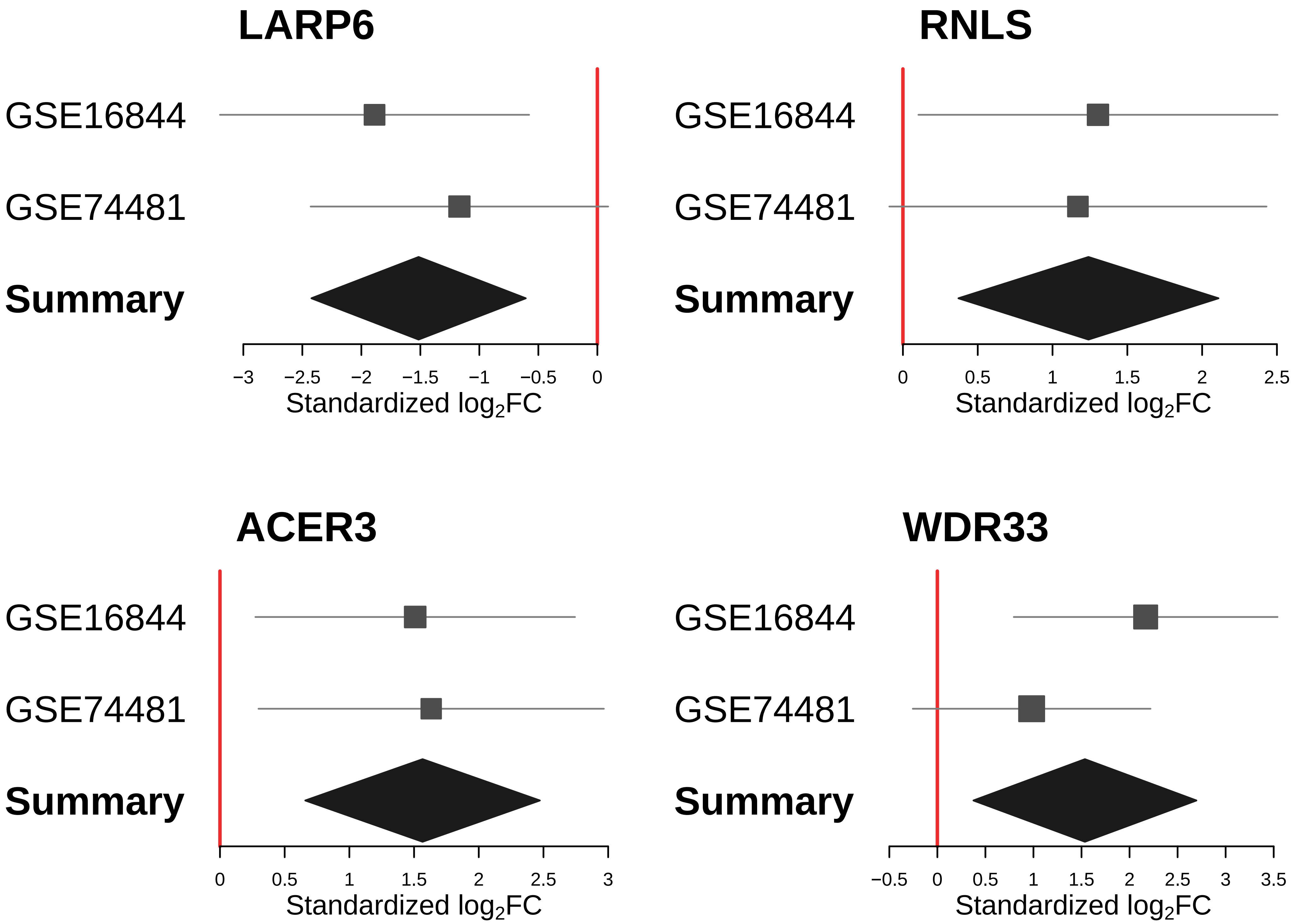

LARP6
GSE16844
GSE74481
Summary
−3 −2.5 −2 −1.5 −1 −0.5 0
Standardized $log_2$FC
RNLS
GSE16844
GSE74481
Summary
0 0.5 1 1.5 2 2.5
Standardized $log_2$FC
ACER3
GSE16844
GSE74481
Summary
0 0.5 1 1.5 2 2.5 3
Standardized $log_2$FC
WDR33
GSE16844
GSE74481
Summary
−0.5 0 0.5 1 1.5 2 2.5 3 3.5
Standardized $log_2$FC

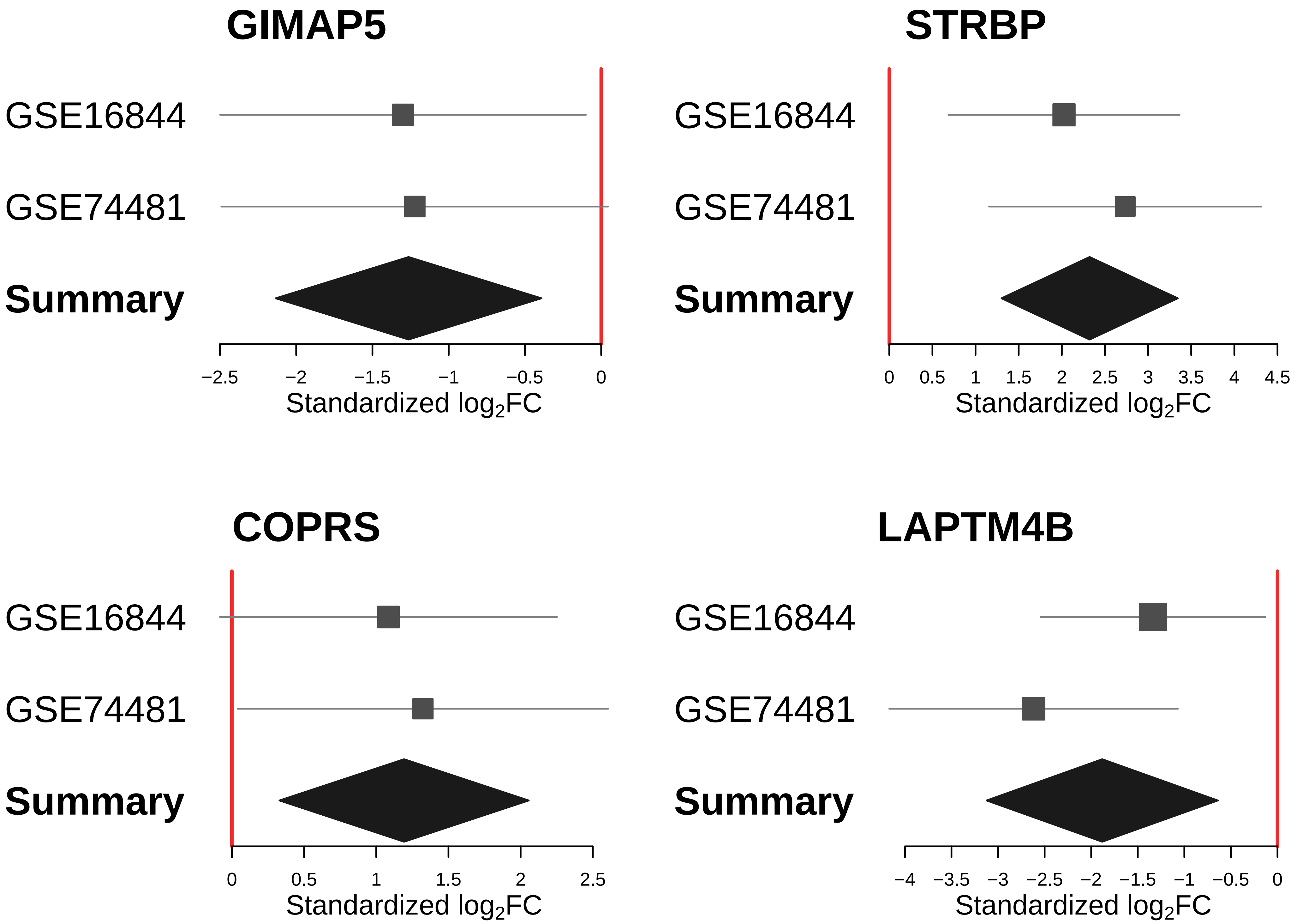
GIMAP5
GSE16844
GSE74481
Summary
−2.5 −2 −1.5 −1 −0.5 0
Standardized log2FC
STRBP
GSE16844
GSE74481
Summary
0 0.5 1 1.5 2 2.5 3 3.5 4 4.5
Standardized log2FC
COPRS
GSE16844
GSE74481
Summary
0 0.5 1 1.5 2 2.5
Standardized log2FC
LAPTM4B
GSE16844
GSE74481
Summary
−4 −3.5 −3 −2.5 −2 −1.5 −1 −0.5 0
Standardized log2FC

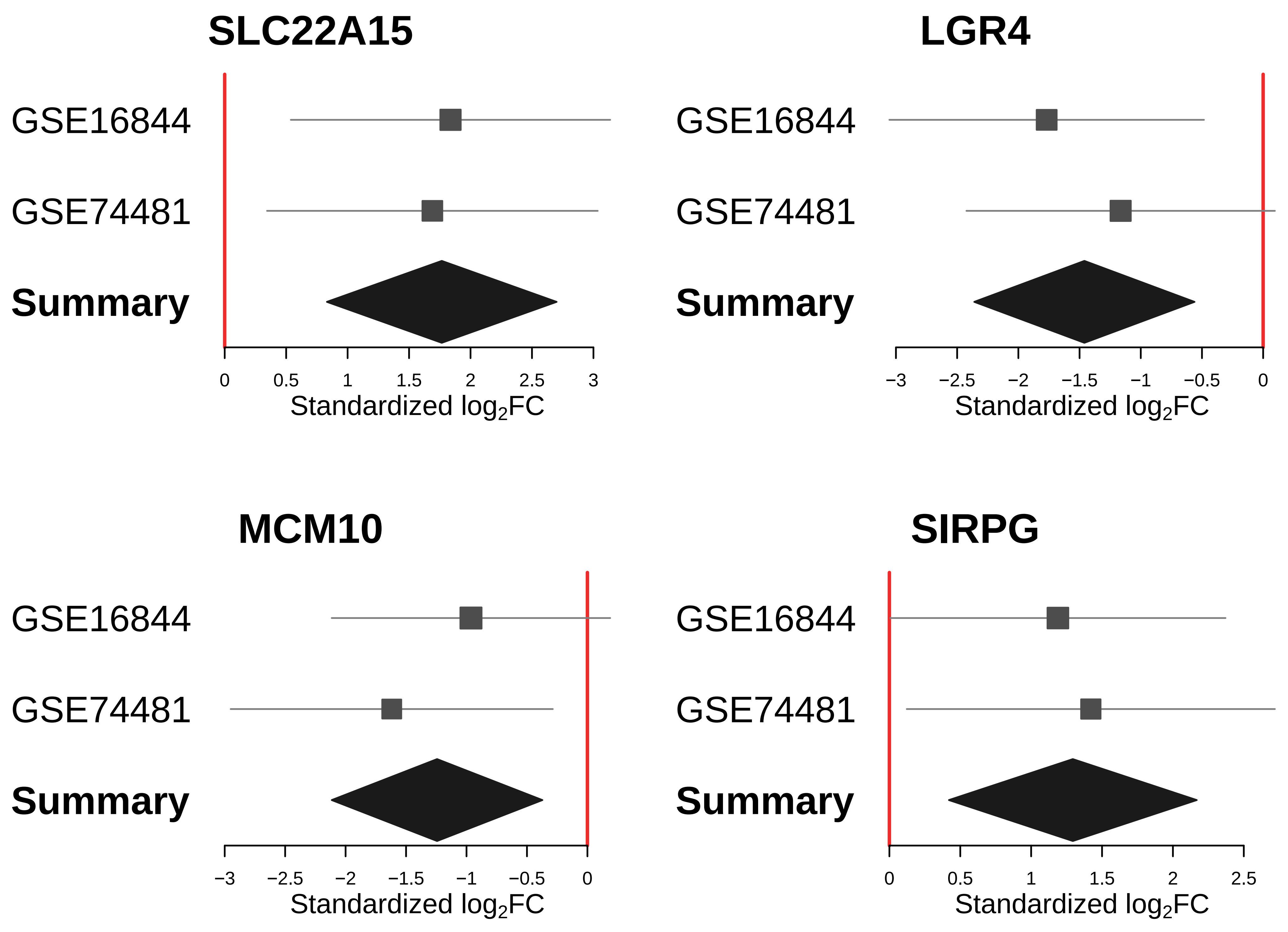
SLC22A15
LGR4
MCM10
SIRPG
GSE16844
GSE74481
Summary
0
0.5
1
1.5
2
2.5
3
−3
−2.5
−2
−1.5
−1
−0.5
Standardized $log_2FC$

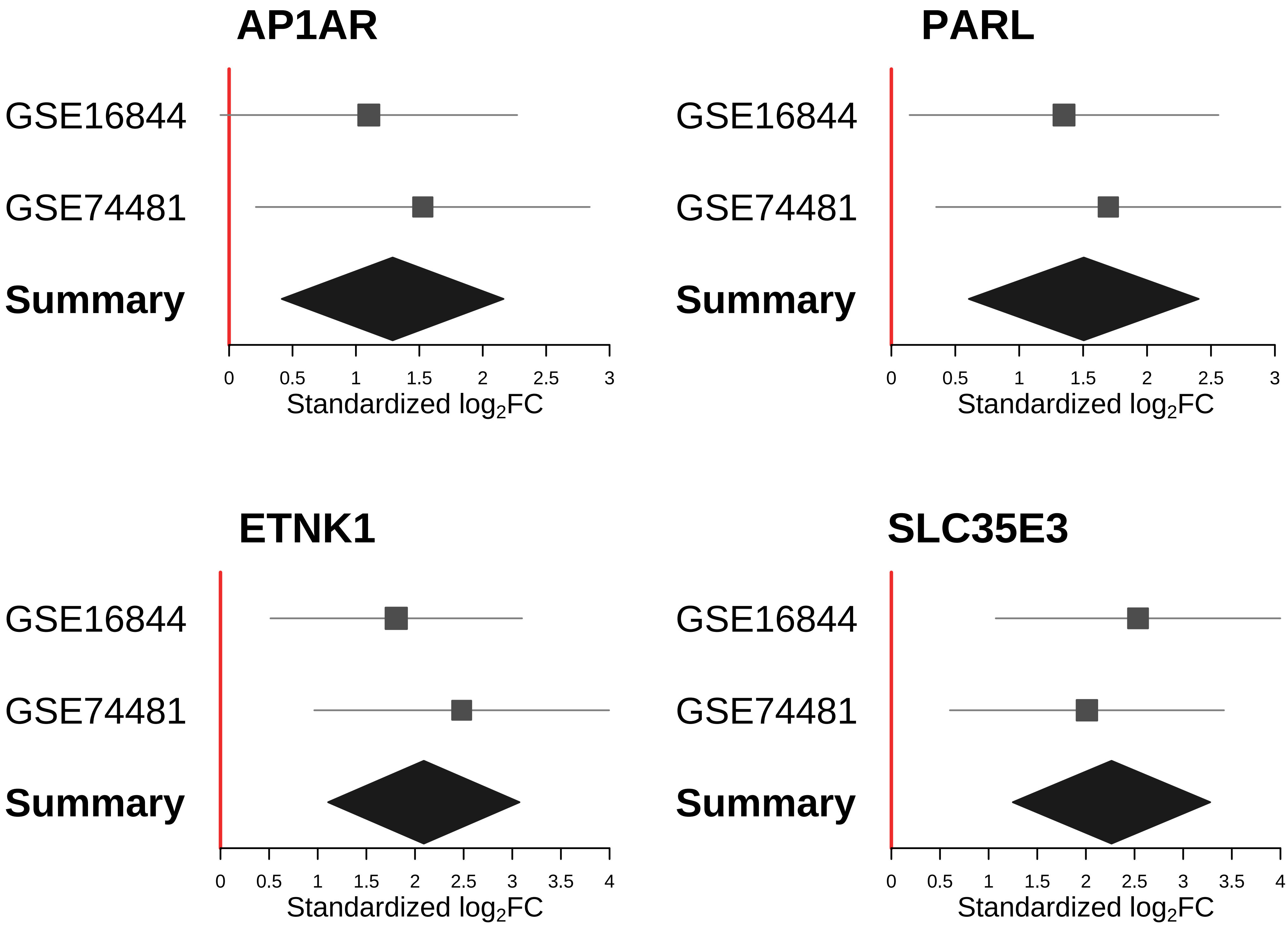
AP1AR
PARL
GSE16844
GSE74481
Summary
0
0.5
1
1.5
2
2.5
3
Standardized log2FC
ETNK1
SLC35E3
3.5
4

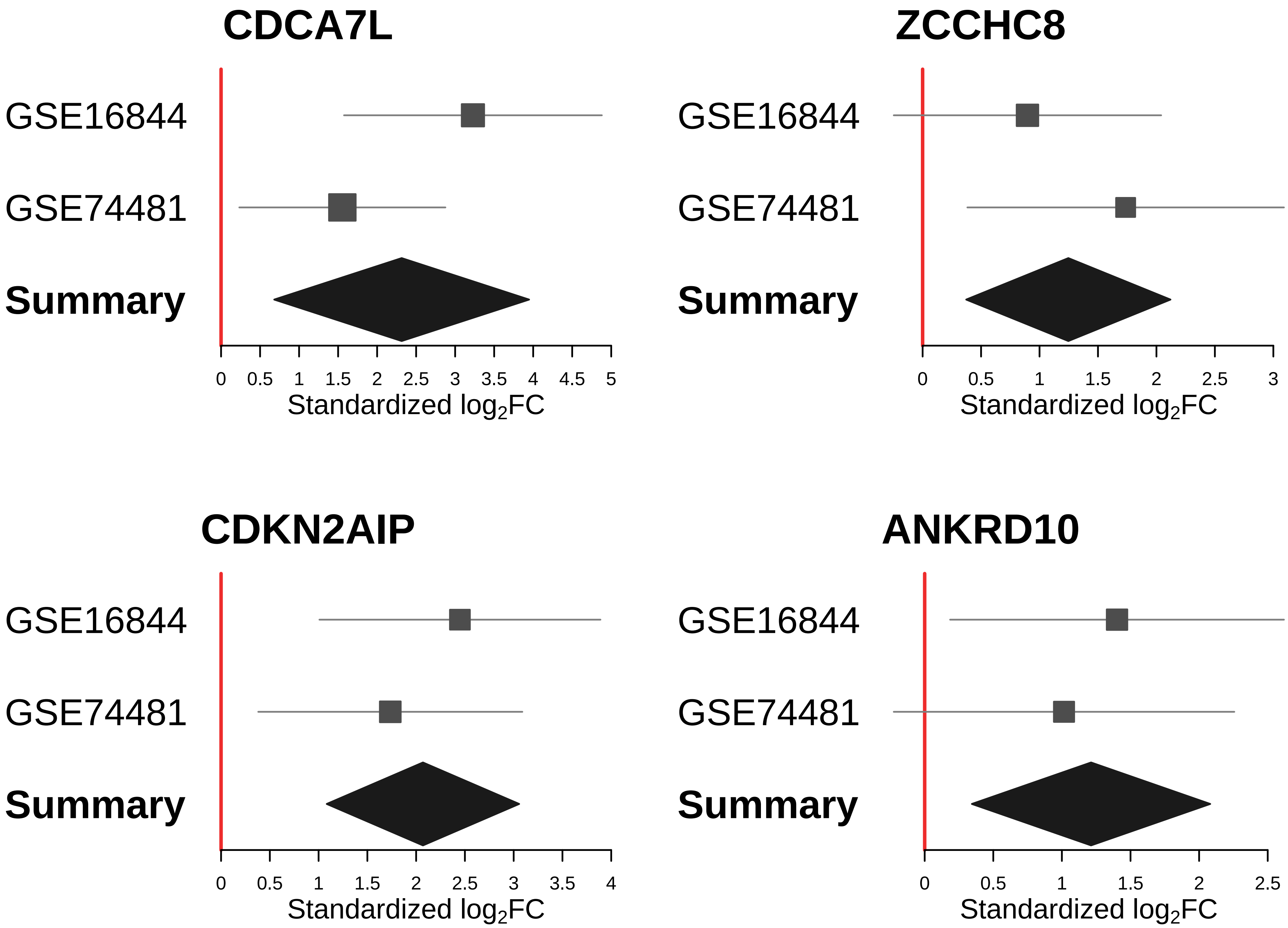

CDCA7L
GSE16844
GSE74481
Summary
0 0.5 1 1.5 2 2.5 3 3.5 4 4.5 5
Standardized $log_2FC$
ZCCHC8
GSE16844
GSE74481
Summary
0 0.5 1 1.5 2 2.5 3
Standardized $log_2FC$
CDKN2AIP
GSE16844
GSE74481
Summary
0 0.5 1 1.5 2 2.5 3 3.5 4
Standardized $log_2FC$
ANKRD10
GSE16844
GSE74481
Summary
0 0.5 1 1.5 2 2.5
Standardized $log_2FC$

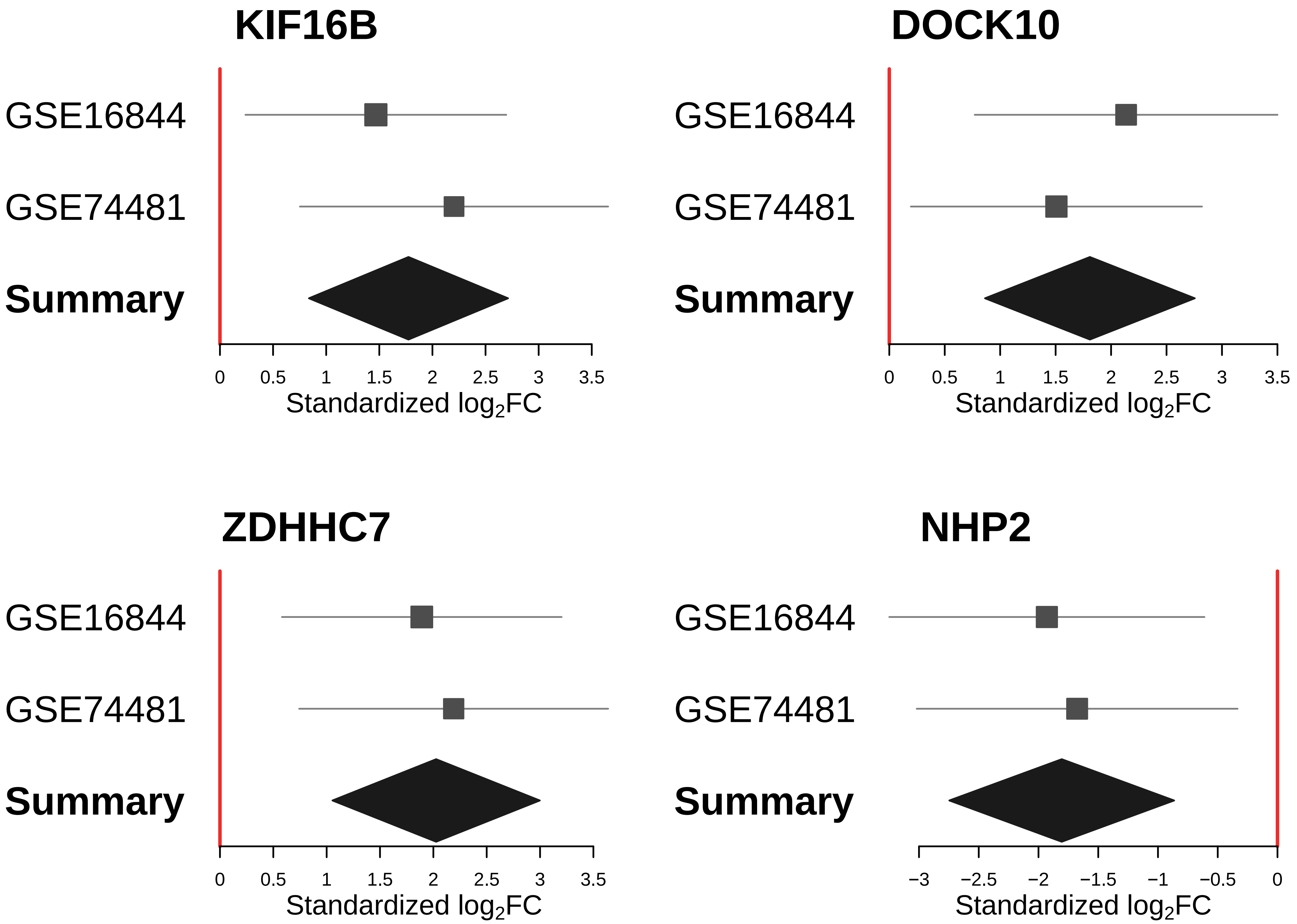
KIF16B
DOCK10
ZDHHC7
NHP2
GSE16844
GSE74481
Summary
0
0.5
1
1.5
2
2.5
3
3.5
−3
−2.5
−2
−1.5
−1
−0.5
Standardized log2FC

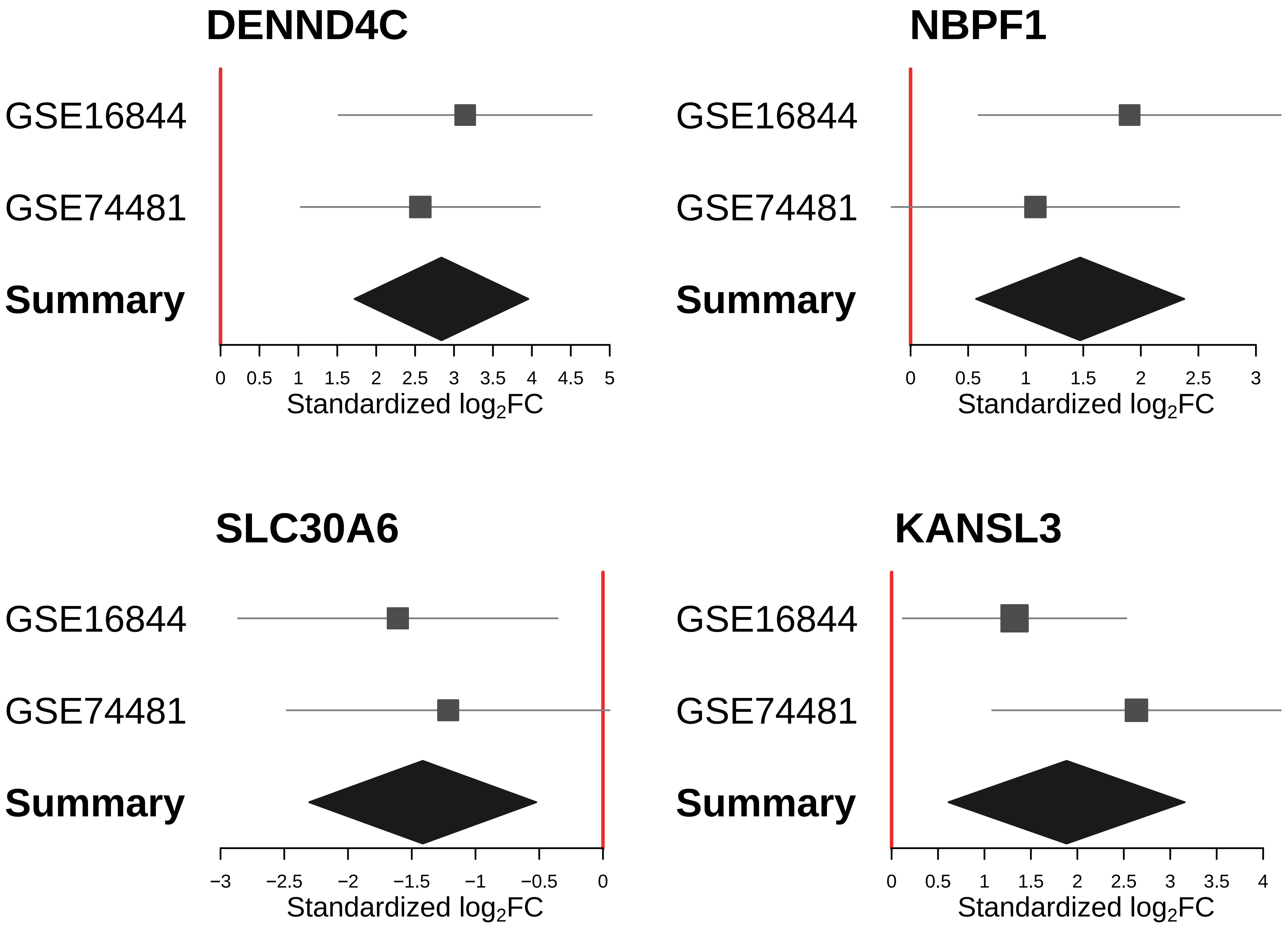

DENND4C
GSE16844
GSE74481
Summary
0 0.5 1 1.5 2 2.5 3 3.5 4 4.5 5
Standardized $log_2FC$
NBPF1
GSE16844
GSE74481
Summary
0 0.5 1 1.5 2 2.5 3
Standardized $log_2FC$
SLC30A6
GSE16844
GSE74481
Summary
−3 −2.5 −2 −1.5 −1 −0.5 0
Standardized $log_2FC$
KANSL3
GSE16844
GSE74481
Summary
0 0.5 1 1.5 2 2.5 3 3.5 4
Standardized $log_2FC$

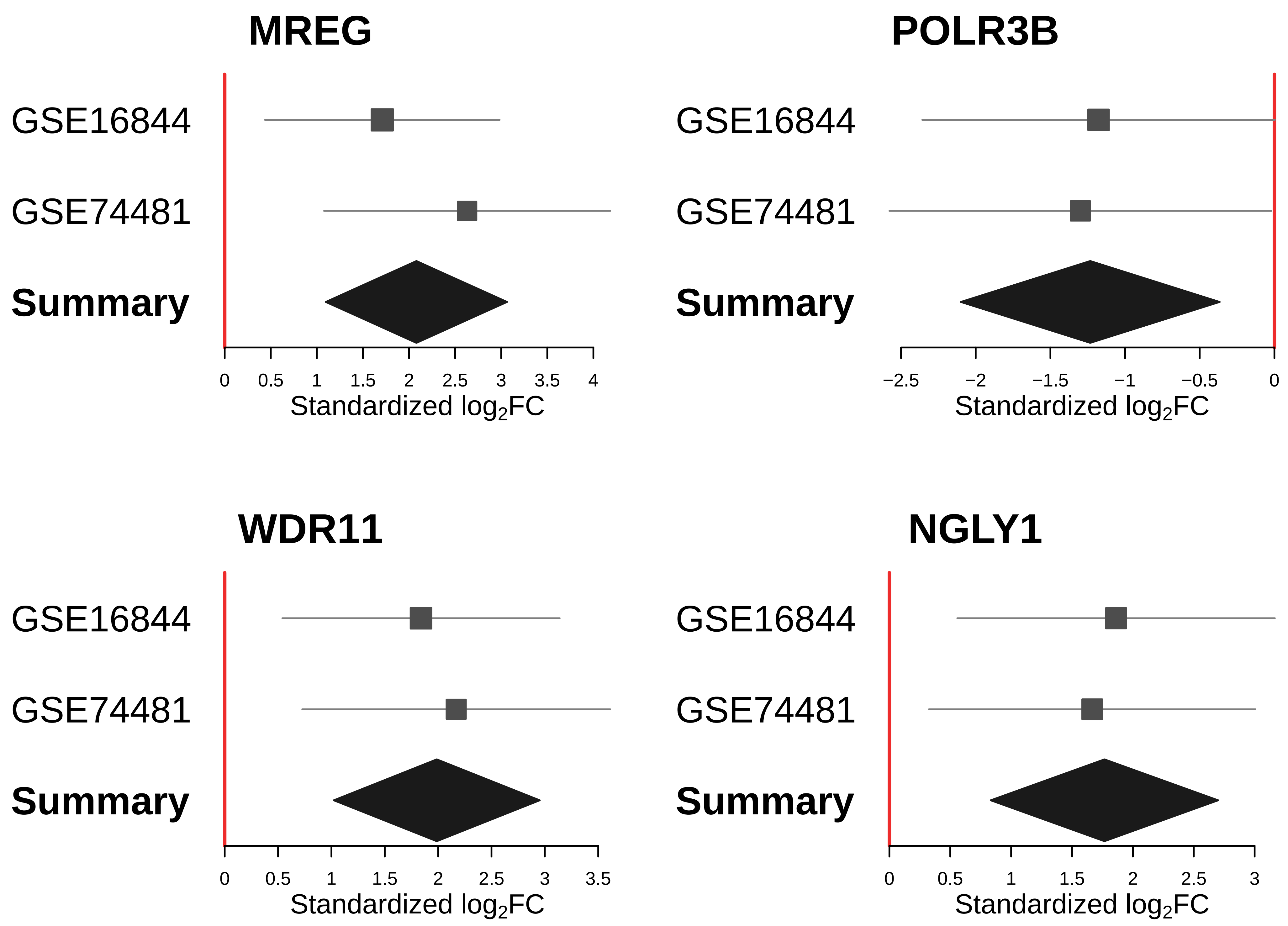

MREG
GSE16844
GSE74481
Summary
0 0.5 1 1.5 2 2.5 3 3.5 4
Standardized log2FC
POLR3B
GSE16844
GSE74481
Summary
−2.5 −2 −1.5 −1 −0.5 0
Standardized log2FC
WDR11
GSE16844
GSE74481
Summary
0 0.5 1 1.5 2 2.5 3 3.5
Standardized log2FC
NGLY1
GSE16844
GSE74481
Summary
0 0.5 1 1.5 2 2.5 3
Standardized log2FC

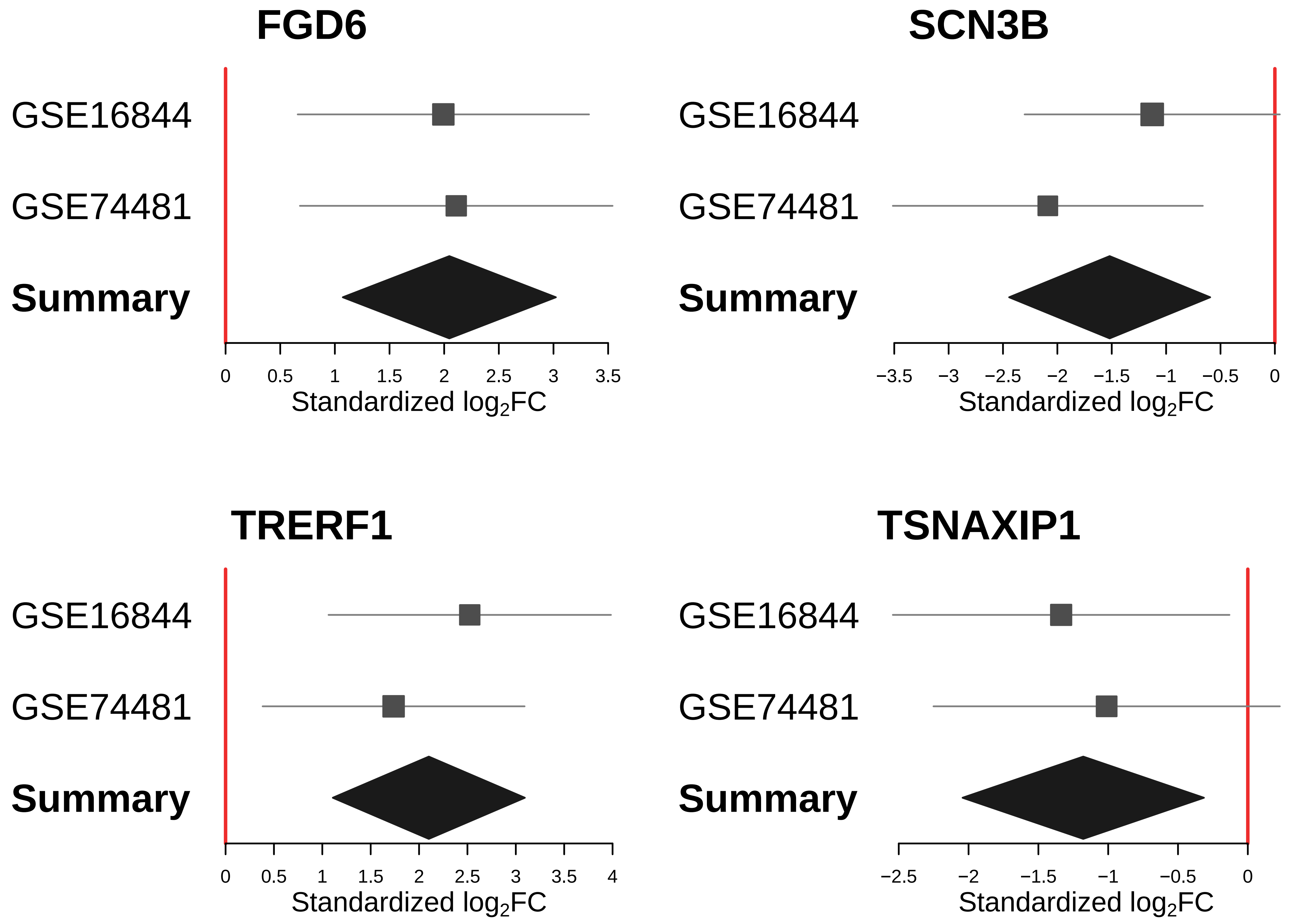
FGD6
GSE16844
GSE74481
Summary
0
0.5
1
1.5
2
2.5
3
3.5
Standardized log2FC
SCN3B
GSE16844
GSE74481
Summary
−3.5
−3
−2.5
−2
−1.5
−1
−0.5
0
Standardized log2FC
TRERF1
GSE16844
GSE74481
Summary
0
0.5
1
1.5
2
2.5
3
3.5
4
Standardized log2FC
TSNAXIP1
GSE16844
GSE74481
Summary
−2.5
−2
−1.5
−1
−0.5
0
Standardized log2FC

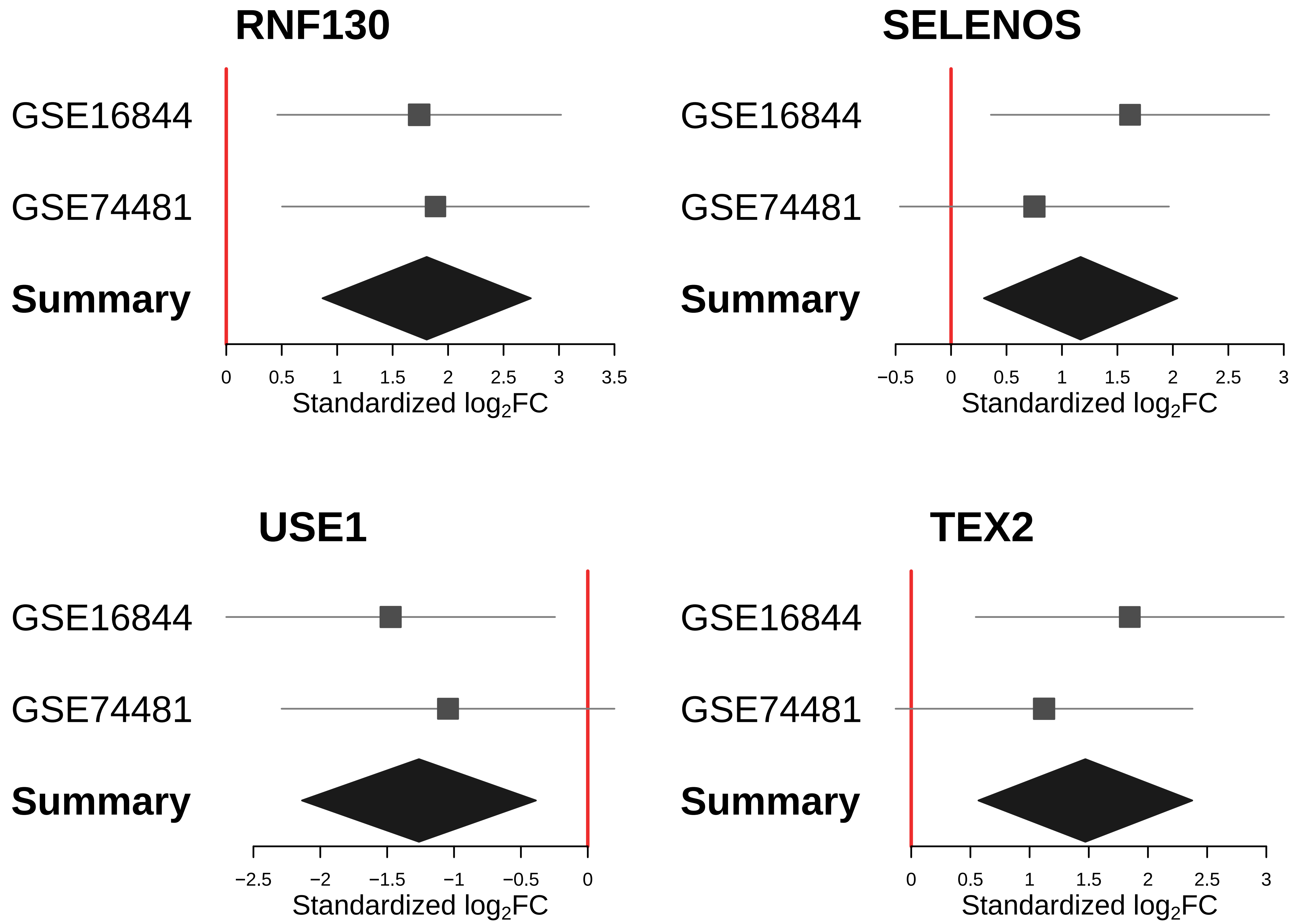

RNF130
GSE16844
GSE74481
Summary
0
0.5
1
1.5
2
2.5
3
3.5
Standardized log2FC
SELENOS
GSE16844
GSE74481
Summary
−0.5
0
0.5
1
1.5
2
2.5
3
Standardized log2FC
USE1
GSE16844
GSE74481
Summary
−2.5
−2
−1.5
−1
−0.5
0
Standardized log2FC
TEX2
GSE16844
GSE74481
Summary
0
0.5
1
1.5
2
2.5
3
Standardized log2FC

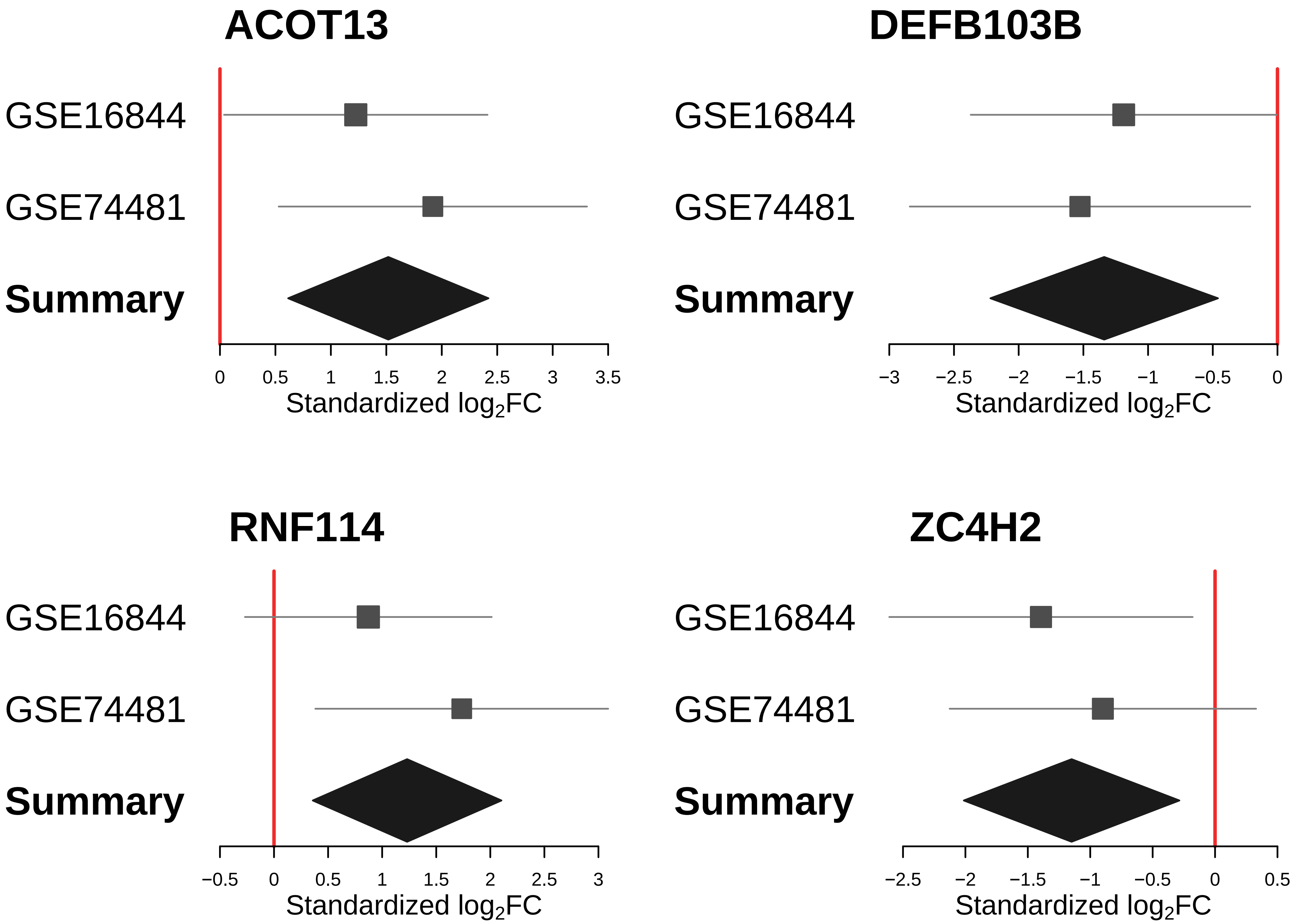
ACOT13
GSE16844
GSE74481
Summary
0 0.5 1 1.5 2 2.5 3 3.5
Standardized log2FC
DEFB103B
GSE16844
GSE74481
Summary
−3 −2.5 −2 −1.5 −1 −0.5 0
Standardized log2FC
RNF114
GSE16844
GSE74481
Summary
−0.5 0 0.5 1 1.5 2 2.5 3
Standardized log2FC
ZC4H2
GSE16844
GSE74481
Summary
−2.5 −2 −1.5 −1 −0.5 0 0.5
Standardized log2FC

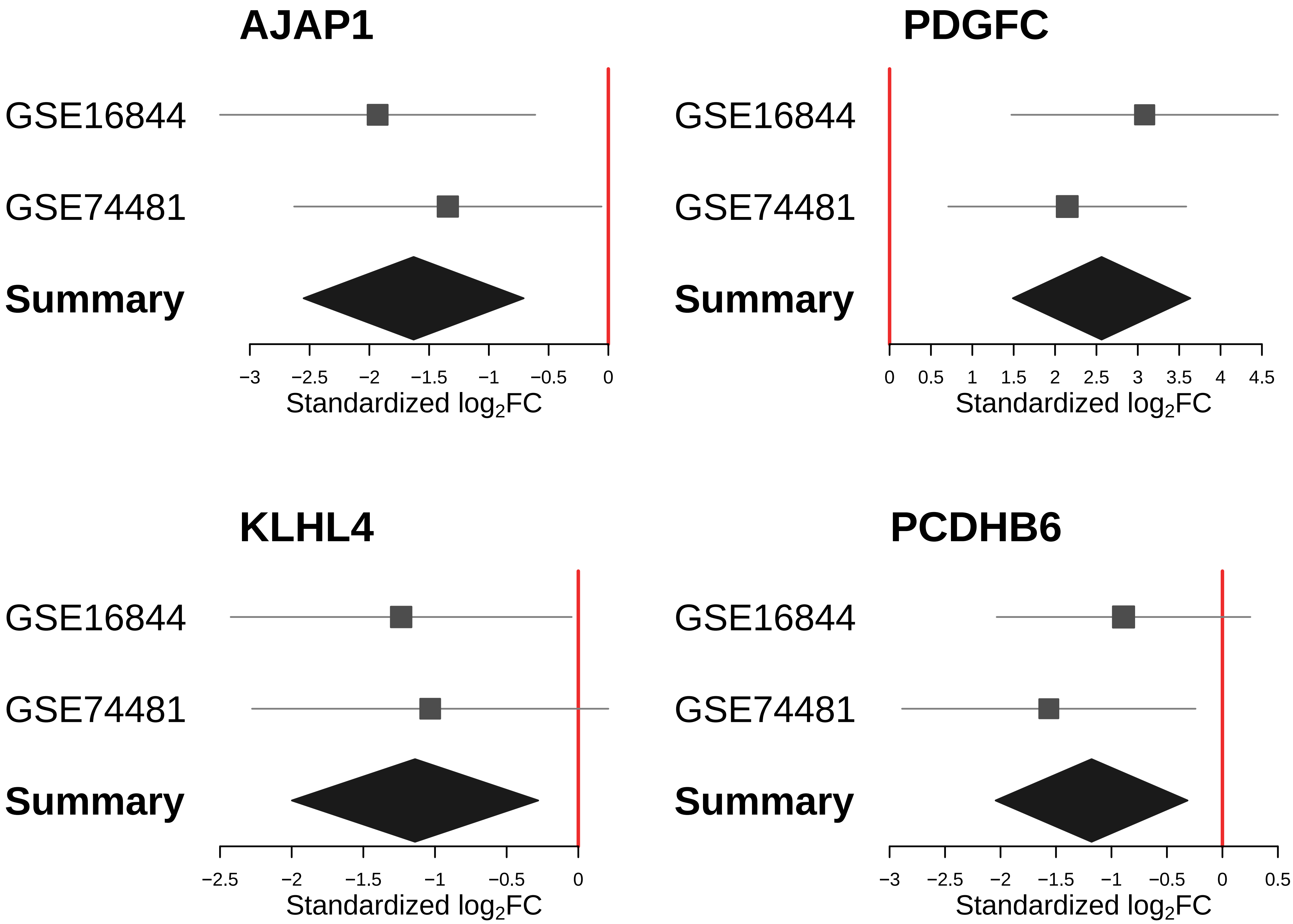
AJAP1
GSE16844
GSE74481
Summary
−3
−2.5
−2
−1.5
−1
−0.5
0
Standardized log2FC
PDGFC
GSE16844
GSE74481
Summary
0
0.5
1
1.5
2
2.5
3
3.5
4
4.5
Standardized log2FC
KLHL4
GSE16844
GSE74481
Summary
−2.5
−2
−1.5
−1
−0.5
0
Standardized log2FC
PCDHB6
GSE16844
GSE74481
Summary
−3
−2.5
−2
−1.5
−1
−0.5
0
0.5
Standardized log2FC

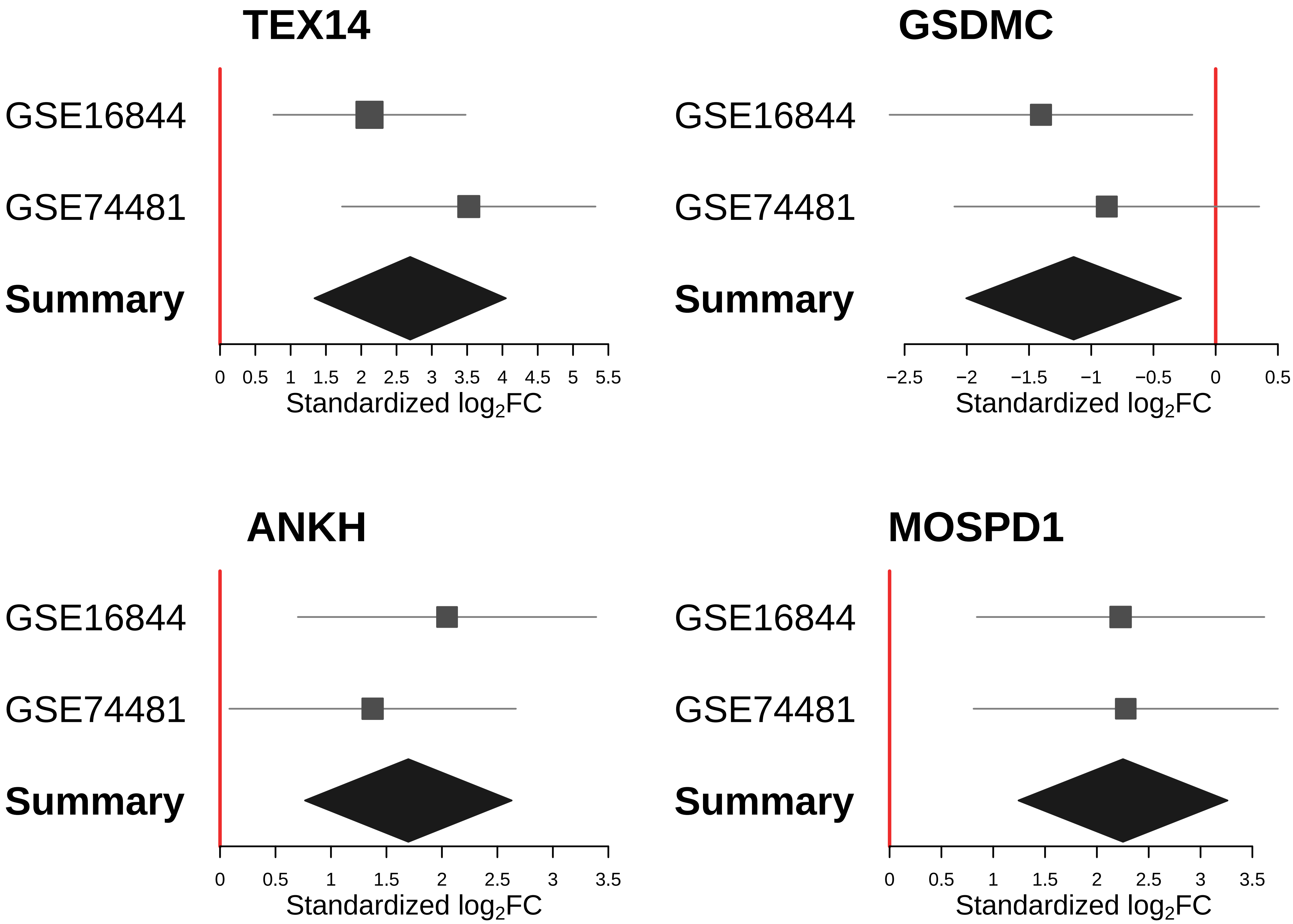

TEX14
GSE16844
GSE74481
Summary
0
0.5
1
1.5
2
2.5
3
3.5
4
4.5
5
5.5
Standardized log2FC
GSDMC
GSE16844
GSE74481
Summary
−2.5
−2
−1.5
−1
−0.5
0
0.5
Standardized log2FC
ANKH
GSE16844
GSE74481
Summary
0
0.5
1
1.5
2
2.5
3
3.5
Standardized log2FC
MOSPD1
GSE16844
GSE74481
Summary
0
0.5
1
1.5
2
2.5
3
3.5
Standardized log2FC

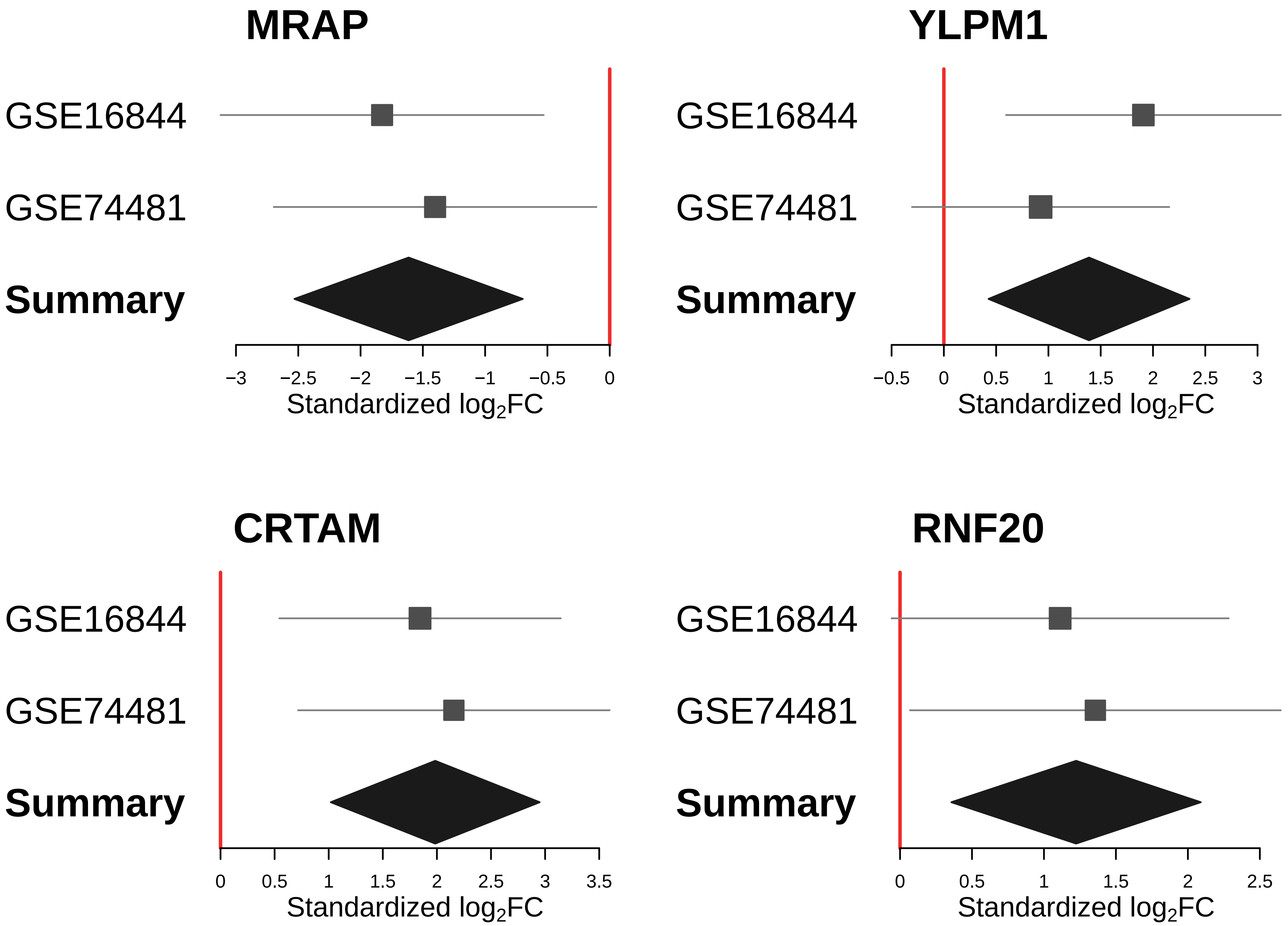

MRAP
GSE16844
GSE74481
Summary
−3 −2.5 −2 −1.5 −1 −0.5 0
Standardized $log_2FC$
YLPM1
GSE16844
GSE74481
Summary
−0.5 0 0.5 1 1.5 2 2.5 3
Standardized $log_2FC$
CRTAM
GSE16844
GSE74481
Summary
0 0.5 1 1.5 2 2.5 3 3.5
Standardized $log_2FC$
RNF20
GSE16844
GSE74481
Summary
0 0.5 1 1.5 2 2.5
Standardized $log_2FC$

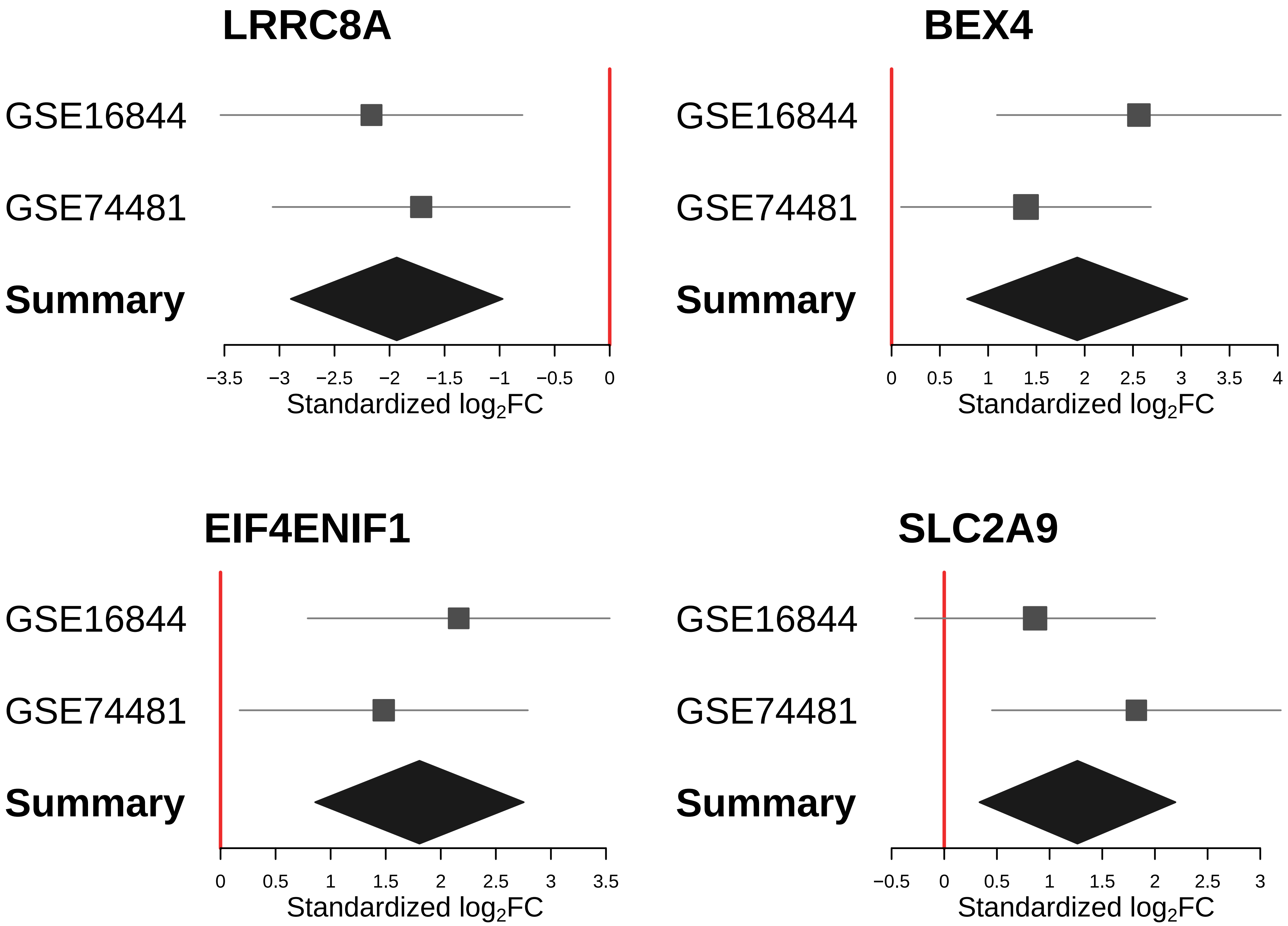
LRRC8A
GSE16844
GSE74481
Summary
−3.5 −3 −2.5 −2 −1.5 −1 −0.5 0
Standardized log2FC
BEX4
GSE16844
GSE74481
Summary
0 0.5 1 1.5 2 2.5 3 3.5 4
Standardized log2FC
EIF4ENIF1
GSE16844
GSE74481
Summary
0 0.5 1 1.5 2 2.5 3 3.5
Standardized log2FC
SLC2A9
GSE16844
GSE74481
Summary
−0.5 0 0.5 1 1.5 2 2.5 3
Standardized log2FC

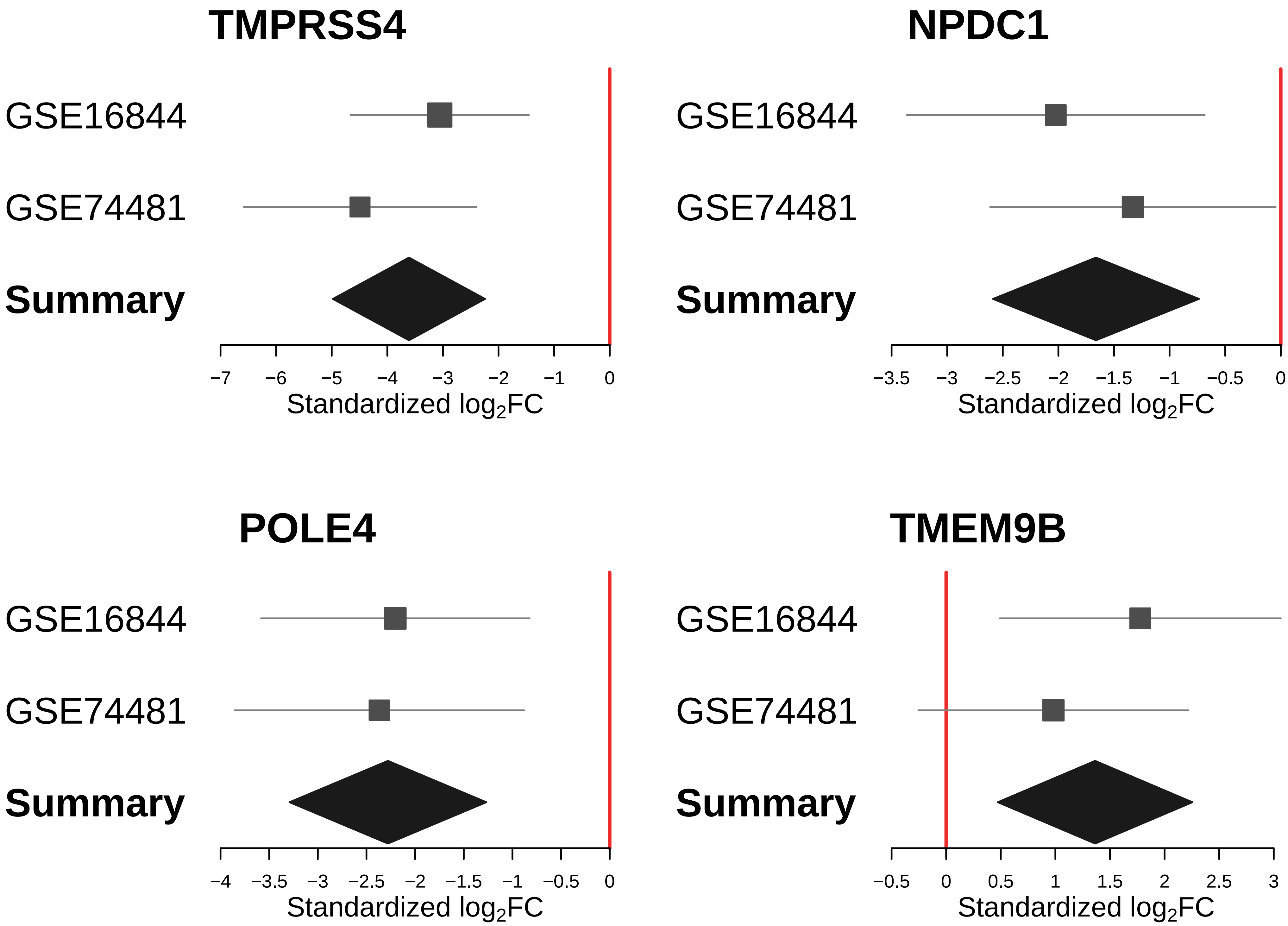
TMPRSS4
GSE16844
GSE74481
Summary
−7 −6 −5 −4 −3 −2 −1 0
Standardized log2FC
NPDC1
GSE16844
GSE74481
Summary
−3.5 −3 −2.5 −2 −1.5 −1 −0.5 0
Standardized log2FC
POLE4
GSE16844
GSE74481
Summary
−4 −3.5 −3 −2.5 −2 −1.5 −1 −0.5 0
Standardized log2FC
TMEM9B
GSE16844
GSE74481
Summary
−0.5 0 0.5 1 1.5 2 2.5 3
Standardized log2FC

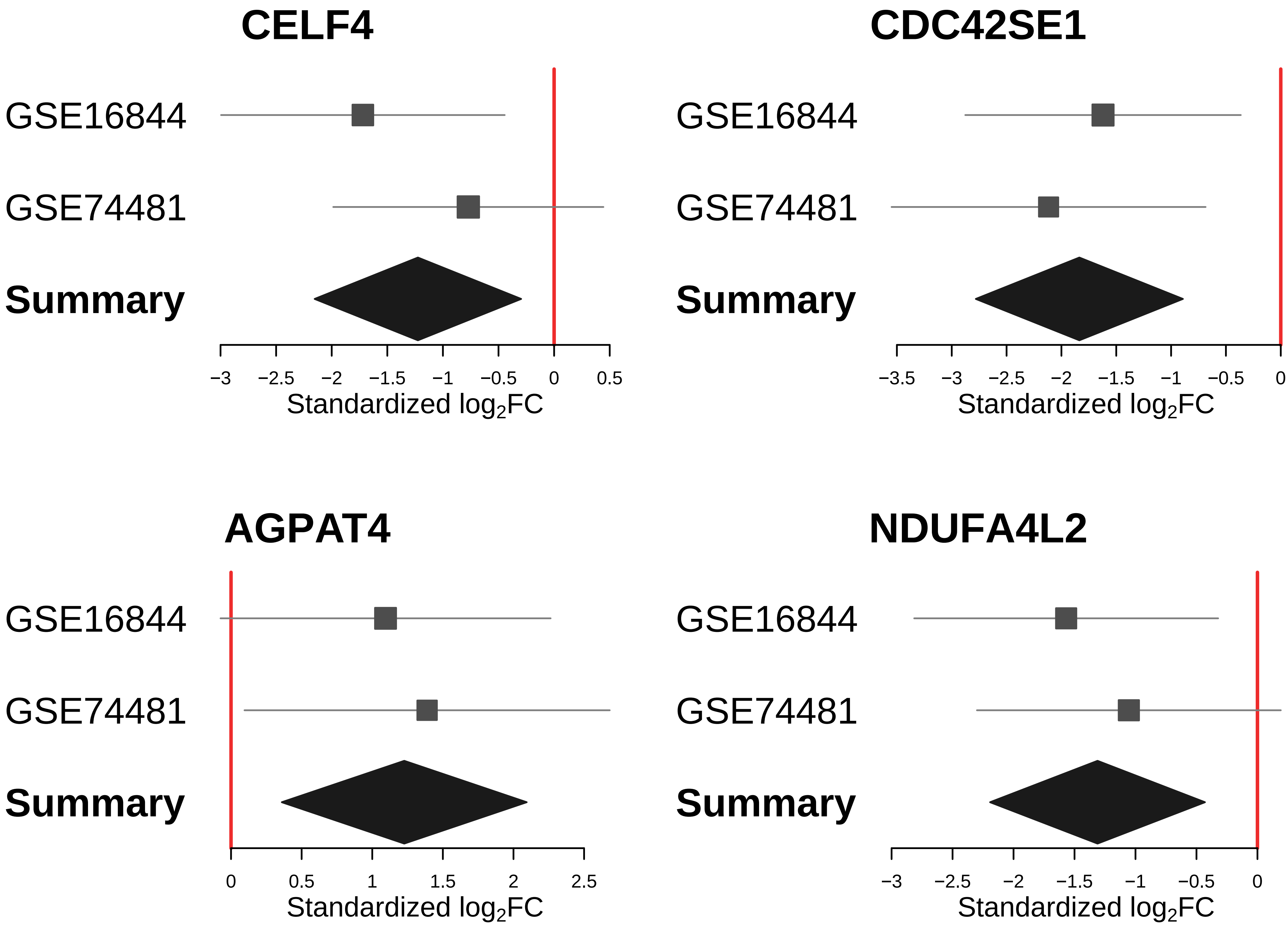

CELF4
GSE16844
GSE74481
Summary
−3 −2.5 −2 −1.5 −1 −0.5 0 0.5
Standardized log2FC
CDC42SE1
GSE16844
GSE74481
Summary
−3.5 −3 −2.5 −2 −1.5 −1 −0.5 0
Standardized log2FC
AGPAT4
GSE16844
GSE74481
Summary
0 0.5 1 1.5 2 2.5
Standardized log2FC
NDUFA4L2
GSE16844
GSE74481
Summary
−3 −2.5 −2 −1.5 −1 −0.5 0
Standardized log2FC

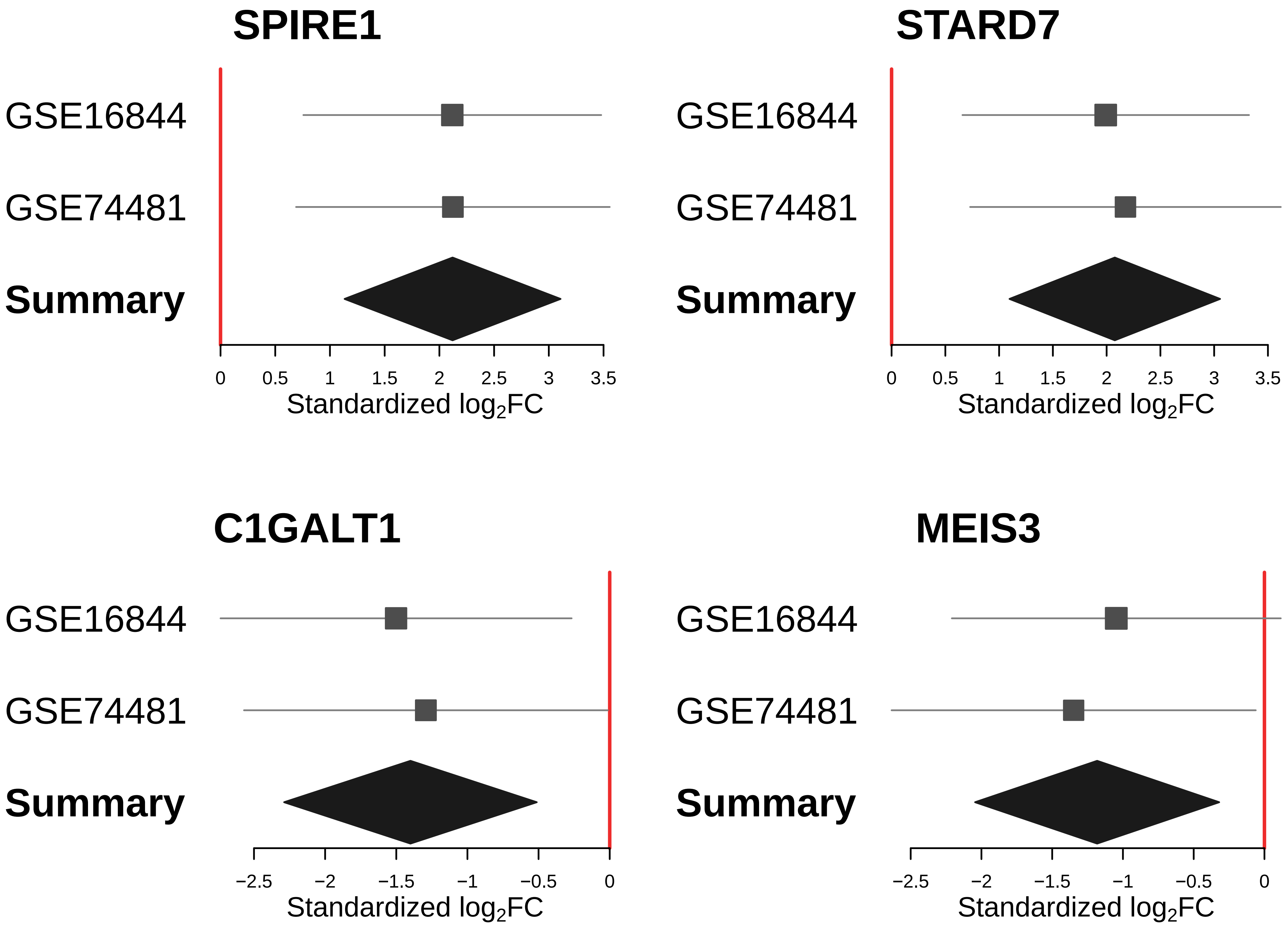
SPIRE1
GSE16844
GSE74481
Summary
0 0.5 1 1.5 2 2.5 3 3.5
Standardized log2FC
STARD7
GSE16844
GSE74481
Summary
0 0.5 1 1.5 2 2.5 3 3.5
Standardized log2FC
C1GALT1
GSE16844
GSE74481
Summary
−2.5 −2 −1.5 −1 −0.5 0
Standardized log2FC
MEIS3
GSE16844
GSE74481
Summary
−2.5 −2 −1.5 −1 −0.5 0
Standardized log2FC

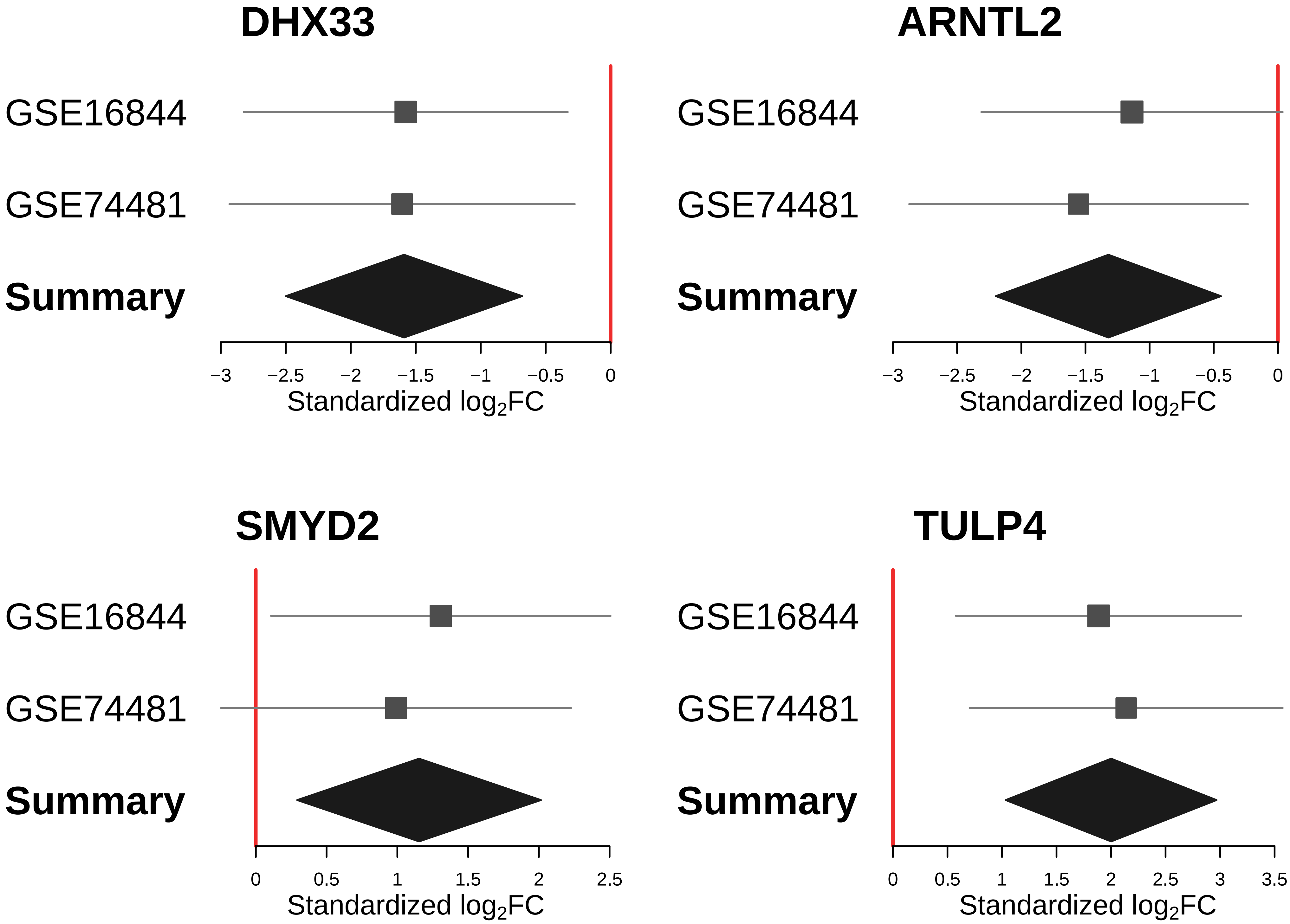
DHX33
GSE16844
GSE74481
Summary
−3
−2.5
−2
−1.5
−1
−0.5
0
Standardized log2FC
ARNTL2
GSE16844
GSE74481
Summary
−3
−2.5
−2
−1.5
−1
−0.5
0
Standardized log2FC
SMYD2
GSE16844
GSE74481
Summary
0
0.5
1
1.5
2
2.5
Standardized log2FC
TULP4
GSE16844
GSE74481
Summary
0
0.5
1
1.5
2
2.5
3
3.5
Standardized log2FC

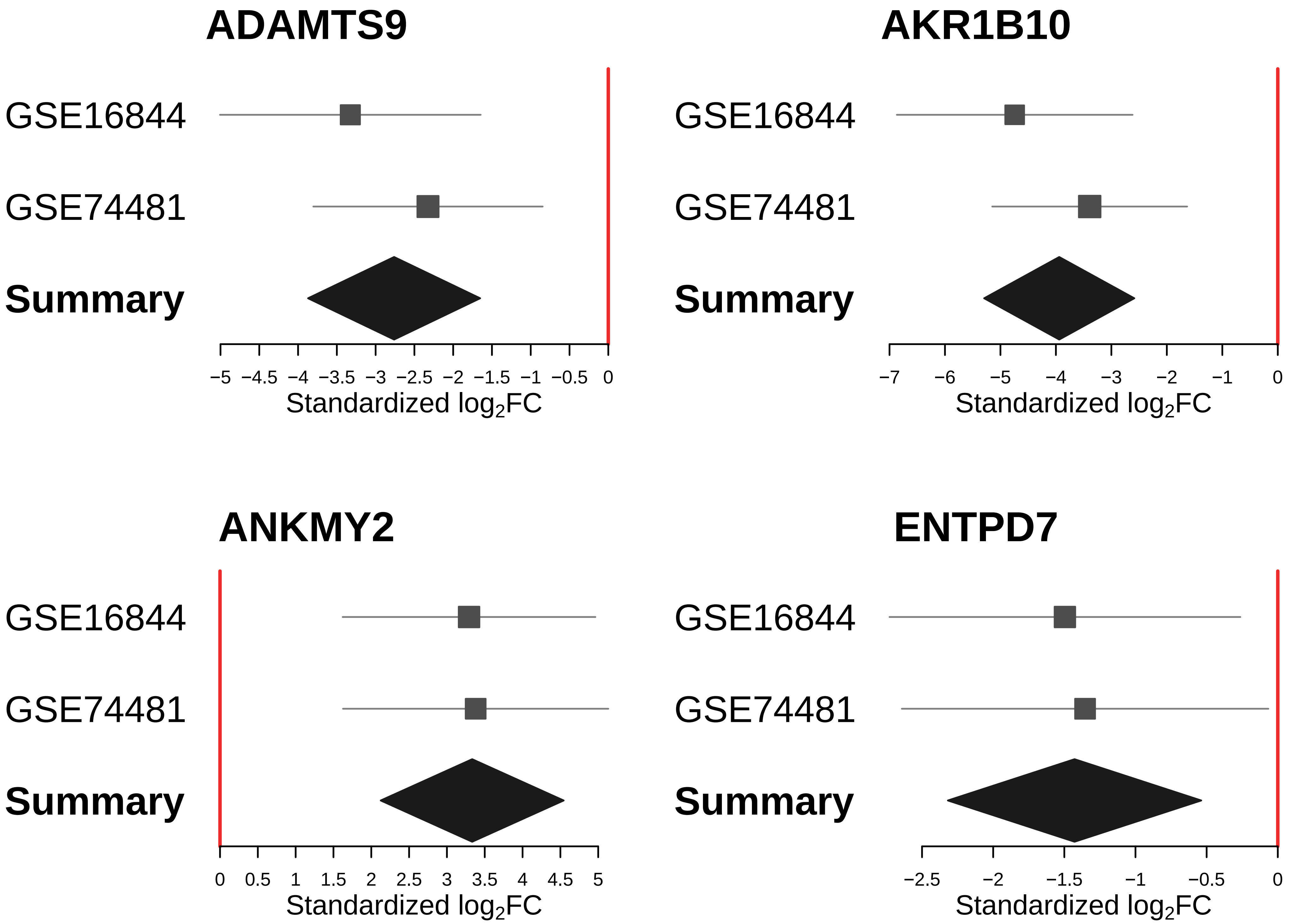
ADAMTS9
GSE16844
GSE74481
Summary
−5 −4.5 −4 −3.5 −3 −2.5 −2 −1.5 −1 −0.5 0
Standardized log2FC
AKR1B10
GSE16844
GSE74481
Summary
−7 −6 −5 −4 −3 −2 −1 0
Standardized log2FC
ANKMY2
GSE16844
GSE74481
Summary
0 0.5 1 1.5 2 2.5 3 3.5 4 4.5 5
Standardized log2FC
ENTPD7
GSE16844
GSE74481
Summary
−2.5 −2 −1.5 −1 −0.5 0
Standardized log2FC

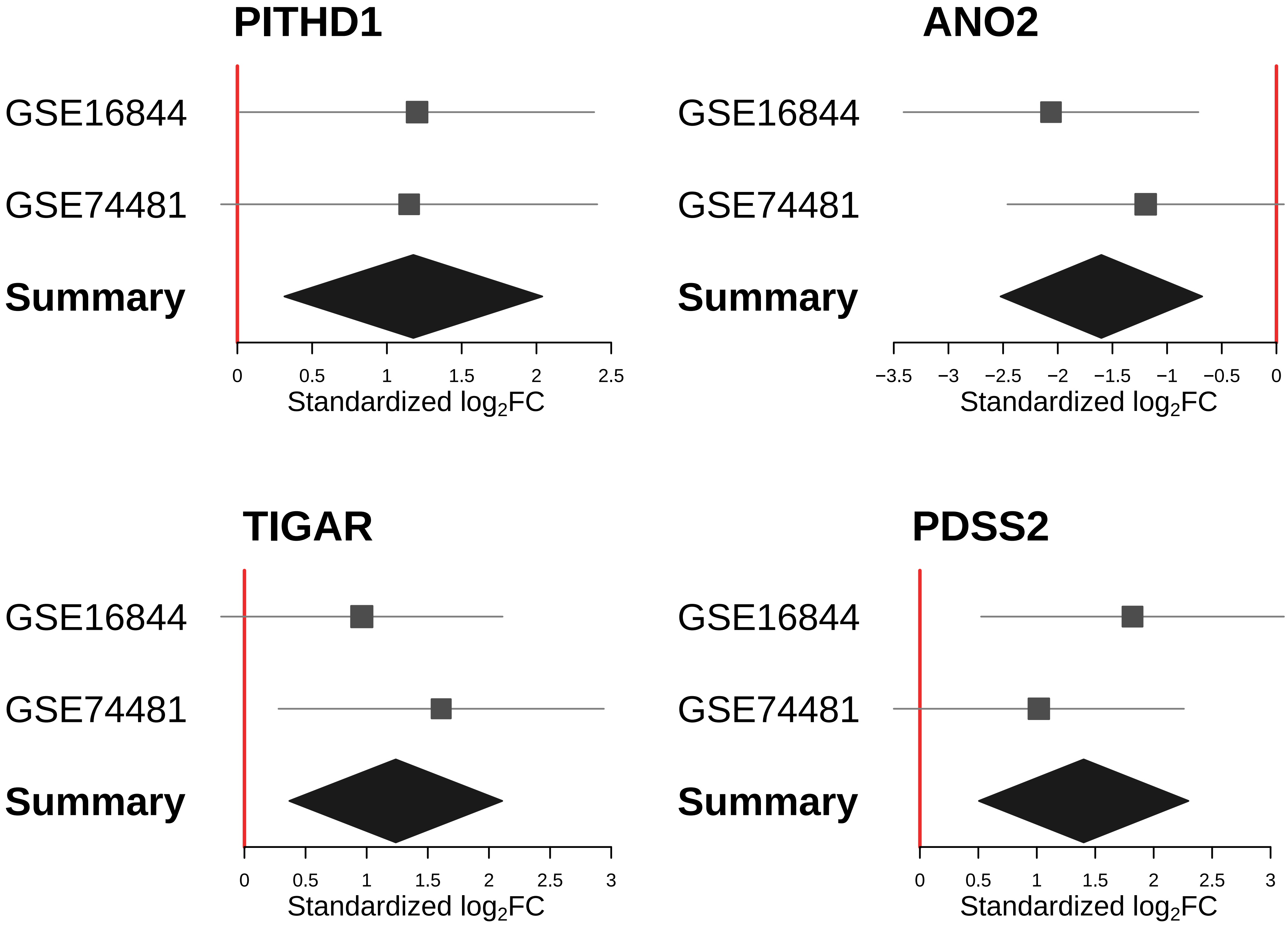
PITHD1
GSE16844
GSE74481
Summary
0
0.5
1
1.5
2
2.5
Standardized log2FC
ANO2
GSE16844
GSE74481
Summary
−3.5
−3
−2.5
−2
−1.5
−1
−0.5
0
Standardized log2FC
TIGAR
GSE16844
GSE74481
Summary
0
0.5
1
1.5
2
2.5
3
Standardized log2FC
PDSS2
GSE16844
GSE74481
Summary
0
0.5
1
1.5
2
2.5
3
Standardized log2FC

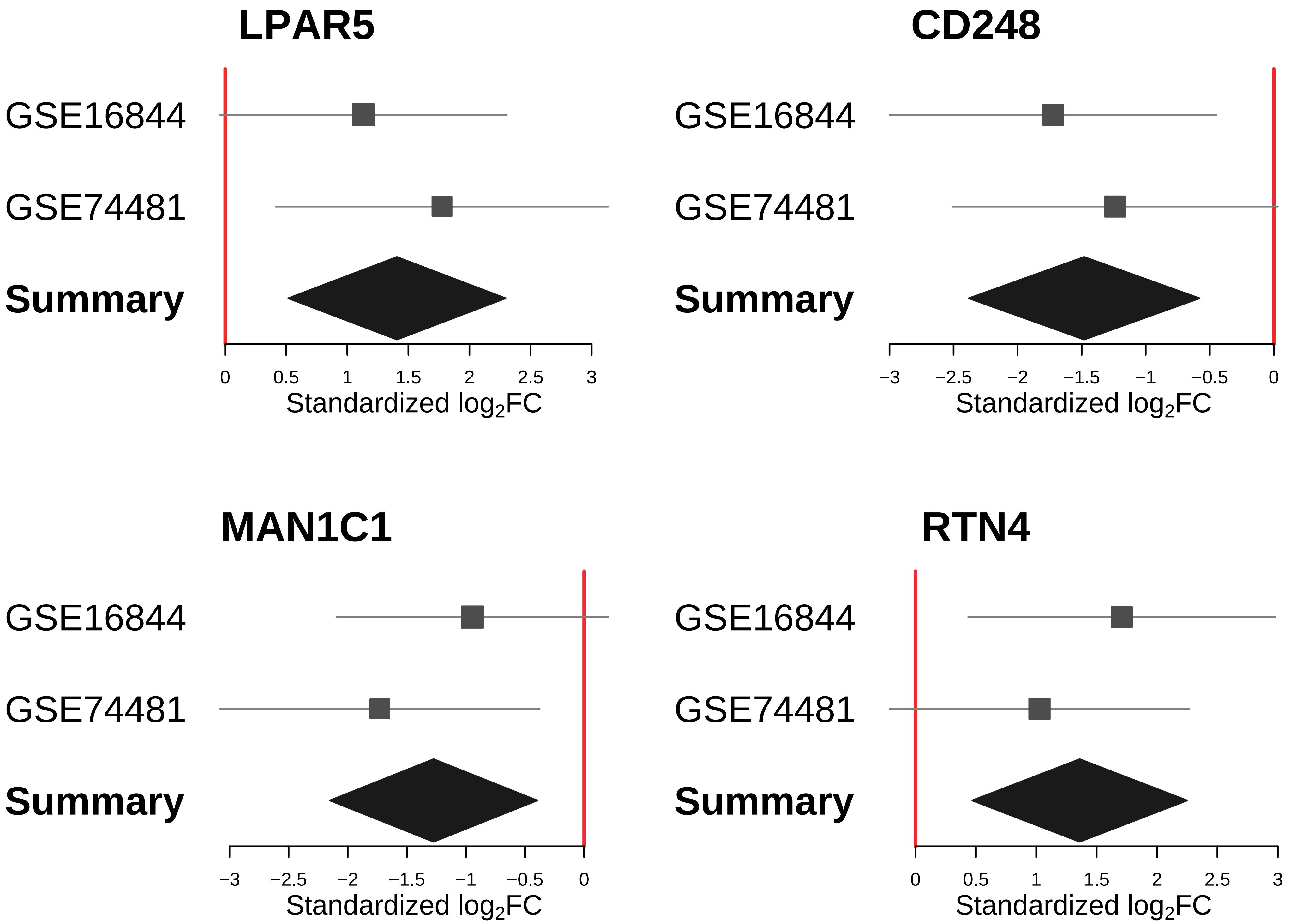

LPAR5
GSE16844
GSE74481
Summary
0
0.5
1
1.5
2
2.5
3
Standardized log2FC
CD248
GSE16844
GSE74481
Summary
−3
−2.5
−2
−1.5
−1
−0.5
0
Standardized log2FC
MAN1C1
GSE16844
GSE74481
Summary
−3
−2.5
−2
−1.5
−1
−0.5
0
Standardized log2FC
RTN4
GSE16844
GSE74481
Summary
0
0.5
1
1.5
2
2.5
3
Standardized log2FC

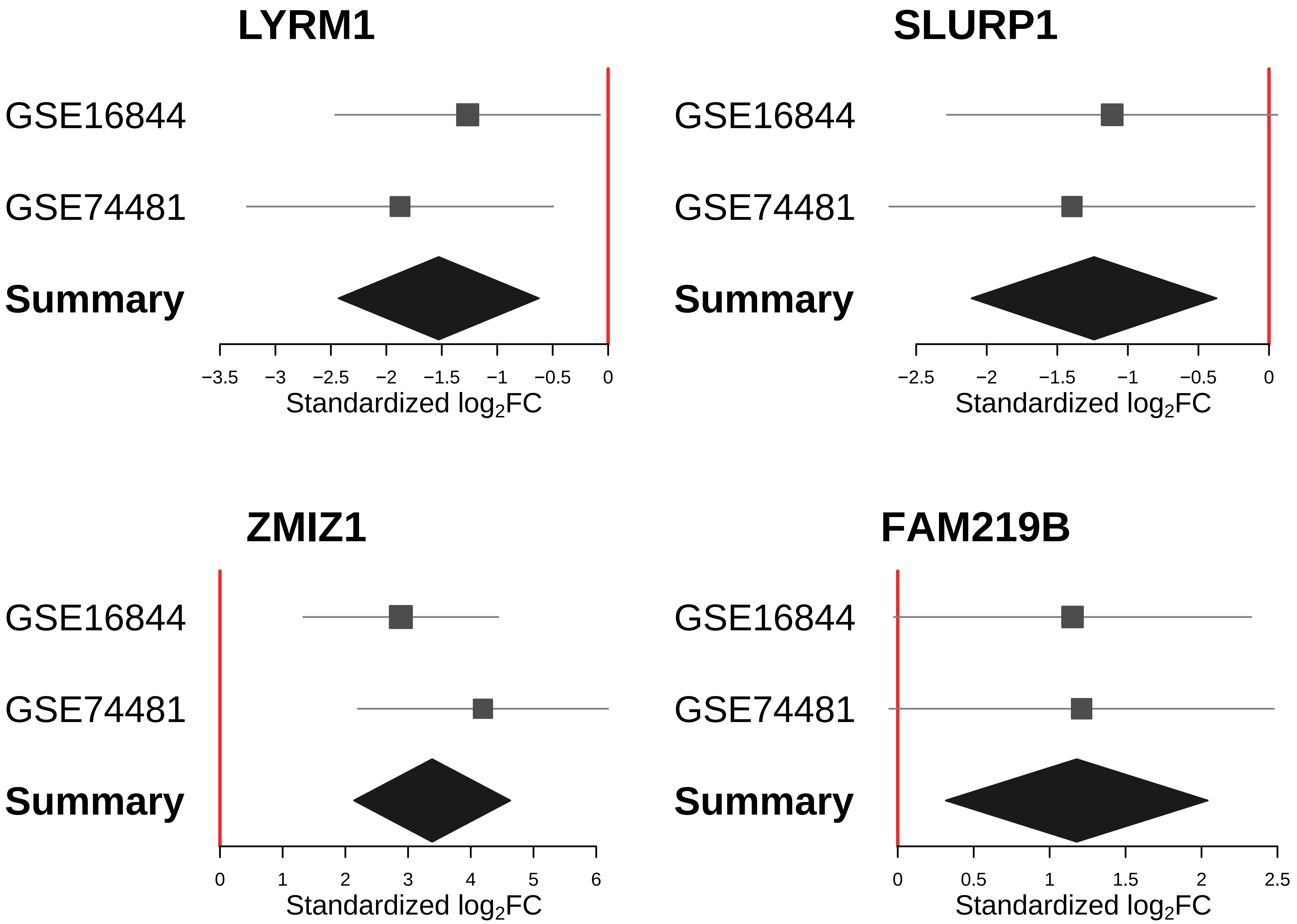
LYRM1
GSE16844
GSE74481
Summary
−3.5 −3 −2.5 −2 −1.5 −1 −0.5 0
Standardized log2FC
SLURP1
GSE16844
GSE74481
Summary
−2.5 −2 −1.5 −1 −0.5 0
Standardized log2FC
ZMIZ1
GSE16844
GSE74481
Summary
0 1 2 3 4 5 6
Standardized log2FC
FAM219B
GSE16844
GSE74481
Summary
0 0.5 1 1.5 2 2.5
Standardized log2FC

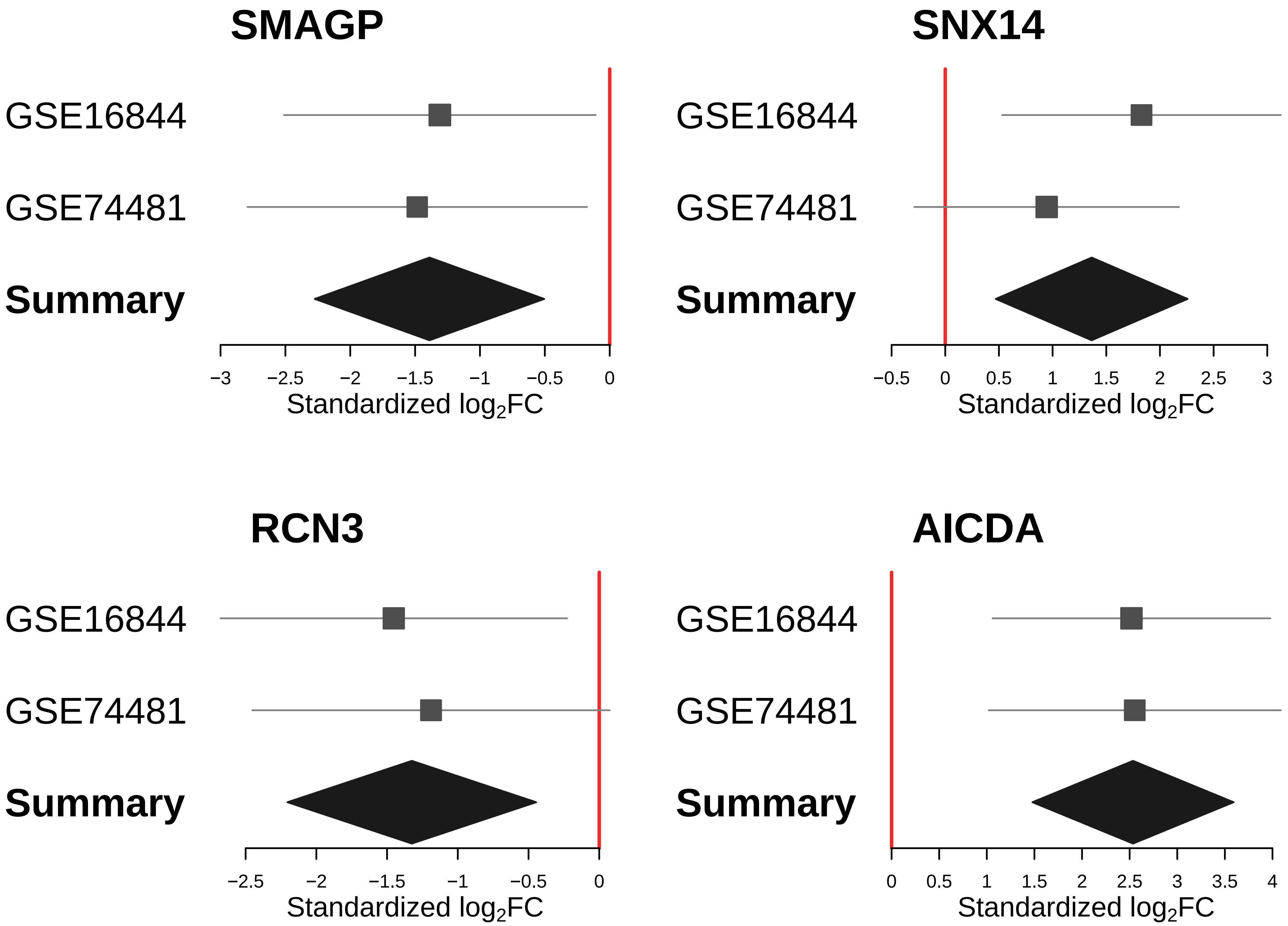
SMAGP
GSE16844
GSE74481
Summary
−3 −2.5 −2 −1.5 −1 −0.5 0
Standardized log2FC
SNX14
GSE16844
GSE74481
Summary
−0.5 0 0.5 1 1.5 2 2.5 3
Standardized log2FC
RCN3
GSE16844
GSE74481
Summary
−2.5 −2 −1.5 −1 −0.5 0
Standardized log2FC
AICDA
GSE16844
GSE74481
Summary
0 0.5 1 1.5 2 2.5 3 3.5 4
Standardized log2FC

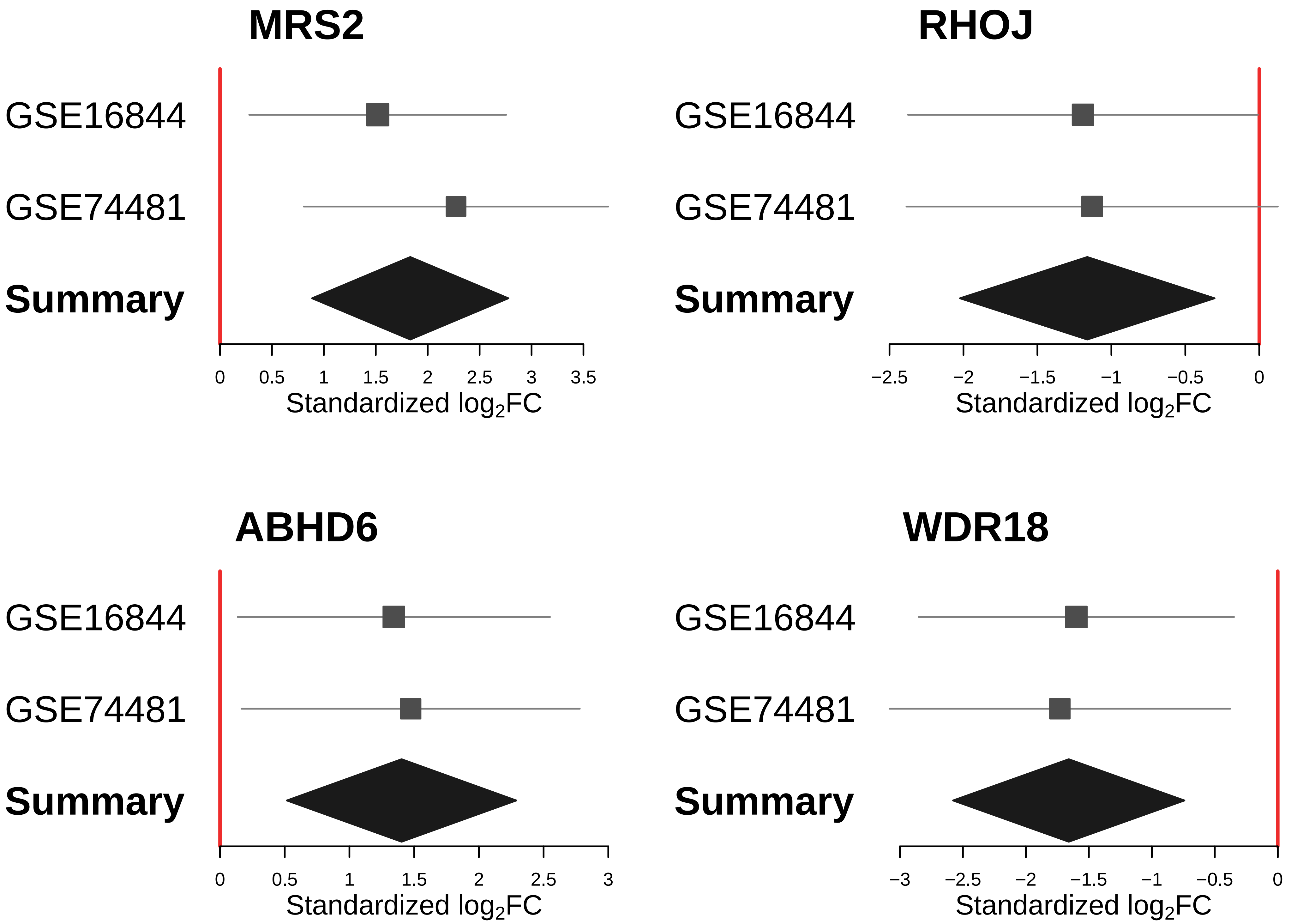
MRS2
GSE16844
GSE74481
Summary
0 0.5 1 1.5 2 2.5 3 3.5
Standardized log2FC
RHOJ
GSE16844
GSE74481
Summary
−2.5 −2 −1.5 −1 −0.5 0
Standardized log2FC
ABHD6
GSE16844
GSE74481
Summary
0 0.5 1 1.5 2 2.5 3
Standardized log2FC
WDR18
GSE16844
GSE74481
Summary
−3 −2.5 −2 −1.5 −1 −0.5 0
Standardized log2FC

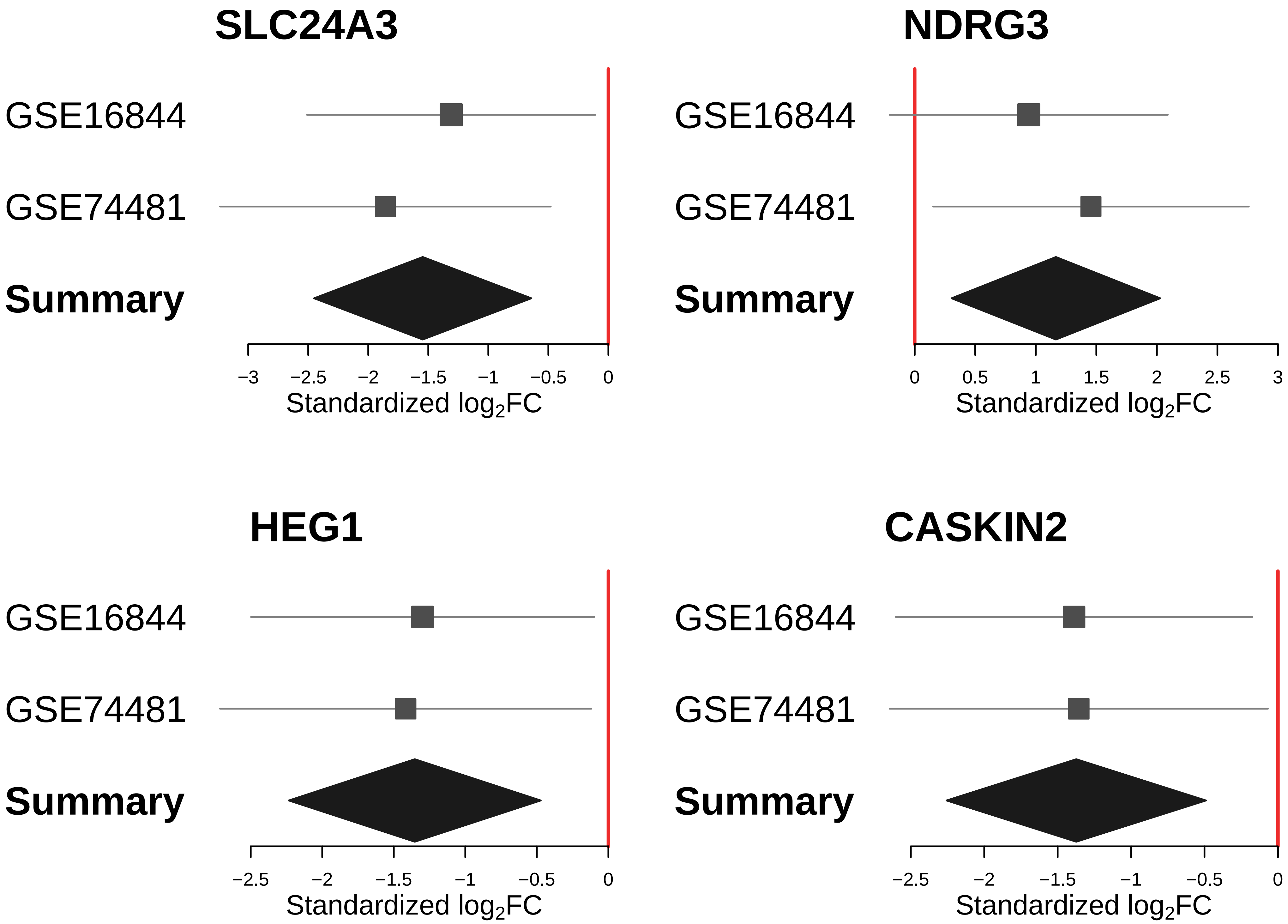
SLC24A3
GSE16844
GSE74481
Summary
−3 −2.5 −2 −1.5 −1 −0.5 0
Standardized log$_2$FC
NDRG3
GSE16844
GSE74481
Summary
0 0.5 1 1.5 2 2.5 3
Standardized log$_2$FC
HEG1
GSE16844
GSE74481
Summary
−2.5 −2 −1.5 −1 −0.5 0
Standardized log$_2$FC
CASKIN2
GSE16844
GSE74481
Summary
−2.5 −2 −1.5 −1 −0.5 0
Standardized log$_2$FC

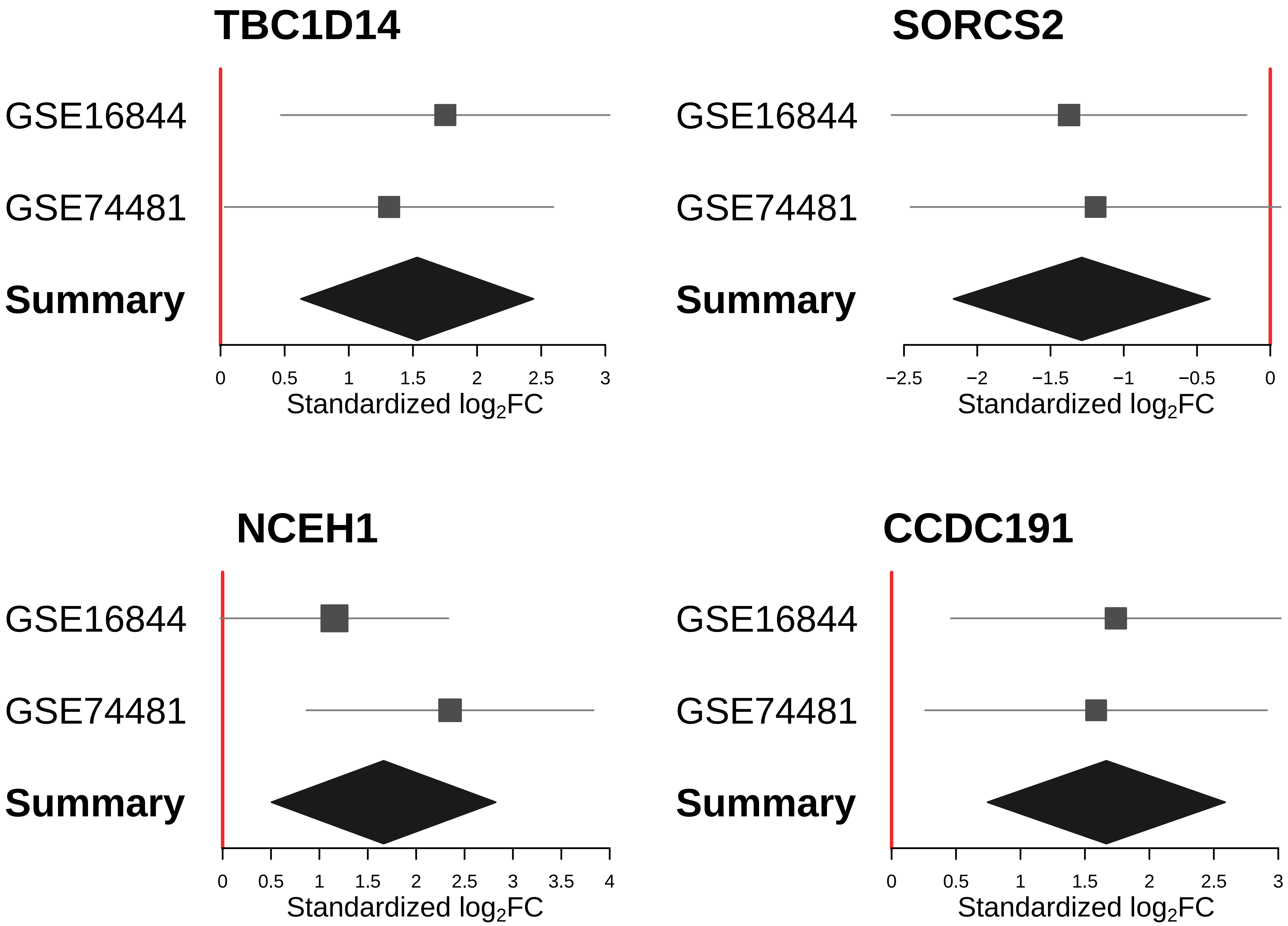
TBC1D14
GSE16844
GSE74481
Summary
0 0.5 1 1.5 2 2.5 3
Standardized log2FC
SORCS2
GSE16844
GSE74481
Summary
−2.5 −2 −1.5 −1 −0.5 0
Standardized log2FC
NCEH1
GSE16844
GSE74481
Summary
0 0.5 1 1.5 2 2.5 3 3.5 4
Standardized log2FC
CCDC191
GSE16844
GSE74481
Summary
0 0.5 1 1.5 2 2.5 3
Standardized log2FC

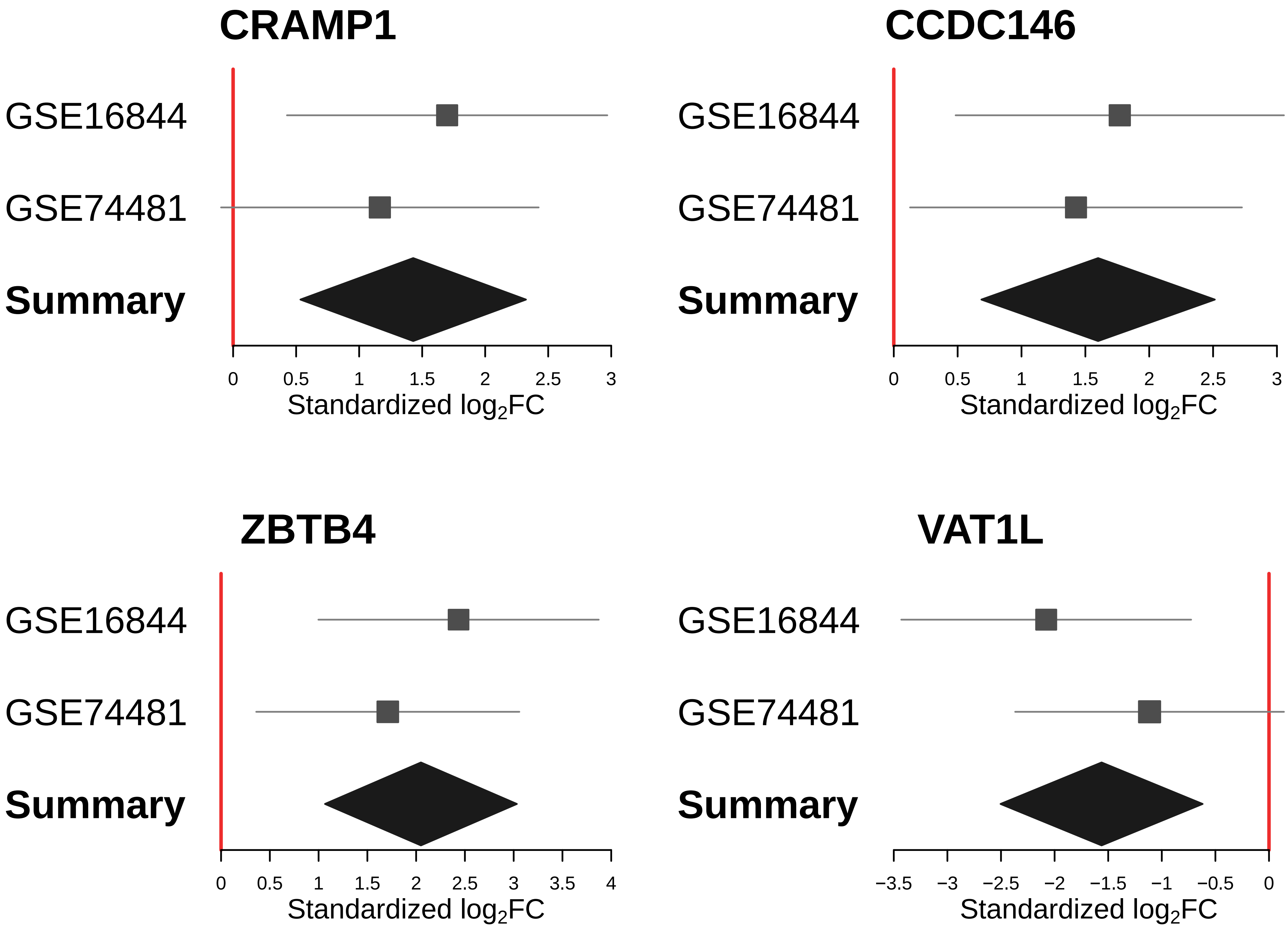
CRAMP1
GSE16844
GSE74481
Summary
0
0.5
1
1.5
2
2.5
3
Standardized log2FC
CCDC146
GSE16844
GSE74481
Summary
0
0.5
1
1.5
2
2.5
3
Standardized log2FC
ZBTB4
GSE16844
GSE74481
Summary
0
0.5
1
1.5
2
2.5
3
3.5
4
Standardized log2FC
VAT1L
GSE16844
GSE74481
Summary
−3.5
−3
−2.5
−2
−1.5
−1
−0.5
0
Standardized log2FC

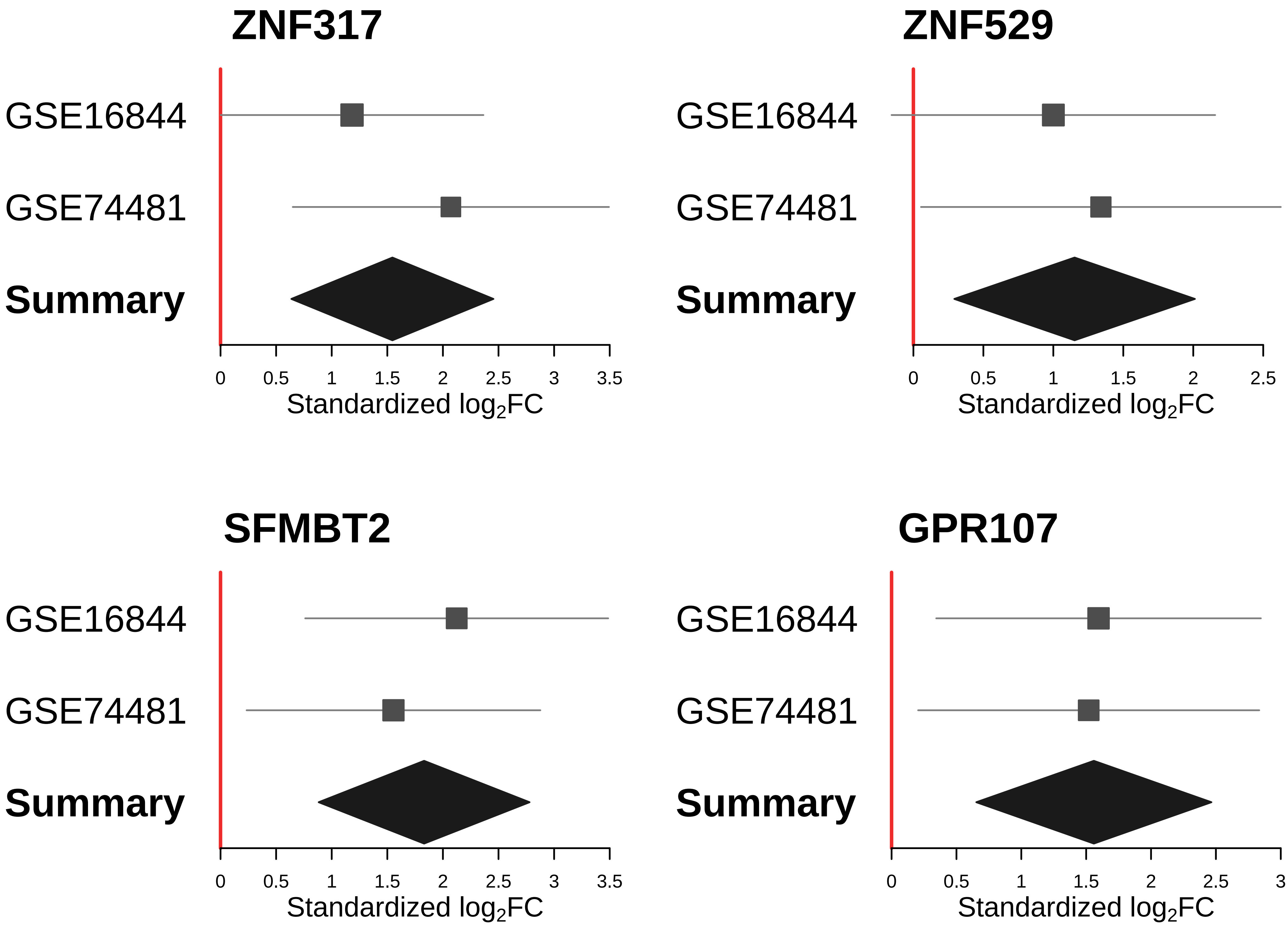
ZNF317
GSE16844
GSE74481
Summary
0
0.5
1
1.5
2
2.5
3
3.5
Standardized log2FC
ZNF529
GSE16844
GSE74481
Summary
0
0.5
1
1.5
2
2.5
Standardized log2FC
SFMBT2
GSE16844
GSE74481
Summary
0
0.5
1
1.5
2
2.5
3
3.5
Standardized log2FC
GPR107
GSE16844
GSE74481
Summary
0
0.5
1
1.5
2
2.5
3
Standardized log2FC

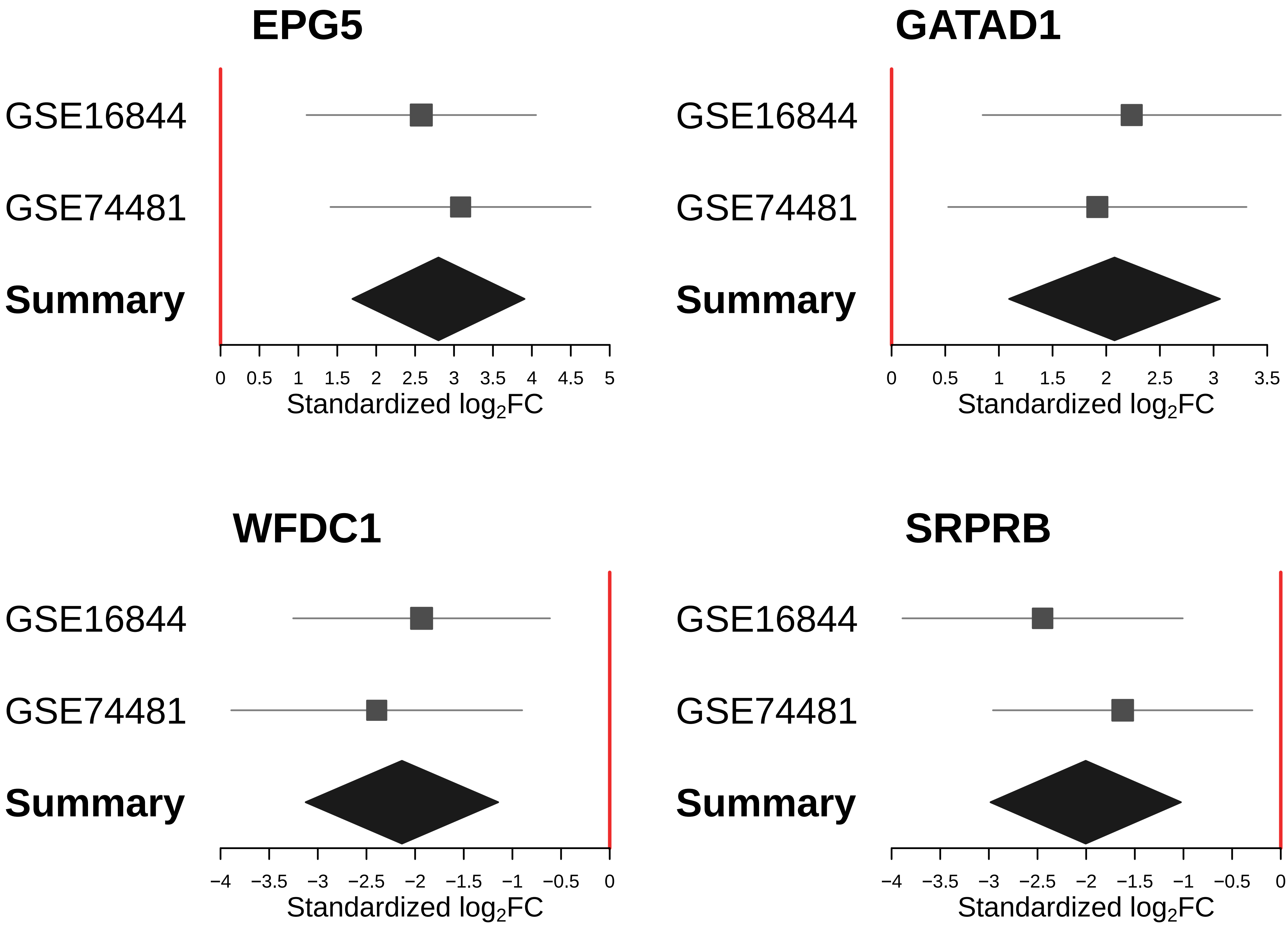

EPG5
GSE16844
GSE74481
Summary
0 0.5 1 1.5 2 2.5 3 3.5 4 4.5 5
Standardized log2FC
GATAD1
GSE16844
GSE74481
Summary
0 0.5 1 1.5 2 2.5 3 3.5
Standardized log2FC
WFDC1
GSE16844
GSE74481
Summary
−4 −3.5 −3 −2.5 −2 −1.5 −1 −0.5 0
Standardized log2FC
SRPRB
GSE16844
GSE74481
Summary
−4 −3.5 −3 −2.5 −2 −1.5 −1 −0.5 0
Standardized log2FC

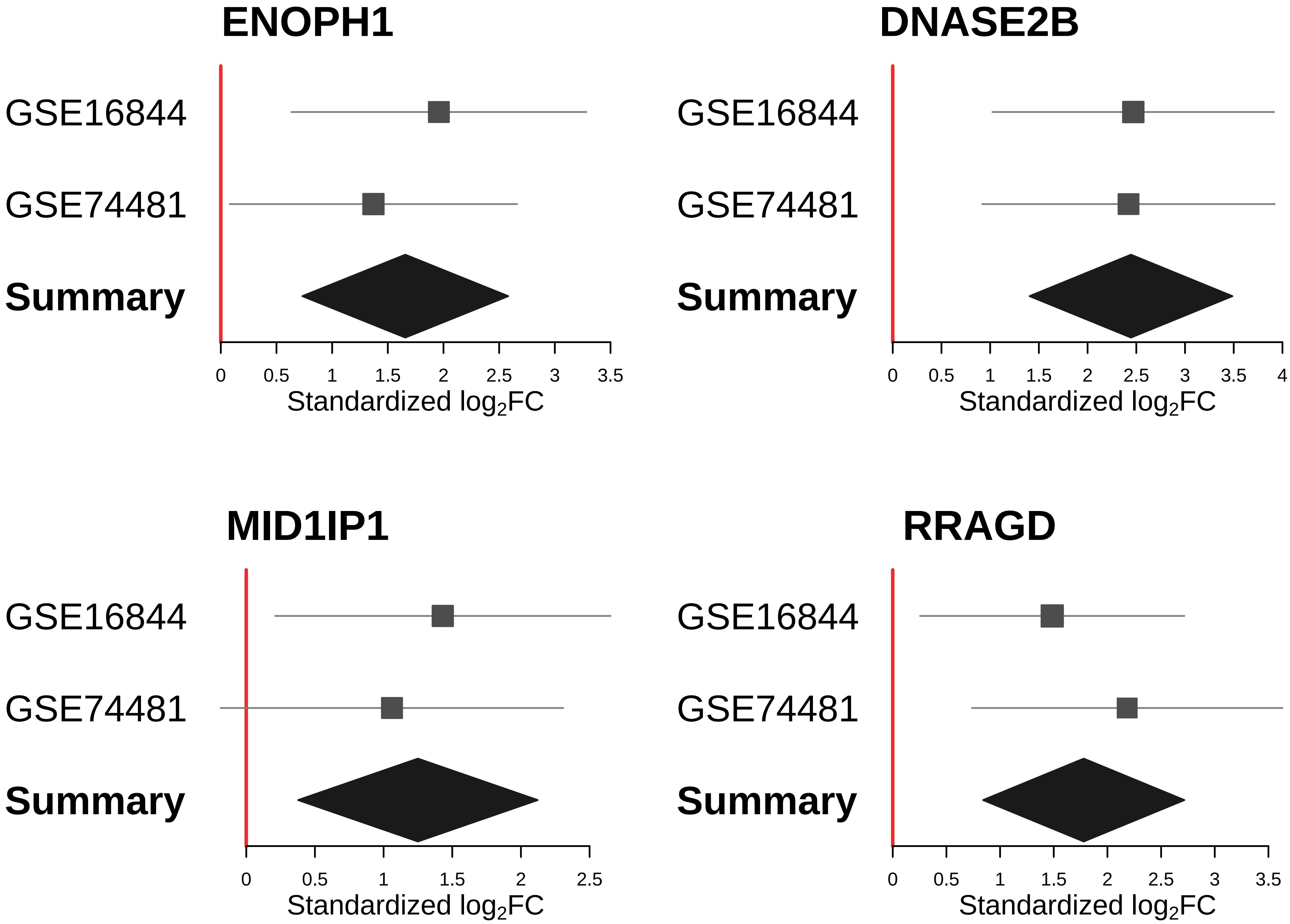
ENOPH1
GSE16844
GSE74481
Summary
0 0.5 1 1.5 2 2.5 3 3.5
Standardized log2FC
DNASE2B
GSE16844
GSE74481
Summary
0 0.5 1 1.5 2 2.5 3 3.5 4
Standardized log2FC
MID1IP1
GSE16844
GSE74481
Summary
0 0.5 1 1.5 2 2.5
Standardized log2FC
RRAGD
GSE16844
GSE74481
Summary
0 0.5 1 1.5 2 2.5 3 3.5
Standardized log2FC

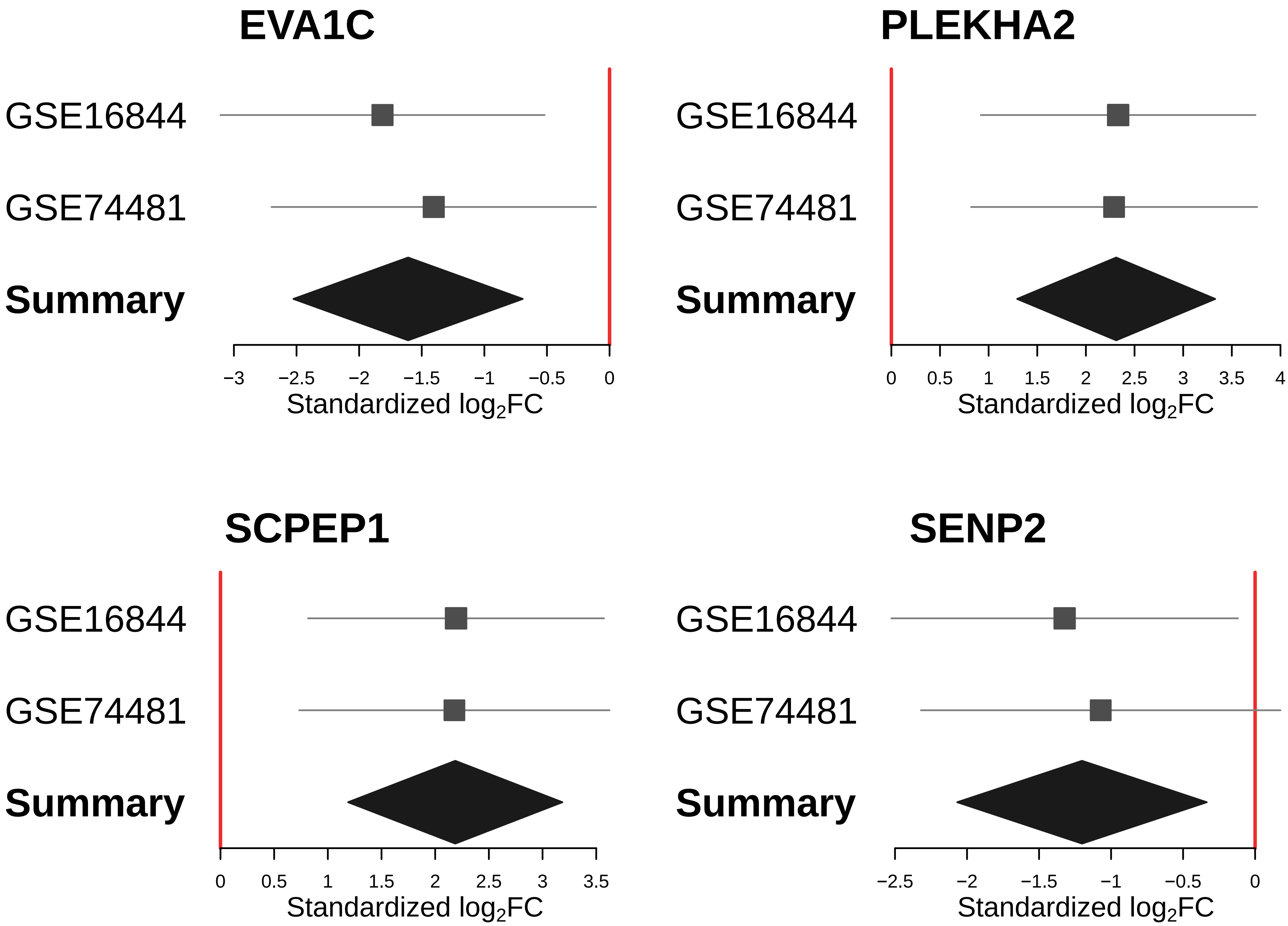
EVA1C
GSE16844
GSE74481
Summary
−3 −2.5 −2 −1.5 −1 −0.5 0
Standardized log2FC
PLEKHA2
GSE16844
GSE74481
Summary
0 0.5 1 1.5 2 2.5 3 3.5 4
Standardized log2FC
SCPEP1
GSE16844
GSE74481
Summary
0 0.5 1 1.5 2 2.5 3 3.5
Standardized log2FC
SENP2
GSE16844
GSE74481
Summary
−2.5 −2 −1.5 −1 −0.5 0
Standardized log2FC

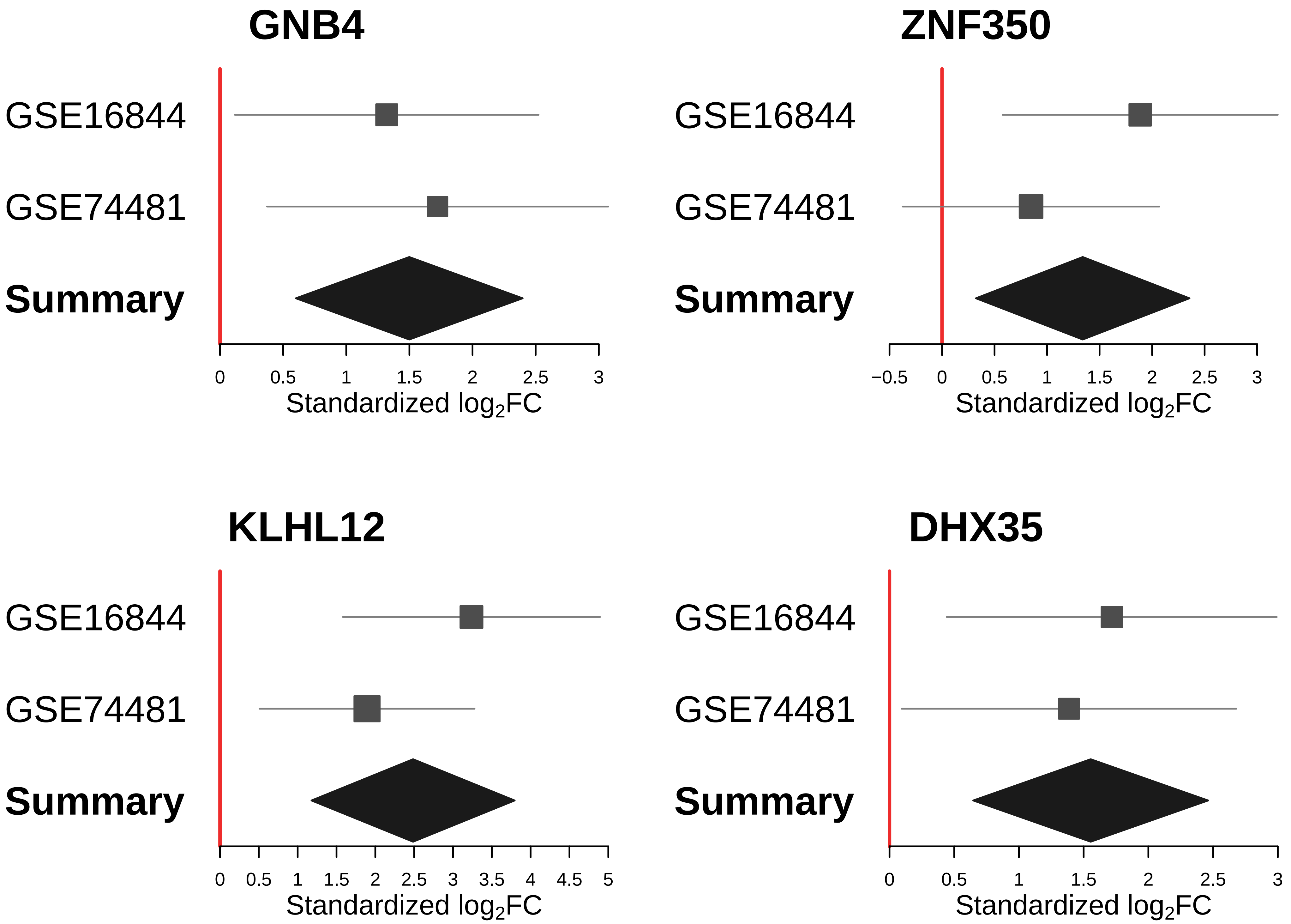
GNB4
GSE16844
GSE74481
Summary
0 0.5 1 1.5 2 2.5 3
Standardized log2FC
ZNF350
GSE16844
GSE74481
Summary
−0.5 0 0.5 1 1.5 2 2.5 3
Standardized log2FC
KLHL12
GSE16844
GSE74481
Summary
0 0.5 1 1.5 2 2.5 3 3.5 4 4.5 5
Standardized log2FC
DHX35
GSE16844
GSE74481
Summary
0 0.5 1 1.5 2 2.5 3
Standardized log2FC

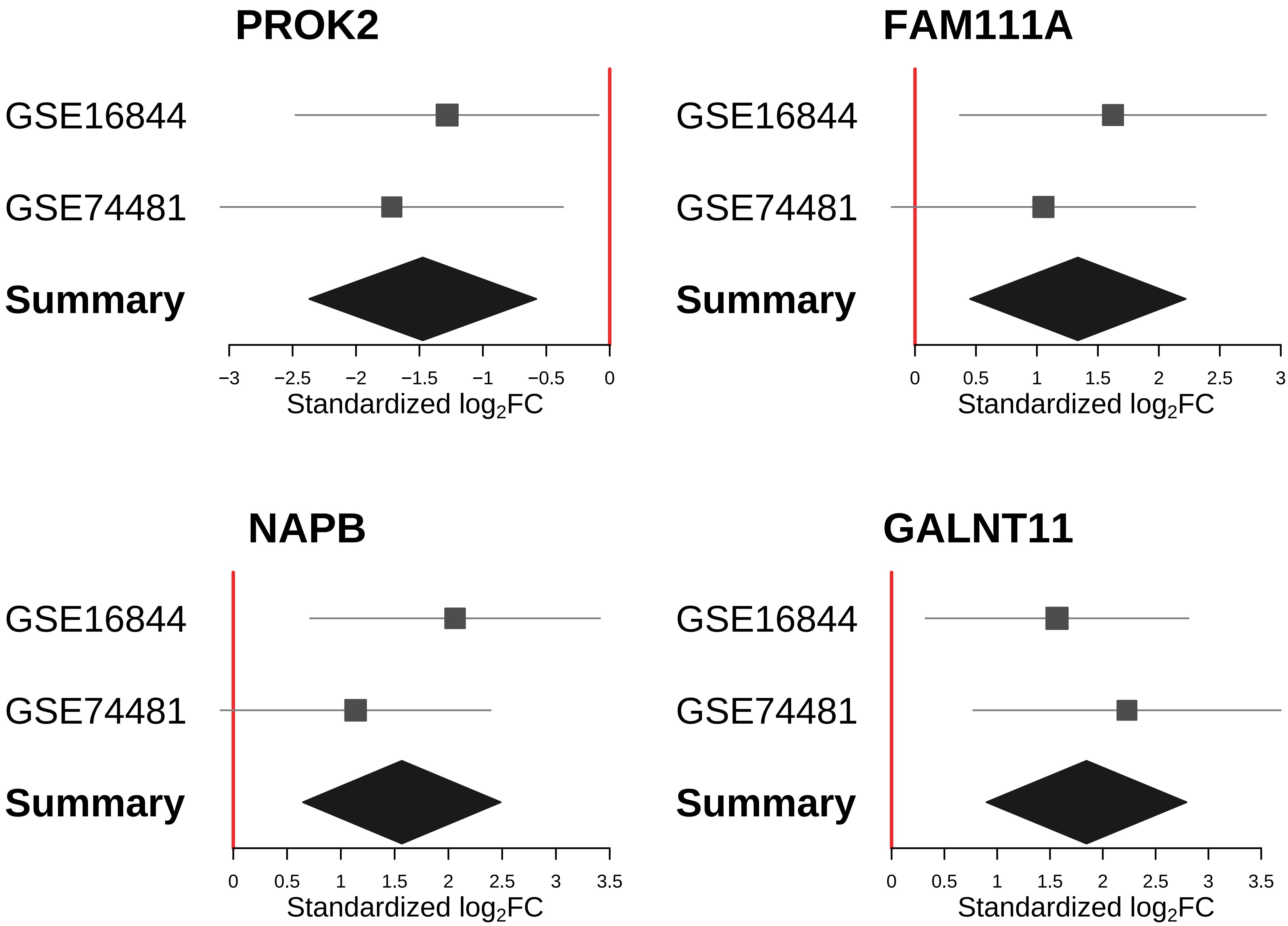
PROK2
GSE16844
GSE74481
Summary
−3
−2.5
−2
−1.5
−1
−0.5
0
Standardized log2FC
FAM111A
GSE16844
GSE74481
Summary
0
0.5
1
1.5
2
2.5
3
Standardized log2FC
NAPB
GSE16844
GSE74481
Summary
0
0.5
1
1.5
2
2.5
3
3.5
Standardized log2FC
GALNT11
GSE16844
GSE74481
Summary
0
0.5
1
1.5
2
2.5
3
3.5
Standardized log2FC

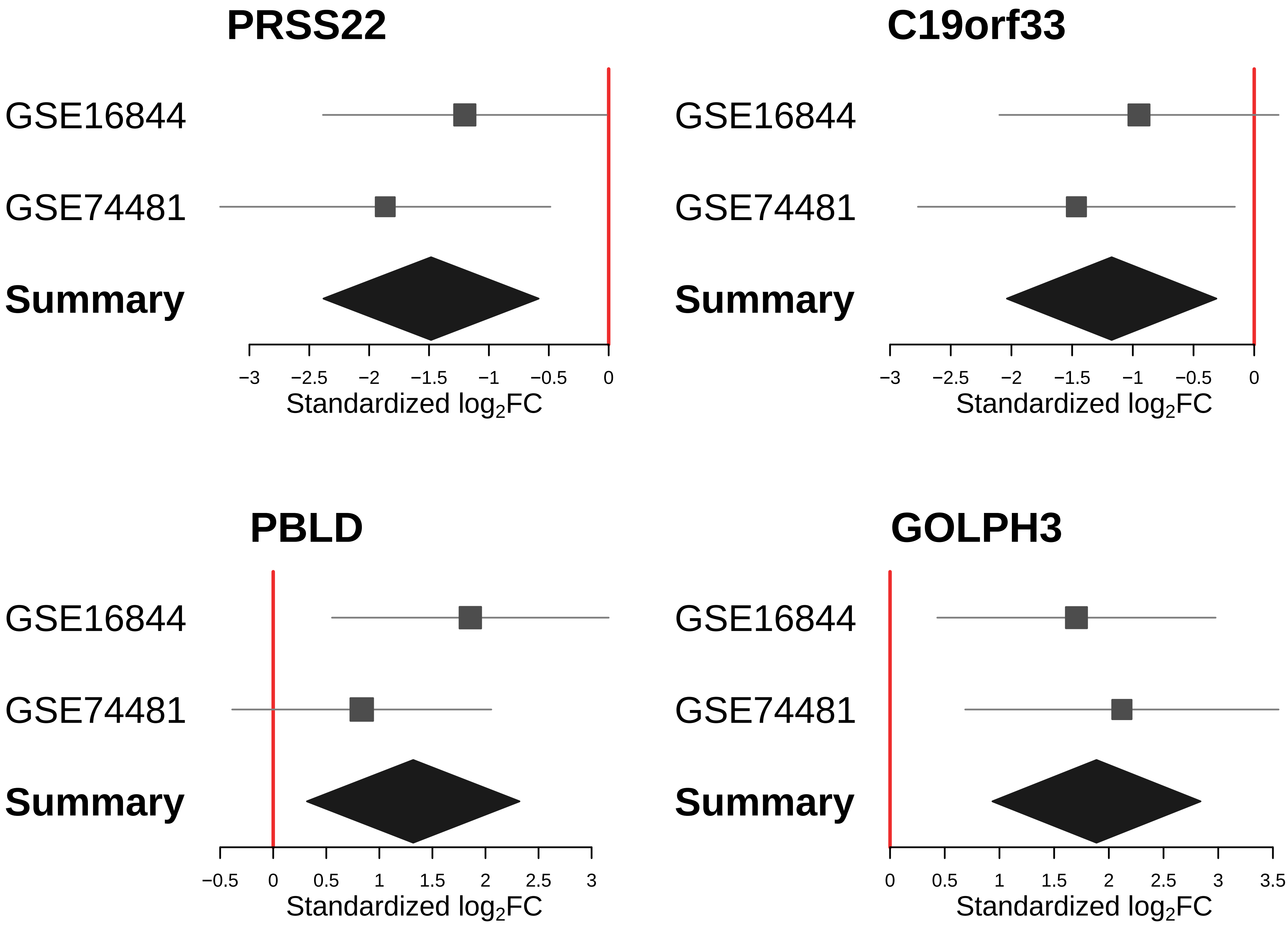

PRSS22
GSE16844
GSE74481
Summary
−3
−2.5
−2
−1.5
−1
−0.5
0
Standardized log2FC
C19orf33
GSE16844
GSE74481
Summary
−3
−2.5
−2
−1.5
−1
−0.5
0
Standardized log2FC
PBLD
GSE16844
GSE74481
Summary
−0.5
0
0.5
1
1.5
2
2.5
3
Standardized log2FC
GOLPH3
GSE16844
GSE74481
Summary
0
0.5
1
1.5
2
2.5
3
3.5
Standardized log2FC

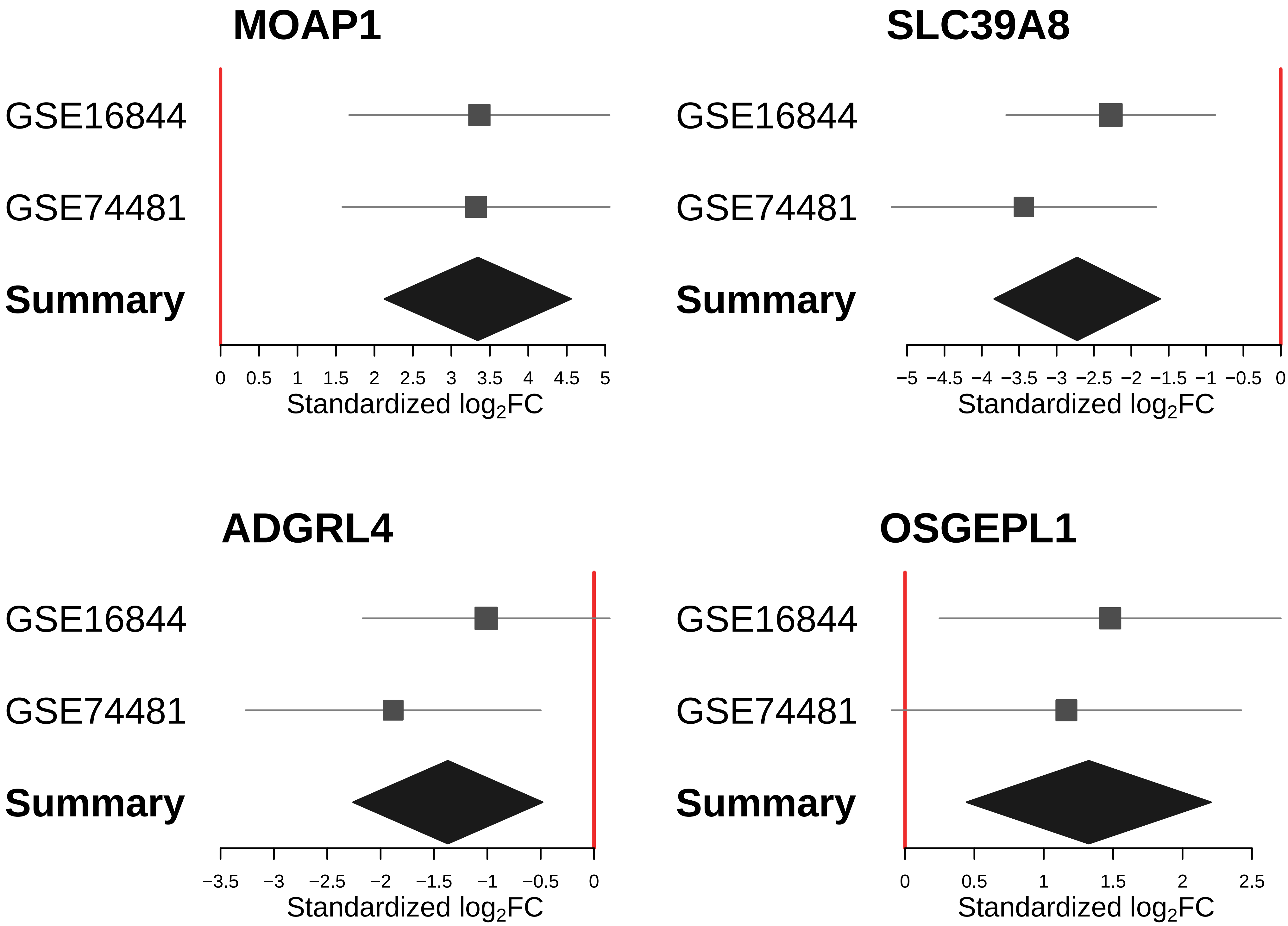

MOAP1
GSE16844
GSE74481
Summary
0
0.5
1
1.5
2
2.5
3
3.5
4
4.5
5
Standardized log2FC
SLC39A8
GSE16844
GSE74481
Summary
−5
−4.5
−4
−3.5
−3
−2.5
−2
−1.5
−1
−0.5
0
Standardized log2FC
ADGRL4
GSE16844
GSE74481
Summary
−3.5
−3
−2.5
−2
−1.5
−1
−0.5
0
Standardized log2FC
OSGEPL1
GSE16844
GSE74481
Summary
0
0.5
1
1.5
2
2.5
Standardized log2FC

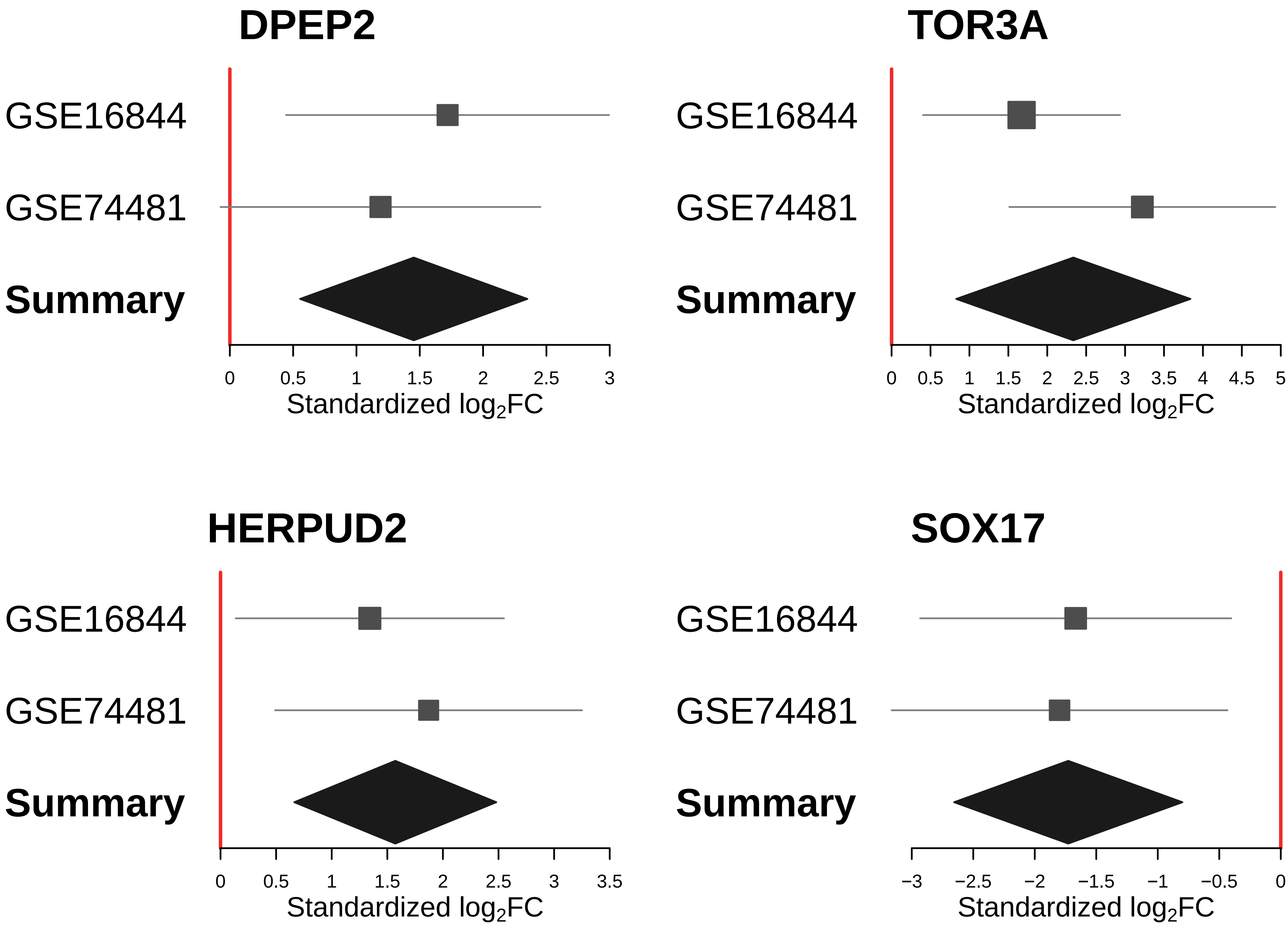
DPEP2
GSE16844
GSE74481
Summary
0 0.5 1 1.5 2 2.5 3
Standardized log2FC
TOR3A
GSE16844
GSE74481
Summary
0 0.5 1 1.5 2 2.5 3 3.5 4 4.5 5
Standardized log2FC
HERPUD2
GSE16844
GSE74481
Summary
0 0.5 1 1.5 2 2.5 3 3.5
Standardized log2FC
SOX17
GSE16844
GSE74481
Summary
−3 −2.5 −2 −1.5 −1 −0.5 0
Standardized log2FC

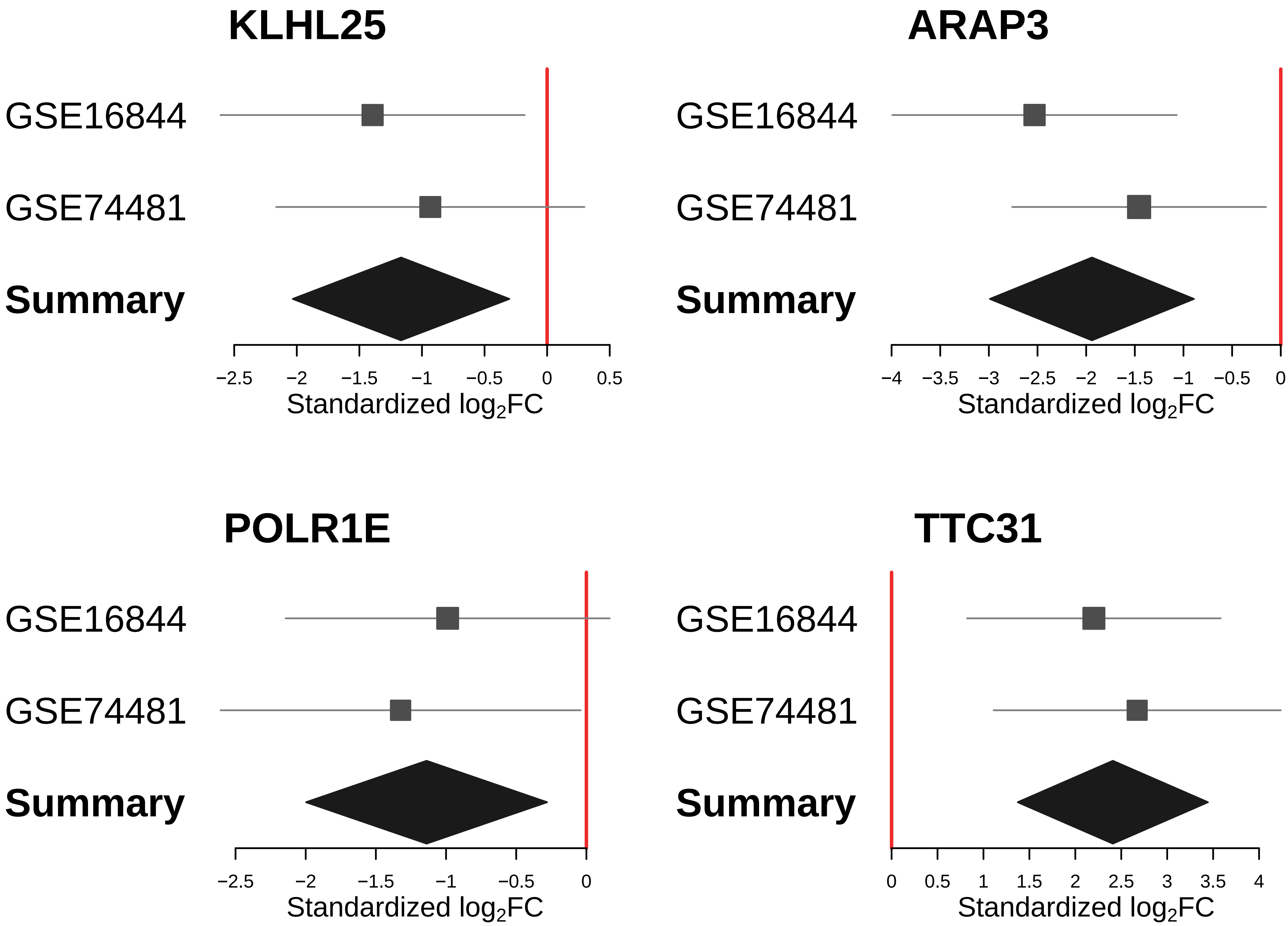
KLHL25
GSE16844
GSE74481
Summary
−2.5 −2 −1.5 −1 −0.5 0 0.5
Standardized log2FC
ARAP3
GSE16844
GSE74481
Summary
−4 −3.5 −3 −2.5 −2 −1.5 −1 −0.5 0
Standardized log2FC
POLR1E
GSE16844
GSE74481
Summary
−2.5 −2 −1.5 −1 −0.5 0
Standardized log2FC
TTC31
GSE16844
GSE74481
Summary
0 0.5 1 1.5 2 2.5 3 3.5 4
Standardized log2FC

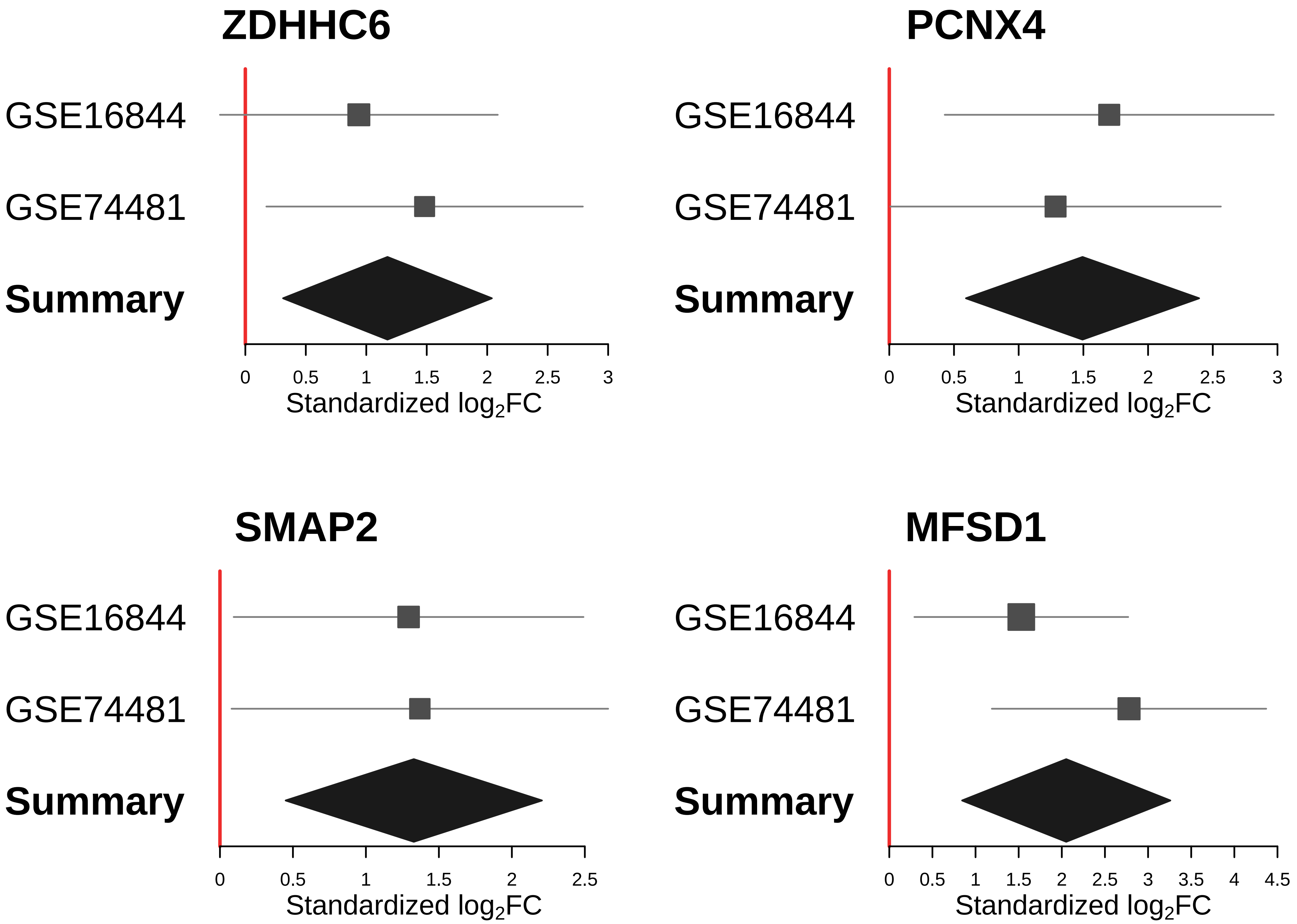
ZDHHC6
GSE16844
GSE74481
Summary
0 0.5 1 1.5 2 2.5 3
Standardized log2FC
PCNX4
GSE16844
GSE74481
Summary
0 0.5 1 1.5 2 2.5 3
Standardized log2FC
SMAP2
GSE16844
GSE74481
Summary
0 0.5 1 1.5 2 2.5
Standardized log2FC
MFSD1
GSE16844
GSE74481
Summary
0 0.5 1 1.5 2 2.5 3 3.5 4 4.5
Standardized log2FC

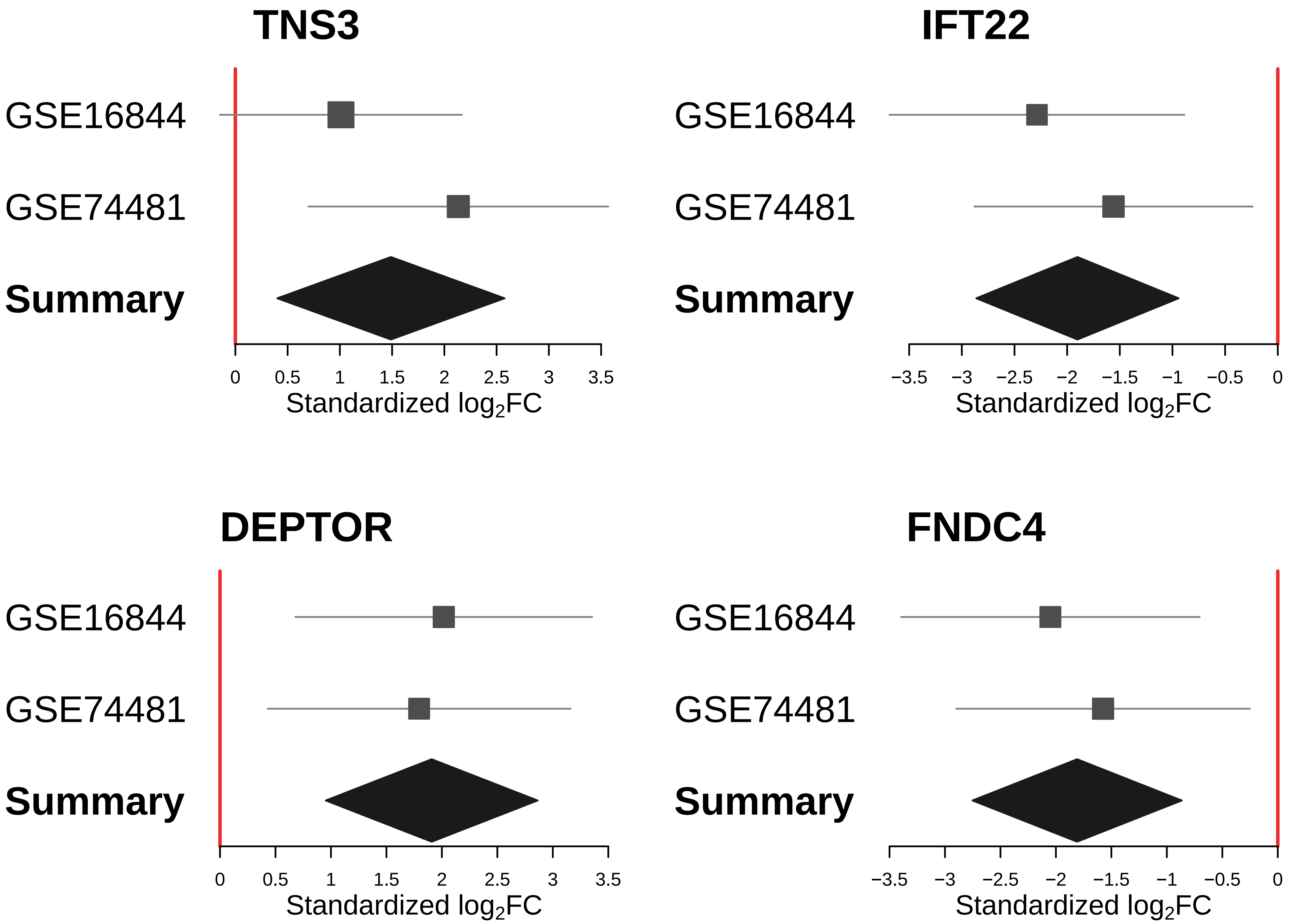
TNS3
GSE16844
GSE74481
Summary
0
0.5
1
1.5
2
2.5
3
3.5
Standardized log2FC
IFT22
GSE16844
GSE74481
Summary
−3.5
−3
−2.5
−2
−1.5
−1
−0.5
0
Standardized log2FC
DEPTOR
GSE16844
GSE74481
Summary
0
0.5
1
1.5
2
2.5
3
3.5
Standardized log2FC
FNDC4
GSE16844
GSE74481
Summary
−3.5
−3
−2.5
−2
−1.5
−1
−0.5
0
Standardized log2FC

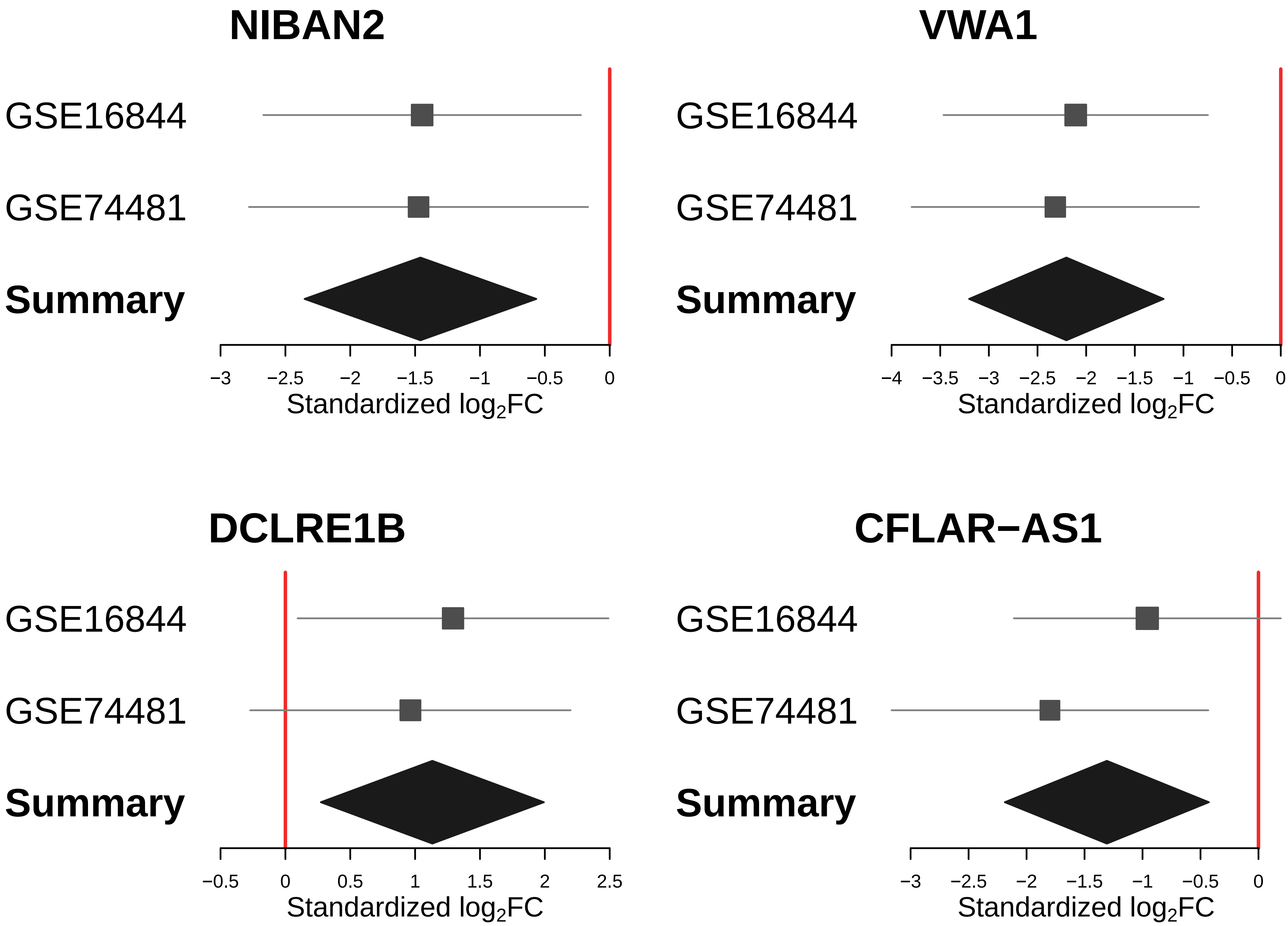

NIBAN2
GSE16844
GSE74481
Summary
−3 −2.5 −2 −1.5 −1 −0.5 0
Standardized log2FC
VWA1
GSE16844
GSE74481
Summary
−4 −3.5 −3 −2.5 −2 −1.5 −1 −0.5 0
Standardized log2FC
DCLRE1B
GSE16844
GSE74481
Summary
−0.5 0 0.5 1 1.5 2 2.5
Standardized log2FC
CFLAR−AS1
GSE16844
GSE74481
Summary
−3 −2.5 −2 −1.5 −1 −0.5 0
Standardized log2FC

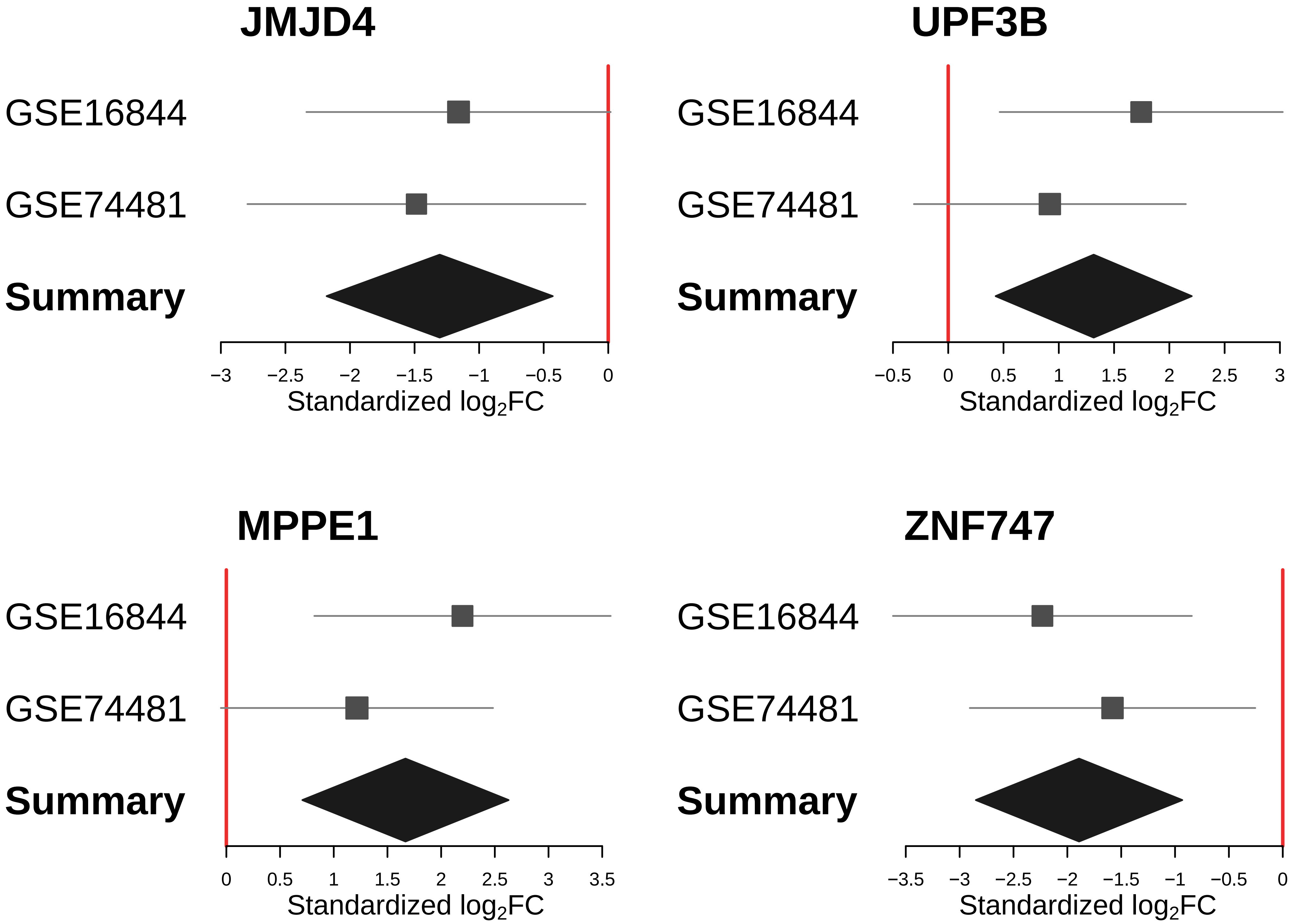
JMJD4
GSE16844
GSE74481
Summary
−3 −2.5 −2 −1.5 −1 −0.5 0
Standardized log2FC
UPF3B
GSE16844
GSE74481
Summary
−0.5 0 0.5 1 1.5 2 2.5 3
Standardized log2FC
MPPE1
GSE16844
GSE74481
Summary
0 0.5 1 1.5 2 2.5 3 3.5
Standardized log2FC
ZNF747
GSE16844
GSE74481
Summary
−3.5 −3 −2.5 −2 −1.5 −1 −0.5 0
Standardized log2FC

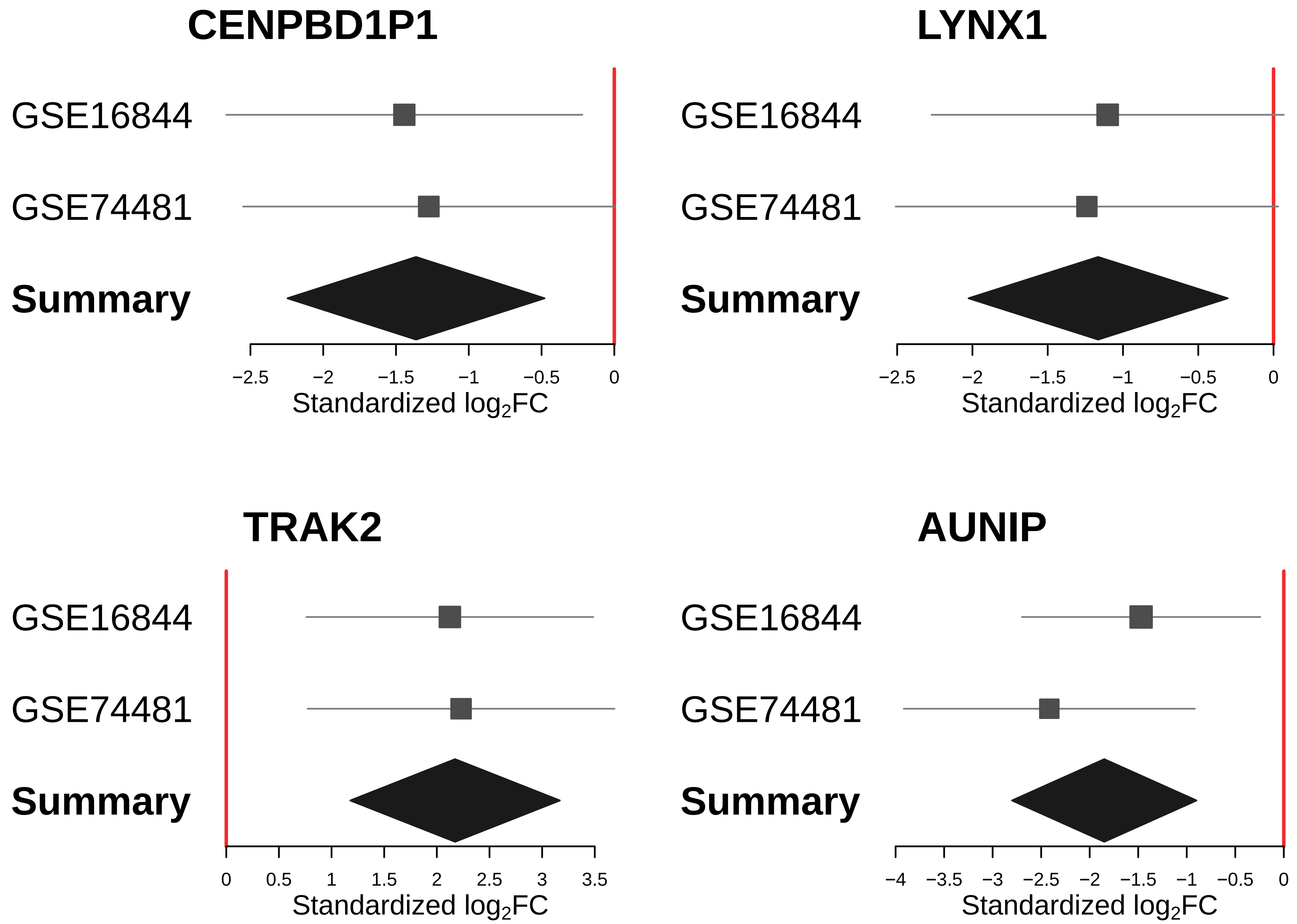
CENPBD1P1
GSE16844
GSE74481
Summary
−2.5 −2 −1.5 −1 −0.5 0
Standardized log2FC
LYNX1
GSE16844
GSE74481
Summary
−2.5 −2 −1.5 −1 −0.5 0
Standardized log2FC
TRAK2
GSE16844
GSE74481
Summary
0 0.5 1 1.5 2 2.5 3 3.5
Standardized log2FC
AUNIP
GSE16844
GSE74481
Summary
−4 −3.5 −3 −2.5 −2 −1.5 −1 −0.5 0
Standardized log2FC

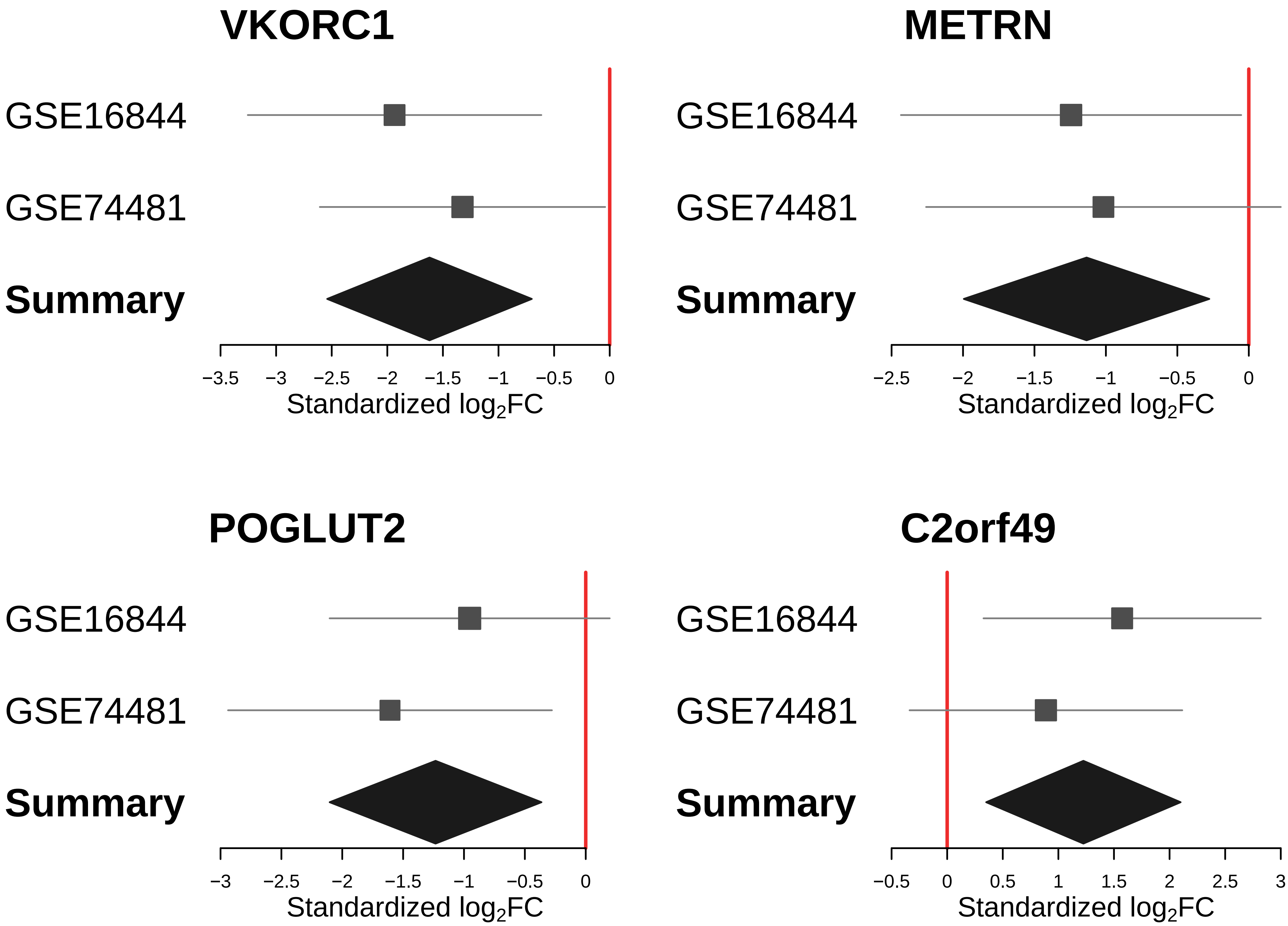

VKORC1
GSE16844
GSE74481
Summary
−3.5 −3 −2.5 −2 −1.5 −1 −0.5 0
Standardized $\log_2$FC
METRN
GSE16844
GSE74481
Summary
−2.5 −2 −1.5 −1 −0.5 0
Standardized $\log_2$FC
POGLUT2
GSE16844
GSE74481
Summary
−3 −2.5 −2 −1.5 −1 −0.5 0
Standardized $\log_2$FC
C2orf49
GSE16844
GSE74481
Summary
−0.5 0 0.5 1 1.5 2 2.5 3
Standardized $\log_2$FC

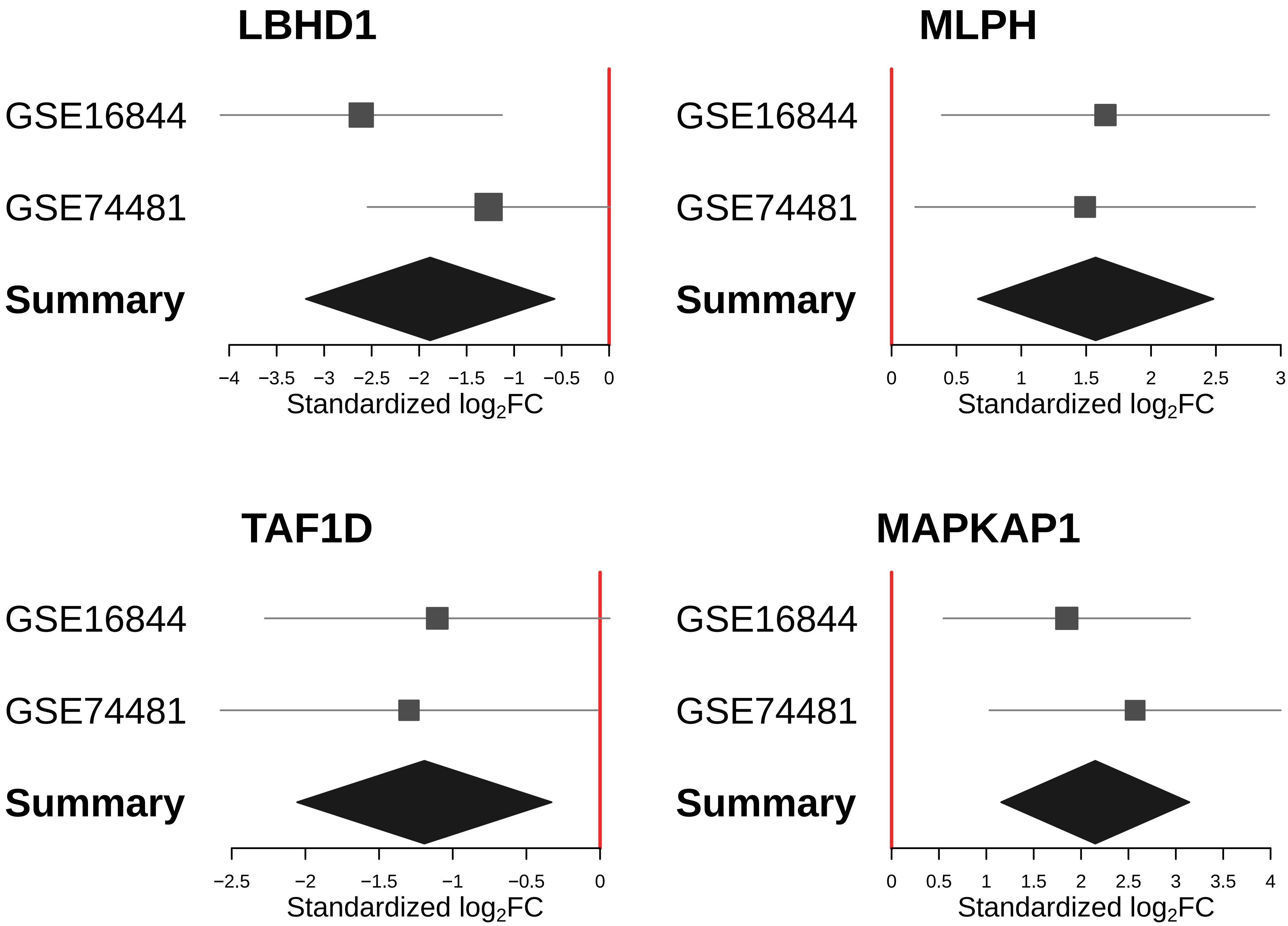
LBHD1
GSE16844
GSE74481
Summary
−4 −3.5 −3 −2.5 −2 −1.5 −1 −0.5 0
Standardized log2FC
MLPH
GSE16844
GSE74481
Summary
0 0.5 1 1.5 2 2.5 3
Standardized log2FC
TAF1D
GSE16844
GSE74481
Summary
−2.5 −2 −1.5 −1 −0.5 0
Standardized log2FC
MAPKAP1
GSE16844
GSE74481
Summary
0 0.5 1 1.5 2 2.5 3 3.5 4
Standardized log2FC

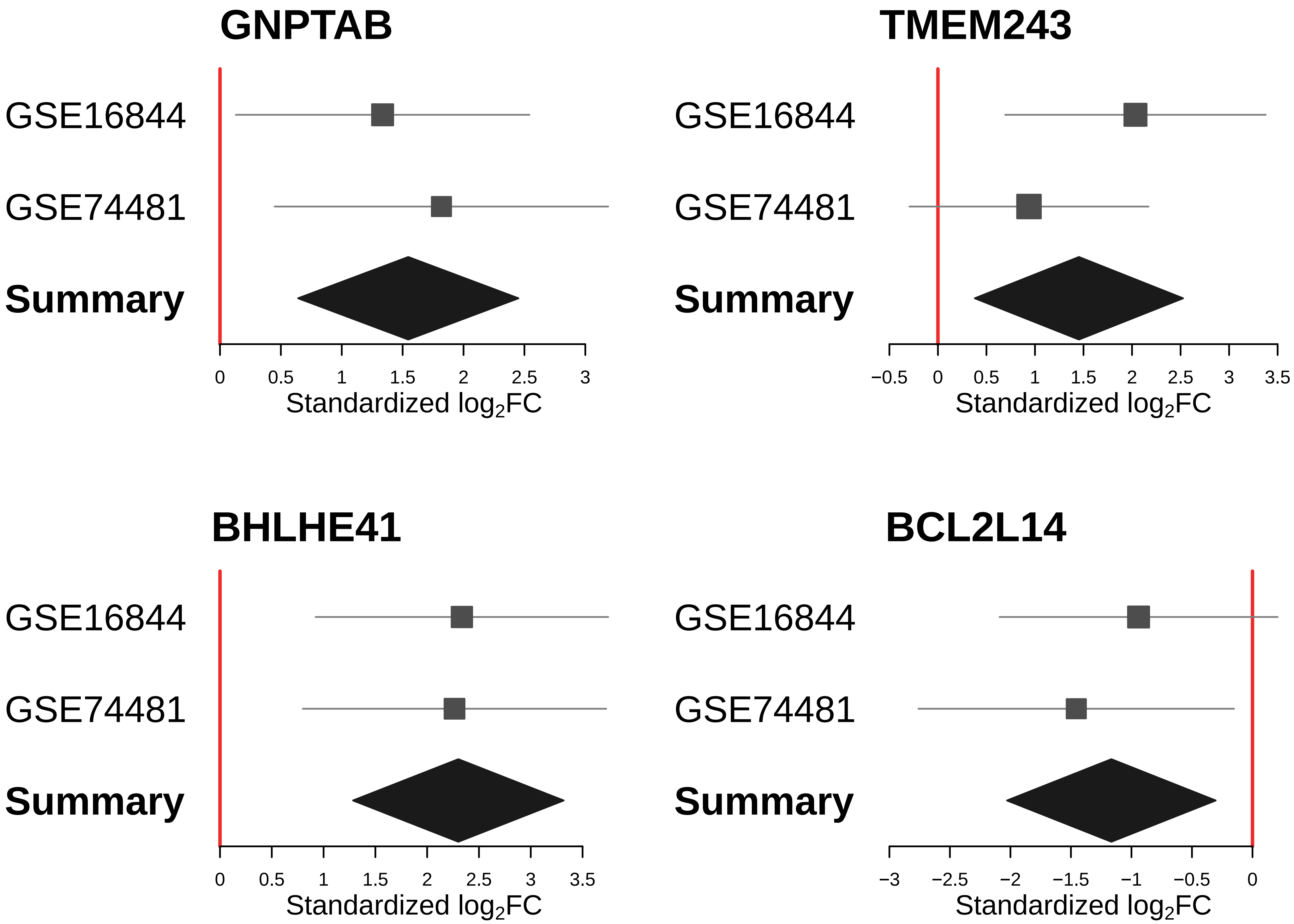
GNPTAB
GSE16844
GSE74481
Summary
0
0.5
1
1.5
2
2.5
3
Standardized log2FC
TMEM243
GSE16844
GSE74481
Summary
−0.5
0
0.5
1
1.5
2
2.5
3
3.5
Standardized log2FC
BHLHE41
GSE16844
GSE74481
Summary
0
0.5
1
1.5
2
2.5
3
3.5
Standardized log2FC
BCL2L14
GSE16844
GSE74481
Summary
−3
−2.5
−2
−1.5
−1
−0.5
0
Standardized log2FC

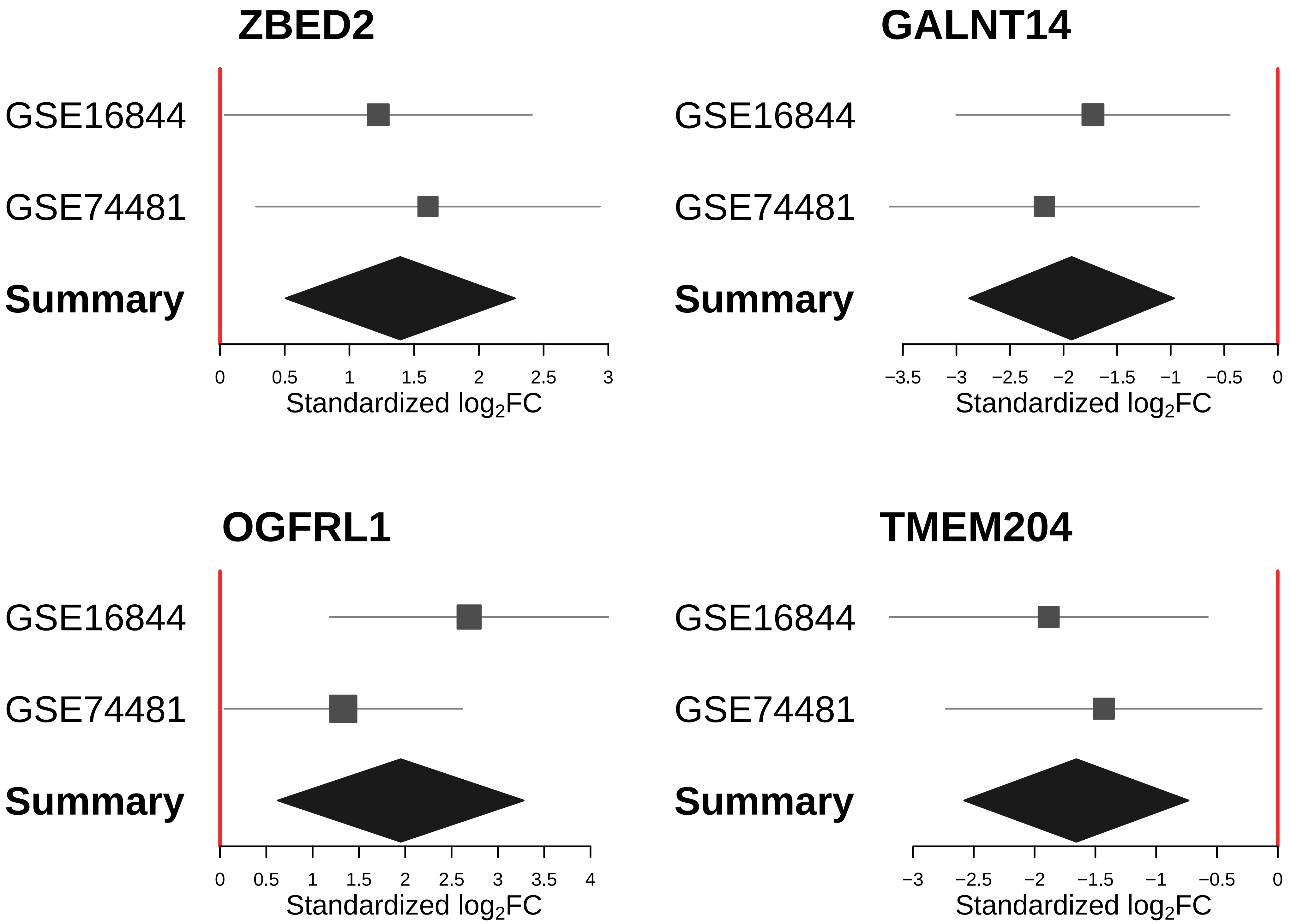
ZBED2
GSE16844
GSE74481
Summary
0
0.5
1
1.5
2
2.5
3
Standardized $log_2FC$
GALNT14
GSE16844
GSE74481
Summary
−3.5
−3
−2.5
−2
−1.5
−1
−0.5
0
Standardized $log_2FC$
OGFRL1
GSE16844
GSE74481
Summary
0
0.5
1
1.5
2
2.5
3
3.5
4
Standardized $log_2FC$
TMEM204
GSE16844
GSE74481
Summary
−3
−2.5
−2
−1.5
−1
−0.5
0
Standardized $log_2FC$

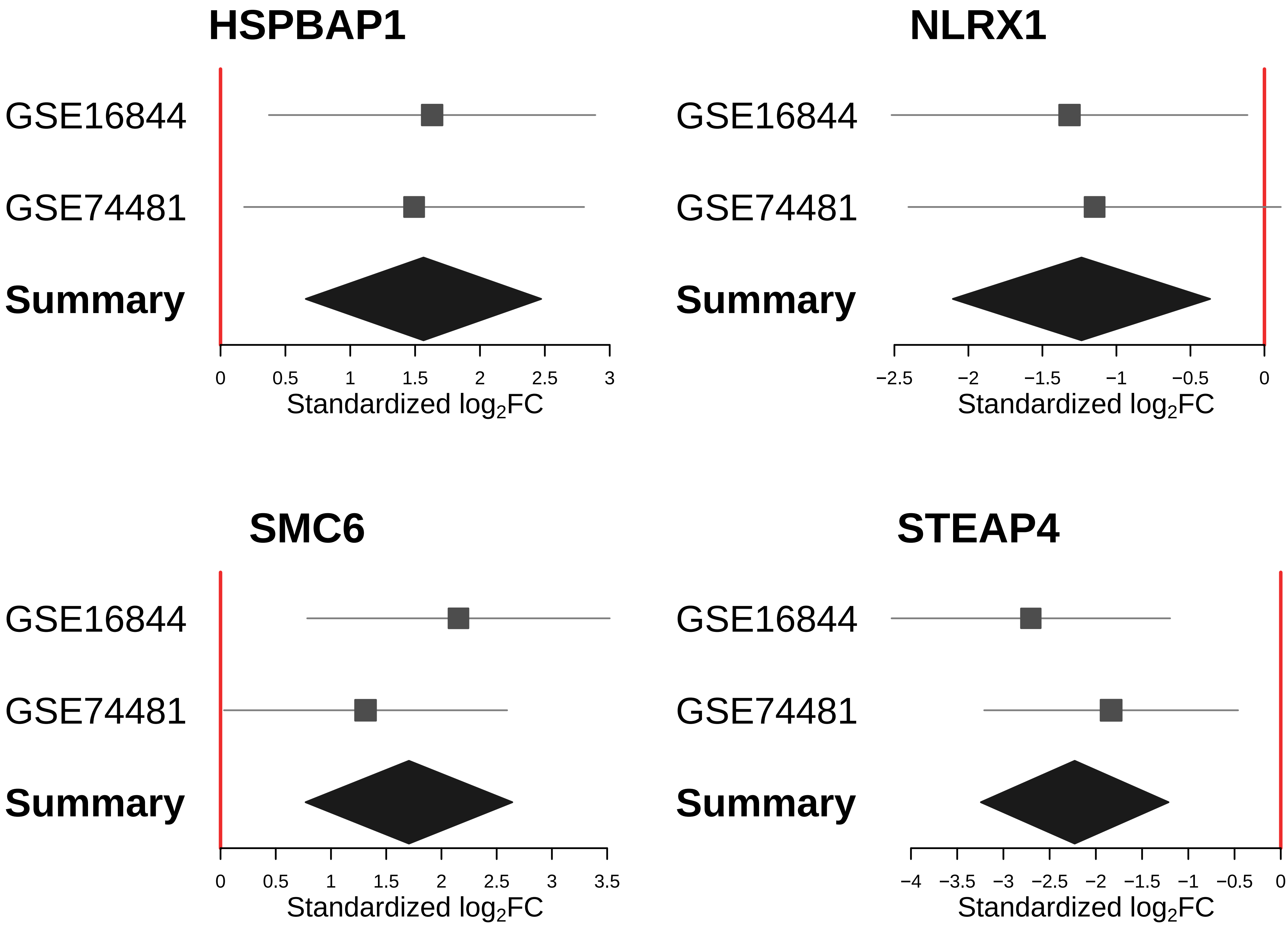
HSPBAP1
GSE16844
GSE74481
Summary
0
0.5
1
1.5
2
2.5
3
Standardized log2FC
NLRX1
GSE16844
GSE74481
Summary
−2.5
−2
−1.5
−1
−0.5
0
Standardized log2FC
SMC6
GSE16844
GSE74481
Summary
0
0.5
1
1.5
2
2.5
3
3.5
Standardized log2FC
STEAP4
GSE16844
GSE74481
Summary
−4
−3.5
−3
−2.5
−2
−1.5
−1
−0.5
0
Standardized log2FC

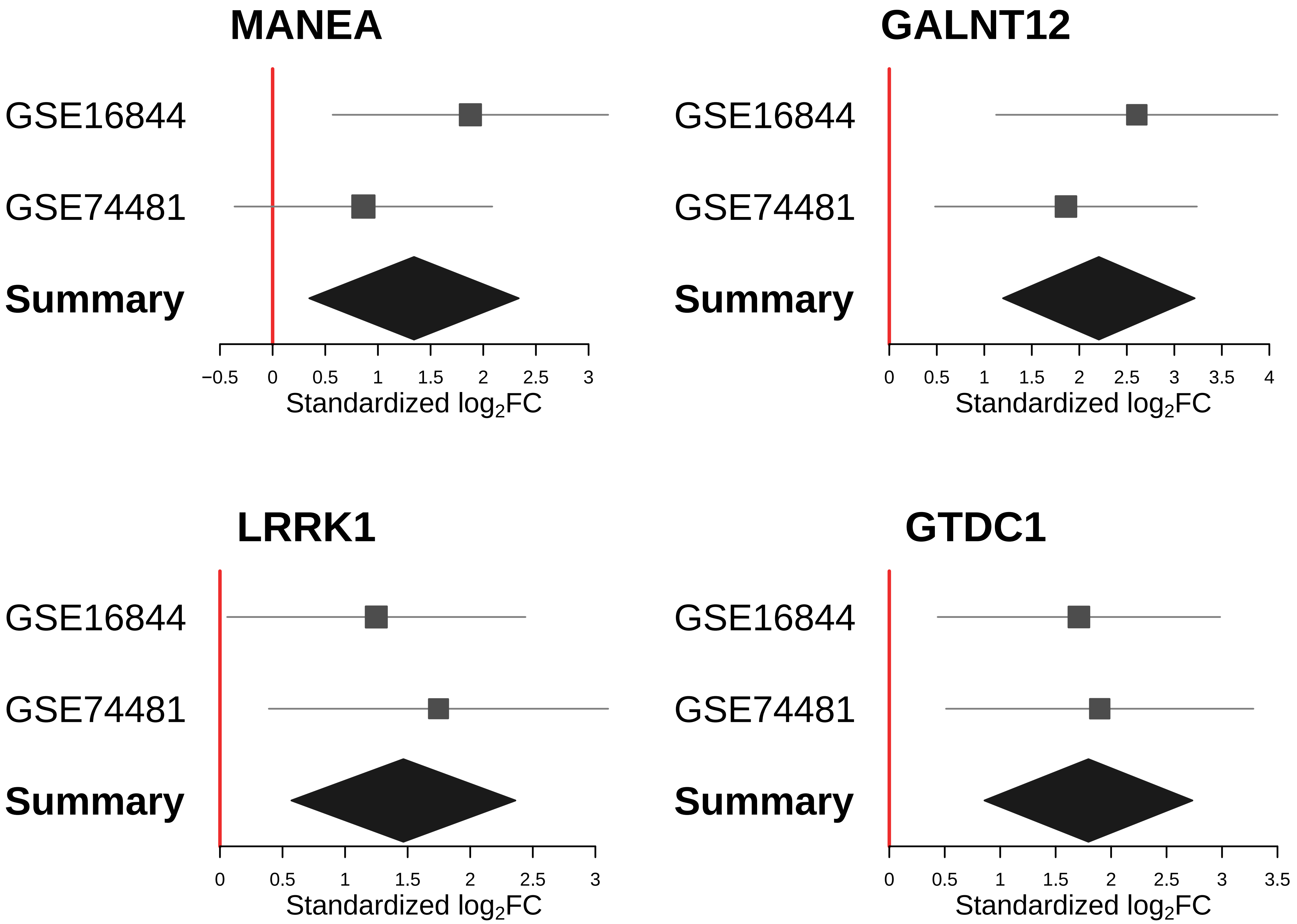
MANEA
GSE16844
GSE74481
Summary
−0.5 0 0.5 1 1.5 2 2.5 3
Standardized log2FC
GALNT12
GSE16844
GSE74481
Summary
0 0.5 1 1.5 2 2.5 3 3.5 4
Standardized log2FC
LRRK1
GSE16844
GSE74481
Summary
0 0.5 1 1.5 2 2.5 3
Standardized log2FC
GTDC1
GSE16844
GSE74481
Summary
0 0.5 1 1.5 2 2.5 3 3.5
Standardized log2FC

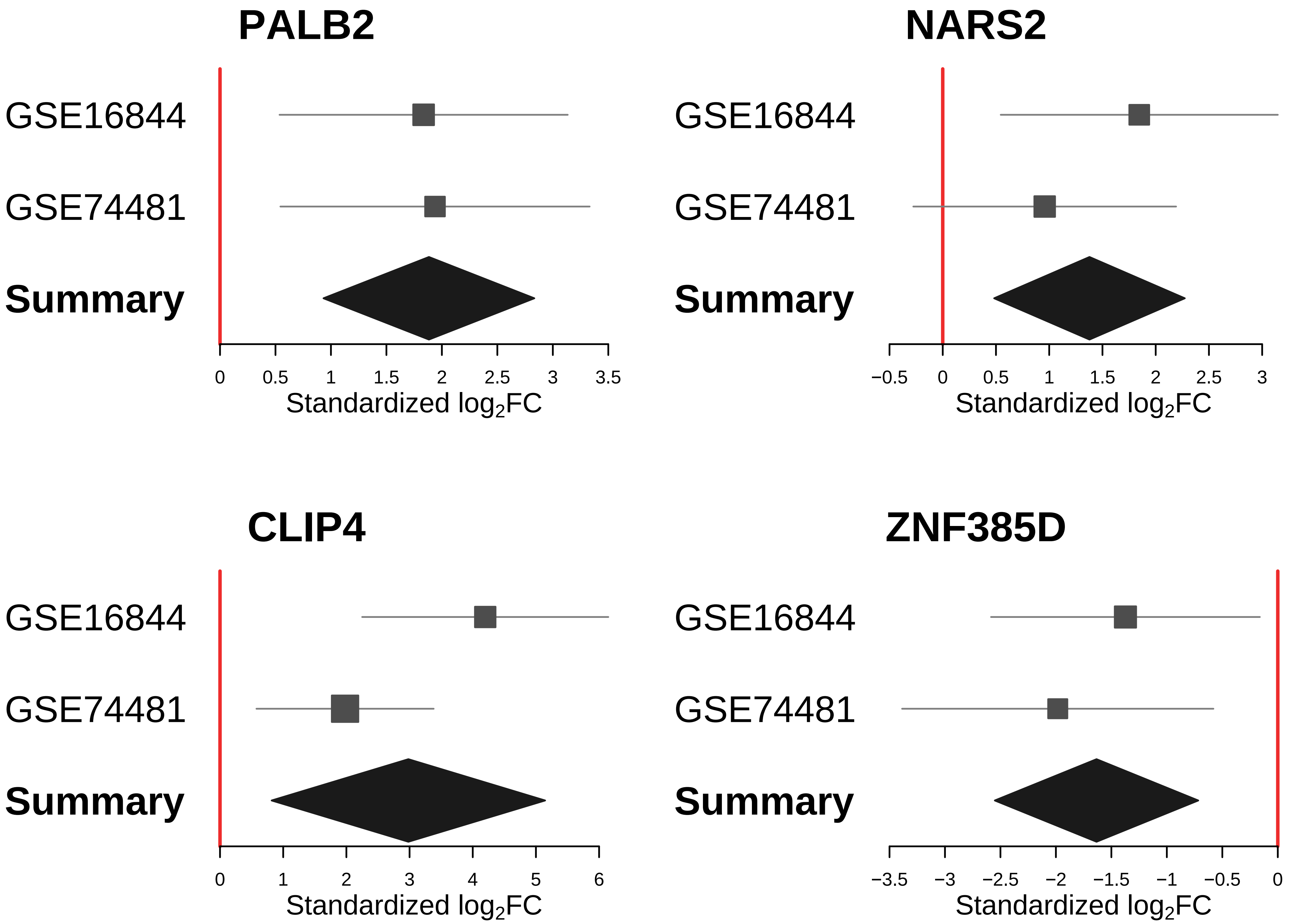
PALB2
GSE16844
GSE74481
Summary
0
0.5
1
1.5
2
2.5
3
3.5
Standardized log2FC
NARS2
GSE16844
GSE74481
Summary
−0.5
0
0.5
1
1.5
2
2.5
3
Standardized log2FC
CLIP4
GSE16844
GSE74481
Summary
0
1
2
3
4
5
6
Standardized log2FC
ZNF385D
GSE16844
GSE74481
Summary
−3.5
−3
−2.5
−2
−1.5
−1
−0.5
0
Standardized log2FC

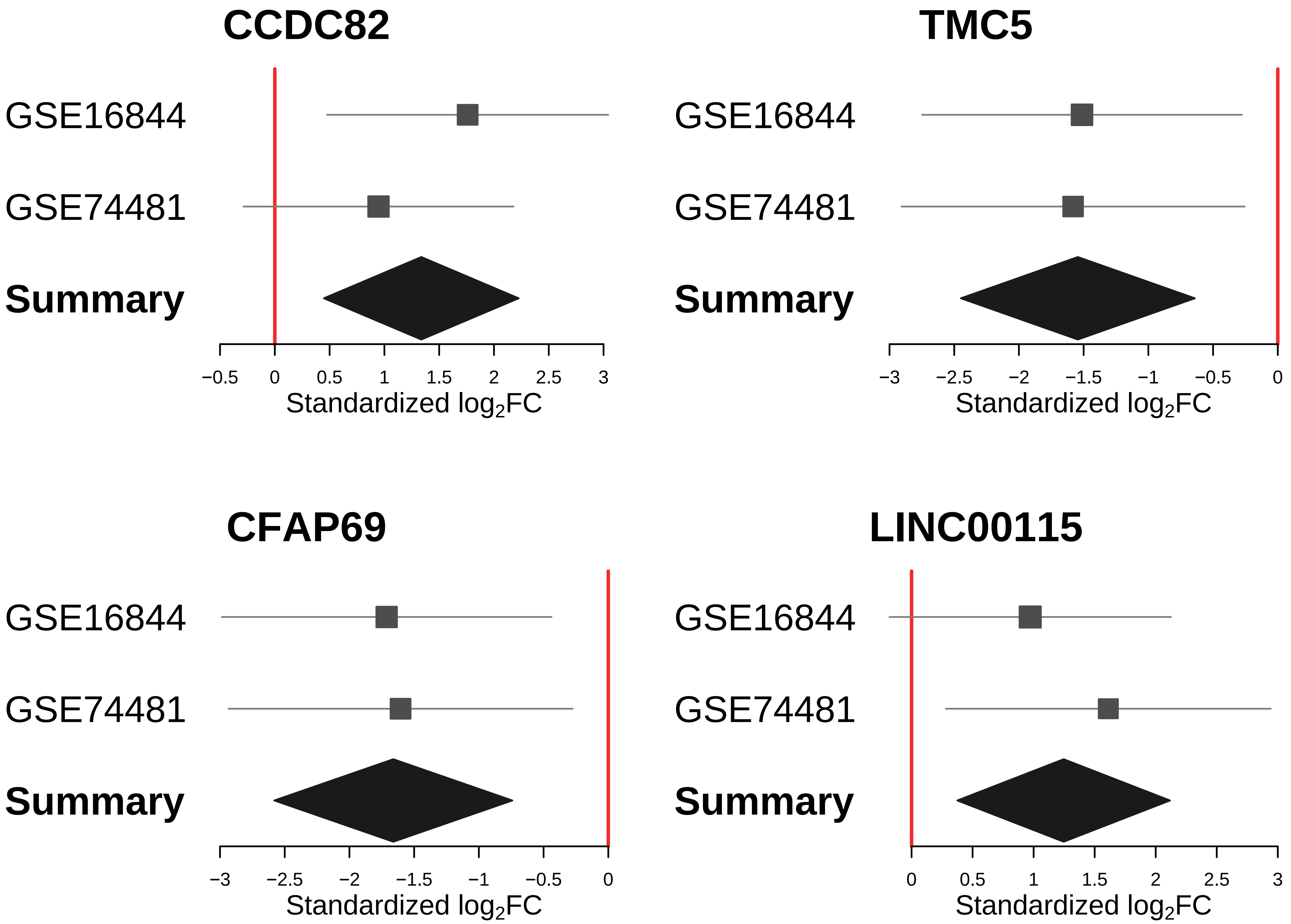
CCDC82
GSE16844
GSE74481
Summary
−0.5 0 0.5 1 1.5 2 2.5 3
Standardized log2FC
TMC5
GSE16844
GSE74481
Summary
−3 −2.5 −2 −1.5 −1 −0.5 0
Standardized log2FC
CFAP69
GSE16844
GSE74481
Summary
−3 −2.5 −2 −1.5 −1 −0.5 0
Standardized log2FC
LINC00115
GSE16844
GSE74481
Summary
0 0.5 1 1.5 2 2.5 3
Standardized log2FC

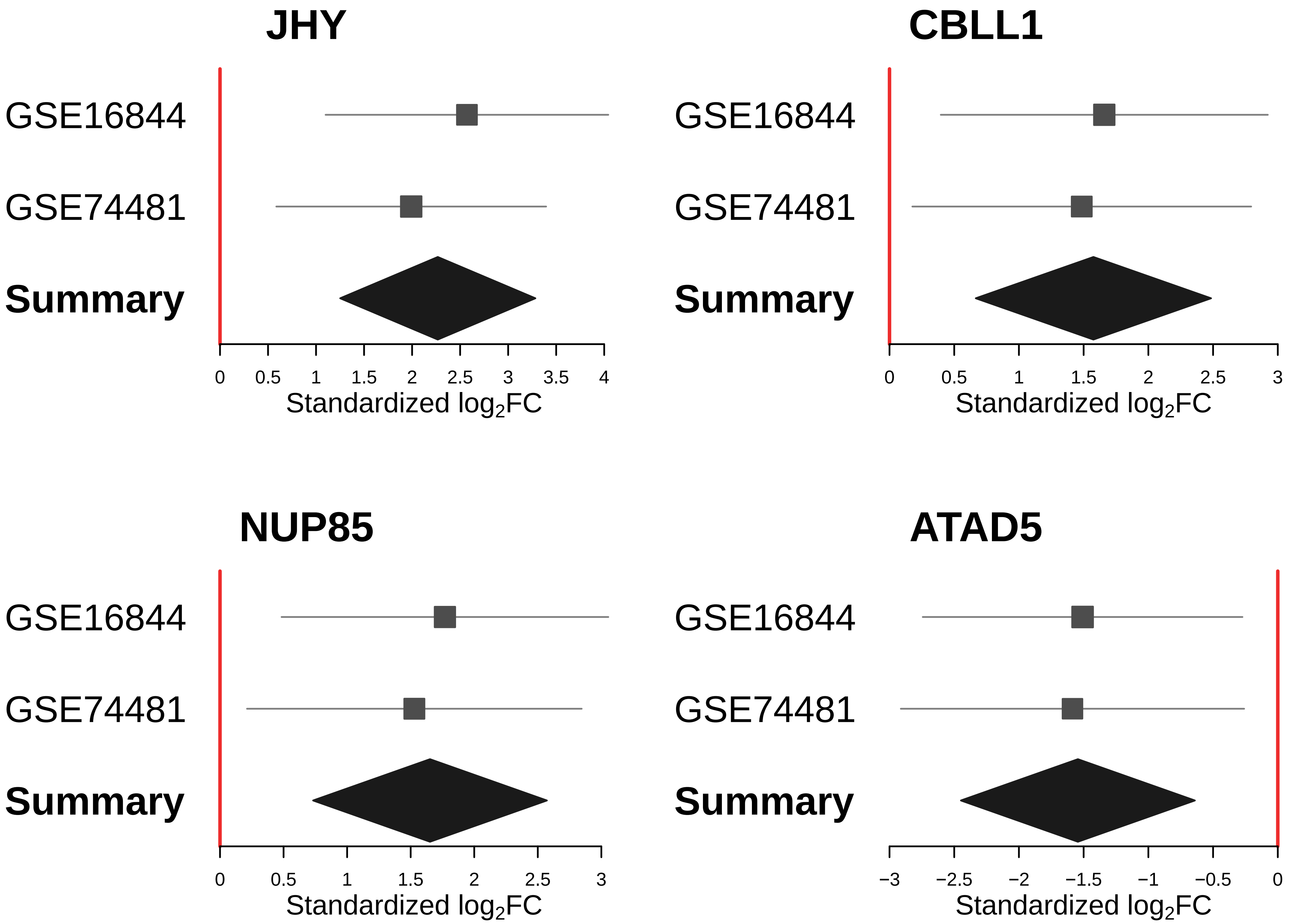
JHY
GSE16844
GSE74481
Summary
0
0.5
1
1.5
2
2.5
3
3.5
4
Standardized log2FC
CBLL1
GSE16844
GSE74481
Summary
0
0.5
1
1.5
2
2.5
3
Standardized log2FC
NUP85
GSE16844
GSE74481
Summary
0
0.5
1
1.5
2
2.5
3
Standardized log2FC
ATAD5
GSE16844
GSE74481
Summary
−3
−2.5
−2
−1.5
−1
−0.5
0
Standardized log2FC

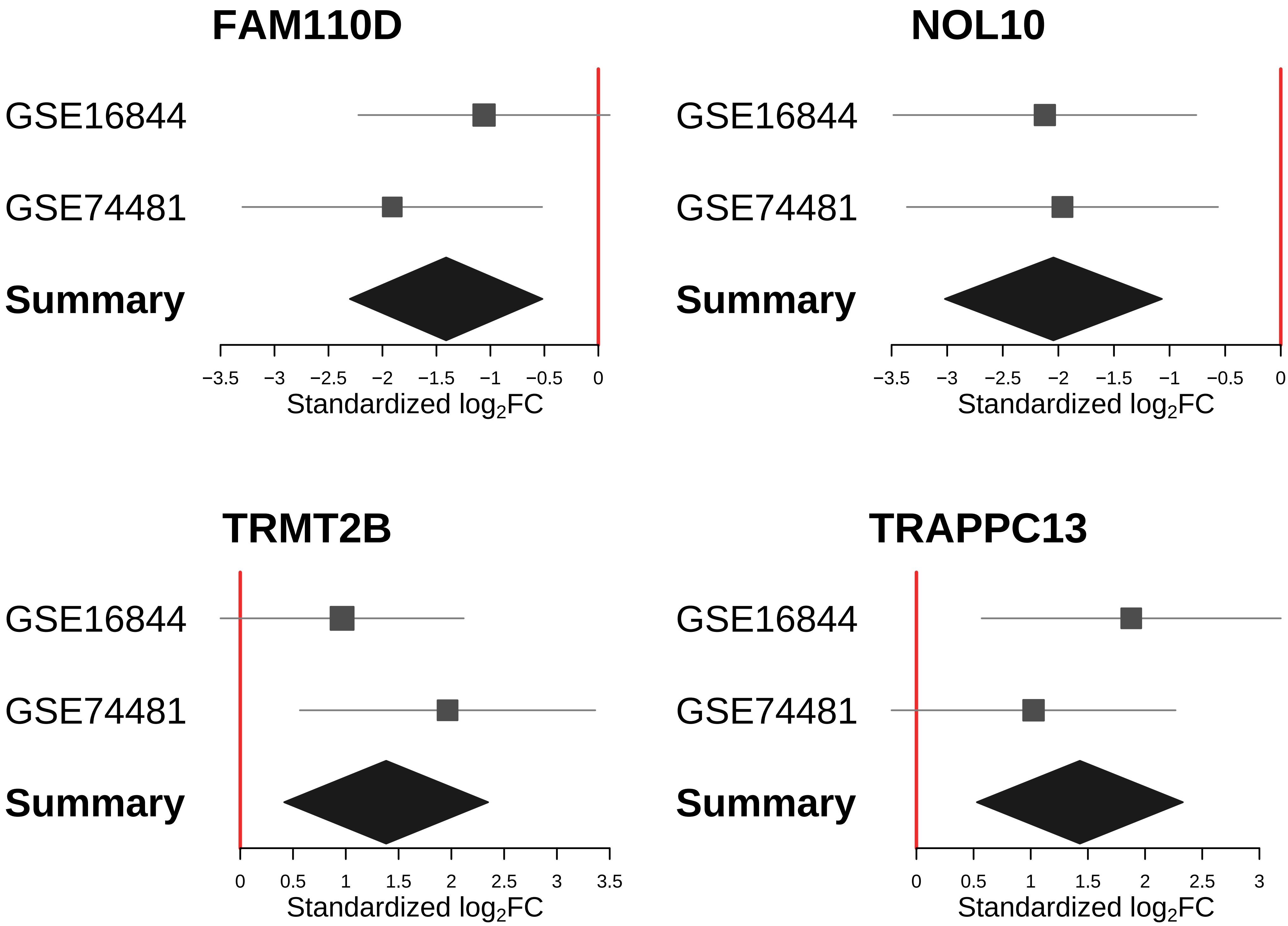

FAM110D
GSE16844
GSE74481
Summary
−3.5
−3
−2.5
−2
−1.5
−1
−0.5
0
Standardized log2FC
NOL10
GSE16844
GSE74481
Summary
−3.5
−3
−2.5
−2
−1.5
−1
−0.5
0
Standardized log2FC
TRMT2B
GSE16844
GSE74481
Summary
0
0.5
1
1.5
2
2.5
3
3.5
Standardized log2FC
TRAPPC13
GSE16844
GSE74481
Summary
0
0.5
1
1.5
2
2.5
3
Standardized log2FC

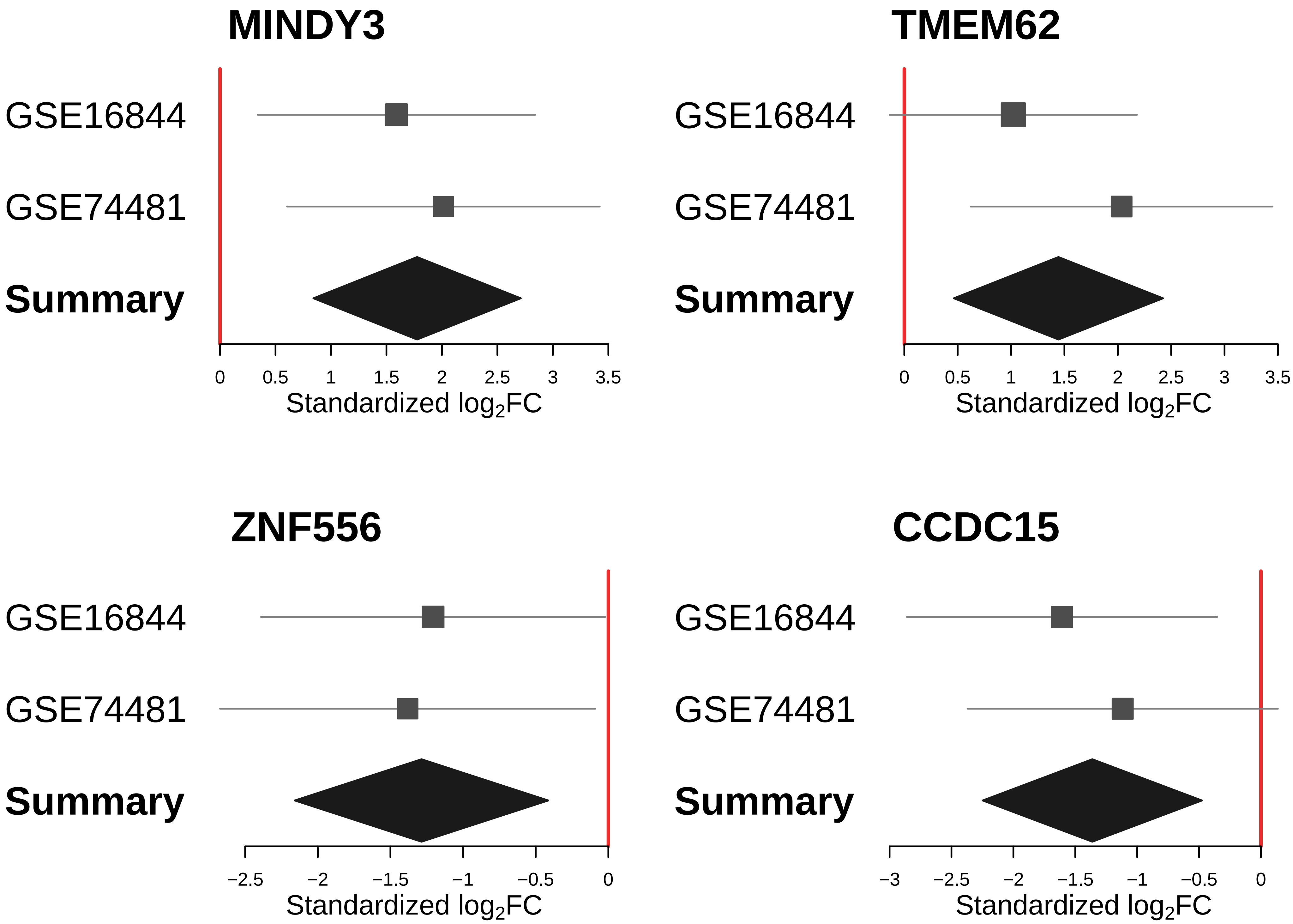
MINDY3
GSE16844
GSE74481
Summary
0 0.5 1 1.5 2 2.5 3 3.5
Standardized log2FC
TMEM62
GSE16844
GSE74481
Summary
0 0.5 1 1.5 2 2.5 3 3.5
Standardized log2FC
ZNF556
GSE16844
GSE74481
Summary
−2.5 −2 −1.5 −1 −0.5 0
Standardized log2FC
CCDC15
GSE16844
GSE74481
Summary
−3 −2.5 −2 −1.5 −1 −0.5 0
Standardized log2FC

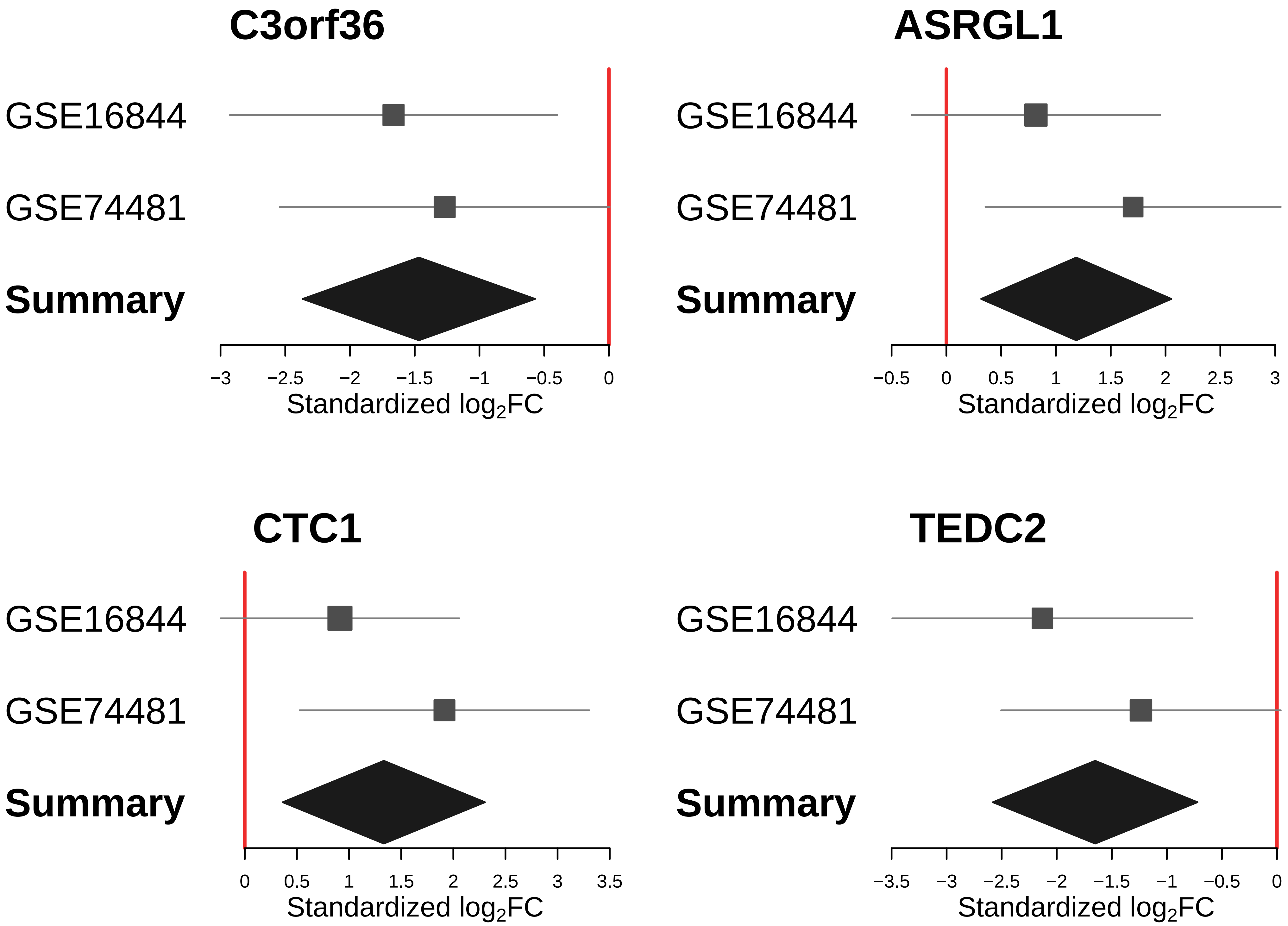
C3orf36
GSE16844
GSE74481
Summary
−3
−2.5
−2
−1.5
−1
−0.5
0
Standardized $log_2FC$
ASRGL1
GSE16844
GSE74481
Summary
−0.5
0
0.5
1
1.5
2
2.5
3
Standardized $log_2FC$
CTC1
GSE16844
GSE74481
Summary
0
0.5
1
1.5
2
2.5
3
3.5
Standardized $log_2FC$
TEDC2
GSE16844
GSE74481
Summary
−3.5
−3
−2.5
−2
−1.5
−1
−0.5
0
Standardized $log_2FC$

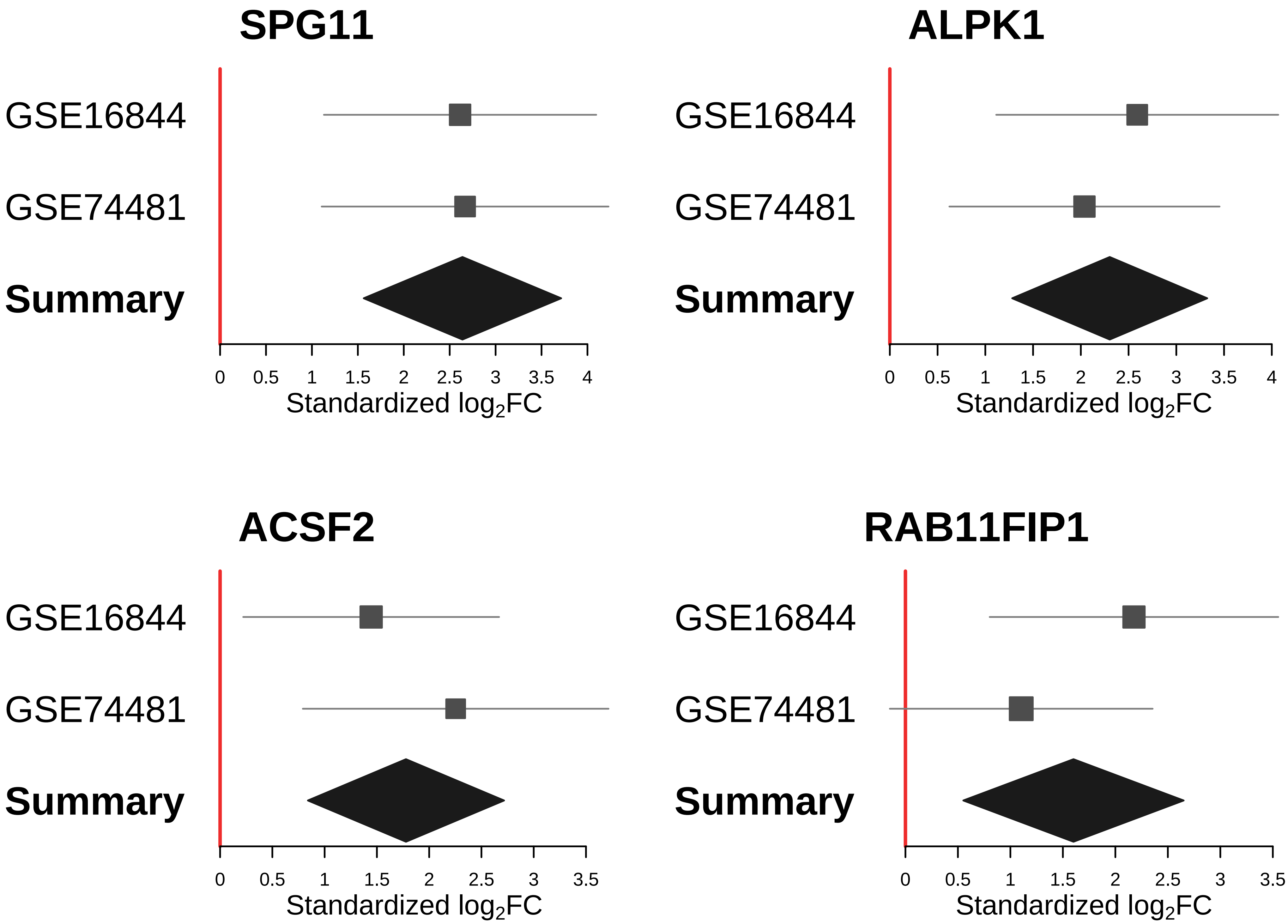
SPG11
GSE16844
GSE74481
Summary
0 0.5 1 1.5 2 2.5 3 3.5 4
Standardized log2FC
ALPK1
GSE16844
GSE74481
Summary
0 0.5 1 1.5 2 2.5 3 3.5 4
Standardized log2FC
ACSF2
GSE16844
GSE74481
Summary
0 0.5 1 1.5 2 2.5 3 3.5
Standardized log2FC
RAB11FIP1
GSE16844
GSE74481
Summary
0 0.5 1 1.5 2 2.5 3 3.5
Standardized log2FC

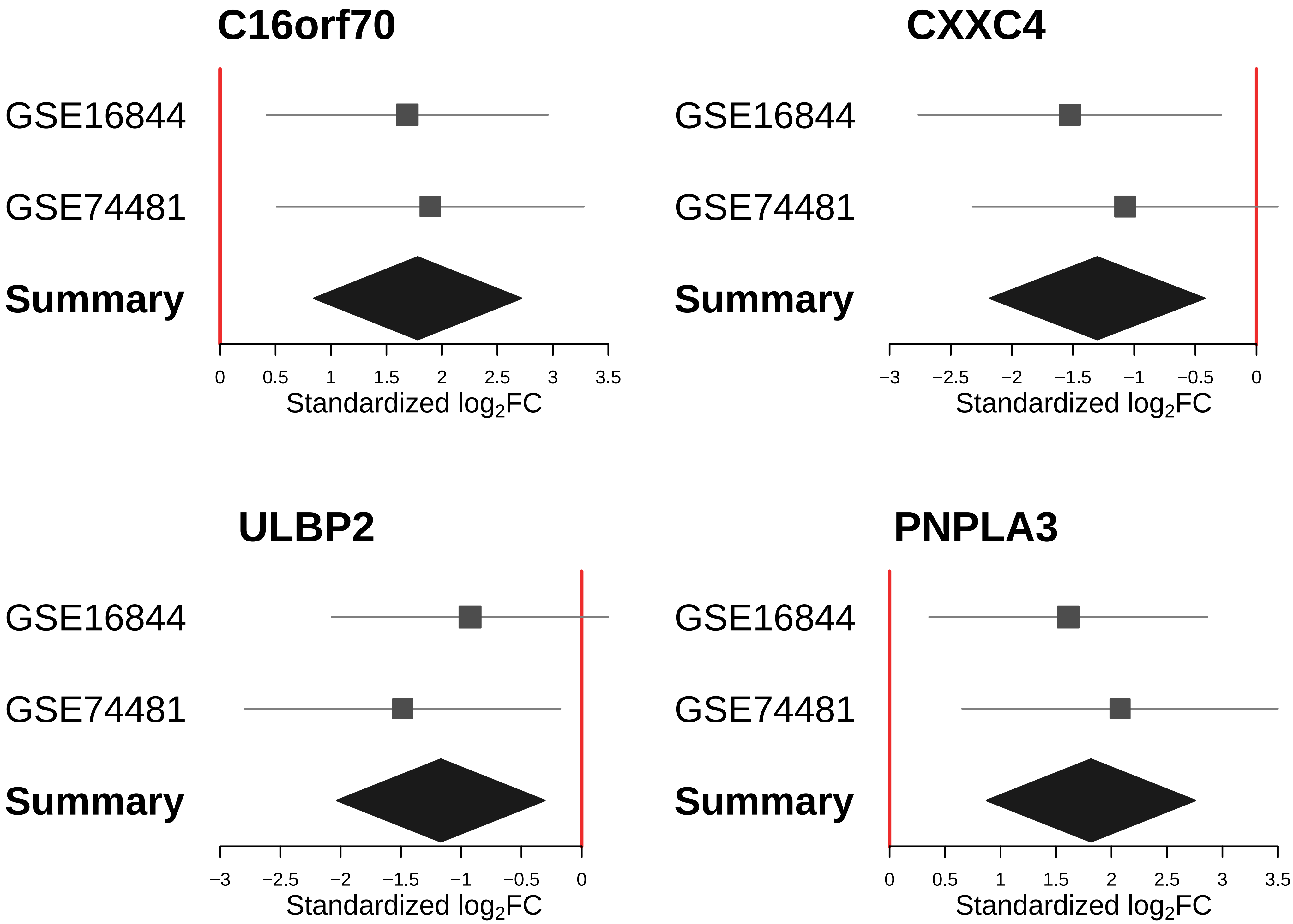
C16orf70
GSE16844
GSE74481
Summary
0 0.5 1 1.5 2 2.5 3 3.5
Standardized log2FC
CXXC4
GSE16844
GSE74481
Summary
−3 −2.5 −2 −1.5 −1 −0.5 0
Standardized log2FC
ULBP2
GSE16844
GSE74481
Summary
−3 −2.5 −2 −1.5 −1 −0.5 0
Standardized log2FC
PNPLA3
GSE16844
GSE74481
Summary
0 0.5 1 1.5 2 2.5 3 3.5
Standardized log2FC

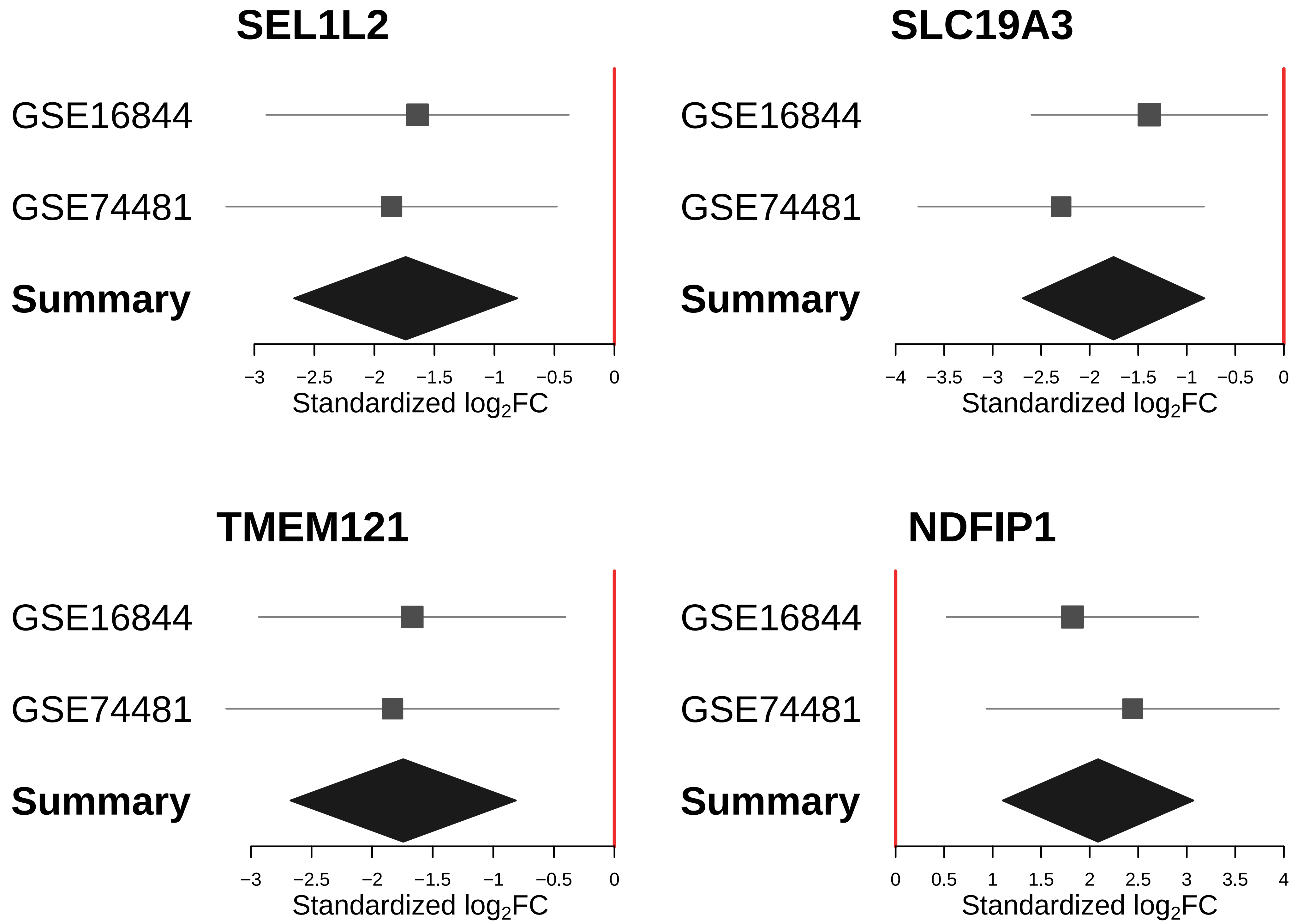
SEL1L2
GSE16844
GSE74481
Summary
−3 −2.5 −2 −1.5 −1 −0.5 0
Standardized log2FC
SLC19A3
GSE16844
GSE74481
Summary
−4 −3.5 −3 −2.5 −2 −1.5 −1 −0.5 0
Standardized log2FC
TMEM121
GSE16844
GSE74481
Summary
−3 −2.5 −2 −1.5 −1 −0.5 0
Standardized log2FC
NDFIP1
GSE16844
GSE74481
Summary
0 0.5 1 1.5 2 2.5 3 3.5 4
Standardized log2FC

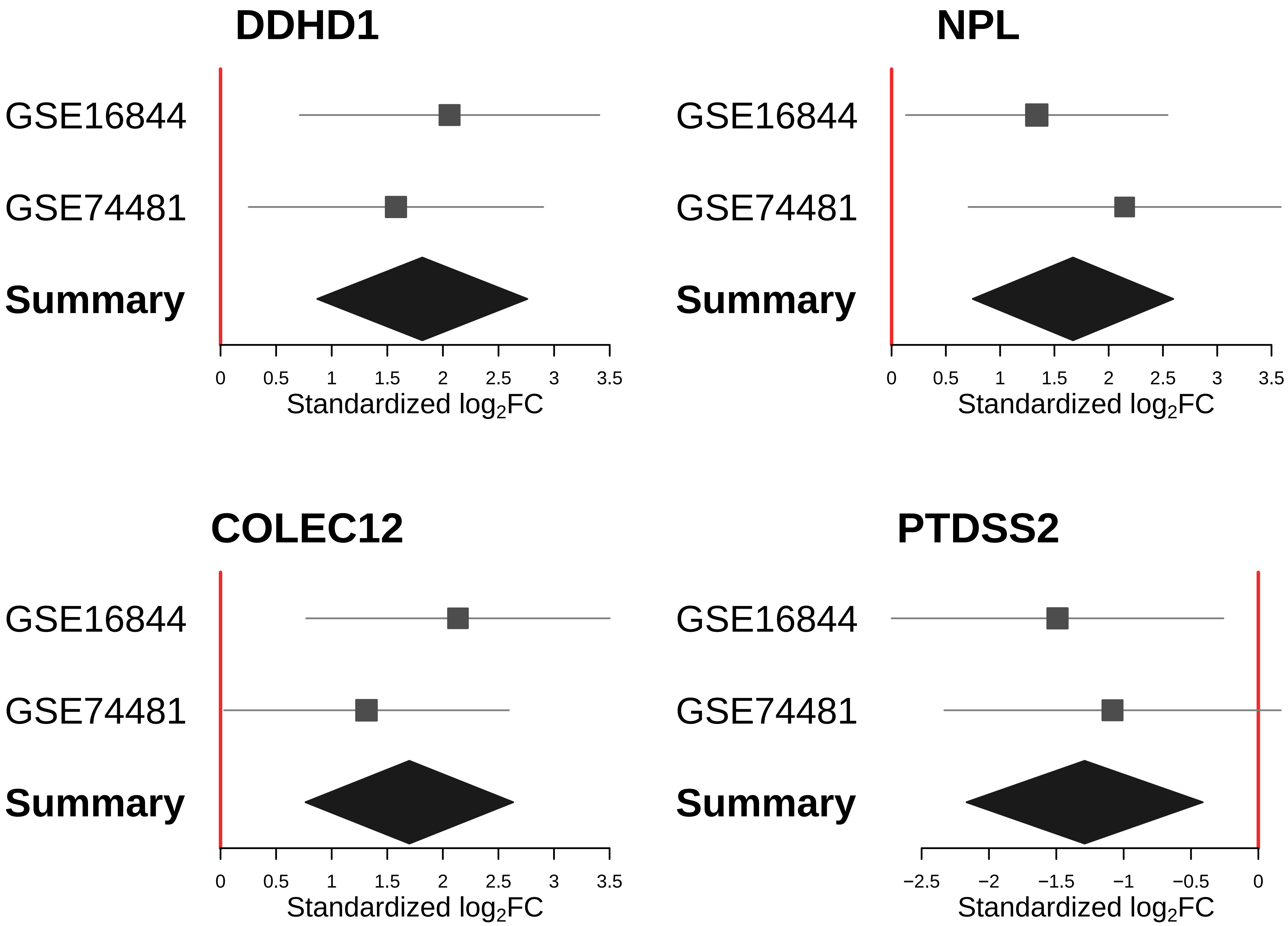

DDHD1
GSE16844
GSE74481
Summary
0
0.5
1
1.5
2
2.5
3
3.5
Standardized log2FC
NPL
GSE16844
GSE74481
Summary
0
0.5
1
1.5
2
2.5
3
3.5
Standardized log2FC
COLEC12
GSE16844
GSE74481
Summary
0
0.5
1
1.5
2
2.5
3
3.5
Standardized log2FC
PTDSS2
GSE16844
GSE74481
Summary
−2.5
−2
−1.5
−1
−0.5
0
Standardized log2FC

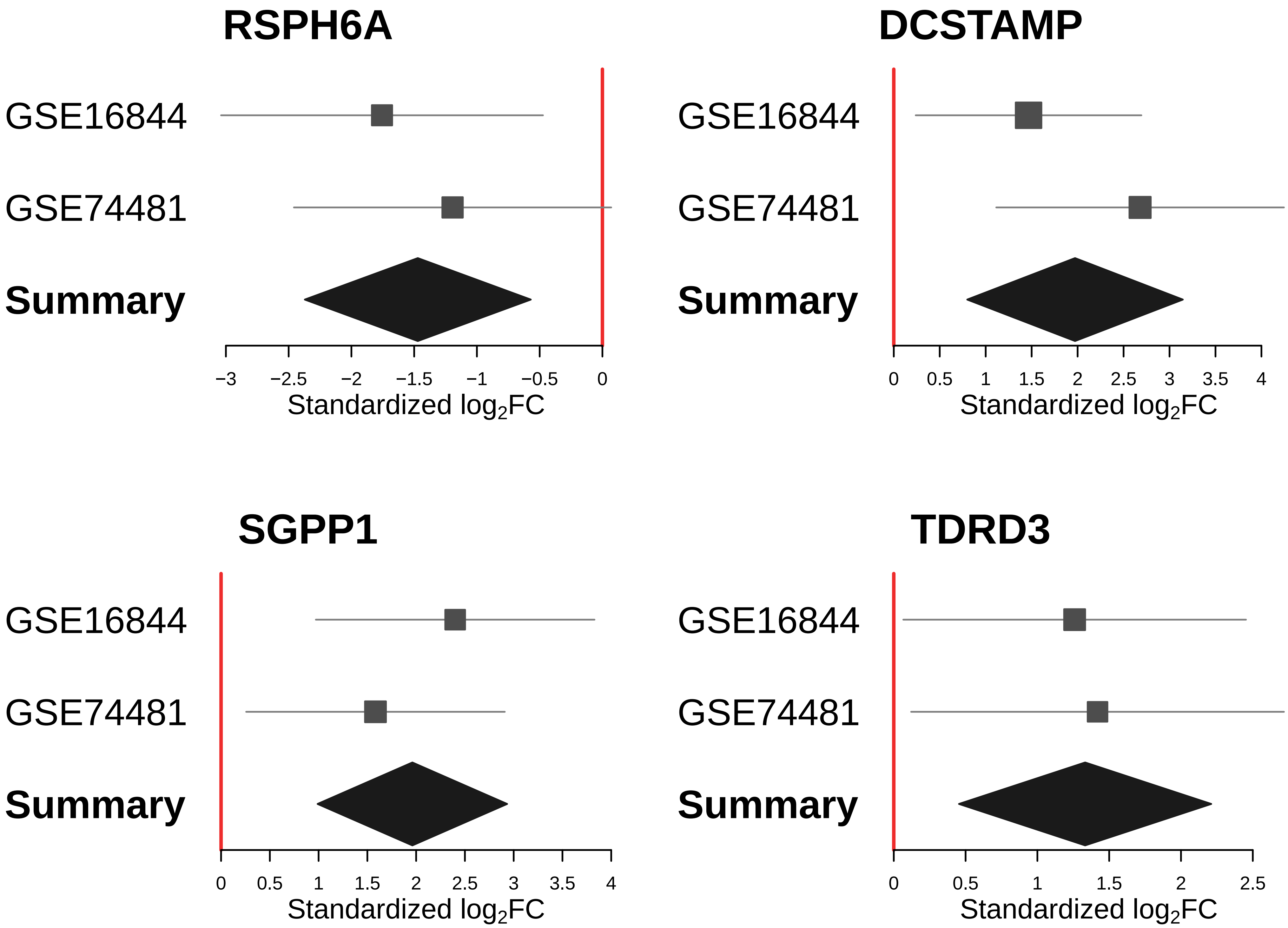
RSPH6A
GSE16844
GSE74481
Summary
−3 −2.5 −2 −1.5 −1 −0.5 0
Standardized log2FC
DCSTAMP
GSE16844
GSE74481
Summary
0 0.5 1 1.5 2 2.5 3 3.5 4
Standardized log2FC
SGPP1
GSE16844
GSE74481
Summary
0 0.5 1 1.5 2 2.5 3 3.5 4
Standardized log2FC
TDRD3
GSE16844
GSE74481
Summary
0 0.5 1 1.5 2 2.5
Standardized log2FC

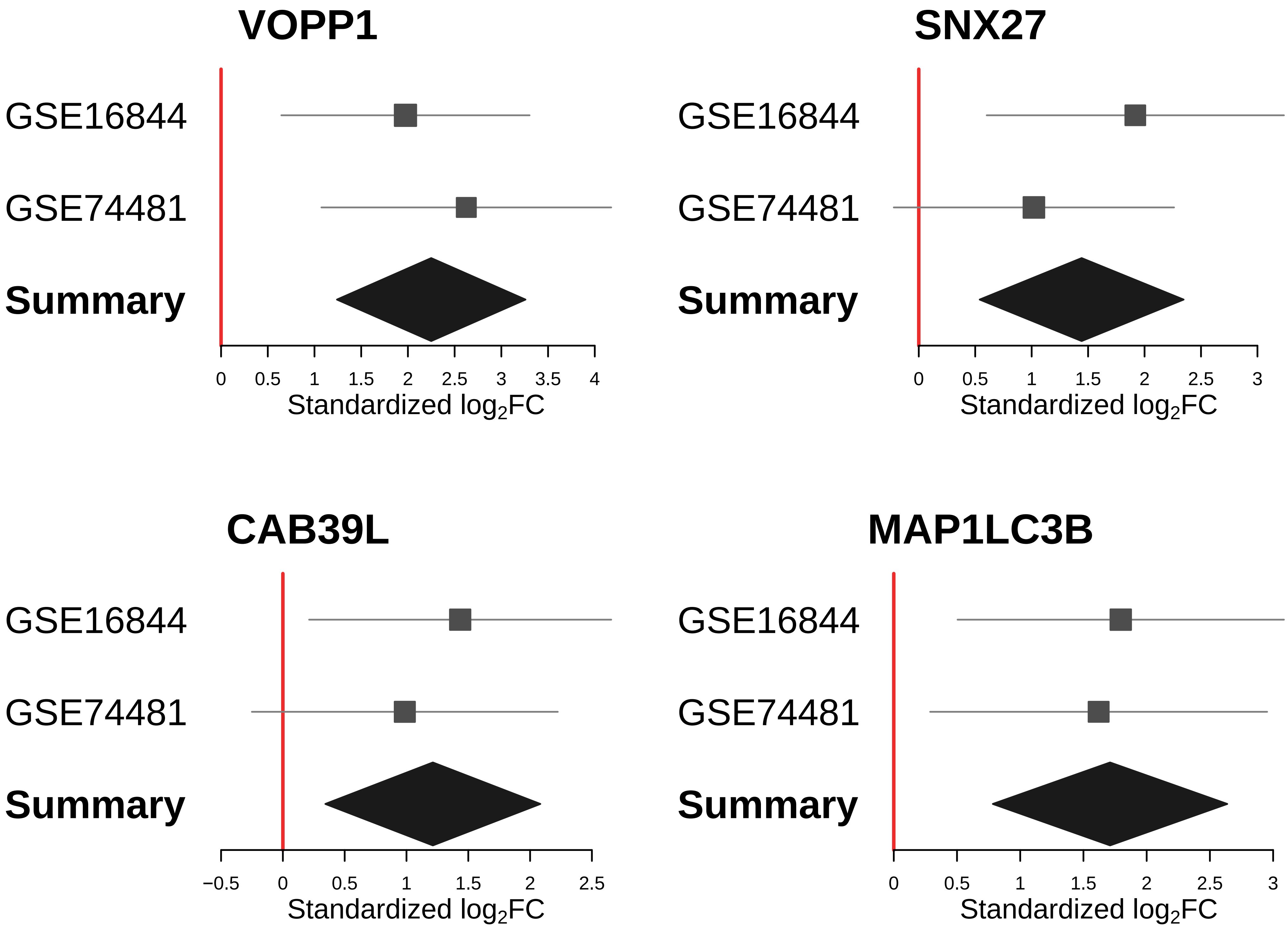
VOPP1
GSE16844
GSE74481
Summary
0 0.5 1 1.5 2 2.5 3 3.5 4
Standardized log2FC
SNX27
GSE16844
GSE74481
Summary
0 0.5 1 1.5 2 2.5 3
Standardized log2FC
CAB39L
GSE16844
GSE74481
Summary
−0.5 0 0.5 1 1.5 2 2.5
Standardized log2FC
MAP1LC3B
GSE16844
GSE74481
Summary
0 0.5 1 1.5 2 2.5 3
Standardized log2FC

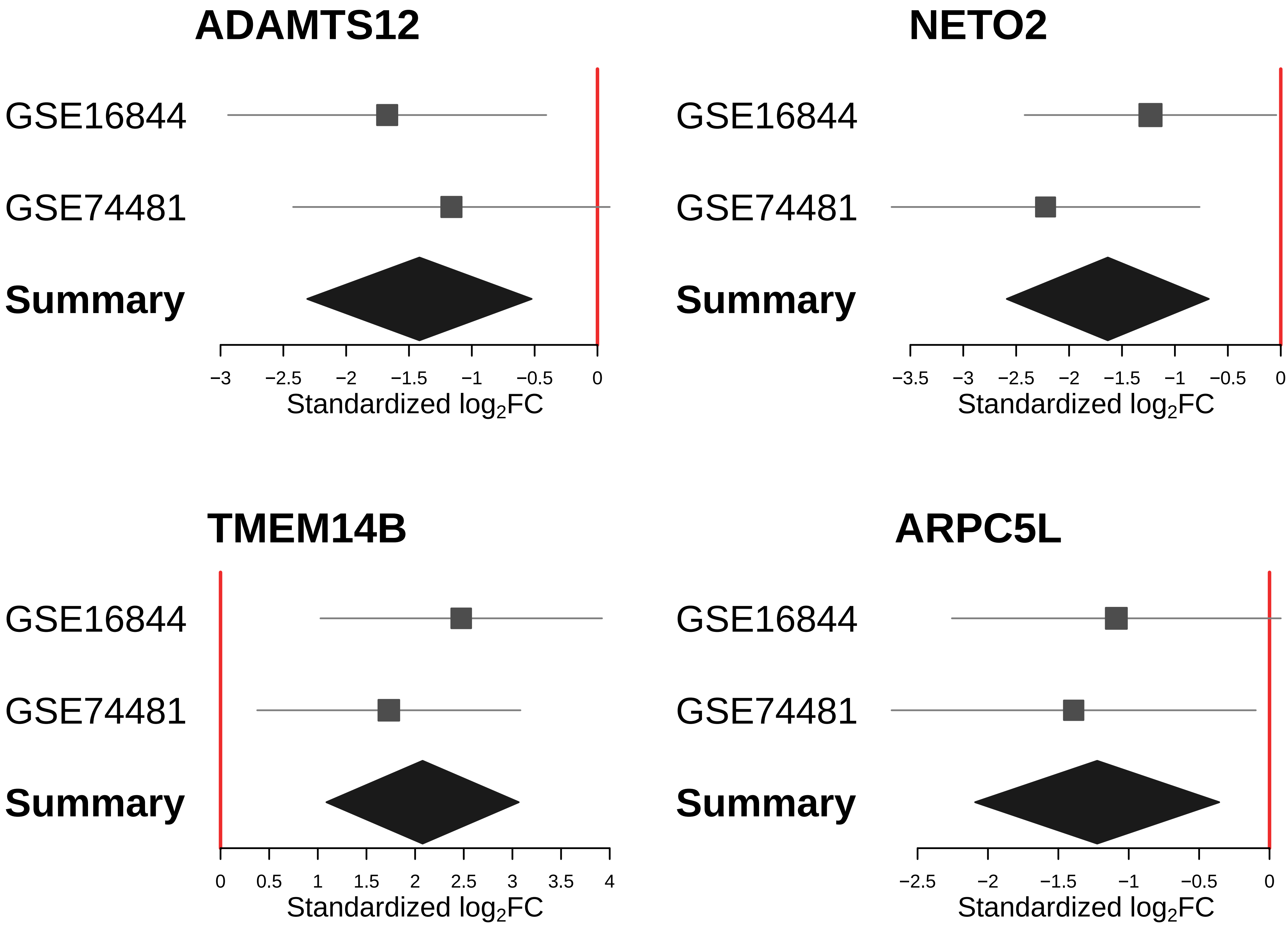
ADAMTS12
GSE16844
GSE74481
Summary
−3 −2.5 −2 −1.5 −1 −0.5 0
Standardized log2FC
NETO2
GSE16844
GSE74481
Summary
−3.5 −3 −2.5 −2 −1.5 −1 −0.5 0
Standardized log2FC
TMEM14B
GSE16844
GSE74481
Summary
0 0.5 1 1.5 2 2.5 3 3.5 4
Standardized log2FC
ARPC5L
GSE16844
GSE74481
Summary
−2.5 −2 −1.5 −1 −0.5 0
Standardized log2FC

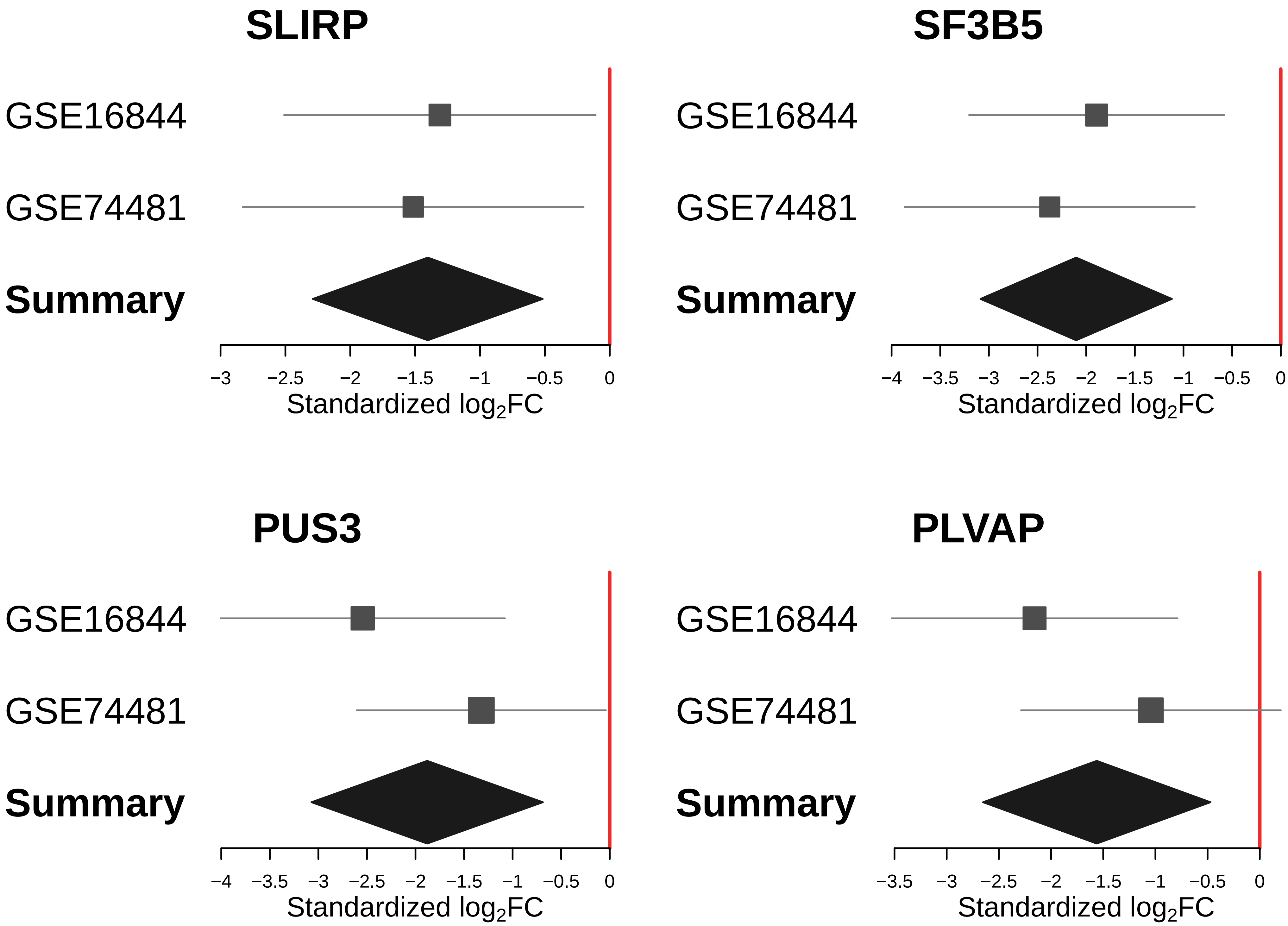

SLIRP
GSE16844
GSE74481
Summary
−3
−2.5
−2
−1.5
−1
−0.5
0
Standardized log2FC
SF3B5
GSE16844
GSE74481
Summary
−4
−3.5
−3
−2.5
−2
−1.5
−1
−0.5
0
Standardized log2FC
PUS3
GSE16844
GSE74481
Summary
−4
−3.5
−3
−2.5
−2
−1.5
−1
−0.5
0
Standardized log2FC
PLVAP
GSE16844
GSE74481
Summary
−3.5
−3
−2.5
−2
−1.5
−1
−0.5
0
Standardized log2FC

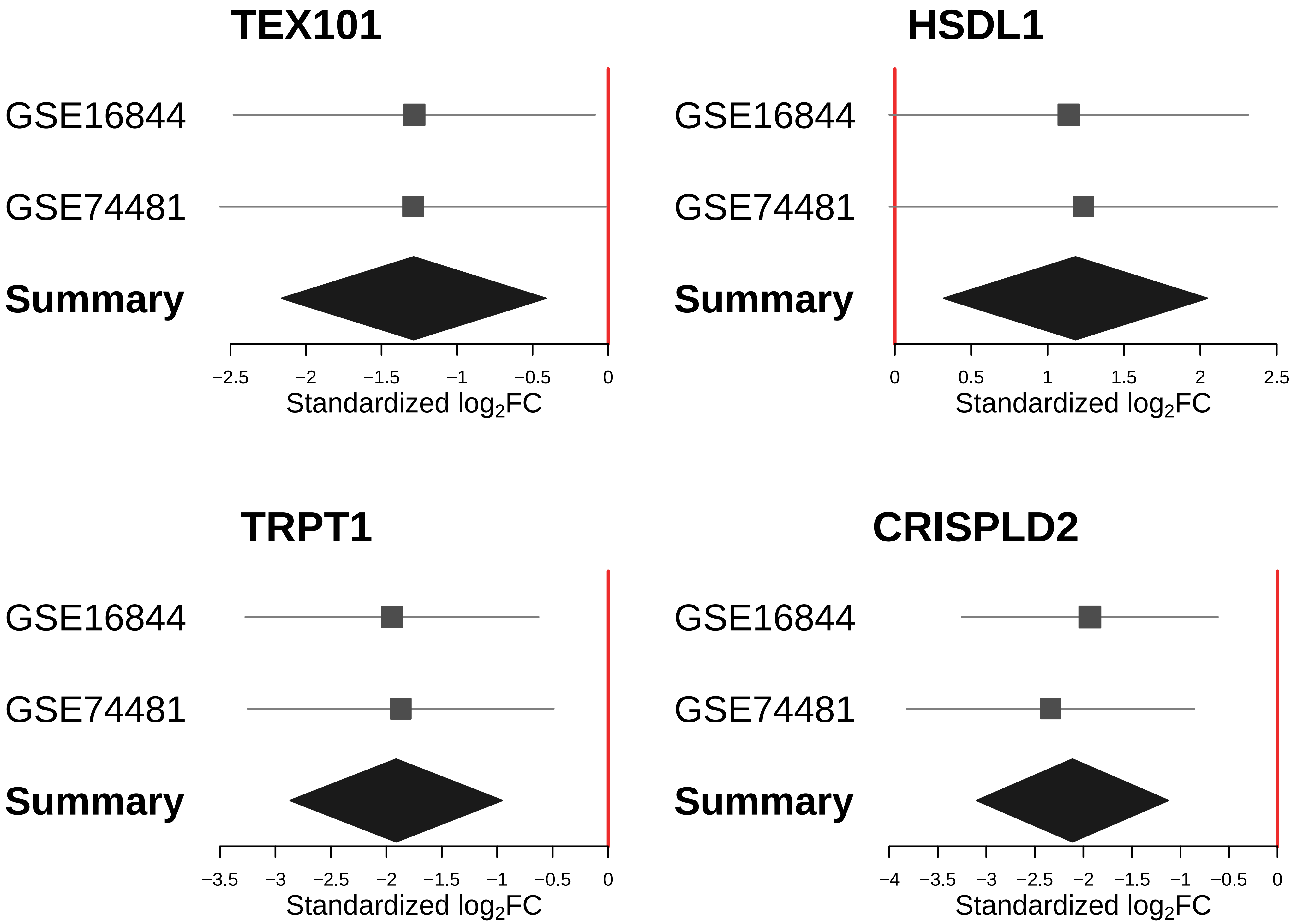
TEX101
GSE16844
GSE74481
Summary
−2.5 −2 −1.5 −1 −0.5 0
Standardized log2FC
HSDL1
GSE16844
GSE74481
Summary
0 0.5 1 1.5 2 2.5
Standardized log2FC
TRPT1
GSE16844
GSE74481
Summary
−3.5 −3 −2.5 −2 −1.5 −1 −0.5 0
Standardized log2FC
CRISPLD2
GSE16844
GSE74481
Summary
−4 −3.5 −3 −2.5 −2 −1.5 −1 −0.5 0
Standardized log2FC

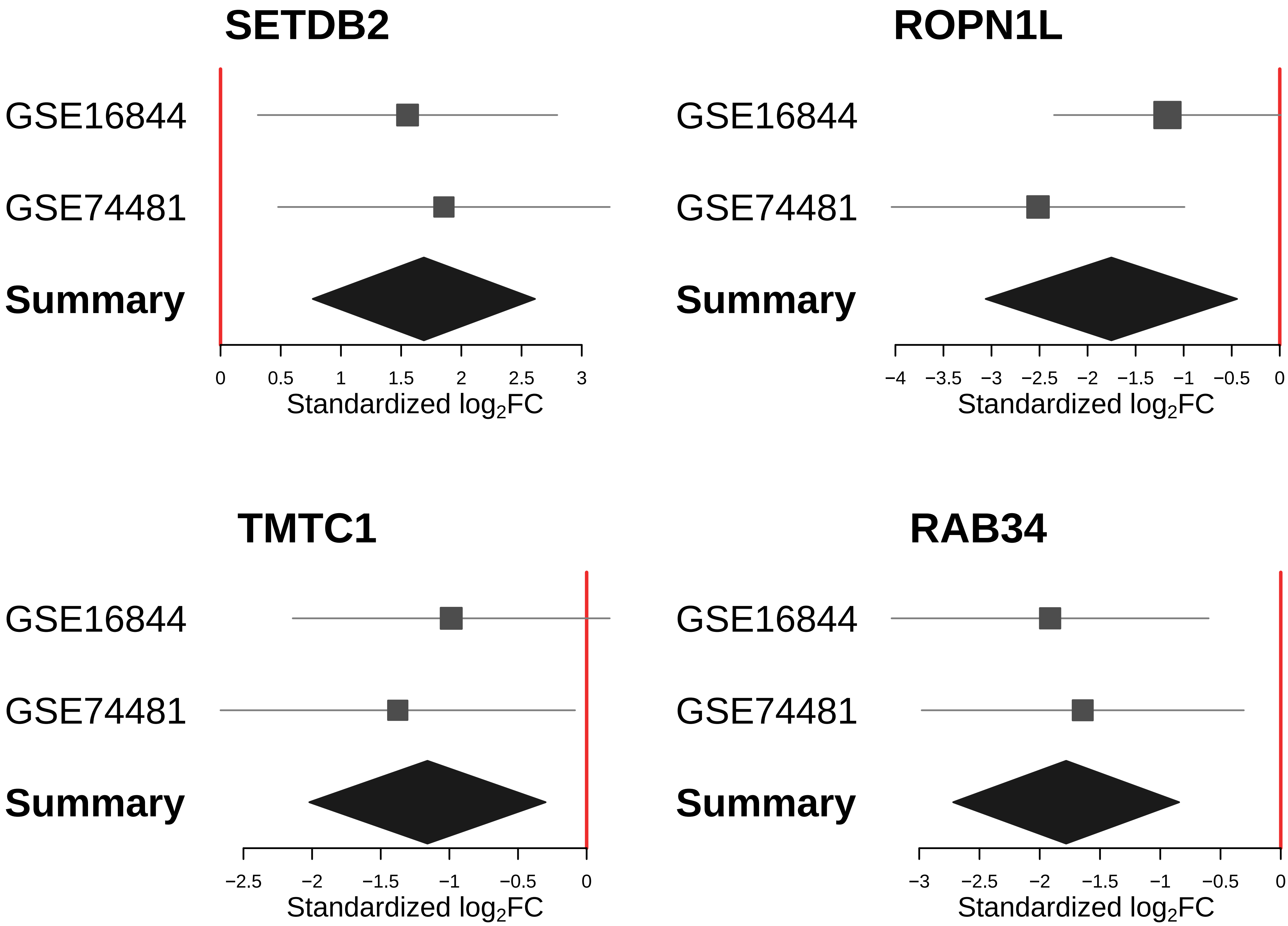
SETDB2
GSE16844
GSE74481
Summary
0
0.5
1
1.5
2
2.5
3
Standardized log2FC
ROPN1L
GSE16844
GSE74481
Summary
−4
−3.5
−3
−2.5
−2
−1.5
−1
−0.5
0
Standardized log2FC
TMTC1
GSE16844
GSE74481
Summary
−2.5
−2
−1.5
−1
−0.5
0
Standardized log2FC
RAB34
GSE16844
GSE74481
Summary
−3
−2.5
−2
−1.5
−1
−0.5
0
Standardized log2FC

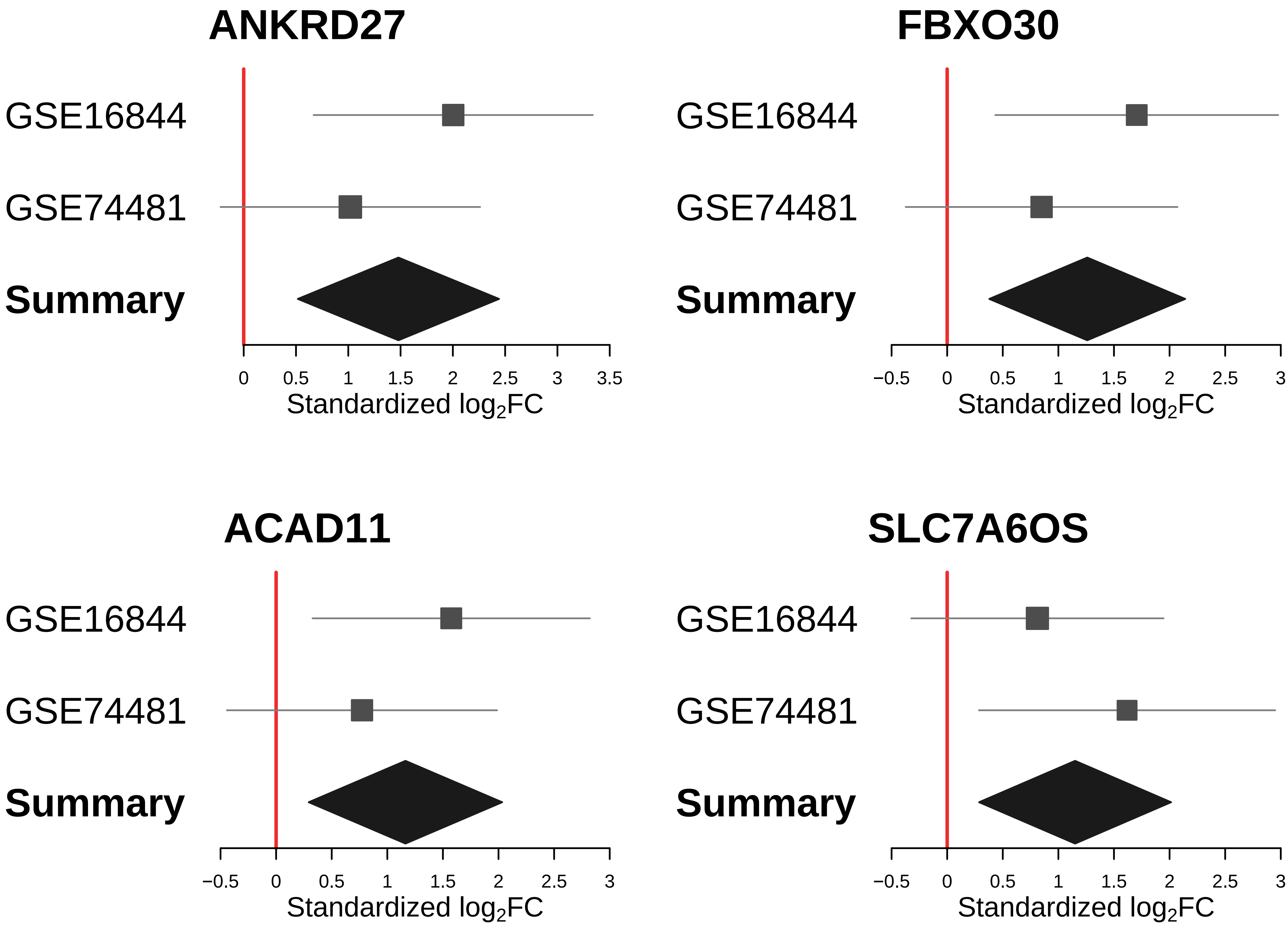
ANKRD27
GSE16844
GSE74481
Summary
0 0.5 1 1.5 2 2.5 3 3.5
Standardized log2FC
FBXO30
GSE16844
GSE74481
Summary
−0.5 0 0.5 1 1.5 2 2.5 3
Standardized log2FC
ACAD11
GSE16844
GSE74481
Summary
−0.5 0 0.5 1 1.5 2 2.5 3
Standardized log2FC
SLC7A6OS
GSE16844
GSE74481
Summary
−0.5 0 0.5 1 1.5 2 2.5 3
Standardized log2FC

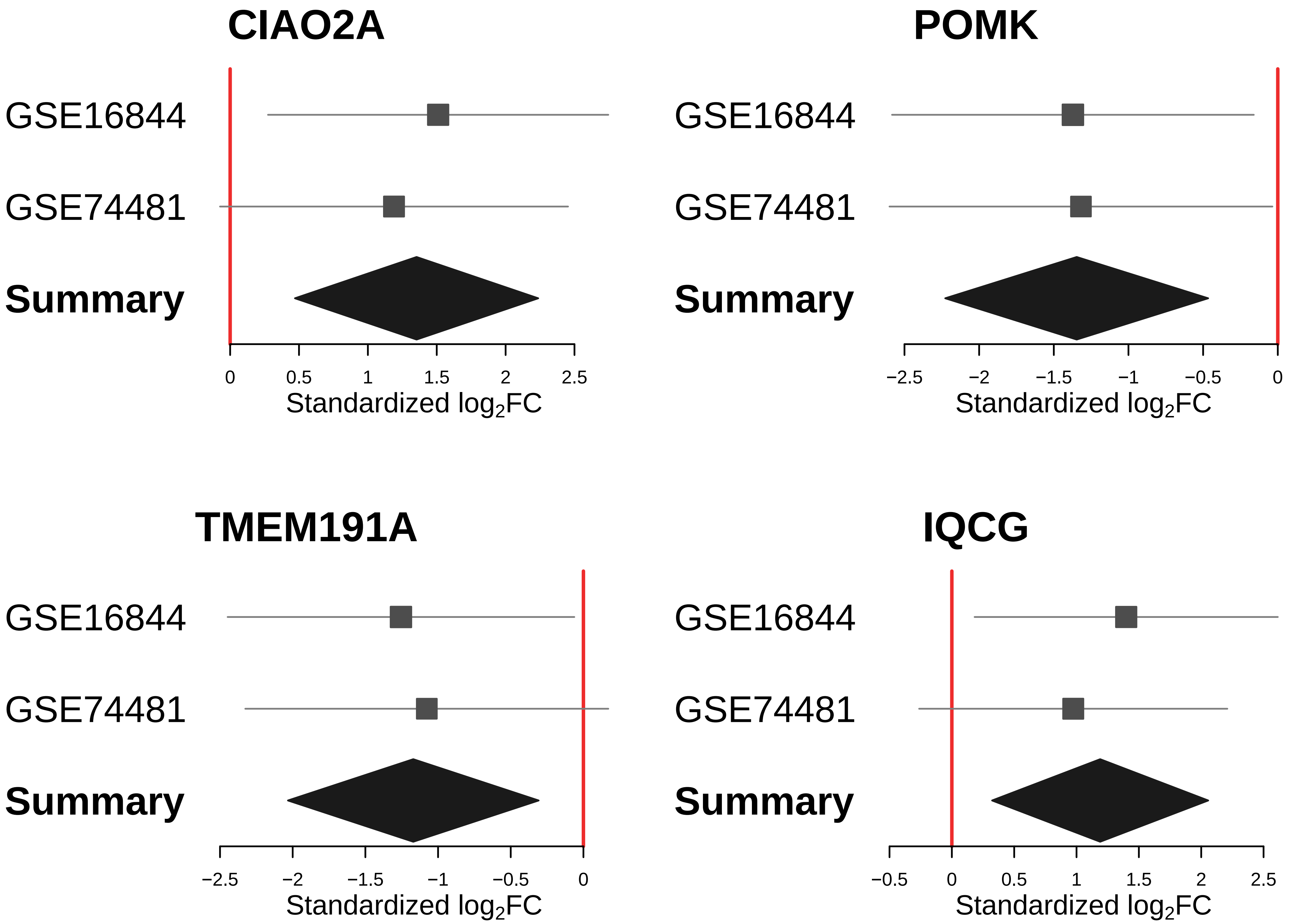

CIAO2A
GSE16844
GSE74481
Summary
0
0.5
1
1.5
2
2.5
Standardized log2FC
POMK
GSE16844
GSE74481
Summary
−2.5
−2
−1.5
−1
−0.5
0
Standardized log2FC
TMEM191A
GSE16844
GSE74481
Summary
−2.5
−2
−1.5
−1
−0.5
0
Standardized log2FC
IQCG
GSE16844
GSE74481
Summary
−0.5
0
0.5
1
1.5
2
2.5
Standardized log2FC

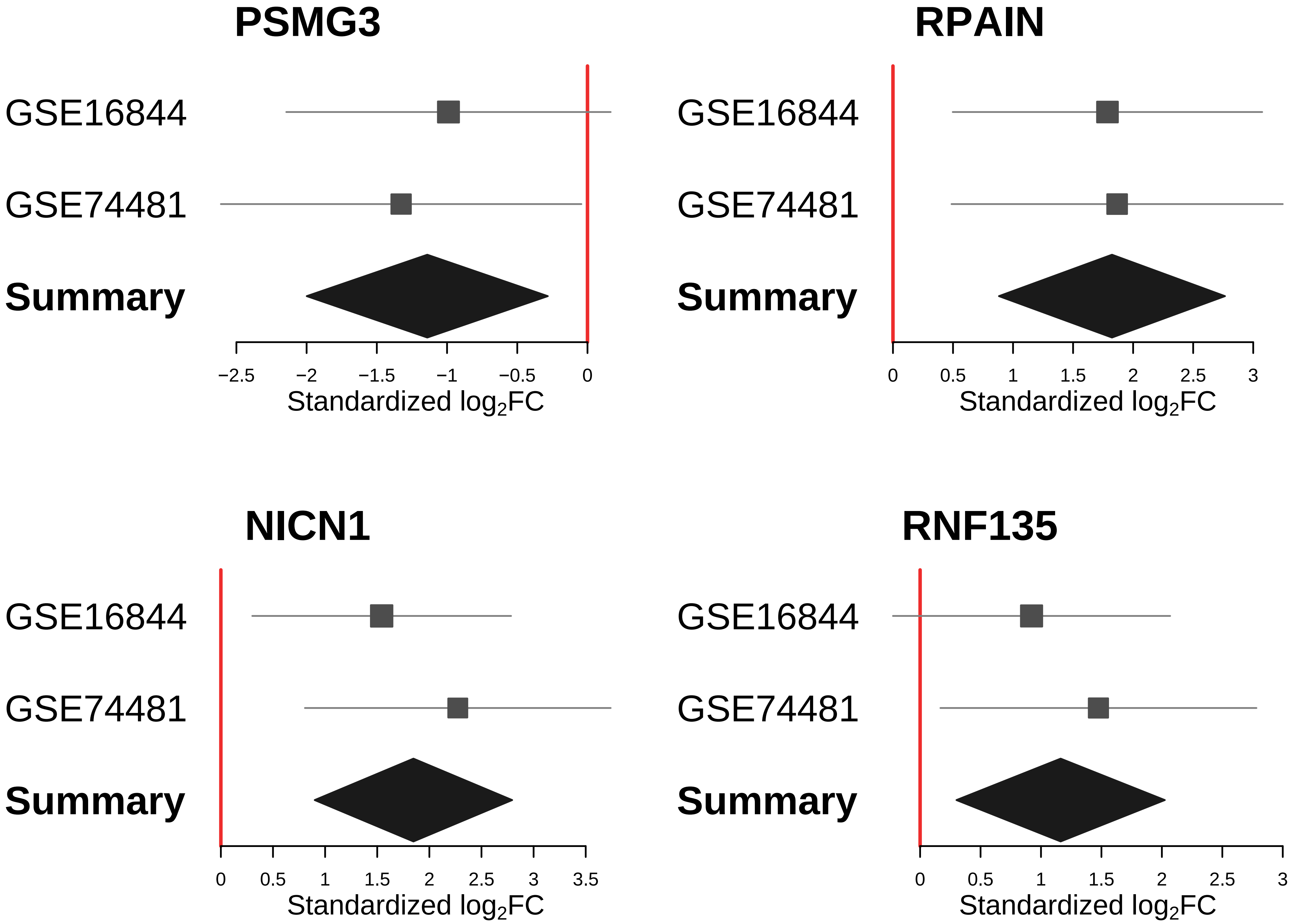
PSMG3
GSE16844
GSE74481
Summary
−2.5 −2 −1.5 −1 −0.5 0
Standardized log2FC
RPAIN
GSE16844
GSE74481
Summary
0 0.5 1 1.5 2 2.5 3
Standardized log2FC
NICN1
GSE16844
GSE74481
Summary
0 0.5 1 1.5 2 2.5 3 3.5
Standardized log2FC
RNF135
GSE16844
GSE74481
Summary
0 0.5 1 1.5 2 2.5 3
Standardized log2FC

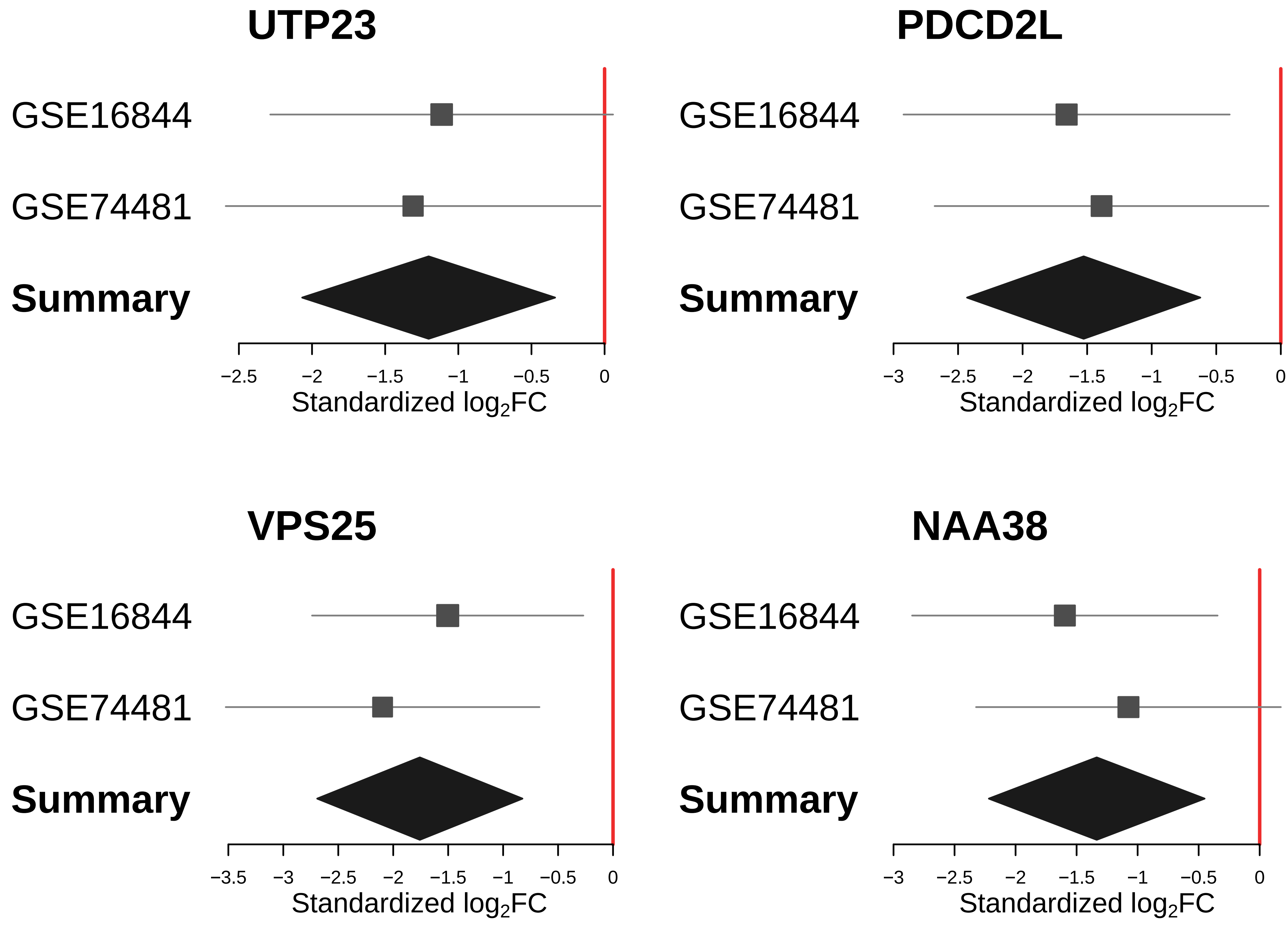
UTP23
GSE16844
GSE74481
Summary
−2.5 −2 −1.5 −1 −0.5 0
Standardized log2FC
PDCD2L
GSE16844
GSE74481
Summary
−3 −2.5 −2 −1.5 −1 −0.5 0
Standardized log2FC
VPS25
GSE16844
GSE74481
Summary
−3.5 −3 −2.5 −2 −1.5 −1 −0.5 0
Standardized log2FC
NAA38
GSE16844
GSE74481
Summary
−3 −2.5 −2 −1.5 −1 −0.5 0
Standardized log2FC

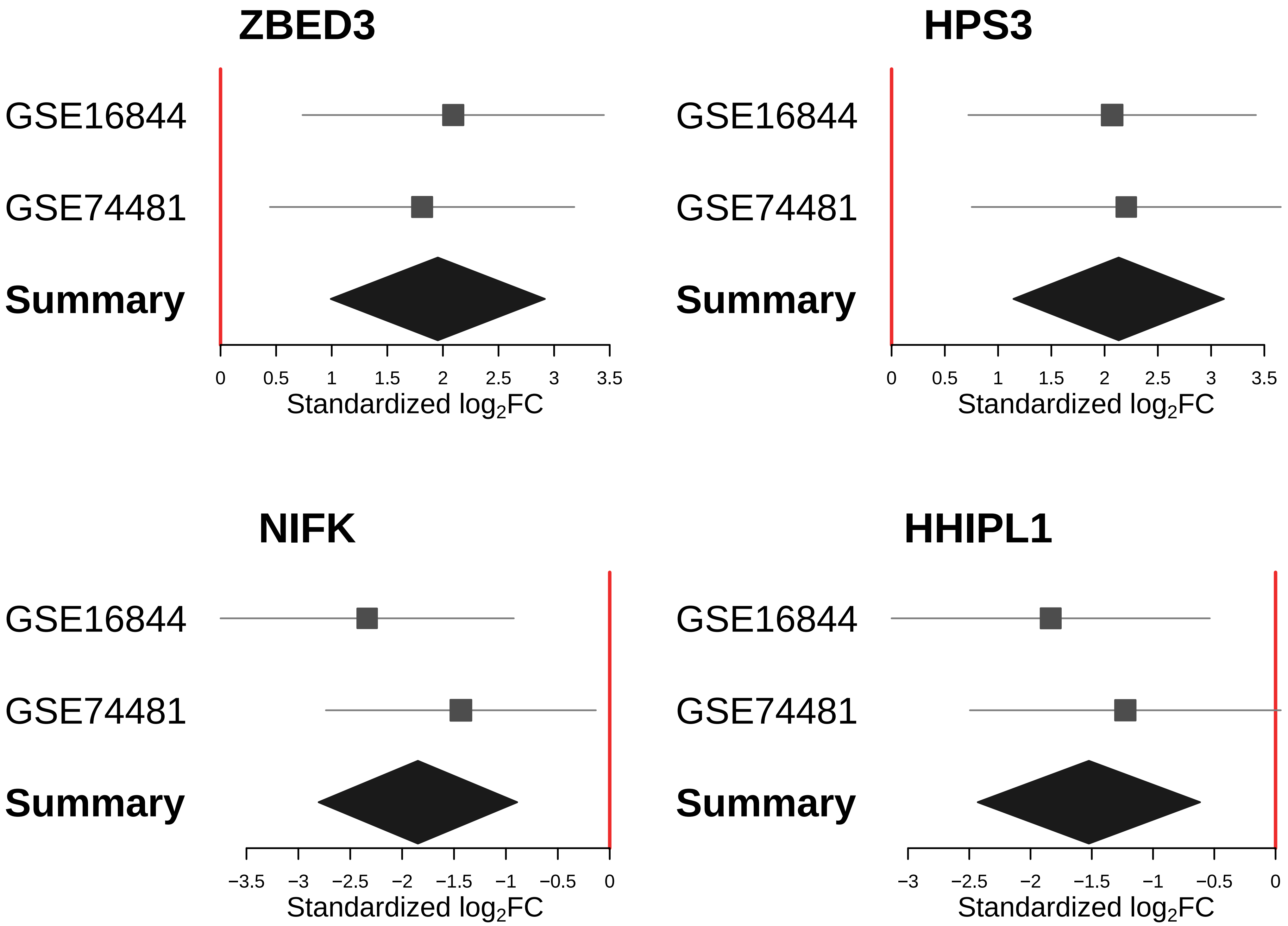
ZBED3
GSE16844
GSE74481
Summary
0
0.5
1
1.5
2
2.5
3
3.5
Standardized log2FC
HPS3
GSE16844
GSE74481
Summary
0
0.5
1
1.5
2
2.5
3
3.5
Standardized log2FC
NIFK
GSE16844
GSE74481
Summary
−3.5
−3
−2.5
−2
−1.5
−1
−0.5
0
Standardized log2FC
HHIPL1
GSE16844
GSE74481
Summary
−3
−2.5
−2
−1.5
−1
−0.5
0
Standardized log2FC

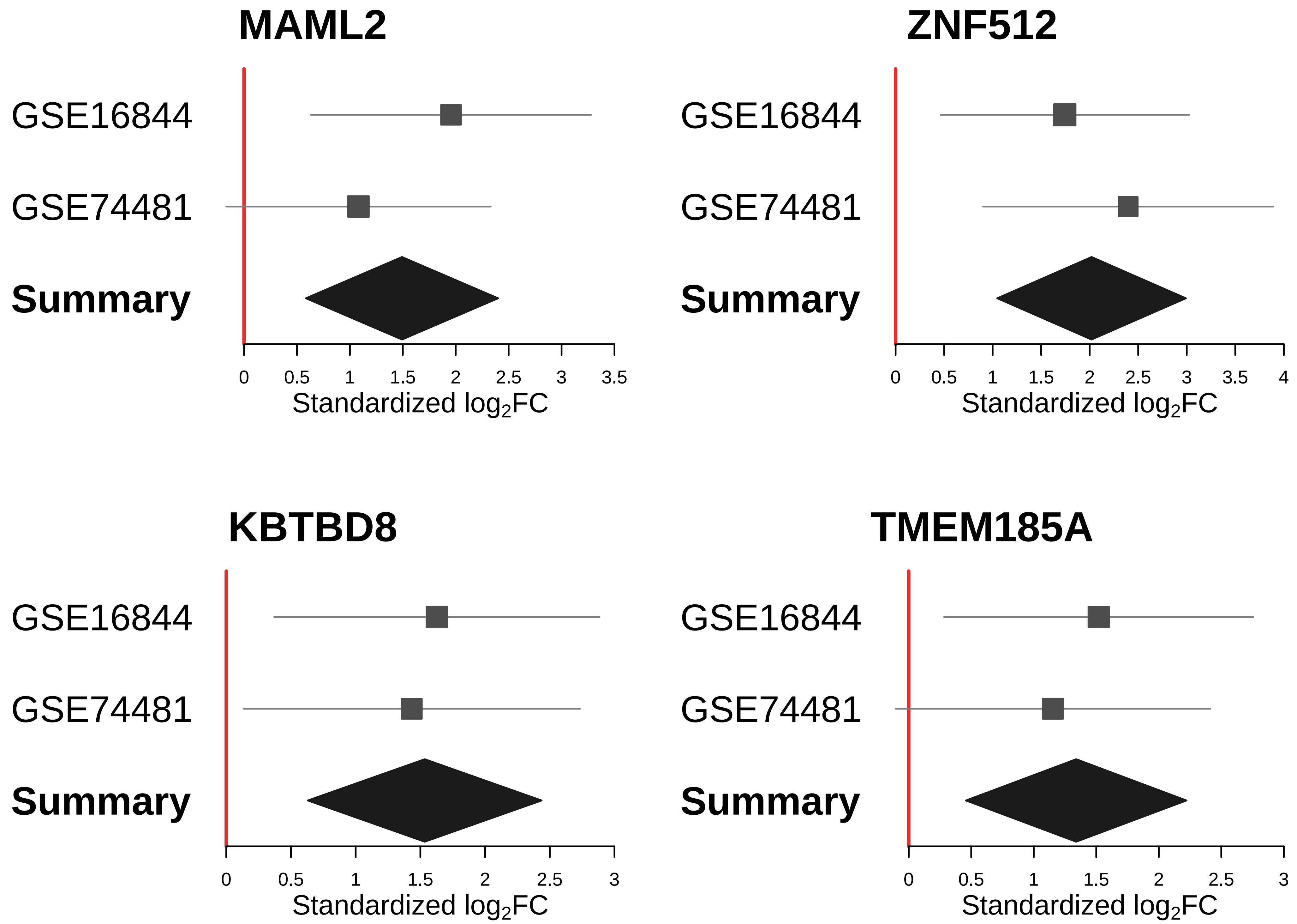
MAML2
GSE16844
GSE74481
Summary
0
0.5
1
1.5
2
2.5
3
3.5
Standardized log2FC
ZNF512
GSE16844
GSE74481
Summary
0
0.5
1
1.5
2
2.5
3
3.5
4
Standardized log2FC
KBTBD8
GSE16844
GSE74481
Summary
0
0.5
1
1.5
2
2.5
3
Standardized log2FC
TMEM185A
GSE16844
GSE74481
Summary
0
0.5
1
1.5
2
2.5
3
Standardized log2FC

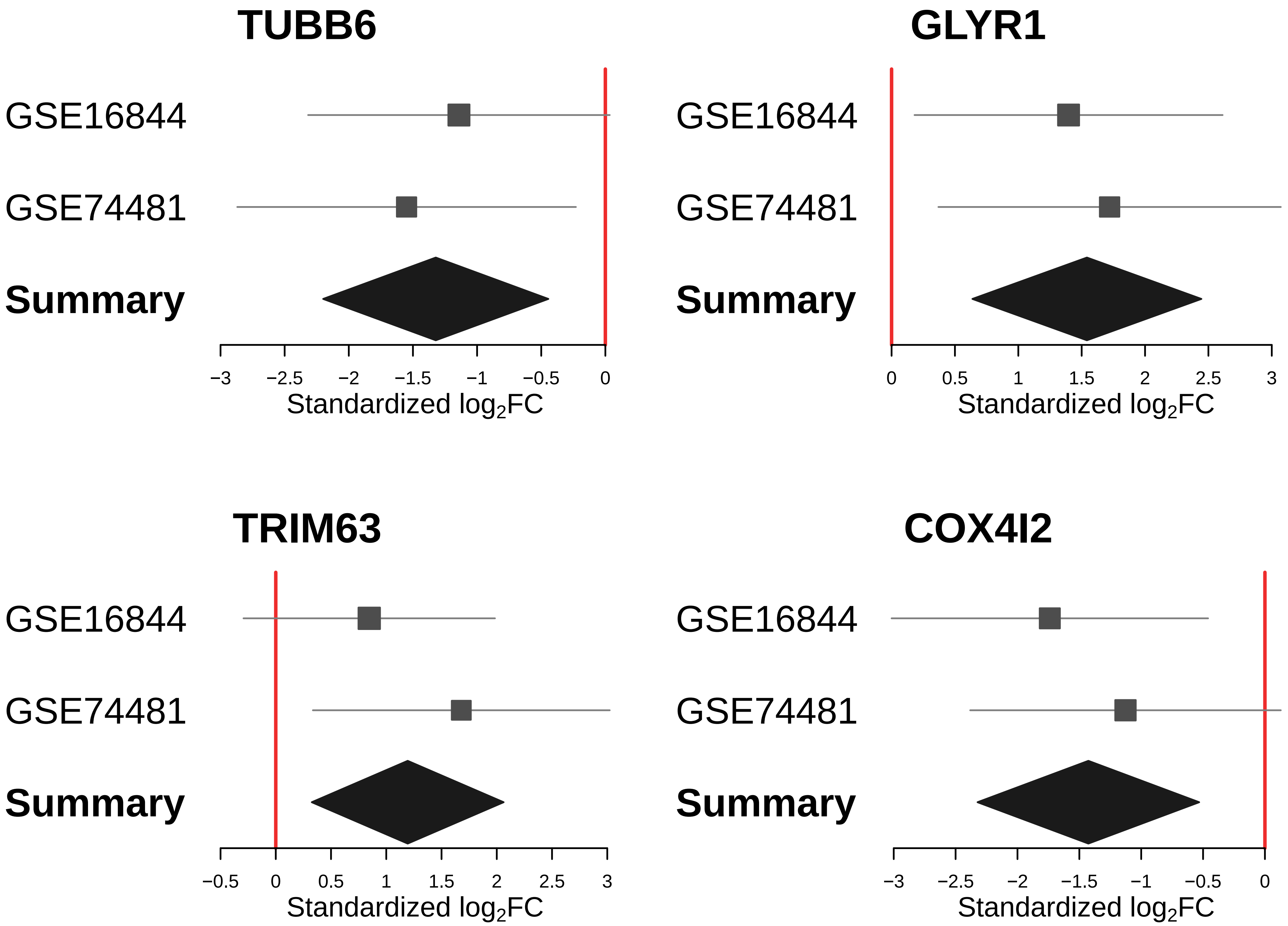
TUBB6
GSE16844
GSE74481
Summary
−3 −2.5 −2 −1.5 −1 −0.5 0
Standardized log2FC
GLYR1
GSE16844
GSE74481
Summary
0 0.5 1 1.5 2 2.5 3
Standardized log2FC
TRIM63
GSE16844
GSE74481
Summary
−0.5 0 0.5 1 1.5 2 2.5 3
Standardized log2FC
COX4I2
GSE16844
GSE74481
Summary
−3 −2.5 −2 −1.5 −1 −0.5 0
Standardized log2FC

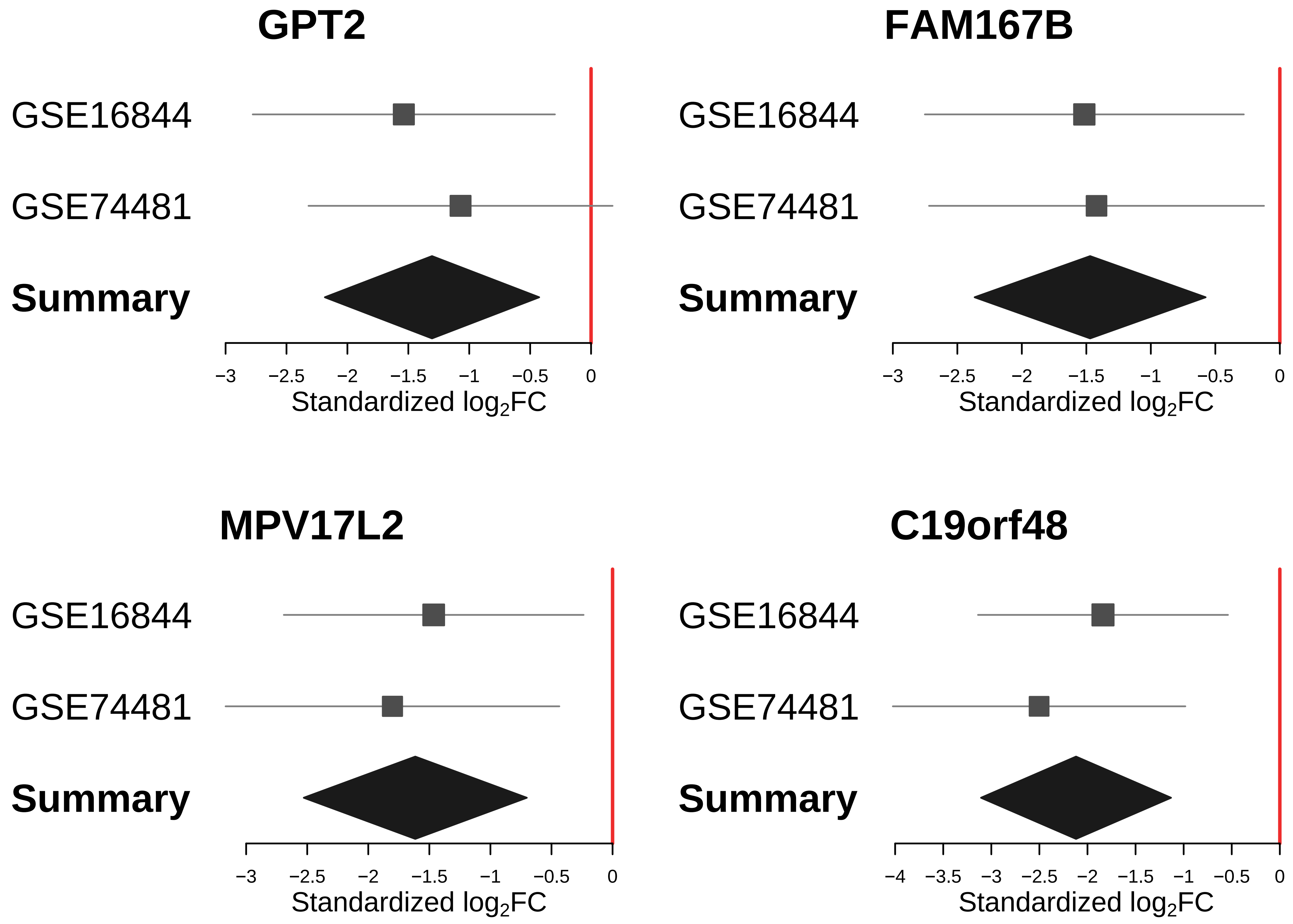
GPT2
GSE16844
GSE74481
Summary
−3 −2.5 −2 −1.5 −1 −0.5 0
Standardized log2FC
FAM167B
GSE16844
GSE74481
Summary
−3 −2.5 −2 −1.5 −1 −0.5 0
Standardized log2FC
MPV17L2
GSE16844
GSE74481
Summary
−3 −2.5 −2 −1.5 −1 −0.5 0
Standardized log2FC
C19orf48
GSE16844
GSE74481
Summary
−4 −3.5 −3 −2.5 −2 −1.5 −1 −0.5 0
Standardized log2FC

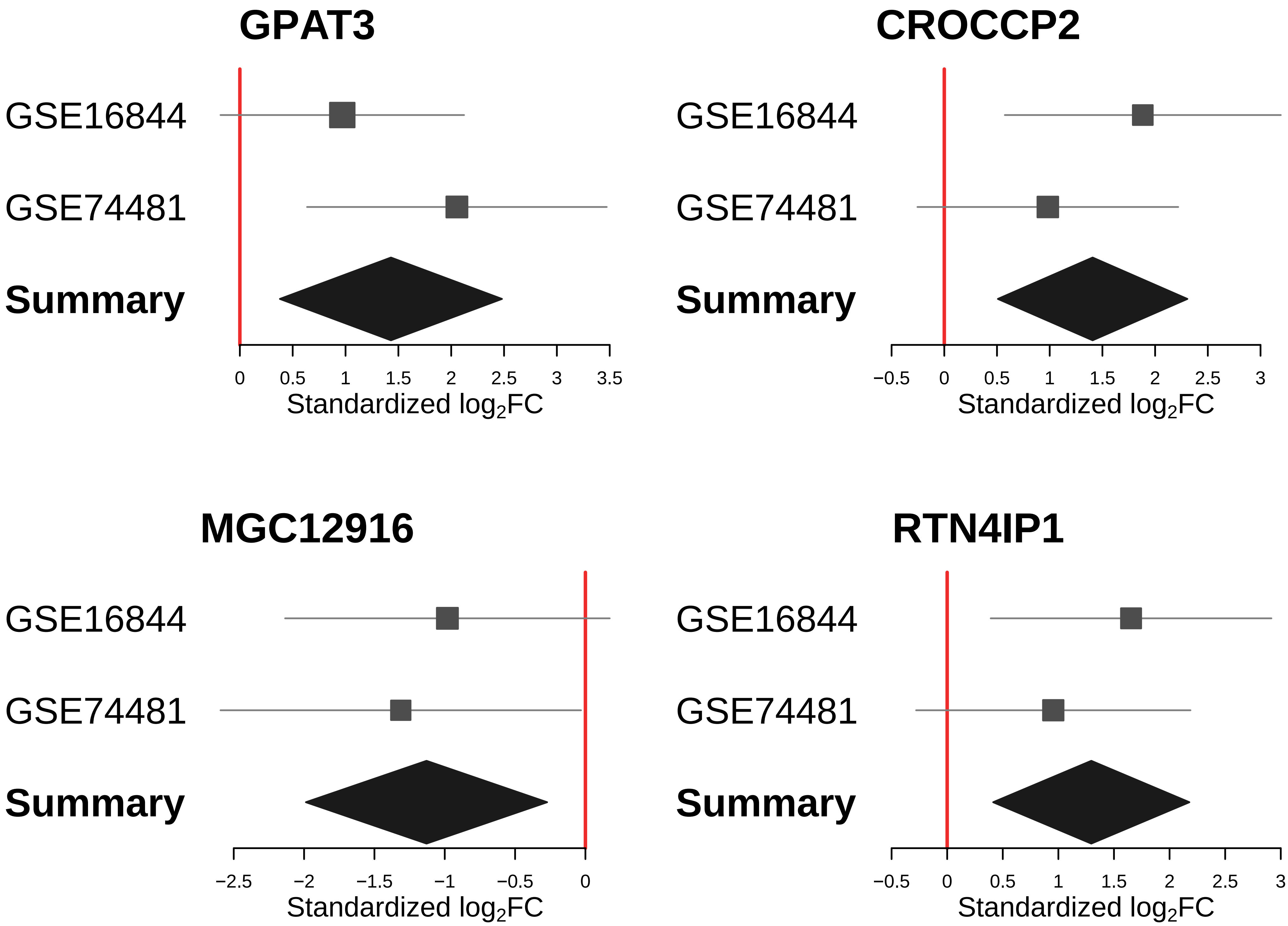
GPAT3
GSE16844
GSE74481
Summary
0
0.5
1
1.5
2
2.5
3
3.5
Standardized log2FC
CROCCP2
GSE16844
GSE74481
Summary
−0.5
0
0.5
1
1.5
2
2.5
3
Standardized log2FC
MGC12916
GSE16844
GSE74481
Summary
−2.5
−2
−1.5
−1
−0.5
0
Standardized log2FC
RTN4IP1
GSE16844
GSE74481
Summary
−0.5
0
0.5
1
1.5
2
2.5
3
Standardized log2FC

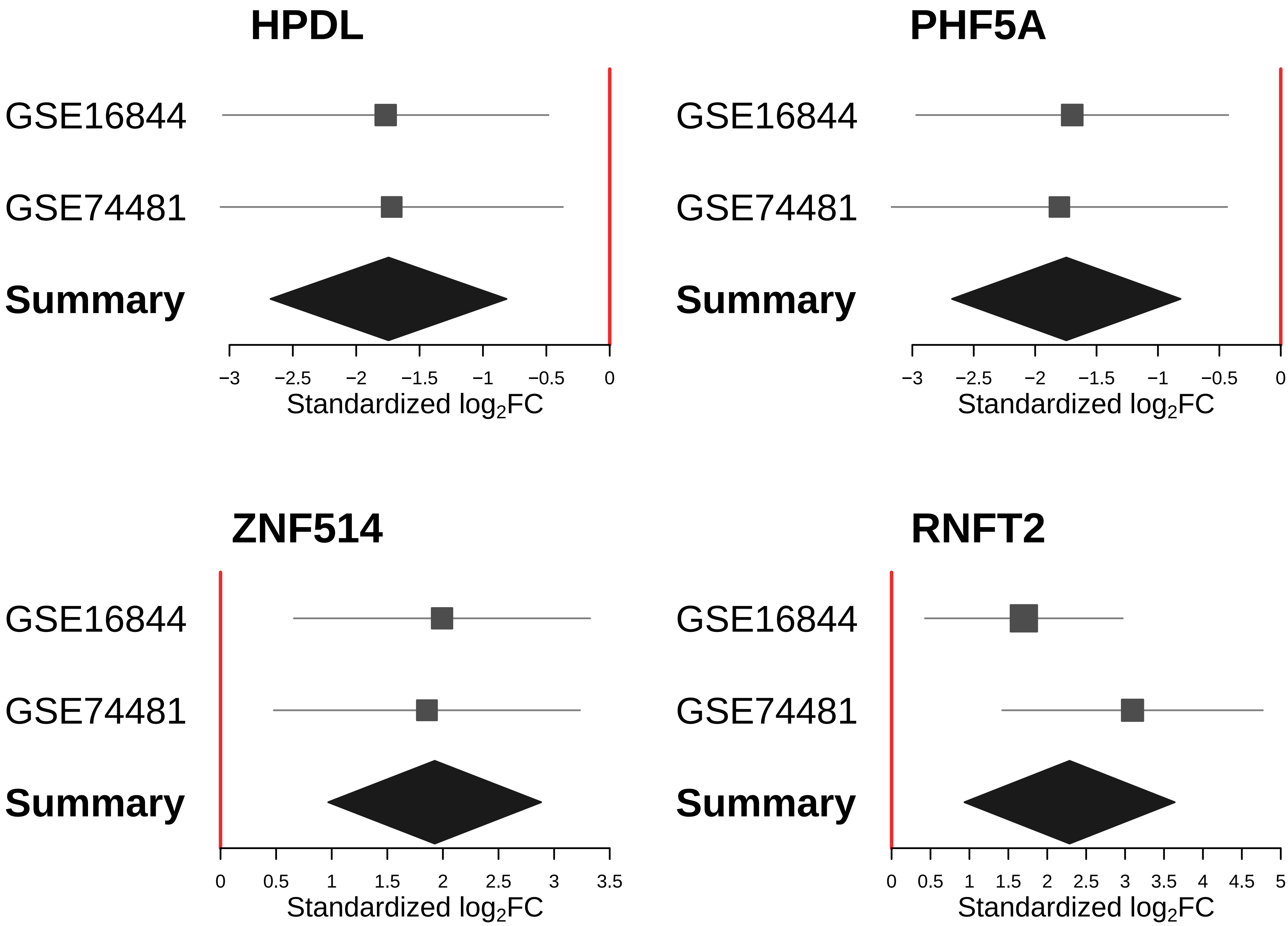
HPDL
GSE16844
GSE74481
Summary
−3
−2.5
−2
−1.5
−1
−0.5
0
Standardized $log_2FC$
PHF5A
GSE16844
GSE74481
Summary
−3
−2.5
−2
−1.5
−1
−0.5
0
Standardized $log_2FC$
ZNF514
GSE16844
GSE74481
Summary
0
0.5
1
1.5
2
2.5
3
3.5
Standardized $log_2FC$
RNFT2
GSE16844
GSE74481
Summary
0
0.5
1
1.5
2
2.5
3
3.5
4
4.5
5
Standardized $log_2FC$

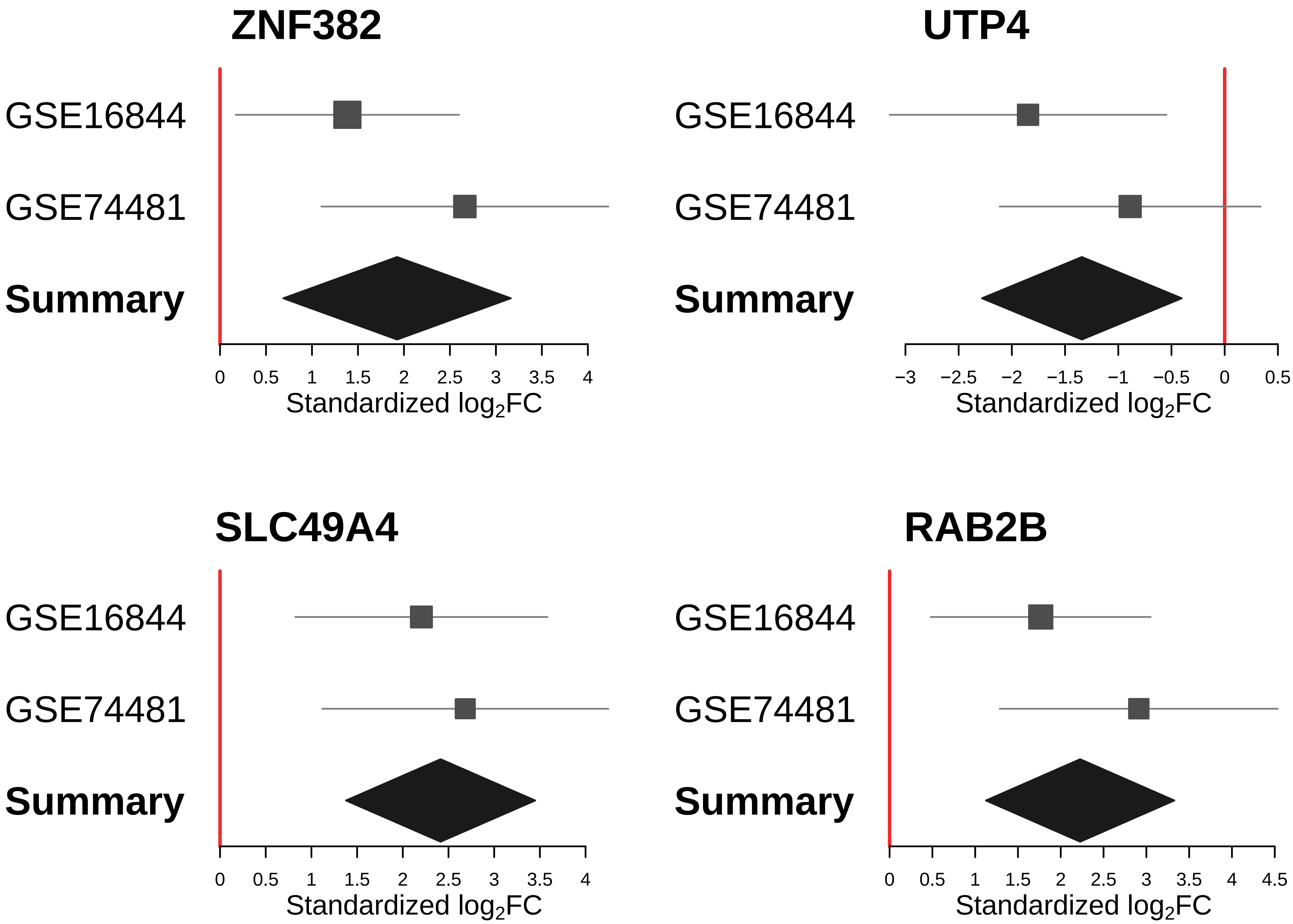
ZNF382
GSE16844
GSE74481
Summary
0 0.5 1 1.5 2 2.5 3 3.5 4
Standardized log2FC
UTP4
GSE16844
GSE74481
Summary
−3 −2.5 −2 −1.5 −1 −0.5 0 0.5
Standardized log2FC
SLC49A4
GSE16844
GSE74481
Summary
0 0.5 1 1.5 2 2.5 3 3.5 4
Standardized log2FC
RAB2B
GSE16844
GSE74481
Summary
0 0.5 1 1.5 2 2.5 3 3.5 4 4.5
Standardized log2FC

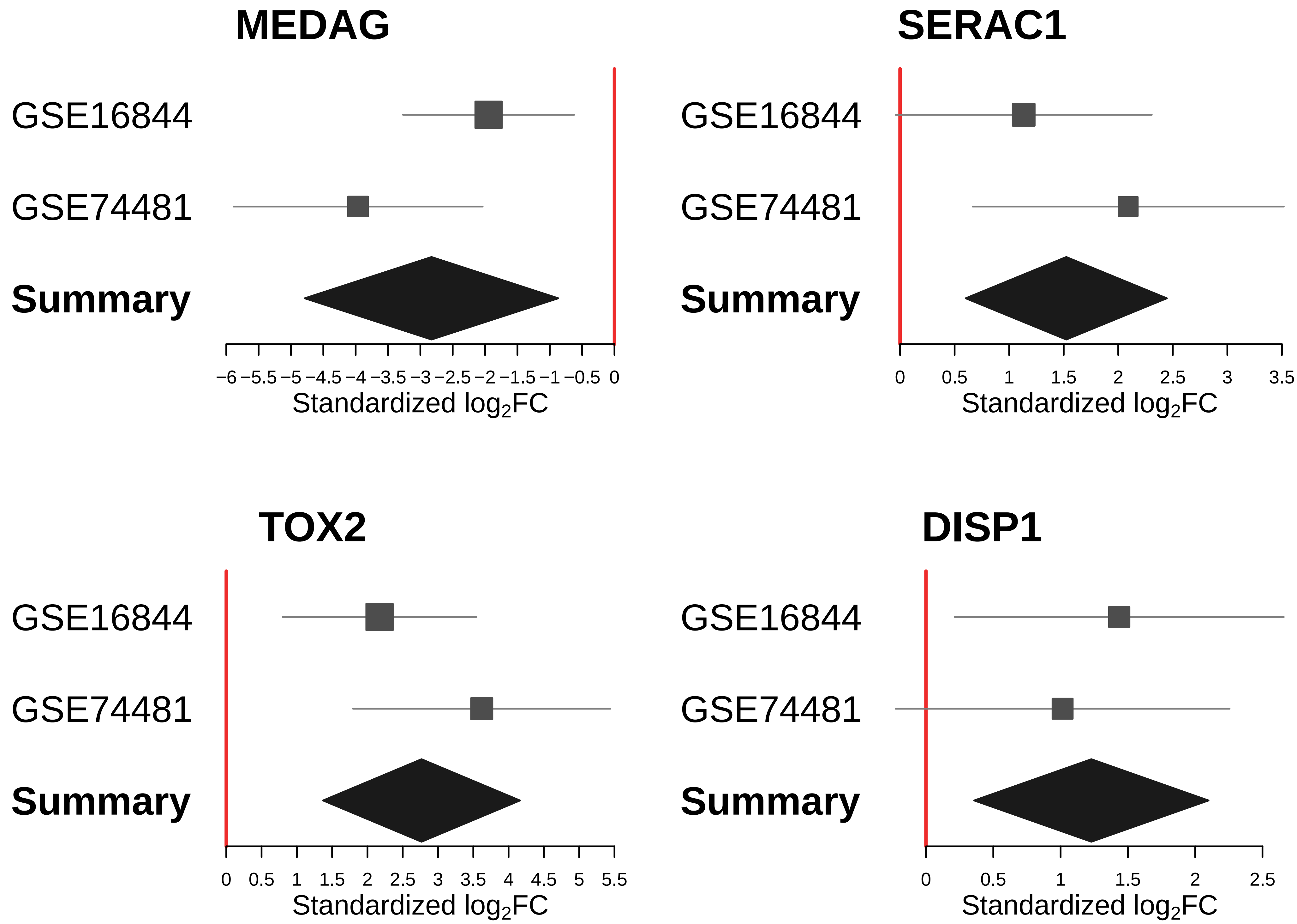
MEDAG
GSE16844
GSE74481
Summary
−6 −5.5 −5 −4.5 −4 −3.5 −3 −2.5 −2 −1.5 −1 −0.5 0
Standardized log2FC
SERAC1
GSE16844
GSE74481
Summary
0 0.5 1 1.5 2 2.5 3 3.5
Standardized log2FC
TOX2
GSE16844
GSE74481
Summary
0 0.5 1 1.5 2 2.5 3 3.5 4 4.5 5 5.5
Standardized log2FC
DISP1
GSE16844
GSE74481
Summary
0 0.5 1 1.5 2 2.5
Standardized log2FC

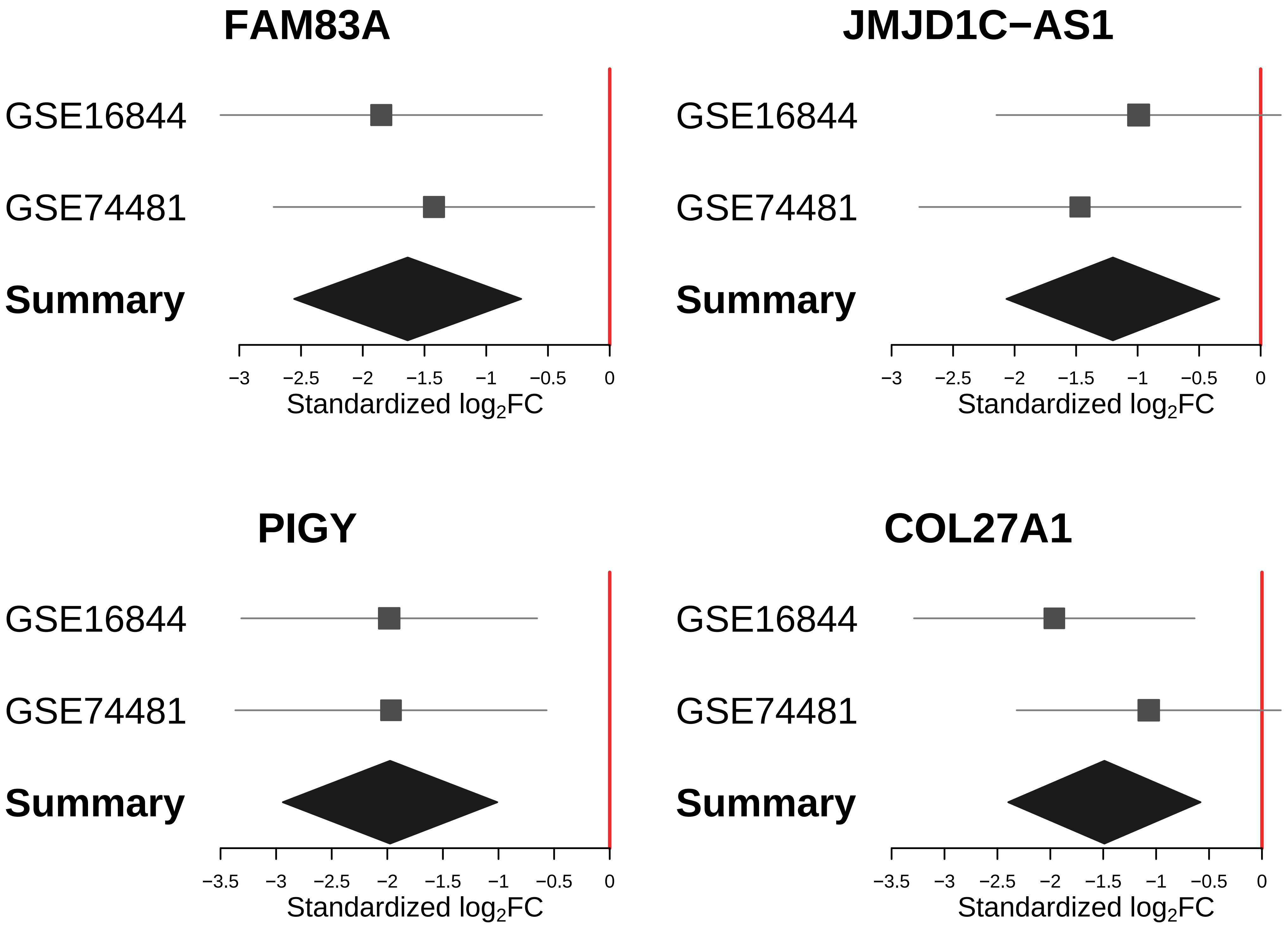
FAM83A
GSE16844
GSE74481
Summary
−3 −2.5 −2 −1.5 −1 −0.5 0
Standardized log2FC
JMJD1C−AS1
GSE16844
GSE74481
Summary
−3 −2.5 −2 −1.5 −1 −0.5 0
Standardized log2FC
PIGY
GSE16844
GSE74481
Summary
−3.5 −3 −2.5 −2 −1.5 −1 −0.5 0
Standardized log2FC
COL27A1
GSE16844
GSE74481
Summary
−3.5 −3 −2.5 −2 −1.5 −1 −0.5 0
Standardized log2FC

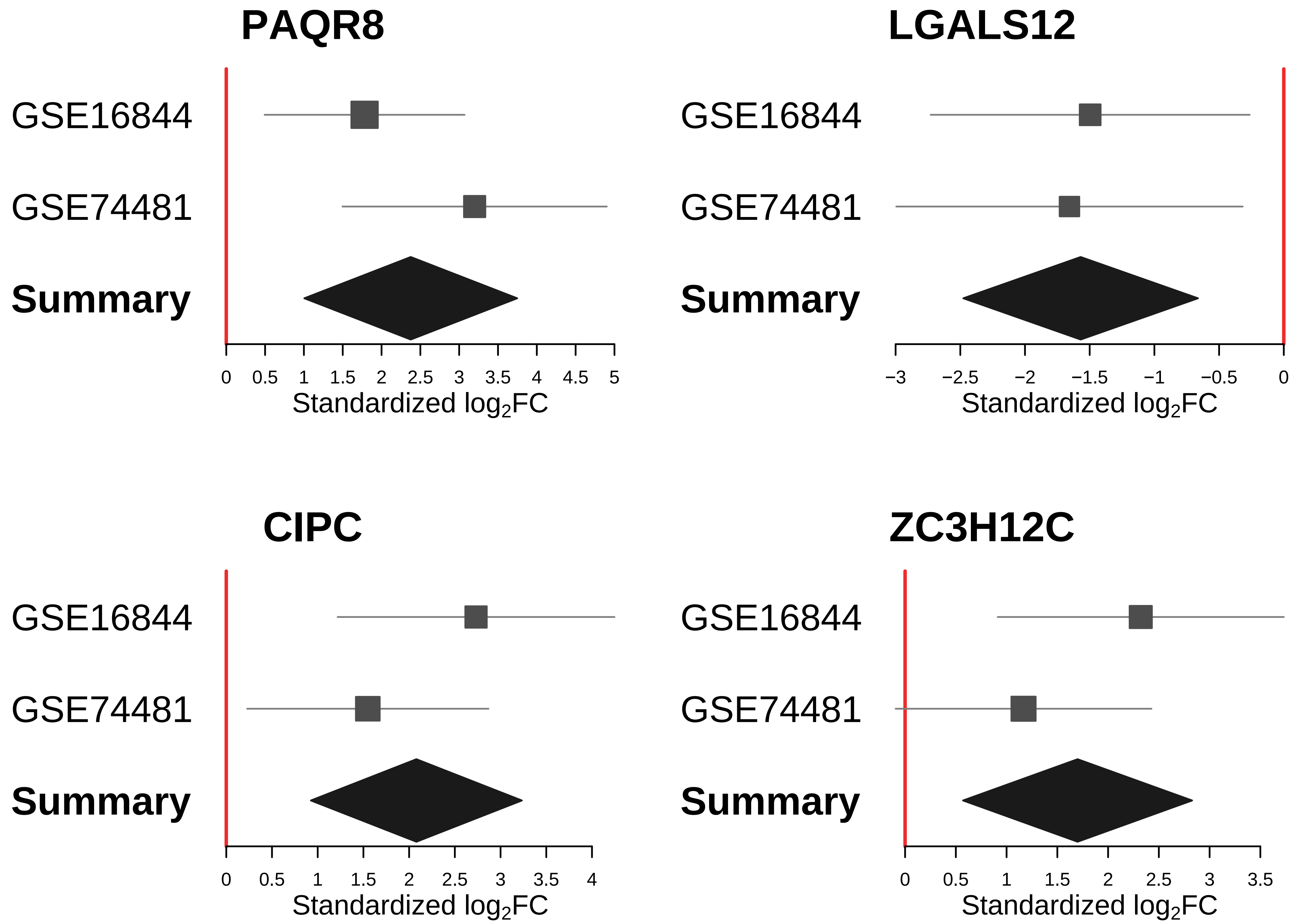

PAQR8
GSE16844
GSE74481
Summary
0 0.5 1 1.5 2 2.5 3 3.5 4 4.5 5
Standardized log2FC
LGALS12
GSE16844
GSE74481
Summary
−3 −2.5 −2 −1.5 −1 −0.5 0
Standardized log2FC
CIPC
GSE16844
GSE74481
Summary
0 0.5 1 1.5 2 2.5 3 3.5 4
Standardized log2FC
ZC3H12C
GSE16844
GSE74481
Summary
0 0.5 1 1.5 2 2.5 3 3.5
Standardized log2FC

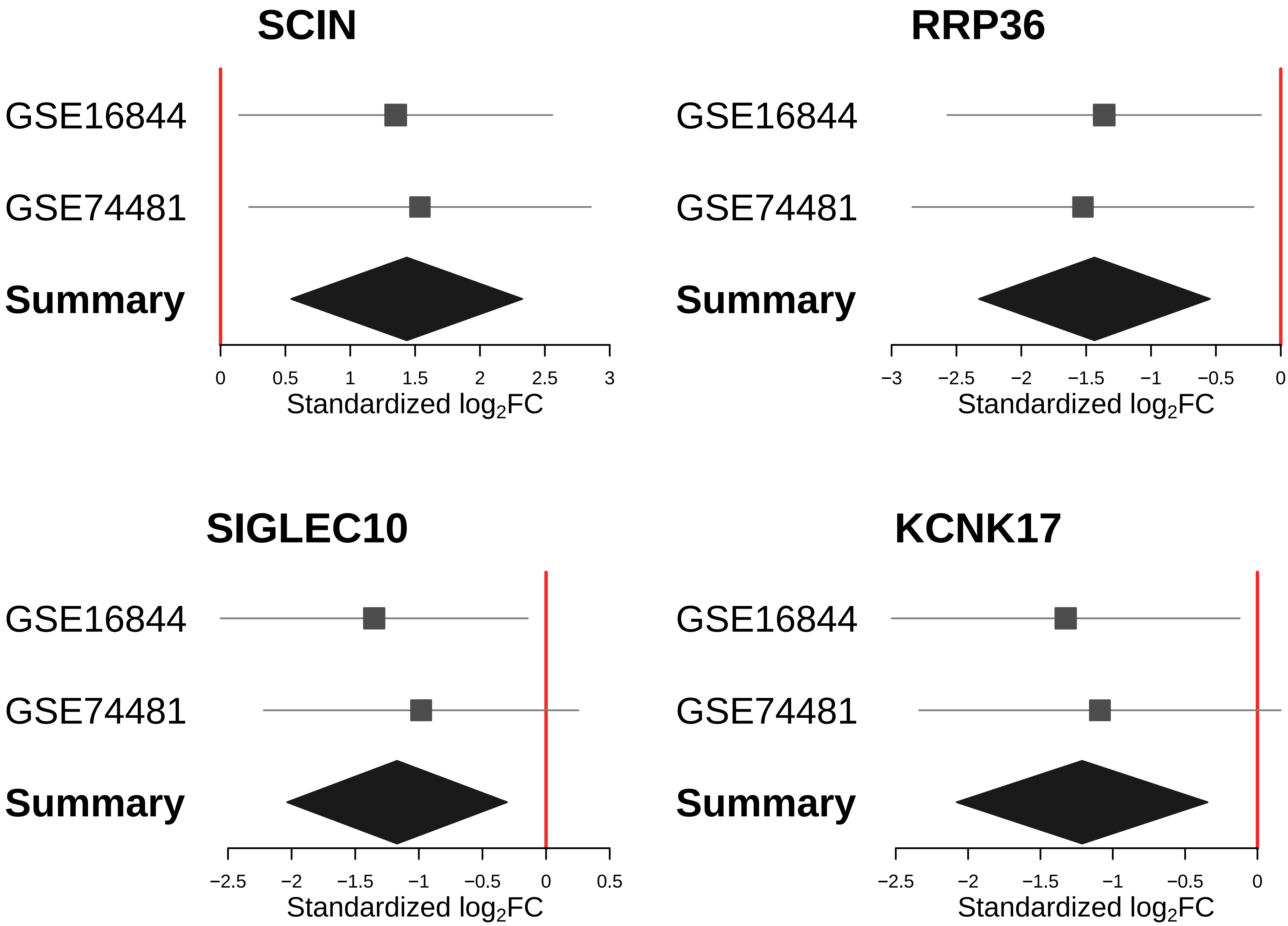
SCIN
GSE16844
GSE74481
Summary
0
0.5
1
1.5
2
2.5
3
Standardized log2FC
RRP36
GSE16844
GSE74481
Summary
−3
−2.5
−2
−1.5
−1
−0.5
0
Standardized log2FC
SIGLEC10
GSE16844
GSE74481
Summary
−2.5
−2
−1.5
−1
−0.5
0
0.5
Standardized log2FC
KCNK17
GSE16844
GSE74481
Summary
−2.5
−2
−1.5
−1
−0.5
0
Standardized log2FC

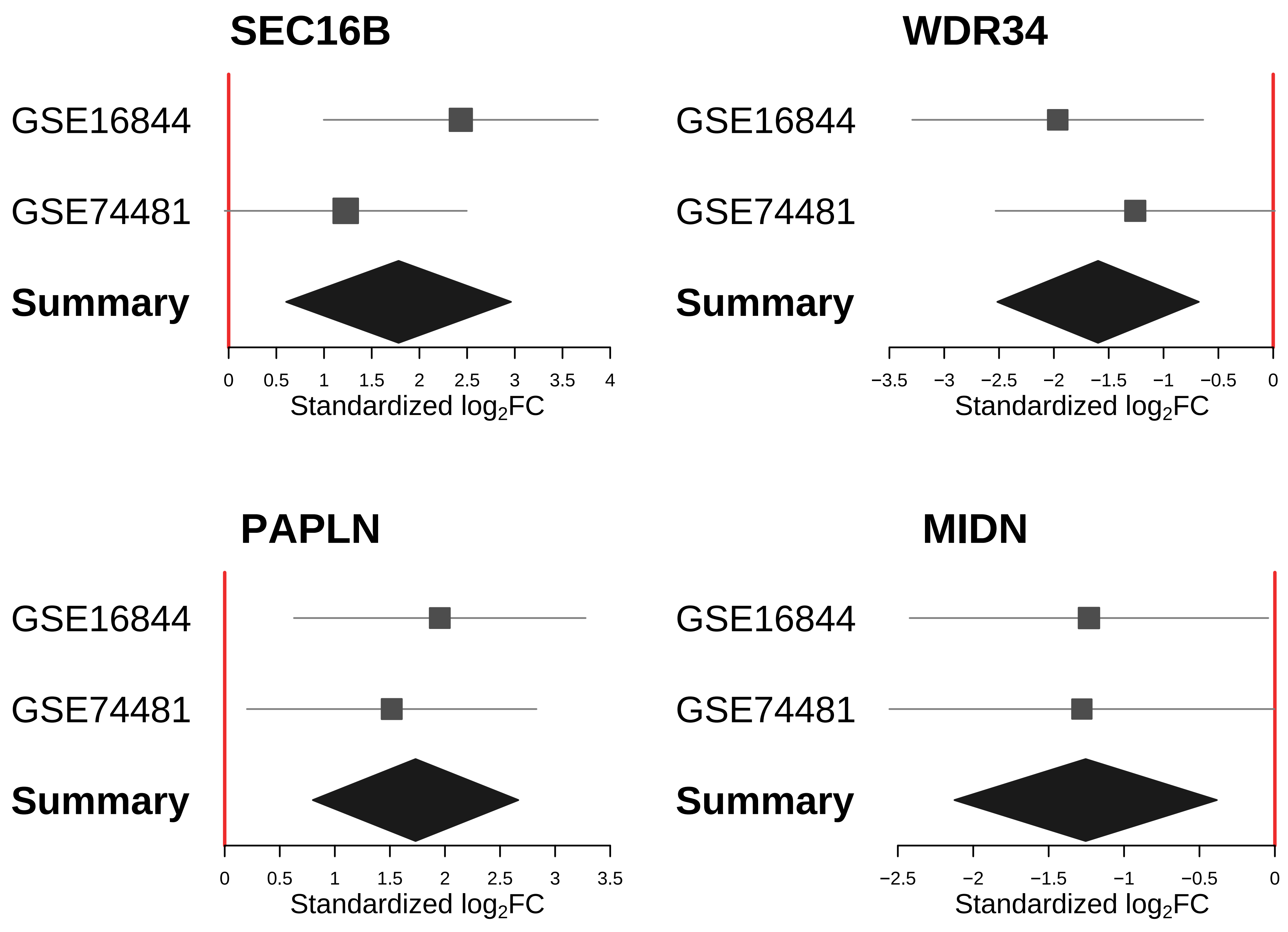
SEC16B
GSE16844
GSE74481
Summary
0 0.5 1 1.5 2 2.5 3 3.5 4
Standardized log2FC
WDR34
GSE16844
GSE74481
Summary
−3.5 −3 −2.5 −2 −1.5 −1 −0.5 0
Standardized log2FC
PAPLN
GSE16844
GSE74481
Summary
0 0.5 1 1.5 2 2.5 3 3.5
Standardized log2FC
MIDN
GSE16844
GSE74481
Summary
−2.5 −2 −1.5 −1 −0.5 0
Standardized log2FC

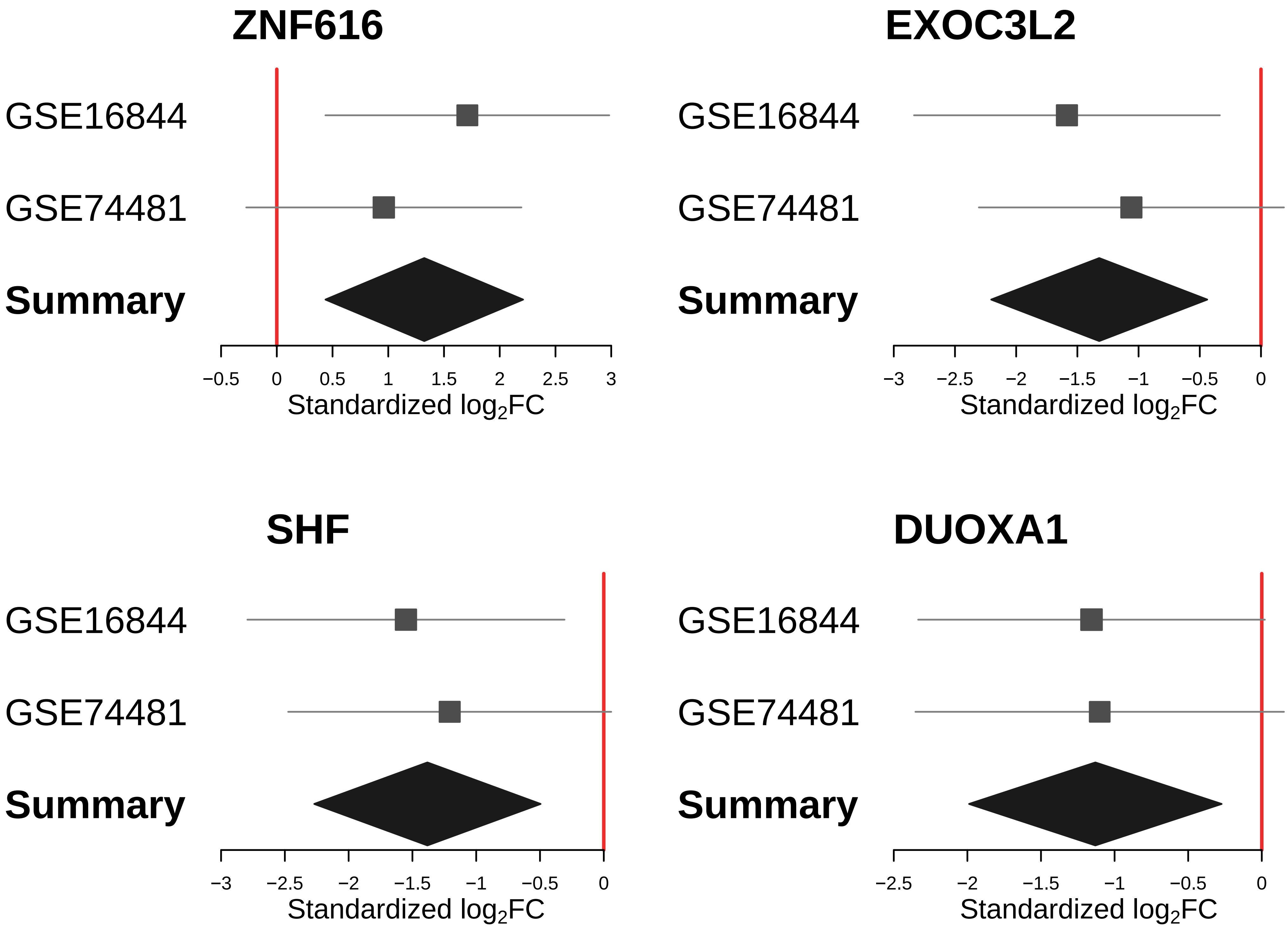
ZNF616
GSE16844
GSE74481
Summary
−0.5 0 0.5 1 1.5 2 2.5 3
Standardized log2FC
EXOC3L2
GSE16844
GSE74481
Summary
−3 −2.5 −2 −1.5 −1 −0.5 0
Standardized log2FC
SHF
GSE16844
GSE74481
Summary
−3 −2.5 −2 −1.5 −1 −0.5 0
Standardized log2FC
DUOXA1
GSE16844
GSE74481
Summary
−2.5 −2 −1.5 −1 −0.5 0
Standardized log2FC

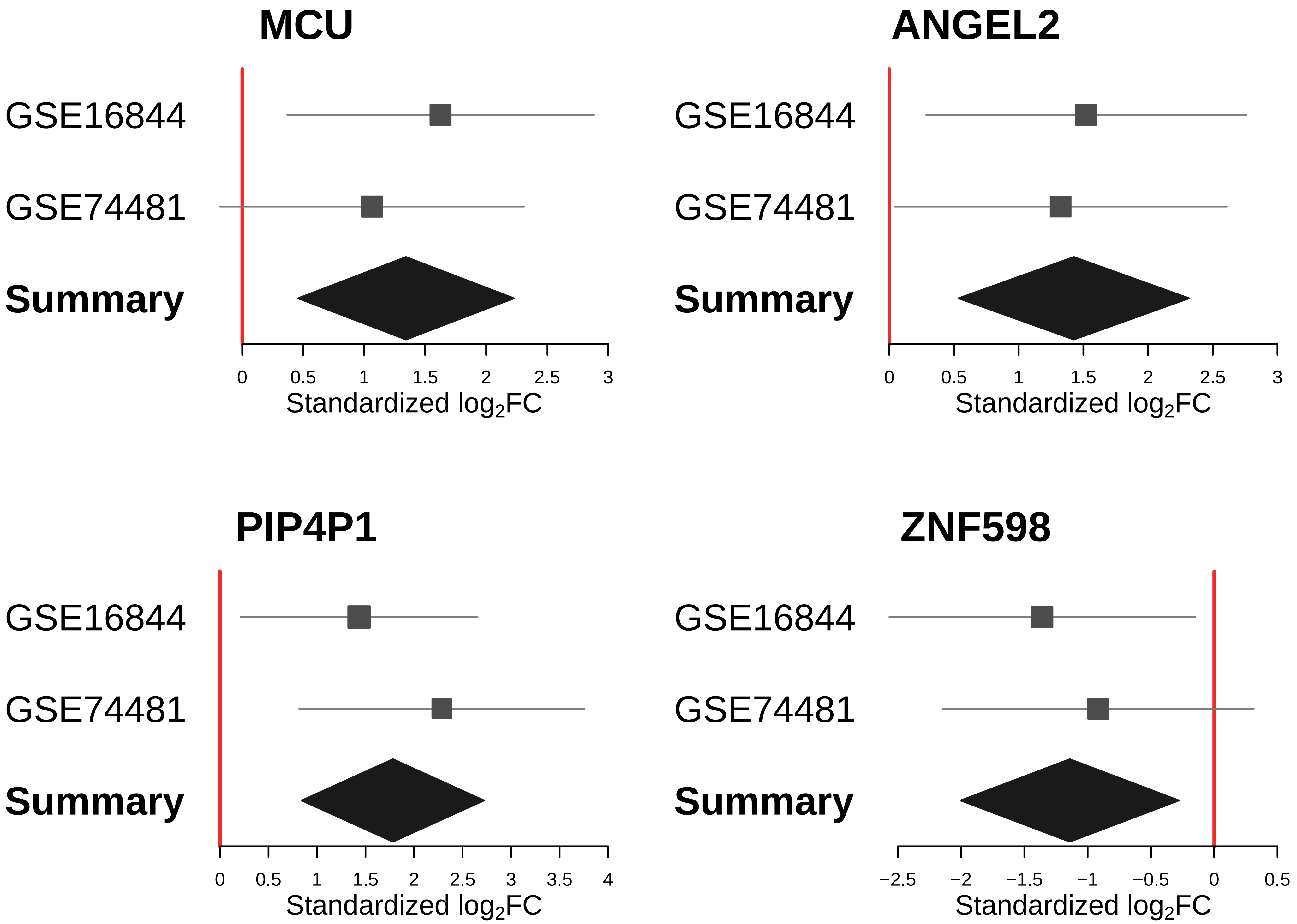
MCU
GSE16844
GSE74481
Summary
0
0.5
1
1.5
2
2.5
3
Standardized log2FC
ANGEL2
GSE16844
GSE74481
Summary
0
0.5
1
1.5
2
2.5
3
Standardized log2FC
PIP4P1
GSE16844
GSE74481
Summary
0
0.5
1
1.5
2
2.5
3
3.5
4
Standardized log2FC
ZNF598
GSE16844
GSE74481
Summary
−2.5
−2
−1.5
−1
−0.5
0
0.5
Standardized log2FC

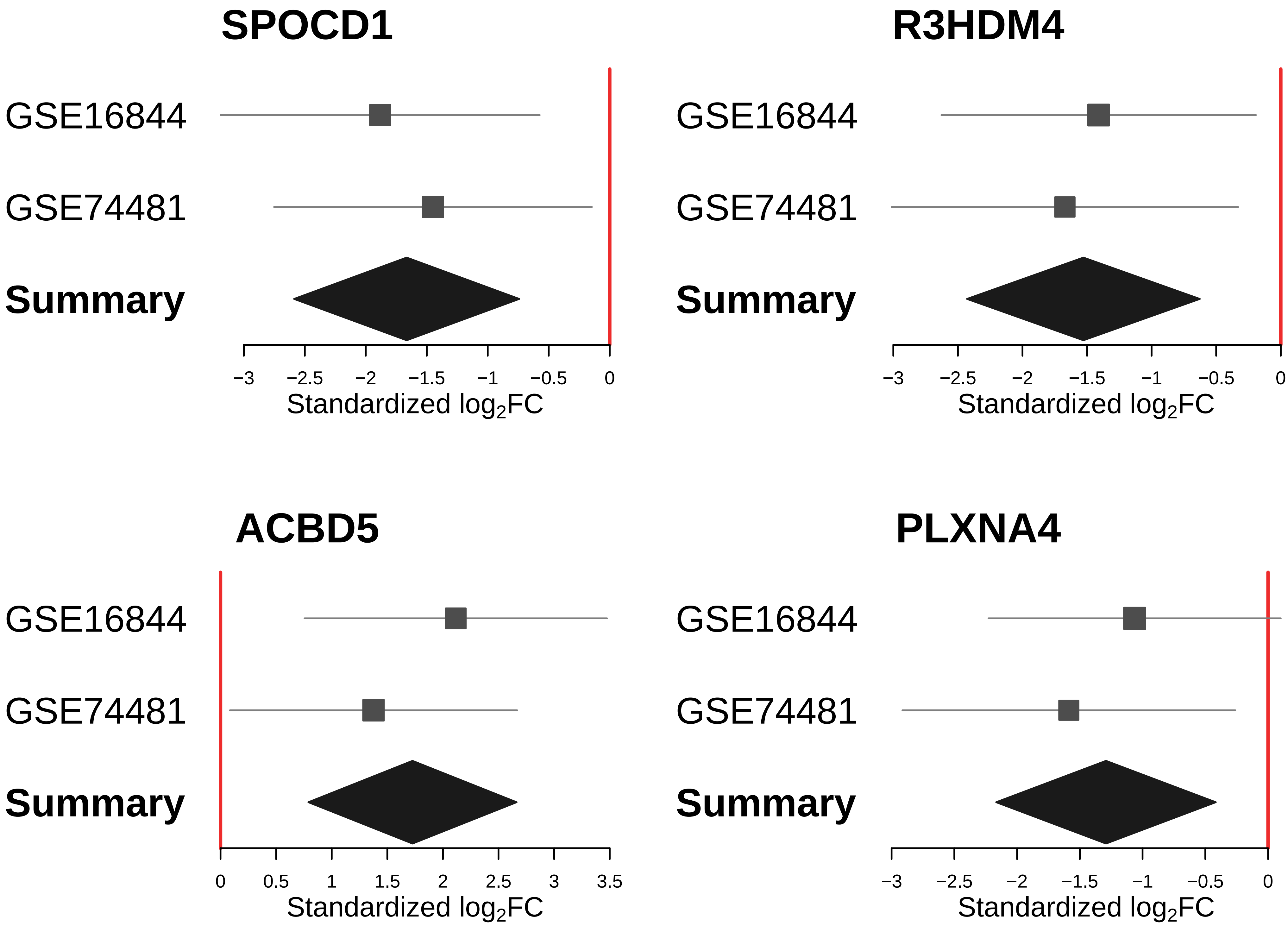
SPOCD1
GSE16844
GSE74481
Summary
−3 −2.5 −2 −1.5 −1 −0.5 0
Standardized log2FC
R3HDM4
GSE16844
GSE74481
Summary
−3 −2.5 −2 −1.5 −1 −0.5 0
Standardized log2FC
ACBD5
GSE16844
GSE74481
Summary
0 0.5 1 1.5 2 2.5 3 3.5
Standardized log2FC
PLXNA4
GSE16844
GSE74481
Summary
−3 −2.5 −2 −1.5 −1 −0.5 0
Standardized log2FC

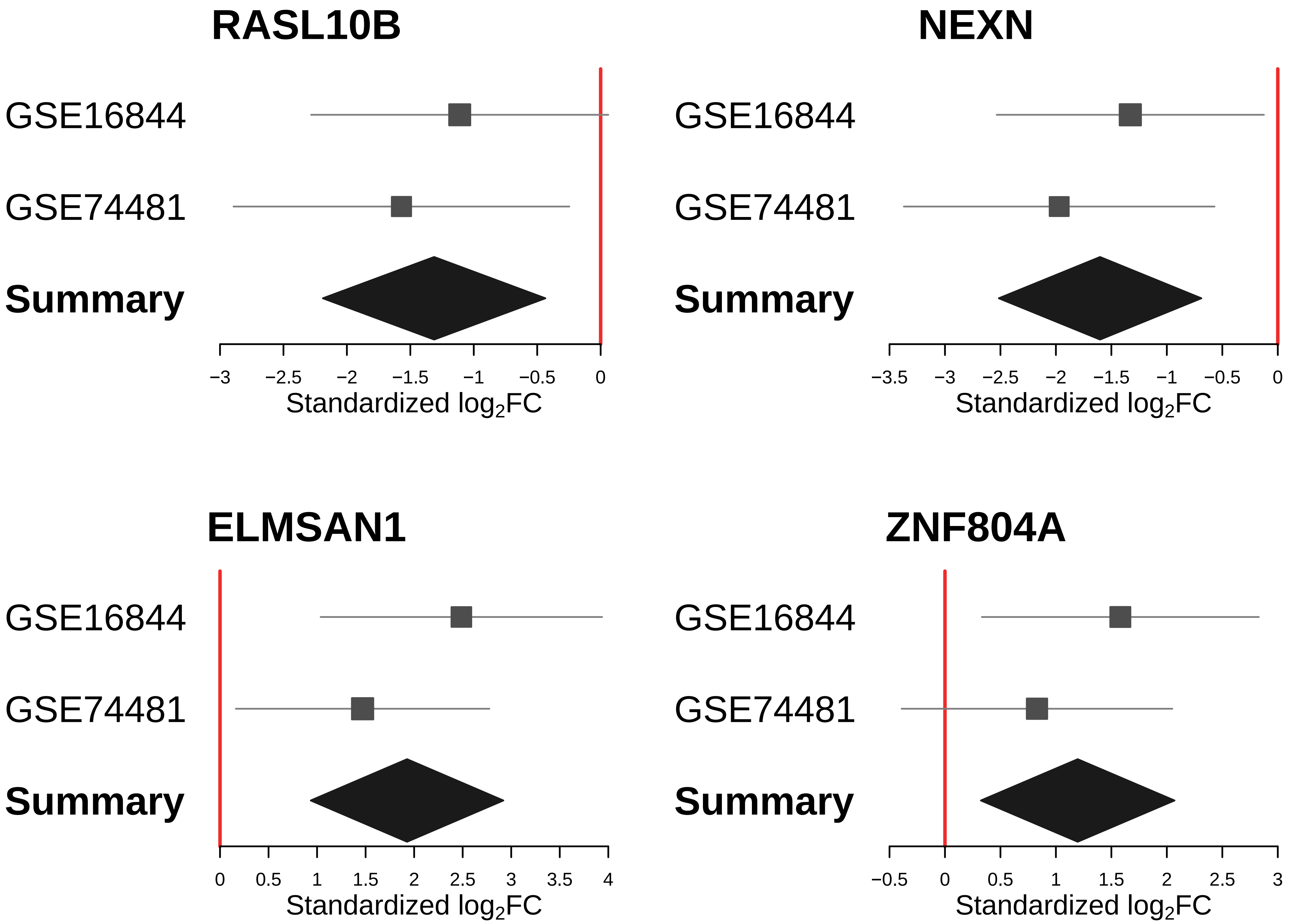
RASL10B
GSE16844
GSE74481
Summary
−3
−2.5
−2
−1.5
−1
−0.5
0
Standardized log2FC
NEXN
GSE16844
GSE74481
Summary
−3.5
−3
−2.5
−2
−1.5
−1
−0.5
0
Standardized log2FC
ELMSAN1
GSE16844
GSE74481
Summary
0
0.5
1
1.5
2
2.5
3
3.5
4
Standardized log2FC
ZNF804A
GSE16844
GSE74481
Summary
−0.5
0
0.5
1
1.5
2
2.5
3
Standardized log2FC

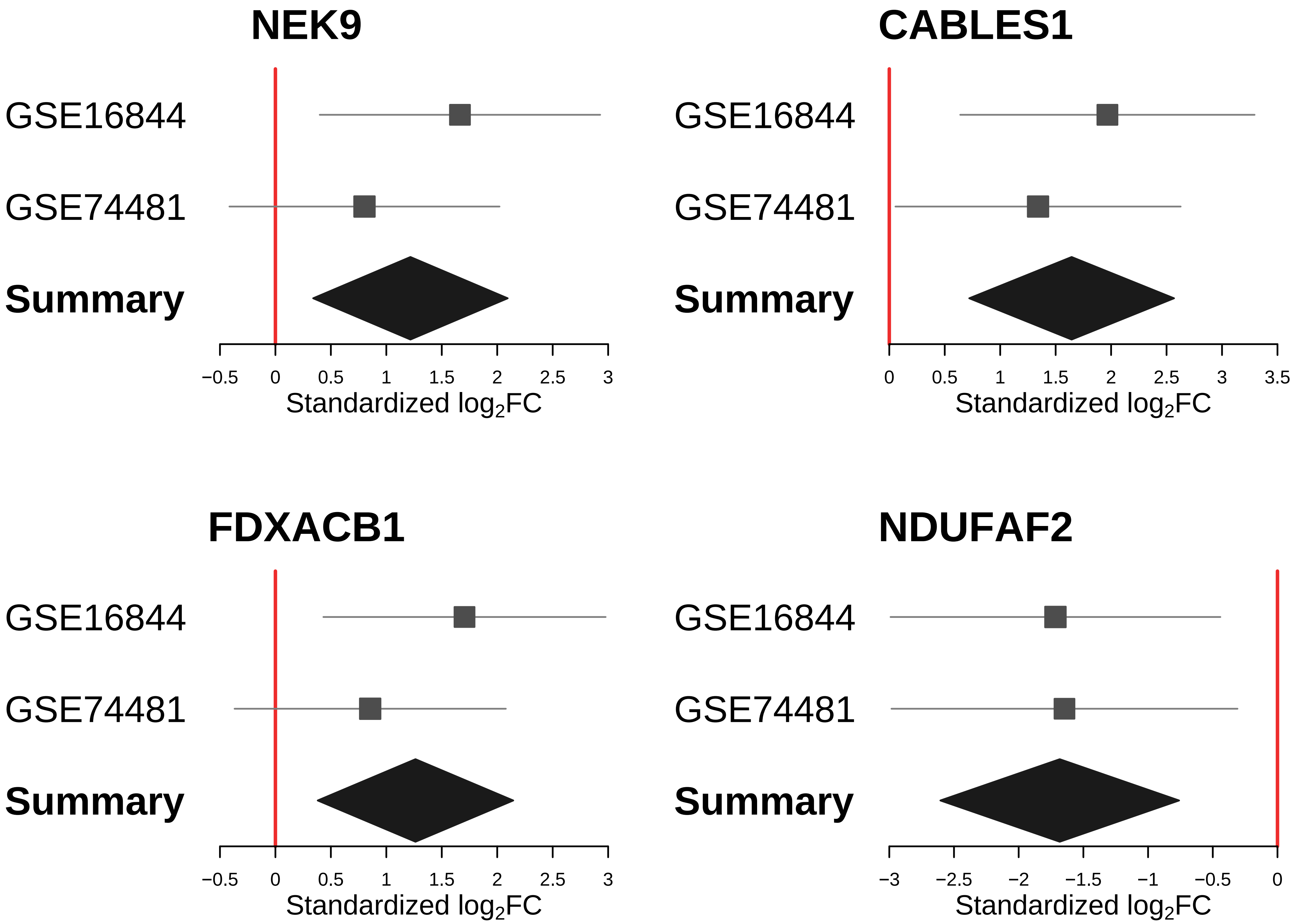
NEK9
GSE16844
GSE74481
Summary
−0.5 0 0.5 1 1.5 2 2.5 3
Standardized log2FC
CABLES1
GSE16844
GSE74481
Summary
0 0.5 1 1.5 2 2.5 3 3.5
Standardized log2FC
FDXACB1
GSE16844
GSE74481
Summary
−0.5 0 0.5 1 1.5 2 2.5 3
Standardized log2FC
NDUFAF2
GSE16844
GSE74481
Summary
−3 −2.5 −2 −1.5 −1 −0.5 0
Standardized log2FC

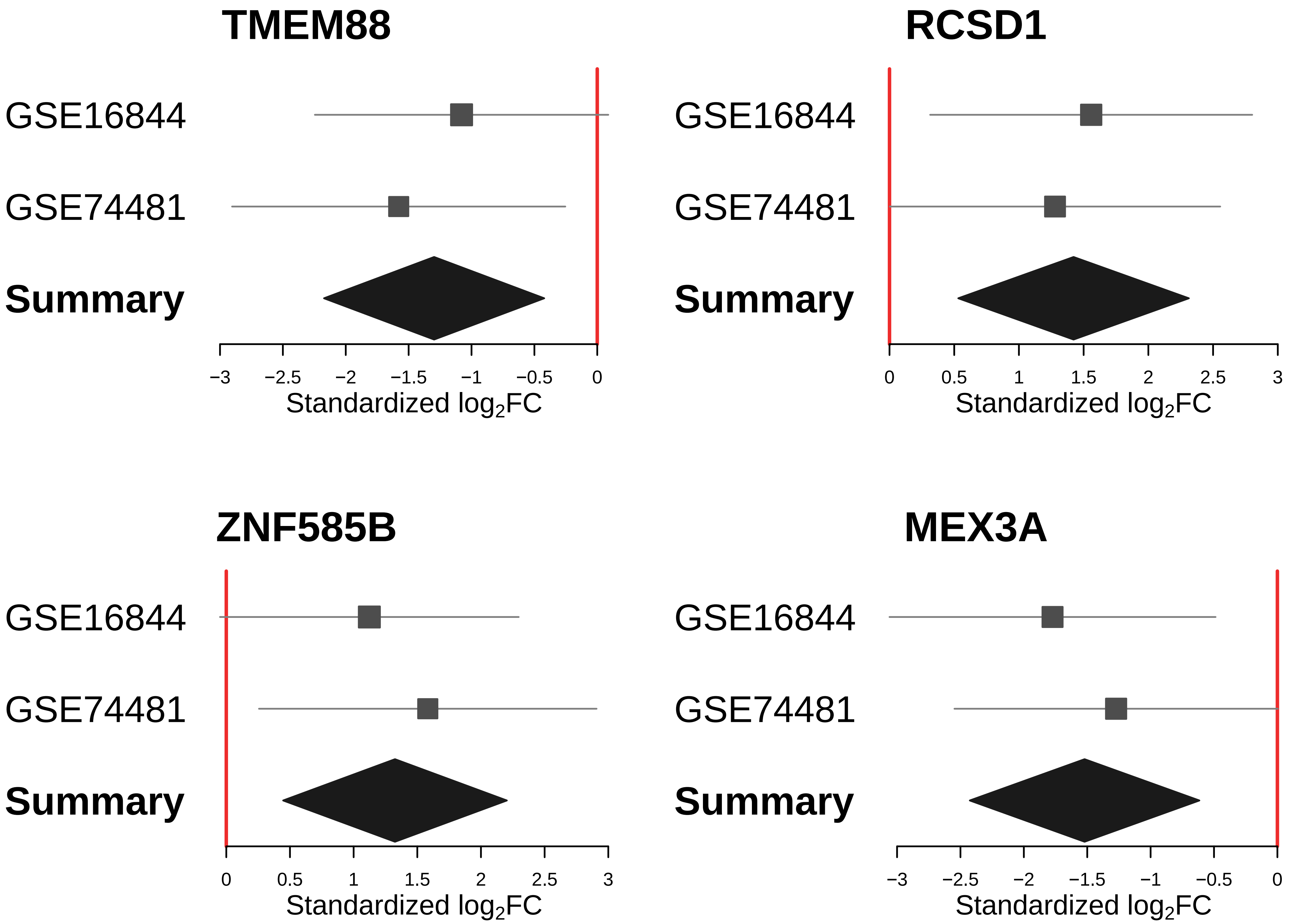
TMEM88
GSE16844
GSE74481
Summary
−3 −2.5 −2 −1.5 −1 −0.5 0
Standardized log2FC
RCSD1
GSE16844
GSE74481
Summary
0 0.5 1 1.5 2 2.5 3
Standardized log2FC
ZNF585B
GSE16844
GSE74481
Summary
0 0.5 1 1.5 2 2.5 3
Standardized log2FC
MEX3A
GSE16844
GSE74481
Summary
−3 −2.5 −2 −1.5 −1 −0.5 0
Standardized log2FC

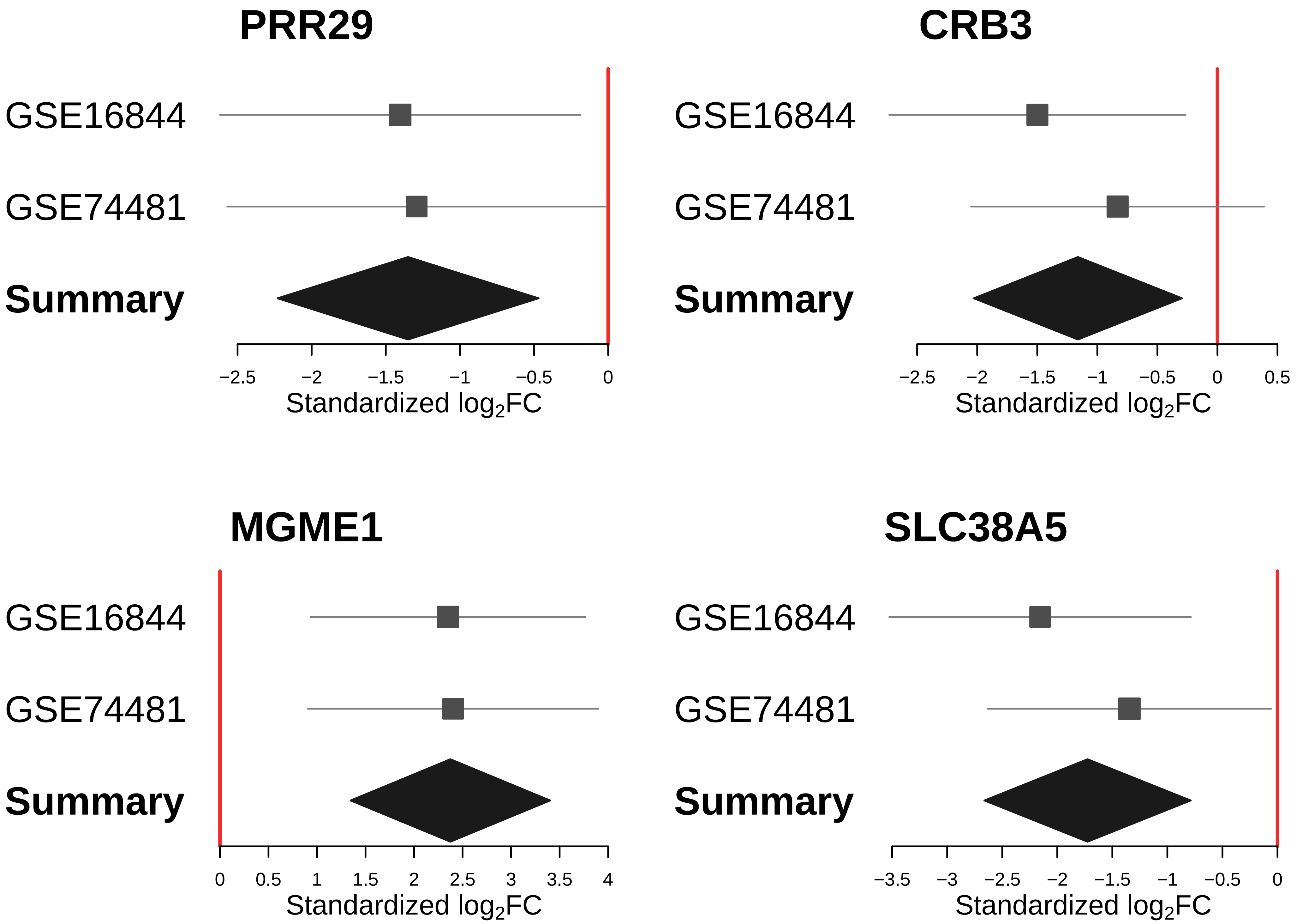
PRR29
GSE16844
GSE74481
Summary
−2.5 −2 −1.5 −1 −0.5 0
Standardized log2FC
CRB3
GSE16844
GSE74481
Summary
−2.5 −2 −1.5 −1 −0.5 0 0.5
Standardized log2FC
MGME1
GSE16844
GSE74481
Summary
0 0.5 1 1.5 2 2.5 3 3.5 4
Standardized log2FC
SLC38A5
GSE16844
GSE74481
Summary
−3.5 −3 −2.5 −2 −1.5 −1 −0.5 0
Standardized log2FC

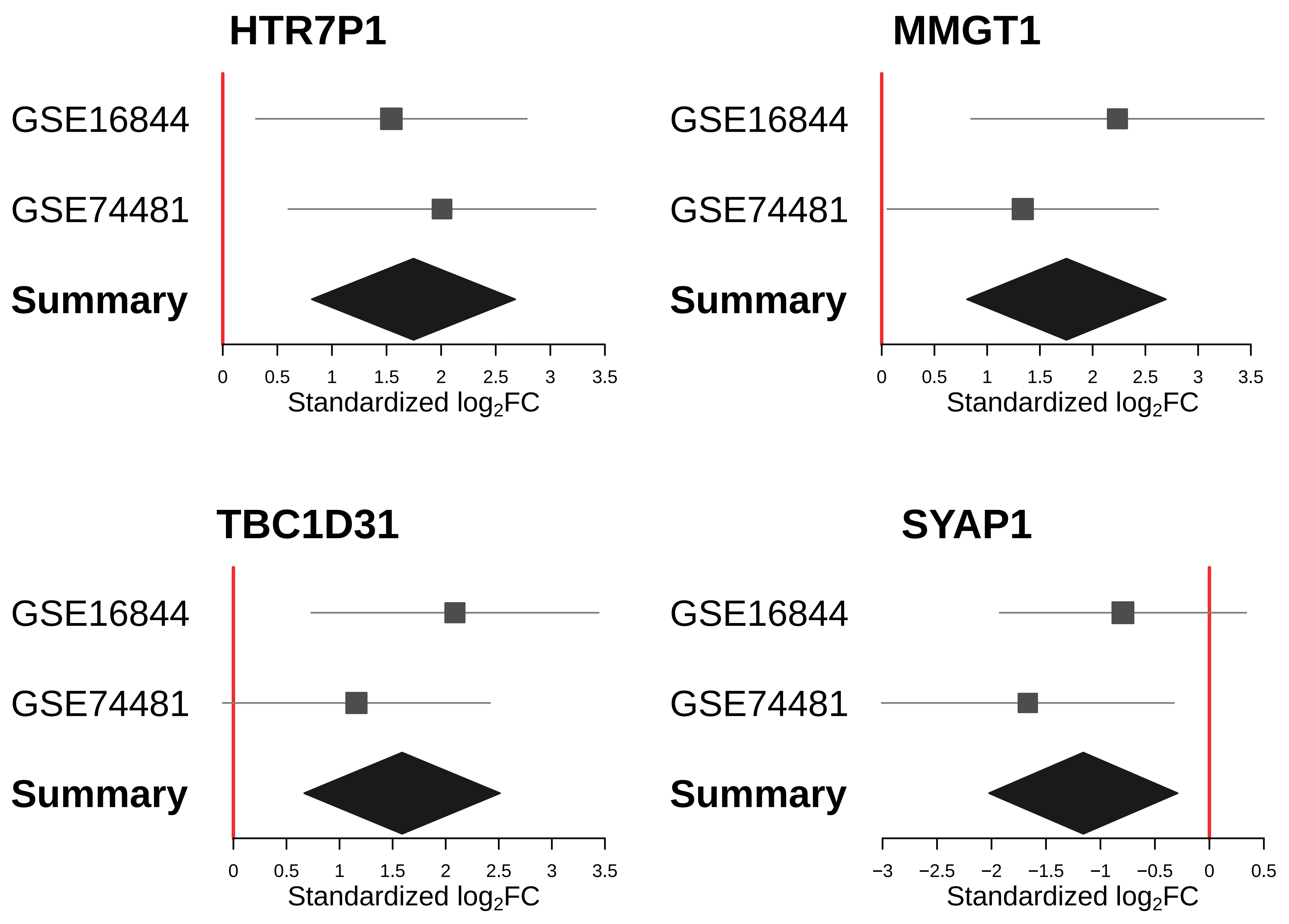

HTR7P1
GSE16844
GSE74481
Summary
0
0.5
1
1.5
2
2.5
3
3.5
Standardized log2FC
MMGT1
GSE16844
GSE74481
Summary
0
0.5
1
1.5
2
2.5
3
3.5
Standardized log2FC
TBC1D31
GSE16844
GSE74481
Summary
0
0.5
1
1.5
2
2.5
3
3.5
Standardized log2FC
SYAP1
GSE16844
GSE74481
Summary
−3
−2.5
−2
−1.5
−1
−0.5
0
0.5
Standardized log2FC

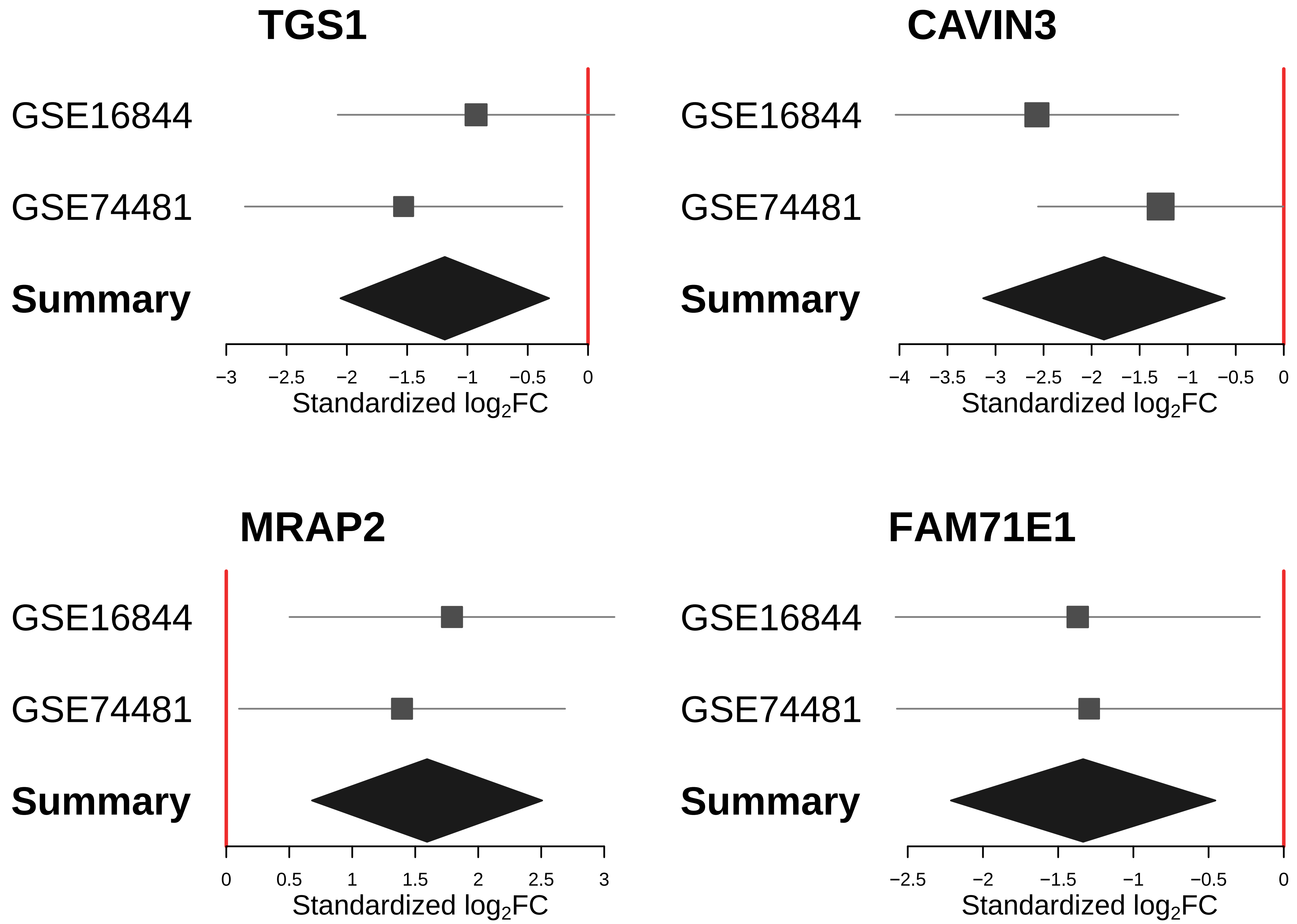
TGS1
GSE16844
GSE74481
Summary
−3 −2.5 −2 −1.5 −1 −0.5 0
Standardized log2FC
CAVIN3
GSE16844
GSE74481
Summary
−4 −3.5 −3 −2.5 −2 −1.5 −1 −0.5 0
Standardized log2FC
MRAP2
GSE16844
GSE74481
Summary
0 0.5 1 1.5 2 2.5 3
Standardized log2FC
FAM71E1
GSE16844
GSE74481
Summary
−2.5 −2 −1.5 −1 −0.5 0
Standardized log2FC

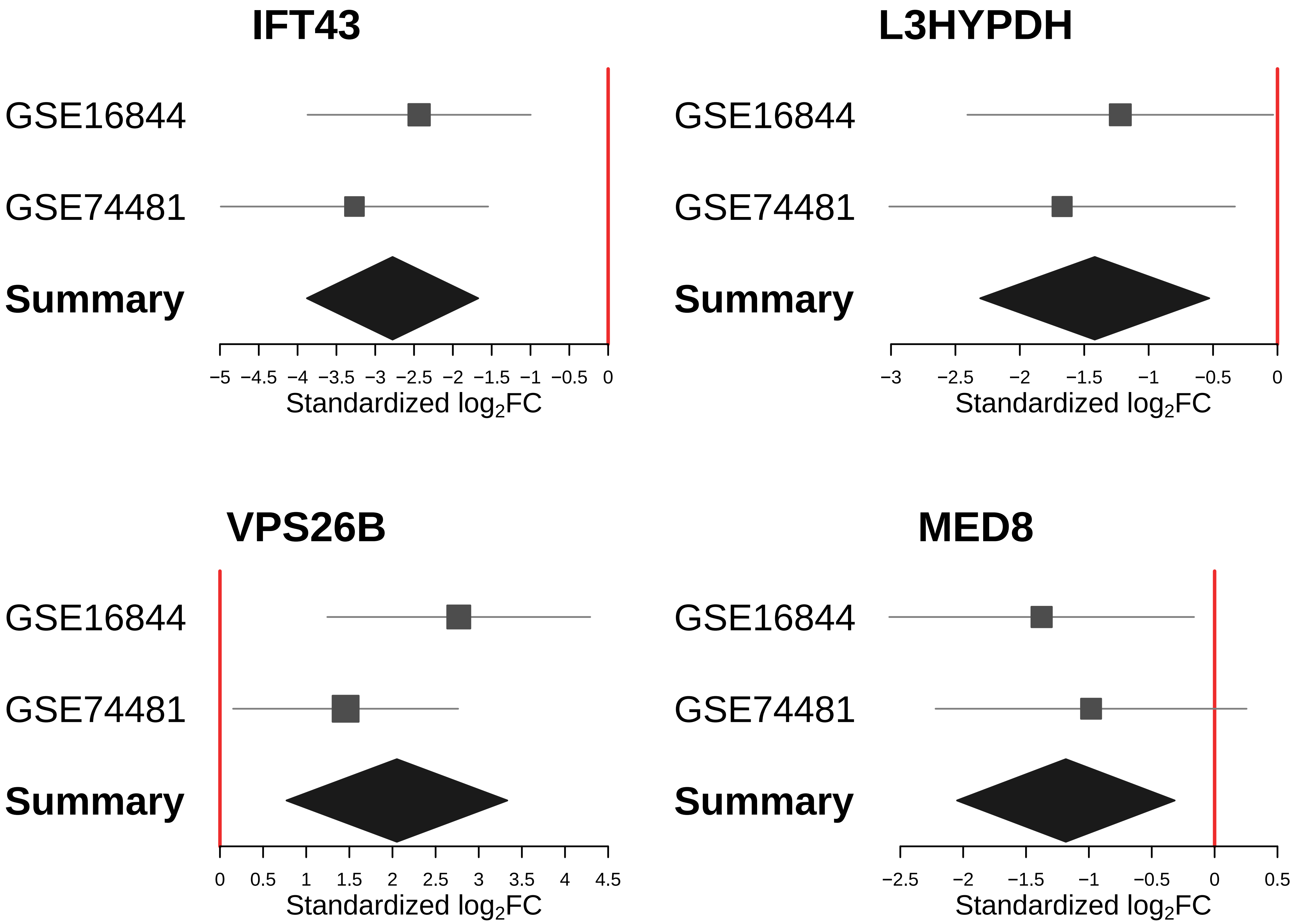
IFT43
GSE16844
GSE74481
Summary
−5 −4.5 −4 −3.5 −3 −2.5 −2 −1.5 −1 −0.5 0
Standardized log2FC
L3HYPDH
GSE16844
GSE74481
Summary
−3 −2.5 −2 −1.5 −1 −0.5 0
Standardized log2FC
VPS26B
GSE16844
GSE74481
Summary
0 0.5 1 1.5 2 2.5 3 3.5 4 4.5
Standardized log2FC
MED8
GSE16844
GSE74481
Summary
−2.5 −2 −1.5 −1 −0.5 0 0.5
Standardized log2FC

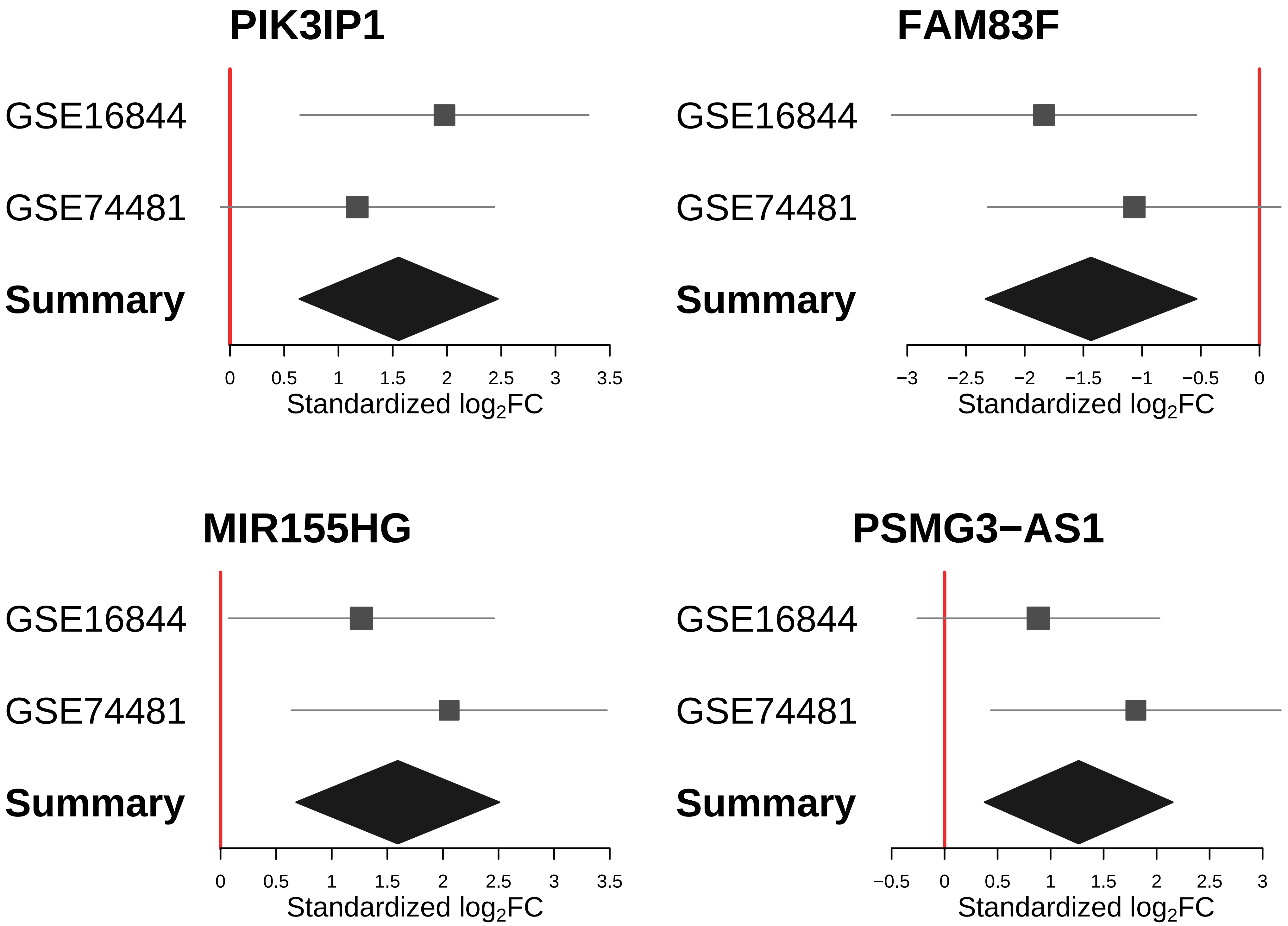

PIK3IP1
GSE16844
GSE74481
Summary
0 0.5 1 1.5 2 2.5 3 3.5
Standardized $log_2FC$
FAM83F
GSE16844
GSE74481
Summary
−3 −2.5 −2 −1.5 −1 −0.5 0
Standardized $log_2FC$
MIR155HG
GSE16844
GSE74481
Summary
0 0.5 1 1.5 2 2.5 3 3.5
Standardized $log_2FC$
PSMG3−AS1
GSE16844
GSE74481
Summary
−0.5 0 0.5 1 1.5 2 2.5 3
Standardized $log_2FC$

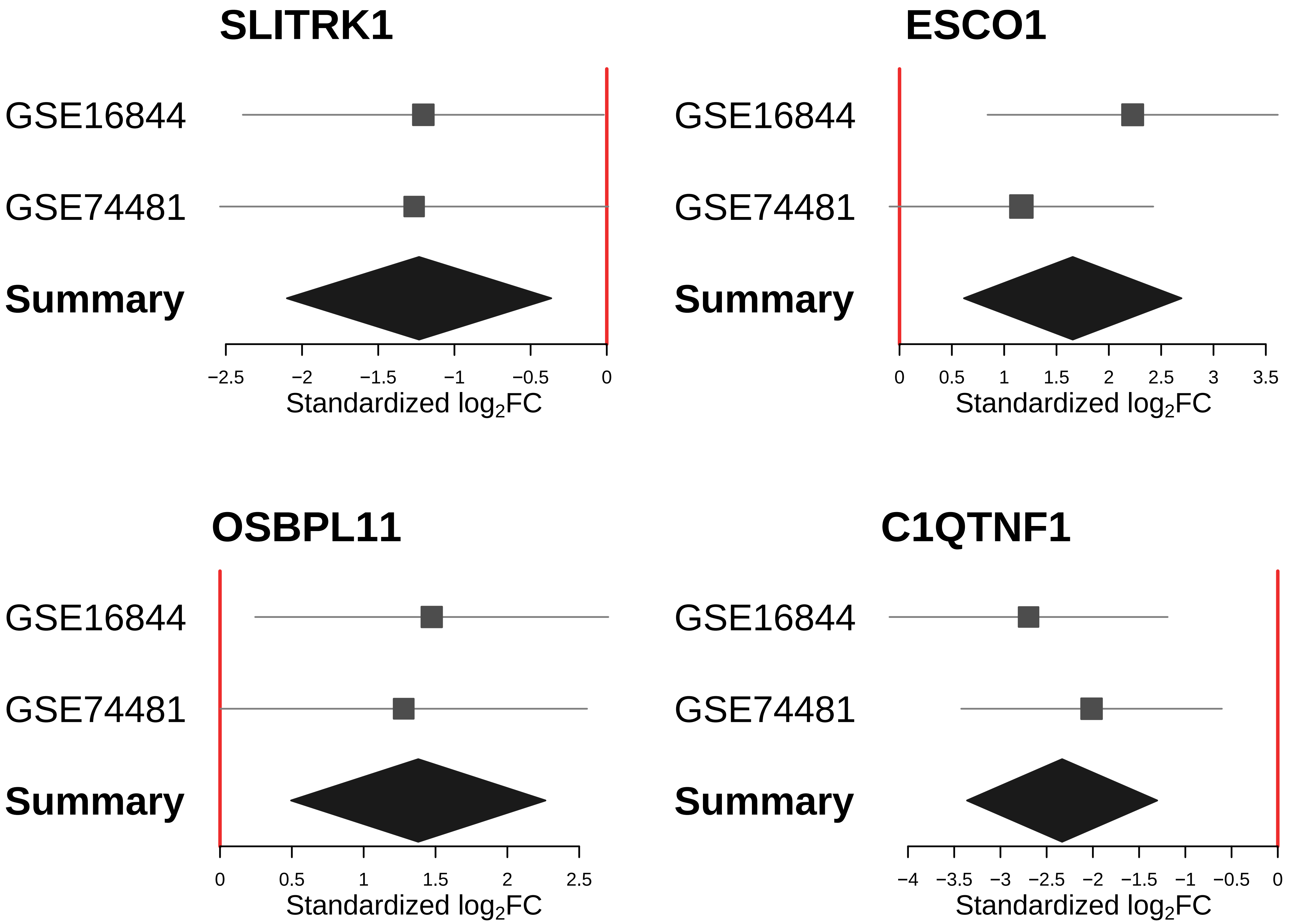
SLITRK1
GSE16844
GSE74481
Summary
−2.5 −2 −1.5 −1 −0.5 0
Standardized log2FC
ESCO1
GSE16844
GSE74481
Summary
0 0.5 1 1.5 2 2.5 3 3.5
Standardized log2FC
OSBPL11
GSE16844
GSE74481
Summary
0 0.5 1 1.5 2 2.5
Standardized log2FC
C1QTNF1
GSE16844
GSE74481
Summary
−4 −3.5 −3 −2.5 −2 −1.5 −1 −0.5 0
Standardized log2FC

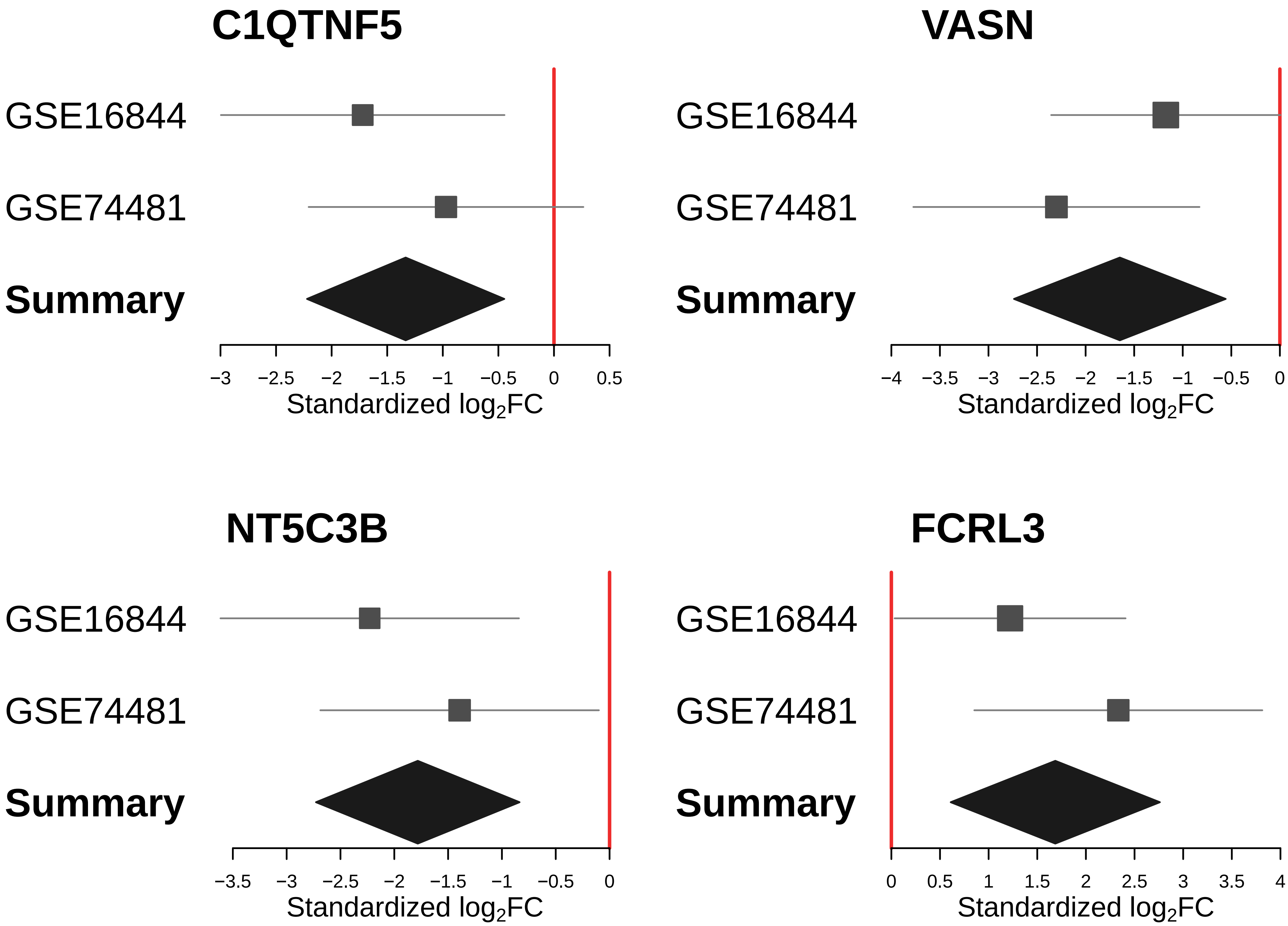

C1QTNF5
GSE16844
GSE74481
Summary
−3 −2.5 −2 −1.5 −1 −0.5 0 0.5
Standardized log2FC
VASN
GSE16844
GSE74481
Summary
−4 −3.5 −3 −2.5 −2 −1.5 −1 −0.5 0
Standardized log2FC
NT5C3B
GSE16844
GSE74481
Summary
−3.5 −3 −2.5 −2 −1.5 −1 −0.5 0
Standardized log2FC
FCRL3
GSE16844
GSE74481
Summary
0 0.5 1 1.5 2 2.5 3 3.5 4
Standardized log2FC

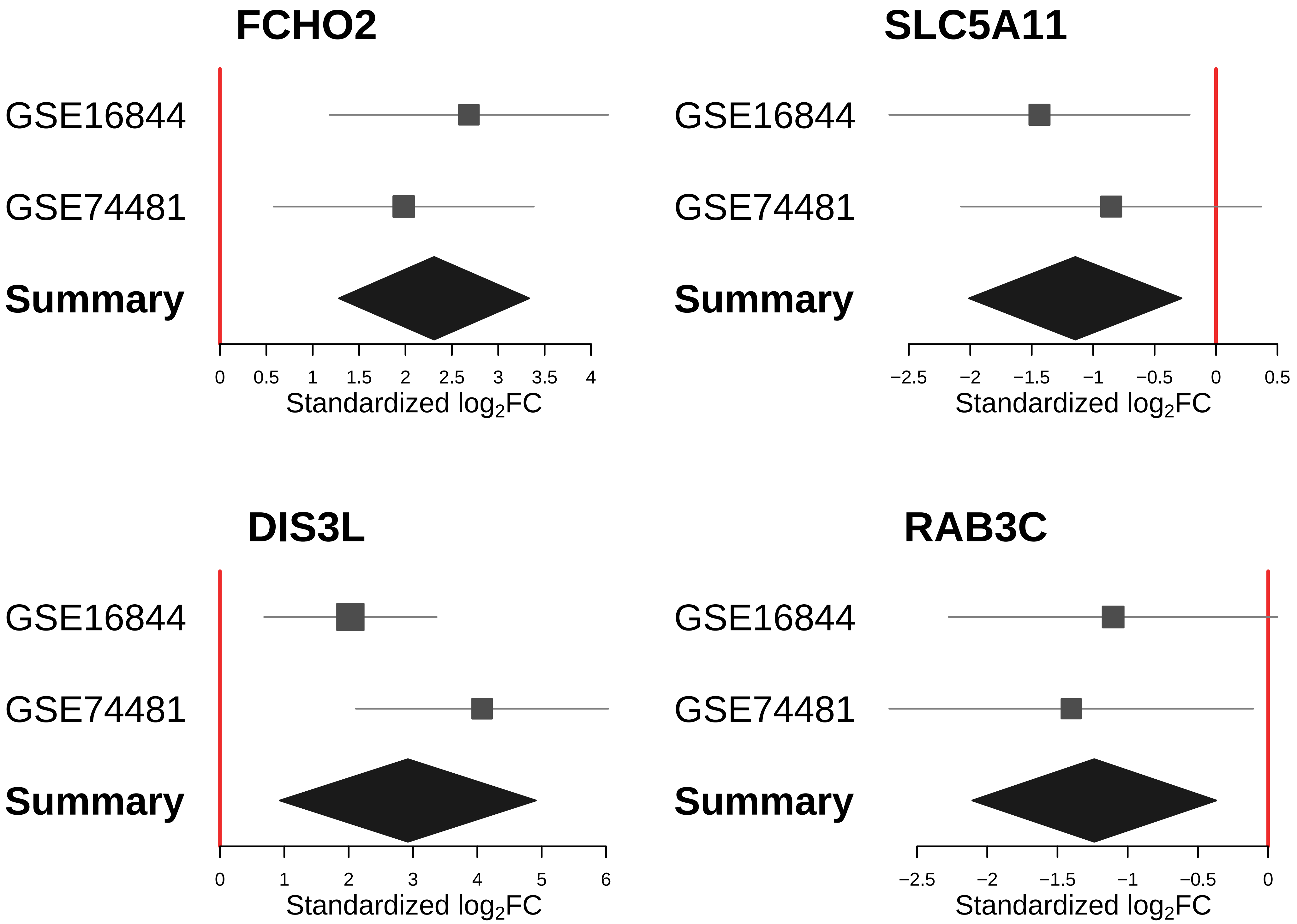
FCHO2
GSE16844
GSE74481
Summary
0 0.5 1 1.5 2 2.5 3 3.5 4
Standardized log2FC
SLC5A11
GSE16844
GSE74481
Summary
−2.5 −2 −1.5 −1 −0.5 0 0.5
Standardized log2FC
DIS3L
GSE16844
GSE74481
Summary
0 1 2 3 4 5 6
Standardized log2FC
RAB3C
GSE16844
GSE74481
Summary
−2.5 −2 −1.5 −1 −0.5 0
Standardized log2FC

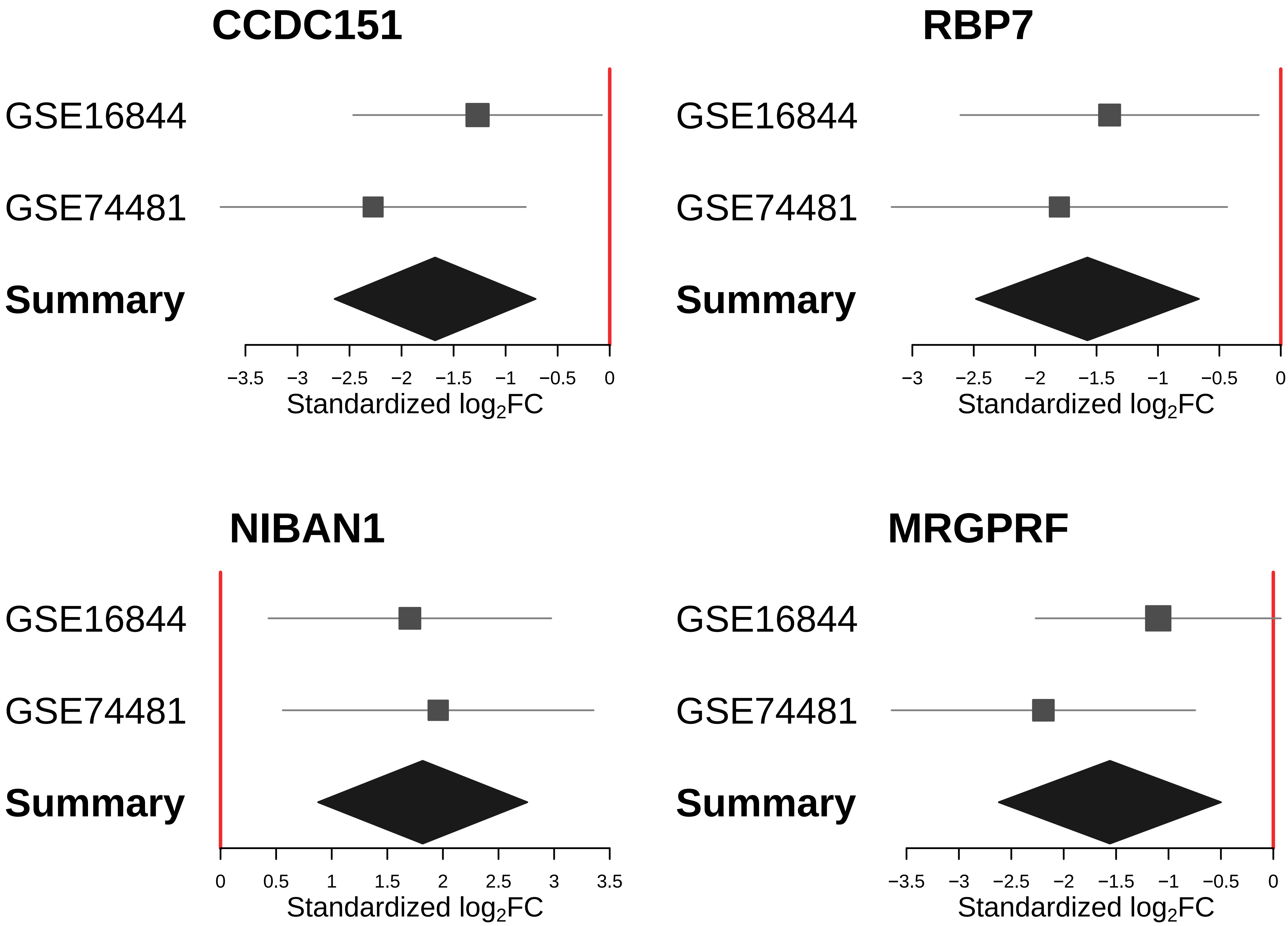
CCDC151
GSE16844
GSE74481
Summary
−3.5 −3 −2.5 −2 −1.5 −1 −0.5 0
Standardized log2FC
RBP7
GSE16844
GSE74481
Summary
−3 −2.5 −2 −1.5 −1 −0.5 0
Standardized log2FC
NIBAN1
GSE16844
GSE74481
Summary
0 0.5 1 1.5 2 2.5 3 3.5
Standardized log2FC
MRGPRF
GSE16844
GSE74481
Summary
−3.5 −3 −2.5 −2 −1.5 −1 −0.5 0
Standardized log2FC

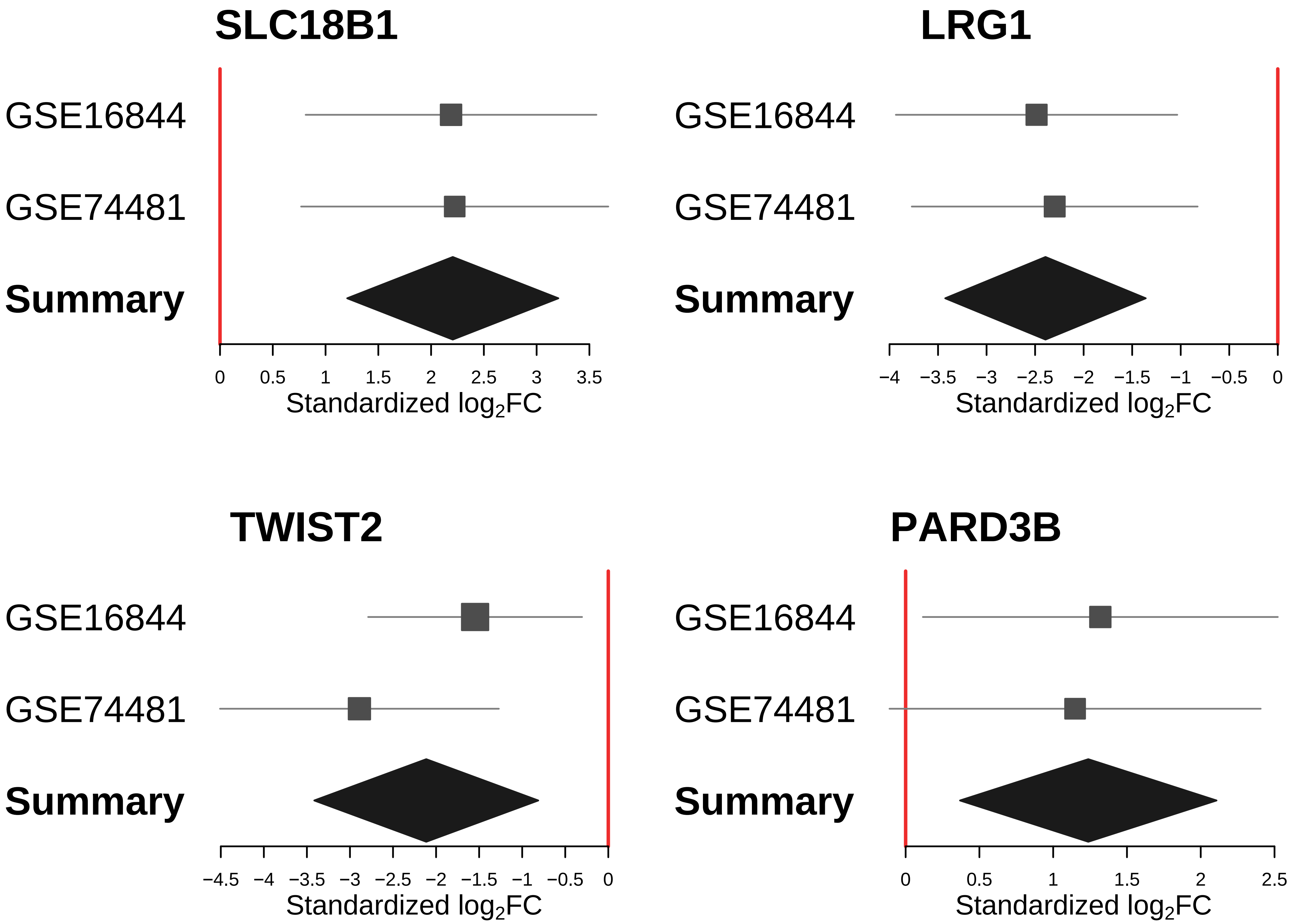
SLC18B1
GSE16844
GSE74481
Summary
0 0.5 1 1.5 2 2.5 3 3.5
Standardized log2FC
LRG1
GSE16844
GSE74481
Summary
−4 −3.5 −3 −2.5 −2 −1.5 −1 −0.5 0
Standardized log2FC
TWIST2
GSE16844
GSE74481
Summary
−4.5 −4 −3.5 −3 −2.5 −2 −1.5 −1 −0.5 0
Standardized log2FC
PARD3B
GSE16844
GSE74481
Summary
0 0.5 1 1.5 2 2.5
Standardized log2FC

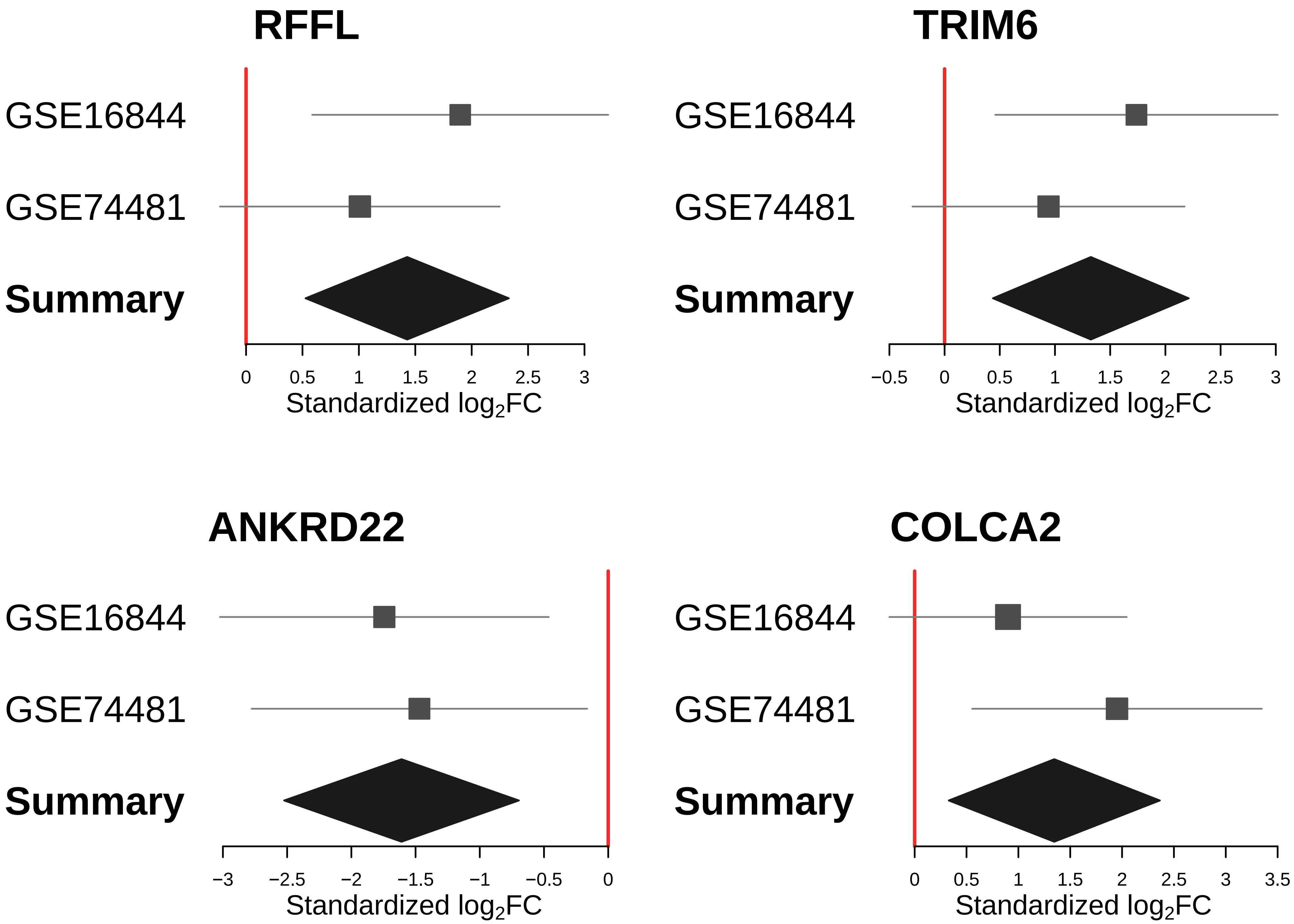
RFFL
GSE16844
GSE74481
Summary
0 0.5 1 1.5 2 2.5 3
Standardized log2FC
TRIM6
GSE16844
GSE74481
Summary
−0.5 0 0.5 1 1.5 2 2.5 3
Standardized log2FC
ANKRD22
GSE16844
GSE74481
Summary
−3 −2.5 −2 −1.5 −1 −0.5 0
Standardized log2FC
COLCA2
GSE16844
GSE74481
Summary
0 0.5 1 1.5 2 2.5 3 3.5
Standardized log2FC

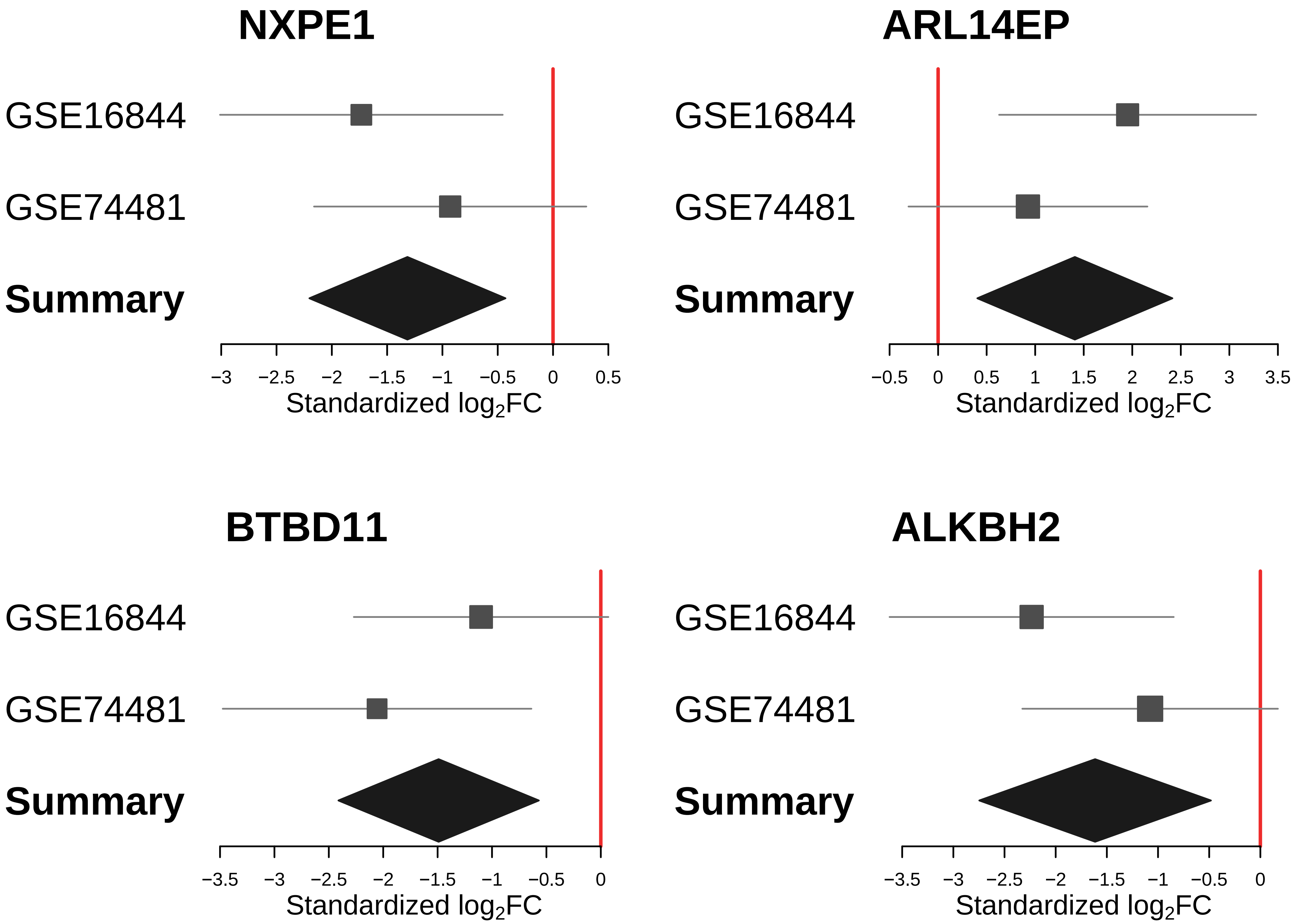
NXPE1
GSE16844
GSE74481
Summary
−3 −2.5 −2 −1.5 −1 −0.5 0 0.5
Standardized log2FC
ARL14EP
GSE16844
GSE74481
Summary
−0.5 0 0.5 1 1.5 2 2.5 3 3.5
Standardized log2FC
BTBD11
GSE16844
GSE74481
Summary
−3.5 −3 −2.5 −2 −1.5 −1 −0.5 0
Standardized log2FC
ALKBH2
GSE16844
GSE74481
Summary
−3.5 −3 −2.5 −2 −1.5 −1 −0.5 0
Standardized log2FC

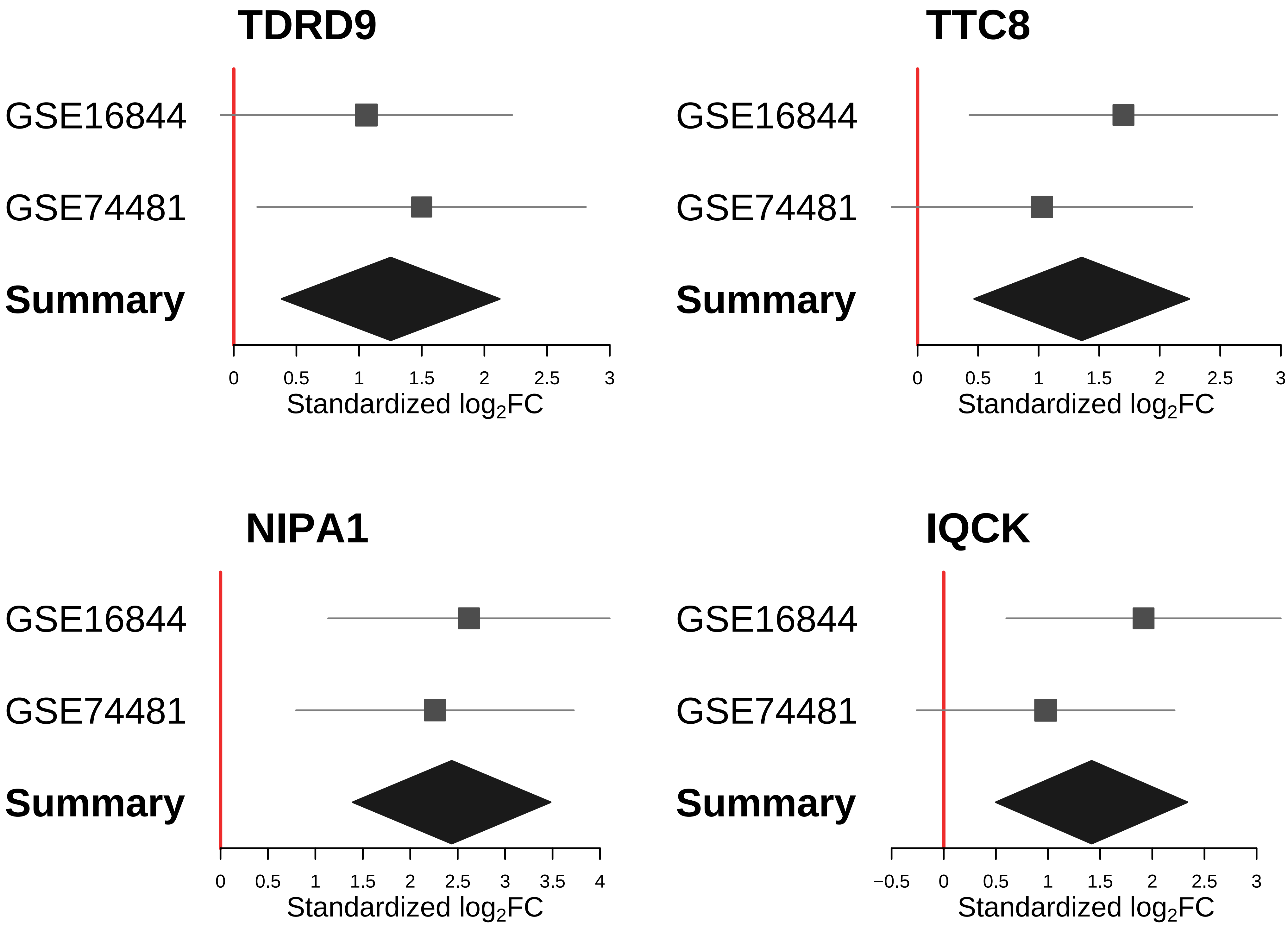

TDRD9
GSE16844
GSE74481
Summary
0 0.5 1 1.5 2 2.5 3
Standardized log2FC
TTC8
GSE16844
GSE74481
Summary
0 0.5 1 1.5 2 2.5 3
Standardized log2FC
NIPA1
GSE16844
GSE74481
Summary
0 0.5 1 1.5 2 2.5 3 3.5 4
Standardized log2FC
IQCK
GSE16844
GSE74481
Summary
−0.5 0 0.5 1 1.5 2 2.5 3
Standardized log2FC

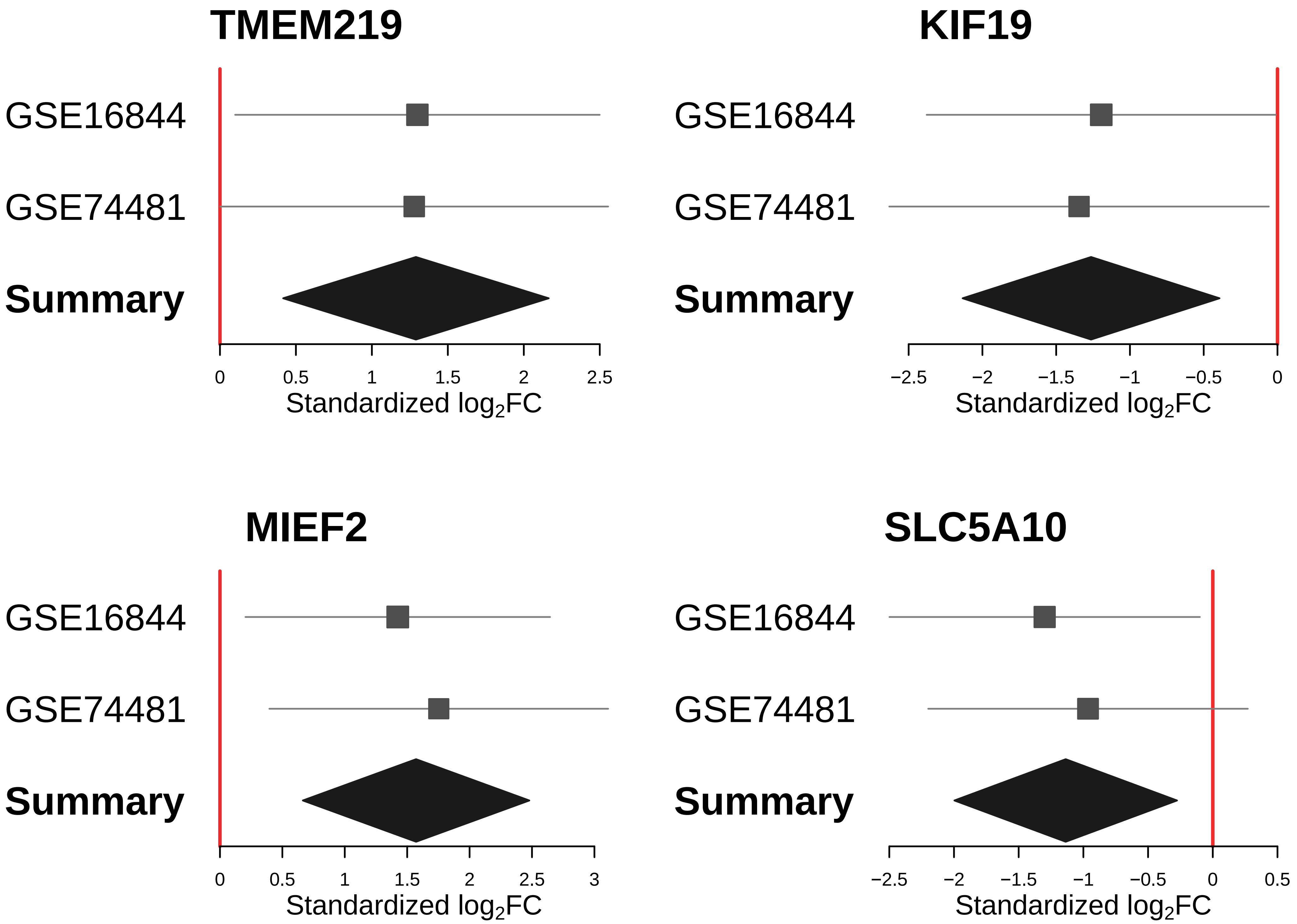
TMEM219
GSE16844
GSE74481
Summary
0
0.5
1
1.5
2
2.5
Standardized log2FC
KIF19
GSE16844
GSE74481
Summary
−2.5
−2
−1.5
−1
−0.5
0
Standardized log2FC
MIEF2
GSE16844
GSE74481
Summary
0
0.5
1
1.5
2
2.5
3
Standardized log2FC
SLC5A10
GSE16844
GSE74481
Summary
−2.5
−2
−1.5
−1
−0.5
0
0.5
Standardized log2FC

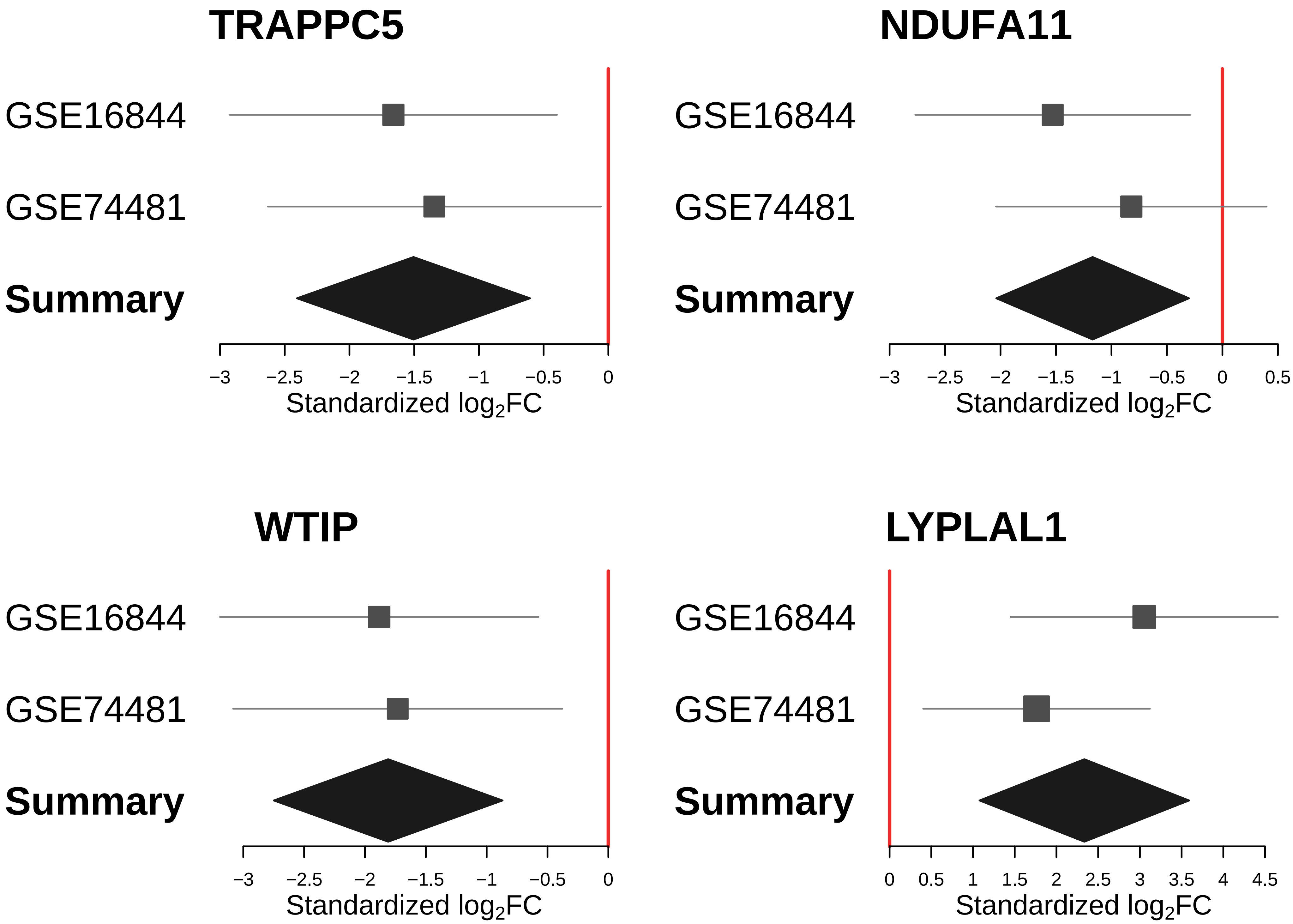
TRAPPC5
GSE16844
GSE74481
Summary
−3 −2.5 −2 −1.5 −1 −0.5 0
Standardized log2FC
NDUFA11
GSE16844
GSE74481
Summary
−3 −2.5 −2 −1.5 −1 −0.5 0 0.5
Standardized log2FC
WTIP
GSE16844
GSE74481
Summary
−3 −2.5 −2 −1.5 −1 −0.5 0
Standardized log2FC
LYPLAL1
GSE16844
GSE74481
Summary
0 0.5 1 1.5 2 2.5 3 3.5 4 4.5
Standardized log2FC

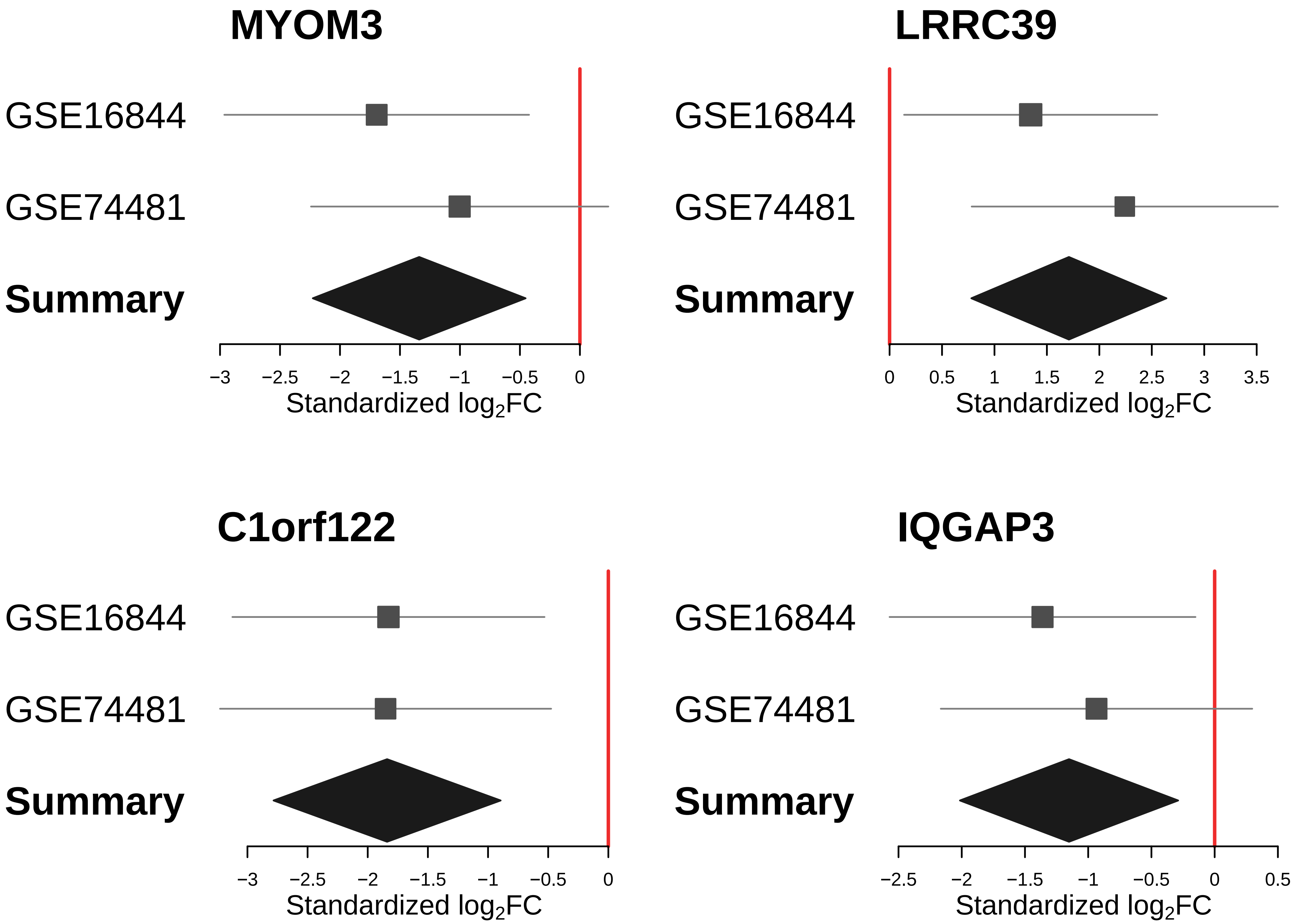
MYOM3
GSE16844
GSE74481
Summary
−3 −2.5 −2 −1.5 −1 −0.5 0
Standardized log2FC
LRRC39
GSE16844
GSE74481
Summary
0 0.5 1 1.5 2 2.5 3 3.5
Standardized log2FC
C1orf122
GSE16844
GSE74481
Summary
−3 −2.5 −2 −1.5 −1 −0.5 0
Standardized log2FC
IQGAP3
GSE16844
GSE74481
Summary
−2.5 −2 −1.5 −1 −0.5 0 0.5
Standardized log2FC

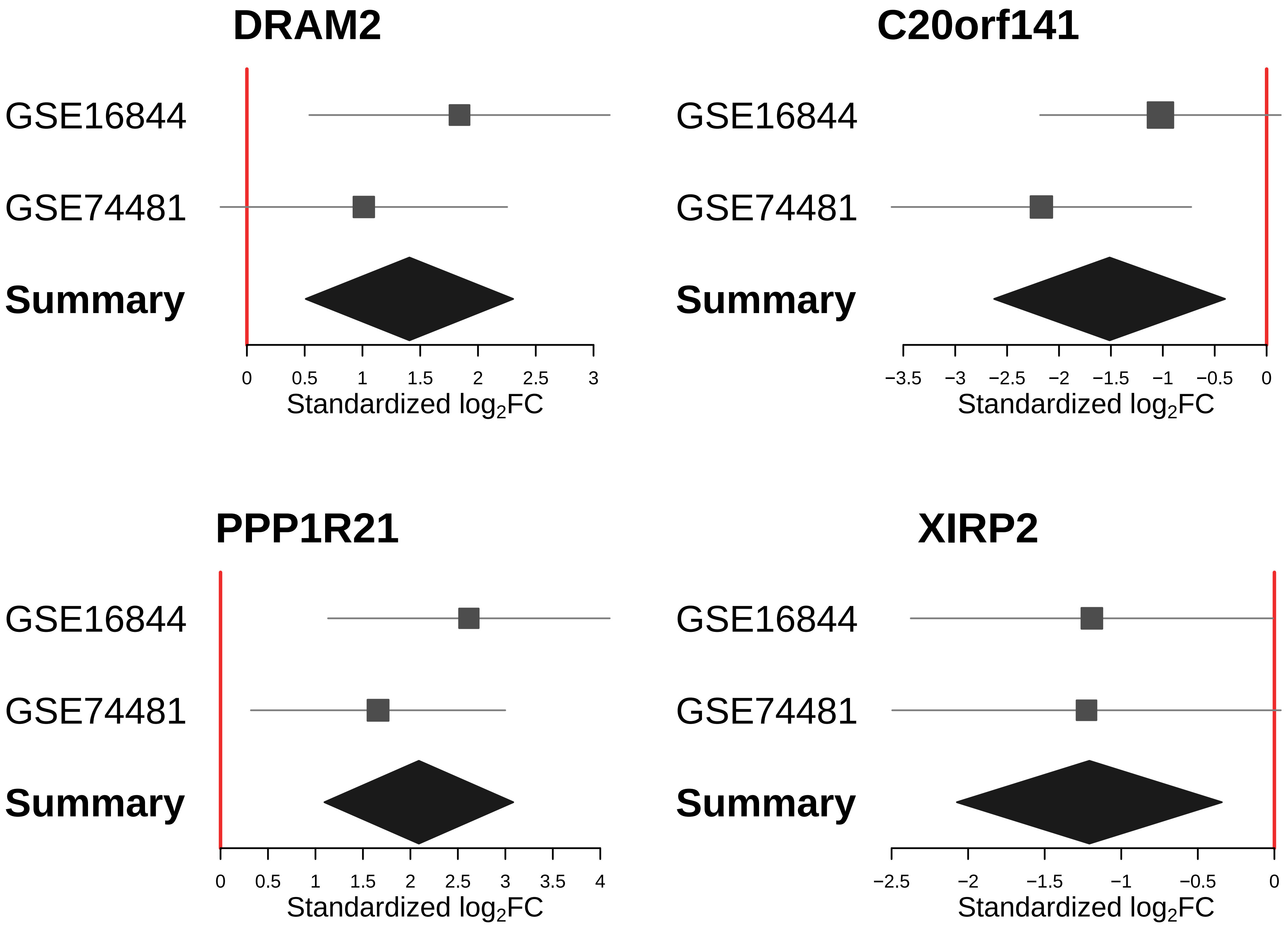

DRAM2
GSE16844
GSE74481
Summary
0 0.5 1 1.5 2 2.5 3
Standardized log2FC
C20orf141
GSE16844
GSE74481
Summary
−3.5 −3 −2.5 −2 −1.5 −1 −0.5 0
Standardized log2FC
PPP1R21
GSE16844
GSE74481
Summary
0 0.5 1 1.5 2 2.5 3 3.5 4
Standardized log2FC
XIRP2
GSE16844
GSE74481
Summary
−2.5 −2 −1.5 −1 −0.5 0
Standardized log2FC

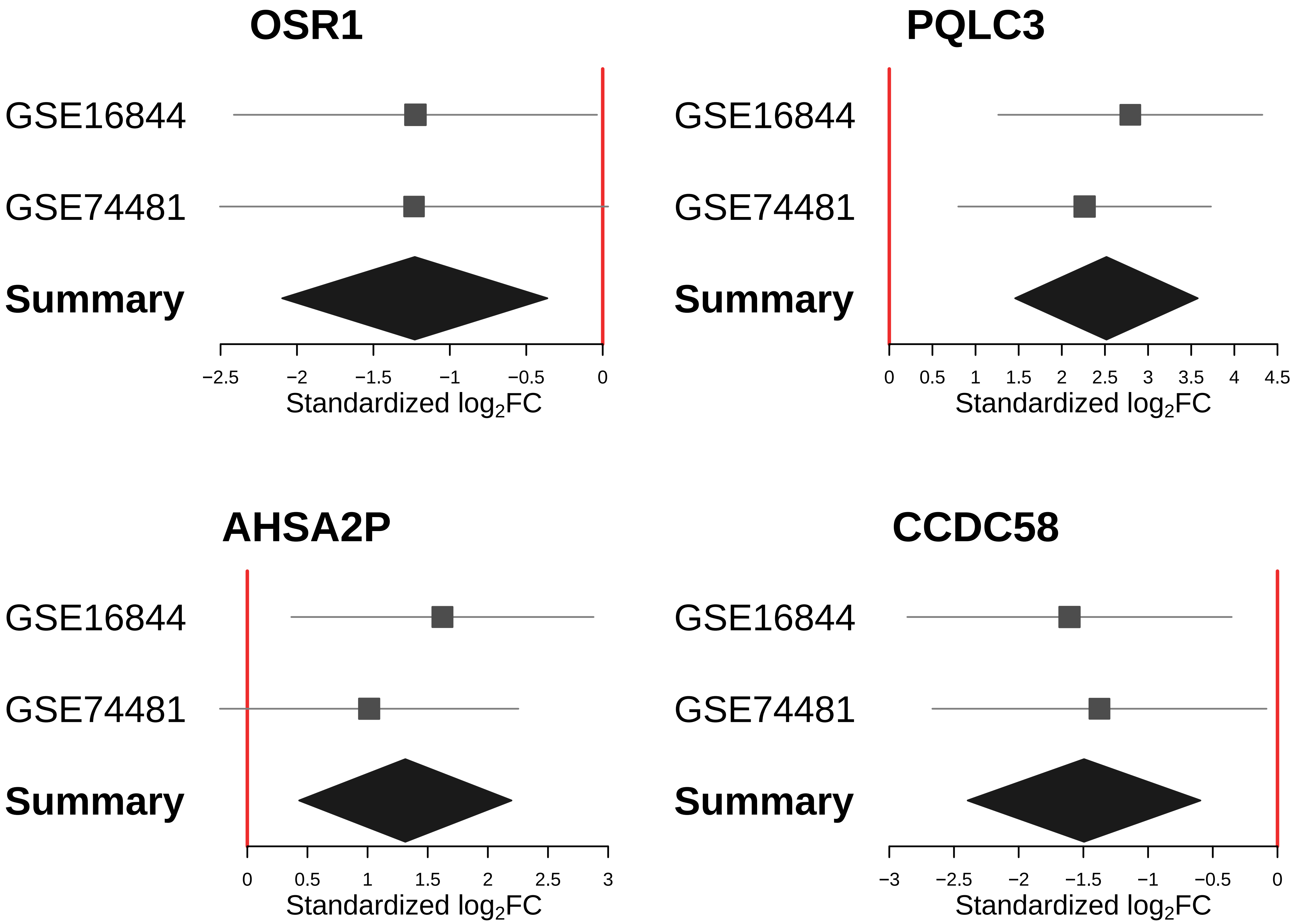

OSR1
GSE16844
GSE74481
Summary
−2.5
−2
−1.5
−1
−0.5
0
Standardized log2FC
PQLC3
GSE16844
GSE74481
Summary
0
0.5
1
1.5
2
2.5
3
3.5
4
4.5
Standardized log2FC
AHSA2P
GSE16844
GSE74481
Summary
0
0.5
1
1.5
2
2.5
3
Standardized log2FC
CCDC58
GSE16844
GSE74481
Summary
−3
−2.5
−2
−1.5
−1
−0.5
0
Standardized log2FC

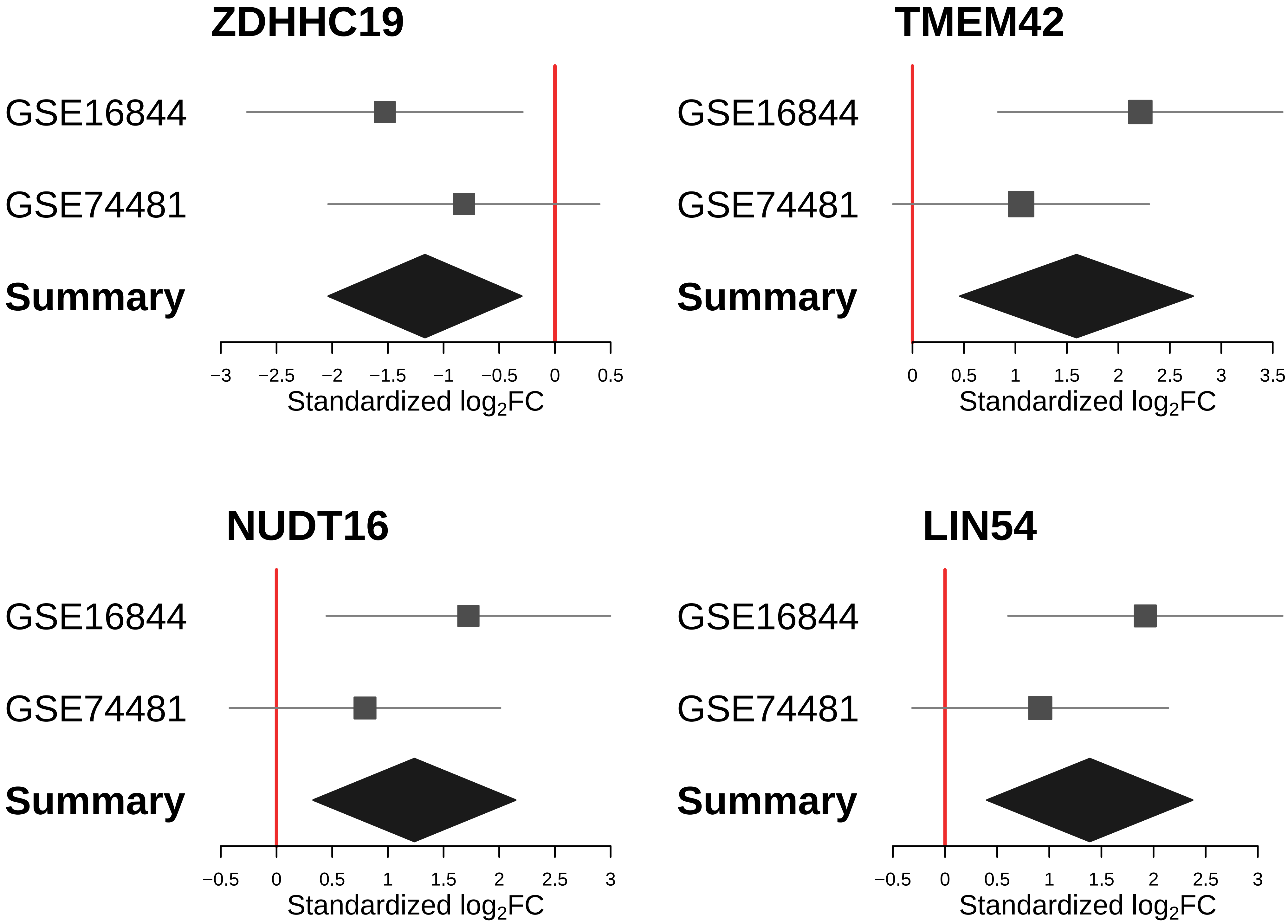
ZDHHC19
GSE16844
GSE74481
Summary
−3
−2.5
−2
−1.5
−1
−0.5
0
0.5
Standardized $log_2FC$
TMEM42
GSE16844
GSE74481
Summary
0
0.5
1
1.5
2
2.5
3
3.5
Standardized $log_2FC$
NUDT16
GSE16844
GSE74481
Summary
−0.5
0
0.5
1
1.5
2
2.5
3
Standardized $log_2FC$
LIN54
GSE16844
GSE74481
Summary
−0.5
0
0.5
1
1.5
2
2.5
3
Standardized $log_2FC$

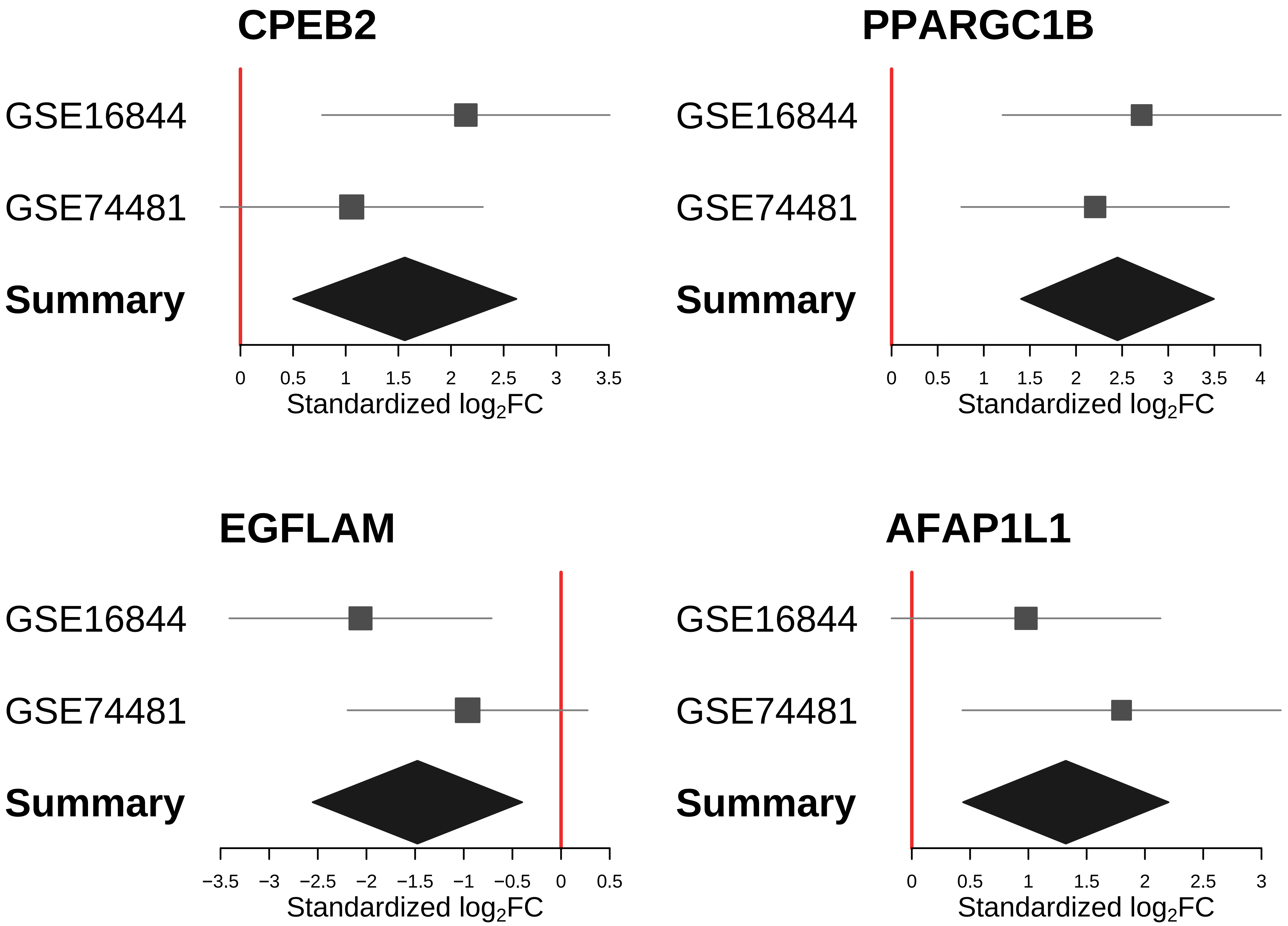
CPEB2
GSE16844
GSE74481
Summary
0
0.5
1
1.5
2
2.5
3
3.5
Standardized log2FC
PPARGC1B
GSE16844
GSE74481
Summary
0
0.5
1
1.5
2
2.5
3
3.5
4
Standardized log2FC
EGFLAM
GSE16844
GSE74481
Summary
−3.5
−3
−2.5
−2
−1.5
−1
−0.5
0
0.5
Standardized log2FC
AFAP1L1
GSE16844
GSE74481
Summary
0
0.5
1
1.5
2
2.5
3
Standardized log2FC

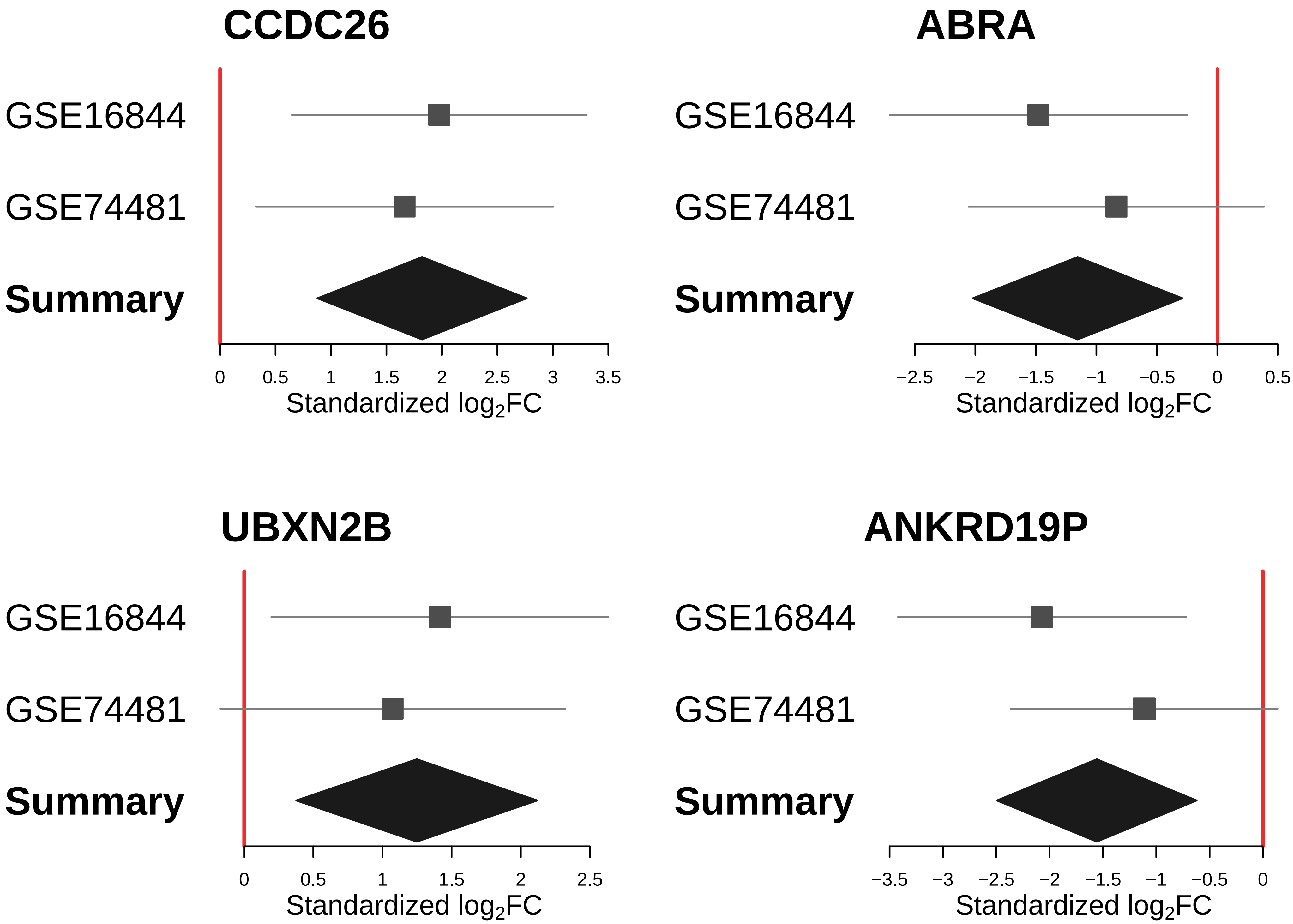
CCDC26
GSE16844
GSE74481
Summary
0
0.5
1
1.5
2
2.5
3
3.5
Standardized log2FC
ABRA
GSE16844
GSE74481
Summary
−2.5
−2
−1.5
−1
−0.5
0
0.5
Standardized log2FC
UBXN2B
GSE16844
GSE74481
Summary
0
0.5
1
1.5
2
2.5
Standardized log2FC
ANKRD19P
GSE16844
GSE74481
Summary
−3.5
−3
−2.5
−2
−1.5
−1
−0.5
0
Standardized log2FC

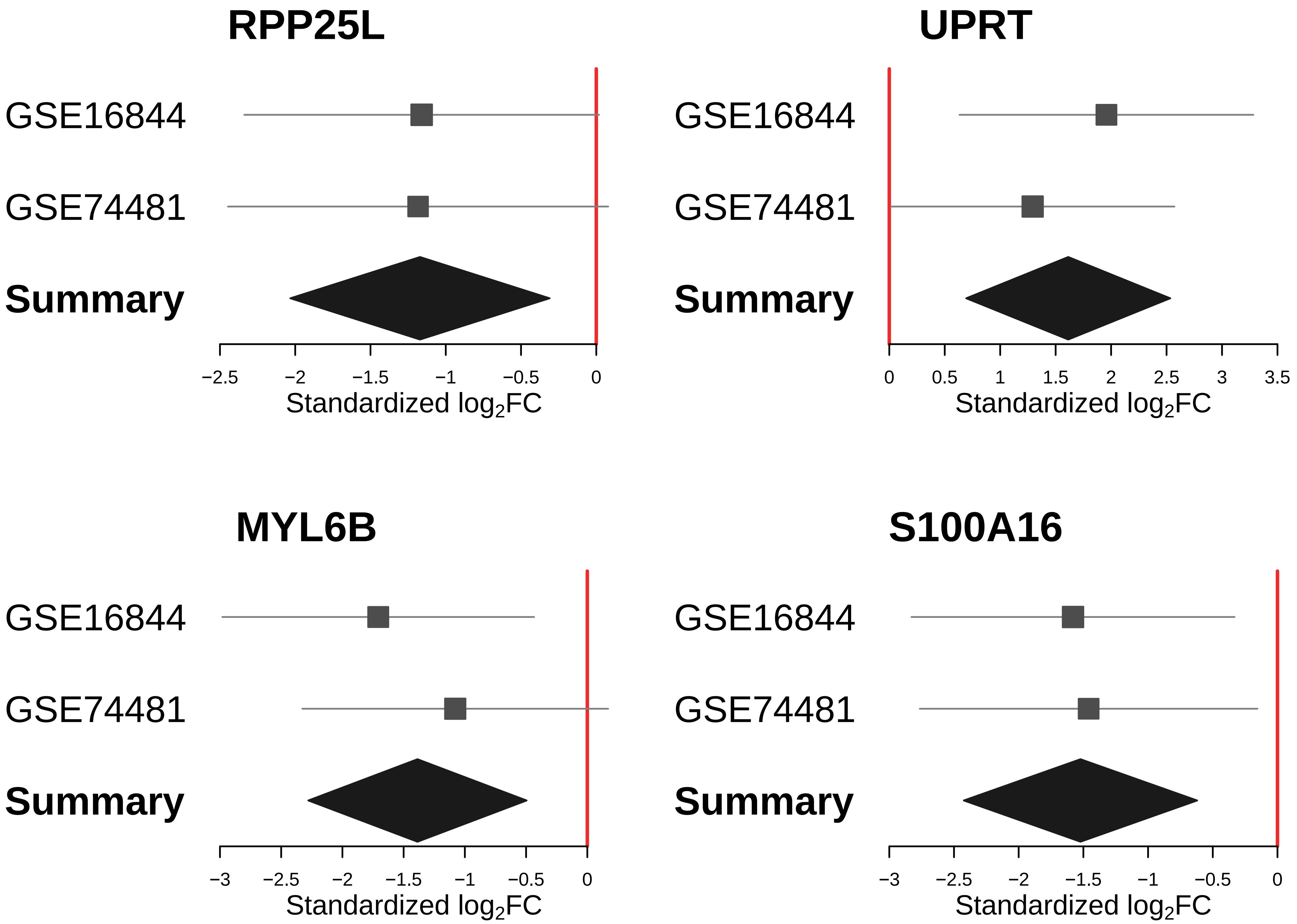
RPP25L
GSE16844
GSE74481
Summary
−2.5
−2
−1.5
−1
−0.5
0
Standardized log2FC
UPRT
GSE16844
GSE74481
Summary
0
0.5
1
1.5
2
2.5
3
3.5
Standardized log2FC
MYL6B
GSE16844
GSE74481
Summary
−3
−2.5
−2
−1.5
−1
−0.5
0
Standardized log2FC
S100A16
GSE16844
GSE74481
Summary
−3
−2.5
−2
−1.5
−1
−0.5
0
Standardized log2FC

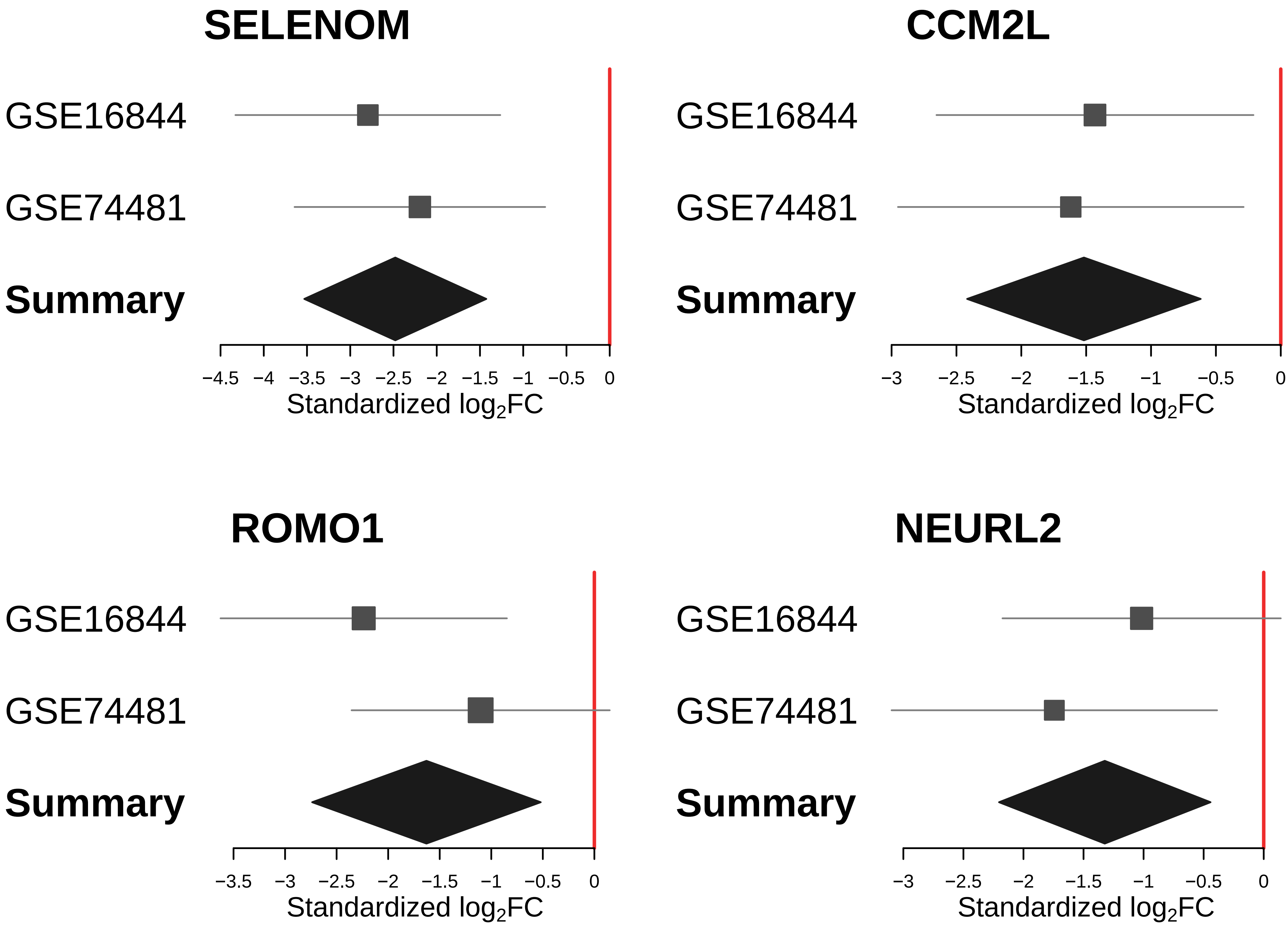
SELENOM
GSE16844
GSE74481
Summary
−4.5 −4 −3.5 −3 −2.5 −2 −1.5 −1 −0.5 0
Standardized $log_2FC$
CCM2L
GSE16844
GSE74481
Summary
−3 −2.5 −2 −1.5 −1 −0.5 0
Standardized $log_2FC$
ROMO1
GSE16844
GSE74481
Summary
−3.5 −3 −2.5 −2 −1.5 −1 −0.5 0
Standardized $log_2FC$
NEURL2
GSE16844
GSE74481
Summary
−3 −2.5 −2 −1.5 −1 −0.5 0
Standardized $log_2FC$

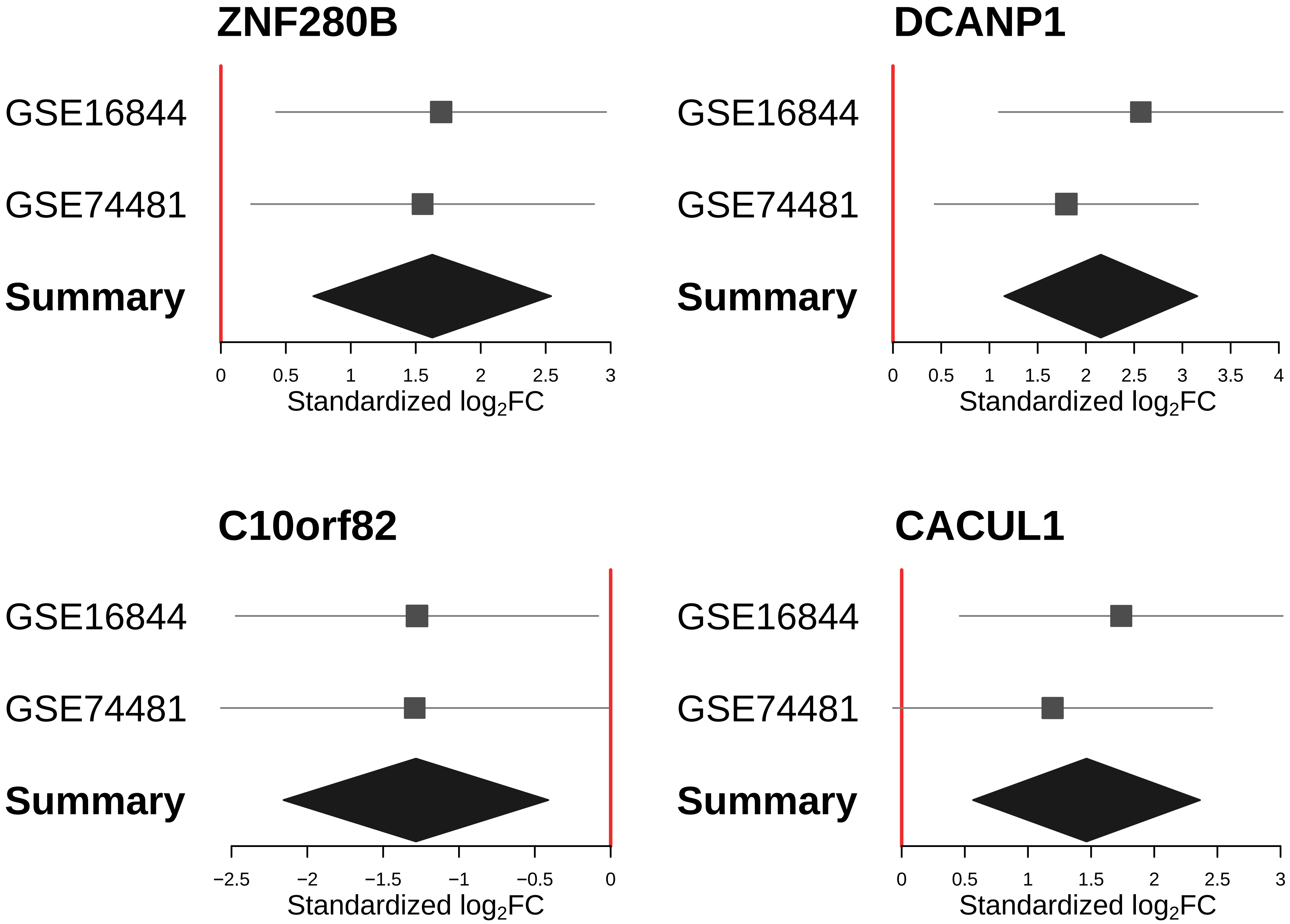
ZNF280B
GSE16844
GSE74481
Summary
0 0.5 1 1.5 2 2.5 3
Standardized log2FC
DCANP1
GSE16844
GSE74481
Summary
0 0.5 1 1.5 2 2.5 3 3.5 4
Standardized log2FC
C10orf82
GSE16844
GSE74481
Summary
−2.5 −2 −1.5 −1 −0.5 0
Standardized log2FC
CACUL1
GSE16844
GSE74481
Summary
0 0.5 1 1.5 2 2.5 3
Standardized log2FC

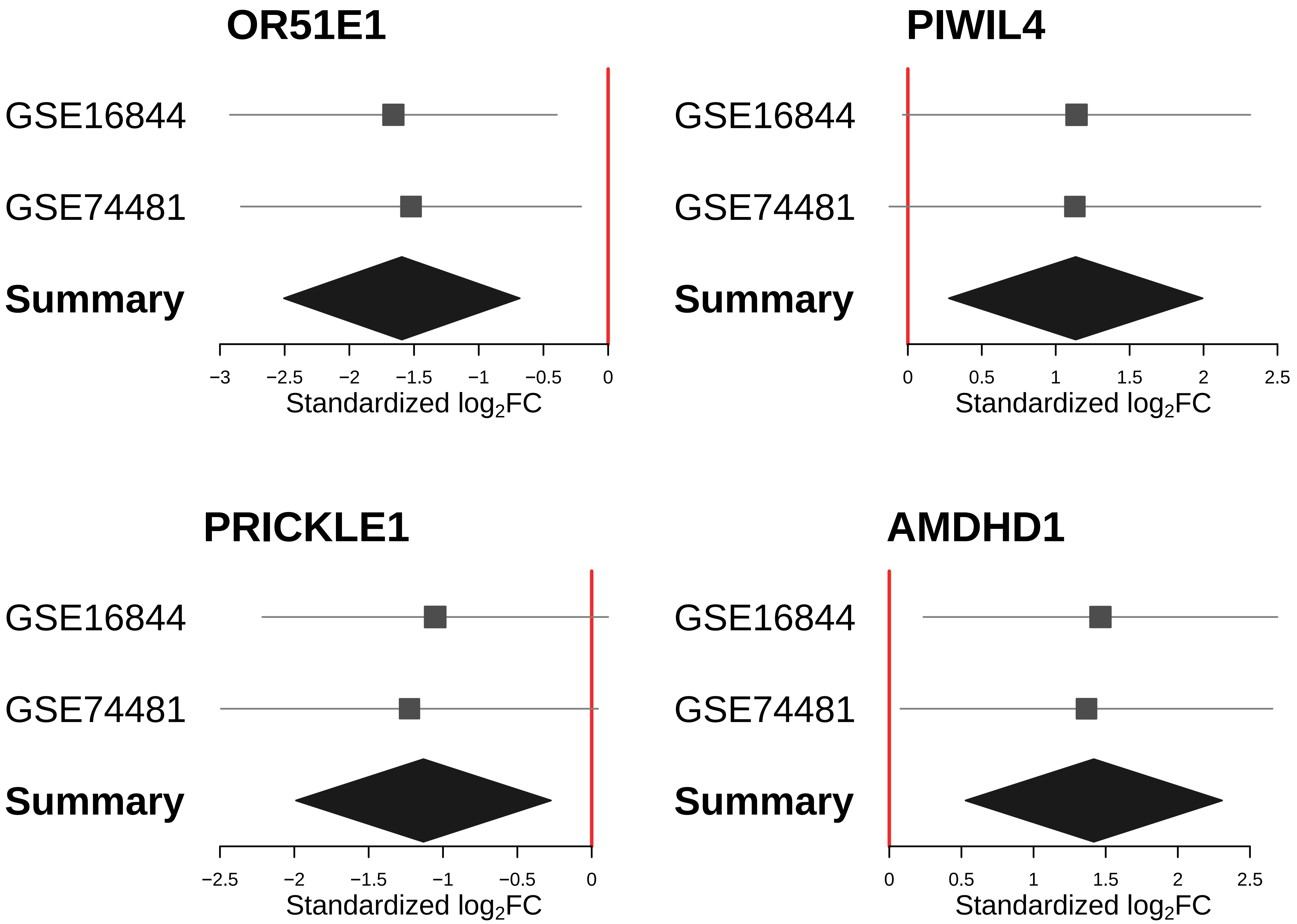
OR51E1
GSE16844
GSE74481
Summary
−3
−2.5
−2
−1.5
−1
−0.5
0
Standardized log2FC
PIWIL4
GSE16844
GSE74481
Summary
0
0.5
1
1.5
2
2.5
Standardized log2FC
PRICKLE1
GSE16844
GSE74481
Summary
−2.5
−2
−1.5
−1
−0.5
0
Standardized log2FC
AMDHD1
GSE16844
GSE74481
Summary
0
0.5
1
1.5
2
2.5
Standardized log2FC

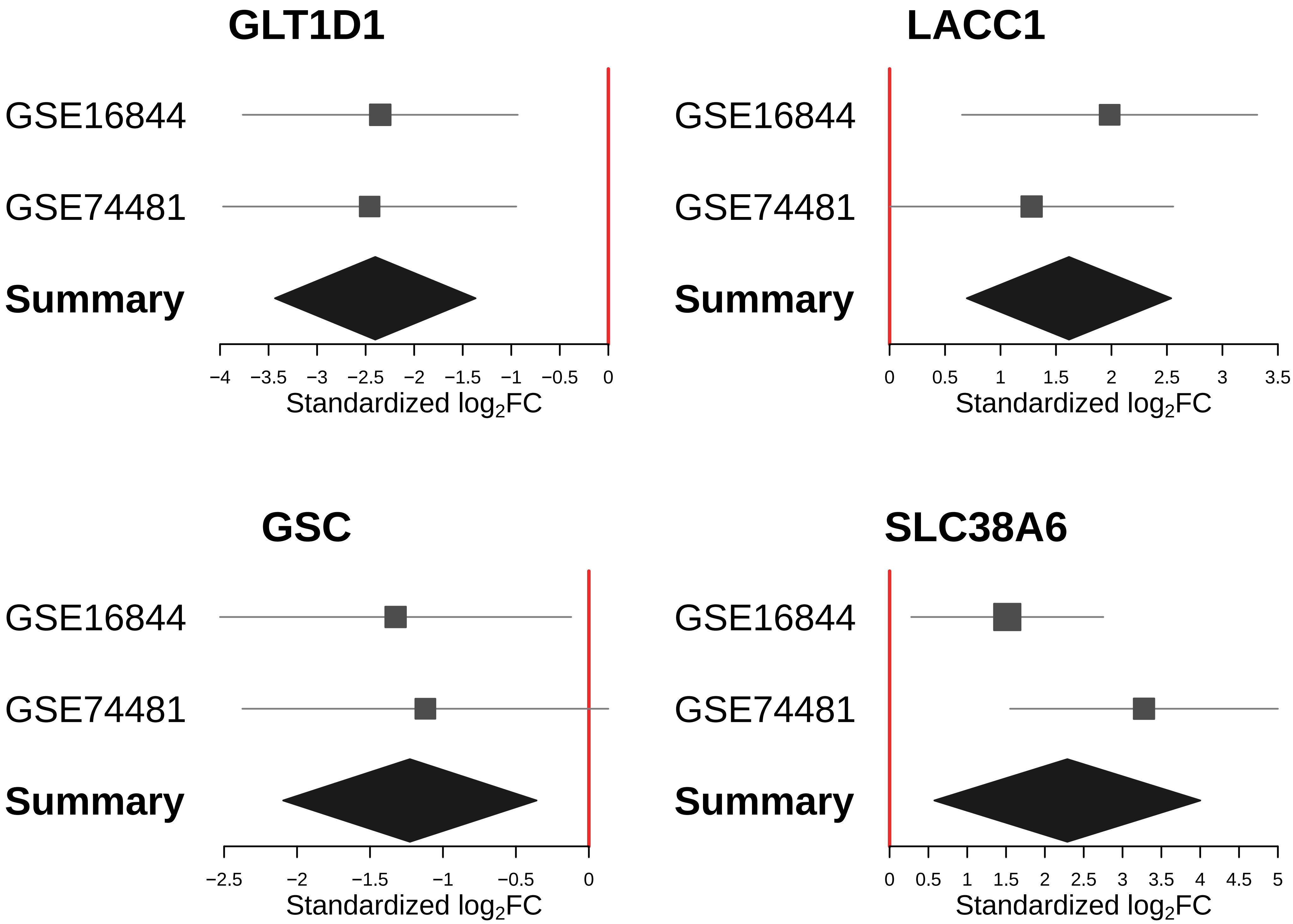
GLT1D1
GSE16844
GSE74481
Summary
−4 −3.5 −3 −2.5 −2 −1.5 −1 −0.5 0
Standardized log2FC
LACC1
GSE16844
GSE74481
Summary
0 0.5 1 1.5 2 2.5 3 3.5
Standardized log2FC
GSC
GSE16844
GSE74481
Summary
−2.5 −2 −1.5 −1 −0.5 0
Standardized log2FC
SLC38A6
GSE16844
GSE74481
Summary
0 0.5 1 1.5 2 2.5 3 3.5 4 4.5 5
Standardized log2FC

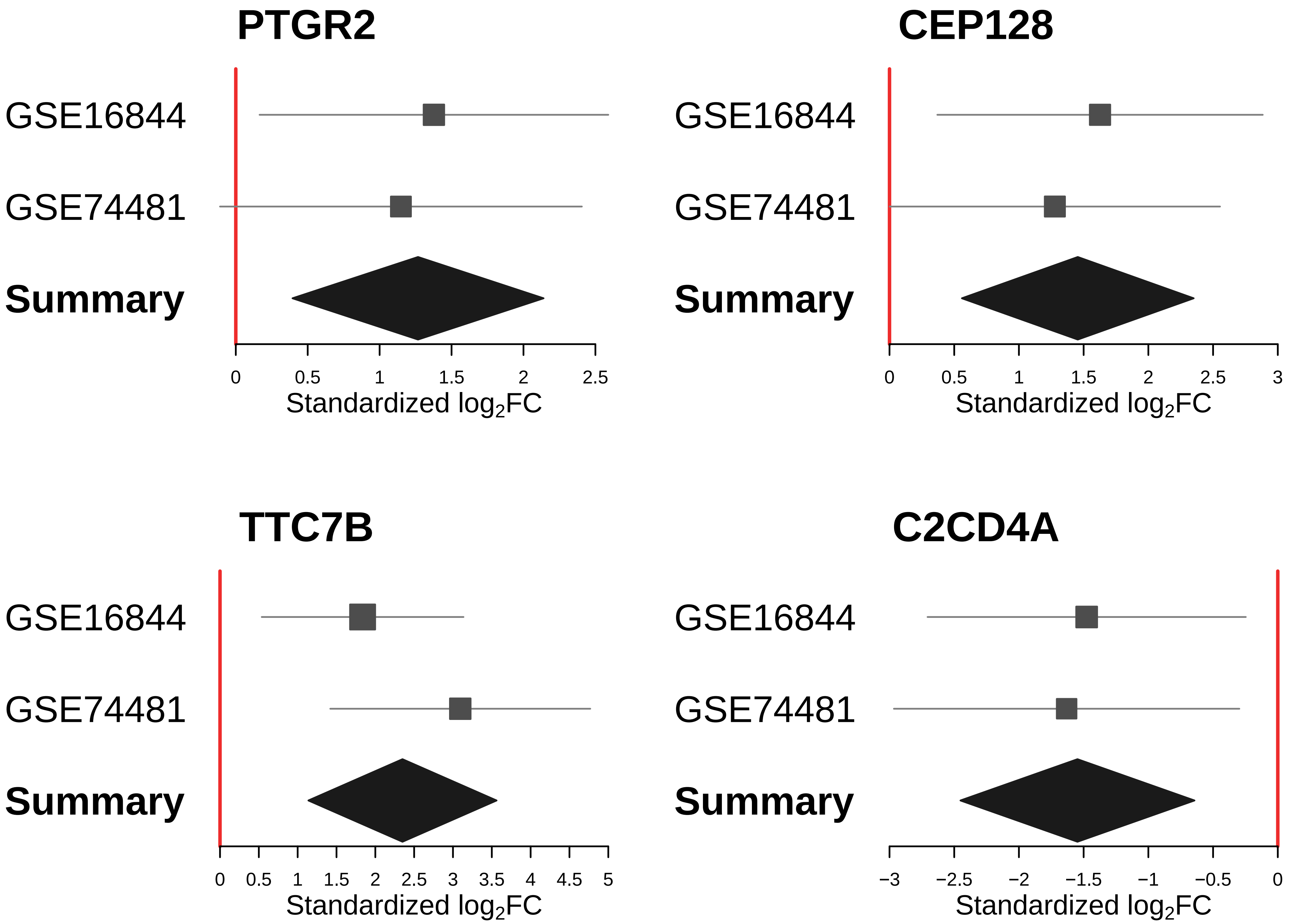
PTGR2
GSE16844
GSE74481
Summary
0 0.5 1 1.5 2 2.5
Standardized log2FC
CEP128
GSE16844
GSE74481
Summary
0 0.5 1 1.5 2 2.5 3
Standardized log2FC
TTC7B
GSE16844
GSE74481
Summary
0 0.5 1 1.5 2 2.5 3 3.5 4 4.5 5
Standardized log2FC
C2CD4A
GSE16844
GSE74481
Summary
−3 −2.5 −2 −1.5 −1 −0.5 0
Standardized log2FC

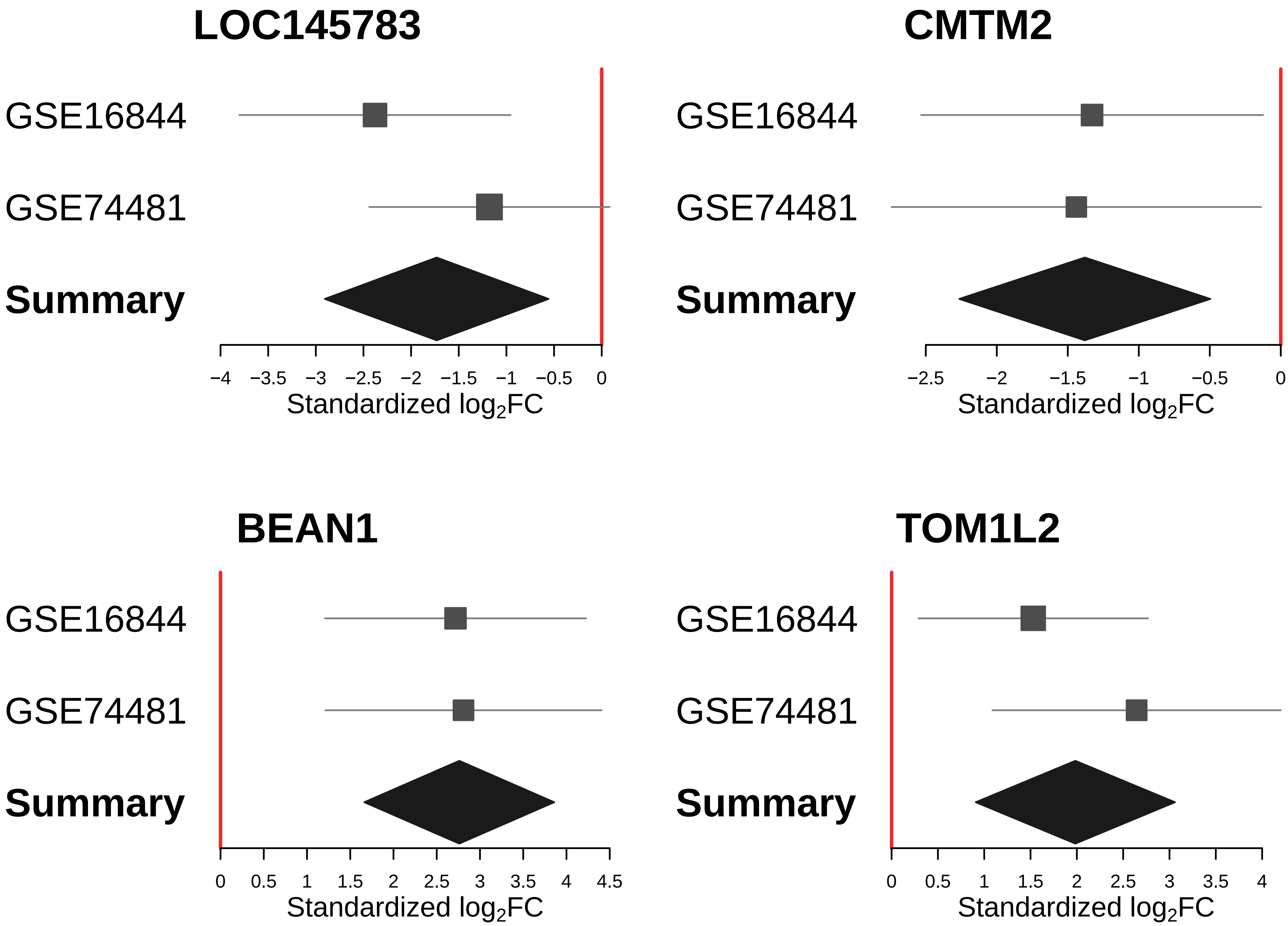
LOC145783
GSE16844
GSE74481
Summary
−4 −3.5 −3 −2.5 −2 −1.5 −1 −0.5 0
Standardized log2FC
CMTM2
GSE16844
GSE74481
Summary
−2.5 −2 −1.5 −1 −0.5 0
Standardized log2FC
BEAN1
GSE16844
GSE74481
Summary
0 0.5 1 1.5 2 2.5 3 3.5 4 4.5
Standardized log2FC
TOM1L2
GSE16844
GSE74481
Summary
0 0.5 1 1.5 2 2.5 3 3.5 4
Standardized log2FC

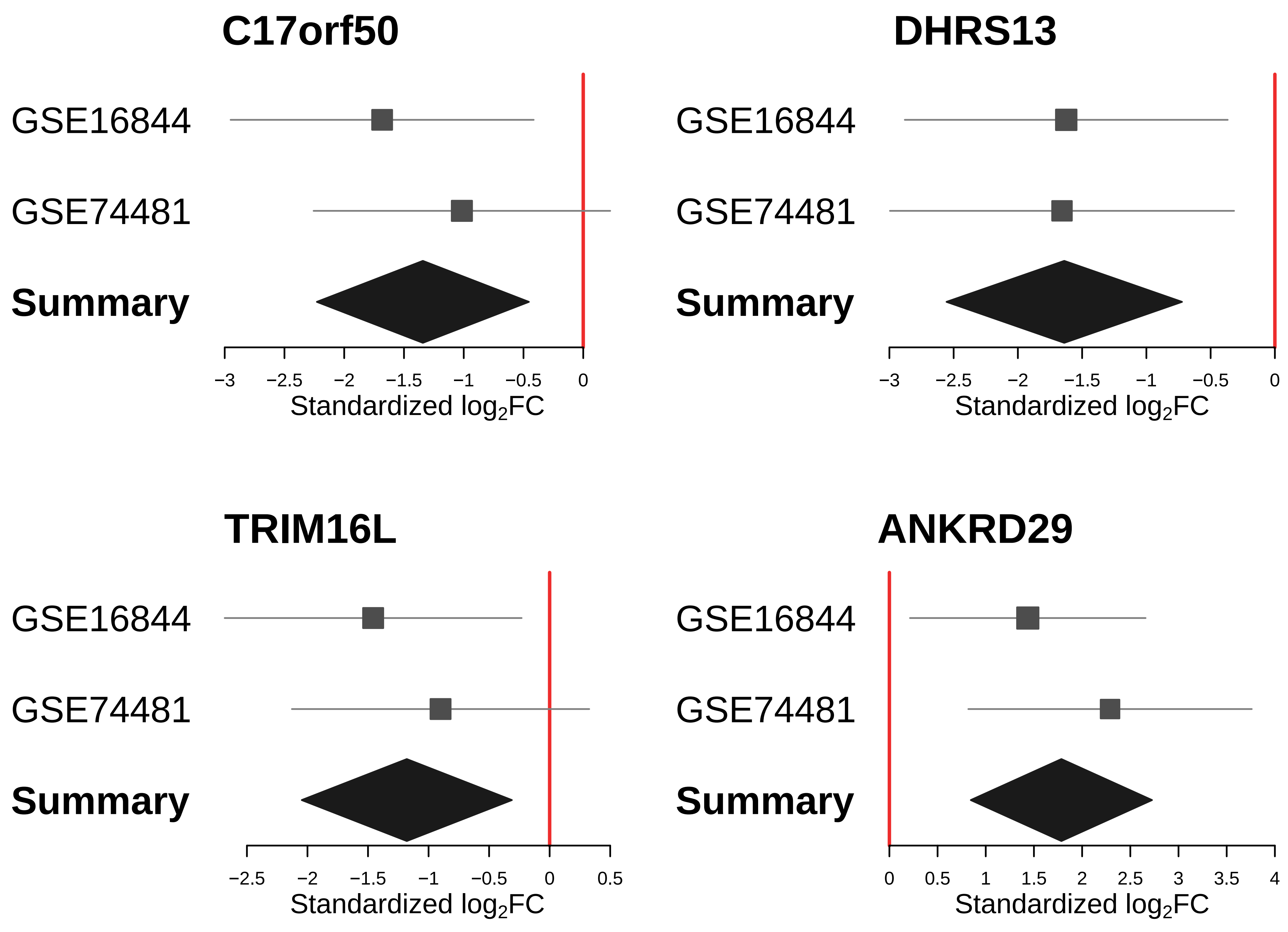

C17orf50
DHRS13
TRIM16L
ANKRD29
GSE16844
GSE74481
Summary
−3
−2.5
−2
−1.5
−1
−0.5
0
0.5
1
1.5
2
2.5
3
3.5
4
Standardized log2FC

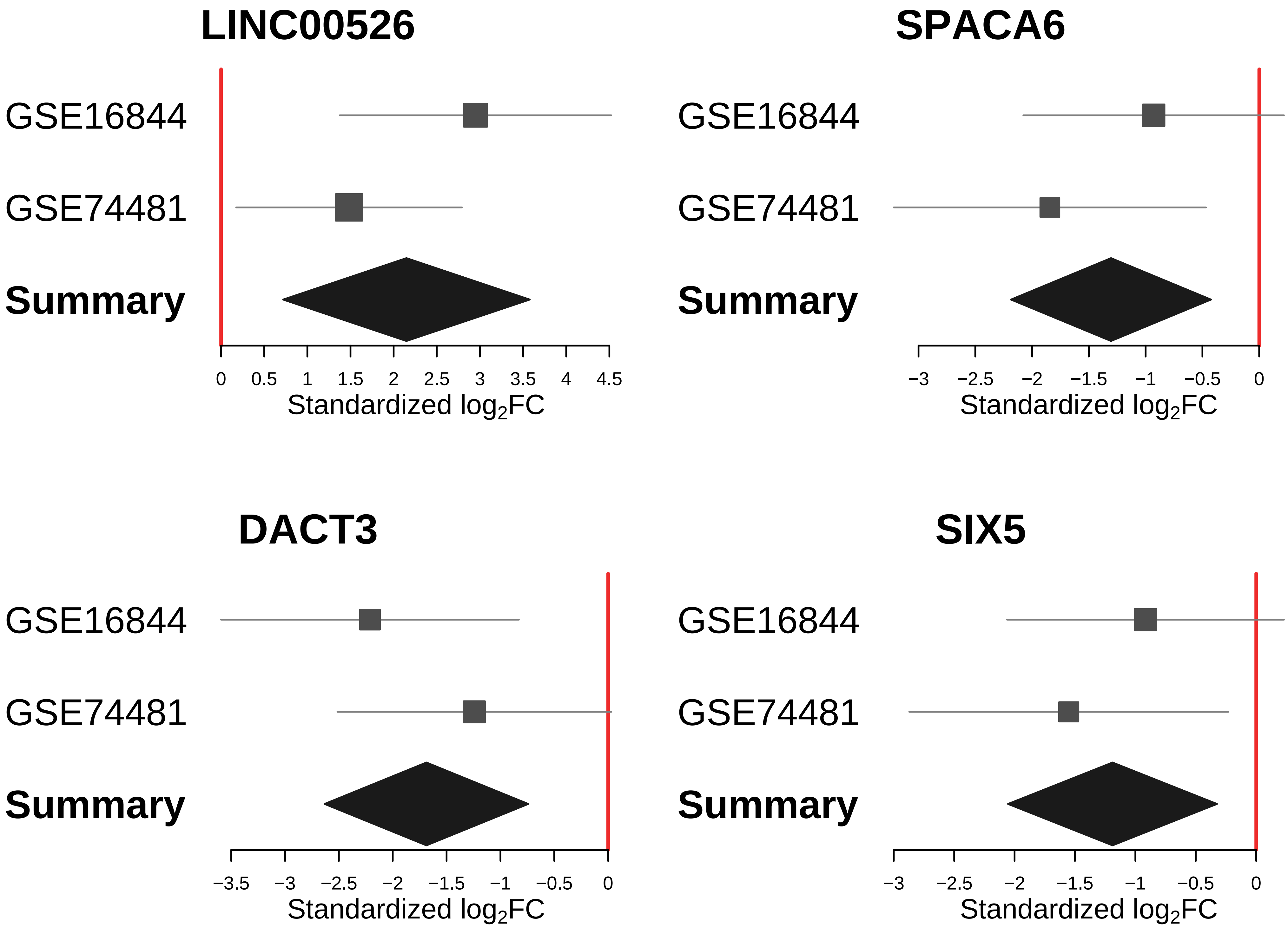
LINC00526
GSE16844
GSE74481
Summary
0 0.5 1 1.5 2 2.5 3 3.5 4 4.5
Standardized log2FC
SPACA6
GSE16844
GSE74481
Summary
−3 −2.5 −2 −1.5 −1 −0.5 0
Standardized log2FC
DACT3
GSE16844
GSE74481
Summary
−3.5 −3 −2.5 −2 −1.5 −1 −0.5 0
Standardized log2FC
SIX5
GSE16844
GSE74481
Summary
−3 −2.5 −2 −1.5 −1 −0.5 0
Standardized log2FC

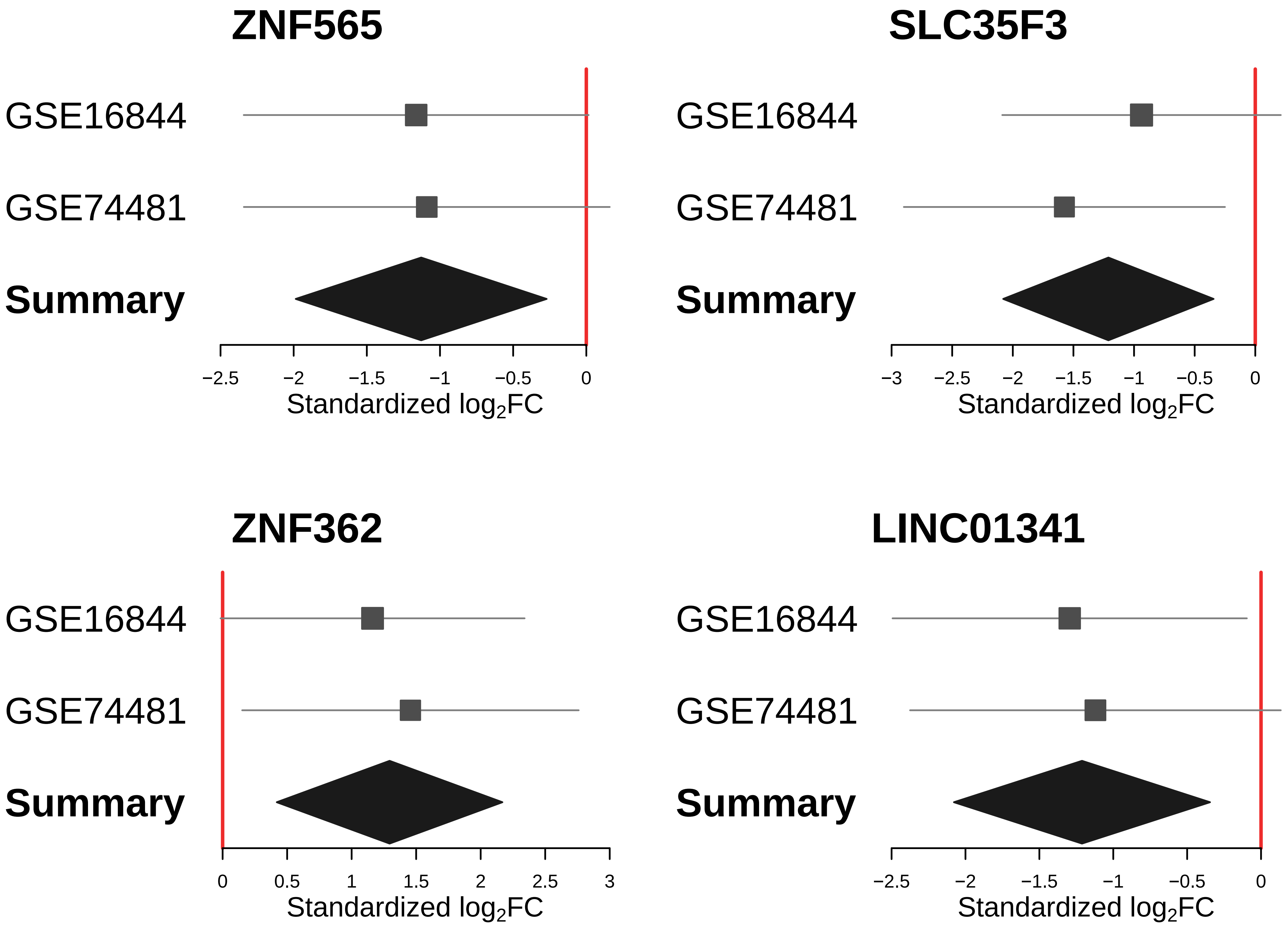

ZNF565
GSE16844
GSE74481
Summary
−2.5 −2 −1.5 −1 −0.5 0
Standardized log2FC
SLC35F3
GSE16844
GSE74481
Summary
−3 −2.5 −2 −1.5 −1 −0.5 0
Standardized log2FC
ZNF362
GSE16844
GSE74481
Summary
0 0.5 1 1.5 2 2.5 3
Standardized log2FC
LINC01341
GSE16844
GSE74481
Summary
−2.5 −2 −1.5 −1 −0.5 0
Standardized log2FC

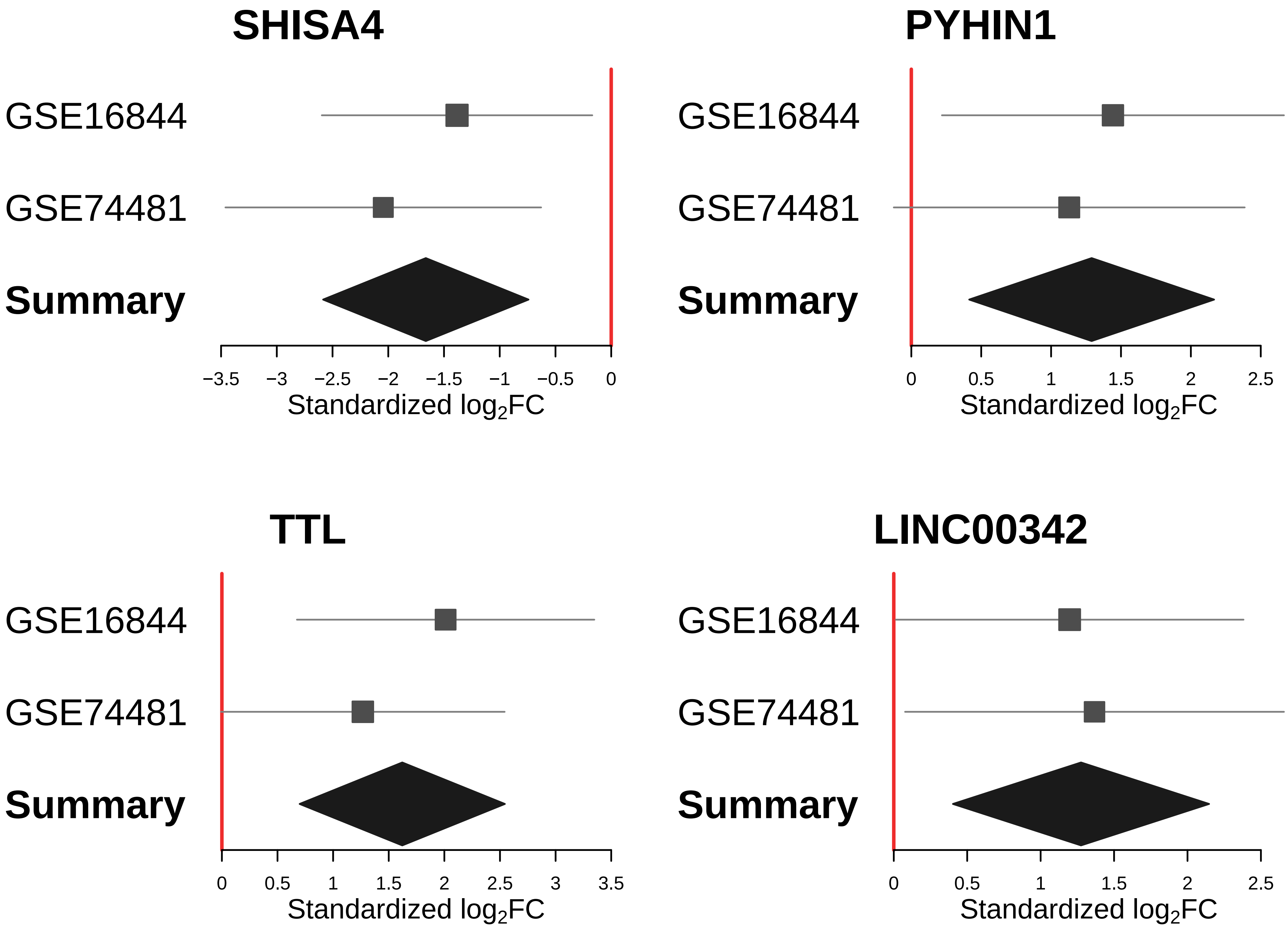
SHISA4
GSE16844
GSE74481
Summary
−3.5 −3 −2.5 −2 −1.5 −1 −0.5 0
Standardized $log_2FC$
PYHIN1
GSE16844
GSE74481
Summary
0 0.5 1 1.5 2 2.5
Standardized $log_2FC$
TTL
GSE16844
GSE74481
Summary
0 0.5 1 1.5 2 2.5 3 3.5
Standardized $log_2FC$
LINC00342
GSE16844
GSE74481
Summary
0 0.5 1 1.5 2 2.5
Standardized $log_2FC$

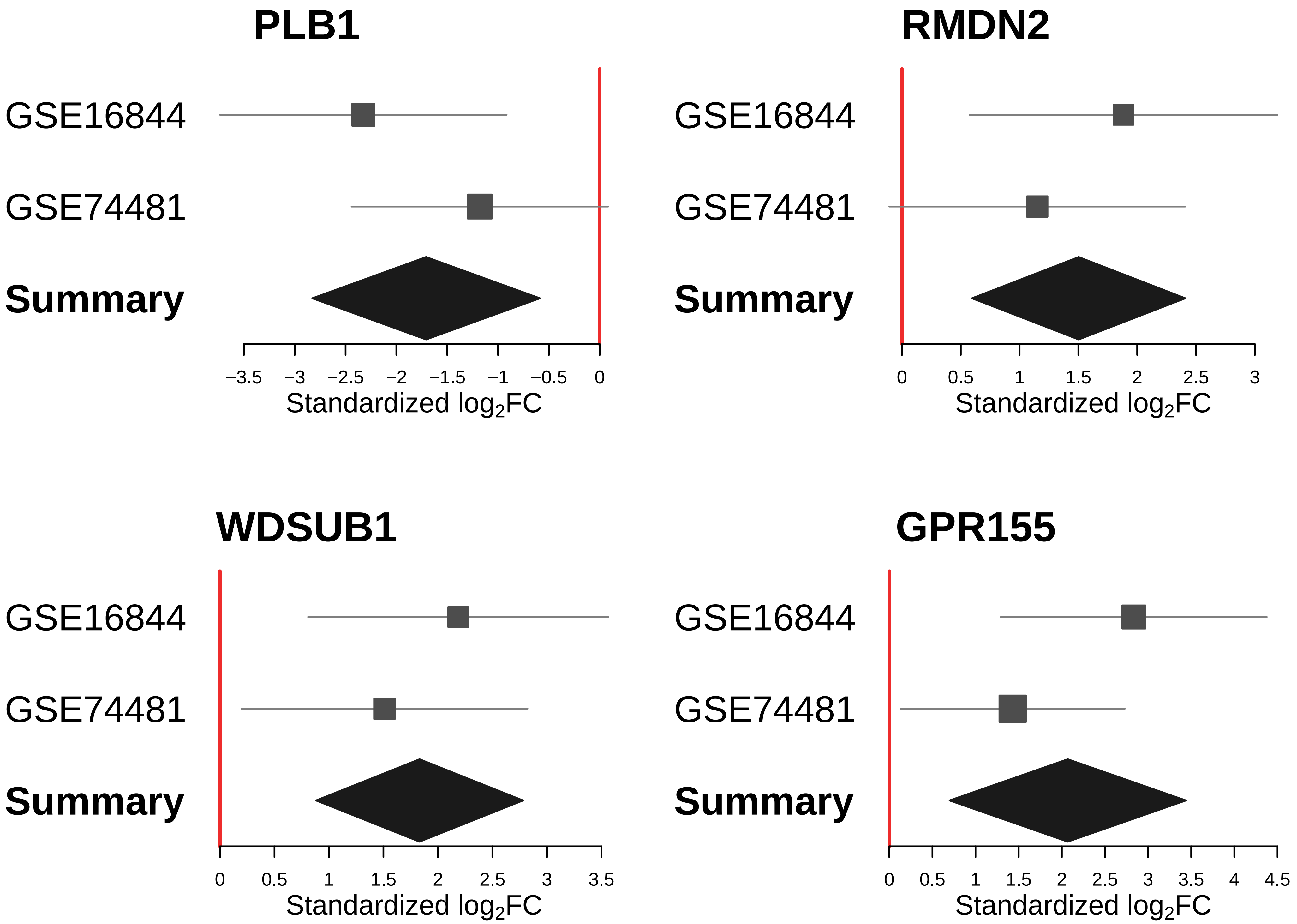
PLB1
GSE16844
GSE74481
Summary
−3.5 −3 −2.5 −2 −1.5 −1 −0.5 0
Standardized log2FC
RMDN2
GSE16844
GSE74481
Summary
0 0.5 1 1.5 2 2.5 3
Standardized log2FC
WDSUB1
GSE16844
GSE74481
Summary
0 0.5 1 1.5 2 2.5 3 3.5
Standardized log2FC
GPR155
GSE16844
GSE74481
Summary
0 0.5 1 1.5 2 2.5 3 3.5 4 4.5
Standardized log2FC

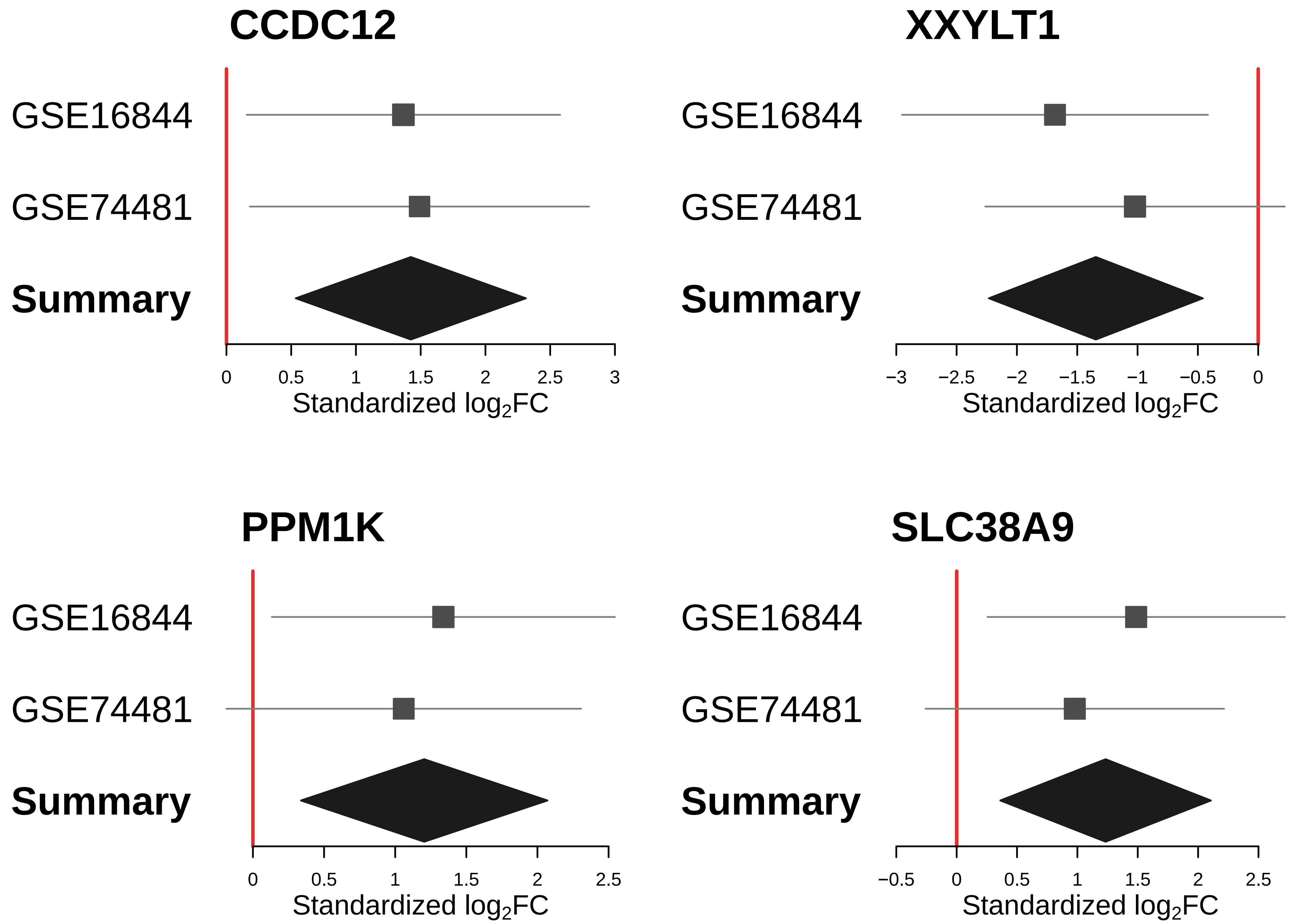

CCDC12
XXYLT1
PPM1K
SLC38A9
GSE16844
GSE74481
Summary
Standardized $log_2FC$
0
0.5
1
1.5
2
2.5
3
−3
−2.5
−2
−1.5
−1
−0.5

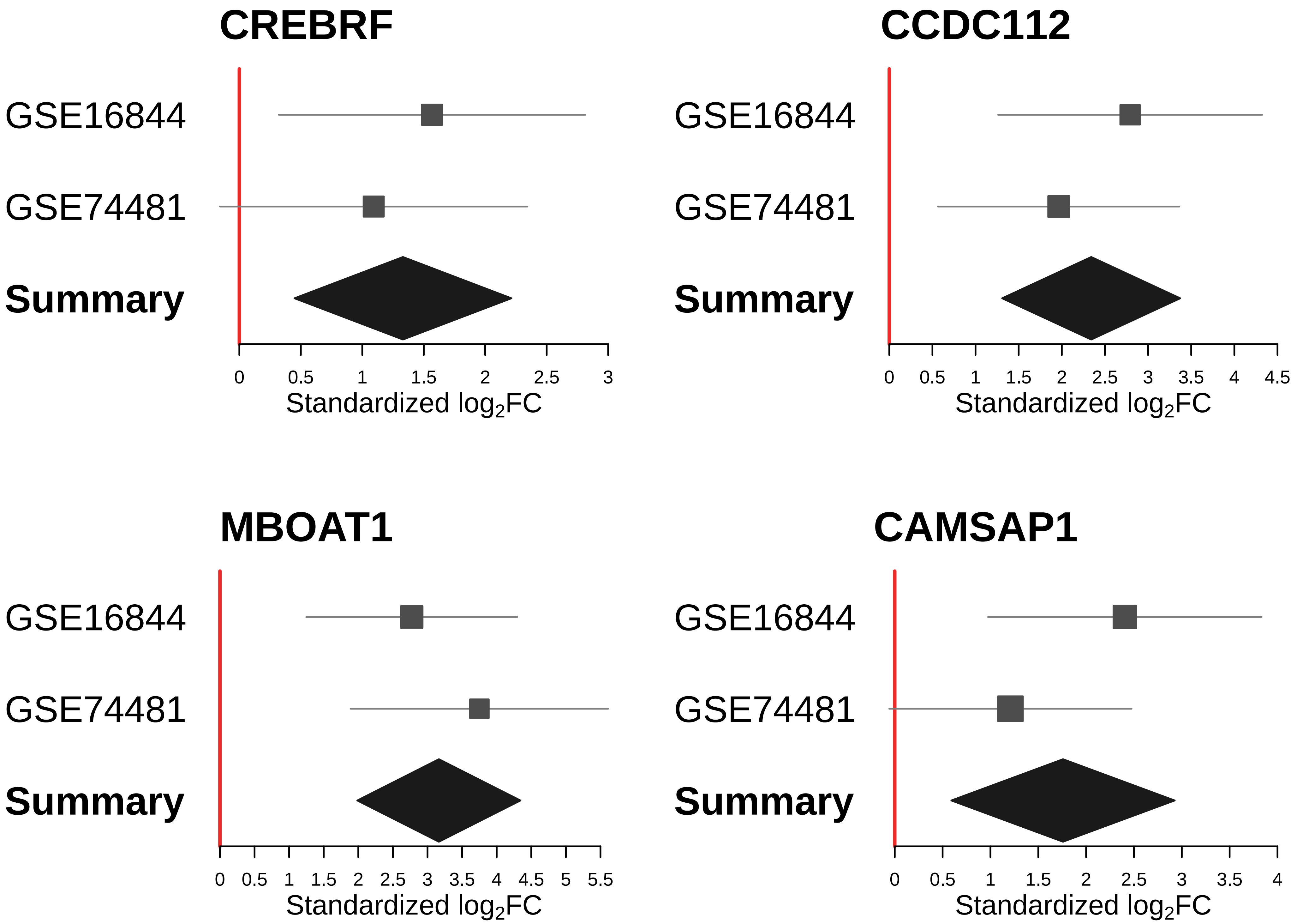
CREBRF
GSE16844
GSE74481
Summary
0 0.5 1 1.5 2 2.5 3
Standardized log2FC
CCDC112
GSE16844
GSE74481
Summary
0 0.5 1 1.5 2 2.5 3 3.5 4 4.5
Standardized log2FC
MBOAT1
GSE16844
GSE74481
Summary
0 0.5 1 1.5 2 2.5 3 3.5 4 4.5 5 5.5
Standardized log2FC
CAMSAP1
GSE16844
GSE74481
Summary
0 0.5 1 1.5 2 2.5 3 3.5 4
Standardized log2FC

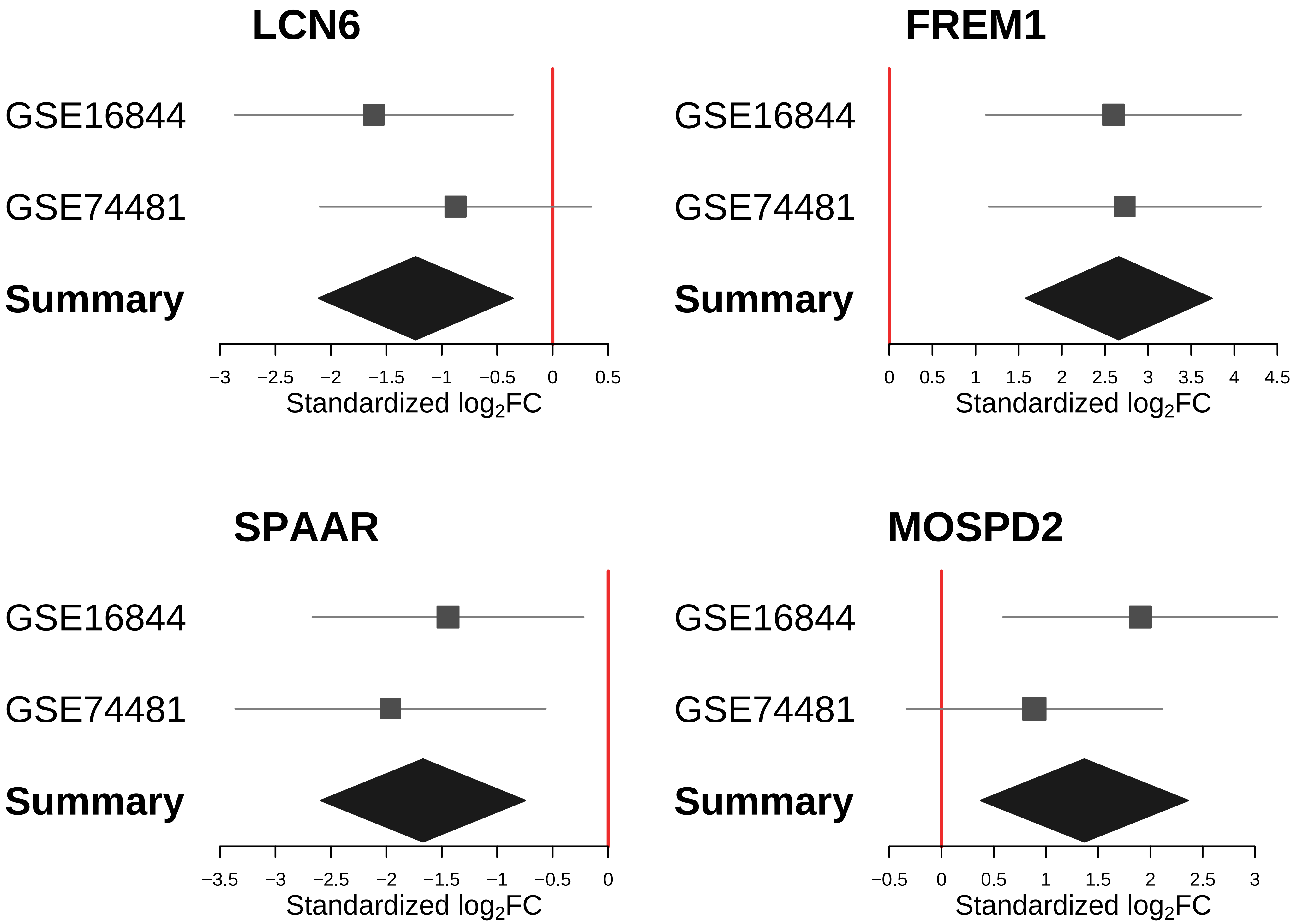
LCN6
GSE16844
GSE74481
Summary
−3
−2.5
−2
−1.5
−1
−0.5
0
0.5
Standardized log2FC
FREM1
GSE16844
GSE74481
Summary
0
0.5
1
1.5
2
2.5
3
3.5
4
4.5
Standardized log2FC
SPAAR
GSE16844
GSE74481
Summary
−3.5
−3
−2.5
−2
−1.5
−1
−0.5
0
Standardized log2FC
MOSPD2
GSE16844
GSE74481
Summary
−0.5
0
0.5
1
1.5
2
2.5
3
Standardized log2FC

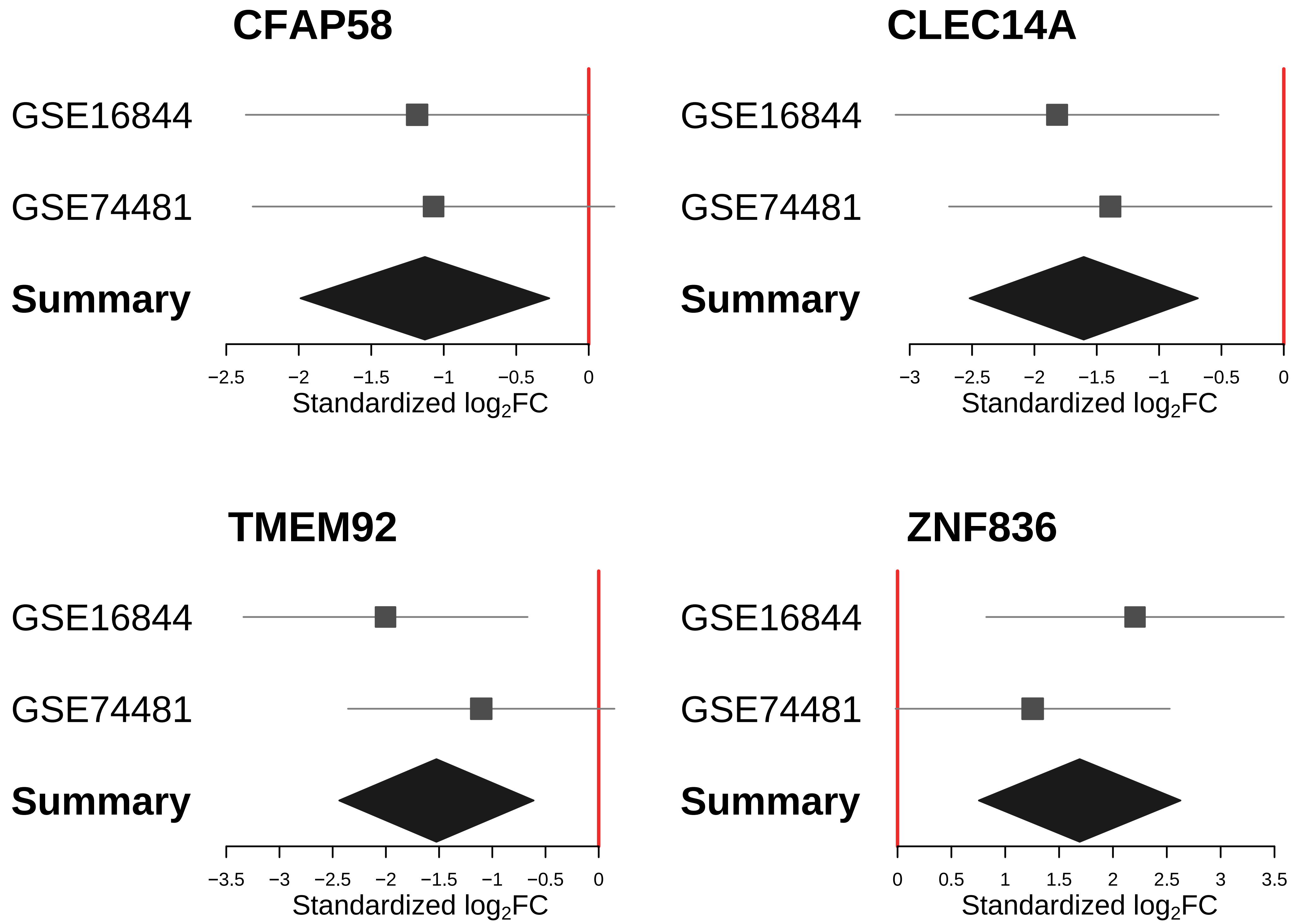
CFAP58
GSE16844
GSE74481
Summary
−2.5 −2 −1.5 −1 −0.5 0
Standardized log2FC
CLEC14A
GSE16844
GSE74481
Summary
−3 −2.5 −2 −1.5 −1 −0.5 0
Standardized log2FC
TMEM92
GSE16844
GSE74481
Summary
−3.5 −3 −2.5 −2 −1.5 −1 −0.5 0
Standardized log2FC
ZNF836
GSE16844
GSE74481
Summary
0 0.5 1 1.5 2 2.5 3 3.5
Standardized log2FC

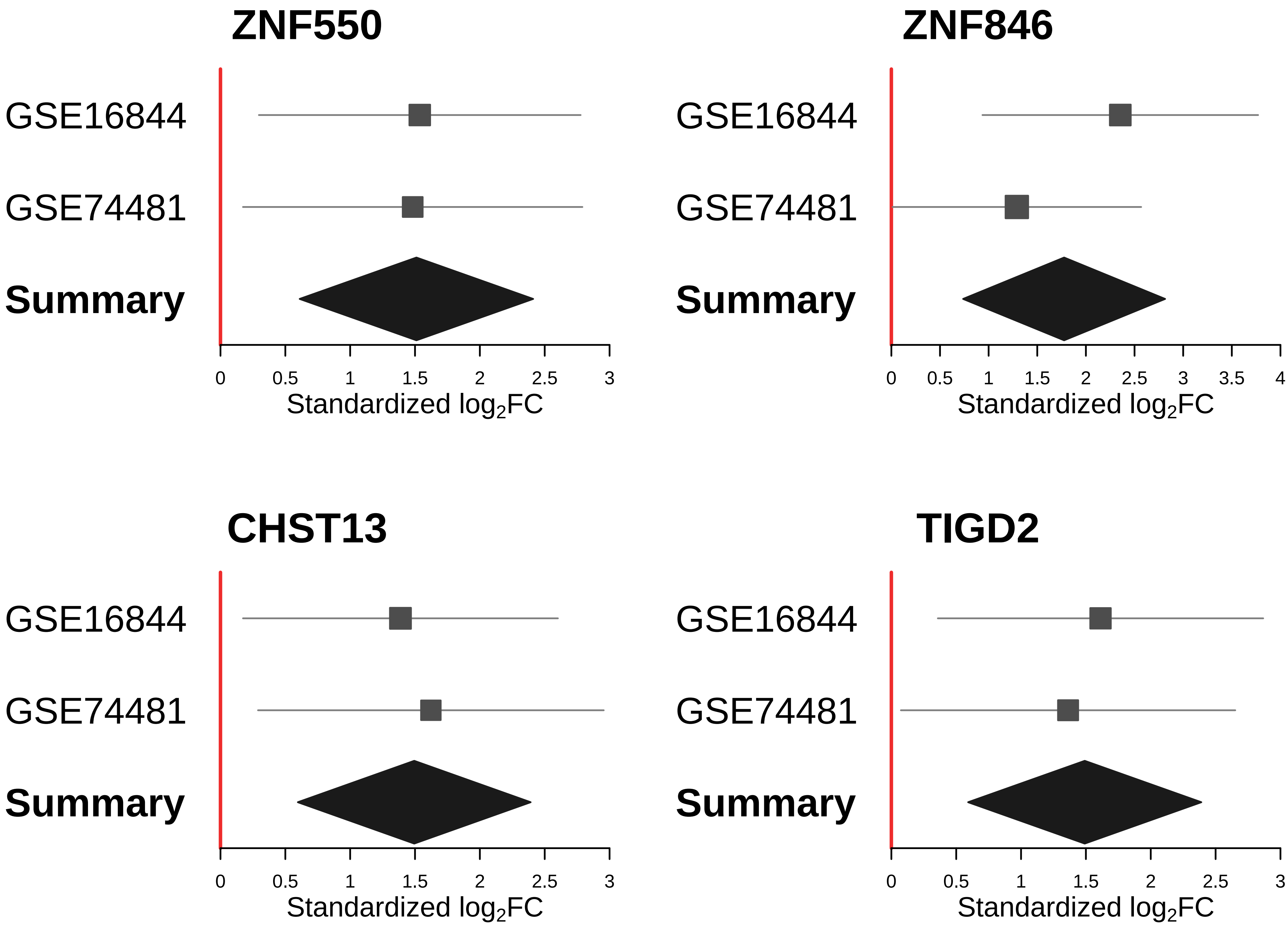
ZNF550
GSE16844
GSE74481
Summary
0 0.5 1 1.5 2 2.5 3
Standardized log2FC
ZNF846
GSE16844
GSE74481
Summary
0 0.5 1 1.5 2 2.5 3 3.5 4
Standardized log2FC
CHST13
GSE16844
GSE74481
Summary
0 0.5 1 1.5 2 2.5 3
Standardized log2FC
TIGD2
GSE16844
GSE74481
Summary
0 0.5 1 1.5 2 2.5 3
Standardized log2FC

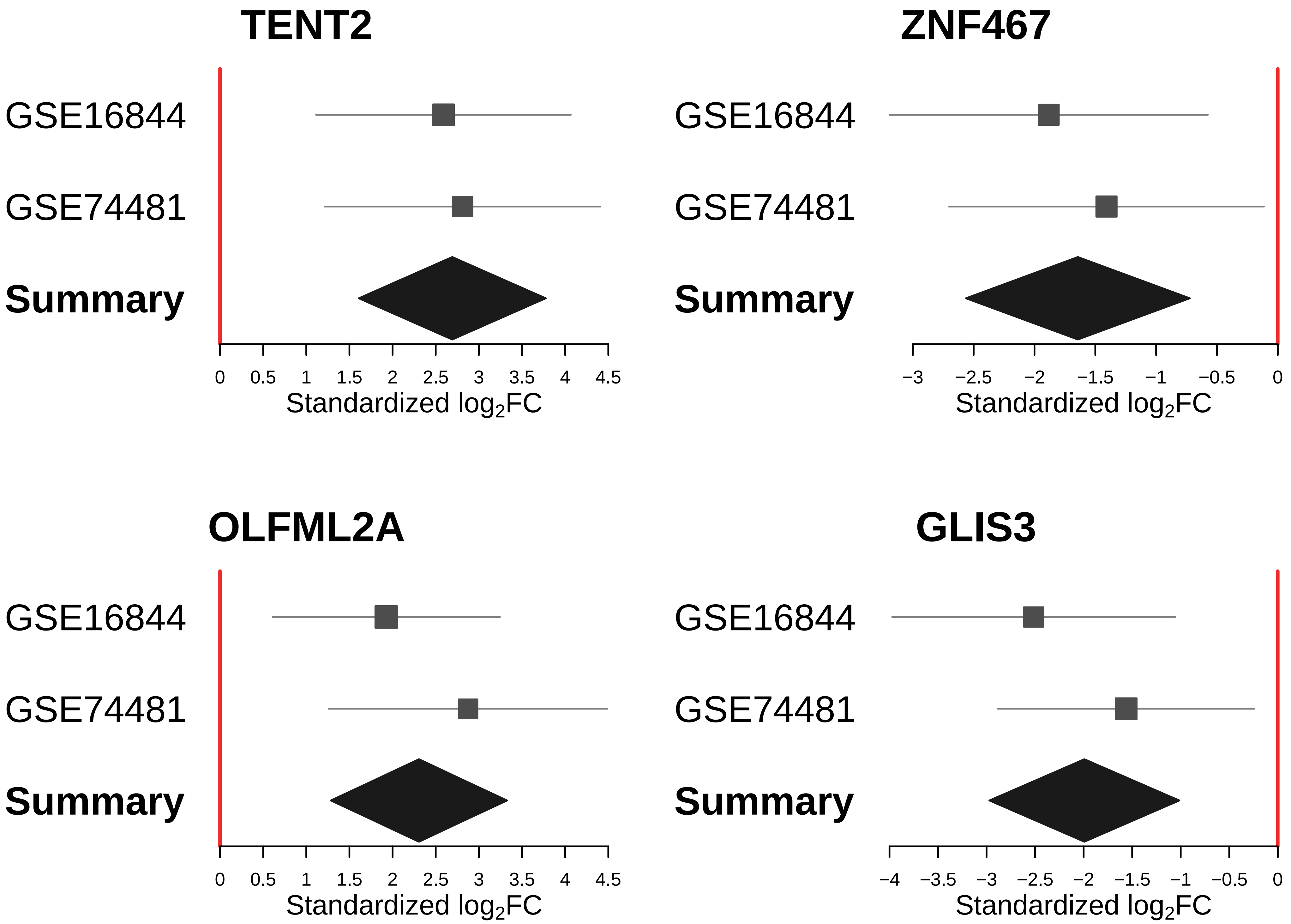

TENT2
GSE16844
GSE74481
Summary
0 0.5 1 1.5 2 2.5 3 3.5 4 4.5
Standardized log2FC
ZNF467
GSE16844
GSE74481
Summary
−3 −2.5 −2 −1.5 −1 −0.5 0
Standardized log2FC
OLFML2A
GSE16844
GSE74481
Summary
0 0.5 1 1.5 2 2.5 3 3.5 4 4.5
Standardized log2FC
GLIS3
GSE16844
GSE74481
Summary
−4 −3.5 −3 −2.5 −2 −1.5 −1 −0.5 0
Standardized log2FC

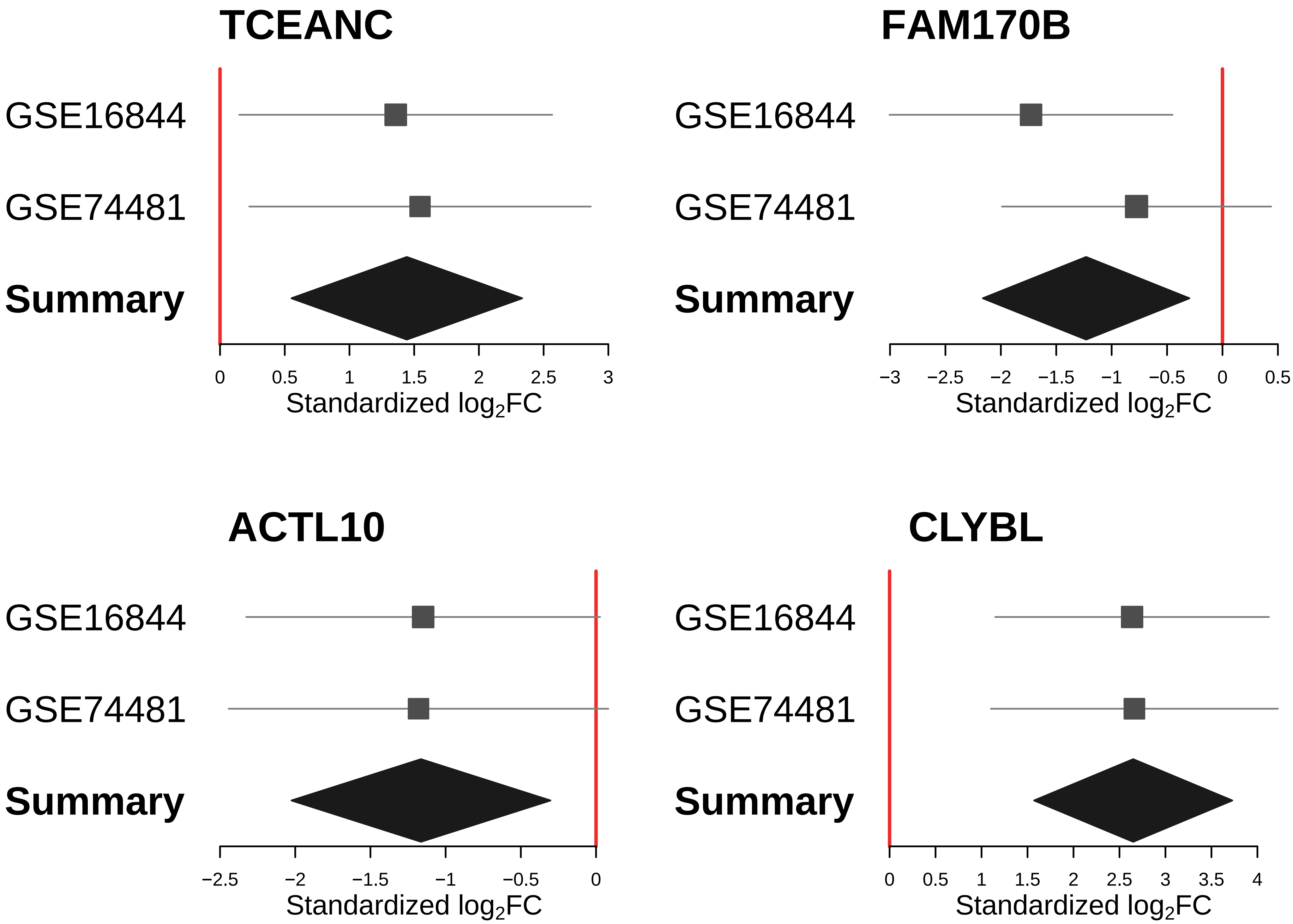
TCEANC
GSE16844
GSE74481
Summary
0
0.5
1
1.5
2
2.5
3
Standardized $log_2FC$
FAM170B
GSE16844
GSE74481
Summary
−3
−2.5
−2
−1.5
−1
−0.5
0
0.5
Standardized $log_2FC$
ACTL10
GSE16844
GSE74481
Summary
−2.5
−2
−1.5
−1
−0.5
0
Standardized $log_2FC$
CLYBL
GSE16844
GSE74481
Summary
0
0.5
1
1.5
2
2.5
3
3.5
4
Standardized $log_2FC$

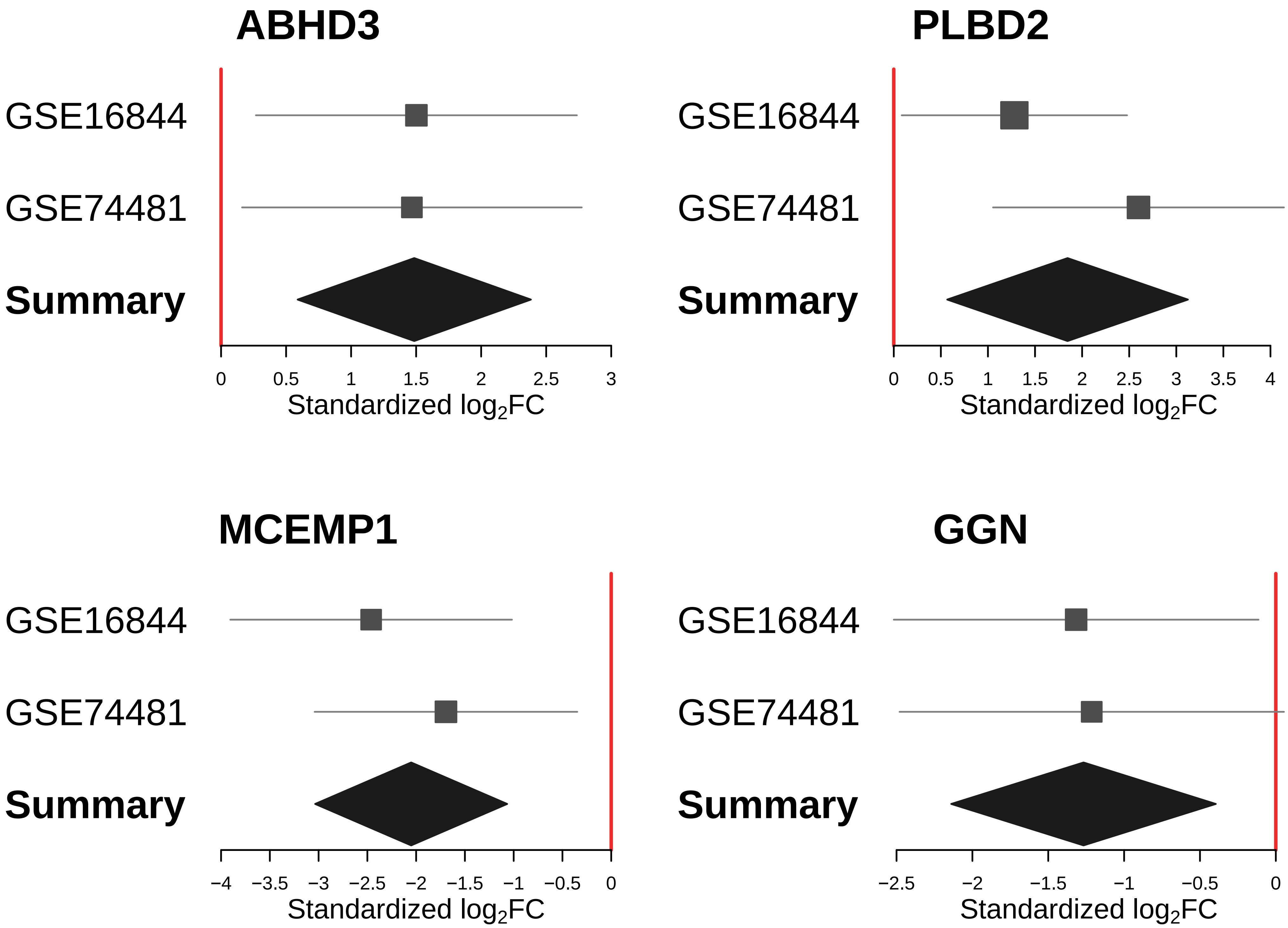
ABHD3
GSE16844
GSE74481
Summary
0
0.5
1
1.5
2
2.5
3
Standardized $log_2FC$
PLBD2
GSE16844
GSE74481
Summary
0
0.5
1
1.5
2
2.5
3
3.5
4
Standardized $log_2FC$
MCEMP1
GSE16844
GSE74481
Summary
−4
−3.5
−3
−2.5
−2
−1.5
−1
−0.5
0
Standardized $log_2FC$
GGN
GSE16844
GSE74481
Summary
−2.5
−2
−1.5
−1
−0.5
0
Standardized $log_2FC$

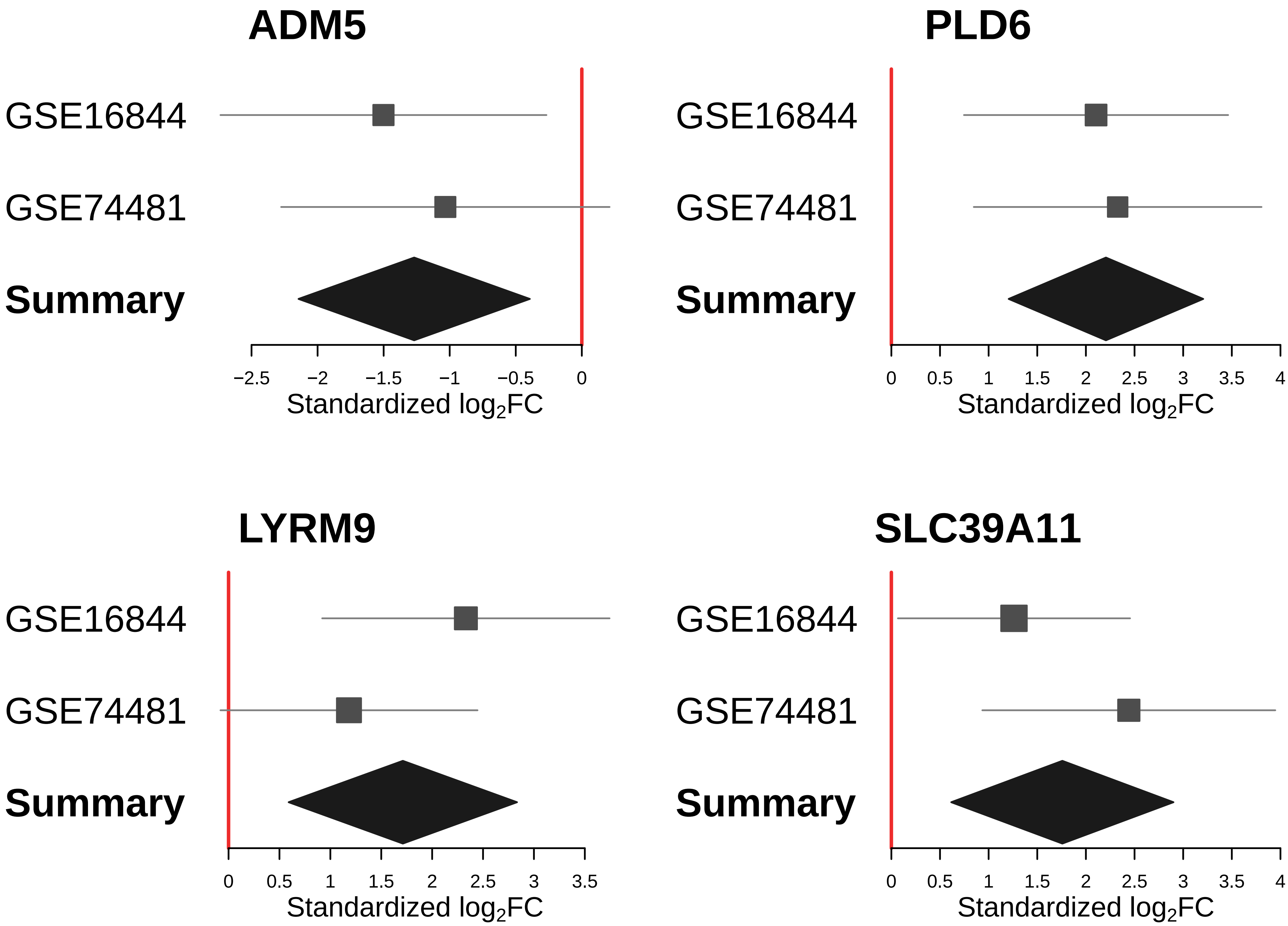

ADM5
GSE16844
GSE74481
Summary
−2.5 −2 −1.5 −1 −0.5 0
Standardized log2FC
PLD6
GSE16844
GSE74481
Summary
0 0.5 1 1.5 2 2.5 3 3.5 4
Standardized log2FC
LYRM9
GSE16844
GSE74481
Summary
0 0.5 1 1.5 2 2.5 3 3.5
Standardized log2FC
SLC39A11
GSE16844
GSE74481
Summary
0 0.5 1 1.5 2 2.5 3 3.5 4
Standardized log2FC

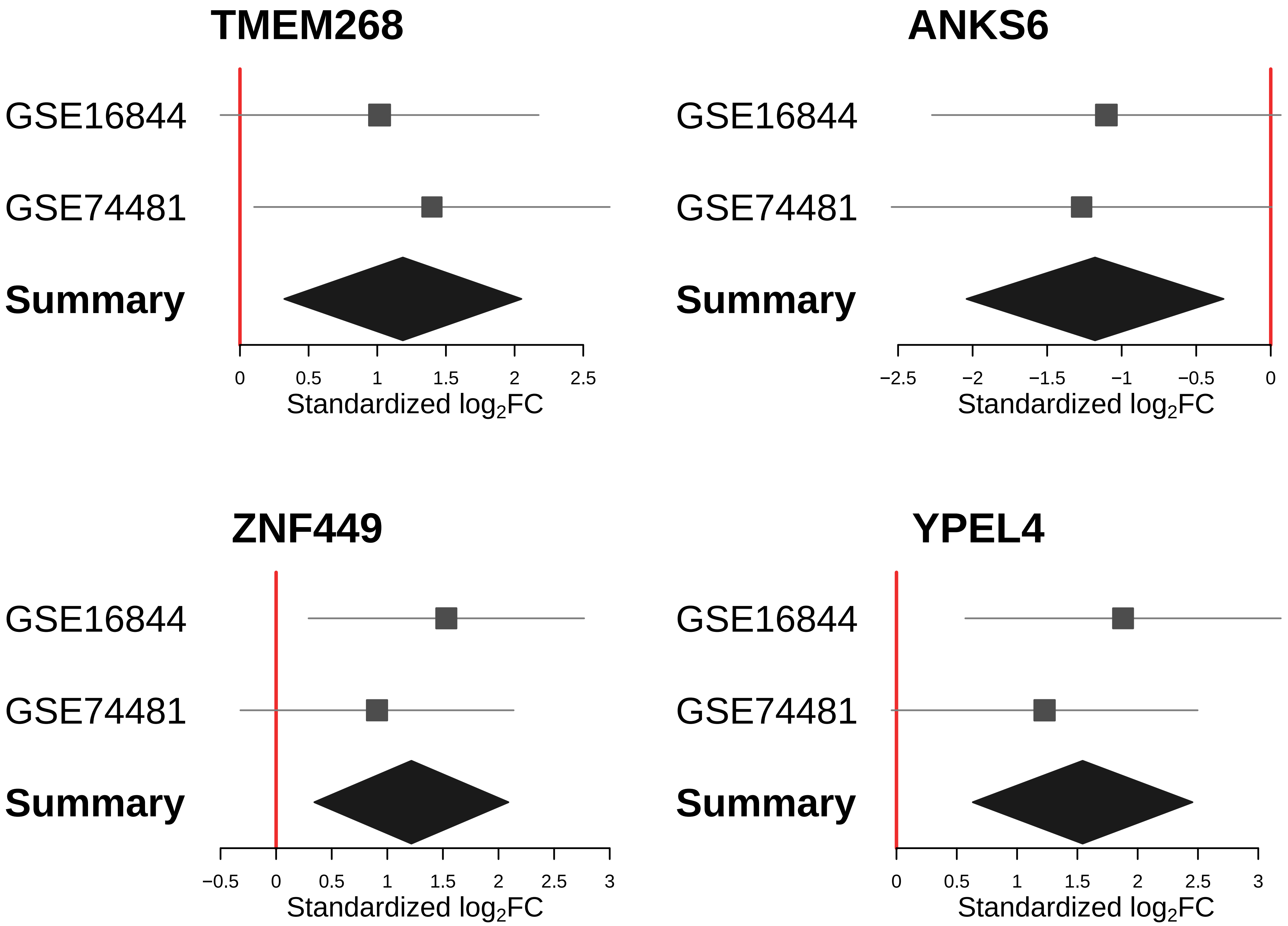
TMEM268
GSE16844
GSE74481
Summary
0
0.5
1
1.5
2
2.5
Standardized log2FC
ANKS6
GSE16844
GSE74481
Summary
−2.5
−2
−1.5
−1
−0.5
0
Standardized log2FC
ZNF449
GSE16844
GSE74481
Summary
−0.5
0
0.5
1
1.5
2
2.5
3
Standardized log2FC
YPEL4
GSE16844
GSE74481
Summary
0
0.5
1
1.5
2
2.5
3
Standardized log2FC

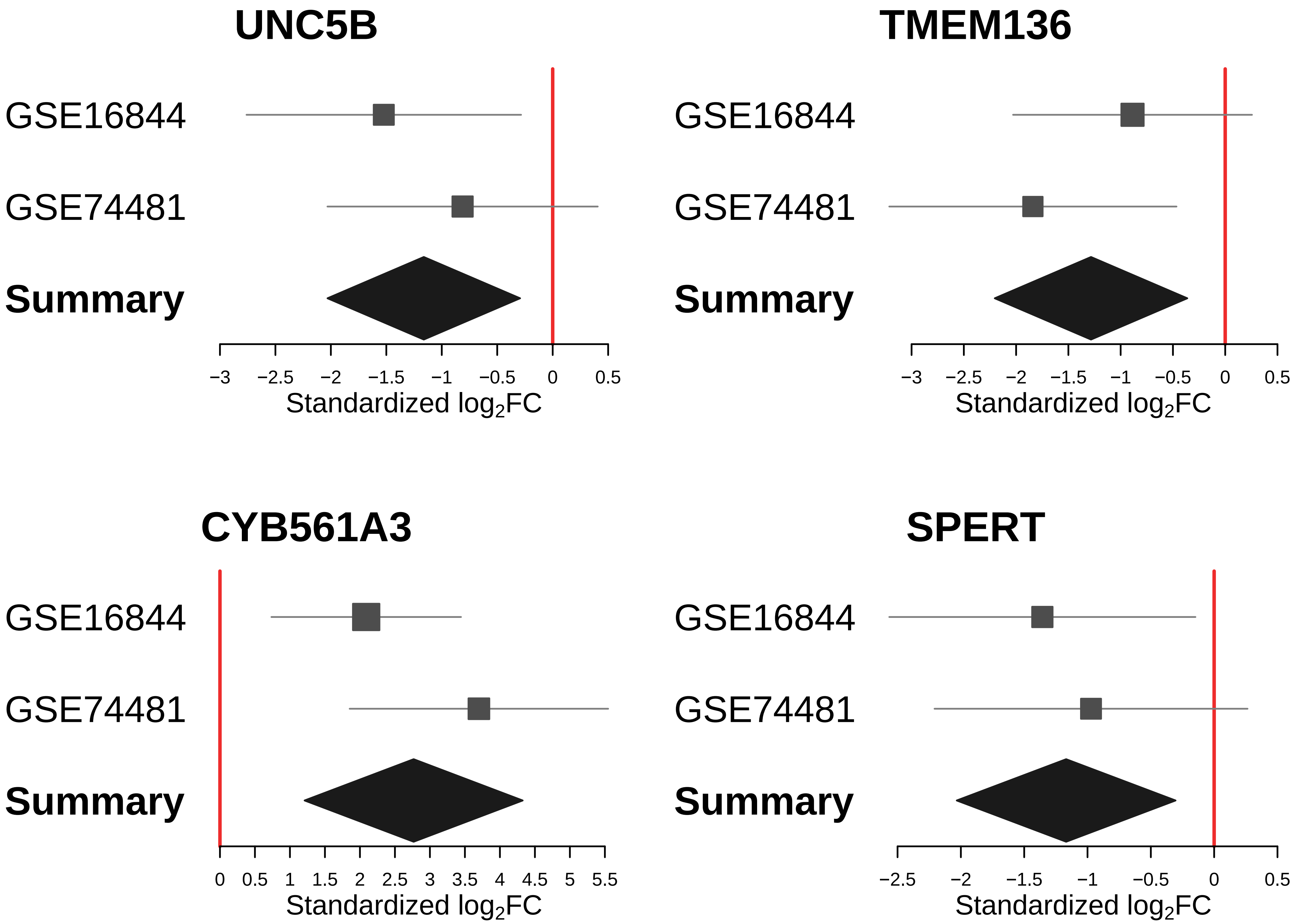
UNC5B
GSE16844
GSE74481
Summary
−3 −2.5 −2 −1.5 −1 −0.5 0 0.5
Standardized log2FC
TMEM136
GSE16844
GSE74481
Summary
−3 −2.5 −2 −1.5 −1 −0.5 0 0.5
Standardized log2FC
CYB561A3
GSE16844
GSE74481
Summary
0 0.5 1 1.5 2 2.5 3 3.5 4 4.5 5 5.5
Standardized log2FC
SPERT
GSE16844
GSE74481
Summary
−2.5 −2 −1.5 −1 −0.5 0 0.5
Standardized log2FC

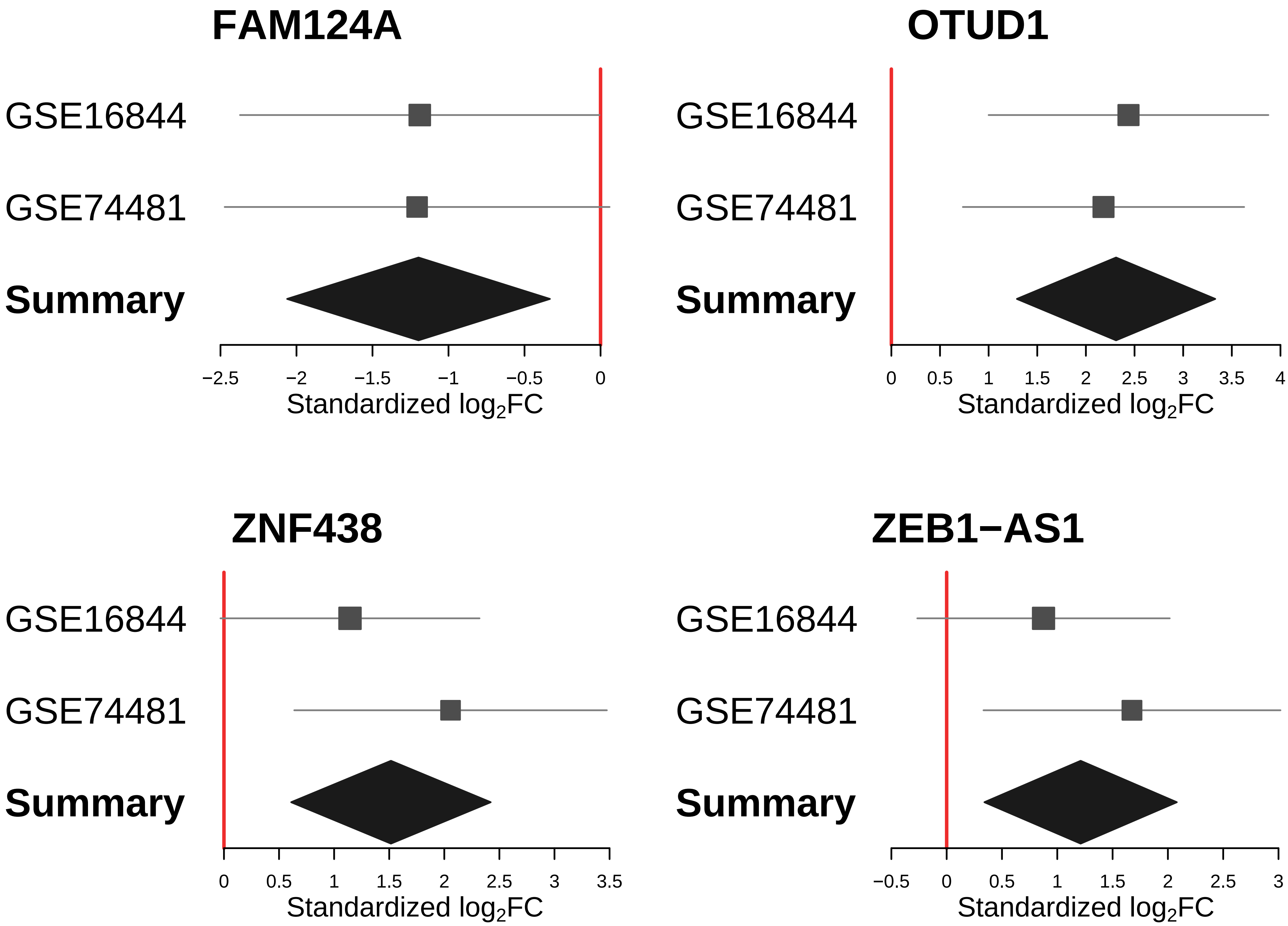

FAM124A
GSE16844
GSE74481
Summary
−2.5
−2
−1.5
−1
−0.5
0
Standardized log2FC
OTUD1
GSE16844
GSE74481
Summary
0
0.5
1
1.5
2
2.5
3
3.5
4
Standardized log2FC
ZNF438
GSE16844
GSE74481
Summary
0
0.5
1
1.5
2
2.5
3
3.5
Standardized log2FC
ZEB1−AS1
GSE16844
GSE74481
Summary
−0.5
0
0.5
1
1.5
2
2.5
3
Standardized log2FC

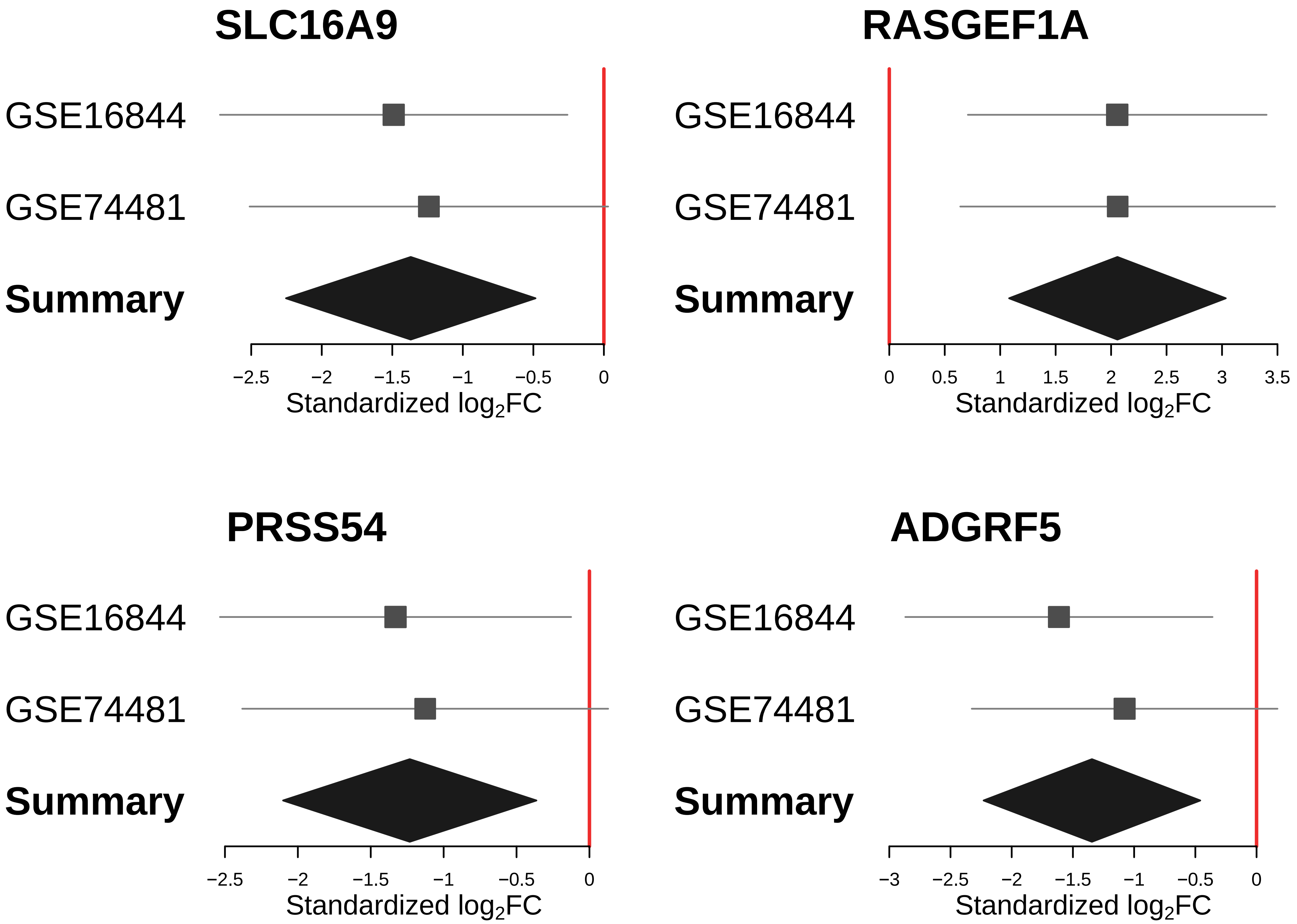
SLC16A9
GSE16844
GSE74481
Summary
−2.5 −2 −1.5 −1 −0.5 0
Standardized log2FC
RASGEF1A
GSE16844
GSE74481
Summary
0 0.5 1 1.5 2 2.5 3 3.5
Standardized log2FC
PRSS54
GSE16844
GSE74481
Summary
−2.5 −2 −1.5 −1 −0.5 0
Standardized log2FC
ADGRF5
GSE16844
GSE74481
Summary
−3 −2.5 −2 −1.5 −1 −0.5 0
Standardized log2FC

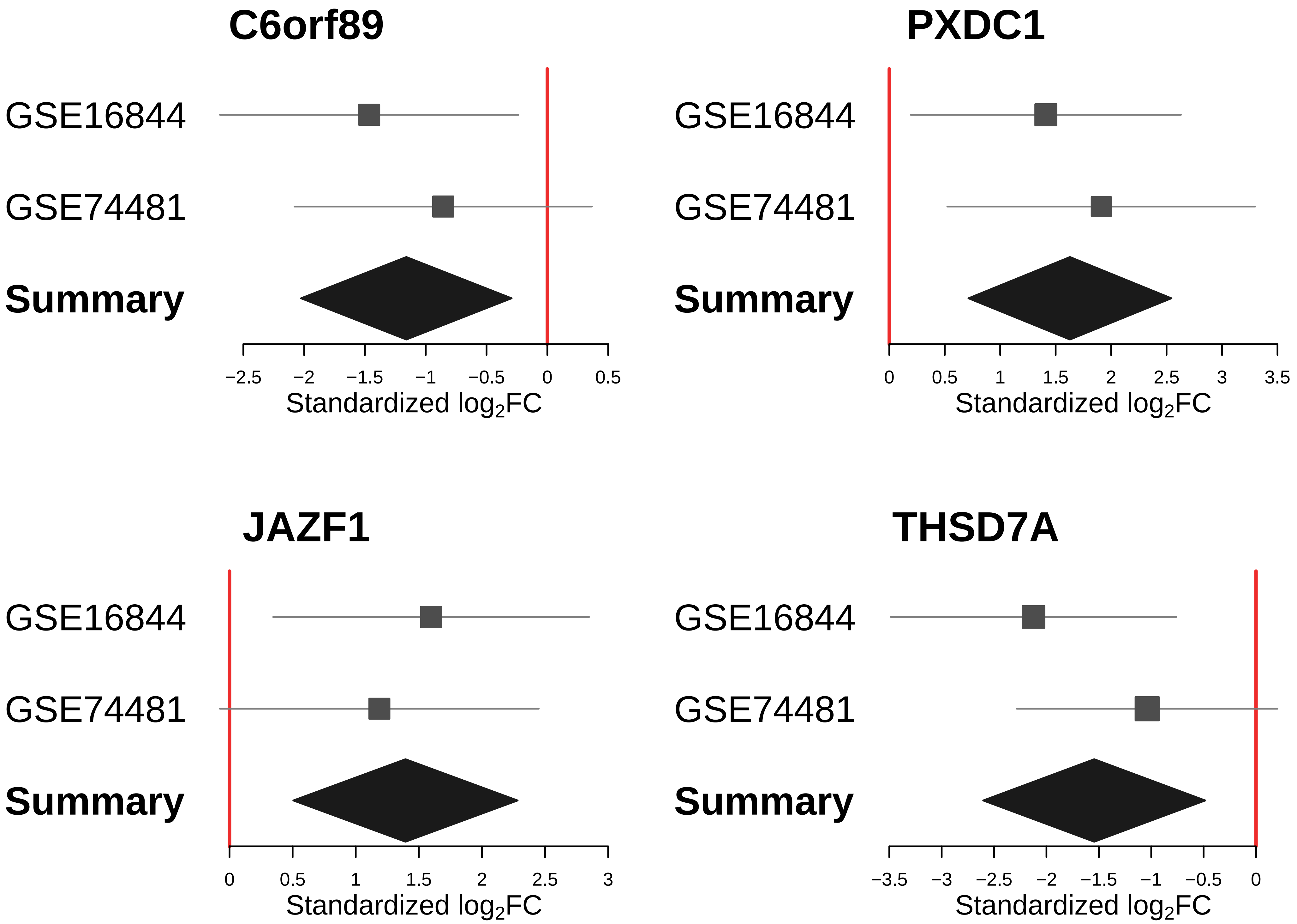
C6orf89
GSE16844
GSE74481
Summary
−2.5
−2
−1.5
−1
−0.5
0
0.5
Standardized log2FC
PXDC1
GSE16844
GSE74481
Summary
0
0.5
1
1.5
2
2.5
3
3.5
Standardized log2FC
JAZF1
GSE16844
GSE74481
Summary
0
0.5
1
1.5
2
2.5
3
Standardized log2FC
THSD7A
GSE16844
GSE74481
Summary
−3.5
−3
−2.5
−2
−1.5
−1
−0.5
0
Standardized log2FC

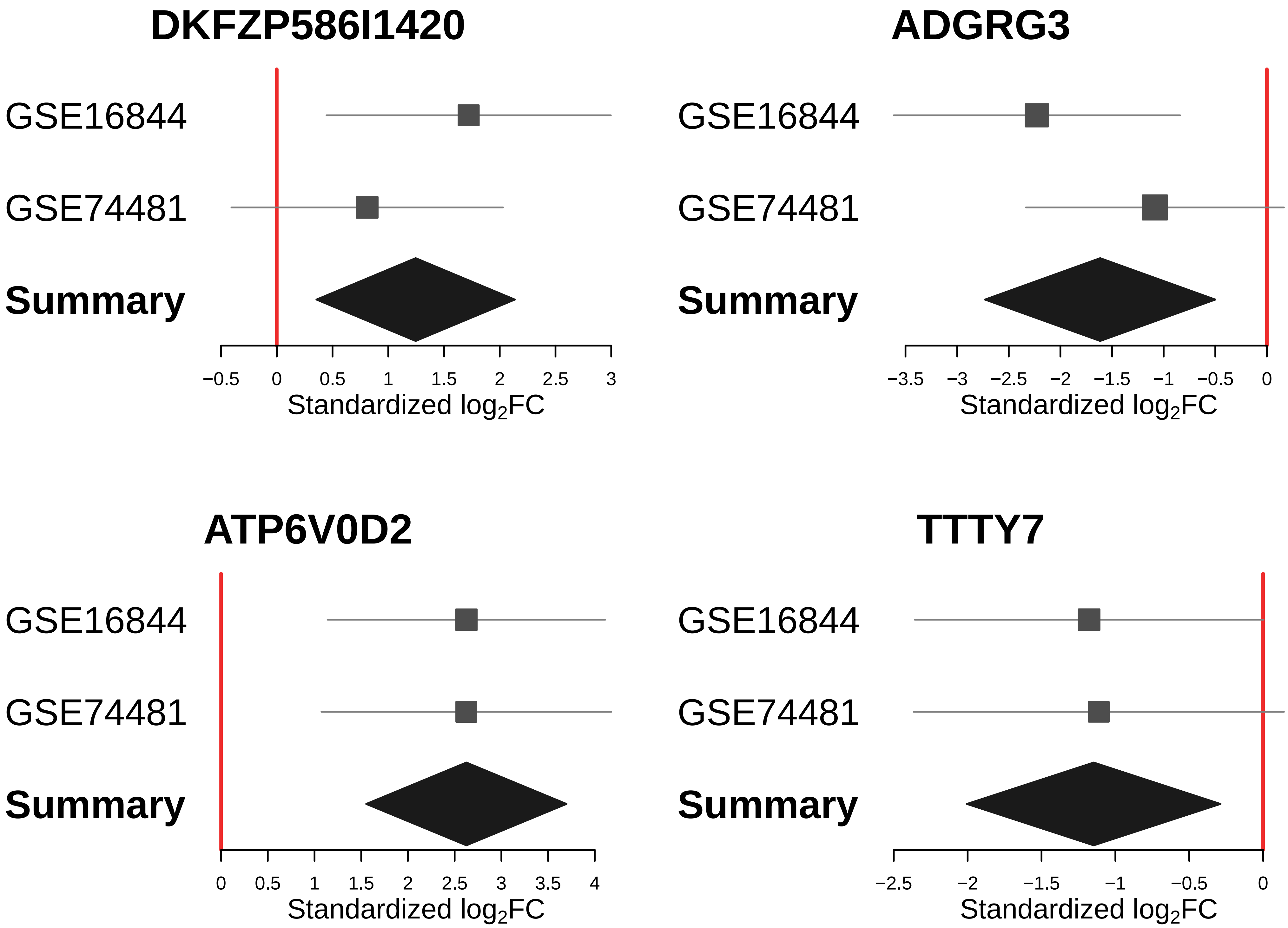
DKFZP586I1420
GSE16844
GSE74481
Summary
−0.5 0 0.5 1 1.5 2 2.5 3
Standardized log2FC
ADGRG3
GSE16844
GSE74481
Summary
−3.5 −3 −2.5 −2 −1.5 −1 −0.5 0
Standardized log2FC
ATP6V0D2
GSE16844
GSE74481
Summary
0 0.5 1 1.5 2 2.5 3 3.5 4
Standardized log2FC
TTTY7
GSE16844
GSE74481
Summary
−2.5 −2 −1.5 −1 −0.5 0
Standardized log2FC

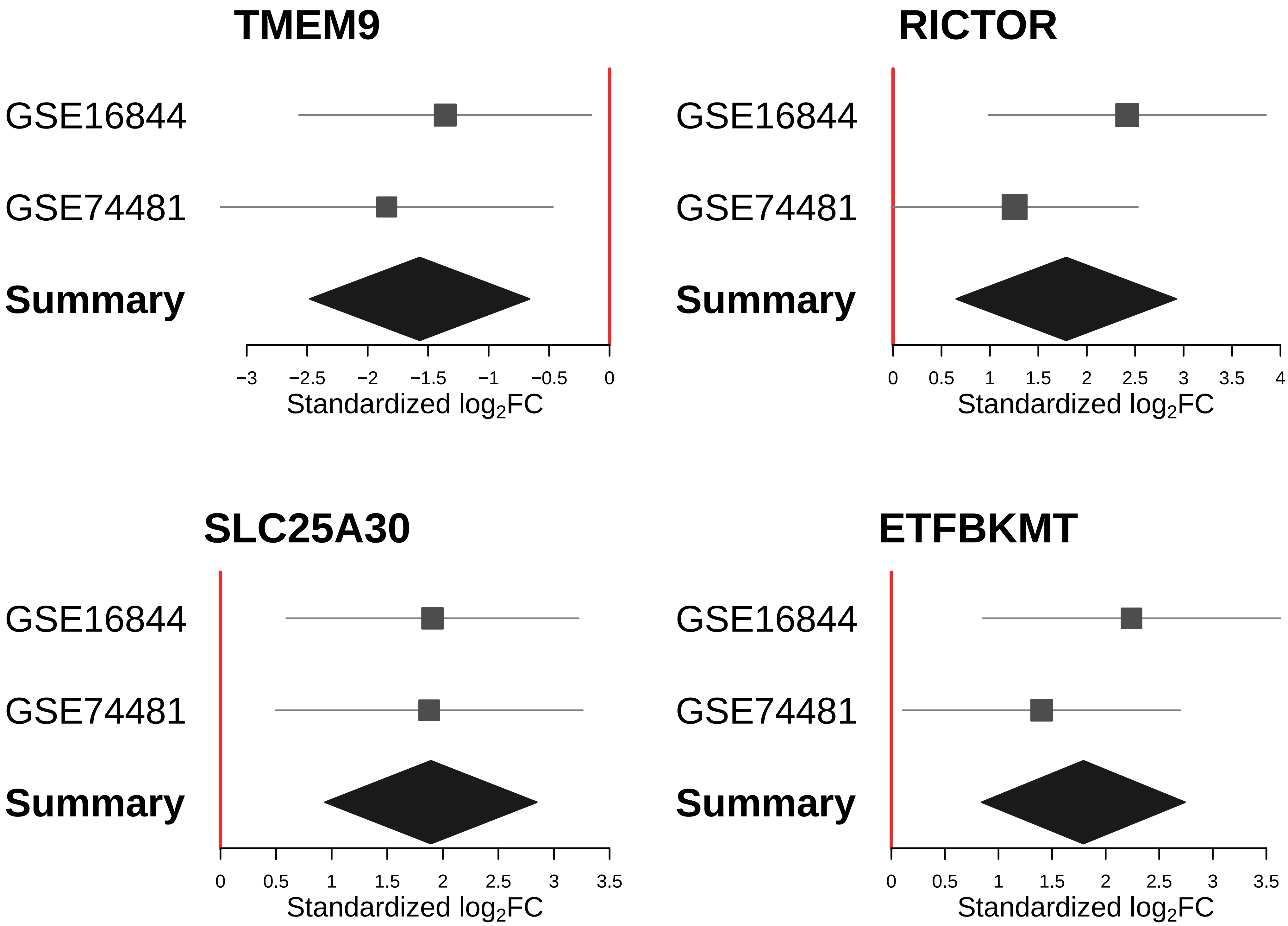
TMEM9
GSE16844
GSE74481
Summary
−3
−2.5
−2
−1.5
−1
−0.5
0
Standardized log2FC
RICTOR
GSE16844
GSE74481
Summary
0
0.5
1
1.5
2
2.5
3
3.5
4
Standardized log2FC
SLC25A30
GSE16844
GSE74481
Summary
0
0.5
1
1.5
2
2.5
3
3.5
Standardized log2FC
ETFBKMT
GSE16844
GSE74481
Summary
0
0.5
1
1.5
2
2.5
3
3.5
Standardized log2FC

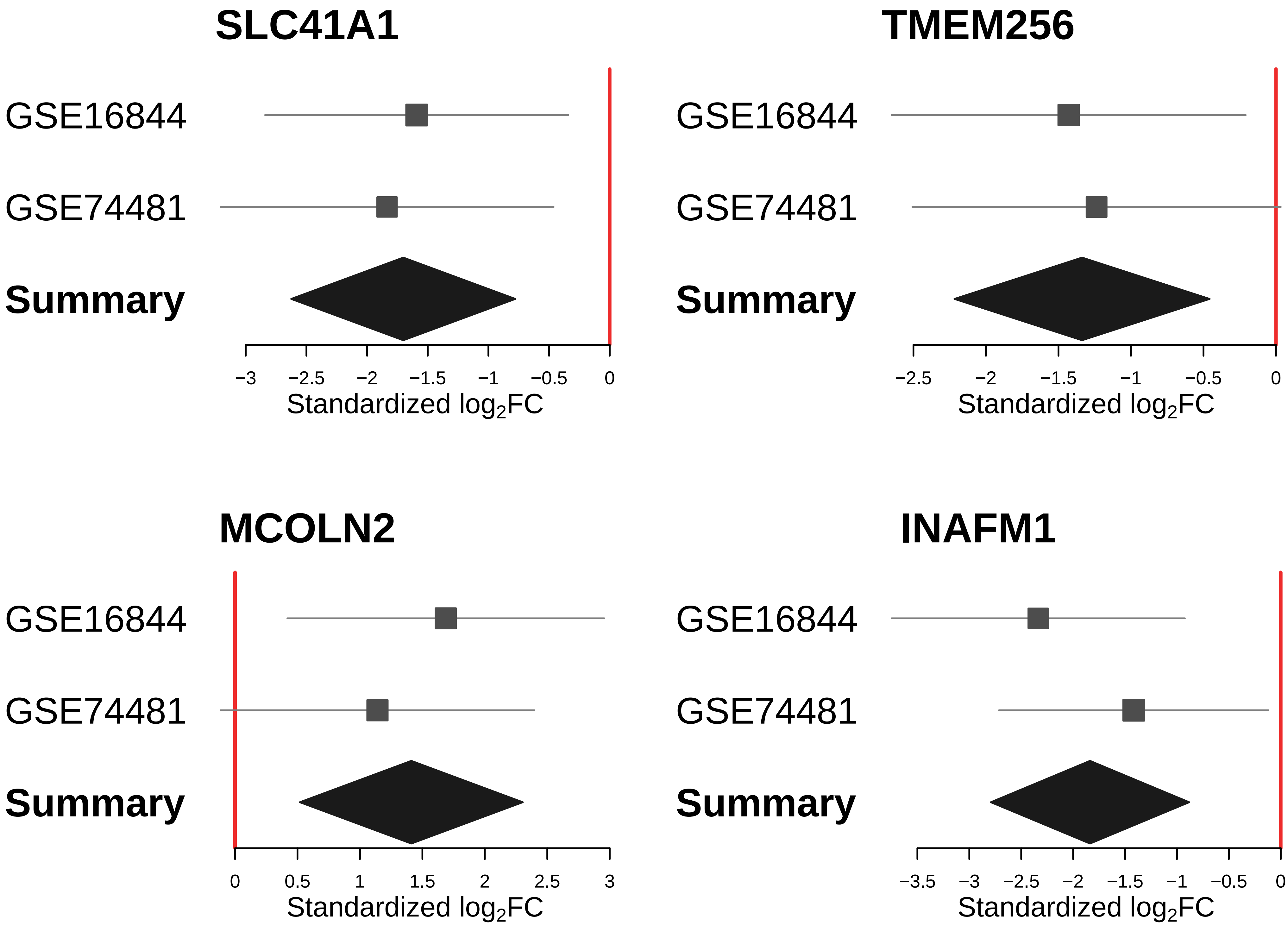

SLC41A1
GSE16844
GSE74481
Summary
−3
−2.5
−2
−1.5
−1
−0.5
0
Standardized log2FC
TMEM256
GSE16844
GSE74481
Summary
−2.5
−2
−1.5
−1
−0.5
0
Standardized log2FC
MCOLN2
GSE16844
GSE74481
Summary
0
0.5
1
1.5
2
2.5
3
Standardized log2FC
INAFM1
GSE16844
GSE74481
Summary
−3.5
−3
−2.5
−2
−1.5
−1
−0.5
0
Standardized log2FC

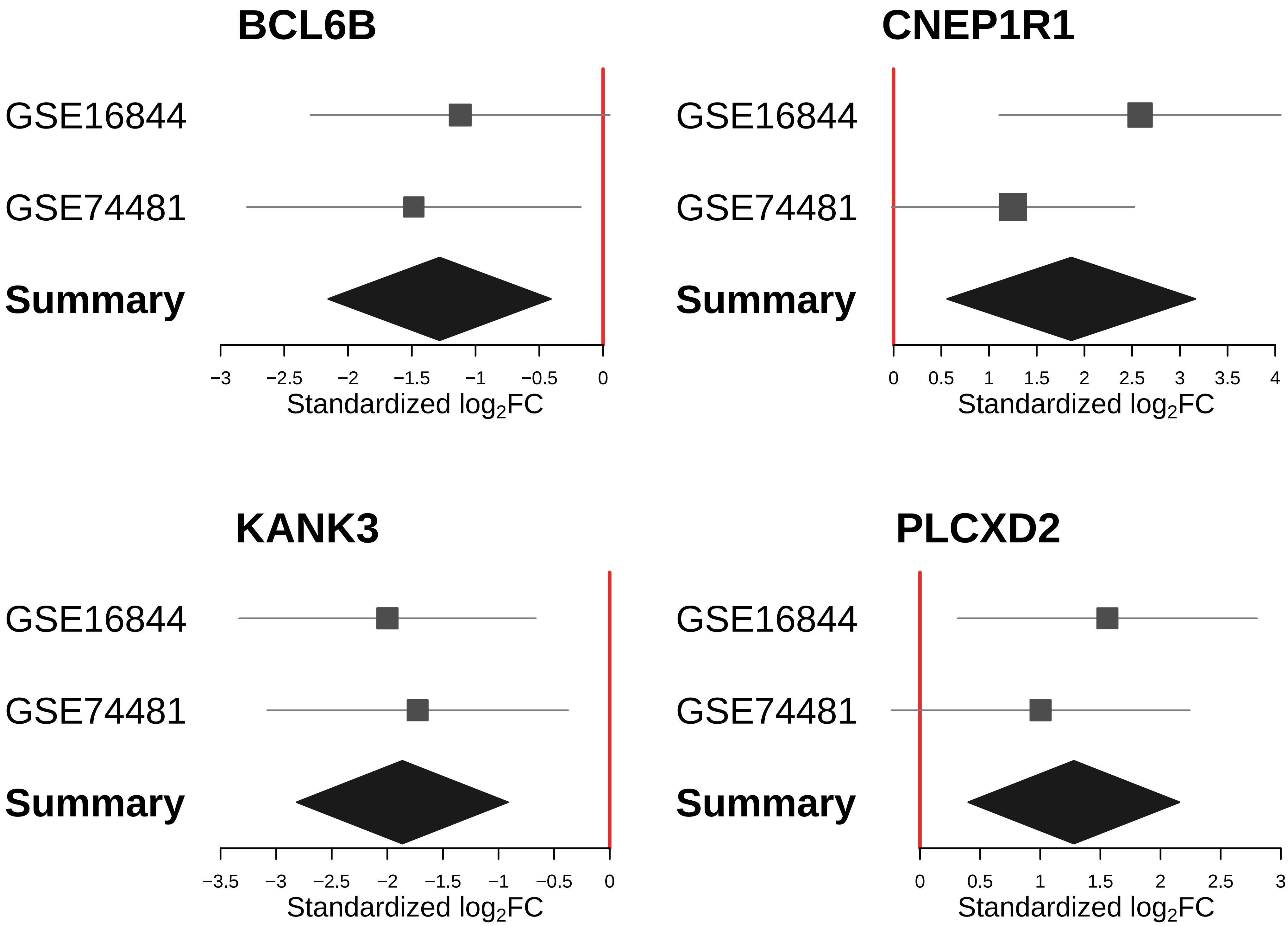

BCL6B
GSE16844
GSE74481
Summary
−3
−2.5
−2
−1.5
−1
−0.5
0
Standardized log2FC
CNEP1R1
GSE16844
GSE74481
Summary
0
0.5
1
1.5
2
2.5
3
3.5
4
Standardized log2FC
KANK3
GSE16844
GSE74481
Summary
−3.5
−3
−2.5
−2
−1.5
−1
−0.5
0
Standardized log2FC
PLCXD2
GSE16844
GSE74481
Summary
0
0.5
1
1.5
2
2.5
3
Standardized log2FC

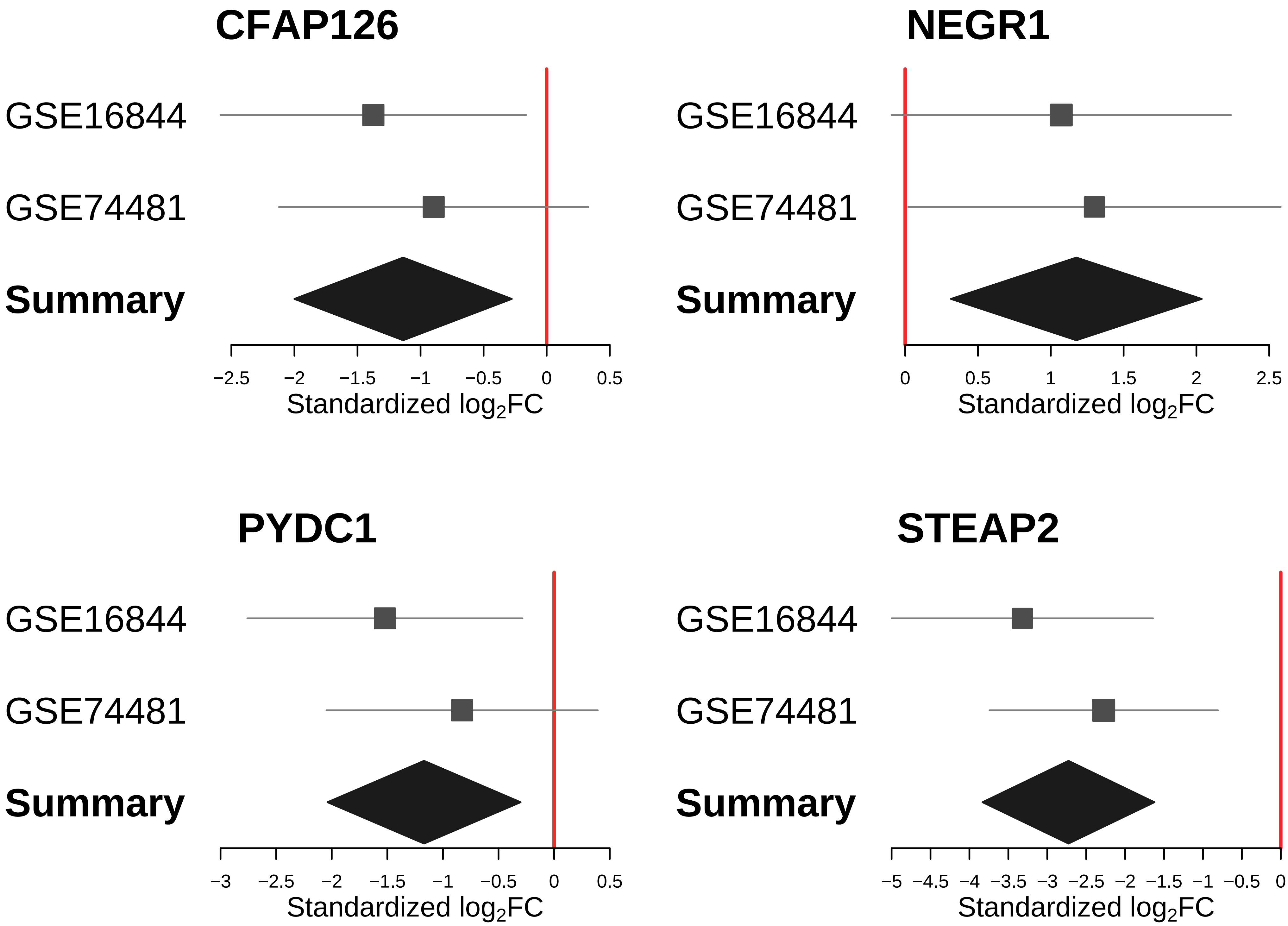
CFAP126
GSE16844
GSE74481
Summary
−2.5 −2 −1.5 −1 −0.5 0 0.5
Standardized log2FC
NEGR1
GSE16844
GSE74481
Summary
0 0.5 1 1.5 2 2.5
Standardized log2FC
PYDC1
GSE16844
GSE74481
Summary
−3 −2.5 −2 −1.5 −1 −0.5 0 0.5
Standardized log2FC
STEAP2
GSE16844
GSE74481
Summary
−5 −4.5 −4 −3.5 −3 −2.5 −2 −1.5 −1 −0.5 0
Standardized log2FC

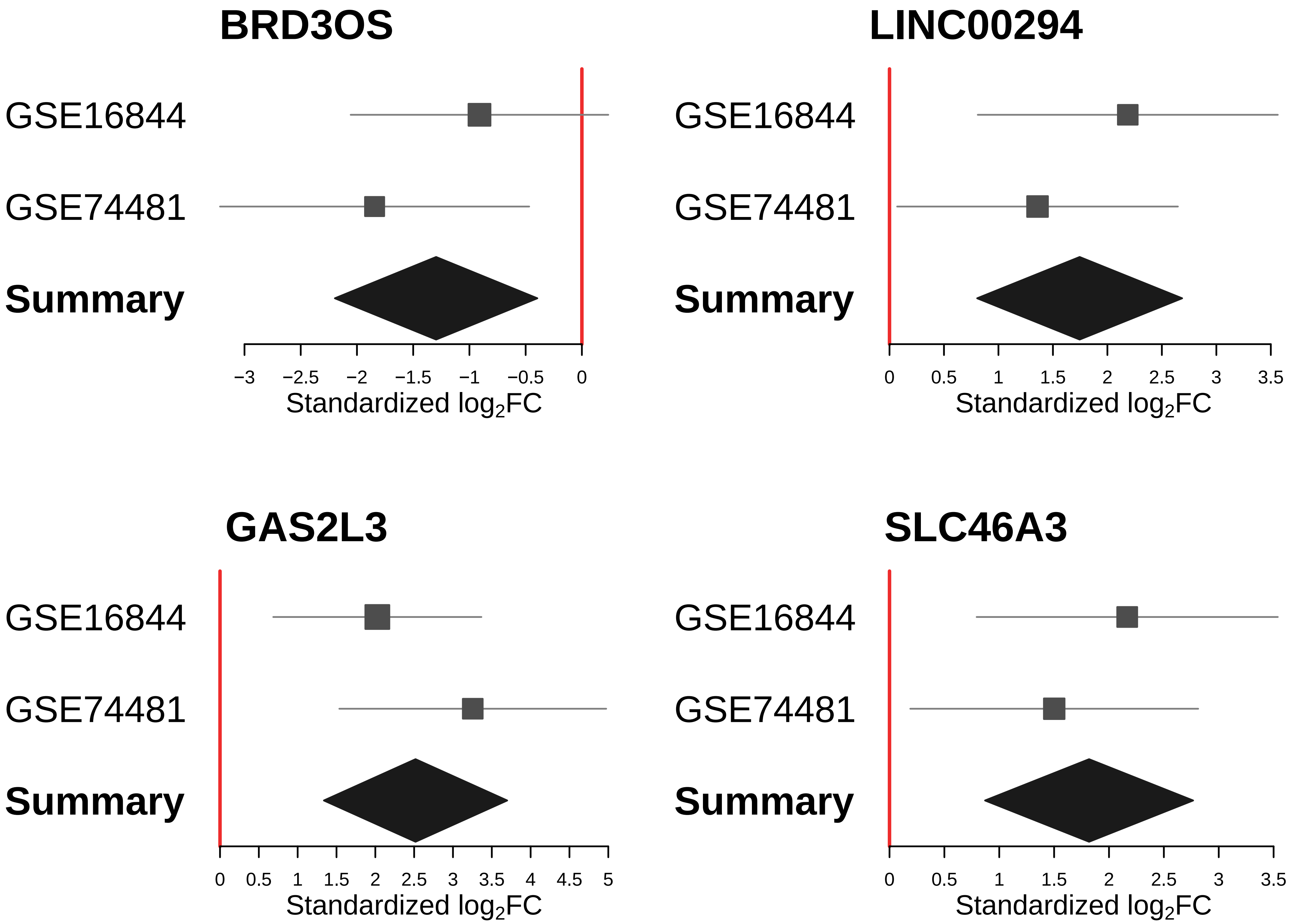
BRD3OS
GSE16844
GSE74481
Summary
−3 −2.5 −2 −1.5 −1 −0.5 0
Standardized log2FC
LINC00294
GSE16844
GSE74481
Summary
0 0.5 1 1.5 2 2.5 3 3.5
Standardized log2FC
GAS2L3
GSE16844
GSE74481
Summary
0 0.5 1 1.5 2 2.5 3 3.5 4 4.5 5
Standardized log2FC
SLC46A3
GSE16844
GSE74481
Summary
0 0.5 1 1.5 2 2.5 3 3.5
Standardized log2FC

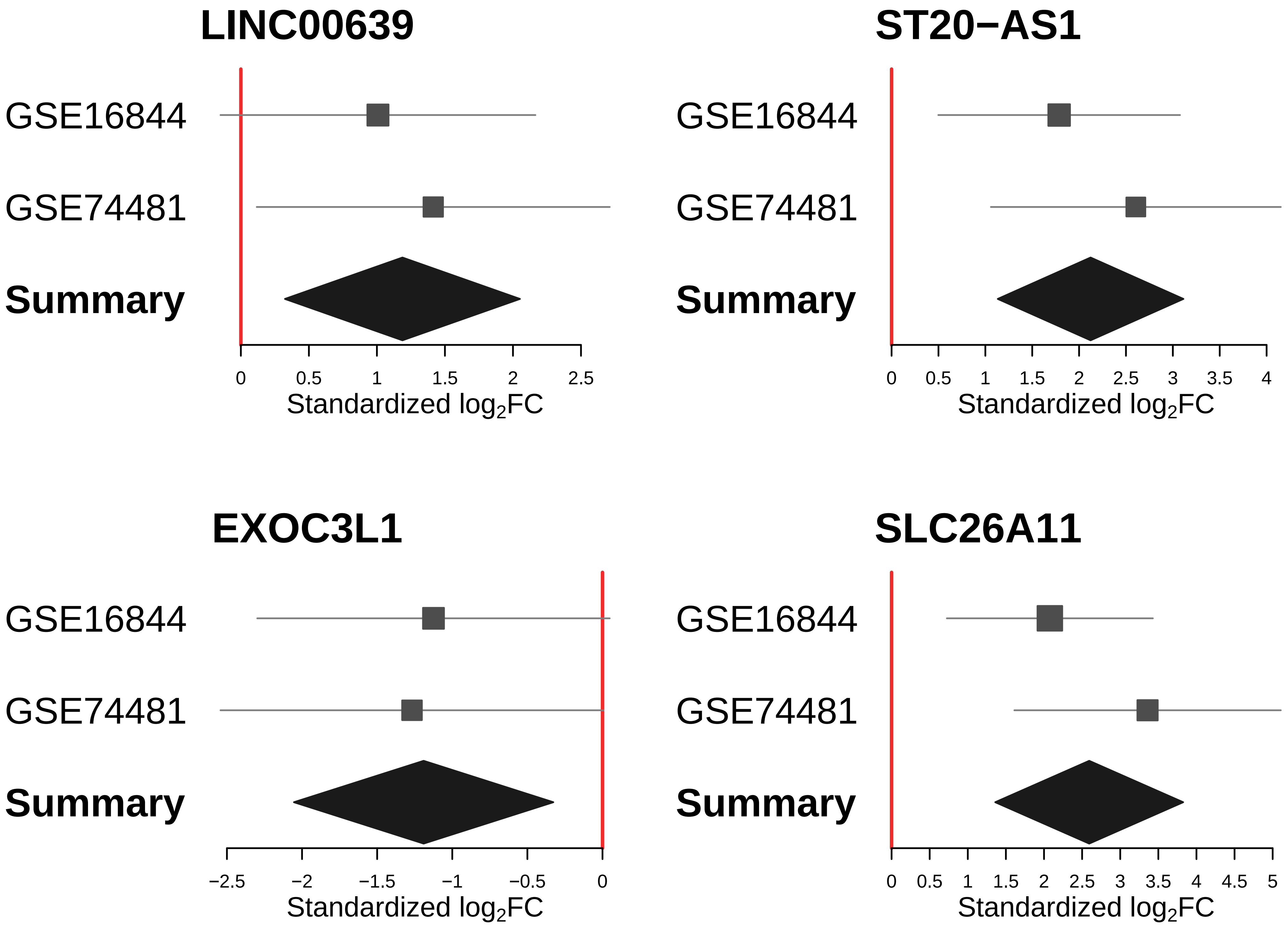
LINC00639
GSE16844
GSE74481
Summary
0
0.5
1
1.5
2
2.5
Standardized log2FC
ST20−AS1
GSE16844
GSE74481
Summary
0
0.5
1
1.5
2
2.5
3
3.5
4
Standardized log2FC
EXOC3L1
GSE16844
GSE74481
Summary
−2.5
−2
−1.5
−1
−0.5
0
Standardized log2FC
SLC26A11
GSE16844
GSE74481
Summary
0
0.5
1
1.5
2
2.5
3
3.5
4
4.5
5
Standardized log2FC

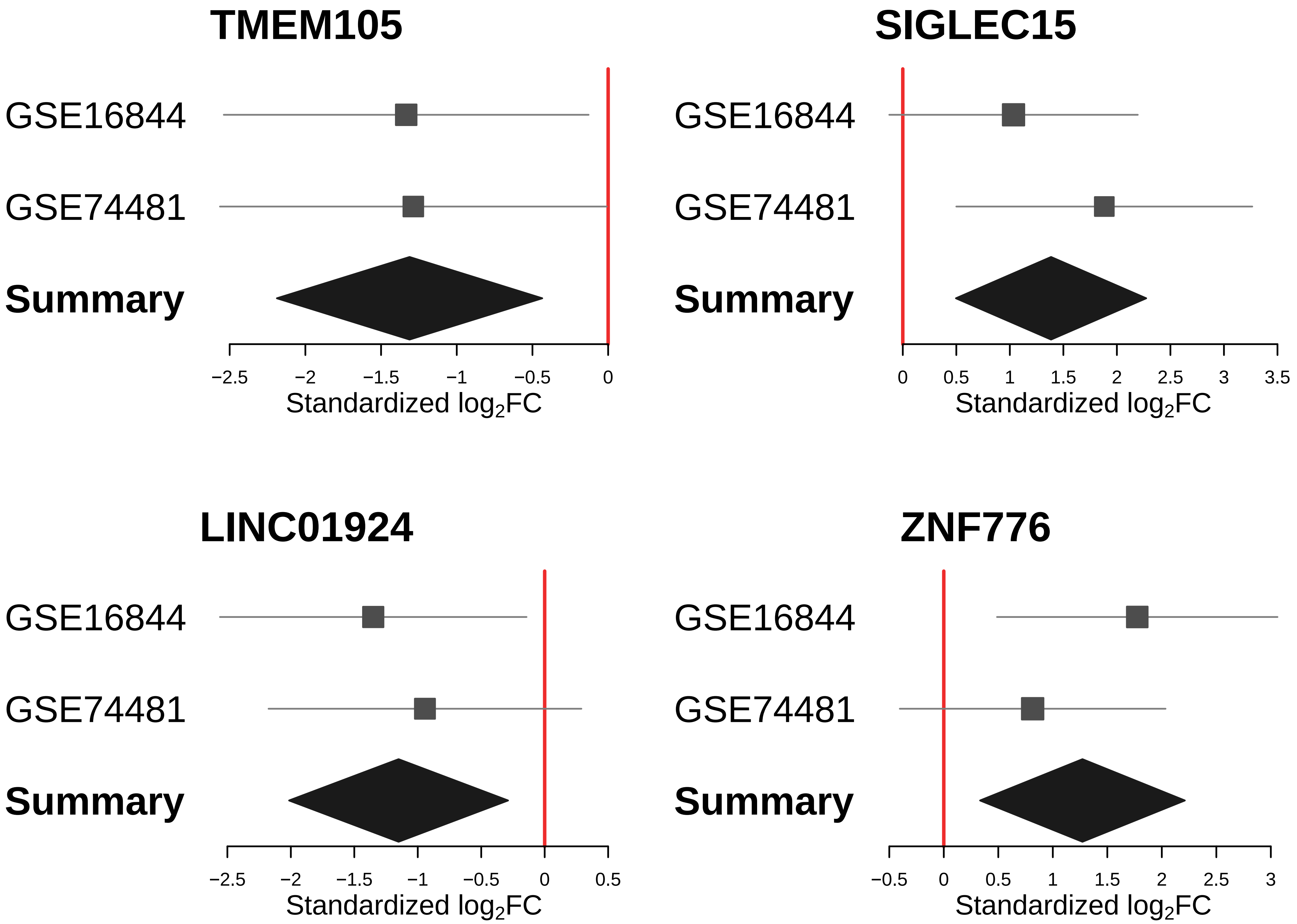
TMEM105
GSE16844
GSE74481
Summary
−2.5
−2
−1.5
−1
−0.5
0
Standardized log2FC
SIGLEC15
GSE16844
GSE74481
Summary
0
0.5
1
1.5
2
2.5
3
3.5
Standardized log2FC
LINC01924
GSE16844
GSE74481
Summary
−2.5
−2
−1.5
−1
−0.5
0
0.5
Standardized log2FC
ZNF776
GSE16844
GSE74481
Summary
−0.5
0
0.5
1
1.5
2
2.5
3
Standardized log2FC

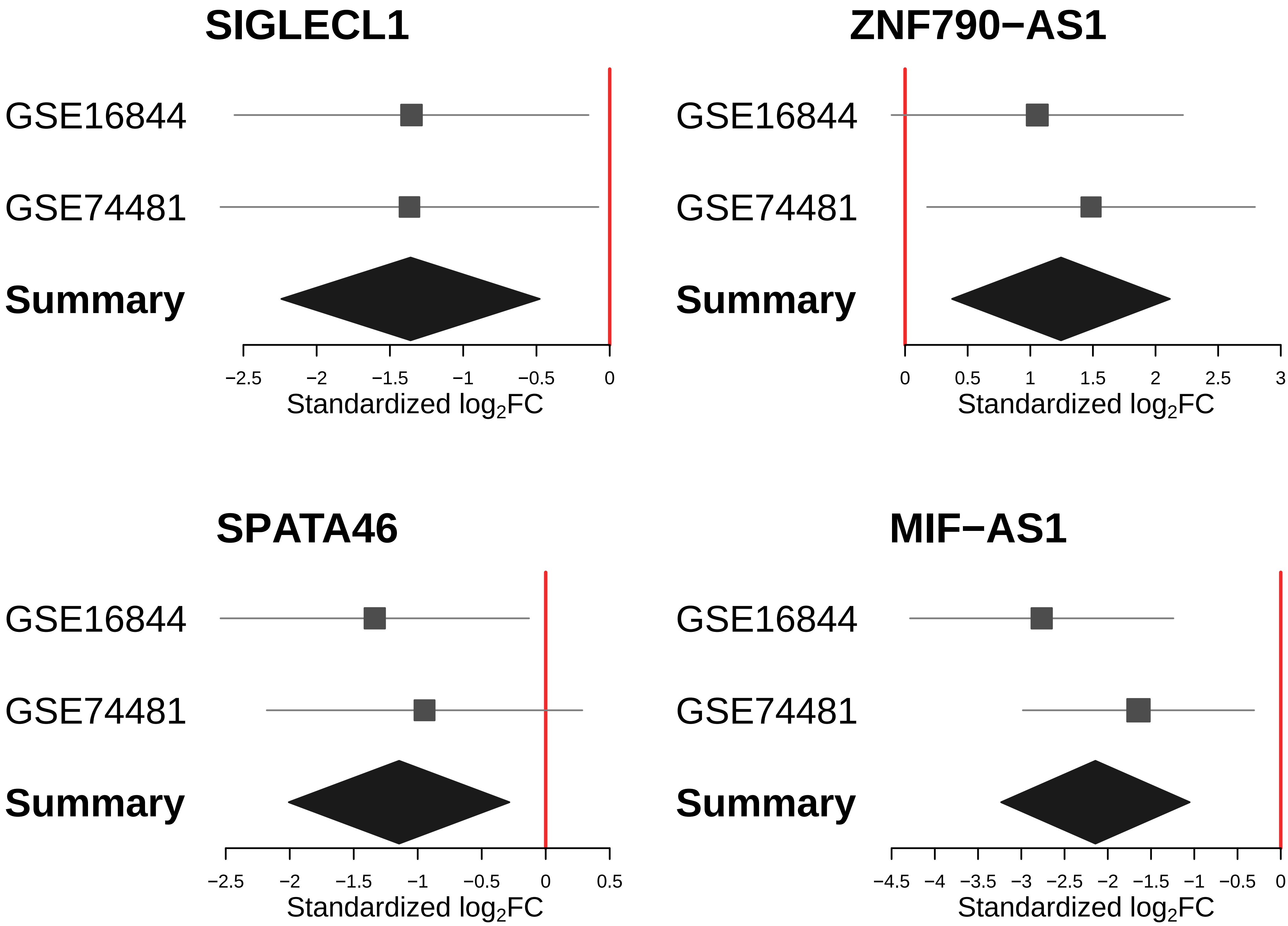
SIGLECL1
GSE16844
GSE74481
Summary
−2.5 −2 −1.5 −1 −0.5 0
Standardized log2FC
ZNF790−AS1
GSE16844
GSE74481
Summary
0 0.5 1 1.5 2 2.5 3
Standardized log2FC
SPATA46
GSE16844
GSE74481
Summary
−2.5 −2 −1.5 −1 −0.5 0 0.5
Standardized log2FC
MIF−AS1
GSE16844
GSE74481
Summary
−4.5 −4 −3.5 −3 −2.5 −2 −1.5 −1 −0.5 0
Standardized log2FC

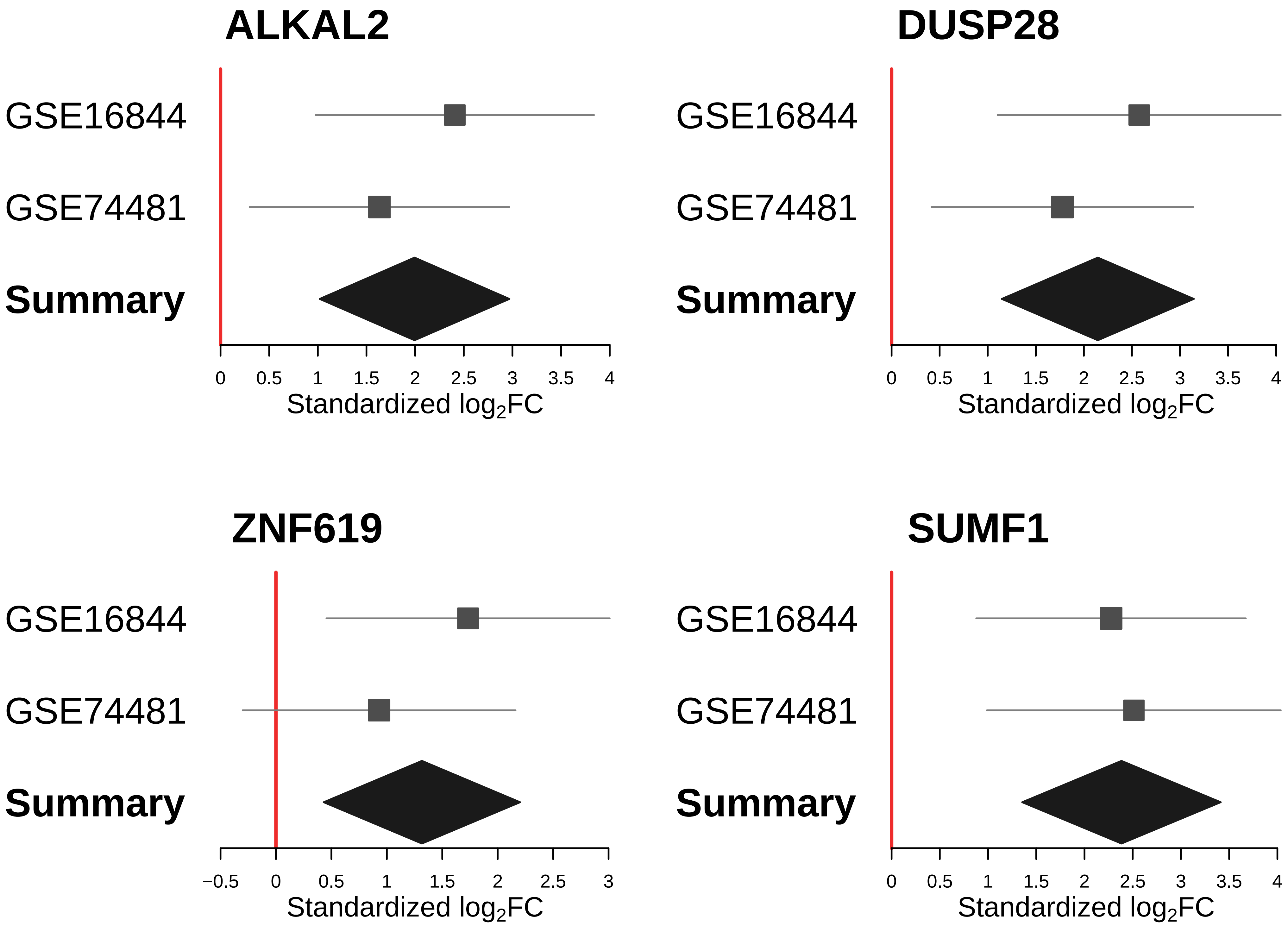
ALKAL2
DUSP28
ZNF619
SUMF1
GSE16844
GSE74481
Summary
Standardized log2FC
0 0.5 1 1.5 2 2.5 3 3.5 4
−0.5 0 0.5 1 1.5 2 2.5 3

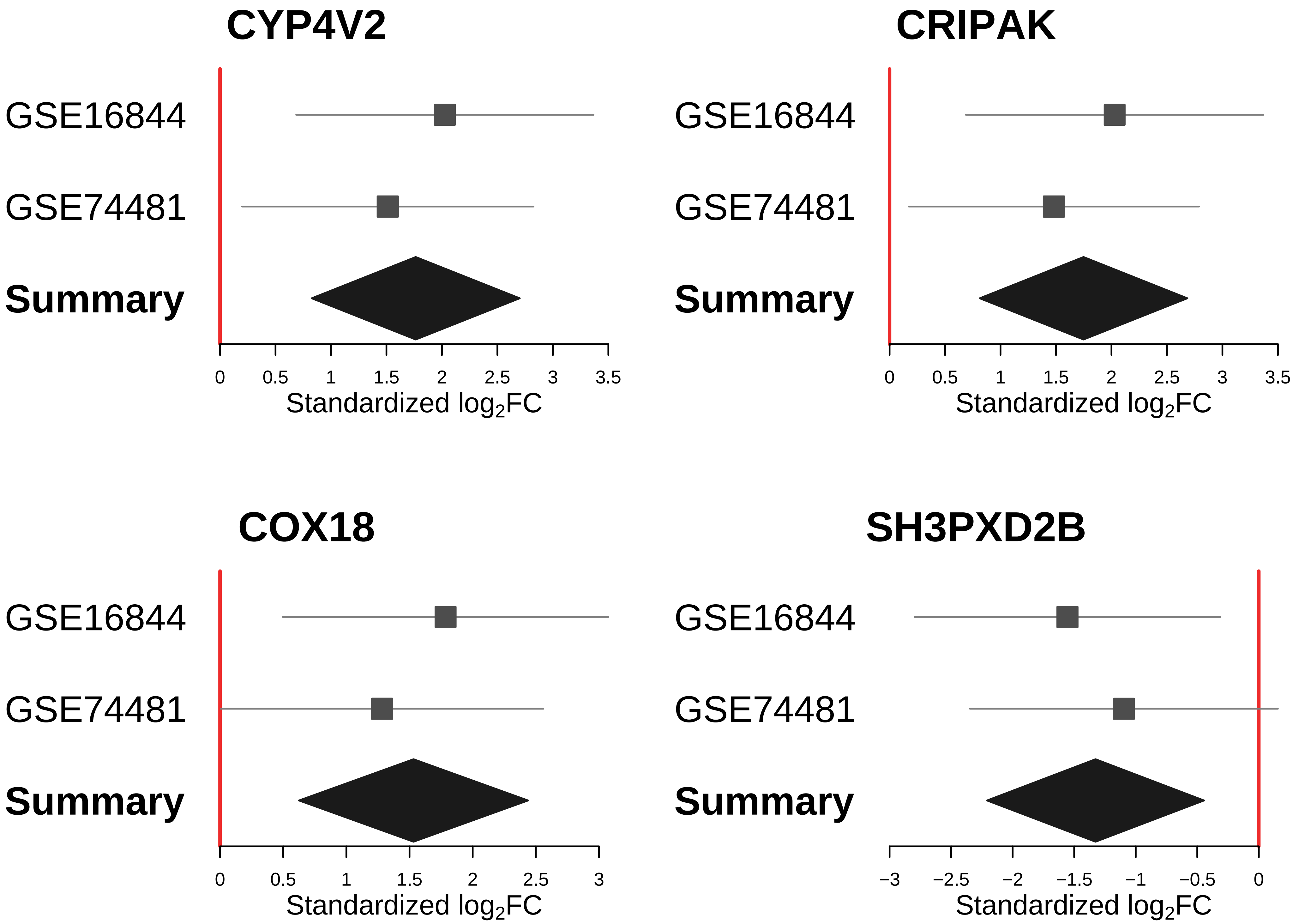
CYP4V2
CRIPAK
COX18
SH3PXD2B
GSE16844
GSE74481
Summary
Standardized log2FC
0
0.5
1
1.5
2
2.5
3
3.5
−3
−2.5
−2
−1.5
−1
−0.5

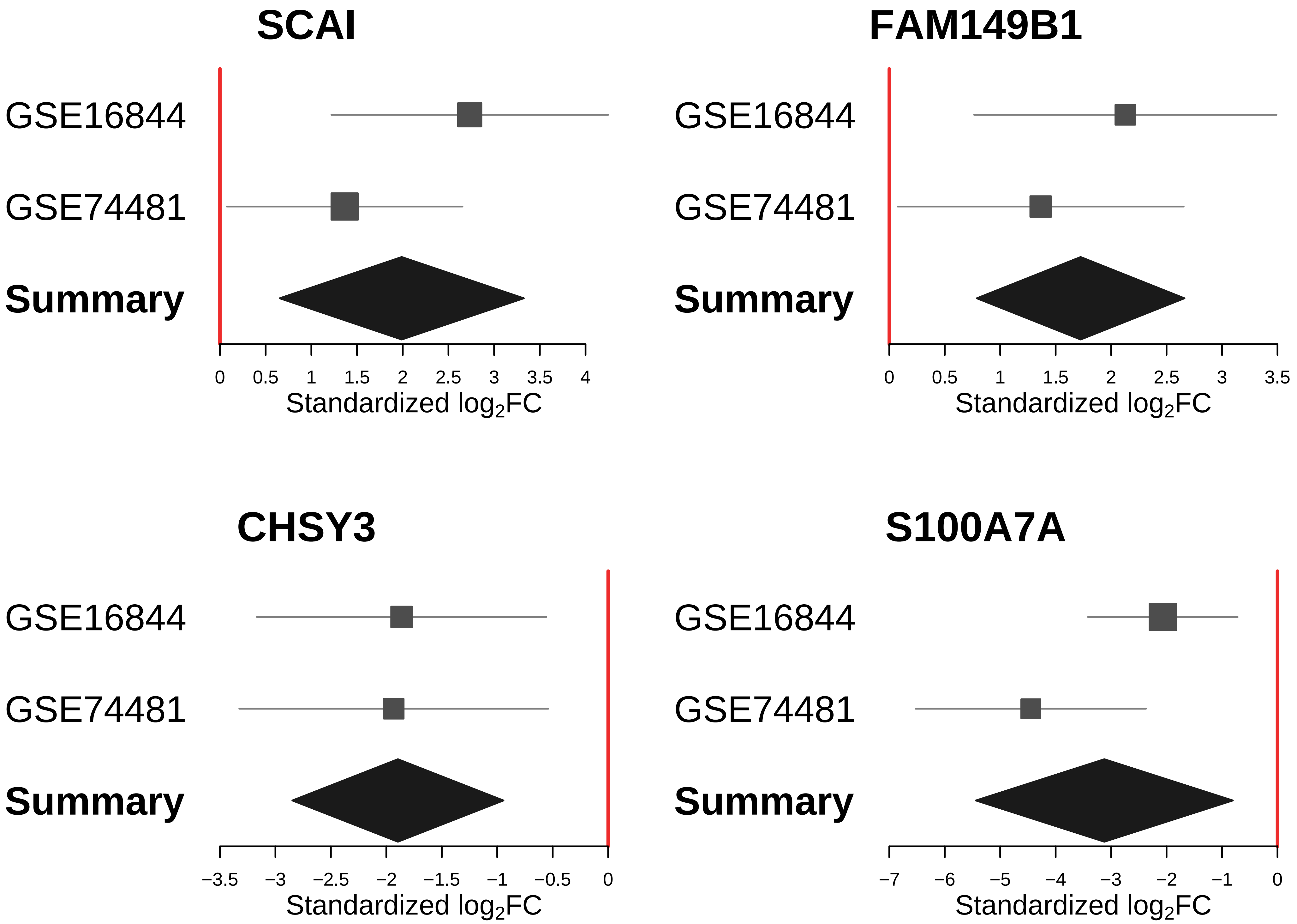
SCAI
GSE16844
GSE74481
Summary
0
0.5
1
1.5
2
2.5
3
3.5
4
Standardized log2FC
FAM149B1
GSE16844
GSE74481
Summary
0
0.5
1
1.5
2
2.5
3
3.5
Standardized log2FC
CHSY3
GSE16844
GSE74481
Summary
−3.5
−3
−2.5
−2
−1.5
−1
−0.5
0
Standardized log2FC
S100A7A
GSE16844
GSE74481
Summary
−7
−6
−5
−4
−3
−2
−1
0
Standardized log2FC

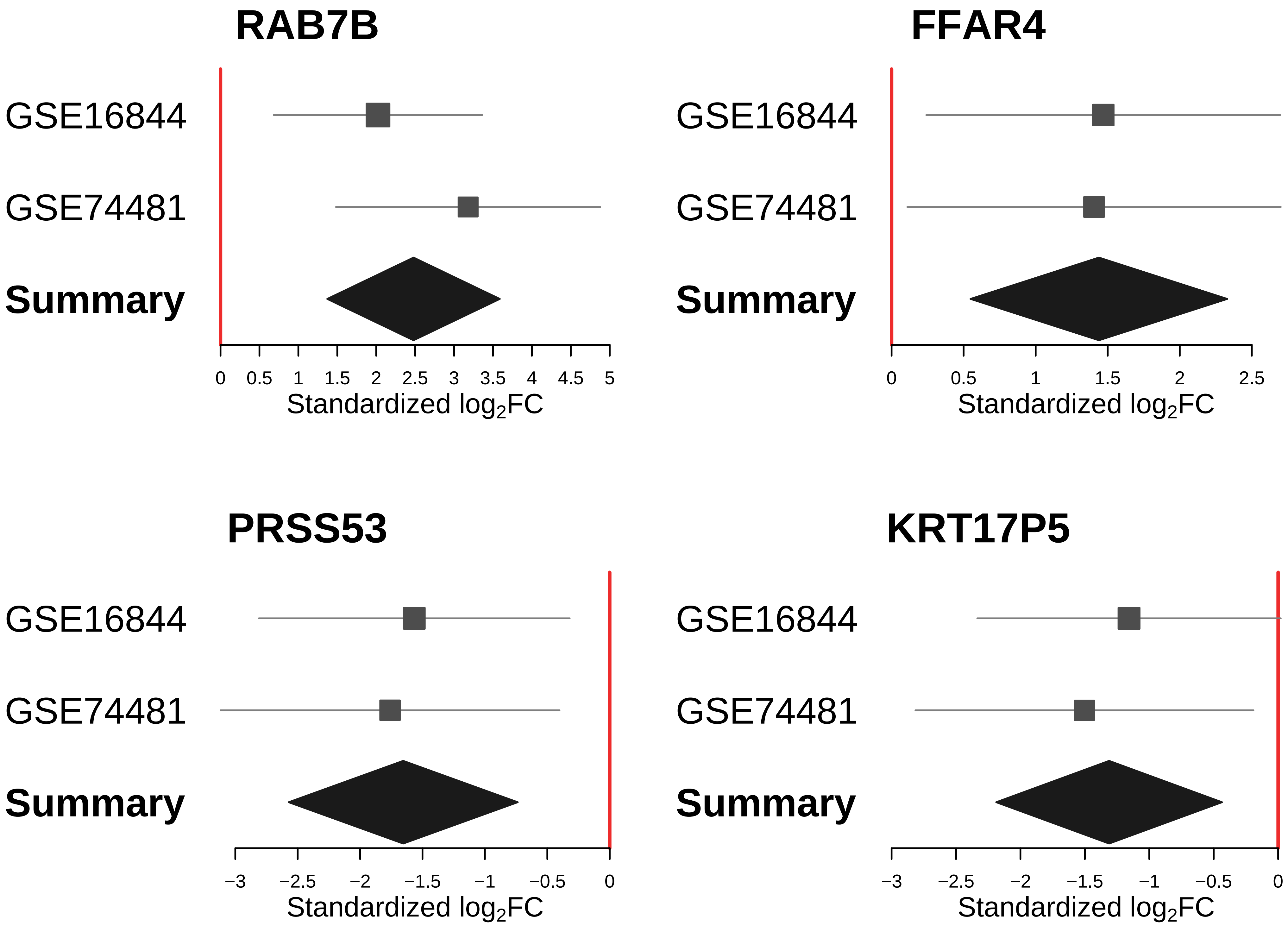
RAB7B
GSE16844
GSE74481
Summary
0 0.5 1 1.5 2 2.5 3 3.5 4 4.5 5
Standardized log2FC
FFAR4
GSE16844
GSE74481
Summary
0 0.5 1 1.5 2 2.5
Standardized log2FC
PRSS53
GSE16844
GSE74481
Summary
−3 −2.5 −2 −1.5 −1 −0.5 0
Standardized log2FC
KRT17P5
GSE16844
GSE74481
Summary
−3 −2.5 −2 −1.5 −1 −0.5 0
Standardized log2FC

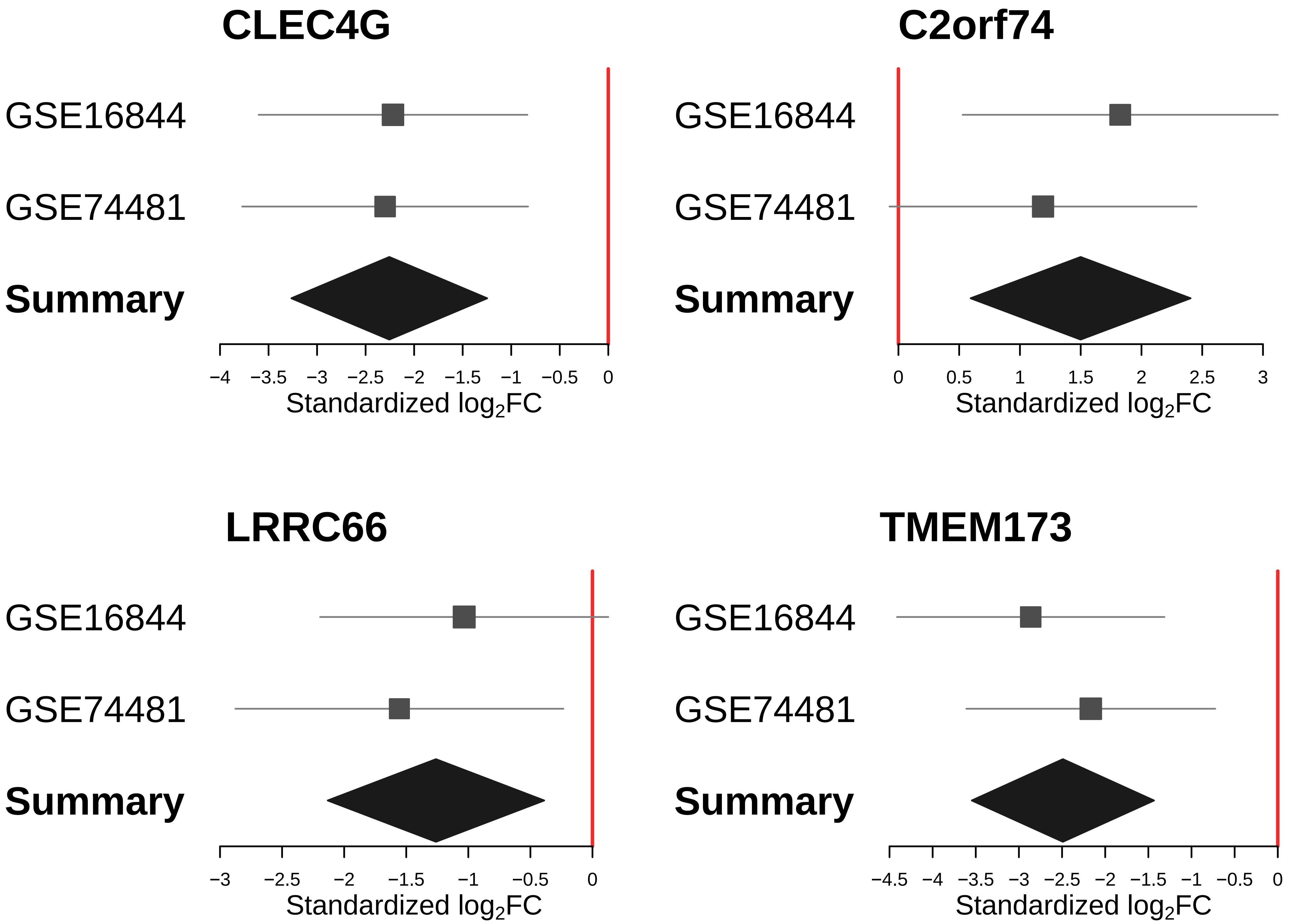

CLEC4G
GSE16844
GSE74481
Summary
−4 −3.5 −3 −2.5 −2 −1.5 −1 −0.5 0
Standardized log2FC
C2orf74
GSE16844
GSE74481
Summary
0 0.5 1 1.5 2 2.5 3
Standardized log2FC
LRRC66
GSE16844
GSE74481
Summary
−3 −2.5 −2 −1.5 −1 −0.5 0
Standardized log2FC
TMEM173
GSE16844
GSE74481
Summary
−4.5 −4 −3.5 −3 −2.5 −2 −1.5 −1 −0.5 0
Standardized log2FC

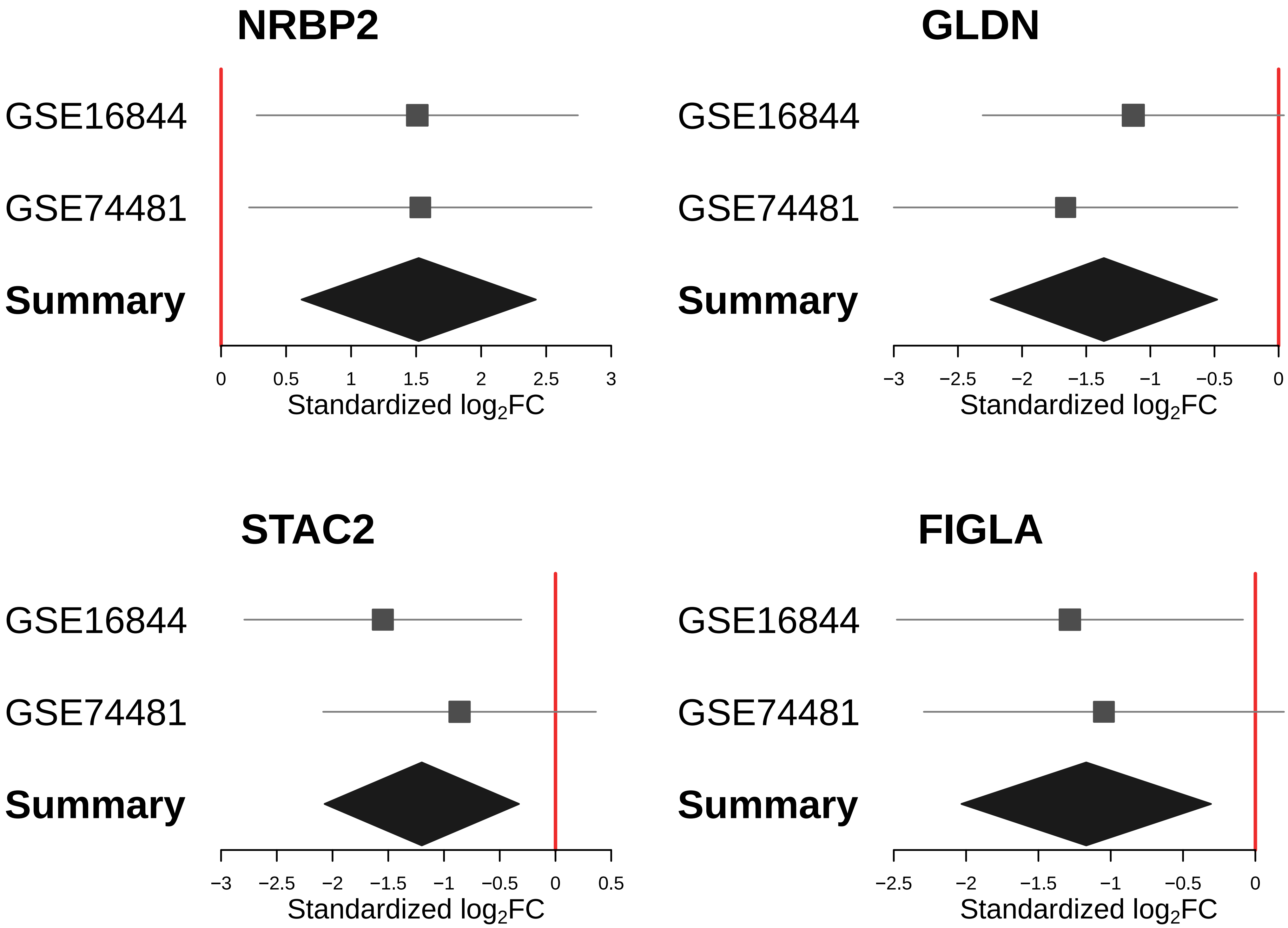
NRBP2
GSE16844
GSE74481
Summary
0 0.5 1 1.5 2 2.5 3
Standardized log2FC
GLDN
GSE16844
GSE74481
Summary
−3 −2.5 −2 −1.5 −1 −0.5 0
Standardized log2FC
STAC2
GSE16844
GSE74481
Summary
−3 −2.5 −2 −1.5 −1 −0.5 0 0.5
Standardized log2FC
FIGLA
GSE16844
GSE74481
Summary
−2.5 −2 −1.5 −1 −0.5 0
Standardized log2FC

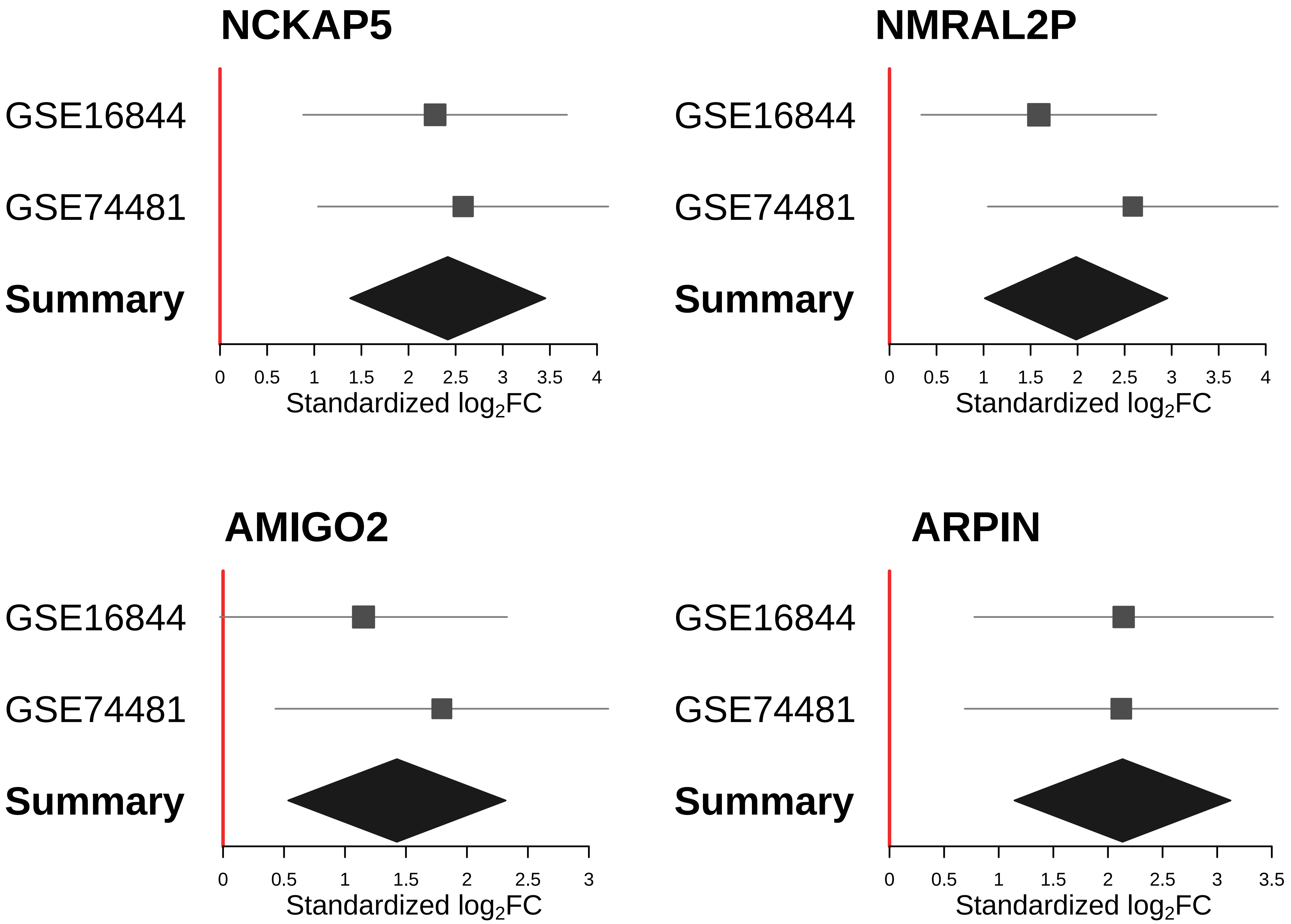

NCKAP5
GSE16844
GSE74481
Summary
0 0.5 1 1.5 2 2.5 3 3.5 4
Standardized log2FC
NMRAL2P
GSE16844
GSE74481
Summary
0 0.5 1 1.5 2 2.5 3 3.5 4
Standardized log2FC
AMIGO2
GSE16844
GSE74481
Summary
0 0.5 1 1.5 2 2.5 3
Standardized log2FC
ARPIN
GSE16844
GSE74481
Summary
0 0.5 1 1.5 2 2.5 3 3.5
Standardized log2FC

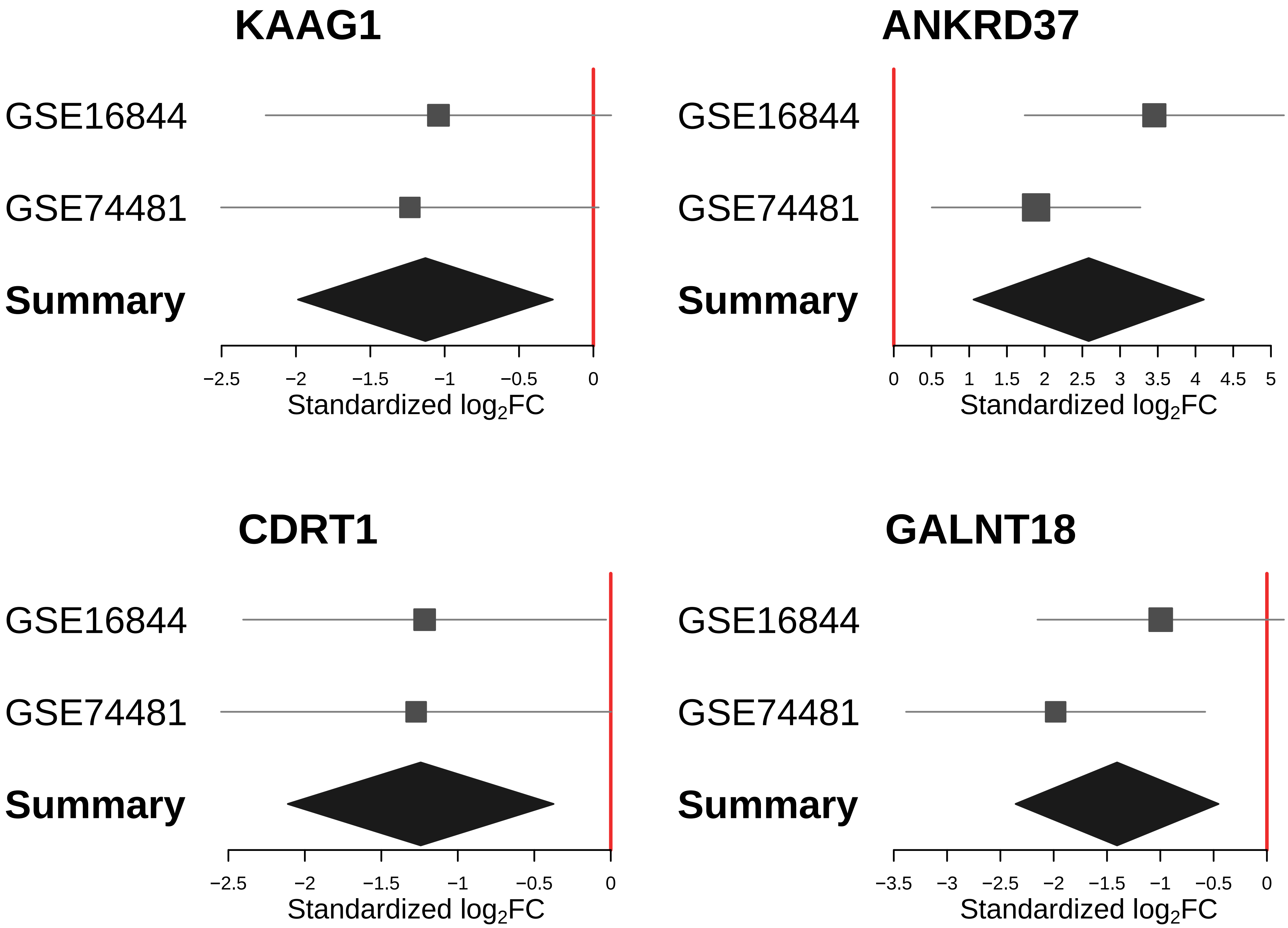
KAAG1
GSE16844
GSE74481
Summary
−2.5 −2 −1.5 −1 −0.5 0
Standardized log2FC
ANKRD37
GSE16844
GSE74481
Summary
0 0.5 1 1.5 2 2.5 3 3.5 4 4.5 5
Standardized log2FC
CDRT1
GSE16844
GSE74481
Summary
−2.5 −2 −1.5 −1 −0.5 0
Standardized log2FC
GALNT18
GSE16844
GSE74481
Summary
−3.5 −3 −2.5 −2 −1.5 −1 −0.5 0
Standardized log2FC

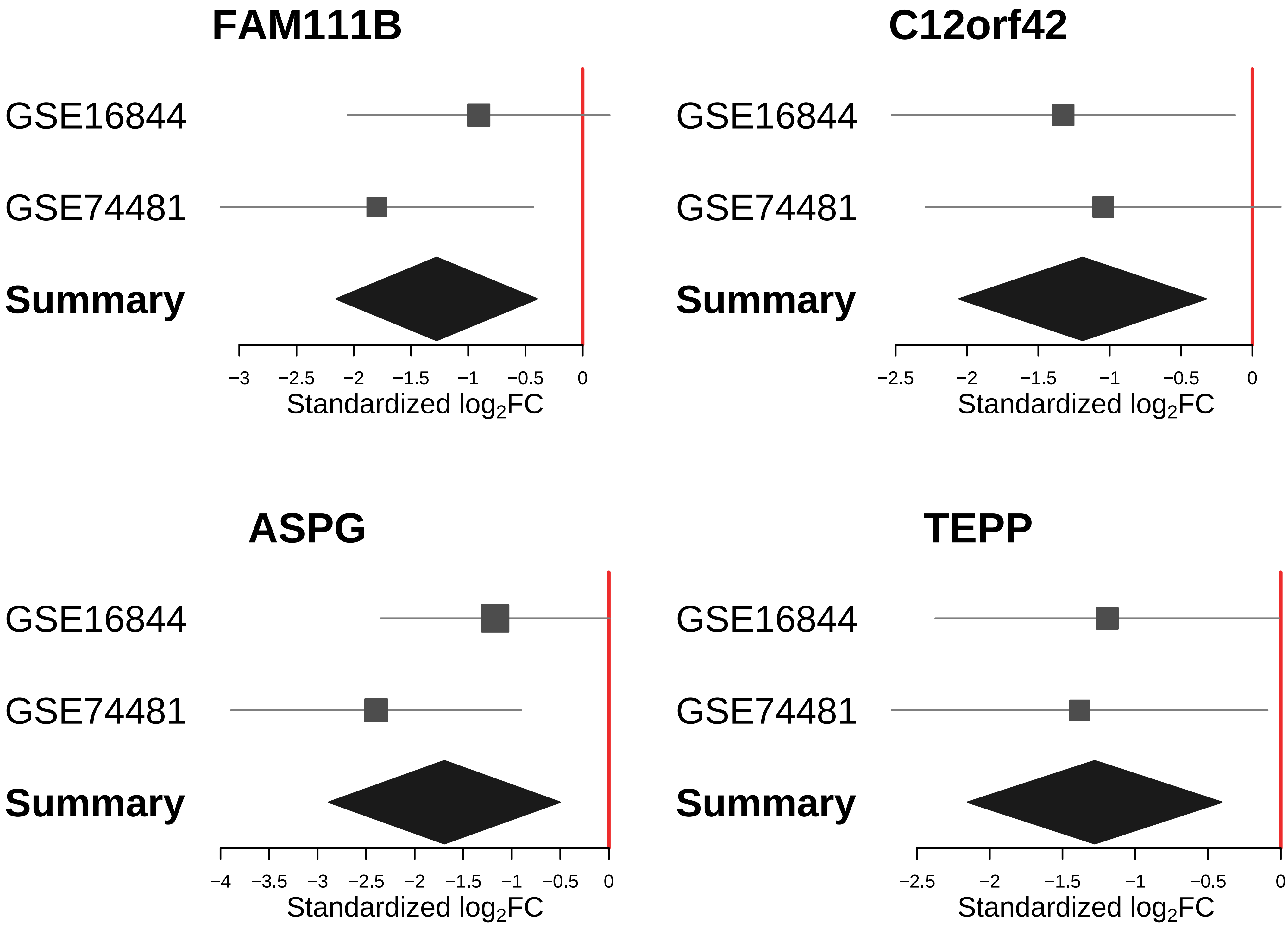
FAM111B
GSE16844
GSE74481
Summary
−3 −2.5 −2 −1.5 −1 −0.5 0
Standardized log2FC
C12orf42
GSE16844
GSE74481
Summary
−2.5 −2 −1.5 −1 −0.5 0
Standardized log2FC
ASPG
GSE16844
GSE74481
Summary
−4 −3.5 −3 −2.5 −2 −1.5 −1 −0.5 0
Standardized log2FC
TEPP
GSE16844
GSE74481
Summary
−2.5 −2 −1.5 −1 −0.5 0
Standardized log2FC

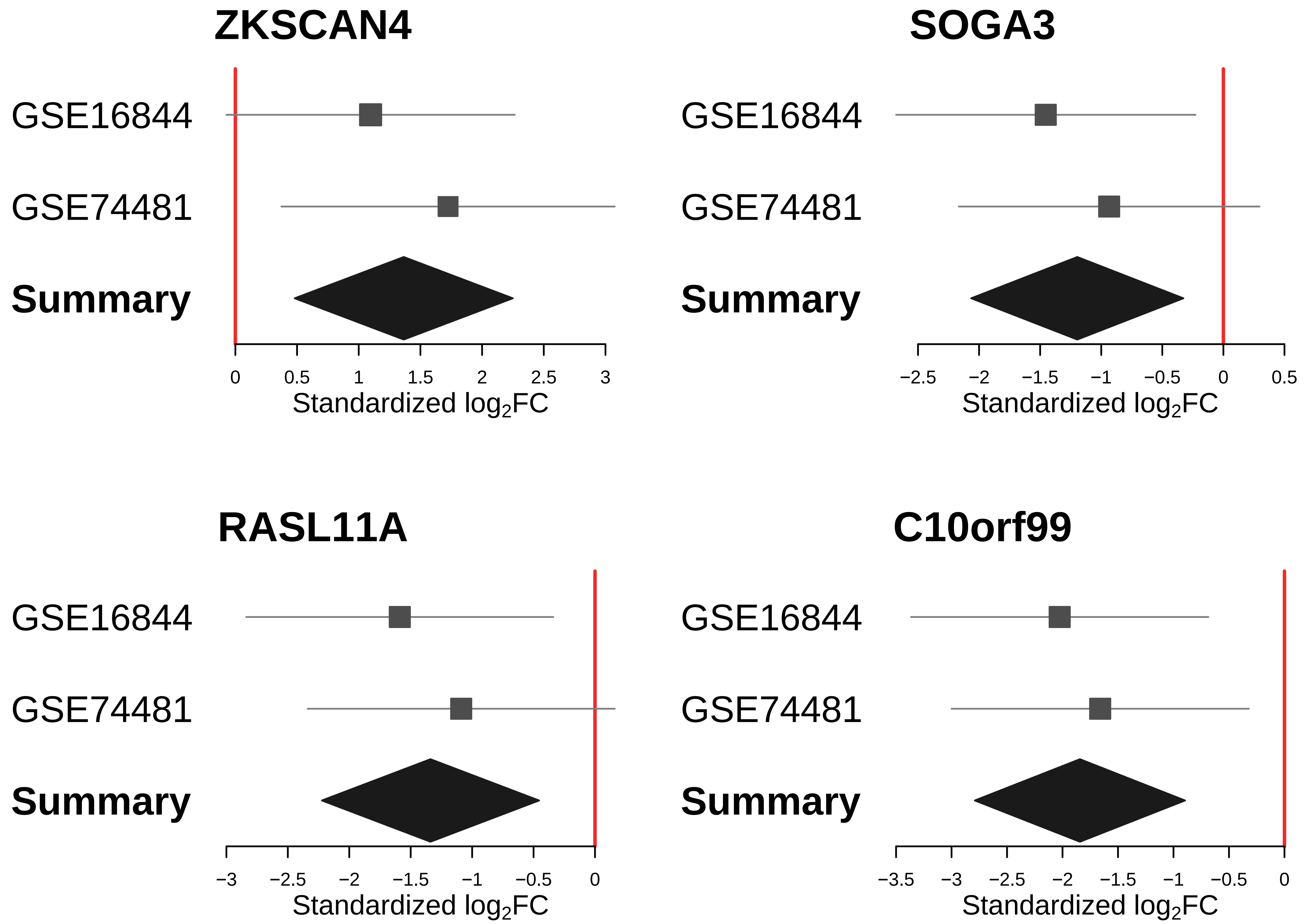

ZKSCAN4
SOGA3
RASL11A
C10orf99
GSE16844
GSE74481
Summary
0
0.5
1
1.5
2
2.5
3
−2.5
−2
−1.5
−1
−0.5
0
0.5
−3
−2.5
−2
−1.5
−1
−0.5
0
−3.5
−3
−2.5
−2
−1.5
−1
−0.5
0
Standardized log2FC

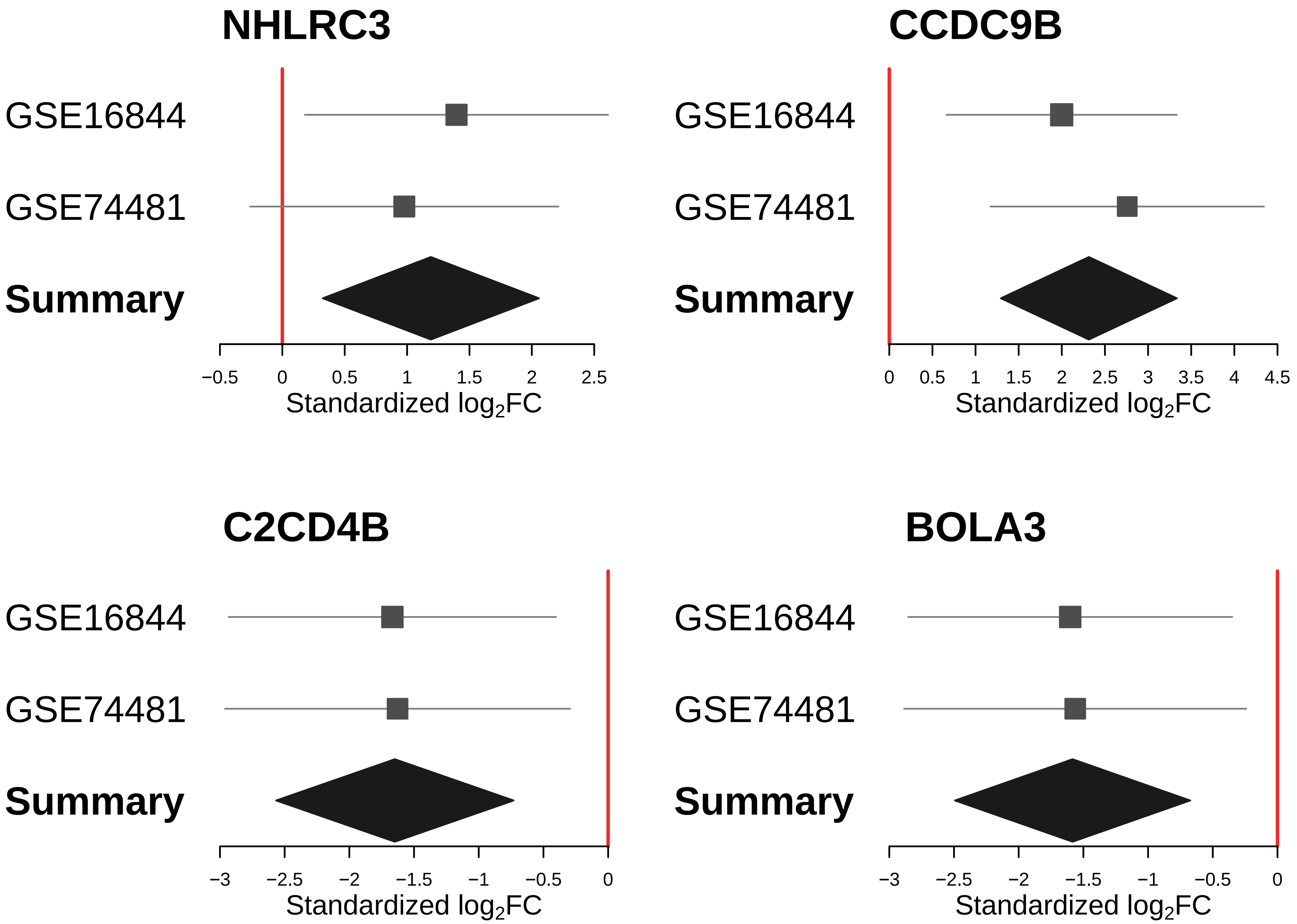
NHLRC3
GSE16844
GSE74481
Summary
−0.5 0 0.5 1 1.5 2 2.5
Standardized log2FC
CCDC9B
GSE16844
GSE74481
Summary
0 0.5 1 1.5 2 2.5 3 3.5 4 4.5
Standardized log2FC
C2CD4B
GSE16844
GSE74481
Summary
−3 −2.5 −2 −1.5 −1 −0.5 0
Standardized log2FC
BOLA3
GSE16844
GSE74481
Summary
−3 −2.5 −2 −1.5 −1 −0.5 0
Standardized log2FC

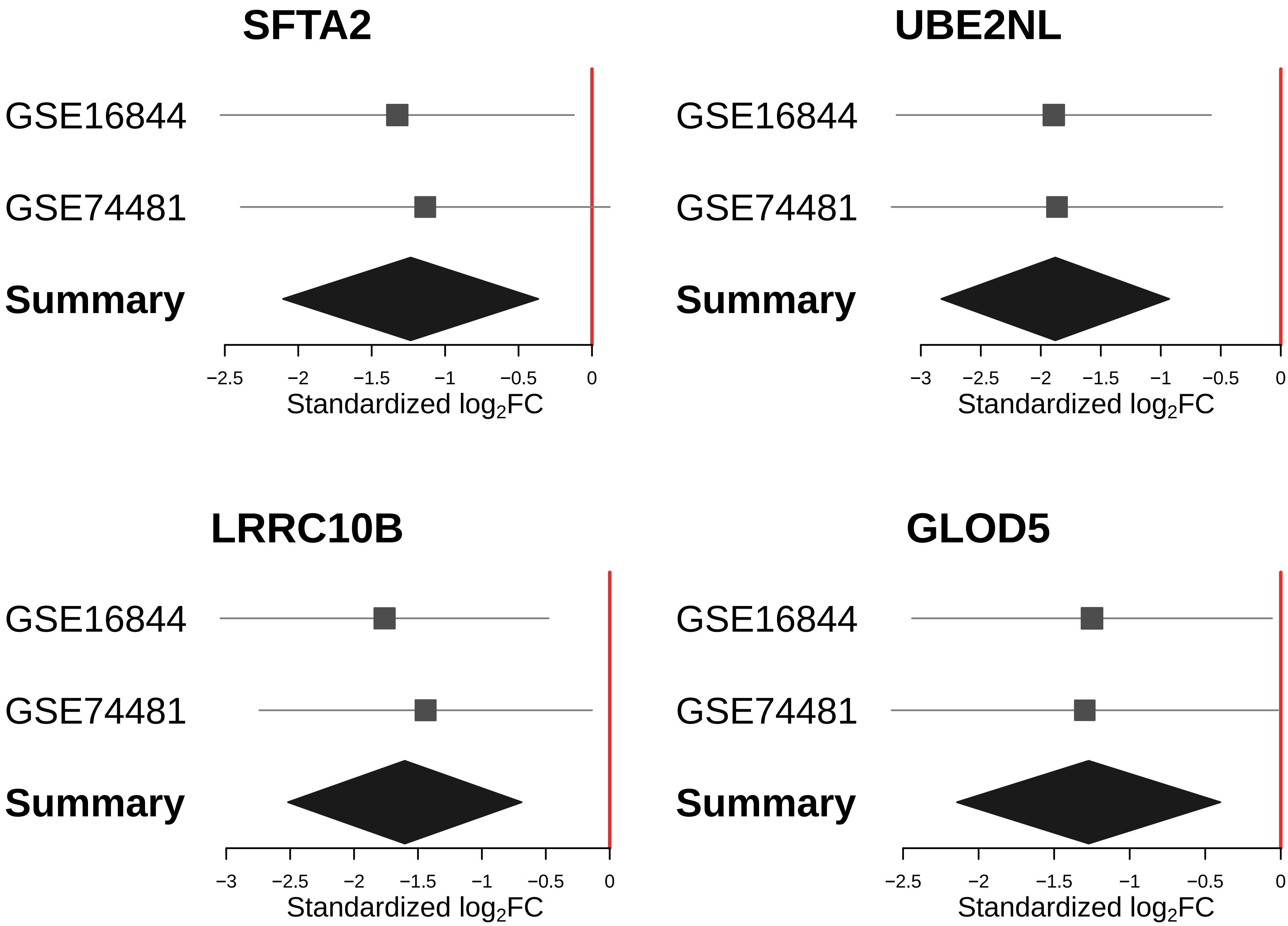
SFTA2
GSE16844
GSE74481
Summary
−2.5 −2 −1.5 −1 −0.5 0
Standardized log2FC
UBE2NL
GSE16844
GSE74481
Summary
−3 −2.5 −2 −1.5 −1 −0.5 0
Standardized log2FC
LRRC10B
GSE16844
GSE74481
Summary
−3 −2.5 −2 −1.5 −1 −0.5 0
Standardized log2FC
GLOD5
GSE16844
GSE74481
Summary
−2.5 −2 −1.5 −1 −0.5 0
Standardized log2FC

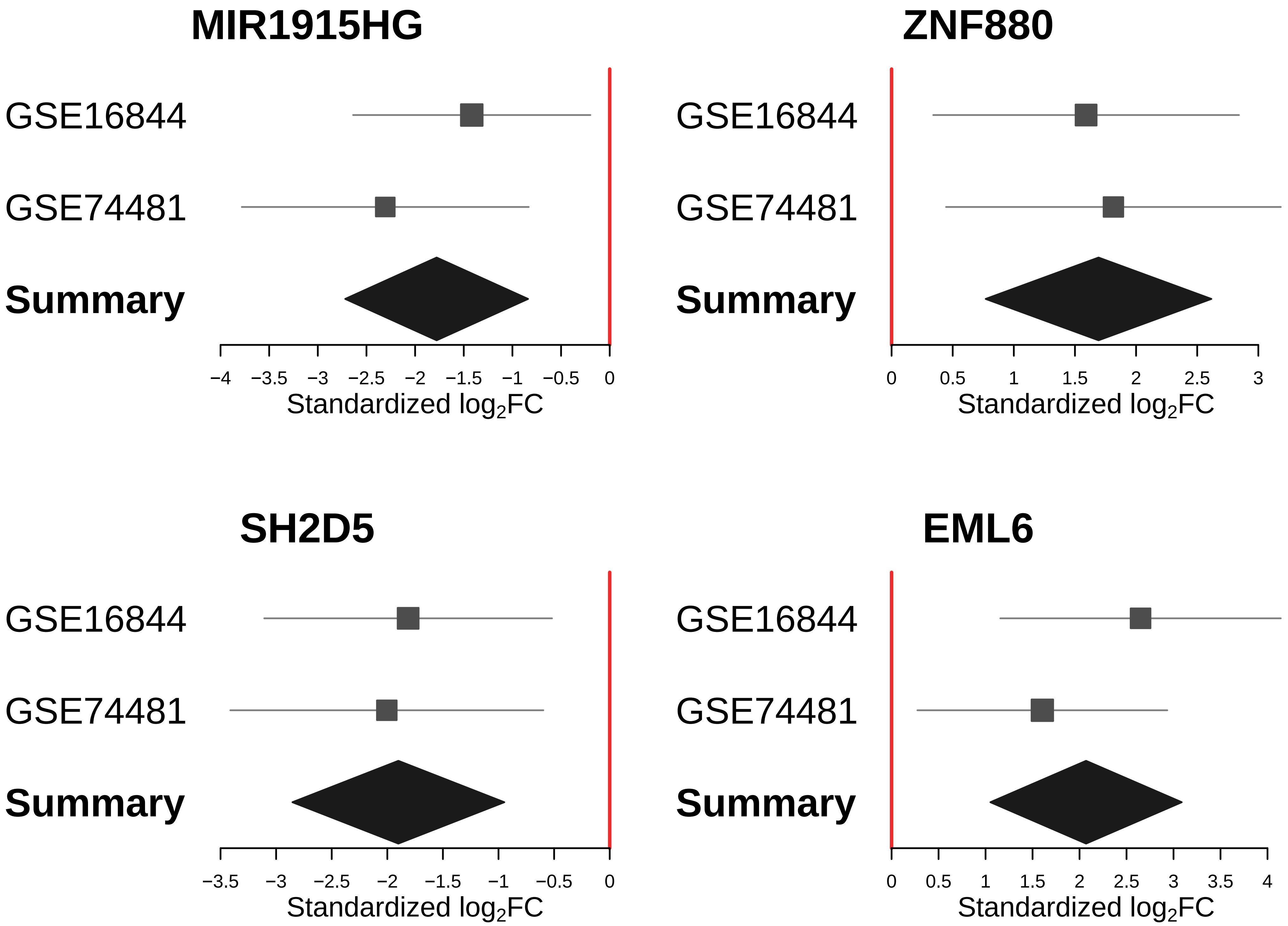
MIR1915HG
GSE16844
GSE74481
Summary
−4 −3.5 −3 −2.5 −2 −1.5 −1 −0.5 0
Standardized log2FC
ZNF880
GSE16844
GSE74481
Summary
0 0.5 1 1.5 2 2.5 3
Standardized log2FC
SH2D5
GSE16844
GSE74481
Summary
−3.5 −3 −2.5 −2 −1.5 −1 −0.5 0
Standardized log2FC
EML6
GSE16844
GSE74481
Summary
0 0.5 1 1.5 2 2.5 3 3.5 4
Standardized log2FC

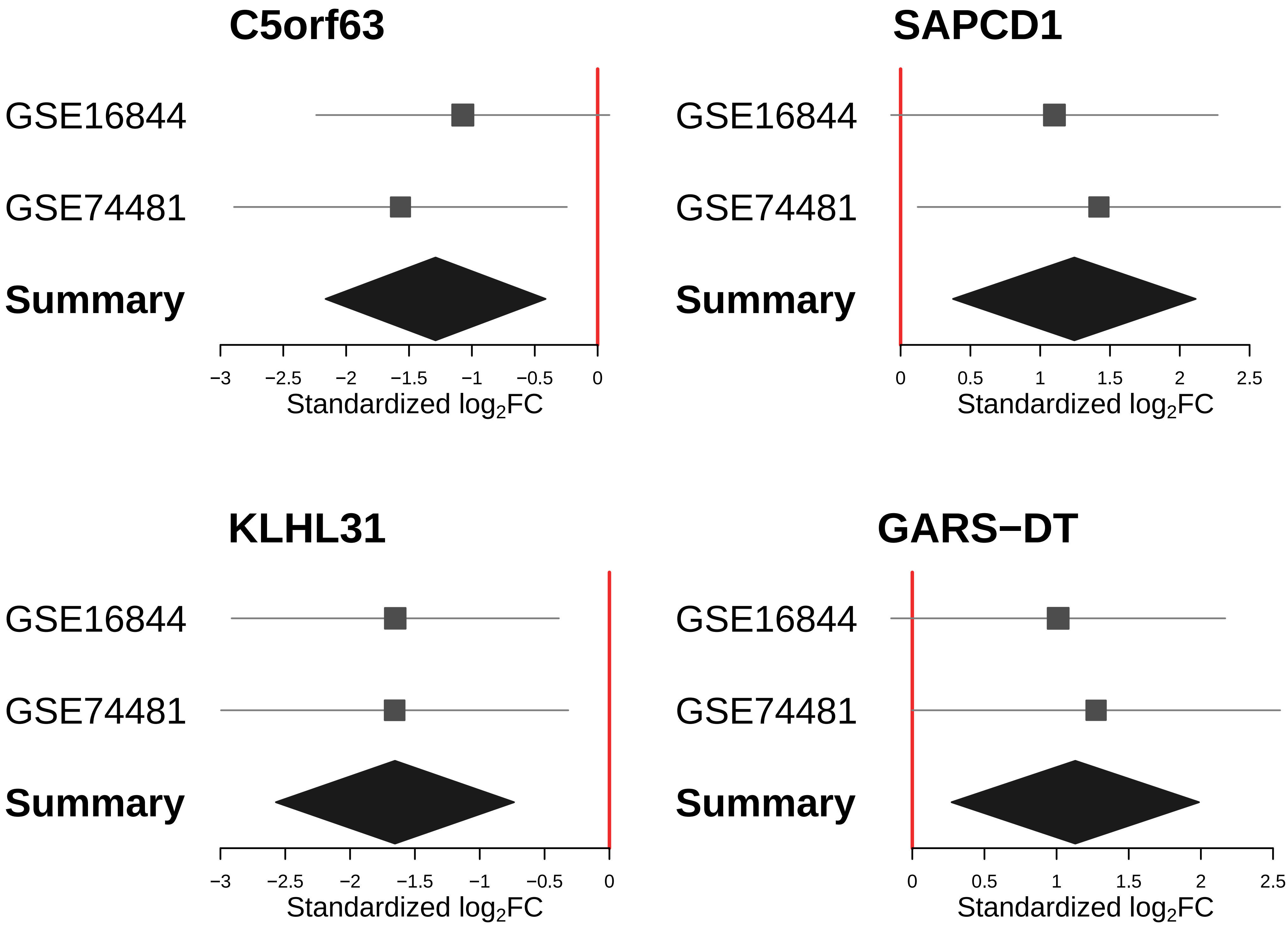
C5orf63
GSE16844
GSE74481
Summary
−3 −2.5 −2 −1.5 −1 −0.5 0
Standardized log2FC
SAPCD1
GSE16844
GSE74481
Summary
0 0.5 1 1.5 2 2.5
Standardized log2FC
KLHL31
GSE16844
GSE74481
Summary
−3 −2.5 −2 −1.5 −1 −0.5 0
Standardized log2FC
GARS−DT
GSE16844
GSE74481
Summary
0 0.5 1 1.5 2 2.5
Standardized log2FC

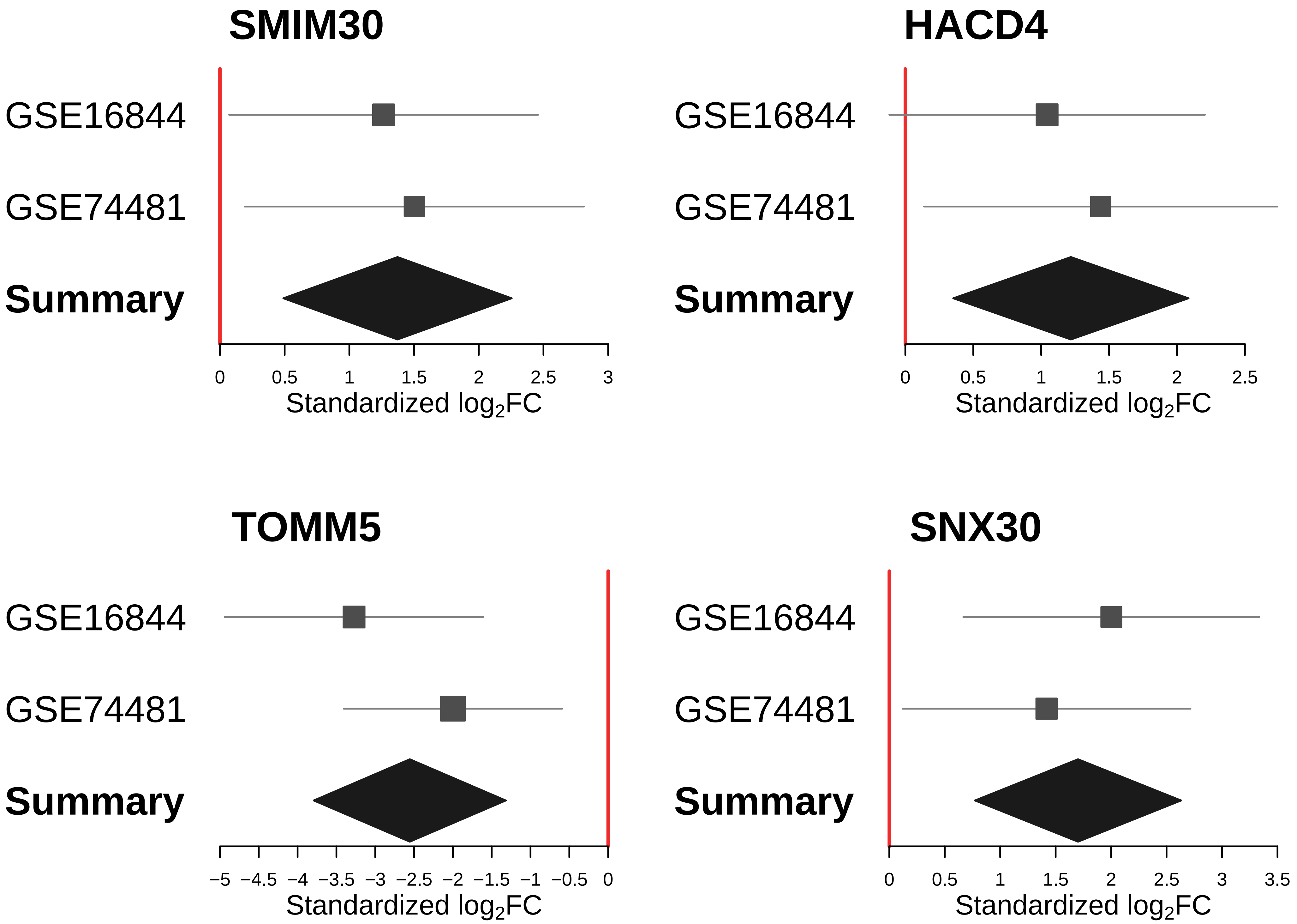
SMIM30
GSE16844
GSE74481
Summary
0 0.5 1 1.5 2 2.5 3
Standardized log2FC
HACD4
GSE16844
GSE74481
Summary
0 0.5 1 1.5 2 2.5
Standardized log2FC
TOMM5
GSE16844
GSE74481
Summary
−5 −4.5 −4 −3.5 −3 −2.5 −2 −1.5 −1 −0.5 0
Standardized log2FC
SNX30
GSE16844
GSE74481
Summary
0 0.5 1 1.5 2 2.5 3 3.5
Standardized log2FC

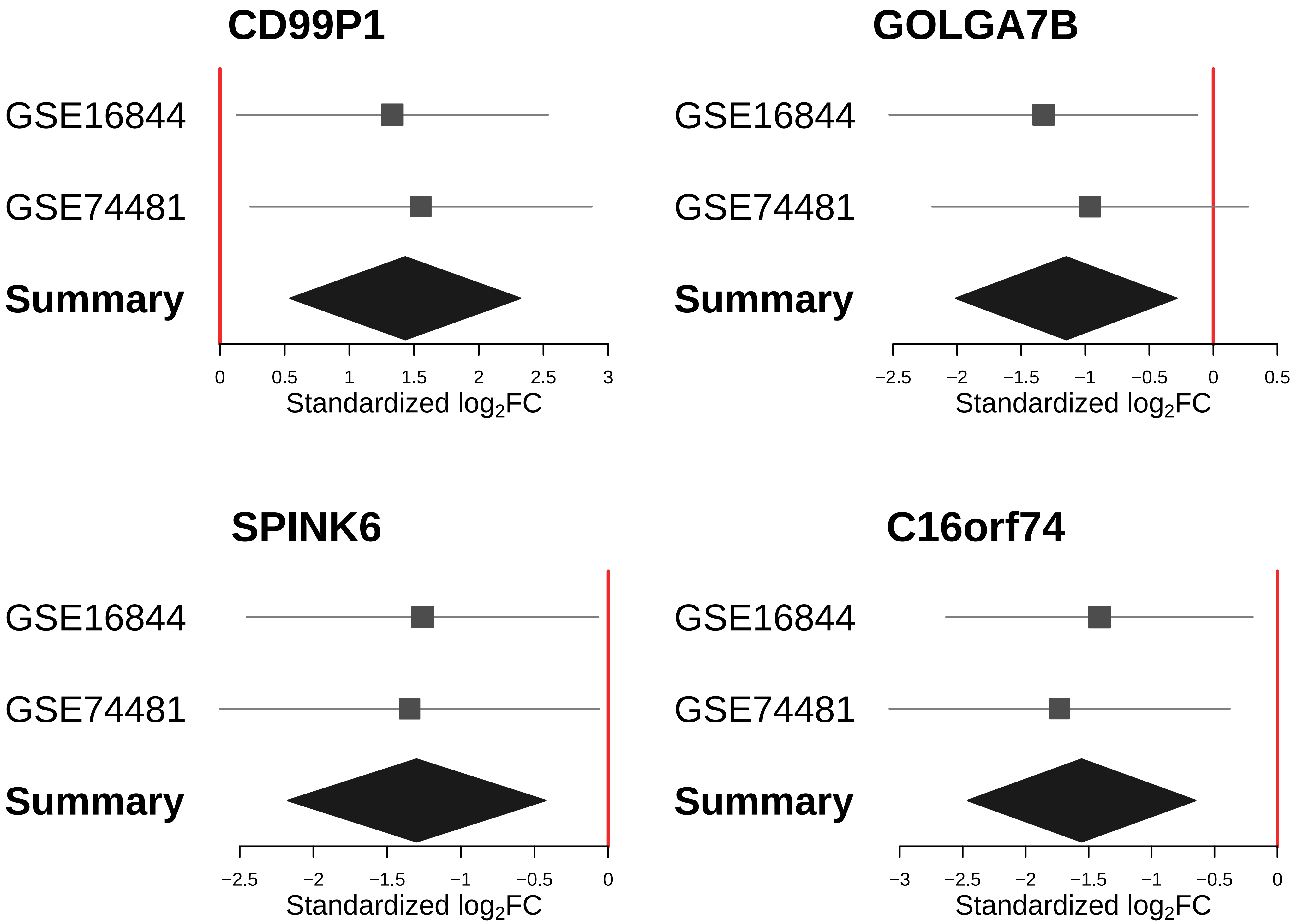
CD99P1
GSE16844
GSE74481
Summary
0
0.5
1
1.5
2
2.5
3
Standardized log2FC
GOLGA7B
GSE16844
GSE74481
Summary
−2.5
−2
−1.5
−1
−0.5
0
0.5
Standardized log2FC
SPINK6
GSE16844
GSE74481
Summary
−2.5
−2
−1.5
−1
−0.5
0
Standardized log2FC
C16orf74
GSE16844
GSE74481
Summary
−3
−2.5
−2
−1.5
−1
−0.5
0
Standardized log2FC

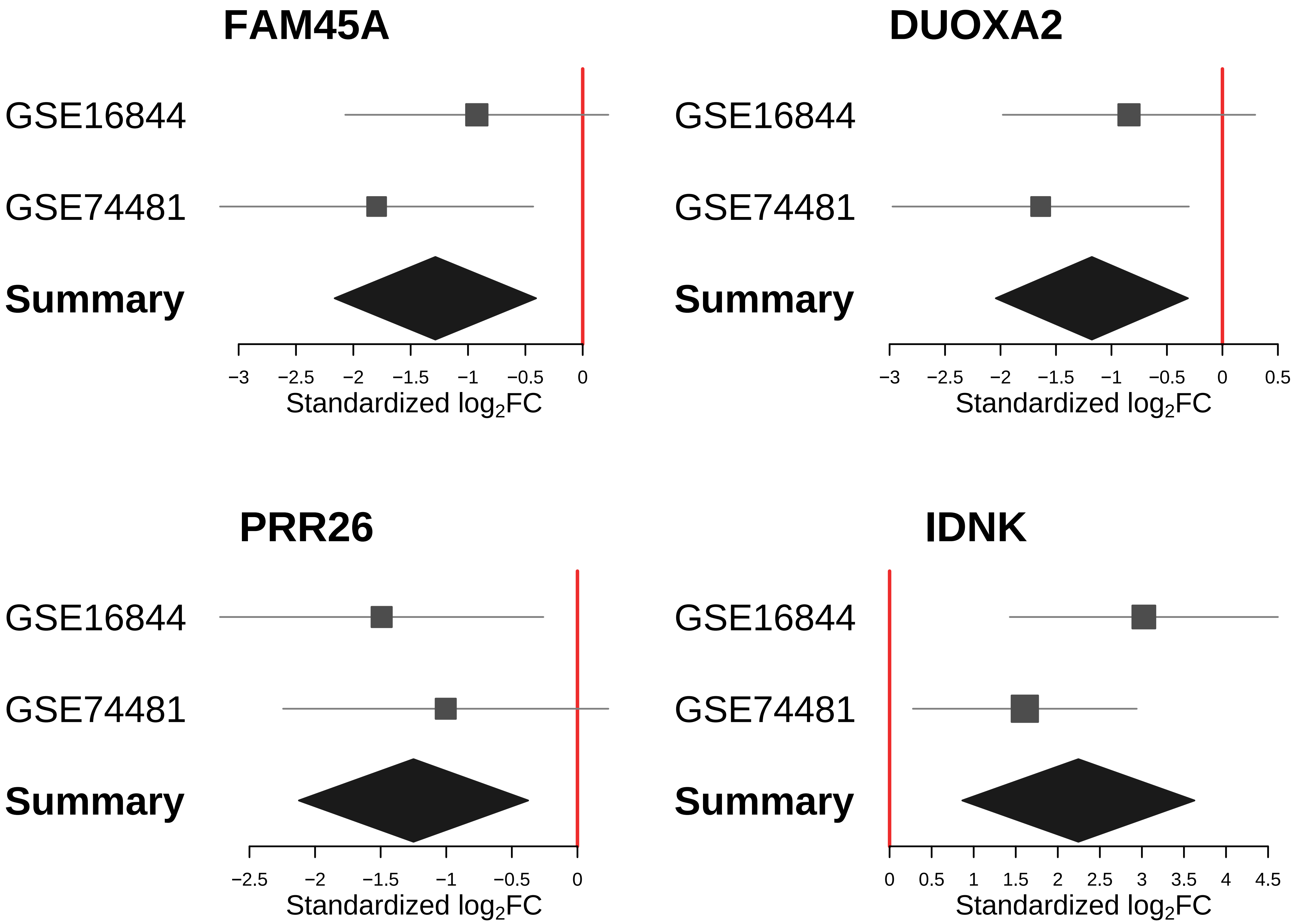

FAM45A
GSE16844
GSE74481
Summary
−3 −2.5 −2 −1.5 −1 −0.5 0
Standardized $log_2FC$
DUOXA2
GSE16844
GSE74481
Summary
−3 −2.5 −2 −1.5 −1 −0.5 0 0.5
Standardized $log_2FC$
PRR26
GSE16844
GSE74481
Summary
−2.5 −2 −1.5 −1 −0.5 0
Standardized $log_2FC$
IDNK
GSE16844
GSE74481
Summary
0 0.5 1 1.5 2 2.5 3 3.5 4 4.5
Standardized $log_2FC$

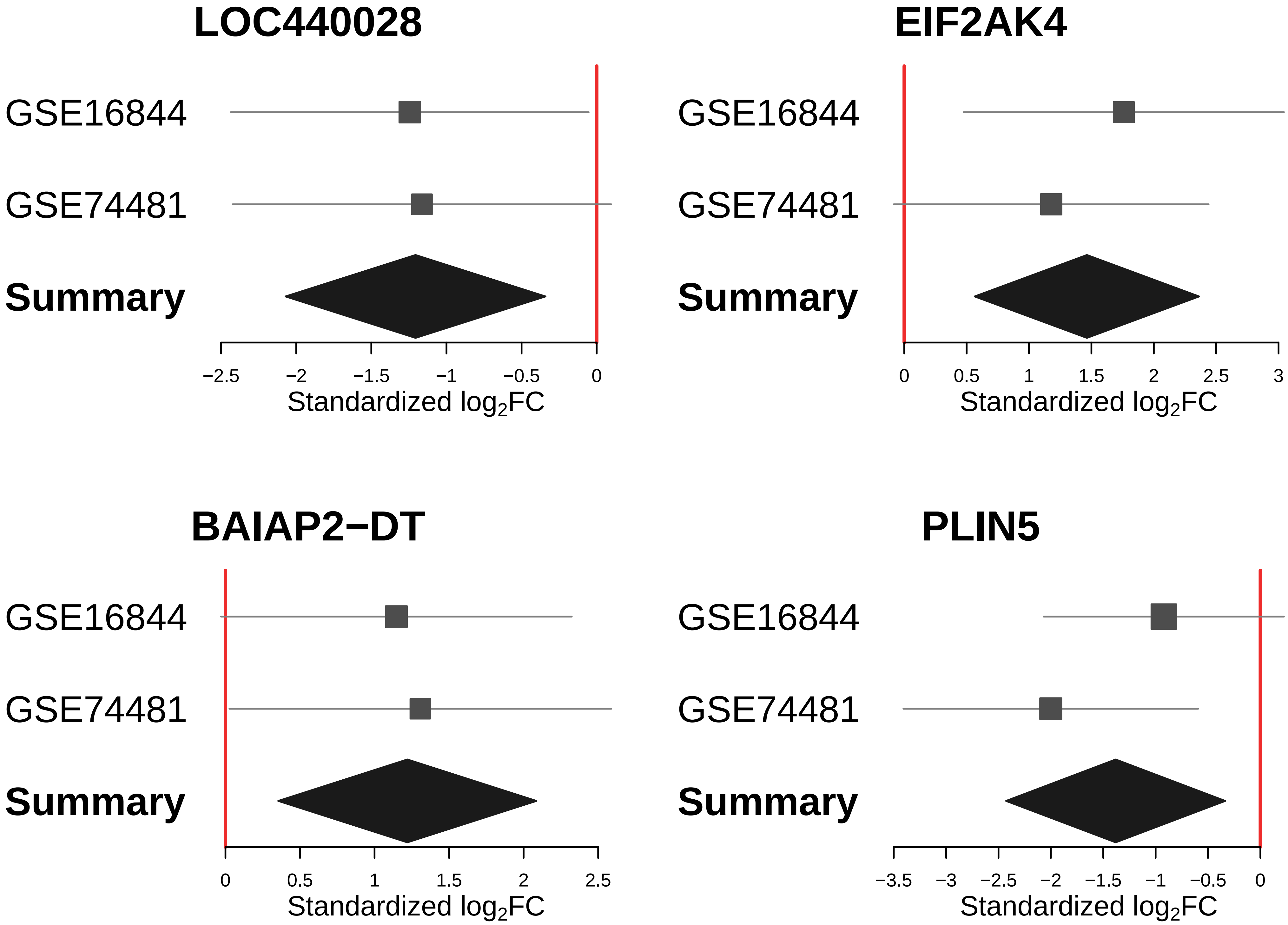
LOC440028
GSE16844
GSE74481
Summary
−2.5
−2
−1.5
−1
−0.5
0
Standardized $log_2FC$
EIF2AK4
GSE16844
GSE74481
Summary
0
0.5
1
1.5
2
2.5
3
Standardized $log_2FC$
BAIAP2−DT
GSE16844
GSE74481
Summary
0
0.5
1
1.5
2
2.5
Standardized $log_2FC$
PLIN5
GSE16844
GSE74481
Summary
−3.5
−3
−2.5
−2
−1.5
−1
−0.5
0
Standardized $log_2FC$

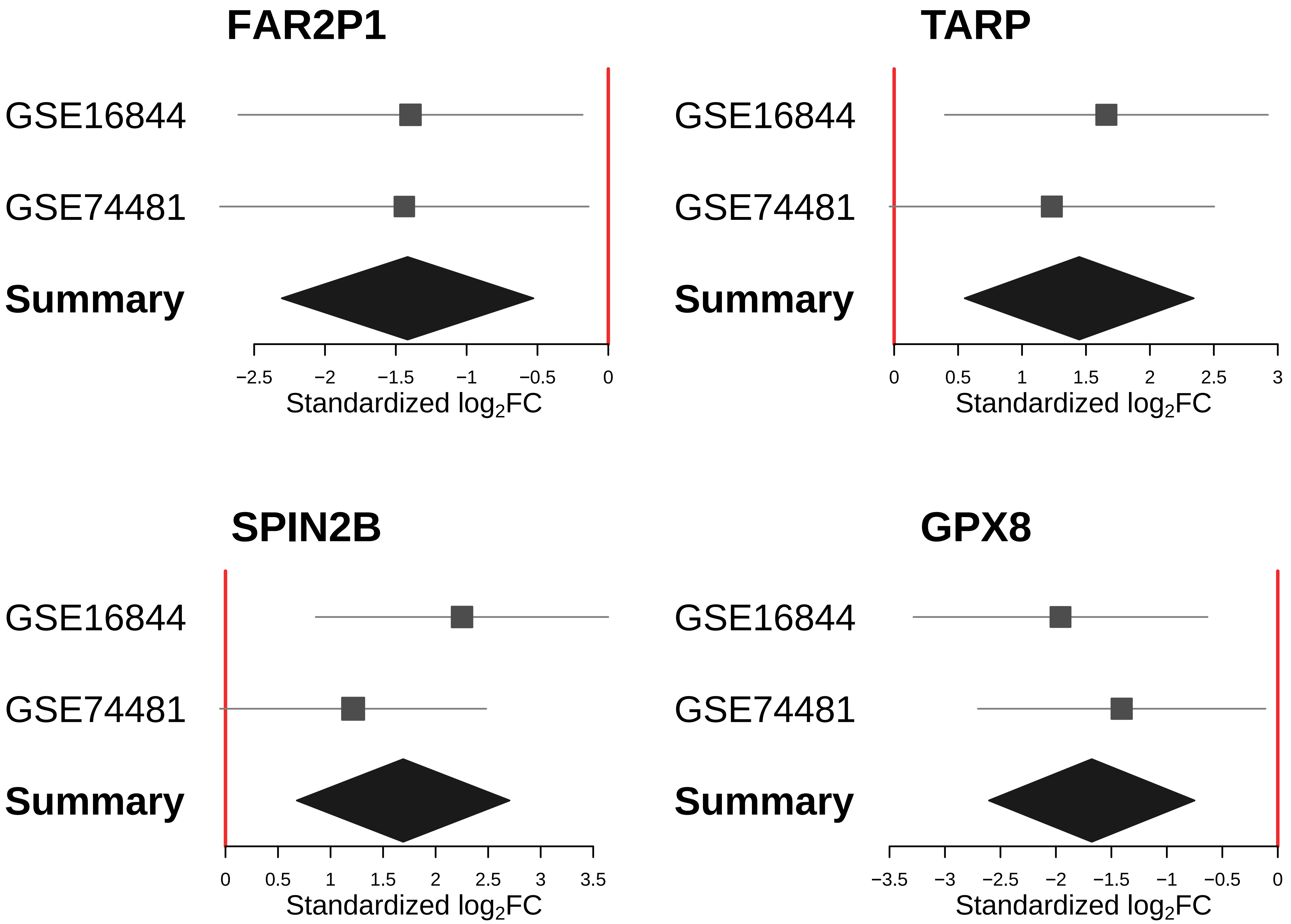

FAR2P1
GSE16844
GSE74481
Summary
−2.5 −2 −1.5 −1 −0.5 0
Standardized log2FC
TARP
GSE16844
GSE74481
Summary
0 0.5 1 1.5 2 2.5 3
Standardized log2FC
SPIN2B
GSE16844
GSE74481
Summary
0 0.5 1 1.5 2 2.5 3 3.5
Standardized log2FC
GPX8
GSE16844
GSE74481
Summary
−3.5 −3 −2.5 −2 −1.5 −1 −0.5 0
Standardized log2FC

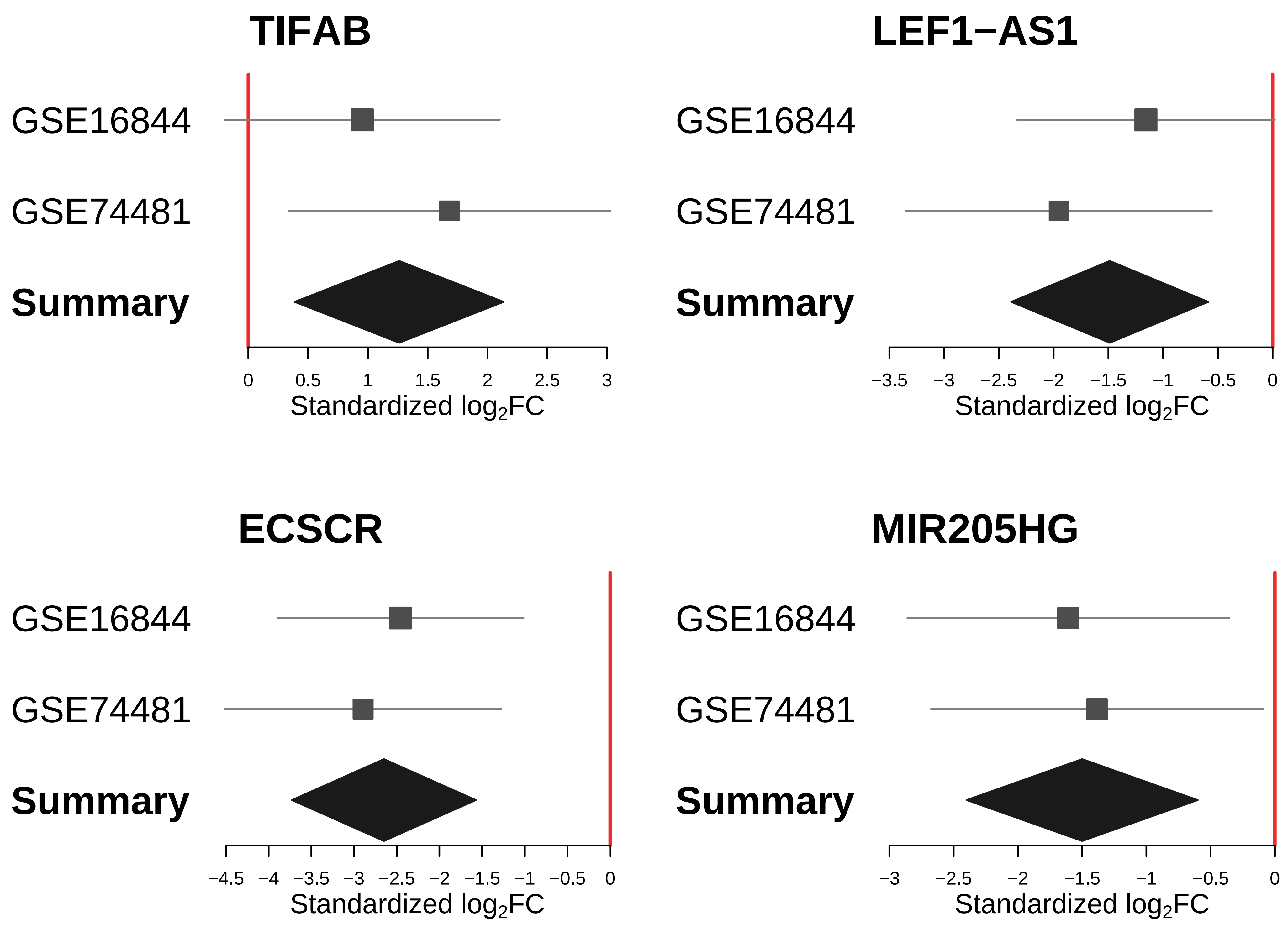

TIFAB
GSE16844
GSE74481
Summary
0
0.5
1
1.5
2
2.5
3
Standardized log2FC
LEF1−AS1
GSE16844
GSE74481
Summary
−3.5
−3
−2.5
−2
−1.5
−1
−0.5
0
Standardized log2FC
ECSCR
GSE16844
GSE74481
Summary
−4.5
−4
−3.5
−3
−2.5
−2
−1.5
−1
−0.5
0
Standardized log2FC
MIR205HG
GSE16844
GSE74481
Summary
−3
−2.5
−2
−1.5
−1
−0.5
0
Standardized log2FC

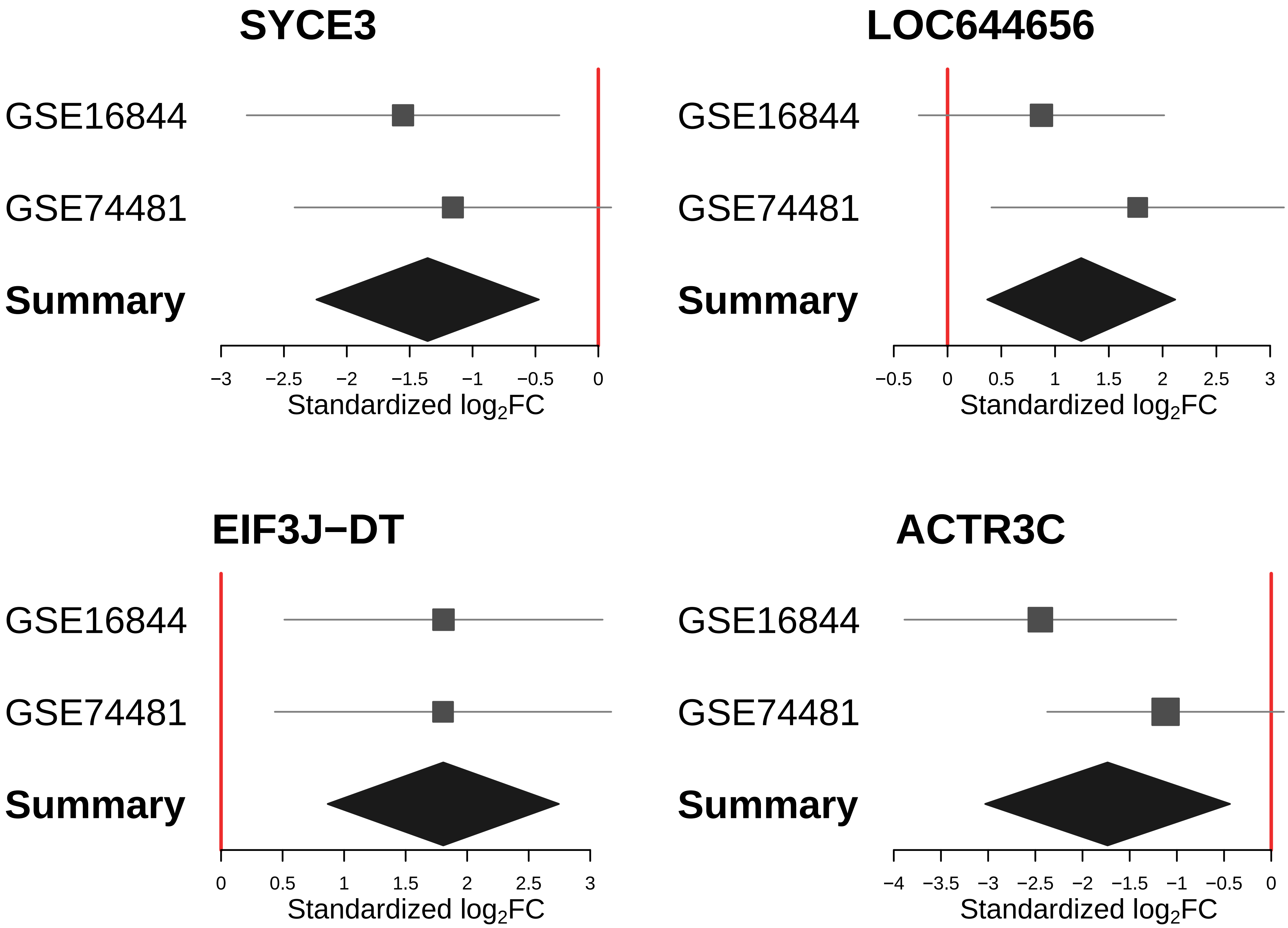

SYCE3
GSE16844
GSE74481
Summary
−3
−2.5
−2
−1.5
−1
−0.5
0
Standardized log2FC
LOC644656
GSE16844
GSE74481
Summary
−0.5
0
0.5
1
1.5
2
2.5
3
Standardized log2FC
EIF3J−DT
GSE16844
GSE74481
Summary
0
0.5
1
1.5
2
2.5
3
Standardized log2FC
ACTR3C
GSE16844
GSE74481
Summary
−4
−3.5
−3
−2.5
−2
−1.5
−1
−0.5
0
Standardized log2FC

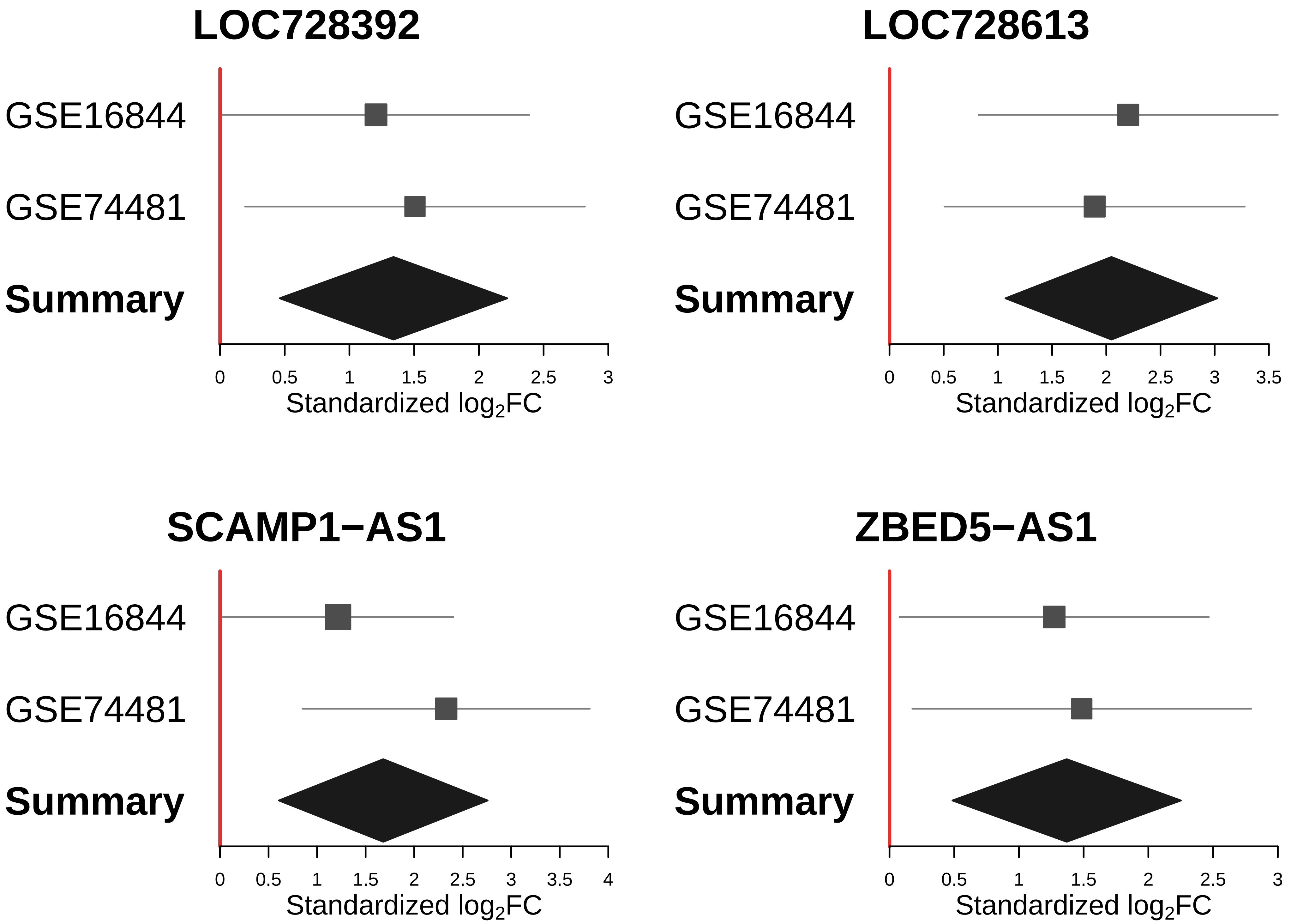
LOC728392
GSE16844
GSE74481
Summary
0 0.5 1 1.5 2 2.5 3
Standardized log2FC
LOC728613
GSE16844
GSE74481
Summary
0 0.5 1 1.5 2 2.5 3 3.5
Standardized log2FC
SCAMP1−AS1
GSE16844
GSE74481
Summary
0 0.5 1 1.5 2 2.5 3 3.5 4
Standardized log2FC
ZBED5−AS1
GSE16844
GSE74481
Summary
0 0.5 1 1.5 2 2.5 3
Standardized log2FC

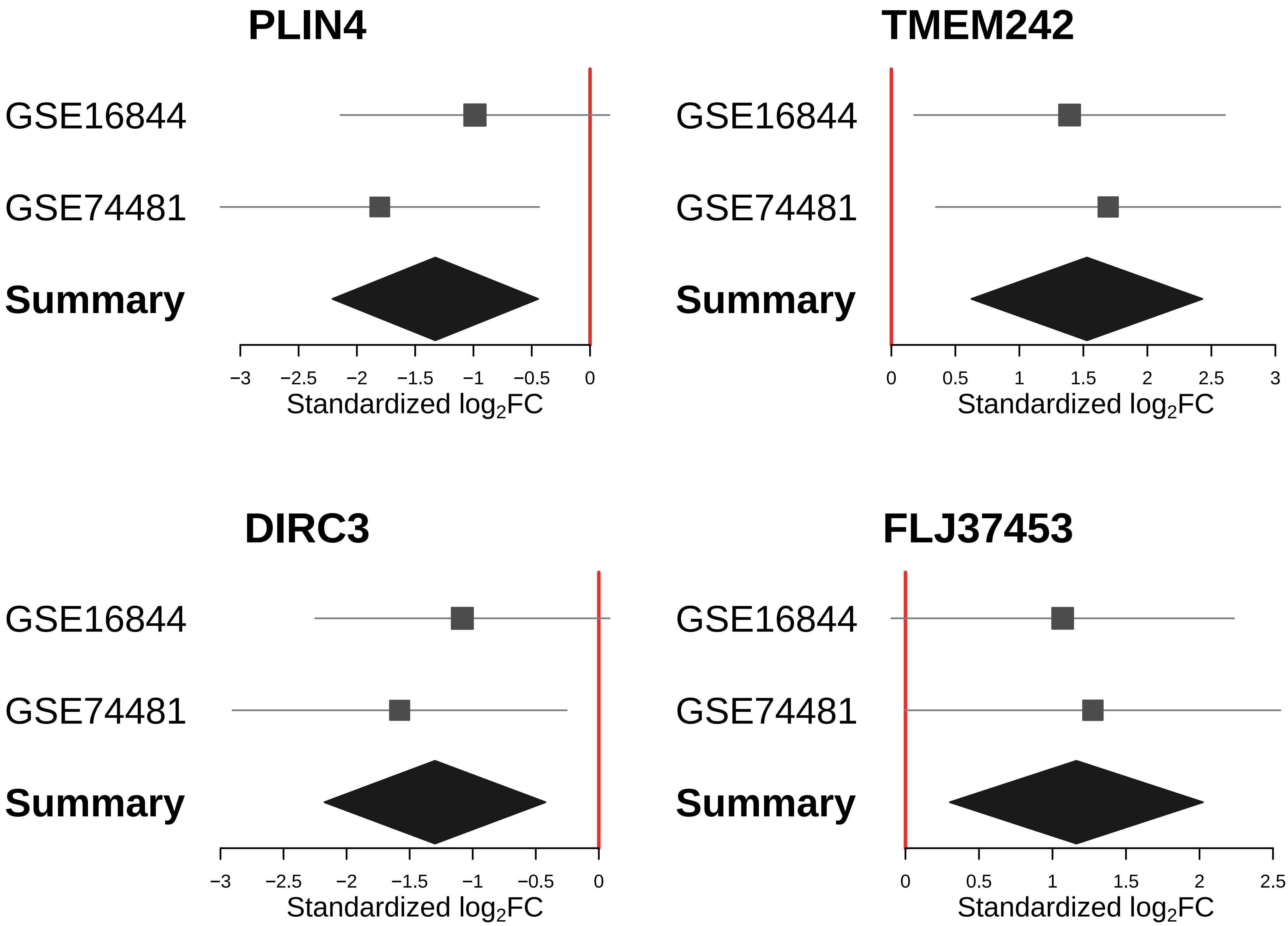
PLIN4
GSE16844
GSE74481
Summary
−3 −2.5 −2 −1.5 −1 −0.5 0
Standardized log2FC
TMEM242
GSE16844
GSE74481
Summary
0 0.5 1 1.5 2 2.5 3
Standardized log2FC
DIRC3
GSE16844
GSE74481
Summary
−3 −2.5 −2 −1.5 −1 −0.5 0
Standardized log2FC
FLJ37453
GSE16844
GSE74481
Summary
0 0.5 1 1.5 2 2.5
Standardized log2FC

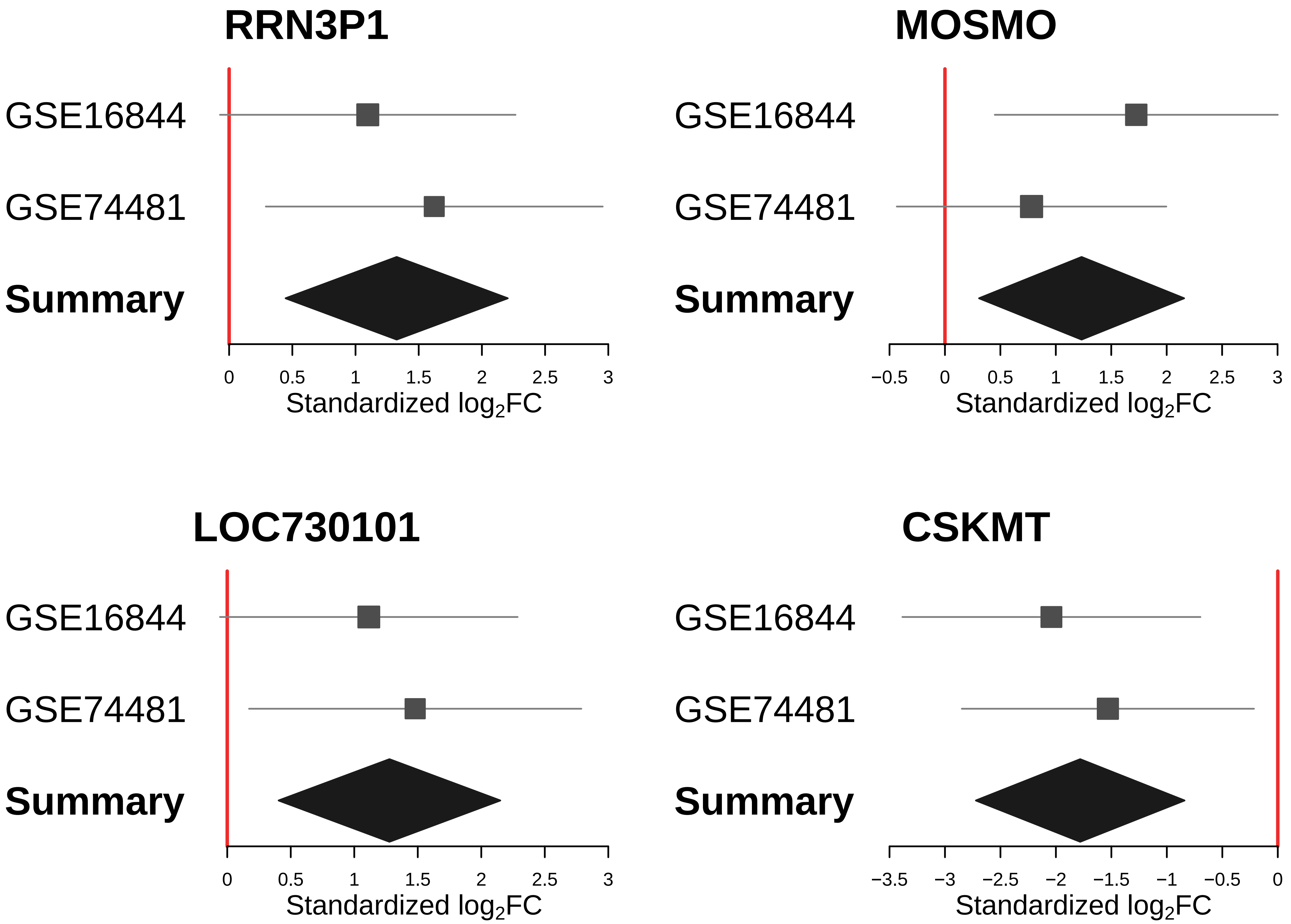
RRN3P1
GSE16844
GSE74481
Summary
0 0.5 1 1.5 2 2.5 3
Standardized log2FC
MOSMO
GSE16844
GSE74481
Summary
−0.5 0 0.5 1 1.5 2 2.5 3
Standardized log2FC
LOC730101
GSE16844
GSE74481
Summary
0 0.5 1 1.5 2 2.5 3
Standardized log2FC
CSKMT
GSE16844
GSE74481
Summary
−3.5 −3 −2.5 −2 −1.5 −1 −0.5 0
Standardized log2FC

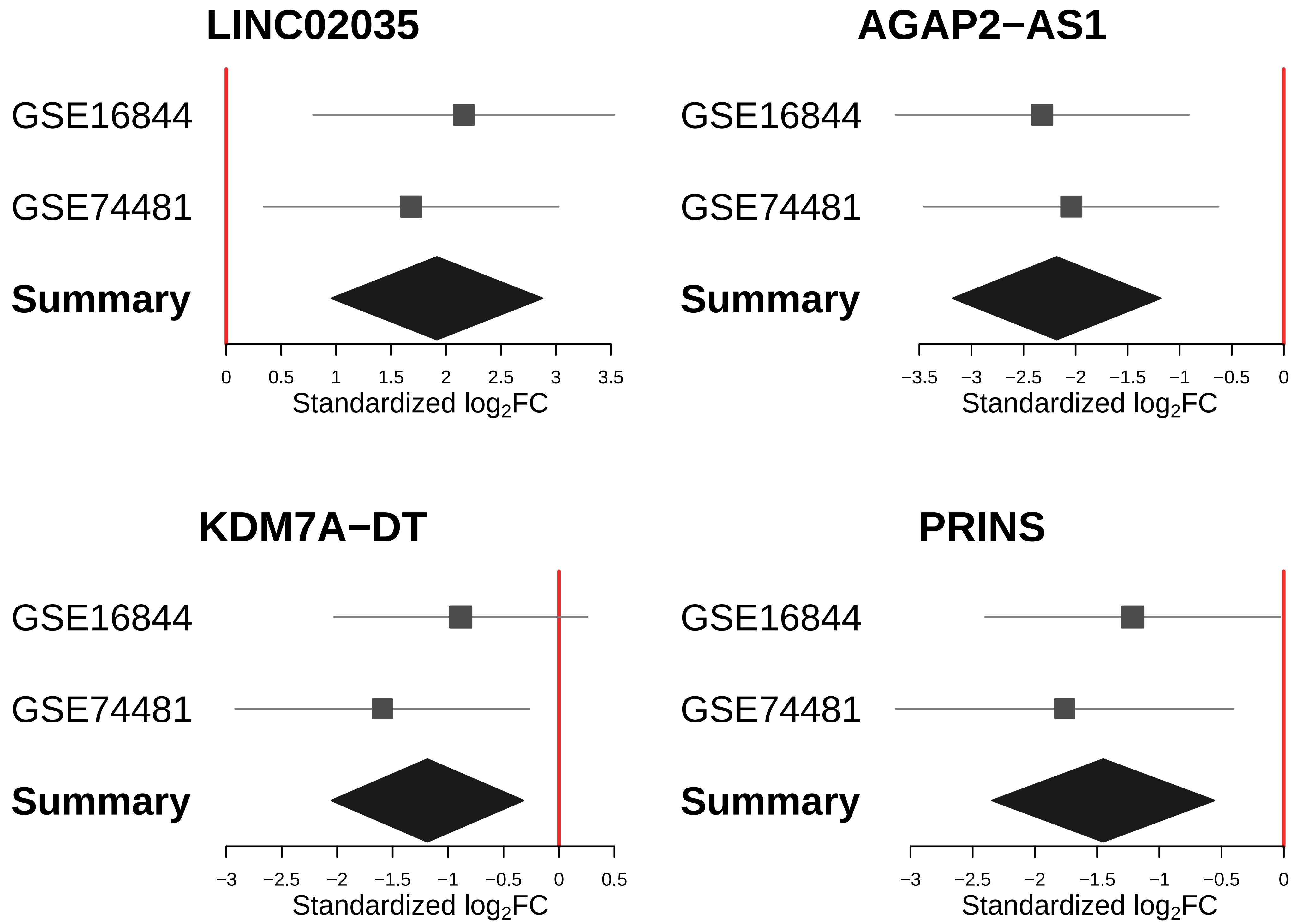
LINC02035
GSE16844
GSE74481
Summary
0 0.5 1 1.5 2 2.5 3 3.5
Standardized log2FC
AGAP2−AS1
GSE16844
GSE74481
Summary
−3.5 −3 −2.5 −2 −1.5 −1 −0.5 0
Standardized log2FC
KDM7A−DT
GSE16844
GSE74481
Summary
−3 −2.5 −2 −1.5 −1 −0.5 0 0.5
Standardized log2FC
PRINS
GSE16844
GSE74481
Summary
−3 −2.5 −2 −1.5 −1 −0.5 0
Standardized log2FC

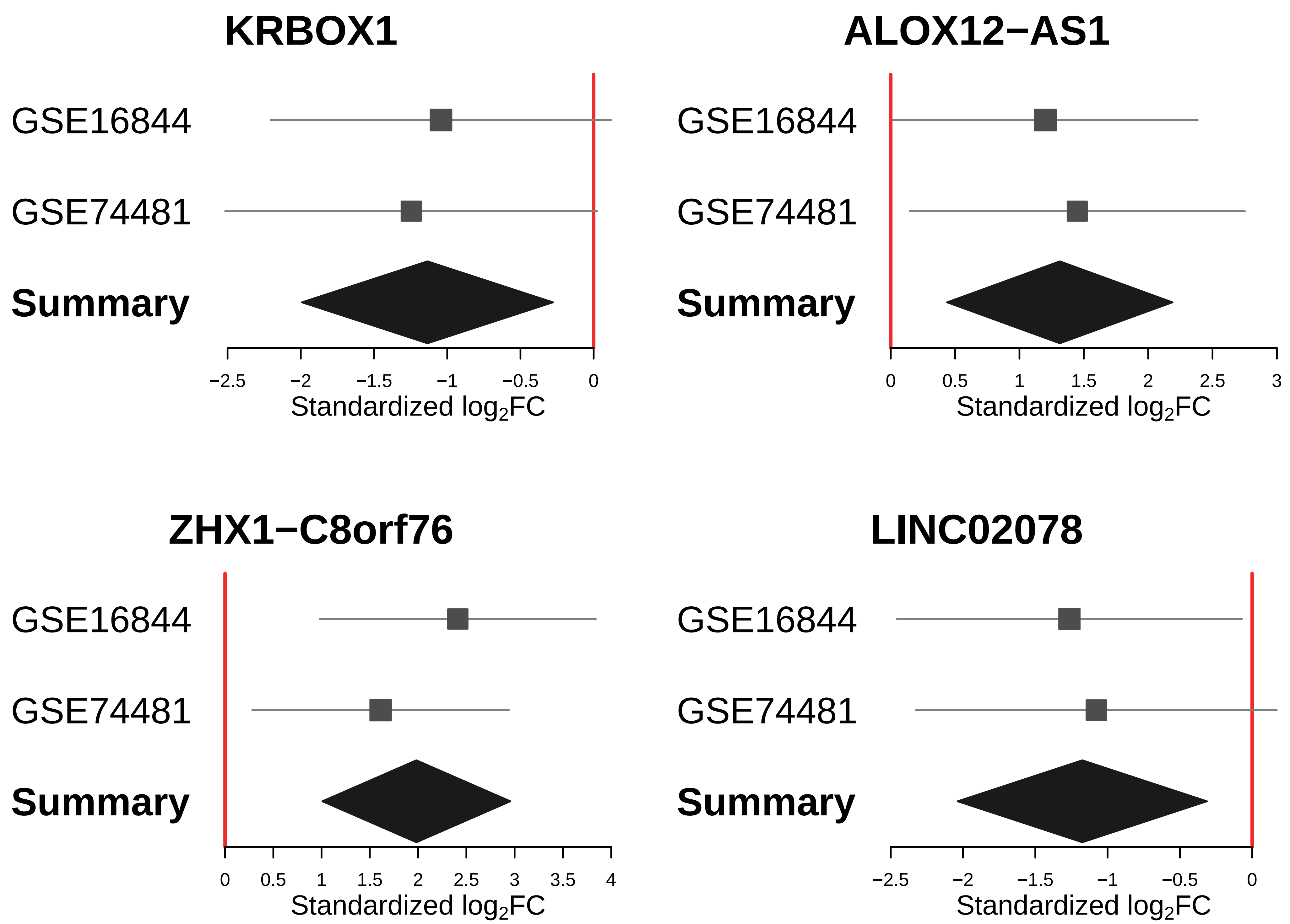
KRBOX1
GSE16844
GSE74481
Summary
−2.5 −2 −1.5 −1 −0.5 0
Standardized log2FC
ALOX12−AS1
GSE16844
GSE74481
Summary
0 0.5 1 1.5 2 2.5 3
Standardized log2FC
ZHX1−C8orf76
GSE16844
GSE74481
Summary
0 0.5 1 1.5 2 2.5 3 3.5 4
Standardized log2FC
LINC02078
GSE16844
GSE74481
Summary
−2.5 −2 −1.5 −1 −0.5 0
Standardized log2FC

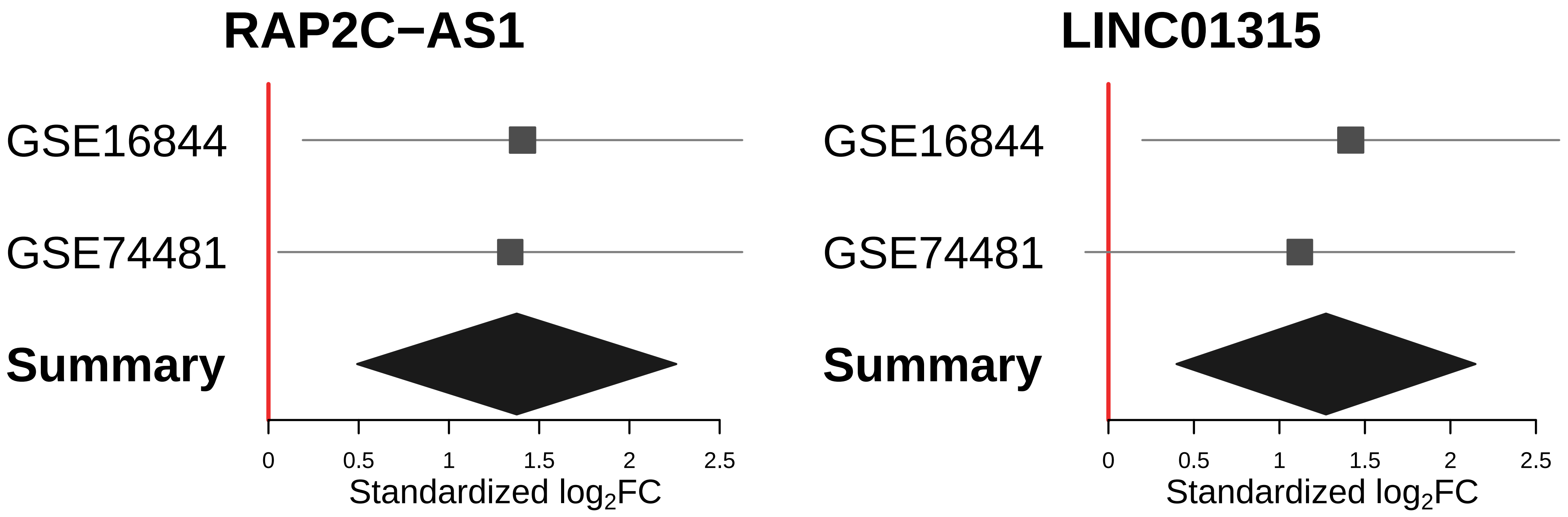
RAP2C−AS1
GSE16844
GSE74481
Summary
0
0.5
1
1.5
2
2.5
Standardized $log_2$FC
LINC01315
GSE16844
GSE74481
Summary
0
0.5
1
1.5
2
2.5
Standardized $log_2$FC
