## Supplementary figures and images for "Reanalysis And Integration Of Public Microarray Datasets Reveals Novel Host Genes Modulated In Leprosy" (from another version of this preprint: version 2, 2019-12-07)

### FigS1.pdf

#### Figure S1 - Enriched biological processes for the LL vs. Control genes (selected by sdef)

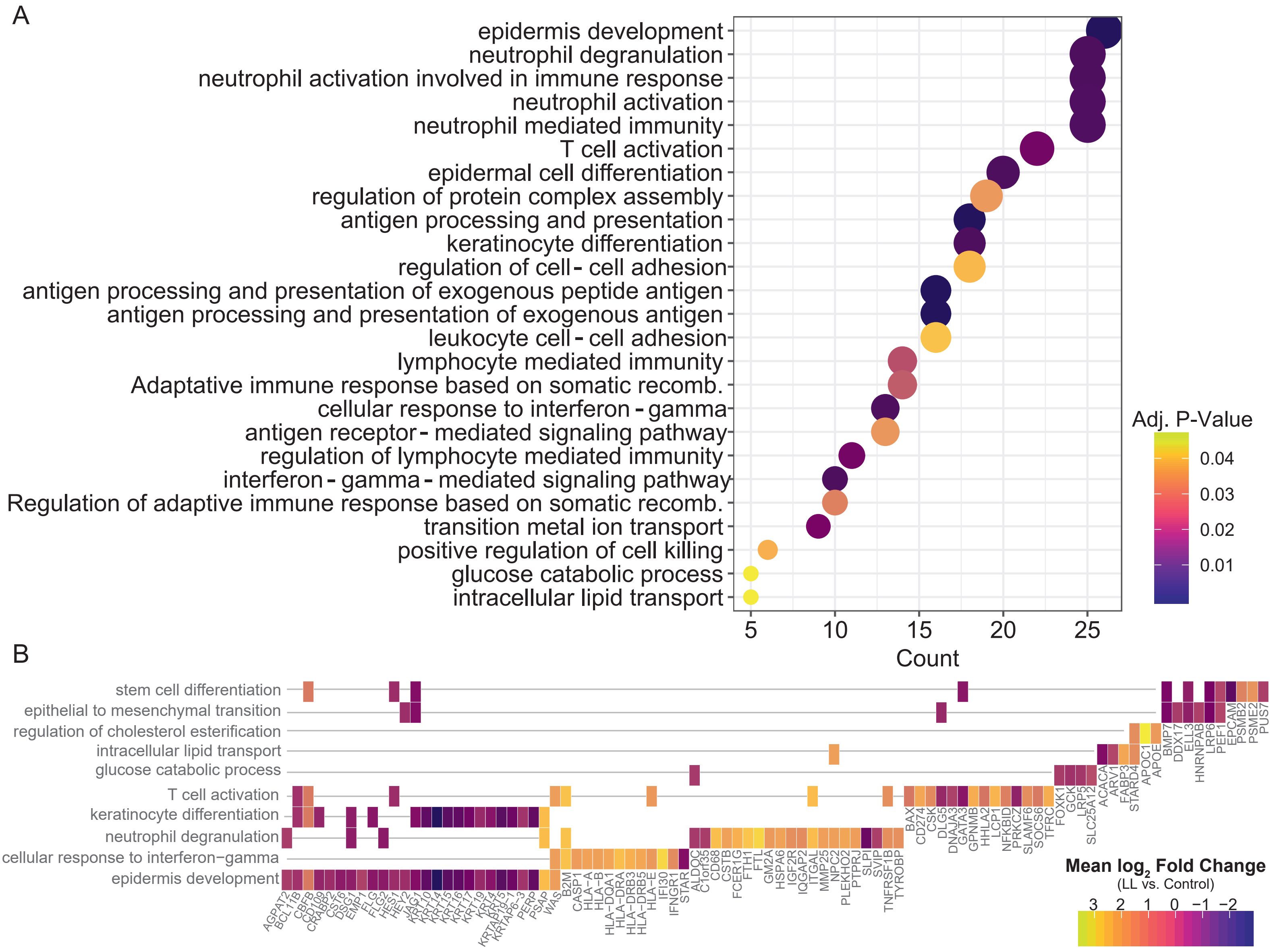
